# Predicting disease-causing variant combinations

**DOI:** 10.1101/520353

**Authors:** Sofia Papadimitriou, Andrea Gazzo, Nassim Versbraegen, Charlotte Nachtegael, Jan Aerts, Yves Moreau, Sonia Van Dooren, Ann Nowé, Guillaume Smits, Tom Lenaerts

## Abstract

Notwithstanding important advances in the context of single-variant pathogenicity identification, novel breakthroughs in discerning the origins of many rare diseases require methods able to identify more complex genetic models. We present here the Variant Combinations Pathogenicity Predictor (VarCoPP), a machine-learning approach that identifies pathogenic variant combinations in gene pairs (bi-locus variant combinations). We show that the results produced by this method are highly accurate and precise, an efficacy that is endorsed when validating the method on recently published independent disease-causing data. Confidence labels of 95% and 99% are identified, representing the probability of a bi-locus combination being a true pathogenic result, providing geneticists with rational markers to evaluate the most relevant pathogenic combinations and limit the search space and time. Finally, VarCoPP has been designed to act as an interpretable method that can provide explanations on why a bi-locus combination is predicted as pathogenic and which biological information is important for that prediction. This work provides an important new step towards the genetic understanding of rare diseases, paving the way to new clinical knowledge and improved patient care.

## INTRODUCTION

Advances in high throughput-sequencing technologies and the application of massive parallel sequencing have revolutionized the field of human genetics, providing a huge amount of information on human genetic variation(1–5). Interpreting this variation has provided important insights into the genetic architecture of many rare diseases, notably those inherited in a mendelian pattern(6–8), and has opened the path to promising preventive, diagnostic and therapeutic strategies(9). The amount of genetic data available also allowed for the development of successful predictive tools integrating genetic, molecular, evolutionary and/or structural information(10–13). Such tools are routinely applied in clinics to identify pathogenic variants potentially associated with a specific disease phenotype. Notwithstanding these advancements, the analysis of a growing number of rare human disorders has highlighted the difficulties in establishing a genotype-phenotype relationship due to non-mendelian patterns of inheritance, incomplete penetrance, phenotypic variability or locus heterogeneity(14–18). The classic concept of one gene leading to a particular phenotype appears to be an oversimplification, since to better explain the situation of an affected individual one often needs to consider more complex genetic models where mutations in multiple genes cause or modulate the development of one or several simultaneous disease phenotypes(15, 19–21). Although the terms “locus” and “gene” can refer to different types of genetic elements, in this paper they are used interchangeably.

Oligogenic or multi-locus genetic patterns have already been discovered for diseases initially considered to be monogenic, for instance phenylketonuria(22) or hereditary non-syndromic deafness(23). These types of diseases may have a central primary causative gene and a network of modifier genes like in Hirschsprung disease(24) and cystic fibrosis(25), or present a spectrum of genetic models from monogenic to polygenic, as in the case of neurodevelopmental disorders(26, 27). Gene-disease network analysis studies further support the notion that a disease phenotype is hardly the result of a mutation in one gene alone, showing that the vast majority of mendelian diseases are actually modulated by multiple genes that are usually involved in similar pathways or cellular and biological processes (28, 29). Along with the cases where a phenotype or syndromic phenotypes can be modulated by several genes, a multi-locus genetic pattern can also be observed in an affected individual where disease-causing monogenic mutations in different genes segregate independently, leading to multiple independent molecular clinical diagnoses (21, 30–33). These multiple diagnoses cases can affect different tissues (distinct), but others can share phenotypes (overlapping), indicating a possible relationship between the involved multi-locus variations at the protein or cellular level. It is evident that in order for the clinical predictive tools to remain valuable for diagnostic purposes, they need an update towards these more elaborate biological and inheritance scenarios. For instance, such tools will need to consider that the nature or frequency of variants observed in oligogenic diseases will be different from those observed in monogenic ones (20, 34). The current work makes this leap, introducing and validating a novel computational approach that predicts the pathogenicity of variant combinations as opposed to single variants, and this within the context of gene pairs.

This leap is made possible by the steady increase in literature reports on disease-causing variant combinations in gene pairs (bi-locus variant combinations) in the last decades, which have been grouped and made publicly available via the online resource DIDA, the Digenic Diseases Database(35). This novel resource collects, organizes and annotates cases where a bi-locus genetic model helped to explain a patient’s phenotypic variability and reduced penetrance, including for example the well-known cases of Bardet-Biedl syndrome (BBS)(36, 37) and retinitis pigmentosa(38). The first version of the database (which will be referred henceforth as DIDAv1) contains 213 manually curated bi-locus variant combinations obtained from independent scientific papers, involving 136 different genes and leading to 44 diseases. These variant combinations can be divided into three different classes of bi-locus diseases (Fig. 1): the first class, referred to as “true digenic class”, requires the presence of variants in two independent genes to trigger the disease, with carriers of the variants found in one gene being unaffected. The second class covers mendelizing variants with modifiers, which is referred to as the “composite class”. In this scenario, the individual carrying the mendelizing variant can present symptoms of the disease, with the extra variant at the second gene modifying the severity of the symptoms or the age of onset. DIDAv1 also contained few cases of a third class, which is referred to as the “dual molecular diagnoses” class. These cases consist of the independent segregation of disease-causing mendelizing variants in two different genes leading to two independent clinical diagnoses. Given their limited number in DIDAv1, they were also grouped into the composite class. An initial study on DIDAv1 revealed that biological features defined at the variant, gene and combination-level are sufficient to differentiate composite from true bi-locus variant combinations, providing novel insights into the properties of disease-causing variant combinations(39).

**Fig. 1.**
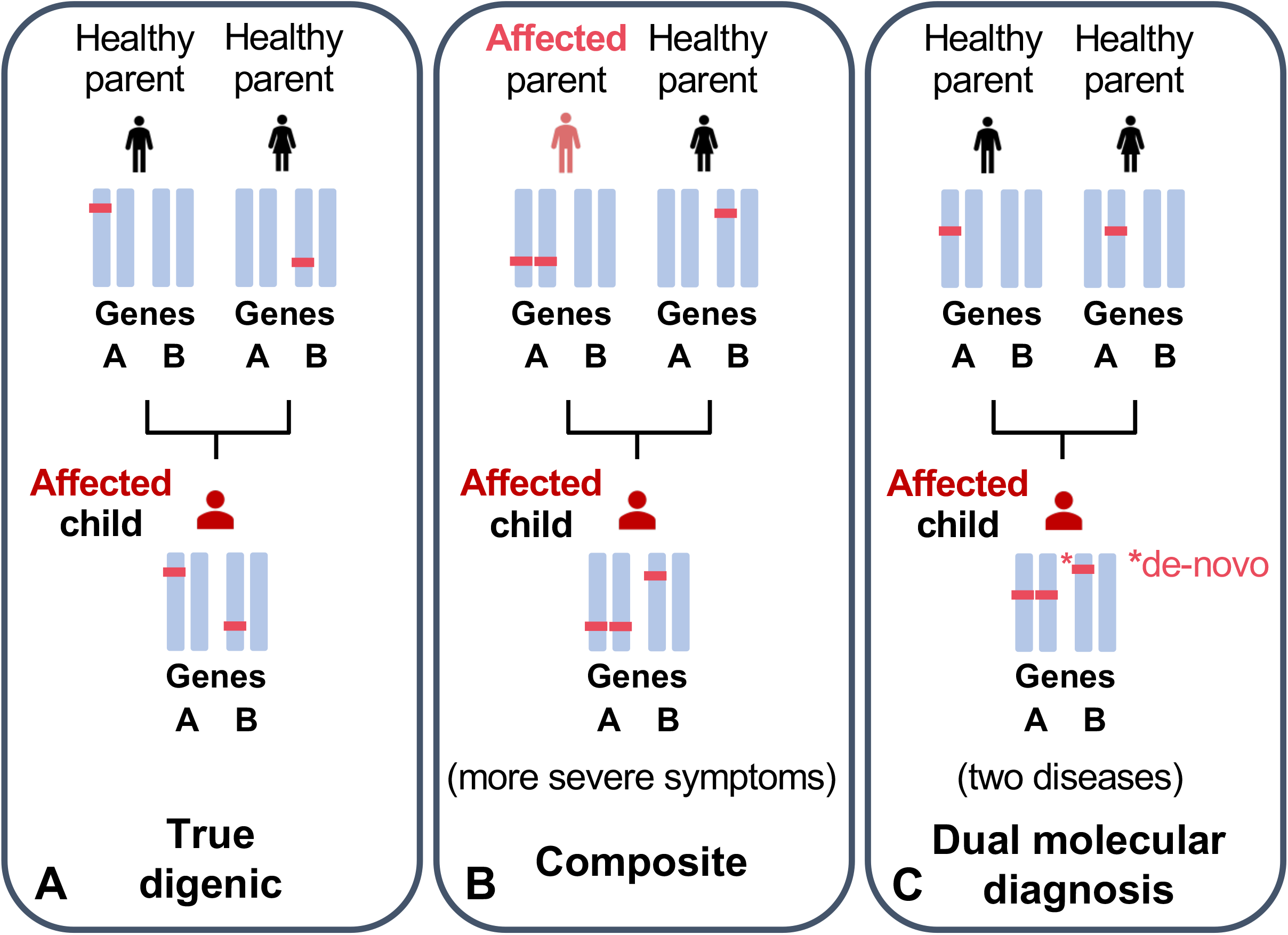
Examples of different cases of disease-causing bi-locus variant combinations present in an individual, and which can be detected by VarCoPP. (A) A “true digenic” case, where mutations on both genes should be present to trigger any symptoms of the disease. Individuals with the mutation in either one of the two genes remain unaffected. (B) One example of a “composite” case, where one mutation at the most pathological gene can be sufficient to show disease symptoms (see affected parent), but the second mutation affects the severity of symptoms or the age of onset. (C) One example of a dual molecular diagnosis case, which concerns the simultaneous aggregation of variants that cause two independent Mendelian diseases, with or without overlapping phenotypes.

Based on the presence of this fully annotated bi-locus disease data in DIDAv1 and the variety of cases it covers, one can hypothesize that the transition from single to variant combination pathogenicity predictors is now possible, starting from variant combinations within gene pairs (which we will refer to in this paper as bi-locus variant combinations). Such a predictor should exclude the non-relevant variant combinations (true negatives, TN), which will be abundantly present in a patient’s exome, and accurately identify the scarce disease-causing ones (true positives, TP). To meet this challenge, we developed VarCoPP (Variant Combination Pathogenicity Predictor), the first pathogenicity predictor for combinations of variants in gene pairs, which is able to accurately identify disease-causing variant combinations using variant, gene and gene pair information. The accuracy and sensitivity of the predictor is also validated on an independent data set consisting of new bi-locus diseases data from novel publications that appeared after the construction of DIDAv1. Moreover, by visualizing how each feature guides the pathogenicity prediction, VarCoPP provides an explanation on why a given bi-locus variant combination is classified as disease causing or not. To further support clinical geneticists in their analysis, statistical scores for each prediction, as well as 95%- and 99%-confidence labels for each evaluated combination, are provided. These labels capture the most relevant variant combinations that should be further analyzed clinically and are potential candidates of relevant information for patient counselling. VarCoPP is available online at: http://varcopp.ibsquare.be/.

## RESULTS

### Curation of the 1KGP data reveals the presence of known disease-causing bi-locus variant combinations

In total, 46% of the individuals in the 1000 Genomes Project (1KGP) carry at least one variant found in DIDAv1 (Fig. 2). The majority of the overlapping variants (86%) are involved in disease-causing variant combinations belonging to the true digenic class, thus explaining their monogenic presence in a control population. Nevertheless, more than 10% of overlapping variants are involved, in DIDAv1, in bi-locus combinations with a composite effect. Most of the variants found in 1KGP (69%) are located in the secondary (modifier) gene of the pair, possibly explaining why the control individuals carrying them could be asymptomatic. However, the rest of the overlapping variants are located in the primary (mendelizing) gene and some of them have been shown to cause disease symptoms in individuals in a dominant monogenic fashion, like the variants c.511C>T and c.637G>A in the WNT10A gene which are involved in tooth agenesis(40), the variant c.670G>A in the PDX1 gene involved in the development of maturity-onset diabetes of the young 4 (MODY 4)(41, 42), and the c.313G>A variant in the SLC7A9 gene involved in nontype I cystinuria(43, 44). It should be noted that MODY could be overlooked as the c.670G>A variant can present incomplete penetrance(41) - also suggested by its frequency in the ExAC database (0.002113) - while incomplete penetrance is also well-known for non-type I cystinuria. Tooth agenesis could also be easily clinically overlooked.

**Fig. 2.**
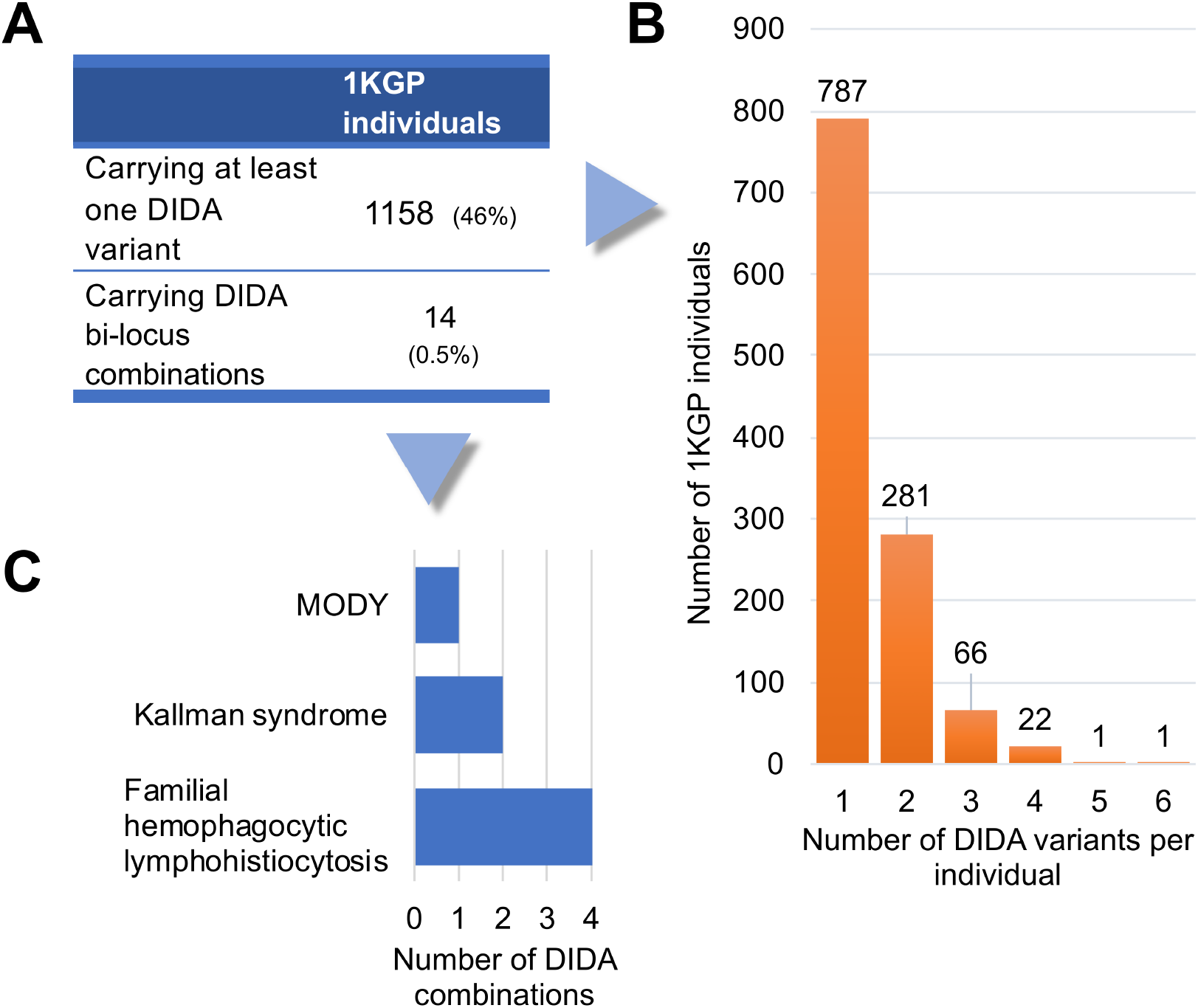
Overlapping variants and bi-locus combinations between DIDA and the 1KGP. (A) Statistics on 1KGP individuals carrying at least one DIDA independent variant or a disease-causing bi-locus combination. (B) Histogram of 1KGP individuals carrying one or more independent DIDA variants. (C) Histogram of the DIDA bi-locus combinations found in 1KGP and the diseases they are leading to.

Intriguingly, we discovered 7 disease-causing bi-locus combinations present in DIDAv1, leading to either MODY(42), Kallman syndrome(45) or familial hemophagocytic lymphohistiocytosis(46) in 14 individuals of the 1KGP (Fig. 2 and *SI Appendix*, Table S1). These combinations were not supported by functional evidence in the original studies and had not been compared with a large control cohort to further statistically ensure their relevance. However, some of the involved pairs were supported by familial evidence in their original papers (see *SI Appendix*, Text S1 for detailed information). From a clinical point of view, the individuals could also be un-diagnosed: a mild Kallman Syndrome could be easily clinically overlooked and, as stated beforehand, the c.670G>A variant for MODY can present incomplete penetrance. Furthermore, the bi-locus combinations could be incompletely penetrant. To ensure that the data used for the construction of VarCoPP does not contain contradicting instances, we removed from our analysis these 14 individuals from the 1KGP neutral set, as well as the 7 incriminated bi-locus combinations from DIDAv1 as a precaution.

### VarCoPP identifies accurately pathogenic variant combinations

Using bi-locus variant combinations randomly selected from individuals of the 1KGP(5) as the neutral set and the bi-locus variant combinations from DIDAv1(35) as the disease causing set, we successfully trained a variant combination pathogenicity predictor (VarCoPP) (see Fig. 3 for a summary of the procedure). We limited the search space to 1KGP variants with up to 3% minor allele frequency (MAF) to match the frequency range observed in DIDAv1, located in or close to exons. Each gene and variant in the combination were ordered in the same way for both data sets, a process necessary for reliability (see Fig. 3A and Materials and Methods). We then annotated our data with information at the variant, gene and gene-pair level, leading to 21 molecular and bioinformatics characteristics (computationally called “features”) per bi-locus combination (see Fig. 3B and *SI Appendix*, Table S2 and S3). This annotated information is then used as input by the predictor to identify the class label of the tested bi-locus combination, i.e. pathogenic or neutral.

**Fig. 3.**
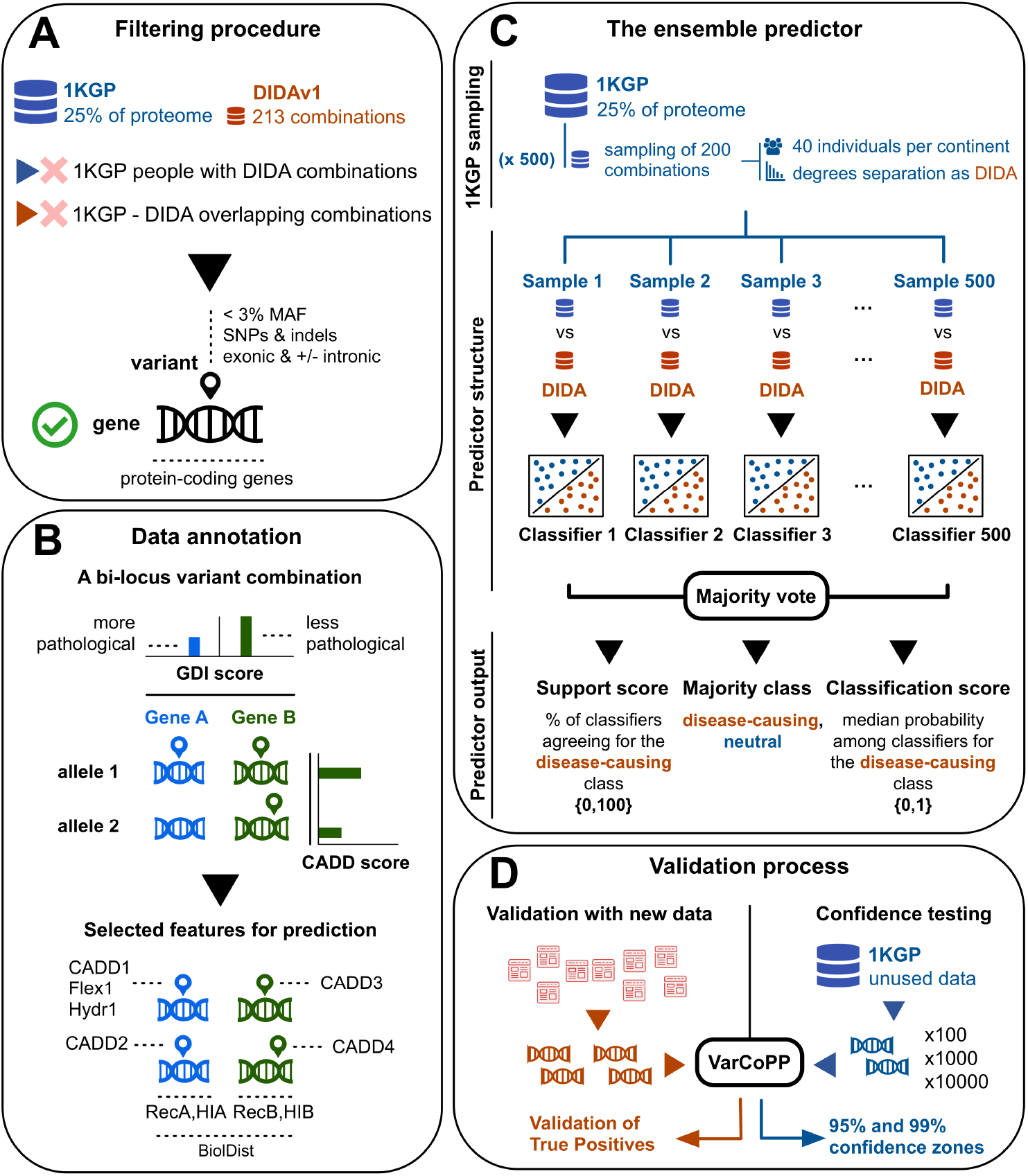
Summary of the methodology procedure for the construction of VarCoPP and the validation process (A) Genes and variants were filtered in the same way for both 1KGP and DIDAv1. Individuals of the 1KGP project carrying DIDAv1 combinations, as well as the overlapping combinations, were filtered out. Exonic variants (SNPs and indels) were used with a MAF frequency of ≤ 3%, including intronic and synonymous variants close to the exon edges (±13 nt). The genes involved in the procedure were only confirmed protein-coding genes, following the gene types present in DIDAv1. (B) A bi-locus variant combination is represented using four variant alleles (two alleles for gene A and two alleles for gene B). Gene A is the gene with the lowest Gene Damage Index (GDI) score, thus with the higher probability of being a pathological gene. Variant alleles inside the same gene were ordered based on their CADD pathogenicity score, with the first variant allele of that gene always having the highest CADD score. The initial number of biological features used for classification was 21, but the final selected and more relevant features were filtered to 11. These included information at the variant level (Flex1 and Hydr1, i.e. flexibility and hydrophobicity amino acid differences of the first variant allele of gene A, as well as CADD1, CADD2, CADD3, CADD4, i.e. the CADD scores of the four alleles of a bi-locus combination), gene level (RecA, RecB, HI_A, HI_B, i.e. gene recessiveness and haploinsufficiency probabilities) and gene pair level (BiolDist, i.e. a metric of biological relatedness between two genes of a pair based on protein-protein interaction information). For a more detailed explanation of the features, see *SI Appendix*, Table S2. (C) After the filtering process, the 1KGP data set contained billions of bi-locus combinations compared to the DIDAv1 set, which contains 200 bilocus combinations. To solve this class imbalance problem, 500 random 1KGP samples, each containing 200 bi-locus combinations, were extracted using two types of stratification: each sample contained an equal amount (40) of bi-locus combinations from individuals of each continent, as well as an equal distribution of degrees of separation between the genes of each pair, following the degrees of separation distribution of DIDAv1. Each 1KGP sample was used against the complete DIDAv1 set to train an individual classifier that gives a class probability for each bi-locus combination. Based on a majority vote among the individual classifiers, the output of VarCoPP for each tested bi-locus combination is the final class (“neutral” or “disease-causing”), a support score (i.e. the percentage of the classifiers agreeing about the pathogenic class, SS) and a classification score (i.e. the median probability among the individual predictors that the bilocus combination is pathogenic, CS). (D) To validate VarCoPP on new disease-causing data we collected 23 bi-locus combinations from independent scientific papers, which included gene pairs not used during the training phase. To perform a confidence testing, we extracted three different random sets of either 100, 1000 and 10000 bi-locus combinations from the 1KGP set, which included gene pairs not used during the training phase of VarCoPP. By exploring the number of False Positives (FPs) predicted with these sets, we defined 95% and 99% confidence zones that provide the minimum SS and CS boundaries above of which a bi-locus combination has 5% or 1% probability of being FP, respectively.

Without going into the technical details (see Materials and Methods), it is important to mention that VarCoPP is an ensemble predictor(47), meaning that it is composed of a large number - 500 - of individual predictors that each try to solve the same task. The individual decisions of the predictors are combined via a majority vote to define the final class: if 50% or more of the predictors agree that a bi-locus combination is disease-causing, then the “pathogenic” class label will be assigned to that combination (Fig. 3C). Our results show that VarCoPP performs very well, achieving a True Positive (TP) rate of 0.88 and False Positive (FP) rate of 0.11 (*SI Appendix*, Fig. S1), meaning that 88% of the disease-causing combinations of DIDAv1 are correctly identified with 11% wrongful assignments of the disease-causing label in non-relevant combinations. The Matthews Correlation Coefficient (MCC), a more robust measure for the predictive quality of binary classifications that takes into account the correlation between observed and predicted results, achieves 74% confirming that the method is highly accurate (*SI Appendix*, Table S4). It is important to also note that these results were obtained using a stratified form of cross-validation on the training data (see Fig. 3C, Materials and Methods), meaning that considerable efforts were made to avoid bias and overfitting in the construction and evaluation of the predictor.

For each variant combination given as input, VarCoPP generates a final majority class label (“pathogenic” or “neutral”) and two prediction scores: a classification score (CS), i.e. the median probability of the variant combination being pathogenic among all pathogenic probabilities provided by each individual predictor of the ensemble, as well as a support score (SS), i.e. the percentage of individual predictors in the ensemble agreeing about the pathogenic label (see Fig. 3C and Materials and Methods for a detailed explanation of those scores). The higher the CS and SS, the more confident the predictor is about the classification of a bi-locus combination as pathogenic. To better split the neutral and disease-causing combinations, the CS threshold for pathogenic combinations was optimized to 0.489 (see Materials and Methods). Consequently, as the predictor is based on a majority vote, a bi-locus variant combination is predicted to be pathogenic when it has SS>50 and CS>0.489 (Fig. 4A). If we plot the predictions of the bi-locus combinations of DIDAv1 based on these two evaluation scores (CS on the x-axis and SS in the y-axis), we see that they are distributed in an S-shaped curve (Fig. 4B). The vast majority of the DIDA data (88%) cluster with high confidence in the right part of the S-shaped curve.

**Fig. 4.**
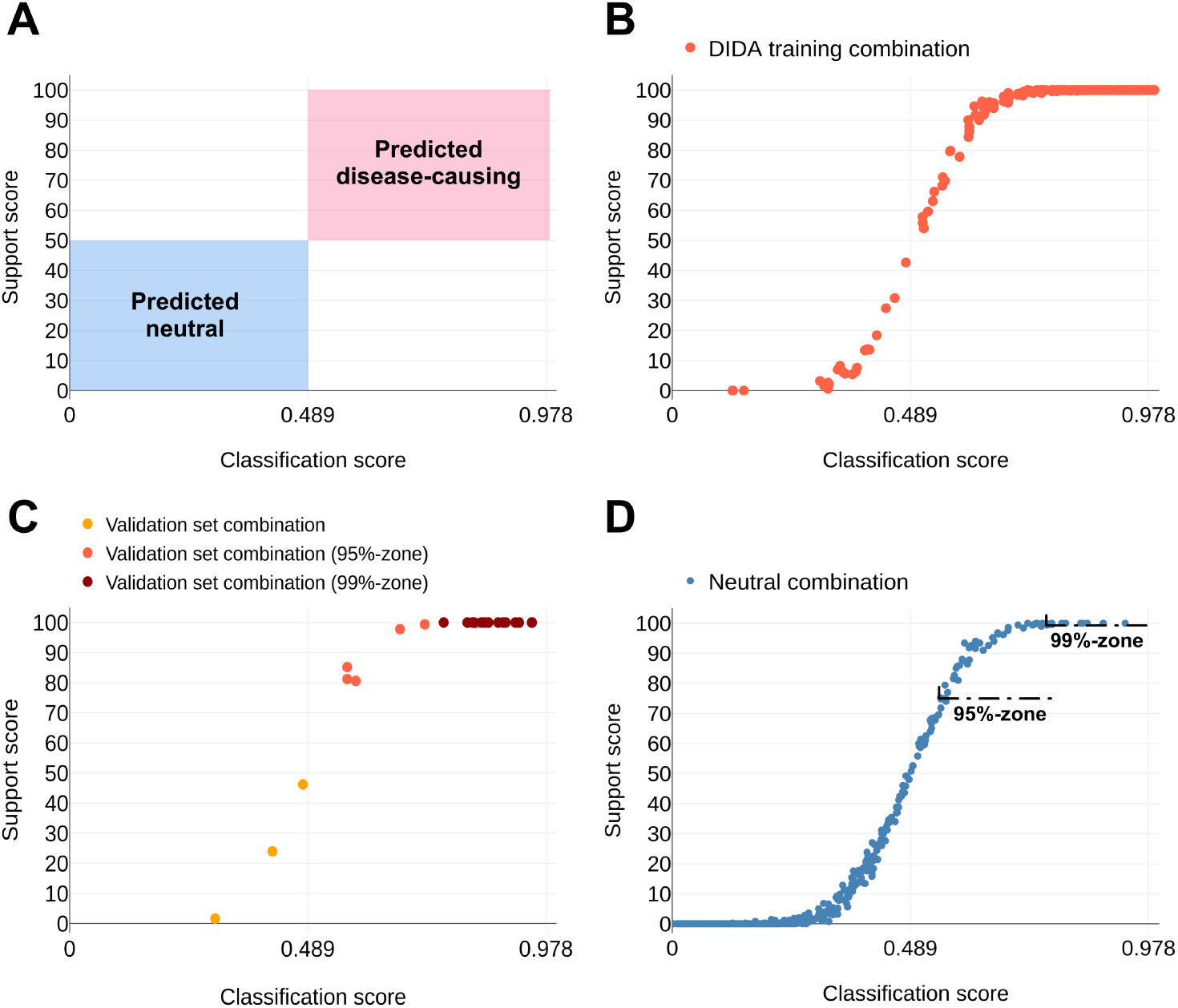
Distribution of the predictions of DIDA and those of the independent test bi-locus combinations based on the classification score (CS) on the x-axis and the support score (SS) on the y-axis. (A). A SS>50 and CS>0.489 was required to label a bi-locus combination as disease-causing. The red box represents the area where a bi-locus combination is predicted as disease-causing, while the blue box represents the area where a bi-locus combination is predicted as neutral. (B) Distribution of disease-causing bi-locus combinations of DIDAv1 during a cross-validation procedure. (C) Distribution of the 23 disease-causing bi-locus combinations of the validation set. (D) Distribution of the 1000 neutral test set combinations. The 95% confidence zone has a minimal boundary of CS=0.55 and SS=75 and contains combinations with 5% probability of being false positives (FPs), while the 99% confidence zone has a minimal boundary of CS=0.74 and SS=100 and contains combinations with 1% probability of being FPs.

### Validation on independent disease-causing data confirms VarCoPP’s predictive success

As the evaluation on independent data provides the best insight into the quality of a predictive method, we validated VarCoPP on a set of 23 new bi-locus disease-causing variant combinations, which were gathered from research articles published after the creation of DIDAv1 (Fig. 3D and *SI Appendix*, Table S5, and Dataset S1). These independent bi-locus variant combinations contained unexplored gene pairs associated with evidence for 10 diseases, which were not previously reported in DIDA, such as Alport syndrome(48) (OMIM: 301050, 203780, 104200), holoprosencephaly(49) (HPE, OMIM: 236100) and Leber Congenital Amaurosis(50) (LCA, OMIM: 204000).

VarCoPP remains very successful on these new data (Fig. 4C): The vast majority of the new bi-locus combinations (20 out of 23) are correctly labelled as pathogenic, with a high confidence (SS>80). Three bi-locus combinations, one leading to CANDLE syndrome(51), and two leading to Alport syndrome(48), were wrongfully predicted as neutral, with support of SS=46.2, 24 and 1.6 respectively. The gene pairs involved seem to be relevant for the studied disease and the genes of the pairs were closely biologically related, indicating that their protein products are most likely directly interacting. However, low CADD variant scores, a single-variant pathogenicity metric(12), as well as some missing gene recessiveness and haploinsufficiency values are most likely the reason why those combinations were misclassified (*SI Appendix*, Text S2). When this missing data becomes available or annotations are improved, VarCoPP might also classify these three cases correctly.

### Statistical confidence zones make it easy to detect the most relevant combinations

It can be expected that even after a standard variant filtering procedure, the number of neutral variant combinations (i.e. True Negatives - TNs) in an individual’s exome will vastly outnumber the number of the real disease-causing ones (i.e. True Positives - TPs). It is therefore highly relevant to estimate how likely it is that a variant combination predicted as pathogenic by VarCoPP is actually a False Positive (FP).

To examine this FP probability, we randomly collected neutral variant combinations from 1KGP individuals, consisting exclusively of gene pairs unknown to VarCoPP, and calculated their prediction scores, i.e. their CS and SS (Fig. 3D). We analyzed three different sets of such random combinations; sets of either 100, 1000 (Fig. 4D) and 10000 combinations, in order to also examine whether the percentage of FPs changes relative to the sample size (see *SI Appendix*, Datasets S2, S3 and S4 respectively).

We observed that, on average, 93% of the combinations are correctly identified as neutral, and of which 72% have a confirmative SS equal to 0, meaning that no predictor in the ensemble classified them as disease-causing (*SI Appendix*, Table S6). The overall fraction of FP combinations that are predicted as disease-causing fluctuates around 7 - 8%. This percentage remains stable even if the sample size changes. Therefore, in general, there is only a 7% chance that a bi-locus variant combination is wrongfully predicted to be disease causing.

Using this insight, it is possible to define stringent confidence zones for the predictions, delimited by specific CS and SS scores, which denote the probability that a bi-locus variant combination is a TP. We define in this manner a 95% confidence zone, containing all predicted variant combinations that have at least CS≥0.55 and SS≥75. Combinations belonging to this zone have at least 95% probability to be a TP disease-causing variant combination. Similarly, we define a 99% confidence zone, which requires at least CS≥0.74 and SS=100, containing all predicted combinations that have a 99% or higher probability of being a TP (*SI Appendix*, Table S6). These confidence zones are useful as the focus can fall directly on the bi-locus variant combinations belonging to one of these two zones and therefore have higher confidence of being relevant. Underlining again the quality of VarCoPP, one can observe that all 20 correctly classified elements in the independent validation set discussed in the previous section belong at least to the 95%-zone, with 15 of those even present in the 99%-zone (Fig. 4C).

Although these confidence zones provide a guarantee on the probability of a variant combination being a TP, the absolute number of combinations falling in those zones increases with the number of variant combinations to be tested. This is also the case when testing single variants with monogenic pathogenicity predictors. A consequence of this observation is that the precision (i.e. the fraction of real disease-causing combinations - TPs - detected among those that were predicted to be disease causing - TPs and FPs) and recall (i.e. the fraction of the real disease-causing combinations predicted correctly as pathogenic over all real disease-causing combinations present in the data set) will be affected: the smaller the fraction of real disease-causing combinations among all tested combinations, the smaller the precision and the bigger the difficulty to recall them all (see *SI Appendix*, Text S3, Fig. S2 and Table S7). As a consequence, it is best to filter down the number of variants and genes as much as possible before testing them for pathogenicity with VarCoPP. Another possibility would be to apply post-VarCoPP FP reducing strategies, as for example, using trio data, to avoid considering further irrelevant combinations already present in an unaffected parent.

### Confidence zones are relevant for the clinical analysis of disease-specific gene panels

With the previously defined 95% and 99% confidence zones and additional filtering steps we can restrict our analysis to the most relevant pathogenic bi-locus variant combinations within full exomes. Yet as there may be still a large absolute number of combinations to consider, one can further reduce the number of combinations by zooming in on those combinations that occur in a subset of genes related to the disease of interest, i.e. to restrict the analysis to well-defined gene panels. However, even by shifting to a gene panel, the current predictive quality of VarCoPP might be altered due to the specific properties of the genes included in that panel.

First, we assessed the expected absolute number of FP combinations in the 95% and 99% confidence zones for different sizes of randomly generated gene panels (ranging from 10 to 300). This analysis provides insight into the number of FP combinations present in each confidence zone that we can expect for a random gene panel of a given size consisted only with neutral variants. That is essential as geneticists do not want to be confronted with a large amount of FPs in these zones given the time and costs associated with analyzing and/or testing them. On the other hand, knowing how many FPs to expect in the confidences zones relative to the size of the gene panel provides a baseline that could be used to quantify differences between healthy patients and those having a specific disease phenotype: if the number of predicted variant combinations present in the confidence zones for a gene panel of a particular size exceeds significantly what is expected for random neutral combinations, then there may be important genetic information in the predicted results that merits future exploration.

The results for random gene panels of different sizes (10, 30, 100 and 300 genes) that contain neutral variant combinations from 1KGP individuals (see Materials and Methods for the details), are shown in Table 1. One can first observe that the percentage of FPs does not fluctuate significantly among the random gene panels, something also observed for the random neutral validation data described before (*SI Appendix*, Table S6). There is only a slight increase in the percentage of FPs in the 95%-zone for the 100 and 300 random gene panels. The strict 99%-zone appears to be more consistent, as for all random gene panels it contains on average less than 1% of neutral bi-locus combinations per individual. The absolute number of FP combinations increases, as expected, with the size of the gene panel; out of the 1312 variant combinations generated on average for a panel of 300 genes per individual, approximately 12 (0.9%) may end up in the 99% confidence zone. Additional evaluations of those cases using knowledge about the disease phenotype or molecular functionalities will most likely further reduce these numbers to acceptable sets of combinations to evaluate clinically or test experimentally.

**Table 1.**
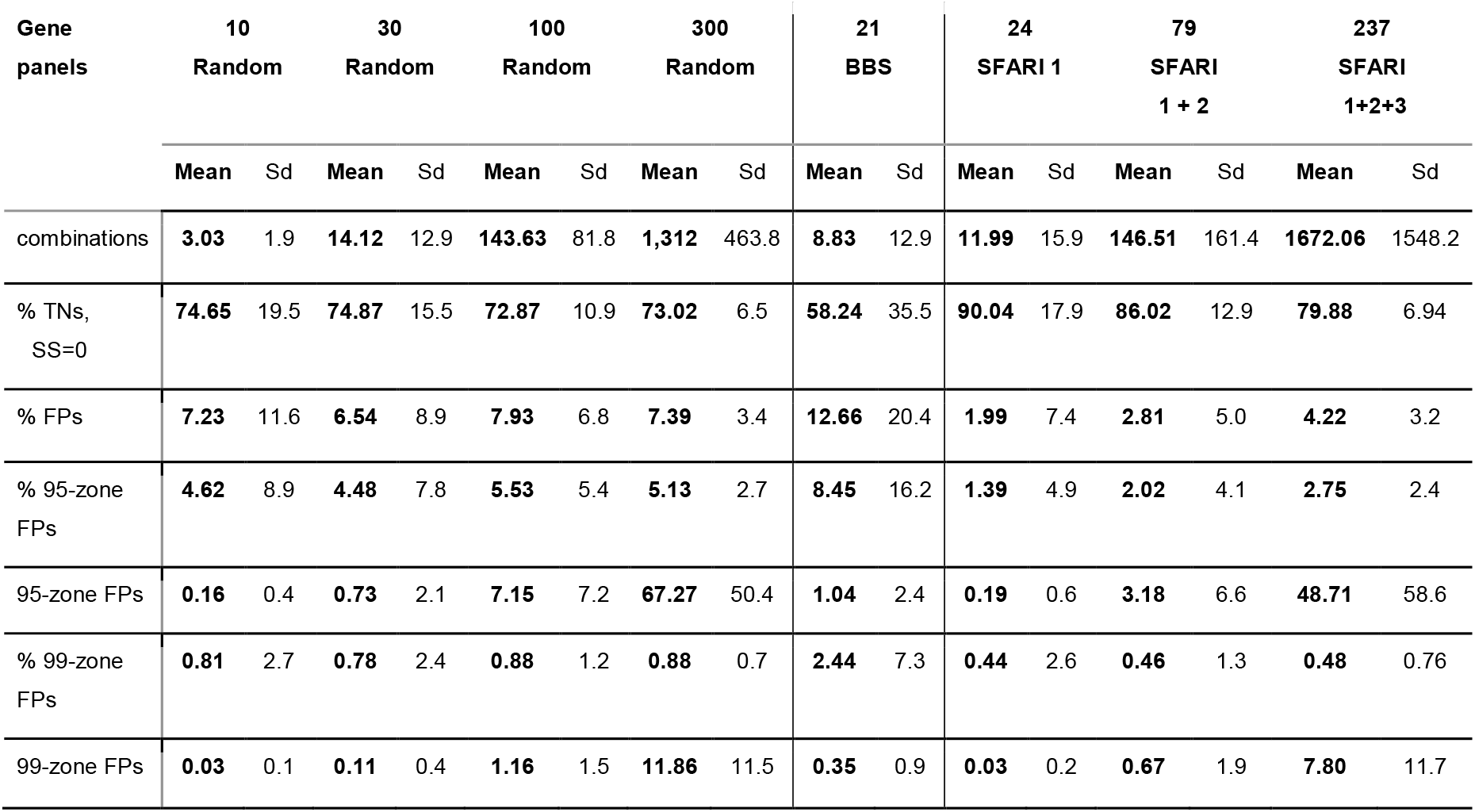
Performance of VarCoPP on independent 10, 30, 100 and 300 random gene panels and on specific gene panels for BBS and autism genes (SFARI category 1 - high confidence, category 2 - strong candidate and category 3 - suggestive evidence), iterated 100 times on 100 random 1KGP individuals (Sd = standard deviation, TNs = True Negatives, FPs = False Positives).

Second, as known disease gene panels have more detrimental properties than randomly selected ones, given that they are known to be associated with a disease, it is important to see how these statistics change for more “pathological” gene panels. We decided here to evaluate, on one hand, a gene panel for a disease known to be caused by bi-locus variants, i.e. Bardet-Biedl syndrome (BBS), and, on the other hand, gene panels for a mono-to-polygenic disease, i.e. autism, using SFARI top gene categories, applied again on neutral combinations of 1KGP individuals (see Materials and Methods). Whereas the first BBS set is expected to generate higher percentages of FPs as most of the genes are present in DIDAv1, we expected to see a reduction in FPs in the latter panels.

As can be observed in Table 1, VarCoPP appears to predict more FPs for the BBS gene panel compared to a gene panel of random genes with similar size. The BBS panel contains highly recessive genes with low haploinsufficiency probabilities (0.19 on average) and whose neutral 1KGP variants have relatively higher CADD scores compared to random genes. However, the defined confidence zones are still clinically relevant as VarCoPP guarantees that on average less than 1 variant combination will be predicted as pathogenic and be present in the strict 99-zone. As a consequence, almost any bi-locus combination present in the 99-zone should be clinically relevant. Such assertion could be tested in the future on new cohorts of BBS demonstrated bi-locus patients.

The gene panels of autism, although larger in size reveal lower FP fractions, as expected. This result is most likely due to the observation that genes in those panels have high haploinsufficiency probabilities (0.40-0.49 on average among the different panels), while the 1KGP variants present in those genes generally have lower CADD scores than average. Hence, the 95- and 99-zones stay quite devoid of false predictions. Together these results show that VarCoPP can be very precise, making it a relevant tool for discovery and diagnosis.

### The synergy of different biological features determines the pathogenicity

VarCoPP combines a number of molecular characteristics at the variant, gene and gene pair level in order to identify which variant combinations are potentially disease-causing. By analyzing how each feature influences the predictions independently, we can gain an idea about their relative importance for the full predictor. Through a feature selection procedure, we determined that a subset of 11 biological features out of the original 21 (see Fig. 3 and *SI Appendix*, Text S4, Fig. S3 and Table S2) is sufficient for making high quality predictions, while at the same time reducing the chance of overfitting.

For each of these 11 features we calculated a Gini importance score(52), which quantifies the importance of a feature proportionally to the number of samples it can successfully differentiate. Figure 5 shows that the CADD score of the first variant allele of gene A (CADD1) and that of the first variant allele of gene B (CADD3), along with the gene recessiveness probabilities (RecA, RecB) are the most important features for separating the two bi-locus combination classes. Their capacity to differentiate between pathogenic and neutral combinations becomes clear by comparing their value distributions between the two sets (*SI Appendix*, Fig. S4).

**Fig. 5.**
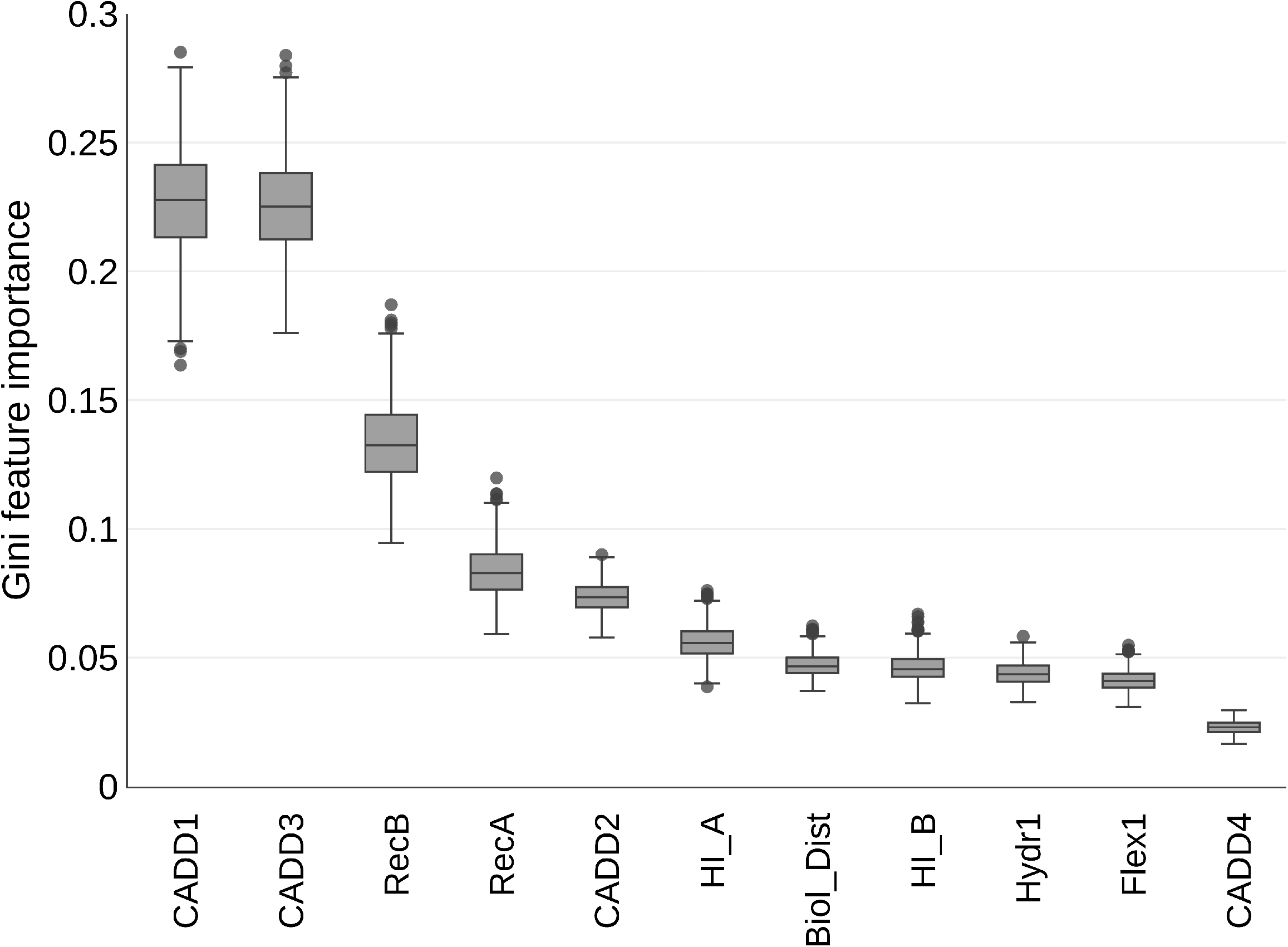
Boxplot of the Gini importance for each feature among all 500 individual predictors of VarCoPP using the training DIDA and 1KGP data.

Although the CADD pathogenicity of variants is important for VarCoPP to classify a variant combination, using CADD1 and CADD3 alone (the CADD scores of the most pathogenic variant alleles of each gene inside a combination) is not sufficient to achieve satisfactory results (see run with 2 features in *SI Appendix*, Fig. S3). By adding information of the genes’ recessiveness (see run with 4 features in *SI Appendix*, Fig. S3) we see an improvement in classification, but it is the addition of the complete biological information (i.e. the 11 selected features) that provides the best performance. Therefore, it is the synergy of all features that contributes to the correct classification of a bi-locus combination, underlining the necessity of developing a tool like VarCoPP compared to solely using combinations based on single-variant pathogenicity information.

### From black-box to white-box predictions that also explain classification decisions

Since it is the synergy between the features that determines whether a particular variant combination in a pair of genes is pathogenic or not, their joint impact on the prediction process should provide an even better understanding of how VarCoPP makes its decisions. Understanding this decision process transforms VarCoPP from a black-box into a white-box predictor, an issue that is becoming more and more important as these artificial decision-makers may have important impact on patients and people in general.

Using a method that follows the decision steps for each new bi-locus combination in each individual predictor inside VarCoPP (see Materials and Methods), we can show the preference of each feature for either the neutral or the disease-causing class in function of the influence of the other features. That preference or decision gradient can be either positive or negative, depending on whether the feature pushes the decision to the pathogenic or neutral class, respectively. For example, in DIDAv1, most disease-causing combinations are between genes that correspond to proteins that are directly or indirectly (i.e. separated by one intermediate protein) interacting. Thus, if the biological distance feature (BiolDist) between the two genes of a variant combination is rather low, meaning that the genes are very close in the protein-protein interaction network, the decision gradient for the biological distance feature will be positive, driving the prediction towards the pathogenic class. Performing that preference analysis for each feature and individual predictor inside VarCoPP gives a distribution of decision gradient values for that feature for every bi-locus variant combination. The simplest way to visualize these values per feature is by using box-plots that reveal both the median and variance among the individual predictors in VarCoPP (as can be seen in Fig. 6). Positive values that vote for the pathogenic class are depicted in red, while negative values voting for the neutral class are depicted in blue.

**Fig. 6.**
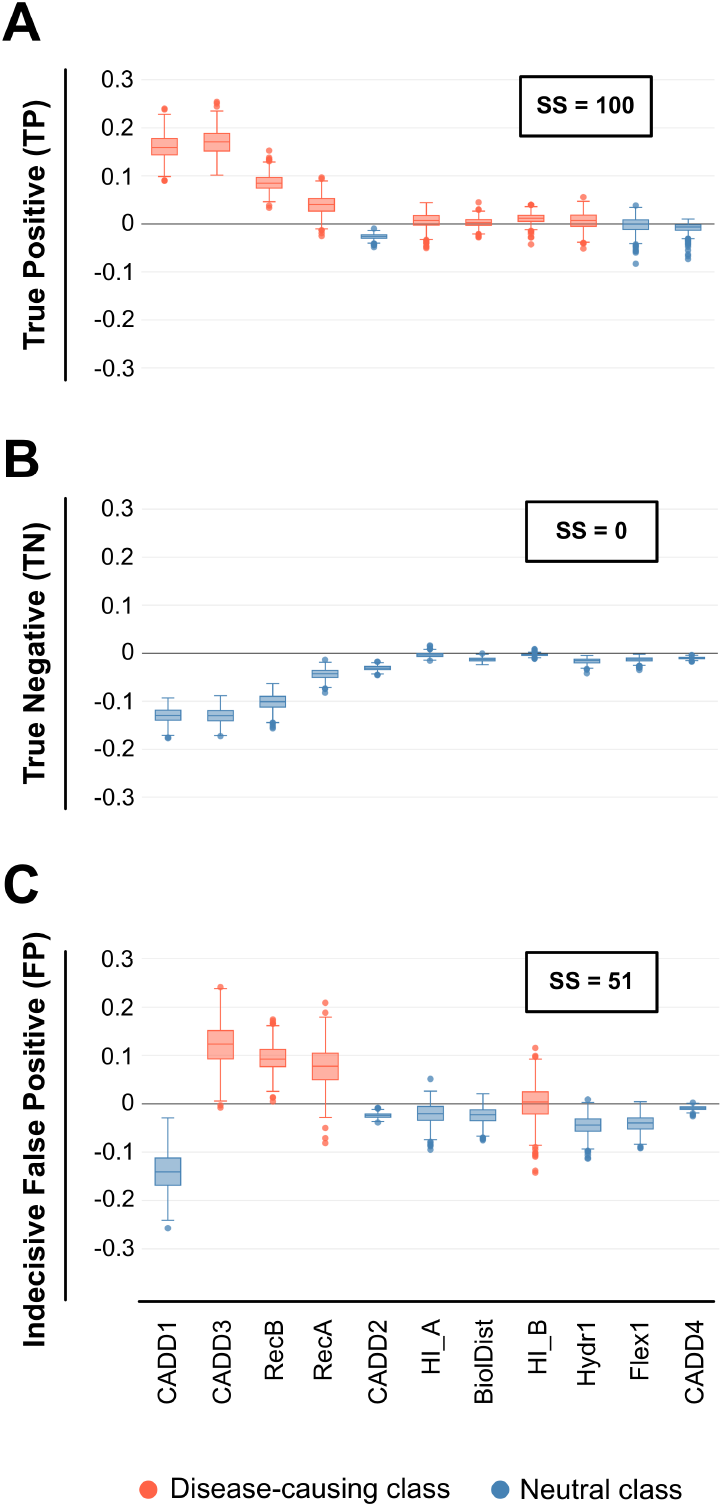
Decision profile (DP) boxplots that show the class preference (or decision) gradients of each feature used for the classification of test bi-locus combinations. Features whose median decision gradient values among all classifiers of VarCoPP fall above 0 on the y-axis are in favor of the disease-causing class (red color), whereas features whose median decision gradient values fall below 0 on the y-axis are in favor of the neutral class (blue color). (A) A DP boxplot for a true positive bi-locus combination with support score (SS)=100 (*SI Appendix*, Dataset S1, testpos_21), where the vast majority of features have a median decision value above 0 (B) A DP boxplot for a true negative bi-locus combination with SS=0 (*SI Appendix*, Dataset S3, testneg_769), where all features have a median decision value below 0 agreeing for the neutral class. (C) An example of an indecisive DP boxplot for a neutral bi-locus combination of the set of 1000 test neutral combinations, which was predicted as disease-causing with of SS=51 (*SI Appendix*, Dataset S3, testneg_358).

The higher the confidence of a prediction, the more clearly features show preference for a particular class. As can be seen in Fig. 6, there is a clear positive or negative preference among the features for cases where there is either full support for the disease-causing (Fig. 6A, SS=100) or neutral (Fig. 6B, SS=0) class for a bi-locus combination. However, in cases where the prediction is ambiguous, like for example in cases where the average support from the individual predictors in VarCoPP is close to the threshold (SS ~ 50%), we observe that such a clear consensus among the features is missing (Fig. 6C).

These visualizations provide a good indication as to why we reach a disease-causing or neutral prediction for a bi-locus variant combination. By examining the actual values of the features that most strongly influence the decision process for a particular combination (i.e. are furthest away from zero, see Fig. 6), we can obtain insight on why this combination gets a class prediction and people can assess their agreement with that prediction. For example, if we see that the CADD feature of a variant drives significantly the prediction towards the pathogenic class, we can most probably expect that the CADD feature value of that variant is relatively high.

## DISCUSSION

This work demonstrates that sufficient genetic knowledge is available to produce pathogenicity predictors capable of differentiating between pathogenic and neutral bi-locus variant combinations. We presented here VarCoPP, a clinically competent predictive tool, which reveals to be precise and sensitive, both in cross validation settings (87% correct predictions) and when tested on new independent data. The performance will further increase by improving the quality of the genetic annotations and more training data becomes available.

VarCoPP provides robust 95% and 99% confidence labels, which constitute an objective assessment of the relevance of newly identified pathogenic bi-locus variant combinations. These zones are important as a form of primary filtering and evaluation of the predictions, while further statistical and biological verification can be performed for those 95% and 99% labeled variant combinations. Such an approach boosts the clinical relevance of VarCoPP, limiting the search space produced by all variant combinations in a gene panel or exome to the most relevant ones and, as a consequence, reducing the required time needed to further explore these relevant results.

Moreover, our method has been designed to produce “white-box” predictions by providing insights into the importance of the biological features in distinguishing disease-causing combinations from neutral ones (Fig. 5). Furthermore, it can provide objective explanations on the class decision made by the predictor for each new bi-locus combination that is being tested (Fig. 6). While the former provides a way to assess the relevance of novel features in further developments of VarCoPP, the latter allows users to assess the relevance of the prediction using their genetic and biological expertise and to capture reasoning differences for different bi-locus instances. Providing such decision transparency for automated systems is highly important given the effect that predictions may have on individuals and society.

Although we can now start to analyze combinations in patient exomes, it is important to keep in mind that the magnitude of the search space increases dramatically when moving to full exome analysis. Although there is only 1% chance of observing a FP in the 99% zone, the absolute number of FP combinations will exponentially increase, a classic problem that is unfortunately encountered in most types of bioinformatics predictors when tested at the exome level. Additional pre- or post-filtering steps to reduce these absolute numbers are thus required, which can be done, for instance, by adding knowledge about the disease or comparing the predictions to genetic information obtained for the parents in trio studies. In line with the former, the study can be limited to gene panels known to be associated with the disease or belonging to the relevant pathways. We demonstrated that such a focus will indeed help in limiting the number of non-relevant bi-locus combinations: using a panel of 150 genes produces potentially 1 non-relevant combination in the 99% confidence zone, that usually corresponds to a percentage of less than 1%, confirming the clinical relevance of our method. Furthermore, rare-diseases recessive gene panels (like BBS) may produce a bit more FPs, in contrast to known haploinsufficient gene panels (like those of neurodevelopmental disorders). Clinical users of VarCoPP should be aware of this issue in the analysis of their target disease.

The results furthermore show that especially the CADD scores of the first variant allele of each gene - an expected observation as we order the variant alleles inside each gene based on pathogenicity - but also the gene recessiveness probabilities seem to be the main drivers of predictions. Although these features independently show great importance, it is the combination of all 11 selected features, including those with a lower effect, that leads to the highest classification accuracy. These results make VarCoPP a clinically important tool that is more informative and accurate than simply selecting potentially relevant variant combinations based solely on monogenic variant pathogenicity scores, such as CADD.

Further expansions of VarCoPP into the oligogenic realm should consider that the variant filtering criteria that were shown to be important for the method, differ from the ‘strict’ criteria that are commonly used to identify pathogenic variants in rare mendelian diseases, i.e. rare exonic variants with a strong monogenic effect. Although the majority of positive cases in DIDA has a MAF of less or equal to 3%, we observed that some variants involved in rare oligogenic diseases can reach for instance a MAF of up to 18%(53). As these are present but constitute exceptions in the current data set, at the moment we restricted ourselves to MAF of 3% for the creation of the neutral data set. Nonetheless, this threshold can be further relaxed in the future as more data on pathogenic bi-locus combinations becomes available.

Similarly, while it is widely presumed that genes involved in the same disease can belong to the same molecular pathway or biological process, this does not necessarily apply to all cases. It is shown in DIDAv1 that for some gene pairs, such as the ANOS1-PROKR2 pair found in many studies associated with Kallman syndrome(45, 54–56), no interaction or co-expression information is known yet, indicating that potentially more complex pathways and cellular mechanisms may be involved to cause disease. Nonetheless, the pairs of genes in the neutral data used by VarCoPP were filtered in such a way that they had the same biological distance distribution as was observed in DIDAv1. As a consequence, this feature has a less important role in the decision process. It remains to be seen whether this stratification should not be relaxed when moving into the realm of oligogenic disease cases, as subsets of genes involved in different pathways may be responsible for the observed phenotype. Yet, relaxing this biological distance normalization will lead to a less “clever” predictor with a higher FP rate, as it would provide an obvious way to learn separating known pathogenic from random neutral bi-locus combinations (*SI Appendix*, Fig. S5).

VarCoPP is a bi-locus variant combination pathogenicity predictor that is trained using combinations involved in known oligogenic diseases. As our predictor is not phenotypically-driven, it could also be used to predict bi-locus combinations involved in cases of dual molecular diagnosis, i.e. cases where several independent monogenic diseases are present in an individual due to segregation of monogenic variants in two unrelated loci. The recent work of Posey *et al*.(21) provides a collection of such dual molecular diagnoses cases. An analysis of 76 cases in that paper revealed that VarCoPP predicted 67 (88%) correctly (see *SI Appendix*, Fig. S6 and Dataset S5). These results are again very promising, especially since dual diagnosis cases are almost completely missing from DIDAv1, which was used to train the predictor. Nonetheless, such cases appear to consist of strong monogenic variants and genes whose nature and properties are different compared to those causing or modulating the diseases contained in DIDAv1. Within the context of another study, expanding on Gazzo *et al*.(39), it is observed that dual diagnosis instances are indeed separated from the other types of bi-locus diseases. Although further developments of VarCoPP should incorporate these cases for training, a distinction should be further made between dual diagnosis instances with distinct and overlapping phenotypes. Especially the latter appear to be relevant for a predictor that aims to find synergies between variants, which is the long-term ambition of VarCoPP.

In conclusion, VarCoPP reveals that the first steps to multi-variant pathogenicity predictions can be taken. Our method shows great predictive ability during cross-validation and using independent validation sets, which may be further improved with the advent of new data and the inclusion of additional biological information. The provision of statistical evaluations, as well as white-box explanations on the obtained results establish VarCoPP as a pioneering clinical tool for the detection of disease-causing variants implicated in more complex genetic patterns. By scoring bi-locus combinations and gene pairs, gene triplets or quadruplets may be identified in exome or gene panel data as causative genetic models for a particular disease, paving the path for the detection of multi-locus signatures derived with machine learning approaches. VarCoPP therefore provides an important leap forward, allowing for more fine-grained pathogenic predictions.

## MATERIALS AND METHODS

An illustrated summary of the Materials and Methods used in this study is presented in Fig. 3. Additional details on each Materials and Methods subsection can be found in the *SI Appendix*, Text S4.

### Data filtering and annotation

We filtered the variants and genes between DIDAv1 and 1KGP so that both sets contained comparable information (Fig. 3A), using exonic and splicing SNPs as well as indels of MAF equal or less than 3%. Individuals in 1KGP that carried disease-causing bi-locus combinations, as well as the corresponding combinations in DIDAv1, were removed (Fig. 2 and *SI Appendix*, Table S1). We annotated both sets based on information at the variant, gene and gene pair level, leading initially to 21 features per entry. After a feature selection procedure this set was reduced to 11 features (Fig. 3B and *SI Appendix*, Text S4, Table S2 and Table S3 for an overview and explanation of the features). Variants and genes inside each bi-locus combination were ordered in both data sets, so that gene A and the first variant allele of each gene in a bi-locus combination were the most pathological ones according to the Gene Damage Index (GDI) score(57) and CADD score(12), respectively (Fig. 3B).

### Stratification of the 1KGP data and training

To train VarCoPP we created 500 balanced sets (Fig. 3C), each consisting of 200 1KGP bi-locus combinations of randomly chosen gene pairs and the 200 disease-causing combinations of DIDAv1. For each 1KGP subset, we included 40 individuals per continent. However, there is no significant difference in performance when the predictor is trained using 1KGP combinations only from individuals of a particular continent against DIDAv1, confirming no population bias (*SI Appendix*, Table S8). Each random control subset contained gene pairs following a degree of separation distribution equal to that of DIDAv1, based on information obtained from the Human Gene Connectome tool(58) (*SI Appendix*, Fig. S5). We used the scikit-learn version 0.18.1 implementation(59) of the RF algorithm(52) as a classifier for each of the 500 balanced sets. Each RF consisted of 100 decision trees using bootstrapping with a maximum tree depth of 10, using the square root of the features for each split. We implemented a Leave-One-Pair-Out stratified cross-validation procedure individually for each predictor(39).

### Validation of VarCoPP

We collected 23 new disease-causing bi-locus combinations derived from independent scientific papers published after the release of DIDAv1 (Fig. 3D and *SI Appendix*, Table S5 and Dataset S1). For confidence testing, we collected different sets of random 100, 1000 and 10000 neutral bi-locus combinations from the 1KGP that were unused during training (Fig. 3D and *SI Appendix*, Datasets S2, S3 and S4). For the gene panel analysis, we created random panels consisting of 10, 30, 100 and 300 genes and tested each gene panel on 100 random 1KGP individuals with 100 iterations. For BBS, we used the 21-gene list obtained from the Genome Diagnostics Nijmegen laboratory (http://www.genomediagnosticsnijmegen.nl/) and for autism, the SFARI gene panels (https://gene.sfari.org/).

### Feature selection and interpretation

We applied a recursive feature elimination procedure(60) on a balanced set with median performance among all sets leading to a performance peak with 10 features (*SI Appendix*, Fig. S3). As no variant features about the second variant allele of gene B remained, we included for interpretability reasons the CADD score of this allele (CADD4), finalizing the number of selected features to 11. To create the decision boxplots per bi-locus combination we used the *treeinterpreter* Python package (https://github.com/andosa/treeinterpreter, Ando Saabas).

VarCoPP can be accessed online at: http://varcopp.ibsquare.be/. This online tool annotates a list of given variants and scores all possible bi-locus variant combinations present in that list.

## ACKNOWLEDGMENTS

We thank all the members of the Interuniversity Institute for Bioinformatics in Brussels, especially the group of people interested in digenic and oligogenic diseases, for their comments and valuable suggestions. This work was supported by the ARC project Deciphering Oligo- and Polygenic Genetic Architecture in Brain Developmental Disorders [to A.G., N.V., C.N. and T.L.]; the European Regional Development Fund (ERDF) and the Brussels-Capital Region-Innoviris within the framework of the Operational Programme 2014–2020 through the ERDF-2020 project ICITY-RDI.BRU [27.002.53.01.4524 to S.P, A.N., S.V.D. and T.L.]; a Fonds de la Recherche Scientifique (F.R.S) - FNRS Fund for Research Training in Industry and Agriculture (FRIA) [to S.P.]; a Vrije Universiteit Brussel, PhD funding [to S.P.]; a Vrije Universiteit Brussel, Reproduction and Genetics and Regenerative Medicine (RGRG) Cluster, Reproduction and Genetics Research Group [to A.G. and S.V.D.]

AUTHOR CONTRIBUTIONS SP, AG, JA, YM, SVD, AN, GS and TL conceived and designed this study; SP, AG, CN, NV, GS and TL developed the predictor; SP, AG, CN, NV, GS and TL analyzed the results; SP, AG, NV, CN, JA, YM, SVD, AN, GS and TL wrote and approved the paper.

