## Supplementary Information for "Predicting disease-causing variant combinations"

**This PDF file includes:**

Supplementary Text S1 to S4
Figs. S1 to S6
Tables S1 to S8
References for SI reference citations

**Other supplementary materials for this manuscript include the following:**
Datasets S1 to S5

### Supplementary Information Text

#### Text S1. Supplementary information for overlapping bi-locus combinations between DIDA and the 1KGP

During an initial screening of bi-locus combinations in the 1000 Genomes Project (1KGP), we discovered at that time 7 bi-locus combinations of DIDA that were also present in 14 individuals of the 1KGP. None of the studies analysing these combinations presented a comparison with the 1KGP. The bi-locus combinations were present in parents or grandparents of 1KGP families and no relative carried extra DIDA bi-locus combinations.

We detected 2 bi-locus combinations leading to Kallman syndrome both derived from the same study(1), which were also present in 5 individuals of the 1KGP. These combinations involved the gene pairs *KISS1R* - *PROKR2* (dd202), which had already been reported to be relevant for digenicity(2), and *GNRHR* - *PROKR2* (dd203). Although *GNRHR* has been involved in digenic cases with other genes (*PROK2* and *FGFR1*) leading to congenital hypogonadotropic hypogonadism(3-5), no other indications of digenicity have been reported between this gene and *PROKR2*. The c.719G>A *GNRHR* heterozygous variant of dd203 has been described previously to be involved in mild symptoms of congenital hypogonadotropic hypogonadism(6). On the other hand the c.991G>A *PROKR2* heterozygous variant of dd203 has already been reported to be implicated in Kallman syndrome with variable phenotypes(7). In the current study, the authors compared cases with a small control cohort of 14 individuals carrying mutations of either *KAL1* and *PROKR2*, therefore did not explicitly check for bi-locus combinations in the control cohort. The bi-locus combinations presented are also not supported by familial or functional evidence.

We also detected in 2 individuals of the 1KGP a bi-locus combination (dd220) leading to maturity-onset diabetes of the young(8) that involved a novel digenic pair *NEUROD1-PDX1*. Both mutations in these genes are heterozygous, in protein-coding areas. In general, it is known that the clinical features of MODY can overlap with those of polygenic diabetes, and several genes have been associated so far with this disease. For the dd220 combination, this novel gene pair can be relevant for MODY, as both genes are required for β-cell development, growth and insulin gene expression participating in a transcription complex that is important for short-range DNA looping(9). Both heterozygous c.723C>G *NEUROD1* and c.670G>A *PDX1* mutations of that bi-locus combination had already known to be individually implicated in forms of obesity and diabetes, respectively(10,11). However, the patient that carried this bi-locus combination was not obese, but showed a serious decrease of insulin intake compared to controls. The c.670G>A *PDX1* variant can present incomplete oligogenic penetrance(11) - also suggested by its frequency in the ExAC database (0.002113) - and, thus, sometimes can be overlooked. At the same time, although studies have shown a contradictory importance of *NEUROD1* with the associated clinical phenotypes being unclear, microRNA analysis showed that silencing this gene led to loss of β-cell proliferation, through the overexpression of miR-24(12). Although the study that presented this combination included a familial analysis, they authors did not provide functional evidence. Furthermore, in the study the authors performed a genetic comparison with a small cohort of 60 control individuals with normal glucose intake, something that could maybe have led to loss of statistical power to show the presence of the bi-locus combination in the control cohort. In general, the relevance of this bi-locus combination may not be unimportant, but it should be noted that for MODY other less-known molecular mechanisms may be involved that make this bi-locus combination present in two individuals of the 1KGP.

Finally, we detected 4 novel bi-locus combinations (dd180, dd188, dd193 and dd196) leading to familial hemophagocytic lymphohistiocytosis (FHL) in 7 individuals, which were all derived from the same study(13). It is worth noticing that one 1KGP individual carried two of these combinations: dd188 and dd193. The variant combinations involve heterozygous mutations in *PRF1*, *UNC13D* and *STXBP2* genes, with three of them (dd180, dd188, dd193) sharing the variant c.272C>T at *PRF1* gene. All of these genes are involved in the lymphocyte cytotoxicity activity for cytotoxic granules and *UNC13D* along with *STXBP2* belong also in the same cytotoxic pathway(14), while *PRF1* assists at a later stage to the delivery and penetration of granzymes(15).
More specifically:

- Combination dd196 involves an intronic c.*12G>A *STXBP2* variant and a protein-coding c.2896C>T *UNC13D* variant. These genes belong in the same degranulation pathway and the authors show that patients with variants in genes of that pathway had earlier age of onset and defective degranulation, compared to those who carried variants in *PRF1* and another gene. With these findings the authors suggest a synergistic deleterious effect of the involved mutations that supports the notion of a digenic inheritance for FHL. Digenic implications of this pair have also been suggested in further murine and human studies(16,17).
- Combination dd180 involves two protein coding c.272C>T *PRF1* and c.3160A>G *UNC13D* variants. On the other hand, combinations dd188 and dd193 involve the gene pair *PRF1 - STXBP2*. Combination dd188 involves two protein-coding variants c.272C>T in *PRF1* and c.1034C>T in *STXBP2*, while dd193 involves the *PRF1* c.272C>T variant and one intronic variant c.795-4C>T in *STXBP2*. The authors present that patients with one mutation in the *PRF1* gene and another mutation at one of the genes of the granulation pathway had later age of onset compared to those carrying mutations in genes of the same cytotoxic pathway and normal degranulation, although they showed decreased perforin expression. Nevertheless, digenic variants in the pair *PRF1 - UNC13D*, but not the exact dd180 combination, have been reported in another later study(17).

As the genes involved in that study are implicated in FHL and digenicity has been suggested in further studies, we cannot conclude that the specific variant combinations that have an overlap between DIDA and 1KGP are not relevant, although in the paper there is a stronger indication of digenicity for the dd196 combination, compared to the rest. It should be noted, however, that the study did not present a familial analysis, neither further functional evidence concerning the digenic cases. Moreover, the authors did not compare their cases with controls or other variant databases for further proof of relevance, but performed a clinical analysis.

#### Text S2. Interpretation of validation cases predicted as neutral

Among the 23 independent variant combinations of the independent validation set, we predicted 3 of them as neutral (Testpos_4, Testpos_10 and Testpos_15). We provide here our interpretation of why we fail to predict them as disease-causing. In general, this can be attributed to missing values of some features for the gene recessiveness or haploinsufficiency, low CADD score values and high gene haploinsufficiency values in some cases. The annotated features for this data set can be further studied at the *SI Appendix*, Dataset S1.

1. Testpos_4, which was predicted with SS = 46.2, carrying novel inherited variants in the *PSMB4*-*PSMB9* gene pair, is involved in the development of CANDLE syndrome, an autoinflammatory disease, and it was detected in two siblings of a familial study. The authors refer that the heterozygous c.44_45insG mutation in *PSB4* causes reduced gene expression, while the second missense c.494G>A mutation in *PSMB9* affects a highly conserved protein residue. Both *PSMB4* and *PSMB9* were predicted to be haploinsufficient in our data. Recessiveness probability of *PSMB9* was not known and therefore we used a median value, something that may have affected the results, as it seems that this feature (RecB), along with the CADD score of the *PSMB9* variant allele (CADD3) that is low in this case, are important for classification. Therefore, we see that this unknown value along with the low value of CADD3 can have an impact on the predictions.
2. Testpos_10, which was predicted with SS = 24, contains variants in the *COL4A4*-*COL4A3* gene pair known in general to be implicated in Alport syndrome, the first one being a heterozygous in-frame c.1293_1310del mutation and the latter being a splicing c.1504+1G>A variant that is predicted to cause the loss of the 5' splice site. The patient carrying this bi-locus combination showed symptoms for heamaturia and proteinuria, but no hearing loss or ocular lessions. In the paper, the authors show that none of the monogenic predictors (SIFT, MutationTaster, Polyphen2) was able to provide a pathogenicity score for these variants. Looking at the CADD values, we saw that these were actually low (2.36 and 2.84). These low scores most probably guide the prediction towards the neutral class.
3. Testpos_15, which was predicted with SS = 1.6, originated from a female patient who showed symptoms of intermittent heamaturia and proteinuria and not hearing loss or ocular lessions, therefore not severe symptoms of the disease. It contains three heterozygous variants, one in the *COL4A5* gene and two in the *COL4A4* gene, both known to be involved in Alport syndrome. Males carrying mutations in this pair show more severe symptoms of the disease, as the *COL4A5* gene lies in the X-chromosome. In this case, although the CADD score of the *COL4A5* variant is relatively high, the very low CADD scores for the *COL4A4* variants (1.70 and -0.22) and the high *COL4A4* haploinsufficiency probability value (0.84) seem to guide the prediction of this bi-locus combination towards the neutral class. We would like to note that in our method, we represent a hemizygous X-linked variant in males similarly to a homozygous variant, to depict the fact that there is no wild-type allele present to compensate in certain cases for the gene and protein function, and thus, this variant, and consequently the bi-locus combination, can be predicted to have a stronger impact in males than females.

#### Text S3. Interpretation of PR curves for confidence zones

As the amount of neutral bi-locus combinations tested increases, the precision drops (*SI Appendix*, Fig. S2*A*), as expected. Clearly, the more combinations need to be checked the more the absolute number of elements in the two confidence zones will increase. Figure S1*B* in *SI Appendix* shows the precision and recall of the 95%-zone and the 99%-zone when VarCoPP is tested on the elements of the positive validation set together with the elements of a collection of either 100, 1000 and 10000 neutral combinations that fall into these zones. As before, increasing the number of neutral cases decreases the precision: the precision of finding all 20 TPs in the 95%-zone drops from ~80% to ~30% when increasing the amount of bi-locus combinations from 100 to 1000. The more stringent 99%-zone improves precision and recall significantly (*SI Appendix,* Fig. S2*B*), yet at the cost of missing some of the real culprits (*SI Appendix*, Table S7).

#### Text S4. Supplementary Materials and Methods

- **Data collection, filtering and annotation**

We used the first version of the Digenic Diseases Database (DIDAv1) as the disease-causing data set and collected information on the bi-locus combinations, genes and variants. 213 bi-locus combinations were present in the database and included single nucleotide variations (SNVs) and small insertions and deletions (indels). These combinations contributed to 44 different diseases. To create the control bi-locus combinations, we used the variant data of the 1000 Genomes Project (1KGP) of Phase 3. The 1KGP contains a broad representation of the human genetic variation, including variants of 2,504 healthy individuals from 26 different populations. For computational reasons, we selected a random 25% subset of the human proteome from the 1KGP, using the list of the curated human proteome from Uniprot(16) and filtered the variants using Highlander (http://sites.uclouvain.be/highlander/). This filtering enabled us to limit the amount of gene pairs and bi-locus combinations present in the control set, in order to be similar to the type of disease-causing combinations composing DIDA.

The filtering process took place in such a way that both sets would contain comparable variants and genes. We used the Ensembl BioMart tool(17) for the Grch37.p13 version of the human genome to obtain information about exon positions and 1KGP Minor Allele Frequency (MAF). We included variants of MAF<=3%, as this variant frequency threshold represents the vast majority of variants in DIDAv1 and in this way we could limit potential noise of the abundance of more common variants from the control dataset. We included exonic SNVs and small indels, as well as intronic variants with a maximum distance of 13 nucleotides (nt) from the exon edge. We also collected synonymous variants close to splicing sites in a maximum distance of 6 nt from the exon boundary, according to the synonymous variants present in DIDAv1. At the gene level, we selected only protein-coding genes from 1KGP, as this type of genes was present in DIDAv1. To further ensure the presence of true protein-coding genes in the control set, we further filtered the genes based on the protein-coding genes of the consensus coding sequence (CCDS) project(18). Finally, we removed from the 1KGP data set 14 individuals who carried disease-causing bi-locus combinations (Fig. 2, and *SI Appendix*, Table S1), as well as the 7 corresponding bi-locus combinations. After the filtering process, 200 disease-causing bi-locus DIDAv1 combinations remained, whereas around 8,000,000 unique gene pairs were present in the 1KGP control set leading to billions of possible combinations to choose.

We annotated both sets based on information at the variant, gene and gene pair level. A summary of the features and a brief explanation is displayed in *SI Appendix*, Table S2 and shown schematically in Fig. 3B. We used the combined annotation dependent depletion (CADD) score(19) as a single-variant deleteriousness metric because it can predict the damaging effect of not only missense variants, but also splicing, nonsense variants, as well as small deletions and insertions and shows good overall performance. We implemented an in-house code to calculate the flexibility and hydrophobicity differences between the wild-type and mutated amino acids, using the flexibility variation scale of Bhaskaran & Ponnuswamy(20) and the Wimley & White whole residue hydrophobicity scale(21) respectively. We also extracted information from Pfam(22) using the Ensembl BioMart tool for the Grch37.p13 version of the human genome(17) to predict whether the variant alleles lie in a conserved protein family domain. For the gene features, we used the gene haploinsufficiency(23) and recessiveness probability(24) retrieved from dbNSFP2.8(25,26). Finally, as a gene pair feature we exploited the biological distance, a metric of biological relatedness in terms of protein-protein interactions between two genes, obtained with the script provided by the Human Gene Connectome tool(27).

- **Representation of a bi-locus variant combination**

We represented a bi-locus variant combination as a vector of categorical and numerical features (see Fig. 3B and *SI Appendix*, Table S3 for numerical representation). At the moment we considered at most two mutated alleles per gene, based on the types of variant combinations present in DIDA. Therefore, as the total number of mutated alleles of a bi-locus combination can range from two to four, depending on the variant zygosity, we encoded each variant-related feature four times (four dimensions), each one representing a different allele. We encoded gene-related features using two dimensions, one for each gene involved in the bi-locus combination. In the end, a set of vectors was created where each vector represented a bi-locus combination and each element in the vector represented a unique feature.

To ensure a fair prediction process, we defined the order of variants and genes inside each bi-locus combination (Fig. 3B) in the same way for both data sets. We used the Gene Damage Index (GDI)(28) to determine the order of the two genes in a bi-locus combination, so that gene A was the gene with the lowest GDI index and, thus, has with a higher probability to be associated with a disease. We also ordered different variant alleles inside the same gene (in cases of tri-allelic or tetra-allelic combinations) using the CADD raw score(19), so that the first variant in that gene would be the one with the highest score.

- **Stratification and sampling of the 1KGP neutral data for training**

The 1KGP neutral set contained a huge number of bi-locus combinations compared to DIDA, leading to a class imbalance problem (Fig. 3C). A recent work has shown that Random Forests (RFs)(29) can outperform other classifiers even when used as base-classifiers to create ensemble predictors, especially in cases where one of the two training sets, *i.e.* the 1KGP set in our study, is much bigger and has to be under sampled(30). Although partitioning of the entire 1KGP data set into smaller samples could offer a good solution(30,31), we were aware that this would pose other computational issues, as it would have resulted in millions of different predictors and would dramatically increase the running time of our method. Therefore, we decided to perform a bagging procedure instead and we created multiple balanced sets, each consisting of 200 1KGP bi-locus combinations of randomly chosen gene pairs and the 200 disease-causing combinations of DIDAv1. We first performed a preliminary analysis to assess the variance of the performance of our method relative to the number of neutral variants it is trained with, by creating different ensemble predictors trained on either 10, 100, 500, 1000, 1500, 2000 and 3500 balanced training sets. Although we did not observe significant performance differences, we finally decided to use 500 balanced sets as they showed small variance and they were the starting point of a performance plateau for bigger sets (*SI Appendix*, Table S4).

The bi-locus control combinations contained one to four variant alleles similarly to those present in DIDAv1. Bi-locus combinations with four non-wild type alleles in two different genes were not included in the 1KGP set, as this type of combinations was not yet present in DIDAv1. As variants from African populations are generally over-represented in the 1KGP set and consequently in our random subsets, we created an equal continent distribution among the individuals for each random 1KGP subset, in order to avoid bias towards variants specific to a particular population. However, preliminary analysis has shown that there is no significant difference in performance when the predictor is trained using 1KGP variant combinations only from individuals of a particular continent against DIDAv2 confirming that the method, being a qualitative machine-learning approach, is not subject to population bias (*SI Appendix*, Table S8). Finally, each random control subset contained gene pairs following a degrees of separation distribution (i.e. number of direct protein interacting connections between two genes) equal to that of DIDAv1, based on information obtained from the Human Gene Connectome tool(27) to again avoid obvious biological relatedness differences between the randomly chosen control pairs and the disease-causing gene pairs (*SI Appendix*, Fig. S5).

- **Training of the ensemble-based machine learning method**

We used the scikit-learn version 0.18.1 implementation(32) of the RF algorithm(29) as a base-classifier for each of the 500 balanced sets. Each RF predictor consisted of 100 decision trees using bootstrapping with a maximum tree depth of 10, while the number of features to consider when looking for the best split at each node was set to the square root of their total number. As most of the used features were numerical, except for the two-class Pfam categorical feature, we selected the default Gini importance option to assess feature importance.

To assess the overall performance between the independent RFs, we implemented a Leave-One-Pair-Out stratified cross-validation procedure individually for each predictor, similarly to the methodology presented in the work of Gazzo *et al.*(33). In this type of cross-validation, we iteratively removed all variant combinations of a particular gene pair and trained the RF predictor with the rest of the combinations. Then, we used the bi-locus combinations of the left-out pair to assess the performance. This procedure was repeated for all gene pairs inside each balanced set and to conclude statistics for the performance among the RF predictors, the results were averaged among all balanced sets.

- **Definition of the Classification Score (CS) and Support Score (SS)**

VarCoPP implements a majority vote to predict the final class ("disease-causing" or "neutral") for a bi-locus combination $\boldsymbol{x}$**,** see Equation (1).

Equation (1):

$$maj.vote\ (\boldsymbol{x}) = \arg\max_{c=N,D} \sum\nolimits_{t=1}^{T} d_{t,c}\ (\boldsymbol{x})$$

Each individual RF $t$, where $t \in \{1,\cdots,T\}$ and $T$ is the total number of RFs, gives a class decision $d_{t,c}(x)$, with class $c$ = {neutral ($N$), disease-causing ($D$)} for an instance $x$, see Equation (2). The majority vote function returns the class that obtained the most votes among the RFs. To make this class decision, each RF calculates a probability $p_t(x)$ for the disease-causing class and a probability 1-$p_t(x)$ for the neutral class. Based on the work of Fan & Lin(34), adjusting the probability decision thresholds can improve the performance of a multi-label classifier, a reasoning that could also be applied to our two-label classification. We identified that the best prediction performance was obtained for a probability threshold of 0.489, which has the best median True Positive / False Positive ratio (0.88/0.11) over all individual RF predictors in the ensemble (*SI Appendix*, Fig. S1). Therefore, the decision function $d_{t,c}(x)$ is defined as follows:

Equation (2):

$$d_{t,c}\ (\boldsymbol{x}) = \begin{cases} 0, & (\ c = N \text{ and } p_t(\boldsymbol{x})\ > 0.489) \text{ and } (c = D \text{ and } p_t(\boldsymbol{x})\ \leq 0.489) \\ 1, & (c = N \text{ and } p_t(\boldsymbol{x})\ \leq 0.489) \text{ and } (c = D \text{ and } p_t(\boldsymbol{x})\ > 0.489) \end{cases}$$

Each final class prediction for a bi-locus combination $x$ is, therefore, supported by a classification score (CS), see Equation (3), that is defined as the median (med) of the disease-causing class probabilities over all 500 independent RFs for that bi-locus combination.

Equation (3):

$$CS\ (\boldsymbol{x}) = \underset{t=1,\ldots,T}{\text{med}} \{p_t\ (\boldsymbol{x})\}$$

The final prediction also provides a support score (SS), see Equation (4), that indicates the percentage of RFs that agree on the disease-causing label for a bi-locus combination $\boldsymbol{x}$.

Equation (4):

$$SS\ (\boldsymbol{x}) = \frac{\sum_{t=1}^{T} d_{t,D}(\boldsymbol{x})}{T} \times\ 100$$

Using these equations, we determined that bi-locus variant combinations predicted as disease-causing with VarCoPP require CS > 0.489 and SS > 50. Higher CS and SS provide a stronger indication of pathogenicity for a bi-locus combination.

- **Validation of VarCoPP using independent disease-causing and neutral data**

As a validation set, we collected 23 new disease-causing bi-locus combinations derived from independent scientific papers published after the release of DIDAv1 (Fig. 3D and *SI Appendix*, Table S5 and Dataset S1). We also collected different sets of random 100, 1000 and 10000 neutral bi-locus combinations from the 1KGP that were unused during training to assess the amount of FPs that we can obtain (Fig. 3D and *SI Appendix*, Datasets S2, S3 and S4), without considering the population of individuals and degrees of separation between genes in a bi-locus combination. However, the genes and variants had similar properties to those used during the training of VarCoPP. The gene pairs that were involved in all these bi-locus combinations had not been exploited during the training phase of the predictor. To assess the predictive ability of VarCoPP on these new data, we created three Precision-Recall (PR) curves using the validation set with the 23 disease-causing combinations and either the 100, 1000 or 10000 random neutral test bi-locus test combinations (*SI Appendix*, Fig. S2). The variant filtering procedure, as well as the order of variants and genes inside each combination, was defined in the same way as for the training sets that were used in this work.

- **Gene panel analysis for assessment of False Positives**

We first assessed the performance of VarCoPP on random lists of genes on multiple 1KGP individuals. We created 100 iterations of random panels consisting of 10, 30, 100 and 300 genes and tested each gene panel on 100 random 1KGP individuals. To create the gene panel specific for Bardet-Biedl syndrome (BBS), we used the publicly available 21-gene list obtained from the Genome Diagnostics Nijmegen laboratory (http://www.genomediagnosticsnijmegen.nl/). We obtained the autism gene panels from the SFARI Gene database (https://gene.sfari.org/) and more specifically the gene panel of scoring category 1 (high confidence, 24 genes), category 2 (strong candidate, 56 genes) and category 3 (suggestive evidence, 158 genes). The variant filtering procedure on these panels was similar with the one performed during the creation of the predictor.

- **Selection of the most relevant biological features**

The initial number of predictor features, based on the representation of each bi-locus combination, was 21 (*SI Appendix*, Table S2). To optimize the model and minimize overfitting, we used a recursive feature elimination (RFE) procedure(35) using a balanced set with median performance among all sets. In each iteration we removed the least important feature based on the Gini importance metric, and calculated statistics about the predictor performance. Based on this procedure, we observed that the use of 10 features led to the first optimal performance peak (*SI Appendix*, Fig. S3). We also observed that in general, information about the fourth allele of a bi-locus combination (gene B, allele B) was removed at the early stages of the feature elimination procedure, indicating that this allele does not contribute significantly to the classification. Although no variant features about the fourth allele remained among the 10 selected ones, we decided for interpretability reasons to include the CADD score of this allele, being the fourth-allele feature that was eliminated last, finalizing the number of selected features to 11.

- **Understanding the contribution of each feature to the predictions using *treeinterpreter***

To gain insight in how each feature contributes to the classification of a bi-locus combination as either disease-causing or neutral, we exploited the *treeinterpreter* Python package (https://github.com/andosa/treeinterpreter, Ando Saabas). This library explores, for each bi-locus combination, information on the order and the way features are used during a decision path from the root of each decision tree to the final leaf. This helps assess how much each feature contributes or "votes" for either the disease-causing or neutral class. The contribution or "vote" for a particular class is represented with a contribution value that is calculated from *treeinterpreter*. A positive feature contribution value means that this feature votes for the disease-causing class, while a negative feature contribution value means that the feature contributes to the neutral class. This also indicates that in the end, the sum of the decision values of all features for a particular bi-locus combination delivers the final classification; positive sum for the disease-causing class and negative sum for the neutral class. As we obtain a contribution value for each decision tree, in order to obtain the feature contribution values for each bi-locus combination inside an individual RF, the contribution values were averaged among all trees of that RF, providing us with a vector called the Decision Profile (DP), which contained average decision values for each feature. Feature contribution values should not be confused with the actual feature values of a bi-locus combination; the former are derived using *treeinterpreter*, whereas the latter are the actual values of the features used for classification. Thus, if a bi-locus combination is represented with a feature vector x = (x1,x2,...,x10), then, using the *treeinterpreter* library, we get a DP vector y=(y1,y2,...,y10) for each individual RF, where $y_i$ is the contribution value of the feature *i*.

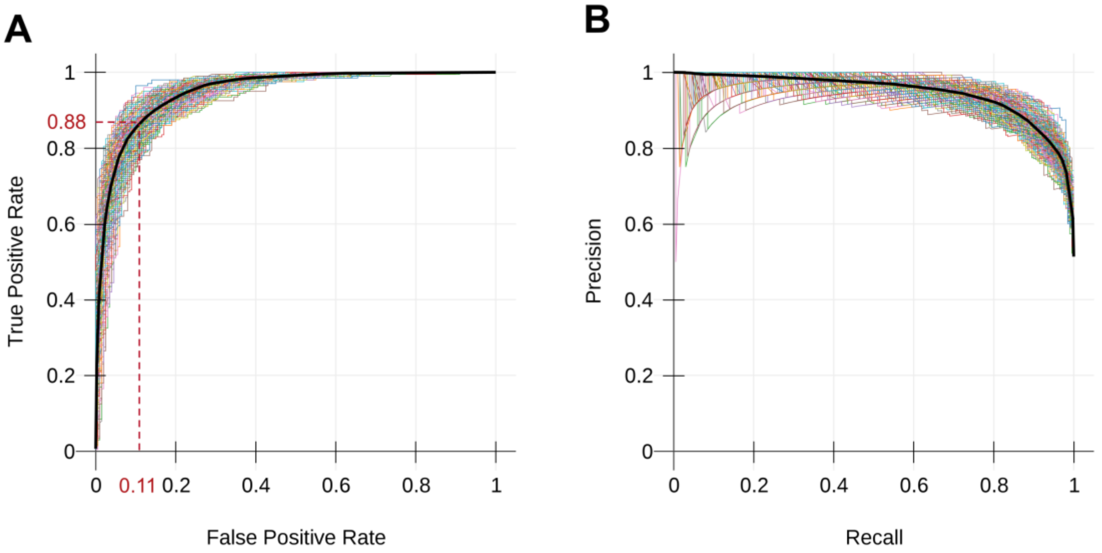


**Fig. S1.** Performance of VarCoPP during cross-validation. The different colored lines represent the curve from each individual predictor of the ensemble, while the black curve is the average curve among all predictors. (A) Receive Operator Characteristic (ROC) curve. The median optimal True Positive (TP) and False Positive (FP) ratio is 0.88 / 0.11 and is associated with a median probability threshold of 0.489 that differentiates disease-causing from neutral combinations. (B) Precision - Recall (PR) curve.

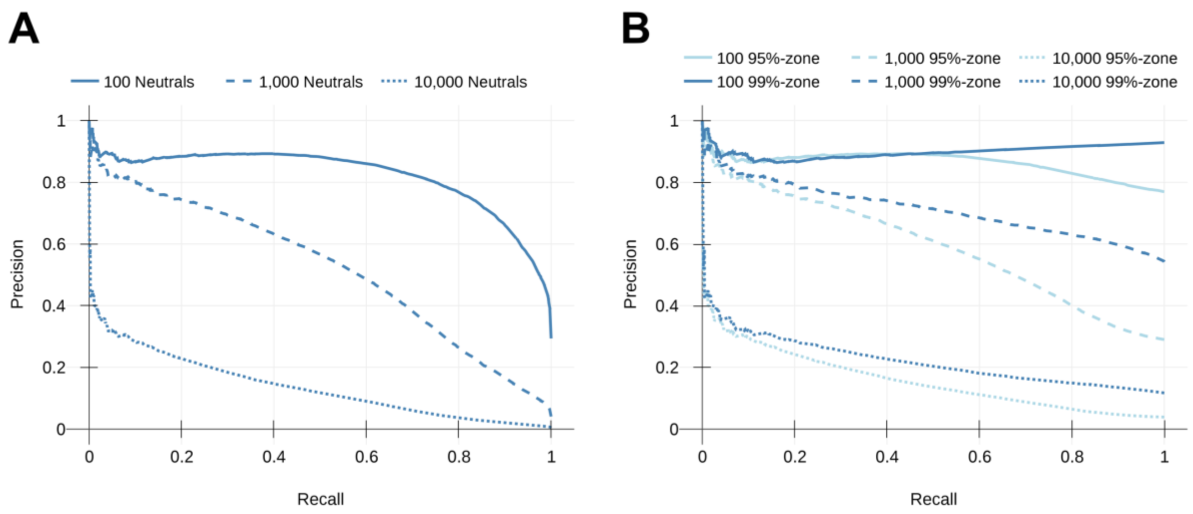


**Fig. S2.** Precision - Recall curve (PR) curves when VarCoPP is tested on the neutral test set and validation set bi-locus combinations. (A) PR using the instances of the validation set together with either the 100 (straight line), 1000 (dashed line) or 10000 test neutral combinations (dotted line). This PR curve contains the average performance among all independent 500 RFs of VarCoPP. (B) PR curve using the validation set instances together with those of either the 100 (straight line), 1000 (dashed line) or 10000 test neutral sets (dark blue), which fall into the 95%-zone (light blue) or 99%-zone (dark blue).

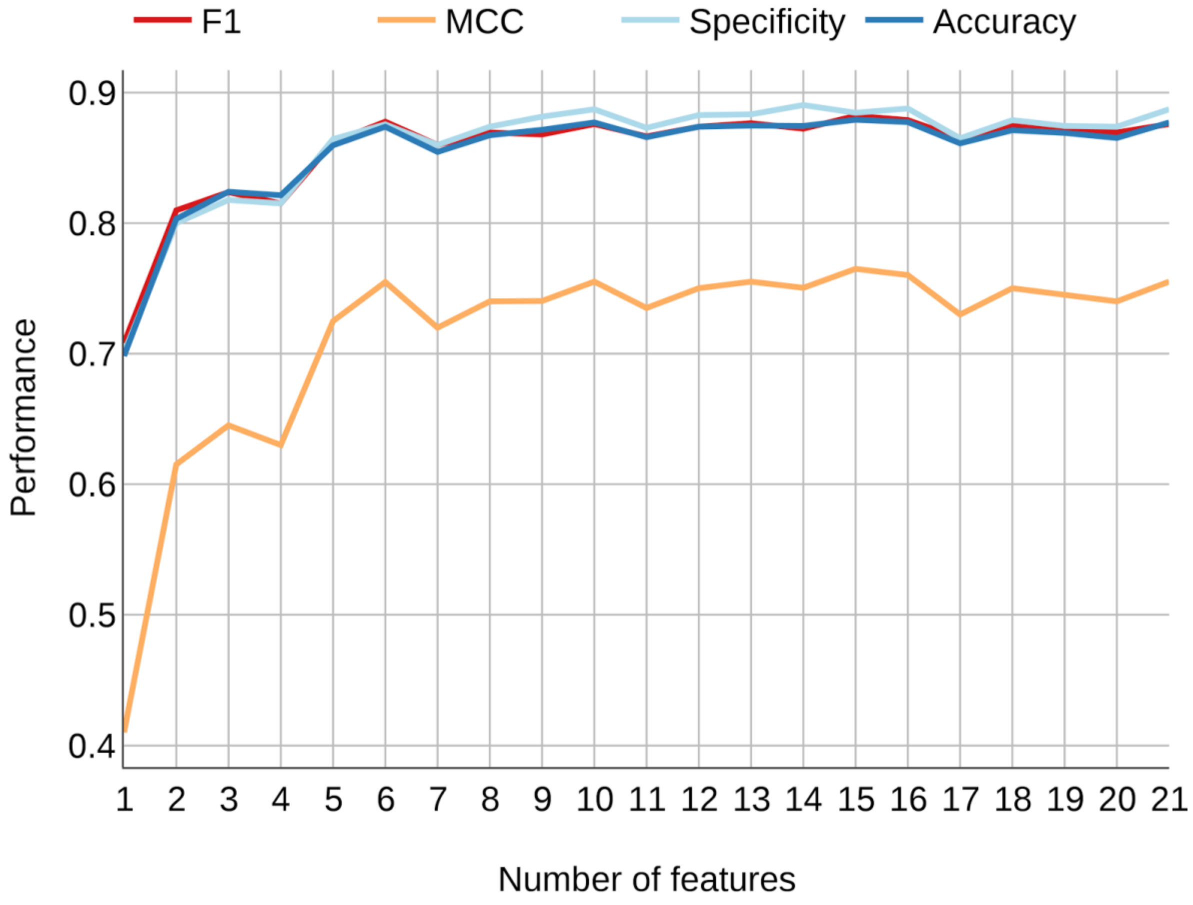


**Fig. S3.** Predictor performance during the recursive feature elimination procedure of a balanced training set with median performance among the balanced sets, after a cross validation procedure. The x-axis represents the number of features included for prediction during each elimination procedure, while the y axis represents the average performance during the cross-validation for that number of features. Performance was measured in terms of accuracy, specificity, F1 score (a balanced measure between precision and recall) and Matthews Correlation Coefficient (a balanced measure of the quality of a two-class classification, resembling a correlation coefficient between the observed and predicted binary classifications). It is shown in that a set of 10 features inferred the first best peak among all performance measures (with slightly better scores than 6 features). A set of 15 features inferred the second-best peak with a slight difference, but in order to reduce the chances of overfitting, we decided to continue with the set of 10. For interpretation reasons, we also included CADD4 (CADD score of the second variant allele in gene B), which was not part of the 10 first selected features, finalizing the number of features to 11.

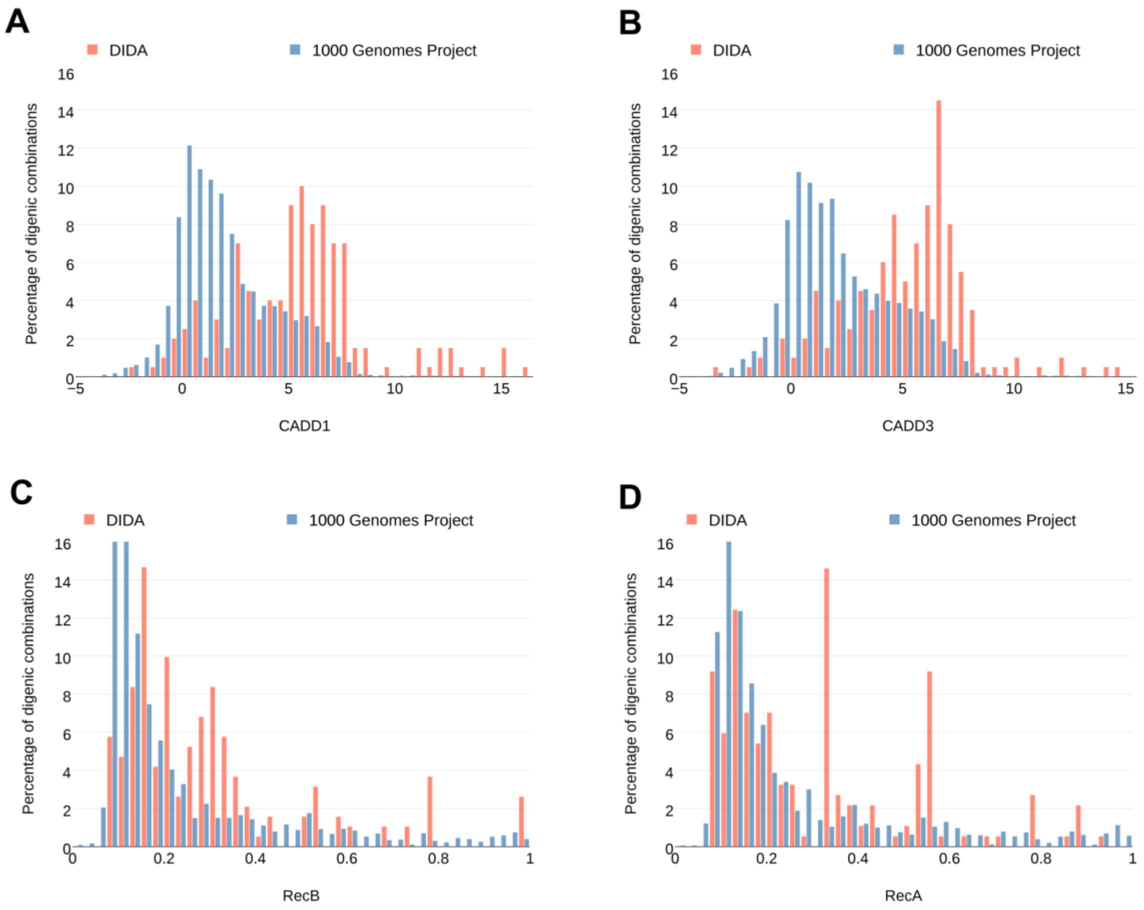


**Fig. S4.** Distribution of feature values for (A) CADD1, (B) CADD3, (C) RecB and (D) RecA between DIDAv1 and the 1000 Genomes Project training sets.

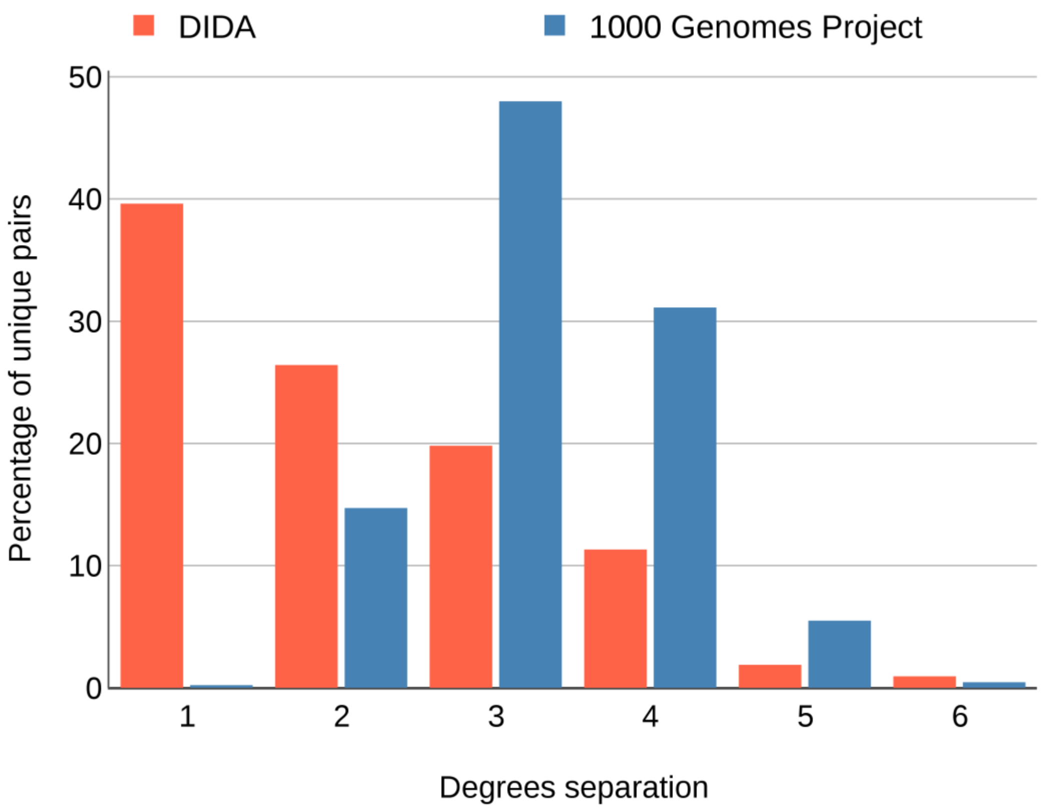


**Fig. S5.** Histogram of the degrees separation for the gene pairs present in the subset of the 1KGP (in blue) used in our analysis and DIDA (in red). Degrees separation depicts how many proteins intervene in the pathway between a pair of genes. Value of 1 indicates that the proteins of two genes in a pair are directly interacting.

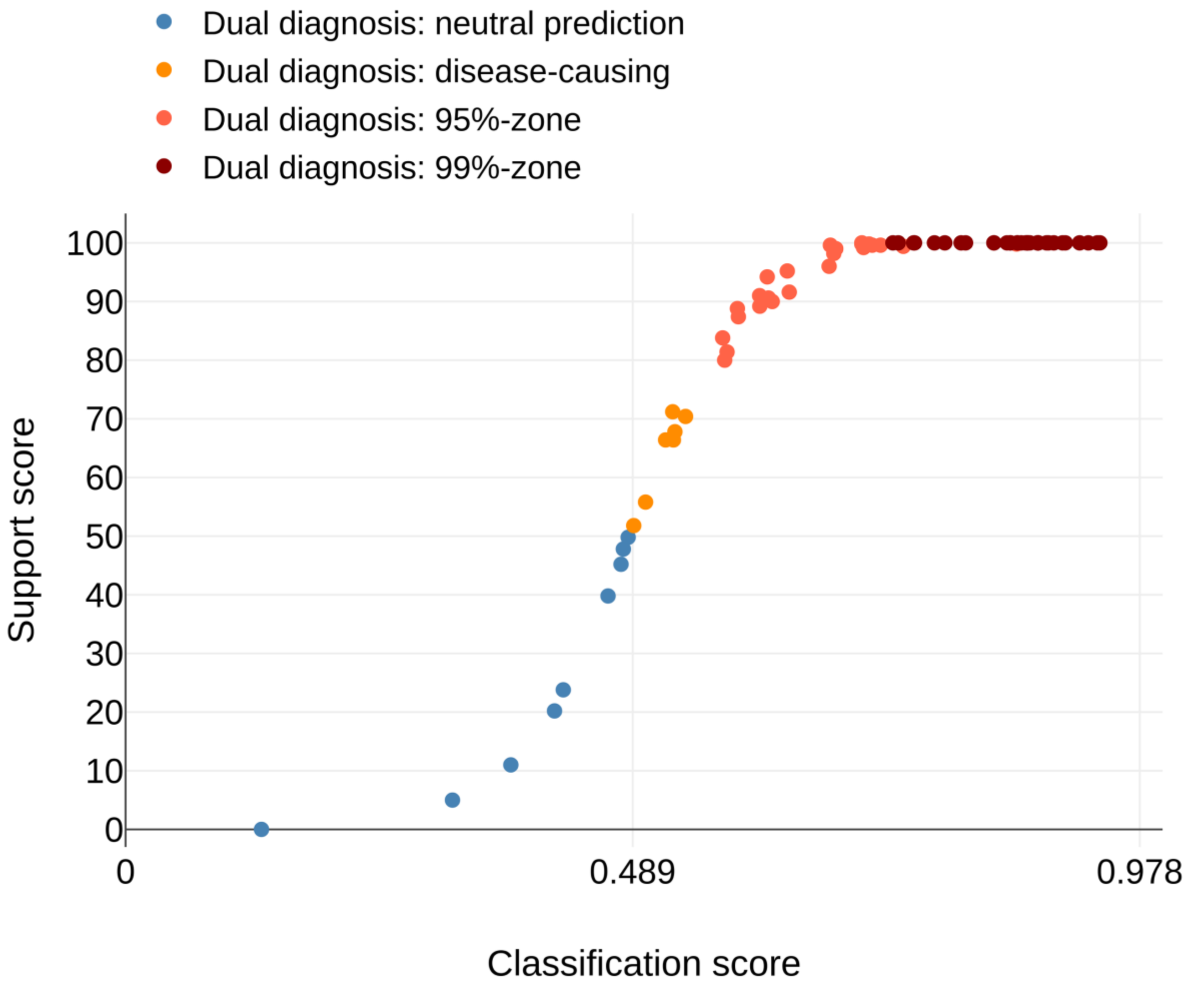


**Fig. S6.** Performance of VarCoPP on 76 Dual Molecular Diagnosis cases, extracted from Posey *et al.*(36).

**Table S1. Information on individuals of the 1KGP carrying DIDA bi-locus combinations.**

| 1KGP individual | Population | Family ID | DIDA ID | Associated disease | PMID |
|---|---|---|---|---|---|
| HG02375 | CDX | N/A | dd203 | Kallmann syndrome | 20022991 |
| HG04042 | STU | ST171 | dd220 | MODY | 25041077 |
| HG00123 | GBR | N/A | dd193 | Familial hemophagocytic lymphohistiocytosis | 24916509 |
| NA12842 | CEU | 1458 | dd180 | Familial hemophagocytic lymphohistiocytosis | 24916509 |
| NA12812 | CEU | 1454 | dd196 | Familial hemophagocytic lymphohistiocytosis | 24916509 |
| NA12342 | CEU | 1330 | dd193 | Familial hemophagocytic lymphohistiocytosis | 24916509 |
| NA12342 | CEU | 1330 | dd188 | Familial hemophagocytic lymphohistiocytosis | 24916509 |
| HG03388 | MSL | SL28 | dd202 | Kallmann syndrome | 20022991 |
| NA19225 | YRI | Y057 | dd202 | Kallmann syndrome | 20022991 |
| NA20807 | TSI | N/A | dd188 | Familial hemophagocytic lymphohistiocytosis | 24916509 |
| HG03410 | MSL | SL35 | dd202 | Kallmann syndrome | 20022991 |
| HG01670 | IBS | | dd193 | Familial hemophagocytic lymphohistiocytosis | 24916509 |
| HG03713 | ITU | IT001 | dd220 | MODY | 25041077 |
| NA19031 | LWK | N/A | dd202 | Kallmann syndrome | 20022991 |
| HG00112 | GBR | N/A | dd188 | Familial hemophagocytic lymphohistiocytosis | 24916509 |

**Table S2. Features used to annotate the 1KGP and DIDA data sets. Those that remained after the feature selection procedure are marked with (*).**

| Feature | Feature Abbreviations | Description | PMID |
|---|---|---|---|
| CADD raw score | CADD1*<br>CADD2*<br>CADD3*<br>CADD4* | First variant allele of gene A<br>Second variant allele of gene A<br>First variant allele of gene B<br>Second variant allele of gene B | 24487276 |
| Pfam protein domain | Pfam1<br>Pfam2<br>Pfam3<br>Pfam4 | First variant allele of gene A<br>Second variant allele of gene A<br>First variant allele of gene B<br>Second variant allele of gene B | 26673716 |
| Amino acid hydrophobicity difference | Hydr1*<br>Hydr2<br>Hydr3<br>Hydr4 | First variant allele of gene A<br>Second variant allele of gene A<br>First variant allele of gene B<br>Second variant allele of gene B | 8836100 |
| Amino acid flexibility difference | Flex1*<br>Flex2<br>Flex3<br>Flex4 | First variant allele of gene A<br>Second variant allele of gene A<br>First variant allele of gene B<br>Second variant allele of gene B | -<br>Bhaskaran & Ponnuswamy 1988 |
| Haploinsufficiency probability | HI_A*<br>HI_B* | Haploinsufficiency probability for gene A<br>Haploinsufficiency probability for gene B | 20976243 |
| Recessiveness probability | RecA*<br>RecB* | Recessiveness probability for gene A<br>Recessiveness probability for gene B | 22344438 |
| Biological distance | Biol_Dist* | Biological relatedness between gene A and gene B | 24694260 |

**Table S3. Computer readable representation of values for the features presented in Table S2.**

| Variant features | Type | Values | Representation in the vector | Explanation |
|---|---|---|---|---|
| CADD raw score<br>(alleles 1 – 4) | Numerical | Raw score<br>NaN<br>Wild-type | [raw score]<br>[CADD median]*<br>[-3] | * Median value of the CADD score of the same type of variants for either DIDA or 1KGP. |
| Amino acid flexibility difference<br>(alleles 1 – 4) | Numerical | Protein variant<br>Intronic/splicing variant<br>NaN (frameshift/loss of stop mutation)<br>Wild-type | [difference value]<br>[0.0]<br>[median value]*<br>[0.0] | * Median value of either DIDA (0.012) or 1KGP (0.0). |
| Amino acid hydrophobicity difference<br>(alleles 1 – 4) | Numerical | Protein variant<br>Intronic/splicing variant<br>NaN (frameshift/loss of stop mutation)<br>Wild-type | [difference value]<br>[0.0]<br>[median value]*<br>[0.0] | *Median value of either DIDA (-0.1) or 1KGP (-0.01). |
| Pfam region<br>(alleles 1 – 4) | Categorical | Yes<br>No<br>NaN<br>Wild-type | [1]<br>[0]<br>[0]<br>[same value as the first allele of the variant] | |
| Haploinsufficiency probability<br>(geneA/B) | Numerical | Known probability<br>NaN | [value]<br>[median = 0.19898] | |
| Recessiveness probability<br>(geneA/B) | Numerical | Known probability<br>NaN | [value]<br>[median = 0.12788] | |
| Biological distance | Numerical | Known distance<br>NaN | [distance]<br>[median]* | *Median for gene pairs with “NaN” pathway for DIDA (4.72) and 1KGP (18.16). |

**Table S4. Mean performance and standard deviation (sd) statistics among all 500 RFs of the ensemble predictor during cross-validation and using the final selection of 11 features.**

| Number of balanced sets | Accuracy | | Precision | | Sensitivity | | MCC | |
|---|---|---|---|---|---|---|---|---|
| | Mean | Sd | Mean | Sd | Mean | Sd | Mean | Sd |
| 10 | 0.884 | 0.014 | 0.892 | 0.017 | 0.876 | 0.014 | 0.77 | 0.022 |
| 50 | 0.874 | 0.013 | 0.884 | 0.014 | 0.863 | 0.015 | 0.750 | 0.024 |
| 100 | 0.875 | 0.016 | 0.883 | 0.019 | 0.865 | 0.016 | 0.751 | 0.031 |
| 500 | 0.878 | 0.014 | 0.886 | 0.016 | 0.868 | 0.015 | 0.757 | 0.026 |
| 500* | 0.877 | 0.014 | 0.880 | 0.016 | 0.874 | 0.015 | 0.756 | 0.027 |
| 1000 | 0.878 | 0.014 | 0.886 | 0.016 | 0.867 | 0.015 | 0.756 | 0.026 |
| 1500 | 0.876 | 0.014 | 0.884 | 0.016 | 0.867 | 0.015 | 0.754 | 0.026 |
| 2000 | 0.877 | 0.014 | 0.885 | 0.016 | 0.867 | 0.015 | 0.755 | 0.026 |
| 3500 | 0.877 | 0.014 | 0.885 | 0.016 | 0.867 | 0.015 | 0.755 | 0.027 |

***** adjusted probability threshold of 0.489

**Table S5. Validation set of 23 new disease-causing bi-locus combinations. For details about the feature annotation of these combinations, see *SI Appendix*, Dataset S1.**

| Combination ID | Gene pair* | Type | Associated disease | PMID |
|---|---|---|---|---|
| testpos_1 | *TRIM54 - TRIM63* | triallelic | Protein aggregate myopathy | 25801283 |
| testpos_2 | *PSMA3 - PSMB8* | diallelic | CANDLE syndrome | 26524591 |
| testpos_3 | *PSMA3 - PSMB8* | diallelic | CANDLE syndrome | 26524591 |
| testpos_4 | *PSMB4 - PSMB9* | diallelic | CANDLE syndrome | 26524591 |
| testpos_5 | *PSMB4 - PSMB8* | diallelic | CANDLE syndrome | 26524591 |
| testpos_6 | *CDK5RAP2 - CEP152* | diallelic | Seckel syndrome | 26436113 |
| testpos_7 | *COL4A4 - COL4A3* | diallelic | Alport sydrome | 25575550 |
| testpos_8 | *COL4A4 - COL4A3* | diallelic | Alport sydrome | 25575550 |
| testpos_9 | *COL4A4 - COL4A3* | diallelic | Alport sydrome | 25575550 |
| testpos_10 | *COL4A4 - COL4A3* | diallelic | Alport sydrome | 25575550 |
| testpos_11 | *COL4A4 - COL4A3* | diallelic | Alport sydrome | 25575550 |
| testpos_12 | *COL4A5 - COL4A4* | diallelic | Alport sydrome | 25575550 |
| testpos_13 | *COL4A5 - COL4A4* | diallelic | Alport sydrome | 25575550 |
| testpos_14 | *COL4A5 - COL4A4* | diallelic | Alport sydrome | 25575550 |
| testpos_15 | *COL4A5 - COL4A4* | triallelic | Alport sydrome | 25575550 |
| testpos_16 | *SEC23A - MAN1B1* | tetrallelic | Overgrowth syndrome | 27148587 |
| testpos_17 | *RP1L1 - C2orf71* | triallelic | Syndromic retinitis pigmentosa | 29295593 |
| testpos_18 | *SHH - DISP1* | diallelic | Holoprosencephaly | 26748417 |
| testpos_19 | *MITF - GJB2* | diallelic | Deafness | 27057829 |
| testpos_20 | *AHI1 - CEP290* | triallelic | Leber congenital amaurosis | 20683928 |
| testpos_21 | *RPE65 - CEP290* | triallelic | Leber congenital amaurosis | 20683928 |
| testpos_22 | *CRB1 - CEP290* | triallelic | Leber congenital amaurosis | 20683928 |
| testpos_23 | *AHI1 - CEP290* | triallelic | Joubert syndrome | 20683928 |

*Gene order based on Gene Damage Index (GDI)

**Table S6. Statistics on the performance of VarCoPP using random test sets of 100, 1000 and 10000 neutral bi-locus combinations derived from the 1KGP (TNs=True Negatives, FPs=False Positives, SS = Support Score, CS = Classification Score).**

| | number of test combinations | | |
|---|---|---|---|
| | **100** | **1000** | **10000** |
| definitive TNs (SS=0) | 67.0% | 72.1% | 72% |
| FPs (SS>50) | 7.0% | 7.7% | 7.2% |
| min. 95%-zone SS | 80.6 | 74.8 | 74.8 |
| min. 95%-zone CS | 0.57 | 0.55 | 0.55 |
| min. 99%-zone SS | 100.0 | 100.0 | 100.0 |
| min. 99%-zone CS | 0.74 | 0.749 | 0.72 |

**Table S7. Statistics on recall on the training data and on the validation set of 23 independent disease-causing bi-locus variant combinations, when applying the 95% and 99% confidence zones.**

| | Overall recall | 95% confidence zone recall | 99% confidence zone recall |
|---|---|---|---|
| Training set (cross-validation) | 0.87 | 0.84 | 0.60 |
| Validation set | 0.87 | 0.87 | 0.60 |

**Table S8. Average performance and standard deviation (sd) statistics among all 500 RFs of the ensemble predictor during cross-validation, using each time 1KGP individuals from one particular continent as training set against DIDA.**

| 1KGP continent | Accuracy | | Precision | | Sensitivity | | MCC | |
|---|---|---|---|---|---|---|---|---|
| | Average | Sd | Average | Sd | Average | Sd | Average | Sd |
| Africa | 0.89 | 0.01 | 0.89 | 0.01 | 0.89 | 0.01 | 0.78 | 0.02 |
| America | 0.87 | 0.01 | 0.84 | 0.02 | 0.91 | 0.01 | 0.74 | 0.04 |
| East Asia | 0.86 | 0.01 | 0.83 | 0.02 | 0.91 | 0.01 | 0.73 | 0.04 |
| Europe | 0.86 | 0.01 | 0.83 | 0.02 | 0.91 | 0.01 | 0.73 | 0.03 |
| South Asia | 0.87 | 0.01 | 0.84 | 0.02 | 0.90 | 0.01 | 0.74 | 0.04 |

**Additional Dataset S1 (separate file)**
Annotated features and VarCoPP prediction scores for the 23 bi-locus combinations of the disease-causing validation set.

**Additional Dataset S2 (separate file)**
Annotated features and VarCoPP prediction scores for the 100 random bi-locus combinations extracted from the 1KGP.

**Additional Dataset S3 (separate file)**
Annotated features and VarCoPP prediction scores for the 1000 random bi-locus combinations extracted from the 1KGP.

**Additional Dataset S4 (separate file)**
Annotated features and VarCoPP prediction scores for the 10000 random bi-locus combinations extracted from the 1KGP.

**Additional Dataset S5 (separate file)**
Annotated features and VarCoPP prediction scores for the 76 Dual Molecular Diagnosis bi-locus combinations extracted from the paper of Posey *et al.*(36)
