## Supplementary material for "Predicting disease-causing variant combinations": Dataset S1

| Combination_ID | Flex1 | Hydr1 | CADD1 | CADD2 | CADD3 | CADD4 | HI_A | RecA | HI_B | RecB | Biol_Dist | Classification_score | Support_score | Predicted_class |
|---|---|---|---|---|---|---|---|---|---|---|---|---|---|---|
| testpos_1 | 0.05 | -0.81 | 6.941602 | -3 | 11.872989 | 11.872989 | 0.13331 | 0.13638 | 0.13419 | 0.38803 | 1.77305 | 0.915205882 | 100 | Disease_causing |
| testpos_2 | 0.02727273 | -0.1989091 | 3.427595 | -3 | 7.077599 | -3 | 0.9944 | 0.40044 | 0.40872 | 0.12788 | 1.6129 | 0.587541478 | 80.6 | Disease_causing |
| testpos_3 | 0.012 | -0.1 | 5.226106 | -3 | 7.077599 | -3 | 0.9944 | 0.40044 | 0.40872 | 0.12788 | 1.6129 | 0.76791886 | 100 | Disease_causing |
| testpos_4 | 0.012 | -0.1 | 5.502773 | -3 | 3.058361 | -3 | 0.86588 | 0.29568 | 0.67676 | 0.12788 | 1.6129 | 0.479021045 | 46.2 | Neutral |
| testpos_5 | 0.04018182 | -0.12 | 11.993666 | -3 | 6.231252 | -3 | 0.86588 | 0.29568 | 0.40872 | 0.12788 | 1.6129 | 0.827555556 | 100 | Disease_causing |
| testpos_6 | -0.14 | 0.37 | 4.179049 | -3 | 8.225553 | -3 | 0.03779 | 0.08467 | 0.16634 | 0.09998 | 1.11111 | 0.57 | 85.2 | Disease_causing |
| testpos_7 | 0.04 | 2.01 | 5.937124 | -3 | 3.734655 | -3 | 0.09807 | 0.24901 | 0.2922 | 0.33981 | 1.11111 | 0.851974559 | 100 | Disease_causing |
| testpos_8 | 0.18 | 0.16 | 4.567777 | -3 | 6.142108 | -3 | 0.09807 | 0.24901 | 0.2922 | 0.33981 | 1.11111 | 0.89534992 | 100 | Disease_causing |
| testpos_9 | 0.012 | -0.1 | 5.09332 | -3 | 5.364796 | -3 | 0.09807 | 0.24901 | 0.2922 | 0.33981 | 1.11111 | 0.922962522 | 100 | Disease_causing |
| testpos_10 | 0.007375 | -0.088625 | 2.363621 | -3 | 2.846208 | -3 | 0.09807 | 0.24901 | 0.2922 | 0.33981 | 1.11111 | 0.416401862 | 24 | Neutral |
| testpos_11 | 0 | 0 | 5.318655 | -3 | 4.596302 | -3 | 0.09807 | 0.24901 | 0.2922 | 0.33981 | 1.11111 | 0.86 | 100 | Disease_causing |
| testpos_12 | 0.15 | 0.06 | 4.389205 | -3 | 5.409798 | -3 | 0.87803 | 0.12788 | 0.09807 | 0.24901 | 1.11111 | 0.678357348 | 97.8 | Disease_causing |
| testpos_13 | 0.03 | 1.22 | 5.023704 | -3 | 6.377085 | -3 | 0.87803 | 0.12788 | 0.09807 | 0.24901 | 1.11111 | 0.729165449 | 99.4 | Disease_causing |
| testpos_14 | 0.01 | 0.8 | 6.344966 | 6.344966 | 6.267977 | -3 | 0.87803 | 0.12788 | 0.09807 | 0.24901 | 1.11111 | 0.89 | 100 | Disease_causing |
| testpos_15 | 0.15 | 0.06 | 4.779126 | -3 | 1.701008 | -0.229803 | 0.87803 | 0.12788 | 0.09807 | 0.24901 | 1.11111 | 0.298919643 | 1.6 | Neutral |
| testpos_16 | -0.16 | -0.08 | 4.466229 | 4.466229 | 7.6294 | 7.6294 | 0.68918 | 0.16819 | 0.31651 | 0.12788 | 9.80316 | 0.83 | 100 | Disease_causing |
| testpos_17 | 0.018 | 0.23454546 | 9.880615 | 2.882466 | 10.509517 | -3 | 0.09178 | 0.12788 | 0.19898 | 0.12788 | 4.72 | 0.88 | 100 | Disease_causing |
| testpos_18 | 0.02 | 0.13 | 5.944149 | -3 | 3.687832 | -3 | 0.49106 | 0.89143 | 0.16068 | 0.11183 | 1.11111 | 0.57 | 81.2 | Disease_causing |
| testpos_19 | 0.18 | -1.05 | 8.035357 | -3 | 6.664169 | -3 | 0.8509 | 0.42299 | 0.42941 | 0.50569 | 10.83333 | 0.94875 | 100 | Disease_causing |
| testpos_20 | -0.19 | -0.51 | 4.805921 | -3 | 9.634885 | -0.026682 | 0.15848 | 0.12754 | 0.48442 | 0.31068 | 1.38889 | 0.816654762 | 100 | Disease_causing |
| testpos_21 | 0.18 | -1.05 | 7.207124 | -3 | 7.541326 | -0.026682 | 0.13937 | 0.25571 | 0.48442 | 0.31068 | 6.98716 | 0.949602142 | 100 | Disease_causing |
| testpos_22 | 0.00963636 | -0.2974545 | 6.795635 | -3 | 5.291384 | -0.026682 | 0.20152 | 0.09895 | 0.48442 | 0.31068 | 10.83333 | 0.84570613 | 100 | Disease_causing |
| testpos_23 | -0.01 | 0.57 | 3.712252 | -3 | 15.015364 | 12.87429 | 0.15848 | 0.12754 | 0.48442 | 0.31068 | 1.38889 | 0.8346875 | 99.8 | Disease_causing |
