## Supplementary material for "Predicting disease-causing variant combinations": Dataset S2

| Combination_ID | Flex1 | Hydr1 | CADD1 | CADD2 | CADD3 | CADD4 | HI_A | RecA | HI_B | RecB | Biol_Dist | Classification_score | Support_score | Predicted_class |
|---|---|---|---|---|---|---|---|---|---|---|---|---|---|---|
| testneg_1 | -0.08 | -0.03 | 7.031819 | -3 | -0.332338 | -3 | 0.64608 | 0.35524 | 0.13708 | 0.81545 | 24.41367 | 0.58 | 86.6 | Disease_causing |
| testneg_2 | 0 | 0 | 0.56241 | -3 | 0.96112 | -3 | 0.9511 | 0.54626 | 0.06331 | 0.2396 | 6.43627 | 0.064892602 | 0 | Neutral |
| testneg_3 | 0.05 | -0.81 | 3.077798 | -3 | -0.157172 | -3 | 0.34295 | 0.12788 | 0.39344 | 0.12788 | 10.41667 | 0.03 | 0 | Neutral |
| testneg_4 | 0 | 0 | -0.153655 | -3 | 1.220563 | -3 | 0.06048 | 0.12788 | 0.47188 | 0.10037 | 30.76653 | 0.02 | 0 | Neutral |
| testneg_5 | 0 | -0.01 | 6.576011 | -3 | -0.811455 | -0.811455 | 0.56605 | 0.15893 | 0.10638 | 0.66554 | 32.89375 | 0.468968254 | 39.2 | Neutral |
| testneg_6 | 0.18 | 0.16 | 0.883022 | -3 | 0.086889 | -3 | 0.1592 | 0.12788 | 0.66693 | 0.0986 | 18.76159 | 0.02 | 0 | Neutral |
| testneg_7 | 0 | 0 | 1.736361 | -3 | 4.866108 | -3 | 0.36461 | 0.11015 | 0.19898 | 0.12788 | 31.94292 | 0.041113194 | 0 | Neutral |
| testneg_8 | -0.17 | 1.22 | -2.338688 | -2.338688 | 6.206483 | -3 | 0.33129 | 0.09957 | 0.18861 | 0.10842 | 18.88889 | 0.28 | 0.6 | Neutral |
| testneg_9 | 0.03 | 0.12 | 0.235146 | -3 | 2.178935 | -3 | 0.16856 | 0.19221 | 0.77847 | 0.10751 | 14.58333 | 0.040694444 | 0 | Neutral |
| testneg_10 | -0.07 | -0.38 | -4.505557 | -3 | 1.217652 | -3 | 0.36858 | 0.09681 | 0.19898 | 0.12788 | 27.3569 | 0.02 | 0 | Neutral |
| testneg_11 | 0 | 0 | 0.687946 | -3 | 3.934145 | 3.934145 | 0.09269 | 0.12788 | 0.25011 | 0.12092 | 10.41667 | 0.25375 | 0 | Neutral |
| testneg_12 | 0.14 | -1.01 | 3.553934 | -3 | 1.268416 | -3 | 0.47997 | 0.10506 | 0.43977 | 0.17284 | 16.8018 | 0.189940476 | 0 | Neutral |
| testneg_13 | 0 | 0 | -0.101683 | -3 | 4.54534 | -3 | 0.72657 | 0.11262 | 0.06713 | 0.11112 | 19.68336 | 0.070833333 | 0 | Neutral |
| testneg_14 | 0.01 | -0.57 | 1.867073 | -3 | -0.574737 | -3 | 0.07999 | 0.23927 | 0.56678 | 0.09955 | 21.51158 | 0.06 | 0 | Neutral |
| testneg_15 | 0 | 0 | 0.062047 | -3 | 0.651966 | -3 | 0.21928 | 0.22085 | 0.28326 | 0.13944 | 10.41667 | 0.062051129 | 0 | Neutral |
| testneg_16 | 0 | 0 | 0.782076 | -3 | 3.633607 | -3 | 0.9193 | 0.14584 | 0.2172 | 0.10891 | 10.83333 | 0.060619048 | 0 | Neutral |
| testneg_17 | 0 | 0 | 0.452369 | -3 | 1.994943 | -3 | 0.09504 | 0.38939 | 0.33148 | 0.11968 | 33.59381 | 0.06 | 0 | Neutral |
| testneg_18 | -0.08 | -0.03 | 3.206814 | -3 | 0.192097 | -3 | 0.7022 | 0.35308 | 0.03651 | 0.06141 | 14.27748 | 0.071572581 | 0 | Neutral |
| testneg_19 | 0.18 | 0.16 | -1.022735 | -3 | 3.032496 | -3 | 0.08896 | 0.12788 | 0.10285 | 0.10186 | 29.16667 | 0.081547619 | 0 | Neutral |
| testneg_20 | 0 | 0 | 0.919505 | -3 | -1.457001 | -3 | 0.19898 | 0.12788 | 0.25119 | 0.13626 | 13.51351 | 0.030215278 | 0 | Neutral |
| testneg_21 | -0.01 | -0.8 | 1.016479 | 1.016479 | -0.307406 | -3 | 0.12649 | 0.12788 | 0.62858 | 0.16645 | 11.93907 | 0.110357143 | 0 | Neutral |
| testneg_22 | 0 | 0 | 3.854758 | -3 | 3.287017 | -3 | 0.54826 | 0.24974 | 0.1594 | 0.22002 | 16.73378 | 0.555296916 | 75.6 | Disease_causing |
| testneg_23 | 0.02 | -0.63 | 0.945927 | -3 | 3.857236 | -3 | 0.21458 | 0.10398 | 0.09602 | 0.10726 | 47.06131 | 0.055 | 0 | Neutral |
| testneg_24 | 0 | 0 | 1.774559 | -3 | 4.928834 | -3 | 0.08628 | 0.11498 | 0.3405 | 0.1816 | 14.26037 | 0.242913603 | 1.2 | Neutral |
| testneg_25 | 0 | 0 | -0.134633 | -3 | 4.068015 | -3 | 0.1617 | 0.17928 | 0.64914 | 0.12481 | 10 | 0.13792803 | 0 | Neutral |
| testneg_26 | 0.04 | -0.23 | 3.457136 | -3 | 1.417365 | 1.417365 | 0.40707 | 0.37988 | 0.61184 | 0.55597 | 18.33333 | 0.413333333 | 19.4 | Neutral |
| testneg_27 | 0 | 0 | 1.095746 | -3 | 0.011454 | -3 | 0.24274 | 0.13034 | 0.05877 | 0.12788 | 17.21847 | 0.01 | 0 | Neutral |
| testneg_28 | 0.07 | -0.31 | 4.510214 | -3 | 1.165526 | -3 | 0.35234 | 0.1303 | 0.63418 | 0.69224 | 12.4985 | 0.230734127 | 0.2 | Neutral |
| testneg_29 | 0.16 | -0.37 | 5.071893 | -1.110728 | 6.382916 | -3 | 0.28817 | 0.10531 | 0.46244 | 0.56748 | 13.79561 | 0.766917448 | 99.4 | Disease_causing |
| testneg_30 | 0.01 | -0.57 | 5.075109 | -3 | -1.970817 | -3 | 0.68863 | 0.09663 | 0.2058 | 0.28174 | 25.18842 | 0.2795 | 0.8 | Neutral |
| testneg_31 | 0 | 0 | 1.605417 | -3 | 1.56158 | -3 | 0.57729 | 0.10831 | 0.0819 | 0.1823 | 13.78642 | 0.05 | 0 | Neutral |
| testneg_32 | 0 | 0 | -0.255524 | -3 | 1.659864 | -3 | 0.60171 | 0.24685 | 0.25938 | 0.12188 | 36.52499 | 0.03 | 0 | Neutral |
| testneg_33 | 0 | 0 | 0.348328 | -3 | 4.886271 | -3 | 0.28878 | 0.16765 | 0.19898 | 0.12788 | 12.29167 | 0.104491827 | 0 | Neutral |
| testneg_34 | 0.03 | 0.12 | 1.354328 | -3 | 0.513578 | -3 | 0.15623 | 0.1447 | 0.19898 | 0.07409 | 22.89111 | 0.01 | 0 | Neutral |
| testneg_35 | 0 | 0 | 0.199574 | -3 | 0.316439 | -3 | 0.0606 | 0.39625 | 0.09531 | 0.09236 | 13.10913 | 0.060995872 | 0 | Neutral |
| testneg_36 | 0.08 | 0.03 | -2.315401 | -3 | -0.163315 | -3 | 0.34295 | 0.12788 | 0.31125 | 0.20061 | 12.63889 | 0.059668011 | 0 | Neutral |
| testneg_37 | -0.03 | 1.03 | -1.918288 | -3 | 1.139058 | -3 | 0.36566 | 0.15312 | 0.27887 | 0.18663 | 18.33333 | 0.109028571 | 0 | Neutral |
| testneg_38 | -0.05 | -0.29 | 5.49525 | -3 | 2.642734 | -3 | 0.54792 | 0.54626 | 0.19898 | 0.12788 | 20.94851 | 0.45 | 32.6 | Neutral |
| testneg_39 | 0.21 | 0.15 | 1.305251 | -3 | 0.468237 | -3 | 0.06304 | 0.12788 | 0.32334 | 0.10525 | 26.34995 | 0.03 | 0 | Neutral |
| testneg_40 | -0.17 | 1.22 | -2.338688 | -2.338688 | 5.650938 | -3 | 0.33129 | 0.09957 | 0.16671 | 0.18311 | 12.29167 | 0.44825 | 31 | Neutral |
| testneg_41 | -0.1 | -1.9 | 5.681253 | -3 | 1.232154 | -3 | 0.11359 | 0.08631 | 0.17768 | 0.10832 | 18.39641 | 0.16 | 0 | Neutral |
| testneg_42 | 0.16 | -1.37 | 2.93759 | -3 | 0.888014 | -3 | 0.17231 | 0.1358 | 0.47451 | 0.3285 | 10.41667 | 0.268464286 | 0.4 | Neutral |
| testneg_43 | 0 | 0 | -0.274465 | -3 | 3.691952 | -3 | 0.04699 | 0.08149 | 0.04694 | 0.44682 | 13.93265 | 0.241550398 | 0.2 | Neutral |
| testneg_44 | -0.15 | 0.28 | 2.759325 | -3 | -0.470538 | -3 | 0.56416 | 0.12788 | 0.12957 | 0.09 | 15.72543 | 0.06 | 0 | Neutral |
| testneg_45 | 0.07 | 0.38 | -2.09754 | -3 | 3.759941 | -3 | 0.11956 | 0.3237 | 0.46058 | 0.12903 | 18.02178 | 0.2229675 | 0 | Neutral |
| testneg_46 | 0.04 | -0.23 | 8.054454 | -3 | 2.843639 | -3 | 0.40632 | 0.1094 | 0.19898 | 0.12788 | 16.43865 | 0.479878049 | 46 | Neutral |
| testneg_47 | 0 | 0 | 1.39998 | -3 | -0.202498 | -3 | 0.161 | 0.11597 | 0.30789 | 0.15135 | 17.21847 | 0.015378571 | 0 | Neutral |
| testneg_48 | -0.07 | -0.38 | -1.690706 | -3 | 1.915589 | -3 | 0.08779 | 0.1777 | 0.15406 | 0.1454 | 18.88889 | 0.150184641 | 0 | Neutral |
| testneg_49 | 0.05 | -0.81 | 4.217475 | -3 | 6.84193 | -3 | 0.1551 | 0.09276 | 0.2734 | 0.2578 | 14.02244 | 0.6901125 | 98.2 | Disease_causing |
| testneg_50 | 0 | 0 | 0.28875 | -3 | -1.730578 | -3 | 0.40179 | 0.12112 | 0.24949 | 0.29709 | 11.63569 | 0.030729443 | 0 | Neutral |
| testneg_51 | -0.06 | -0.18 | 0.867502 | -3 | -0.001612 | -3 | 0.71511 | 0.34803 | 0.34205 | 0.11195 | 10.83333 | 0.03 | 0 | Neutral |
| testneg_52 | 0 | 0 | 2.123289 | -3 | 6.559099 | -3 | 0.19898 | 0.12788 | 0.12857 | 0.09376 | 14.16667 | 0.17035401 | 0.2 | Neutral |
| testneg_53 | 0.02 | 0.45 | 5.194939 | -3 | 0.64921 | -3 | 0.03631 | 0.80564 | 0.17016 | 0.12748 | 20.38798 | 0.34 | 3.4 | Neutral |
| testneg_54 | 0 | 0 | -0.299858 | -3 | 3.523914 | -3 | 0.29284 | 0.11262 | 0.53698 | 0.10653 | 20.33859 | 0.035 | 0 | Neutral |
| testneg_55 | 0 | 0 | 0.532295 | -3 | 1.65767 | 1.65767 | 0.755 | 0.12788 | 0.89992 | 0.10231 | 10.83333 | 0.02 | 0 | Neutral |
| testneg_56 | 0 | 0 | 0.15975 | -3 | 2.390877 | -3 | 0.19898 | 0.16603 | 0.43691 | 0.12794 | 18.88889 | 0.037773585 | 0 | Neutral |
| testneg_57 | -0.09 | 0.3 | 0.631708 | -3 | 3.753987 | -3 | 0.34546 | 0.0932 | 0.06736 | 0.06344 | 18.33333 | 0.106694444 | 0 | Neutral |
| testneg_58 | 0 | 0 | 1.687239 | -3 | 3.300854 | -3 | 0.16134 | 0.1847 | 0.43977 | 0.17284 | 9.31758 | 0.253769046 | 1.2 | Neutral |
| testneg_59 | 0 | 0 | -0.32829 | -3 | 0.046064 | -3 | 0.18509 | 0.12831 | 0.2543 | 0.15588 | 11.95767 | 0.030158275 | 0 | Neutral |
| testneg_60 | -0.15 | 1.06 | 1.213874 | -3 | 3.451445 | -3 | 0.04543 | 0.28103 | 0.49064 | 0.12294 | 10.41667 | 0.399333333 | 11.2 | Neutral |
| testneg_61 | 0 | 0 | -0.771216 | -3 | 3.115437 | -3 | 0.37986 | 0.14229 | 0.20666 | 0.13449 | 9.66667 | 0.165759958 | 0.2 | Neutral |
| testneg_62 | 0.14 | -0.37 | 1.92089 | -3 | -0.912757 | -3 | 0.10556 | 0.28423 | 0.07211 | 0.12934 | 25.2381 | 0.060442005 | 0 | Neutral |
| testneg_63 | 0 | 0.32 | 1.693254 | -3 | 0.586242 | -3 | 0.18047 | 0.2356 | 0.559 | 0.2286 | 14.26037 | 0.096509798 | 0 | Neutral |
| testneg_64 | 0 | 0 | 2.157422 | -3 | 6.118303 | -3 | 0.85508 | 0.12634 | 0.8376 | 0.24333 | 10.41667 | 0.248078588 | 0.6 | Neutral |
| testneg_65 | 0.19 | -0.25 | 7.226253 | -3 | 0.722185 | -3 | 0.83798 | 0.14433 | 0.18866 | 0.12788 | 31.08131 | 0.319375 | 3.6 | Neutral |
| testneg_66 | 0.04 | -0.23 | 3.009853 | -3 | 1.957496 | -3 | 0.50741 | 0.17915 | 0.06395 | 0.0759 | 29.3644 | 0.08 | 0 | Neutral |
| testneg_67 | 0.01 | 0.8 | 7.175154 | -3 | 4.133728 | -3 | 0.57188 | 0.34038 | 0.13794 | 0.1141 | 10.22898 | 0.738712121 | 99.4 | Disease_causing |
| testneg_68 | -0.1 | -1.9 | -0.947091 | -3 | -0.701339 | -3 | 0.19898 | 0.12788 | 0.46735 | 0.10674 | 34.29705 | 0.05 | 0 | Neutral |
| testneg_69 | 0 | 0 | -0.68826 | -3 | 1.068497 | -3 | 0.32076 | 0.14229 | 0.51278 | 0.35859 | 4.72222 | 0.045858325 | 0 | Neutral |
| testneg_70 | 0 | 0 | 0.270932 | -3 | 5.315177 | -3 | 0.16404 | 0.09378 | 0.12091 | 0.13585 | 17.33727 | 0.217081391 | 0.4 | Neutral |
| testneg_71 | 0.00037879 | 0.07787879 | 1.536103 | -3 | 4.170871 | -3 | 0.17786 | 0.15314 | 0.19898 | 0.12788 | 18.22929 | 0.123868298 | 0 | Neutral |
| testneg_72 | -0.02 | -0.45 | 5.051631 | -3 | -0.435286 | -3 | 0.08531 | 0.12788 | 0.18341 | 0.13901 | 20.18807 | 0.188932692 | 0.2 | Neutral |
| testneg_73 | -0.07 | -0.38 | 3.483266 | -3 | 1.417365 | 1.417365 | 0.25856 | 0.17179 | 0.61184 | 0.55597 | 12.6082 | 0.29979451 | 2.2 | Neutral |
| testneg_74 | 0.06 | 0.18 | 3.600146 | -3 | -0.823621 | -0.823621 | 0.3954 | 0.17605 | 0.27382 | 0.16898 | 16.8018 | 0.207201378 | 0 | Neutral |
| testneg_75 | -0.01 | -0.8 | 0.738714 | -3 | 4.947111 | -3 | 0.17158 | 0.1149 | 0.12036 | 0.0873 | 20.57907 | 0.07 | 0 | Neutral |
| testneg_76 | 0 | 0 | 1.673515 | -3 | 1.462347 | -3 | 0.56554 | 0.76978 | 0.11972 | 0.09438 | 12.87785 | 0.04 | 0 | Neutral |
| testneg_77 | 0 | 0 | 0.269653 | -0.735798 | 1.379202 | -3 | 0.7219 | 0.41509 | 0.10566 | 0.12788 | 10.35825 | 0.02 | 0 | Neutral |
| testneg_78 | 0.06 | -0.57 | 3.37237 | -3 | 1.917158 | 1.917158 | 0.11447 | 0.12378 | 0.11618 | 0.20784 | 18.33333 | 0.27 | 1 | Neutral |
| testneg_79 | -0.01 | -0.79 | 2.403241 | -1.897594 | 2.243237 | -3 | 0.19898 | 0.12788 | 0.43977 | 0.17284 | 13.51351 | 0.111547619 | 0 | Neutral |
| testneg_80 | -0.07 | -0.38 | 4.05433 | -3 | 2.32114 | -3 | 0.45415 | 0.12971 | 0.13331 | 0.13638 | 10.83333 | 0.227615025 | 0.4 | Neutral |
| testneg_81 | 0 | 0 | 1.411308 | -3 | -1.122085 | -3 | 0.73253 | 0.09747 | 0.0774 | 0.1702 | 44.46023 | 0.100486111 | 0 | Neutral |
| testneg_82 | -0.07 | -0.38 | 0.879664 | -3 | 3.540511 | 2.589938 | 0.33163 | 0.26601 | 0.16616 | 0.11073 | 12.29167 | 0.176795793 | 0 | Neutral |
| testneg_83 | -0.07 | -0.38 | 1.894127 | -3 | 5.586363 | -3 | 0.62004 | 0.10842 | 0.65909 | 0.31908 | 12.66667 | 0.372361111 | 10.2 | Neutral |
| testneg_84 | -0.02 | 0.63 | 3.191277 | -3 | -0.749376 | -0.749376 | 0.37074 | 0.11532 | 0.51437 | 0.17173 | 26.50889 | 0.140361729 | 0 | Neutral |
| testneg_85 | 0.18 | -1.05 | 5.390189 | -3 | 5.342136 | -3 | 0.15041 | 0.43468 | 0.1496 | 0.16692 | 17.21847 | 0.919755516 | 100 | Disease_causing |
| testneg_86 | 0.12 | -1.14 | 3.586622 | -3 | 3.337227 | -3 | 0.25555 | 0.12094 | 0.29247 | 0.20085 | 10.83333 | 0.503311688 | 55.4 | Disease_causing |
| testneg_87 | -0.15 | -0.04 | -0.451132 | -3 | 3.652963 | -3 | 0.12048 | 0.2381 | 0.82269 | 0.15732 | 26.62774 | 0.461044372 | 39 | Neutral |
| testneg_88 | -0.22 | 2.66 | 0.984298 | -3 | 0.21945 | 0.21945 | 0.34406 | 0.09894 | 0.81566 | 0.22678 | 21.96727 | 0.198650794 | 0 | Neutral |
| testneg_89 | 0.08 | 0.03 | 0.231468 | -3 | 1.433381 | -3 | 0.21811 | 0.18489 | 0.12543 | 0.12788 | 27.91246 | 0.06 | 0 | Neutral |
| testneg_90 | 0 | 0 | 0.387542 | -3 | 0.878873 | -3 | 0.76693 | 0.19444 | 0.62197 | 0.12788 | 10 | 0.01 | 0 | Neutral |
| testneg_91 | 0 | 0 | 1.372857 | -3 | 0.853858 | -3 | 0.14378 | 0.41915 | 0.35513 | 0.21966 | 13.08333 | 0.095372517 | 0 | Neutral |
| testneg_92 | 0 | 0 | 1.144569 | -3 | 1.957825 | -3 | 0.73985 | 0.13844 | 0.02518 | 0.07165 | 12.72727 | 0.010769231 | 0 | Neutral |
| testneg_93 | 0 | 0 | 0.389761 | -3 | 0.423127 | -3 | 0.73601 | 0.26791 | 0.25643 | 0.12788 | 20.32634 | 0.010784445 | 0 | Neutral |
| testneg_94 | 0.14 | -0.69 | 0.584676 | -3 | 0.323125 | -3 | 0.31701 | 0.09986 | 0.07258 | 0.06248 | 20.26596 | 0.015 | 0 | Neutral |
| testneg_95 | 0.01 | 0.8 | 2.326649 | -3 | 0.265271 | -3 | 0.20914 | 0.10089 | 0.45405 | 0.10896 | 23.75274 | 0.010129958 | 0 | Neutral |
| testneg_96 | -0.07 | -0.38 | 3.41492 | -3 | 1.324006 | -3 | 0.22162 | 0.11997 | 0.03441 | 0.10663 | 18.35253 | 0.03 | 0 | Neutral |
| testneg_97 | 0.08 | 0.03 | -1.290562 | -3 | -0.707182 | -3 | 0.19898 | 0.12895 | 0.06789 | 0.09707 | 10.22898 | 0.020158521 | 0 | Neutral |
| testneg_98 | 0.03 | 0.1 | -0.942639 | -3 | 3.566632 | -3 | 0.05996 | 0.12788 | 0.9581 | 0.12788 | 60.20998 | 0.110407925 | 0 | Neutral |
| testneg_99 | 0 | 0.32 | 2.93612 | -3 | 2.383861 | 1.519228 | 0.05657 | 0.14812 | 0.12322 | 0.1246 | 21.57457 | 0.16 | 0 | Neutral |
| testneg_100 | -0.03 | -0.1 | 2.260957 | -3 | 5.167963 | -3 | 0.36054 | 0.12788 | 0.36541 | 0.27749 | 15.72733 | 0.405459293 | 22.6 | Neutral |
