## Supplementary material for "Predicting disease-causing variant combinations": Dataset S3

| Combination_ID | Flex1 | Hydr1 | CADD1 | CADD2 | CADD3 | CADD4 | HI_A | RecA | HI_B | RecB | Biol_Dist | Classification_score | Support_score | Predicted_class |
|---|---|---|---|---|---|---|---|---|---|---|---|---|---|---|
| testneg_1 | 0.06 | 0.18 | 1.976668 | -3 | 4.068145 | -3 | 0.13001 | 0.13176 | 0.0406 | 0.12788 | 18.88889 | 0.125 | 0 | Neutral |
| testneg_2 | -0.01 | 0.57 | 1.80646 | -3 | 0.931031 | -3 | 0.32111 | 0.12788 | 0.23939 | 0.25284 | 24.729 | 0.075119048 | 0 | Neutral |
| testneg_3 | 0.00055944 | 0.00363636 | 1.360315 | -3 | 5.745573 | -3 | 0.29482 | 0.15726 | 0.08027 | 0.12788 | 12.70833 | 0.141683261 | 0 | Neutral |
| testneg_4 | 0.07 | 0.33 | 5.070028 | -3 | 1.614385 | -3 | 0.23005 | 0.37216 | 0.1336 | 0.15669 | 6.2037 | 0.464910041 | 41.2 | Neutral |
| testneg_5 | 0 | 0 | 0.017266 | -3 | 6.011632 | -3 | 0.1228 | 0.10482 | 0.16968 | 0.12788 | 18.00969 | 0.090333704 | 0 | Neutral |
| testneg_6 | 0 | 0 | -0.43649 | -3 | -0.133106 | -0.133106 | 0.71968 | 0.54266 | 0.07096 | 0.09965 | 12.31545 | 0.032946429 | 0 | Neutral |
| testneg_7 | 0.14 | -1.01 | 3.175447 | -3 | 4.581463 | -3 | 0.18925 | 0.32479 | 0.16741 | 0.10762 | 7.00579 | 0.362488095 | 9 | Neutral |
| testneg_8 | 0.03 | 0.1 | -0.895222 | -3 | 6.710008 | -3 | 0.84468 | 0.55945 | 0.10921 | 0.34026 | 20.3131 | 0.57922619 | 82.6 | Disease_causing |
| testneg_9 | 0 | 0 | 1.075324 | 1.075324 | 6.497557 | -3 | 0.49934 | 0.11069 | 0.18706 | 0.10259 | 17.95745 | 0.19 | 0 | Neutral |
| testneg_10 | 0 | -0.32 | 0.363708 | -3 | 0.141395 | -3 | 0.20231 | 0.12788 | 0.07961 | 0.11474 | 23.47349 | 0.01 | 0 | Neutral |
| testneg_11 | 0.07 | -1.12 | 4.283389 | -3 | -0.659954 | -3 | 0.19989 | 0.16777 | 0.19898 | 0.12788 | 17.15067 | 0.08 | 0 | Neutral |
| testneg_12 | 0.05 | -0.81 | 3.968796 | -3 | -1.617473 | -3 | 0.35475 | 0.12788 | 0.19898 | 0.12788 | 17.77778 | 0.025 | 0 | Neutral |
| testneg_13 | 0.21 | 0.15 | 2.151895 | -3 | 1.022243 | -3 | 0.11791 | 0.39607 | 0.13099 | 0.17453 | 21.41309 | 0.265357143 | 0.2 | Neutral |
| testneg_14 | 0.21 | 0.15 | 3.072922 | -3 | 1.249863 | -3 | 0.17301 | 0.14089 | 0.19898 | 0.1054 | 10.41667 | 0.06 | 0 | Neutral |
| testneg_15 | 0.19 | 0.51 | 2.754442 | -3 | 1.697009 | -3 | 0.48825 | 0.10951 | 0.52771 | 0.10974 | 25.50457 | 0.05 | 0 | Neutral |
| testneg_16 | 0.14 | -1.01 | 7.140458 | -3 | 5.126166 | -3 | 0.24823 | 0.12788 | 0.65857 | 0.26294 | 19.10253 | 0.779791667 | 100 | Disease_causing |
| testneg_17 | 0.22 | -2.66 | 3.35038 | -3 | 1.30662 | -3 | 0.73985 | 0.13844 | 0.40264 | 0.12788 | 10.83333 | 0.13 | 0 | Neutral |
| testneg_18 | 0.09 | -0.3 | 7.198996 | -3 | -0.454393 | -1.150939 | 0.12463 | 0.12788 | 0.03443 | 0.1032 | 10.69344 | 0.25 | 1.8 | Neutral |
| testneg_19 | -0.02 | 0.36 | 5.302115 | -3 | 5.860091 | -3 | 0.26972 | 0.12484 | 0.75481 | 0.12788 | 22.71762 | 0.44 | 32.4 | Neutral |
| testneg_20 | 0.02 | 0.13 | 5.790455 | -3 | 3.399462 | -3 | 0.10895 | 0.10442 | 0.05472 | 0.12788 | 21.42455 | 0.26 | 2 | Neutral |
| testneg_21 | -0.07 | -0.38 | -0.007494 | -3 | 0.11497 | -3 | 0.46919 | 0.60081 | 0.16355 | 0.20781 | 13.75 | 0.16 | 0 | Neutral |
| testneg_22 | 0.03 | -1.03 | 1.493352 | -3 | 1.917158 | 1.917158 | 0.17364 | 0.18111 | 0.11618 | 0.20784 | 11.875 | 0.187915522 | 0 | Neutral |
| testneg_23 | 0.14 | -0.37 | 2.102067 | -3 | 0.275543 | 0.275543 | 0.95557 | 0.38833 | 0.4738 | 0.14744 | 6.00877 | 0.124343407 | 0 | Neutral |
| testneg_24 | 0.14 | -0.69 | 4.370948 | -3 | 1.830118 | -3 | 0.87268 | 0.21837 | 0.55367 | 0.57993 | 11.25 | 0.39 | 17.6 | Neutral |
| testneg_25 | 0 | 0 | -0.044046 | -3 | 0.331991 | -3 | 0.47544 | 0.17658 | 0.11571 | 0.17324 | 11.875 | 0.080414621 | 0 | Neutral |
| testneg_26 | 0 | -0.32 | -0.590938 | -3 | 0.219522 | -3 | 0.06513 | 0.12353 | 0.19898 | 0.09533 | 58.95202 | 0.06 | 0 | Neutral |
| testneg_27 | -0.06 | -0.18 | 1.759866 | -3 | 3.717179 | -3 | 0.7249 | 0.41559 | 0.15808 | 0.14548 | 5.71262 | 0.350824183 | 8 | Neutral |
| testneg_28 | 0 | 0 | 2.53141 | 2.53141 | 3.085424 | -3 | 0.1123 | 0.1568 | 0.43282 | 0.10804 | 22.92553 | 0.24 | 0.6 | Neutral |
| testneg_29 | 0 | 0 | 1.799258 | -3 | 2.780842 | -3 | 0.204 | 0.15424 | 0.80034 | 0.10518 | 10.22898 | 0.042874125 | 0 | Neutral |
| testneg_30 | 0 | 0 | 0.309568 | -3 | 4.232024 | -3 | 0.54724 | 0.12788 | 0.47019 | 0.12788 | 18.33333 | 0.051030303 | 0 | Neutral |
| testneg_31 | 0 | -0.01 | 1.028355 | -3 | 2.550986 | -3 | 0.87835 | 0.13522 | 0.08408 | 0.12788 | 12.29167 | 0.03 | 0 | Neutral |
| testneg_32 | -0.21 | -0.15 | 2.657036 | -3 | -0.238112 | -3 | 0.12408 | 0.08828 | 0.0611 | 0.12135 | 5 | 0.07 | 0 | Neutral |
| testneg_33 | 0 | 0 | 0.71683 | -3 | 1.329031 | -3 | 0.42583 | 0.10901 | 0.22489 | 0.16646 | 19.09347 | 0.039473684 | 0 | Neutral |
| testneg_34 | 0 | -0.01 | 7.652567 | 7.652567 | 0.390032 | -3 | 0.73253 | 0.09747 | 0.15689 | 0.28307 | 18.6768 | 0.769657029 | 99.4 | Disease_causing |
| testneg_35 | -0.01 | -0.8 | 0.738714 | -3 | -1.655921 | -3 | 0.17158 | 0.1149 | 0.19898 | 0.12788 | 18.61427 | 0.01 | 0 | Neutral |
| testneg_36 | 0.07 | 0.7 | -2.193757 | -3 | 2.597499 | -3 | 0.18795 | 0.09487 | 0.07864 | 0.12788 | 17.57227 | 0.053333333 | 0 | Neutral |
| testneg_37 | -0.08 | -0.03 | 0.950149 | -3 | 3.05843 | 3.05843 | 0.31318 | 0.11262 | 0.74136 | 0.1049 | 18.33333 | 0.14 | 0 | Neutral |
| testneg_38 | 0.02 | 0.45 | 1.808811 | -3 | 1.042833 | -3 | 0.18485 | 0.40044 | 0.19898 | 0.12788 | 10 | 0.04019638 | 0 | Neutral |
| testneg_39 | 0.1 | 1.9 | 3.737058 | -3 | 1.329031 | -3 | 0.28191 | 0.20693 | 0.22489 | 0.16646 | 10.41667 | 0.34 | 5 | Neutral |
| testneg_40 | -0.002 | 0.19418182 | 10.570212 | -3 | 2.364247 | 2.364247 | 0.09577 | 0.2436 | 0.09532 | 0.35459 | 16.668 | 0.76 | 100 | Disease_causing |
| testneg_41 | -0.07 | -0.38 | 0.649388 | -3 | 3.635653 | -3 | 0.30951 | 0.64738 | 0.11501 | 0.13748 | 12.72648 | 0.414250264 | 22 | Neutral |
| testneg_42 | -0.06 | -0.18 | 2.632596 | -3 | -0.105139 | -3 | 0.9662 | 0.28104 | 0.08489 | 0.12788 | 5.75579 | 0.04 | 0 | Neutral |
| testneg_43 | 0.17 | 0.38 | 1.596978 | -3 | 1.518871 | -3 | 0.15581 | 0.10782 | 0.52953 | 0.10241 | 29.58002 | 0.02 | 0 | Neutral |
| testneg_44 | 0 | 0 | -0.027704 | -3 | 0.098075 | -3 | 0.45299 | 0.16306 | 0.44329 | 0.12788 | 10.41667 | 0.02 | 0 | Neutral |
| testneg_45 | 0.02 | -0.36 | 4.167444 | -3 | 0.329621 | 0.329621 | 0.87806 | 0.38966 | 0.4129 | 0.11172 | 18.33333 | 0.1 | 0 | Neutral |
| testneg_46 | 0 | 0 | -0.153757 | -3 | 0.573225 | 0.573225 | 0.23027 | 0.37171 | 0.73947 | 0.2058 | 30.95732 | 0.16 | 0 | Neutral |
| testneg_47 | 0 | 0 | 0.371766 | -3 | 6.099347 | -3 | 0.3095 | 0.13502 | 0.12253 | 0.10814 | 9.36366 | 0.103687888 | 0 | Neutral |
| testneg_48 | -0.06 | -0.18 | 3.18158 | -3 | -1.197202 | -1.197202 | 0.23439 | 0.09253 | 0.08688 | 0.08809 | 23.69465 | 0.04 | 0 | Neutral |
| testneg_49 | 0.04 | -0.23 | 4.017255 | -3 | 0.944088 | -3 | 0.19898 | 0.16603 | 0.71025 | 0.1632 | 9.29788 | 0.230357143 | 0.2 | Neutral |
| testneg_50 | 0 | -0.32 | 2.645906 | -3 | 5.007683 | -3 | 0.20914 | 0.10089 | 0.07415 | 0.12788 | 13.51351 | 0.135 | 0 | Neutral |
| testneg_51 | 0 | 0 | 1.050504 | -3 | 1.999648 | 1.999648 | 0.46792 | 0.11262 | 0.12626 | 0.10279 | 23.08943 | 0.04 | 0 | Neutral |
| testneg_52 | -0.1 | -1.9 | -0.234016 | -0.234016 | 3.557588 | -3 | 0.04436 | 0.12788 | 0.80034 | 0.10518 | 28.18275 | 0.141714286 | 0 | Neutral |
| testneg_53 | 0.03 | 0.12 | 2.210824 | -3 | 1.996983 | 1.996983 | 0.38171 | 0.15977 | 0.09553 | 0.11637 | 23.60682 | 0.063333333 | 0 | Neutral |
| testneg_54 | -0.02 | -0.41 | 0.728673 | -3 | 0.945458 | 0.230086 | 0.50562 | 0.2196 | 0.19898 | 0.08275 | 12.07839 | 0.031666667 | 0 | Neutral |
| testneg_55 | 0.05 | -0.81 | -0.646746 | -3 | 1.996725 | -3 | 0.12073 | 0.10969 | 0.76942 | 0.13267 | 19.8494 | 0.06 | 0 | Neutral |
| testneg_56 | 0 | 0 | 0.231295 | -3 | 2.310369 | -3 | 0.19898 | 0.12788 | 0.54895 | 0.15082 | 33.40179 | 0.066898952 | 0 | Neutral |
| testneg_57 | 0.07 | -0.31 | 6.107142 | -3 | -0.843404 | -3 | 0.12554 | 0.10854 | 0.11587 | 0.12001 | 10.83333 | 0.14 | 0 | Neutral |
| testneg_58 | 0.15 | -0.28 | 0.552009 | 0.552009 | 3.751338 | -3 | 0.53172 | 0.12788 | 0.12933 | 0.12788 | 17.21847 | 0.130954545 | 0 | Neutral |
| testneg_59 | -0.14 | -0.54 | 1.697775 | -3 | 1.682022 | -3 | 0.39774 | 0.10813 | 0.06895 | 0.06354 | 16.85532 | 0.020218183 | 0 | Neutral |
| testneg_60 | 0.09 | -0.25 | -0.159404 | -3 | 3.413304 | -3 | 0.95319 | 0.20297 | 0.31626 | 0.12936 | 4.22222 | 0.14 | 0 | Neutral |
| testneg_61 | -0.07 | -0.01 | 3.375538 | -3 | 2.013858 | -3 | 0.23439 | 0.09253 | 0.24536 | 0.38555 | 8.97898 | 0.238656463 | 0.4 | Neutral |
| testneg_62 | 0 | -0.01 | 7.652567 | -3 | 1.524896 | -3 | 0.73253 | 0.09747 | 0.15526 | 0.08511 | 19.09347 | 0.3885 | 15.2 | Neutral |
| testneg_63 | 0 | -0.32 | -0.565398 | -0.565398 | 1.222548 | 1.222548 | 0.19898 | 0.12788 | 0.06705 | 0.12788 | 31.8712 | 0.020281332 | 0 | Neutral |
| testneg_64 | -0.03 | -0.12 | 0.226482 | -3 | 1.340083 | -3 | 0.34279 | 0.09378 | 0.28227 | 0.10211 | 10.83333 | 0.010206094 | 0 | Neutral |
| testneg_65 | 0.03 | 0.12 | 0.90434 | -3 | 1.015573 | -3 | 0.53424 | 0.08484 | 0.16974 | 0.80184 | 10 | 0.08045098 | 0 | Neutral |
| testneg_66 | -0.02 | -0.45 | 3.106183 | -3 | 0.281314 | -3 | 0.99267 | 0.1069 | 0.09048 | 0.62605 | 11.21556 | 0.17 | 0 | Neutral |
| testneg_67 | -0.14 | -0.54 | 3.467861 | 0.55511 | -0.915676 | -3 | 0.37293 | 0.25159 | 0.05767 | 0.07325 | 12.35574 | 0.13 | 0 | Neutral |
| testneg_68 | -0.09 | 0.3 | 0.850367 | -3 | 2.97091 | -3 | 0.29024 | 0.1472 | 0.0961 | 0.09786 | 34.16556 | 0.072880952 | 0 | Neutral |
| testneg_69 | -0.03 | -0.1 | 2.834756 | -3 | 0.191856 | -3 | 0.13447 | 0.12788 | 0.13749 | 0.17824 | 17.21847 | 0.121966846 | 0 | Neutral |
| testneg_70 | -0.03 | -0.1 | 3.202429 | -3 | 6.472745 | 5.290372 | 0.39964 | 0.22293 | 0.49276 | 0.10648 | 17.77778 | 0.595370215 | 88 | Disease_causing |
| testneg_71 | 0.14 | -0.37 | 5.019921 | -3 | 0.341374 | -3 | 0.41953 | 0.44852 | 0.10646 | 0.1069 | 19.2812 | 0.160230082 | 0 | Neutral |
| testneg_72 | 0.06 | -0.57 | 1.830175 | -3 | 7.49852 | -3 | 0.11351 | 0.10667 | 0.3112 | 0.33703 | 10.70652 | 0.530579139 | 67.4 | Disease_causing |
| testneg_73 | 0.02618182 | -0.2483636 | 11.046933 | -3 | -0.429687 | -3 | 0.4113 | 0.2303 | 0.06296 | 0.09823 | 29.86111 | 0.42 | 21.6 | Neutral |
| testneg_74 | 0 | 0 | -0.12637 | -3 | -0.445161 | -0.445161 | 0.22761 | 0.14229 | 0.09728 | 0.12788 | 36.37944 | 0.03 | 0 | Neutral |
| testneg_75 | 0.17 | 0.38 | 3.132136 | -3 | 2.341037 | 2.341037 | 0.0105 | 0.12788 | 0.95535 | 0.12788 | 20.11942 | 0.170057471 | 0 | Neutral |
| testneg_76 | 0 | 0 | 0.957081 | -3 | 0.126173 | -3 | 0.12259 | 0.1041 | 0.07652 | 0.30451 | 11.93907 | 0.05 | 0 | Neutral |
| testneg_77 | 0.02 | -0.68 | 4.168346 | -3 | 0.635125 | -3 | 0.30042 | 0.13517 | 0.06187 | 0.36607 | 29.16667 | 0.258688187 | 0.8 | Neutral |
| testneg_78 | -0.03 | -0.1 | 4.494734 | -3 | 4.841698 | -3 | 0.51675 | 0.11055 | 0.29356 | 0.46298 | 27.14503 | 0.577269778 | 81.6 | Disease_causing |
| testneg_79 | 0.04 | -0.23 | 4.454617 | -3 | 0.655763 | 0.655763 | 0.7639 | 0.10408 | 0.09087 | 0.12788 | 12.12963 | 0.04 | 0 | Neutral |
| testneg_80 | -0.08 | -0.03 | 1.052459 | -3 | 1.848117 | -3 | 0.67331 | 0.10118 | 0.10249 | 0.1399 | 18.33333 | 0.080055556 | 0 | Neutral |
| testneg_81 | 0.03 | 0.12 | 4.71937 | -3 | 1.73624 | -3 | 0.16965 | 0.25991 | 0.74488 | 0.12788 | 24.1857 | 0.14 | 0 | Neutral |
| testneg_82 | -0.06 | -0.18 | 0.947918 | -3 | 1.21293 | -3 | 0.25236 | 0.11809 | 0.07189 | 0.11325 | 12.81915 | 0.01 | 0 | Neutral |
| testneg_83 | 0.03 | -1.03 | 4.595163 | -3 | -0.01145 | -0.01145 | 0.26916 | 0.13285 | 0.78462 | 0.12258 | 23.24074 | 0.078819444 | 0 | Neutral |
| testneg_84 | 0 | 0 | 0.412754 | -3 | 4.959501 | -3 | 0.9327 | 0.12788 | 0.14301 | 0.12788 | 8.97898 | 0.050467391 | 0 | Neutral |
| testneg_85 | 0.01 | 0.79 | 2.557398 | -3 | -0.474175 | -3 | 0.961 | 0.25991 | 0.3414 | 0.13575 | 17.46678 | 0.124543655 | 0 | Neutral |
| testneg_86 | -0.05 | 0.44 | 5.838894 | -3 | 1.102485 | -3 | 0.10917 | 0.12788 | 0.25731 | 0.12662 | 34.72222 | 0.172132353 | 0 | Neutral |
| testneg_87 | 0.21 | 0.15 | 1.305251 | -3 | 1.056 | 1.056 | 0.06304 | 0.12788 | 0.21153 | 0.11357 | 26.34995 | 0.04 | 0 | Neutral |
| testneg_88 | 0 | 0 | -0.692897 | -3 | 6.106898 | -3 | 0.57953 | 0.15513 | 0.15989 | 0.12788 | 19.41377 | 0.151415776 | 0 | Neutral |
| testneg_89 | 0 | 0 | 3.603309 | -3 | 3.916127 | -3 | 0.26192 | 0.15452 | 0.08876 | 0.09037 | 11.25 | 0.2 | 0 | Neutral |
| testneg_90 | 0.18 | 0.16 | -0.110087 | -3 | 6.510328 | -3 | 0.82126 | 0.17937 | 0.53216 | 0.57092 | 4.44444 | 0.51922459 | 61 | Disease_causing |
| testneg_91 | 0.05 | -0.81 | 6.33079 | -3 | 7.563765 | -1.914326 | 0.31906 | 0.1705 | 0.2124 | 0.17288 | 19.83528 | 0.93 | 100 | Disease_causing |
| testneg_92 | -0.0087273 | -0.0077576 | 1.035348 | -3 | -0.472766 | -3 | 0.87268 | 0.21837 | 0.15119 | 0.08823 | 26.34995 | 0.023333333 | 0 | Neutral |
| testneg_93 | 0.02 | -0.36 | 0.993581 | -3 | -0.691124 | -3 | 0.04508 | 0.12788 | 0.65284 | 0.2592 | 18.88889 | 0.12 | 0 | Neutral |
| testneg_94 | -0.17 | -0.38 | 5.724603 | -3 | -0.811455 | -0.811455 | 0.20918 | 0.10825 | 0.10638 | 0.66554 | 32.89375 | 0.372222222 | 12.2 | Neutral |
| testneg_95 | 0 | 0 | 1.012812 | -3 | -0.976851 | -3 | 0.53313 | 0.14072 | 0.27197 | 0.12346 | 18.88889 | 0.010267214 | 0 | Neutral |
| testneg_96 | 0 | 0 | 0.984412 | -3 | 6.53007 | -3 | 0.34931 | 0.11906 | 0.13185 | 0.51629 | 15.39358 | 0.353157897 | 11.2 | Neutral |
| testneg_97 | 0 | 0 | -0.134529 | -3 | 0.096105 | -3 | 0.46919 | 0.60081 | 0.20448 | 0.12788 | 25.78212 | 0.06 | 0 | Neutral |
| testneg_98 | 0 | 0 | 0.802919 | 0.802919 | 0.099399 | -3 | 0.12042 | 0.11067 | 0.48817 | 0.10601 | 18.6768 | 0.03 | 0 | Neutral |
| testneg_99 | 0.09 | -2.17 | 6.188658 | -3 | 0.518309 | -3 | 0.13203 | 0.12092 | 0.06331 | 0.2396 | 12.78881 | 0.4 | 20.2 | Neutral |
| testneg_100 | 0 | 0 | -0.153917 | -3 | 2.71786 | -3 | 0.1589 | 0.17874 | 0.04912 | 0.08574 | 18.88889 | 0.055 | 0 | Neutral |
| testneg_101 | 0.03 | 0.12 | -1.295182 | -3 | 5.481047 | -0.323214 | 0.26913 | 0.1087 | 0.15432 | 0.095 | 20.27778 | 0.102055834 | 0 | Neutral |
| testneg_102 | 0.04 | -0.23 | 1.86404 | -3 | -1.158834 | -3 | 0.21843 | 0.85214 | 0.12739 | 0.12788 | 21.64934 | 0.10059419 | 0 | Neutral |
| testneg_103 | 0.06 | -0.57 | 2.442942 | -3 | -0.045315 | -3 | 0.09048 | 0.62605 | 0.10052 | 0.10074 | 10.93937 | 0.09 | 0 | Neutral |
| testneg_104 | -0.18 | -0.16 | 1.260137 | -3 | 2.966798 | -3 | 0.1214 | 0.0899 | 0.5184 | 0.11889 | 19.44444 | 0.1 | 0 | Neutral |
| testneg_105 | 0.03 | -1.03 | 6.284786 | -3 | 7.462634 | -3 | 0.1675 | 0.09413 | 0.0571 | 0.10747 | 21.10568 | 0.56 | 79.4 | Disease_causing |
| testneg_106 | 0.05 | -0.81 | 3.823198 | -3 | -0.272804 | -0.272804 | 0.6992 | 0.1804 | 0.17839 | 0.12009 | 11.82515 | 0.07 | 0 | Neutral |
| testneg_107 | 0 | 0 | 0.617174 | -3 | 3.775158 | -3 | 0.28416 | 0.27805 | 0.07297 | 0.12788 | 19.9179 | 0.12 | 0 | Neutral |

| | | | | | | | | | | | | | | |
|---|---|---|---|---|---|---|---|---|---|---|---|---|---|---|
| testneg_108 | 0 | 0 | -0.101683 | -3 | 2.531153 | -3 | 0.72657 | 0.11262 | 0.36833 | 0.10433 | 7.07344 | 0.01685904 | 0 | Neutral |
| testneg_109 | 0 | 0 | -1.230837 | -1.230837 | 1.902366 | 1.902366 | 0.14821 | 0.11262 | 0.12231 | 0.23155 | 10.22898 | 0.090894118 | 0 | Neutral |
| testneg_110 | 0.06 | -0.57 | -2.190906 | -3 | 3.641057 | -3 | 0.03137 | 0.12788 | 0.055 | 0.20858 | 29.00394 | 0.29169287 | 1.4 | Neutral |
| testneg_111 | -0.1 | -1.9 | -1.487847 | -3 | 6.497505 | -3 | 0.06728 | 0.20664 | 0.13702 | 0.12788 | 25.02318 | 0.410962963 | 20.8 | Neutral |
| testneg_112 | 0.19 | 0.51 | -0.093394 | -0.093394 | 11.796446 | -3 | 0.34587 | 0.29822 | 0.13626 | 0.12788 | 43.2869 | 0.521240602 | 62.6 | Disease_causing |
| testneg_113 | -0.03 | -0.12 | 1.215025 | -3 | 2.478795 | 1.782913 | 0.09028 | 0.10643 | 0.1327 | 0.09885 | 17.91076 | 0.05 | 0 | Neutral |
| testneg_114 | 0.18 | -1.05 | 2.305068 | -3 | 2.441441 | -3 | 0.19898 | 0.12788 | 0.08224 | 0.29179 | 32.98611 | 0.24 | 0.2 | Neutral |
| testneg_115 | 0.14 | -0.37 | 1.658258 | -3 | 1.806186 | 1.806186 | 0.3414 | 0.13575 | 0.19021 | 0.09379 | 24.97903 | 0.049444444 | 0 | Neutral |
| testneg_116 | 0 | 0 | -0.183807 | -3 | 2.420323 | -3 | 0.13906 | 0.09925 | 0.16436 | 0.19109 | 24.68926 | 0.093894919 | 0 | Neutral |
| testneg_117 | 0.14 | -0.69 | 3.934145 | -3 | 1.27566 | -3 | 0.25011 | 0.12092 | 0.18535 | 0.12788 | 11.77095 | 0.039041667 | 0 | Neutral |
| testneg_118 | 0 | -0.01 | 1.373536 | -3 | -0.668638 | -3 | 0.17258 | 0.12788 | 0.37131 | 0.10994 | 18.7347 | 0 | 0 | Neutral |
| testneg_119 | -0.09 | 1.07 | 5.012896 | -3 | 4.092943 | -3 | 0.47166 | 0.13548 | 0.83421 | 0.18817 | 20.83333 | 0.582818763 | 85 | Disease_causing |
| testneg_120 | 0 | 0 | 1.928458 | -3 | 1.232154 | -3 | 0.45152 | 0.34457 | 0.17768 | 0.10832 | 21.94444 | 0.04 | 0 | Neutral |
| testneg_121 | 0.06 | 0.18 | 1.528609 | -3 | 3.591789 | -3 | 0.20249 | 0.10327 | 0.0925 | 0.08222 | 17.77778 | 0.047791667 | 0 | Neutral |
| testneg_122 | 0.01 | -1.44 | 2.300256 | -3 | 0.452369 | -3 | 0.14996 | 0.09121 | 0.09504 | 0.38939 | 22.08102 | 0.149616279 | 0 | Neutral |
| testneg_123 | 0.03 | -0.85 | 1.940625 | -3 | 0.637019 | -3 | 0.85829 | 0.4292 | 0.16392 | 0.12788 | 10.83333 | 0.042083333 | 0 | Neutral |
| testneg_124 | 0 | 0 | 0.617174 | -3 | 1.790743 | -3 | 0.28416 | 0.27805 | 0.7823 | 0.5949 | 10.83333 | 0.080872526 | 0 | Neutral |
| testneg_125 | 0.14 | -1.01 | 3.030637 | -3 | 0.820981 | 0.427242 | 0.26482 | 0.12788 | 0.13173 | 0.12788 | 14.91327 | 0.03 | 0 | Neutral |
| testneg_126 | 0 | 0 | -0.961702 | -0.961702 | 0.894948 | -3 | 0.46979 | 0.1705 | 0.2236 | 0.09422 | 19.09347 | 0.02 | 0 | Neutral |
| testneg_127 | 0 | 0 | 1.98291 | -3 | 4.370033 | 4.08472 | 0.68918 | 0.16819 | 0.10213 | 0.22954 | 12.38369 | 0.532197989 | 67.2 | Disease_causing |
| testneg_128 | 0 | 0.32 | -0.420859 | -3 | 2.61063 | -3 | 0.0597 | 0.18092 | 0.04627 | 0.05223 | 10 | 0.095111111 | 0 | Neutral |
| testneg_129 | 0.14 | -1.01 | 2.403577 | -3 | 3.958582 | -3 | 0.99417 | 0.17365 | 0.39163 | 0.2681 | 11.25 | 0.532527943 | 67.2 | Disease_causing |
| testneg_130 | 0 | 0 | 1.200491 | -3 | 2.51727 | -3 | 0.34198 | 0.1113 | 0.06196 | 0.10881 | 11.25 | 0.010093074 | 0 | Neutral |
| testneg_131 | 0.07 | 0.38 | -1.728128 | -3 | 4.534835 | -3 | 0.37635 | 0.16791 | 0.19898 | 0.12788 | 10.41667 | 0.181835128 | 0 | Neutral |
| testneg_132 | 0.05 | 0.29 | 2.610797 | -3 | 2.283549 | 2.283549 | 0.00971 | 0.0791 | 0.17786 | 0.15314 | 17.80091 | 0.2 | 0.2 | Neutral |
| testneg_133 | 0 | 0 | 1.119261 | -0.18335 | -0.4837 | -3 | 0.11858 | 0.11551 | 0.23276 | 0.31401 | 24.04977 | 0.07 | 0 | Neutral |
| testneg_134 | 0.08 | -1.2 | 1.68139 | -3 | 3.003558 | -0.365462 | 0.4094 | 0.1944 | 0.19898 | 0.12788 | 19.47505 | 0.14 | 0 | Neutral |
| testneg_135 | -0.15 | 1.06 | 3.233126 | -3 | 6.118134 | -3 | 0.66422 | 0.99377 | 0.05754 | 0.10881 | 11.875 | 0.45 | 34.2 | Neutral |
| testneg_136 | -0.08 | -0.03 | 2.148373 | -3 | -2.342151 | -3 | 0.21376 | 0.12708 | 0.11006 | 0.14352 | 30.53632 | 0.07 | 0 | Neutral |
| testneg_137 | -0.17 | 1.22 | -0.126067 | -3 | 3.427531 | -3 | 0.19898 | 0.09171 | 0.09527 | 0.09424 | 60.48989 | 0.22 | 0.2 | Neutral |
| testneg_138 | -0.02 | 0.36 | 0.416033 | -3 | 5.004967 | -3 | 0.43563 | 0.22365 | 0.1594 | 0.22002 | 16.79207 | 0.507092534 | 59 | Disease_causing |
| testneg_139 | 0 | 0 | 1.255675 | -3 | 7.302547 | -3 | 0.20675 | 0.13709 | 0.09195 | 0.14651 | 8.97898 | 0.46187167 | 37 | Neutral |
| testneg_140 | 0 | 0 | 0.387274 | -3 | 0.465329 | -3 | 0.35412 | 0.10819 | 0.11131 | 0.68435 | 18.33333 | 0.060480437 | 0 | Neutral |
| testneg_141 | -0.09 | 0.3 | 0.246063 | -3 | 2.642734 | -3 | 0.46091 | 0.13852 | 0.19898 | 0.12788 | 23.44851 | 0.062055556 | 0 | Neutral |
| testneg_142 | 0.05 | -0.81 | 2.157622 | -3 | 0.30427 | -3 | 0.35967 | 0.09418 | 0.47246 | 0.47221 | 5.69444 | 0.07 | 0 | Neutral |
| testneg_143 | 0 | 0 | 0.839823 | -3 | -0.907503 | -3 | 0.26997 | 0.63944 | 0.05315 | 0.12788 | 28.83293 | 0.07 | 0 | Neutral |
| testneg_144 | 0 | 0 | -0.535508 | -3 | 1.56019 | -3 | 0.01196 | 0.12788 | 0.25487 | 0.13098 | 19.44444 | 0.06 | 0 | Neutral |
| testneg_145 | -0.18 | -0.16 | 2.830718 | -3 | 3.447686 | -3 | 0.11229 | 0.08897 | 0.19898 | 0.1054 | 18.16092 | 0.13 | 0 | Neutral |
| testneg_146 | -0.03 | -0.1 | 2.078666 | -3 | 0.723398 | 0.723398 | 0.17576 | 0.12193 | 0.19898 | 0.14957 | 27.4024 | 0.07 | 0 | Neutral |
| testneg_147 | -0.15 | -0.04 | 0.577305 | -3 | 0.731991 | -3 | 0.17725 | 0.12788 | 0.13141 | 0.1123 | 23.65426 | 0.04 | 0 | Neutral |
| testneg_148 | 0.03 | -1.03 | 3.226562 | -3 | -2.80816 | -3 | 0.17597 | 0.12788 | 0.19898 | 0.0989 | 21.33705 | 0.030873851 | 0 | Neutral |
| testneg_149 | 0.02 | -0.63 | 0.695636 | 0.695636 | -0.10396 | -3 | 0.12003 | 0.12788 | 0.4722 | 0.13321 | 55.7521 | 0.15 | 0 | Neutral |
| testneg_150 | 0 | 0 | 1.103204 | -3 | -1.260717 | -3 | 0.57199 | 0.11262 | 0.30428 | 0.12788 | 20.46346 | 0.01 | 0 | Neutral |
| testneg_151 | 0.2 | -1.26 | 6.540654 | -3 | 0.86879 | 0.86879 | 0.19898 | 0.12788 | 0.50303 | 0.23508 | 17.29167 | 0.4 | 23.8 | Neutral |
| testneg_152 | 0.16 | -0.37 | 5.705013 | -3 | 0.313134 | -3 | 0.07211 | 0.12934 | 0.11983 | 0.24162 | 18.86983 | 0.403996969 | 19 | Neutral |
| testneg_153 | -0.0087273 | 0.00581818 | 10.530907 | -3 | 2.00954 | -3 | 0.29024 | 0.1472 | 0.17808 | 0.12788 | 14.24934 | 0.51771875 | 59.6 | Disease_causing |
| testneg_154 | 0.07 | 0.01 | 1.091058 | -3 | -0.329218 | -0.329218 | 0.26816 | 0.13347 | 0.13532 | 0.19904 | 12.7356 | 0.070122969 | 0 | Neutral |
| testneg_155 | 0 | 0 | 0.235716 | -3 | 2.049333 | -3 | 0.42033 | 0.17821 | 0.19898 | 0.12788 | 16.8018 | 0.013917749 | 0 | Neutral |
| testneg_156 | -0.19 | -0.51 | -0.445993 | -0.445993 | 0.661515 | -3 | 0.18324 | 0.13041 | 0.07534 | 0.09292 | 5.75579 | 0.052916667 | 0 | Neutral |
| testneg_157 | 0 | 0 | -0.258914 | -3 | -1.094626 | -3 | 0.15623 | 0.1447 | 0.3783 | 0.08224 | 17.21847 | 0.01 | 0 | Neutral |
| testneg_158 | -0.14 | 0.37 | 3.473048 | -3 | 1.065508 | -3 | 0.14485 | 0.16945 | 0.84236 | 0.14093 | 16.42913 | 0.188375 | 0 | Neutral |
| testneg_159 | -0.15 | -0.04 | 3.490664 | -3 | -1.480749 | -1.509154 | 0.29362 | 0.22177 | 0.19898 | 0.12788 | 23.22567 | 0.1 | 0 | Neutral |
| testneg_160 | 0.08 | -1.2 | 1.332661 | -3 | 1.630979 | -3 | 0.03664 | 0.06179 | 0.35532 | 0.24703 | 49.93377 | 0.150634058 | 0 | Neutral |
| testneg_161 | 0.16 | 0.08 | 1.309209 | -3 | 4.017316 | -3.785865 | 0.84588 | 0.14896 | 0.04593 | 0.09315 | 17.77778 | 0.154740725 | 0 | Neutral |
| testneg_162 | 0 | 0 | 1.56019 | -3 | -0.674729 | -3 | 0.25487 | 0.13098 | 0.24765 | 0.32858 | 18.33333 | 0.051254963 | 0 | Neutral |
| testneg_163 | -0.02 | 0.36 | 1.311291 | -3 | 0.75187 | -3 | 0.07577 | 0.09128 | 0.19898 | 0.535 | 18.33333 | 0.070145658 | 0 | Neutral |
| testneg_164 | 0 | 0 | 0.112197 | 0.112197 | 0.801954 | -3 | 0.1123 | 0.1568 | 0.09661 | 0.09981 | 12.31231 | 0.020382653 | 0 | Neutral |
| testneg_165 | -0.18 | -0.16 | 3.107738 | -3 | 1.849259 | -3 | 0.49012 | 0.12138 | 0.41261 | 0.12096 | 18.6768 | 0.070297619 | 0 | Neutral |
| testneg_166 | -0.02 | 0.36 | 2.655425 | -3 | 2.025501 | -3 | 0.19883 | 0.09434 | 0.12253 | 0.10814 | 9.23878 | 0.030253442 | 0 | Neutral |
| testneg_167 | -0.07 | -0.7 | 3.736514 | -3 | 0.212096 | -3 | 0.19898 | 0.12788 | 0.88456 | 0.11674 | 10.41667 | 0.04352381 | 0 | Neutral |
| testneg_168 | 0.01 | 0.8 | 2.947434 | -3 | -1.188116 | -3 | 0.27915 | 0.27706 | 0.25203 | 0.12788 | 19.8494 | 0.080289352 | 0 | Neutral |
| testneg_169 | 0.14 | -0.37 | 0.586641 | -3 | 0.574036 | -3 | 0.07571 | 0.09432 | 0.19898 | 0.12788 | 6.73689 | 0.01 | 0 | Neutral |
| testneg_170 | -0.05 | -0.29 | -0.803439 | -3 | 1.149331 | -3 | 0.50276 | 0.09638 | 0.15582 | 0.14216 | 21.15756 | 0.08 | 0 | Neutral |
| testneg_171 | 0.03 | -1.03 | 2.347484 | -3 | 3.81823 | -3 | 0.16091 | 0.11571 | 0.1755 | 0.10455 | 12.46106 | 0.071260639 | 0 | Neutral |
| testneg_172 | 0 | 0 | 1.145496 | -3 | 3.977929 | -3 | 0.08437 | 0.97198 | 0.06048 | 0.12788 | 20.68173 | 0.210438998 | 0 | Neutral |
| testneg_173 | -0.06 | -0.18 | 1.285578 | -3 | 6.945269 | -3 | 0.087 | 0.12788 | 0.09972 | 0.13992 | 18.04698 | 0.4838125 | 48.6 | Neutral |
| testneg_174 | 0 | 0 | -0.173789 | -3 | 0.866833 | -3 | 0.37275 | 0.16262 | 0.14941 | 0.0838 | 26.66667 | 0.020508621 | 0 | Neutral |
| testneg_175 | -0.07 | -0.7 | 3.730837 | -3 | 4.799959 | -0.069304 | 0.50461 | 0.11034 | 0.32531 | 0.12351 | 13.37927 | 0.229968254 | 0.6 | Neutral |
| testneg_176 | -0.01 | 0.57 | -1.839743 | -3 | 0.131182 | -3 | 0.12446 | 0.08908 | 0.44429 | 0.20468 | 17.97948 | 0.059935897 | 0 | Neutral |
| testneg_177 | 0 | 0 | 0.666829 | -3 | 4.280436 | -3 | 0.30951 | 0.64738 | 0.40116 | 0.43982 | 11.25 | 0.430494359 | 30 | Neutral |
| testneg_178 | -0.01 | 0.57 | 1.661207 | -3 | 0.944356 | -3 | 0.13555 | 0.16663 | 0.32532 | 0.09651 | 10 | 0.04 | 0 | Neutral |
| testneg_179 | 0.00109091 | -0.0100909 | 1.971045 | -3 | 0.968236 | -3 | 0.14457 | 0.11614 | 0.34184 | 0.1964 | 7.641 | 0.030462252 | 0 | Neutral |
| testneg_180 | 0 | -0.01 | 0.36151 | -3 | 1.183797 | -3 | 0.28878 | 0.16765 | 0.99765 | 0.95064 | 4.44444 | 0.135799898 | 0 | Neutral |
| testneg_181 | 0 | 0 | 1.365392 | -3 | 2.511594 | -3 | 0.72062 | 0.14321 | 0.15432 | 0.095 | 17.21847 | 0.021279178 | 0 | Neutral |
| testneg_182 | 0.17 | -1.22 | 6.579463 | -3 | 1.401482 | -3 | 0.08999 | 0.0989 | 0.53496 | 0.13586 | 18.33333 | 0.279211091 | 1.6 | Neutral |
| testneg_183 | -0.07 | -0.01 | 2.087185 | -3 | 1.613914 | -3 | 0.06392 | 0.12788 | 0.07558 | 0.12788 | 26.9055 | 0.021843884 | 0 | Neutral |
| testneg_184 | 0.03 | 0.1 | 5.653067 | -3 | -0.691406 | -3 | 0.13704 | 0.10054 | 0.04046 | 0.06599 | 18.33333 | 0.09 | 0 | Neutral |
| testneg_185 | -0.1 | -1.9 | 2.083029 | 0.83037 | 3.849548 | -3 | 0.16929 | 0.12788 | 0.36257 | 0.26115 | 13.93265 | 0.429588747 | 28.2 | Neutral |
| testneg_186 | 0.00672727 | 0.04872727 | -0.132461 | -3 | 7.817244 | -3 | 0.16002 | 0.10481 | 0.19898 | 0.46867 | 18.76159 | 0.442749351 | 33.4 | Neutral |
| testneg_187 | -0.02 | -0.45 | 3.127301 | -3 | 1.559263 | -3 | 0.1621 | 0.1641 | 0.1662 | 0.16982 | 28.46606 | 0.285 | 1.8 | Neutral |
| testneg_188 | -0.07 | -0.01 | 2.087185 | 2.087185 | -1.722922 | -3 | 0.06392 | 0.12788 | 0.06704 | 0.1068 | 23.68839 | 0.09 | 0 | Neutral |
| testneg_189 | -0.23 | 1.86 | 4.535556 | -3 | 1.349086 | -3 | 0.20673 | 0.13236 | 0.17199 | 0.10586 | 13.23582 | 0.195833333 | 0 | Neutral |
| testneg_190 | 0 | 0 | 0.101625 | -3 | 3.299396 | -3 | 0.6323 | 0.09928 | 0.12547 | 0.09267 | 22.08561 | 0.071263889 | 0 | Neutral |
| testneg_191 | -0.02 | 0.28 | 3.559312 | -3 | 4.082236 | -3 | 0.39964 | 0.22293 | 0.47322 | 0.11118 | 20.13889 | 0.307295038 | 2.6 | Neutral |
| testneg_192 | 0 | 0 | 0.844548 | -3 | 5.658839 | -3 | 0.9237 | 0.55859 | 0.42899 | 0.31045 | 10 | 0.337982684 | 9.2 | Neutral |
| testneg_193 | 0 | 0 | 1.678033 | -3 | -0.196592 | -0.196592 | 0.33957 | 0.12788 | 0.32733 | 0.16869 | 17.77778 | 0.034520412 | 0 | Neutral |
| testneg_194 | 0.16 | -0.37 | 5.42275 | -3 | 0.268956 | -3 | 0.44816 | 0.56655 | 0.18013 | 0.25346 | 10.83333 | 0.560609848 | 75 | Disease_causing |
| testneg_195 | 0.21 | 0.15 | 4.319063 | -3 | -1.950188 | -3 | 0.1647 | 0.12076 | 0.09273 | 0.08316 | 19.1153 | 0.052 | 0 | Neutral |
| testneg_196 | 0 | -0.32 | -0.324056 | -3 | -0.560044 | -3 | 0.08999 | 0.0989 | 0.19898 | 0.16603 | 12.70833 | 0.050215278 | 0 | Neutral |
| testneg_197 | -0.06 | -0.18 | 4.193908 | -3 | -0.658155 | -3 | 0.23088 | 0.12788 | 0.05087 | 0.09682 | 12.88604 | 0.04 | 0 | Neutral |
| testneg_198 | 0 | 0 | 1.217695 | -3 | 0.196323 | -3 | 0.27629 | 0.11166 | 0.13693 | 0.12788 | 17.43377 | 0 | 0 | Neutral |
| testneg_199 | 0.03 | 1.22 | 1.399794 | -3 | 1.00433 | -3 | 0.49425 | 0.25538 | 0.06254 | 0.23561 | 16.60044 | 0.170662698 | 0 | Neutral |
| testneg_200 | -0.06 | -0.18 | 3.18158 | -3 | 2.661704 | 2.661704 | 0.23439 | 0.09253 | 0.23691 | 0.1015 | 26.07351 | 0.15 | 0 | Neutral |
| testneg_201 | 0.14 | -1.01 | 5.641638 | -3 | 3.074698 | -3 | 0.57517 | 0.10873 | 0.8238 | 0.38836 | 10.30286 | 0.584918915 | 85.6 | Disease_causing |
| testneg_202 | 0 | 0 | 0.717597 | -3 | -0.361677 | -3 | 0.9327 | 0.12788 | 0.85759 | 0.1031 | 25.93403 | 0.020086682 | 0 | Neutral |
| testneg_203 | -0.04 | 0.23 | 4.913745 | -3 | 0.429957 | -3 | 0.12912 | 0.09693 | 0.16737 | 0.12788 | 17.21847 | 0.07 | 0 | Neutral |
| testneg_204 | 0.05 | -0.81 | 6.106102 | -3 | 1.311634 | -3 | 0.34288 | 0.24605 | 0.93853 | 0.93439 | 10 | 0.531666667 | 68.2 | Disease_causing |
| testneg_205 | 0 | -0.01 | 0.78013 | -3 | -0.332338 | -3 | 0.14643 | 0.12788 | 0.13708 | 0.81545 | 24.96922 | 0.090286555 | 0 | Neutral |
| testneg_206 | -0.06 | -0.18 | 4.107901 | -3 | -0.718609 | -3 | 0.16797 | 0.11809 | 0.18371 | 0.10748 | 9.0992 | 0.03 | 0 | Neutral |
| testneg_207 | 0 | -0.01 | 0.651236 | -3 | -1.902903 | -3 | 0.16826 | 0.11262 | 0.1254 | 0.08697 | 10 | 0.000115611 | 0 | Neutral |
| testneg_208 | 0 | -0.01 | 2.889805 | -3 | 3.748941 | -3 | 0.80979 | 0.5595 | 0.19898 | 0.09322 | 20.59685 | 0.220540936 | 0.2 | Neutral |
| testneg_209 | 0.04 | -0.23 | 4.364671 | -3 | 1.464574 | 1.464574 | 0.1247 | 0.10331 | 0.60603 | 0.12788 | 24.083 | 0.064219298 | 0 | Neutral |
| testneg_210 | 0.09 | -0.25 | 2.752111 | -3 | -0.445161 | -0.445161 | 0.10556 | 0.28423 | 0.09728 | 0.12788 | 47.91519 | 0.084040476 | 0 | Neutral |
| testneg_211 | 0.06 | -0.57 | 0.402174 | -3 | 2.577846 | -3 | 0.22892 | 0.10495 | 0.12819 | 0.1229 | 31.13776 | 0.02 | 0 | Neutral |
| testneg_212 | -0.2 | 1.98 | -2.111831 | -3 | 1.124718 | -3 | 0.10486 | 0.08083 | 0.55043 | 0.12112 | 13.14653 | 0.19 | 0 | Neutral |
| testneg_213 | 0 | 0 | 0.123596 | -3 | -0.938971 | -3 | 0.53292 | 0.32174 | 0.07552 | 0.12788 | 19.60216 | 0.03 | 0 | Neutral |
| testneg_214 | -0.21 | -0.15 | 2.709016 | -3 | 3.067866 | -3 | 0.17576 | 0.12193 | 0.04831 | 0.12788 | 36.00451 | 0.11 | 0 | Neutral |
| testneg_215 | 0 | 0 | 0.342134 | -3 | 2.956376 | -3 | 0.64661 | 0.49283 | 0.12176 | 0.09356 | 10.11674 | 0.051682903 | 0 | Neutral |

| | | | | | | | | | | | | | | |
|---|---|---|---|---|---|---|---|---|---|---|---|---|---|---|
| testneg_216 | 0.15 | -0.28 | 0.907 | -3 | -1.707001 | -3 | 0.54015 | 0.13176 | 0.1087 | 0.08523 | 12.40196 | 0.04 | 0 | Neutral |
| testneg_217 | 0.09 | -2.17 | 6.595057 | -3 | 2.73719 | -3 | 0.59066 | 0.10874 | 0.1454 | 0.10285 | 19.09347 | 0.41 | 18.4 | Neutral |
| testneg_218 | 0.02 | 0.45 | 5.906217 | -3 | 0.278169 | -3 | 0.72163 | 0.10596 | 0.10963 | 0.12788 | 16.8018 | 0.17 | 0 | Neutral |
| testneg_219 | 0 | 0 | 1.890439 | -3 | 3.159116 | -3 | 0.64188 | 0.09494 | 0.54891 | 0.55746 | 5 | 0.156658293 | 0 | Neutral |
| testneg_220 | -0.07 | -0.38 | 2.626972 | -3 | -1.989097 | -3 | 0.47775 | 0.10103 | 0.11094 | 0.12788 | 18.88889 | 0.02 | 0 | Neutral |
| testneg_221 | 0 | 0 | 0.062047 | -3 | 2.379482 | -3 | 0.21928 | 0.22085 | 0.6854 | 0.12563 | 11.875 | 0.05 | 0 | Neutral |
| testneg_222 | -0.06 | -0.18 | 2.578731 | -3 | 3.857783 | -3 | 0.18283 | 0.11376 | 0.09527 | 0.12788 | 4.72222 | 0.1 | 0 | Neutral |
| testneg_223 | 0.2 | -1.26 | 3.203428 | -3 | 5.357019 | -3 | 0.35325 | 0.12614 | 0.19898 | 0.11773 | 12.29167 | 0.21 | 0.2 | Neutral |
| testneg_224 | 0.21 | 0.15 | 3.567896 | -3 | 1.412972 | -3 | 0.22443 | 0.10657 | 0.10565 | 0.08739 | 10.41667 | 0.054166667 | 0 | Neutral |
| testneg_225 | 0.07 | 0.7 | 4.036956 | -3 | 4.466961 | -3 | 0.19898 | 0.12788 | 0.49644 | 0.10958 | 11.25 | 0.240238179 | 0.8 | Neutral |
| testneg_226 | 0 | -0.01 | 2.187535 | -3 | 5.955324 | -3 | 0.93654 | 0.38765 | 0.24287 | 0.11626 | 11.25 | 0.16196746 | 0 | Neutral |
| testneg_227 | 0.14 | -0.37 | 5.21715 | 1.307491 | 6.869474 | -3 | 0.20657 | 0.12788 | 0.47188 | 0.10037 | 10.41667 | 0.59 | 86 | Disease_causing |
| testneg_228 | 0 | 0 | 1.204698 | -3 | 1.183756 | -3 | 0.6513 | 0.15254 | 0.98118 | 0.10229 | 10.09538 | 0.02 | 0 | Neutral |
| testneg_229 | 0 | 0 | 0.038426 | -3 | 0.823839 | -3 | 0.54909 | 0.85056 | 0.13031 | 0.85678 | 24.01158 | 0.258063241 | 0.2 | Neutral |
| testneg_230 | -0.07 | -0.38 | 4.03608 | -3 | 6.995977 | -3 | 0.39459 | 0.12788 | 0.16462 | 0.18467 | 10.41667 | 0.757793956 | 99 | Disease_causing |
| testneg_231 | 0 | 0 | -0.02198 | -0.02198 | 1.498072 | -3 | 0.15509 | 0.1326 | 0.22113 | 0.10673 | 23.6233 | 0.02 | 0 | Neutral |
| testneg_232 | -0.03 | -0.12 | 1.627305 | -3 | 2.196162 | -3 | 0.08448 | 0.17914 | 0.12977 | 0.1128 | 12.40248 | 0.050311982 | 0 | Neutral |
| testneg_233 | 0 | -0.01 | 8.815204 | -3 | -0.749587 | -3 | 0.51398 | 0.38821 | 0.84318 | 0.11313 | 13.99351 | 0.421884058 | 21.4 | Neutral |
| testneg_234 | -0.05 | -0.29 | -1.669304 | -3 | -0.293928 | -3 | 0.19898 | 0.12788 | 0.103 | 0.403 | 16.8018 | 0.1 | 0 | Neutral |
| testneg_235 | -0.07 | -0.33 | 0.148752 | -3 | 3.811358 | -3 | 0.93468 | 0.12788 | 0.58985 | 0.10504 | 10.83333 | 0.08 | 0 | Neutral |
| testneg_236 | 0 | 0 | 0.004869 | -3 | 0.332253 | -3 | 0.16134 | 0.1847 | 0.23461 | 0.11809 | 16.44639 | 0.01 | 0 | Neutral |
| testneg_237 | -0.02 | -0.13 | 1.214257 | -3 | 5.509658 | -3 | 0.43459 | 0.12595 | 0.16485 | 0.13045 | 13.87828 | 0.158120197 | 0 | Neutral |
| testneg_238 | 0.03 | -1.03 | 2.514378 | -3 | 1.005118 | -3 | 0.3793 | 0.24105 | 0.10502 | 0.11262 | 20.34389 | 0.0505 | 0 | Neutral |
| testneg_239 | 0 | -0.32 | -0.565398 | -3 | -1.227124 | -3 | 0.19898 | 0.12788 | 0.10914 | 0.76607 | 10.30286 | 0.090673401 | 0 | Neutral |
| testneg_240 | 0.11 | -1.95 | 3.776207 | -3 | 0.616957 | -3 | 0.18765 | 0.27794 | 0.11748 | 0.12788 | 9.59364 | 0.10125 | 0 | Neutral |
| testneg_241 | 0.03 | 0.12 | 0.565832 | -3 | 2.186439 | -3 | 0.1105 | 0.44945 | 0.05476 | 0.12788 | 20.208 | 0.09092328 | 0 | Neutral |
| testneg_242 | 0.03 | 0.12 | 0.235146 | -3 | 0.470474 | -3 | 0.16856 | 0.19221 | 0.31824 | 0.6682 | 14.16667 | 0.12672105 | 0 | Neutral |
| testneg_243 | -0.02 | 0.28 | -0.009422 | -3 | 5.068794 | -3 | 0.16275 | 0.12788 | 0.04627 | 0.05223 | 10 | 0.096317016 | 0 | Neutral |
| testneg_244 | 0.19 | -0.27 | 3.485229 | -3 | 0.421557 | 0.421557 | 0.22872 | 0.15588 | 0.82507 | 0.19615 | 14.10753 | 0.19 | 0 | Neutral |
| testneg_245 | 0.07 | 0.38 | -0.999113 | -3 | 5.537244 | -3 | 0.25961 | 0.07098 | 0.19898 | 0.09972 | 16.84579 | 0.094880952 | 0 | Neutral |
| testneg_246 | 0 | 0 | -0.450207 | -3 | 0.242595 | -3 | 0.17083 | 0.31043 | 0.11358 | 0.12788 | 19.44444 | 0.03 | 0 | Neutral |
| testneg_247 | 0 | 0 | 0.353546 | -3 | 2.114299 | -3 | 0.28191 | 0.20693 | 0.42666 | 0.12788 | 10.21277 | 0.03 | 0 | Neutral |
| testneg_248 | 0 | 0 | 1.371909 | -3 | 2.045629 | -3 | 0.47733 | 0.08388 | 0.23005 | 0.37216 | 19.76242 | 0.070115986 | 0 | Neutral |
| testneg_249 | -0.06 | -0.18 | 0.491194 | 0.491194 | 5.21592 | -3 | 0.72124 | 0.09906 | 0.83798 | 0.14433 | 11.74429 | 0.262365569 | 0.6 | Neutral |
| testneg_250 | -0.01 | -0.79 | 4.248239 | -3 | -0.805253 | -3 | 0.16061 | 0.12426 | 0.19898 | 0.12788 | 29.34685 | 0.03 | 0 | Neutral |
| testneg_251 | 0 | -0.32 | 3.269143 | 3.269143 | -0.45893 | -3 | 0.11534 | 0.12281 | 0.059 | 0.12788 | 15.37063 | 0.4 | 19.2 | Neutral |
| testneg_252 | 0.21 | 0.15 | 4.319063 | -3 | 0.05039 | -3 | 0.1647 | 0.12076 | 0.19898 | 0.12788 | 20.74074 | 0.04 | 0 | Neutral |
| testneg_253 | 0.14 | -0.69 | -0.760208 | -3 | 2.890453 | -3 | 0.47443 | 0.11062 | 0.90308 | 0.28373 | 10 | 0.161237088 | 0 | Neutral |
| testneg_254 | 0.22 | -2.66 | 6.137657 | -3 | 0.362333 | -3 | 0.40232 | 0.13342 | 0.7413 | 0.18974 | 10.20833 | 0.368970375 | 13.2 | Neutral |
| testneg_255 | 0.08 | -1.2 | 1.68139 | -3 | 4.2288 | -3 | 0.4094 | 0.1944 | 0.18706 | 0.10259 | 17.91366 | 0.191163385 | 0 | Neutral |
| testneg_256 | 0.04 | -0.23 | -2.488973 | -3 | 1.129155 | -3 | 0.0513 | 0.33622 | 0.16835 | 0.09282 | 20.83333 | 0.13 | 0 | Neutral |
| testneg_257 | 0.07 | -0.31 | 1.843162 | -3 | -1.060552 | -3 | 0.30869 | 0.11948 | 0.2559 | 0.10848 | 21.08313 | 0.01 | 0 | Neutral |
| testneg_258 | 0.11 | -0.19 | 3.662322 | -3 | 0.5524 | -3 | 0.49745 | 0.1103 | 0.44644 | 0.10813 | 19.10253 | 0.03 | 0 | Neutral |
| testneg_259 | 0 | 0 | 0.173288 | -3 | 2.1029 | -3 | 0.61211 | 0.11059 | 0.40232 | 0.13342 | 11.93907 | 0.07 | 0 | Neutral |
| testneg_260 | 0.14 | -0.69 | 0.767974 | -3 | 2.458226 | -3 | 0.09531 | 0.11449 | 0.97005 | 0.31407 | 10.41667 | 0.15 | 0 | Neutral |
| testneg_261 | -0.01 | -0.8 | 1.016479 | 1.016479 | 2.074047 | -3 | 0.12649 | 0.12788 | 0.09353 | 0.12788 | 16.8018 | 0.04 | 0 | Neutral |
| testneg_262 | -0.09 | 0.67 | 0.082232 | -3 | 1.897319 | -3 | 0.08475 | 0.19889 | 0.39225 | 0.10625 | 16.48772 | 0.076688713 | 0 | Neutral |
| testneg_263 | 0 | 0 | 3.603309 | -3 | 0.2182 | -3 | 0.26192 | 0.15452 | 0.19898 | 0.12788 | 34.45733 | 0.04 | 0 | Neutral |
| testneg_264 | 0.06 | -0.57 | 5.215375 | -3 | 0.430807 | -3 | 0.11818 | 0.10561 | 0.21122 | 0.12788 | 17.5 | 0.05975 | 0 | Neutral |
| testneg_265 | 0.03 | -1.03 | 4.65196 | -3 | 0.476145 | -3 | 0.1014 | 0.12788 | 0.22645 | 0.1704 | 29.64276 | 0.206666667 | 0 | Neutral |
| testneg_266 | 0 | -0.32 | 2.002476 | -3 | 3.374642 | -3 | 0.15958 | 0.41628 | 0.33264 | 0.10493 | 6.0461 | 0.119328378 | 0 | Neutral |
| testneg_267 | -0.05 | -0.29 | -1.753034 | -3 | -0.888141 | -0.888141 | 0.06845 | 0.13843 | 0.37637 | 0.12788 | 12.70833 | 0.03 | 0 | Neutral |
| testneg_268 | 0 | 0 | 1.39998 | -3 | -0.907393 | -3 | 0.161 | 0.11597 | 0.05332 | 0.12788 | 13.51351 | 0.000168852 | 0 | Neutral |
| testneg_269 | 0 | 0.32 | -2.144521 | -3 | 0.297516 | 0.297516 | 0.49246 | 0.11033 | 0.19294 | 0.10464 | 12.40248 | 0.03 | 0 | Neutral |
| testneg_270 | -0.08 | -0.03 | -0.385913 | -3 | 1.27985 | -3 | 0.0752 | 0.74467 | 0.09334 | 0.10027 | 13.08587 | 0.090557276 | 0 | Neutral |
| testneg_271 | -0.06 | -0.18 | 1.648992 | -3 | -0.908797 | -3 | 0.73601 | 0.26791 | 0.12282 | 0.20617 | 10.41667 | 0.092592585 | 0 | Neutral |
| testneg_272 | -0.06 | 0.57 | 1.183447 | -3 | 4.766859 | -3 | 0.0146 | 0.03045 | 0.20632 | 0.12788 | 19.81145 | 0.125 | 0 | Neutral |
| testneg_273 | -0.01 | -0.79 | 1.796129 | -3 | 2.326926 | -3 | 0.16052 | 0.12788 | 0.07558 | 0.12788 | 26.9055 | 0.02 | 0 | Neutral |
| testneg_274 | 0.03 | 1.22 | 2.297873 | -3 | 5.621498 | 2.331744 | 0.31168 | 0.12788 | 0.18122 | 0.16895 | 23.44399 | 0.486180836 | 48 | Neutral |
| testneg_275 | 0.09 | -0.25 | 1.459193 | -3 | 4.73993 | -3 | 0.22817 | 0.10634 | 0.40081 | 0.15418 | 12.29167 | 0.244019231 | 0.4 | Neutral |
| testneg_276 | 0.16 | 0.08 | 1.309209 | -3 | 3.497767 | 0.990877 | 0.84588 | 0.14896 | 0.75123 | 0.20667 | 18.33333 | 0.321649884 | 0.8 | Neutral |
| testneg_277 | 0 | 0 | 1.687316 | -3 | 7.674169 | -3 | 0.26434 | 0.12788 | 0.8426 | 0.47836 | 16.48941 | 0.368690476 | 15.4 | Neutral |
| testneg_278 | -0.02 | -0.45 | 1.073382 | -3 | 3.576466 | -0.697584 | 0.31542 | 0.26739 | 0.19898 | 0.12788 | 61.13734 | 0.146071261 | 0 | Neutral |
| testneg_279 | 0 | 0 | 2.055296 | -3 | 0.978375 | -3 | 0.15092 | 0.09074 | 0.62381 | 0.12788 | 22.94152 | 0.01 | 0 | Neutral |
| testneg_280 | 0.04 | -0.23 | 7.16104 | -3 | -0.946844 | -3 | 0.25961 | 0.07098 | 0.19477 | 0.12788 | 24.68926 | 0.227083333 | 0.6 | Neutral |
| testneg_281 | 0 | -0.01 | 0.862118 | -3 | 0.092333 | -3 | 0.50403 | 0.21568 | 0.09522 | 0.20323 | 13.46817 | 0.085282363 | 0 | Neutral |
| testneg_282 | 0.14 | -1.01 | 2.081126 | -3 | 7.426793 | -3 | 0.2663 | 0.32806 | 0.69254 | 0.10021 | 10.2717 | 0.45 | 35.4 | Neutral |
| testneg_283 | 0.03 | -0.85 | -0.678462 | -3 | 2.918405 | -3 | 0.67216 | 0.17808 | 0.836 | 0.15993 | 10.41667 | 0.201153266 | 0 | Neutral |
| testneg_284 | 0.03 | 0.12 | 2.051799 | -3 | -0.296252 | -3 | 0.14624 | 0.10369 | 0.09323 | 0.16732 | 34.2576 | 0.09 | 0 | Neutral |
| testneg_285 | 0 | 0 | 1.405787 | -3 | 2.631127 | -3 | 0.33304 | 0.15588 | 0.16068 | 0.12788 | 10.83333 | 0.031944345 | 0 | Neutral |
| testneg_286 | 0.03 | 1.22 | 1.359632 | -3 | 5.604435 | -3 | 0.38171 | 0.15977 | 0.10261 | 0.0933 | 14.5639 | 0.242230145 | 0.4 | Neutral |
| testneg_287 | 0.05 | -0.81 | 3.455713 | -3 | 1.137849 | -3 | 0.12175 | 0.21177 | 0.15147 | 0.12788 | 12.37434 | 0.1 | 0 | Neutral |
| testneg_288 | 0 | 0 | -0.171544 | -3 | 2.43732 | -3 | 0.19898 | 0.12788 | 0.66422 | 0.99377 | 10.83333 | 0.098390777 | 0 | Neutral |
| testneg_289 | -0.01 | 1.44 | 0.099948 | -3 | 2.667661 | -3 | 0.19898 | 0.56316 | 0.10498 | 0.08421 | 19.44444 | 0.216005952 | 0 | Neutral |
| testneg_290 | -0.08 | -0.03 | 4.604598 | -3 | 2.124149 | -3 | 0.09767 | 0.39937 | 0.14482 | 0.08592 | 15.15306 | 0.180666799 | 0 | Neutral |
| testneg_291 | 0 | 0 | 1.972347 | -3 | -1.814506 | -3 | 0.17181 | 0.10294 | 0.05999 | 0.12788 | 27.73693 | 0.01 | 0 | Neutral |
| testneg_292 | 0 | 0 | 0.484541 | -3 | -0.049164 | -0.049164 | 0.48023 | 0.12788 | 0.12322 | 0.1246 | 26.17936 | 0.02 | 0 | Neutral |
| testneg_293 | 0 | 0 | 0.861365 | -3 | -0.127262 | -3 | 0.37953 | 0.11262 | 0.29523 | 0.27623 | 16.66667 | 0.020327957 | 0 | Neutral |
| testneg_294 | 0.14 | -1.01 | 1.424004 | -3 | -0.532627 | -3 | 0.08932 | 0.12788 | 0.12173 | 0.12788 | 15 | 0.01 | 0 | Neutral |
| testneg_295 | -0.01 | 1.44 | 5.577538 | -3 | 2.400472 | -3 | 0.1196 | 0.12788 | 0.04627 | 0.05223 | 26.34995 | 0.200300082 | 0.4 | Neutral |
| testneg_296 | -0.12 | 1.14 | -1.180653 | -3 | -1.950188 | -3 | 0.04192 | 0.06306 | 0.09273 | 0.08316 | 13.51351 | 0.11 | 0 | Neutral |
| testneg_297 | 0 | 0 | 2.100416 | -3 | 3.36473 | -3 | 0.22255 | 0.10057 | 0.16616 | 0.11073 | 19.97695 | 0.030217494 | 0 | Neutral |
| testneg_298 | -0.02 | -0.45 | 1.073382 | -3 | -0.392743 | -3 | 0.31542 | 0.26739 | 0.16275 | 0.12788 | 22.12374 | 0.03 | 0 | Neutral |
| testneg_299 | 0.01 | -0.57 | 5.075109 | -3 | 0.914524 | -3 | 0.68863 | 0.09663 | 0.31238 | 0.14393 | 44.00858 | 0.150276611 | 0 | Neutral |
| testneg_300 | -0.03 | -1.22 | 4.583105 | -3 | 3.009853 | -3 | 0.06366 | 0.12788 | 0.50741 | 0.17915 | 29.67756 | 0.44151109 | 31.2 | Neutral |
| testneg_301 | 0 | -0.01 | 0.834318 | -3 | 0.437223 | -3 | 0.31726 | 0.17249 | 0.0684 | 0.08803 | 14.43558 | 0.01 | 0 | Neutral |
| testneg_302 | 0.03 | -1.03 | 4.689762 | -3 | 3.940451 | -3 | 0.08268 | 0.09939 | 0.10756 | 0.0892 | 17.21847 | 0.16 | 0 | Neutral |
| testneg_303 | 0 | 0 | 3.319009 | -3 | 3.763769 | -3 | 0.22892 | 0.10495 | 0.32558 | 0.32796 | 20.33859 | 0.470239295 | 42.4 | Neutral |
| testneg_304 | 0.08 | 0.03 | -1.587151 | -3 | 3.628039 | -3 | 0.15486 | 0.28768 | 0.055 | 0.20858 | 11.25 | 0.464738698 | 38.8 | Neutral |
| testneg_305 | -0.03 | -0.1 | 4.57636 | -3 | -1.941492 | -3 | 0.11773 | 0.11669 | 0.37711 | 0.12788 | 41.27033 | 0.050704545 | 0 | Neutral |
| testneg_306 | -0.04 | 0.23 | 1.498072 | -0.580461 | 1.139387 | -3 | 0.22113 | 0.10673 | 0.62197 | 0.12788 | 21.57895 | 0.01 | 0 | Neutral |
| testneg_307 | 0 | -0.01 | 0.491559 | -3 | 0.284081 | -3 | 0.64319 | 0.12788 | 0.95535 | 0.12788 | 12.37434 | 0.01 | 0 | Neutral |
| testneg_308 | -0.14 | 0.37 | 0.450722 | -3 | 0.236592 | -3 | 0.16078 | 0.39498 | 0.40234 | 0.10353 | 10.41667 | 0.08 | 0 | Neutral |
| testneg_309 | 0.18 | -1.05 | 5.2613 | -3 | -2.091153 | -3 | 0.13696 | 0.09566 | 0.69696 | 0.10507 | 9.54828 | 0.11 | 0 | Neutral |
| testneg_310 | 0 | 0 | -0.052835 | -3 | 2.100416 | -3 | 0.19898 | 0.13434 | 0.22255 | 0.10057 | 12.76316 | 0.01 | 0 | Neutral |
| testneg_311 | 0 | 0 | 0.914524 | -3 | 3.26904 | -3 | 0.31238 | 0.14393 | 0.18554 | 0.12299 | 29.37445 | 0.048612698 | 0 | Neutral |
| testneg_312 | 0.17 | 0.38 | 3.132136 | -3 | 4.985443 | -3 | 0.0105 | 0.12788 | 0.09618 | 0.10294 | 32.09372 | 0.172446277 | 0 | Neutral |
| testneg_313 | 0 | 0 | -0.117724 | -3 | 5.366894 | 1.280316 | 0.17529 | 0.09662 | 0.43762 | 0.12788 | 35.72379 | 0.083380952 | 0 | Neutral |
| testneg_314 | 0.03 | -1.03 | 0.896883 | -3 | 1.982936 | -3 | 0.02124 | 0.05186 | 0.07044 | 0.11679 | 21.52409 | 0.05 | 0 | Neutral |
| testneg_315 | 0 | 0 | 1.550971 | -3 | -0.249119 | -3 | 0.73311 | 0.10866 | 0.14638 | 0.12788 | 10 | 9.52E-05 | 0 | Neutral |
| testneg_316 | -0.07 | -0.38 | 1.434265 | -3 | 7.289398 | -3 | 0.69563 | 0.12788 | 0.57188 | 0.34038 | 4.72222 | 0.445434018 | 33.8 | Neutral |
| testneg_317 | 0 | 0 | 0.358974 | -3 | 2.864701 | -3 | 0.27197 | 0.12346 | 0.98535 | 0.15071 | 18.88889 | 0.074332371 | 0 | Neutral |
| testneg_318 | -0.04 | -2.01 | 5.756532 | -3 | 4.046865 | -3 | 0.15992 | 0.12788 | 0.42666 | 0.12788 | 11.0461 | 0.42 | 24.4 | Neutral |
| testneg_319 | -0.03 | -0.1 | 0.212913 | -3 | 0.046701 | 0.046701 | 0.11391 | 0.19434 | 0.06611 | 0.08834 | 13.51351 | 0.05 | 0 | Neutral |
| testneg_320 | 0.15 | -0.28 | -0.431211 | -3 | -0.229739 | -3 | 0.36676 | 0.37749 | 0.29372 | 0.07609 | 16.0119 | 0.05 | 0 | Neutral |
| testneg_321 | -0.08 | -0.03 | 3.833967 | -3 | 4.113708 | -3 | 0.17855 | 0.11262 | 0.46244 | 0.56748 | 10.41667 | 0.565519119 | 77 | Disease_causing |
| testneg_322 | 0.07 | 0.7 | 4.525325 | -3 | 2.598643 | -3 | 0.11564 | 0.1184 | 0.09515 | 0.07784 | 20.96973 | 0.092166667 | 0 | Neutral |
| testneg_323 | -0.09 | 0.25 | 4.068872 | -3 | -0.01214 | -3 | 0.19898 | 0.12788 | 0.16966 | 0.61198 | 6.99956 | 0.20597619 | 0 | Neutral |

| | | | | | | | | | | | | | | |
|---|---|---|---|---|---|---|---|---|---|---|---|---|---|---|
| testneg_324 | 0.19 | -0.27 | 4.230056 | -3 | 0.368341 | -3 | 0.08489 | 0.12788 | 0.77847 | 0.10751 | 14.25869 | 0.048888889 | 0 | Neutral |
| testneg_325 | 0.02 | -0.36 | 1.517891 | -3 | -3.068024 | -3 | 0.42755 | 0.09484 | 0.41089 | 0.1897 | 11.66667 | 0.049373503 | 0 | Neutral |
| testneg_326 | 0 | 0 | 1.433534 | -3 | 2.698075 | -3 | 0.30275 | 0.12788 | 0.32558 | 0.32796 | 7.78889 | 0.083303846 | 0 | Neutral |
| testneg_327 | 0 | -0.01 | 1.961035 | -3 | -2.166097 | -3 | 0.55105 | 0.43208 | 0.41269 | 0.19135 | 11.25 | 0.082408029 | 0 | Neutral |
| testneg_328 | -0.02 | -0.45 | 2.354402 | -3 | 2.449599 | -3 | 0.0538 | 0.0823 | 0.18578 | 0.27854 | 11.33956 | 0.209089286 | 0 | Neutral |
| testneg_329 | 0 | 0 | -0.22828 | -3 | 6.382916 | -3 | 0.31694 | 0.13475 | 0.46244 | 0.56748 | 11.25 | 0.373645833 | 16 | Neutral |
| testneg_330 | 0.1 | 1.9 | 3.737058 | -3 | 2.742872 | -3 | 0.28191 | 0.20693 | 0.13195 | 0.57267 | 10 | 0.61371502 | 92.6 | Disease_causing |
| testneg_331 | 0.14 | -0.37 | 7.532012 | -3 | 1.87784 | -3 | 0.99704 | 0.17463 | 0.17793 | 0.10516 | 8.97898 | 0.519594595 | 61 | Disease_causing |
| testneg_332 | 0 | 0 | 1.235264 | -3 | 3.759941 | -3 | 0.56904 | 0.14839 | 0.46058 | 0.12903 | 16.56344 | 0.085261905 | 0 | Neutral |
| testneg_333 | 0 | 0 | 0.415281 | -3 | -0.056564 | -3 | 0.1472 | 0.12788 | 0.01642 | 0.12788 | 10.38652 | 0.000259784 | 0 | Neutral |
| testneg_334 | 0 | 0 | 0.849647 | -3 | 1.13018 | 1.13018 | 0.96651 | 0.28104 | 0.50505 | 0.11563 | 9.1693 | 0.010246951 | 0 | Neutral |
| testneg_335 | 0.06 | -0.57 | 4.767867 | -3 | 0.938815 | -3 | 0.161 | 0.11597 | 0.33292 | 0.17111 | 17.29026 | 0.179160819 | 0 | Neutral |
| testneg_336 | -0.08 | -0.03 | 3.206814 | -3 | 3.685438 | -3 | 0.7022 | 0.35308 | 0.16545 | 0.10002 | 18.88889 | 0.24 | 0 | Neutral |
| testneg_337 | 0 | 0 | -0.014148 | -3 | 2.833945 | -3 | 0.24274 | 0.13034 | 0.21623 | 0.42074 | 17.46936 | 0.134862233 | 0 | Neutral |
| testneg_338 | 0 | -0.32 | 1.011617 | -3 | 4.182656 | -3 | 0.19898 | 0.0989 | 0.12933 | 0.12788 | 23.67719 | 0.052967391 | 0 | Neutral |
| testneg_339 | 0.18 | -1.05 | 4.57164 | -3 | 6.905837 | -3 | 0.11335 | 0.14225 | 0.27004 | 0.17052 | 13.04361 | 0.842759259 | 100 | Disease_causing |
| testneg_340 | 0 | 0 | -0.100586 | -3 | 1.272278 | -3 | 0.14668 | 0.25103 | 0.05324 | 0.19446 | 10.41667 | 0.100076923 | 0 | Neutral |
| testneg_341 | -0.16 | 1.37 | 3.207028 | -3 | 7.586474 | -3 | 0.03169 | 0.10615 | 0.13961 | 0.09565 | 16.53165 | 0.48 | 49.2 | Neutral |
| testneg_342 | 0 | 0 | 2.30425 | -3 | 0.825237 | -3 | 0.46792 | 0.11262 | 0.41089 | 0.1897 | 18.33333 | 0.03308908 | 0 | Neutral |
| testneg_343 | -0.08 | -0.03 | 1.155408 | -3 | 3.679774 | -3 | 0.04161 | 0.12788 | 0.06982 | 0.10226 | 13.51351 | 0.090053191 | 0 | Neutral |
| testneg_344 | 0.04 | -0.23 | 4.939659 | -3 | -2.304927 | -2.304927 | 0.13563 | 0.12378 | 0.02665 | 0.05082 | 19.8494 | 0.05 | 0 | Neutral |
| testneg_345 | 0.05 | 0.29 | 2.245213 | -3 | -0.390566 | -3 | 0.1516 | 0.10704 | 0.08283 | 0.11797 | 13.02635 | 0.01 | 0 | Neutral |
| testneg_346 | 0 | 0 | 1.849259 | -3 | 0.970904 | -3 | 0.41261 | 0.12096 | 0.06313 | 0.12788 | 21.99535 | 0.01 | 0 | Neutral |
| testneg_347 | 0.08 | -1.2 | 5.79639 | 0.475838 | 4.83533 | -3 | 0.25516 | 0.12788 | 0.23975 | 0.13861 | 24.8779 | 0.61 | 87.8 | Disease_causing |
| testneg_348 | 0.05 | -0.81 | 2.681817 | -3 | 0.182103 | -3 | 0.11956 | 0.3237 | 0.16355 | 0.20781 | 20.27778 | 0.275 | 0 | Neutral |
| testneg_349 | 0 | 0 | 1.329031 | -3 | 2.768215 | -3 | 0.22489 | 0.16646 | 0.26928 | 0.12774 | 17.62769 | 0.040366667 | 0 | Neutral |
| testneg_350 | -0.08 | -0.03 | 2.285785 | -3 | -0.335332 | -3 | 0.9511 | 0.54626 | 0.50388 | 0.12295 | 14.83527 | 0.05 | 0 | Neutral |
| testneg_351 | 0 | 0 | -0.406057 | -3 | -1.549264 | -3 | 0.7413 | 0.18974 | 0.53172 | 0.12788 | 18.46847 | 0.010846154 | 0 | Neutral |
| testneg_352 | 0.03 | -1.03 | 4.474669 | -3 | 0.657265 | -3 | 0.12446 | 0.08908 | 0.10385 | 0.09775 | 27.33797 | 0.040747023 | 0 | Neutral |
| testneg_353 | -0.000303 | 0.00015152 | -0.119638 | -3 | 3.341558 | -3 | 0.16379 | 0.31462 | 0.19102 | 0.12788 | 17.62359 | 0.095884615 | 0 | Neutral |
| testneg_354 | 0 | 0 | -0.424911 | -3 | -0.075115 | -3 | 0.11797 | 0.12788 | 0.21576 | 0.17241 | 11.25 | 0.030188134 | 0 | Neutral |
| testneg_355 | -0.11 | 0.19 | 3.504228 | 3.504228 | 3.779532 | -3 | 0.47751 | 0.2062 | 0.11373 | 0.10739 | 4.72222 | 0.53 | 67.8 | Disease_causing |
| testneg_356 | 0.14 | -1.01 | 1.869027 | -3 | 1.034706 | -3 | 0.113 | 0.12788 | 0.08896 | 0.12788 | 26.63957 | 0.02 | 0 | Neutral |
| testneg_357 | 0.04 | -0.23 | 7.386316 | -3 | 6.243256 | -3 | 0.14785 | 0.09084 | 0.29804 | 0.12768 | 13.51351 | 0.54 | 67.8 | Disease_causing |
| testneg_358 | 0 | 0 | 2.30653 | -3 | 4.681218 | -3 | 0.49106 | 0.89143 | 0.08401 | 0.16123 | 6.073 | 0.490680644 | 50.8 | Disease_causing |
| testneg_359 | 0 | -0.01 | 1.887787 | -3 | 1.996983 | 1.996983 | 0.51398 | 0.38821 | 0.09553 | 0.11637 | 20.78048 | 0.07 | 0 | Neutral |
| testneg_360 | -0.05 | 0.81 | 4.111365 | 4.111365 | 4.857402 | -3 | 0.1462 | 0.18896 | 0.31648 | 0.12788 | 20.43247 | 0.622876112 | 94 | Disease_causing |
| testneg_361 | 0.12 | -1.16 | 4.470041 | -3 | 6.151216 | 2.096257 | 0.06841 | 0.10927 | 0.34324 | 0.085 | 19.09347 | 0.31 | 2.6 | Neutral |
| testneg_362 | 0.08 | 0.03 | 1.284293 | -3 | 5.396431 | -3 | 0.19898 | 0.13162 | 0.19452 | 0.79779 | 10.83333 | 0.449634511 | 34.8 | Neutral |
| testneg_363 | 0 | 0.32 | 2.196162 | -3 | 0.435447 | -3 | 0.12977 | 0.1128 | 0.21941 | 0.46348 | 11.98582 | 0.051030093 | 0 | Neutral |
| testneg_364 | 0 | 0 | 0.537223 | -3 | 5.001837 | -3 | 0.08186 | 0.11813 | 0.28201 | 0.33127 | 20.83333 | 0.266953093 | 2 | Neutral |
| testneg_365 | 0 | 0 | 0.046064 | -3 | -1.973513 | -1.973513 | 0.2543 | 0.15588 | 0.67796 | 0.10874 | 19.44444 | 0.02 | 0 | Neutral |
| testneg_366 | 0 | 0 | -0.048403 | -3 | 3.994855 | -3 | 0.19898 | 0.12788 | 0.1311 | 0.12667 | 30.2652 | 0.0604 | 0 | Neutral |
| testneg_367 | -0.07 | -0.38 | 6.969629 | -3 | 3.628039 | -3 | 0.76186 | 0.10052 | 0.055 | 0.20858 | 10.41667 | 0.668296367 | 96.8 | Disease_causing |
| testneg_368 | 0 | 0 | 1.433601 | -3 | 2.710461 | -3 | 0.85984 | 0.40044 | 0.15202 | 0.12722 | 24.77407 | 0.07 | 0 | Neutral |
| testneg_369 | 0 | -0.01 | 2.262596 | -3 | 4.091311 | -3 | 0.28732 | 0.11223 | 0.06729 | 0.17051 | 27.4024 | 0.338209486 | 3.4 | Neutral |
| testneg_370 | 0.03 | 0.12 | 1.249184 | -3 | 3.122928 | 2.114299 | 0.40592 | 0.22108 | 0.37637 | 0.12788 | 10 | 0.154703149 | 0 | Neutral |
| testneg_371 | 0.07 | 0.38 | 0.140559 | -3 | -2.747914 | -3 | 0.68029 | 0.12788 | 0.03651 | 0.06141 | 13.75 | 0.03 | 0 | Neutral |
| testneg_372 | -0.04 | 1.61 | 2.828286 | -3 | -0.093852 | -3 | 0.85075 | 0.16646 | 0.12173 | 0.12788 | 16.98582 | 0.1 | 0 | Neutral |
| testneg_373 | 0.02 | -0.68 | 3.397361 | -3 | 0.819861 | -1.346902 | 0.06897 | 0.08445 | 0.05005 | 0.07051 | 26.46391 | 0.05 | 0 | Neutral |
| testneg_374 | 0.18 | 0.16 | -0.887539 | -3 | 0.692563 | -3 | 0.05996 | 0.12788 | 0.39648 | 0.97288 | 29.56338 | 0.16052139 | 0 | Neutral |
| testneg_375 | 0 | 0 | -0.696429 | -3 | 1.280927 | -3 | 0.26476 | 0.08217 | 0.47217 | 0.15544 | 30.87719 | 0.053276698 | 0 | Neutral |
| testneg_376 | 0 | 0 | -0.001616 | -3 | 3.267547 | -3 | 0.90767 | 0.12788 | 0.06027 | 0.11574 | 10.83333 | 0.063081274 | 0 | Neutral |
| testneg_377 | 0 | -0.01 | 0.183879 | -3 | 3.548514 | -3 | 0.19898 | 0.12788 | 0.19623 | 0.08363 | 32.91082 | 0.030476752 | 0 | Neutral |
| testneg_378 | -0.02 | 0.36 | 3.437388 | -3 | -0.416655 | -0.416655 | 0.22645 | 0.1704 | 0.05453 | 0.12788 | 28.44683 | 0.07 | 0 | Neutral |
| testneg_379 | 0 | -0.01 | 1.790566 | -3 | 0.425868 | -3 | 0.59827 | 0.11479 | 0.19898 | 0.12788 | 48.97798 | 0.01 | 0 | Neutral |
| testneg_380 | 0 | 0 | 0.36941 | -3 | 4.276849 | -3 | 0.15406 | 0.1454 | 0.0778 | 0.0728 | 18.88889 | 0.083333333 | 0 | Neutral |
| testneg_381 | -0.2 | 1.26 | 0.425452 | -3 | 3.254561 | -3 | 0.23859 | 0.1202 | 0.14904 | 0.13261 | 13.75 | 0.330138889 | 3 | Neutral |
| testneg_382 | 0.14 | -0.69 | 3.81721 | -3 | -0.692188 | -3 | 0.35945 | 0.12025 | 0.12269 | 0.08105 | 32.36768 | 0.03 | 0 | Neutral |
| testneg_383 | -0.05 | -0.29 | 4.740942 | -3 | -0.749376 | -0.749376 | 0.28063 | 0.40063 | 0.51437 | 0.17173 | 10.83333 | 0.386845238 | 17 | Neutral |
| testneg_384 | -0.05 | -0.29 | 0.110557 | -3 | 1.713048 | -3 | 0.11703 | 0.09364 | 0.42872 | 0.12788 | 26.53988 | 0.02 | 0 | Neutral |
| testneg_385 | 0.04 | -0.23 | 2.455519 | -3 | 1.907441 | 1.907441 | 0.04668 | 0.12788 | 0.20708 | 0.12788 | 18.33333 | 0.07 | 0 | Neutral |
| testneg_386 | 0 | 0 | -0.074927 | -3 | -0.162621 | -3 | 0.99538 | 0.40044 | 0.08105 | 0.12788 | 20.27778 | 0.06 | 0 | Neutral |
| testneg_387 | 0.18 | -1.05 | 5.028255 | -3 | 0.354645 | -3 | 0.17843 | 0.15468 | 0.25893 | 0.10353 | 13.88464 | 0.091111111 | 0 | Neutral |
| testneg_388 | -0.02 | -0.41 | 2.191142 | -3 | 4.1333 | -3 | 0.39225 | 0.10625 | 0.80314 | 0.42344 | 20 | 0.325777433 | 3.2 | Neutral |
| testneg_389 | 0 | 0 | 0.040296 | -3 | 1.956887 | -3 | 0.25856 | 0.17179 | 0.1087 | 0.08523 | 23.45564 | 0.023333333 | 0 | Neutral |
| testneg_390 | 0.01 | 0.79 | 1.728142 | -3 | 6.039671 | -3 | 0.34467 | 0.13076 | 0.11301 | 0.10716 | 35.76137 | 0.1695 | 0 | Neutral |
| testneg_391 | -0.01 | -0.8 | 0.738714 | -3 | 1.131706 | -3 | 0.17158 | 0.1149 | 0.43921 | 0.153 | 10.22898 | 0.032763889 | 0 | Neutral |
| testneg_392 | 0.07 | 0.33 | 3.81823 | -3 | 2.65323 | -3 | 0.1755 | 0.10455 | 0.14485 | 0.16945 | 12.46106 | 0.360202803 | 9.4 | Neutral |
| testneg_393 | 0.07 | -0.31 | 1.028884 | -3 | 1.219225 | -3 | 0.93468 | 0.12788 | 0.29548 | 0.10732 | 12.89073 | 0.02 | 0 | Neutral |
| testneg_394 | 0.01 | 0.8 | 1.447055 | -3 | 2.019145 | -3 | 0.60959 | 0.76078 | 0.93064 | 0.10933 | 10.41667 | 0.09 | 0 | Neutral |
| testneg_395 | 0.16 | -0.37 | 2.034234 | -3 | -0.889855 | -0.889855 | 0.09583 | 0.09376 | 0.042 | 0.07218 | 17.88396 | 0.02 | 0 | Neutral |
| testneg_396 | 0.18 | -1.05 | 7.618716 | -3 | 0.009894 | -3 | 0.38445 | 0.58278 | 0.07996 | 0.08354 | 21.68678 | 0.4 | 18 | Neutral |
| testneg_397 | -0.06 | 0.57 | 4.857661 | -3 | 6.282799 | -3 | 0.08579 | 0.6068 | 0.27528 | 0.12788 | 10.83333 | 0.669014375 | 96.4 | Disease_causing |
| testneg_398 | 0.09 | -0.3 | -0.408308 | -3 | 0.04439 | -3 | 0.28165 | 0.15912 | 0.40081 | 0.15418 | 16.42913 | 0.099502706 | 0 | Neutral |
| testneg_399 | 0.07 | 0.38 | -3.186924 | -3 | 0.286557 | -3 | 0.19898 | 0.09822 | 0.18305 | 0.12743 | 29.34685 | 0.02 | 0 | Neutral |
| testneg_400 | 0.21 | 0.15 | 4.605481 | -3 | 6.559155 | -3 | 0.53888 | 0.09904 | 0.18431 | 0.13863 | 17.21847 | 0.606242007 | 86.4 | Disease_causing |
| testneg_401 | 0 | 0 | -0.326201 | -3 | 3.730217 | -3 | 0.19898 | 0.12788 | 0.16968 | 0.12788 | 28.36964 | 0.060232527 | 0 | Neutral |
| testneg_402 | 0.18 | -1.05 | 6.439341 | -3 | 0.811439 | -3 | 0.61098 | 0.11179 | 0.25758 | 0.2262 | 13.03942 | 0.372015766 | 17.6 | Neutral |
| testneg_403 | 0 | 0 | 0.941338 | -3 | 4.869758 | -3 | 0.35401 | 0.08438 | 0.2218 | 0.11533 | 12.56338 | 0.030425725 | 0 | Neutral |
| testneg_404 | 0.04 | -0.23 | 5.956747 | -3 | 5.497384 | -3 | 0.19898 | 0.15588 | 0.25462 | 0.12788 | 8.97898 | 0.638495689 | 91 | Disease_causing |
| testneg_405 | 0.14 | -1.01 | 1.797795 | -3 | 2.35249 | -3 | 0.19898 | 0.12788 | 0.28685 | 0.1118 | 18.33333 | 0.02083658 | 0 | Neutral |
| testneg_406 | 0.15 | -0.28 | 0.552009 | -3 | 1.40988 | -3 | 0.53172 | 0.12788 | 0.19898 | 0.12788 | 20.5518 | 0.02 | 0 | Neutral |
| testneg_407 | 0.14 | -1.01 | 2.75054 | -3 | 4.478975 | -3 | 0.13691 | 0.11058 | 0.85781 | 0.22949 | 15.31368 | 0.400763628 | 19.6 | Neutral |
| testneg_408 | -0.03 | -0.1 | 0.212913 | 0.212913 | 1.729145 | -3 | 0.11391 | 0.19434 | 0.13575 | 0.12788 | 12.46106 | 0.049398865 | 0 | Neutral |
| testneg_409 | 0.14 | -0.37 | 3.662416 | -3 | 0.295109 | -3 | 0.83176 | 0.23122 | 0.35142 | 0.18303 | 10 | 0.256555556 | 0.4 | Neutral |
| testneg_410 | -0.15 | 0.82 | 2.031124 | -3 | 1.534492 | -3 | 0.08628 | 0.11498 | 0.10183 | 0.11847 | 33.22048 | 0.070052632 | 0 | Neutral |
| testneg_411 | 0.14 | -0.37 | -0.486677 | -3 | -0.483923 | -3 | 0.10496 | 0.21491 | 0.23574 | 0.12788 | 12.46106 | 0.05 | 0 | Neutral |
| testneg_412 | 0.14 | -0.69 | 1.692426 | -3 | 6.279292 | 6.279292 | 0.09306 | 0.09084 | 0.09488 | 0.10426 | 21.33768 | 0.41 | 15.8 | Neutral |
| testneg_413 | 0 | 0 | 1.636749 | -3 | 1.790281 | -3 | 0.1109 | 0.17295 | 0.02059 | 0.066 | 24.65364 | 0.03 | 0 | Neutral |
| testneg_414 | 0.02 | 0.45 | 1.771445 | -3 | 4.126674 | -3 | 0.82126 | 0.17937 | 0.08579 | 0.6068 | 10 | 0.41359375 | 22.6 | Neutral |
| testneg_415 | 0.03 | 0.12 | 4.271499 | 0.486085 | 3.907702 | -3 | 0.71844 | 0.20204 | 0.14498 | 0.33896 | 4.72222 | 0.747487956 | 100 | Disease_causing |
| testneg_416 | 0 | 0 | 0.453554 | -3 | 1.091058 | -3 | 0.70349 | 0.12788 | 0.26816 | 0.13347 | 12.7356 | 0.050303658 | 0 | Neutral |
| testneg_417 | 0 | 0 | 1.504538 | -3 | 1.048529 | -3 | 0.19898 | 0.12788 | 0.54625 | 0.11856 | 11.7735 | 0 | 0 | Neutral |
| testneg_418 | 0.09 | -0.3 | 3.770204 | -3 | 6.36376 | -3 | 0.41647 | 0.11239 | 0.25697 | 0.13883 | 29.27707 | 0.49341733 | 52.4 | Disease_causing |
| testneg_419 | 0 | 0 | 0.419427 | -3 | 3.781815 | -3 | 0.31244 | 0.4615 | 0.01704 | 0.05734 | 16.08774 | 0.095088068 | 0 | Neutral |
| testneg_420 | -0.07 | -0.38 | 5.152033 | -3 | 2.093152 | -3 | 0.34546 | 0.0932 | 0.19898 | 0.12788 | 18.88889 | 0.129772727 | 0 | Neutral |
| testneg_421 | -0.02 | 0.63 | 4.506874 | -3 | 3.198443 | -3 | 0.30942 | 0.13105 | 0.09353 | 0.12788 | 19.09347 | 0.21 | 0 | Neutral |
| testneg_422 | -0.03 | -0.1 | 1.673611 | -3 | 2.12021 | 2.12021 | 0.10554 | 0.12788 | 0.07698 | 0.12788 | 13.51351 | 0.05 | 0 | Neutral |
| testneg_423 | 0 | 0 | -0.163315 | -3 | 2.327743 | -3 | 0.31125 | 0.20061 | 0.0538 | 0.0823 | 20.74074 | 0.044308333 | 0 | Neutral |
| testneg_424 | 0.03 | 0.12 | -1.295182 | -3 | 0.469894 | -3 | 0.26913 | 0.1087 | 0.06067 | 0.07796 | 29.03465 | 0.02 | 0 | Neutral |
| testneg_425 | 0 | 0 | 0.839553 | -3 | 0.606925 | -3 | 0.16835 | 0.09282 | 0.36541 | 0.27749 | 21.38889 | 0.041380952 | 0 | Neutral |
| testneg_426 | 0 | -0.01 | 1.940661 | -3 | 4.759697 | -3 | 0.25151 | 0.12788 | 0.19898 | 0.10335 | 11.5289 | 0.050341435 | 0 | Neutral |
| testneg_427 | -0.02 | 0.36 | 0.523312 | 0.523312 | 7.660132 | -3 | 0.41987 | 0.67373 | 0.34324 | 0.085 | 17.21847 | 0.424166667 | 25.8 | Neutral |
| testneg_428 | 0.16 | -0.37 | -0.68469 | -3 | 2.070069 | -3 | 0.34248 | 0.15472 | 0.42748 | 0.10231 | 18.9586 | 0.040354232 | 0 | Neutral |
| testneg_429 | 0.07 | 0.7 | 3.069764 | -3 | -1.985489 | -1.985489 | 0.25474 | 0.11262 | 0.97005 | 0.31407 | 16.8018 | 0.147136364 | 0 | Neutral |
| testneg_430 | 0 | 0 | 0.768207 | -3 | 2.216369 | -3 | 0.30669 | 0.11489 | 0.1287 | 0.10423 | 12.38369 | 0.000305504 | 0 | Neutral |
| testneg_431 | 0.05 | 0.29 | 0.113571 | -3 | -0.491857 | -3 | 0.02844 | 0.12788 | 0.07598 | 0.08456 | 13.51351 | 0.03 | 0 | Neutral |

| | | | | | | | | | | | | | | |
|---|---|---|---|---|---|---|---|---|---|---|---|---|---|---|
| testneg_432 | -0.05 | -0.29 | -1.315358 | -3 | 3.39754 | -3 | 0.58984 | 0.15588 | 0.62203 | 0.15583 | 10 | 0.246395876 | 0 | Neutral |
| testneg_433 | -0.16 | -0.08 | 6.095834 | -3 | 1.385775 | -3 | 0.11818 | 0.10561 | 0.05027 | 0.10981 | 11.66667 | 0.2301 | 1 | Neutral |
| testneg_434 | -0.08 | -0.03 | 0.395961 | -3 | 5.21715 | 1.307491 | 0.23027 | 0.37171 | 0.20657 | 0.12788 | 10.41667 | 0.249235589 | 0.4 | Neutral |
| testneg_435 | -0.08 | -0.03 | 2.60662 | -3 | 0.670876 | 0.670876 | 0.98853 | 0.88685 | 0.09645 | 0.13239 | 16.577 | 0.23 | 0 | Neutral |
| testneg_436 | -0.04 | -2.01 | 5.568814 | -3 | 6.7522 | -3 | 0.77364 | 0.12676 | 0.19237 | 0.53191 | 10.41667 | 0.78 | 99.6 | Disease_causing |
| testneg_437 | 0.04 | -0.23 | 3.570813 | -3 | 4.111008 | -0.002658 | 0.68054 | 0.09208 | 0.1743 | 0.12788 | 17.82772 | 0.170941939 | 0 | Neutral |
| testneg_438 | 0.05 | -0.81 | 5.57218 | -3 | 0.152708 | -3 | 0.65674 | 0.10572 | 0.06744 | 0.12788 | 39.64236 | 0.120757143 | 0 | Neutral |
| testneg_439 | -0.02 | -0.45 | 2.197733 | -3 | 1.785708 | -3 | 0.19969 | 0.1163 | 0.47975 | 0.18568 | 18.33333 | 0.074856527 | 0 | Neutral |
| testneg_440 | 0.21 | 0.15 | 6.448066 | -3 | 3.316205 | -3 | 0.40707 | 0.37988 | 0.7823 | 0.5949 | 10.41667 | 0.721172027 | 98.4 | Disease_causing |
| testneg_441 | 0.18 | 0.16 | 1.009156 | -3 | -0.214978 | -3 | 0.19898 | 0.12788 | 0.39192 | 0.09135 | 17.21847 | 0.01087037 | 0 | Neutral |
| testneg_442 | 0 | 0 | 0.438443 | 0.438443 | 0.816556 | -3 | 0.36857 | 0.36135 | 0.07195 | 0.10356 | 18.33333 | 0.03 | 0 | Neutral |
| testneg_443 | 0 | -0.01 | 0.018319 | -3 | -0.326958 | -3 | 0.3828 | 0.5569 | 0.50388 | 0.12295 | 16.9194 | 0.030646991 | 0 | Neutral |
| testneg_444 | 0.09 | -0.3 | 1.673428 | 1.673428 | 5.931499 | -3 | 0.32352 | 0.10243 | 0.11019 | 0.08132 | 26.62774 | 0.18 | 0 | Neutral |
| testneg_445 | 0.19 | -0.25 | 5.711497 | -3 | 1.477143 | -3 | 0.08186 | 0.11813 | 0.0978 | 0.26338 | 20.83333 | 0.402083333 | 22.6 | Neutral |
| testneg_446 | 0.16 | -0.37 | 3.233391 | -3 | 4.445287 | -3 | 0.2674 | 0.28725 | 0.1215 | 0.12788 | 10.83333 | 0.370966184 | 14 | Neutral |
| testneg_447 | 0.16 | -0.37 | 5.06041 | -3 | 2.474428 | -3 | 0.81054 | 0.28104 | 0.16709 | 0.08659 | 10.09538 | 0.236470245 | 0.2 | Neutral |
| testneg_448 | -0.2 | 1.26 | 0.425452 | -3 | 4.429067 | -3 | 0.23859 | 0.1202 | 0.19898 | 0.09824 | 16.88725 | 0.222666667 | 0 | Neutral |
| testneg_449 | -0.2 | 1.26 | 1.53776 | 1.53776 | 11.793086 | -3 | 0.29455 | 0.12388 | 0.35532 | 0.24703 | 11.25 | 0.596148148 | 87.6 | Disease_causing |
| testneg_450 | 0.00030303 | 0.01522727 | -0.370487 | -3 | 0.148561 | -3 | 0.61872 | 0.10479 | 0.2559 | 0.10848 | 21.63869 | 0.020164634 | 0 | Neutral |
| testneg_451 | 0.02 | -0.63 | 3.746637 | 1.966897 | 5.898353 | -3 | 0.39262 | 0.12788 | 0.42919 | 0.1092 | 11.25 | 0.3925 | 18.6 | Neutral |
| testneg_452 | 0 | 0 | -0.052835 | -3 | 6.188044 | -3 | 0.19898 | 0.13434 | 0.19898 | 0.12788 | 4.72222 | 0.135658601 | 0 | Neutral |
| testneg_453 | 0.16 | -1.37 | 6.076778 | -3 | 2.547548 | -3 | 0.33818 | 0.12788 | 0.41647 | 0.11239 | 26.84685 | 0.24 | 0.8 | Neutral |
| testneg_454 | 0.19 | 0.51 | 0.552815 | 0.552815 | 3.467861 | -3 | 0.20417 | 0.10681 | 0.37293 | 0.25159 | 10.83333 | 0.330728438 | 4.4 | Neutral |
| testneg_455 | 0 | 0 | 0.386087 | 0.386087 | 3.489285 | -3 | 0.26927 | 0.10752 | 0.05079 | 0.0924 | 26.5391 | 0.06 | 0 | Neutral |
| testneg_456 | 0 | 0 | 0.447219 | -3 | 1.626257 | -3 | 0.13884 | 0.12788 | 0.30837 | 0.19754 | 18.33333 | 0.028194875 | 0 | Neutral |
| testneg_457 | 0 | 0 | 0.038348 | -3 | -0.018405 | -3 | 0.71968 | 0.54266 | 0.25957 | 0.06969 | 10.83333 | 0.02 | 0 | Neutral |
| testneg_458 | 0.2 | -1.98 | 4.022119 | -3 | -0.137655 | -3 | 0.20692 | 0.70649 | 0.1926 | 0.12788 | 21.95857 | 0.18 | 0 | Neutral |
| testneg_459 | 0.22 | -2.66 | 5.261584 | -3 | -0.440113 | -3 | 0.49425 | 0.25538 | 0.53592 | 0.11293 | 10 | 0.21 | 0 | Neutral |
| testneg_460 | 0.18 | -1.05 | 3.963944 | -3 | 2.300256 | -3 | 0.12379 | 0.12788 | 0.14996 | 0.09121 | 27.62738 | 0.07 | 0 | Neutral |
| testneg_461 | 0 | 0 | 2.313183 | -3 | 3.997628 | 3.997628 | 0.04032 | 0.12788 | 0.66799 | 0.12788 | 18.88889 | 0.31 | 1.8 | Neutral |
| testneg_462 | 0 | 0 | 1.39106 | -3 | -0.84394 | -3 | 0.74136 | 0.1049 | 0.54895 | 0.15082 | 10.41667 | 0.02 | 0 | Neutral |
| testneg_463 | 0 | 0 | 0.389761 | -3 | 4.182553 | -3 | 0.73601 | 0.26791 | 0.3038 | 0.11278 | 18.6768 | 0.08 | 0 | Neutral |
| testneg_464 | 0.18 | 0.16 | -0.812162 | -3 | 0.275603 | -3 | 0.64922 | 0.10176 | 0.36417 | 0.21506 | 18.7315 | 0.095028999 | 0 | Neutral |
| testneg_465 | 0.06 | 0.18 | 1.528609 | -3 | 0.922605 | -3 | 0.20249 | 0.10327 | 0.2152 | 0.12788 | 18.33333 | 0 | 0 | Neutral |
| testneg_466 | 0 | 0 | 1.151992 | -3 | 3.621482 | -3 | 0.17141 | 0.1104 | 0.1004 | 0.11409 | 21.84557 | 0.041701389 | 0 | Neutral |
| testneg_467 | -0.02 | 0.28 | 2.739091 | -3 | 1.442384 | -3 | 0.23439 | 0.09253 | 0.25092 | 0.10843 | 19.09347 | 0.022050725 | 0 | Neutral |
| testneg_468 | 0.09 | -0.25 | 1.300233 | -3 | -0.851109 | -3 | 0.13119 | 0.12788 | 0.10362 | 0.12788 | 29.57012 | 0.01 | 0 | Neutral |
| testneg_469 | 0 | -0.01 | 7.854373 | -3 | 3.171839 | -1.24052 | 0.4113 | 0.2303 | 0.22582 | 0.12788 | 30.56819 | 0.609698718 | 91.8 | Disease_causing |
| testneg_470 | 0 | 0 | -0.060754 | -3 | 1.645021 | 1.645021 | 0.70946 | 0.11809 | 0.19898 | 0.14957 | 10.41667 | 0.051233333 | 0 | Neutral |
| testneg_471 | 0.22 | -2.66 | 4.55974 | -3 | 4.257584 | -3 | 0.16078 | 0.39498 | 0.05005 | 0.07051 | 11.00193 | 0.34 | 5.2 | Neutral |
| testneg_472 | 0 | 0 | -0.181123 | -3 | -1.182385 | -3 | 0.12867 | 0.11159 | 0.23879 | 0.12788 | 18.41436 | 0 | 0 | Neutral |
| testneg_473 | 0.01 | 0.79 | 0.841519 | -3 | 7.196151 | -3 | 0.81054 | 0.28104 | 0.12147 | 0.1486 | 11.77748 | 0.534186173 | 68.4 | Disease_causing |
| testneg_474 | 0.22 | -2.66 | 5.898121 | -3 | -0.436022 | -3 | 0.1296 | 0.164 | 0.24823 | 0.12788 | 13.51351 | 0.217708333 | 0.2 | Neutral |
| testneg_475 | 0 | 0 | 0.0275 | -3 | 5.815666 | -3 | 0.25179 | 0.09355 | 0.34109 | 0.2167 | 17.21847 | 0.240617195 | 1 | Neutral |
| testneg_476 | -0.15 | 0.28 | 3.501223 | -3 | 0.13329 | -3 | 0.46339 | 0.10423 | 0.05444 | 0.12788 | 13.46801 | 0.07 | 0 | Neutral |
| testneg_477 | 0 | 0 | -0.549073 | -3 | 5.127855 | -3 | 0.38016 | 0.14503 | 0.28449 | 0.10981 | 20.43633 | 0.077677798 | 0 | Neutral |
| testneg_478 | 0.05 | 0.29 | 5.633216 | -3 | 0.423127 | -3 | 0.98339 | 0.18221 | 0.25643 | 0.12788 | 14.20309 | 0.23 | 0.2 | Neutral |
| testneg_479 | -0.07 | -0.38 | 1.476163 | -3 | -0.586534 | -0.586534 | 0.19898 | 0.09395 | 0.09439 | 0.1884 | 28.54866 | 0.09 | 0 | Neutral |
| testneg_480 | 0 | 0 | 0.889458 | -3 | 0.461043 | -3 | 0.48937 | 0.22085 | 0.62864 | 0.11103 | 20.27778 | 0.02 | 0 | Neutral |
| testneg_481 | 0 | 0 | 1.928458 | -3 | 0.743493 | -3 | 0.45152 | 0.34457 | 0.11703 | 0.09364 | 19.09347 | 0.03 | 0 | Neutral |
| testneg_482 | 0 | 0 | 1.609578 | -3 | 5.845448 | 5.845448 | 0.19898 | 0.07094 | 0.06926 | 0.08399 | 28.92534 | 0.36 | 5.6 | Neutral |
| testneg_483 | -0.01 | 1.44 | 1.762821 | 1.762821 | 1.546894 | -3 | 0.1375 | 0.112 | 0.10314 | 0.08894 | 20.83333 | 0.1 | 0 | Neutral |
| testneg_484 | -0.04 | -2.01 | 5.819271 | -3 | 1.022048 | -3 | 0.29356 | 0.47581 | 0.27637 | 0.0872 | 10.22898 | 0.315 | 3.2 | Neutral |
| testneg_485 | 0.03 | -1.03 | 7.275646 | -3 | 7.190251 | -3 | 0.19898 | 0.12788 | 0.99704 | 0.17463 | 10.41667 | 0.770640351 | 99.8 | Disease_causing |
| testneg_486 | -0.09 | 2.17 | 1.005409 | 1.005409 | 1.036843 | -3 | 0.21153 | 0.11357 | 0.18102 | 0.6537 | 18.88889 | 0.21 | 0 | Neutral |
| testneg_487 | -0.02 | 0.36 | 3.195501 | -3 | -0.699644 | -3 | 0.22899 | 0.31462 | 0.3828 | 0.5569 | 10.41667 | 0.30275246 | 3.8 | Neutral |
| testneg_488 | -0.02 | 0.68 | 1.270877 | -3 | 3.866074 | -3 | 0.37292 | 0.11979 | 0.04829 | 0.09848 | 24.16296 | 0.1 | 0 | Neutral |
| testneg_489 | -0.1 | -1.9 | 3.182803 | -3 | 1.083169 | -3 | 0.15911 | 0.11022 | 0.26621 | 0.10639 | 17.53344 | 0.05 | 0 | Neutral |
| testneg_490 | -0.0125 | 0.08833333 | 1.658635 | -3 | 0.35706 | -3 | 0.15582 | 0.14216 | 0.12921 | 0.11605 | 11.25 | 0.018542373 | 0 | Neutral |
| testneg_491 | -0.03 | -0.1 | 0.447373 | -3 | -2.351627 | -3 | 0.37065 | 0.12788 | 0.16646 | 0.16629 | 26.81293 | 0.09 | 0 | Neutral |
| testneg_492 | 0 | 0 | 1.548509 | -3 | 5.567762 | 5.567762 | 0.06711 | 0.09765 | 0.74641 | 0.66512 | 6.15923 | 0.460208333 | 36.8 | Neutral |
| testneg_493 | 0 | 0 | 0.268006 | -3 | 3.832502 | -3 | 0.68568 | 0.65219 | 0.16392 | 0.12788 | 11.25 | 0.12 | 0 | Neutral |
| testneg_494 | 0.03 | -1.03 | 3.556604 | -3 | -0.361677 | -3 | 0.55043 | 0.12112 | 0.85759 | 0.1031 | 26.25896 | 0.06 | 0 | Neutral |
| testneg_495 | -0.18 | -0.16 | 1.665933 | -3 | 2.17123 | -3 | 0.19668 | 0.16042 | 0.18597 | 0.12788 | 18.6768 | 0.098638889 | 0 | Neutral |
| testneg_496 | 0 | 0 | 0.261473 | -3 | 2.797846 | -3 | 0.11006 | 0.14352 | 0.10485 | 0.09565 | 13.51351 | 0.04 | 0 | Neutral |
| testneg_497 | 0 | 0 | 0.605243 | -3 | 4.71458 | -3 | 0.40803 | 0.47186 | 0.52299 | 0.12788 | 10.41667 | 0.121191729 | 0 | Neutral |
| testneg_498 | 0 | 0 | 1.926886 | -3 | -1.803416 | -3 | 0.18431 | 0.13863 | 0.62381 | 0.12788 | 13.41972 | 0.01 | 0 | Neutral |
| testneg_499 | 0.04 | -0.23 | 4.806318 | -3 | 5.877636 | -3 | 0.20951 | 0.16278 | 0.54891 | 0.55746 | 10 | 0.852058679 | 100 | Disease_causing |
| testneg_500 | 0.07 | -0.31 | -1.289628 | -3 | 1.820594 | -3 | 0.20116 | 0.10008 | 0.16061 | 0.12426 | 29.34685 | 0.01 | 0 | Neutral |
| testneg_501 | -0.14 | 0.37 | 2.640854 | -3 | 1.808824 | 1.808824 | 0.20072 | 0.08606 | 0.2445 | 0.53069 | 10 | 0.191621573 | 0 | Neutral |
| testneg_502 | -0.15 | 1.06 | 4.179359 | -3 | 2.722615 | -3 | 0.19898 | 0.12788 | 0.60026 | 0.35923 | 10.41667 | 0.476811665 | 43.6 | Neutral |
| testneg_503 | 0 | 0 | -0.064925 | -3 | -0.545857 | -3 | 0.25924 | 0.19965 | 0.12488 | 0.12788 | 28.38164 | 0.03 | 0 | Neutral |
| testneg_504 | 0 | 0 | -0.961702 | -0.961702 | 1.096466 | -3 | 0.46979 | 0.1705 | 0.49745 | 0.1259 | 19.44444 | 0.02 | 0 | Neutral |
| testneg_505 | -0.07 | -0.38 | 2.857565 | -3 | -1.064743 | -3 | 0.23088 | 0.12788 | 0.13624 | 0.09458 | 12.46937 | 0.02 | 0 | Neutral |
| testneg_506 | 0 | 0 | 0.036223 | -3 | -0.307406 | -0.307406 | 0.9355 | 0.40418 | 0.62858 | 0.16645 | 5.75579 | 0.053832071 | 0 | Neutral |
| testneg_507 | 0.01 | -1.44 | 2.20135 | -3 | 5.114664 | -3 | 0.05677 | 0.06758 | 0.15192 | 0.14302 | 18.33333 | 0.306549254 | 3 | Neutral |
| testneg_508 | -0.03 | -0.1 | 7.121043 | -3 | 2.765662 | -0.837354 | 0.35945 | 0.12025 | 0.10438 | 0.20558 | 22.43494 | 0.622400514 | 91.6 | Disease_causing |
| testneg_509 | 0.22 | -2.66 | 4.869758 | -3 | -0.603839 | -3 | 0.2218 | 0.11533 | 0.16485 | 0.13045 | 41.13269 | 0.1 | 0 | Neutral |
| testneg_510 | -0.03 | -0.1 | 0.428176 | -3 | 2.71494 | -3 | 0.24274 | 0.13034 | 0.11264 | 0.11163 | 17.86874 | 0.03 | 0 | Neutral |
| testneg_511 | -0.04 | 0.23 | 0.791681 | -3 | -0.483923 | -3 | 0.15689 | 0.28307 | 0.23574 | 0.12788 | 16.42913 | 0.03 | 0 | Neutral |
| testneg_512 | 0.09 | -1.07 | 6.398555 | -3 | 0.34022 | -3 | 0.18488 | 0.53379 | 0.19898 | 0.12788 | 13.5211 | 0.34 | 8 | Neutral |
| testneg_513 | -0.11 | 1.95 | 3.697209 | -3 | 3.022813 | -3 | 0.2288 | 0.15849 | 0.17301 | 0.10048 | 26.18816 | 0.269732587 | 0.2 | Neutral |
| testneg_514 | 0 | 0 | -0.050489 | -3 | 0.829043 | -3 | 0.13395 | 0.29754 | 0.23141 | 0.12788 | 42.60854 | 0.022747439 | 0 | Neutral |
| testneg_515 | 0 | 0 | -0.156459 | -3 | 2.489772 | -3 | 0.10774 | 0.13117 | 0.3962 | 0.10523 | 18.33333 | 0.020357143 | 0 | Neutral |
| testneg_516 | 0 | 0 | 1.384899 | 1.384899 | 4.623093 | -3 | 0.56315 | 0.13761 | 0.0774 | 0.1702 | 30.55556 | 0.444281615 | 34.2 | Neutral |
| testneg_517 | -0.01 | 0.57 | 1.436065 | -3 | 3.677193 | -3 | 0.31773 | 0.10816 | 0.29958 | 0.11203 | 14.20922 | 0.062163185 | 0 | Neutral |
| testneg_518 | 0.04 | -0.23 | 7.022414 | -3 | 0.34099 | -3 | 0.89972 | 0.10766 | 0.33057 | 0.10549 | 20.74851 | 0.25 | 1.4 | Neutral |
| testneg_519 | 0 | 0 | 0.080104 | -3 | 3.500008 | -3 | 0.73947 | 0.2058 | 0.12513 | 0.12171 | 32.14779 | 0.113920868 | 0 | Neutral |
| testneg_520 | -0.03 | -0.1 | 6.945269 | -3 | 1.999625 | -3 | 0.09972 | 0.13992 | 0.58587 | 0.09866 | 14.87554 | 0.37 | 11.8 | Neutral |
| testneg_521 | -0.07 | -0.38 | 6.440704 | -3 | 5.84756 | -3 | 0.12131 | 0.12533 | 0.37292 | 0.11979 | 49.88402 | 0.43 | 31.2 | Neutral |
| testneg_522 | 0 | 0 | 0.18237 | -3 | -0.763372 | -3 | 0.60251 | 0.39112 | 0.19979 | 0.09328 | 18.61427 | 0.02 | 0 | Neutral |
| testneg_523 | 0 | 0 | 0.339282 | -3 | 1.479041 | -3 | 0.2547 | 0.07589 | 0.16098 | 0.12788 | 22.7957 | 0.01 | 0 | Neutral |
| testneg_524 | 0.02 | -0.68 | 2.213031 | -3 | 0.461547 | -3 | 0.99976 | 0.28689 | 0.055 | 0.20858 | 10.14695 | 0.120540936 | 0 | Neutral |
| testneg_525 | 0 | 0 | 1.446796 | -3 | 0.883022 | -3 | 0.24528 | 0.37575 | 0.1592 | 0.12788 | 28.26517 | 0.035964286 | 0 | Neutral |
| testneg_526 | 0.01 | 0.79 | -0.188304 | -3 | 2.551786 | -3 | 0.16418 | 0.12788 | 0.02329 | 0.07615 | 16.8018 | 0.030211111 | 0 | Neutral |
| testneg_527 | -0.04 | 0.23 | -0.28312 | -3 | -2.635405 | -3 | 0.06513 | 0.12353 | 0.09964 | 0.10384 | 43.628 | 0.03 | 0 | Neutral |
| testneg_528 | 0.05236364 | -0.4594545 | 9.443557 | -3 | -0.313035 | -3 | 0.07237 | 0.12788 | 0.07406 | 0.08059 | 43.07223 | 0.28 | 3.6 | Neutral |
| testneg_529 | 0 | 0 | 0.009894 | -3 | 1.052948 | -3 | 0.07996 | 0.08354 | 0.13708 | 0.81545 | 38.06813 | 0.1 | 0 | Neutral |
| testneg_530 | -0.15 | 1.06 | 1.213874 | -3 | 3.781815 | -3 | 0.04543 | 0.28103 | 0.01704 | 0.05734 | 22.00588 | 0.339821429 | 3.2 | Neutral |
| testneg_531 | 0.18 | -1.05 | 2.305068 | -3 | 4.028947 | -3 | 0.19898 | 0.12788 | 0.52957 | 0.18687 | 31.25 | 0.392017943 | 13.6 | Neutral |
| testneg_532 | 0 | 0 | 1.975201 | -3 | 0.194184 | -3 | 0.34495 | 0.16861 | 0.18744 | 0.09392 | 10 | 0.011338141 | 0 | Neutral |
| testneg_533 | 0 | 0 | 0.038426 | 0.038426 | 2.854321 | -3 | 0.54909 | 0.85056 | 0.45281 | 0.21621 | 10.83333 | 0.215522817 | 0 | Neutral |
| testneg_534 | 0.15 | -0.28 | 0.268824 | -3 | 5.504738 | -3 | 0.9924 | 0.37896 | 0.48585 | 0.66989 | 10.83333 | 0.551882267 | 74.8 | Disease_causing |
| testneg_535 | 0 | 0 | 0.739398 | -3 | 3.073373 | -3 | 0.48906 | 0.11262 | 0.96396 | 0.7628 | 31.20854 | 0.167794118 | 0 | Neutral |
| testneg_536 | -0.15 | 0.28 | 3.6541 | -3 | 4.202319 | -3 | 0.19898 | 0.37148 | 0.19898 | 0.30319 | 24.44444 | 0.807725088 | 100 | Disease_causing |
| testneg_537 | 0 | 0 | 1.125734 | -3 | 1.541854 | -3 | 0.12576 | 0.08486 | 0.1065 | 0.12788 | 17.29012 | 0.01 | 0 | Neutral |
| testneg_538 | -0.03 | -0.1 | 1.133171 | -3 | 0.521707 | -3 | 0.34279 | 0.09378 | 0.09402 | 0.12788 | 15.90686 | 0.01 | 0 | Neutral |
| testneg_539 | -0.15 | -0.04 | 1.334446 | 1.334446 | 0.231552 | -3 | 0.15514 | 0.14852 | 0.03811 | 0.05629 | 76.49098 | 0.16 | 0 | Neutral |

| | | | | | | | | | | | | | | |
|---|---|---|---|---|---|---|---|---|---|---|---|---|---|---|
| testneg_540 | -0.0014545 | -0.1627273 | 13.584842 | -3 | 7.655162 | -3 | 0.45152 | 0.34457 | 0.11775 | 0.10812 | 23.65801 | 0.838758333 | 100 | Disease_causing |
| testneg_541 | 0.04 | -0.23 | 6.877035 | 6.877035 | 1.196584 | -3 | 0.12131 | 0.12533 | 0.62273 | 0.10215 | 46.82931 | 0.6 | 93.4 | Disease_causing |
| testneg_542 | -0.08 | -0.03 | 4.020214 | -3 | 3.803915 | -3 | 0.94229 | 0.26731 | 0.31773 | 0.10816 | 13.93265 | 0.259375 | 0.8 | Neutral |
| testneg_543 | -0.06 | -0.18 | 2.444685 | -3 | -2.103676 | -3 | 0.16462 | 0.18467 | 0.23975 | 0.13861 | 10.41667 | 0.11 | 0 | Neutral |
| testneg_544 | 0.22 | -2.66 | 5.315177 | -3 | 4.276849 | -3 | 0.12091 | 0.13585 | 0.0778 | 0.0728 | 10.83333 | 0.342414467 | 7.8 | Neutral |
| testneg_545 | 0.14 | -0.69 | 1.912827 | -3 | 2.577299 | -3 | 0.1203 | 0.13872 | 0.19898 | 0.12788 | 18.91775 | 0.05 | 0 | Neutral |
| testneg_546 | -0.19 | -0.51 | 3.156196 | -3 | 0.044337 | -3 | 0.34225 | 0.12788 | 0.11775 | 0.10812 | 21.22411 | 0.05 | 0 | Neutral |
| testneg_547 | -0.14 | 1.85 | -0.371966 | -3 | 3.316487 | -3 | 0.12971 | 0.11434 | 0.18697 | 0.12232 | 12.83966 | 0.200607143 | 0 | Neutral |
| testneg_548 | -0.15 | -0.04 | 0.591299 | -3 | 3.320502 | -3 | 0.07293 | 0.12989 | 0.19898 | 0.12788 | 10.28409 | 0.13 | 0 | Neutral |
| testneg_549 | -0.08 | -0.03 | 0.461196 | -3 | -1.097042 | -3 | 0.39305 | 0.18672 | 0.10703 | 0.11611 | 24.38551 | 0.03 | 0 | Neutral |
| testneg_550 | 0 | 0 | -0.4932 | -3 | -0.811454 | -0.811454 | 0.20379 | 0.43705 | 0.72912 | 0.18218 | 10 | 0.080702248 | 0 | Neutral |
| testneg_551 | 0.04 | -0.23 | 6.104155 | -3 | -0.213236 | -3 | 0.34859 | 0.09799 | 0.13099 | 0.17453 | 20.95603 | 0.37 | 13.2 | Neutral |
| testneg_552 | 0 | -0.01 | 2.241825 | -3 | -2.165884 | -3 | 0.46318 | 0.14229 | 0.12957 | 0.12788 | 23.40643 | 0.02 | 0 | Neutral |
| testneg_553 | -0.07 | -0.01 | 2.625488 | -3 | -1.389876 | -3 | 0.87803 | 0.12788 | 0.05564 | 0.06933 | 10 | 0.021211436 | 0 | Neutral |
| testneg_554 | -0.03 | 1.03 | 1.888775 | -3 | -0.285825 | -3 | 0.09633 | 0.12788 | 0.15016 | 0.08826 | 20 | 0.03 | 0 | Neutral |
| testneg_555 | 0 | 0 | 1.069188 | -3 | 4.857402 | -3 | 0.68729 | 0.10641 | 0.31648 | 0.12788 | 34.39014 | 0.048645833 | 0 | Neutral |
| testneg_556 | 0 | 0 | 1.799258 | -3 | 3.726036 | -3 | 0.204 | 0.15424 | 0.4334 | 0.09454 | 18.17595 | 0.089918485 | 0 | Neutral |
| testneg_557 | 0 | 0 | 1.376454 | -3 | 2.905355 | -3 | 0.1679 | 0.24232 | 0.04305 | 0.06354 | 13.51351 | 0.041422414 | 0 | Neutral |
| testneg_558 | -0.14 | 1.01 | 4.425381 | -3 | 2.667661 | -3 | 0.15192 | 0.14302 | 0.10498 | 0.08421 | 11.875 | 0.220371628 | 0 | Neutral |
| testneg_559 | 0.04 | -0.89 | -0.296252 | -3 | -1.158834 | -3 | 0.09323 | 0.16732 | 0.12739 | 0.12788 | 44.89949 | 0.050526316 | 0 | Neutral |
| testneg_560 | 0.04 | -0.23 | 0.703459 | -3 | 3.336222 | -3 | 0.04483 | 0.59766 | 0.19898 | 0.12788 | 16.8018 | 0.23 | 0 | Neutral |
| testneg_561 | 0.05 | -0.81 | 2.681817 | -3 | -1.141522 | -1.141522 | 0.11956 | 0.3237 | 0.09555 | 0.09639 | 12.70833 | 0.0775 | 0 | Neutral |
| testneg_562 | 0 | 0 | 0.402508 | 0.402508 | 7.104846 | -3 | 0.29794 | 0.11106 | 0.29098 | 0.10482 | 12.70833 | 0.210662879 | 1.2 | Neutral |
| testneg_563 | 0 | -0.32 | -0.680064 | -3 | 1.311144 | -3 | 0.19898 | 0.10512 | 0.34324 | 0.085 | 19.18549 | 0.000467391 | 0 | Neutral |
| testneg_564 | 0.01 | -0.57 | 4.174712 | -3 | 1.908053 | -3 | 0.18979 | 0.07948 | 0.50287 | 0.15426 | 21.49912 | 0.171194113 | 0 | Neutral |
| testneg_565 | -0.18 | -0.16 | 3.107738 | -3 | 2.22273 | -3 | 0.49012 | 0.12138 | 0.09353 | 0.12788 | 18.6768 | 0.09 | 0 | Neutral |
| testneg_566 | 0.01 | 0.8 | 0.851144 | -3 | 3.379098 | -3 | 0.10004 | 0.12788 | 0.08437 | 0.97198 | 11.54167 | 0.248680556 | 0.4 | Neutral |
| testneg_567 | 0.07 | 0.38 | 2.371801 | -3 | 1.129332 | -3 | 0.1254 | 0.08697 | 0.37094 | 0.10762 | 20.98109 | 0.01008042 | 0 | Neutral |
| testneg_568 | -0.03 | -1.22 | 1.025171 | -3 | -0.231245 | -3 | 0.46664 | 0.09008 | 0.1077 | 0.12788 | 18.88889 | 0.03 | 0 | Neutral |
| testneg_569 | 0 | 0 | -0.50303 | -3 | 2.947434 | -3 | 0.15192 | 0.14302 | 0.27915 | 0.27706 | 6.87938 | 0.123799274 | 0.2 | Neutral |
| testneg_570 | -0.03 | -0.12 | 0.965044 | -3 | -0.072487 | -3 | 0.17264 | 0.09426 | 0.21118 | 0.10074 | 18.70407 | 0.01 | 0 | Neutral |
| testneg_571 | -0.08 | -0.03 | 4.744051 | -3 | -0.373009 | -3 | 0.19052 | 0.12788 | 0.2755 | 0.12788 | 10.17556 | 0.05 | 0 | Neutral |
| testneg_572 | 0.08 | 0.03 | -1.489251 | -3 | 5.314042 | -3 | 0.07141 | 0.12788 | 0.13615 | 0.12788 | 18.49559 | 0.11 | 0 | Neutral |
| testneg_573 | 0 | -0.01 | 0.741645 | -3 | 5.007683 | -3 | 0.04606 | 0.20522 | 0.07415 | 0.12788 | 19.09347 | 0.201339286 | 0.2 | Neutral |
| testneg_574 | -0.09 | 0.25 | 4.334907 | -3 | 3.592336 | -3 | 0.15263 | 0.12788 | 0.03297 | 0.16627 | 10 | 0.494179664 | 52.6 | Disease_causing |
| testneg_575 | 0 | 0 | 0.083867 | -3 | 3.522629 | -3 | 0.30789 | 0.15135 | 0.13693 | 0.12788 | 17.01711 | 0.065070017 | 0 | Neutral |
| testneg_576 | 0 | 0 | -0.181123 | -3 | 3.087541 | -3 | 0.12867 | 0.11159 | 0.06344 | 0.12788 | 10.41667 | 0.031414686 | 0 | Neutral |
| testneg_577 | 0 | 0 | -0.029976 | -3 | 0.484682 | -3 | 0.11731 | 0.06348 | 0.19898 | 0.12788 | 30.17377 | 0.01 | 0 | Neutral |
| testneg_578 | 0.14 | -0.69 | 5.324969 | -3 | 0.986599 | -3 | 0.835 | 0.13588 | 0.59066 | 0.10874 | 10.09538 | 0.1676874 | 0 | Neutral |
| testneg_579 | 0 | 0 | -0.489156 | -3 | 4.004011 | -3 | 0.05887 | 0.10046 | 0.37736 | 0.11686 | 24.2635 | 0.069444444 | 0 | Neutral |
| testneg_580 | 0.01 | -1.44 | 1.987251 | -3 | 0.54521 | -3 | 0.05183 | 0.18779 | 0.3718 | 0.2248 | 30.0984 | 0.267503608 | 0 | Neutral |
| testneg_581 | 0.15 | -0.28 | -1.897147 | -3 | 5.105753 | 1.261526 | 0.00971 | 0.0791 | 0.25637 | 0.37408 | 19.68254 | 0.335812068 | 3.6 | Neutral |
| testneg_582 | 0 | 0 | 0.847229 | -3 | 4.547064 | -3 | 0.14785 | 0.09084 | 0.02617 | 0.12788 | 45.40963 | 0.0525 | 0 | Neutral |
| testneg_583 | 0 | 0 | 1.257814 | -3 | 5.418765 | -3 | 0.23278 | 0.11027 | 0.47246 | 0.47221 | 11.95767 | 0.268107065 | 0.8 | Neutral |
| testneg_584 | 0 | 0 | -0.075115 | -3 | 1.323985 | -3 | 0.21576 | 0.17241 | 0.22998 | 0.08738 | 26.9055 | 0.02 | 0 | Neutral |
| testneg_585 | -0.18 | -0.16 | -0.253658 | -3 | 7.264005 | -3 | 0.07293 | 0.12989 | 0.1856 | 0.12788 | 7.07344 | 0.43 | 26 | Neutral |
| testneg_586 | 0 | 0 | -1.230837 | -1.230837 | 0.527682 | -3 | 0.14821 | 0.11262 | 0.10803 | 0.14652 | 10.41667 | 0.04 | 0 | Neutral |
| testneg_587 | 0 | -0.32 | -0.085941 | -3 | 5.82246 | -3 | 0.50276 | 0.09638 | 0.8376 | 0.24333 | 21.85185 | 0.299489964 | 2.4 | Neutral |
| testneg_588 | 0 | 0 | -0.090459 | -3 | -0.052835 | -3 | 0.15581 | 0.19351 | 0.19898 | 0.13434 | 6.59836 | 0.074593313 | 0 | Neutral |
| testneg_589 | 0.07 | -0.31 | 1.550876 | -3 | 0.416268 | -3 | 0.44615 | 0.09968 | 0.34412 | 0.08672 | 23.33333 | 0.01 | 0 | Neutral |
| testneg_590 | 0.04 | -0.23 | 7.683159 | -3 | 0.382515 | -3 | 0.89972 | 0.10766 | 0.19898 | 0.12788 | 27.17637 | 0.32 | 4 | Neutral |
| testneg_591 | 0.06054545 | -0.3370909 | 8.1928 | -3 | -0.316889 | -3 | 0.12073 | 0.10969 | 0.06957 | 0.07707 | 16.88659 | 0.274337945 | 1.2 | Neutral |
| testneg_592 | 0.07 | 0.38 | -1.041977 | -3 | 4.880745 | -3 | 0.49665 | 0.20425 | 0.08723 | 0.09503 | 26.66667 | 0.222537879 | 0 | Neutral |
| testneg_593 | 0.03 | -1.03 | 2.626617 | -3 | 2.398801 | -3 | 0.19898 | 0.12788 | 0.18175 | 0.13527 | 10 | 0.144272727 | 0 | Neutral |
| testneg_594 | 0.07 | 0.38 | -2.378865 | -3 | 2.02727 | -3 | 0.19668 | 0.16042 | 0.82269 | 0.15732 | 37.64126 | 0.130246951 | 0 | Neutral |
| testneg_595 | 0 | -0.01 | 0.730013 | -3 | -0.006282 | -0.006282 | 0.14785 | 0.09084 | 0.50287 | 0.15426 | 20.47167 | 0.051281214 | 0 | Neutral |
| testneg_596 | 0.01 | -0.57 | 0.911948 | -3 | 5.229818 | -3 | 0.19898 | 0.12788 | 0.0961 | 0.09786 | 17.21847 | 0.08 | 0 | Neutral |
| testneg_597 | 0.22 | -2.66 | 5.898121 | -3 | 0.97365 | -3 | 0.1296 | 0.164 | 0.17301 | 0.10048 | 18.6768 | 0.23 | 0.2 | Neutral |
| testneg_598 | 0.09 | -0.3 | -1.411613 | -3 | -1.286656 | -1.286656 | 0.13696 | 0.09566 | 0.10208 | 0.08867 | 22.95764 | 0.03 | 0 | Neutral |
| testneg_599 | 0.14 | -0.37 | 6.412568 | -3 | 4.004671 | -3 | 0.41193 | 0.10133 | 0.21884 | 0.16743 | 10.41667 | 0.74 | 99.4 | Disease_causing |
| testneg_600 | 0.21 | 0.15 | 6.222747 | -3 | 2.167581 | -3 | 0.1602 | 0.11262 | 0.8238 | 0.38836 | 18.19575 | 0.398668085 | 18.4 | Neutral |
| testneg_601 | 0 | 0 | 1.347564 | -3 | -1.422633 | -3 | 0.70946 | 0.11809 | 0.07598 | 0.08456 | 13.97163 | 0.01 | 0 | Neutral |
| testneg_602 | 0.07 | -0.31 | 1.843162 | -3 | 1.802906 | -3 | 0.30869 | 0.11948 | 0.33886 | 0.15644 | 11.25 | 0.05 | 0 | Neutral |
| testneg_603 | 0.05 | 1.52 | 6.426626 | -3 | 2.800312 | -3 | 0.2663 | 0.32806 | 0.09259 | 0.11165 | 12.35574 | 0.51 | 60.4 | Disease_causing |
| testneg_604 | -0.03 | -0.12 | 0.38962 | -3 | 3.68705 | -3 | 0.49745 | 0.1103 | 0.25827 | 0.21386 | 13.51351 | 0.222453342 | 0.2 | Neutral |
| testneg_605 | -0.03 | -0.1 | 0.224925 | -3 | 1.628888 | -3 | 0.03676 | 0.12788 | 0.04047 | 0.12788 | 18.33333 | 0.05 | 0 | Neutral |
| testneg_606 | 0 | 0 | -1.158589 | -3 | 4.500743 | -3 | 0.16362 | 0.11879 | 0.11337 | 0.09363 | 20.30197 | 0.06 | 0 | Neutral |
| testneg_607 | 0 | 0 | 0.291054 | -3 | 1.959161 | -3 | 0.28499 | 0.10637 | 0.18312 | 0.12788 | 13.51351 | 0 | 0 | Neutral |
| testneg_608 | 0.07 | 0.01 | -0.898947 | -3 | 2.74015 | -3 | 0.14248 | 0.12788 | 0.80314 | 0.42344 | 28.41653 | 0.14 | 0 | Neutral |
| testneg_609 | 0 | 0.73 | 1.972252 | -3 | 5.126283 | -3 | 0.07543 | 0.10211 | 0.68423 | 0.08163 | 17.53344 | 0.123645833 | 0 | Neutral |
| testneg_610 | -0.08 | -0.03 | 2.053783 | -3 | 3.870047 | -3 | 0.17285 | 0.09314 | 0.85829 | 0.4292 | 5.63643 | 0.300978469 | 3.2 | Neutral |
| testneg_611 | 0.01 | -1.44 | 2.300256 | 2.300256 | -0.115816 | -3 | 0.14996 | 0.09121 | 0.18554 | 0.12299 | 19.21665 | 0.12525 | 0 | Neutral |
| testneg_612 | 0 | 0 | -0.235365 | -3 | 0.298115 | -3 | 0.99535 | 0.38424 | 0.5481 | 0.10207 | 17.87363 | 0.040287149 | 0 | Neutral |
| testneg_613 | 0 | 0 | -0.379805 | -3 | 0.24062 | -3 | 0.12977 | 0.1128 | 0.18351 | 0.11213 | 10.30286 | 0.000121969 | 0 | Neutral |
| testneg_614 | -0.08 | -0.03 | 4.744051 | -3 | 3.968663 | 2.364808 | 0.19052 | 0.12788 | 0.16621 | 0.49845 | 7.41082 | 0.660573016 | 94 | Disease_causing |
| testneg_615 | 0.19 | 0.51 | 3.28427 | -3 | 0.073933 | -3 | 0.08982 | 0.15279 | 0.13521 | 0.10126 | 10.83333 | 0.09 | 0 | Neutral |
| testneg_616 | -0.02 | -0.13 | 3.836263 | -3 | 0.425501 | -3 | 0.19898 | 0.12788 | 0.45011 | 0.1751 | 17.21847 | 0.170333333 | 0 | Neutral |
| testneg_617 | -0.03 | -0.1 | 2.158998 | -3 | -0.111883 | -0.111883 | 0.81659 | 0.10424 | 0.06313 | 0.12788 | 15.15647 | 0.02 | 0 | Neutral |
| testneg_618 | -0.09 | 0.3 | 0.559345 | -3 | 5.194579 | -3 | 0.37542 | 0.19444 | 0.49684 | 0.29035 | 12.70833 | 0.532423049 | 65 | Disease_causing |
| testneg_619 | 0 | 0 | 0.984324 | -3 | 5.214636 | -3 | 0.21998 | 0.20297 | 0.12253 | 0.10814 | 10.73171 | 0.128214286 | 0 | Neutral |
| testneg_620 | 0.05 | 0.29 | 0.333377 | -3 | 3.446417 | 3.446417 | 0.32181 | 0.07041 | 0.29013 | 0.11368 | 20.95603 | 0.19 | 0 | Neutral |
| testneg_621 | 0.18 | -1.05 | 5.887296 | -3 | 2.556329 | -3 | 0.13918 | 0.12788 | 0.33256 | 0.14909 | 16.78342 | 0.410757846 | 27 | Neutral |
| testneg_622 | -0.03 | -0.1 | -0.316956 | -3 | 2.589372 | -3 | 0.14956 | 0.10585 | 0.11295 | 0.14675 | 14.64539 | 0.091376679 | 0 | Neutral |
| testneg_623 | 0.02 | -0.36 | 1.517891 | -3 | 7.617556 | -3 | 0.42755 | 0.09484 | 0.0935 | 0.0819 | 30.10881 | 0.28 | 0.8 | Neutral |
| testneg_624 | 0.04 | -0.23 | 3.388612 | -3 | 5.007683 | -3 | 0.32992 | 0.33227 | 0.07415 | 0.12788 | 17.21847 | 0.397392539 | 20.6 | Neutral |
| testneg_625 | 0 | 0 | -0.399895 | -3 | 1.096703 | 1.096703 | 0.5632 | 0.28442 | 0.66215 | 0.14538 | 10.83333 | 0.07094697 | 0 | Neutral |
| testneg_626 | 0.04 | -0.23 | -1.237459 | -3 | 0.879763 | 0.879763 | 0.21807 | 0.12788 | 0.68569 | 0.12463 | 12.76316 | 0.010244048 | 0 | Neutral |
| testneg_627 | 0.16 | -0.37 | 3.967892 | -3 | 1.493352 | -3 | 0.19898 | 0.12788 | 0.17364 | 0.18111 | 18.9733 | 0.220380952 | 0 | Neutral |
| testneg_628 | 0 | 0 | 0.632009 | -3 | 1.613091 | -3 | 0.16689 | 0.11395 | 0.53922 | 0.12788 | 16.8018 | 0 | 0 | Neutral |
| testneg_629 | 0 | 0 | -0.22016 | -0.22016 | 4.227814 | -3 | 0.17911 | 0.58893 | 0.15676 | 0.12788 | 40.30585 | 0.180198623 | 0 | Neutral |
| testneg_630 | 0 | 0 | -0.259811 | -3 | 7.706281 | -3 | 0.70964 | 0.12663 | 0.89972 | 0.10766 | 18.52388 | 0.23 | 1.2 | Neutral |
| testneg_631 | 0.03 | -0.85 | 0.416481 | -3 | 2.660158 | -3 | 0.04792 | 0.12788 | 0.18243 | 0.11102 | 25.6804 | 0.06 | 0 | Neutral |
| testneg_632 | -0.18 | -0.16 | 2.338395 | -3 | -0.062057 | -3 | 0.25744 | 0.2162 | 0.26489 | 0.10549 | 38.56209 | 0.077261366 | 0 | Neutral |
| testneg_633 | 0.17 | -1.22 | 4.037788 | -3 | -0.274976 | -3 | 0.08213 | 0.08913 | 0.26767 | 0.25506 | 18.44444 | 0.230801282 | 0.2 | Neutral |
| testneg_634 | 0.21 | 0.15 | 4.83533 | -3 | 0.191986 | -3 | 0.23975 | 0.13861 | 0.03235 | 0.0948 | 18.33333 | 0.08 | 0 | Neutral |
| testneg_635 | 0.14 | -1.01 | 4.075251 | -3 | 5.030268 | -3 | 0.13001 | 0.13176 | 0.39932 | 0.10693 | 30.23348 | 0.23893315 | 0.6 | Neutral |
| testneg_636 | 0 | 0 | 1.203329 | -3 | 7.372849 | -3 | 0.42585 | 0.18081 | 0.31732 | 0.27963 | 39.13978 | 0.547089044 | 69.6 | Disease_causing |
| testneg_637 | -0.05 | -0.29 | 2.013485 | 2.013485 | 1.782871 | -3 | 0.87803 | 0.12788 | 0.61834 | 0.10784 | 12.16808 | 0.07 | 0 | Neutral |
| testneg_638 | 0.16 | -1.37 | 6.47044 | -3 | 1.29795 | -3 | 0.32065 | 0.19642 | 0.5184 | 0.11889 | 10.41667 | 0.344166667 | 9.8 | Neutral |
| testneg_639 | -0.09 | 0.3 | -2.556301 | -3 | 1.9182 | 1.9182 | 0.12379 | 0.12788 | 0.43762 | 0.12788 | 21.27038 | 0.07 | 0 | Neutral |
| testneg_640 | 0 | 0 | 0.030464 | -3 | 6.985856 | -3 | 0.33795 | 0.17455 | 0.91587 | 0.10818 | 14.84841 | 0.276861569 | 2.4 | Neutral |
| testneg_641 | 0 | 0 | 1.335647 | -3 | 4.599697 | -3 | 0.54876 | 0.22335 | 0.13601 | 0.19952 | 5.69444 | 0.336513542 | 8.8 | Neutral |
| testneg_642 | 0 | 0 | -0.456898 | -3 | 0.410723 | 0.410723 | 0.06397 | 0.12788 | 0.19721 | 0.12485 | 33.95189 | 0.02 | 0 | Neutral |
| testneg_643 | -0.09 | 0.67 | 0.082232 | -3 | 2.11694 | -3 | 0.08475 | 0.19889 | 0.40403 | 0.12788 | 12.53918 | 0.100801282 | 0 | Neutral |
| testneg_644 | 0.14 | -0.69 | 3.700286 | -3 | 0.984871 | -3 | 0.11687 | 0.09069 | 0.21201 | 0.20962 | 18.30089 | 0.165692068 | 0 | Neutral |
| testneg_645 | -0.02 | 0.63 | 2.813311 | -3 | -0.061795 | -3 | 0.10809 | 0.09057 | 0.34698 | 0.12788 | 20.86969 | 0.03080303 | 0 | Neutral |
| testneg_646 | 0.06 | -0.57 | 2.053078 | -3 | -0.163315 | -3 | 0.07014 | 0.53339 | 0.31125 | 0.20061 | 12.22222 | 0.239263889 | 0 | Neutral |
| testneg_647 | 0.15 | -0.82 | 2.320954 | -3 | 1.372125 | -3 | 0.51994 | 0.37283 | 0.45075 | 0.12232 | 12.30981 | 0.08 | 0 | Neutral |

| | | | | | | | | | | | | | | |
|---|---|---|---|---|---|---|---|---|---|---|---|---|---|---|
| testneg_648 | -0.19 | -0.51 | 1.29341 | -3 | 1.6888 | -3 | 0.11773 | 0.11669 | 0.226 | 0.28784 | 35.25602 | 0.13 | 0 | Neutral |
| testneg_649 | 0.03 | -1.03 | 2.793619 | -3 | 0.272659 | 0.272659 | 0.13011 | 0.14157 | 0.17093 | 0.23281 | 11.61113 | 0.16 | 0 | Neutral |
| testneg_650 | 0.01 | 0.8 | 1.656526 | -3 | 0.03956 | -3 | 0.19898 | 0.09972 | 0.07961 | 0.11474 | 20.32051 | 0.01 | 0 | Neutral |
| testneg_651 | 0 | 0 | 0.785121 | -3 | -2.410599 | -3 | 0.10004 | 0.12788 | 0.59691 | 0.11049 | 15.77966 | 0.01 | 0 | Neutral |
| testneg_652 | 0 | 0 | 0.342408 | -3 | 6.817459 | -3 | 0.5255 | 0.15968 | 0.06622 | 0.12378 | 24.08925 | 0.271769231 | 2.8 | Neutral |
| testneg_653 | 0.18 | -1.05 | 6.639367 | -3 | -2.188525 | -2.188525 | 0.36324 | 0.15729 | 0.07282 | 0.12788 | 19.48711 | 0.274166667 | 2.2 | Neutral |
| testneg_654 | 0.04 | -0.23 | 4.740439 | -3 | 5.898475 | -3 | 0.28156 | 0.11579 | 0.35504 | 0.12788 | 22.41883 | 0.31081729 | 6.8 | Neutral |
| testneg_655 | 0 | 0 | 0.191318 | -3 | 4.014482 | -3 | 0.96113 | 0.24974 | 0.95992 | 0.10958 | 17.77778 | 0.105695825 | 0 | Neutral |
| testneg_656 | 0.07 | 0.38 | -0.459078 | -3 | -0.055675 | -3 | 0.19214 | 0.12788 | 0.35051 | 0.11664 | 29.09389 | 0.01 | 0 | Neutral |
| testneg_657 | 0.01 | 0.79 | 5.361706 | -3 | 6.89099 | 6.89099 | 0.835 | 0.13588 | 0.33418 | 0.09979 | 19.7626 | 0.73 | 100 | Disease_causing |
| testneg_658 | 0.05 | -0.81 | 3.142994 | -3 | 2.548387 | -3 | 0.26668 | 0.11598 | 0.05221 | 0.12788 | 13.14653 | 0.064779762 | 0 | Neutral |
| testneg_659 | 0 | 0 | 1.323079 | -3 | 1.65767 | 1.65767 | 0.5103 | 0.10923 | 0.89992 | 0.10231 | 19.13153 | 0.02 | 0 | Neutral |
| testneg_660 | 0.05 | -0.81 | 2.270715 | -3 | 2.199525 | -3 | 0.04441 | 0.13603 | 0.1909 | 0.14279 | 16.48772 | 0.150127428 | 0 | Neutral |
| testneg_661 | 0.07 | 0.01 | 2.947718 | -3 | 6.296362 | -3 | 0.5068 | 0.12276 | 0.19861 | 0.12057 | 20.89414 | 0.228055556 | 0.6 | Neutral |
| testneg_662 | 0.06 | -0.57 | 4.857454 | -3 | -0.646746 | -3 | 0.19898 | 0.12788 | 0.12073 | 0.10969 | 19.85862 | 0.04 | 0 | Neutral |
| testneg_663 | 0 | 0 | 0.147043 | -3 | 5.156881 | -3 | 0.14668 | 0.12788 | 0.8169 | 0.23301 | 11.25 | 0.192019969 | 0 | Neutral |
| testneg_664 | 0 | -0.01 | 1.062204 | -3 | 5.980097 | -3 | 0.73731 | 0.26107 | 0.27164 | 0.10265 | 19.44444 | 0.11220333 | 0 | Neutral |
| testneg_665 | 0 | 0 | 0.61552 | -3 | 7.268055 | -3 | 0.53654 | 0.22867 | 0.072 | 0.3038 | 12.71739 | 0.540925198 | 68.8 | Disease_causing |
| testneg_666 | 0.16 | -0.37 | 1.179018 | -3 | 2.702357 | -3 | 0.62645 | 0.12788 | 0.94674 | 0.12413 | 19.37853 | 0.06 | 0 | Neutral |
| testneg_667 | -0.02 | 0.68 | 1.730125 | -3 | 1.764776 | -3 | 0.10671 | 0.10867 | 0.6975 | 0.53794 | 10.41667 | 0.07 | 0 | Neutral |
| testneg_668 | 0.18 | -1.05 | 2.976331 | -3 | 7.113432 | -3 | 0.08775 | 0.12788 | 0.25485 | 0.12788 | 18.81606 | 0.382794118 | 17.8 | Neutral |
| testneg_669 | -0.15 | -0.04 | 0.939929 | -3 | 2.796855 | -3 | 0.18701 | 0.12788 | 0.2288 | 0.15849 | 13.51351 | 0.185637395 | 0 | Neutral |
| testneg_670 | 0 | 0 | -0.060754 | -3 | 1.366001 | -3 | 0.70946 | 0.11809 | 0.04305 | 0.06354 | 16.95817 | 0.02 | 0 | Neutral |
| testneg_671 | 0 | -0.01 | 0.522695 | -3 | 3.17007 | -3 | 0.84616 | 0.1471 | 0.12377 | 0.12788 | 29.08136 | 0.089443452 | 0 | Neutral |
| testneg_672 | -0.05 | -0.29 | 2.310369 | -3 | 3.668266 | -3 | 0.54895 | 0.15082 | 0.63645 | 0.14596 | 11.25 | 0.33026944 | 5.4 | Neutral |
| testneg_673 | 0 | 0 | 0.922275 | -3 | 1.8794 | -3 | 0.26586 | 0.10825 | 0.54711 | 0.11809 | 11.875 | 0 | 0 | Neutral |
| testneg_674 | 0 | 0 | 1.252726 | -3 | -0.434903 | -3 | 0.54231 | 0.15254 | 0.40981 | 0.21807 | 29.16667 | 0.082369048 | 0 | Neutral |
| testneg_675 | 0 | 0 | 0.276668 | -3 | 3.719978 | -3 | 0.35376 | 0.09478 | 0.63134 | 0.12788 | 20.83333 | 0.0595625 | 0 | Neutral |
| testneg_676 | 0.21 | 0.15 | 6.924355 | -3 | 1.871099 | -3 | 0.38002 | 0.22508 | 0.87611 | 0.16759 | 13.75309 | 0.509552018 | 58.6 | Disease_causing |
| testneg_677 | -0.15 | -0.04 | 0.012434 | -3 | -2.769769 | -3 | 0.08629 | 0.12788 | 0.05291 | 0.08633 | 18.6768 | 0.06 | 0 | Neutral |
| testneg_678 | 0 | 0 | 0.300937 | -3 | -0.649424 | -3 | 0.46318 | 0.14229 | 0.09515 | 0.07784 | 15.32878 | 0.01 | 0 | Neutral |
| testneg_679 | 0.17 | 0.38 | 3.132136 | -3 | 5.050499 | 5.050499 | 0.0105 | 0.12788 | 0.12819 | 0.1229 | 30.16935 | 0.41 | 16.6 | Neutral |
| testneg_680 | 0.15 | -0.28 | 0.118932 | 0.118932 | -2.464297 | -2.464297 | 0.19567 | 0.09606 | 0.11524 | 0.12788 | 15.37063 | 0.02 | 0 | Neutral |
| testneg_681 | -0.18 | 1.05 | 4.595565 | -3 | 1.778612 | -3 | 0.19898 | 0.12788 | 0.0968 | 0.15592 | 13.02635 | 0.296693723 | 1.8 | Neutral |
| testneg_682 | 0 | 0 | -0.725688 | -3 | 2.550986 | -3 | 0.17285 | 0.09314 | 0.08408 | 0.12788 | 18.33333 | 0.02 | 0 | Neutral |
| testneg_683 | -0.02 | -0.45 | 5.514554 | -3 | 0.542687 | -3 | 0.19898 | 0.12788 | 0.5184 | 0.11889 | 41.57338 | 0.130545906 | 0 | Neutral |
| testneg_684 | 0 | 0 | 1.527665 | -3 | 1.432285 | -3 | 0.33713 | 0.12788 | 0.40663 | 0.24974 | 10.22898 | 0.02027027 | 0 | Neutral |
| testneg_685 | 0.03 | 0.12 | 2.135113 | -3 | -1.648545 | -3 | 0.40391 | 0.13431 | 0.79857 | 0.12788 | 4.72222 | 0.02 | 0 | Neutral |
| testneg_686 | -0.03 | -0.1 | 2.445411 | -3 | 0.639164 | -3 | 0.22179 | 0.09831 | 0.2521 | 0.1784 | 21.38889 | 0.09 | 0 | Neutral |
| testneg_687 | 0 | -0.32 | 0.345315 | -3 | 2.944773 | -3 | 0.09964 | 0.10384 | 0.42666 | 0.12788 | 17.26546 | 0.020741758 | 0 | Neutral |
| testneg_688 | 0 | 0 | 0.103447 | 0.103447 | 0.327712 | -3 | 0.15813 | 0.61197 | 0.57517 | 0.10873 | 26.13031 | 0.07 | 0 | Neutral |
| testneg_689 | 0.21 | 0.15 | 2.619874 | -3 | 2.151559 | -3 | 0.68423 | 0.08163 | 0.82591 | 0.12207 | 16.70011 | 0.07 | 0 | Neutral |
| testneg_690 | 0.22 | -2.66 | 2.115161 | -3 | 1.272946 | -3 | 0.34859 | 0.09799 | 0.12977 | 0.1128 | 10.30286 | 0.062575758 | 0 | Neutral |
| testneg_691 | -0.02 | -0.45 | 3.546645 | -3 | 0.118414 | -3 | 0.18502 | 0.21329 | 0.20513 | 0.09801 | 14.39795 | 0.060513158 | 0 | Neutral |
| testneg_692 | 0.21 | 0.15 | 4.605481 | -3 | 3.484642 | -3 | 0.53888 | 0.09904 | 0.10485 | 0.09565 | 13.51351 | 0.215708333 | 0 | Neutral |
| testneg_693 | 0.2 | -1.26 | 1.20337 | -3 | 0.528801 | -3 | 0.03072 | 0.27473 | 0.12136 | 0.27707 | 39.60168 | 0.307333333 | 1 | Neutral |
| testneg_694 | 0 | 0 | 2.314807 | -3 | -0.370487 | -3 | 0.44615 | 0.09968 | 0.61872 | 0.10479 | 18.33333 | 0.01 | 0 | Neutral |
| testneg_695 | -0.15 | 1.06 | 2.17146 | -3 | 2.83231 | -3 | 0.07447 | 0.23492 | 0.13001 | 0.13176 | 19.44444 | 0.41308658 | 18.4 | Neutral |
| testneg_696 | 0 | 0 | 0.379738 | -3 | -0.232414 | -3 | 0.11731 | 0.06348 | 0.2168 | 0.11297 | 43.96721 | 0.01 | 0 | Neutral |
| testneg_697 | 0 | 0 | 0.484587 | -3 | 3.05843 | 3.05843 | 0.30056 | 0.12272 | 0.74136 | 0.1049 | 38.83469 | 0.12 | 0 | Neutral |
| testneg_698 | -0.22 | 2.66 | -1.189105 | -3 | 1.01961 | -3 | 0.25561 | 0.11004 | 0.53016 | 0.10373 | 18.6768 | 0.16 | 0 | Neutral |
| testneg_699 | 0.04 | -0.23 | 0.666384 | -3 | 0.150956 | -3 | 0.30669 | 0.11489 | 0.10703 | 0.11611 | 24.5082 | 0.01 | 0 | Neutral |
| testneg_700 | -0.15 | -0.04 | 3.116374 | -3 | 1.849259 | 1.849259 | 0.19898 | 0.12788 | 0.41261 | 0.12096 | 27.67636 | 0.11 | 0 | Neutral |
| testneg_701 | 0.02 | -0.63 | -2.196989 | -3 | 0.165993 | -3 | 0.02759 | 0.12458 | 0.2339 | 0.0998 | 16.8018 | 0.04 | 0 | Neutral |
| testneg_702 | 0 | 0 | -0.136916 | -3 | -0.328402 | -3 | 0.04853 | 0.4186 | 0.09281 | 0.13929 | 17.37434 | 0.233309693 | 0 | Neutral |
| testneg_703 | 0.03 | 0.1 | 6.254981 | -3 | 0.035553 | 0.035553 | 0.95993 | 0.15588 | 0.15414 | 0.17421 | 10.41667 | 0.466988095 | 42.4 | Neutral |
| testneg_704 | 0 | 0 | 0.598467 | -3 | 4.530931 | -3 | 0.52836 | 0.12785 | 0.16409 | 0.12788 | 13.94832 | 0.053711709 | 0 | Neutral |
| testneg_705 | 0.04 | -0.23 | 6.188656 | -3 | 1.869068 | -3 | 0.29624 | 0.1067 | 0.65717 | 0.16156 | 21.32417 | 0.399324252 | 21.2 | Neutral |
| testneg_706 | 0 | 0 | 1.078588 | -3 | 2.225367 | 0.533924 | 0.11329 | 0.12116 | 0.15839 | 0.10522 | 14.78889 | 0.01 | 0 | Neutral |
| testneg_707 | -0.01 | -0.79 | -1.595595 | -3 | 0.252727 | -3 | 0.53172 | 0.12788 | 0.13985 | 0.08349 | 10.26553 | 0.03 | 0 | Neutral |
| testneg_708 | -0.02 | -0.45 | 5.786078 | -3 | -0.049164 | -0.049164 | 0.42306 | 0.86891 | 0.12322 | 0.1246 | 20.83333 | 0.357840909 | 6.8 | Neutral |
| testneg_709 | 0.03 | 0.1 | 4.732985 | -3 | -0.500727 | -3 | 0.23506 | 0.24555 | 0.07684 | 0.09138 | 16.8018 | 0.08 | 0 | Neutral |
| testneg_710 | -0.03 | -0.1 | 4.57636 | -3 | 4.6688 | -3 | 0.11773 | 0.11669 | 0.06723 | 0.10296 | 46.50157 | 0.188911765 | 0.2 | Neutral |
| testneg_711 | 0 | 0 | -0.136163 | -3 | 0.070243 | -3 | 0.0513 | 0.33622 | 0.14486 | 0.33725 | 18.33333 | 0.251972037 | 0 | Neutral |
| testneg_712 | 0 | -0.32 | 2.076532 | -3 | -0.972856 | -3 | 0.08775 | 0.12788 | 0.19898 | 0.12788 | 13.46801 | 0.01 | 0 | Neutral |
| testneg_713 | 0 | 0 | 0.184247 | -3 | 5.537244 | -3 | 0.47787 | 0.25991 | 0.19898 | 0.09972 | 10.41667 | 0.12 | 0 | Neutral |
| testneg_714 | 0.05 | -0.81 | 5.57218 | -3 | 0.400947 | -3 | 0.65674 | 0.10572 | 0.0328 | 0.0691 | 41.03395 | 0.12 | 0 | Neutral |
| testneg_715 | 0 | 0 | 0.462587 | -3 | 1.901205 | -3 | 0.26014 | 0.12378 | 0.20914 | 0.10089 | 18.08385 | 0 | 0 | Neutral |
| testneg_716 | -0.08 | -0.03 | 6.549241 | -3 | 1.970058 | -3 | 0.04142 | 0.09564 | 0.06067 | 0.07796 | 29.19177 | 0.26 | 1 | Neutral |
| testneg_717 | 0 | 0 | 0.254162 | -3 | 0.760045 | -3 | 0.94807 | 0.14977 | 0.10176 | 0.12788 | 8.97898 | 0.02 | 0 | Neutral |
| testneg_718 | 0 | 0 | 1.283706 | -3 | 1.384899 | 1.384899 | 0.16856 | 0.19221 | 0.56315 | 0.13761 | 21.81093 | 0.102916667 | 0 | Neutral |
| testneg_719 | 0 | 0 | 1.170621 | -3 | 1.932311 | 1.932311 | 0.84616 | 0.1471 | 0.97673 | 0.22976 | 10.83333 | 0.07892316 | 0 | Neutral |
| testneg_720 | 0.21 | 0.15 | 1.912237 | -3 | 3.967152 | -3 | 0.12511 | 0.11897 | 0.95992 | 0.10958 | 25.98109 | 0.110733333 | 0 | Neutral |
| testneg_721 | 0.04 | -0.23 | 4.146945 | -3 | 3.82459 | -3 | 0.16965 | 0.25991 | 0.09893 | 0.1266 | 26.02198 | 0.282857143 | 2.2 | Neutral |
| testneg_722 | -0.03 | 1.03 | 3.441233 | -3 | 5.617032 | -3 | 0.22046 | 0.12788 | 0.18328 | 0.12788 | 18.63948 | 0.306044444 | 3.4 | Neutral |
| testneg_723 | 0 | 0 | 1.372857 | -3 | 0.454215 | -3 | 0.14378 | 0.41915 | 0.13815 | 0.40001 | 11.13095 | 0.140190374 | 0 | Neutral |
| testneg_724 | 0.07 | 0.01 | -0.323717 | -3 | 1.697009 | -3 | 0.23691 | 0.1015 | 0.52771 | 0.10974 | 28.65272 | 0.01 | 0 | Neutral |
| testneg_725 | 0 | 0 | 2.189786 | -3 | -0.220959 | -3 | 0.54876 | 0.22335 | 0.49083 | 0.14252 | 11.71717 | 0.061309034 | 0 | Neutral |
| testneg_726 | 0 | 0 | -0.22828 | -3 | 0.944356 | -3 | 0.31694 | 0.13475 | 0.32532 | 0.09651 | 17.77778 | 0.01 | 0 | Neutral |
| testneg_727 | -0.03 | -0.1 | -0.76618 | -3 | 2.652503 | -3 | 0.1503 | 0.12788 | 0.08059 | 0.09576 | 38.4322 | 0.03005618 | 0 | Neutral |
| testneg_728 | 0 | 0 | 0.819832 | -3 | 3.193004 | -3 | 0.19898 | 0.12788 | 0.16926 | 0.11871 | 28.94043 | 0.037014286 | 0 | Neutral |
| testneg_729 | 0 | 0 | 0.229422 | -3 | 2.16064 | -3 | 0.35428 | 0.12971 | 0.16734 | 0.15176 | 17.88889 | 0.06013377 | 0 | Neutral |
| testneg_730 | -0.04 | 0.23 | 0.791681 | -3 | 0.021357 | 0.021357 | 0.15689 | 0.28307 | 0.17935 | 0.36953 | 12.70833 | 0.198652899 | 0 | Neutral |
| testneg_731 | -0.08 | -0.03 | 0.20058 | -3 | 0.840567 | -3 | 0.1897 | 0.12459 | 0.11746 | 0.1471 | 26.29328 | 0.070037037 | 0 | Neutral |
| testneg_732 | -0.1 | -1.9 | 3.828665 | -3 | 4.671506 | -3 | 0.61491 | 0.11954 | 0.15856 | 0.10994 | 10.73722 | 0.230980808 | 0.2 | Neutral |
| testneg_733 | -0.17 | 1.22 | -2.338688 | -2.338688 | 0.97098 | -3 | 0.33129 | 0.09957 | 0.75415 | 0.29199 | 14.16667 | 0.142356062 | 0 | Neutral |
| testneg_734 | 0 | 0 | -0.181123 | -3 | 3.300854 | -3 | 0.12867 | 0.11159 | 0.43977 | 0.17284 | 19.09347 | 0.144030897 | 0 | Neutral |
| testneg_735 | 0.21 | 0.15 | 1.631249 | -3 | -0.856582 | -3 | 0.47166 | 0.13548 | 0.2074 | 0.09696 | 20.5518 | 0.04 | 0 | Neutral |
| testneg_736 | -0.03 | -0.1 | 2.436626 | -3 | 0.840567 | -3 | 0.26181 | 0.12875 | 0.11746 | 0.1471 | 29.86111 | 0.07422619 | 0 | Neutral |
| testneg_737 | 0 | -0.01 | 0.567087 | -3 | 1.042194 | -3 | 0.46757 | 0.14122 | 0.19898 | 0.12788 | 29.45989 | 0.010102614 | 0 | Neutral |
| testneg_738 | 0.02 | -0.36 | -0.01145 | -0.01145 | 5.955605 | -3 | 0.78462 | 0.12258 | 0.16311 | 0.09662 | 21.86269 | 0.134158748 | 0 | Neutral |
| testneg_739 | -0.17 | -0.38 | 0.128896 | -3 | 0.991883 | -3 | 0.67331 | 0.10118 | 0.26816 | 0.13347 | 20.86969 | 0.12 | 0 | Neutral |
| testneg_740 | 0 | -0.01 | 2.187535 | -3 | 1.386411 | 0.561005 | 0.93654 | 0.38765 | 0.09527 | 0.09424 | 27.40959 | 0.050178798 | 0 | Neutral |
| testneg_741 | 0.04 | -0.89 | -0.296252 | -3 | 0.978606 | -3 | 0.09323 | 0.16732 | 0.04046 | 0.06599 | 25.92054 | 0.052258403 | 0 | Neutral |
| testneg_742 | -0.01 | -0.8 | 7.383324 | -3 | -1.289628 | -3 | 0.22761 | 0.14229 | 0.20116 | 0.10008 | 12.67392 | 0.38 | 13.8 | Neutral |
| testneg_743 | 0 | 0 | 0.098641 | -3 | 2.390877 | -3 | 0.07121 | 0.06917 | 0.43691 | 0.12794 | 10.25938 | 0.02 | 0 | Neutral |
| testneg_744 | -0.18 | 1.05 | 2.40944 | 1.660423 | -1.715522 | -3 | 0.17075 | 0.12788 | 0.19898 | 0.12788 | 20.71879 | 0.12 | 0 | Neutral |
| testneg_745 | -0.15 | -0.04 | 2.006498 | -3 | 1.173281 | -3 | 0.13647 | 0.11806 | 0.04983 | 0.08759 | 20.42553 | 0.040350985 | 0 | Neutral |
| testneg_746 | 0 | 0 | 0.39745 | -3 | 0.201766 | -3 | 0.11404 | 0.12788 | 0.16388 | 0.14439 | 23.45603 | 0.042228395 | 0 | Neutral |
| testneg_747 | -0.02 | 0.36 | 2.804789 | -3 | 2.896502 | -3 | 0.53313 | 0.14072 | 0.18096 | 0.12788 | 38.65749 | 0.100615098 | 0 | Neutral |
| testneg_748 | 0 | 0 | -1.556747 | -3 | -1.024972 | -3 | 0.52768 | 0.12788 | 0.23955 | 0.97212 | 16.25133 | 0.08 | 0 | Neutral |
| testneg_749 | 0.07 | 0.38 | -2.70564 | -3 | 5.901884 | -3 | 0.94229 | 0.26731 | 0.09184 | 0.38254 | 11.25 | 0.448078573 | 34 | Neutral |
| testneg_750 | 0 | -0.01 | 2.631127 | -3 | 6.440155 | -3 | 0.16068 | 0.12788 | 0.10161 | 0.04472 | 18.58924 | 0.19 | 0.2 | Neutral |
| testneg_751 | 0.21 | 0.15 | 4.635288 | -3 | 0.623338 | 0.623338 | 0.38749 | 0.1076 | 0.09707 | 0.12788 | 19.3286 | 0.07 | 0 | Neutral |
| testneg_752 | 0 | 0 | -0.399895 | -3 | -0.501499 | -3 | 0.5632 | 0.28442 | 0.16716 | 0.10951 | 25.9387 | 0.0301547 | 0 | Neutral |
| testneg_753 | -0.16 | -0.08 | 3.702072 | -3 | 2.117525 | -3 | 0.57019 | 0.38533 | 0.37463 | 0.23021 | 11.25 | 0.550653884 | 75.2 | Disease_causing |
| testneg_754 | -0.03 | -0.12 | 1.782773 | -3 | 3.381673 | 3.381673 | 0.42544 | 0.11809 | 0.23578 | 0.12788 | 16.8018 | 0.171111111 | 0 | Neutral |
| testneg_755 | 0 | 0 | 1.103204 | -3 | 0.930193 | -3 | 0.57199 | 0.11262 | 0.09439 | 0.07415 | 20.78308 | 0.02 | 0 | Neutral |

| | | | | | | | | | | | | | | |
|---|---|---|---|---|---|---|---|---|---|---|---|---|---|---|
| testneg_756 | -0.05 | -0.29 | 3.715243 | -3 | -0.01248 | -3 | 0.25516 | 0.12788 | 0.61231 | 0.13362 | 19.09347 | 0.1 | 0 | Neutral |
| testneg_757 | 0.09 | -0.3 | 5.118069 | -3 | 1.01469 | -3 | 0.33818 | 0.12788 | 0.39019 | 0.18821 | 10.83333 | 0.278961153 | 2 | Neutral |
| testneg_758 | 0.04 | -0.23 | 4.420773 | 2.05038 | 3.036954 | -3 | 0.09416 | 0.17821 | 0.36541 | 0.27749 | 21.94444 | 0.745036363 | 99.8 | Disease_causing |
| testneg_759 | -0.03 | -0.1 | 3.331301 | -3 | 6.971224 | -3 | 0.17527 | 0.13941 | 0.20632 | 0.12788 | 24.83921 | 0.433272306 | 29.8 | Neutral |
| testneg_760 | -0.17 | -0.38 | 2.902612 | -3 | 0.188594 | -3 | 0.37463 | 0.23021 | 0.35499 | 0.12052 | 30.96351 | 0.11 | 0 | Neutral |
| testneg_761 | 0.02 | -0.36 | 1.451698 | -3 | 1.995788 | -3 | 0.18979 | 0.07948 | 0.19043 | 0.07855 | 20.41417 | 0.01 | 0 | Neutral |
| testneg_762 | 0.03 | 1.22 | 4.319571 | -3 | 5.496455 | -3 | 0.36324 | 0.15729 | 0.17301 | 0.10048 | 18.6768 | 0.427696914 | 27.4 | Neutral |
| testneg_763 | 0 | 0 | 0.332693 | -3 | 3.56934 | -3 | 0.53292 | 0.32174 | 0.25884 | 0.12788 | 19.60216 | 0.098722252 | 0 | Neutral |
| testneg_764 | 0 | 0 | -0.237972 | -3 | 0.144369 | -3 | 0.21972 | 0.12788 | 0.12403 | 0.12788 | 19.86796 | 0.001666667 | 0 | Neutral |
| testneg_765 | -0.09 | 0.3 | -0.522358 | -3 | 0.385855 | 0.385855 | 0.12836 | 0.09736 | 0.10153 | 0.13343 | 16.8018 | 0.110083333 | 0 | Neutral |
| testneg_766 | 0 | 0 | 0.431179 | -3 | -0.592626 | -3 | 0.47751 | 0.2062 | 0.143 | 0.11236 | 14.95396 | 0.020408333 | 0 | Neutral |
| testneg_767 | 0 | 0 | -0.481968 | -0.481968 | 3.470004 | -3 | 0.67331 | 0.10118 | 0.18067 | 0.14759 | 12.87484 | 0.167547414 | 0 | Neutral |
| testneg_768 | -0.15 | -0.04 | 1.218539 | -3 | 1.720276 | -3 | 0.16068 | 0.12788 | 0.10781 | 0.09901 | 29.34685 | 0.04 | 0 | Neutral |
| testneg_769 | 0 | 0 | 0.297806 | -3 | -1.259442 | -3 | 0.19898 | 0.10362 | 0.24185 | 0.12788 | 14.16667 | 0 | 0 | Neutral |
| testneg_770 | 0 | 0 | -0.136916 | -3 | 4.774952 | -3 | 0.04853 | 0.4186 | 0.10472 | 0.10014 | 15 | 0.21 | 0 | Neutral |
| testneg_771 | 0.04 | -1.61 | 3.074697 | -3 | 0.566382 | -3 | 0.37282 | 0.07943 | 0.14349 | 0.27976 | 15.25941 | 0.183452381 | 0 | Neutral |
| testneg_772 | 0.18 | -1.05 | 6.665788 | -3 | 2.01147 | -3 | 0.20431 | 0.17304 | 0.07887 | 0.07487 | 31.22387 | 0.31 | 3.8 | Neutral |
| testneg_773 | 0 | 0 | -0.194983 | -3 | 0.734729 | -3 | 0.84238 | 0.12559 | 0.58587 | 0.09866 | 24.2785 | 0.02 | 0 | Neutral |
| testneg_774 | 0 | 0 | 0.015054 | -3 | 1.969356 | -3 | 0.12196 | 0.12788 | 0.30876 | 0.28825 | 11.25 | 0.039605742 | 0 | Neutral |
| testneg_775 | 0 | 0 | -0.344792 | -3 | 0.847539 | -3 | 0.19898 | 0.58572 | 0.65564 | 0.11468 | 10.29532 | 0.04 | 0 | Neutral |
| testneg_776 | -0.09 | 0.3 | 0.912537 | -3 | 3.867932 | -3 | 0.2049 | 0.12678 | 0.1725 | 0.09643 | 23.40224 | 0.099547063 | 0 | Neutral |
| testneg_777 | 0.07 | 0.38 | 1.224856 | -3 | 3.399462 | -3 | 0.10176 | 0.12788 | 0.05472 | 0.12788 | 33.21306 | 0.06 | 0 | Neutral |
| testneg_778 | -0.03 | -0.1 | 4.076957 | 4.076957 | 2.322245 | -3 | 0.38324 | 0.25193 | 0.30734 | 0.12788 | 12.70833 | 0.510219732 | 61.4 | Disease_causing |
| testneg_779 | 0 | 0 | -0.181123 | -3 | -0.008346 | -3 | 0.12867 | 0.11159 | 0.14862 | 0.12788 | 22.20588 | 0.0025 | 0 | Neutral |
| testneg_780 | 0 | 0 | 0.916236 | -3 | 0.747659 | -3 | 0.37499 | 0.17803 | 0.08572 | 0.13089 | 4.44444 | 0.03 | 0 | Neutral |
| testneg_781 | 0.14 | -1.01 | 0.712749 | -3 | 4.593525 | -3 | 0.12203 | 0.13515 | 0.57564 | 0.12788 | 10.41667 | 0.143541667 | 0 | Neutral |
| testneg_782 | 0 | 0 | 1.095746 | -3 | 3.271853 | -3 | 0.24274 | 0.13034 | 0.60152 | 0.12166 | 10.24108 | 0.031356061 | 0 | Neutral |
| testneg_783 | 0 | 0 | -0.445161 | -0.445161 | 3.377364 | -3 | 0.09728 | 0.12788 | 0.82148 | 0.14847 | 34.16548 | 0.14225 | 0 | Neutral |
| testneg_784 | 0 | 0 | 0.245964 | -3 | 5.928464 | -3 | 0.74099 | 0.44443 | 0.11316 | 0.12788 | 10.22898 | 0.14114532 | 0 | Neutral |
| testneg_785 | -0.05 | 0.81 | -1.696114 | -3 | -0.094022 | -3 | 0.19898 | 0.82835 | 0.8256 | 0.11062 | 30.04169 | 0.13 | 0 | Neutral |
| testneg_786 | -0.09 | 0.3 | 0.631708 | -3 | 0.20058 | -3 | 0.34546 | 0.0932 | 0.1897 | 0.12459 | 29.16667 | 0.020801282 | 0 | Neutral |
| testneg_787 | -0.05 | -0.29 | -2.038213 | -3 | 0.44595 | -3 | 0.64256 | 0.12241 | 0.15019 | 0.10342 | 18.09741 | 0.02 | 0 | Neutral |
| testneg_788 | 0 | 0 | 0.522479 | -3 | -0.448326 | -0.562184 | 0.24949 | 0.29709 | 0.03466 | 0.19518 | 27.47562 | 0.160438679 | 0 | Neutral |
| testneg_789 | -0.02 | 0.63 | 5.105103 | -3 | -0.176493 | -3 | 0.61211 | 0.11059 | 0.95978 | 0.09493 | 9.78453 | 0.12 | 0 | Neutral |
| testneg_790 | -0.04 | 0.23 | -1.561199 | -3 | -0.143224 | -3 | 0.07617 | 0.07182 | 0.12176 | 0.09356 | 18.57252 | 0.03 | 0 | Neutral |
| testneg_791 | 0 | 0 | 0.050569 | -3 | 2.251984 | -3 | 0.16263 | 0.53493 | 0.15685 | 0.17839 | 17.11441 | 0.190330313 | 0.2 | Neutral |
| testneg_792 | 0.16 | -0.37 | 1.160968 | -3 | 1.520495 | -3 | 0.98565 | 0.23629 | 0.50921 | 0.21466 | 10.41667 | 0.14 | 0 | Neutral |
| testneg_793 | 0.04 | -0.23 | 1.625356 | -3 | -1.390634 | -3 | 0.5065 | 0.20697 | 0.29672 | 0.10485 | 21.2963 | 0.04 | 0 | Neutral |
| testneg_794 | 0 | -0.01 | 3.197244 | -3 | -0.421907 | -3 | 0.09611 | 0.07772 | 0.49276 | 0.10648 | 18.6768 | 0.03 | 0 | Neutral |
| testneg_795 | -0.08 | -0.03 | 2.788816 | -3 | -0.545857 | -3 | 0.19898 | 0.14896 | 0.12488 | 0.12788 | 34.54497 | 0.04 | 0 | Neutral |
| testneg_796 | 0 | 0 | 0.720343 | -3 | 2.23589 | -3 | 0.73395 | 0.1705 | 0.17978 | 0.20186 | 54.20074 | 0.18 | 0 | Neutral |
| testneg_797 | 0 | 0 | 0.094735 | -3 | 2.493678 | -3 | 0.024 | 0.07726 | 0.50057 | 0.10408 | 46.59562 | 0.05 | 0 | Neutral |
| testneg_798 | 0.21 | 0.15 | 1.266844 | -3 | 2.044018 | -3 | 0.7257 | 0.64394 | 0.63431 | 0.09837 | 12.34649 | 0.11 | 0 | Neutral |
| testneg_799 | 0.01 | 0.8 | 1.727114 | -3 | 1.648624 | 0.910063 | 0.19898 | 0.12788 | 0.54248 | 0.12333 | 18.6768 | 0.02 | 0 | Neutral |
| testneg_800 | 0.03 | 0.1 | -0.895222 | -3 | 2.753234 | -3 | 0.84468 | 0.55945 | 0.24519 | 0.12788 | 14.16667 | 0.11615942 | 0 | Neutral |
| testneg_801 | 0.05 | 0.29 | 1.902852 | -3 | 2.72063 | -3 | 0.56966 | 0.68625 | 0.65141 | 0.18046 | 7.01832 | 0.273422502 | 1.8 | Neutral |
| testneg_802 | 0 | 0.32 | 0.471907 | -3 | 11.056573 | -3 | 0.53424 | 0.08484 | 0.10914 | 0.76607 | 10.41667 | 0.457810724 | 34 | Neutral |
| testneg_803 | -0.07 | 0.31 | 6.012591 | -3 | 0.722185 | -3 | 0.16663 | 0.12788 | 0.18866 | 0.12788 | 28.0507 | 0.17725 | 0 | Neutral |
| testneg_804 | 0 | 0 | 0.507327 | -3 | 1.759394 | -3 | 0.11172 | 0.10034 | 0.03297 | 0.16627 | 15.05208 | 0.05620979 | 0 | Neutral |
| testneg_805 | 0 | 0 | 0.596959 | -3 | 1.534507 | -3 | 0.18925 | 0.32479 | 0.98118 | 0.10229 | 6.54109 | 0.03 | 0 | Neutral |
| testneg_806 | 0.09 | -0.3 | 2.237174 | -3 | -3.138593 | -3 | 0.14349 | 0.27976 | 0.06254 | 0.23561 | 10.13746 | 0.179861035 | 0 | Neutral |
| testneg_807 | 0 | 0 | 1.063047 | -3 | 2.891817 | -3 | 0.17855 | 0.11262 | 0.12436 | 0.11074 | 17.33333 | 0.01 | 0 | Neutral |
| testneg_808 | 0 | -0.01 | 2.23556 | -3 | -0.800423 | -3 | 0.98578 | 0.91075 | 0.17639 | 0.12919 | 4.72222 | 0.07 | 0 | Neutral |
| testneg_809 | 0.14 | -1.01 | 5.127855 | -3 | -2.664317 | -3 | 0.28449 | 0.10981 | 0.16321 | 0.09621 | 18.88889 | 0.07 | 0 | Neutral |
| testneg_810 | 0.04 | -0.23 | 7.535006 | -3 | 0.520101 | -3 | 0.0893 | 0.11207 | 0.4118 | 0.1154 | 19.09347 | 0.24011236 | 1.2 | Neutral |
| testneg_811 | -0.14 | 1.01 | -1.162831 | -1.162831 | 0.894917 | -3 | 0.13398 | 0.12788 | 0.10723 | 0.33098 | 38.74699 | 0.15 | 0 | Neutral |
| testneg_812 | -0.04 | 0.23 | -0.271338 | -3 | 5.751908 | -3 | 0.16797 | 0.12788 | 0.27115 | 0.14766 | 12.46106 | 0.320586265 | 6.6 | Neutral |
| testneg_813 | 0 | 0 | 0.195879 | -3 | 6.733255 | -3 | 0.05887 | 0.10046 | 0.37424 | 0.12788 | 22.8627 | 0.17 | 0 | Neutral |
| testneg_814 | 0.17 | 0.38 | 2.123391 | 2.123391 | 3.398658 | -3 | 0.04574 | 0.05824 | 0.06971 | 0.09918 | 25.55475 | 0.18025 | 0 | Neutral |
| testneg_815 | 0.18 | -1.05 | 6.446705 | -3 | 4.93471 | -3 | 0.18013 | 0.25346 | 0.09442 | 0.1153 | 18.75947 | 0.58687499 | 81 | Disease_causing |
| testneg_816 | 0.15 | -0.28 | 0.230515 | -3 | 1.23113 | -3 | 0.19898 | 0.1092 | 0.16973 | 0.17555 | 12.46106 | 0.06 | 0 | Neutral |
| testneg_817 | 0 | -0.32 | 5.730226 | -3 | 0.727897 | -3 | 0.13704 | 0.10054 | 0.34209 | 0.09515 | 17.77778 | 0.11 | 0 | Neutral |
| testneg_818 | 0 | 0 | 1.264197 | -3 | 1.500634 | -3 | 0.37542 | 0.19444 | 0.15736 | 0.5002 | 10.41667 | 0.068963565 | 0 | Neutral |
| testneg_819 | 0 | -0.32 | 4.048611 | 0.199688 | -0.085229 | -3 | 0.07237 | 0.12788 | 0.44644 | 0.10813 | 27.82051 | 0.08 | 0 | Neutral |
| testneg_820 | 0.05 | -0.81 | 5.789217 | -3 | 0.041172 | -3 | 0.26668 | 0.11598 | 0.01369 | 0.12788 | 21.9824 | 0.110192308 | 0 | Neutral |
| testneg_821 | 0.14 | -0.37 | 2.120517 | -3 | 4.142101 | -3 | 0.14457 | 0.11614 | 0.12885 | 0.61089 | 11.25 | 0.378183183 | 13.2 | Neutral |
| testneg_822 | 0 | 0 | 1.826458 | 1.826458 | 0.927961 | -3 | 0.46401 | 0.24608 | 0.7351 | 0.16019 | 4.44444 | 0.149390595 | 0 | Neutral |
| testneg_823 | 0.03 | -1.03 | 3.246994 | -3 | 0.472476 | -3 | 0.12633 | 0.19996 | 0.37424 | 0.12788 | 16.8018 | 0.081464286 | 0 | Neutral |
| testneg_824 | 0 | 0 | 0.153038 | -3 | 0.837529 | -3 | 0.92659 | 0.13588 | 0.26913 | 0.1087 | 23.89992 | 0.020502874 | 0 | Neutral |
| testneg_825 | -0.04 | -2.01 | 2.777008 | -3 | 0.706599 | -3 | 0.09093 | 0.28643 | 0.1077 | 0.12788 | 18.88889 | 0.11 | 0 | Neutral |
| testneg_826 | -0.03 | -0.1 | 0.907219 | -3 | 1.26259 | 1.26259 | 0.19898 | 0.12788 | 0.44722 | 0.11641 | 7.00579 | 0.010093491 | 0 | Neutral |
| testneg_827 | 0 | 0 | -0.260149 | -3 | -0.187363 | -3 | 0.16418 | 0.12788 | 0.90649 | 0.3543 | 5.65275 | 0.021588235 | 0 | Neutral |
| testneg_828 | -0.07 | -0.38 | 2.097779 | -3 | 0.700369 | -3 | 0.81054 | 0.28104 | 0.13195 | 0.57267 | 4.44444 | 0.12023823 | 0 | Neutral |
| testneg_829 | 0.02 | -0.36 | 3.235017 | 3.235017 | 1.491454 | -3 | 0.30329 | 0.14582 | 0.1078 | 0.17394 | 10.41667 | 0.689298518 | 97.6 | Disease_causing |
| testneg_830 | 0 | 0 | -0.216528 | -3 | 6.591382 | -3 | 0.40612 | 0.45135 | 0.46386 | 0.16229 | 18.33333 | 0.530083859 | 64 | Disease_causing |
| testneg_831 | 0.08 | 0.03 | 1.747877 | -3 | 1.377039 | -3 | 0.90811 | 0.17448 | 0.81838 | 0.09725 | 11.25 | 0.053875 | 0 | Neutral |
| testneg_832 | 0.07 | 0.7 | 3.047307 | -3 | 0.842221 | -3 | 0.12091 | 0.13585 | 0.07923 | 0.08035 | 14.40298 | 0.041111111 | 0 | Neutral |
| testneg_833 | 0 | 0 | 1.076963 | -3 | 3.58202 | -3 | 0.20011 | 0.27949 | 0.09633 | 0.12788 | 10.41667 | 0.077970033 | 0 | Neutral |
| testneg_834 | 0 | 0 | -0.272804 | -0.272804 | 0.924684 | -3 | 0.17839 | 0.12009 | 0.43283 | 0.23984 | 16.89285 | 0.038852073 | 0 | Neutral |
| testneg_835 | 0 | 0 | 1.829958 | -3 | 3.303858 | 3.303858 | 0.47775 | 0.10103 | 0.4558 | 0.9277 | 18.88889 | 0.31969697 | 4.6 | Neutral |
| testneg_836 | 0.07 | 0.33 | 3.81823 | -3 | 0.268278 | -3 | 0.1755 | 0.10455 | 0.09669 | 0.12526 | 19.18549 | 0.03 | 0 | Neutral |
| testneg_837 | 0 | 0 | 3.298555 | 1.465212 | -0.954908 | -3 | 0.24791 | 0.09675 | 0.10285 | 0.10886 | 21.38889 | 0.09 | 0 | Neutral |
| testneg_838 | 0 | -0.01 | 1.609618 | -3 | 1.091905 | -3 | 0.67038 | 0.1794 | 0.10058 | 0.13882 | 18.55981 | 0.081240836 | 0 | Neutral |
| testneg_839 | -0.07 | -0.38 | 1.821777 | -3 | 0.618993 | 0.618993 | 0.11187 | 0.12788 | 0.04207 | 0.12788 | 22.61299 | 0.02 | 0 | Neutral |
| testneg_840 | 0.01 | 0.8 | 3.453134 | -3 | 1.213652 | -3 | 0.16497 | 0.74918 | 0.11458 | 0.16496 | 11.66667 | 0.363782062 | 9.8 | Neutral |
| testneg_841 | -0.07 | -0.38 | 5.176715 | -3 | 0.256955 | -3 | 0.206 | 0.14631 | 0.1846 | 0.08682 | 16.8018 | 0.11 | 0 | Neutral |
| testneg_842 | 0 | 0 | 1.144493 | -3 | 4.738569 | -3 | 0.13794 | 0.40936 | 0.04161 | 0.12788 | 19.09347 | 0.166253205 | 0 | Neutral |
| testneg_843 | -0.06 | -0.18 | 1.465212 | -3 | 0.889233 | -3 | 0.24791 | 0.09675 | 0.50392 | 0.12788 | 12.29167 | 0.000299254 | 0 | Neutral |
| testneg_844 | 0.19 | -0.27 | 3.485229 | -3 | -0.296252 | -3 | 0.22872 | 0.15588 | 0.09323 | 0.16732 | 23.76984 | 0.267586158 | 0.6 | Neutral |
| testneg_845 | 0 | -0.01 | 1.062204 | -3 | -0.323717 | -3 | 0.73731 | 0.26107 | 0.23691 | 0.1015 | 20.07035 | 0.01 | 0 | Neutral |
| testneg_846 | 0 | 0 | 0.346053 | -3 | -1.444998 | -3 | 0.3047 | 0.11929 | 0.10561 | 0.12788 | 18.33333 | 0.00050641 | 0 | Neutral |
| testneg_847 | -0.02 | 0.68 | 5.845448 | -3 | 0.265756 | -3 | 0.06926 | 0.08399 | 0.96396 | 0.7628 | 18.89436 | 0.33 | 4.2 | Neutral |
| testneg_848 | 0.16 | 0.08 | -2.804234 | -3 | 1.435364 | -3 | 0.57349 | 0.09764 | 0.20357 | 0.12788 | 17.30496 | 0.04 | 0 | Neutral |
| testneg_849 | 0 | 0 | 1.411308 | -3 | 0.623338 | 0.623338 | 0.73253 | 0.09747 | 0.09707 | 0.12788 | 25.27596 | 0.021562993 | 0 | Neutral |
| testneg_850 | -0.0036364 | 0.11719697 | 0.781229 | 0.781229 | 1.058268 | -3 | 0.07321 | 0.12788 | 0.77981 | 0.62662 | 25.30416 | 0.14 | 0 | Neutral |
| testneg_851 | 0.01 | -0.57 | 4.598146 | -3 | 4.311522 | -3 | 0.08531 | 0.12788 | 0.08857 | 0.83499 | 11.13095 | 0.561941585 | 74 | Disease_causing |
| testneg_852 | 0 | 0 | 2.410424 | -3 | 0.092429 | -3 | 0.92514 | 0.19444 | 0.12259 | 0.1041 | 11.93907 | 0.01546875 | 0 | Neutral |
| testneg_853 | 0.01 | 0.8 | 0.586242 | -3 | 6.861369 | -3 | 0.559 | 0.2286 | 0.66863 | 0.10981 | 30.33972 | 0.33 | 3.8 | Neutral |
| testneg_854 | 0 | 0 | 2.434662 | 0.418118 | 0.273635 | -3 | 0.19898 | 0.12788 | 0.18578 | 0.27854 | 10 | 0.09094983 | 0 | Neutral |
| testneg_855 | -0.06 | 0.57 | 1.042833 | -3 | 2.317175 | -3 | 0.19898 | 0.12788 | 0.1909 | 0.14279 | 16.07106 | 0.086666667 | 0 | Neutral |
| testneg_856 | 0.07 | 0.7 | 4.067705 | -3 | 0.576122 | -3 | 0.14525 | 0.14744 | 0.18341 | 0.13901 | 12.37434 | 0.150161328 | 0 | Neutral |
| testneg_857 | -0.06 | -0.18 | 0.975671 | -3 | 0.243489 | -3 | 0.29739 | 0.10893 | 0.05476 | 0.12788 | 22.708 | 0.01 | 0 | Neutral |
| testneg_858 | 0 | 0 | 0.342408 | -3 | 0.631528 | -3 | 0.5255 | 0.15968 | 0.11688 | 0.13208 | 26.62482 | 0.109448761 | 0 | Neutral |
| testneg_859 | -0.14 | 0.37 | 3.496726 | -3 | -0.284893 | -3 | 0.25904 | 0.16134 | 0.24038 | 0.15296 | 20.83333 | 0.219375902 | 0 | Neutral |
| testneg_860 | 0.15 | 0.04 | -0.330052 | -3 | 0.957084 | -3 | 0.40398 | 0.12788 | 0.12987 | 0.18163 | 10.41667 | 0.07228181 | 0 | Neutral |
| testneg_861 | 0.09 | -0.3 | 3.200815 | -3 | 3.279189 | -3 | 0.98312 | 0.12788 | 0.2183 | 0.10789 | 15.01941 | 0.12 | 0 | Neutral |
| testneg_862 | 0.04 | -0.23 | 1.114692 | -3 | 1.342541 | -3 | 0.05124 | 0.07099 | 0.24524 | 0.10803 | 15.41667 | 0.03 | 0 | Neutral |
| testneg_863 | 0.01 | 0.8 | 1.582141 | -3 | -0.483923 | -3 | 0.34117 | 0.12167 | 0.23574 | 0.12788 | 16.8018 | 0.01 | 0 | Neutral |

| | | | | | | | | | | | | | | |
|---|---|---|---|---|---|---|---|---|---|---|---|---|---|---|
| testneg_864 | 0.18 | -1.05 | 2.222067 | -3 | 1.84534 | 1.84534 | 0.09462 | 0.11807 | 0.40996 | 0.57507 | 25.74388 | 0.222797619 | 0 | Neutral |
| testneg_865 | -0.14 | -0.54 | -0.251883 | -3 | 0.139533 | -3 | 0.39258 | 0.12788 | 0.2521 | 0.1784 | 27.17755 | 0.085069444 | 0 | Neutral |
| testneg_866 | 0 | 0 | 0.027672 | -3 | -0.330275 | -0.330275 | 0.16091 | 0.90971 | 0.10462 | 0.1278 | 10.74877 | 0.12 | 0 | Neutral |
| testneg_867 | 0 | 0 | 0.044696 | -3 | 1.003907 | -3 | 0.16125 | 0.08406 | 0.60079 | 0.11198 | 28.64708 | 0.01 | 0 | Neutral |
| testneg_868 | -0.08 | -0.03 | 3.966289 | -3 | 3.39754 | -3 | 0.61718 | 0.1648 | 0.62203 | 0.15583 | 4.72222 | 0.503888674 | 55.8 | Disease_causing |
| testneg_869 | 0 | 0 | -0.002747 | -3 | -0.196592 | -0.196592 | 0.07394 | 0.09408 | 0.32733 | 0.16869 | 15.41667 | 0.046217391 | 0 | Neutral |
| testneg_870 | 0.05 | -0.81 | 5.703258 | -3 | 0.787669 | -3 | 0.60812 | 0.19286 | 0.09332 | 0.53331 | 11.95833 | 0.473712607 | 46 | Neutral |
| testneg_871 | 0.21 | 0.15 | 0.02187 | -3 | 2.66851 | -3 | 0.34279 | 0.09378 | 0.84238 | 0.12559 | 10.41667 | 0.08 | 0 | Neutral |
| testneg_872 | -0.03 | -0.1 | -0.369846 | -3 | 2.25379 | -3 | 0.33467 | 0.12788 | 0.059 | 0.12788 | 27.51545 | 0.020566188 | 0 | Neutral |
| testneg_873 | -0.15 | -0.04 | 2.692574 | -3 | 3.710256 | -3 | 0.45628 | 0.15819 | 0.89661 | 0.17382 | 11.25 | 0.473440735 | 44 | Neutral |
| testneg_874 | 0.03072727 | -0.4116364 | 2.905355 | -3 | 4.202319 | -3 | 0.04305 | 0.06354 | 0.19898 | 0.30319 | 13.51351 | 0.444483491 | 34.6 | Neutral |
| testneg_875 | -0.08 | -0.03 | 3.758994 | -3 | 2.365185 | -3 | 0.22169 | 0.11262 | 0.14731 | 0.12788 | 31.08891 | 0.06 | 0 | Neutral |
| testneg_876 | 0.05 | -0.81 | 3.770242 | -0.305877 | 0.60694 | -3 | 0.22179 | 0.09831 | 0.07387 | 0.12788 | 25.82919 | 0.04 | 0 | Neutral |
| testneg_877 | 0 | 0 | 0.258619 | -3 | 0.857797 | -3 | 0.30823 | 0.17188 | 0.83798 | 0.14433 | 14.56463 | 0.037584228 | 0 | Neutral |
| testneg_878 | 0.04 | -0.23 | 0.007971 | -3 | 7.813348 | -3 | 0.12591 | 0.09845 | 0.08836 | 0.0911 | 13.51351 | 0.31 | 2.8 | Neutral |
| testneg_879 | -0.2 | 1.26 | 5.385549 | -3 | 1.418046 | -3 | 0.13126 | 0.11628 | 0.42872 | 0.12788 | 21.417 | 0.226857143 | 0.4 | Neutral |
| testneg_880 | -0.03 | -0.12 | 1.183797 | -3 | -0.410022 | -3 | 0.99765 | 0.95064 | 0.17419 | 0.33842 | 4.72222 | 0.182582721 | 0 | Neutral |
| testneg_881 | 0.17 | 0.38 | 2.901134 | -3 | 1.807438 | -3 | 0.82014 | 0.14054 | 0.80034 | 0.10518 | 17.21847 | 0.08 | 0 | Neutral |
| testneg_882 | 0 | 0 | -0.674729 | -3 | 1.666396 | -3 | 0.24765 | 0.32858 | 0.19898 | 0.10512 | 18.88889 | 0.03 | 0 | Neutral |
| testneg_883 | -0.1 | -1.9 | 0.700131 | -3 | 0.459406 | -3 | 0.46056 | 0.12785 | 0.03981 | 0.09432 | 10.22898 | 0.04146875 | 0 | Neutral |
| testneg_884 | -0.03 | -0.1 | 2.574694 | -3 | 1.614653 | -3 | 0.59871 | 0.34719 | 0.54895 | 0.15082 | 4.72222 | 0.159492424 | 0 | Neutral |
| testneg_885 | -0.18 | -0.16 | 0.369791 | -3 | -3.138593 | -3 | 0.66227 | 0.11002 | 0.06254 | 0.23561 | 18.82957 | 0.128055556 | 0 | Neutral |
| testneg_886 | 0 | 0.32 | 0.66894 | -3 | 3.303858 | -3 | 0.81212 | 0.14915 | 0.4558 | 0.9277 | 11.25 | 0.246813254 | 0.4 | Neutral |
| testneg_887 | -0.06 | -0.18 | 0.491194 | 0.491194 | -1.968465 | -3 | 0.72124 | 0.09906 | 0.65048 | 0.12778 | 13.51351 | 0.020188134 | 0 | Neutral |
| testneg_888 | 0 | 0 | 1.708897 | -3 | 0.310587 | 0.310587 | 0.5202 | 0.16306 | 0.2132 | 0.12788 | 11.26161 | 0.020325225 | 0 | Neutral |
| testneg_889 | 0 | -0.01 | 7.854373 | -3 | 0.434159 | -3 | 0.4113 | 0.2303 | 0.10168 | 0.12788 | 32.29167 | 0.437592593 | 27.6 | Neutral |
| testneg_890 | 0 | 0 | 1.788475 | -3 | 4.030653 | -3 | 0.37986 | 0.14229 | 0.48585 | 0.66989 | 10.75027 | 0.30923736 | 4 | Neutral |
| testneg_891 | 0 | 0 | 0.699729 | -3 | 1.902977 | 1.902977 | 0.76391 | 0.14229 | 0.18504 | 0.0822 | 13.27141 | 0.030485893 | 0 | Neutral |
| testneg_892 | 0.04 | -0.23 | -0.492173 | -3 | 2.494251 | -3 | 0.36225 | 0.1054 | 0.10314 | 0.08894 | 17.18434 | 0.020265845 | 0 | Neutral |
| testneg_893 | 0.06 | -0.57 | 3.795863 | -3 | -2.576045 | -2.576045 | 0.68055 | 0.12788 | 0.25275 | 0.12788 | 13.75 | 0.030893614 | 0 | Neutral |
| testneg_894 | 0 | 0 | 1.584574 | -3 | 7.596582 | -3 | 0.29362 | 0.22177 | 0.11391 | 0.12788 | 7.58745 | 0.414722222 | 26.4 | Neutral |
| testneg_895 | 0.03 | -1.03 | 5.957726 | -3 | 5.272058 | -3 | 0.44103 | 0.10698 | 0.16262 | 0.15786 | 27.51634 | 0.649313492 | 92.6 | Disease_causing |
| testneg_896 | 0.21 | 0.15 | 2.391994 | -3 | 3.087586 | -3 | 0.18785 | 0.14723 | 0.3828 | 0.5569 | 23.10245 | 0.395946292 | 13.4 | Neutral |
| testneg_897 | 0.18 | -1.05 | 7.035477 | -3 | 0.405386 | -3 | 0.13918 | 0.12788 | 0.08476 | 0.12788 | 14.90842 | 0.203125 | 1.4 | Neutral |
| testneg_898 | 0 | -0.01 | 2.776983 | -3 | 2.752736 | -3 | 0.24977 | 0.11262 | 0.15556 | 0.12841 | 21.8656 | 0.04 | 0 | Neutral |
| testneg_899 | 0 | 0 | 0.103447 | 0.103447 | 3.659658 | -3 | 0.15813 | 0.61197 | 0.21596 | 0.08969 | 19.12782 | 0.121192542 | 0 | Neutral |
| testneg_900 | 0.06 | -0.57 | 3.841536 | -3 | 4.877158 | -3 | 0.08002 | 0.08792 | 0.78211 | 0.35815 | 21.38889 | 0.479037378 | 45.8 | Neutral |
| testneg_901 | 0.14 | -1.01 | 4.346301 | -3 | 0.090223 | -3 | 0.44541 | 0.13979 | 0.34248 | 0.15472 | 19.42906 | 0.2025 | 0 | Neutral |
| testneg_902 | 0.14 | -0.69 | 3.53144 | 0.866429 | 4.494271 | -3 | 0.45405 | 0.10896 | 0.12453 | 0.17736 | 12.76316 | 0.631317793 | 93.4 | Disease_causing |
| testneg_903 | 0.21 | 0.15 | 6.222747 | -3 | 4.727625 | -3 | 0.1602 | 0.11262 | 0.13575 | 0.12788 | 29.52861 | 0.35 | 12.8 | Neutral |
| testneg_904 | -0.03 | 1.03 | 0.958989 | -3 | 0.774689 | 0.774689 | 0.39436 | 0.16446 | 0.24219 | 0.17458 | 18.6768 | 0.127630348 | 0 | Neutral |
| testneg_905 | 0.01 | -1.44 | 5.905772 | -3 | 4.136929 | -0.007379 | 0.55441 | 0.08704 | 0.19898 | 0.12788 | 30.5642 | 0.324444444 | 4.4 | Neutral |
| testneg_906 | 0.2 | -1.26 | 3.163156 | -3 | 5.705013 | -3 | 0.08954 | 0.63401 | 0.07211 | 0.12934 | 16.0119 | 0.460333333 | 38.8 | Neutral |
| testneg_907 | 0 | 0 | 1.914232 | -3 | 0.991883 | -3 | 0.77196 | 0.14089 | 0.26816 | 0.13347 | 13.01854 | 0.06 | 0 | Neutral |
| testneg_908 | 0.07 | 0.01 | 3.996475 | -3 | -2.106325 | -3 | 0.34698 | 0.12788 | 0.59812 | 0.19387 | 10.83333 | 0.149783163 | 0 | Neutral |
| testneg_909 | 0 | 0 | 0.821308 | -3 | -1.180653 | -3 | 0.64608 | 0.35524 | 0.04192 | 0.06306 | 17.44725 | 0.02 | 0 | Neutral |
| testneg_910 | -0.0088889 | 0.30555556 | 11.415361 | -3 | 0.371631 | -3 | 0.13275 | 0.12788 | 0.08337 | 0.18096 | 17.25464 | 0.50628046 | 60 | Disease_causing |
| testneg_911 | 0.02 | -0.63 | 2.847602 | -3 | -0.223089 | -3 | 0.15704 | 0.72174 | 0.33163 | 0.26601 | 10.21277 | 0.308819444 | 3 | Neutral |
| testneg_912 | 0 | -0.32 | 0.455675 | -3 | 6.2831 | -3 | 0.30951 | 0.12788 | 0.0626 | 0.1393 | 19.70284 | 0.363166667 | 11.4 | Neutral |
| testneg_913 | -0.04 | 0.23 | -0.52031 | -3 | 5.128101 | -3 | 0.12554 | 0.10854 | 0.41655 | 0.49485 | 10.83333 | 0.33141292 | 4.8 | Neutral |
| testneg_914 | 0.07 | 0.38 | -2.70564 | -3 | 1.385917 | -3 | 0.94229 | 0.26731 | 0.98641 | 0.83983 | 10.08333 | 0.170166667 | 0 | Neutral |
| testneg_915 | 0 | 0 | 0.194425 | -3 | -0.614161 | -0.614161 | 0.19898 | 0.10106 | 0.17796 | 0.09522 | 27.95796 | 0.01 | 0 | Neutral |
| testneg_916 | 0 | 0 | 0.094735 | -3 | 1.205837 | -3 | 0.024 | 0.07726 | 0.07071 | 0.11499 | 33.26542 | 0.04 | 0 | Neutral |
| testneg_917 | 0 | 0 | 1.587314 | -3 | 0.664858 | -3 | 0.20457 | 0.12378 | 0.19898 | 0.09822 | 19.09347 | 0 | 0 | Neutral |
| testneg_918 | 0 | 0 | 2.30425 | -3 | -0.46506 | -3 | 0.46792 | 0.11262 | 0.48615 | 0.14369 | 13.8296 | 0.020629891 | 0 | Neutral |
| testneg_919 | -0.07 | -0.38 | 1.176788 | -3 | -0.4837 | -3 | 0.4558 | 0.9277 | 0.23276 | 0.31401 | 12.29167 | 0.210345238 | 0.2 | Neutral |
| testneg_920 | -0.2 | 1.26 | 0.425452 | -3 | 1.700368 | -3 | 0.23859 | 0.1202 | 0.09211 | 0.08973 | 10 | 0.11 | 0 | Neutral |
| testneg_921 | 0 | 0 | -0.725688 | -3 | 6.223663 | -3 | 0.17285 | 0.09314 | 0.34698 | 0.12788 | 10 | 0.085627413 | 0 | Neutral |
| testneg_922 | 0.05 | -0.81 | 0.668367 | -3 | 7.673652 | -3 | 0.03656 | 0.07074 | 0.09059 | 0.12032 | 26.69591 | 0.28 | 1.2 | Neutral |
| testneg_923 | 0 | 0 | 0.889458 | 0.889458 | 1.926886 | -3 | 0.48937 | 0.22085 | 0.18431 | 0.13863 | 14.47072 | 0.134820904 | 0 | Neutral |
| testneg_924 | -0.07 | -0.33 | 0.985586 | -3 | 0.723398 | 0.723398 | 0.12077 | 0.11244 | 0.19898 | 0.14957 | 18.57759 | 0.06 | 0 | Neutral |
| testneg_925 | 0 | 0 | 0.365268 | -3 | -0.710033 | -3 | 0.53627 | 0.12788 | 0.12689 | 0.13376 | 18.60878 | 0.086579689 | 0 | Neutral |
| testneg_926 | 0.07 | -0.31 | -0.344853 | -3 | 1.113746 | -3 | 0.50276 | 0.09638 | 0.70616 | 0.09323 | 25.58201 | 0.022777778 | 0 | Neutral |
| testneg_927 | 0 | 0 | 0.930011 | 0.930011 | 2.125818 | -3 | 0.12786 | 0.72066 | 0.15713 | 0.3507 | 18.81242 | 0.287871462 | 1.2 | Neutral |
| testneg_928 | 0 | 0 | 1.519976 | -3 | 0.116245 | -3 | 0.19898 | 0.12788 | 0.8238 | 0.38836 | 10 | 0.02036039 | 0 | Neutral |
| testneg_929 | 0 | -0.01 | 0.465839 | -3 | 3.222237 | -3 | 0.07875 | 0.11082 | 0.78515 | 0.35093 | 21.6561 | 0.150565476 | 0 | Neutral |
| testneg_930 | -0.0236364 | 0.26490909 | 11.240696 | -3 | 1.223596 | -3 | 0.10241 | 0.08165 | 0.24519 | 0.12788 | 18.33333 | 0.37 | 10.8 | Neutral |
| testneg_931 | 0.14 | -0.37 | 0.198379 | -3 | -2.124954 | -3 | 0.03991 | 0.08342 | 0.34153 | 0.08855 | 10.83333 | 0.04 | 0 | Neutral |
| testneg_932 | 0 | -0.01 | 0.558245 | -3 | 1.631249 | -3 | 0.19898 | 0.10362 | 0.47166 | 0.13548 | 22.91417 | 0.047881579 | 0 | Neutral |
| testneg_933 | 0.02563636 | 0.22745455 | 12.72993 | -3 | 4.893835 | 4.893835 | 0.39019 | 0.18821 | 0.0778 | 0.0728 | 11.92982 | 0.71 | 99.4 | Disease_causing |
| testneg_934 | 0 | 0 | 0.648536 | -3 | 5.91019 | -3 | 0.31124 | 0.08875 | 0.16462 | 0.18467 | 10.83333 | 0.275720033 | 1.4 | Neutral |
| testneg_935 | 0.06 | -0.57 | 1.240379 | -3 | 3.287232 | -3 | 0.64591 | 0.38907 | 0.20448 | 0.12788 | 11.875 | 0.131316667 | 0 | Neutral |
| testneg_936 | 0 | -0.32 | 1.947087 | -3 | 1.26259 | 1.26259 | 0.31004 | 0.09727 | 0.44722 | 0.11641 | 12.38369 | 0.010111735 | 0 | Neutral |
| testneg_937 | 0 | 0 | -0.108752 | -3 | 1.223596 | -3 | 0.48075 | 0.20313 | 0.24519 | 0.12788 | 4.44444 | 0.01 | 0 | Neutral |
| testneg_938 | 0 | 0 | 4.825663 | 1.802862 | 0.271343 | -3 | 0.16393 | 0.08897 | 0.10168 | 0.12788 | 29.45309 | 0.15 | 0 | Neutral |
| testneg_939 | 0.03 | 0.12 | 0.932751 | -3 | 0.453787 | -3 | 0.15612 | 0.11465 | 0.17843 | 0.15468 | 21.83511 | 0.053088901 | 0 | Neutral |
| testneg_940 | 0.14 | -1.01 | 0.139802 | -3 | -0.20288 | -3 | 0.40234 | 0.10353 | 0.22901 | 0.12788 | 12.37434 | 0.01 | 0 | Neutral |
| testneg_941 | 0 | -0.32 | 3.667889 | -3 | 1.069555 | -3 | 0.84572 | 0.78546 | 0.19898 | 0.12788 | 5 | 0.110909091 | 0 | Neutral |
| testneg_942 | -0.18 | -0.16 | 2.173009 | -3 | 1.730475 | -3 | 0.13119 | 0.12788 | 0.73601 | 0.26791 | 29.45989 | 0.14 | 0 | Neutral |
| testneg_943 | 0 | 0 | 0.628811 | -3 | 1.861556 | -3 | 0.07874 | 0.12788 | 0.19423 | 0.14002 | 26.92123 | 0.05 | 0 | Neutral |
| testneg_944 | 0 | 0 | 2.124149 | -3 | 3.793394 | -3 | 0.14482 | 0.08592 | 0.23955 | 0.97212 | 12.01128 | 0.257938455 | 0.6 | Neutral |
| testneg_945 | 0 | 0 | 0.540495 | -3 | -3.629472 | -3 | 0.78286 | 0.19831 | 0.15612 | 0.11465 | 18.76159 | 0.019299133 | 0 | Neutral |
| testneg_946 | 0.02 | -0.63 | -2.337287 | -3 | 5.862785 | -3 | 0.04946 | 0.12788 | 0.04829 | 0.09848 | 35.83027 | 0.159269624 | 0 | Neutral |
| testneg_947 | 0 | -0.01 | 1.844079 | -3 | -0.463513 | -3 | 0.54777 | 0.25991 | 0.94531 | 0.91608 | 5 | 0.081821475 | 0 | Neutral |
| testneg_948 | 0 | 0 | -0.041573 | -3 | 3.567601 | -3 | 0.06095 | 0.08179 | 0.04655 | 0.10225 | 12.29167 | 0.061782967 | 0 | Neutral |
| testneg_949 | 0.07 | 0.38 | 2.246546 | -3 | 1.231291 | -3 | 0.32022 | 0.09529 | 0.20577 | 0.12653 | 10 | 0.01 | 0 | Neutral |
| testneg_950 | -0.06 | -0.18 | 0.041839 | -3 | -0.870639 | -3 | 0.54211 | 0.12592 | 0.05494 | 0.20364 | 10.01954 | 0.075636364 | 0 | Neutral |
| testneg_951 | 0.17 | 0.38 | 3.17007 | -3 | 0.645032 | -3 | 0.12377 | 0.12788 | 0.16547 | 0.16092 | 18.66667 | 0.176891026 | 0 | Neutral |
| testneg_952 | 0.01 | -1.44 | 4.610927 | -3 | 1.531669 | -3 | 0.67796 | 0.10874 | 0.59321 | 0.59981 | 12.70833 | 0.260926871 | 0.4 | Neutral |
| testneg_953 | 0.14 | -0.37 | 6.711275 | -3 | -0.680715 | -3 | 0.1647 | 0.12076 | 0.40264 | 0.12788 | 12.63889 | 0.170416667 | 0.8 | Neutral |
| testneg_954 | 0 | 0 | 1.107432 | -3 | -0.274976 | -3 | 0.5726 | 0.1165 | 0.26767 | 0.25506 | 12.37434 | 0.020833333 | 0 | Neutral |
| testneg_955 | 0 | 0 | 0.256955 | -3 | 7.426793 | -3 | 0.1846 | 0.08682 | 0.69254 | 0.10021 | 33.77923 | 0.2 | 0.4 | Neutral |
| testneg_956 | 0 | 0 | -0.155612 | -3 | 0.406982 | -3 | 0.07141 | 0.12788 | 0.13255 | 0.0962 | 19.86444 | 0.000741758 | 0 | Neutral |
| testneg_957 | -0.0403636 | 0.23963636 | 5.990837 | -3 | 5.907423 | -3 | 0.18633 | 0.14783 | 0.00451 | 0.2022 | 20.26637 | 0.8 | 100 | Disease_causing |
| testneg_958 | -0.14 | 1.01 | 4.464407 | -3 | -2.650454 | -3 | 0.05904 | 0.09282 | 0.01438 | 0.03448 | 33.53535 | 0.14 | 0 | Neutral |
| testneg_959 | 0.14 | -1.01 | 0.727619 | -3 | 1.343895 | -3 | 0.57975 | 0.1248 | 0.35137 | 0.09161 | 24.8605 | 0.03 | 0 | Neutral |
| testneg_960 | 0 | 0 | 0.949965 | -3 | 1.402346 | -3 | 0.11796 | 0.12788 | 0.52299 | 0.12788 | 21.94444 | 0.01 | 0 | Neutral |
| testneg_961 | 0 | 0 | -0.280416 | -3 | 4.602995 | -3 | 0.21728 | 0.1132 | 0.04484 | 0.95426 | 18.33333 | 0.282605723 | 0.8 | Neutral |
| testneg_962 | 0.16 | -1.37 | 0.561 | -3 | 0.807044 | -3 | 0.09184 | 0.38254 | 0.0778 | 0.0728 | 10.41667 | 0.08 | 0 | Neutral |
| testneg_963 | 0.02 | -0.36 | 7.397319 | -3 | 0.294506 | -3 | 0.19355 | 0.18675 | 0.0961 | 0.09786 | 10.41667 | 0.398444444 | 17.6 | Neutral |
| testneg_964 | 0 | 0 | 0.926316 | -3 | 0.498535 | -3 | 0.34495 | 0.16861 | 0.63798 | 0.19967 | 5.74956 | 0.026475541 | 0 | Neutral |
| testneg_965 | 0 | 0 | 7.325251 | -3 | -1.367472 | -3 | 0.16601 | 0.14015 | 0.049 | 0.11324 | 58.22673 | 0.300591954 | 5 | Neutral |
| testneg_966 | -0.06 | 0.57 | 3.421338 | -3 | 0.517404 | -3 | 0.32201 | 0.10713 | 0.16418 | 0.12788 | 10 | 0.040271392 | 0 | Neutral |
| testneg_967 | 0.08 | 0.03 | -0.040342 | -3 | 0.157379 | -3 | 0.1834 | 0.12788 | 0.45997 | 0.15162 | 37.98701 | 0.070686869 | 0 | Neutral |
| testneg_968 | 0.03 | -1.03 | 3.202376 | -3 | 0.054931 | -3 | 0.25854 | 0.18247 | 0.35499 | 0.12052 | 58.92857 | 0.11 | 0 | Neutral |
| testneg_969 | 0.05 | 0.29 | 3.783684 | -3 | 3.584096 | -3 | 0.39399 | 0.12788 | 0.18127 | 0.12788 | 30.00722 | 0.137221491 | 0 | Neutral |
| testneg_970 | -0.05 | -0.29 | 4.656948 | -3 | 4.106182 | -3 | 0.91587 | 0.10818 | 0.06394 | 0.24464 | 12.89073 | 0.532132924 | 66.6 | Disease_causing |
| testneg_971 | 0 | 0 | 0.617174 | -3 | 6.765808 | 4.740762 | 0.28416 | 0.27805 | 0.13532 | 0.19904 | 13.32524 | 0.679791667 | 96.2 | Disease_causing |

| | | | | | | | | | | | | | |
|---|---|---|---|---|---|---|---|---|---|---|---|---|---|
| testneg_972 | 0 | 0 | 0.190697 | -3 | 4.157365 | -3 | 0.38503 | 0.11286 | 0.29672 | 0.10485 | 10.26553 | 0.020651867 | 0 Neutral |
| testneg_973 | 0.21 | 0.15 | 2.233647 | -3 | 2.535807 | -3 | 0.27175 | 0.14094 | 0.23844 | 0.27772 | 8.11707 | 0.256127996 | 0.2 Neutral |
| testneg_974 | 0.07 | 0.01 | 2.863307 | -3 | 2.341037 | 2.341037 | 0.32022 | 0.40347 | 0.95535 | 0.12788 | 11.25 | 0.206666667 | 0 Neutral |
| testneg_975 | -0.02 | -0.45 | 1.511803 | -3 | 0.615335 | -3 | 0.09306 | 0.09084 | 0.70872 | 0.09884 | 18.9446 | 0.02 | 0 Neutral |
| testneg_976 | 0.14 | -1.01 | 7.178388 | -3 | 5.05614 | -3 | 0.11905 | 0.23747 | 0.09442 | 0.1153 | 16.8018 | 0.65775 | 95.2 Disease_causing |
| testneg_977 | 0 | 0 | -0.064795 | -3 | 3.39754 | -3 | 0.4566 | 0.14201 | 0.62203 | 0.15583 | 8.45975 | 0.167939466 | 0 Neutral |
| testneg_978 | 0 | 0 | 0.470493 | -3 | -0.907393 | -3 | 0.42802 | 0.12096 | 0.05332 | 0.12788 | 20.20599 | 0.01 | 0 Neutral |
| testneg_979 | 0.21 | 0.15 | 4.528664 | -3 | 0.953557 | -3 | 0.49106 | 0.89143 | 0.09522 | 0.20323 | 9.35754 | 0.36 | 9 Neutral |
| testneg_980 | 0.21 | 0.15 | 2.317295 | -3 | 1.681428 | -3 | 0.99178 | 0.93124 | 0.59691 | 0.11049 | 15.95477 | 0.155 | 0 Neutral |
| testneg_981 | 0.22 | -2.66 | 4.505271 | -3 | 3.683685 | -0.787396 | 0.10472 | 0.10014 | 0.09633 | 0.12788 | 12.29167 | 0.210502137 | 0 Neutral |
| testneg_982 | -0.07 | -0.38 | 1.821777 | -3 | 3.289605 | -3 | 0.11187 | 0.12788 | 0.25758 | 0.2262 | 10.41667 | 0.214238095 | 0.2 Neutral |
| testneg_983 | -0.21 | -0.15 | -0.83994 | -3 | 1.132066 | -3 | 0.95319 | 0.20297 | 0.99317 | 0.60884 | 9.79167 | 0.17466092 | 0 Neutral |
| testneg_984 | -0.09 | 0.3 | 2.137709 | -3 | 0.712486 | -3 | 0.11956 | 0.3237 | 0.10161 | 0.04472 | 20.90186 | 0.076485507 | 0 Neutral |
| testneg_985 | 0.01485714 | -0.0434286 | 1.535215 | -3 | 3.725973 | -3 | 0.26323 | 0.2399 | 0.27058 | 0.13363 | 11.16442 | 0.382748192 | 16.4 Neutral |
| testneg_986 | -0.01 | 1.44 | 2.603186 | -3 | 2.859591 | -3 | 0.96341 | 0.18404 | 0.1576 | 0.12788 | 12.40248 | 0.2 | 0 Neutral |
| testneg_987 | 0 | 0 | 4.336736 | -3 | 3.806625 | -3 | 0.27275 | 0.11752 | 0.29372 | 0.07609 | 26.34921 | 0.110261006 | 0 Neutral |
| testneg_988 | 0 | 0 | -0.680715 | -3 | -2.067049 | -3 | 0.40264 | 0.12788 | 0.19199 | 0.16616 | 17.77778 | 0.048698741 | 0 Neutral |
| testneg_989 | 0.22 | -2.66 | 3.94836 | -3 | 3.385856 | -3 | 0.19898 | 0.12788 | 0.18013 | 0.25346 | 18.09499 | 0.5515865 | 71.8 Disease_causing |
| testneg_990 | 0 | 0 | 2.064931 | -3 | 4.416059 | -3 | 0.30122 | 0.09165 | 0.53216 | 0.84775 | 13.83267 | 0.262915836 | 1 Neutral |
| testneg_991 | -0.01 | -0.79 | 5.611956 | -3 | 5.072901 | -3 | 0.14187 | 0.1353 | 0.16436 | 0.19109 | 26.34995 | 0.749395137 | 100 Disease_causing |
| testneg_992 | 0.16 | 0.08 | 1.986437 | -3 | -1.644193 | -1.644193 | 0.69666 | 0.11055 | 0.19898 | 0.06726 | 48.38929 | 0.03 | 0 Neutral |
| testneg_993 | 0.00454545 | -0.1357273 | 0.208107 | -3 | 6.223663 | -3 | 0.10863 | 0.098 | 0.34698 | 0.12788 | 18.61427 | 0.12 | 0 Neutral |
| testneg_994 | -0.09 | 2.17 | 4.614663 | -3 | 0.2182 | -3 | 0.20604 | 0.10821 | 0.19898 | 0.12788 | 41.30276 | 0.0945 | 0 Neutral |
| testneg_995 | 0 | -0.01 | 7.854373 | -3 | 5.823976 | -3 | 0.4113 | 0.2303 | 0.06562 | 0.12788 | 29.86111 | 0.69 | 98.6 Disease_causing |
| testneg_996 | 0 | -0.32 | 3.779532 | -3 | 0.568251 | -3 | 0.11373 | 0.10739 | 0.08932 | 0.32882 | 11.25 | 0.200302137 | 0 Neutral |
| testneg_997 | 0 | 0 | 1.190149 | -3 | 1.148696 | -3 | 0.16841 | 0.11143 | 0.50592 | 0.0952 | 21.49849 | 0.000837456 | 0 Neutral |
| testneg_998 | 0.17 | 0.38 | 2.311167 | 2.311167 | 0.461043 | -3 | 0.05413 | 0.05691 | 0.62864 | 0.11103 | 11.76005 | 0.150178571 | 0.2 Neutral |
| testneg_999 | -0.15 | 0.82 | 4.208794 | -3 | 6.211178 | -3 | 0.07665 | 0.31408 | 0.14668 | 0.17962 | 18.051 | 0.884924862 | 100 Disease_causing |
| testneg_1000 | -0.19 | -0.51 | 1.521336 | -3 | 1.457871 | -3 | 0.39553 | 0.11991 | 0.08837 | 0.09327 | 16.69872 | 0.04 | 0 Neutral |
