## Supplementary material for "Predicting disease-causing variant combinations": Dataset S4

| Combination_ID | Flex1 | Hydr1 | CADD1 | CADD2 | CADD3 | CADD4 | HI_A | RecA | HI_B | RecB | Biol_Dist | Classification_score | Support_score | Predicted_class |
|---|---|---|---|---|---|---|---|---|---|---|---|---|---|---|
| testneg_1 | -0.16 | -0.08 | 1.845825 | -3 | 1.871099 | -3 | 0.69249 | 0.12788 | 0.87611 | 0.16759 | 17.3118 | 0.116625 | 0 | Neutral |
| testneg_2 | 0 | 0 | 0.719031 | -3 | -0.223591 | -0.223591 | 0.9924 | 0.37896 | 0.48705 | 0.09993 | 9.76428 | 0.03 | 0 | Neutral |
| testneg_3 | 0 | 0 | -0.364413 | -3 | 1.557286 | -3 | 0.16965 | 0.25991 | 0.27705 | 0.22346 | 12.31894 | 0.079217686 | 0 | Neutral |
| testneg_4 | -0.06 | -0.18 | 0.556885 | -3 | 0.553148 | -3 | 0.78664 | 0.11725 | 0.07427 | 0.09451 | 16.92102 | 0.01 | 0 | Neutral |
| testneg_5 | 0.05 | -0.81 | 6.108615 | -3 | 1.919463 | -3 | 0.0984 | 0.1489 | 0.24736 | 0.12777 | 18.0784 | 0.28 | 2.6 | Neutral |
| testneg_6 | 0 | 0 | -0.034419 | -0.034419 | 3.398658 | -3 | 0.62203 | 0.12788 | 0.06971 | 0.09918 | 9.88018 | 0.070502346 | 0 | Neutral |
| testneg_7 | 0 | -0.01 | 0.866411 | -3 | 1.182833 | -3 | 0.13011 | 0.14157 | 0.89661 | 0.17382 | 17.21847 | 0.03245 | 0 | Neutral |
| testneg_8 | 0 | 0 | -1.230837 | -1.230837 | 8.821972 | -3 | 0.14821 | 0.11262 | 0.05546 | 0.28027 | 12.81915 | 0.426537213 | 26 | Neutral |
| testneg_9 | 0.02 | -0.63 | 6.009451 | -3 | 5.486545 | -3 | 0.32406 | 0.10985 | 0.04142 | 0.09564 | 20.27778 | 0.3925 | 19.4 | Neutral |
| testneg_10 | 0.02 | -0.63 | 3.746637 | -3 | 2.950441 | -3 | 0.39262 | 0.12788 | 0.05245 | 0.13458 | 24.1453 | 0.253095238 | 0.6 | Neutral |
| testneg_11 | -0.18 | -0.16 | 2.826632 | -3 | 1.932209 | -3 | 0.52388 | 0.25831 | 0.54662 | 0.22085 | 17.49759 | 0.308992174 | 2.6 | Neutral |
| testneg_12 | 0 | 0 | 0.890402 | -3 | 1.921721 | -3 | 0.21376 | 0.12708 | 0.16613 | 0.12378 | 19.09347 | 0.000111735 | 0 | Neutral |
| testneg_13 | 0 | 0 | -0.32829 | -3 | -0.468268 | -3 | 0.18509 | 0.12831 | 0.49777 | 0.14244 | 25.32134 | 0.04191944 | 0 | Neutral |
| testneg_14 | 0.02 | 0.45 | 5.680826 | 1.976922 | 1.763064 | -3 | 0.18013 | 0.25346 | 0.11873 | 0.10499 | 35.84358 | 0.36 | 11 | Neutral |
| testneg_15 | 0.03 | 1.22 | 5.09045 | -3 | 0.502731 | -3 | 0.10724 | 0.49624 | 0.75033 | 0.64204 | 5 | 0.406090401 | 21 | Neutral |
| testneg_16 | 0.08 | 0.03 | 2.365497 | -3 | 6.16876 | -3 | 0.61491 | 0.11954 | 0.35984 | 0.11129 | 9.47046 | 0.133888889 | 0 | Neutral |
| testneg_17 | 0.03 | 0.12 | 2.051799 | -3 | 1.396704 | -3 | 0.14624 | 0.10369 | 0.12161 | 0.24178 | 11.25 | 0.061569301 | 0 | Neutral |
| testneg_18 | 0.21 | 0.15 | 7.3961 | -3 | 5.639599 | -3 | 0.21289 | 0.08709 | 0.53157 | 0.50154 | 4.72222 | 0.708431996 | 96.2 | Disease_causing |
| testneg_19 | 0.15 | -0.28 | -0.216905 | -3 | 0.867502 | -3 | 0.493 | 0.35835 | 0.71511 | 0.34803 | 10 | 0.179321429 | 0 | Neutral |
| testneg_20 | 0.01 | 0.79 | 3.142297 | 3.142297 | 2.278399 | -3 | 0.26758 | 0.10473 | 0.26489 | 0.10549 | 20.49061 | 0.39 | 14.2 | Neutral |
| testneg_21 | 0.03 | 0.12 | 1.995674 | -3 | -0.668726 | -3 | 0.16345 | 0.11716 | 0.27924 | 0.12788 | 24.71767 | 0.000566396 | 0 | Neutral |
| testneg_22 | 0 | -0.32 | 1.514021 | -3 | 5.537244 | -3 | 0.02791 | 0.0854 | 0.19898 | 0.09972 | 18.88889 | 0.1 | 0 | Neutral |
| testneg_23 | 0 | 0 | 0.103447 | 0.103447 | 3.125318 | -3 | 0.15813 | 0.61197 | 0.78536 | 0.12709 | 16.04167 | 0.110871212 | 0 | Neutral |
| testneg_24 | 0.01 | 0.8 | -0.214756 | -3 | 5.126283 | -3 | 0.07428 | 0.07645 | 0.68423 | 0.08163 | 21.69549 | 0.113101852 | 0 | Neutral |
| testneg_25 | 0.18 | -1.05 | 5.931475 | -3 | 1.173845 | -3 | 0.19494 | 0.14451 | 0.88689 | 0.41514 | 10.83333 | 0.448582251 | 35.2 | Neutral |
| testneg_26 | 0.17 | 0.38 | 0.812539 | -3 | 0.576852 | -3 | 0.19898 | 0.12788 | 0.07571 | 0.09432 | 16.88659 | 0.02 | 0 | Neutral |
| testneg_27 | 0.04 | -0.23 | 3.457136 | -3 | 1.698695 | -3 | 0.40707 | 0.37988 | 0.59302 | 0.10897 | 21.70397 | 0.1255 | 0 | Neutral |
| testneg_28 | -0.09 | 0.3 | 3.208069 | -3 | -0.772668 | -3 | 0.24327 | 0.12192 | 0.17643 | 0.10767 | 18.20544 | 0.023541667 | 0 | Neutral |
| testneg_29 | -0.14 | -0.54 | 0.83037 | -3 | 5.507024 | -3 | 0.16929 | 0.12788 | 0.21587 | 0.11701 | 19.81927 | 0.16 | 0 | Neutral |
| testneg_30 | 0 | 0 | 0.335364 | -3 | 3.992072 | -3 | 0.87839 | 0.33588 | 0.05767 | 0.07325 | 17.88889 | 0.094275641 | 0 | Neutral |
| testneg_31 | 0.07 | -0.31 | 1.843162 | -3 | 1.120506 | -3 | 0.30869 | 0.11948 | 0.16547 | 0.16092 | 11.25 | 0.062874304 | 0 | Neutral |
| testneg_32 | 0 | 0 | -0.626667 | -3 | 0.320279 | -3 | 0.21933 | 0.18619 | 0.43158 | 0.10472 | 11.58784 | 0.010188746 | 0 | Neutral |
| testneg_33 | 0.02 | -0.28 | -3.482963 | -3 | 1.218246 | -3 | 0.19898 | 0.12788 | 0.20448 | 0.12788 | 43.23486 | 0.010257086 | 0 | Neutral |
| testneg_34 | 0.07 | 0.38 | -0.473879 | -3 | 4.945871 | -3 | 0.0695 | 0.12788 | 0.055 | 0.20858 | 16.42913 | 0.308552557 | 3.4 | Neutral |
| testneg_35 | -0.02 | 0.63 | -2.507892 | -3 | 1.681199 | -3 | 0.13565 | 0.46572 | 0.52932 | 0.65749 | 10 | 0.193925382 | 0 | Neutral |
| testneg_36 | -0.08 | -0.03 | 0.630495 | 0.630495 | -0.6438 | -3 | 0.14419 | 0.1668 | 0.43158 | 0.10472 | 18.6768 | 0.06 | 0 | Neutral |
| testneg_37 | 0 | 0 | 1.712042 | -3 | -1.847326 | -3 | 0.31004 | 0.09727 | 0.07595 | 0.11822 | 13.02486 | 0.000250655 | 0 | Neutral |
| testneg_38 | 0.05 | 0.29 | 0.660357 | -3 | 0.001281 | -3 | 0.61718 | 0.1648 | 0.07675 | 0.11464 | 16.42913 | 0.03 | 0 | Neutral |
| testneg_39 | -0.08 | -0.03 | 1.53322 | -3 | 2.135175 | -3 | 0.78537 | 0.33817 | 0.47192 | 0.11305 | 27.91667 | 0.060042373 | 0 | Neutral |
| testneg_40 | 0.00036364 | -0.0270909 | 2.070789 | -3 | 7.564785 | -3 | 0.91139 | 0.65138 | 0.13749 | 0.17824 | 19.67873 | 0.62 | 90.2 | Disease_causing |
| testneg_41 | 0.05 | -0.81 | 3.247775 | -3 | 2.579167 | -3 | 0.47888 | 0.43756 | 0.1294 | 0.10964 | 22.59553 | 0.200651179 | 0 | Neutral |
| testneg_42 | -0.08 | -0.03 | 3.600262 | -3 | 1.917158 | 1.917158 | 0.22944 | 0.09039 | 0.11618 | 0.20784 | 20.42005 | 0.273842857 | 0.2 | Neutral |
| testneg_43 | 0 | 0 | 0.741693 | -3 | -1.062235 | -3 | 0.48906 | 0.11262 | 0.29172 | 0.08143 | 31.59206 | 0.01 | 0 | Neutral |
| testneg_44 | 0.19 | -0.25 | 7.226253 | 0.342775 | -1.14689 | -3 | 0.83798 | 0.14433 | 0.14704 | 0.12788 | 11.74429 | 0.38 | 12.8 | Neutral |
| testneg_45 | 0.22 | -2.66 | 7.2081 | -3 | 2.702357 | -3 | 0.65734 | 0.1013 | 0.94674 | 0.12413 | 26.22294 | 0.4 | 14 | Neutral |
| testneg_46 | 0.03 | 0.12 | 2.051799 | -3 | -0.071864 | -0.071864 | 0.14624 | 0.10369 | 0.97005 | 0.31407 | 5.73716 | 0.07 | 0 | Neutral |
| testneg_47 | 0.02 | -0.68 | 0.188403 | -3 | -0.454393 | -0.454393 | 0.39111 | 0.15887 | 0.03443 | 0.1032 | 19.09347 | 0.035 | 0 | Neutral |
| testneg_48 | 0 | 0 | 0.386178 | -3 | -0.187157 | -3 | 0.04441 | 0.13603 | 0.09871 | 0.09286 | 17.21847 | 0.04 | 0 | Neutral |
| testneg_49 | 0.08 | 0.03 | -2.363283 | -2.363283 | 0.010179 | -3 | 0.05846 | 0.12788 | 0.19898 | 0.12788 | 19.3328 | 0.04 | 0 | Neutral |
| testneg_50 | 0 | 0 | 0.429192 | -3 | 4.7721 | -3 | 0.50224 | 0.4059 | 0.45014 | 0.12788 | 16.8018 | 0.129782828 | 0 | Neutral |
| testneg_51 | 0.02 | -0.63 | 1.351912 | -3 | -0.270292 | -3 | 0.18584 | 0.11114 | 0.22998 | 0.08738 | 30.36721 | 0.01 | 0 | Neutral |
| testneg_52 | -0.0036364 | 0.11719697 | 0.781229 | 0.781229 | -0.833975 | -3 | 0.07321 | 0.12788 | 0.20849 | 0.12788 | 54.20623 | 0.06 | 0 | Neutral |
| testneg_53 | -0.23 | 1.04 | 5.857535 | -3 | 0.285978 | -3 | 0.17532 | 0.36561 | 0.19898 | 0.09824 | 16.84579 | 0.33983871 | 6 | Neutral |
| testneg_54 | 0.06 | -0.57 | 2.146864 | -3 | 1.754249 | -3 | 0.19898 | 0.09316 | 0.07275 | 0.09562 | 20.02084 | 0.02 | 0 | Neutral |
| testneg_55 | 0.17 | 0.38 | 1.603588 | -3 | 0.062343 | -3 | 0.10496 | 0.21491 | 0.19898 | 0.11691 | 21.13681 | 0.060259613 | 0 | Neutral |
| testneg_56 | -0.05 | -0.29 | -1.375282 | -3 | 1.940661 | -3 | 0.19898 | 0.12788 | 0.25151 | 0.12788 | 13.51472 | 0.01 | 0 | Neutral |
| testneg_57 | -0.07 | -0.01 | -1.595702 | -3 | 2.422515 | -3 | 0.82131 | 0.12788 | 0.14379 | 0.12788 | 22.06349 | 0.040712094 | 0 | Neutral |
| testneg_58 | 0.02 | -0.68 | -0.505763 | -3 | 0.722971 | -3 | 0.11585 | 0.12788 | 0.19898 | 0.12788 | 32.42286 | 0.01 | 0 | Neutral |
| testneg_59 | 0 | 0 | 0.147043 | -3 | 1.623152 | 1.623152 | 0.14668 | 0.12788 | 0.19898 | 0.12788 | 20 | 0.010047619 | 0 | Neutral |
| testneg_60 | 0.02 | -0.28 | -1.20584 | -3 | -1.752013 | -3 | 0.53216 | 0.57092 | 0.0609 | 0.47188 | 12.37434 | 0.187851732 | 0 | Neutral |
| testneg_61 | 0.17 | 0.38 | 1.691672 | 1.691672 | 0.110548 | -3 | 0.30789 | 0.15135 | 0.28326 | 0.13944 | 10 | 0.157371951 | 0 | Neutral |
| testneg_62 | 0.19 | -0.27 | 2.160683 | -3 | 1.105669 | -3 | 0.13433 | 0.09578 | 0.44385 | 0.1311 | 10.76923 | 0.04 | 0 | Neutral |
| testneg_63 | 0.15 | 0.04 | 1.146136 | -3 | 3.025435 | -3 | 0.08186 | 0.11813 | 0.25674 | 0.83834 | 20.83333 | 0.244294691 | 0.2 | Neutral |
| testneg_64 | -0.02 | -0.45 | 2.978114 | -3 | 2.442702 | -3 | 0.18047 | 0.2356 | 0.95535 | 0.12788 | 10.83333 | 0.19 | 0 | Neutral |
| testneg_65 | -0.14 | -0.54 | 3.948867 | -3 | 0.142928 | -3 | 0.21236 | 0.12788 | 0.17843 | 0.15468 | 12.81915 | 0.150135135 | 0 | Neutral |
| testneg_66 | 0 | 0 | -0.961702 | -0.961702 | 1.259325 | -3 | 0.46979 | 0.1705 | 0.16394 | 0.08512 | 10.83333 | 0.03 | 0 | Neutral |
| testneg_67 | 0 | 0 | 1.45834 | -3 | 1.325183 | -3 | 0.64701 | 0.94571 | 0.3437 | 0.5169 | 10 | 0.144781197 | 0 | Neutral |
| testneg_68 | -0.02 | 0.68 | 2.752736 | -3 | 2.650636 | 0.593681 | 0.15556 | 0.12841 | 0.82595 | 0.52905 | 13.89287 | 0.220325441 | 0 | Neutral |
| testneg_69 | 0 | 0 | 0.31374 | -3 | 0.491194 | 0.491194 | 0.12582 | 0.12788 | 0.72124 | 0.09906 | 13.51351 | 0.01 | 0 | Neutral |
| testneg_70 | 0 | 0 | 0.347529 | -3 | 0.373317 | -3 | 0.22443 | 0.10657 | 0.06895 | 0.06354 | 10.41667 | 0 | 0 | Neutral |
| testneg_71 | -0.16 | -0.08 | 1.819138 | -3 | 1.164691 | -3 | 0.02791 | 0.0854 | 0.09972 | 0.2187 | 20 | 0.21 | 0 | Neutral |
| testneg_72 | 0.15 | -0.28 | 1.567816 | -3 | 0.707942 | -3 | 0.80979 | 0.5595 | 0.21731 | 0.11795 | 10.41667 | 0.05 | 0 | Neutral |
| testneg_73 | 0 | 0 | 0.181615 | -3 | 1.80072 | -3 | 0.34698 | 0.12788 | 0.24287 | 0.11626 | 18.88889 | 0 | 0 | Neutral |
| testneg_74 | -0.05 | -0.29 | 0.901946 | -3 | -2.190834 | -3 | 0.15055 | 0.22097 | 0.29672 | 0.10485 | 9.02778 | 0.03 | 0 | Neutral |
| testneg_75 | -0.01 | 0.57 | 5.88281 | 5.88281 | 2.136098 | -3 | 0.17322 | 0.10311 | 0.13374 | 0.12788 | 18.70486 | 0.6 | 91.2 | Disease_causing |
| testneg_76 | 0.03 | 0.1 | 0.896188 | -3 | 0.621832 | -3 | 0.42544 | 0.11809 | 0.10244 | 0.10107 | 26.9055 | 0.02 | 0 | Neutral |
| testneg_77 | 0 | 0 | 0.705852 | -3 | 2.296124 | -3 | 0.16689 | 0.11395 | 0.22258 | 0.10471 | 18.6768 | 0.000208916 | 0 | Neutral |
| testneg_78 | 0.18 | -1.05 | 5.667747 | -3 | 1.24588 | -3 | 0.19898 | 0.12788 | 0.13535 | 0.13562 | 6.99956 | 0.275 | 1.6 | Neutral |
| testneg_79 | 0.16 | -0.37 | 6.175848 | -3 | 0.273635 | -3 | 0.19898 | 0.12788 | 0.18578 | 0.27854 | 9.71453 | 0.416367216 | 28 | Neutral |
| testneg_80 | 0.00109091 | -0.0100909 | 1.971045 | -3 | 2.104429 | -3 | 0.14457 | 0.11614 | 0.30585 | 0.10254 | 10.26553 | 0.01 | 0 | Neutral |
| testneg_81 | 0 | -0.01 | 1.221125 | -3 | 6.493314 | -3 | 0.36793 | 0.15588 | 0.33166 | 0.11755 | 11.25 | 0.19 | 0.2 | Neutral |
| testneg_82 | -0.07 | -0.33 | 0.222047 | -3 | 2.252372 | -3 | 0.19898 | 0.12788 | 0.20513 | 0.09801 | 12.26954 | 0.01 | 0 | Neutral |
| testneg_83 | -0.05 | -0.29 | 1.38401 | -3 | 0.402535 | -3 | 0.37212 | 0.11323 | 0.11791 | 0.39607 | 12.7356 | 0.06 | 0 | Neutral |
| testneg_84 | -0.03 | -1.22 | 3.652963 | -3 | 6.969695 | -3 | 0.82269 | 0.15732 | 0.82595 | 0.52905 | 27.1833 | 0.66 | 96.2 | Disease_causing |
| testneg_85 | 0.05 | -0.81 | 6.717504 | -3 | 5.990881 | 0.282726 | 0.19898 | 0.12788 | 0.10314 | 0.12173 | 18.33333 | 0.579517345 | 76.4 | Disease_causing |
| testneg_86 | -0.1 | -1.9 | 2.35005 | -3 | 0.574875 | 0.574875 | 0.64256 | 0.12241 | 0.13468 | 0.09235 | 50.55558 | 0.06 | 0 | Neutral |
| testneg_87 | 0.06 | -0.57 | 4.943137 | -3 | 5.497218 | -3 | 0.15767 | 0.12788 | 0.19898 | 0.12788 | 17.21847 | 0.32 | 8.4 | Neutral |
| testneg_88 | -0.02 | -0.45 | 3.420583 | -3 | 2.339087 | -3 | 0.80034 | 0.10518 | 0.64432 | 0.10409 | 10.30286 | 0.081202381 | 0 | Neutral |
| testneg_89 | -0.06 | -0.18 | -0.580228 | -3 | 0.00733 | -3 | 0.13331 | 0.09729 | 0.13779 | 0.0945 | 17.21847 | 0.01 | 0 | Neutral |
| testneg_90 | 0.02 | 0.45 | 3.151664 | 3.151664 | 0.482528 | -3 | 0.10161 | 0.13484 | 0.19322 | 0.12788 | 18.73594 | 0.401666667 | 20.6 | Neutral |
| testneg_91 | -0.06 | -0.18 | -1.695325 | -3 | 0.545969 | -3 | 0.36052 | 0.2858 | 0.19898 | 0.12788 | 15.5381 | 0.03 | 0 | Neutral |
| testneg_92 | 0 | -0.01 | 0.632891 | -3 | 3.235017 | -3 | 0.28856 | 0.1544 | 0.30329 | 0.14582 | 32.49855 | 0.207701686 | 0 | Neutral |
| testneg_93 | 0.21 | 0.15 | 5.359975 | -3 | 5.080174 | -3 | 0.12196 | 0.12788 | 0.1215 | 0.12788 | 11.25 | 0.431125 | 32 | Neutral |
| testneg_94 | 0 | 0 | 1.510278 | -3 | 1.313527 | -3 | 0.19898 | 0.07875 | 0.06112 | 0.06901 | 18.40909 | 0.01 | 0 | Neutral |
| testneg_95 | 0.18 | 0.16 | -1.868425 | -3 | 3.132136 | -3 | 0.00971 | 0.0791 | 0.0105 | 0.12788 | 19.13548 | 0.12 | 0 | Neutral |
| testneg_96 | -0.08 | -0.03 | 3.717179 | -3 | 6.497236 | -3 | 0.15808 | 0.14548 | 0.17725 | 0.12788 | 10.6968 | 0.47 | 45.2 | Neutral |
| testneg_97 | -0.03 | -1.22 | 0.713058 | -3 | 0.570404 | -3 | 0.14375 | 0.12788 | 0.07571 | 0.09432 | 10.41667 | 0.020093491 | 0 | Neutral |
| testneg_98 | 0.01 | 0.79 | 0.643195 | -3 | -0.274976 | -3 | 0.54777 | 0.25991 | 0.26767 | 0.25506 | 11.58108 | 0.110681555 | 0 | Neutral |
| testneg_99 | 0 | 0 | 0.197229 | -3 | 0.565057 | -3 | 0.99994 | 0.51636 | 0.28474 | 0.12788 | 17.40099 | 0.06 | 0 | Neutral |
| testneg_100 | 0.19 | 0.51 | 5.539608 | -3 | 4.557208 | -3 | 0.35957 | 0.11476 | 0.59864 | 0.14106 | 22.5469 | 0.523156091 | 65.2 | Disease_causing |
| testneg_101 | -0.15 | -0.04 | 3.490664 | -3 | 2.14685 | -3 | 0.29362 | 0.22177 | 0.12161 | 0.24178 | 12.83951 | 0.45 | 35.8 | Neutral |
| testneg_102 | 0.14 | -1.01 | 2.75054 | -3 | -1.080758 | -3 | 0.13691 | 0.11058 | 0.18869 | 0.23984 | 22.91823 | 0.121125 | 0 | Neutral |
| testneg_103 | 0 | 0 | 0.555194 | -3 | 1.392857 | -3 | 0.91921 | 0.32331 | 0.19898 | 0.08714 | 18.33333 | 0.010909662 | 0 | Neutral |
| testneg_104 | 0 | -0.32 | 1.05419 | -3 | 4.225636 | -3 | 0.45152 | 0.34457 | 0.10781 | 0.09901 | 29.9024 | 0.16 | 0 | Neutral |
| testneg_105 | -0.09 | 0.3 | 2.113451 | -3 | 0.134423 | -3 | 0.19749 | 0.11482 | 0.07415 | 0.12788 | 13.37927 | 0.010289916 | 0 | Neutral |
| testneg_106 | -0.01 | 1.44 | -0.133106 | -0.133106 | 2.158998 | -3 | 0.07096 | 0.09965 | 0.81659 | 0.10424 | 11.11802 | 0.12 | 0 | Neutral |
| testneg_107 | 0 | 0 | -0.099687 | -3 | 3.013004 | 3.013004 | 0.09972 | 0.13992 | 0.15537 | 0.19705 | 10.83333 | 0.258111794 | 2 | Neutral |
| testneg_108 | 0.01 | -1.44 | 3.616919 | -3 | 1.821777 | -3 | 0.19898 | 0.12788 | 0.11187 | 0.12788 | 21.58514 | 0.058988782 | 0 | Neutral |
| testneg_109 | 0.05 | -0.81 | 4.343218 | -3 | 0.023521 | -3 | 0.56176 | 0.11451 | 0.16002 | 0.14644 | 10 | 0.104764835 | 0 | Neutral |
| testneg_110 | -0.08 | -0.03 | 3.219123 | -3 | 6.188656 | -3 | 0.31924 | 0.11145 | 0.29624 | 0.1067 | 13.02635 | 0.238739316 | 0.8 | Neutral |
| testneg_111 | -0.09 | 0.3 | 0.912537 | -3 | 2.549873 | -3 | 0.2049 | 0.12678 | 0.04484 | 0.95426 | 16.8018 | 0.172923597 | 0 | Neutral |
| testneg_112 | 0 | 0 | 3.192398 | -3 | -1.543536 | -1.543536 | 0.19898 | 0.12788 | 0.06744 | 0.12788 | 17.21847 | 0.02 | 0 | Neutral |
| testneg_113 | 0 | 0 | 1.872374 | -3 | 6.395815 | -3 | 0.28907 | 0.12788 | 0.52005 | 0.12788 | 22.27594 | 0.13 | 0 | Neutral |
| testneg_114 | 0 | 0 | 0.740874 | -3 | -0.573377 | -1.816133 | 0.1424 | 0.12788 | 0.42366 | 0.14372 | 26.42807 | 0.03260786 | 0 | Neutral |
| testneg_115 | 0.03 | -1.03 | 5.102429 | -3 | -0.392743 | -3 | 0.10723 | 0.11163 | 0.16275 | 0.12788 | 19.21569 | 0.06 | 0 | Neutral |
| testneg_116 | -0.0167273 | 0.08290909 | 13.012264 | 4.777924 | 7.144915 | -3 | 0.24331 | 0.11911 | 0.07415 | 0.12788 | 11.74429 | 0.879775379 | 100 | Disease_causing |
| testneg_117 | 0.21 | 0.15 | 6.917265 | -3 | 1.083169 | -3 | 0.18633 | 0.14783 | 0.26621 | 0.10639 | 10.83333 | 0.295833333 | 4.8 | Neutral |
| testneg_118 | 0 | 0 | 0.741693 | -3 | 0.13879 | -3 | 0.48906 | 0.11262 | 0.113 | 0.12788 | 40.72752 | 0.010106395 | 0 | Neutral |
| testneg_119 | 0 | -0.01 | 7.854373 | -3 | 0.381493 | 0.381493 | 0.4113 | 0.2303 | 0.14432 | 0.16919 | 19.44444 | 0.581892857 | 83.8 | Disease_causing |
| testneg_120 | 0 | 0 | 0.162648 | -3 | 7.056082 | -3 | 0.38002 | 0.22508 | 0.83114 | 0.60631 | 11.25 | 0.521006127 | 60 | Disease_causing |
| testneg_121 | 0 | 0 | 0.376038 | -3 | -0.290062 | -3 | 0.18485 | 0.40044 | 0.13468 | 0.09235 | 39.64383 | 0.04 | 0 | Neutral |

| | | | | | | | | | | | | | | |
|---|---|---|---|---|---|---|---|---|---|---|---|---|---|---|
| testneg_122 | -0.02 | 0.63 | 3.191277 | -3 | -1.097042 | -3 | 0.37074 | 0.11532 | 0.10703 | 0.11611 | 43.53811 | 0.04 | 0 | Neutral |
| testneg_123 | 0 | 0 | 0.496132 | -3 | 1.991433 | -3 | 0.16252 | 0.11134 | 0.25758 | 0.2262 | 12.07437 | 0.020899659 | 0 | Neutral |
| testneg_124 | 0 | 0 | 0.889458 | -3 | -2.581564 | -3 | 0.48937 | 0.22085 | 0.19898 | 0.12788 | 27.35225 | 0.03 | 0 | Neutral |
| testneg_125 | 0.12 | -1.16 | 4.470041 | -3 | 0.952842 | -3 | 0.06841 | 0.10927 | 0.13535 | 0.13562 | 12.30981 | 0.141423784 | 0 | Neutral |
| testneg_126 | 0.16 | -1.37 | 6.261588 | -3 | 0.603761 | -3 | 0.3693 | 0.12788 | 0.53784 | 0.5574 | 10.41667 | 0.435214286 | 31.8 | Neutral |
| testneg_127 | 0.22 | -2.66 | 6.181114 | -3 | -0.031182 | -3 | 0.55384 | 0.26913 | 0.14511 | 0.12788 | 10.83333 | 0.275166667 | 1.6 | Neutral |
| testneg_128 | -0.03 | -0.1 | 6.245625 | -3 | 2.650409 | -3 | 0.05535 | 0.12788 | 0.1324 | 0.09503 | 26.79941 | 0.29 | 2.4 | Neutral |
| testneg_129 | 0.05 | 0.29 | -1.569633 | -3 | 0.07066 | -3 | 0.02759 | 0.12458 | 0.08903 | 0.12788 | 23.13568 | 0.04 | 0 | Neutral |
| testneg_130 | 0.21 | 0.15 | 0.833236 | -3 | 1.840603 | -3 | 0.4172 | 0.11643 | 0.83421 | 0.18817 | 11.25 | 0.08 | 0 | Neutral |
| testneg_131 | 0 | 0 | -0.235365 | -3 | 0.945866 | -3 | 0.99535 | 0.38424 | 0.23537 | 0.33997 | 9.91694 | 0.084479776 | 0 | Neutral |
| testneg_132 | 0 | 0 | 1.190149 | -3 | 2.254589 | -3 | 0.16841 | 0.11143 | 0.63497 | 0.23021 | 21.43373 | 0.053056339 | 0 | Neutral |
| testneg_133 | 0.15 | 0.06 | 7.140042 | -3 | 11.093155 | 6.526897 | 0.04142 | 0.09564 | 0.16692 | 0.09243 | 43.6335 | 0.69 | 99.4 | Disease_causing |
| testneg_134 | -0.07 | -0.7 | -0.548079 | -3 | 1.565943 | -3 | 0.19898 | 0.12788 | 0.17796 | 0.09522 | 23.58961 | 0.02 | 0 | Neutral |
| testneg_135 | 0 | 0 | 1.875005 | -3 | -0.454534 | -3 | 0.50562 | 0.2196 | 0.45628 | 0.15819 | 14.16667 | 0.075053914 | 0 | Neutral |
| testneg_136 | 0.14 | -1.01 | 5.616603 | -3 | 0.506986 | -3 | 0.12836 | 0.09736 | 0.65857 | 0.26294 | 18.88889 | 0.329547817 | 6.4 | Neutral |
| testneg_137 | 0.04 | -0.23 | 4.939659 | -3 | 1.719272 | -3 | 0.13563 | 0.12378 | 0.024 | 0.07726 | 33.53535 | 0.05 | 0 | Neutral |
| testneg_138 | 0 | 0 | 0.452231 | -3 | 4.368284 | -3 | 0.46691 | 0.07886 | 0.78253 | 0.1114 | 13.93735 | 0.040741758 | 0 | Neutral |
| testneg_139 | 0 | 0 | 0.609221 | -3 | 5.752691 | -3 | 0.56904 | 0.14839 | 0.38756 | 0.08351 | 13.19331 | 0.096018707 | 0 | Neutral |
| testneg_140 | 0 | 0 | 2.254535 | -3 | 0.973671 | -3 | 0.21728 | 0.1132 | 0.47246 | 0.47221 | 8.97898 | 0.035411731 | 0 | Neutral |
| testneg_141 | 0.05 | -0.81 | 2.157622 | -3 | 0.29558 | -3 | 0.35967 | 0.09418 | 0.87806 | 0.38966 | 6.94444 | 0.060386166 | 0 | Neutral |
| testneg_142 | 0.04 | -0.23 | 2.305272 | -3 | 1.414729 | -3 | 0.18882 | 0.14637 | 0.33026 | 0.30576 | 13.73701 | 0.120075188 | 0 | Neutral |
| testneg_143 | -0.08 | -0.03 | 3.635933 | -3 | -1.158589 | -3 | 0.21265 | 0.1284 | 0.16362 | 0.11879 | 12.72648 | 0.03 | 0 | Neutral |
| testneg_144 | 0 | 0 | -0.548861 | -3 | 1.367687 | -3 | 0.25179 | 0.09355 | 0.2439 | 0.08848 | 11.01208 | 0.01 | 0 | Neutral |
| testneg_145 | 0 | 0.32 | 2.93612 | -3 | -0.109177 | -3 | 0.05657 | 0.14812 | 0.49278 | 0.09047 | 12.38369 | 0.1 | 0 | Neutral |
| testneg_146 | -0.07 | -0.33 | 1.282461 | -3 | 1.687316 | -3 | 0.59131 | 0.66216 | 0.26434 | 0.12788 | 10.26553 | 0.04 | 0 | Neutral |
| testneg_147 | 0.14 | -1.01 | 3.372659 | -3 | 1.902977 | 1.902977 | 0.15263 | 0.12788 | 0.18504 | 0.0822 | 10.83333 | 0.09 | 0 | Neutral |
| testneg_148 | -0.21 | -0.15 | -1.316616 | -3 | 1.144584 | -3 | 0.07347 | 0.08817 | 0.56817 | 0.10766 | 28.16398 | 0.08 | 0 | Neutral |
| testneg_149 | -0.14 | 1.01 | 5.758583 | 0.133907 | 3.470215 | -3 | 0.19898 | 0.12788 | 0.17412 | 0.10595 | 11.24844 | 0.485217391 | 48.6 | Neutral |
| testneg_150 | 0.03 | 0.12 | 0.199359 | 0.199359 | 0.273693 | -3 | 0.07626 | 0.12788 | 0.2755 | 0.12788 | 23.68839 | 0.01 | 0 | Neutral |
| testneg_151 | 0 | 0 | -0.064925 | -3 | 6.81698 | 0.240475 | 0.25924 | 0.19965 | 0.19898 | 0.535 | 10 | 0.576469298 | 79.2 | Disease_causing |
| testneg_152 | 0 | 0 | -0.179321 | -3 | 0.709734 | -3 | 0.14101 | 0.14318 | 0.04851 | 0.12788 | 16.48772 | 0.010241518 | 0 | Neutral |
| testneg_153 | 0 | -0.01 | 0.639661 | -3 | 1.399164 | -3 | 0.16263 | 0.53493 | 0.61332 | 0.1004 | 10.22898 | 0.03 | 0 | Neutral |
| testneg_154 | -0.01 | 1.44 | -3.047221 | -3 | 2.746161 | -3 | 0.7219 | 0.41509 | 0.17793 | 0.10516 | 21.54762 | 0.202211111 | 0 | Neutral |
| testneg_155 | 0 | 0 | 1.15789 | 1.15789 | -2.464297 | -2.464297 | 0.68439 | 0.09371 | 0.11524 | 0.12788 | 10 | 0.03 | 0 | Neutral |
| testneg_156 | 0 | 0 | 1.175399 | -3 | 3.089647 | -3 | 0.37065 | 0.12788 | 0.03324 | 0.12788 | 19.90393 | 0.036904762 | 0 | Neutral |
| testneg_157 | 0.07 | 0.01 | -0.676534 | -0.676534 | 0.951477 | 0.951477 | 0.16262 | 0.15786 | 0.27083 | 0.14313 | 10.41667 | 0.0896875 | 0 | Neutral |
| testneg_158 | 0 | 0 | 0.965606 | -3 | 0.278187 | -3 | 0.14341 | 0.10134 | 0.21587 | 0.11701 | 18.98413 | 0 | 0 | Neutral |
| testneg_159 | -0.08 | -0.03 | 2.32778 | -3 | 4.526132 | -3 | 0.74488 | 0.12788 | 0.04668 | 0.12788 | 23.98456 | 0.12 | 0 | Neutral |
| testneg_160 | -0.02 | 0.63 | 4.506874 | -3 | 3.054544 | -3 | 0.30942 | 0.13105 | 0.32022 | 0.40347 | 10.83333 | 0.558521297 | 75.2 | Disease_causing |
| testneg_161 | 0 | 0 | 0.847539 | -3 | 1.096737 | 1.096737 | 0.65564 | 0.11468 | 0.23071 | 0.09582 | 12.29167 | 0.01 | 0 | Neutral |
| testneg_162 | -0.16 | -0.08 | 2.98751 | -3 | 2.61063 | -3 | 0.18022 | 0.12788 | 0.04627 | 0.05223 | 27.51545 | 0.103333333 | 0 | Neutral |
| testneg_163 | 0 | 0 | 0.569405 | -3 | 5.893689 | -3 | 0.47544 | 0.17658 | 0.21383 | 0.11171 | 13.22427 | 0.126203934 | 0 | Neutral |
| testneg_164 | 0 | 0 | 0.383952 | -3 | 2.740758 | -3 | 0.28629 | 0.77234 | 0.17325 | 0.17964 | 7.00579 | 0.203527516 | 0 | Neutral |
| testneg_165 | -0.08 | -0.03 | 6.557579 | -3 | 3.677559 | -3 | 0.16741 | 0.10762 | 0.12855 | 0.11656 | 12.47902 | 0.43 | 29.4 | Neutral |
| testneg_166 | 0.14 | -1.01 | 0.712749 | -3 | 3.703007 | 2.033448 | 0.12203 | 0.13515 | 0.13708 | 0.81545 | 24.41367 | 0.463999369 | 39 | Neutral |
| testneg_167 | -0.03 | -0.1 | -0.369846 | -3 | 1.161522 | -3 | 0.33467 | 0.12788 | 0.52765 | 0.20417 | 26.65756 | 0.07035401 | 0 | Neutral |
| testneg_168 | -0.1 | -1.9 | -0.298153 | -3 | -0.625744 | -3 | 0.12099 | 0.11502 | 0.19898 | 0.12788 | 18.22113 | 0.04 | 0 | Neutral |
| testneg_169 | 0.09 | -1.07 | 4.373261 | -3 | -0.676534 | -0.676534 | 0.29356 | 0.47581 | 0.16262 | 0.15786 | 29.16667 | 0.359733974 | 8.6 | Neutral |
| testneg_170 | 0 | 0 | 0.09433 | -3 | -1.060307 | -3 | 0.18925 | 0.32479 | 0.19423 | 0.14002 | 13.54137 | 0.087521083 | 0 | Neutral |
| testneg_171 | 0.14 | -0.69 | -0.111883 | -0.111883 | 2.074047 | -3 | 0.06313 | 0.12788 | 0.09353 | 0.12788 | 21.99535 | 0.04 | 0 | Neutral |
| testneg_172 | -0.09 | 0.67 | 0.082232 | -3 | 0.389489 | -3 | 0.08475 | 0.19889 | 0.11858 | 0.11551 | 13.02635 | 0.07 | 0 | Neutral |
| testneg_173 | -0.08 | -0.03 | 0.405561 | -3 | 7.974494 | -3 | 0.39132 | 0.11179 | 0.11316 | 0.12788 | 8.97898 | 0.354436312 | 10.6 | Neutral |
| testneg_174 | -0.06 | -0.18 | 1.398945 | -3 | 0.681825 | -3 | 0.95993 | 0.15588 | 0.30837 | 0.19754 | 13.06326 | 0.061947832 | 0 | Neutral |
| testneg_175 | -0.17 | -0.38 | -1.129477 | -3 | 0.914524 | -3 | 0.11006 | 0.14352 | 0.31238 | 0.14393 | 35.87042 | 0.140041667 | 0 | Neutral |
| testneg_176 | 0 | 0 | 0.47609 | -3 | 0.866086 | -3 | 0.59131 | 0.66216 | 0.19898 | 0.12788 | 20.52239 | 0.04 | 0 | Neutral |
| testneg_177 | -0.07 | -0.38 | 0.816148 | -3 | 5.882463 | -3 | 0.26347 | 0.16383 | 0.06099 | 0.12788 | 15.10488 | 0.23036445 | 0.2 | Neutral |
| testneg_178 | 0.02 | 0.45 | 3.151664 | -3 | 0.712749 | -3 | 0.10161 | 0.13484 | 0.12203 | 0.13515 | 10.21277 | 0.13 | 0 | Neutral |
| testneg_179 | 0.09 | -0.25 | -0.159404 | -3 | 3.331298 | -3 | 0.95319 | 0.20297 | 0.12682 | 0.11357 | 4.56414 | 0.13191954 | 0 | Neutral |
| testneg_180 | -0.16 | -0.08 | 1.042194 | -3 | 6.136401 | -3 | 0.19898 | 0.12788 | 0.3238 | 0.18333 | 18.76159 | 0.475241071 | 44 | Neutral |
| testneg_181 | 0 | 0 | 0.567808 | -3 | 1.070987 | -3 | 0.03631 | 0.80564 | 0.52768 | 0.12788 | 16.668 | 0.140839161 | 0 | Neutral |
| testneg_182 | 0 | 0 | 0.44595 | -3 | -0.850513 | -3 | 0.15019 | 0.10342 | 0.06982 | 0.10226 | 18.78762 | 0.000177116 | 0 | Neutral |
| testneg_183 | 0.06 | -0.57 | 3.841536 | -3 | 0.79796 | -3 | 0.08002 | 0.08792 | 0.57466 | 0.12788 | 20.55556 | 0.041136364 | 0 | Neutral |
| testneg_184 | -0.15 | -0.04 | 3.116374 | -3 | 0.97098 | -3 | 0.19898 | 0.12788 | 0.75415 | 0.29199 | 25.80569 | 0.171109185 | 0.2 | Neutral |
| testneg_185 | -0.06 | -0.18 | 0.101783 | -3 | 0.179304 | -3 | 0.06196 | 0.10881 | 0.56817 | 0.10766 | 30.98291 | 0.03 | 0 | Neutral |
| testneg_186 | 0.09 | -0.3 | -0.668726 | -3 | -0.674759 | -0.674759 | 0.27924 | 0.12788 | 0.02782 | 0.06218 | 24.4774 | 0.02 | 0 | Neutral |
| testneg_187 | -0.02 | 0.36 | 4.076116 | -3 | 0.48203 | -3 | 0.1331 | 0.12788 | 0.1287 | 0.10423 | 21.92139 | 0.04 | 0 | Neutral |
| testneg_188 | 0.21 | 0.15 | 1.854082 | -3 | 3.381673 | 3.381673 | 0.20084 | 0.0851 | 0.23578 | 0.12788 | 17.36842 | 0.212168831 | 0.2 | Neutral |
| testneg_189 | -0.09 | 0.3 | -1.867658 | -3 | 2.68936 | -3 | 0.19433 | 0.12788 | 0.10703 | 0.11611 | 30.95458 | 0.050989927 | 0 | Neutral |
| testneg_190 | 0.04 | 2.01 | -3.135825 | -3 | 2.757779 | -3 | 0.35422 | 0.11185 | 0.99948 | 0.5595 | 29.46766 | 0.269484307 | 0.2 | Neutral |
| testneg_191 | -0.04 | 0.89 | 0.428266 | -3 | 3.59862 | -3 | 0.05323 | 0.32175 | 0.2016 | 0.13956 | 29.86111 | 0.504736575 | 56.6 | Disease_causing |
| testneg_192 | 0.03 | -1.03 | 4.768167 | -3 | -0.481271 | -3 | 0.91587 | 0.10818 | 0.09332 | 0.53331 | 11.49684 | 0.232386538 | 0 | Neutral |
| testneg_193 | 0.09 | -0.3 | 0.571254 | -3 | 0.586954 | -3 | 0.143 | 0.11236 | 0.4118 | 0.1154 | 17.24563 | 0.01 | 0 | Neutral |
| testneg_194 | 0 | 0 | -0.14981 | -3 | 4.492087 | -3 | 0.10176 | 0.12788 | 0.37424 | 0.12788 | 12.81421 | 0.05 | 0 | Neutral |
| testneg_195 | 0 | -0.01 | 1.723095 | 1.723095 | 0.393827 | -3 | 0.91293 | 0.12788 | 0.07516 | 0.15472 | 4.44444 | 0.111385157 | 0 | Neutral |
| testneg_196 | 0 | 0 | -0.258437 | -3 | 2.5258 | -2.07002 | 0.07044 | 0.11679 | 0.48801 | 0.09118 | 18.47604 | 0.02 | 0 | Neutral |
| testneg_197 | -0.03 | -0.1 | 2.246735 | -3 | 2.296298 | -3 | 0.11827 | 0.23034 | 0.96475 | 0.27545 | 4.72222 | 0.294347673 | 1.8 | Neutral |
| testneg_198 | 0 | 0 | 1.465695 | -3 | -0.099491 | -3 | 0.94822 | 0.14005 | 0.06218 | 0.10661 | 27.43237 | 0.03 | 0 | Neutral |
| testneg_199 | -0.03 | -0.1 | 1.538056 | 0.552009 | 0.452369 | -3 | 0.53172 | 0.12788 | 0.09504 | 0.38939 | 22.63149 | 0.16 | 0 | Neutral |
| testneg_200 | 0 | 0 | -0.137475 | -0.137475 | 1.210569 | -3 | 0.19898 | 0.12788 | 0.19898 | 0.12788 | 19.44444 | 0.000374126 | 0 | Neutral |
| testneg_201 | 0.04 | -0.23 | -0.124406 | -3 | 2.574553 | -3 | 0.11096 | 0.10074 | 0.04525 | 0.12788 | 52.08333 | 0.030691919 | 0 | Neutral |
| testneg_202 | -0.07 | -0.38 | 2.623579 | -3 | 0.730641 | -3 | 0.59812 | 0.19387 | 0.04606 | 0.20522 | 19.44444 | 0.170576923 | 0 | Neutral |
| testneg_203 | 0.02 | 0.45 | 3.982621 | -3 | -1.14689 | -3 | 0.70627 | 0.11276 | 0.14704 | 0.12788 | 32.33639 | 0.05 | 0 | Neutral |
| testneg_204 | -0.03 | -0.1 | 1.738251 | -3 | 0.285371 | -3 | 0.17303 | 0.18733 | 0.85793 | 0.80177 | 10.41667 | 0.15 | 0 | Neutral |
| testneg_205 | -0.17 | 1.22 | -2.338688 | -2.338688 | 0.58387 | -3 | 0.33129 | 0.09957 | 0.13244 | 0.14875 | 19.01709 | 0.167857143 | 0 | Neutral |
| testneg_206 | 0.08 | 0.03 | -0.765287 | -3 | 8.109614 | -3 | 0.21002 | 0.12788 | 0.25896 | 0.45481 | 14.00313 | 0.44795098 | 35.8 | Neutral |
| testneg_207 | -0.08 | -0.03 | 1.87433 | -3 | 1.308546 | -3 | 0.95857 | 0.12137 | 0.12545 | 0.08963 | 16.84579 | 0.02 | 0 | Neutral |
| testneg_208 | 0.18 | 0.16 | 1.009156 | -3 | 1.5447 | -3 | 0.19898 | 0.12788 | 0.08932 | 0.32882 | 19.09347 | 0.128882094 | 0 | Neutral |
| testneg_209 | 0 | 0 | -0.294826 | -3 | 3.7974 | 2.311448 | 0.63818 | 0.12788 | 0.1231 | 0.3662 | 71.36923 | 0.442188645 | 24.8 | Neutral |
| testneg_210 | 0 | 0 | 1.052989 | -3 | -0.669222 | -3 | 0.9874 | 0.15588 | 0.22225 | 0.11186 | 10 | 0.010037037 | 0 | Neutral |
| testneg_211 | -0.03 | -0.1 | 1.942918 | -3 | 0.323793 | -3 | 0.11187 | 0.10744 | 0.05315 | 0.12788 | 24.25984 | 0.02 | 0 | Neutral |
| testneg_212 | 0.18 | -1.05 | 5.383128 | -3 | 0.933869 | -3 | 0.08999 | 0.0989 | 0.9862 | 0.4528 | 5 | 0.30964886 | 3.2 | Neutral |
| testneg_213 | 0 | 0 | 0.590474 | -3 | -2.240872 | -3 | 0.14484 | 0.14928 | 0.14175 | 0.12788 | 10.83333 | 0.02 | 0 | Neutral |
| testneg_214 | 0.08 | 0.03 | 1.082238 | -3 | 2.686945 | -2.460553 | 0.27062 | 0.12378 | 0.08499 | 0.08678 | 20.83333 | 0.04 | 0 | Neutral |
| testneg_215 | 0 | 0 | 1.531907 | -3 | 7.187505 | -3 | 0.19898 | 0.14509 | 0.31344 | 0.12788 | 10.1991 | 0.29 | 4.4 | Neutral |
| testneg_216 | -0.08 | -0.03 | -0.472766 | -3 | 2.82342 | -3 | 0.15119 | 0.08823 | 0.16462 | 0.18467 | 26.34995 | 0.14516129 | 0 | Neutral |
| testneg_217 | -0.16 | -0.08 | 0.294338 | -3 | 0.782319 | -3 | 0.86306 | 0.10962 | 0.42235 | 0.12474 | 17.21847 | 0.06 | 0 | Neutral |
| testneg_218 | 0.03 | 0.1 | -1.932308 | -3 | -1.08606 | -3 | 0.06474 | 0.23607 | 0.28269 | 0.12505 | 12.95455 | 0.06 | 0 | Neutral |
| testneg_219 | 0.07 | -0.31 | 0.573002 | -3 | 0.494557 | -3 | 0.34205 | 0.11195 | 0.06476 | 0.07491 | 13.24405 | 0.000765017 | 0 | Neutral |
| testneg_220 | 0 | 0 | 1.172534 | -3 | 0.058546 | -3 | 0.04056 | 0.55024 | 0.18998 | 0.26095 | 10.83333 | 0.230936195 | 0 | Neutral |
| testneg_221 | 0.15 | 0.06 | -0.490791 | -3 | 4.431523 | -3 | 0.06953 | 0.17439 | 0.14007 | 0.10647 | 12.29167 | 0.197138905 | 0 | Neutral |
| testneg_222 | 0.16 | -0.37 | 2.530692 | -3 | 1.081088 | -3 | 0.11329 | 0.12116 | 0.21201 | 0.20962 | 11.5657 | 0.074642857 | 0 | Neutral |
| testneg_223 | 0 | 0 | 0.177066 | -3 | 2.892104 | -3 | 0.30275 | 0.12788 | 0.39344 | 0.12788 | 13.46631 | 0.01 | 0 | Neutral |
| testneg_224 | 0 | 0 | -0.024839 | -3 | 1.592541 | -3 | 0.17935 | 0.36953 | 0.65981 | 0.47389 | 19.44444 | 0.163333333 | 0 | Neutral |
| testneg_225 | 0.01 | -1.44 | 2.15882 | -3 | 3.206814 | -3 | 0.90767 | 0.12788 | 0.7022 | 0.35308 | 11.25 | 0.255618291 | 0.4 | Neutral |
| testneg_226 | -0.06 | -0.18 | 2.727897 | -3 | 0.089943 | -3 | 0.18047 | 0.2356 | 0.09556 | 0.10109 | 16.84579 | 0.043801215 | 0 | Neutral |
| testneg_227 | 0.15 | -0.28 | 0.118932 | 0.118932 | 1.739888 | -3 | 0.19567 | 0.09606 | 0.10031 | 0.10724 | 15.19178 | 0.02 | 0 | Neutral |
| testneg_228 | 0.21 | 0.15 | 5.657785 | -3 | 1.802906 | -3 | 0.16497 | 0.74918 | 0.33886 | 0.15644 | 19.44444 | 0.587923813 | 86.8 | Disease_causing |
| testneg_229 | 0 | -0.01 | 1.480753 | -3 | 1.20271 | -3 | 0.26264 | 0.13812 | 0.57991 | 0.2033 | 10.83333 | 0.026801388 | 0 | Neutral |
| testneg_230 | 0.07 | 0.38 | -2.492731 | -3 | 0.245113 | -3 | 0.24832 | 0.84371 | 0.46399 | 0.15334 | 5 | 0.16 | 0 | Neutral |
| testneg_231 | 0.01 | -1.3 | 5.149724 | -3 | 4.162376 | -3 | 0.29679 | 0.12788 | 0.71511 | 0.34803 | 9.82456 | 0.671206941 | 97 | Disease_causing |
| testneg_232 | -0.04 | 0.23 | -0.77433 | -3 | -0.106366 | -3 | 0.51167 | 0.13142 | 0.19898 | 0.12788 | 12.31894 | 0.026156938 | 0 | Neutral |
| testneg_233 | 0.03 | 0.12 | 1.186176 | -3 | 0.828564 | -3 | 0.19898 | 0.12788 | 0.7062 | 0.83002 | 9.67871 | 0.06 | 0 | Neutral |
| testneg_234 | 0 | 0 | 0.221415 | -3 | 1.091905 | 0.580057 | 0.89972 | 0.10766 | 0.10058 | 0.13882 | 16.80958 | 0.074008117 | 0 | Neutral |
| testneg_235 | -0.07 | 0.31 | -0.564575 | -3 | 0.017266 | -3 | 0.68029 | 0.12788 | 0.1228 | 0.10482 | 16.84579 | 0.02017564 | 0 | Neutral |
| testneg_236 | 0 | 0 | 2.049487 | -3 | -0.027869 | -3 | 0.54349 | 0.60885 | 0.19898 | 0.12788 | 5 | 0.02 | 0 | Neutral |
| testneg_237 | 0.07 | 0.7 | 5.078544 | -3 | 0.242603 | -3 | 0.2098 | 0.44672 | 0.32333 | 0.23356 | 12.70833 | 0.421111111 | 26.2 | Neutral |
| testneg_238 | 0 | 0 | -0.433937 | -0.433937 | 0.980675 | -3 | 0.84572 | 0.78546 | 0.35137 | 0.09161 | 8.45238 | 0.030098049 | 0 | Neutral |
| testneg_239 | 0 | 0 | -0.274465 | -3 | -1.728128 | -3 | 0.04699 | 0.08149 | 0.37635 | 0.16791 | 5 | 0.08 | 0 | Neutral |
| testneg_240 | -0.03 | -0.12 | 5.938598 | -3 | 0.941706 | -3 | 0.10835 | 0.12788 | 0.15294 | 0.12788 | 19.09021 | 0.18 | 0 | Neutral |
| testneg_241 | 0 | 0 | 0.840063 | -3 | 1.29806 | -3 | 0.08683 | 0.26961 | 0.27887 | 0.19826 | 9.77836 | 0.09024359 | 0 | Neutral |
| testneg_242 | 0 | 0 | 1.590262 | -3 | 0.590072 | -3 | 0.44385 | 0.1311 | 0.0819 | 0.1823 | 11.91142 | 0.053210784 | 0 | Neutral |
| testneg_243 | 0.12 | -1.16 | 4.936092 | -3 | 0.994999 | 0.994999 | 0.2118 | 0.2009 | 0.51282 | 0.12075 | 20.27778 | 0.131 | 0 | Neutral |

| | | | | | | | | | | | | | | |
|---|---|---|---|---|---|---|---|---|---|---|---|---|---|---|
| testneg_244 | 0.18 | -1.05 | 6.443034 | -3 | 3.355696 | -3 | 0.1846 | 0.375 | 0.16098 | 0.12788 | 35.94982 | 0.480416667 | 49 | Neutral |
| testneg_245 | 0 | 0 | -0.137475 | -0.137475 | 0.014287 | -3 | 0.19898 | 0.12788 | 0.30103 | 0.1462 | 20 | 0.040757143 | 0 | Neutral |
| testneg_246 | -0.09 | 0.67 | 0.082232 | -3 | -2.177154 | -4.48597 | 0.08475 | 0.19889 | 0.10362 | 0.12788 | 19.90373 | 0.077161172 | 0 | Neutral |
| testneg_247 | 0.14 | -0.69 | 3.352085 | -3 | 0.35706 | -3 | 0.19898 | 0.12788 | 0.12921 | 0.11605 | 23.97163 | 0.020248771 | 0 | Neutral |
| testneg_248 | 0.04 | -0.23 | 0.982052 | -3 | 6.161965 | -3 | 0.09211 | 0.08973 | 0.09569 | 0.09246 | 22.78986 | 0.12 | 0 | Neutral |
| testneg_249 | 0.07 | -0.31 | 2.441441 | -3 | 2.174391 | 1.770701 | 0.08224 | 0.29179 | 0.10213 | 0.22954 | 10.41667 | 0.32664652 | 4.2 | Neutral |
| testneg_250 | 0.17 | 0.38 | 1.490909 | -3 | 0.270884 | 0.270884 | 0.90259 | 0.31502 | 0.24536 | 0.38555 | 4.72222 | 0.210643225 | 0 | Neutral |
| testneg_251 | -0.08 | -0.03 | 0.750291 | -3 | 0.005583 | -3 | 0.09275 | 0.14911 | 0.59066 | 0.10874 | 20.83333 | 0.050204167 | 0 | Neutral |
| testneg_252 | -0.09 | 0.3 | 3.208069 | -3 | 1.698695 | -3 | 0.24327 | 0.12192 | 0.59302 | 0.10897 | 22.83743 | 0.04 | 0 | Neutral |
| testneg_253 | -0.15 | 0.82 | 2.970884 | -3 | -0.176493 | -3 | 0.29362 | 0.22177 | 0.95978 | 0.09493 | 10.38482 | 0.152609649 | 0 | Neutral |
| testneg_254 | -0.03 | -0.1 | 5.112742 | -3 | 0.5942 | -3 | 0.54231 | 0.15254 | 0.07748 | 0.11591 | 11.25723 | 0.131458333 | 0 | Neutral |
| testneg_255 | 0 | 0 | -0.342253 | -3 | 3.945475 | -3 | 0.63363 | 0.14795 | 0.27115 | 0.14766 | 17.28707 | 0.266022727 | 0.6 | Neutral |
| testneg_256 | 0 | 0.32 | -0.420859 | -3 | -1.509154 | -3 | 0.0597 | 0.18092 | 0.19898 | 0.12788 | 20.78544 | 0.06 | 0 | Neutral |
| testneg_257 | -0.05 | -0.29 | -1.086101 | -3 | 0.434159 | -3 | 0.20557 | 0.10131 | 0.10168 | 0.12788 | 20.91877 | 0.02 | 0 | Neutral |
| testneg_258 | 0.01 | 0.8 | 5.619978 | -3 | 6.905837 | -3 | 0.19898 | 0.82835 | 0.27004 | 0.17052 | 12.29167 | 0.929522222 | 100 | Disease_causing |
| testneg_259 | 0.09 | -0.3 | 5.443984 | -3 | 1.7235 | -3 | 0.28271 | 0.60265 | 0.19898 | 0.10778 | 20.85753 | 0.338874005 | 3.8 | Neutral |
| testneg_260 | 0.05 | -0.81 | -0.762363 | -3 | 3.277724 | -3 | 0.57239 | 0.16622 | 0.10438 | 0.20558 | 23.89693 | 0.38814881 | 7.8 | Neutral |
| testneg_261 | 0 | 0 | 0.280928 | -3 | 2.581966 | -3 | 0.28211 | 0.25332 | 0.10738 | 0.0974 | 27.4024 | 0.052678571 | 0 | Neutral |
| testneg_262 | 0.21 | 0.15 | 1.506819 | -3 | 5.71339 | -3 | 0.18744 | 0.09392 | 0.0778 | 0.0728 | 17.66703 | 0.12 | 0 | Neutral |
| testneg_263 | 0.00109091 | -0.0100909 | 1.971045 | -3 | 4.003489 | -3 | 0.14457 | 0.11614 | 0.48504 | 0.12788 | 9.85816 | 0.065 | 0 | Neutral |
| testneg_264 | 0.07 | 0.38 | 1.419746 | -3 | 5.8239 | -3 | 0.1192 | 0.10391 | 0.3112 | 0.33703 | 10.11674 | 0.371805556 | 11.6 | Neutral |
| testneg_265 | 0 | 0 | 1.648624 | -3 | 2.423922 | -3 | 0.54248 | 0.12333 | 0.28334 | 0.10368 | 18.6768 | 0.020082584 | 0 | Neutral |
| testneg_266 | 0.02 | 0.41 | 8.488711 | -3 | 6.189768 | -3 | 0.19102 | 0.12788 | 0.25448 | 0.15814 | 16.8018 | 0.782581349 | 100 | Disease_causing |
| testneg_267 | 0 | 0 | 0.001837 | -3 | 1.381361 | -3 | 0.36486 | 0.14268 | 0.20379 | 0.43705 | 17.21847 | 0.056352764 | 0 | Neutral |
| testneg_268 | 0 | 0 | 0.397741 | -3 | 3.608471 | -3 | 0.05378 | 0.12788 | 0.65048 | 0.12778 | 23.38932 | 0.09 | 0 | Neutral |
| testneg_269 | 0.04 | 2.01 | -0.602009 | -3 | 0.450881 | -3 | 0.17529 | 0.09662 | 0.13328 | 0.10117 | 33.62403 | 0.070360332 | 0 | Neutral |
| testneg_270 | 0.22 | -2.66 | 6.272129 | -3 | 4.286469 | -3 | 0.65674 | 0.10572 | 0.18328 | 0.12788 | 41.02985 | 0.369555336 | 13.6 | Neutral |
| testneg_271 | -0.02 | -0.41 | 2.303477 | -3 | 1.84534 | -3 | 0.53216 | 0.84775 | 0.40996 | 0.57507 | 11.95767 | 0.223205882 | 0.2 | Neutral |
| testneg_272 | -0.05 | -0.29 | 0.368341 | -3 | 1.451698 | -3 | 0.77847 | 0.10751 | 0.18979 | 0.07948 | 11.25 | 0.010111903 | 0 | Neutral |
| testneg_273 | 0.09 | -0.3 | 6.525765 | -3 | 0.25161 | -3 | 0.18866 | 0.12788 | 0.213 | 0.09535 | 38.18837 | 0.176824324 | 0.4 | Neutral |
| testneg_274 | 0.06 | -0.57 | 0.402174 | -3 | 2.210341 | -3 | 0.22892 | 0.10495 | 0.38695 | 0.12579 | 11.69931 | 0.01 | 0 | Neutral |
| testneg_275 | 0.21 | 0.15 | 1.623618 | -3 | 0.562804 | -3 | 0.10083 | 0.17511 | 0.11692 | 0.08324 | 19.09347 | 0.050591071 | 0 | Neutral |
| testneg_276 | -0.03 | -0.1 | 4.022456 | -3 | 5.070556 | -3 | 0.56416 | 0.12788 | 0.10185 | 0.12788 | 27.34174 | 0.249857143 | 1.2 | Neutral |
| testneg_277 | 0 | 0 | 0.346053 | -3 | 3.230275 | -3 | 0.3047 | 0.11929 | 0.19898 | 0.10996 | 19.09347 | 0.020820076 | 0 | Neutral |
| testneg_278 | -0.06 | 0.57 | 1.295804 | -3 | 1.655515 | 0.884796 | 0.64795 | 0.21581 | 0.26337 | 0.12771 | 5 | 0.05 | 0 | Neutral |
| testneg_279 | -0.2 | 1.26 | 1.53776 | 1.53776 | 3.542729 | -3 | 0.29455 | 0.12388 | 0.09273 | 0.08316 | 18.78762 | 0.21 | 0 | Neutral |
| testneg_280 | -0.03 | 1.03 | -0.361103 | -3 | 0.829043 | -3 | 0.26982 | 0.12788 | 0.23141 | 0.12788 | 24.68926 | 0.020303309 | 0 | Neutral |
| testneg_281 | -0.1 | -1.9 | 1.091771 | -3 | 0.309305 | -3 | 0.7174 | 0.15815 | 0.16973 | 0.17555 | 23.19121 | 0.150409091 | 0.2 | Neutral |
| testneg_282 | -0.06 | -0.18 | 1.730485 | -3 | 2.531439 | -3 | 0.30275 | 0.12788 | 0.16616 | 0.11073 | 15.34131 | 0.010310544 | 0 | Neutral |
| testneg_283 | 0.02 | -0.36 | -0.250215 | -3 | 0.870324 | -3 | 0.14705 | 0.08702 | 0.16355 | 0.12788 | 17.21847 | 0.000282078 | 0 | Neutral |
| testneg_284 | -0.15 | -0.04 | 0.591299 | -3 | 2.392808 | -3 | 0.07293 | 0.12989 | 0.09176 | 0.07645 | 22.92402 | 0.080871212 | 0 | Neutral |
| testneg_285 | 0 | 0 | 1.445184 | 1.445184 | -0.895031 | -3 | 0.33469 | 0.17432 | 0.07631 | 0.12788 | 10 | 0.03 | 0 | Neutral |
| testneg_286 | 0.09 | -0.3 | 5.619423 | -3 | 0.840434 | -3 | 0.82455 | 0.2093 | 0.48936 | 0.10963 | 13.99351 | 0.23 | 0.4 | Neutral |
| testneg_287 | 0.04 | -0.23 | 4.115116 | -3 | 1.285558 | 1.285558 | 0.09532 | 0.35459 | 0.18122 | 0.16895 | 29.724 | 0.469710505 | 42.2 | Neutral |
| testneg_288 | 0.07 | 0.33 | 1.928887 | -3 | 5.395477 | -3 | 0.05324 | 0.19446 | 0.35908 | 0.18099 | 10.83333 | 0.664988047 | 96.8 | Disease_causing |
| testneg_289 | 0.02 | -0.68 | 1.222356 | -3 | 0.212473 | -3 | 0.17631 | 0.14648 | 0.61098 | 0.11179 | 34.79902 | 0.02 | 0 | Neutral |
| testneg_290 | 0.02 | -0.63 | 2.090818 | -3 | 2.519804 | -3 | 0.12974 | 0.11886 | 0.56605 | 0.15893 | 19.44444 | 0.130121951 | 0 | Neutral |
| testneg_291 | 0 | 0 | 0.461408 | -3 | 9.117416 | -3 | 0.16797 | 0.11809 | 0.44953 | 0.14403 | 10.54354 | 0.400620726 | 19 | Neutral |
| testneg_292 | 0 | 0 | 0.391614 | -3 | 14.354326 | -3 | 0.9449 | 0.9378 | 0.17006 | 0.14718 | 10 | 0.513330228 | 59.8 | Disease_causing |
| testneg_293 | -0.02 | -0.45 | -0.076098 | -3 | 4.985443 | -3 | 0.1878 | 0.08946 | 0.09618 | 0.10294 | 13.51351 | 0.074212415 | 0 | Neutral |
| testneg_294 | 0 | 0 | 0.943238 | -3 | 0.939477 | -3 | 0.19898 | 0.58572 | 0.12497 | 0.09805 | 21.53664 | 0.053333333 | 0 | Neutral |
| testneg_295 | -0.05 | -0.29 | 5.329404 | -3 | 4.082785 | -3 | 0.78741 | 0.22085 | 0.07204 | 0.07708 | 10.22222 | 0.393043531 | 19 | Neutral |
| testneg_296 | 0 | 0 | 0.337778 | -3 | 0.865658 | -3 | 0.24287 | 0.11626 | 0.64108 | 0.51829 | 19.07944 | 0.0325 | 0 | Neutral |
| testneg_297 | 0 | 0 | 2.652503 | -3 | 3.206825 | -3 | 0.08059 | 0.09576 | 0.55197 | 0.11305 | 35.60847 | 0.06 | 0 | Neutral |
| testneg_298 | -0.01 | -0.79 | 5.941706 | -3 | 1.995788 | -3 | 0.32065 | 0.19642 | 0.19043 | 0.07855 | 13.72549 | 0.320416667 | 4.2 | Neutral |
| testneg_299 | 0.19 | -0.25 | 0.670876 | -3 | 6.431621 | 6.431621 | 0.09645 | 0.13239 | 0.26676 | 0.76568 | 16.94934 | 0.652449673 | 96 | Disease_causing |
| testneg_300 | 0.06 | -0.57 | 0.983026 | -3 | 3.022813 | -3 | 0.02791 | 0.0854 | 0.17301 | 0.10048 | 12.31404 | 0.07 | 0 | Neutral |
| testneg_301 | 0 | 0 | 0.57285 | -3 | 0.3368 | -3 | 0.20501 | 0.6166 | 0.08837 | 0.10939 | 18.88889 | 0.05 | 0 | Neutral |
| testneg_302 | -0.01 | -0.8 | 3.501031 | -3 | 3.002533 | 3.002533 | 0.25421 | 0.16235 | 0.14304 | 0.10715 | 44.58333 | 0.26 | 0.4 | Neutral |
| testneg_303 | -0.16 | -0.08 | -1.975003 | -1.975003 | 6.958432 | -3 | 0.08465 | 0.15428 | 0.20632 | 0.12788 | 20.2584 | 0.529015152 | 68.8 | Disease_causing |
| testneg_304 | 0.09 | -0.3 | 5.693708 | -3 | 3.958232 | -3 | 0.28271 | 0.60265 | 0.44479 | 0.20004 | 18.88889 | 0.927100754 | 100 | Disease_causing |
| testneg_305 | -0.02 | -0.45 | 6.768699 | -3 | 3.063227 | -3 | 0.99531 | 0.86203 | 0.05153 | 0.12788 | 10.41667 | 0.616375 | 92.2 | Disease_causing |
| testneg_306 | -0.14 | 1.01 | 6.237563 | -3 | 5.204992 | -3 | 0.16973 | 0.17555 | 0.21118 | 0.10074 | 13.51351 | 0.66988537 | 96 | Disease_causing |
| testneg_307 | 0 | -0.01 | 0.36151 | -3 | 1.574468 | -3 | 0.28878 | 0.16765 | 0.12121 | 0.4142 | 10 | 0.090174268 | 0 | Neutral |
| testneg_308 | 0.18 | -1.05 | 5.037392 | -3 | 0.254036 | -3 | 0.18224 | 0.10419 | 0.14601 | 0.12788 | 12.01754 | 0.06 | 0 | Neutral |
| testneg_309 | 0 | 0 | 0.509496 | -3 | -0.207193 | -3 | 0.54724 | 0.12788 | 0.41383 | 0.119 | 9.66667 | 0.000795833 | 0 | Neutral |
| testneg_310 | -0.09 | 0.3 | 0.840416 | -3 | 3.854582 | -3 | 0.2058 | 0.28174 | 0.44953 | 0.14403 | 11.25 | 0.473014368 | 43 | Neutral |
| testneg_311 | 0 | 0 | 1.545946 | -3 | 0.253504 | -3 | 0.03844 | 0.12788 | 0.53325 | 0.48065 | 12.32809 | 0.08 | 0 | Neutral |
| testneg_312 | 0 | 0 | 1.011117 | -3 | 5.360167 | -3 | 0.15524 | 0.10607 | 0.09918 | 0.11061 | 34.79823 | 0.06 | 0 | Neutral |
| testneg_313 | 0.14 | -0.69 | -0.176192 | -3 | -2.044408 | -3 | 0.21843 | 0.85214 | 0.10084 | 0.12788 | 10 | 0.090082645 | 0 | Neutral |
| testneg_314 | 0 | 0 | 0.542661 | -3 | 0.032222 | -3 | 0.32531 | 0.12351 | 0.14941 | 0.0838 | 46.50072 | 0.000224719 | 0 | Neutral |
| testneg_315 | 0 | 0 | 0.546525 | -3 | 3.002533 | 3.002533 | 0.12545 | 0.08963 | 0.14304 | 0.10715 | 44.88078 | 0.12 | 0 | Neutral |
| testneg_316 | 0 | 0 | -0.090459 | -0.090459 | 2.076639 | -3 | 0.15581 | 0.19351 | 0.50505 | 0.11563 | 16.8018 | 0.02 | 0 | Neutral |
| testneg_317 | 0.06 | 0.18 | 3.024681 | -3 | 0.185804 | -3 | 0.40557 | 0.11865 | 0.08306 | 0.12788 | 13.02635 | 0.020671759 | 0 | Neutral |
| testneg_318 | -0.06 | -0.18 | 5.5303 | -3 | 0.433941 | 0.433941 | 0.07467 | 0.09224 | 0.30475 | 0.21064 | 10.25326 | 0.38 | 17.4 | Neutral |
| testneg_319 | -0.08 | -0.03 | 1.518871 | -3 | 2.019586 | -3 | 0.52953 | 0.10241 | 0.44488 | 0.31694 | 12.01128 | 0.08 | 0 | Neutral |
| testneg_320 | 0 | 0 | 0.379383 | -3 | 1.645376 | -3 | 0.33127 | 0.49952 | 0.38184 | 0.10478 | 18.46051 | 0.023770476 | 0 | Neutral |
| testneg_321 | -0.03 | -0.12 | 1.627305 | -3 | 0.051776 | -3 | 0.08448 | 0.17914 | 0.53157 | 0.50154 | 10 | 0.170717391 | 0 | Neutral |
| testneg_322 | -0.18 | 1.05 | -0.907393 | -3 | 2.355419 | -3 | 0.05332 | 0.12788 | 0.82595 | 0.52905 | 10.00351 | 0.258714286 | 0.4 | Neutral |
| testneg_323 | 0.01 | 0.8 | 4.81138 | -3 | -0.044729 | -3 | 0.284 | 0.15339 | 0.15989 | 0.12788 | 24.54084 | 0.09 | 0 | Neutral |
| testneg_324 | 0.03 | 0.12 | 5.213775 | -3 | 3.043853 | -3 | 0.10671 | 0.10867 | 0.58985 | 0.10504 | 13.06078 | 0.21 | 0.4 | Neutral |
| testneg_325 | 0.14 | -1.01 | 3.493531 | -3 | 0.416226 | -3 | 0.09997 | 0.1797 | 0.32111 | 0.12788 | 27.65104 | 0.099481481 | 0 | Neutral |
| testneg_326 | 0.03 | -1.03 | 0.315829 | -3 | -2.240872 | -3 | 0.15612 | 0.11465 | 0.14175 | 0.12788 | 29.5012 | 0.014461207 | 0 | Neutral |
| testneg_327 | 0.05 | -0.81 | 1.982936 | -3 | -0.307406 | -3 | 0.07044 | 0.11679 | 0.62858 | 0.16645 | 12.40248 | 0.07 | 0 | Neutral |
| testneg_328 | 0.07 | 0.7 | 5.90163 | -3 | -1.097042 | -3 | 0.15856 | 0.10994 | 0.10703 | 0.11611 | 32.91244 | 0.120237722 | 0 | Neutral |
| testneg_329 | -0.02 | -0.41 | 0.433545 | -3 | 0.586954 | -3 | 0.06996 | 0.35923 | 0.4118 | 0.1154 | 19.44444 | 0.070261986 | 0 | Neutral |
| testneg_330 | -0.0430909 | 0.216 | 9.272843 | -3 | 3.489496 | -3 | 0.24544 | 0.0894 | 0.69254 | 0.10021 | 24.8846 | 0.5005 | 56.6 | Disease_causing |
| testneg_331 | 0.03 | 0.12 | 0.607888 | -3 | 3.313271 | -3 | 0.57349 | 0.09764 | 0.12147 | 0.1486 | 12.40248 | 0.223121212 | 0 | Neutral |
| testneg_332 | -0.04 | 0.23 | 3.260847 | -3 | 2.393948 | 2.393948 | 0.27605 | 0.12788 | 0.70819 | 0.5975 | 9.10066 | 0.352426471 | 7 | Neutral |
| testneg_333 | 0.03 | 0.12 | -0.072035 | -3 | 2.780623 | -3 | 0.28227 | 0.24183 | 0.13143 | 0.90853 | 17.4261 | 0.341194171 | 4.8 | Neutral |
| testneg_334 | -0.1 | -1.9 | 1.091771 | -3 | -0.734901 | -3 | 0.7174 | 0.15815 | 0.64721 | 0.8758 | 10.41667 | 0.16 | 0 | Neutral |
| testneg_335 | -0.06 | -0.18 | 1.398739 | -3 | 0.040505 | -3 | 0.54113 | 0.11017 | 0.16926 | 0.11871 | 14.33332 | 0.01 | 0 | Neutral |
| testneg_336 | 0 | 0 | 0.571304 | -3 | 2.811366 | -3 | 0.99448 | 0.67288 | 0.61718 | 0.1648 | 6.21747 | 0.150234466 | 0 | Neutral |
| testneg_337 | -0.03 | -0.1 | 4.062248 | -3 | 4.631076 | -3 | 0.08779 | 0.1777 | 0.75906 | 0.09702 | 10.88561 | 0.402460784 | 17 | Neutral |
| testneg_338 | 0.14 | -1.01 | 1.041001 | -3 | 0.817193 | -3 | 0.36858 | 0.09681 | 0.4719 | 0.33344 | 11.25 | 0.082145249 | 0 | Neutral |
| testneg_339 | 0 | 0 | 1.323079 | -3 | 4.00403 | -3 | 0.5103 | 0.10923 | 0.07887 | 0.07487 | 36.60187 | 0.08 | 0 | Neutral |
| testneg_340 | -0.15 | -0.04 | 4.866108 | -3 | 1.931574 | -3 | 0.19898 | 0.12788 | 0.49278 | 0.09047 | 29.40656 | 0.097083333 | 0 | Neutral |
| testneg_341 | 0.21 | 0.15 | 6.428561 | -3 | 1.091905 | 1.091905 | 0.1602 | 0.11262 | 0.10058 | 0.13882 | 24.44575 | 0.26610119 | 2.2 | Neutral |
| testneg_342 | -0.17 | -0.38 | 1.808824 | 1.808824 | -0.405389 | -0.405389 | 0.2445 | 0.53069 | 0.14996 | 0.09121 | 15.57324 | 0.14 | 0 | Neutral |
| testneg_343 | 0 | 0 | 1.491195 | -3 | 2.398979 | 2.398979 | 0.03391 | 0.09745 | 0.93246 | 0.12788 | 33.95664 | 0.1 | 0 | Neutral |
| testneg_344 | 0 | -0.32 | 1.978841 | -3 | 0.393827 | -3 | 0.19898 | 0.12788 | 0.07516 | 0.15472 | 4.55594 | 0.07 | 0 | Neutral |
| testneg_345 | 0.14 | -1.01 | 2.16064 | -3 | -1.031314 | -3 | 0.16734 | 0.15176 | 0.27409 | 0.10785 | 24.02248 | 0.04 | 0 | Neutral |
| testneg_346 | 0 | -0.32 | -2.345951 | -3 | -0.384446 | -3 | 0.13199 | 0.14263 | 0.36829 | 0.11457 | 30.55556 | 0.03 | 0 | Neutral |
| testneg_347 | 0.15 | 0.04 | 0.261359 | -3 | 1.13087 | -3 | 0.84616 | 0.1471 | 0.2172 | 0.10891 | 10.83333 | 0.040565742 | 0 | Neutral |
| testneg_348 | 0 | 0 | 1.447389 | 1.447389 | 4.672344 | -3 | 0.14419 | 0.1668 | 0.80034 | 0.10518 | 13.51351 | 0.18966572 | 0 | Neutral |
| testneg_349 | -0.03 | -0.1 | 6.211821 | -3 | 0.828564 | -3 | 0.1215 | 0.12788 | 0.7062 | 0.83002 | 10.41667 | 0.422514468 | 31.2 | Neutral |
| testneg_350 | 0 | 0 | -0.134529 | -3 | 0.679359 | -3 | 0.46919 | 0.60081 | 0.16544 | 0.1313 | 20.83333 | 0.110537613 | 0 | Neutral |
| testneg_351 | -0.07 | -0.38 | -1.026052 | -3 | 4.13176 | -3 | 0.16469 | 0.16042 | 0.226 | 0.28784 | 12.18072 | 0.477659075 | 45.6 | Neutral |
| testneg_352 | 0 | -0.01 | 0.631034 | -3 | 0.932751 | -3 | 0.19898 | 0.12788 | 0.15612 | 0.11465 | 13.46801 | 0 | 0 | Neutral |
| testneg_353 | -0.02 | -0.45 | 2.958368 | -3 | 5.984963 | -3 | 0.1413 | 0.74044 | 0.11114 | 0.09582 | 10.83333 | 0.439931507 | 29.6 | Neutral |
| testneg_354 | -0.19 | 0.25 | 4.16391 | -3 | 7.320532 | -3 | 0.46058 | 0.12903 | 0.19898 | 0.12788 | 31.61597 | 0.519212121 | 63.4 | Disease_causing |
| testneg_355 | 0 | -0.01 | 1.233202 | -3 | 2.387251 | 2.294938 | 0.22034 | 0.12788 | 0.09284 | 0.10978 | 16.25133 | 0.044363165 | 0 | Neutral |
| testneg_356 | -0.05 | 0.81 | 0.558914 | -3 | 1.937622 | -3 | 0.90308 | 0.28373 | 0.24087 | 0.27985 | 5 | 0.121684783 | 0 | Neutral |
| testneg_357 | -0.07 | -0.01 | 1.263257 | -3 | 6.82383 | -3 | 0.11773 | 0.11669 | 0.22233 | 0.31758 | 45.43062 | 0.436339286 | 29.8 | Neutral |
| testneg_358 | -0.08 | -0.03 | 1.994638 | -3 | 1.420779 | -3 | 0.16468 | 0.11275 | 0.14386 | 0.12788 | 43.75 | 0.01 | 0 | Neutral |
| testneg_359 | -0.15 | -0.04 | 1.221649 | -3 | 0.196995 | -3 | 0.03676 | 0.12788 | 0.18438 | 0.13272 | 9.862 | 0.16 | 0 | Neutral |
| testneg_360 | 0 | 0 | 1.230202 | -3 | 0.442489 | -3 | 0.54826 | 0.24974 | 0.0478 | 0.12788 | 21.9774 | 0.03 | 0 | Neutral |
| testneg_361 | -0.14 | 1.85 | 2.577417 | 1.192851 | 6.877035 | -3 | 0.26181 | 0.12875 | 0.12131 | 0.12533 | 50.71736 | 0.4 | 16.2 | Neutral |
| testneg_362 | 0.22 | -2.66 | 6.776625 | -3 | 1.855162 | -3 | 0.17039 | 0.12788 | 0.33294 | 0.16155 | 16.60044 | 0.438392857 | 31.4 | Neutral |
| testneg_363 | -0.15 | -0.04 | 3.431501 | -3 | -1.171244 | -1.171244 | 0.1289 | 0.14878 | 0.19898 | 0.12788 | 25.94517 | 0.11 | 0 | Neutral |
| testneg_364 | 0.14 | -1.01 | 1.85098 | -1.813393 | 2.706663 | -3 | 0.06957 | 0.07707 | 0.09555 | 0.09639 | 27.63501 | 0.05 | 0 | Neutral |
| testneg_365 | 0 | 0 | 2.132856 | -3 | 0.0676 | -3 | 0.07963 | 0.23713 | 0.06731 | 0.2149 | 18.88889 | 0.162168367 | 0 | Neutral |

| | | | | | | | | | | | | | | |
|---|---|---|---|---|---|---|---|---|---|---|---|---|---|---|
| testneg_366 | 0.04 | -0.23 | 7.27275 | -3 | 7.789724 | -3 | 0.38445 | 0.58278 | 0.1254 | 0.08697 | 10 | 0.83 | 100 | Disease_causing |
| testneg_367 | -0.21 | -0.15 | -0.83994 | -3 | 0.037077 | -3 | 0.95319 | 0.20297 | 0.09004 | 0.07829 | 9.75999 | 0.075666667 | 0 | Neutral |
| testneg_368 | 0.15 | -0.28 | 5.363862 | -3 | 3.408834 | -3 | 0.04208 | 0.12788 | 0.19479 | 0.12788 | 18.6768 | 0.258484526 | 1 | Neutral |
| testneg_369 | -0.19 | -0.51 | 0.918079 | 0.918079 | 1.541854 | -3 | 0.83039 | 0.1727 | 0.1065 | 0.12788 | 8.97898 | 0.09 | 0 | Neutral |
| testneg_370 | -0.08 | -0.03 | 1.138753 | -3 | -2.452264 | -2.452264 | 0.94859 | 0.75045 | 0.16259 | 0.102 | 8.97898 | 0.04 | 0 | Neutral |
| testneg_371 | -0.14 | 0.37 | 0.940198 | -3 | -1.755229 | -3 | 0.09105 | 0.27434 | 0.05216 | 0.23851 | 68.01935 | 0.320805153 | 0.4 | Neutral |
| testneg_372 | 0 | 0 | 0.81674 | -3 | 1.240379 | -3 | 0.84238 | 0.12559 | 0.64591 | 0.38907 | 10.83333 | 0.03025641 | 0 | Neutral |
| testneg_373 | 0 | 0 | 2.104224 | -3 | -0.846787 | -3 | 0.37499 | 0.17803 | 0.03558 | 0.05202 | 20.51768 | 0.02 | 0 | Neutral |
| testneg_374 | 0 | 0 | 0.191318 | -3 | 1.781508 | -3 | 0.96113 | 0.24974 | 0.04699 | 0.08149 | 10.83333 | 0.02 | 0 | Neutral |
| testneg_375 | 0 | 0 | 0.946307 | -3 | 0.64288 | -3 | 0.18081 | 0.1258 | 0.57188 | 0.34038 | 10 | 0.020650794 | 0 | Neutral |
| testneg_376 | 0.14 | -0.69 | 5.93352 | -3 | 3.457136 | 3.457136 | 0.99689 | 0.28104 | 0.40707 | 0.37988 | 13.0557 | 0.871198916 | 100 | Disease_causing |
| testneg_377 | 0.03 | 0.12 | -1.487172 | -1.487172 | 0.852208 | -3 | 0.103 | 0.403 | 0.11237 | 0.08916 | 19.37853 | 0.07 | 0 | Neutral |
| testneg_378 | 0.14 | -0.37 | 3.308288 | -3 | 2.997859 | -2.209574 | 0.48585 | 0.66989 | 0.06397 | 0.12788 | 34.3967 | 0.254305556 | 0.8 | Neutral |
| testneg_379 | 0 | 0 | 0.560537 | -3 | 1.764776 | -3 | 0.32804 | 0.10857 | 0.6975 | 0.53794 | 10.41667 | 0.020525892 | 0 | Neutral |
| testneg_380 | 0.12 | -1.14 | -0.006272 | -0.006272 | -0.09216 | -3 | 0.07335 | 0.12788 | 0.39932 | 0.10693 | 13.40128 | 0.03 | 0 | Neutral |
| testneg_381 | 0 | 0 | -0.061652 | -3 | 0.034534 | 0.034534 | 0.83074 | 0.14769 | 0.62381 | 0.9477 | 13.51532 | 0.100465003 | 0 | Neutral |
| testneg_382 | 0.15 | -0.28 | -0.431211 | -3 | 0.552815 | 0.552815 | 0.36676 | 0.37749 | 0.20417 | 0.10681 | 10.83333 | 0.06 | 0 | Neutral |
| testneg_383 | 0 | -0.32 | 2.22695 | -3 | 0.328203 | -3 | 0.1273 | 0.12788 | 0.3414 | 0.13575 | 17.46678 | 0.0458171 | 0 | Neutral |
| testneg_384 | 0 | 0 | -0.009858 | -3 | 1.649381 | -3 | 0.31664 | 0.14181 | 0.22686 | 0.24974 | 11.25 | 0.040100317 | 0 | Neutral |
| testneg_385 | 0 | 0 | 0.291236 | -3 | 5.126283 | -3 | 0.32531 | 0.12351 | 0.68423 | 0.08163 | 13.51351 | 0.047525048 | 0 | Neutral |
| testneg_386 | 0 | 0 | 0.619175 | 0.619175 | 1.304689 | -3 | 0.68138 | 0.20944 | 0.06134 | 0.08207 | 24.36385 | 0.05 | 0 | Neutral |
| testneg_387 | 0.15 | -0.28 | 1.144976 | -3 | 0.433643 | -3 | 0.53592 | 0.11293 | 0.1601 | 0.12546 | 18.76882 | 0.02 | 0 | Neutral |
| testneg_388 | 0.07 | 0.7 | -2.193757 | -3 | 1.68744 | -3 | 0.18795 | 0.09487 | 0.67549 | 0.1273 | 17.42866 | 0.03022547 | 0 | Neutral |
| testneg_389 | 0.18 | 0.16 | 1.157565 | -3 | 3.369604 | -3 | 0.72124 | 0.09906 | 0.10249 | 0.09681 | 17.21847 | 0.12 | 0 | Neutral |
| testneg_390 | 0.02 | 0.13 | 4.492963 | -3 | 1.668921 | -1.582259 | 0.54648 | 0.19042 | 0.15994 | 0.11186 | 10.41667 | 0.09 | 0 | Neutral |
| testneg_391 | -0.03 | -0.12 | 6.051034 | -3 | 7.448756 | -3 | 0.91587 | 0.10818 | 0.16409 | 0.12788 | 13.01219 | 0.689891304 | 99.4 | Disease_causing |
| testneg_392 | 0 | 0 | 0.29846 | -3 | 0.443157 | -3 | 0.07543 | 0.10211 | 0.20011 | 0.27949 | 10.41667 | 0.03 | 0 | Neutral |
| testneg_393 | 0.00172727 | -0.1280909 | 1.232154 | -3 | 0.467995 | -3 | 0.17768 | 0.10832 | 0.11237 | 0.08916 | 27.27269 | 0.01 | 0 | Neutral |
| testneg_394 | -0.02 | -0.41 | -0.086855 | -3 | 1.024964 | -3 | 0.59827 | 0.11479 | 0.19898 | 0.08486 | 49.00822 | 0.02 | 0 | Neutral |
| testneg_395 | -0.07 | -0.38 | 0.816148 | -3 | 3.391965 | -3 | 0.26347 | 0.16383 | 0.19898 | 0.12788 | 14.03595 | 0.094560606 | 0 | Neutral |
| testneg_396 | 0.07 | 0.38 | 2.405373 | -3 | -0.044091 | -3 | 0.19898 | 0.13495 | 0.07534 | 0.09292 | 10 | 0.01225 | 0 | Neutral |
| testneg_397 | -0.01 | -0.79 | -0.439987 | -3 | -1.097876 | -3 | 0.12403 | 0.12788 | 0.12317 | 0.12345 | 29.16667 | 0.02 | 0 | Neutral |
| testneg_398 | 0.15 | 0.06 | 2.197563 | -3 | -0.160038 | -3 | 0.11114 | 0.09582 | 0.60079 | 0.11198 | 31.30092 | 0.02 | 0 | Neutral |
| testneg_399 | 0 | 0 | -0.264755 | -3 | 1.837239 | -3 | 0.09024 | 0.08559 | 0.1109 | 0.12788 | 8.97898 | 0.01 | 0 | Neutral |
| testneg_400 | -0.08 | -0.03 | 1.562114 | -3 | 2.035708 | -3 | 0.78356 | 0.13377 | 0.10278 | 0.08667 | 34.70057 | 0.04 | 0 | Neutral |
| testneg_401 | 0.05 | 0.29 | 3.783684 | -3 | 2.706663 | -3 | 0.39399 | 0.12788 | 0.09555 | 0.09639 | 23.10245 | 0.090769231 | 0 | Neutral |
| testneg_402 | 0.18 | -1.05 | 7.147075 | -3 | -1.111616 | -3 | 0.25989 | 0.11651 | 0.05444 | 0.12788 | 33.62656 | 0.22 | 0.4 | Neutral |
| testneg_403 | 0 | 0 | 0.865385 | -3 | 0.734765 | -3 | 0.3238 | 0.18333 | 0.07204 | 0.07708 | 16.42857 | 0.01 | 0 | Neutral |
| testneg_404 | 0 | 0 | 0.154631 | -3 | 1.672333 | 1.672333 | 0.25195 | 0.09584 | 0.15338 | 0.13614 | 8.50246 | 0.07 | 0 | Neutral |
| testneg_405 | 0 | 0 | 1.471577 | -3 | 2.90416 | -3 | 0.33923 | 0.12032 | 0.11618 | 0.20784 | 18.88889 | 0.1115625 | 0 | Neutral |
| testneg_406 | 0.23 | -1.04 | 4.434358 | -3 | -0.044659 | -3 | 0.10236 | 0.09622 | 0.53216 | 0.57092 | 18.77407 | 0.253333333 | 0.2 | Neutral |
| testneg_407 | -0.07 | -0.38 | 2.623579 | -3 | 6.165327 | -3 | 0.59812 | 0.19387 | 0.12112 | 0.11353 | 11.09451 | 0.299491071 | 4 | Neutral |
| testneg_408 | -0.05 | -0.29 | 0.116519 | 0.116519 | -0.756067 | -3 | 0.2207 | 0.09998 | 0.58124 | 0.31208 | 18.88889 | 0.09 | 0 | Neutral |
| testneg_409 | 0 | 0 | -0.364413 | -3 | 3.555867 | -3 | 0.16965 | 0.25991 | 0.71968 | 0.54266 | 12.72648 | 0.328190476 | 7.2 | Neutral |
| testneg_410 | 0 | 0 | 0.760575 | -3 | 1.068896 | -3 | 0.1878 | 0.08946 | 0.25448 | 0.15814 | 16.8018 | 0.03005102 | 0 | Neutral |
| testneg_411 | -0.21 | -0.15 | -0.976851 | -3 | 3.129189 | -3 | 0.27197 | 0.12346 | 0.51137 | 0.19235 | 10.83333 | 0.213501068 | 0 | Neutral |
| testneg_412 | 0.08 | 0.03 | 1.214028 | -3 | 1.26451 | -3 | 0.08136 | 0.10647 | 0.11439 | 0.21573 | 31.73046 | 0.083287582 | 0 | Neutral |
| testneg_413 | -0.04 | 0.23 | 0.421557 | 0.421557 | 2.91359 | -3 | 0.82507 | 0.19615 | 0.19898 | 0.12788 | 15.36471 | 0.09 | 0 | Neutral |
| testneg_414 | 0.06 | -0.57 | 0.237935 | -3 | 2.263733 | -3 | 0.16362 | 0.11879 | 0.16091 | 0.90971 | 22.80197 | 0.132040043 | 0 | Neutral |
| testneg_415 | 0.05 | -0.81 | 0.668367 | -3 | -0.669222 | -3 | 0.03656 | 0.07074 | 0.22225 | 0.11186 | 28.64035 | 0.03 | 0 | Neutral |
| testneg_416 | 0.03 | -0.85 | -1.390553 | -3 | 1.27985 | -3 | 0.08789 | 0.6361 | 0.09334 | 0.10027 | 10.22898 | 0.09 | 0 | Neutral |
| testneg_417 | -0.17 | 1.22 | 2.946635 | 0.502493 | 0.078385 | -3 | 0.19898 | 0.12788 | 0.52914 | 0.25278 | 8.45975 | 0.290823853 | 0.6 | Neutral |
| testneg_418 | -0.02 | 0.63 | 2.900715 | -3 | -2.137796 | -3 | 0.09211 | 0.08973 | 0.13698 | 0.12788 | 18.6768 | 0.05 | 0 | Neutral |
| testneg_419 | -0.03 | -0.12 | 1.731383 | -3 | 1.26254 | -3 | 0.1617 | 0.17928 | 0.35189 | 0.12788 | 16.84579 | 0.03 | 0 | Neutral |
| testneg_420 | 0 | 0 | 0.777519 | -3 | 1.624942 | 1.624942 | 0.75071 | 0.63704 | 0.16208 | 0.10597 | 11.875 | 0.04 | 0 | Neutral |
| testneg_421 | 0.03 | 1.22 | 7.358991 | -3 | 3.142585 | -3 | 0.03109 | 0.10705 | 0.14704 | 0.12788 | 13.37927 | 0.468188574 | 41 | Neutral |
| testneg_422 | 0 | 0 | 1.343568 | -3 | 5.779595 | -3 | 0.27187 | 0.41985 | 0.22046 | 0.12788 | 10.83333 | 0.169895833 | 0 | Neutral |
| testneg_423 | 0.07 | 0.01 | 1.526464 | -3 | 4.806063 | -3 | 0.04208 | 0.12788 | 0.22239 | 0.12885 | 10.41667 | 0.090564388 | 0 | Neutral |
| testneg_424 | 0.02 | 0.41 | -2.259592 | -3 | 6.062062 | -3 | 0.06311 | 0.14802 | 0.17458 | 0.12788 | 46.55069 | 0.266283432 | 0.2 | Neutral |
| testneg_425 | -0.09 | 0.3 | 0.912537 | -3 | 1.457871 | -3 | 0.2049 | 0.12678 | 0.08837 | 0.09327 | 17.21847 | 0.011458333 | 0 | Neutral |
| testneg_426 | 0.15 | 0.06 | 6.549059 | -3 | 2.193743 | 0.613871 | 0.2663 | 0.32806 | 0.19898 | 0.46867 | 6.98716 | 0.779671592 | 100 | Disease_causing |
| testneg_427 | 0.01666667 | -0.1875 | 0.933869 | -3 | 0.803533 | -3 | 0.9862 | 0.4528 | 0.15267 | 0.12788 | 16.05412 | 0.04 | 0 | Neutral |
| testneg_428 | 0 | 0 | 0.123542 | -3 | 0.271296 | -3 | 0.99915 | 0.96333 | 0.0978 | 0.26338 | 5 | 0.180915079 | 0 | Neutral |
| testneg_429 | 0.09 | -0.3 | 2.616434 | -3 | 2.808035 | -3 | 0.21998 | 0.20297 | 0.15055 | 0.22097 | 19.44444 | 0.428269551 | 30.4 | Neutral |
| testneg_430 | 0.07 | 0.01 | 0.631605 | -3 | -0.845539 | -3 | 0.14524 | 0.10162 | 0.16663 | 0.12788 | 20.83333 | 0.01 | 0 | Neutral |
| testneg_431 | -0.07 | -0.33 | 2.870116 | -3 | 0.615391 | -3 | 0.17325 | 0.17964 | 0.19898 | 0.19444 | 12.29167 | 0.214151786 | 0.2 | Neutral |
| testneg_432 | 0.15 | -0.28 | 0.118932 | 0.118932 | -0.630792 | -3 | 0.19567 | 0.09606 | 0.26181 | 0.12875 | 24.93861 | 0.02 | 0 | Neutral |
| testneg_433 | 0 | 0 | 1.531907 | -3 | 7.124928 | -3 | 0.19898 | 0.14509 | 0.15155 | 0.09923 | 7.98689 | 0.314188034 | 4.6 | Neutral |
| testneg_434 | -0.15 | 1.06 | 5.228347 | -3 | -0.707182 | -3 | 0.20491 | 0.22087 | 0.06789 | 0.09707 | 14.16667 | 0.229444444 | 0.2 | Neutral |
| testneg_435 | 0 | -0.01 | 1.961035 | -3 | 3.872922 | -3 | 0.55105 | 0.43208 | 0.1675 | 0.09413 | 12.32704 | 0.128184524 | 0 | Neutral |
| testneg_436 | 0.00172727 | -0.1280909 | 1.232154 | -3 | 1.071267 | -3 | 0.17768 | 0.10832 | 0.06526 | 0.12788 | 27.27269 | 0.01 | 0 | Neutral |
| testneg_437 | 0.17 | -1.22 | 4.037788 | -3 | 0.634119 | -3 | 0.08213 | 0.08913 | 0.41647 | 0.11239 | 19.09347 | 0.06 | 0 | Neutral |
| testneg_438 | -0.03 | -1.22 | 3.683701 | -3 | 6.431695 | -3 | 0.61008 | 0.17538 | 0.17764 | 0.09681 | 21.55172 | 0.39 | 17.8 | Neutral |
| testneg_439 | 0 | 0 | 1.240526 | -3 | -0.05742 | -3 | 0.68729 | 0.10641 | 0.07996 | 0.08354 | 24.24888 | 0.02 | 0 | Neutral |
| testneg_440 | -0.09 | 1.07 | 5.048579 | -3 | 3.447686 | -3 | 0.45927 | 0.74955 | 0.19898 | 0.1054 | 6.94444 | 0.448512294 | 37.8 | Neutral |
| testneg_441 | -0.05 | -0.29 | 1.344634 | -3 | 5.076843 | -3 | 0.33256 | 0.14909 | 0.45281 | 0.21621 | 5 | 0.445318483 | 34.6 | Neutral |
| testneg_442 | 0 | 0 | 1.145768 | -3 | 1.649124 | -3 | 0.22658 | 0.20646 | 0.37635 | 0.16791 | 10.3336 | 0.071665564 | 0 | Neutral |
| testneg_443 | 0.04 | -0.23 | 1.605283 | -3 | -0.481968 | -3 | 0.7257 | 0.64394 | 0.67331 | 0.10118 | 12.29167 | 0.05 | 0 | Neutral |
| testneg_444 | -0.08 | -0.03 | 2.540588 | -3 | -0.687752 | -3 | 0.32089 | 0.1035 | 0.40116 | 0.43982 | 12.18339 | 0.082470519 | 0 | Neutral |
| testneg_445 | -0.06 | 0.57 | 0.883365 | -3 | 0.885979 | -3 | 0.10004 | 0.12788 | 0.1653 | 0.0977 | 16.46847 | 0.02 | 0 | Neutral |
| testneg_446 | 0.14 | -1.01 | 3.46287 | -3 | 4.389466 | -3 | 0.49512 | 0.08696 | 0.12321 | 0.08084 | 10.29033 | 0.17 | 0 | Neutral |
| testneg_447 | 0 | 0 | 1.675065 | -3 | 0.6576 | -3 | 0.17285 | 0.09314 | 0.12147 | 0.1486 | 11.81921 | 0.030527778 | 0 | Neutral |
| testneg_448 | 0.01 | -1.3 | 6.159898 | -3 | -0.224622 | -3 | 0.1475 | 0.4323 | 0.12642 | 0.11218 | 18.88889 | 0.3 | 2.2 | Neutral |
| testneg_449 | 0.14 | -1.85 | 2.23079 | -3 | 3.994855 | -3 | 0.17806 | 0.12788 | 0.1311 | 0.12667 | 22.95764 | 0.117262694 | 0 | Neutral |
| testneg_450 | -0.05 | -0.29 | 1.421123 | -3 | 0.100154 | -3 | 0.9924 | 0.37896 | 0.21122 | 0.12686 | 10 | 0.033333333 | 0 | Neutral |
| testneg_451 | 0 | 0 | -0.029976 | -3 | 1.949203 | -3 | 0.11731 | 0.06348 | 0.61332 | 0.1004 | 19.29699 | 0.01 | 0 | Neutral |
| testneg_452 | 0 | 0 | -0.396403 | -3 | 1.212491 | -3 | 0.42585 | 0.18081 | 0.1509 | 0.27707 | 18.88889 | 0.091418626 | 0 | Neutral |
| testneg_453 | 0.02 | -0.36 | 0.993581 | -3 | 0.353411 | -3 | 0.04508 | 0.12788 | 0.80889 | 0.40016 | 18.88889 | 0.12 | 0 | Neutral |
| testneg_454 | -0.05 | -0.29 | 1.135406 | -3 | -0.39694 | -3 | 0.88414 | 0.13991 | 0.21807 | 0.12788 | 19.92611 | 0.011263298 | 0 | Neutral |
| testneg_455 | 0 | 0 | 1.736361 | -3 | 1.445664 | -3 | 0.36461 | 0.11015 | 0.14038 | 0.21177 | 12.09558 | 0.02082483 | 0 | Neutral |
| testneg_456 | 0 | 0 | 2.1641 | 2.1641 | 1.012349 | -3 | 0.84682 | 0.35105 | 0.27789 | 0.17365 | 21.53664 | 0.220166667 | 0.8 | Neutral |
| testneg_457 | 0.07 | 0.38 | 2.92632 | -3 | 1.139144 | -3 | 0.47442 | 0.11873 | 0.37711 | 0.12788 | 20.61588 | 0.03 | 0 | Neutral |
| testneg_458 | 0 | 0 | 0.839823 | -3 | 1.305896 | -3 | 0.26997 | 0.63944 | 0.1871 | 0.10907 | 19.44444 | 0.05 | 0 | Neutral |
| testneg_459 | 0 | 0 | 0.461263 | -3 | 0.857442 | -3 | 0.55193 | 0.22384 | 0.83041 | 0.14229 | 10.23791 | 0.041202552 | 0 | Neutral |
| testneg_460 | 0 | 0 | 0.173288 | -3 | -1.00023 | -3 | 0.61211 | 0.11059 | 0.43158 | 0.10472 | 12.35574 | 0.01 | 0 | Neutral |
| testneg_461 | 0.05 | -0.81 | 3.217486 | 3.217486 | 2.478986 | -3 | 0.74699 | 0.12286 | 0.13483 | 0.12788 | 20 | 0.396497696 | 20 | Neutral |
| testneg_462 | 0.04 | -0.23 | 2.30123 | -3 | 0.323865 | -3 | 0.23152 | 0.09962 | 0.63737 | 0.18081 | 14.11214 | 0.045142857 | 0 | Neutral |
| testneg_463 | -0.02 | 0.63 | 5.422854 | -3 | 2.363756 | -3 | 0.64152 | 0.15239 | 0.12121 | 0.4142 | 21.95402 | 0.607791667 | 88.2 | Disease_causing |
| testneg_464 | -0.07 | -0.38 | -1.097042 | -3 | 0.922605 | -3 | 0.10703 | 0.11611 | 0.2152 | 0.12788 | 33.60688 | 0.01 | 0 | Neutral |
| testneg_465 | 0 | 0 | 0.795894 | -3 | 4.640934 | -3 | 0.99448 | 0.67288 | 0.961 | 0.25991 | 10.41667 | 0.263629893 | 1.6 | Neutral |
| testneg_466 | 0.14 | -1.01 | 2.399127 | 2.100416 | 7.46225 | -3 | 0.22255 | 0.10057 | 0.27577 | 0.09812 | 19.49285 | 0.37 | 12 | Neutral |
| testneg_467 | 0 | 0 | -0.159191 | -3 | -0.490791 | -3 | 0.68029 | 0.12788 | 0.06953 | 0.17439 | 10.83333 | 0.068205046 | 0 | Neutral |
| testneg_468 | 0 | 0 | -0.090459 | -3 | 1.849176 | -3 | 0.15581 | 0.19351 | 0.56824 | 0.10646 | 9.66667 | 0.010477273 | 0 | Neutral |
| testneg_469 | -0.02 | 0.36 | 1.313372 | -3 | 2.622736 | -3 | 0.12836 | 0.09736 | 0.18404 | 0.14494 | 14.22149 | 0.1 | 0 | Neutral |
| testneg_470 | -0.02 | 0.68 | -0.892172 | -3 | 4.310552 | -3 | 0.21807 | 0.12788 | 0.13433 | 0.09578 | 20.68176 | 0.096 | 0 | Neutral |
| testneg_471 | 0.22 | -2.66 | 3.810217 | -3 | 4.004011 | -3 | 0.10562 | 0.09329 | 0.37736 | 0.11686 | 29.41758 | 0.19 | 0 | Neutral |
| testneg_472 | 0 | -0.01 | 0.229126 | -3 | 0.113253 | -3 | 0.25561 | 0.11004 | 0.38072 | 0.15993 | 27.4024 | 0.070060241 | 0 | Neutral |
| testneg_473 | -0.03 | -0.12 | 3.299396 | -3 | 1.414521 | -3 | 0.12547 | 0.09267 | 0.40081 | 0.15418 | 22.08561 | 0.159399995 | 0 | Neutral |
| testneg_474 | 0 | -0.01 | 0.646466 | -3 | 1.839296 | -3 | 0.14635 | 0.13588 | 0.19898 | 0.12788 | 30.55556 | 0.01 | 0 | Neutral |
| testneg_475 | 0.14 | 0.54 | 3.592951 | 3.592951 | 1.939281 | -3 | 0.18122 | 0.16895 | 0.07297 | 0.12788 | 20.01226 | 0.463365004 | 41.2 | Neutral |
| testneg_476 | 0.18 | -1.05 | 4.237406 | -3 | 1.500923 | -3 | 0.19898 | 0.12788 | 0.25473 | 0.07863 | 23.68839 | 0.05 | 0 | Neutral |
| testneg_477 | 0.06 | -0.57 | 3.838835 | -3 | 0.435681 | -3 | 0.70964 | 0.12663 | 0.0146 | 0.03045 | 10.7048 | 0.04 | 0 | Neutral |
| testneg_478 | 0.00916667 | -0.1625 | 0.801954 | -3 | 3.566467 | -3 | 0.09661 | 0.09981 | 0.17548 | 0.11384 | 13.51351 | 0.031776181 | 0 | Neutral |
| testneg_479 | 0 | 0 | 2.01155 | -3 | 1.280424 | -3 | 0.54648 | 0.19042 | 0.23427 | 0.10875 | 10.83333 | 0.011478792 | 0 | Neutral |
| testneg_480 | 0.03 | 0.12 | 3.782185 | -3 | 0.737804 | -3 | 0.83732 | 0.19325 | 0.39576 | 0.10761 | 27.4024 | 0.07 | 0 | Neutral |
| testneg_481 | 0 | -0.01 | 0.952222 | -3 | 0.97905 | -3 | 0.08373 | 0.09191 | 0.49278 | 0.09047 | 18.33333 | 0.01 | 0 | Neutral |
| testneg_482 | 0 | 0 | -0.136916 | -3 | 3.19575 | -3 | 0.04853 | 0.4186 | 0.98853 | 0.88685 | 15 | 0.389583333 | 14.8 | Neutral |
| testneg_483 | 0.16 | 0.08 | -0.169987 | -3 | 3.66939 | -3 | 0.99445 | 0.10396 | 0.66422 | 0.99377 | 10.41667 | 0.298153941 | 0.4 | Neutral |
| testneg_484 | -0.04 | 1.61 | 5.680307 | -3 | -0.467119 | -3 | 0.02617 | 0.12788 | 0.55441 | 0.08704 | 52.37015 | 0.21 | 0 | Neutral |
| testneg_485 | 0 | 0 | 1.817109 | -3 | -0.942639 | -3 | 0.51622 | 0.11809 | 0.05996 | 0.12788 | 64.94565 | 0.09 | 0 | Neutral |
| testneg_486 | 0.01 | 0.79 | 4.3808 | 2.330651 | 1.729145 | -3 | 0.36858 | 0.09681 | 0.13575 | 0.12788 | 13.19331 | 0.22 | 1.8 | Neutral |
| testneg_487 | 0 | 0 | 0.184064 | -3 | 1.821777 | -3 | 0.04436 | 0.12788 | 0.11187 | 0.12788 | 11.00193 | 0.03 | 0 | Neutral |

| | | | | | | | | | | | | | | |
|---|---|---|---|---|---|---|---|---|---|---|---|---|---|---|
| testneg_488 | 0.14 | -1.01 | 0.599729 | -3 | 5.616992 | -3 | 0.38762 | 0.12788 | 0.13413 | 0.12788 | 34.50745 | 0.12 | 0 | Neutral |
| testneg_489 | 0.04 | -1.36 | 2.710461 | -3 | 0.734765 | -3 | 0.15202 | 0.12722 | 0.07204 | 0.07708 | 43.81612 | 0.03 | 0 | Neutral |
| testneg_490 | 0.02 | -0.63 | -0.055258 | -3 | 5.133476 | 5.133476 | 0.16139 | 0.09116 | 0.20417 | 0.10681 | 22.85088 | 0.33 | 2.4 | Neutral |
| testneg_491 | 0.14 | -0.37 | 4.085318 | -3 | 0.473367 | 0.052969 | 0.3828 | 0.5569 | 0.62381 | 0.12788 | 9.44444 | 0.146742424 | 0 | Neutral |
| testneg_492 | -0.04 | 0.23 | -0.274997 | -3 | 2.079972 | -3 | 0.37501 | 0.0997 | 0.25549 | 0.12378 | 23.33333 | 0.02 | 0 | Neutral |
| testneg_493 | 0 | -0.01 | 1.723095 | 1.723095 | 1.441246 | -3 | 0.91293 | 0.12788 | 0.33292 | 0.17111 | 4.22222 | 0.101341502 | 0 | Neutral |
| testneg_494 | 0.14 | -1.85 | -1.690089 | -3 | 1.463492 | -3 | 0.20918 | 0.10825 | 0.32606 | 0.10251 | 12.80035 | 0.041224336 | 0 | Neutral |
| testneg_495 | 0 | 0 | -0.17286 | -3 | 6.218251 | -3 | 0.95682 | 0.22744 | 0.204 | 0.15424 | 9.82124 | 0.315260784 | 6.8 | Neutral |
| testneg_496 | 0 | 0 | 1.941287 | -3 | 4.623974 | -3 | 0.24977 | 0.11262 | 0.09866 | 0.104 | 23.78498 | 0.061295019 | 0 | Neutral |
| testneg_497 | 0 | 0 | 1.313175 | -3 | 1.091905 | -3 | 0.03953 | 0.05763 | 0.10058 | 0.13882 | 37.25578 | 0.104369766 | 0 | Neutral |
| testneg_498 | 0.06 | 0.18 | 4.598509 | -3 | 4.387792 | -3 | 0.11351 | 0.10667 | 0.21118 | 0.10074 | 19.09347 | 0.181582341 | 0 | Neutral |
| testneg_499 | -0.1 | -1.9 | 2.083029 | 0.83037 | -0.592626 | -3 | 0.16929 | 0.12788 | 0.143 | 0.11236 | 17.97143 | 0.06 | 0 | Neutral |
| testneg_500 | 0.15 | -0.28 | 1.374995 | -3 | 1.902977 | 1.902977 | 0.26727 | 0.12788 | 0.18504 | 0.0822 | 12.32809 | 0.04 | 0 | Neutral |
| testneg_501 | 0 | 0 | 0.484541 | -3 | 2.53183 | -3 | 0.48023 | 0.12788 | 0.38207 | 0.35052 | 12.2619 | 0.083160034 | 0 | Neutral |
| testneg_502 | 0 | 0 | 0.418262 | -3 | -0.125432 | -3 | 0.69007 | 0.09893 | 0.30926 | 0.11955 | 21.38889 | 0.010104784 | 0 | Neutral |
| testneg_503 | 0 | 0 | 0.693679 | -3 | -1.462645 | -3 | 0.13331 | 0.13638 | 0.19898 | 0.09069 | 18.88889 | 0.01 | 0 | Neutral |
| testneg_504 | -0.02 | -0.45 | 4.751068 | -3 | 0.415846 | -3 | 0.9449 | 0.9378 | 0.2755 | 0.12788 | 16.8018 | 0.177328431 | 0 | Neutral |
| testneg_505 | -0.05 | -0.29 | 0.720622 | -3 | 6.087637 | -3 | 0.98321 | 0.27035 | 0.098 | 0.08791 | 9.31758 | 0.17975 | 0 | Neutral |
| testneg_506 | -0.02 | -0.41 | -1.58691 | -3 | -0.687336 | -3 | 0.10617 | 0.15182 | 0.09997 | 0.09515 | 18.88889 | 0.05 | 0 | Neutral |
| testneg_507 | -0.02 | -0.41 | 3.378414 | -3 | 0.050569 | -3 | 0.39724 | 0.11058 | 0.16263 | 0.53493 | 8.97898 | 0.14086724 | 0 | Neutral |
| testneg_508 | 0.03 | 0.12 | 0.332787 | -3 | -2.855052 | -3 | 0.08011 | 0.12788 | 0.16545 | 0.10002 | 56.1036 | 0.05 | 0 | Neutral |
| testneg_509 | 0 | 0 | 0.442945 | -3 | 0.775551 | -3 | 0.19499 | 0.12111 | 0.0903 | 0.09372 | 18.66667 | 0 | 0 | Neutral |
| testneg_510 | -0.15 | -0.06 | 5.168376 | -3 | 2.135113 | -3 | 0.30275 | 0.12788 | 0.40391 | 0.13431 | 14.92465 | 0.296055139 | 1.8 | Neutral |
| testneg_511 | 0.04 | -0.23 | 4.454617 | -3 | 0.465329 | 0.465329 | 0.7639 | 0.10408 | 0.11131 | 0.68435 | 13.04617 | 0.219207996 | 0 | Neutral |
| testneg_512 | 0 | 0 | 0.182861 | -3 | 4.25337 | -3 | 0.25474 | 0.11262 | 0.05027 | 0.10981 | 28.12163 | 0.050516106 | 0 | Neutral |
| testneg_513 | 0 | 0 | 0.440209 | -3 | 1.375928 | -3 | 0.13083 | 0.23231 | 0.19898 | 0.10996 | 16.8018 | 0.011742424 | 0 | Neutral |
| testneg_514 | 0.02 | -0.63 | -2.196989 | -3 | 2.022849 | 2.022849 | 0.02759 | 0.12458 | 0.20708 | 0.12788 | 27.4024 | 0.08 | 0 | Neutral |
| testneg_515 | 0.14 | -0.37 | 5.904299 | -3 | -1.662468 | -3 | 0.16716 | 0.10951 | 0.16692 | 0.09243 | 34.44813 | 0.100851393 | 0 | Neutral |
| testneg_516 | 0 | 0 | -0.509361 | -3 | 0.65387 | -3 | 0.71814 | 0.72461 | 0.03505 | 0.0907 | 10.22898 | 0.03 | 0 | Neutral |
| testneg_517 | 0.02 | -0.63 | 2.302571 | -3 | 7.812451 | 7.812451 | 0.09179 | 0.12788 | 0.7219 | 0.41509 | 24.22512 | 0.637757143 | 95.2 | Disease_causing |
| testneg_518 | 0.14 | -0.37 | 2.120517 | -3 | -2.285677 | -3 | 0.14457 | 0.11614 | 0.08911 | 0.09298 | 12.76316 | 0.01 | 0 | Neutral |
| testneg_519 | 0 | 0 | 1.474347 | -3 | 1.710229 | -3 | 0.88414 | 0.13991 | 0.13767 | 0.11503 | 10.41667 | 0.01 | 0 | Neutral |
| testneg_520 | 0 | 0 | 0.140815 | -3 | 1.567225 | -3 | 0.51278 | 0.35859 | 0.32406 | 0.10985 | 12.70833 | 0.02 | 0 | Neutral |
| testneg_521 | 0 | 0 | 0.996313 | -3 | 1.366812 | -3 | 0.25744 | 0.2162 | 0.19898 | 0.12788 | 17.77778 | 0.010501508 | 0 | Neutral |
| testneg_522 | -0.03 | -1.22 | 3.205092 | -3 | 6.639474 | -3 | 0.18559 | 0.09173 | 0.13173 | 0.12788 | 20.74074 | 0.280064103 | 4 | Neutral |
| testneg_523 | 0.01 | -0.57 | -1.439791 | -3 | -2.012697 | -3 | 0.37711 | 0.12788 | 0.08027 | 0.12788 | 17.21847 | 0.0105 | 0 | Neutral |
| testneg_524 | 0.05 | 0.29 | 2.54094 | -3 | 7.496918 | -3 | 0.32484 | 0.11621 | 0.30521 | 0.36067 | 21.72903 | 0.518482926 | 62.4 | Disease_causing |
| testneg_525 | 0 | -0.01 | 0.855893 | -3 | 0.200692 | -3 | 0.26916 | 0.13285 | 0.09445 | 0.12788 | 23.7963 | 0.02 | 0 | Neutral |
| testneg_526 | 0 | 0 | 0.461408 | -3 | -0.071864 | -3 | 0.16797 | 0.11809 | 0.97005 | 0.31407 | 10.43432 | 0.020331442 | 0 | Neutral |
| testneg_527 | 0.14 | -0.37 | 4.741413 | -3 | 4.343075 | -3 | 0.21115 | 0.17723 | 0.57991 | 0.2033 | 23.65565 | 0.799016175 | 100 | Disease_causing |
| testneg_528 | -0.05 | -0.29 | 0.06448 | -3 | 0.509453 | -3 | 0.30684 | 0.12788 | 0.11587 | 0.12001 | 20.65799 | 0.00125 | 0 | Neutral |
| testneg_529 | -0.21 | -0.15 | -0.83994 | -3 | -0.586534 | -0.586534 | 0.95319 | 0.20297 | 0.09439 | 0.1884 | 20.14775 | 0.19 | 0 | Neutral |
| testneg_530 | 0 | 0 | 0.214112 | -3 | 6.158659 | -3 | 0.17301 | 0.14089 | 0.22258 | 0.10471 | 18.33333 | 0.110443806 | 0 | Neutral |
| testneg_531 | 0 | 0 | -0.027704 | -3 | 0.052371 | 0.052371 | 0.45299 | 0.16306 | 0.82107 | 0.27926 | 11.85281 | 0.08 | 0 | Neutral |
| testneg_532 | -0.01 | -0.8 | -0.697584 | -3 | -1.062235 | -3 | 0.19898 | 0.12788 | 0.29172 | 0.08143 | 23.68839 | 0.011625 | 0 | Neutral |
| testneg_533 | 0 | 0 | 0.731896 | -3 | 7.50603 | -3 | 0.10236 | 0.09622 | 0.16311 | 0.09662 | 39.47801 | 0.26 | 1.2 | Neutral |
| testneg_534 | 0.18 | -1.05 | 3.274763 | -3 | 5.342136 | -3 | 0.49512 | 0.08696 | 0.1496 | 0.16692 | 10.29033 | 0.549058146 | 70.2 | Disease_causing |
| testneg_535 | 0 | -0.01 | 1.723095 | 1.723095 | 0.00589 | -3 | 0.91293 | 0.12788 | 0.38503 | 0.11286 | 6.75553 | 0.030250623 | 0 | Neutral |
| testneg_536 | 0.03 | 0.12 | -1.186234 | -3 | 0.877291 | -3 | 0.35843 | 0.15758 | 0.86166 | 0.14254 | 10.22898 | 0.054207461 | 0 | Neutral |
| testneg_537 | -0.08 | -0.03 | -0.534744 | -3 | 0.591702 | -3 | 0.07923 | 0.08035 | 0.43283 | 0.23984 | 20.59286 | 0.080050505 | 0 | Neutral |
| testneg_538 | -0.05 | -0.29 | 4.62142 | -3 | 4.558299 | -3 | 0.09269 | 0.12788 | 0.20357 | 0.12788 | 16.42857 | 0.251073589 | 1.4 | Neutral |
| testneg_539 | -0.1 | -1.9 | 3.072235 | -3 | 2.561326 | -3 | 0.04866 | 0.05972 | 0.20765 | 0.27124 | 11.14078 | 0.378750527 | 9.2 | Neutral |
| testneg_540 | 0.04 | -0.23 | 3.025327 | -3 | -1.596835 | -3 | 0.94807 | 0.14977 | 0.17443 | 0.19181 | 5.73716 | 0.219833333 | 0 | Neutral |
| testneg_541 | 0 | 0 | 1.181359 | -3 | 6.076778 | -3 | 0.46401 | 0.24608 | 0.33818 | 0.12788 | 11.90227 | 0.163229575 | 0.2 | Neutral |
| testneg_542 | 0.21 | 0.15 | -0.608116 | -3 | 7.39468 | -3 | 0.0925 | 0.08222 | 0.1417 | 0.1398 | 17.77778 | 0.381362553 | 12.8 | Neutral |
| testneg_543 | 0 | 0 | 1.539579 | -3 | 0.86879 | -3 | 0.18459 | 0.33541 | 0.50303 | 0.23508 | 11.25 | 0.092444939 | 0 | Neutral |
| testneg_544 | 0.02218182 | 0.39763636 | 0.700961 | -3 | 5.842275 | -3 | 0.1296 | 0.164 | 0.80889 | 0.40016 | 18.88889 | 0.549996503 | 73.4 | Disease_causing |
| testneg_545 | 0 | 0 | 2.28293 | -3 | 0.152832 | 0.152832 | 0.08332 | 0.09046 | 0.04829 | 0.09848 | 10.24108 | 0.02 | 0 | Neutral |
| testneg_546 | 0 | -0.73 | 2.726423 | -3 | 1.07196 | -3 | 0.14273 | 0.1162 | 0.19898 | 0.12788 | 17.57576 | 0.0118267 | 0 | Neutral |
| testneg_547 | 0 | 0 | -0.031197 | -3 | 7.890352 | -3 | 0.1017 | 0.11809 | 0.23879 | 0.12788 | 22.05174 | 0.2825 | 2.4 | Neutral |
| testneg_548 | -0.17 | -0.38 | 3.968918 | -3 | -1.359951 | -3 | 0.19898 | 0.12788 | 0.47816 | 0.66526 | 27.51545 | 0.23 | 0 | Neutral |
| testneg_549 | 0 | 0 | 0.383044 | -3 | -0.908797 | -3 | 0.63819 | 0.32258 | 0.12282 | 0.20617 | 12.79101 | 0.085469031 | 0 | Neutral |
| testneg_550 | 0.00055944 | 0.00363636 | 1.360315 | -3 | 1.211987 | -3 | 0.29482 | 0.15726 | 0.18373 | 0.12788 | 30.55556 | 0.012106482 | 0 | Neutral |
| testneg_551 | -0.0042857 | -0.2491429 | 9.525107 | -3 | -0.273874 | -3 | 0.04032 | 0.12788 | 0.0501 | 0.12788 | 31.53736 | 0.33 | 7.8 | Neutral |
| testneg_552 | 0 | 0 | 0.922275 | -3 | 3.383357 | -3 | 0.26586 | 0.10825 | 0.90813 | 0.10329 | 5.73099 | 0.027479642 | 0 | Neutral |
| testneg_553 | 0.02 | 0.45 | 2.331456 | -3 | 1.902366 | -3 | 0.43762 | 0.12788 | 0.12231 | 0.23155 | 12.37434 | 0.079797684 | 0 | Neutral |
| testneg_554 | 0 | 0 | 1.782835 | -3 | 3.916231 | -3 | 0.96704 | 0.16878 | 0.26812 | 0.30371 | 9.67871 | 0.303753325 | 4 | Neutral |
| testneg_555 | 0.03 | -0.85 | 5.714497 | -3 | 2.304438 | -3 | 0.098 | 0.08791 | 0.05767 | 0.07325 | 18.6768 | 0.146071429 | 0 | Neutral |
| testneg_556 | 0.09 | -0.67 | 4.020687 | -3 | -0.656574 | -3 | 0.34001 | 0.12788 | 0.09531 | 0.09236 | 13.41166 | 0.03 | 0 | Neutral |
| testneg_557 | 0 | 0 | 0.755726 | -3 | 0.65387 | -3 | 0.08454 | 0.22908 | 0.03505 | 0.0907 | 16.8018 | 0.020615106 | 0 | Neutral |
| testneg_558 | -0.08 | -0.03 | 2.428041 | -3 | -0.351099 | -3 | 0.07606 | 0.13082 | 0.19898 | 0.12788 | 12.46106 | 0.02 | 0 | Neutral |
| testneg_559 | 0 | 0 | 1.178544 | -3 | 1.27985 | -3 | 0.19898 | 0.12788 | 0.09334 | 0.10027 | 44.32245 | 0.001428571 | 0 | Neutral |
| testneg_560 | 0 | 0 | 3.977929 | -3 | 0.527496 | -3 | 0.06048 | 0.12788 | 0.06723 | 0.10296 | 31.32208 | 0.05 | 0 | Neutral |
| testneg_561 | 0 | 0 | 0.764567 | -3 | -2.844353 | -3 | 0.06027 | 0.11574 | 0.07872 | 0.12788 | 10.22898 | 0.02 | 0 | Neutral |
| testneg_562 | 0.07 | -0.31 | 1.708283 | 1.708283 | 1.042979 | -3 | 0.19898 | 0.12788 | 0.84682 | 0.35105 | 20.62504 | 0.17 | 0 | Neutral |
| testneg_563 | 0.04 | -0.23 | 7.597341 | -3 | 0.362935 | -3 | 0.136 | 0.17348 | 0.12488 | 0.12788 | 44.58452 | 0.35 | 10.4 | Neutral |
| testneg_564 | -0.02 | 0.68 | 0.349669 | -3 | 3.858939 | 1.41921 | 0.19898 | 0.12788 | 0.51852 | 0.11202 | 5.97008 | 0.107480392 | 0 | Neutral |
| testneg_565 | 0.21 | 0.15 | 1.631249 | -3 | 2.022849 | -3 | 0.47166 | 0.13548 | 0.20708 | 0.12788 | 21.94444 | 0.05 | 0 | Neutral |
| testneg_566 | 0 | -0.32 | 1.05419 | -3 | 1.688334 | -3 | 0.45152 | 0.34457 | 0.40283 | 0.34309 | 12.29167 | 0.158203222 | 0 | Neutral |
| testneg_567 | 0 | -0.01 | 1.723095 | 1.723095 | -0.032364 | -3 | 0.91293 | 0.12788 | 0.16524 | 0.12788 | 4.30556 | 0.030816667 | 0 | Neutral |
| testneg_568 | 0.22 | -2.66 | 7.366549 | -3 | 1.192651 | -3 | 0.143 | 0.11236 | 0.65952 | 0.49708 | 14.95396 | 0.4 | 18 | Neutral |
| testneg_569 | 0 | -0.01 | 0.952222 | -3 | 3.953111 | 3.373436 | 0.08373 | 0.09191 | 0.20657 | 0.12788 | 17.77778 | 0.175555556 | 0 | Neutral |
| testneg_570 | 0 | 0 | 1.550971 | -3 | 0.772688 | -3 | 0.73311 | 0.10866 | 0.40021 | 0.21702 | 10.83333 | 0.016439245 | 0 | Neutral |
| testneg_571 | 0.04 | -0.23 | 2.511043 | -3 | 2.078125 | -3 | 0.07204 | 0.07708 | 0.84365 | 0.12788 | 15.5381 | 0.02 | 0 | Neutral |
| testneg_572 | 0.00055944 | 0.00363636 | 1.360315 | -3 | 3.078755 | 1.112437 | 0.29482 | 0.15726 | 0.62172 | 0.18135 | 11.25 | 0.195055547 | 0 | Neutral |
| testneg_573 | -0.21 | -0.15 | 0.913651 | -3 | 1.351912 | -3 | 0.10285 | 0.11721 | 0.18584 | 0.11114 | 19.66924 | 0.06 | 0 | Neutral |
| testneg_574 | -0.06 | 0.57 | 0.172291 | -0.066175 | 0.571926 | -3 | 0.0138 | 0.06769 | 0.15406 | 0.1454 | 19.44444 | 0.120320513 | 0 | Neutral |
| testneg_575 | -0.03 | -0.1 | 2.058438 | -3 | -1.294754 | -3 | 0.0749 | 0.12788 | 0.13369 | 0.09791 | 17.88352 | 0.010087538 | 0 | Neutral |
| testneg_576 | 0.02 | -0.68 | 6.639247 | -3 | 2.083519 | -3 | 0.27407 | 0.1136 | 0.14375 | 0.12788 | 18.33333 | 0.288511722 | 3.8 | Neutral |
| testneg_577 | 0 | -0.01 | 0.846806 | -3 | 1.262077 | -3 | 0.20344 | 0.1705 | 0.11351 | 0.10667 | 12.27567 | 0.01 | 0 | Neutral |
| testneg_578 | 0.14 | -0.37 | 1.599249 | -3 | 4.764927 | -3 | 0.08714 | 0.08945 | 0.07631 | 0.12788 | 20.83333 | 0.1 | 0 | Neutral |
| testneg_579 | 0.05 | -0.81 | 5.789217 | -3 | 6.043672 | -3 | 0.26668 | 0.11598 | 0.3025 | 0.2499 | 11.90227 | 0.837795603 | 100 | Disease_causing |
| testneg_580 | 0 | 0 | 0.722971 | -3 | 2.656717 | -3 | 0.19898 | 0.12788 | 0.79857 | 0.12788 | 22.40741 | 0.02 | 0 | Neutral |
| testneg_581 | -0.03 | -0.1 | 1.347962 | -3 | 0.390651 | -3 | 0.0418 | 0.12788 | 0.07275 | 0.12788 | 23.54743 | 0.05 | 0 | Neutral |
| testneg_582 | 0.22 | -2.66 | 7.430811 | -3 | 1.51996 | -3 | 0.11359 | 0.08631 | 0.09105 | 0.27434 | 61.07469 | 0.39 | 14.4 | Neutral |
| testneg_583 | 0.01 | 0.8 | 5.875511 | -3 | 7.667083 | -3 | 0.19898 | 0.12788 | 0.13369 | 0.09791 | 24.14517 | 0.65 | 95.6 | Disease_causing |
| testneg_584 | -0.18 | -0.16 | 0.910161 | -3 | -1.58983 | -3 | 0.93468 | 0.12788 | 0.21677 | 0.1681 | 18.88889 | 0.12 | 0 | Neutral |
| testneg_585 | 0 | 0 | 0.175869 | -3 | 1.492764 | -3 | 0.43897 | 0.12441 | 0.10822 | 0.10942 | 32.48276 | 0.010755656 | 0 | Neutral |
| testneg_586 | -0.0081119 | -0.0026573 | 1.451979 | -3 | 1.074985 | -3 | 0.26727 | 0.12788 | 0.74558 | 0.10643 | 12.47414 | 0.01 | 0 | Neutral |
| testneg_587 | 0 | 0 | 1.640374 | -3 | 1.445664 | -3 | 0.53203 | 0.12788 | 0.14038 | 0.21177 | 20.83333 | 0.07 | 0 | Neutral |
| testneg_588 | 0 | 0 | 1.233889 | -3 | 2.053049 | -3 | 0.10228 | 0.22182 | 0.26586 | 0.10825 | 22.34729 | 0.03 | 0 | Neutral |
| testneg_589 | 0.19 | -0.27 | 1.873078 | -3 | 5.294804 | -3 | 0.0626 | 0.5373 | 0.12446 | 0.08908 | 30.93485 | 0.314318327 | 1.8 | Neutral |
| testneg_590 | 0.01 | 0.79 | 1.807734 | -3 | 1.342384 | -3 | 0.14273 | 0.1162 | 0.36813 | 0.09414 | 10.41667 | 0.010184246 | 0 | Neutral |
| testneg_591 | 0.1 | 1.9 | -0.979272 | -3 | 0.788897 | -3 | 0.1589 | 0.17874 | 0.80849 | 0.21975 | 10.41667 | 0.2 | 0 | Neutral |
| testneg_592 | 0.14 | -1.01 | 6.678677 | -3 | -2.112814 | -3 | 0.86373 | 0.11969 | 0.26982 | 0.12788 | 19.09347 | 0.23 | 1.2 | Neutral |
| testneg_593 | 0 | 0 | -0.259811 | -3 | 5.385549 | -3 | 0.70964 | 0.12663 | 0.13126 | 0.11628 | 18.33333 | 0.07 | 0 | Neutral |
| testneg_594 | 0 | 0 | 0.432421 | -3 | -0.049164 | -0.049164 | 0.0922 | 0.07873 | 0.12322 | 0.1246 | 35.88644 | 0.010628019 | 0 | Neutral |
| testneg_595 | 0.07 | 0.38 | 2.405373 | -3 | 0.101783 | -3 | 0.19898 | 0.13495 | 0.06196 | 0.10881 | 18.77199 | 0.01076141 | 0 | Neutral |
| testneg_596 | 0 | 0 | 0.103447 | 0.103447 | 3.086284 | -3 | 0.15813 | 0.61197 | 0.13099 | 0.17453 | 26.50014 | 0.360204957 | 9.4 | Neutral |
| testneg_597 | 0.02 | -0.68 | 7.177031 | -3 | 3.423277 | -3 | 0.55043 | 0.12112 | 0.13322 | 0.11297 | 34.72222 | 0.453611111 | 39 | Neutral |
| testneg_598 | 0.22 | -2.66 | 3.552335 | -3 | 0.726148 | -3 | 0.99338 | 0.14229 | 0.10782 | 0.14795 | 10.83333 | 0.239219697 | 0 | Neutral |
| testneg_599 | -0.06 | 0.57 | 0.172291 | -3 | 3.267694 | -3 | 0.0138 | 0.06769 | 0.21576 | 0.17241 | 18.33333 | 0.232460317 | 0.2 | Neutral |
| testneg_600 | -0.03 | -0.12 | 0.194478 | -3 | -1.455534 | -3 | 0.11006 | 0.14352 | 0.10314 | 0.08894 | 30.53632 | 0.030590278 | 0 | Neutral |
| testneg_601 | 0.03 | 0.1 | 4.582242 | -3 | 5.391094 | -3 | 0.06247 | 0.14184 | 0.37424 | 0.12788 | 10 | 0.414569589 | 22.8 | Neutral |
| testneg_602 | 0.01127273 | -0.5378182 | 13.369443 | -3 | 0.960962 | -3 | 0.10719 | 0.08618 | 0.14525 | 0.14744 | 18.73163 | 0.38 | 10.6 | Neutral |
| testneg_603 | -0.1 | -1.9 | 0.96317 | -3 | 5.791651 | -3 | 0.20379 | 0.35591 | 0.22415 | 0.70109 | 13.18841 | 0.663287082 | 95.6 | Disease_causing |
| testneg_604 | -0.21 | -0.15 | 0.197741 | -3 | -0.068708 | -3 | 0.31923 | 0.17492 | 0.25077 | 0.06261 | 12.60353 | 0.07 | 0 | Neutral |
| testneg_605 | 0 | -0.32 | 1.964014 | -3 | 2.67715 | -3 | 0.98442 | 0.44966 | 0.07999 | 0.10413 | 11.83382 | 0.079583333 | 0 | Neutral |
| testneg_606 | -0.02 | -0.41 | 6.179947 | -3 | 1.156713 | 1.156713 | 0.14021 | 0.43531 | 0.36458 | 0.16397 | 18.33333 | 0.696666667 | 99 | Disease_causing |
| testneg_607 | 0 | 0 | -0.181123 | -3 | 0.781229 | 0.781229 | 0.12867 | 0.11159 | 0.07321 | 0.12788 | 37.88019 | 0.02 | 0 | Neutral |
| testneg_608 | 0 | 0 | 0.39745 | 0.39745 | 2.374684 | -3 | 0.11404 | 0.12788 | 0.33314 | 0.47999 | 9.94383 | 0.123909524 | 0 | Neutral |
| testneg_609 | 0.14 | -0.69 | 2.480925 | -3 | 6.481793 | -3 | 0.17478 | 0.1501 | 0.1417 | 0.1398 | 10 | 0.594133333 | 87.2 | Disease_causing |

| | | | | | | | | | | | | | | |
|---|---|---|---|---|---|---|---|---|---|---|---|---|---|---|
| testneg_610 | 0.22 | -2.66 | 5.527307 | -3 | -1.599196 | -3 | 0.1589 | 0.17874 | 0.37711 | 0.12788 | 19.09347 | 0.23 | 0 | Neutral |
| testneg_611 | -0.21 | -0.15 | 5.077905 | -3 | 5.453129 | -3 | 0.18022 | 0.12788 | 0.1134 | 0.12788 | 30.48139 | 0.370669118 | 12.6 | Neutral |
| testneg_612 | 0.14 | -1.01 | 0.599729 | -3 | 7.146764 | -3 | 0.38762 | 0.12788 | 0.18147 | 0.12788 | 31.84073 | 0.25 | 1.4 | Neutral |
| testneg_613 | -0.07 | -0.33 | -2.373259 | -3 | 1.845442 | -3 | 0.04525 | 0.12788 | 0.99293 | 0.79067 | 10.83333 | 0.18 | 0 | Neutral |
| testneg_614 | -0.01 | -0.8 | 4.589251 | -3 | 3.124356 | -3 | 0.34694 | 0.20397 | 0.20557 | 0.10131 | 15.85471 | 0.251473793 | 1.2 | Neutral |
| testneg_615 | 0 | 0 | -0.211085 | -3 | 2.022849 | 2.022849 | 0.04543 | 0.28103 | 0.20708 | 0.12788 | 18.88889 | 0.11 | 0 | Neutral |
| testneg_616 | 0 | 0 | 0.419995 | -3 | 0.661515 | -3 | 0.7174 | 0.15815 | 0.07534 | 0.09292 | 10.41667 | 0.010508621 | 0 | Neutral |
| testneg_617 | -0.07 | -0.38 | 5.775288 | -3 | 2.598788 | -3 | 0.16663 | 0.12788 | 0.19898 | 0.10307 | 34.09722 | 0.210916667 | 0.2 | Neutral |
| testneg_618 | 0.14 | -0.37 | 4.001731 | -3 | 3.9441 | -3 | 0.12912 | 0.09693 | 0.35779 | 0.12788 | 13.51351 | 0.185093213 | 0 | Neutral |
| testneg_619 | -0.21 | -0.15 | -1.299109 | -3 | -0.659954 | -3 | 0.28038 | 0.11814 | 0.19898 | 0.12788 | 23.54743 | 0.05 | 0 | Neutral |
| testneg_620 | -0.16 | -0.08 | 2.94383 | -3 | 1.136968 | -3 | 0.51707 | 0.45685 | 0.31744 | 0.10986 | 13.97697 | 0.147682441 | 0 | Neutral |
| testneg_621 | 0.2 | -1.26 | -0.469793 | -3 | 5.734507 | -3 | 0.0811 | 0.09449 | 0.06731 | 0.2149 | 19.44444 | 0.360981703 | 9 | Neutral |
| testneg_622 | 0 | 0 | 0.111854 | -3 | 0.127388 | -3 | 0.14462 | 0.11239 | 0.21967 | 0.09467 | 18.78886 | 0 | 0 | Neutral |
| testneg_623 | 0.18 | -1.05 | 6.296843 | -3 | 0.877291 | -3 | 0.20327 | 0.11174 | 0.86166 | 0.14254 | 13.51351 | 0.21 | 0.4 | Neutral |
| testneg_624 | 0.02 | -0.63 | 0.544663 | -3 | -0.867421 | -3 | 0.09767 | 0.39937 | 0.23385 | 0.12788 | 10.83333 | 0.06 | 0 | Neutral |
| testneg_625 | 0 | 0 | 1.720142 | -3 | 0.50158 | -3 | 0.64701 | 0.94571 | 0.12885 | 0.61089 | 10.41667 | 0.182738095 | 0 | Neutral |
| testneg_626 | 0.19 | -0.25 | -0.332338 | -3 | 2.397456 | -3 | 0.13708 | 0.81545 | 0.17443 | 0.19181 | 24.41367 | 0.370229908 | 6.6 | Neutral |
| testneg_627 | 0 | 0 | -0.233364 | -3 | -1.245289 | -3 | 0.26323 | 0.2399 | 0.11043 | 0.12788 | 17.77778 | 0.020548048 | 0 | Neutral |
| testneg_628 | 0.03 | -1.03 | 7.541227 | -3 | -1.280796 | -3 | 0.13482 | 0.12788 | 0.06743 | 0.20381 | 16.8018 | 0.448507937 | 33 | Neutral |
| testneg_629 | 0.04 | -0.23 | 3.89714 | -3 | 4.19568 | -3 | 0.33022 | 0.13818 | 0.35513 | 0.21966 | 12.74587 | 0.646646096 | 92.2 | Disease_causing |
| testneg_630 | 0 | 0 | 1.436812 | -3 | 1.287073 | -3 | 0.29302 | 0.17394 | 0.06744 | 0.12788 | 19.44716 | 0.010769231 | 0 | Neutral |
| testneg_631 | 0.22 | -2.66 | 4.55974 | -3 | -0.513734 | -3 | 0.16078 | 0.39498 | 0.256 | 0.20036 | 6.01729 | 0.377694444 | 11.8 | Neutral |
| testneg_632 | 0.02 | 0.41 | 0.799457 | -3 | 6.598988 | -3 | 0.08714 | 0.08945 | 0.1454 | 0.10285 | 19.09347 | 0.18 | 0 | Neutral |
| testneg_633 | -0.03 | 1.03 | 0.958989 | -3 | 2.355419 | -3 | 0.39436 | 0.16446 | 0.82595 | 0.52905 | 9.09528 | 0.195463823 | 0 | Neutral |
| testneg_634 | 0.03 | 1.22 | 5.227225 | -3 | 2.350147 | -3 | 0.04866 | 0.05972 | 0.21983 | 0.10146 | 26.34995 | 0.18 | 0 | Neutral |
| testneg_635 | 0 | 0 | 0.013815 | -3 | 0.177501 | -3 | 0.47888 | 0.43756 | 0.0925 | 0.08222 | 18.77551 | 0.03 | 0 | Neutral |
| testneg_636 | 0 | 0 | -0.104866 | -3 | 5.632523 | -3 | 0.2698 | 0.18815 | 0.17764 | 0.09681 | 18.42945 | 0.148266667 | 0 | Neutral |
| testneg_637 | 0 | -0.01 | 1.284608 | -3 | 1.087392 | -3 | 0.57975 | 0.1248 | 0.20502 | 0.12788 | 26.9055 | 0.01 | 0 | Neutral |
| testneg_638 | 0.14 | -0.37 | -0.136533 | -3 | 2.23696 | 2.23696 | 0.05932 | 0.09551 | 0.68865 | 0.19414 | 19.06433 | 0.19 | 0 | Neutral |
| testneg_639 | 0 | -0.01 | 2.663664 | -3 | 3.257494 | -3 | 0.19898 | 0.26604 | 0.09356 | 0.2448 | 18.33333 | 0.430668325 | 30.6 | Neutral |
| testneg_640 | -0.08 | -0.03 | 5.820832 | -3 | 4.946403 | -3 | 0.1004 | 0.11409 | 0.27175 | 0.14094 | 16.50674 | 0.615570652 | 88.8 | Disease_causing |
| testneg_641 | 0.14 | -1.01 | 5.146768 | -3 | 1.80646 | -3 | 0.10379 | 0.34799 | 0.32111 | 0.12788 | 18.04862 | 0.24025 | 0 | Neutral |
| testneg_642 | 0 | 0 | -0.22828 | -3 | 2.02257 | -3 | 0.31694 | 0.13475 | 0.46946 | 0.08281 | 13.3358 | 0.01 | 0 | Neutral |
| testneg_643 | 0.14 | -0.37 | 3.758883 | -3 | 1.26259 | 1.26259 | 0.50461 | 0.11034 | 0.44722 | 0.11641 | 10.15609 | 0.053333333 | 0 | Neutral |
| testneg_644 | 0.05 | 0.29 | 0.113571 | -3 | 0.588209 | -3 | 0.02844 | 0.12788 | 0.07431 | 0.11513 | 13.51351 | 0.03 | 0 | Neutral |
| testneg_645 | -0.14 | 0.69 | 0.233598 | 0.233598 | 0.231661 | -3 | 0.25804 | 0.11876 | 0.25637 | 0.37408 | 18.23037 | 0.120403069 | 0 | Neutral |
| testneg_646 | 0.09 | -0.3 | 5.573517 | -3 | -1.949243 | -3 | 0.09416 | 0.17821 | 0.08499 | 0.08678 | 19.44444 | 0.21 | 0 | Neutral |
| testneg_647 | 0.14 | -0.37 | 0.864504 | -3 | 3.619507 | 3.619507 | 0.67216 | 0.17808 | 0.67676 | 0.12788 | 19.23304 | 0.3 | 1.2 | Neutral |
| testneg_648 | -0.02 | 0.36 | 3.195501 | -3 | 0.278169 | -3 | 0.22899 | 0.31462 | 0.10963 | 0.12788 | 18.6768 | 0.080627451 | 0 | Neutral |
| testneg_649 | 0 | 0 | 1.229821 | -3 | 2.688597 | 2.688597 | 0.46091 | 0.13852 | 0.19898 | 0.12788 | 25.73468 | 0.08925 | 0 | Neutral |
| testneg_650 | 0.06 | -0.57 | 3.156075 | -3 | 0.870324 | -3 | 0.44164 | 0.13588 | 0.16355 | 0.12788 | 27.61996 | 0.040641026 | 0 | Neutral |
| testneg_651 | 0 | 0 | 0.089405 | -3 | 0.651966 | -3 | 0.06301 | 0.32138 | 0.28326 | 0.13944 | 11.25 | 0.129588745 | 0 | Neutral |
| testneg_652 | -0.05 | -0.29 | 3.494755 | -3 | 4.752866 | -3 | 0.21201 | 0.20962 | 0.75728 | 0.34838 | 11.96702 | 0.773932745 | 99.2 | Disease_causing |
| testneg_653 | 0.00233333 | -0.0513333 | 3.565578 | -3 | 1.392438 | -3 | 0.99689 | 0.28104 | 0.21118 | 0.10074 | 9.74183 | 0.110589681 | 0 | Neutral |
| testneg_654 | -0.07 | -0.38 | 2.184344 | -3 | 2.475186 | -3 | 0.05472 | 0.12788 | 0.9581 | 0.12788 | 28.90523 | 0.090339862 | 0 | Neutral |
| testneg_655 | -0.09 | 0.3 | 0.782249 | -3 | 0.483862 | -3 | 0.5127 | 0.12263 | 0.21551 | 0.12788 | 12.94465 | 0.02 | 0 | Neutral |
| testneg_656 | 0.08 | 0.03 | -0.431527 | -3 | 0.615335 | -3 | 0.19772 | 0.09261 | 0.70872 | 0.09884 | 19.375 | 0.02 | 0 | Neutral |
| testneg_657 | 0.21 | 0.15 | 2.022528 | -3 | 1.623152 | 1.623152 | 0.57953 | 0.15513 | 0.19898 | 0.12788 | 30.92949 | 0.07 | 0 | Neutral |
| testneg_658 | 0.06054545 | -0.3370909 | 8.1928 | -3 | -0.51508 | -3 | 0.12073 | 0.10969 | 0.06667 | 0.08666 | 5.26625 | 0.304423077 | 3.2 | Neutral |
| testneg_659 | 0.07 | -0.31 | 1.843162 | -3 | 0.840434 | -3 | 0.30869 | 0.11948 | 0.48936 | 0.10963 | 13.99351 | 0 | 0 | Neutral |
| testneg_660 | -0.0190909 | 0.62818182 | 7.974744 | -3 | 1.926664 | -3 | 0.087 | 0.12788 | 0.32334 | 0.10525 | 25.12484 | 0.4 | 19.4 | Neutral |
| testneg_661 | -0.22 | 2.66 | 0.427319 | -3 | 5.58705 | -3 | 0.11096 | 0.10074 | 0.0538 | 0.0823 | 50.69444 | 0.271142857 | 0 | Neutral |
| testneg_662 | 0 | 0 | 0.365268 | -3 | -0.057557 | -3 | 0.53627 | 0.12788 | 0.06729 | 0.17051 | 19.44444 | 0.095807605 | 0 | Neutral |
| testneg_663 | -0.05 | -0.29 | 1.012343 | -3 | 4.54534 | -3 | 0.93035 | 0.31722 | 0.06713 | 0.11112 | 24.8779 | 0.132403846 | 0 | Neutral |
| testneg_664 | 0.15 | 0.06 | 1.47438 | -3 | 3.020834 | -3 | 0.61343 | 0.11838 | 0.19894 | 0.27411 | 13.08751 | 0.223665715 | 0 | Neutral |
| testneg_665 | 0.14 | -1.01 | -0.270969 | -3 | 0.696041 | -3 | 0.04441 | 0.13603 | 0.78515 | 0.35093 | 11.25 | 0.14 | 0 | Neutral |
| testneg_666 | 0 | 0 | -0.961702 | -0.961702 | 7.536912 | -3 | 0.46979 | 0.1705 | 0.10042 | 0.09296 | 18.55633 | 0.41 | 21.6 | Neutral |
| testneg_667 | -0.12 | 1.14 | 2.652229 | -3 | -1.390634 | -3 | 0.04393 | 0.12788 | 0.29672 | 0.10485 | 13.2927 | 0.11 | 0 | Neutral |
| testneg_668 | 0.14 | -0.69 | 7.000202 | -3 | 2.097152 | -3 | 0.19898 | 0.12788 | 0.17199 | 0.10586 | 11.22671 | 0.358545455 | 12.2 | Neutral |
| testneg_669 | -0.03 | 1.03 | 2.078465 | -3 | 0.124841 | -3 | 0.99665 | 0.12884 | 0.1834 | 0.12788 | 29.92514 | 0.07 | 0 | Neutral |
| testneg_670 | -0.15 | -0.04 | 0.939929 | -3 | 1.516949 | -3 | 0.18701 | 0.12788 | 0.87202 | 0.42207 | 17.21847 | 0.091683007 | 0 | Neutral |
| testneg_671 | 0.23 | -1.86 | 4.232372 | -3 | 1.87267 | -3 | 0.87268 | 0.21837 | 0.17478 | 0.1501 | 10.83333 | 0.324726307 | 4.6 | Neutral |
| testneg_672 | 0 | 0 | 4.240578 | -3 | -0.061002 | -0.061002 | 0.17903 | 0.37232 | 0.35561 | 0.08845 | 23.38719 | 0.100978823 | 0 | Neutral |
| testneg_673 | -0.04 | -2.01 | 3.61412 | -3 | 0.626243 | 0.626243 | 0.18256 | 0.13499 | 0.34412 | 0.08672 | 14.16667 | 0.072103604 | 0 | Neutral |
| testneg_674 | 0 | 0 | -0.961702 | -0.961702 | -0.390908 | -3 | 0.46979 | 0.1705 | 0.07652 | 0.30451 | 10.41667 | 0.085767249 | 0 | Neutral |
| testneg_675 | 0 | 0 | -0.535508 | -0.785566 | 9.272843 | -3 | 0.01196 | 0.12788 | 0.24544 | 0.0894 | 22.06714 | 0.323333333 | 3.4 | Neutral |
| testneg_676 | 0 | 0 | 0.026725 | -3 | 2.950441 | -3 | 0.55527 | 0.12788 | 0.05245 | 0.13458 | 17.66021 | 0.177178473 | 0 | Neutral |
| testneg_677 | 0.09 | -0.3 | 1.673428 | 1.673428 | 0.955677 | 0.955677 | 0.32352 | 0.10243 | 0.10034 | 0.10906 | 18.27928 | 0.05 | 0 | Neutral |
| testneg_678 | -0.03 | -0.12 | 1.324962 | -3 | 0.05475 | -3 | 0.38184 | 0.10478 | 0.13468 | 0.09235 | 40.33827 | 0.01 | 0 | Neutral |
| testneg_679 | 0.18 | -1.05 | 5.614334 | -3 | 1.760309 | -3 | 0.75033 | 0.64204 | 0.11618 | 0.20784 | 10 | 0.587032609 | 81.8 | Disease_causing |
| testneg_680 | 0.14 | -1.01 | 6.277915 | -3 | 0.211586 | 0.211586 | 0.12036 | 0.0873 | 0.1119 | 0.24875 | 19.24159 | 0.339518141 | 8.2 | Neutral |
| testneg_681 | -0.02 | 0.36 | 3.195501 | -3 | -0.35214 | -3 | 0.22899 | 0.31462 | 0.11134 | 0.13818 | 20.18018 | 0.232678571 | 0 | Neutral |
| testneg_682 | 0.08 | 0.03 | -0.129461 | -1.481985 | -3.9524 | -3 | 0.18416 | 0.13292 | 0.50276 | 0.09638 | 13.65398 | 0.03 | 0 | Neutral |
| testneg_683 | 0.02 | 0.45 | 0.215947 | -3 | 0.229134 | -3 | 0.0418 | 0.12788 | 0.09515 | 0.07784 | 13.51351 | 0.031416667 | 0 | Neutral |
| testneg_684 | -0.14 | 0.69 | 5.249654 | -3 | 2.22565 | -3 | 0.1247 | 0.10331 | 0.25159 | 0.12788 | 28.57769 | 0.17 | 0 | Neutral |
| testneg_685 | -0.07 | -0.38 | 3.896611 | -3 | 0.383334 | -3 | 0.57349 | 0.09764 | 0.04935 | 0.06599 | 10 | 0.05 | 0 | Neutral |
| testneg_686 | 0 | 0 | 1.511019 | 0.030464 | 0.131914 | -3 | 0.33795 | 0.17455 | 0.23879 | 0.12788 | 19.17258 | 0.01 | 0 | Neutral |
| testneg_687 | -0.04 | 0.23 | 1.397648 | -3 | 4.093803 | -3 | 0.11006 | 0.20781 | 0.35234 | 0.1303 | 12.4985 | 0.316463585 | 4.8 | Neutral |
| testneg_688 | 0.03 | -1.03 | 5.503167 | -3 | 2.12021 | 2.12021 | 0.13961 | 0.09565 | 0.07698 | 0.12788 | 11.74429 | 0.216333333 | 0.2 | Neutral |
| testneg_689 | 0 | -0.01 | 1.594546 | -3 | 1.744838 | -3 | 0.33304 | 0.15588 | 0.9902 | 0.19761 | 19.96169 | 0.071229382 | 0 | Neutral |
| testneg_690 | 0 | 0 | 1.482956 | -3 | -0.384446 | -3 | 0.2852 | 0.12788 | 0.36829 | 0.11457 | 10.83333 | 9.17E-05 | 0 | Neutral |
| testneg_691 | -0.16 | -0.08 | 0.679359 | -3 | 4.55192 | -3 | 0.16544 | 0.1313 | 0.13458 | 0.17634 | 11.25 | 0.456838859 | 36.8 | Neutral |
| testneg_692 | 0 | -0.01 | 0.342228 | -3 | 2.503067 | -3 | 0.07684 | 0.09138 | 0.56025 | 0.12788 | 16.8018 | 0.02 | 0 | Neutral |
| testneg_693 | -0.14 | 1.01 | 1.408725 | -3 | 3.249291 | -3 | 0.40283 | 0.34309 | 0.25758 | 0.2262 | 14.73008 | 0.434546685 | 27 | Neutral |
| testneg_694 | 0.09 | -0.3 | 5.693708 | -3 | 2.013485 | -3 | 0.28271 | 0.60265 | 0.87803 | 0.12788 | 10.41667 | 0.3807168 | 11.2 | Neutral |
| testneg_695 | 0 | 0 | -0.203227 | -3 | -0.644713 | -3 | 0.58827 | 0.57942 | 0.14511 | 0.12788 | 5.69444 | 0.02 | 0 | Neutral |
| testneg_696 | 0.14 | -1.01 | 5.779693 | -3 | 1.630994 | -3 | 0.92514 | 0.19444 | 0.30429 | 0.29938 | 4.44444 | 0.622799784 | 89.2 | Disease_causing |
| testneg_697 | 0.14 | -0.69 | 4.284627 | -3 | 3.161443 | -3 | 0.9327 | 0.12788 | 0.26961 | 0.09979 | 11.97842 | 0.179730827 | 0 | Neutral |
| testneg_698 | -0.07 | -0.7 | 0.044039 | -3 | -0.808449 | -0.808449 | 0.36486 | 0.14268 | 0.13258 | 0.9258 | 10.22898 | 0.120175456 | 0 | Neutral |
| testneg_699 | -0.17 | 1.22 | 6.396974 | -3 | 2.051799 | -3 | 0.41743 | 0.78545 | 0.14624 | 0.10369 | 5 | 0.515 | 61.8 | Disease_causing |
| testneg_700 | 0 | -0.01 | 0.684698 | -3 | 1.216081 | -3 | 0.09661 | 0.09981 | 0.08973 | 0.07662 | 13.08587 | 0.000431494 | 0 | Neutral |
| testneg_701 | 0.04 | -0.89 | 1.703642 | -3 | -0.885623 | -3 | 0.25764 | 0.53109 | 0.51037 | 0.10999 | 18.88889 | 0.06 | 0 | Neutral |
| testneg_702 | 0 | 0 | 0.496899 | -3 | 4.875419 | -3 | 0.17325 | 0.17964 | 0.08393 | 0.1123 | 16.54706 | 0.122664116 | 0 | Neutral |
| testneg_703 | 0 | 0 | 0.467241 | -3 | -0.354855 | -3 | 0.14948 | 0.12788 | 0.14524 | 0.10162 | 20.83333 | 0.000229915 | 0 | Neutral |
| testneg_704 | -0.03 | 1.03 | 1.699863 | -3 | 3.138075 | -3 | 0.06048 | 0.12788 | 0.82156 | 0.14621 | 20.18737 | 0.2 | 0 | Neutral |
| testneg_705 | 0 | -0.01 | 1.129155 | -3 | 5.892768 | 3.098857 | 0.16835 | 0.09282 | 0.19898 | 0.12788 | 18.6768 | 0.18 | 0.2 | Neutral |
| testneg_706 | 0 | 0 | 0.990948 | -3 | 5.477048 | 2.079972 | 0.31859 | 0.11262 | 0.25549 | 0.12378 | 18.6768 | 0.12 | 0 | Neutral |
| testneg_707 | 0.14 | -1.01 | 5.471755 | -3 | 1.414521 | 0.04439 | 0.1784 | 0.12788 | 0.40081 | 0.15418 | 11.875 | 0.364826856 | 12.4 | Neutral |
| testneg_708 | -0.03 | -0.1 | 1.276737 | -3 | 1.340083 | -3 | 0.19564 | 0.59275 | 0.28227 | 0.10211 | 10.83333 | 0.06020999 | 0 | Neutral |
| testneg_709 | 0.03 | 0.12 | 4.329007 | -3 | 0.896883 | -3 | 0.95682 | 0.22744 | 0.02124 | 0.05186 | 13.23122 | 0.070487377 | 0 | Neutral |
| testneg_710 | 0 | 0 | -0.10912 | -3 | 4.170859 | -3 | 0.85984 | 0.40044 | 0.09998 | 0.09022 | 9.57237 | 0.086875 | 0 | Neutral |
| testneg_711 | 0.07 | 0.38 | 5.554338 | -3 | 0.019644 | -3 | 0.49745 | 0.1259 | 0.42899 | 0.31045 | 21.75706 | 0.36 | 8 | Neutral |
| testneg_712 | 0.03 | -1.03 | 3.488962 | -3 | 2.410093 | -0.045995 | 0.59245 | 0.12788 | 0.1743 | 0.12788 | 17.78631 | 0.091143697 | 0 | Neutral |
| testneg_713 | -0.02 | 0.63 | 3.574998 | -3 | 2.439723 | -3 | 0.32974 | 0.12245 | 0.16431 | 0.09173 | 13.04617 | 0.091094624 | 0 | Neutral |
| testneg_714 | 0 | 0 | 1.663039 | -3 | -0.360768 | -0.360768 | 0.11395 | 0.26806 | 0.47147 | 0.09503 | 12.89568 | 0.03 | 0 | Neutral |
| testneg_715 | 0.07 | 0.38 | -1.485747 | -3 | 0.230409 | 0.230409 | 0.13029 | 0.12788 | 0.16116 | 0.08314 | 11.92061 | 0.02 | 0 | Neutral |
| testneg_716 | 0 | 0 | -0.187157 | -3 | 7.530767 | -3 | 0.09871 | 0.09286 | 0.1897 | 0.12459 | 13.14653 | 0.26 | 1.8 | Neutral |
| testneg_717 | 0 | -0.01 | 1.594546 | -3 | 1.063136 | -3 | 0.33304 | 0.15588 | 0.29268 | 0.16012 | 18.33333 | 0.069866473 | 0 | Neutral |
| testneg_718 | 0 | 0.32 | 2.212176 | -3 | 0.955677 | 0.955677 | 0.63233 | 0.16248 | 0.10034 | 0.10906 | 12.06621 | 0.030182059 | 0 | Neutral |
| testneg_719 | -0.03 | 1.03 | 1.03165 | -3 | 3.224653 | -3 | 0.17873 | 0.12378 | 0.4129 | 0.11172 | 43.87043 | 0.080278201 | 0 | Neutral |
| testneg_720 | 0.05 | -0.81 | 1.982936 | -3 | 2.541931 | -3 | 0.07044 | 0.11679 | 0.07558 | 0.12788 | 18.88889 | 0.03 | 0 | Neutral |
| testneg_721 | 0.04 | -0.23 | 6.602112 | -3 | 0.744952 | -3 | 0.32897 | 0.10658 | 0.38279 | 0.10578 | 20.8805 | 0.194166667 | 0.4 | Neutral |
| testneg_722 | 0.21 | 0.15 | 3.309253 | 2.147578 | -1.085568 | -3 | 0.47335 | 0.12788 | 0.22901 | 0.12788 | 11.92982 | 0.17 | 0.4 | Neutral |
| testneg_723 | 0.07 | -0.31 | 2.407105 | -3 | 1.757173 | -3 | 0.21095 | 0.10574 | 0.21726 | 0.23214 | 13.75 | 0.063445693 | 0 | Neutral |
| testneg_724 | 0 | 0 | 0.759983 | -3 | 1.762821 | 1.762821 | 0.13505 | 0.09019 | 0.1375 | 0.112 | 13.02635 | 0.02 | 0 | Neutral |
| testneg_725 | 0 | -0.01 | 0.639661 | -3 | 0.327712 | -3 | 0.16263 | 0.53493 | 0.57517 | 0.10873 | 13.51351 | 0.03 | 0 | Neutral |
| testneg_726 | -0.17 | -0.38 | 3.567601 | -3 | 0.350371 | -3 | 0.04655 | 0.10225 | 0.05221 | 0.12788 | 16.668 | 0.12 | 0 | Neutral |
| testneg_727 | 0 | 0 | 0.012846 | -3 | 6.303575 | -3 | 0.07995 | 0.12788 | 0.11237 | 0.08916 | 23.15811 | 0.128166667 | 0 | Neutral |
| testneg_728 | -0.06 | -0.18 | -1.695325 | -3 | 4.797748 | -3 | 0.36052 | 0.2858 | 0.98535 | 0.15071 | 10.83333 | 0.428252628 | 26.6 | Neutral |
| testneg_729 | 0 | 0 | -0.164302 | -3 | 7.237047 | -3 | 0.35475 | 0.12788 | 0.16485 | 0.13045 | 32.36768 | 0.252203422 | 1.8 | Neutral |
| testneg_730 | 0.05 | 0.29 | 1.340083 | -3 | 3.369604 | -3 | 0.28227 | 0.10211 | 0.10249 | 0.09681 | 10.83333 | 0.062478355 | 0 | Neutral |
| testneg_731 | -0.04 | 0.23 | 4.755795 | -3 | 2.849519 | -3 | 0.06877 | 0.17777 | 0.14486 | 0.33725 | 22.7381 | 0.609251962 | 90.6 | Disease_causing |

| | | | | | | | | | | | | | | |
|---|---|---|---|---|---|---|---|---|---|---|---|---|---|---|
| testneg_732 | 0.15 | 0.06 | 6.577474 | -3 | -0.110087 | -3 | 0.08646 | 0.24855 | 0.82126 | 0.17937 | 16.23032 | 0.46 | 37 | Neutral |
| testneg_733 | -0.15 | 0.28 | 0.27384 | -3 | 4.773876 | -3 | 0.17173 | 0.15363 | 0.25453 | 0.1177 | 11.03941 | 0.273346417 | 0.8 | Neutral |
| testneg_734 | 0 | 0 | 0.510124 | -3 | 3.561616 | -3 | 0.10161 | 0.13484 | 0.10919 | 0.105 | 10.41667 | 0.058197115 | 0 | Neutral |
| testneg_735 | 0.21 | 0.15 | 4.039152 | -3 | -0.011228 | -3 | 0.21623 | 0.42074 | 0.31339 | 0.10191 | 30.23693 | 0.12775 | 0 | Neutral |
| testneg_736 | -0.09 | 0.3 | 1.799707 | -3 | 1.302273 | -3 | 0.98442 | 0.44966 | 0.1947 | 0.11413 | 12.63272 | 0.050405844 | 0 | Neutral |
| testneg_737 | 0.18 | -1.05 | 7.936771 | -3 | 0.936874 | -3 | 0.32022 | 0.40347 | 0.16061 | 0.12426 | 13.86082 | 0.458844812 | 34.2 | Neutral |
| testneg_738 | 0.21 | 0.15 | -0.131021 | -3 | -0.064527 | -3 | 0.35325 | 0.12614 | 0.12642 | 0.11218 | 10.83333 | 0.03625 | 0 | Neutral |
| testneg_739 | 0.01 | -0.57 | 0.723491 | -3 | 3.916127 | -3 | 0.55315 | 0.09195 | 0.08876 | 0.09037 | 17.21847 | 0.090522714 | 0 | Neutral |
| testneg_740 | 0 | 0 | 0.415464 | -3 | 0.774689 | 0.774689 | 0.18671 | 0.12788 | 0.24219 | 0.17458 | 16.02705 | 0.040513158 | 0 | Neutral |
| testneg_741 | 0 | -0.32 | 3.107095 | -3 | 3.422566 | -3 | 0.19898 | 0.40608 | 0.16462 | 0.18467 | 10.41667 | 0.670778549 | 94.8 | Disease_causing |
| testneg_742 | 0.09 | -2.17 | 3.628039 | 3.628039 | 2.137668 | -3 | 0.055 | 0.20858 | 0.34184 | 0.1964 | 5 | 0.857729751 | 100 | Disease_causing |
| testneg_743 | 0.09 | -0.3 | 3.117713 | -3 | 4.560265 | -3 | 0.58827 | 0.57942 | 0.43459 | 0.12595 | 10 | 0.400408732 | 22.6 | Neutral |
| testneg_744 | 0 | 0 | 1.107432 | -3 | 2.908555 | -3 | 0.5726 | 0.1165 | 0.14175 | 0.12788 | 10.83333 | 0.03 | 0 | Neutral |
| testneg_745 | 0.14 | -0.37 | -0.136533 | -3 | 1.970058 | -3 | 0.05932 | 0.09551 | 0.06067 | 0.07796 | 29.539 | 0.04 | 0 | Neutral |
| testneg_746 | -0.15 | -0.04 | 0.720517 | -3 | 4.789497 | -3 | 0.1109 | 0.17295 | 0.16646 | 0.16629 | 44.59115 | 0.625543521 | 92.4 | Disease_causing |
| testneg_747 | 0.04 | -0.23 | 1.605283 | -3 | -1.729368 | -3 | 0.7257 | 0.64394 | 0.34295 | 0.12788 | 10.41667 | 0.04 | 0 | Neutral |
| testneg_748 | 0 | 0 | 0.393827 | -3 | -1.645867 | -3 | 0.07516 | 0.15472 | 0.11608 | 0.09123 | 12.29167 | 0.02 | 0 | Neutral |
| testneg_749 | 0.09 | -0.3 | 2.911406 | -3 | 4.051473 | -3 | 0.25421 | 0.16235 | 0.50461 | 0.11034 | 31.16398 | 0.240813725 | 0.2 | Neutral |
| testneg_750 | 0 | -0.01 | 1.149331 | -3 | 4.280436 | -3 | 0.15582 | 0.14216 | 0.40116 | 0.43982 | 10.83333 | 0.292299332 | 3.6 | Neutral |
| testneg_751 | 0.08 | 0.03 | -1.360462 | -3 | 1.837985 | -3 | 0.28537 | 0.12788 | 0.36541 | 0.27749 | 18.33333 | 0.08 | 0 | Neutral |
| testneg_752 | 0.08 | 0.03 | -1.571869 | -3 | 4.894593 | -3 | 0.54489 | 0.09921 | 0.13743 | 0.12788 | 14.07895 | 0.112131579 | 0 | Neutral |
| testneg_753 | -0.0005303 | -0.0306061 | 0.082842 | -3 | -0.436881 | -3 | 0.00971 | 0.0791 | 0.27004 | 0.17052 | 11.875 | 0.090140196 | 0 | Neutral |
| testneg_754 | 0 | 0 | 0.802919 | 0.802919 | 3.022813 | -3 | 0.12042 | 0.11067 | 0.17301 | 0.10048 | 20.27778 | 0.06 | 0 | Neutral |
| testneg_755 | 0 | 0 | 1.504538 | -3 | 3.886171 | -3 | 0.19898 | 0.12788 | 0.19898 | 0.12788 | 29.34704 | 0.040570669 | 0 | Neutral |
| testneg_756 | 0.04 | 2.01 | 2.884683 | -3 | -0.323214 | -3 | 0.11957 | 0.08975 | 0.15432 | 0.095 | 18.11231 | 0.08 | 0 | Neutral |
| testneg_757 | -0.0174545 | 0.25218182 | 7.014767 | -3 | 4.674123 | -3 | 0.01369 | 0.12788 | 0.95992 | 0.10958 | 30.03248 | 0.42 | 23.6 | Neutral |
| testneg_758 | 0.03 | 0.1 | 6.634872 | -3 | 0.34432 | -3 | 0.20112 | 0.17491 | 0.52957 | 0.18687 | 10.83333 | 0.513561366 | 60.6 | Disease_causing |
| testneg_759 | 0 | 0 | 1.66597 | -3 | 3.700286 | -3 | 0.66027 | 0.23984 | 0.11687 | 0.09069 | 24.2785 | 0.11 | 0 | Neutral |
| testneg_760 | 0 | 0 | 1.076963 | -3 | 0.810517 | -3 | 0.20011 | 0.27949 | 0.08729 | 0.10515 | 10 | 0.01 | 0 | Neutral |
| testneg_761 | 0.16 | 0.08 | -0.169987 | -3 | 4.325287 | -3 | 0.99445 | 0.10396 | 0.16973 | 0.17555 | 11.09572 | 0.28327381 | 0.4 | Neutral |
| testneg_762 | -0.02 | 0.68 | 0.385771 | -3 | 1.314868 | -3 | 0.11077 | 0.10076 | 0.18744 | 0.09392 | 5.73099 | 0.02 | 0 | Neutral |
| testneg_763 | -0.08 | -0.03 | -0.514498 | -3 | 0.323125 | -3 | 0.81305 | 0.17045 | 0.07258 | 0.06248 | 22.95764 | 0.04 | 0 | Neutral |
| testneg_764 | 0.11 | -0.19 | 2.886405 | -3 | -0.215823 | -3 | 0.62966 | 0.14896 | 0.1556 | 0.11007 | 16.03108 | 0.05 | 0 | Neutral |
| testneg_765 | 0.06 | -0.57 | 0.983026 | -3 | 2.036657 | 2.036657 | 0.02791 | 0.0854 | 0.91691 | 0.25567 | 12.70833 | 0.16 | 0 | Neutral |
| testneg_766 | 0.09 | -0.3 | 2.911406 | -3 | 0.261922 | -3 | 0.25421 | 0.16235 | 0.12403 | 0.12788 | 29.86111 | 0.06 | 0 | Neutral |
| testneg_767 | 0 | 0 | 1.135324 | -3 | -0.486478 | -3 | 0.79692 | 0.51979 | 0.64608 | 0.35524 | 10.41667 | 0.070357369 | 0 | Neutral |
| testneg_768 | 0 | 0 | 0.044696 | -3 | 2.089989 | -3 | 0.16125 | 0.08406 | 0.99941 | 0.50288 | 14.42474 | 0.085439068 | 0 | Neutral |
| testneg_769 | -0.02 | -0.45 | 3.26904 | -3 | 3.612522 | -3 | 0.18554 | 0.12299 | 0.10183 | 0.11847 | 23.90643 | 0.100524852 | 0 | Neutral |
| testneg_770 | -0.02 | 0.36 | 4.444198 | -3 | 7.419656 | -3 | 0.18165 | 0.49827 | 0.19898 | 0.09316 | 5 | 0.649772727 | 94.4 | Disease_causing |
| testneg_771 | 0 | -0.32 | 1.514021 | -3 | 0.146321 | -3 | 0.02791 | 0.0854 | 0.11827 | 0.23034 | 12.31894 | 0.1 | 0 | Neutral |
| testneg_772 | 0 | 0 | -0.480488 | -3 | -1.66039 | -3 | 0.11797 | 0.12788 | 0.06957 | 0.07707 | 29.68935 | 0.010522529 | 0 | Neutral |
| testneg_773 | 0.15 | -0.28 | 0.268824 | -3 | -0.041068 | -3 | 0.9924 | 0.37896 | 0.19898 | 0.07875 | 23.79887 | 0.08 | 0 | Neutral |
| testneg_774 | 0 | 0.32 | -0.420859 | -3 | 1.987251 | -4.39493 | 0.0597 | 0.18092 | 0.05183 | 0.18779 | 19.90385 | 0.203166667 | 0 | Neutral |
| testneg_775 | 0.07 | -0.31 | 2.733966 | -3 | -1.086101 | -1.086101 | 0.34205 | 0.11195 | 0.20557 | 0.10131 | 21.47432 | 0.01 | 0 | Neutral |
| testneg_776 | 0 | 0 | 1.66597 | -3 | 4.959501 | -3 | 0.66027 | 0.23984 | 0.14301 | 0.12788 | 18.37615 | 0.114102564 | 0 | Neutral |
| testneg_777 | 0 | 0 | 0.112197 | 0.112197 | 1.33986 | -3 | 0.1123 | 0.1568 | 0.19199 | 0.16616 | 17.77778 | 0.101878342 | 0 | Neutral |
| testneg_778 | -0.05 | 0.81 | 2.856976 | -3 | 4.649335 | -3 | 0.19898 | 0.12788 | 0.37424 | 0.12788 | 13.89558 | 0.19477406 | 0.2 | Neutral |
| testneg_779 | -0.02 | 0.63 | -1.218292 | -1.411613 | 0.979179 | -3 | 0.13696 | 0.09566 | 0.63233 | 0.16248 | 17.55483 | 0.060490431 | 0 | Neutral |
| testneg_780 | -0.09 | 0.25 | 5.036276 | -3 | 5.893689 | -3 | 0.22944 | 0.09039 | 0.21383 | 0.11171 | 18.85932 | 0.368276807 | 12.4 | Neutral |
| testneg_781 | 0 | 0 | 0.188212 | -3 | -0.298555 | -0.298555 | 0.34859 | 0.09799 | 0.10314 | 0.12173 | 20.40047 | 0.011222148 | 0 | Neutral |
| testneg_782 | 0.02 | -0.28 | 1.428565 | -3 | 5.928152 | -3 | 0.78664 | 0.11725 | 0.68228 | 0.09762 | 9.98555 | 0.106538961 | 0 | Neutral |
| testneg_783 | 0 | 0 | 1.916985 | -3 | 3.277943 | -3 | 0.84031 | 0.16749 | 0.10317 | 0.14954 | 10.41667 | 0.241696385 | 0 | Neutral |
| testneg_784 | 0 | 0 | -0.880286 | -3 | 4.528664 | -3 | 0.17083 | 0.31043 | 0.49106 | 0.89143 | 10.41667 | 0.442923629 | 33 | Neutral |
| testneg_785 | 0.11 | -0.19 | 2.886405 | -3 | 1.655615 | -3 | 0.62966 | 0.14896 | 0.06667 | 0.08666 | 10.41667 | 0.06 | 0 | Neutral |
| testneg_786 | 0.16 | -1.37 | 6.076778 | -3 | 1.573155 | -3 | 0.33818 | 0.12788 | 0.06296 | 0.09823 | 10.41667 | 0.22 | 0.4 | Neutral |
| testneg_787 | -0.02 | -0.13 | 3.206825 | -3 | 1.789477 | -3 | 0.55197 | 0.11305 | 0.85482 | 0.1072 | 6.0461 | 0.04 | 0 | Neutral |
| testneg_788 | 0.02 | -0.28 | 0.888018 | -3 | 4.823598 | -3 | 0.65284 | 0.2592 | 0.18122 | 0.16895 | 19.44444 | 0.491626709 | 51.2 | Disease_causing |
| testneg_789 | 0.19 | -0.27 | 1.4509 | -3 | 0.104192 | -3 | 0.64566 | 0.22552 | 0.67644 | 0.11323 | 20.65799 | 0.06 | 0 | Neutral |
| testneg_790 | 0.07 | 0.38 | 0.500759 | -3 | 5.342136 | -3 | 0.12497 | 0.09805 | 0.1496 | 0.16692 | 25.55556 | 0.32 | 4.2 | Neutral |
| testneg_791 | 0 | 0 | 0.74063 | -3 | 3.087586 | -3 | 0.07394 | 0.09408 | 0.3828 | 0.5569 | 15.10472 | 0.142013889 | 0 | Neutral |
| testneg_792 | 0 | -0.01 | 2.187535 | -3 | 1.256694 | -3 | 0.93654 | 0.38765 | 0.53325 | 0.48065 | 4.72222 | 0.082059228 | 0 | Neutral |
| testneg_793 | 0 | -0.01 | 7.652567 | -3 | 0.313674 | -3 | 0.73253 | 0.09747 | 0.26812 | 0.30371 | 13.51351 | 0.447083333 | 34.4 | Neutral |
| testneg_794 | 0 | 0 | 1.28208 | -3 | 6.003616 | -3 | 0.38375 | 0.09256 | 0.33166 | 0.11755 | 20.83333 | 0.061951621 | 0 | Neutral |
| testneg_795 | 0.008 | 0.24890909 | 4.814261 | -3 | 1.035434 | -3 | 0.08775 | 0.12788 | 0.4558 | 0.9277 | 11.08369 | 0.235235806 | 0.6 | Neutral |
| testneg_796 | 0.02 | -0.28 | 1.428565 | -3 | 4.19883 | -3 | 0.78664 | 0.11725 | 0.28685 | 0.1118 | 10.15099 | 0.044448713 | 0 | Neutral |
| testneg_797 | 0 | 0 | 1.104198 | -3 | 1.233889 | -3 | 0.81169 | 0.11096 | 0.10228 | 0.22182 | 29.16667 | 0.090470085 | 0 | Neutral |
| testneg_798 | 0.05 | 0.29 | -0.614223 | -3 | 3.048743 | -3 | 0.20502 | 0.10447 | 0.96475 | 0.27545 | 18.96175 | 0.161602247 | 0 | Neutral |
| testneg_799 | -0.14 | -0.54 | -0.251883 | -3 | 3.328978 | -3 | 0.39258 | 0.12788 | 0.13001 | 0.13176 | 34.2081 | 0.19284143 | 0 | Neutral |
| testneg_800 | -0.18 | 1.05 | 1.851573 | -3 | 0.071089 | -3 | 0.13295 | 0.12914 | 0.94531 | 0.91608 | 16.88659 | 0.16 | 0 | Neutral |
| testneg_801 | 0.09125 | -0.2675 | 10.838582 | -3 | 1.430219 | -3 | 0.29478 | 0.12788 | 0.21731 | 0.11795 | 5.69444 | 0.41 | 22.2 | Neutral |
| testneg_802 | -0.1 | -1.9 | 4.518999 | -3 | 1.389804 | -3 | 0.90955 | 0.10096 | 0.10703 | 0.11611 | 46.16159 | 0.100501253 | 0 | Neutral |
| testneg_803 | 0.08 | 0.03 | -1.709673 | -3 | 1.936037 | -3 | 0.16708 | 0.11594 | 0.12951 | 0.08688 | 14.27224 | 0.02 | 0 | Neutral |
| testneg_804 | -0.02 | -0.41 | -1.58691 | -3 | 3.713951 | -3 | 0.10617 | 0.15182 | 0.1846 | 0.08682 | 19.09347 | 0.145 | 0 | Neutral |
| testneg_805 | 0 | 0 | 0.515297 | -3 | -2.338688 | -2.338688 | 0.29417 | 0.10965 | 0.33129 | 0.09957 | 17.21847 | 0 | 0 | Neutral |
| testneg_806 | 0.18 | -1.05 | 4.646906 | -3 | -0.133819 | -3 | 0.09508 | 0.09834 | 0.13023 | 0.29692 | 13.19973 | 0.266833333 | 1 | Neutral |
| testneg_807 | 0.07 | -0.31 | 1.374903 | -3 | 6.179556 | -3 | 0.50649 | 0.11394 | 0.7062 | 0.83002 | 9.88506 | 0.41 | 20.6 | Neutral |
| testneg_808 | 0 | 0 | 1.554921 | -3 | 1.038417 | -3 | 0.10379 | 0.34799 | 0.32897 | 0.10658 | 14.79675 | 0.03 | 0 | Neutral |
| testneg_809 | -0.02 | 0.68 | 0.250053 | -3 | -0.295414 | -3 | 0.36541 | 0.27749 | 0.09741 | 0.11013 | 18.33333 | 0.040296013 | 0 | Neutral |
| testneg_810 | 0 | -0.01 | 0.306573 | -3 | 2.503507 | -3 | 0.03898 | 0.09943 | 0.14349 | 0.27976 | 26.28502 | 0.122916667 | 0 | Neutral |
| testneg_811 | -0.15 | 1.06 | 6.731764 | -3 | 1.447263 | -3 | 0.91619 | 0.10042 | 0.19531 | 0.18996 | 23.78562 | 0.46 | 41.2 | Neutral |
| testneg_812 | 0 | 0 | 0.061034 | -3 | 1.1308 | -3 | 0.08779 | 0.1777 | 0.19898 | 0.11185 | 35.81824 | 0.020710702 | 0 | Neutral |
| testneg_813 | -0.19 | -0.51 | 2.183162 | 1.811863 | 2.566854 | -3 | 0.22078 | 0.12788 | 0.42797 | 0.17974 | 16.88659 | 0.3 | 2.6 | Neutral |
| testneg_814 | 0.03 | 0.1 | 5.424976 | -3 | 1.383675 | -1.104207 | 0.14956 | 0.10585 | 0.16966 | 0.61198 | 10.22898 | 0.375315859 | 14.2 | Neutral |
| testneg_815 | 0 | 0 | 0.634596 | -3 | -0.705378 | -3 | 0.54489 | 0.09921 | 0.2852 | 0.12788 | 19.13226 | 0.01 | 0 | Neutral |
| testneg_816 | 0.02 | -0.36 | 3.235017 | -3 | 0.185181 | -3 | 0.30329 | 0.14582 | 0.82156 | 0.14621 | 4.86423 | 0.093084344 | 0 | Neutral |
| testneg_817 | 0 | 0 | -0.156459 | -3 | -0.9279 | -3 | 0.10774 | 0.13117 | 0.61867 | 0.15658 | 16.0119 | 0.050607296 | 0 | Neutral |
| testneg_818 | -0.01 | 0.57 | 1.471683 | -3 | 6.259581 | -3 | 0.05677 | 0.06758 | 0.17843 | 0.15468 | 18.88889 | 0.350797414 | 7 | Neutral |
| testneg_819 | 0 | -0.32 | 2.337712 | -3 | 3.997807 | -3 | 0.14273 | 0.1162 | 0.19403 | 0.09278 | 13.51598 | 0.069266667 | 0 | Neutral |
| testneg_820 | 0.03 | 0.1 | 2.021264 | -3 | -0.605051 | -3 | 0.07098 | 0.39918 | 0.48144 | 0.46642 | 7.46424 | 0.210411765 | 0 | Neutral |
| testneg_821 | 0 | 0 | 3.520044 | -3 | 3.529157 | -3 | 0.36324 | 0.15729 | 0.03811 | 0.05629 | 51.93219 | 0.149217852 | 0 | Neutral |
| testneg_822 | 0 | -0.01 | 0.636505 | -3 | 4.704757 | -3 | 0.84238 | 0.12559 | 0.12855 | 0.11656 | 18.88889 | 0.044282343 | 0 | Neutral |
| testneg_823 | 0 | 0 | 2.044018 | -3 | 3.027149 | -3 | 0.63431 | 0.09837 | 0.10812 | 0.12788 | 15.06206 | 0.050339862 | 0 | Neutral |
| testneg_824 | 0 | 0 | -0.074927 | -3 | 2.038093 | -3 | 0.99538 | 0.40044 | 0.58705 | 0.1604 | 10 | 0.095548885 | 0 | Neutral |
| testneg_825 | 0 | 0 | 2.25466 | -3 | -1.848417 | -3 | 0.22489 | 0.16646 | 0.12036 | 0.18651 | 10.41667 | 0.089232955 | 0 | Neutral |
| testneg_826 | 0.08 | 0.03 | -1.968383 | -3 | -0.299109 | -3 | 0.20379 | 0.43705 | 0.82846 | 0.13042 | 10 | 0.09 | 0 | Neutral |
| testneg_827 | 0.08 | 0.03 | 0.695461 | -3 | 1.956938 | -3 | 0.15989 | 0.12788 | 0.19898 | 0.06726 | 44.32245 | 0.02 | 0 | Neutral |
| testneg_828 | 0.08 | 0.03 | -0.126022 | -3 | 3.987084 | -3 | 0.00971 | 0.0791 | 0.1056 | 0.12788 | 19.28571 | 0.110185185 | 0 | Neutral |
| testneg_829 | 0 | 0 | 0.97071 | -3 | 3.683685 | -3 | 0.17855 | 0.11262 | 0.09633 | 0.12788 | 18.33333 | 0.04 | 0 | Neutral |
| testneg_830 | 0.01 | 0.79 | 1.807734 | -3 | -2.444182 | -3 | 0.14273 | 0.1162 | 0.25637 | 0.37408 | 19.68254 | 0.088720085 | 0 | Neutral |
| testneg_831 | 0 | 0 | 0.278729 | -3 | -0.157172 | -3 | 0.95682 | 0.22744 | 0.39344 | 0.12788 | 10.23791 | 0.01 | 0 | Neutral |
| testneg_832 | 0 | 0.32 | 4.911711 | -3 | 2.22273 | -3 | 0.13278 | 0.20513 | 0.09353 | 0.12788 | 18.70407 | 0.18 | 0 | Neutral |
| testneg_833 | 0 | 0 | -0.247515 | -3 | 2.055296 | 2.055296 | 0.3769 | 0.12788 | 0.15092 | 0.09074 | 20.8819 | 0.04 | 0 | Neutral |
| testneg_834 | 0 | 0 | 1.545946 | -3 | -1.528753 | -1.528753 | 0.03844 | 0.12788 | 0.10482 | 0.12788 | 31.65793 | 0.04 | 0 | Neutral |
| testneg_835 | 0 | 0 | 0.116245 | -3 | 0.177663 | -0.172859 | 0.8238 | 0.38836 | 0.31646 | 0.09844 | 10 | 0.010784615 | 0 | Neutral |
| testneg_836 | 0 | -0.32 | 1.050891 | -3 | 3.355918 | -3 | 0.85759 | 0.1031 | 0.09515 | 0.07784 | 43.30005 | 0.061597222 | 0 | Neutral |
| testneg_837 | 0.07 | 0.38 | 1.20841 | -3 | 7.190251 | -3 | 0.28211 | 0.25332 | 0.99704 | 0.17463 | 10.41667 | 0.611672619 | 88.8 | Disease_causing |
| testneg_838 | -0.08 | -0.03 | 0.310186 | -3 | 1.990571 | -3 | 0.68863 | 0.09663 | 0.9581 | 0.12788 | 31.96359 | 0.030455487 | 0 | Neutral |
| testneg_839 | 0.03 | 1.22 | 0.949523 | 0.949523 | -1.247889 | -3 | 0.10056 | 0.12608 | 0.27577 | 0.09812 | 10.1991 | 0.0775 | 0 | Neutral |
| testneg_840 | -0.03 | -0.1 | -0.76618 | -3 | 1.070904 | -3 | 0.1503 | 0.12788 | 0.40361 | 0.17821 | 20.55556 | 0.070203093 | 0 | Neutral |
| testneg_841 | 0.18 | -1.05 | 7.116856 | -3 | 1.879759 | -3 | 0.25831 | 0.10988 | 0.30311 | 0.11121 | 12.69209 | 0.27 | 2.6 | Neutral |
| testneg_842 | 0 | 0 | 2.184916 | -3 | 0.965044 | -3 | 0.18512 | 0.12788 | 0.17264 | 0.09426 | 18.70407 | 0.000540936 | 0 | Neutral |
| testneg_843 | 0.15 | -0.28 | -0.431211 | -3 | 0.692756 | -3 | 0.36676 | 0.37749 | 0.17443 | 0.19181 | 10 | 0.142117424 | 0 | Neutral |
| testneg_844 | -0.15 | -0.04 | -0.451132 | -3 | 3.009853 | -3 | 0.12048 | 0.2381 | 0.50741 | 0.17915 | 25.06207 | 0.416603896 | 23.8 | Neutral |
| testneg_845 | 0.01 | 0.79 | -0.188304 | -3 | 2.555582 | -3 | 0.16418 | 0.12788 | 0.65012 | 0.21208 | 5.36194 | 0.100461676 | 0 | Neutral |
| testneg_846 | -0.09 | 0.3 | -2.714151 | -3 | 0.295868 | -3 | 0.47443 | 0.11062 | 0.15989 | 0.12788 | 19.09347 | 0.03 | 0 | Neutral |
| testneg_847 | 0.02 | -0.63 | -0.055258 | -3 | 0.789 | -3 | 0.16139 | 0.09116 | 0.82156 | 0.14621 | 20.27778 | 0.035800439 | 0 | Neutral |
| testneg_848 | 0 | 0 | 0.917232 | -3 | 0.603563 | -3 | 0.25556 | 0.09342 | 0.10058 | 0.13882 | 21.00597 | 0.07 | 0 | Neutral |
| testneg_849 | 0.14 | -1.01 | 7.142186 | -3 | 2.10021 | -3 | 0.16841 | 0.11143 | 0.15526 | 0.08511 | 21.40647 | 0.26 | 2 | Neutral |
| testneg_850 | 0.04 | -0.23 | 5.286462 | -3 | 0.495052 | -3 | 0.26913 | 0.1087 | 0.90112 | 0.20622 | 20.83333 | 0.2290625 | 0.8 | Neutral |
| testneg_851 | 0.14 | -1.01 | 6.136401 | -3 | -0.323214 | -3 | 0.3238 | 0.18333 | 0.15432 | 0.095 | 12.29167 | 0.23 | 1 | Neutral |
| testneg_852 | 0 | 0 | 0.358974 | -3 | 4.566812 | -3 | 0.27197 | 0.12346 | 0.16926 | 0.11871 | 18.33333 | 0.031186811 | 0 | Neutral |
| testneg_853 | 0 | 0 | -0.489156 | -0.489156 | -0.573377 | -3 | 0.05887 | 0.10046 | 0.42366 | 0.14372 | 34.32374 | 0.072517483 | 0 | Neutral |

| | | | | | | | | | | | | | | |
|---|---|---|---|---|---|---|---|---|---|---|---|---|---|---|
| testneg_854 | -0.04 | 0.23 | 2.272008 | -3 | 0.347565 | -3 | 0.23913 | 0.12788 | 0.63233 | 0.16248 | 18.28557 | 0.07020202 | 0 | Neutral |
| testneg_855 | 0.04 | -0.23 | 5.956747 | -3 | 1.720938 | -3 | 0.19898 | 0.15588 | 0.28856 | 0.1544 | 20.71483 | 0.523159523 | 63 | Disease_causing |
| testneg_856 | -0.15 | -0.04 | 2.846922 | -3 | 2.66819 | -3 | 0.18081 | 0.1258 | 0.06922 | 0.07026 | 16.8018 | 0.08 | 0 | Neutral |
| testneg_857 | 0 | 0 | 1.545946 | -3 | 1.16373 | -3 | 0.03844 | 0.12788 | 0.28416 | 0.27805 | 21.99301 | 0.10369453 | 0 | Neutral |
| testneg_858 | -0.02 | 0.63 | -3.022434 | -3 | -0.682676 | -3 | 0.13565 | 0.46572 | 0.10803 | 0.14652 | 10.41667 | 0.163313693 | 0 | Neutral |
| testneg_859 | -0.03 | -0.1 | 3.350172 | -3 | 4.276849 | -3 | 0.13331 | 0.13638 | 0.0778 | 0.0728 | 18.88889 | 0.17 | 0 | Neutral |
| testneg_860 | 0 | 0 | 0.261658 | -3 | -0.489156 | -3 | 0.20055 | 0.12788 | 0.05887 | 0.10046 | 41.07816 | 0.01 | 0 | Neutral |
| testneg_861 | 0.02 | -0.28 | 2.60503 | -1.689038 | 3.751338 | -3 | 0.64256 | 0.12241 | 0.12933 | 0.12788 | 16.84579 | 0.099494949 | 0 | Neutral |
| testneg_862 | 0.07 | 0.7 | 5.31634 | -3 | 0.888608 | -3 | 0.11261 | 0.14664 | 0.62203 | 0.15583 | 10.22898 | 0.34 | 7.6 | Neutral |
| testneg_863 | 0.07 | 0.38 | 1.944832 | -3 | 1.7235 | -3 | 0.69254 | 0.10021 | 0.19898 | 0.10778 | 20.56995 | 0.02 | 0 | Neutral |
| testneg_864 | 0 | -0.01 | 3.05843 | 3.05843 | -0.780137 | -3 | 0.74136 | 0.1049 | 0.06958 | 0.16086 | 18.33333 | 0.53 | 67 | Disease_causing |
| testneg_865 | -0.05 | -0.29 | -1.851788 | -3 | -2.855052 | -3 | 0.78741 | 0.22085 | 0.16545 | 0.10002 | 14.19394 | 0.04 | 0 | Neutral |
| testneg_866 | 0 | -0.01 | 0.558245 | -3 | 1.660713 | -3 | 0.19898 | 0.10362 | 0.65981 | 0.47389 | 12.29167 | 0.024967949 | 0 | Neutral |
| testneg_867 | -0.03 | -0.1 | -0.369846 | -3 | 0.800175 | -3 | 0.33467 | 0.12788 | 0.213 | 0.09535 | 30.13814 | 0.01 | 0 | Neutral |
| testneg_868 | 0 | 0 | -0.505055 | -3 | 4.752866 | -3 | 0.99859 | 0.12788 | 0.75728 | 0.34838 | 4.72222 | 0.171729457 | 0 | Neutral |
| testneg_869 | 0 | 0 | -0.317625 | -3 | 0.728741 | -3 | 0.33127 | 0.49952 | 0.79857 | 0.12788 | 11.25 | 0.03 | 0 | Neutral |
| testneg_870 | 0.00159091 | -0.0003788 | 0.952714 | -3 | 2.23589 | -3 | 0.19979 | 0.09328 | 0.17978 | 0.20186 | 47.09376 | 0.080345238 | 0 | Neutral |
| testneg_871 | 0.18 | -1.05 | 6.199054 | -3 | 4.729977 | -3 | 0.15989 | 0.12788 | 0.05095 | 0.08353 | 25.90917 | 0.3 | 3 | Neutral |
| testneg_872 | 0.07 | 0.38 | 2.257725 | -3 | 6.36911 | -3 | 0.47888 | 0.43756 | 0.17412 | 0.10595 | 9.41366 | 0.343455285 | 5 | Neutral |
| testneg_873 | 0 | 0 | 0.585506 | -3 | 5.626664 | -3 | 0.17146 | 0.19882 | 0.07092 | 0.20005 | 5 | 0.477309368 | 45.2 | Neutral |
| testneg_874 | 0 | -0.01 | 0.242595 | -3 | 0.188403 | -3 | 0.11358 | 0.12788 | 0.39111 | 0.15887 | 27.95796 | 0.06 | 0 | Neutral |
| testneg_875 | 0 | 0 | 0.943238 | -3 | -1.057731 | -3 | 0.19898 | 0.58572 | 0.22901 | 0.12788 | 11.95767 | 0.03 | 0 | Neutral |
| testneg_876 | 0.02 | 0.41 | 0.154673 | -3 | -0.605051 | -3 | 0.04484 | 0.95426 | 0.48144 | 0.46642 | 12.49191 | 0.323884058 | 1.2 | Neutral |
| testneg_877 | 0 | 0 | 4.037148 | -3 | 1.087862 | -3 | 0.92832 | 0.16306 | 0.32606 | 0.10251 | 10.41667 | 0.079583333 | 0 | Neutral |
| testneg_878 | -0.02 | 0.68 | 5.845448 | -3 | -0.037427 | -3 | 0.06926 | 0.08399 | 0.07118 | 0.08861 | 28.95646 | 0.164111111 | 0 | Neutral |
| testneg_879 | -0.01 | 1.44 | 0.989458 | -3 | 2.622206 | -3 | 0.1481 | 0.1117 | 0.8169 | 0.23301 | 16.8018 | 0.170172414 | 0 | Neutral |
| testneg_880 | 0 | 0 | 0.854548 | -3 | 0.632795 | -3 | 0.04483 | 0.59766 | 0.06775 | 0.09511 | 10.83333 | 0.12 | 0 | Neutral |
| testneg_881 | -0.23 | 1.04 | 1.436839 | -3 | 0.015616 | -3 | 0.19898 | 0.12788 | 0.05999 | 0.12788 | 22.47062 | 0.1 | 0 | Neutral |
| testneg_882 | -0.07 | -0.38 | -1.898366 | -1.898366 | -0.445162 | -3 | 0.11983 | 0.24162 | 0.13173 | 0.12788 | 16.57655 | 0.054608974 | 0 | Neutral |
| testneg_883 | 0 | 0 | 0.044696 | -3 | -0.889855 | -0.889855 | 0.16125 | 0.08406 | 0.042 | 0.07218 | 15.45808 | 0.01 | 0 | Neutral |
| testneg_884 | 0.05 | -0.81 | 3.964534 | -3 | 5.457427 | -3 | 0.16326 | 0.09256 | 0.45343 | 0.12014 | 19.09347 | 0.200526557 | 0 | Neutral |
| testneg_885 | -0.15 | 0.28 | -0.668638 | -0.668638 | 2.276796 | -3 | 0.37131 | 0.10994 | 0.06939 | 0.08981 | 24.01158 | 0.07 | 0 | Neutral |
| testneg_886 | -0.02 | 0.68 | 5.284085 | -3 | -2.062509 | -3 | 0.08509 | 0.12788 | 0.20084 | 0.0851 | 25.29456 | 0.12 | 0 | Neutral |
| testneg_887 | 0 | 0 | -0.310339 | -3 | 5.894587 | -3 | 0.87518 | 0.43416 | 0.15364 | 0.08789 | 10.83333 | 0.124325397 | 0 | Neutral |
| testneg_888 | 0 | 0 | 0.617174 | -3 | 1.498072 | -3 | 0.28416 | 0.27805 | 0.22113 | 0.10673 | 5.04453 | 0.01 | 0 | Neutral |
| testneg_889 | 0.01 | 0.8 | -0.725111 | -0.725111 | 2.212176 | -3 | 0.05116 | 0.08996 | 0.63233 | 0.16248 | 18.28557 | 0.1409938 | 0 | Neutral |
| testneg_890 | 0.01 | -0.57 | 1.155339 | 1.155339 | 4.150455 | -3 | 0.10977 | 0.31695 | 0.14767 | 0.09559 | 13.86082 | 0.231345029 | 0.6 | Neutral |
| testneg_891 | 0.04 | -0.23 | 1.940606 | -3 | 2.951507 | -3 | 0.03991 | 0.08342 | 0.18127 | 0.12788 | 15.59524 | 0.06 | 0 | Neutral |
| testneg_892 | -0.06 | -0.18 | 2.504795 | -3 | 5.06041 | -3 | 0.0146 | 0.03045 | 0.81054 | 0.28104 | 13.00964 | 0.358448052 | 8.8 | Neutral |
| testneg_893 | -0.14 | 0.69 | 2.798182 | 2.798182 | 5.894587 | -3 | 0.31043 | 0.09688 | 0.15364 | 0.08789 | 11.25 | 0.47375 | 44.4 | Neutral |
| testneg_894 | 0 | 0 | -0.134529 | -3 | 0.245964 | -3 | 0.46919 | 0.60081 | 0.74099 | 0.44443 | 20.27778 | 0.145762938 | 0 | Neutral |
| testneg_895 | 0.21 | 0.15 | 0.789411 | -3 | 0.196499 | 0.196499 | 0.13067 | 0.1005 | 0.29098 | 0.10482 | 17.88673 | 0.03 | 0 | Neutral |
| testneg_896 | -0.05 | -0.29 | -0.427621 | -3 | 2.317196 | -3 | 0.40803 | 0.47186 | 0.05851 | 0.0724 | 11.87609 | 0.06 | 0 | Neutral |
| testneg_897 | 0 | 0 | 2.165318 | -3 | 1.473689 | -3 | 0.19989 | 0.16777 | 0.21677 | 0.1681 | 31.69228 | 0.114696385 | 0 | Neutral |
| testneg_898 | 0.18 | -1.05 | 4.454666 | -3 | -0.351099 | -3 | 0.09462 | 0.11807 | 0.19898 | 0.12788 | 23.81804 | 0.041451645 | 0 | Neutral |
| testneg_899 | 0 | -0.01 | 0.549483 | -3 | 0.663866 | 0.663866 | 0.46084 | 0.88583 | 0.08283 | 0.11797 | 20.46784 | 0.12 | 0 | Neutral |
| testneg_900 | 0.08 | 0.03 | -1.587151 | -3 | 2.689471 | -3 | 0.15486 | 0.28768 | 0.83798 | 0.14433 | 24.27204 | 0.281102444 | 0.4 | Neutral |
| testneg_901 | 0.19 | -0.25 | 5.093198 | -3 | 0.232341 | -3 | 0.99859 | 0.12788 | 0.19898 | 0.12788 | 18.29457 | 0.12 | 0 | Neutral |
| testneg_902 | 0.03 | -1.03 | 0.839368 | 0.839368 | 3.749392 | -3 | 0.18785 | 0.14723 | 0.10317 | 0.14954 | 24.49846 | 0.420416667 | 26.8 | Neutral |
| testneg_903 | 0 | -0.01 | 0.473989 | -3 | -0.435392 | -3 | 0.19898 | 0.12788 | 0.19898 | 0.12788 | 33.83469 | 0 | 0 | Neutral |
| testneg_904 | -0.04 | 0.23 | -2.175398 | -3 | 1.167527 | -3 | 0.10228 | 0.22182 | 0.7639 | 0.10408 | 30.05952 | 0.08 | 0 | Neutral |
| testneg_905 | 0.03 | 0.12 | 5.829899 | -3 | 0.338128 | -3 | 0.06196 | 0.10881 | 0.32558 | 0.32796 | 10.83333 | 0.425709291 | 26.6 | Neutral |
| testneg_906 | 0 | 0 | 0.148263 | -3 | 0.070243 | -3 | 0.13482 | 0.12788 | 0.14486 | 0.33725 | 18.6768 | 0.051717172 | 0 | Neutral |
| testneg_907 | 0.07 | 0.38 | 5.023091 | -3 | -0.169548 | -3 | 0.19898 | 0.09972 | 0.3834 | 0.3731 | 5.61526 | 0.30213812 | 3 | Neutral |
| testneg_908 | 0.07 | 0.38 | -0.473879 | -3 | 0.211586 | 0.211586 | 0.0695 | 0.12788 | 0.1119 | 0.24875 | 23.35178 | 0.12 | 0 | Neutral |
| testneg_909 | 0 | 0 | -0.258437 | -3 | 1.420779 | -3 | 0.07044 | 0.11679 | 0.14386 | 0.12788 | 27.62269 | 0.01 | 0 | Neutral |
| testneg_910 | 0.18 | 0.16 | 1.885526 | -3 | 2.550881 | -3 | 0.2141 | 0.19929 | 0.19898 | 0.12788 | 14.64539 | 0.110744828 | 0 | Neutral |
| testneg_911 | -0.07 | -0.38 | 3.319584 | -3 | 1.649124 | -3 | 0.99543 | 0.12788 | 0.37635 | 0.16791 | 21.52409 | 0.196287879 | 0 | Neutral |
| testneg_912 | 0 | 0 | 1.315096 | -3 | 6.609854 | -3 | 0.26972 | 0.12484 | 0.18096 | 0.12788 | 45.16728 | 0.13 | 0 | Neutral |
| testneg_913 | 0.02 | 0.45 | -0.031677 | -3 | 1.623152 | 1.623152 | 0.30669 | 0.11489 | 0.19898 | 0.12788 | 20.95603 | 0.020409699 | 0 | Neutral |
| testneg_914 | 0 | 0 | 0.382766 | -3 | -0.147137 | -3 | 0.6201 | 0.12788 | 0.0597 | 0.18092 | 10.41667 | 0.051172851 | 0 | Neutral |
| testneg_915 | 0.22 | -2.66 | 5.126283 | -3 | 7.172848 | -3 | 0.68423 | 0.08163 | 0.24011 | 0.28559 | 17.03372 | 0.664128205 | 94 | Disease_causing |
| testneg_916 | 0.03 | 1.22 | 1.399794 | -3 | 9.402773 | -3 | 0.49425 | 0.25538 | 0.07118 | 0.08861 | 11.875 | 0.478247821 | 46 | Neutral |
| testneg_917 | 0 | 0 | 0.854548 | -3 | 6.938887 | -3 | 0.04483 | 0.59766 | 0.0907 | 0.10273 | 29.44444 | 0.4195 | 20.8 | Neutral |
| testneg_918 | 0 | 0 | 0.902021 | -3 | 3.068089 | -3 | 0.37499 | 0.17803 | 0.10278 | 0.08667 | 23.33333 | 0.078967162 | 0 | Neutral |
| testneg_919 | -0.12 | 1.16 | 6.464782 | 6.464782 | 0.459406 | -3 | 0.98833 | 0.48581 | 0.03981 | 0.09432 | 17.21847 | 0.61 | 92.8 | Disease_causing |
| testneg_920 | 0 | 0 | -0.749587 | -3 | -0.045491 | -3 | 0.84318 | 0.11313 | 0.09456 | 0.14056 | 12.31231 | 0.054907407 | 0 | Neutral |
| testneg_921 | 0.02 | 0.45 | 2.583598 | -3 | 6.352744 | -3 | 0.09058 | 0.50106 | 0.18373 | 0.12788 | 43.33333 | 0.425602273 | 24 | Neutral |
| testneg_922 | -0.04 | 0.23 | 3.260847 | -3 | 0.612807 | -3 | 0.27605 | 0.12788 | 0.2559 | 0.10848 | 29.28445 | 0.02 | 0 | Neutral |
| testneg_923 | 0.04 | -0.23 | 0.604758 | -3 | 2.11694 | -3 | 0.56605 | 0.15893 | 0.40403 | 0.12788 | 18.77939 | 0.04 | 0 | Neutral |
| testneg_924 | 0.18 | 0.16 | 0.697887 | -3 | 2.649004 | -3 | 0.15119 | 0.08823 | 0.8376 | 0.24333 | 12.46106 | 0.143791063 | 0 | Neutral |
| testneg_925 | 0.07 | 0.38 | 0.581582 | -3 | 1.393735 | -3 | 0.06636 | 0.1079 | 0.55441 | 0.08704 | 11.25 | 0.02 | 0 | Neutral |
| testneg_926 | 0 | -0.01 | 0.505705 | -3 | 3.319838 | -1.897594 | 0.73395 | 0.1705 | 0.19898 | 0.12788 | 13.02635 | 0.065298822 | 0 | Neutral |
| testneg_927 | -0.07 | -0.38 | 3.731276 | -3 | 0.933098 | -3 | 0.82455 | 0.2093 | 0.48298 | 0.1128 | 20.96973 | 0.077911765 | 0 | Neutral |
| testneg_928 | 0 | 0 | 1.539176 | 1.539176 | -0.691406 | -3 | 0.91208 | 0.52258 | 0.04046 | 0.06599 | 11.25 | 0.04 | 0 | Neutral |
| testneg_929 | -0.14 | 0.37 | 2.327743 | -3 | 3.566467 | -3 | 0.0538 | 0.0823 | 0.17548 | 0.11384 | 18.6768 | 0.16 | 0 | Neutral |
| testneg_930 | -0.01 | -0.8 | 6.193509 | -3 | 0.355939 | -3 | 0.31817 | 0.17348 | 0.15562 | 0.10384 | 29.65497 | 0.264166667 | 1 | Neutral |
| testneg_931 | 0 | 0 | 1.870152 | -3 | 0.382515 | -3 | 0.25025 | 0.18924 | 0.19898 | 0.12788 | 18.26149 | 0.01 | 0 | Neutral |
| testneg_932 | 0 | 0 | 0.159372 | 0.159372 | 6.195353 | -3 | 0.05228 | 0.12788 | 0.28532 | 0.10663 | 15.07524 | 0.130938416 | 0 | Neutral |
| testneg_933 | 0.16 | -1.37 | 2.488273 | -3 | 2.95462 | -3 | 0.49629 | 0.11989 | 0.10809 | 0.09057 | 24.71852 | 0.090226244 | 0 | Neutral |
| testneg_934 | -0.01 | -0.8 | 4.547515 | -3 | 0.90918 | -3 | 0.20491 | 0.22087 | 0.098 | 0.08791 | 19.09347 | 0.094684874 | 0 | Neutral |
| testneg_935 | 0.07 | 0.7 | 2.352012 | -3 | 3.725973 | -3 | 0.45354 | 0.09957 | 0.27058 | 0.13363 | 9.68746 | 0.267631046 | 0.6 | Neutral |
| testneg_936 | -0.19 | 0.25 | 0.502289 | -3 | 3.56934 | -3 | 0.57349 | 0.09764 | 0.25884 | 0.12788 | 16.8018 | 0.149375 | 0 | Neutral |
| testneg_937 | 0.22 | -2.66 | 6.983534 | -3 | -0.811454 | -0.811454 | 0.62645 | 0.12788 | 0.72912 | 0.18218 | 26.95989 | 0.357172439 | 9.6 | Neutral |
| testneg_938 | 0 | 0 | 2.057663 | -3 | -0.117585 | -3 | 0.11187 | 0.10744 | 0.56025 | 0.12788 | 13.51351 | 0.01 | 0 | Neutral |
| testneg_939 | 0.07 | -1.12 | 1.763972 | -3 | 2.51125 | -3 | 0.20914 | 0.10089 | 0.31344 | 0.12788 | 13.51351 | 0.03 | 0 | Neutral |
| testneg_940 | 0.07 | 0.33 | -0.896362 | -3 | -1.400944 | -3 | 0.9924 | 0.37896 | 0.16061 | 0.12426 | 13.53335 | 0.07 | 0 | Neutral |
| testneg_941 | 0.03 | -1.03 | 2.660158 | -3 | 2.477978 | -3 | 0.18243 | 0.11102 | 0.24519 | 0.12788 | 17.77778 | 0.04 | 0 | Neutral |
| testneg_942 | 0.04 | -0.23 | 1.062269 | -3 | 1.504305 | -3 | 0.24184 | 0.06921 | 0.22233 | 0.31758 | 10.83333 | 0.059392857 | 0 | Neutral |
| testneg_943 | 0 | -0.01 | 7.854373 | -3 | -1.970817 | -3 | 0.4113 | 0.2303 | 0.2058 | 0.28174 | 24.27489 | 0.581547619 | 84.8 | Disease_causing |
| testneg_944 | -0.15 | -0.04 | 2.900501 | -3 | 2.01894 | -3 | 0.31906 | 0.1705 | 0.09508 | 0.09834 | 18.33333 | 0.11 | 0 | Neutral |
| testneg_945 | 0.14 | -1.01 | 5.679483 | -3 | 1.372699 | -3 | 0.34467 | 0.13076 | 0.06257 | 0.12788 | 29.63392 | 0.17 | 0.2 | Neutral |
| testneg_946 | 0.03 | 0.12 | 3.603334 | -3 | -0.908867 | -3 | 0.15954 | 0.12788 | 0.33332 | 0.12106 | 17.02146 | 0.02 | 0 | Neutral |
| testneg_947 | 0 | 0 | 0.299785 | -3 | 1.148851 | -3 | 0.64152 | 0.15239 | 0.18671 | 0.12788 | 18.62915 | 0.01 | 0 | Neutral |
| testneg_948 | 0 | 0 | -0.648338 | -3 | -0.459078 | -0.459078 | 0.37344 | 0.11262 | 0.19214 | 0.12788 | 19.43262 | 0.01 | 0 | Neutral |
| testneg_949 | -0.03 | 1.03 | -2.025835 | -3 | 3.132136 | -3 | 0.52457 | 0.38957 | 0.0105 | 0.12788 | 22.18656 | 0.199381579 | 0 | Neutral |
| testneg_950 | 0 | 0 | 1.217695 | -3 | 5.529935 | -3 | 0.27629 | 0.11166 | 0.21596 | 0.08969 | 19.44444 | 0.06 | 0 | Neutral |
| testneg_951 | 0.14 | -0.37 | 0.198379 | -3 | 0.807877 | 0.807877 | 0.03991 | 0.08342 | 0.10751 | 0.08512 | 22.62411 | 0.05 | 0 | Neutral |
| testneg_952 | 0 | 0 | 1.169905 | -3 | -0.267626 | -3 | 0.51398 | 0.38821 | 0.19898 | 0.12788 | 17.62359 | 0.016025641 | 0 | Neutral |
| testneg_953 | 0.2 | -1.26 | 2.073808 | -3 | -0.272804 | -0.272804 | 0.08683 | 0.26961 | 0.17839 | 0.12009 | 12.53918 | 0.084479167 | 0 | Neutral |
| testneg_954 | 0 | -0.01 | 0.655486 | -3 | 4.695019 | -3 | 0.20772 | 0.16918 | 0.29357 | 0.79524 | 20.27778 | 0.481093089 | 47.4 | Neutral |
| testneg_955 | 0.21 | 0.15 | 4.516879 | -3 | 5.305114 | -3 | 0.1296 | 0.164 | 0.11501 | 0.13748 | 21.38889 | 0.674846796 | 95 | Disease_causing |
| testneg_956 | 0.07 | -0.31 | -0.565166 | -3 | 5.049328 | -3 | 0.9832 | 0.39904 | 0.22234 | 0.09474 | 19.84262 | 0.171141636 | 0 | Neutral |
| testneg_957 | 0.08 | 0.03 | -0.240285 | -3 | 7.056082 | -3 | 0.35082 | 0.12788 | 0.83114 | 0.60631 | 11.25 | 0.406681818 | 20.6 | Neutral |
| testneg_958 | 0 | 0 | 0.240863 | -3 | 0.925103 | -3 | 0.22816 | 0.10878 | 0.0749 | 0.12788 | 16.07106 | 0.000122846 | 0 | Neutral |
| testneg_959 | 0.14 | -1.01 | 6.92218 | -3 | 5.203439 | -3 | 0.15767 | 0.12788 | 0.17969 | 0.11473 | 13.51351 | 0.51 | 58.2 | Disease_causing |
| testneg_960 | 0.04 | -0.23 | 0.321069 | -3 | 2.38149 | -3 | 0.08408 | 0.12788 | 0.10511 | 0.12788 | 12.37434 | 0.02 | 0 | Neutral |
| testneg_961 | -0.09 | 0.3 | -1.493022 | -3 | -0.017058 | -3 | 0.25159 | 0.12788 | 0.08529 | 0.12376 | 23.86495 | 0.0225 | 0 | Neutral |
| testneg_962 | 0.05 | -0.81 | -0.842653 | -3 | -0.48519 | -3 | 0.06708 | 0.09234 | 0.19898 | 0.12788 | 28.89754 | 0.015 | 0 | Neutral |
| testneg_963 | 0 | 0 | 0.057075 | -3 | 0.381813 | -3 | 0.97504 | 0.18545 | 0.33127 | 0.49952 | 10.41667 | 0.053738184 | 0 | Neutral |
| testneg_964 | 0.09 | -0.3 | 1.334595 | -3 | 1.730022 | -3 | 0.35843 | 0.15758 | 0.0472 | 0.12788 | 10.41667 | 0.03 | 0 | Neutral |
| testneg_965 | -0.05 | -1.52 | 1.347937 | -3 | 2.145747 | -3 | 0.9827 | 0.16837 | 0.98535 | 0.15071 | 18.86584 | 0.15009434 | 0 | Neutral |
| testneg_966 | 0.05 | -0.81 | 1.039181 | 1.039181 | 0.147975 | -3 | 0.1551 | 0.09276 | 0.06898 | 0.11659 | 41.3996 | 0.04 | 0 | Neutral |
| testneg_967 | 0 | 0 | 0.537223 | -3 | 10.513159 | -3 | 0.08186 | 0.11813 | 0.0961 | 0.09786 | 12.29167 | 0.34 | 6.4 | Neutral |
| testneg_968 | 0 | -0.01 | 0.730013 | -3 | 0.281717 | -3 | 0.14785 | 0.09084 | 0.68436 | 0.10157 | 16.8018 | 0.01 | 0 | Neutral |
| testneg_969 | 0.21 | 0.15 | 6.210114 | -3 | 0.775551 | -3 | 0.59864 | 0.14106 | 0.0903 | 0.09372 | 21.99134 | 0.21 | 0.4 | Neutral |
| testneg_970 | 0.03 | 0.1 | -0.042671 | -3 | 1.126466 | -3 | 0.20951 | 0.16278 | 0.37275 | 0.16262 | 10.41667 | 0.09 | 0 | Neutral |
| testneg_971 | 0 | -0.32 | 0.969562 | 0.345116 | 1.298472 | -3 | 0.12121 | 0.4142 | 0.75415 | 0.29199 | 11.875 | 0.196944444 | 0 | Neutral |
| testneg_972 | 0 | 0 | 0.106009 | -3 | 5.955605 | -3 | 0.2877 | 0.18291 | 0.16311 | 0.09662 | 21.18098 | 0.168281997 | 0 | Neutral |
| testneg_973 | 0.16 | 0.08 | 3.022813 | -3 | 2.246903 | -3 | 0.17301 | 0.10048 | 0.1019 | 0.17526 | 18.12633 | 0.270005952 | 0.6 | Neutral |
| testneg_974 | -0.17 | -0.38 | -0.92068 | -3 | 2.965582 | -3 | 0.28038 | 0.11814 | 0.19898 | 0.15276 | 23.34606 | 0.197362741 | 0 | Neutral |
| testneg_975 | -0.01 | 1.44 | -1.973513 | -3 | 3.544108 | -3 | 0.67796 | 0.10874 | 0.13974 | 0.10225 | 19.09347 | 0.164242081 | 0 | Neutral |

| | | | | | | | | | | | | | | |
|---|---|---|---|---|---|---|---|---|---|---|---|---|---|---|
| testneg_976 | 0.04 | -0.23 | 1.605283 | -3 | 3.997628 | -3 | 0.7257 | 0.64394 | 0.66799 | 0.12788 | 11.25 | 0.199678442 | 0 | Neutral |
| testneg_977 | 0 | 0 | -0.10912 | -3 | 5.041302 | -3 | 0.85984 | 0.40044 | 0.52005 | 0.12788 | 14.22149 | 0.103376894 | 0 | Neutral |
| testneg_978 | -0.21 | -0.15 | -0.217187 | -3 | 1.655973 | -3 | 0.19898 | 0.11262 | 0.1977 | 0.09728 | 13.14653 | 0.041964286 | 0 | Neutral |
| testneg_979 | 0.15 | -0.28 | -0.961423 | -3 | -0.086254 | -3 | 0.99417 | 0.17365 | 0.07387 | 0.12788 | 25.46373 | 0.072428847 | 0 | Neutral |
| testneg_980 | 0.03 | 0.12 | 2.305859 | -3 | 3.828961 | -3 | 0.31924 | 0.11145 | 0.06723 | 0.10296 | 17.21847 | 0.070874416 | 0 | Neutral |
| testneg_981 | 0.18 | 0.16 | 3.553675 | -3 | 0.156645 | -3 | 0.39553 | 0.11991 | 0.11802 | 0.09304 | 14.2398 | 0.03 | 0 | Neutral |
| testneg_982 | 0.01 | 0.79 | 2.037729 | -3 | 0.143544 | -3 | 0.58984 | 0.15588 | 0.17768 | 0.10832 | 17.90495 | 0.03005102 | 0 | Neutral |
| testneg_983 | -0.01 | 0.57 | 0.205271 | -3 | 1.472715 | -3 | 0.06311 | 0.14802 | 0.29624 | 0.1067 | 28.97432 | 0.05 | 0 | Neutral |
| testneg_984 | 0.03 | 0.12 | 0.199359 | 0.199359 | 1.294129 | -3 | 0.07626 | 0.12788 | 0.14498 | 0.33896 | 10.80591 | 0.100235588 | 0 | Neutral |
| testneg_985 | -0.15 | -0.06 | 5.384158 | -3 | 1.594546 | -3 | 0.15 | 0.18963 | 0.09951 | 0.14299 | 19.24301 | 0.482421569 | 48.4 | Neutral |
| testneg_986 | 0 | 0 | -0.281959 | -3 | 0.022532 | -3 | 0.0684 | 0.08803 | 0.04192 | 0.06306 | 19.36238 | 0.014166667 | 0 | Neutral |
| testneg_987 | 0 | 0 | -0.330275 | -0.330275 | 5.467754 | -3 | 0.10462 | 0.1278 | 0.05877 | 0.12788 | 13.07136 | 0.111483516 | 0 | Neutral |
| testneg_988 | 0 | 0 | -0.238028 | -3 | 0.811043 | -3 | 0.17039 | 0.12788 | 0.82148 | 0.14847 | 20.32634 | 0.030211694 | 0 | Neutral |
| testneg_989 | 0.02 | -0.63 | 2.090818 | -3 | 2.297873 | -3 | 0.12974 | 0.11886 | 0.31168 | 0.12788 | 12.28264 | 0.01 | 0 | Neutral |
| testneg_990 | -0.07 | -0.38 | -0.783661 | -3 | 2.067877 | -3 | 0.02059 | 0.066 | 0.11094 | 0.12788 | 20 | 0.050061728 | 0 | Neutral |
| testneg_991 | 0 | 0 | 1.491195 | -3 | -0.416655 | -0.416655 | 0.03391 | 0.09745 | 0.05453 | 0.12788 | 58.09011 | 0.08 | 0 | Neutral |
| testneg_992 | 0.03 | -1.03 | 5.578387 | -3 | -1.031314 | -3 | 0.15704 | 0.72174 | 0.27409 | 0.10785 | 21.52546 | 0.30921875 | 1.8 | Neutral |
| testneg_993 | 0.14 | -1.01 | 3.382399 | -3 | 4.65096 | 4.65096 | 0.97252 | 0.8859 | 0.98578 | 0.91075 | 4.44444 | 0.68 | 97 | Disease_causing |
| testneg_994 | -0.02 | 0.63 | -1.996459 | -3 | 6.687775 | -3 | 0.04988 | 0.25503 | 0.07092 | 0.20005 | 10 | 0.629636438 | 93.2 | Disease_causing |
| testneg_995 | 0 | -0.01 | 1.246339 | -3 | 0.673252 | 0.673252 | 0.54826 | 0.24974 | 0.11897 | 0.3919 | 10.43529 | 0.130134914 | 0 | Neutral |
| testneg_996 | -0.07 | -0.38 | -0.785726 | -3 | 0.819832 | -3 | 0.14956 | 0.10585 | 0.19898 | 0.12788 | 25.16423 | 0.01 | 0 | Neutral |
| testneg_997 | 0.09 | -0.3 | 2.387342 | -3 | 2.60503 | -3 | 0.07428 | 0.52467 | 0.64256 | 0.12241 | 16.42913 | 0.175833333 | 0 | Neutral |
| testneg_998 | 0.21 | 0.15 | 7.308333 | -3 | -0.195051 | -3 | 0.10822 | 0.10942 | 0.36417 | 0.21506 | 21.54177 | 0.367885698 | 11.4 | Neutral |
| testneg_999 | 0.03 | 0.1 | -0.942639 | -3 | 1.345014 | -3 | 0.05996 | 0.12788 | 0.3437 | 0.5169 | 30.41667 | 0.111666667 | 0 | Neutral |
| testneg_1000 | 0.03 | 0.12 | -2.609079 | -3 | 2.372066 | -3 | 0.19898 | 0.09533 | 0.65981 | 0.47389 | 18.82075 | 0.128679258 | 0 | Neutral |
| testneg_1001 | 0 | 0 | 0.50666 | -3 | 1.847923 | -3 | 0.72062 | 0.14321 | 0.13118 | 0.08646 | 13.51351 | 0.01 | 0 | Neutral |
| testneg_1002 | 0.22 | -2.66 | 5.764485 | -3 | -0.180177 | -3 | 0.27175 | 0.14094 | 0.1743 | 0.12788 | 13.30616 | 0.18 | 0 | Neutral |
| testneg_1003 | 0 | 0 | 2.30653 | -3 | -0.771216 | -3 | 0.49106 | 0.89143 | 0.37986 | 0.14229 | 5 | 0.13 | 0 | Neutral |
| testneg_1004 | 0.03 | 0.1 | -1.628421 | -1.628421 | 1.681428 | -3 | 0.54441 | 0.08891 | 0.59691 | 0.11049 | 17.47024 | 0.03 | 0 | Neutral |
| testneg_1005 | 0.07 | -0.31 | 3.295096 | -3 | 1.952626 | -3 | 0.89959 | 0.55078 | 0.2074 | 0.09696 | 17.21847 | 0.120394118 | 0 | Neutral |
| testneg_1006 | -0.03 | -0.1 | 1.681063 | -3 | 2.378747 | -3 | 0.31906 | 0.1705 | 0.7482 | 0.18812 | 10.41667 | 0.160984997 | 0 | Neutral |
| testneg_1007 | -0.03 | -0.1 | 3.306597 | -3 | 6.677811 | -3 | 0.21802 | 0.10187 | 0.06099 | 0.12788 | 14.47445 | 0.33 | 7.6 | Neutral |
| testneg_1008 | 0 | 0 | -0.095889 | -3 | 4.968773 | -3 | 0.14643 | 0.12788 | 0.39932 | 0.10693 | 18.50844 | 0.04 | 0 | Neutral |
| testneg_1009 | 0.14 | -0.69 | 0.584676 | -3 | 0.636487 | -3 | 0.31701 | 0.09986 | 0.84236 | 0.14093 | 10.41667 | 0.04 | 0 | Neutral |
| testneg_1010 | 0 | 0 | 1.196382 | -3 | -2.43178 | -3 | 0.11818 | 0.10561 | 0.10893 | 0.12072 | 12.67869 | 0.00625 | 0 | Neutral |
| testneg_1011 | -0.15 | 1.06 | 3.233126 | -3 | -1.487172 | -1.487172 | 0.66422 | 0.99377 | 0.103 | 0.403 | 4.44444 | 0.350284455 | 6.6 | Neutral |
| testneg_1012 | -0.09 | 0.3 | 3.208069 | -3 | 1.458682 | -3 | 0.24327 | 0.12192 | 0.20577 | 0.12653 | 10.41667 | 0.04 | 0 | Neutral |
| testneg_1013 | 0 | 0 | 0.030464 | -3 | 2.470276 | -3 | 0.33795 | 0.17455 | 0.06841 | 0.12788 | 17.21847 | 0.030960118 | 0 | Neutral |
| testneg_1014 | 0 | -0.01 | 0.242595 | -3 | 2.864158 | -3 | 0.11358 | 0.12788 | 0.61718 | 0.1648 | 10.83333 | 0.086464286 | 0 | Neutral |
| testneg_1015 | 0 | 0 | 1.683026 | -3 | 6.431621 | 6.431621 | 0.84106 | 0.63405 | 0.26676 | 0.76568 | 10.41667 | 0.561075306 | 75.8 | Disease_causing |
| testneg_1016 | 0.02 | -0.36 | 2.305765 | -3 | 1.157706 | -3 | 0.11055 | 0.1127 | 0.09951 | 0.14299 | 18.21104 | 0.06 | 0 | Neutral |
| testneg_1017 | 0 | 0 | 4.941822 | -3 | -0.809518 | -3 | 0.08789 | 0.6361 | 0.47187 | 0.14227 | 13.21003 | 0.340258621 | 5.8 | Neutral |
| testneg_1018 | 0.03 | -1.03 | 4.496184 | -3 | -3.590514 | -3 | 0.44644 | 0.10813 | 0.47322 | 0.11118 | 18.50855 | 0.06 | 0 | Neutral |
| testneg_1019 | 0 | 0 | 1.125734 | -3 | 0.634406 | -3 | 0.12576 | 0.08486 | 0.56824 | 0.10646 | 4.44444 | 0.000212206 | 0 | Neutral |
| testneg_1020 | 0.14 | -1.01 | 2.831948 | -3 | -0.264337 | -3 | 0.46399 | 0.15334 | 0.2547 | 0.07589 | 10 | 0.05 | 0 | Neutral |
| testneg_1021 | 0.18 | -1.05 | 2.232408 | -3 | 4.805738 | -3 | 0.22046 | 0.12788 | 0.03441 | 0.10663 | 19.09347 | 0.13 | 0 | Neutral |
| testneg_1022 | 0 | 0 | -0.843404 | -3 | 5.393767 | -3 | 0.11587 | 0.12001 | 0.0882 | 0.12788 | 18.88889 | 0.095416667 | 0 | Neutral |
| testneg_1023 | 0.12 | -1.14 | 3.580218 | 0.751664 | -0.100475 | -3 | 0.07387 | 0.12788 | 0.11261 | 0.13673 | 20.78454 | 0.238942308 | 0 | Neutral |
| testneg_1024 | 0.04 | -0.23 | 7.16104 | -3 | 3.206825 | 2.04209 | 0.25961 | 0.07098 | 0.55197 | 0.11305 | 16.84579 | 0.404613095 | 21.2 | Neutral |
| testneg_1025 | 0 | 0 | 0.478518 | -3 | 0.856523 | -3 | 0.89959 | 0.55078 | 0.25077 | 0.06261 | 10.83333 | 0.02 | 0 | Neutral |
| testneg_1026 | -0.14 | 0.69 | 6.185375 | -3 | 5.258433 | -3 | 0.27629 | 0.11166 | 0.30474 | 0.12788 | 17.63514 | 0.548944444 | 75.4 | Disease_causing |
| testneg_1027 | 0.16 | -1.37 | 1.079819 | -3 | -1.817551 | -3 | 0.12079 | 0.62023 | 0.44541 | 0.13979 | 10.38179 | 0.20135142 | 0 | Neutral |
| testneg_1028 | 0 | 0 | 1.550481 | -3 | 3.576633 | -3 | 0.30739 | 0.16306 | 0.20431 | 0.17304 | 18.04499 | 0.283240977 | 2 | Neutral |
| testneg_1029 | 0 | 0 | 1.66597 | -3 | -0.121817 | -3 | 0.66027 | 0.23984 | 0.33821 | 0.39417 | 18.37615 | 0.090811688 | 0 | Neutral |
| testneg_1030 | 0.07 | 0.38 | 1.79061 | -3 | 0.736335 | -3 | 0.12112 | 0.11353 | 0.98641 | 0.83983 | 11.19856 | 0.09 | 0 | Neutral |
| testneg_1031 | 0 | -0.01 | 1.940661 | -3 | 6.226032 | -3 | 0.25151 | 0.12788 | 0.75033 | 0.64204 | 10.73171 | 0.359892336 | 13.4 | Neutral |
| testneg_1032 | 0 | 0 | 0.616139 | -3 | 1.219225 | -3 | 0.47997 | 0.10506 | 0.29548 | 0.10732 | 20.83333 | 0.01 | 0 | Neutral |
| testneg_1033 | -0.03 | -0.1 | 1.347962 | -3 | -0.40895 | -3 | 0.0418 | 0.12788 | 0.15646 | 0.12788 | 17.21847 | 0.04 | 0 | Neutral |
| testneg_1034 | 0 | 0 | 0.09433 | -3 | 3.41952 | -3 | 0.18925 | 0.32479 | 0.98833 | 0.48581 | 5 | 0.292 | 3.6 | Neutral |
| testneg_1035 | -0.03 | -0.1 | 5.381477 | -3 | 2.002267 | -3 | 0.34698 | 0.12788 | 0.31621 | 0.12788 | 18.33333 | 0.193819444 | 0.2 | Neutral |
| testneg_1036 | 0 | 0 | 1.071372 | -3 | 0.815573 | -3 | 0.15822 | 0.12788 | 0.19898 | 0.12788 | 12.62186 | 0 | 0 | Neutral |
| testneg_1037 | 0.01166667 | -0.0684804 | 2.8284 | -3 | -1.566121 | -1.566121 | 0.20417 | 0.10681 | 0.12003 | 0.12788 | 40.15723 | 0.02 | 0 | Neutral |
| testneg_1038 | 0 | 0 | 1.048522 | -3 | 1.836976 | -3 | 0.6992 | 0.1804 | 0.84491 | 0.30945 | 10.41667 | 0.045916015 | 0 | Neutral |
| testneg_1039 | -0.07 | -0.38 | 1.821777 | -3 | 1.948911 | -3 | 0.11187 | 0.12788 | 0.2442 | 0.11401 | 16.43865 | 0.01 | 0 | Neutral |
| testneg_1040 | 0 | 0 | 1.45834 | -3 | 1.606239 | -3 | 0.64701 | 0.94571 | 0.89002 | 0.12788 | 10 | 0.060997322 | 0 | Neutral |
| testneg_1041 | 0.06 | 0.18 | 2.453732 | -3 | 2.58225 | 1.226373 | 0.18744 | 0.09392 | 0.19883 | 0.09434 | 18.04327 | 0.04 | 0 | Neutral |
| testneg_1042 | 0 | 0 | 1.23629 | -3 | 3.036177 | -3 | 0.6258 | 0.16461 | 0.10835 | 0.12788 | 18.88889 | 0.065232323 | 0 | Neutral |
| testneg_1043 | 0 | 0 | -0.639059 | -3 | -0.040956 | -3 | 0.99178 | 0.93124 | 0.29525 | 0.12788 | 25 | 0.12 | 0 | Neutral |
| testneg_1044 | 0.02 | 0.45 | 1.791247 | -3 | 5.894587 | -3 | 0.39193 | 0.11262 | 0.15364 | 0.08789 | 10 | 0.139570175 | 0 | Neutral |
| testneg_1045 | -0.12 | 1.16 | 3.020638 | -3 | -2.479355 | -3 | 0.29277 | 0.12788 | 0.17853 | 0.32085 | 13.8437 | 0.1946337 | 0 | Neutral |
| testneg_1046 | 0.08 | 0.03 | -1.968383 | -3 | -0.440113 | -3 | 0.20379 | 0.43705 | 0.53592 | 0.11293 | 10.83333 | 0.07 | 0 | Neutral |
| testneg_1047 | 0.09 | -0.25 | 5.708885 | -3 | 2.698795 | 2.698795 | 0.19898 | 0.12788 | 0.17806 | 0.12788 | 23.68839 | 0.296589899 | 2.6 | Neutral |
| testneg_1048 | 0 | -0.32 | -0.849297 | -0.849297 | 2.751884 | -3 | 0.23691 | 0.1015 | 0.1533 | 0.13168 | 25 | 0.097727273 | 0 | Neutral |
| testneg_1049 | -0.07 | -0.38 | -1.415783 | -3 | 3.559501 | -3 | 0.13278 | 0.20513 | 0.3834 | 0.3731 | 6.58998 | 0.479034852 | 46.2 | Neutral |
| testneg_1050 | 0 | 0 | 1.217986 | -3 | 0.17445 | -0.215509 | 0.19898 | 0.12788 | 0.15025 | 0.4872 | 43.61658 | 0.086606079 | 0 | Neutral |
| testneg_1051 | 0 | -0.01 | 2.426894 | -3 | 0.968355 | -0.77772 | 0.06513 | 0.12353 | 0.13433 | 0.09578 | 64.39103 | 0.08 | 0 | Neutral |
| testneg_1052 | -0.01 | 1.44 | 1.762821 | 1.762821 | 3.355037 | -3 | 0.1375 | 0.112 | 0.17443 | 0.19181 | 10.41667 | 0.349988901 | 7.8 | Neutral |
| testneg_1053 | -0.06 | -0.18 | 1.657765 | -3 | 0.034534 | 0.034534 | 0.31923 | 0.17492 | 0.62381 | 0.9477 | 9.75627 | 0.150106383 | 0 | Neutral |
| testneg_1054 | 0.02 | 0.45 | 1.791247 | -3 | 2.935612 | -3 | 0.39193 | 0.11262 | 0.1109 | 0.12788 | 18.6768 | 0.04 | 0 | Neutral |
| testneg_1055 | -0.0190909 | 0.62818182 | 7.974744 | -3 | 5.805734 | -3 | 0.087 | 0.12788 | 0.20502 | 0.12788 | 25.12484 | 0.60993007 | 87.2 | Disease_causing |
| testneg_1056 | -0.08 | -0.03 | -0.573221 | -3 | -2.432125 | -3 | 0.20604 | 0.10821 | 0.19898 | 0.12788 | 25.58105 | 0.01 | 0 | Neutral |
| testneg_1057 | 0 | -0.01 | 1.828167 | -3 | 0.035498 | -3 | 0.25474 | 0.11262 | 0.7639 | 0.10408 | 18.03715 | 0.000218638 | 0 | Neutral |
| testneg_1058 | 0 | 0 | 0.342408 | -3 | 1.423217 | -3 | 0.5255 | 0.15968 | 0.84365 | 0.12788 | 12.5595 | 0.02 | 0 | Neutral |
| testneg_1059 | 0.21 | 0.15 | 6.719042 | -3 | 0.125077 | -3 | 0.07874 | 0.12788 | 0.99564 | 0.25794 | 16.66667 | 0.324166667 | 6.4 | Neutral |
| testneg_1060 | 0.02618182 | -0.2483636 | 11.046933 | -3 | 1.500259 | -3 | 0.4113 | 0.2303 | 0.1787 | 0.1327 | 30.55556 | 0.575166667 | 82 | Disease_causing |
| testneg_1061 | 0 | 0 | 1.05584 | -3 | 3.959602 | -3 | 0.71463 | 0.29732 | 0.15446 | 0.08618 | 18.36275 | 0.0819375 | 0 | Neutral |
| testneg_1062 | 0.21 | 0.15 | 5.850445 | -3 | 7.294015 | -3 | 0.25236 | 0.11809 | 0.07118 | 0.08861 | 14.3695 | 0.602775822 | 88.6 | Disease_causing |
| testneg_1063 | 0 | 0 | 0.297628 | -3 | 5.679638 | -3 | 0.13244 | 0.14875 | 0.61834 | 0.10784 | 34.02638 | 0.147685714 | 0 | Neutral |
| testneg_1064 | 0 | -0.01 | 0.583496 | -3 | -0.433937 | -3 | 0.40663 | 0.24974 | 0.84572 | 0.78546 | 6.76439 | 0.082272621 | 0 | Neutral |
| testneg_1065 | 0.08 | 0.03 | -1.877026 | -3 | -0.448326 | -3 | 0.25758 | 0.2262 | 0.03466 | 0.19518 | 28.58673 | 0.196694678 | 0 | Neutral |
| testneg_1066 | 0 | 0 | 0.039637 | -3 | 1.885526 | -3 | 0.20304 | 0.31462 | 0.11233 | 0.28224 | 6.43275 | 0.110533428 | 0 | Neutral |
| testneg_1067 | 0.07 | 0.38 | -2.492731 | -3 | 4.821873 | -3 | 0.24832 | 0.84371 | 0.78664 | 0.11725 | 10 | 0.230302041 | 0 | Neutral |
| testneg_1068 | 0.16 | -0.37 | 5.827202 | -3 | -0.356956 | -3 | 0.13275 | 0.12788 | 0.11945 | 0.10135 | 34.55685 | 0.11 | 0 | Neutral |
| testneg_1069 | 0 | 0 | -0.313877 | -3 | 1.606772 | -3 | 0.19898 | 0.12788 | 0.35401 | 0.08438 | 17.56627 | 0 | 0 | Neutral |
| testneg_1070 | 0.03 | -1.03 | -0.674759 | -0.674759 | 4.068145 | -3 | 0.02782 | 0.06218 | 0.0406 | 0.12788 | 20.83333 | 0.12 | 0 | Neutral |
| testneg_1071 | 0 | 0 | 0.178815 | -3 | 1.672333 | 1.672333 | 0.5471 | 0.11262 | 0.15338 | 0.13614 | 11.73188 | 0.080155108 | 0 | Neutral |
| testneg_1072 | 0.07 | 0.33 | 0.081615 | 0.081615 | 5.007683 | -3 | 0.14491 | 0.10508 | 0.07415 | 0.12788 | 12.76276 | 0.1 | 0 | Neutral |
| testneg_1073 | 0.14 | -1.01 | 0.29463 | 0.29463 | 0.852085 | -3 | 0.18507 | 0.12788 | 0.56554 | 0.76978 | 10 | 0.13 | 0 | Neutral |
| testneg_1074 | 0.03 | -1.03 | 2.113936 | -3 | 2.055296 | -3 | 0.4132 | 0.12788 | 0.15092 | 0.09074 | 37.23015 | 0.021072829 | 0 | Neutral |
| testneg_1075 | 0 | 0 | 0.951603 | -3 | 11.53432 | -3 | 0.13535 | 0.13562 | 0.13126 | 0.11628 | 10.41667 | 0.4002 | 19.2 | Neutral |
| testneg_1076 | 0 | 0 | 5.068716 | -3 | -0.565166 | -3 | 0.12424 | 0.09575 | 0.9832 | 0.39904 | 18.25269 | 0.21 | 0.4 | Neutral |
| testneg_1077 | 0.16 | -0.37 | 3.233391 | -3 | 3.809453 | -3 | 0.2674 | 0.28725 | 0.14462 | 0.11239 | 11.90227 | 0.254627674 | 0.6 | Neutral |
| testneg_1078 | 0.03 | -1.03 | 2.793619 | -3 | -0.121102 | -0.121102 | 0.13011 | 0.14157 | 0.78935 | 0.17351 | 10.22898 | 0.13 | 0 | Neutral |
| testneg_1079 | 0 | 0 | -0.008063 | -3 | 1.074751 | -3 | 0.31824 | 0.6682 | 0.57991 | 0.2033 | 12.29167 | 0.095769023 | 0 | Neutral |
| testneg_1080 | 0 | 0 | 0.383044 | -3 | 0.734798 | -3 | 0.63819 | 0.32258 | 0.18102 | 0.6537 | 19.70069 | 0.124382619 | 0 | Neutral |
| testneg_1081 | 0 | 0 | 1.313175 | -3 | -0.124213 | -0.124213 | 0.03953 | 0.05763 | 0.74641 | 0.66512 | 26.978 | 0.12 | 0 | Neutral |
| testneg_1082 | 0.05 | 0.29 | 3.783684 | -3 | -2.237532 | -3 | 0.39399 | 0.12788 | 0.42836 | 0.12788 | 33.44125 | 0.04 | 0 | Neutral |
| testneg_1083 | 0 | 0 | 0.648544 | -3 | -0.124213 | -0.124213 | 0.88689 | 0.41514 | 0.74641 | 0.66512 | 4.72222 | 0.070378571 | 0 | Neutral |
| testneg_1084 | 0.14 | -1.85 | 5.795051 | -3 | 3.793394 | -3 | 0.08531 | 0.12788 | 0.23955 | 0.97212 | 10.83333 | 0.632037037 | 89.6 | Disease_causing |
| testneg_1085 | 0 | 0 | -0.237972 | -3 | 6.712641 | -3 | 0.21972 | 0.12788 | 0.25731 | 0.12662 | 22.58084 | 0.15 | 0.2 | Neutral |
| testneg_1086 | 0 | 0 | -0.049164 | -0.049164 | 0.614275 | -3 | 0.12322 | 0.1246 | 0.14472 | 0.11809 | 21.01901 | 0.01 | 0 | Neutral |
| testneg_1087 | 0 | 0.32 | -0.420859 | -3 | 0.973671 | -3 | 0.0597 | 0.18092 | 0.47246 | 0.47221 | 4.72222 | 0.150168079 | 0 | Neutral |
| testneg_1088 | -0.02 | 0.36 | 5.485163 | -3 | 3.506817 | -3 | 0.27407 | 0.1136 | 0.23088 | 0.21602 | 10.41667 | 0.675785948 | 96.4 | Disease_causing |
| testneg_1089 | 0.22 | -2.66 | 6.497626 | -3 | -0.529774 | -3 | 0.29356 | 0.47581 | 0.26878 | 0.09179 | 39.41832 | 0.32 | 3 | Neutral |
| testneg_1090 | -0.02 | -0.13 | -0.203425 | -3 | 2.439678 | -3 | 0.1276 | 0.30104 | 0.25059 | 0.42498 | 11.98582 | 0.284024533 | 2.2 | Neutral |
| testneg_1091 | 0 | 0 | 0.786542 | -3 | 0.959564 | 0.959564 | 0.7257 | 0.64394 | 0.1846 | 0.375 | 20 | 0.20389001 | 0.2 | Neutral |
| testneg_1092 | -0.21 | -0.15 | -0.491857 | -3 | 2.502927 | -3 | 0.07598 | 0.08456 | 0.21122 | 0.12788 | 30.90278 | 0.090121612 | 0 | Neutral |
| testneg_1093 | 0.05 | 1.52 | 3.5315 | -3 | 0.219141 | -3 | 0.24452 | 0.21203 | 0.93284 | 0.71489 | 10.83333 | 0.277235294 | 0.8 | Neutral |
| testneg_1094 | -0.07 | -0.33 | 1.795553 | -3 | 0.359116 | -3 | 0.07681 | 0.22865 | 0.19898 | 0.12788 | 12.23396 | 0.05 | 0 | Neutral |
| testneg_1095 | 0.03 | 0.1 | -0.187467 | -0.187467 | -0.390908 | -3 | 0.12314 | 0.11709 | 0.07652 | 0.30451 | 10 | 0.08 | 0 | Neutral |
| testneg_1096 | 0.11 | -0.19 | 2.575251 | -3 | 1.628587 | -3 | 0.8426 | 0.47836 | 0.67644 | 0.11323 | 22.11633 | 0.09 | 0 | Neutral |
| testneg_1097 | -0.03 | -0.12 | 1.215025 | -3 | 1.176476 | -3 | 0.09028 | 0.10643 | 0.64432 | 0.10409 | 10.30286 | 0.01012502 | 0 | Neutral |

| | | | | | | | | | | | | | | |
|---|---|---|---|---|---|---|---|---|---|---|---|---|---|---|
| testneg_1098 | 0 | 0 | 0.342431 | -3 | 1.541854 | -3 | 0.41647 | 0.11239 | 0.1065 | 0.12788 | 13.51351 | 0.00023823 | 0 | Neutral |
| testneg_1099 | 0.21 | 0.15 | 2.671237 | -3 | 0.734882 | -3 | 0.36852 | 0.0903 | 0.62029 | 0.09082 | 58.0547 | 0.073530021 | 0 | Neutral |
| testneg_1100 | 0.06 | -0.57 | 1.830175 | -3 | -0.214127 | -3 | 0.11351 | 0.10667 | 0.18578 | 0.27854 | 12.58152 | 0.06 | 0 | Neutral |
| testneg_1101 | -0.07 | -0.38 | -1.803731 | -3 | 0.118932 | 0.118932 | 0.07963 | 0.23713 | 0.19567 | 0.09606 | 24.93861 | 0.073642857 | 0 | Neutral |
| testneg_1102 | -0.02 | 0.63 | 3.191277 | -3 | 0.366387 | -3 | 0.37074 | 0.11532 | 0.41102 | 0.0974 | 30.98533 | 0.030385154 | 0 | Neutral |
| testneg_1103 | -0.08 | -0.03 | 2.43732 | -3 | 0.892722 | -3 | 0.66422 | 0.99377 | 0.10244 | 0.10107 | 4.72222 | 0.08 | 0 | Neutral |
| testneg_1104 | 0 | 0 | 2.263609 | 1.253567 | 0.5524 | -3 | 0.4811 | 0.13398 | 0.44644 | 0.10813 | 11.25 | 0.03 | 0 | Neutral |
| testneg_1105 | -0.05 | -0.29 | 0.06448 | -3 | 0.586936 | 0.586936 | 0.30684 | 0.12788 | 0.3405 | 0.1816 | 31.43288 | 0.08 | 0 | Neutral |
| testneg_1106 | 0 | -0.01 | 1.315543 | -3 | 0.483862 | -3 | 0.28358 | 0.38765 | 0.21551 | 0.12788 | 17.77778 | 0.02 | 0 | Neutral |
| testneg_1107 | 0.23 | -1.86 | 4.93441 | -3 | 3.121773 | -1.290749 | 0.11564 | 0.09588 | 0.44479 | 0.20004 | 22.11955 | 0.528223769 | 68.6 | Disease_causing |
| testneg_1108 | -0.08 | -0.03 | -0.48043 | -3 | 4.515915 | -3 | 0.27843 | 0.54325 | 0.49644 | 0.10958 | 18.33333 | 0.189327311 | 0 | Neutral |
| testneg_1109 | 0 | -0.32 | 1.872697 | 1.872697 | 0.821913 | -3 | 0.84468 | 0.55945 | 0.82714 | 0.7562 | 12.70833 | 0.224029214 | 0.6 | Neutral |
| testneg_1110 | 0 | 0 | -0.68826 | -3 | -0.395207 | -3 | 0.32076 | 0.14229 | 0.2098 | 0.44672 | 11.875 | 0.050860258 | 0 | Neutral |
| testneg_1111 | -0.08 | -0.03 | 0.49816 | -3 | -0.775943 | -0.775943 | 0.95069 | 0.1747 | 0.13696 | 0.09566 | 8.49181 | 0.021226045 | 0 | Neutral |
| testneg_1112 | 0.14 | -0.37 | 6.399462 | 6.399462 | 2.507265 | -3 | 0.19721 | 0.12485 | 0.98581 | 0.27035 | 11.875 | 0.800793651 | 100 | Disease_causing |
| testneg_1113 | -0.02 | -0.45 | 2.626432 | -3 | 6.902692 | -3 | 0.17471 | 0.12788 | 0.13356 | 0.09963 | 25.47879 | 0.284166667 | 1.8 | Neutral |
| testneg_1114 | 0 | 0 | 1.716798 | 1.716798 | 0.195289 | -3 | 0.70349 | 0.12788 | 0.09972 | 0.2187 | 11.25 | 0.121247449 | 0 | Neutral |
| testneg_1115 | 0.12 | -1.14 | -0.006272 | -0.006272 | 0.152832 | 0.152832 | 0.07335 | 0.12788 | 0.04829 | 0.09848 | 13.51351 | 0.04 | 0 | Neutral |
| testneg_1116 | 0 | 0 | 1.28208 | -3 | 5.477048 | -3 | 0.38375 | 0.09256 | 0.25549 | 0.12378 | 28.84995 | 0.060130264 | 0 | Neutral |
| testneg_1117 | 0 | 0 | 1.519976 | -3 | -0.736509 | -3 | 0.19898 | 0.12788 | 0.16068 | 0.12788 | 10 | 0 | 0 | Neutral |
| testneg_1118 | 0.07 | 0.38 | 0.878803 | -3 | 1.105669 | -3 | 0.07837 | 0.12788 | 0.44385 | 0.1311 | 10.11674 | 0.030091982 | 0 | Neutral |
| testneg_1119 | 0.04 | -0.23 | -1.417681 | -3 | -0.771684 | -3 | 0.04519 | 0.12788 | 0.21383 | 0.11171 | 18.97014 | 0.03 | 0 | Neutral |
| testneg_1120 | -0.02 | 0.63 | 0.394668 | -3 | -0.711858 | -3 | 0.30209 | 0.12788 | 0.12176 | 0.09356 | 19.51037 | 0.017810245 | 0 | Neutral |
| testneg_1121 | 0 | 0 | 0.974649 | -3 | 1.182833 | -3 | 0.16663 | 0.12788 | 0.89661 | 0.17382 | 12.70833 | 0.010603106 | 0 | Neutral |
| testneg_1122 | -0.14 | 1.85 | 0.923069 | -3 | -0.457896 | -3 | 0.06711 | 0.09765 | 0.22998 | 0.08738 | 19.30596 | 0.11 | 0 | Neutral |
| testneg_1123 | -0.1 | -1.9 | -1.487847 | -3 | 6.762042 | -3 | 0.06728 | 0.20664 | 0.09445 | 0.12788 | 25.39869 | 0.443244048 | 32.2 | Neutral |
| testneg_1124 | 0 | 0 | 1.010818 | -3 | 1.084274 | -3 | 0.36793 | 0.15588 | 0.07606 | 0.13082 | 17.21847 | 0.030820901 | 0 | Neutral |
| testneg_1125 | -0.06 | 0.57 | -2.307199 | -3 | 2.565143 | -3 | 0.07098 | 0.39918 | 0.81305 | 0.17045 | 9.74181 | 0.282224387 | 0.4 | Neutral |
| testneg_1126 | 0.22 | -2.66 | 3.992923 | -3 | -0.435286 | -0.435286 | 0.06344 | 0.12788 | 0.18341 | 0.13901 | 11.25 | 0.181334623 | 0 | Neutral |
| testneg_1127 | 0 | -0.32 | 6.40665 | -3 | 0.27272 | -3 | 0.29828 | 0.12788 | 0.02967 | 0.06605 | 30.43571 | 0.17 | 0.6 | Neutral |
| testneg_1128 | -0.03 | -1.22 | 3.414233 | -3 | 4.485767 | -3 | 0.12424 | 0.09575 | 0.4113 | 0.2303 | 36.26448 | 0.511458333 | 61.2 | Disease_causing |
| testneg_1129 | 0 | 0 | 0.245964 | -3 | -1.483403 | -3 | 0.74099 | 0.44443 | 0.45927 | 0.74955 | 6.94444 | 0.090282946 | 0 | Neutral |
| testneg_1130 | -0.05 | -0.29 | 2.477566 | -3 | -1.566121 | -3 | 0.29451 | 0.13582 | 0.12003 | 0.12788 | 37.26752 | 0.02 | 0 | Neutral |
| testneg_1131 | 0 | 0 | 1.512216 | 1.241234 | 0.981816 | -3 | 0.6258 | 0.16461 | 0.19416 | 0.0901 | 23.00062 | 0.04 | 0 | Neutral |
| testneg_1132 | 0 | 0 | 0.332849 | -3 | 0.723398 | 0.723398 | 0.99595 | 0.23021 | 0.19898 | 0.14957 | 13.86082 | 0.090355346 | 0 | Neutral |
| testneg_1133 | 0 | 0.32 | 3.465539 | 1.154108 | 6.840515 | -3 | 0.13139 | 0.11068 | 0.77364 | 0.12676 | 18.66667 | 0.408688889 | 21.2 | Neutral |
| testneg_1134 | 0.04 | -0.23 | 0.407117 | -3 | 1.448132 | -3 | 0.17728 | 0.46704 | 0.694 | 0.09974 | 23.81445 | 0.07 | 0 | Neutral |
| testneg_1135 | 0 | 0 | -0.237401 | -3 | 6.216009 | -3 | 0.07537 | 0.1399 | 0.19898 | 0.12788 | 23.33333 | 0.161179067 | 0 | Neutral |
| testneg_1136 | 0 | 0 | 0.727897 | -3 | 8.001358 | -3 | 0.34209 | 0.09515 | 0.10895 | 0.11082 | 23.42347 | 0.284583333 | 2.4 | Neutral |
| testneg_1137 | 0.03 | -1.03 | 3.976603 | -3 | -0.026681 | -3 | 0.64125 | 0.28792 | 0.17006 | 0.14718 | 20.54017 | 0.22 | 0.2 | Neutral |
| testneg_1138 | -0.05 | 0.81 | 1.880391 | -3 | 1.68118 | -3 | 0.17818 | 0.15728 | 0.17412 | 0.10595 | 19.79888 | 0.040808608 | 0 | Neutral |
| testneg_1139 | 0.04 | -0.23 | -2.488973 | -3 | 2.228689 | -3 | 0.0513 | 0.33622 | 0.41571 | 0.11118 | 11.96702 | 0.14 | 0 | Neutral |
| testneg_1140 | -0.05 | -0.29 | 1.124718 | -3 | -0.496606 | -3 | 0.55043 | 0.12112 | 0.57564 | 0.12788 | 10.83333 | 0.01 | 0 | Neutral |
| testneg_1141 | 0 | 0 | 0.584441 | -3 | 3.479092 | -3 | 0.97678 | 0.09885 | 0.22113 | 0.10673 | 12.01754 | 0.043958333 | 0 | Neutral |
| testneg_1142 | 0 | -0.01 | 2.002215 | -3 | 1.613322 | -3 | 0.95682 | 0.22744 | 0.39132 | 0.11179 | 4.32527 | 0.01 | 0 | Neutral |
| testneg_1143 | 0 | 0 | 1.005512 | -3 | 7.16059 | -3 | 0.56176 | 0.11451 | 0.2183 | 0.10789 | 15.59524 | 0.19 | 0.6 | Neutral |
| testneg_1144 | 0 | 0 | 0.802919 | 0.802919 | 4.079016 | -3 | 0.12042 | 0.11067 | 0.30837 | 0.19754 | 10.44025 | 0.317369505 | 6 | Neutral |
| testneg_1145 | 0.21 | 0.15 | 4.118971 | -3 | 0.194184 | -3 | 0.126 | 0.14076 | 0.18744 | 0.09392 | 19.33369 | 0.06 | 0 | Neutral |
| testneg_1146 | 0.21 | 0.15 | 3.945475 | -3 | 0.783079 | -3 | 0.27115 | 0.14766 | 0.03391 | 0.09745 | 35.90108 | 0.071602564 | 0 | Neutral |
| testneg_1147 | 0.18 | -1.05 | 4.053851 | -3 | 0.598467 | -3 | 0.41743 | 0.78545 | 0.52836 | 0.12785 | 13.53166 | 0.182030229 | 0 | Neutral |
| testneg_1148 | -0.03 | -0.1 | 0.325459 | -3 | 0.152832 | -3 | 0.05027 | 0.10981 | 0.04829 | 0.09848 | 22.70405 | 0.06 | 0 | Neutral |
| testneg_1149 | 0 | -0.01 | 0.746602 | -3 | 2.823466 | -3 | 0.20304 | 0.31462 | 0.227 | 0.14389 | 11.54762 | 0.18433331 | 0 | Neutral |
| testneg_1150 | 0 | 0 | 0.386087 | 0.386087 | -0.794725 | -3 | 0.26927 | 0.10752 | 0.11153 | 0.10767 | 29.16667 | 0.01 | 0 | Neutral |
| testneg_1151 | 0 | 0 | 0.386087 | 0.386087 | 0.385855 | -3 | 0.26927 | 0.10752 | 0.10153 | 0.13343 | 12.28162 | 0.086607143 | 0 | Neutral |
| testneg_1152 | -0.04 | 1.61 | 6.77188 | -3 | 0.894817 | -3 | 0.19898 | 0.13162 | 0.99317 | 0.60884 | 10.41667 | 0.438333333 | 28.2 | Neutral |
| testneg_1153 | 0.15 | -0.28 | -1.464692 | -3 | 0.752432 | -3 | 0.49629 | 0.11989 | 0.19898 | 0.1347 | 22.3462 | 0.09 | 0 | Neutral |
| testneg_1154 | 0 | 0 | 0.602359 | -3 | 1.222548 | 1.222548 | 0.26758 | 0.10473 | 0.06705 | 0.12788 | 35.42833 | 0.02 | 0 | Neutral |
| testneg_1155 | 0 | 0 | 1.9846 | -3 | 0.694709 | -3 | 0.64701 | 0.94571 | 0.71956 | 0.12519 | 12.05022 | 0.07 | 0 | Neutral |
| testneg_1156 | 0.07 | -0.31 | 1.218458 | -3 | 0.437812 | -3 | 0.13119 | 0.12788 | 0.10723 | 0.33098 | 28.1759 | 0.101024009 | 0 | Neutral |
| testneg_1157 | 0.16 | -1.37 | -0.103493 | -3 | 3.546645 | -3 | 0.12835 | 0.15143 | 0.18502 | 0.21329 | 10.83333 | 0.376098173 | 13 | Neutral |
| testneg_1158 | 0.03 | -1.03 | 2.18442 | -3 | 6.531553 | -3 | 0.19898 | 0.12788 | 0.13555 | 0.16663 | 17.81908 | 0.512178211 | 61 | Disease_causing |
| testneg_1159 | 0 | 0 | 0.086346 | -3 | 0.211278 | -3 | 0.18671 | 0.12788 | 0.25447 | 0.12788 | 42.90702 | 0.000140957 | 0 | Neutral |
| testneg_1160 | 0.15 | -0.28 | -1.379902 | -3 | 2.483347 | -3 | 0.19898 | 0.56316 | 0.06841 | 0.10927 | 12.74476 | 0.125086207 | 0 | Neutral |
| testneg_1161 | 0.15 | -0.28 | 1.770462 | -3 | -0.058596 | -3 | 0.05754 | 0.10881 | 0.1144 | 0.09798 | 10.22898 | 0.03 | 0 | Neutral |
| testneg_1162 | 0 | 0 | 2.676827 | -3 | 5.113079 | -3 | 0.29362 | 0.22177 | 0.9581 | 0.12788 | 12.73545 | 0.246749623 | 0.6 | Neutral |
| testneg_1163 | 0.02 | -0.68 | -0.505763 | -3 | 0.784892 | -3 | 0.11585 | 0.12788 | 0.47192 | 0.11305 | 22.95764 | 0.010513158 | 0 | Neutral |
| testneg_1164 | 0 | 0 | 2.041141 | -3 | 2.079972 | -3 | 0.92235 | 0.62242 | 0.25549 | 0.12378 | 8.73053 | 0.03 | 0 | Neutral |
| testneg_1165 | -0.05 | -0.29 | -1.769506 | -3 | -0.578808 | -3 | 0.15341 | 0.12272 | 0.94531 | 0.91608 | 4.72222 | 0.090495272 | 0 | Neutral |
| testneg_1166 | -0.03 | -0.12 | -0.132037 | -3 | 1.572393 | -3 | 0.22794 | 0.10688 | 0.72946 | 0.28603 | 10.44434 | 0.041531469 | 0 | Neutral |
| testneg_1167 | 0.03 | 0.12 | 6.044603 | -3 | 5.588603 | -3 | 0.28485 | 0.12788 | 0.12782 | 0.08502 | 18.76159 | 0.435119048 | 32 | Neutral |
| testneg_1168 | 0 | 0 | 1.649381 | -3 | 1.551505 | -3 | 0.22686 | 0.24974 | 0.05413 | 0.05691 | 11.25 | 0.011481331 | 0 | Neutral |
| testneg_1169 | 0 | 0 | 0.56241 | -3 | 0.714207 | -3 | 0.9511 | 0.54626 | 0.04346 | 0.10785 | 5.69444 | 0.02 | 0 | Neutral |
| testneg_1170 | 0.09 | -1.07 | 4.333168 | -3 | 3.793394 | -3 | 0.19898 | 0.12788 | 0.23955 | 0.97212 | 11.08369 | 0.555460377 | 73 | Disease_causing |
| testneg_1171 | 0.05 | -0.81 | -0.765143 | -0.765143 | -0.858883 | -3 | 0.31627 | 0.23288 | 0.10914 | 0.76607 | 6.0461 | 0.180089286 | 0 | Neutral |
| testneg_1172 | 0.07 | 0.7 | 4.681701 | -3 | 0.781173 | -3 | 0.10571 | 0.12788 | 0.78253 | 0.1114 | 36.72882 | 0.07 | 0 | Neutral |
| testneg_1173 | 0 | -0.01 | -0.302033 | -3 | 2.148349 | -3 | 0.19898 | 0.12788 | 0.13794 | 0.1141 | 19.09347 | 0.000322581 | 0 | Neutral |
| testneg_1174 | 0 | 0 | 0.039637 | -3 | -1.352736 | -3 | 0.20304 | 0.31462 | 0.27976 | 0.12613 | 16.8018 | 0.010085476 | 0 | Neutral |
| testneg_1175 | 0.02 | -0.63 | 5.797862 | -3 | 0.852085 | -3 | 0.15713 | 0.3507 | 0.56554 | 0.76978 | 18.81242 | 0.599642857 | 86.4 | Disease_causing |
| testneg_1176 | 0.03 | 0.12 | 6.140208 | -3 | 1.409648 | -3 | 0.40179 | 0.12112 | 0.88914 | 0.39065 | 10 | 0.394666667 | 17.4 | Neutral |
| testneg_1177 | 0.01 | 0.8 | 4.932002 | -3 | 4.206678 | -3 | 0.21998 | 0.20297 | 0.09093 | 0.28643 | 20.27778 | 0.761631944 | 99.8 | Disease_causing |
| testneg_1178 | 0.03890909 | -0.3345455 | 12.061237 | -3 | -0.700053 | -3 | 0.19898 | 0.12788 | 0.19898 | 0.12788 | 22.49197 | 0.33 | 6.6 | Neutral |
| testneg_1179 | 0.07 | 0.38 | -3.901266 | -3 | 0.228109 | -3 | 0.30669 | 0.11489 | 0.19898 | 0.13057 | 12.38369 | 0.021060363 | 0 | Neutral |
| testneg_1180 | 0.14 | -1.01 | 4.0507 | -3 | 1.154108 | -3 | 0.25989 | 0.11651 | 0.13139 | 0.11068 | 27.05467 | 0.03 | 0 | Neutral |
| testneg_1181 | 0.18 | 0.16 | 5.112733 | -3 | -2.287729 | -3 | 0.03676 | 0.12788 | 0.1837 | 0.27351 | 6.89287 | 0.331666667 | 5.6 | Neutral |
| testneg_1182 | 0.00375758 | 0.04909091 | 1.721395 | -3 | 5.704027 | -3 | 0.20417 | 0.10681 | 0.09504 | 0.38939 | 25.0282 | 0.36782326 | 10.6 | Neutral |
| testneg_1183 | 0.14 | -1.01 | 2.941823 | -3 | 2.057447 | -3 | 0.10285 | 0.10186 | 0.13523 | 0.12788 | 19.09347 | 0.049208791 | 0 | Neutral |
| testneg_1184 | -0.07 | -0.33 | 0.406963 | -3 | 0.989458 | -3 | 0.48825 | 0.10951 | 0.1481 | 0.1117 | 27.81922 | 0.010571895 | 0 | Neutral |
| testneg_1185 | 0.22 | -2.66 | 6.421996 | -3 | -1.168846 | -3 | 0.26337 | 0.12771 | 0.19898 | 0.12788 | 20.41417 | 0.18 | 0 | Neutral |
| testneg_1186 | 0 | 0 | 0.004158 | -3 | 3.138075 | -3 | 0.06789 | 0.09707 | 0.82156 | 0.14621 | 5.69444 | 0.085477778 | 0 | Neutral |
| testneg_1187 | 0.02 | -0.36 | 7.102968 | -3 | 5.67151 | -3 | 0.19898 | 0.12788 | 0.30942 | 0.13105 | 14.16667 | 0.665897436 | 95.4 | Disease_causing |
| testneg_1188 | -0.03 | -0.1 | 3.548299 | 0.884796 | 5.718258 | -3 | 0.26337 | 0.12771 | 0.08143 | 0.44857 | 14.20309 | 0.68 | 98.2 | Disease_causing |
| testneg_1189 | 0 | 0 | 0.395779 | -3 | 2.518162 | -3 | 0.46723 | 0.08832 | 0.13447 | 0.12788 | 12.32809 | 0.021088235 | 0 | Neutral |
| testneg_1190 | 0.01 | 0.8 | -0.214756 | -3 | 3.442352 | -3 | 0.07428 | 0.07645 | 0.04946 | 0.12788 | 33.28611 | 0.09 | 0 | Neutral |
| testneg_1191 | 0 | 0 | 5.011748 | -3 | 4.702387 | -3 | 0.33135 | 0.73068 | 0.08729 | 0.10515 | 10 | 0.435002708 | 30.2 | Neutral |
| testneg_1192 | 0.07 | 0.38 | 0.878803 | -3 | 4.875592 | -3 | 0.07837 | 0.12788 | 0.12269 | 0.08105 | 29.18318 | 0.08 | 0 | Neutral |
| testneg_1193 | -0.06 | 0.57 | 2.929719 | -3 | 3.701009 | -3 | 0.34288 | 0.24605 | 0.12642 | 0.11218 | 7.0922 | 0.251234115 | 1 | Neutral |
| testneg_1194 | 0 | 0 | 0.547246 | -3 | 0.197229 | -3 | 0.81169 | 0.11096 | 0.99994 | 0.51636 | 4.72222 | 0.060211676 | 0 | Neutral |
| testneg_1195 | 0 | 0 | 0.374605 | -3 | -0.164302 | -3 | 0.45112 | 0.12083 | 0.35475 | 0.12788 | 20.98109 | 0.01 | 0 | Neutral |
| testneg_1196 | 0 | 0 | 0.966924 | -3 | -1.162831 | -1.162831 | 0.19898 | 0.12788 | 0.13398 | 0.12788 | 13.51351 | 0 | 0 | Neutral |
| testneg_1197 | -0.07 | -0.33 | -2.373259 | -3 | 1.019702 | -3 | 0.04525 | 0.12788 | 0.98238 | 0.10873 | 10.83333 | 0.07 | 0 | Neutral |
| testneg_1198 | 0.04 | 2.01 | 3.781815 | -3 | 0.885023 | -3 | 0.01704 | 0.05734 | 0.19898 | 0.12788 | 26.45033 | 0.111583333 | 0 | Neutral |
| testneg_1199 | -0.08 | -0.03 | -0.042605 | -3 | -0.344576 | -3 | 0.87806 | 0.38966 | 0.25422 | 0.10102 | 6.94444 | 0.020454545 | 0 | Neutral |
| testneg_1200 | 0 | -0.01 | 0.684698 | -3 | 0.359117 | -3 | 0.09661 | 0.09981 | 0.05564 | 0.06933 | 20.27778 | 0.01 | 0 | Neutral |
| testneg_1201 | 0.09 | -0.25 | 2.341371 | -3 | 2.191197 | -3 | 0.87268 | 0.21837 | 0.18667 | 0.10488 | 10.41667 | 0.07 | 0 | Neutral |
| testneg_1202 | -0.17 | -0.38 | 1.850031 | -3 | 0.445189 | -3 | 0.25904 | 0.16134 | 0.12784 | 0.11244 | 12.30556 | 0.07 | 0 | Neutral |
| testneg_1203 | -0.02 | 0.68 | 3.109082 | -3 | -1.624227 | -3 | 0.31244 | 0.4615 | 0.19237 | 0.53191 | 10 | 0.340254902 | 6.8 | Neutral |
| testneg_1204 | 0.05 | 0.29 | 3.613738 | -3 | 1.372857 | -3 | 0.52388 | 0.25831 | 0.14378 | 0.41915 | 19.28571 | 0.341081923 | 6.8 | Neutral |
| testneg_1205 | 0.04 | -0.23 | 1.130255 | -3 | 5.085097 | -3 | 0.1679 | 0.24232 | 0.30103 | 0.1462 | 21.38889 | 0.518928909 | 61.8 | Disease_causing |
| testneg_1206 | -0.05 | -0.29 | 1.872244 | -3 | 1.275959 | -3 | 0.19494 | 0.14451 | 0.2058 | 0.28174 | 13.79474 | 0.094252871 | 0 | Neutral |
| testneg_1207 | 0 | 0 | 1.305769 | -3 | 1.721437 | -3 | 0.16468 | 0.11275 | 0.13873 | 0.12788 | 23.68839 | 0 | 0 | Neutral |
| testneg_1208 | 0 | 0 | 0.433941 | 0.433941 | -1.646194 | -3 | 0.30475 | 0.21064 | 0.10362 | 0.09649 | 12.32809 | 0.03 | 0 | Neutral |
| testneg_1209 | 0.14 | -1.01 | 4.293539 | -3 | 0.782368 | -3 | 0.11891 | 0.11186 | 0.31651 | 0.12788 | 12.1005 | 0.03 | 0 | Neutral |
| testneg_1210 | 0.15 | 0.06 | 5.619784 | -3 | 5.258974 | 1.22874 | 0.38762 | 0.12788 | 0.19898 | 0.28128 | 34.50745 | 0.703527778 | 98.6 | Disease_causing |
| testneg_1211 | -0.08 | -0.03 | 4.88562 | -3 | 2.147565 | -3 | 0.12974 | 0.11886 | 0.12121 | 0.4142 | 10 | 0.359374045 | 9.6 | Neutral |
| testneg_1212 | 0 | 0 | 0.013394 | -3 | -0.330967 | -3 | 0.12582 | 0.12788 | 0.20657 | 0.12788 | 10.28409 | 0 | 0 | Neutral |
| testneg_1213 | 0 | -0.01 | 8.815204 | -3 | 3.136838 | -3 | 0.51398 | 0.38821 | 0.07999 | 0.10413 | 12.77946 | 0.65775 | 97.4 | Disease_causing |
| testneg_1214 | -0.01 | 0.57 | 0.88667 | -3 | 3.191277 | -3 | 0.06419 | 0.35525 | 0.37074 | 0.11532 | 26.27832 | 0.190060976 | 0 | Neutral |
| testneg_1215 | 0.05 | -0.81 | -0.842653 | -3 | 3.650561 | -3 | 0.06708 | 0.09234 | 0.0961 | 0.21256 | 16.8018 | 0.235265698 | 0 | Neutral |
| testneg_1216 | 0.14 | -0.69 | 1.091597 | -0.390123 | 1.143193 | -3 | 0.09657 | 0.58605 | 0.65564 | 0.11468 | 10.41667 | 0.102538462 | 0 | Neutral |
| testneg_1217 | 0 | 0 | 1.375504 | -3 | 8.488711 | -3 | 0.19898 | 0.12788 | 0.19102 | 0.12788 | 23.36946 | 0.305914286 | 3.2 | Neutral |
| testneg_1218 | 0.15 | -0.28 | 1.351643 | -3 | 8.199987 | -3 | 0.86306 | 0.10962 | 0.09488 | 0.10426 | 11.25 | 0.32 | 3.6 | Neutral |
| testneg_1219 | 0.09 | -0.3 | 4.680216 | -3 | 3.749392 | -3 | 0.18055 | 0.12564 | 0.10317 | 0.14954 | 17.77778 | 0.457176218 | 39.6 | Neutral |

| | | | | | | | | | | | | | | |
|---|---|---|---|---|---|---|---|---|---|---|---|---|---|---|
| testneg_1220 | -0.03 | -0.1 | 0.805377 | -3 | -0.355202 | -3 | 0.3769 | 0.12788 | 0.21802 | 0.10187 | 18.88889 | 0.00061553 | 0 | Neutral |
| testneg_1221 | 0 | 0 | 0.763292 | -3 | 0.175869 | -3 | 0.12596 | 0.18924 | 0.43897 | 0.12441 | 22.40427 | 0.020234244 | 0 | Neutral |
| testneg_1222 | -0.19 | 0.25 | -2.608658 | -3 | 4.534864 | -3 | 0.19898 | 0.12788 | 0.13615 | 0.12788 | 25.91122 | 0.184781746 | 0 | Neutral |
| testneg_1223 | -0.03 | 0.85 | 3.717434 | -3 | 2.706309 | -3 | 0.17006 | 0.14718 | 0.50592 | 0.0952 | 18.88889 | 0.166658801 | 0 | Neutral |
| testneg_1224 | -0.02 | -0.13 | 0.687351 | -3 | 4.805369 | -3 | 0.51675 | 0.11055 | 0.22415 | 0.70109 | 19.69114 | 0.327004167 | 3.6 | Neutral |
| testneg_1225 | 0 | 0 | 1.720206 | -3 | 0.889233 | -1.06889 | 0.6992 | 0.1804 | 0.50392 | 0.12788 | 12.29167 | 0.01 | 0 | Neutral |
| testneg_1226 | 0 | -0.01 | 6.857724 | 6.857724 | 0.742409 | -3 | 0.18224 | 0.10419 | 0.12949 | 0.44208 | 17.77778 | 0.829797619 | 100 | Disease_causing |
| testneg_1227 | -0.03 | -0.1 | 0.271296 | -3 | 0.945927 | -3 | 0.0978 | 0.26338 | 0.21458 | 0.10398 | 39.93056 | 0.05 | 0 | Neutral |
| testneg_1228 | 0 | 0 | 1.07486 | -3 | 2.344705 | -3 | 0.18697 | 0.12232 | 0.20577 | 0.12653 | 10.41667 | 0.01 | 0 | Neutral |
| testneg_1229 | 0.21 | 0.15 | 3.945475 | -3 | -1.643119 | -1.643119 | 0.27115 | 0.14766 | 0.1535 | 0.12382 | 33.05556 | 0.071035714 | 0 | Neutral |
| testneg_1230 | -0.2 | 1.26 | -3.127268 | -3 | 0.792263 | -3 | 0.03502 | 0.08119 | 0.25447 | 0.12788 | 39.20331 | 0.16 | 0 | Neutral |
| testneg_1231 | 0 | -0.01 | 2.661909 | -3 | 0.829687 | -3 | 0.92776 | 0.09677 | 0.06841 | 0.12788 | 16.55807 | 0.020521421 | 0 | Neutral |
| testneg_1232 | 0.03 | 0.12 | 1.527626 | -3 | 0.342122 | -3 | 0.20501 | 0.6166 | 0.74919 | 0.85508 | 10.41667 | 0.161457875 | 0 | Neutral |
| testneg_1233 | 0 | 0 | 0.376038 | -3 | 0.113964 | -3 | 0.18485 | 0.40044 | 0.64914 | 0.12481 | 10 | 0.020178121 | 0 | Neutral |
| testneg_1234 | -0.01 | -0.79 | 3.572092 | -3 | 0.01486 | -3 | 0.339 | 0.13169 | 0.2903 | 0.11176 | 12.58475 | 0.030345238 | 0 | Neutral |
| testneg_1235 | 0.21 | 0.15 | 3.311456 | -3 | -0.444318 | -3 | 0.04483 | 0.59766 | 0.52278 | 0.41207 | 10.41667 | 0.45 | 33.8 | Neutral |
| testneg_1236 | 0.09 | -0.67 | 6.356297 | -3 | 2.777008 | -3 | 0.24966 | 0.12788 | 0.09093 | 0.28643 | 13.07874 | 0.6452 | 92.8 | Disease_causing |
| testneg_1237 | 0.04 | -0.23 | 8.064683 | -3 | 1.448212 | 1.448212 | 0.227 | 0.14389 | 0.87803 | 0.12788 | 19.28571 | 0.4098 | 20.6 | Neutral |
| testneg_1238 | 0 | 0 | -0.216528 | -3 | 6.399462 | -3 | 0.40612 | 0.45135 | 0.19721 | 0.12485 | 9.71235 | 0.201811124 | 0.8 | Neutral |
| testneg_1239 | -0.04 | -2.01 | 5.968963 | -3 | 1.829958 | -3 | 0.33256 | 0.14909 | 0.47775 | 0.10103 | 19.44444 | 0.319486842 | 4 | Neutral |
| testneg_1240 | -0.03 | 0.85 | -2.027152 | -3 | 5.266428 | -3 | 0.04436 | 0.12788 | 0.58994 | 0.11602 | 20.22479 | 0.170213152 | 0 | Neutral |
| testneg_1241 | -0.1 | -1.9 | -0.369606 | -3 | 0.314664 | -3 | 0.24452 | 0.21203 | 0.08224 | 0.29179 | 18.33333 | 0.19 | 0 | Neutral |
| testneg_1242 | 0 | 0 | 1.961488 | -3 | 6.046726 | -3 | 0.29044 | 0.12142 | 0.13559 | 0.1946 | 13.14653 | 0.382630822 | 14 | Neutral |
| testneg_1243 | -0.21 | -0.15 | 0.71504 | -3 | 0.728537 | -3 | 0.21894 | 0.09041 | 0.08521 | 0.11239 | 18.76159 | 0.04 | 0 | Neutral |
| testneg_1244 | 0 | 0 | 1.813076 | -3 | 4.754501 | -0.006195 | 0.53996 | 0.14577 | 0.1837 | 0.27351 | 10.41667 | 0.316243283 | 4.4 | Neutral |
| testneg_1245 | 0 | 0.32 | 2.93612 | -3 | 1.393294 | -3 | 0.05657 | 0.14812 | 0.19898 | 0.09927 | 24.11907 | 0.09828125 | 0 | Neutral |
| testneg_1246 | -0.06 | 0.57 | 1.112541 | -3 | 0.278951 | -3 | 0.07982 | 0.12788 | 0.08975 | 0.07399 | 39.38605 | 0.02 | 0 | Neutral |
| testneg_1247 | -0.09 | 0.3 | 0.782249 | -3 | 4.752866 | -3 | 0.5127 | 0.12263 | 0.75728 | 0.34838 | 10.91667 | 0.294561688 | 2 | Neutral |
| testneg_1248 | -0.07 | -0.38 | 3.896611 | -3 | -0.072198 | -0.072198 | 0.57349 | 0.09764 | 0.1025 | 0.3268 | 10 | 0.223267351 | 0 | Neutral |
| testneg_1249 | -0.04 | -2.01 | 2.571576 | -3 | -0.049164 | -0.049164 | 0.37183 | 0.10564 | 0.12322 | 0.1246 | 30.87252 | 0.05 | 0 | Neutral |
| testneg_1250 | -0.19 | 0.27 | 1.951681 | -3 | -0.15609 | -3 | 0.1875 | 0.10316 | 0.19898 | 0.10335 | 19.44444 | 0.05 | 0 | Neutral |
| testneg_1251 | 0.21 | 0.15 | 1.478329 | -3 | 1.0263 | 0.14403 | 0.27614 | 0.12085 | 0.08183 | 0.11603 | 17.47657 | 0.03 | 0 | Neutral |
| testneg_1252 | 0.15 | 0.06 | 6.054442 | -3 | 1.167634 | -3 | 0.32335 | 0.10604 | 0.89178 | 0.23115 | 17.21847 | 0.314212121 | 4.6 | Neutral |
| testneg_1253 | -0.02 | -0.41 | -1.730578 | -3 | 0.300088 | -3 | 0.24949 | 0.29709 | 0.36494 | 0.12763 | 10 | 0.03 | 0 | Neutral |
| testneg_1254 | 0 | 0 | -0.961702 | -0.961702 | -1.239866 | -3 | 0.46979 | 0.1705 | 0.12173 | 0.12788 | 15.80978 | 0.02 | 0 | Neutral |
| testneg_1255 | -0.03 | -0.1 | 0.988046 | -3 | 1.860726 | -3 | 0.19898 | 0.12788 | 0.5901 | 0.19444 | 10.83333 | 0.050564944 | 0 | Neutral |
| testneg_1256 | 0.02 | 0.41 | 1.045047 | 0.621284 | 4.404174 | -3 | 0.0832 | 0.09323 | 0.09828 | 0.07733 | 13.51351 | 0.110324692 | 0 | Neutral |
| testneg_1257 | 0.01 | -0.57 | 0.809099 | -3 | -0.055901 | -3 | 0.6532 | 0.12788 | 0.33418 | 0.09979 | 21.94393 | 0.02 | 0 | Neutral |
| testneg_1258 | -0.07 | -0.38 | 2.393948 | 2.393948 | -1.223654 | -3 | 0.70819 | 0.5975 | 0.09353 | 0.12788 | 16.42913 | 0.15 | 0.2 | Neutral |
| testneg_1259 | -0.14 | -0.54 | 0.83037 | -3 | 0.329118 | 0.329118 | 0.16929 | 0.12788 | 0.51707 | 0.45685 | 9.01239 | 0.095436508 | 0 | Neutral |
| testneg_1260 | 0 | 0 | 0.269466 | -3 | 4.110069 | -3 | 0.33795 | 0.17455 | 0.24536 | 0.38555 | 10.41667 | 0.339663374 | 9.6 | Neutral |
| testneg_1261 | 0.04 | -0.23 | 2.128783 | -3 | 4.488254 | -3 | 0.08047 | 0.06746 | 0.19898 | 0.08855 | 32.22765 | 0.060355091 | 0 | Neutral |
| testneg_1262 | 0 | 0 | -0.536837 | -3 | 5.068794 | -3 | 0.02617 | 0.12788 | 0.04627 | 0.05223 | 49.12114 | 0.093333333 | 0 | Neutral |
| testneg_1263 | 0 | 0 | 0.191318 | -3 | 5.996034 | -3 | 0.96113 | 0.24974 | 0.08306 | 0.12788 | 19.09347 | 0.150082645 | 0 | Neutral |
| testneg_1264 | 0.05 | 0.29 | 1.287073 | -3 | 1.413393 | -3 | 0.06744 | 0.12788 | 0.2293 | 0.12788 | 19.23156 | 0.01 | 0 | Neutral |
| testneg_1265 | 0.01 | 0.8 | 4.324034 | -3 | -0.303923 | -0.303923 | 0.39015 | 0.23372 | 0.04593 | 0.09315 | 10.41667 | 0.092742424 | 0 | Neutral |
| testneg_1266 | 0.09 | -0.3 | 4.602859 | -3 | 2.028242 | -3 | 0.5065 | 0.20697 | 0.27058 | 0.13363 | 10.83333 | 0.348352381 | 6.8 | Neutral |
| testneg_1267 | -0.07 | -0.38 | -1.066068 | -3 | 1.6302 | -3 | 0.04515 | 0.14318 | 0.23468 | 0.10874 | 19.09347 | 0.06 | 0 | Neutral |
| testneg_1268 | -0.02 | 0.63 | 2.082622 | -3 | 1.139058 | -3 | 0.14335 | 0.14499 | 0.27887 | 0.18663 | 21.10002 | 0.110385185 | 0 | Neutral |
| testneg_1269 | 0 | -0.01 | 1.894323 | -3 | 1.76699 | -3 | 0.14482 | 0.08592 | 0.05027 | 0.10981 | 22.16924 | 0.010455038 | 0 | Neutral |
| testneg_1270 | -0.01 | -0.79 | 5.611956 | -3 | -0.361103 | -3 | 0.14187 | 0.1353 | 0.26982 | 0.12788 | 16.8018 | 0.15834595 | 0 | Neutral |
| testneg_1271 | 0.16 | -0.37 | 0.701088 | -3 | -0.384224 | -3 | 0.99538 | 0.40044 | 0.53698 | 0.10653 | 12.33727 | 0.08 | 0 | Neutral |
| testneg_1272 | 0.14 | -1.01 | -0.384327 | -3 | 0.076953 | -3 | 0.18507 | 0.12788 | 0.10187 | 0.12339 | 28.47222 | 0.020513158 | 0 | Neutral |
| testneg_1273 | 0 | 0 | 0.007168 | -3 | -0.677314 | -3 | 0.29356 | 0.47581 | 0.3783 | 0.08224 | 19.44444 | 0.03 | 0 | Neutral |
| testneg_1274 | 0 | 0 | -0.061342 | -3 | 1.736396 | -3 | 0.07626 | 0.12788 | 0.19898 | 0.46867 | 18.76159 | 0.056805556 | 0 | Neutral |
| testneg_1275 | -0.01 | -0.79 | 0.065789 | -3 | 5.09904 | -3 | 0.18233 | 0.14781 | 0.26621 | 0.10639 | 10.83333 | 0.142111111 | 0 | Neutral |
| testneg_1276 | 0 | 0 | -0.480488 | -3 | 1.902977 | 1.902977 | 0.11797 | 0.12788 | 0.18504 | 0.0822 | 18.88889 | 0.03016313 | 0 | Neutral |
| testneg_1277 | 0 | 0 | 1.759937 | -3 | 4.055487 | -3 | 0.24247 | 0.19222 | 0.10458 | 0.10499 | 33.04507 | 0.134448764 | 0 | Neutral |
| testneg_1278 | -0.04 | -2.01 | 2.438483 | -3 | 4.202971 | -3 | 0.31924 | 0.11145 | 0.53292 | 0.32174 | 19.60216 | 0.408802859 | 19 | Neutral |
| testneg_1279 | 0.09 | -0.3 | 2.387342 | -3 | 1.599689 | -3 | 0.07428 | 0.52467 | 0.17057 | 0.12788 | 10.83333 | 0.100837272 | 0 | Neutral |
| testneg_1280 | 0.07 | 0.38 | 0.559242 | -3 | 0.51352 | -3 | 0.87803 | 0.12788 | 0.09527 | 0.09424 | 39.45884 | 0.03 | 0 | Neutral |
| testneg_1281 | 0 | 0 | 0.483032 | -3 | 0.905929 | -3 | 0.39724 | 0.11058 | 0.28817 | 0.10531 | 8.75817 | 0 | 0 | Neutral |
| testneg_1282 | -0.19 | -0.51 | 2.183162 | -3 | 6.223377 | -3 | 0.22078 | 0.12788 | 0.143 | 0.11236 | 21.84055 | 0.210613354 | 0 | Neutral |
| testneg_1283 | 0 | 0 | 1.600017 | -3 | 0.853858 | -3 | 0.07994 | 0.1101 | 0.35513 | 0.21966 | 10.83333 | 0.020725641 | 0 | Neutral |
| testneg_1284 | 0 | 0 | 1.519976 | -3 | 6.381213 | -3 | 0.19898 | 0.12788 | 0.28156 | 0.11579 | 15.01941 | 0.1 | 0 | Neutral |
| testneg_1285 | 0 | 0 | 1.323079 | -3 | 0.660523 | -3 | 0.5103 | 0.10923 | 0.06897 | 0.08445 | 36.30782 | 0.011372913 | 0 | Neutral |
| testneg_1286 | 0.15 | 0.06 | 3.129189 | -3 | 4.489998 | -3 | 0.51137 | 0.19235 | 0.03378 | 0.07001 | 10.83333 | 0.270052632 | 1.2 | Neutral |
| testneg_1287 | 0.01 | 0.8 | -0.214756 | -3 | 4.76216 | -3 | 0.07428 | 0.07645 | 0.23126 | 0.16717 | 16.76869 | 0.245616939 | 0 | Neutral |
| testneg_1288 | -0.07 | -0.38 | 1.681972 | -3 | 1.474779 | 1.474779 | 0.4563 | 0.12971 | 0.0882 | 0.12788 | 20.89606 | 0.03 | 0 | Neutral |
| testneg_1289 | -0.15 | 0.28 | 3.6541 | -3 | -0.247762 | -3 | 0.19898 | 0.37148 | 0.44654 | 0.10907 | 11.36293 | 0.16 | 0 | Neutral |
| testneg_1290 | 0.05 | 0.29 | 1.381522 | -3 | 0.715358 | -3 | 0.11325 | 0.17021 | 0.74488 | 0.12788 | 13.73701 | 0.04026084 | 0 | Neutral |
| testneg_1291 | 0.21 | 0.15 | 1.912237 | -3 | 1.562114 | -3 | 0.12511 | 0.11897 | 0.78356 | 0.13377 | 34.70057 | 0.09 | 0 | Neutral |
| testneg_1292 | 0.07 | 0.7 | 5.438117 | -3 | 0.513578 | 0.513578 | 0.12463 | 0.12788 | 0.19898 | 0.07409 | 15.02472 | 0.1 | 0 | Neutral |
| testneg_1293 | 0.03 | 0.12 | 2.135113 | -3 | -2.304927 | -2.304927 | 0.40391 | 0.13431 | 0.02665 | 0.05082 | 20.27778 | 0.020074028 | 0 | Neutral |
| testneg_1294 | 0 | 0 | 1.887991 | -3 | -0.275748 | -3 | 0.18869 | 0.23984 | 0.13779 | 0.0945 | 20.91877 | 0.021676659 | 0 | Neutral |
| testneg_1295 | -0.21 | -0.15 | 4.120211 | -3 | 2.029753 | -3 | 0.44086 | 0.173 | 0.11006 | 0.20781 | 12.33727 | 0.388702564 | 15.2 | Neutral |
| testneg_1296 | 0 | 0 | 1.436812 | -3 | 1.791247 | -3 | 0.29302 | 0.17394 | 0.39193 | 0.11262 | 12.57218 | 0.002592968 | 0 | Neutral |
| testneg_1297 | -0.03 | -0.12 | -2.124954 | -3 | 4.46199 | -3 | 0.34153 | 0.08855 | 0.77364 | 0.12676 | 18.33333 | 0.082373985 | 0 | Neutral |
| testneg_1298 | 0.17 | 0.38 | 0.96383 | -3 | 5.495238 | 5.492392 | 0.13195 | 0.57267 | 0.08857 | 0.83499 | 10 | 0.76 | 100 | Disease_causing |
| testneg_1299 | 0.14 | -1.01 | 1.799629 | -3 | 1.291726 | -3 | 0.30085 | 0.18374 | 0.29525 | 0.12788 | 18.75 | 0.04 | 0 | Neutral |
| testneg_1300 | 0 | -0.01 | 5.0589 | -3 | 2.390968 | -3 | 0.09658 | 0.11286 | 0.07999 | 0.10413 | 25.15581 | 0.101041667 | 0 | Neutral |
| testneg_1301 | -0.08 | -0.03 | 1.50021 | -3 | 2.095355 | -3 | 0.27618 | 0.11809 | 0.46339 | 0.10423 | 14.27661 | 0.01 | 0 | Neutral |
| testneg_1302 | -0.05 | -0.29 | 1.33986 | -3 | 2.507265 | -3 | 0.19199 | 0.16616 | 0.98581 | 0.27035 | 12.34649 | 0.218422664 | 0.2 | Neutral |
| testneg_1303 | 0.02 | -0.63 | 4.472023 | -3 | -0.431211 | -3 | 0.11956 | 0.3237 | 0.36676 | 0.37749 | 12.29167 | 0.427621212 | 25.6 | Neutral |
| testneg_1304 | -0.03 | 1.03 | 5.582884 | -3 | 0.66227 | -3 | 0.34314 | 0.18668 | 0.23141 | 0.12788 | 16.84579 | 0.26 | 2.2 | Neutral |
| testneg_1305 | -0.18 | 1.05 | -0.907393 | -3 | 2.316264 | -3 | 0.05332 | 0.12788 | 0.29804 | 0.12768 | 13.51351 | 0.19 | 0 | Neutral |
| testneg_1306 | -0.07 | -0.38 | 3.673027 | -3 | 1.67748 | -3 | 0.19989 | 0.16777 | 0.42797 | 0.17974 | 20.30635 | 0.308541647 | 3.8 | Neutral |
| testneg_1307 | 0.16 | -1.37 | 3.503407 | -3 | 4.74013 | -3 | 0.68233 | 0.58998 | 0.48585 | 0.66989 | 10.3336 | 0.819910714 | 100 | Disease_causing |
| testneg_1308 | 0.14 | -0.37 | 6.450038 | -3 | 3.803348 | -3 | 0.53798 | 0.10163 | 0.67263 | 0.09781 | 18.88889 | 0.370064659 | 11.6 | Neutral |
| testneg_1309 | 0 | 0 | -0.164302 | -3 | 0.204096 | -3 | 0.35475 | 0.12788 | 0.08531 | 0.12788 | 18.33333 | 0.01 | 0 | Neutral |
| testneg_1310 | 0.09 | -0.3 | 5.235545 | -3 | 0.270884 | -3 | 0.09998 | 0.09022 | 0.24536 | 0.38555 | 8.84474 | 0.307100598 | 3.4 | Neutral |
| testneg_1311 | 0.00018182 | -0.2654545 | 12.863729 | -3 | 5.964649 | 1.801091 | 0.43918 | 0.46357 | 0.39459 | 0.12788 | 11.25 | 0.838453213 | 100 | Disease_causing |
| testneg_1312 | 0.07 | 0.33 | 1.074305 | -3 | 2.519144 | -3 | 0.25884 | 0.12788 | 0.1496 | 0.16692 | 13.51351 | 0.129340463 | 0 | Neutral |
| testneg_1313 | -0.03 | -0.1 | 3.646993 | -3 | 5.338598 | -3 | 0.7062 | 0.83002 | 0.22179 | 0.09831 | 4.72222 | 0.364260913 | 13 | Neutral |
| testneg_1314 | 0.2 | -1.26 | 6.373102 | -3 | 0.10268 | -3 | 0.05754 | 0.10881 | 0.34193 | 0.12788 | 10.22898 | 0.17 | 0 | Neutral |
| testneg_1315 | 0.04 | -0.23 | 3.107857 | -3 | -1.416622 | -3 | 0.11316 | 0.12788 | 0.30475 | 0.21064 | 10.25326 | 0.169083333 | 0.2 | Neutral |
| testneg_1316 | 0.00563636 | -0.1407273 | 2.08939 | -3 | -2.106562 | -2.106562 | 0.09633 | 0.12788 | 0.27083 | 0.14313 | 18.86401 | 0.052833333 | 0 | Neutral |
| testneg_1317 | -0.15 | -0.04 | 2.846922 | -3 | 5.315177 | -3 | 0.18081 | 0.1258 | 0.12091 | 0.13585 | 10.41667 | 0.499302975 | 53.6 | Disease_causing |
| testneg_1318 | 0 | 0 | 1.550971 | -3 | 1.695811 | -3 | 0.73311 | 0.10866 | 0.10371 | 0.08655 | 10.22898 | 0.01 | 0 | Neutral |
| testneg_1319 | 0 | 0 | 0.764227 | -3 | 1.591111 | -3 | 0.25025 | 0.18924 | 0.07015 | 0.0889 | 24.2629 | 0.02 | 0 | Neutral |
| testneg_1320 | 0.17 | 0.38 | 1.603588 | -3 | 3.269216 | -3 | 0.10496 | 0.21491 | 0.2734 | 0.2578 | 11.25 | 0.478546794 | 46.6 | Neutral |
| testneg_1321 | 0 | 0 | 0.006704 | -3 | 2.716468 | -3 | 0.30925 | 0.10335 | 0.28066 | 0.18241 | 14.85392 | 0.06951756 | 0 | Neutral |
| testneg_1322 | -0.08 | -0.03 | 3.577743 | 2.147578 | 6.451231 | -3 | 0.47335 | 0.12788 | 0.41383 | 0.119 | 5.69444 | 0.462635468 | 40.2 | Neutral |
| testneg_1323 | 0 | 0 | -0.155612 | -3 | 1.695225 | -3 | 0.07141 | 0.12788 | 0.33294 | 0.16155 | 19.37853 | 0.07 | 0 | Neutral |
| testneg_1324 | -0.02 | 0.28 | 2.201231 | -3 | -2.628134 | -3 | 0.20231 | 0.12788 | 0.26971 | 0.08509 | 33.12684 | 0.01 | 0 | Neutral |
| testneg_1325 | 0.01 | 0.79 | 3.635653 | -3 | -2.342961 | -3 | 0.11501 | 0.13748 | 0.1533 | 0.13168 | 18.33333 | 0.113666667 | 0 | Neutral |
| testneg_1326 | 0 | 0 | -0.100514 | -3 | 3.652682 | -3 | 0.16469 | 0.16042 | 0.31238 | 0.14393 | 26.80135 | 0.292331598 | 3 | Neutral |
| testneg_1327 | -0.02 | 0.63 | 4.67013 | -3 | 7.527814 | -3 | 0.10004 | 0.12788 | 0.84596 | 0.10633 | 14.30976 | 0.509375 | 55.4 | Disease_causing |
| testneg_1328 | -0.21 | -0.15 | -0.470437 | -3 | -0.454393 | -0.454393 | 0.25904 | 0.16134 | 0.03443 | 0.1032 | 20.83333 | 0.08 | 0 | Neutral |
| testneg_1329 | -0.01 | -0.79 | 5.611956 | -3 | 3.332054 | -3 | 0.14187 | 0.1353 | 0.12051 | 0.08503 | 20.41417 | 0.29 | 1.6 | Neutral |
| testneg_1330 | 0 | 0 | 1.397791 | -3 | 0.96317 | -3 | 0.70872 | 0.09884 | 0.20379 | 0.35591 | 12.30072 | 0.030930146 | 0 | Neutral |
| testneg_1331 | 0 | 0 | 1.687932 | -3 | 0.994999 | 0.994999 | 0.16299 | 0.10454 | 0.51282 | 0.12075 | 21.3735 | 0.01 | 0 | Neutral |
| testneg_1332 | 0.02 | -0.68 | 1.520121 | -3 | 1.158382 | -3 | 0.27187 | 0.41985 | 0.19087 | 0.44337 | 10 | 0.164297784 | 0 | Neutral |
| testneg_1333 | -0.02 | 0.63 | 3.191277 | -3 | 3.266081 | -3 | 0.37074 | 0.11532 | 0.09556 | 0.10109 | 34.85049 | 0.119384164 | 0 | Neutral |
| testneg_1334 | 0 | 0 | 1.519976 | -3 | 7.038327 | -3 | 0.19898 | 0.12788 | 0.30367 | 0.41964 | 5.19399 | 0.426067308 | 28.8 | Neutral |
| testneg_1335 | 0.07 | -0.31 | 3.295096 | -3 | 1.751331 | -3 | 0.89959 | 0.55078 | 0.06647 | 0.09129 | 4.72222 | 0.097291667 | 0 | Neutral |
| testneg_1336 | 0 | 0 | 0.930011 | -3 | 3.213612 | -3 | 0.12786 | 0.72066 | 0.26323 | 0.2399 | 10 | 0.298359649 | 2.6 | Neutral |
| testneg_1337 | 0.02 | 0.13 | 2.814701 | -3 | 0.234126 | -3 | 0.96842 | 0.23432 | 0.54792 | 0.54626 | 10 | 0.19290497 | 0 | Neutral |
| testneg_1338 | -0.14 | -0.54 | 0.812392 | -3 | 2.359354 | -3 | 0.19898 | 0.10512 | 0.35908 | 0.18099 | 19.84985 | 0.135643939 | 0 | Neutral |
| testneg_1339 | 0 | 0 | 1.861867 | -3 | 3.328368 | -3 | 0.32963 | 0.1035 | 0.14731 | 0.12788 | 19.50564 | 0.03 | 0 | Neutral |
| testneg_1340 | 0 | -0.01 | 3.197244 | -3 | 4.216279 | -3 | 0.09611 | 0.07772 | 0.41089 | 0.1897 | 10.22898 | 0.393501832 | 17.4 | Neutral |
| testneg_1341 | 0 | 0 | 1.235264 | -3 | 2.425778 | -3 | 0.56904 | 0.14839 | 0.11613 | 0.1109 | 18.88889 | 0.030339862 | 0 | Neutral |

| | | | | | | | | | | | | | |
|---|---|---|---|---|---|---|---|---|---|---|---|---|---|
| testneg_1342 | 0 | 0 | 0.205888 | -3 | 5.479934 | 0.979716 | 0.08922 | 0.10539 | 0.39932 | 0.10693 | 10.11674 | 0.09 | 0 Neutral |
| testneg_1343 | 0.02 | 0.45 | 5.389515 | -3 | 3.479077 | -3 | 0.19898 | 0.12788 | 0.17487 | 0.1672 | 26.46332 | 0.578569264 | 82 Disease_causing |
| testneg_1344 | 0 | 0 | 1.75727 | -3 | 2.12697 | -3 | 0.16797 | 0.11809 | 0.16053 | 0.15049 | 19.96647 | 0.05 | 0 Neutral |
| testneg_1345 | 0.04 | -0.23 | 3.24838 | -3 | 5.587644 | -3 | 0.68055 | 0.12788 | 0.6185 | 0.09316 | 19.92207 | 0.190238095 | 0 Neutral |
| testneg_1346 | 0 | 0 | 0.119948 | -3 | 3.26904 | -3 | 0.14632 | 0.12662 | 0.18554 | 0.12299 | 8.97898 | 0.020648457 | 0 Neutral |
| testneg_1347 | 0.17 | 0.38 | 4.078311 | -3 | 1.290794 | -3 | 0.20951 | 0.16278 | 0.84365 | 0.12788 | 10.83333 | 0.121210195 | 0 Neutral |
| testneg_1348 | 0.08 | 0.03 | -1.290562 | -3 | 2.347933 | -3 | 0.19898 | 0.12895 | 0.43158 | 0.10472 | 18.88889 | 0.04 | 0 Neutral |
| testneg_1349 | 0.02 | -0.63 | 1.81834 | -3 | 0.744952 | -3 | 0.18096 | 0.12788 | 0.38279 | 0.10578 | 45.26721 | 0.01 | 0 Neutral |
| testneg_1350 | 0 | 0 | 0.902021 | -3 | 0.543748 | -3 | 0.37499 | 0.17803 | 0.34153 | 0.08855 | 11.25 | 0.01 | 0 Neutral |
| testneg_1351 | -0.03 | -0.12 | 1.782773 | -3 | 7.351473 | -3 | 0.42544 | 0.11809 | 0.17155 | 0.18585 | 16.84579 | 0.495943096 | 52.4 Disease_causing |
| testneg_1352 | 0 | 0 | 0.699953 | -3 | 1.1308 | -3 | 0.54231 | 0.15254 | 0.19898 | 0.11185 | 19.03739 | 0.010417391 | 0 Neutral |
| testneg_1353 | 0 | 0 | 2.434662 | 0.418118 | 4.106182 | -3 | 0.19898 | 0.12788 | 0.06394 | 0.24464 | 10 | 0.380014104 | 14.4 Neutral |
| testneg_1354 | -0.04 | 0.23 | -0.28312 | -3 | 1.083169 | -3 | 0.06513 | 0.12353 | 0.26621 | 0.10639 | 45.41667 | 0.02 | 0 Neutral |
| testneg_1355 | 0.02 | 0.45 | 0.245416 | -3 | 1.715304 | -3 | 0.10004 | 0.12788 | 0.07447 | 0.23492 | 9.36989 | 0.060370879 | 0 Neutral |
| testneg_1356 | -0.0087273 | 0.00581818 | 10.530907 | -3 | 3.002533 | -3 | 0.29024 | 0.1472 | 0.14304 | 0.10715 | 51.41534 | 0.56 | 79.6 Disease_causing |
| testneg_1357 | 0 | -0.01 | -0.496432 | -3 | 2.531153 | -3 | 0.06247 | 0.14184 | 0.36833 | 0.10433 | 5.74956 | 0.04 | 0 Neutral |
| testneg_1358 | -0.09 | 0.3 | -0.355202 | -3 | 6.080162 | -3 | 0.21802 | 0.10187 | 0.50448 | 0.50327 | 6.94453 | 0.402387489 | 20.6 Neutral |
| testneg_1359 | -0.03 | -0.1 | 0.224925 | -3 | 5.998771 | -3 | 0.03676 | 0.12788 | 0.61231 | 0.13362 | 20 | 0.349464286 | 5.4 Neutral |
| testneg_1360 | 0.14 | -0.69 | 5.461184 | -3 | 6.151341 | -3 | 0.29145 | 0.11884 | 0.47733 | 0.08388 | 34.76048 | 0.330634058 | 7.4 Neutral |
| testneg_1361 | 0 | 0 | 0.600565 | -3 | 0.892238 | -3 | 0.99564 | 0.25794 | 0.20655 | 0.15317 | 13.3392 | 0.07 | 0 Neutral |
| testneg_1362 | -0.03 | -0.1 | 0.904827 | -3 | 0.567562 | -3 | 0.19898 | 0.12788 | 0.17046 | 0.21736 | 18.76882 | 0.052481974 | 0 Neutral |
| testneg_1363 | 0.02 | 0.45 | 5.697436 | -3 | -0.296252 | -3 | 0.0832 | 0.09323 | 0.09323 | 0.16732 | 44.89949 | 0.350637446 | 6 Neutral |
| testneg_1364 | 0 | -0.32 | 1.285139 | -3 | -0.361103 | -3 | 0.54895 | 0.15082 | 0.26982 | 0.12788 | 10.80591 | 0.02 | 0 Neutral |
| testneg_1365 | -0.03 | -0.1 | 2.445411 | -3 | 6.489682 | 0.0552 | 0.22179 | 0.09831 | 0.13532 | 0.19904 | 18.88889 | 0.52901379 | 65.8 Disease_causing |
| testneg_1366 | 0.22 | -2.66 | 1.967583 | -3 | 4.102055 | -3 | 0.09749 | 0.82924 | 0.24823 | 0.12788 | 35.99537 | 0.34 | 1.6 Neutral |
| testneg_1367 | -0.04 | 0.23 | 2.272008 | -3 | -0.577508 | -3 | 0.23913 | 0.12788 | 0.1 | 0.22633 | 10.30286 | 0.083301282 | 0 Neutral |
| testneg_1368 | 0.04 | -0.23 | 2.504423 | -3 | 1.349086 | -3 | 0.14482 | 0.08592 | 0.17199 | 0.10586 | 20.78035 | 0.01 | 0 Neutral |
| testneg_1369 | 0 | 0 | -0.200482 | -3 | 2.927777 | -3 | 0.3889 | 0.10831 | 0.65564 | 0.11468 | 12.52798 | 0.020196154 | 0 Neutral |
| testneg_1370 | 0.16 | -0.37 | 2.851691 | -3 | 7.325251 | -3 | 0.21623 | 0.42074 | 0.16601 | 0.14015 | 24.09297 | 0.797566964 | 100 Disease_causing |
| testneg_1371 | -0.03 | -0.1 | 1.820578 | -3 | 0.211586 | 0.211586 | 0.31507 | 0.61057 | 0.1119 | 0.24875 | 9.33732 | 0.22110101 | 0.4 Neutral |
| testneg_1372 | 0 | 0 | 0.67738 | -3 | -1.532457 | -3 | 0.97678 | 0.09885 | 0.53784 | 0.5574 | 5 | 0.044418215 | 0 Neutral |
| testneg_1373 | 0.17 | 0.38 | 2.311167 | -3 | 0.829195 | -3 | 0.05413 | 0.05691 | 0.68423 | 0.08163 | 17.21847 | 0.07 | 0 Neutral |
| testneg_1374 | 0 | 0 | -0.757789 | -0.757789 | 5.102298 | -3 | 0.40491 | 0.17596 | 0.10213 | 0.17266 | 12.69104 | 0.411302309 | 22.6 Neutral |
| testneg_1375 | 0 | -0.32 | 2.54814 | -3 | 3.263788 | -3 | 0.85592 | 0.20297 | 0.19898 | 0.12788 | 18.33333 | 0.114343434 | 0 Neutral |
| testneg_1376 | 0.09 | -0.67 | 2.624257 | -3 | 0.049803 | -3 | 0.19043 | 0.07855 | 0.21726 | 0.23214 | 20.41417 | 0.092198077 | 0 Neutral |
| testneg_1377 | 0 | 0 | 0.229773 | -3 | 1.624942 | 1.624942 | 0.13275 | 0.12788 | 0.16208 | 0.10597 | 20.77313 | 0.02 | 0 Neutral |
| testneg_1378 | -0.02 | 0.28 | 1.683599 | -3 | 6.687775 | -3 | 0.3047 | 0.11929 | 0.07092 | 0.20005 | 10.83333 | 0.479587302 | 46.6 Neutral |
| testneg_1379 | 0.08 | 0.03 | -1.360462 | -3 | 3.381638 | -3 | 0.28537 | 0.12788 | 0.07552 | 0.12788 | 13.02635 | 0.091213235 | 0 Neutral |
| testneg_1380 | 0 | 0 | 0.447219 | -3 | 1.902977 | 1.902977 | 0.13884 | 0.12788 | 0.18504 | 0.0822 | 10.83333 | 0.030116112 | 0 Neutral |
| testneg_1381 | 0 | 0 | -0.445161 | -0.445161 | 2.374684 | -3 | 0.09728 | 0.12788 | 0.33314 | 0.47999 | 39.09389 | 0.123210784 | 0 Neutral |
| testneg_1382 | 0.06 | -0.57 | 1.240379 | -3 | -1.873728 | -1.873728 | 0.64591 | 0.38907 | 0.19898 | 0.12788 | 12.29167 | 0.03 | 0 Neutral |
| testneg_1383 | -0.04 | -2.01 | 5.42516 | -3 | 0.182103 | -3 | 0.2141 | 0.19929 | 0.16355 | 0.20781 | 10 | 0.55 | 74.8 Disease_causing |
| testneg_1384 | 0 | 0 | 1.302273 | -3 | 1.729145 | -3 | 0.1947 | 0.11413 | 0.13575 | 0.12788 | 14.02665 | 0 | 0 Neutral |
| testneg_1385 | -0.03 | -0.1 | 3.149921 | -3 | -2.242853 | -3 | 0.29357 | 0.79524 | 0.06358 | 0.5364 | 13.86017 | 0.369953271 | 9.2 Neutral |
| testneg_1386 | -0.03 | -1.22 | 3.62024 | -3 | -0.274976 | -3 | 0.52388 | 0.25831 | 0.26767 | 0.25506 | 12.57784 | 0.307441378 | 2.2 Neutral |
| testneg_1387 | -0.02 | -0.45 | 2.925716 | -3 | 1.074461 | -3 | 0.13906 | 0.09925 | 0.14379 | 0.12788 | 17.46936 | 0.020165301 | 0 Neutral |
| testneg_1388 | 0 | 0 | 1.317673 | -3 | 0.925856 | -3 | 0.45415 | 0.12971 | 0.08628 | 0.11498 | 19.89554 | 0.01 | 0 Neutral |
| testneg_1389 | 0.07 | -0.31 | 1.028884 | -3 | 5.687452 | 3.618842 | 0.93468 | 0.12788 | 0.30241 | 0.104 | 10.41667 | 0.25 | 0.2 Neutral |
| testneg_1390 | -0.17 | -0.38 | 0.941167 | -3 | 3.048051 | -3 | 0.32299 | 0.12788 | 0.07631 | 0.12788 | 28.82035 | 0.090481061 | 0 Neutral |
| testneg_1391 | -0.03 | -0.1 | -0.039157 | -3 | 2.477618 | -3 | 0.04606 | 0.20522 | 0.44103 | 0.10698 | 33.16257 | 0.14 | 0 Neutral |
| testneg_1392 | -0.07 | -0.38 | 3.802792 | -3 | 2.542698 | 1.556018 | 0.71844 | 0.20204 | 0.11748 | 0.12788 | 10.66104 | 0.181404762 | 0 Neutral |
| testneg_1393 | 0.09 | -0.3 | 5.344639 | -3 | 1.491454 | -3 | 0.6992 | 0.1804 | 0.1078 | 0.17394 | 10.41667 | 0.542083333 | 70.8 Disease_causing |
| testneg_1394 | -0.15 | 1.06 | 3.713115 | -3 | -0.032364 | -3 | 0.46919 | 0.60081 | 0.16524 | 0.12788 | 13.75 | 0.208 | 0 Neutral |
| testneg_1395 | -0.03 | -1.22 | 4.490043 | -3 | 6.87277 | -3 | 0.7342 | 0.11023 | 0.09724 | 0.09172 | 12.03949 | 0.41 | 23 Neutral |
| testneg_1396 | 0 | 0 | -0.502553 | -3 | 1.190149 | -3 | 0.94822 | 0.14005 | 0.16841 | 0.11143 | 22.80186 | 0.030763376 | 0 Neutral |
| testneg_1397 | 0.16 | -1.37 | -0.103493 | -3 | 1.112437 | -3 | 0.12835 | 0.15143 | 0.62172 | 0.18135 | 8.44735 | 0.123095238 | 0 Neutral |
| testneg_1398 | -0.05 | -0.29 | 1.139387 | -3 | 2.849519 | -3 | 0.62197 | 0.12788 | 0.14486 | 0.33725 | 11.69956 | 0.140637161 | 0 Neutral |
| testneg_1399 | -0.04 | 0.23 | 1.835412 | -3 | 0.031448 | -3 | 0.10799 | 0.10754 | 0.3783 | 0.08224 | 17.21847 | 0.01 | 0 Neutral |
| testneg_1400 | 0.07 | -0.31 | 2.407105 | -3 | -1.763617 | -3 | 0.21095 | 0.10574 | 0.19433 | 0.12788 | 18.6768 | 0.01 | 0 Neutral |
| testneg_1401 | 0 | 0 | 1.172534 | -3 | -1.63682 | -3 | 0.04056 | 0.55024 | 0.13505 | 0.09019 | 16.48772 | 0.11 | 0 Neutral |
| testneg_1402 | 0.15 | -0.28 | -0.255381 | -3 | 0.459406 | -3 | 0.0917 | 0.13124 | 0.03981 | 0.09432 | 29.9024 | 0.03 | 0 Neutral |
| testneg_1403 | 0.02 | 0.45 | 5.488128 | -3 | 0.219231 | -3 | 0.99998 | 0.9588 | 0.68423 | 0.08163 | 16.69775 | 0.33 | 1.8 Neutral |
| testneg_1404 | -0.01 | -0.79 | -3.816017 | -3 | 1.418333 | -3 | 0.06095 | 0.08179 | 0.72235 | 0.14785 | 6.94444 | 0.07 | 0 Neutral |
| testneg_1405 | 0.15 | -0.28 | 1.45686 | -3 | 5.03801 | -3 | 0.38552 | 0.14172 | 0.18964 | 0.34036 | 10.41667 | 0.403439874 | 20.8 Neutral |
| testneg_1406 | 0.16 | -0.37 | 4.518381 | -3 | 0.608612 | -3 | 0.21201 | 0.20962 | 0.09259 | 0.11165 | 12.38369 | 0.08 | 0 Neutral |
| testneg_1407 | -0.09 | 0.3 | -3.629472 | -3 | 4.387695 | -3 | 0.15612 | 0.11465 | 0.18341 | 0.13901 | 30.06401 | 0.270178571 | 0.4 Neutral |
| testneg_1408 | -0.09 | 0.67 | 2.302703 | -3 | 0.80567 | -3 | 0.19898 | 0.11185 | 0.94531 | 0.91608 | 33.06212 | 0.109166667 | 0 Neutral |
| testneg_1409 | 0.04 | -0.23 | 1.625356 | -3 | 2.327743 | -3 | 0.5065 | 0.20697 | 0.0538 | 0.0823 | 10.41667 | 0.07 | 0 Neutral |
| testneg_1410 | 0.08 | 0.03 | 1.884645 | -3 | 1.614723 | 1.614723 | 0.10285 | 0.11721 | 0.74944 | 0.1263 | 5 | 0.03 | 0 Neutral |
| testneg_1411 | -0.0036364 | 0.11719697 | 0.781229 | 0.781229 | 1.87595 | -3 | 0.07321 | 0.12788 | 0.39883 | 0.12346 | 36.4913 | 0.04 | 0 Neutral |
| testneg_1412 | 0.04 | -1.61 | 3.074697 | -3 | 1.137955 | -3 | 0.37282 | 0.07943 | 0.16462 | 0.18467 | 22.92553 | 0.18 | 0 Neutral |
| testneg_1413 | -0.05 | -0.29 | 5.703118 | -3 | 4.00772 | 0.58285 | 0.61491 | 0.11954 | 0.48825 | 0.10951 | 19.73311 | 0.35 | 7.4 Neutral |
| testneg_1414 | 0 | 0 | 0.046064 | -3 | -0.034875 | -3 | 0.2543 | 0.15588 | 0.2547 | 0.07589 | 10 | 0.01 | 0 Neutral |
| testneg_1415 | 0 | 0 | 1.56019 | -3 | 0.012846 | -3 | 0.25487 | 0.13098 | 0.07995 | 0.12788 | 16.01716 | 0.01 | 0 Neutral |
| testneg_1416 | 0.07 | 0.38 | 2.43609 | -3 | 4.172643 | -3 | 0.02146 | 0.10351 | 0.10893 | 0.12072 | 20 | 0.110579832 | 0 Neutral |
| testneg_1417 | 0.06 | -0.57 | 5.777528 | -3 | 0.389489 | -3 | 0.17672 | 0.14229 | 0.11858 | 0.11551 | 19.09347 | 0.17 | 0 Neutral |
| testneg_1418 | -0.03 | -0.1 | 2.059294 | 2.059294 | 7.504294 | -3 | 0.66711 | 0.31094 | 0.0968 | 0.15592 | 21.85185 | 0.67 | 97.4 Disease_causing |
| testneg_1419 | -0.05 | -0.29 | -0.242457 | -0.242457 | 5.778555 | -3 | 0.48662 | 0.23781 | 0.16636 | 0.12384 | 13.51351 | 0.24 | 0 Neutral |
| testneg_1420 | 0.21 | 0.15 | 4.730594 | -3 | 2.970766 | -3 | 0.15192 | 0.14302 | 0.10285 | 0.10886 | 20 | 0.19 | 0 Neutral |
| testneg_1421 | 0 | 0 | 0.655162 | -3 | 2.996727 | -3 | 0.26756 | 0.15588 | 0.19898 | 0.17518 | 23.35313 | 0.217122111 | 0 Neutral |
| testneg_1422 | 0 | 0 | 0.137649 | -3 | -0.079539 | -3 | 0.19898 | 0.12788 | 0.47816 | 0.66526 | 16.98432 | 0.04031592 | 0 Neutral |
| testneg_1423 | 0.08 | 0.03 | -0.431527 | -3 | -0.244679 | -3 | 0.19772 | 0.09261 | 0.16381 | 0.09851 | 18.33032 | 0.02 | 0 Neutral |
| testneg_1424 | 0.02 | 0.45 | 3.732818 | -3 | -0.415118 | -3 | 0.59321 | 0.59981 | 0.1144 | 0.12788 | 18.56611 | 0.121534091 | 0 Neutral |
| testneg_1425 | 0 | 0 | -0.125656 | -3 | 5.486225 | -3 | 0.08213 | 0.08913 | 0.10963 | 0.12788 | 19.09347 | 0.09 | 0 Neutral |
| testneg_1426 | -0.0022727 | -0.0332576 | 2.427467 | -3 | 0.416226 | -3 | 0.43866 | 0.12121 | 0.32111 | 0.12788 | 25.84011 | 0.01 | 0 Neutral |
| testneg_1427 | 0 | 0 | 1.836976 | -3 | 6.015888 | -3 | 0.84491 | 0.30945 | 0.11237 | 0.08916 | 16.88659 | 0.147267512 | 0 Neutral |
| testneg_1428 | 0 | 0 | 0.614162 | -3 | 4.668523 | -3 | 0.54211 | 0.12592 | 0.10565 | 0.08739 | 12.52798 | 0.065833333 | 0 Neutral |
| testneg_1429 | -0.19 | -0.51 | 1.521336 | -3 | 0.037077 | 0.037077 | 0.39553 | 0.11991 | 0.09004 | 0.07829 | 12.67985 | 0.040374126 | 0 Neutral |
| testneg_1430 | -0.01 | -0.79 | -1.10605 | -3 | -0.720759 | -0.720759 | 0.23401 | 0.12374 | 0.17758 | 0.15983 | 16.13039 | 0.070348485 | 0 Neutral |
| testneg_1431 | 0 | -0.32 | 4.72762 | -3 | 1.366318 | -3 | 0.21376 | 0.12708 | 0.78536 | 0.12709 | 4.72222 | 0.07 | 0 Neutral |
| testneg_1432 | 0.01 | -0.57 | 6.141111 | -3 | -0.180006 | -3 | 0.21983 | 0.10146 | 0.09806 | 0.12788 | 27.46106 | 0.148544974 | 0 Neutral |
| testneg_1433 | 0 | -0.01 | 1.061542 | -3 | -0.844526 | -3 | 0.15623 | 0.1447 | 0.28128 | 0.12883 | 17.21847 | 0.00021055 | 0 Neutral |
| testneg_1434 | -0.07 | -0.38 | 2.28341 | -3 | 0.097356 | -3 | 0.27723 | 0.13229 | 0.14485 | 0.16945 | 12.46106 | 0.091676828 | 0 Neutral |
| testneg_1435 | 0 | 0 | 1.181844 | -3 | 3.423277 | -3 | 0.94822 | 0.14005 | 0.13322 | 0.11297 | 38.19876 | 0.071984197 | 0 Neutral |
| testneg_1436 | 0.04 | -0.89 | 6.423333 | -3 | 1.902977 | 1.902977 | 0.16962 | 0.12788 | 0.18504 | 0.0822 | 10.41667 | 0.28 | 1.4 Neutral |
| testneg_1437 | -0.09 | 0.3 | 0.912537 | -3 | -0.262132 | -0.262132 | 0.2049 | 0.12678 | 0.11788 | 0.12788 | 17.21847 | 0.02 | 0 Neutral |
| testneg_1438 | 0.08 | 0.03 | -0.61726 | -0.61726 | 1.01324 | -3 | 0.20673 | 0.13236 | 0.45354 | 0.09957 | 10.23506 | 0.020513158 | 0 Neutral |
| testneg_1439 | -0.08 | -0.03 | 1.562114 | -3 | 0.148971 | -3 | 0.78356 | 0.13377 | 0.17007 | 0.12771 | 33.81526 | 0.024583333 | 0 Neutral |
| testneg_1440 | 0 | 0 | 1.54969 | -3 | -2.286968 | -3 | 0.46723 | 0.08832 | 0.17039 | 0.12788 | 10.11902 | 0.01 | 0 Neutral |
| testneg_1441 | 0 | -0.01 | 2.040282 | -3 | 6.292417 | -3 | 0.08437 | 0.97198 | 0.16663 | 0.12788 | 11.875 | 0.337983229 | 4.6 Neutral |
| testneg_1442 | 0 | 0 | 0.477383 | -3 | 4.912842 | -3 | 0.18431 | 0.13863 | 0.42899 | 0.31045 | 12.70833 | 0.281316502 | 4.2 Neutral |
| testneg_1443 | -0.14 | 0.37 | 4.115885 | -3 | 0.992466 | -3 | 0.09356 | 0.2448 | 0.17697 | 0.12974 | 17.33607 | 0.16 | 0 Neutral |
| testneg_1444 | 0.11 | -0.19 | 3.662322 | -3 | 2.15326 | -3 | 0.49745 | 0.1103 | 0.13029 | 0.12788 | 8.97898 | 0.069192589 | 0 Neutral |
| testneg_1445 | -0.08 | -0.03 | 2.222857 | -3 | 0.223637 | -3 | 0.19898 | 0.11185 | 0.12784 | 0.11244 | 35.26269 | 0.01 | 0 Neutral |
| testneg_1446 | -0.04 | -2.01 | 0.355171 | -3 | 0.649476 | -3 | 0.19898 | 0.12788 | 0.08184 | 0.10206 | 16.76768 | 0.03 | 0 Neutral |
| testneg_1447 | -0.07 | -0.7 | 1.388133 | -3 | 0.942465 | -3 | 0.35142 | 0.18303 | 0.24536 | 0.38555 | 10.41667 | 0.110678161 | 0 Neutral |
| testneg_1448 | -0.01 | -0.8 | 1.792644 | -3 | -0.55851 | -3 | 0.19898 | 0.12788 | 0.06874 | 0.12788 | 10.41667 | 0.01 | 0 Neutral |
| testneg_1449 | 0 | 0 | 1.844569 | -3 | -0.614161 | -0.614161 | 0.14735 | 0.12788 | 0.17796 | 0.09522 | 10.41667 | 0.01 | 0 Neutral |
| testneg_1450 | 0.02 | 0.45 | 6.053701 | -3 | 5.514783 | -3 | 0.43918 | 0.46357 | 0.13173 | 0.12788 | 15.51141 | 0.732980991 | 99.4 Disease_causing |
| testneg_1451 | 0.21 | 0.15 | 7.974494 | -3 | 0.242998 | -3 | 0.11316 | 0.12788 | 0.16078 | 0.39498 | 10.22898 | 0.44 | 30.2 Neutral |
| testneg_1452 | 0 | 0 | 1.128119 | -3 | -0.501932 | -3 | 0.83074 | 0.14769 | 0.64914 | 0.12481 | 10.41667 | 0.01 | 0 Neutral |
| testneg_1453 | 0 | 0 | 1.68348 | -3 | 0.211324 | -3 | 0.78286 | 0.19831 | 0.50063 | 0.154 | 10.83333 | 0.044017119 | 0 Neutral |
| testneg_1454 | 0.14 | -1.01 | 1.878171 | -3 | -0.456172 | -3 | 0.13011 | 0.14157 | 0.19898 | 0.1054 | 10.22898 | 0.02 | 0 Neutral |
| testneg_1455 | 0 | 0 | 0.097356 | -3 | -0.907467 | -3 | 0.14485 | 0.16945 | 0.61098 | 0.11179 | 28.07205 | 0.021483516 | 0 Neutral |
| testneg_1456 | 0 | 0 | 0.116062 | -3 | -0.298555 | -0.298555 | 0.4811 | 0.13398 | 0.10314 | 0.12173 | 10.83333 | 0.022390845 | 0 Neutral |
| testneg_1457 | 0.17 | 0.38 | 1.490909 | -3 | 3.28103 | -3 | 0.90259 | 0.31502 | 0.27077 | 0.10337 | 18.88889 | 0.18511646 | 0 Neutral |
| testneg_1458 | 0 | 0 | 0.740665 | -3 | 0.27289 | -3 | 0.23278 | 0.11027 | 0.14668 | 0.25103 | 12.37434 | 0.020431494 | 0 Neutral |
| testneg_1459 | 0 | 0 | 1.367025 | -3 | 0.662178 | -3 | 0.34467 | 0.13076 | 0.82513 | 0.10875 | 27.68948 | 0.01 | 0 Neutral |
| testneg_1460 | 0.05 | 0.29 | 2.987197 | -3 | 3.181816 | -3 | 0.88414 | 0.13991 | 0.08529 | 0.12376 | 19.9164 | 0.159440476 | 0 Neutral |
| testneg_1461 | 0 | 0 | 1.730475 | -3 | 2.738014 | -3 | 0.73601 | 0.26791 | 0.09987 | 0.09248 | 20.74751 | 0.058902047 | 0 Neutral |
| testneg_1462 | 0 | 0 | 2.044018 | -3 | 0.167824 | -3 | 0.63431 | 0.09837 | 0.11264 | 0.11163 | 21.43894 | 0.013666667 | 0 Neutral |
| testneg_1463 | 0.18 | -1.05 | 5.2613 | -3 | 0.146321 | -0.03734 | 0.13696 | 0.09566 | 0.11827 | 0.23034 | 20.98039 | 0.304298835 | 2.4 Neutral |

| | | | | | | | | | | | | | | |
|---|---|---|---|---|---|---|---|---|---|---|---|---|---|---|
| testneg_1464 | 0.05 | -0.81 | 1.356695 | -3 | 0.83151 | -3 | 0.12455 | 0.0998 | 0.16485 | 0.13045 | 13.51351 | 0.01 | 0 | Neutral |
| testneg_1465 | 0 | 0 | 0.388922 | -3 | 5.393665 | 5.393665 | 0.98107 | 0.70722 | 0.03809 | 0.16044 | 10.41667 | 0.466744987 | 41.6 | Neutral |
| testneg_1466 | 0.04 | 2.01 | 1.240851 | -3 | 0.857156 | -3 | 0.23859 | 0.1202 | 0.50063 | 0.154 | 10.31609 | 0.128523284 | 0 | Neutral |
| testneg_1467 | -0.06 | 0.57 | 0.709477 | -3 | 0.1877 | -3 | 0.19898 | 0.12788 | 0.76138 | 0.16258 | 24.50725 | 0.070310993 | 0 | Neutral |
| testneg_1468 | 0.15 | -0.28 | 0.411846 | -3 | 7.264005 | 7.264005 | 0.10133 | 0.32155 | 0.1856 | 0.12788 | 11.875 | 0.72 | 100 | Disease_causing |
| testneg_1469 | 0.17 | -1.22 | 5.736168 | -3 | 0.71504 | -3 | 0.33245 | 0.45921 | 0.21894 | 0.09041 | 22.27418 | 0.24 | 0.4 | Neutral |
| testneg_1470 | -0.05 | -0.29 | -2.166097 | -3 | 1.123147 | -3 | 0.41269 | 0.19135 | 0.59947 | 0.14339 | 19.44444 | 0.1 | 0 | Neutral |
| testneg_1471 | 0 | 0 | 0.006016 | -3 | 0.665428 | -3 | 0.43459 | 0.12595 | 0.20546 | 0.10734 | 16.8018 | 0.000335804 | 0 | Neutral |
| testneg_1472 | 0 | 0 | 1.840893 | -3 | -0.092248 | -0.092248 | 0.18465 | 0.11262 | 0.09445 | 0.12788 | 18.2557 | 0.01 | 0 | Neutral |
| testneg_1473 | 0 | 0 | 1.394643 | -3 | -0.347591 | -3 | 0.49629 | 0.11989 | 0.32094 | 0.36254 | 19.11765 | 0.051344462 | 0 | Neutral |
| testneg_1474 | 0 | 0 | 1.860726 | -3 | 1.183533 | -3 | 0.5901 | 0.19444 | 0.11042 | 0.09881 | 21.53664 | 0.021417617 | 0 | Neutral |
| testneg_1475 | 0 | 0 | 1.383368 | -3 | 7.405 | -3 | 0.19898 | 0.13057 | 0.21288 | 0.10049 | 18.33333 | 0.28 | 3.6 | Neutral |
| testneg_1476 | 0.05 | 0.29 | 0.41534 | -0.313968 | 2.66819 | -3 | 0.24966 | 0.12788 | 0.06922 | 0.07026 | 13.44572 | 0.03 | 0 | Neutral |
| testneg_1477 | -0.02 | 0.28 | 1.9684 | -3 | 7.214721 | -3 | 0.19898 | 0.07409 | 0.78515 | 0.35093 | 25.12484 | 0.39 | 15.2 | Neutral |
| testneg_1478 | -0.03 | -0.1 | 5.114664 | -3 | -0.238795 | -3 | 0.15192 | 0.14302 | 0.24832 | 0.21196 | 11.96702 | 0.367416667 | 11.2 | Neutral |
| testneg_1479 | 0 | 0 | 0.009894 | -3 | 2.260209 | -3 | 0.07996 | 0.08354 | 0.12965 | 0.75487 | 14.75193 | 0.09 | 0 | Neutral |
| testneg_1480 | 0.05 | -0.81 | 3.064064 | -3 | -0.163315 | -3 | 0.5481 | 0.10207 | 0.31125 | 0.20061 | 18.24519 | 0.14 | 0 | Neutral |
| testneg_1481 | 0 | 0 | 0.990948 | -3 | 1.514563 | -3 | 0.31859 | 0.11262 | 0.60152 | 0.12166 | 5.69444 | 0 | 0 | Neutral |
| testneg_1482 | 0.02 | 0.13 | 4.128054 | -3 | 0.649412 | -3 | 0.18498 | 0.12788 | 0.3394 | 0.5826 | 13.51351 | 0.192470017 | 0 | Neutral |
| testneg_1483 | 0.18 | -1.05 | 7.147075 | -3 | 1.295804 | -3 | 0.25989 | 0.11651 | 0.64795 | 0.21581 | 24.44444 | 0.41 | 20.4 | Neutral |
| testneg_1484 | 0.05 | 0.29 | -0.995638 | -3 | -0.022585 | -3 | 0.09669 | 0.12526 | 0.2543 | 0.12929 | 12.38369 | 0.01 | 0 | Neutral |
| testneg_1485 | 0 | 0 | 1.169086 | -3 | 5.494664 | -3 | 0.70616 | 0.09323 | 0.19898 | 0.12788 | 25.55152 | 0.070047619 | 0 | Neutral |
| testneg_1486 | 0.04 | -0.23 | 3.374375 | -3 | 0.860167 | -3 | 0.11564 | 0.1184 | 0.14767 | 0.09559 | 20.89414 | 0.02 | 0 | Neutral |
| testneg_1487 | -0.01 | 1.44 | 1.646184 | -3 | 1.963259 | -3 | 0.19698 | 0.1459 | 0.18964 | 0.34036 | 10.41667 | 0.161026316 | 0 | Neutral |
| testneg_1488 | -0.000303 | -0.0069697 | 0.145016 | -3 | 1.952626 | -3 | 0.42621 | 0.49093 | 0.2074 | 0.09696 | 17.21847 | 0.03 | 0 | Neutral |
| testneg_1489 | 0.08 | 0.03 | 2.172209 | -3 | -0.005763 | -3 | 0.14904 | 0.13261 | 0.20657 | 0.12788 | 6.99956 | 0.02 | 0 | Neutral |
| testneg_1490 | -0.01 | -0.79 | 1.377684 | -3 | 0.303179 | -3 | 0.09406 | 0.19858 | 0.14628 | 0.09488 | 17.21847 | 0.041314446 | 0 | Neutral |
| testneg_1491 | 0.01 | -1.3 | 5.149724 | -3 | 5.066913 | -3 | 0.29679 | 0.12788 | 0.08041 | 0.21802 | 10 | 0.725250815 | 98.8 | Disease_causing |
| testneg_1492 | 0.03 | -1.03 | 2.760537 | -3 | 2.37713 | -3 | 0.23126 | 0.16717 | 0.06704 | 0.1068 | 18.76159 | 0.099146825 | 0 | Neutral |
| testneg_1493 | 0 | 0 | 0.281322 | -3 | 2.494344 | -3 | 0.11335 | 0.14225 | 0.06344 | 0.12788 | 10.83333 | 0.030527778 | 0 | Neutral |
| testneg_1494 | -0.08 | -0.03 | 6.039402 | -3 | 0.590696 | -3 | 0.35244 | 0.31462 | 0.10809 | 0.09057 | 20.86969 | 0.273458333 | 2 | Neutral |
| testneg_1495 | 0 | 0 | 0.447219 | -3 | -0.877392 | -3 | 0.13884 | 0.12788 | 0.1311 | 0.12667 | 16.48772 | 0 | 0 | Neutral |
| testneg_1496 | 0.21 | 0.15 | 2.859152 | -3 | 1.24588 | 1.24588 | 0.33713 | 0.12788 | 0.13535 | 0.13562 | 10.22898 | 0.136151961 | 0 | Neutral |
| testneg_1497 | 0.04 | -0.23 | 0.914436 | -3 | -1.182618 | -3 | 0.01642 | 0.12788 | 0.16462 | 0.18467 | 14.04114 | 0.101291667 | 0 | Neutral |
| testneg_1498 | 0 | 0 | -0.801037 | -3 | 0.6838 | 0.6838 | 0.2049 | 0.12678 | 0.67327 | 0.23543 | 26.84685 | 0.070154581 | 0 | Neutral |
| testneg_1499 | 0.01 | 0.8 | 2.60181 | -3 | 2.296804 | -3 | 0.45014 | 0.12788 | 0.14668 | 0.17962 | 22.93219 | 0.158035714 | 0 | Neutral |
| testneg_1500 | 0 | 0 | 1.031442 | -3 | 5.240577 | -3 | 0.2761 | 0.24559 | 0.14525 | 0.14744 | 19.44444 | 0.439942785 | 31.6 | Neutral |
| testneg_1501 | -0.09 | 0.3 | 1.16938 | -3 | 0.539166 | -3 | 0.32974 | 0.12245 | 0.19898 | 0.12788 | 22.95376 | 0.010096475 | 0 | Neutral |
| testneg_1502 | 0 | -0.01 | 1.45889 | -3 | 4.837307 | -3 | 0.30209 | 0.14229 | 0.18705 | 0.43522 | 19.34591 | 0.315626884 | 4.6 | Neutral |
| testneg_1503 | 0.19 | -0.27 | 1.4509 | -3 | 1.435364 | -3 | 0.64566 | 0.22552 | 0.20357 | 0.12788 | 16.0119 | 0.05021164 | 0 | Neutral |
| testneg_1504 | 0.01 | -1.44 | 2.788395 | -3 | 5.245171 | -3 | 0.23567 | 0.12788 | 0.10914 | 0.76607 | 20.12164 | 0.537196429 | 70.2 | Disease_causing |
| testneg_1505 | 0 | -0.01 | -0.190389 | -3 | 1.293654 | -3 | 0.19898 | 0.07875 | 0.03324 | 0.12788 | 21.40066 | 0.02 | 0 | Neutral |
| testneg_1506 | 0.04 | 2.01 | -0.074864 | -3 | 1.062694 | -3 | 0.16468 | 0.11275 | 0.41655 | 0.49485 | 20.83333 | 0.16 | 0 | Neutral |
| testneg_1507 | -0.04 | 0.23 | 1.471012 | -3 | 0.80255 | -3 | 0.41102 | 0.0974 | 0.25957 | 0.06969 | 22.62411 | 0.01 | 0 | Neutral |
| testneg_1508 | 0.09 | -0.3 | 1.668712 | -3 | 3.793394 | -3 | 0.69696 | 0.10507 | 0.23955 | 0.97212 | 4.64133 | 0.307838456 | 1.2 | Neutral |
| testneg_1509 | -0.15 | -0.06 | -0.659954 | -3 | 1.696599 | -3 | 0.19898 | 0.12788 | 0.4334 | 0.09454 | 19.06745 | 0.05 | 0 | Neutral |
| testneg_1510 | 0.01763636 | -0.2005455 | 10.652511 | -3 | 3.577883 | 1.114692 | 0.42583 | 0.10901 | 0.05124 | 0.07099 | 27.4024 | 0.442486652 | 32.6 | Neutral |
| testneg_1511 | -0.06 | -0.18 | 3.18158 | -3 | 2.697859 | -3 | 0.23439 | 0.09253 | 0.64188 | 0.09494 | 18.6768 | 0.080833333 | 0 | Neutral |
| testneg_1512 | 0.04 | -0.23 | 1.122786 | -3 | -0.330887 | -3 | 0.19898 | 0.08029 | 0.44612 | 0.24054 | 18.6768 | 0.042007556 | 0 | Neutral |
| testneg_1513 | -0.08 | -0.03 | -0.057252 | -0.057252 | -0.272308 | -3 | 0.83176 | 0.23122 | 0.2288 | 0.15849 | 12.55779 | 0.100681818 | 0 | Neutral |
| testneg_1514 | -0.07 | -0.38 | 1.102328 | -3 | 5.180313 | -3 | 0.74271 | 0.10951 | 0.19898 | 0.12788 | 20.42553 | 0.09 | 0 | Neutral |
| testneg_1515 | 0.04 | -0.23 | 3.118463 | -3 | 2.367361 | -3 | 0.1846 | 0.375 | 0.15676 | 0.12788 | 19.27198 | 0.210212787 | 0 | Neutral |
| testneg_1516 | 0 | 0 | 0.191318 | -3 | 0.310587 | 0.310587 | 0.96113 | 0.24974 | 0.2132 | 0.12788 | 10.83333 | 0.021683635 | 0 | Neutral |
| testneg_1517 | 0 | 0 | -0.235365 | -3 | 0.510124 | -3 | 0.99535 | 0.38424 | 0.10161 | 0.13484 | 10.41667 | 0.115964286 | 0 | Neutral |
| testneg_1518 | 0 | 0 | 1.610282 | -3 | 0.127079 | -3 | 0.2663 | 0.32806 | 0.3012 | 0.10317 | 9.78453 | 0.010422436 | 0 | Neutral |
| testneg_1519 | 0.14 | -1.01 | 4.762479 | -3 | 4.116365 | -3 | 0.01573 | 0.10031 | 0.17843 | 0.15468 | 12.38369 | 0.504292766 | 56.6 | Disease_causing |
| testneg_1520 | -0.07 | 0.31 | 0.183303 | 0.183303 | 0.44932 | -3 | 0.12079 | 0.62023 | 0.05216 | 0.23851 | 14.94636 | 0.225797683 | 0 | Neutral |
| testneg_1521 | 0.03 | 0.1 | 3.638601 | -3 | 0.684698 | -3 | 0.9149 | 0.27793 | 0.09661 | 0.09981 | 16.54575 | 0.06008547 | 0 | Neutral |
| testneg_1522 | 0.2 | -1.26 | 4.721241 | -3 | -1.509499 | -3 | 0.40557 | 0.11865 | 0.85781 | 0.22949 | 29.86111 | 0.22 | 0.2 | Neutral |
| testneg_1523 | 0 | 0 | -0.02198 | -0.02198 | 0.616917 | -3 | 0.15509 | 0.1326 | 0.10781 | 0.09901 | 27.4024 | 0.03 | 0 | Neutral |
| testneg_1524 | 0.07 | 0.01 | 2.16506 | -3 | 0.57285 | -3 | 0.16379 | 0.31462 | 0.20501 | 0.6166 | 12.77241 | 0.169063875 | 0 | Neutral |
| testneg_1525 | 0 | 0 | 0.096056 | -3 | -1.462797 | -3 | 0.41655 | 0.49485 | 0.06547 | 0.09123 | 17.21847 | 0.021002506 | 0 | Neutral |
| testneg_1526 | -0.05 | -0.29 | 4.987047 | -3 | 4.765066 | -0.160038 | 0.31726 | 0.17249 | 0.60079 | 0.11198 | 34.83293 | 0.369027778 | 12.2 | Neutral |
| testneg_1527 | 0.06 | -0.57 | 5.777528 | -3 | 5.82456 | -3 | 0.17672 | 0.14229 | 0.33166 | 0.11755 | 18.88889 | 0.50527033 | 57.6 | Disease_causing |
| testneg_1528 | -0.21 | -0.15 | 5.060273 | -3 | 3.516778 | -3 | 0.06254 | 0.23561 | 0.30789 | 0.15135 | 17.01711 | 0.7090633 | 99.6 | Disease_causing |
| testneg_1529 | 0 | 0 | 0.188212 | -3 | 2.013802 | -3 | 0.34859 | 0.09799 | 0.22239 | 0.12885 | 20.95603 | 0.01 | 0 | Neutral |
| testneg_1530 | -0.01 | -0.79 | 0.652225 | -3 | 1.123147 | -3 | 0.04694 | 0.44682 | 0.59947 | 0.14339 | 14.92208 | 0.240417391 | 0 | Neutral |
| testneg_1531 | -0.05 | -0.29 | -1.484621 | -3 | 0.941259 | -3 | 0.38171 | 0.15977 | 0.19832 | 0.19622 | 14.22149 | 0.080395921 | 0 | Neutral |
| testneg_1532 | -0.16 | -0.08 | 1.251399 | -3 | -1.024193 | -3 | 0.20329 | 0.10718 | 0.0406 | 0.12788 | 12.29167 | 0.05 | 0 | Neutral |
| testneg_1533 | 0.02 | -0.36 | -0.01145 | -0.01145 | -1.247889 | -3 | 0.78462 | 0.12258 | 0.27577 | 0.09812 | 16.84579 | 0.010625 | 0 | Neutral |
| testneg_1534 | 0.14 | -1.01 | 2.621366 | -3 | 0.500561 | -3 | 0.81899 | 0.12788 | 0.66693 | 0.0986 | 11.25 | 0.037654639 | 0 | Neutral |
| testneg_1535 | -0.02 | 0.63 | 0.385855 | -3 | 3.442155 | -3 | 0.10153 | 0.13343 | 0.19898 | 0.12788 | 29.57517 | 0.09 | 0 | Neutral |
| testneg_1536 | 0 | 0 | 0.161405 | -3 | 1.294927 | -0.153636 | 0.64566 | 0.22552 | 0.35486 | 0.11584 | 11.69872 | 0.011758865 | 0 | Neutral |
| testneg_1537 | 0.14 | -1.01 | 3.161443 | -3 | -1.356362 | -1.912416 | 0.26961 | 0.09979 | 0.10362 | 0.12788 | 22.92553 | 0.04 | 0 | Neutral |
| testneg_1538 | -0.02 | -0.13 | 3.258128 | -3 | -0.245869 | -3 | 0.56307 | 0.10536 | 0.28156 | 0.13431 | 13.06915 | 0.09 | 0 | Neutral |
| testneg_1539 | -0.07 | -0.38 | -1.29721 | -3 | 0.454738 | -3 | 0.23385 | 0.12788 | 0.44103 | 0.10698 | 11.08369 | 0.01 | 0 | Neutral |
| testneg_1540 | 0.14 | -1.01 | 2.672539 | -3 | 5.282991 | -3 | 0.46371 | 0.11637 | 0.40234 | 0.10353 | 6.57011 | 0.162063312 | 0 | Neutral |
| testneg_1541 | 0 | 0 | 0.085558 | -3 | 3.561616 | -3 | 0.14643 | 0.12788 | 0.10919 | 0.105 | 10.41667 | 0.032902647 | 0 | Neutral |
| testneg_1542 | 0 | -0.32 | 4.776173 | -3 | 3.905772 | -3 | 0.25487 | 0.13098 | 0.19898 | 0.07875 | 34.87912 | 0.179263901 | 0 | Neutral |
| testneg_1543 | 0.21 | 0.15 | 0.789411 | -3 | -0.11997 | -3 | 0.13067 | 0.1005 | 0.44612 | 0.24054 | 18.6768 | 0.084166667 | 0 | Neutral |
| testneg_1544 | 0 | 0 | 1.320932 | -3 | -1.632697 | -3 | 0.39015 | 0.23372 | 0.08184 | 0.10206 | 10.83333 | 0.02 | 0 | Neutral |
| testneg_1545 | 0.01 | -1.44 | 0.737497 | -3 | 4.974068 | -3 | 0.97678 | 0.09885 | 0.18423 | 0.10808 | 20.43813 | 0.0958125 | 0 | Neutral |
| testneg_1546 | 0 | 0 | 5.922404 | -3 | 6.013954 | -3 | 0.08393 | 0.1123 | 0.22233 | 0.31758 | 27.06274 | 0.697070804 | 97.4 | Disease_causing |
| testneg_1547 | 0.22 | -2.66 | 7.324107 | -3 | 1.46728 | -3 | 0.33057 | 0.10549 | 0.09707 | 0.12788 | 29.05204 | 0.285 | 3.6 | Neutral |
| testneg_1548 | 0 | 0 | -0.50303 | -3 | 1.237005 | -3 | 0.15192 | 0.14302 | 0.38942 | 0.345 | 10.22898 | 0.05230977 | 0 | Neutral |
| testneg_1549 | 0.21 | 0.15 | 1.098308 | -3 | 2.494251 | -3 | 0.09346 | 0.12788 | 0.10314 | 0.08894 | 19.04744 | 0.05 | 0 | Neutral |
| testneg_1550 | -0.07 | -0.01 | 4.711487 | -3 | -0.435682 | -3 | 0.23292 | 0.15913 | 0.30103 | 0.1462 | 18.88889 | 0.200284389 | 0 | Neutral |
| testneg_1551 | -0.04 | 0.23 | -0.587078 | -3 | 1.875915 | -3 | 0.47872 | 0.09856 | 0.79857 | 0.12788 | 18.41494 | 0.02 | 0 | Neutral |
| testneg_1552 | 0.14 | -0.37 | 3.499396 | -3 | -0.085229 | -3 | 0.69249 | 0.12788 | 0.44644 | 0.10813 | 18.46106 | 0.031051282 | 0 | Neutral |
| testneg_1553 | 0.01 | -1.3 | 2.010656 | -3 | 2.435525 | -3 | 0.06848 | 0.09963 | 0.18096 | 0.12788 | 30.16992 | 0.04 | 0 | Neutral |
| testneg_1554 | 0 | 0 | 0.200125 | -3 | -1.171244 | -1.171244 | 0.13863 | 0.2579 | 0.19898 | 0.12788 | 10.22898 | 0.012526683 | 0 | Neutral |
| testneg_1555 | 0 | 0 | -0.415246 | -3 | 1.253567 | -3 | 0.06027 | 0.11574 | 0.4811 | 0.13398 | 5.69444 | 0.07 | 0 | Neutral |
| testneg_1556 | 0.15 | -0.82 | 4.135893 | -3 | 0.076953 | -3 | 0.08628 | 0.11498 | 0.10187 | 0.12339 | 23.64499 | 0.041077598 | 0 | Neutral |
| testneg_1557 | 0 | -0.01 | 0.56153 | -3 | 5.108849 | -3 | 0.08509 | 0.12788 | 0.49064 | 0.12294 | 19.09347 | 0.06 | 0 | Neutral |
| testneg_1558 | 0.04 | -0.23 | 4.939659 | -3 | 0.256959 | -3 | 0.13563 | 0.12378 | 0.46946 | 0.08281 | 4.63186 | 0.05 | 0 | Neutral |
| testneg_1559 | 0 | 0 | 1.003967 | -3 | 5.00736 | -3 | 0.12835 | 0.15143 | 0.18842 | 0.12378 | 19.28571 | 0.130657507 | 0 | Neutral |
| testneg_1560 | 0 | 0 | 2.078803 | -3 | -0.019609 | -3 | 0.40074 | 0.36296 | 0.69563 | 0.12788 | 16.76075 | 0.03 | 0 | Neutral |
| testneg_1561 | 0 | 0 | 0.388922 | -3 | 2.493678 | -3 | 0.98107 | 0.70722 | 0.50057 | 0.10408 | 21.6702 | 0.0901 | 0 | Neutral |
| testneg_1562 | 0.03 | 0.12 | 3.15417 | -3 | 5.957349 | -3 | 0.25764 | 0.53109 | 0.46142 | 0.11057 | 18.55089 | 0.415448718 | 27 | Neutral |
| testneg_1563 | 0 | -0.01 | -0.496432 | -3 | 0.452231 | -3 | 0.06247 | 0.14184 | 0.46691 | 0.07886 | 12.06235 | 0.02 | 0 | Neutral |
| testneg_1564 | 0 | 0 | 1.362887 | -3 | -1.996459 | -3 | 0.38527 | 0.24812 | 0.04988 | 0.25503 | 10.41667 | 0.073451025 | 0 | Neutral |
| testneg_1565 | 0 | -0.01 | 1.210317 | -3 | 2.268243 | -3 | 0.19898 | 0.11262 | 0.85345 | 0.13067 | 33.85548 | 0.02 | 0 | Neutral |
| testneg_1566 | 0 | 0 | 0.634596 | -3 | 2.682586 | -3 | 0.54489 | 0.09921 | 0.10383 | 0.08853 | 23.50608 | 0.052195308 | 0 | Neutral |
| testneg_1567 | -0.05 | 0.81 | 3.717561 | -3 | 6.562934 | -3 | 0.09661 | 0.09981 | 0.12965 | 0.75487 | 15.67982 | 0.65 | 95.6 | Disease_causing |
| testneg_1568 | 0.07 | 0.38 | -1.041977 | -3 | -0.388631 | -3 | 0.49665 | 0.20425 | 0.19227 | 0.1096 | 18.09524 | 0.051326174 | 0 | Neutral |
| testneg_1569 | -0.01 | -0.8 | 2.607371 | -3 | -0.44217 | -3 | 0.53239 | 0.17509 | 0.33148 | 0.11968 | 11.90227 | 0.04 | 0 | Neutral |
| testneg_1570 | 0 | 0 | 1.317673 | -3 | 2.503584 | -3 | 0.45415 | 0.12971 | 0.11359 | 0.08631 | 11.30221 | 0.02 | 0 | Neutral |
| testneg_1571 | -0.02 | 0.63 | 2.928857 | 2.928857 | 1.199598 | -3 | 0.11534 | 0.12281 | 0.13647 | 0.11806 | 23.53679 | 0.36 | 9.2 | Neutral |
| testneg_1572 | -0.07 | -0.38 | 2.097779 | -3 | -0.483554 | -3 | 0.81054 | 0.28104 | 0.19898 | 0.56316 | 11.55371 | 0.131341737 | 0 | Neutral |
| testneg_1573 | -0.01 | -0.79 | 2.619944 | -3 | 1.144976 | -3 | 0.9924 | 0.37896 | 0.53592 | 0.11293 | 10 | 0.080713762 | 0 | Neutral |
| testneg_1574 | 0 | -0.01 | 5.580492 | -3 | 0.312443 | -3 | 0.07347 | 0.08817 | 0.3834 | 0.3731 | 17.21847 | 0.341794118 | 7.6 | Neutral |
| testneg_1575 | 0 | 0 | 0.664346 | -3 | 6.878081 | -3 | 0.22169 | 0.11262 | 0.28227 | 0.10211 | 21.82136 | 0.164020468 | 0 | Neutral |
| testneg_1576 | 0.06 | -0.57 | 4.926259 | -3 | -0.354552 | -3 | 0.19898 | 0.12788 | 0.09815 | 0.12788 | 22.89111 | 0.05 | 0 | Neutral |
| testneg_1577 | 0.09 | -0.3 | 5.374733 | -3 | -0.513734 | -3 | 0.20055 | 0.12788 | 0.256 | 0.20036 | 16.95767 | 0.362795455 | 11.6 | Neutral |
| testneg_1578 | 0.16 | -1.37 | 2.791385 | -3 | 0.925236 | -3 | 0.59853 | 0.75237 | 0.30925 | 0.10335 | 15.51622 | 0.13 | 0 | Neutral |
| testneg_1579 | 0.07 | 0.38 | -0.115877 | -3 | 0.916588 | -3 | 0.44385 | 0.1311 | 0.06603 | 0.08071 | 10.22898 | 0.03 | 0 | Neutral |
| testneg_1580 | 0 | -0.32 | -0.156596 | -3 | 1.237062 | -3 | 0.26878 | 0.09179 | 0.09473 | 0.08327 | 55.44203 | 0.05 | 0 | Neutral |
| testneg_1581 | -0.03 | 1.03 | 2.97112 | -3 | 1.135919 | -3 | 0.5901 | 0.19444 | 0.86513 | 0.31355 | 11.25 | 0.236026786 | 0.2 | Neutral |
| testneg_1582 | 0.04 | 2.01 | 1.269375 | -3 | 2.908082 | -3 | 0.54489 | 0.09921 | 0.18246 | 0.12791 | 17.97975 | 0.133095238 | 0 | Neutral |
| testneg_1583 | -0.03 | -0.1 | 0.212913 | -3 | 0.953839 | -3 | 0.11391 | 0.19434 | 0.11229 | 0.08897 | 26.04838 | 0.042386364 | 0 | Neutral |
| testneg_1584 | 0 | 0 | 2.165318 | -3 | 0.951477 | 0.951477 | 0.19989 | 0.16777 | 0.27083 | 0.14313 | 43.37913 | 0.098510101 | 0 | Neutral |
| testneg_1585 | 0.21 | 0.15 | 6.051557 | -3 | 2.089989 | -3 | 0.13482 | 0.12788 | 0.99941 | 0.50288 | 17.07858 | 0.38 | 15.2 | Neutral |

| | | | | | | | | | | | | | | |
|---|---|---|---|---|---|---|---|---|---|---|---|---|---|---|
| testneg_1586 | 0 | 0 | 1.51996 | -3 | 0.615391 | -3 | 0.09105 | 0.27434 | 0.19898 | 0.19444 | 50.43279 | 0.167973485 | 0 | Neutral |
| testneg_1587 | 0 | 0 | -1.230621 | -3 | 4.875592 | -3 | 0.64254 | 0.10729 | 0.12269 | 0.08105 | 20.33859 | 0.075409425 | 0 | Neutral |
| testneg_1588 | -0.03 | -0.12 | -0.618073 | -3 | 1.944832 | -3 | 0.39863 | 0.44944 | 0.69254 | 0.10021 | 18.88889 | 0.059491342 | 0 | Neutral |
| testneg_1589 | 0 | 0 | 0.147975 | -0.123782 | 5.691093 | -3 | 0.06898 | 0.11659 | 0.06729 | 0.17051 | 39.9725 | 0.361330357 | 8.4 | Neutral |
| testneg_1590 | 0.04 | -0.23 | 2.522879 | -3 | 4.751068 | -3 | 0.99998 | 0.86471 | 0.9449 | 0.9378 | 4.44444 | 0.54 | 71.8 | Disease_causing |
| testneg_1591 | 0.05 | -0.81 | 0.820857 | -3 | 5.862225 | -3 | 0.0887 | 0.10047 | 0.21587 | 0.11701 | 32.82772 | 0.090377493 | 0 | Neutral |
| testneg_1592 | 0.07 | 0.38 | 4.587402 | -3 | 6.972724 | -3 | 0.23567 | 0.12788 | 0.44953 | 0.14403 | 19.01053 | 0.654658156 | 94.2 | Disease_causing |
| testneg_1593 | 0 | 0 | 0.038426 | -3 | -0.046909 | -3 | 0.54909 | 0.85056 | 0.13143 | 0.90853 | 21.25875 | 0.260799612 | 0.2 | Neutral |
| testneg_1594 | 0 | 0 | 0.330621 | -3 | 0.48851 | -3 | 0.41383 | 0.119 | 0.17793 | 0.10516 | 10.80038 | 0.01 | 0 | Neutral |
| testneg_1595 | 0.03 | -1.03 | 2.432661 | -3 | 2.631274 | -3 | 0.1541 | 0.1077 | 0.14458 | 0.14062 | 29.4946 | 0.122060724 | 0 | Neutral |
| testneg_1596 | 0.21 | 0.15 | 5.422894 | -3 | -0.280066 | -3 | 0.36709 | 0.19019 | 0.13562 | 0.13216 | 45.0343 | 0.284047619 | 1.4 | Neutral |
| testneg_1597 | 0.14 | -0.37 | 4.82283 | -3 | 5.915401 | -3 | 0.08999 | 0.0989 | 0.16777 | 0.1749 | 10.83333 | 0.729615029 | 99.2 | Disease_causing |
| testneg_1598 | 0 | 0 | 0.191318 | -3 | 0.599729 | -3 | 0.96113 | 0.24974 | 0.38762 | 0.12788 | 22.54725 | 0.020909091 | 0 | Neutral |
| testneg_1599 | 0.16 | 0.08 | -0.169987 | -3 | 1.28563 | -3 | 0.99445 | 0.10396 | 0.25029 | 0.12267 | 18.88889 | 0.05 | 0 | Neutral |
| testneg_1600 | -0.15 | -0.04 | 3.431501 | -3 | 0.190526 | -3 | 0.1289 | 0.14878 | 0.1834 | 0.12788 | 29.36393 | 0.11 | 0 | Neutral |
| testneg_1601 | -0.06 | -0.18 | 3.107593 | -3 | 1.831606 | -3 | 0.18512 | 0.12788 | 0.25896 | 0.45481 | 14.00313 | 0.205823198 | 0 | Neutral |
| testneg_1602 | 0.03 | 1.22 | 2.753966 | -3 | 5.537244 | -3 | 0.1551 | 0.09276 | 0.19898 | 0.09972 | 13.10593 | 0.191009615 | 0.2 | Neutral |
| testneg_1603 | 0 | 0 | 0.699953 | -3 | 0.201826 | -3 | 0.54231 | 0.15254 | 0.26337 | 0.12771 | 10.41667 | 0.010155833 | 0 | Neutral |
| testneg_1604 | 0.03 | 0.1 | 4.671506 | -3 | 1.614723 | 1.614723 | 0.15856 | 0.10994 | 0.74944 | 0.1263 | 10 | 0.08 | 0 | Neutral |
| testneg_1605 | 0 | 0 | 0.1162 | -3 | 0.245964 | -3 | 0.17141 | 0.1104 | 0.74099 | 0.44443 | 11.35684 | 0.02 | 0 | Neutral |
| testneg_1606 | 0 | 0 | 0.085558 | -3 | 0.513687 | -3 | 0.14643 | 0.12788 | 0.05153 | 0.12788 | 18.33333 | 0.000817308 | 0 | Neutral |
| testneg_1607 | -0.09 | 0.3 | 1.16938 | -3 | 0.712749 | -3 | 0.32974 | 0.12245 | 0.12203 | 0.13515 | 10.41667 | 0.080704088 | 0 | Neutral |
| testneg_1608 | -0.03 | 1.03 | 1.290195 | 1.290195 | 5.488868 | -3 | 0.10761 | 0.12246 | 0.10458 | 0.10499 | 35.34208 | 0.21 | 0 | Neutral |
| testneg_1609 | 0 | 0 | 0.085674 | -3 | -1.013068 | -1.27012 | 0.57975 | 0.1248 | 0.40076 | 0.11341 | 16.42913 | 0.01 | 0 | Neutral |
| testneg_1610 | -0.03 | -0.1 | 3.735674 | -3 | 0.774689 | 0.774689 | 0.13023 | 0.29692 | 0.24219 | 0.17458 | 6.00877 | 0.348050105 | 8.4 | Neutral |
| testneg_1611 | -0.16 | -0.08 | 3.87308 | -3 | 0.580978 | -3 | 0.18925 | 0.32479 | 0.19898 | 0.11773 | 6.03357 | 0.15 | 0 | Neutral |
| testneg_1612 | -0.09 | 0.3 | -0.859467 | -3 | 1.246046 | -3 | 0.84616 | 0.1471 | 0.31621 | 0.12788 | 11.14837 | 0.04 | 0 | Neutral |
| testneg_1613 | -0.08 | -0.03 | 0.536745 | -3 | 3.746869 | -3 | 0.34295 | 0.12788 | 0.10244 | 0.10107 | 10.41667 | 0.08009434 | 0 | Neutral |
| testneg_1614 | 0 | -0.01 | 0.655486 | 0.655486 | 1.39708 | -3 | 0.20772 | 0.16918 | 0.25637 | 0.37408 | 22.77778 | 0.18 | 0 | Neutral |
| testneg_1615 | 0 | 0 | 0.699729 | -3 | 0.939929 | -3 | 0.76391 | 0.14229 | 0.18701 | 0.12788 | 13.51351 | 0.01 | 0 | Neutral |
| testneg_1616 | 0 | 0.73 | 5.650938 | -3 | 0.576122 | -3 | 0.16671 | 0.18311 | 0.18341 | 0.13901 | 19.44444 | 0.37 | 12.6 | Neutral |
| testneg_1617 | 0.08 | 0.03 | 1.747877 | -3 | 0.254162 | -3 | 0.90811 | 0.17448 | 0.94807 | 0.14977 | 10.20833 | 0.091 | 0 | Neutral |
| testneg_1618 | 0 | -0.01 | 1.961035 | -3 | -0.384176 | -3 | 0.55105 | 0.43208 | 0.31238 | 0.14393 | 26.80135 | 0.106939727 | 0 | Neutral |
| testneg_1619 | 0.22 | -2.66 | 6.235574 | -3 | 1.887378 | -3 | 0.15842 | 0.1294 | 0.98565 | 0.23629 | 15.72887 | 0.376666667 | 14.6 | Neutral |
| testneg_1620 | 0 | 0 | 1.592805 | -3 | 6.281692 | -1.296157 | 0.25025 | 0.18924 | 0.56013 | 0.29233 | 20.83333 | 0.516293832 | 60.4 | Disease_causing |
| testneg_1621 | 0.14 | -0.37 | 0.198379 | -3 | 2.729204 | -3 | 0.03991 | 0.08342 | 0.62339 | 0.19592 | 10.41667 | 0.17035628 | 0 | Neutral |
| testneg_1622 | -0.07 | -0.38 | 1.941375 | -3 | -1.356524 | -3 | 0.23962 | 0.10029 | 0.09602 | 0.10726 | 13.02635 | 0.01 | 0 | Neutral |
| testneg_1623 | 0.11 | -0.19 | -0.25166 | -1.810379 | 2.960128 | -3 | 0.05677 | 0.06758 | 0.19898 | 0.12788 | 17.5 | 0.060468991 | 0 | Neutral |
| testneg_1624 | 0 | 0 | 0.22996 | -1.285414 | 0.292982 | -3 | 0.0693 | 0.09293 | 0.15677 | 0.18537 | 18.6768 | 0.041452922 | 0 | Neutral |
| testneg_1625 | 0 | 0 | 0.380797 | 0.380797 | -0.05178 | -3 | 0.61011 | 0.09584 | 0.19898 | 0.12788 | 19.36941 | 0.0119375 | 0 | Neutral |
| testneg_1626 | 0.03 | 1.22 | 3.228457 | -3 | -0.112406 | -3 | 0.07681 | 0.22865 | 0.94807 | 0.14977 | 4.53393 | 0.23813529 | 0 | Neutral |
| testneg_1627 | 0.14 | -1.01 | 6.843378 | -3 | -0.140634 | -3 | 0.23624 | 0.12788 | 0.19898 | 0.12788 | 12.63889 | 0.21 | 1.8 | Neutral |
| testneg_1628 | -0.03 | -0.12 | 1.697358 | -3 | -1.214088 | -3 | 0.25989 | 0.11651 | 0.04192 | 0.06306 | 32.18757 | 0.010198506 | 0 | Neutral |
| testneg_1629 | 0 | 0 | 0.81674 | -3 | 5.198624 | -3 | 0.84238 | 0.12559 | 0.21967 | 0.09467 | 19.7952 | 0.050483997 | 0 | Neutral |
| testneg_1630 | 0 | 0.32 | -0.20856 | -3 | 3.035676 | -3 | 0.19898 | 0.12788 | 0.74919 | 0.85508 | 17.21847 | 0.163083895 | 0 | Neutral |
| testneg_1631 | 0.08 | 0.03 | -0.61726 | -0.61726 | 0.996116 | -3 | 0.20673 | 0.13236 | 0.0478 | 0.12788 | 14.95527 | 0.030270468 | 0 | Neutral |
| testneg_1632 | 0 | 0 | 0.237368 | -3 | 0.672473 | -0.063333 | 0.1602 | 0.11262 | 0.493 | 0.35835 | 19.90803 | 0.070794466 | 0 | Neutral |
| testneg_1633 | 0 | 0 | -0.061652 | -3 | 2.733966 | -3 | 0.83074 | 0.14769 | 0.34205 | 0.11195 | 18.61111 | 0.04 | 0 | Neutral |
| testneg_1634 | 0.07 | 0.01 | 2.724401 | -3 | 1.300713 | -3 | 0.16778 | 0.82322 | 0.23088 | 0.12788 | 12.46937 | 0.105 | 0 | Neutral |
| testneg_1635 | 0 | 0 | 1.563064 | 1.563064 | 5.295382 | -3 | 0.61027 | 0.14229 | 0.19898 | 0.09584 | 12.65937 | 0.141037879 | 0 | Neutral |
| testneg_1636 | 0 | 0 | 0.036223 | -3 | 0.851312 | -3 | 0.9355 | 0.40418 | 0.19898 | 0.12788 | 4.44444 | 0.000844182 | 0 | Neutral |
| testneg_1637 | 0 | 0 | 0.233061 | -3 | 3.124222 | -3 | 0.07828 | 0.76259 | 0.20249 | 0.10327 | 10.41667 | 0.111230735 | 0 | Neutral |
| testneg_1638 | 0 | 0 | 1.840893 | -3 | 0.486089 | -3 | 0.18465 | 0.11262 | 0.10613 | 0.11203 | 12.35574 | 0.000166667 | 0 | Neutral |
| testneg_1639 | -0.03 | 1.03 | -1.584877 | -3 | 1.999184 | -3 | 0.01409 | 0.11307 | 0.84236 | 0.14093 | 10.83333 | 0.14 | 0 | Neutral |
| testneg_1640 | -0.07 | -0.38 | 5.176715 | -3 | 0.292294 | -3 | 0.206 | 0.14631 | 0.08184 | 0.10206 | 10 | 0.13 | 0 | Neutral |
| testneg_1641 | 0 | 0 | 1.56019 | -3 | -0.030607 | -3 | 0.25487 | 0.13098 | 0.17146 | 0.19882 | 11.25 | 0.034579612 | 0 | Neutral |
| testneg_1642 | 0.01300699 | -0.0248951 | 2.462213 | -3 | 0.233598 | 0.233598 | 0.32016 | 0.11638 | 0.25804 | 0.11876 | 20.36138 | 0.01 | 0 | Neutral |
| testneg_1643 | -0.1 | -1.9 | 0.353374 | -3 | 6.211178 | -3 | 0.16393 | 0.08897 | 0.14668 | 0.17962 | 37.07139 | 0.38 | 11.2 | Neutral |
| testneg_1644 | 0.07 | 0.33 | 3.909213 | -3 | -0.362138 | -0.362138 | 0.30926 | 0.11955 | 0.09964 | 0.10384 | 29.9024 | 0.03 | 0 | Neutral |
| testneg_1645 | -0.02 | 0.63 | -0.628934 | -3 | 0.130473 | -3 | 0.17576 | 0.12193 | 0.41261 | 0.12096 | 13.51351 | 0.01 | 0 | Neutral |
| testneg_1646 | 0.21 | 0.15 | 2.233647 | -3 | -1.155982 | -3 | 0.27175 | 0.14094 | 0.03942 | 0.12788 | 23.97333 | 0.05 | 0 | Neutral |
| testneg_1647 | -0.01 | -0.79 | 3.446663 | -3 | 1.178684 | -3 | 0.34225 | 0.12788 | 0.12974 | 0.11886 | 10.83333 | 0.031306818 | 0 | Neutral |
| testneg_1648 | 0 | 0 | 2.66657 | -3 | 9.585319 | -3 | 0.99948 | 0.5595 | 0.0895 | 0.12788 | 18.19225 | 0.48 | 47.4 | Neutral |
| testneg_1649 | 0.01 | 0.79 | -0.500727 | -3 | 3.507953 | -3 | 0.07684 | 0.09138 | 0.78253 | 0.1114 | 10.41667 | 0.1 | 0 | Neutral |
| testneg_1650 | 0 | 0 | 1.125734 | -3 | 1.084274 | -3 | 0.12576 | 0.08486 | 0.07606 | 0.13082 | 16.04971 | 0.02 | 0 | Neutral |
| testneg_1651 | 0.18 | -1.05 | 6.08905 | -3 | 1.746374 | -3 | 0.13555 | 0.16663 | 0.1077 | 0.12788 | 4.44444 | 0.277 | 2.6 | Neutral |
| testneg_1652 | 0 | 0 | 0.016935 | -3 | 3.28103 | -3 | 0.11584 | 0.19704 | 0.27077 | 0.10337 | 18.33333 | 0.086719488 | 0 | Neutral |
| testneg_1653 | 0.18 | -1.05 | 4.57164 | -3 | 3.05843 | 3.05843 | 0.11335 | 0.14225 | 0.74136 | 0.1049 | 18.33333 | 0.309166667 | 0.2 | Neutral |
| testneg_1654 | -0.14 | -0.54 | 3.948867 | -3 | 2.830718 | -3 | 0.21236 | 0.12788 | 0.11229 | 0.08897 | 20.45259 | 0.110217494 | 0 | Neutral |
| testneg_1655 | -0.07 | -0.38 | 6.836452 | -3 | 3.01628 | 3.01628 | 0.19898 | 0.09316 | 0.35532 | 0.24703 | 18.33333 | 0.705 | 98.4 | Disease_causing |
| testneg_1656 | 0.07 | -0.31 | -1.241467 | -3 | 0.084517 | -3 | 0.2578 | 0.09796 | 0.25179 | 0.09355 | 18.73163 | 0.01 | 0 | Neutral |
| testneg_1657 | 0 | 0 | 1.315096 | -3 | 0.90228 | -3 | 0.26972 | 0.12484 | 0.10058 | 0.13882 | 17.27427 | 0.046046721 | 0 | Neutral |
| testneg_1658 | 0 | 0 | 0.013394 | -3 | 1.024238 | -3 | 0.12582 | 0.12788 | 0.0893 | 0.11207 | 13.51351 | 9.64E-05 | 0 | Neutral |
| testneg_1659 | 0 | 0 | 0.318608 | -3 | -1.124679 | -3 | 0.11373 | 0.10739 | 0.83798 | 0.14433 | 8.45975 | 0.02 | 0 | Neutral |
| testneg_1660 | 0 | 0 | -0.054636 | -3 | 2.753563 | -3 | 0.5632 | 0.28442 | 0.25009 | 0.11017 | 9.56261 | 0.04 | 0 | Neutral |
| testneg_1661 | 0 | 0 | 0.359449 | -3 | 5.907423 | -3 | 0.06218 | 0.10661 | 0.00451 | 0.2022 | 39.07476 | 0.34 | 7 | Neutral |
| testneg_1662 | 0.02 | -0.28 | -1.087398 | -3 | 0.827167 | -3 | 0.22794 | 0.10688 | 0.07211 | 0.12934 | 22.00647 | 0.010250627 | 0 | Neutral |
| testneg_1663 | 0 | 0 | 0.584441 | -3 | 1.454709 | -3 | 0.97678 | 0.09885 | 0.12488 | 0.10849 | 10.41667 | 0.010228639 | 0 | Neutral |
| testneg_1664 | 0 | -0.01 | 0.291492 | -3 | 0.779287 | -3 | 0.07064 | 0.11699 | 0.18067 | 0.14759 | 13.29151 | 0.02266954 | 0 | Neutral |
| testneg_1665 | 0 | 0 | -0.123782 | -0.652914 | 0.57285 | -3 | 0.06898 | 0.11659 | 0.20501 | 0.6166 | 39.27806 | 0.083596932 | 0 | Neutral |
| testneg_1666 | 0.19 | -0.27 | 2.303838 | -3 | 4.902265 | -3 | 0.17677 | 0.67006 | 0.1059 | 0.1265 | 18.33333 | 0.32 | 3.8 | Neutral |
| testneg_1667 | 0.03 | 0.1 | 0.13559 | -3 | 0.249407 | -3 | 0.19898 | 0.12788 | 0.07652 | 0.30451 | 16.8018 | 0.073333333 | 0 | Neutral |
| testneg_1668 | 0 | 0 | 0.068014 | 0.068014 | 6.27863 | -3 | 0.20869 | 0.13493 | 0.96396 | 0.7628 | 5.9904 | 0.419469246 | 25.4 | Neutral |
| testneg_1669 | 0.22 | -2.66 | 4.098464 | -3 | 1.286474 | -3 | 0.31124 | 0.08875 | 0.75033 | 0.64204 | 4.72222 | 0.2726909 | 0.4 | Neutral |
| testneg_1670 | 0.01 | 0.8 | 5.156941 | 3.013154 | 1.092413 | -3 | 0.28156 | 0.13431 | 0.96396 | 0.7628 | 12.81915 | 0.68 | 97 | Disease_causing |
| testneg_1671 | 0.07 | -0.31 | 2.407105 | -3 | -4.511894 | -3 | 0.21095 | 0.10574 | 0.10011 | 0.08014 | 21.53664 | 0.02 | 0 | Neutral |
| testneg_1672 | 0.15 | -0.28 | 1.300713 | -3 | 0.560505 | -3 | 0.23088 | 0.12788 | 0.99564 | 0.25794 | 14.82512 | 0.08 | 0 | Neutral |
| testneg_1673 | 0.04 | -0.23 | 3.570813 | -3 | 1.845442 | -3 | 0.68054 | 0.09208 | 0.99293 | 0.79067 | 10.03745 | 0.21 | 0 | Neutral |
| testneg_1674 | 0.05 | -0.81 | 1.441797 | -3 | 6.526232 | -3 | 0.14 | 0.12151 | 0.2016 | 0.13956 | 18.88889 | 0.359285714 | 11.8 | Neutral |
| testneg_1675 | 0.02 | -0.68 | 6.639247 | -3 | -1.902903 | -3 | 0.27407 | 0.1136 | 0.1254 | 0.08697 | 12.29167 | 0.22 | 0.6 | Neutral |
| testneg_1676 | 0 | 0 | 1.861867 | -3 | -0.047736 | -3 | 0.32963 | 0.1035 | 0.19898 | 0.82835 | 13.03571 | 0.050792779 | 0 | Neutral |
| testneg_1677 | -0.17 | -0.38 | -0.915939 | -3 | 7.324107 | -3 | 0.17322 | 0.10311 | 0.33057 | 0.10549 | 13.37793 | 0.33 | 4.8 | Neutral |
| testneg_1678 | -0.08 | -0.03 | -0.741718 | -3 | 1.902366 | -3 | 0.06218 | 0.10661 | 0.12231 | 0.23155 | 27.61073 | 0.122142081 | 0 | Neutral |
| testneg_1679 | -0.02 | 0.36 | -1.22394 | -3 | 0.504888 | -3 | 0.50063 | 0.154 | 0.06476 | 0.07491 | 18.72565 | 0.04 | 0 | Neutral |
| testneg_1680 | 0.11 | -1.95 | 0.609952 | 0.609952 | 4.051032 | -3 | 0.06224 | 0.12788 | 0.99178 | 0.93124 | 20.98109 | 0.47639652 | 43.2 | Neutral |
| testneg_1681 | 0 | 0 | 0.497873 | -3 | 3.369604 | -3 | 0.33049 | 0.12788 | 0.10249 | 0.09681 | 11.92061 | 0.04 | 0 | Neutral |
| testneg_1682 | 0.14 | -1.01 | 5.928464 | -3 | -0.554043 | -3 | 0.11316 | 0.12788 | 0.81659 | 0.10424 | 18.41137 | 0.110540936 | 0 | Neutral |
| testneg_1683 | -0.04 | 0.23 | 1.095448 | -3 | 0.803253 | -3 | 0.13794 | 0.40936 | 0.05637 | 0.06544 | 20.66855 | 0.06 | 0 | Neutral |
| testneg_1684 | 0 | 0 | 1.123147 | -3 | 5.090552 | -3 | 0.59947 | 0.14339 | 0.10185 | 0.12788 | 33.72427 | 0.123777174 | 0 | Neutral |
| testneg_1685 | -0.01 | 0.57 | 0.464102 | -3 | 2.560172 | -0.265696 | 0.14668 | 0.25103 | 0.53292 | 0.32174 | 10.83333 | 0.313098726 | 3.8 | Neutral |
| testneg_1686 | -0.09 | 0.3 | -2.629762 | -3 | 1.346164 | -3 | 0.19898 | 0.12788 | 0.17639 | 0.12919 | 16.55041 | 0.03 | 0 | Neutral |
| testneg_1687 | 0 | 0 | -0.196789 | -3 | 1.473689 | -3 | 0.1192 | 0.10391 | 0.21677 | 0.1681 | 19.01713 | 0.040930736 | 0 | Neutral |
| testneg_1688 | 0.05 | 0.29 | 0.313674 | -3 | 2.670295 | -3 | 0.26812 | 0.30371 | 0.09807 | 0.24901 | 4.72222 | 0.280411451 | 0.4 | Neutral |
| testneg_1689 | -0.03 | -0.12 | 0.803064 | -3 | 1.555282 | -3 | 0.08684 | 0.08917 | 0.67796 | 0.10874 | 19.09347 | 0.02 | 0 | Neutral |
| testneg_1690 | 0.00442308 | -0.005641 | 2.309347 | -3 | -0.552345 | -3 | 0.80629 | 0.12788 | 0.23308 | 0.77395 | 10 | 0.063178733 | 0 | Neutral |
| testneg_1691 | -0.03 | -0.1 | 1.531669 | -3 | 1.27566 | -3 | 0.59321 | 0.59981 | 0.18535 | 0.12788 | 12.45484 | 0.05 | 0 | Neutral |
| testneg_1692 | 0.05 | 0.29 | 0.312568 | -3 | 1.790743 | -3 | 0.18264 | 0.10206 | 0.7823 | 0.5949 | 18.6768 | 0.060227273 | 0 | Neutral |
| testneg_1693 | -0.03 | -0.1 | 0.968512 | -3 | 2.007803 | -3 | 0.40074 | 0.36296 | 0.06647 | 0.09129 | 27.34767 | 0.06 | 0 | Neutral |
| testneg_1694 | -0.02 | 0.36 | 0.953839 | -3 | -1.168846 | -3 | 0.11229 | 0.08897 | 0.19898 | 0.12788 | 30.73984 | 0.01 | 0 | Neutral |
| testneg_1695 | -0.16 | 1.37 | 4.353927 | -3 | 5.557447 | -3 | 0.03072 | 0.27473 | 0.34633 | 0.47228 | 11.92982 | 0.829055336 | 100 | Disease_causing |
| testneg_1696 | 0.07 | 0.7 | -1.101027 | -3 | 0.979262 | -3 | 0.14477 | 0.18932 | 0.14038 | 0.21177 | 11.30228 | 0.122344615 | 0 | Neutral |
| testneg_1697 | -0.18 | 1.05 | 1.978065 | -3 | 1.9313 | -3 | 0.08011 | 0.12788 | 0.68568 | 0.65219 | 30.45796 | 0.186875 | 0 | Neutral |
| testneg_1698 | -0.04 | 0.23 | 2.067026 | -3 | 2.025629 | -3 | 0.19898 | 0.40608 | 0.06344 | 0.12788 | 10.83333 | 0.07 | 0 | Neutral |
| testneg_1699 | -0.07 | -0.38 | 6.068773 | -3 | -1.487172 | -1.487172 | 0.08509 | 0.12788 | 0.103 | 0.403 | 16.8018 | 0.47 | 44.2 | Neutral |
| testneg_1700 | 0 | 0 | 1.62409 | -3 | -0.055664 | -3 | 0.20457 | 0.12378 | 0.38002 | 0.22508 | 19.646 | 0.03 | 0 | Neutral |
| testneg_1701 | -0.08 | -0.03 | 6.10531 | -3 | 1.406345 | -3 | 0.23506 | 0.24555 | 0.7062 | 0.83002 | 10.22898 | 0.635648419 | 92.8 | Disease_causing |
| testneg_1702 | 0 | 0 | 1.657251 | -3 | -0.780137 | -3 | 0.17973 | 0.12278 | 0.06958 | 0.16086 | 11.875 | 0.063775124 | 0 | Neutral |
| testneg_1703 | -0.07 | -0.38 | 3.089818 | -3 | 0.292982 | -3 | 0.09026 | 0.09447 | 0.15677 | 0.18537 | 13.51351 | 0.159222048 | 0 | Neutral |
| testneg_1704 | 0.04 | -0.23 | 1.620184 | -3 | -1.065416 | -3 | 0.20502 | 0.10447 | 0.12161 | 0.24178 | 42.41609 | 0.071663898 | 0 | Neutral |
| testneg_1705 | 0.09 | -0.3 | 1.673428 | 1.673428 | 5.061069 | -3 | 0.32352 | 0.10243 | 0.2734 | 0.2578 | 10.22898 | 0.431157521 | 26 | Neutral |
| testneg_1706 | 0.02 | -0.63 | -0.730881 | -3 | 1.836976 | -3 | 0.26476 | 0.08217 | 0.84491 | 0.30945 | 19.82456 | 0.07 | 0 | Neutral |
| testneg_1707 | -0.08 | -0.03 | 5.960756 | -3 | 5.673256 | -3 | 0.2477 | 0.11928 | 0.04519 | 0.12788 | 4.72222 | 0.539852941 | 66.2 | Disease_causing |

| | | | | | | | | | | | | | | |
|---|---|---|---|---|---|---|---|---|---|---|---|---|---|---|
| testneg_1708 | 0 | 0 | 1.314085 | -3 | 3.775158 | -3 | 0.10236 | 0.09622 | 0.07297 | 0.12788 | 28.66971 | 0.062861111 | 0 | Neutral |
| testneg_1709 | 0 | 0.32 | 0.353578 | -3 | 6.561184 | -3 | 0.62645 | 0.12788 | 0.46056 | 0.12785 | 18.76159 | 0.161111111 | 0 | Neutral |
| testneg_1710 | 0.15 | -0.28 | -0.177889 | -3 | 1.474779 | 1.474779 | 0.29965 | 0.26486 | 0.0882 | 0.12788 | 10.83333 | 0.060346476 | 0 | Neutral |
| testneg_1711 | 0 | -0.01 | -0.041384 | -3 | 1.374903 | -3 | 0.00451 | 0.2022 | 0.50649 | 0.11394 | 30.0251 | 0.070206383 | 0 | Neutral |
| testneg_1712 | -0.09 | 0.3 | -2.446934 | -3 | -0.175371 | -3 | 0.46664 | 0.09008 | 0.21399 | 0.10708 | 21.5133 | 0.04 | 0 | Neutral |
| testneg_1713 | 0.21 | 0.15 | 6.548309 | -3 | 0.146037 | -0.782399 | 0.16777 | 0.1749 | 0.19898 | 0.12788 | 20.27778 | 0.231212945 | 1.2 | Neutral |
| testneg_1714 | 0.04 | -0.23 | 6.293797 | -3 | 2.390877 | -3 | 0.11072 | 0.38259 | 0.43691 | 0.12794 | 10.83333 | 0.481837607 | 48.6 | Neutral |
| testneg_1715 | -0.03 | -0.1 | 1.285186 | -3 | -1.650482 | -3 | 0.35957 | 0.11476 | 0.05095 | 0.08353 | 21.13948 | 0.02 | 0 | Neutral |
| testneg_1716 | -0.02 | -0.45 | 4.751068 | -3 | -0.061002 | -0.061002 | 0.9449 | 0.9378 | 0.35561 | 0.08845 | 22.83164 | 0.19 | 0 | Neutral |
| testneg_1717 | -0.08 | -0.03 | 2.560236 | -3 | 0.233598 | 0.233598 | 0.17728 | 0.46704 | 0.25804 | 0.11876 | 29.16667 | 0.070442308 | 0 | Neutral |
| testneg_1718 | -0.08 | -0.03 | 2.560236 | -3 | 4.494511 | -3 | 0.17728 | 0.46704 | 0.19898 | 0.12788 | 10.41667 | 0.279904762 | 1.2 | Neutral |
| testneg_1719 | 0 | 0 | 1.630014 | -3 | 2.08939 | -3 | 0.1109 | 0.17295 | 0.09633 | 0.12788 | 26.52342 | 0.04 | 0 | Neutral |
| testneg_1720 | 0 | 0 | 1.069792 | -3 | 1.715954 | -3 | 0.18869 | 0.23984 | 0.13126 | 0.11628 | 18.33333 | 0.010909091 | 0 | Neutral |
| testneg_1721 | 0 | 0 | 5.099298 | -3 | 6.60851 | -3 | 0.18498 | 0.12788 | 0.06541 | 0.12012 | 26.84685 | 0.418392857 | 24.4 | Neutral |
| testneg_1722 | 0 | 0 | 0.376038 | -3 | 6.320292 | -3 | 0.18485 | 0.40044 | 0.32188 | 0.08974 | 20.27778 | 0.223479853 | 0.4 | Neutral |
| testneg_1723 | 0 | -0.32 | 6.71239 | -3 | 3.132136 | -3 | 0.68568 | 0.65219 | 0.0105 | 0.12788 | 20.67497 | 0.549443956 | 74.8 | Disease_causing |
| testneg_1724 | 0 | 0 | 1.587314 | -3 | 1.928887 | -3.226454 | 0.20457 | 0.12378 | 0.05324 | 0.19446 | 18.33333 | 0.051452381 | 0 | Neutral |
| testneg_1725 | 0.18 | -1.05 | 2.729607 | -3 | 0.999127 | -3 | 0.29739 | 0.10893 | 0.08183 | 0.11603 | 6.94444 | 0.03 | 0 | Neutral |
| testneg_1726 | -0.18 | -0.16 | -0.566382 | -3 | 2.754446 | -3 | 0.23152 | 0.09962 | 0.63798 | 0.19967 | 20.69982 | 0.19 | 0 | Neutral |
| testneg_1727 | 0.07 | 0.38 | -3.472929 | -3 | 5.824607 | -3 | 0.4097 | 0.36733 | 0.03391 | 0.09745 | 33.40108 | 0.244375 | 0.2 | Neutral |
| testneg_1728 | -0.05 | -0.29 | 4.882717 | -3 | -0.08357 | -3 | 0.62339 | 0.19592 | 0.19898 | 0.09533 | 17.18434 | 0.09 | 0 | Neutral |
| testneg_1729 | 0.09 | -0.3 | 1.960992 | -3 | 2.97091 | -3 | 0.15486 | 0.28768 | 0.0961 | 0.09786 | 18.88889 | 0.110342745 | 0 | Neutral |
| testneg_1730 | 0.07 | 0.01 | 5.804215 | -3 | 0.127388 | -3 | 0.08067 | 0.34914 | 0.21967 | 0.09467 | 47.31826 | 0.23975 | 0.4 | Neutral |
| testneg_1731 | 0 | 0 | 0.619175 | 0.619175 | 6.983534 | -3 | 0.68138 | 0.20944 | 0.62645 | 0.12788 | 27.07912 | 0.303704283 | 4.6 | Neutral |
| testneg_1732 | 0 | 0 | 1.23713 | -3 | -0.081322 | -3 | 0.22489 | 0.16646 | 0.54489 | 0.09921 | 20.19985 | 0.02 | 0 | Neutral |
| testneg_1733 | 0 | 0 | -0.961702 | -0.961702 | -1.247889 | -3 | 0.46979 | 0.1705 | 0.27577 | 0.09812 | 15.80978 | 0.02 | 0 | Neutral |
| testneg_1734 | -0.0118182 | -0.0598182 | 4.426423 | -3 | 0.797761 | -3 | 0.08041 | 0.21802 | 0.71276 | 0.88182 | 10.41667 | 0.368854839 | 9.6 | Neutral |
| testneg_1735 | -0.18 | 1.05 | 2.778725 | 2.778725 | 0.603563 | -3 | 0.33528 | 0.29463 | 0.10058 | 0.13882 | 17.08333 | 0.5 | 56 | Disease_causing |
| testneg_1736 | -0.09 | 0.25 | 3.484844 | -3 | 0.628936 | -3 | 0.19898 | 0.12788 | 0.15152 | 0.12645 | 14.31225 | 0.03 | 0 | Neutral |
| testneg_1737 | -0.21 | -0.15 | 0.197741 | -3 | 4.786205 | -3 | 0.31923 | 0.17492 | 0.70872 | 0.09884 | 9.18689 | 0.250846154 | 0.6 | Neutral |
| testneg_1738 | 0 | 0 | 1.255675 | -3 | 1.214666 | -3 | 0.20675 | 0.13709 | 0.40021 | 0.21702 | 10.41667 | 0.021748214 | 0 | Neutral |
| testneg_1739 | 0.09 | -0.3 | 3.49868 | -3 | 4.404978 | -3 | 0.18282 | 0.12022 | 0.03505 | 0.0907 | 17.21847 | 0.129800662 | 0 | Neutral |
| testneg_1740 | 0.15 | -0.28 | 1.477409 | -3 | 4.83533 | -3 | 0.55527 | 0.12788 | 0.23975 | 0.13861 | 10.83333 | 0.288055556 | 1.6 | Neutral |
| testneg_1741 | -0.01 | -0.8 | 7.038327 | -3 | 2.132862 | -3 | 0.30367 | 0.41964 | 0.11692 | 0.08324 | 17.21847 | 0.483685897 | 49 | Neutral |
| testneg_1742 | 0.08 | 0.03 | -1.481985 | -3 | 2.642734 | -3 | 0.18416 | 0.13292 | 0.19898 | 0.12788 | 13.08943 | 0.06 | 0 | Neutral |
| testneg_1743 | 0.03 | 1.22 | 6.59818 | -3 | 0.196499 | -3 | 0.20152 | 0.09895 | 0.29098 | 0.10482 | 17.88673 | 0.21 | 0.2 | Neutral |
| testneg_1744 | 0 | 0 | -0.417855 | -3 | 4.741449 | -3 | 0.99543 | 0.12788 | 0.10241 | 0.08165 | 21.52409 | 0.12 | 0 | Neutral |
| testneg_1745 | 0.03 | 1.22 | 2.297873 | -3 | 1.432122 | -3 | 0.31168 | 0.12788 | 0.06301 | 0.32138 | 12.29167 | 0.124852941 | 0 | Neutral |
| testneg_1746 | -0.14 | 1.01 | 3.095581 | -3 | -0.534744 | -3 | 0.41269 | 0.19135 | 0.07923 | 0.08035 | 24.20397 | 0.1 | 0 | Neutral |
| testneg_1747 | -0.05 | -0.29 | -2.69333 | -3 | 4.453269 | -3 | 0.0984 | 0.1489 | 0.20507 | 0.11381 | 15.27491 | 0.2 | 0.2 | Neutral |
| testneg_1748 | 0.05 | 0.29 | 3.783684 | -3 | 1.135659 | -3 | 0.39399 | 0.12788 | 0.21303 | 0.09348 | 22.5469 | 0.03 | 0 | Neutral |
| testneg_1749 | -0.02 | 0.28 | 3.559312 | -3 | 0.389761 | -3 | 0.39964 | 0.22293 | 0.73601 | 0.26791 | 17.77778 | 0.259035326 | 0.4 | Neutral |
| testneg_1750 | 0.01 | 0.8 | 0.282453 | -3 | 1.333773 | -3 | 0.42366 | 0.14372 | 0.15294 | 0.12788 | 28.47222 | 0.03 | 0 | Neutral |
| testneg_1751 | 0 | 0 | 1.012321 | -3 | 4.901007 | -3 | 0.21805 | 0.12777 | 0.227 | 0.14389 | 17.29129 | 0.198130647 | 0 | Neutral |
| testneg_1752 | 0.17 | 0.38 | 0.177046 | -3 | 1.163857 | -3 | 0.21029 | 0.27753 | 0.09067 | 0.10671 | 21.53707 | 0.07 | 0 | Neutral |
| testneg_1753 | 0 | 0 | 1.831141 | -3 | 6.549241 | -3 | 0.99531 | 0.86203 | 0.04142 | 0.09564 | 4.44444 | 0.251076087 | 0.4 | Neutral |
| testneg_1754 | 0.01 | 0.79 | -1.428127 | -3 | 1.104931 | -3 | 0.11564 | 0.1184 | 0.06708 | 0.09112 | 21.38889 | 0.02019295 | 0 | Neutral |
| testneg_1755 | -0.02 | -0.41 | 3.657869 | -3 | 0.851144 | -3 | 0.18834 | 0.12788 | 0.10004 | 0.12788 | 9.81765 | 0.030762408 | 0 | Neutral |
| testneg_1756 | 0.08 | 0.03 | 1.795418 | -3 | 3.966623 | -3 | 0.11055 | 0.19829 | 0.16737 | 0.12788 | 14.56463 | 0.243666667 | 0.2 | Neutral |
| testneg_1757 | -0.08 | -0.03 | 2.342086 | -3 | 2.51331 | -3 | 0.5901 | 0.19444 | 0.14601 | 0.12788 | 5 | 0.061666667 | 0 | Neutral |
| testneg_1758 | -0.07 | -0.38 | 1.656921 | -3 | -0.750318 | -3 | 0.19898 | 0.12788 | 0.26068 | 0.46457 | 20.28889 | 0.08 | 0 | Neutral |
| testneg_1759 | -0.08 | -0.03 | 5.967322 | -3 | 0.307774 | 0.307774 | 0.25312 | 0.12788 | 0.19898 | 0.0522 | 13.51351 | 0.15 | 0 | Neutral |
| testneg_1760 | -0.02 | 0.63 | 2.451821 | -3 | 5.494664 | -3 | 0.25025 | 0.18924 | 0.19898 | 0.12788 | 22.25709 | 0.280865424 | 0.4 | Neutral |
| testneg_1761 | 0 | 0 | 1.371909 | -3 | 0.3368 | -3 | 0.47733 | 0.08388 | 0.08837 | 0.10939 | 29.61905 | 0.02 | 0 | Neutral |
| testneg_1762 | 0 | 0 | 0.789516 | 0.600565 | 2.08807 | -3 | 0.99564 | 0.25794 | 0.80264 | 0.11567 | 10 | 0.051266026 | 0 | Neutral |
| testneg_1763 | 0 | -0.01 | 0.601639 | -3 | -0.33067 | -3 | 0.08373 | 0.09191 | 0.17065 | 0.11746 | 23.19121 | 0.01 | 0 | Neutral |
| testneg_1764 | 0 | 0 | 1.185651 | -3 | 0.216154 | -3 | 0.87813 | 0.23021 | 0.17808 | 0.12788 | 11.25 | 0.01023953 | 0 | Neutral |
| testneg_1765 | 0 | 0 | -0.301372 | -3 | 0.413122 | -3 | 0.11019 | 0.08132 | 0.32531 | 0.12351 | 27.37041 | 0.01 | 0 | Neutral |
| testneg_1766 | 0.21 | 0.15 | 3.266344 | -3 | 7.835698 | -3 | 0.12977 | 0.1128 | 0.01704 | 0.05734 | 9.21141 | 0.409416667 | 21 | Neutral |
| testneg_1767 | 0 | -0.01 | 0.611606 | -3 | 1.999648 | 1.999648 | 0.9149 | 0.27793 | 0.12626 | 0.10279 | 15.85874 | 0.042071429 | 0 | Neutral |
| testneg_1768 | 0 | 0 | 0.007192 | -3 | 3.779532 | -3 | 0.13011 | 0.14157 | 0.11373 | 0.10739 | 10.22898 | 0.08127537 | 0 | Neutral |
| testneg_1769 | 0 | -0.01 | -0.496432 | -3 | -0.625744 | -3 | 0.06247 | 0.14184 | 0.19898 | 0.12788 | 10 | 0.02 | 0 | Neutral |
| testneg_1770 | 0.14 | 0.54 | 5.412626 | -3 | -1.722922 | -3 | 0.24327 | 0.12192 | 0.06704 | 0.1068 | 19.37853 | 0.14 | 0 | Neutral |
| testneg_1771 | -0.07 | -0.38 | 4.428125 | -3 | 1.096737 | 1.096737 | 0.76693 | 0.19444 | 0.23071 | 0.09582 | 11.875 | 0.08 | 0 | Neutral |
| testneg_1772 | 0 | 0 | -0.470651 | -3 | 0.413122 | -3 | 0.28063 | 0.40063 | 0.32531 | 0.12351 | 18.04026 | 0.020575808 | 0 | Neutral |
| testneg_1773 | -0.15 | 0.82 | 5.378906 | -3 | 0.313674 | -3 | 0.16326 | 0.09256 | 0.26812 | 0.30371 | 13.51351 | 0.369010417 | 11 | Neutral |
| testneg_1774 | 0 | 0 | 1.000182 | -3 | 6.097788 | -3 | 0.82455 | 0.2093 | 0.22944 | 0.09039 | 20 | 0.123758366 | 0 | Neutral |
| testneg_1775 | -0.07 | -0.38 | 2.063043 | -3 | 1.323079 | -3 | 0.90296 | 0.28521 | 0.5103 | 0.10923 | 18.71486 | 0.03 | 0 | Neutral |
| testneg_1776 | -0.02 | -0.45 | 6.344462 | -3 | 4.745256 | -2.031435 | 0.29624 | 0.1067 | 0.09306 | 0.12788 | 21.34897 | 0.410211111 | 25.6 | Neutral |
| testneg_1777 | 0.01 | -1.44 | 4.523358 | -3 | 7.485974 | -3 | 0.21201 | 0.20962 | 0.0968 | 0.15592 | 12.30981 | 0.835038647 | 100 | Disease_causing |
| testneg_1778 | 0.02 | 0.45 | -0.664965 | -3 | 0.72503 | -3 | 0.12786 | 0.72066 | 0.56862 | 0.12788 | 19.31241 | 0.1 | 0 | Neutral |
| testneg_1779 | 0.14 | -0.69 | 2.526774 | -3 | 0.426905 | -3 | 0.16797 | 0.12788 | 0.19531 | 0.18996 | 28.84995 | 0.08 | 0 | Neutral |
| testneg_1780 | 0.07 | 0.7 | 5.438117 | -3 | 5.582498 | -3 | 0.12463 | 0.12788 | 0.85829 | 0.4292 | 11.11011 | 0.649635679 | 93.8 | Disease_causing |
| testneg_1781 | 0 | 0 | 5.815666 | 5.815666 | 1.550276 | -3 | 0.34109 | 0.2167 | 0.18142 | 0.21216 | 10.83333 | 0.877801222 | 100 | Disease_causing |
| testneg_1782 | -0.15 | -0.04 | 2.846922 | -3 | 2.790651 | -3 | 0.18081 | 0.1258 | 0.19898 | 0.10116 | 10.83333 | 0.11 | 0 | Neutral |
| testneg_1783 | 0 | 0 | 0.917232 | -3 | 0.264978 | -3 | 0.25556 | 0.09342 | 0.56817 | 0.10766 | 36.21309 | 0.01 | 0 | Neutral |
| testneg_1784 | -0.03 | -1.22 | 2.92942 | -3 | 1.213226 | -3 | 0.15107 | 0.10457 | 0.19898 | 0.12788 | 10 | 0.04 | 0 | Neutral |
| testneg_1785 | 0.22 | -2.66 | 6.248531 | -3 | 0.892974 | 0.892974 | 0.12839 | 0.09503 | 0.06986 | 0.12788 | 15.16525 | 0.15 | 0 | Neutral |
| testneg_1786 | 0 | 0 | 0.094005 | -3 | 1.282461 | -3 | 0.64302 | 0.11262 | 0.59131 | 0.66216 | 9.89855 | 0.023651629 | 0 | Neutral |
| testneg_1787 | 0.03 | -1.03 | 3.976603 | -3 | -0.767748 | -3 | 0.64125 | 0.28792 | 0.16116 | 0.08314 | 14.04389 | 0.078285714 | 0 | Neutral |
| testneg_1788 | 0 | 0 | 0.29846 | -3 | 5.417151 | -3 | 0.07543 | 0.10211 | 0.12259 | 0.1041 | 12.77241 | 0.061547619 | 0 | Neutral |
| testneg_1789 | -0.03 | -0.1 | 0.852085 | -3 | 4.833511 | -3 | 0.56554 | 0.76978 | 0.19898 | 0.07312 | 26.95989 | 0.17 | 0 | Neutral |
| testneg_1790 | 0 | 0 | 1.716798 | -3 | 2.412319 | -3 | 0.70349 | 0.12788 | 0.79692 | 0.51979 | 5 | 0.06 | 0 | Neutral |
| testneg_1791 | 0.04 | -0.23 | 0.493221 | -3 | 1.551505 | -3 | 0.13325 | 0.12788 | 0.05413 | 0.05691 | 19.84929 | 0.01 | 0 | Neutral |
| testneg_1792 | 0.15 | -0.28 | 2.017059 | -3 | 5.443984 | -3 | 0.27275 | 0.11752 | 0.28271 | 0.60265 | 11.25 | 0.427475455 | 26 | Neutral |
| testneg_1793 | -0.09 | 0.3 | 2.569627 | -3 | 2.950487 | -3 | 0.54909 | 0.85056 | 0.14495 | 0.37127 | 39.04612 | 0.463630952 | 38.2 | Neutral |
| testneg_1794 | 0 | 0 | 1.34425 | -3 | -0.111883 | -0.111883 | 0.16299 | 0.10454 | 0.06313 | 0.12788 | 15.70637 | 0.000322581 | 0 | Neutral |
| testneg_1795 | -0.01 | 1.3 | 1.784506 | -3 | 0.27272 | -3 | 0.27629 | 0.11166 | 0.02967 | 0.06605 | 28.13539 | 0.050034965 | 0 | Neutral |
| testneg_1796 | 0 | 0 | -0.920517 | -3 | -2.237532 | -3 | 0.6975 | 0.53794 | 0.42836 | 0.12788 | 11.875 | 0.03 | 0 | Neutral |
| testneg_1797 | -0.04 | -2.01 | 1.963276 | -3 | 0.877291 | -3 | 0.21355 | 0.12788 | 0.86166 | 0.14254 | 13.37927 | 0.072 | 0 | Neutral |
| testneg_1798 | 0.03 | 0.1 | 0.502322 | -3 | -2.702977 | -3 | 0.13696 | 0.09566 | 0.19898 | 0.56316 | 22.92484 | 0.09 | 0 | Neutral |
| testneg_1799 | 0 | 0 | 0.245964 | -3 | 0.902524 | -3 | 0.74099 | 0.44443 | 0.21941 | 0.46348 | 10 | 0.074439871 | 0 | Neutral |
| testneg_1800 | 0.02 | 0.45 | 3.429131 | -3 | 1.061712 | -3 | 0.03499 | 0.12971 | 0.19898 | 0.12788 | 22.03773 | 0.07 | 0 | Neutral |
| testneg_1801 | 0.09 | -0.25 | 1.841476 | -3 | 0.975747 | -3 | 0.13295 | 0.12914 | 0.53447 | 0.10345 | 13.46801 | 0.01 | 0 | Neutral |
| testneg_1802 | 0 | 0 | 0.402508 | 0.402508 | 2.6592 | -3 | 0.29794 | 0.11106 | 0.24524 | 0.10803 | 16.875 | 0.02 | 0 | Neutral |
| testneg_1803 | 0 | -0.01 | 2.438214 | -3 | 0.240706 | -3 | 0.16741 | 0.10762 | 0.25831 | 0.10988 | 5.65457 | 0 | 0 | Neutral |
| testneg_1804 | -0.05 | 0.81 | 1.623152 | -3 | 4.845098 | -3 | 0.19898 | 0.12788 | 0.09093 | 0.28643 | 30.55556 | 0.343086805 | 6.4 | Neutral |
| testneg_1805 | 0 | -0.01 | 1.801091 | -3 | 5.700045 | -3 | 0.39459 | 0.12788 | 0.12682 | 0.11357 | 18.21789 | 0.070691303 | 0 | Neutral |
| testneg_1806 | 0 | 0 | 0.036223 | -3 | 0.105909 | -3 | 0.9355 | 0.40418 | 0.24765 | 0.32858 | 5.69444 | 0.066085274 | 0 | Neutral |
| testneg_1807 | -0.03 | -0.1 | -0.76618 | -3 | 1.252341 | -3 | 0.1503 | 0.12788 | 0.38072 | 0.15993 | 12.29167 | 0.07 | 0 | Neutral |
| testneg_1808 | -0.07 | -0.33 | 1.810073 | -3 | 0.734682 | -3 | 0.9662 | 0.28104 | 0.12576 | 0.08486 | 9.67871 | 0.020173893 | 0 | Neutral |
| testneg_1809 | 0.05 | 0.29 | 2.273291 | -3 | 0.252727 | -3 | 0.19898 | 0.12788 | 0.13985 | 0.08349 | 12.34649 | 0.003333333 | 0 | Neutral |
| testneg_1810 | -0.08 | -0.03 | 2.788201 | -3 | -0.199284 | -3 | 0.25009 | 0.11017 | 0.11688 | 0.13208 | 23.07457 | 0.08 | 0 | Neutral |
| testneg_1811 | 0.18 | -1.05 | 7.047951 | 3.942193 | -0.669222 | -3 | 0.14668 | 0.12788 | 0.22225 | 0.11186 | 5 | 0.6 | 87.2 | Disease_causing |
| testneg_1812 | 0.14 | -1.01 | 1.85098 | -3 | 5.387063 | -3 | 0.06957 | 0.07707 | 0.10244 | 0.10107 | 29.21005 | 0.11 | 0 | Neutral |
| testneg_1813 | 0 | 0 | 0.894121 | -3 | 1.270956 | -3 | 0.02146 | 0.10351 | 0.39019 | 0.18821 | 11.25 | 0.071483516 | 0 | Neutral |
| testneg_1814 | 0.07 | 0.38 | 1.715954 | -3 | 7.532012 | -3 | 0.13126 | 0.11628 | 0.99704 | 0.17463 | 10.41667 | 0.427438388 | 27.6 | Neutral |
| testneg_1815 | -0.02 | -0.45 | 5.568518 | -3 | 5.100385 | -3 | 0.72657 | 0.11262 | 0.11418 | 0.2009 | 33.06414 | 0.66525 | 96.2 | Disease_causing |
| testneg_1816 | -0.16 | -0.08 | 5.960242 | -3 | 0.483862 | -3 | 0.1304 | 0.12788 | 0.21551 | 0.12788 | 19.44444 | 0.181785714 | 0 | Neutral |
| testneg_1817 | -0.15 | 0.28 | 2.650636 | -3 | 3.355302 | -3 | 0.82595 | 0.52905 | 0.21348 | 0.10616 | 10.41667 | 0.248700397 | 0.2 | Neutral |
| testneg_1818 | -0.08 | -0.03 | 7.833352 | -3 | 0.947631 | -3 | 0.08067 | 0.34914 | 0.19898 | 0.12788 | 34.72222 | 0.47 | 40.8 | Neutral |
| testneg_1819 | 0 | -0.01 | 1.129155 | -3 | 5.213355 | -3 | 0.16835 | 0.09282 | 0.37094 | 0.10762 | 21.36401 | 0.040144632 | 0 | Neutral |
| testneg_1820 | 0.22 | -2.66 | 5.257983 | -3 | 2.655355 | 2.655355 | 0.12689 | 0.13376 | 0.16436 | 0.19109 | 24.62124 | 0.556458333 | 74.4 | Disease_causing |
| testneg_1821 | 0 | 0 | 1.024939 | -3 | 6.883386 | -3 | 0.17529 | 0.09662 | 0.1594 | 0.22002 | 33.62403 | 0.36724445 | 12.8 | Neutral |
| testneg_1822 | -0.16 | 1.37 | 2.25398 | -3 | 3.390375 | -3 | 0.32022 | 0.40347 | 0.25462 | 0.12788 | 18.6768 | 0.35 | 4 | Neutral |
| testneg_1823 | 0 | 0 | 0.505411 | -3 | 1.237933 | -3 | 0.19898 | 0.09431 | 0.04063 | 0.05848 | 38.94231 | 0.01 | 0 | Neutral |
| testneg_1824 | -0.03 | -0.1 | 4.162376 | -3 | 1.888775 | -3 | 0.71511 | 0.34803 | 0.09633 | 0.12788 | 10.83333 | 0.14125 | 0 | Neutral |
| testneg_1825 | 0 | -0.32 | 1.514021 | -3 | 2.24203 | -3 | 0.02791 | 0.0854 | 0.08059 | 0.09576 | 38.98776 | 0.05 | 0 | Neutral |
| testneg_1826 | 0 | 0 | 0.739398 | -3 | -2.576045 | -2.576045 | 0.48906 | 0.11262 | 0.25275 | 0.12788 | 23.47941 | 0.01 | 0 | Neutral |
| testneg_1827 | 0.02 | -0.36 | -0.551863 | -3 | -0.133236 | -3 | 0.51037 | 0.10999 | 0.18328 | 0.12788 | 21.81773 | 0.02 | 0 | Neutral |
| testneg_1828 | 0 | 0 | 1.201989 | -3 | -0.116772 | -3 | 0.2231 | 0.22973 | 0.16741 | 0.10762 | 13.77192 | 0.010121431 | 0 | Neutral |
| testneg_1829 | 0 | -0.01 | 2.227487 | -3 | 0.556885 | -3 | 0.85984 | 0.40044 | 0.78664 | 0.11725 | 10.83333 | 0.020084087 | 0 | Neutral |

| | | | | | | | | | | | | | | |
|---|---|---|---|---|---|---|---|---|---|---|---|---|---|---|
| testneg_1830 | -0.02 | 0.68 | 3.109082 | -3 | -0.3227 | -3 | 0.31244 | 0.4615 | 0.25893 | 0.10353 | 10.45412 | 0.104491694 | 0 | Neutral |
| testneg_1831 | 0.04 | -0.23 | 3.599958 | -3 | -0.445993 | -0.445993 | 0.40491 | 0.17596 | 0.18324 | 0.13041 | 11.25 | 0.08 | 0 | Neutral |
| testneg_1832 | -0.07 | -0.38 | 3.994714 | -3 | 5.372461 | -3 | 0.339 | 0.13169 | 0.13796 | 0.27869 | 22.0923 | 0.703057079 | 99.2 | Disease_causing |
| testneg_1833 | 0 | 0 | -0.378226 | -3 | 5.924715 | -3 | 0.99668 | 0.30319 | 0.61924 | 0.11925 | 12.33727 | 0.136376391 | 0 | Neutral |
| testneg_1834 | 0 | 0 | -0.13661 | -3 | -0.078944 | -3 | 0.23624 | 0.12788 | 0.13255 | 0.0962 | 14.51389 | 0.000164634 | 0 | Neutral |
| testneg_1835 | 0.07 | -0.31 | 5.981941 | -3 | 0.079033 | -3 | 0.9541 | 0.96732 | 0.25827 | 0.21386 | 10.22898 | 0.55 | 68.6 | Disease_causing |
| testneg_1836 | 0.03 | -0.85 | -1.390553 | -3 | 0.589367 | -3 | 0.08789 | 0.6361 | 0.49064 | 0.12294 | 14.16667 | 0.090259784 | 0 | Neutral |
| testneg_1837 | -0.0083217 | -0.1418881 | -0.105139 | -3 | 0.721819 | -3 | 0.08489 | 0.12788 | 0.45342 | 0.14389 | 12.76592 | 0.04 | 0 | Neutral |
| testneg_1838 | -0.09 | 0.25 | 4.068872 | -3 | 0.610442 | -3 | 0.19898 | 0.12788 | 0.17786 | 0.15314 | 12.19458 | 0.168416612 | 0 | Neutral |
| testneg_1839 | 0.04 | -0.23 | 2.046649 | -3 | 0.436031 | -3 | 0.1014 | 0.12788 | 0.10977 | 0.31695 | 17.77909 | 0.1 | 0 | Neutral |
| testneg_1840 | -0.02 | -0.45 | 3.493501 | -3 | -2.655397 | -3 | 0.19898 | 0.10482 | 0.06296 | 0.09823 | 55.83765 | 0.05 | 0 | Neutral |
| testneg_1841 | 0.02 | -0.36 | 2.067777 | -3 | 0.234129 | -3 | 0.12408 | 0.08828 | 0.25758 | 0.2262 | 10.83333 | 0.049772727 | 0 | Neutral |
| testneg_1842 | 0.07 | 0.01 | 0.509453 | -3 | 2.141722 | -3 | 0.11587 | 0.12001 | 0.42872 | 0.12788 | 19.90385 | 0.010201927 | 0 | Neutral |
| testneg_1843 | 0 | -0.01 | -0.496432 | -3 | -0.401457 | -3 | 0.06247 | 0.14184 | 0.15155 | 0.09923 | 8.97898 | 0.02 | 0 | Neutral |
| testneg_1844 | 0.07 | 0.38 | -2.083041 | -3 | -1.203781 | -3 | 0.10383 | 0.08853 | 0.29523 | 0.27623 | 13.11828 | 0.061391414 | 0 | Neutral |
| testneg_1845 | 0 | 0 | 0.928885 | -3 | -0.10769 | -2.106562 | 0.39724 | 0.11058 | 0.27083 | 0.14313 | 17.77778 | 0.020186673 | 0 | Neutral |
| testneg_1846 | 0.03 | 0.12 | 3.066776 | -3 | -0.002747 | -3 | 0.2877 | 0.18291 | 0.07394 | 0.09408 | 21.52409 | 0.050107577 | 0 | Neutral |
| testneg_1847 | 0 | 0 | 1.358901 | -3 | 0.048986 | -3 | 0.26264 | 0.13812 | 0.0773 | 0.14062 | 10.41667 | 0.045667915 | 0 | Neutral |
| testneg_1848 | 0.06 | -0.57 | -0.888141 | -3 | 2.310369 | 2.310369 | 0.37637 | 0.12788 | 0.54895 | 0.15082 | 4.72222 | 0.121142857 | 0 | Neutral |
| testneg_1849 | 0 | 0 | 0.865243 | -3 | 0.390074 | -3 | 0.19898 | 0.25991 | 0.48615 | 0.14369 | 5.20285 | 0.041941777 | 0 | Neutral |
| testneg_1850 | 0.04 | 2.01 | 1.520495 | -3 | -1.047288 | -3 | 0.50921 | 0.21466 | 0.52005 | 0.12788 | 14.78155 | 0.13 | 0 | Neutral |
| testneg_1851 | 0.01 | 0.8 | -0.725111 | -0.725111 | 0.094735 | -3 | 0.05116 | 0.08996 | 0.024 | 0.07726 | 30.54981 | 0.06 | 0 | Neutral |
| testneg_1852 | 0.21 | 0.15 | 1.912237 | -3 | 4.625597 | -3 | 0.12511 | 0.11897 | 0.58292 | 0.87725 | 21.51158 | 0.39 | 13.2 | Neutral |
| testneg_1853 | 0 | 0 | 0.975549 | -3 | 3.384606 | -3 | 0.91619 | 0.10042 | 0.13001 | 0.13176 | 34.24591 | 0.150113636 | 0 | Neutral |
| testneg_1854 | 0.14 | -1.01 | 0.318992 | -3 | 1.572604 | -3 | 0.66799 | 0.12788 | 0.21967 | 0.09467 | 20.26596 | 0.03 | 0 | Neutral |
| testneg_1855 | 0 | 0 | -0.633568 | -3 | 0.465329 | 0.465329 | 0.23027 | 0.37171 | 0.11131 | 0.68435 | 7.46424 | 0.199933333 | 0 | Neutral |
| testneg_1856 | 0.03 | -1.03 | 4.395754 | -3 | 6.003823 | 5.545276 | 0.23879 | 0.12788 | 0.13369 | 0.09791 | 18.06147 | 0.51 | 61.6 | Disease_causing |
| testneg_1857 | 0 | 0 | 0.245113 | -3 | -0.317598 | -3 | 0.46399 | 0.15334 | 0.59321 | 0.59981 | 4.44444 | 0.038859127 | 0 | Neutral |
| testneg_1858 | -0.07 | -0.38 | 1.619888 | -3 | -1.752013 | -3 | 0.19477 | 0.12788 | 0.0609 | 0.47188 | 19.09347 | 0.100073529 | 0 | Neutral |
| testneg_1859 | -0.08 | -0.03 | 1.483649 | 1.483649 | 5.359975 | -3 | 0.47678 | 0.38085 | 0.12196 | 0.12788 | 21.52529 | 0.272352103 | 0.8 | Neutral |
| testneg_1860 | 0 | 0 | 0.864386 | -3 | 2.273291 | -3 | 0.18998 | 0.26095 | 0.19898 | 0.12788 | 10.83333 | 0.03 | 0 | Neutral |
| testneg_1861 | 0 | 0 | 1.212869 | -3 | 2.906492 | 0.554192 | 0.43282 | 0.10804 | 0.09323 | 0.16732 | 39.27199 | 0.174068342 | 0 | Neutral |
| testneg_1862 | 0 | 0 | -0.535508 | -3 | 1.175993 | -3 | 0.01196 | 0.12788 | 0.06781 | 0.09009 | 23.99208 | 0.04 | 0 | Neutral |
| testneg_1863 | 0 | -0.01 | -0.390012 | -3 | 0.322589 | -3 | 0.33292 | 0.17111 | 0.15992 | 0.12788 | 10.5 | 0.01 | 0 | Neutral |
| testneg_1864 | 0 | 0 | -0.338241 | -3 | 1.68139 | -3 | 0.26451 | 0.09729 | 0.4094 | 0.1944 | 10.41667 | 0.02375 | 0 | Neutral |
| testneg_1865 | 0.21 | 0.15 | 2.146258 | -3 | 1.00919 | -3 | 0.62339 | 0.19592 | 0.3414 | 0.13575 | 17.46678 | 0.15134465 | 0 | Neutral |
| testneg_1866 | 0 | 0 | 0.42612 | -3 | 3.99034 | -3 | 0.10133 | 0.32155 | 0.12957 | 0.12788 | 21.46199 | 0.177513158 | 0 | Neutral |
| testneg_1867 | 0.08 | 0.03 | -1.481985 | -3 | 1.841607 | -3 | 0.18416 | 0.13292 | 0.17977 | 0.12788 | 40.38409 | 0.04 | 0 | Neutral |
| testneg_1868 | 0 | 0 | -0.549073 | -3 | -0.692965 | -3 | 0.38016 | 0.14503 | 0.19898 | 0.12788 | 28.30044 | 0.010645833 | 0 | Neutral |
| testneg_1869 | 0.08 | 0.03 | -1.242275 | -3 | 0.480554 | -3 | 0.7597 | 0.12788 | 0.31067 | 0.35414 | 4.72222 | 0.080555556 | 0 | Neutral |
| testneg_1870 | 0 | 0 | 0.213657 | -3 | -0.126067 | -3 | 0.3769 | 0.12788 | 0.19898 | 0.09171 | 23.23468 | 0.000540936 | 0 | Neutral |
| testneg_1871 | 0 | 0 | 1.806647 | -3 | 3.199326 | -3 | 0.21728 | 0.1132 | 0.24038 | 0.15296 | 18.88889 | 0.129488324 | 0 | Neutral |
| testneg_1872 | 0.19 | 0.51 | 4.472767 | -3 | 3.046889 | -3 | 0.12853 | 0.08565 | 0.1066 | 0.12788 | 16.60044 | 0.19 | 0 | Neutral |
| testneg_1873 | -0.01 | -0.8 | 3.446913 | -3 | 1.81348 | -3 | 0.32531 | 0.12351 | 0.08332 | 0.09046 | 10.24108 | 0.04 | 0 | Neutral |
| testneg_1874 | -0.07 | -0.01 | 2.291931 | -3 | 1.599412 | -3 | 0.99998 | 0.86471 | 0.13099 | 0.17453 | 10.41667 | 0.182890467 | 0 | Neutral |
| testneg_1875 | 0 | 0 | 2.044018 | -3 | 3.896665 | -3 | 0.63431 | 0.09837 | 0.18305 | 0.12743 | 22.85088 | 0.081125 | 0 | Neutral |
| testneg_1876 | 0 | 0 | 0.305495 | -3 | 1.773131 | -3 | 0.98255 | 0.98129 | 0.12848 | 0.65153 | 10.83333 | 0.202505952 | 0 | Neutral |
| testneg_1877 | 0.02 | -0.28 | 1.782871 | -3 | 1.381201 | -3 | 0.61834 | 0.10784 | 0.2016 | 0.13956 | 32.78013 | 0.05025 | 0 | Neutral |
| testneg_1878 | -0.07 | -0.38 | 0.882446 | -3 | 3.725973 | -3 | 0.19301 | 0.12955 | 0.27058 | 0.13363 | 14.91442 | 0.22331205 | 0.2 | Neutral |
| testneg_1879 | 0.05 | 0.29 | 1.0486 | -3 | 0.379358 | -3 | 0.38375 | 0.09256 | 0.19898 | 0.10996 | 12.13941 | 0.01 | 0 | Neutral |
| testneg_1880 | 0 | 0 | 2.097362 | -3 | -2.511821 | -3 | 0.50063 | 0.154 | 0.13873 | 0.12788 | 21.26172 | 0.03 | 0 | Neutral |
| testneg_1881 | 0 | 0 | 2.119414 | -3 | 1.940661 | -3 | 0.25744 | 0.2162 | 0.25151 | 0.12788 | 28.24632 | 0.03 | 0 | Neutral |
| testneg_1882 | 0 | 0 | 2.100416 | -3 | -1.775505 | -3 | 0.22255 | 0.10057 | 0.17303 | 0.18733 | 14.56604 | 0.04 | 0 | Neutral |
| testneg_1883 | -0.06 | -0.18 | -1.413197 | -3 | -2.689919 | -3 | 0.11172 | 0.10034 | 0.10066 | 0.17927 | 15.05208 | 0.080702007 | 0 | Neutral |
| testneg_1884 | -0.22 | 2.66 | 6.020917 | -3 | 0.865975 | -3 | 0.1589 | 0.10761 | 0.2306 | 0.0952 | 8.05556 | 0.27 | 2.6 | Neutral |
| testneg_1885 | 0.02 | 0.45 | 5.194939 | -3 | 3.130587 | -3 | 0.03631 | 0.80564 | 0.05027 | 0.10981 | 19.44444 | 0.41 | 19.6 | Neutral |
| testneg_1886 | 0.09 | -0.25 | -0.159404 | -3 | 0.317105 | -3 | 0.95319 | 0.20297 | 0.19898 | 0.12788 | 20.77349 | 0.05 | 0 | Neutral |
| testneg_1887 | 0 | 0 | 0.034609 | -3 | 1.369093 | -3 | 0.42033 | 0.17821 | 0.23292 | 0.15913 | 10.41667 | 0.06135147 | 0 | Neutral |
| testneg_1888 | -0.02 | -0.41 | 0.579456 | 0.579456 | 0.54521 | -3 | 0.47678 | 0.38085 | 0.3718 | 0.2248 | 29.86111 | 0.200066917 | 0 | Neutral |
| testneg_1889 | 0.19 | -0.25 | 3.925425 | -3 | -2.062376 | -2.062376 | 0.24239 | 0.111 | 0.11315 | 0.12788 | 12.58766 | 0.04 | 0 | Neutral |
| testneg_1890 | 0.03 | 0.12 | 2.467924 | -3 | 1.491454 | -3 | 0.84031 | 0.16749 | 0.1078 | 0.17394 | 4.44444 | 0.098094828 | 0 | Neutral |
| testneg_1891 | 0.05 | -0.81 | 6.05191 | -3 | 1.624358 | -3 | 0.38184 | 0.10478 | 0.16248 | 0.14352 | 12.40248 | 0.3 | 4.8 | Neutral |
| testneg_1892 | 0 | -0.01 | 0.492211 | -3 | 6.733255 | -3 | 0.20673 | 0.13236 | 0.37424 | 0.12788 | 11.98582 | 0.192746212 | 0.6 | Neutral |
| testneg_1893 | 0 | 0 | 0.272863 | -3 | -0.877392 | -3 | 0.19898 | 0.12788 | 0.1311 | 0.12667 | 16.48772 | 0 | 0 | Neutral |
| testneg_1894 | -0.07 | -0.38 | 0.868703 | -3 | 2.385228 | -3 | 0.06526 | 0.09018 | 0.09093 | 0.28643 | 21.38889 | 0.160802083 | 0 | Neutral |
| testneg_1895 | 0 | 0 | 0.244786 | -3 | 2.299454 | -3 | 0.08975 | 0.07399 | 0.19898 | 0.09316 | 18.82075 | 0.01 | 0 | Neutral |
| testneg_1896 | 0.19 | -0.27 | 0.86086 | -3 | 0.553729 | -3 | 0.12591 | 0.09845 | 0.44329 | 0.12788 | 10.30286 | 0.02 | 0 | Neutral |
| testneg_1897 | 0 | -0.01 | 0.494045 | -3 | 4.697901 | -3 | 0.1304 | 0.12788 | 0.11791 | 0.39607 | 21.41309 | 0.304496819 | 3.4 | Neutral |
| testneg_1898 | 0.17 | 0.38 | 4.608358 | -3 | 3.566467 | -3 | 0.04912 | 0.08574 | 0.17548 | 0.11384 | 29.9024 | 0.17 | 0 | Neutral |
| testneg_1899 | -0.08 | -0.03 | 2.768711 | 2.768711 | 0.838677 | -3 | 0.38324 | 0.25193 | 0.25884 | 0.12788 | 10.22898 | 0.319166667 | 4.6 | Neutral |
| testneg_1900 | 0.11 | -0.19 | 3.662322 | -3 | 0.573225 | -3 | 0.49745 | 0.1103 | 0.73947 | 0.2058 | 25.18369 | 0.146034091 | 0 | Neutral |
| testneg_1901 | -0.02 | 0.36 | 0.553539 | -3 | 6.591428 | -3 | 0.14511 | 0.12788 | 0.0501 | 0.12788 | 23.28544 | 0.206666667 | 0 | Neutral |
| testneg_1902 | 0.03 | 0.12 | 3.15417 | -3 | -0.360768 | -0.360768 | 0.25764 | 0.53109 | 0.47147 | 0.09503 | 19.11368 | 0.11 | 0 | Neutral |
| testneg_1903 | 0 | 0 | 0.990948 | -3 | 5.412025 | -3 | 0.31859 | 0.11262 | 0.1174 | 0.12788 | 13.79561 | 0.058796791 | 0 | Neutral |
| testneg_1904 | -0.0236364 | 0.26490909 | 11.240696 | -3 | 3.099902 | -3 | 0.10241 | 0.08165 | 0.15822 | 0.12788 | 15.88099 | 0.51 | 60.2 | Disease_causing |
| testneg_1905 | -0.02 | 0.68 | -0.240423 | -3 | 4.293539 | -3 | 0.07864 | 0.12788 | 0.11891 | 0.11186 | 17.28802 | 0.096676446 | 0 | Neutral |
| testneg_1906 | 0 | 0 | 0.388922 | -3 | 0.827687 | -3 | 0.98107 | 0.70722 | 0.39576 | 0.10761 | 16.8018 | 0.030331105 | 0 | Neutral |
| testneg_1907 | 0.05 | 0.29 | 3.783684 | -3 | -1.3044 | -3 | 0.39399 | 0.12788 | 0.15217 | 0.12544 | 22.5469 | 0.03 | 0 | Neutral |
| testneg_1908 | -0.07 | -0.38 | 0.734729 | -3 | -0.165977 | -3 | 0.58587 | 0.09866 | 0.32144 | 0.11774 | 15.29221 | 0.01 | 0 | Neutral |
| testneg_1909 | 0.07 | 0.7 | 1.763064 | -3 | 1.89715 | -3 | 0.11873 | 0.10499 | 0.25957 | 0.06969 | 25.63139 | 0.01 | 0 | Neutral |
| testneg_1910 | 0.03 | 0.12 | 3.471969 | -3 | 2.859591 | -3 | 0.99859 | 0.12788 | 0.1576 | 0.12788 | 10.83333 | 0.15 | 0 | Neutral |
| testneg_1911 | 0 | 0 | -0.057988 | -3 | 2.022784 | -3 | 0.15713 | 0.3507 | 0.84616 | 0.1471 | 24.96124 | 0.13623612 | 0 | Neutral |
| testneg_1912 | -0.06 | -0.18 | -0.439044 | -3 | 0.542604 | -3 | 0.3797 | 0.1257 | 0.54213 | 0.09044 | 18.80317 | 0.01 | 0 | Neutral |
| testneg_1913 | 0 | 0 | -0.989153 | -3 | 1.687515 | -3 | 0.3783 | 0.08224 | 0.53424 | 0.08484 | 18.88889 | 0.01 | 0 | Neutral |
| testneg_1914 | -0.04 | 1.36 | -1.284127 | -3 | 5.518112 | -3 | 0.11096 | 0.10074 | 0.32977 | 0.15427 | 43.75115 | 0.34 | 6.8 | Neutral |
| testneg_1915 | 0 | 0 | 1.333395 | -3 | -0.140634 | -3 | 0.12259 | 0.1041 | 0.19898 | 0.12788 | 11.93907 | 0 | 0 | Neutral |
| testneg_1916 | 0.03 | -1.03 | 0.221 | -0.888141 | 0.953201 | -3 | 0.37637 | 0.12788 | 0.11688 | 0.13208 | 12.84759 | 0.061875 | 0 | Neutral |
| testneg_1917 | 0 | -0.32 | 0.284563 | -3 | 6.154212 | -3 | 0.39436 | 0.16446 | 0.09288 | 0.16617 | 13.51351 | 0.520259075 | 64.6 | Disease_causing |
| testneg_1918 | -0.08 | -0.03 | 0.630495 | 0.630495 | -0.049164 | -0.049164 | 0.14419 | 0.1668 | 0.12322 | 0.1246 | 31.84685 | 0.07 | 0 | Neutral |
| testneg_1919 | -0.18 | 1.05 | 2.778725 | 2.778725 | 2.663229 | -3 | 0.33528 | 0.29463 | 0.25275 | 0.12788 | 12.29167 | 0.45 | 35.6 | Neutral |
| testneg_1920 | 0 | 0 | 0.085558 | -3 | 1.260137 | -3 | 0.14643 | 0.12788 | 0.1214 | 0.0899 | 18.88889 | 0.000294181 | 0 | Neutral |
| testneg_1921 | 0.006 | -0.1058182 | 1.290794 | -3 | 4.176231 | -3 | 0.84365 | 0.12788 | 0.55315 | 0.09195 | 8.97898 | 0.06 | 0 | Neutral |
| testneg_1922 | -0.09 | 1.07 | 5.012896 | -3 | 3.730704 | -3 | 0.47166 | 0.13548 | 0.37951 | 0.19435 | 21.38889 | 0.631136218 | 95.2 | Disease_causing |
| testneg_1923 | 0 | 0 | 0.036223 | -3 | 5.422386 | -3 | 0.9355 | 0.40418 | 0.10723 | 0.33098 | 18.92511 | 0.416748717 | 26 | Neutral |
| testneg_1924 | 0 | 0 | 1.107432 | -3 | 2.793253 | -3 | 0.5726 | 0.1165 | 0.17903 | 0.37232 | 10.41667 | 0.100745301 | 0 | Neutral |
| testneg_1925 | 0 | 0 | -0.445161 | -0.445161 | 4.710694 | -3 | 0.09728 | 0.12788 | 0.19898 | 0.12788 | 49.3589 | 0.07 | 0 | Neutral |
| testneg_1926 | 0.21 | 0.15 | 6.256857 | -3 | -1.66039 | -3 | 0.65674 | 0.10572 | 0.06957 | 0.07707 | 54.58928 | 0.17 | 0 | Neutral |
| testneg_1927 | 0.03 | 0.12 | 1.131121 | -3 | 5.049155 | -3 | 0.96704 | 0.20571 | 0.30925 | 0.10335 | 8.97898 | 0.141929223 | 0 | Neutral |
| testneg_1928 | -0.02 | -0.45 | -0.197057 | -3 | 1.232806 | -3 | 0.16797 | 0.12788 | 0.49641 | 0.12614 | 16.84579 | 0.003541667 | 0 | Neutral |
| testneg_1929 | 0.18 | -1.05 | 6.918655 | 6.918655 | 1.326969 | -3 | 0.15155 | 0.09923 | 0.12642 | 0.11218 | 22.91823 | 0.59 | 88.4 | Disease_causing |
| testneg_1930 | 0.00442308 | -0.005641 | 2.309347 | -3 | 0.049172 | 0.049172 | 0.80629 | 0.12788 | 0.22034 | 0.12788 | 9.862 | 0.01 | 0 | Neutral |
| testneg_1931 | 0 | 0 | 0.939339 | -3 | 0.818598 | -3 | 0.19898 | 0.14509 | 0.12739 | 0.12788 | 10.1991 | 0.00022619 | 0 | Neutral |
| testneg_1932 | -0.03 | -0.12 | 3.715248 | -3 | -0.272804 | -0.272804 | 0.83067 | 0.10583 | 0.17839 | 0.12009 | 12.65937 | 0.03 | 0 | Neutral |
| testneg_1933 | -0.02 | -0.13 | -0.203425 | -3 | -0.269735 | -0.269735 | 0.1276 | 0.30104 | 0.12461 | 0.1241 | 21.38889 | 0.051026316 | 0 | Neutral |
| testneg_1934 | -0.16 | 1.37 | 0.955677 | -3 | 2.028242 | -3 | 0.10034 | 0.10906 | 0.27058 | 0.13363 | 9.88601 | 0.25 | 0 | Neutral |
| testneg_1935 | 0 | 0 | 1.539176 | 1.539176 | 0.557877 | -3 | 0.91208 | 0.52258 | 0.25209 | 0.10029 | 9.74181 | 0.03 | 0 | Neutral |
| testneg_1936 | -0.21 | -0.15 | -0.83994 | -3 | -2.188525 | -2.188525 | 0.95319 | 0.20297 | 0.07282 | 0.12788 | 5.36688 | 0.07701457 | 0 | Neutral |
| testneg_1937 | 0.07 | 0.38 | -2.242853 | -3 | 3.13682 | -3 | 0.06358 | 0.5364 | 0.29828 | 0.12788 | 21.4497 | 0.22 | 0 | Neutral |
| testneg_1938 | 0.00290909 | 0.03636364 | 8.244795 | -0.412994 | 5.301011 | -3 | 0.04393 | 0.12788 | 0.45342 | 0.14389 | 17.52038 | 0.684316317 | 97 | Disease_causing |
| testneg_1939 | -0.18 | -0.16 | 4.070803 | -3 | 2.891351 | -3 | 0.37951 | 0.19435 | 0.93246 | 0.12788 | 10.83333 | 0.297121493 | 1 | Neutral |
| testneg_1940 | 0.21 | 0.15 | 1.631249 | -3 | 3.355302 | -3 | 0.47166 | 0.13548 | 0.21348 | 0.10616 | 5.97222 | 0.117638889 | 0 | Neutral |
| testneg_1941 | 0.01 | 0.79 | 5.588206 | -3 | -0.347591 | -3 | 0.16662 | 0.11306 | 0.32094 | 0.36254 | 26.53465 | 0.340294567 | 6.2 | Neutral |
| testneg_1942 | -0.08 | -0.03 | 3.929955 | -3 | 4.006946 | -3 | 0.72163 | 0.10596 | 0.42872 | 0.12788 | 19.90385 | 0.199188222 | 0 | Neutral |
| testneg_1943 | 0.17 | 0.38 | 2.59555 | -3 | -0.726772 | -3 | 0.79692 | 0.51979 | 0.1072 | 0.63446 | 10.22898 | 0.283661131 | 0 | Neutral |
| testneg_1944 | -0.07 | -0.38 | 2.28341 | -3 | 4.676207 | -3 | 0.27723 | 0.13229 | 0.1871 | 0.10907 | 16.84579 | 0.115563853 | 0 | Neutral |
| testneg_1945 | 0 | 0 | 2.221516 | -3 | 3.992923 | -3 | 0.16952 | 0.10613 | 0.06344 | 0.12788 | 24.67046 | 0.070590278 | 0 | Neutral |
| testneg_1946 | 0.03 | 0.1 | 5.405937 | -3 | 0.672589 | -3 | 0.13555 | 0.16663 | 0.07912 | 0.09055 | 20.31414 | 0.180047619 | 0.2 | Neutral |
| testneg_1947 | 0.03 | 0.12 | -0.183476 | -3 | 1.436437 | -3 | 0.34279 | 0.09378 | 0.71844 | 0.20204 | 10.41667 | 0.036093282 | 0 | Neutral |
| testneg_1948 | 0.08 | 0.03 | 1.617731 | -3 | 0.619175 | 0.619175 | 0.87038 | 0.43956 | 0.68138 | 0.20944 | 12.70833 | 0.147087787 | 0 | Neutral |
| testneg_1949 | -0.07 | -0.38 | -0.657494 | -3 | 0.753435 | -3 | 0.04056 | 0.55024 | 0.11187 | 0.12788 | 5.22072 | 0.13047726 | 0 | Neutral |
| testneg_1950 | 0.21 | 0.15 | 7.822266 | -3 | 0.185181 | -3 | 0.23005 | 0.37216 | 0.82156 | 0.14621 | 10.41667 | 0.46 | 37.4 | Neutral |
| testneg_1951 | -0.03 | -1.22 | 0.713058 | -3 | -1.646111 | -3 | 0.14375 | 0.12788 | 0.06939 | 0.08981 | 19.44444 | 0.03 | 0 | Neutral |

| | | | | | | | | | | | | | | |
|---|---|---|---|---|---|---|---|---|---|---|---|---|---|---|
| testneg_1952 | 0.16 | 0.08 | 1.340788 | -3 | 7.674169 | -3 | 0.08999 | 0.0989 | 0.8426 | 0.47836 | 10.41667 | 0.43 | 27.4 | Neutral |
| testneg_1953 | -0.07 | -0.7 | 0.044039 | -3 | -0.490467 | -3 | 0.36486 | 0.14268 | 0.0931 | 0.12788 | 34.48699 | 0.03 | 0 | Neutral |
| testneg_1954 | 0.07 | 0.01 | 0.98096 | -3 | 1.847923 | -3 | 0.5632 | 0.28442 | 0.13118 | 0.08646 | 18.6768 | 0.032188672 | 0 | Neutral |
| testneg_1955 | -0.16 | -0.08 | 2.388305 | -3 | -0.409484 | -3 | 0.13275 | 0.12788 | 0.22162 | 0.11997 | 9.97411 | 0.060182715 | 0 | Neutral |
| testneg_1956 | 0.04 | -0.23 | 7.508064 | -3 | 0.580057 | -3 | 0.9989 | 0.8746 | 0.10058 | 0.13882 | 17.08333 | 0.56 | 77.6 | Disease_causing |
| testneg_1957 | 0 | 0 | 0.227306 | -3 | -0.251883 | -3 | 0.30412 | 0.54626 | 0.39258 | 0.12788 | 21.81093 | 0.05 | 0 | Neutral |
| testneg_1958 | -0.08 | -0.03 | 5.150137 | -3 | -3.291659 | -3 | 0.44067 | 0.11201 | 0.11957 | 0.08975 | 16.63108 | 0.094761905 | 0 | Neutral |
| testneg_1959 | 0.18 | -1.05 | 5.931475 | -3 | 3.405014 | -3 | 0.19494 | 0.14451 | 0.32423 | 0.08104 | 24.22833 | 0.29 | 3 | Neutral |
| testneg_1960 | -0.07 | -0.38 | 1.684021 | -3 | 5.975539 | -3 | 0.64922 | 0.10176 | 0.35142 | 0.18303 | 18.17595 | 0.354710893 | 6.8 | Neutral |
| testneg_1961 | 0.04 | -0.23 | 1.9237 | -3 | 3.779532 | -3 | 0.60793 | 0.18264 | 0.11373 | 0.10739 | 11.25 | 0.142561688 | 0 | Neutral |
| testneg_1962 | 0 | 0 | -0.086282 | -3 | 4.030631 | -3 | 0.1014 | 0.12788 | 0.07617 | 0.07182 | 21.91231 | 0.08 | 0 | Neutral |
| testneg_1963 | -0.08 | -0.03 | 0.950149 | -3 | 1.681061 | -3 | 0.31318 | 0.11262 | 0.14403 | 0.30507 | 18.33333 | 0.065277778 | 0 | Neutral |
| testneg_1964 | -0.18 | -0.16 | -0.253658 | -3 | 1.572604 | -3 | 0.07293 | 0.12989 | 0.21967 | 0.09467 | 11.08369 | 0.08 | 0 | Neutral |
| testneg_1965 | 0.12 | -1.14 | 0.80533 | -3 | 1.932311 | 1.932311 | 0.44479 | 0.20004 | 0.97673 | 0.22976 | 12.35574 | 0.202959207 | 0 | Neutral |
| testneg_1966 | 0 | 0 | 1.444927 | -3 | 0.551681 | -3 | 0.25421 | 0.16235 | 0.17065 | 0.11746 | 35.23902 | 0.02 | 0 | Neutral |
| testneg_1967 | 0.17 | 0.38 | 2.7281 | -3 | 0.112197 | 0.112197 | 0.33163 | 0.26601 | 0.1123 | 0.1568 | 17.77778 | 0.24 | 0 | Neutral |
| testneg_1968 | 0.07 | 0.38 | -0.923815 | -3 | 3.840401 | -3 | 0.69249 | 0.12788 | 0.12917 | 0.15037 | 16.577 | 0.291987335 | 0 | Neutral |
| testneg_1969 | 0.05 | 0.29 | -0.673818 | -3 | 5.495658 | -3 | 0.76138 | 0.16258 | 0.15294 | 0.12788 | 18.33333 | 0.182820588 | 0 | Neutral |
| testneg_1970 | 0.04 | -0.23 | 7.117744 | -3 | -0.113155 | -3 | 0.39774 | 0.10813 | 0.14941 | 0.0838 | 33.09579 | 0.219162776 | 0.6 | Neutral |
| testneg_1971 | -0.03 | -0.12 | 0.991332 | -3 | 6.532451 | -3 | 0.15954 | 0.12788 | 0.19832 | 0.19622 | 14.78343 | 0.452194075 | 33.8 | Neutral |
| testneg_1972 | 0.14 | -0.37 | 6.747174 | -3 | 2.51727 | -3 | 0.34546 | 0.0932 | 0.06196 | 0.10881 | 11.25 | 0.35 | 9.8 | Neutral |
| testneg_1973 | 0 | 0 | 0.788459 | -3 | 4.072619 | 4.072619 | 0.18559 | 0.09173 | 0.16388 | 0.14439 | 12.29167 | 0.344054989 | 8.2 | Neutral |
| testneg_1974 | 0.18 | -1.05 | 7.432303 | -3 | 3.173176 | -3 | 0.16962 | 0.12788 | 0.17226 | 0.13001 | 11.92061 | 0.487139146 | 49.6 | Neutral |
| testneg_1975 | 0.14 | -1.85 | 3.507668 | -3 | 7.530767 | -3 | 0.88689 | 0.41514 | 0.1897 | 0.12459 | 8.45275 | 0.57 | 76.6 | Disease_causing |
| testneg_1976 | 0 | -0.01 | 1.806927 | -3 | 6.628636 | -3 | 0.17973 | 0.12278 | 0.51577 | 0.19561 | 6.32937 | 0.409728936 | 22.8 | Neutral |
| testneg_1977 | -0.01 | -0.8 | 2.652072 | -3 | 6.766427 | -3 | 0.51675 | 0.11055 | 0.10422 | 0.0994 | 26.44809 | 0.269398693 | 1.4 | Neutral |
| testneg_1978 | 0.04 | 2.01 | 3.382187 | -3 | 4.133728 | -3 | 0.50687 | 0.11283 | 0.13794 | 0.1141 | 17.21847 | 0.206963086 | 0 | Neutral |
| testneg_1979 | 0.04 | -0.23 | 0.436559 | -3 | -2.660026 | -3 | 0.14638 | 0.12788 | 0.37293 | 0.25159 | 10.41667 | 0.053144731 | 0 | Neutral |
| testneg_1980 | 0.2 | -1.26 | 1.378526 | -3 | 4.135613 | -3 | 0.12879 | 0.12788 | 0.62172 | 0.18135 | 14.27748 | 0.313411034 | 2.2 | Neutral |
| testneg_1981 | 0.14 | -1.01 | 0.701935 | -3 | 3.628292 | -3 | 0.09569 | 0.09246 | 0.15562 | 0.10384 | 23.2873 | 0.070344551 | 0 | Neutral |
| testneg_1982 | 0 | 0 | -0.378129 | -3 | 2.260209 | -3 | 0.85252 | 0.83576 | 0.12965 | 0.75487 | 11.875 | 0.207704374 | 0 | Neutral |
| testneg_1983 | 0 | 0 | 0.719031 | -3 | 2.393948 | 2.393948 | 0.9924 | 0.37896 | 0.70819 | 0.5975 | 9.81793 | 0.220462963 | 0 | Neutral |
| testneg_1984 | -0.16 | -0.08 | 2.465346 | -3 | 0.503836 | -3 | 0.25989 | 0.11651 | 0.19898 | 0.07875 | 36.14331 | 0.05 | 0 | Neutral |
| testneg_1985 | 0.11 | -1.95 | -1.183124 | -3 | -0.797179 | -3 | 0.13565 | 0.46572 | 0.1485 | 0.30134 | 13.75 | 0.2358553 | 0 | Neutral |
| testneg_1986 | 0 | 0 | 5.995443 | -3 | 0.752432 | -3 | 0.19898 | 0.14645 | 0.19898 | 0.1347 | 16.98166 | 0.3203125 | 5.6 | Neutral |
| testneg_1987 | -0.18 | 1.05 | 0.012106 | -3 | 1.01469 | -3 | 0.42755 | 0.09484 | 0.39019 | 0.18821 | 11.66667 | 0.129383117 | 0 | Neutral |
| testneg_1988 | -0.14 | 1.01 | 1.951553 | -3 | 3.19614 | -3 | 0.91377 | 0.5142 | 0.16309 | 0.12788 | 13.08659 | 0.241126984 | 0 | Neutral |
| testneg_1989 | 0 | 0 | 1.975201 | -3 | 1.366812 | -3 | 0.34495 | 0.16861 | 0.19898 | 0.12788 | 10.41667 | 0.010325225 | 0 | Neutral |
| testneg_1990 | 0 | 0 | -0.51617 | -3 | 0.053683 | -3 | 0.19898 | 0.12788 | 0.06895 | 0.06354 | 10.41667 | 0.005679945 | 0 | Neutral |
| testneg_1991 | 0.05 | -0.81 | 5.140271 | -3 | 6.986119 | -3 | 0.09542 | 0.14806 | 0.30742 | 0.30362 | 11.25 | 0.910354167 | 100 | Disease_causing |
| testneg_1992 | -0.01 | -0.79 | 1.796129 | -3 | 1.410512 | 0.502731 | 0.16052 | 0.12788 | 0.75033 | 0.64204 | 16.84579 | 0.081225913 | 0 | Neutral |
| testneg_1993 | 0.01 | -1.44 | 5.580166 | -3 | 0.293781 | -3 | 0.07681 | 0.22865 | 0.09273 | 0.08316 | 17.1181 | 0.22025 | 0.4 | Neutral |
| testneg_1994 | -0.15 | 0.28 | 3.501223 | -3 | 2.058739 | -3 | 0.46339 | 0.10423 | 0.19898 | 0.12788 | 12.85432 | 0.0925 | 0 | Neutral |
| testneg_1995 | 0 | 0 | 1.594568 | -3 | 1.88229 | -3 | 0.31318 | 0.11262 | 0.64922 | 0.10176 | 28.27549 | 0.01 | 0 | Neutral |
| testneg_1996 | 0.14 | -0.37 | 2.51438 | -3 | 4.33297 | -3 | 0.06118 | 0.08724 | 0.06331 | 0.2396 | 10.73741 | 0.374804563 | 11.4 | Neutral |
| testneg_1997 | 0 | 0 | 1.781646 | -3 | 0.645321 | -3 | 0.17258 | 0.12788 | 0.0538 | 0.23912 | 18.64268 | 0.05 | 0 | Neutral |
| testneg_1998 | 0.09 | -0.3 | 4.376111 | -3 | 6.277858 | -0.036663 | 0.03631 | 0.80564 | 0.09978 | 0.12788 | 11.66667 | 0.549388504 | 74.4 | Disease_causing |
| testneg_1999 | 0.03 | -1.03 | 3.590596 | -3 | 7.58963 | -3 | 0.1205 | 0.23508 | 0.10285 | 0.10886 | 19.44444 | 0.596602254 | 86.8 | Disease_causing |
| testneg_2000 | -0.01 | -0.79 | 1.879131 | -3 | 3.981688 | -3 | 0.59131 | 0.66216 | 0.19547 | 0.10692 | 5.73099 | 0.162886071 | 0 | Neutral |
| testneg_2001 | 0 | -0.32 | 6.451231 | -3 | 1.999648 | 1.999648 | 0.41383 | 0.119 | 0.12626 | 0.10279 | 23.47862 | 0.312111111 | 4.4 | Neutral |
| testneg_2002 | -0.16 | 1.37 | 3.207028 | -3 | 2.008385 | -3 | 0.03169 | 0.10615 | 0.07275 | 0.12788 | 28.62307 | 0.198935829 | 0 | Neutral |
| testneg_2003 | -0.19 | -0.51 | 6.05963 | 2.214477 | 5.956926 | -3 | 0.22162 | 0.11997 | 0.10423 | 0.12788 | 21.01754 | 0.59 | 87.2 | Disease_causing |
| testneg_2004 | 0 | -0.01 | 0.549483 | -3 | 5.871334 | -3 | 0.46084 | 0.88583 | 0.82148 | 0.14847 | 4.72222 | 0.329344082 | 6.8 | Neutral |
| testneg_2005 | 0.19 | 0.51 | 2.941559 | -3 | 1.541854 | -3 | 0.30474 | 0.12788 | 0.1065 | 0.12788 | 13.51351 | 0.05 | 0 | Neutral |
| testneg_2006 | -0.19 | 0.25 | 1.575308 | -3 | 7.140042 | -3 | 0.20655 | 0.15317 | 0.04142 | 0.09564 | 20.90643 | 0.440472689 | 29.4 | Neutral |
| testneg_2007 | 0 | 0 | 1.304791 | -3 | 6.256857 | -3 | 0.96704 | 0.20571 | 0.65674 | 0.10572 | 28.07329 | 0.150429825 | 0 | Neutral |
| testneg_2008 | 0 | 0 | 0.287181 | -3 | 2.091639 | -3 | 0.93834 | 0.11088 | 0.67327 | 0.23543 | 22.06714 | 0.071687822 | 0 | Neutral |
| testneg_2009 | 0.12 | -1.16 | 6.340508 | -3 | 1.326414 | -3 | 0.339 | 0.13169 | 0.39863 | 0.44944 | 10.08333 | 0.583590248 | 82.8 | Disease_causing |
| testneg_2010 | 0.2 | -1.98 | 2.423357 | -3 | 0.752432 | -3 | 0.13083 | 0.23231 | 0.19898 | 0.1347 | 20.17299 | 0.2 | 0 | Neutral |
| testneg_2011 | 0.04 | -0.23 | 6.519516 | -3 | -0.025657 | -2.808208 | 0.28211 | 0.25332 | 0.16259 | 0.102 | 20.83333 | 0.309833333 | 4.8 | Neutral |
| testneg_2012 | 0 | 0 | 5.510751 | -3 | 2.891491 | -3 | 0.35428 | 0.12971 | 0.39087 | 0.10381 | 9.67871 | 0.30825 | 2.6 | Neutral |
| testneg_2013 | 0 | 0 | 1.630014 | -3 | 3.229354 | -3 | 0.1109 | 0.17295 | 0.1743 | 0.12788 | 32.36846 | 0.093717105 | 0 | Neutral |
| testneg_2014 | 0.03 | -1.03 | 4.994758 | -3 | 1.640857 | -3 | 0.1664 | 0.10113 | 0.064 | 0.24721 | 32.29167 | 0.274476805 | 1.6 | Neutral |
| testneg_2015 | -0.09 | 0.3 | -2.556301 | -3 | 5.354249 | -3 | 0.12379 | 0.12788 | 0.64914 | 0.12481 | 20.15927 | 0.191715686 | 0 | Neutral |
| testneg_2016 | -0.02 | -0.45 | 5.696978 | -3 | -1.258796 | -1.258796 | 0.32974 | 0.12245 | 0.7055 | 0.11261 | 10.06328 | 0.153888889 | 0 | Neutral |
| testneg_2017 | 0.16 | 0.08 | 3.167319 | -3 | 4.85825 | -3 | 0.25195 | 0.09584 | 0.16275 | 0.12788 | 11.95767 | 0.211586168 | 0.2 | Neutral |
| testneg_2018 | -0.02 | -0.45 | 5.696978 | -3 | 3.986941 | -3 | 0.32974 | 0.12245 | 0.11522 | 0.11925 | 17.88889 | 0.345418129 | 6 | Neutral |
| testneg_2019 | 0.04 | -0.23 | 0.34099 | -3 | 2.324167 | -3 | 0.33057 | 0.10549 | 0.47274 | 0.11928 | 12.33727 | 0.01 | 0 | Neutral |
| testneg_2020 | 0 | -0.01 | -0.052748 | -3 | 0.88775 | 0.88775 | 0.91139 | 0.65138 | 0.32649 | 0.15357 | 10.83333 | 0.102718568 | 0 | Neutral |
| testneg_2021 | 0 | -0.73 | 2.726423 | -3 | 1.152738 | -3 | 0.14273 | 0.1162 | 0.08067 | 0.34914 | 30.90278 | 0.14 | 0 | Neutral |
| testneg_2022 | -0.09 | 1.07 | 6.071016 | -3 | 0.130932 | -3 | 0.24791 | 0.09675 | 0.19898 | 0.12788 | 18.76159 | 0.206583333 | 0.2 | Neutral |
| testneg_2023 | 0 | 0 | -0.030607 | -3 | 5.597555 | -3 | 0.17146 | 0.19882 | 0.07406 | 0.08059 | 21.31912 | 0.162834416 | 0 | Neutral |
| testneg_2024 | 0.07 | 0.38 | 1.845239 | -3 | 4.060263 | -3 | 0.21728 | 0.1132 | 0.15125 | 0.08976 | 28.47222 | 0.060391337 | 0 | Neutral |
| testneg_2025 | 0.15 | -0.82 | 3.321668 | -3 | 1.932311 | 1.932311 | 0.32977 | 0.15427 | 0.97673 | 0.22976 | 9.90471 | 0.293166667 | 2.2 | Neutral |
| testneg_2026 | 0 | 0 | -0.399895 | -3 | -0.314397 | -3 | 0.5632 | 0.28442 | 0.10208 | 0.08867 | 19.5422 | 0.031026316 | 0 | Neutral |
| testneg_2027 | 0 | 0 | -0.090459 | -0.090459 | -1.08606 | -3 | 0.15581 | 0.19351 | 0.28269 | 0.12505 | 20.83333 | 0.020145416 | 0 | Neutral |
| testneg_2028 | 0.09 | -0.25 | -0.159404 | -0.904157 | 4.369274 | -3 | 0.95319 | 0.20297 | 0.28416 | 0.27805 | 12.40248 | 0.426442163 | 23.4 | Neutral |
| testneg_2029 | 0.14 | -0.37 | 0.198379 | -3 | -1.097042 | -3 | 0.03991 | 0.08342 | 0.10703 | 0.11611 | 32.91244 | 0.040283816 | 0 | Neutral |
| testneg_2030 | 0.02 | -0.36 | 2.305765 | -3 | 1.323079 | -3 | 0.11055 | 0.1127 | 0.5103 | 0.10923 | 32.52813 | 0.01 | 0 | Neutral |
| testneg_2031 | 0 | 0 | 0.067126 | -3 | 4.354155 | -3 | 0.18869 | 0.23984 | 0.10244 | 0.10107 | 10.83333 | 0.130188948 | 0 | Neutral |
| testneg_2032 | 0.22 | -2.66 | 4.869758 | -3 | 0.442489 | -2.924962 | 0.2218 | 0.11533 | 0.0478 | 0.12788 | 31.14232 | 0.09 | 0 | Neutral |
| testneg_2033 | 0 | 0 | 1.992408 | -3 | 0.60982 | 0.60982 | 0.19898 | 0.25991 | 0.23844 | 0.27772 | 11.28725 | 0.130632206 | 0 | Neutral |
| testneg_2034 | 0 | 0 | 0.82819 | 0.82819 | -0.456898 | -3 | 0.19898 | 0.12788 | 0.06397 | 0.12788 | 22.13058 | 0.03 | 0 | Neutral |
| testneg_2035 | 0.04 | -0.23 | 4.771654 | -3 | 1.9182 | -3 | 0.6201 | 0.12788 | 0.43762 | 0.12788 | 10.83333 | 0.070266714 | 0 | Neutral |
| testneg_2036 | 0 | 0 | 0.435323 | -3 | 0.626243 | 0.626243 | 0.20527 | 0.10053 | 0.34412 | 0.08672 | 13.02635 | 0.000227273 | 0 | Neutral |
| testneg_2037 | 0.07 | 0.38 | -0.575902 | -3 | 1.223517 | -3 | 0.69666 | 0.11055 | 0.19898 | 0.12788 | 22.35904 | 0.02051501 | 0 | Neutral |
| testneg_2038 | 0.09 | -0.25 | -3.307114 | -3 | 1.675753 | -3 | 0.14495 | 0.37127 | 0.24736 | 0.12777 | 34.21139 | 0.080666996 | 0 | Neutral |
| testneg_2039 | 0.07 | -1.12 | 4.403102 | -3 | -0.01145 | -0.01145 | 0.19301 | 0.10828 | 0.78462 | 0.12258 | 11.69624 | 0.060147619 | 0 | Neutral |
| testneg_2040 | 0 | 0 | 0.371766 | -3 | 0.191986 | -3 | 0.3095 | 0.13502 | 0.03235 | 0.0948 | 13.51598 | 0.01 | 0 | Neutral |
| testneg_2041 | -0.02 | -0.45 | 1.744503 | -3 | 1.776558 | -3 | 0.17301 | 0.14089 | 0.42214 | 0.21008 | 18.33333 | 0.083616883 | 0 | Neutral |
| testneg_2042 | 0.05 | 0.29 | 2.297641 | -3 | 1.366001 | -3 | 0.80629 | 0.12788 | 0.04305 | 0.06354 | 10.22898 | 0.02 | 0 | Neutral |
| testneg_2043 | 0 | 0 | 0.346053 | -3 | 6.966984 | -3 | 0.3047 | 0.11929 | 0.08143 | 0.44857 | 18.88889 | 0.417335907 | 22 | Neutral |
| testneg_2044 | -0.15 | 0.28 | 5.116499 | -3 | -1.784808 | -3 | 0.0597 | 0.18092 | 0.11133 | 0.12788 | 19.66924 | 0.23 | 0.2 | Neutral |
| testneg_2045 | -0.01 | -0.8 | 1.765078 | -3 | 4.444325 | -3 | 0.36461 | 0.11015 | 0.28474 | 0.12788 | 16.98432 | 0.065 | 0 | Neutral |
| testneg_2046 | 0.14 | -0.69 | 2.950487 | -3 | 0.782368 | -3 | 0.14495 | 0.37127 | 0.31651 | 0.12788 | 33.48065 | 0.128808933 | 0 | Neutral |
| testneg_2047 | 0 | 0 | 0.769846 | -3 | 3.891578 | -3 | 0.19898 | 0.10512 | 0.78253 | 0.1114 | 20.04401 | 0.043452381 | 0 | Neutral |
| testneg_2048 | 0 | 0 | 1.075324 | 1.075324 | -0.219149 | -3 | 0.49934 | 0.11069 | 0.16409 | 0.12788 | 12.52798 | 0.03 | 0 | Neutral |
| testneg_2049 | 0 | 0 | 2.652503 | -3 | 0.577403 | -3 | 0.08059 | 0.09576 | 0.13815 | 0.40001 | 23.61582 | 0.128928571 | 0 | Neutral |
| testneg_2050 | -0.07 | -0.33 | 1.282461 | -3 | -0.431211 | -3 | 0.59131 | 0.66216 | 0.36676 | 0.37749 | 10 | 0.144208333 | 0.2 | Neutral |
| testneg_2051 | 0.05 | -0.81 | 3.217486 | -3 | 6.080162 | -3 | 0.74699 | 0.12286 | 0.50448 | 0.50327 | 11.59461 | 0.587647013 | 82.6 | Disease_causing |
| testneg_2052 | -0.03 | -0.1 | 2.296124 | -3 | 0.338128 | -3 | 0.22258 | 0.10471 | 0.32558 | 0.32796 | 18.33333 | 0.083547619 | 0 | Neutral |
| testneg_2053 | 0.19 | 0.51 | 3.650653 | -3 | 0.288206 | -3 | 0.07558 | 0.12788 | 0.25462 | 0.12788 | 27.4024 | 0.05 | 0 | Neutral |
| testneg_2054 | 0.06 | -0.57 | 2.083519 | -3 | -1.329864 | -3 | 0.14375 | 0.12788 | 0.16966 | 0.61198 | 10.41667 | 0.080465368 | 0 | Neutral |
| testneg_2055 | -0.0072727 | -0.2709091 | 4.824644 | -3 | -0.052442 | -3 | 0.10723 | 0.11163 | 0.19898 | 0.09201 | 19.77124 | 0.04 | 0 | Neutral |
| testneg_2056 | -0.01 | -0.8 | 3.547766 | -3 | -0.842215 | -3 | 0.05124 | 0.07099 | 0.21876 | 0.12373 | 18.88889 | 0.070120282 | 0 | Neutral |
| testneg_2057 | -0.07 | -0.01 | 4.449247 | 4.449247 | 0.67523 | -3 | 0.35779 | 0.12788 | 0.60603 | 0.12788 | 19.44444 | 0.48 | 45.2 | Neutral |
| testneg_2058 | -0.03 | -0.1 | 6.889081 | -3 | 5.829171 | -3 | 0.13559 | 0.1946 | 0.05079 | 0.0924 | 28.38695 | 0.591010101 | 85.6 | Disease_causing |
| testneg_2059 | -0.17 | -0.38 | 5.556132 | -3 | -1.646194 | -3 | 0.03109 | 0.10705 | 0.10362 | 0.09649 | 17.01711 | 0.19 | 0 | Neutral |
| testneg_2060 | 0.02 | -0.36 | 1.715391 | -3 | -2.464297 | -2.464297 | 0.60959 | 0.76078 | 0.11524 | 0.12788 | 10 | 0.048434343 | 0 | Neutral |
| testneg_2061 | 0.03 | -1.03 | 4.387695 | -3 | -1.989097 | -3 | 0.18341 | 0.13901 | 0.11094 | 0.12788 | 18.33333 | 0.06 | 0 | Neutral |
| testneg_2062 | -0.03 | -0.12 | 0.345153 | -3 | 1.851573 | -3 | 0.15785 | 0.35538 | 0.13295 | 0.12914 | 27.40684 | 0.063150183 | 0 | Neutral |
| testneg_2063 | -0.03 | 0.85 | -0.534952 | -3 | 0.654023 | -3 | 0.12203 | 0.13515 | 0.63645 | 0.14596 | 18.88889 | 0.070093705 | 0 | Neutral |
| testneg_2064 | -0.08 | -0.03 | 0.536745 | -3 | 1.441424 | -3 | 0.34295 | 0.12788 | 0.20231 | 0.12788 | 12.52798 | 0.01 | 0 | Neutral |
| testneg_2065 | 0.09 | -0.3 | 2.387342 | -3 | 1.280424 | -3 | 0.07428 | 0.52467 | 0.23427 | 0.10875 | 10.41667 | 0.08 | 0 | Neutral |
| testneg_2066 | -0.07 | -0.38 | 6.836452 | -3 | 3.274992 | -3 | 0.19898 | 0.09316 | 0.37637 | 0.12788 | 11.25 | 0.459450549 | 39.6 | Neutral |
| testneg_2067 | 0 | 0 | 2.548364 | -3 | 5.522589 | -3 | 0.21376 | 0.12708 | 0.19355 | 0.18675 | 12.90858 | 0.408038837 | 21.4 | Neutral |
| testneg_2068 | 0.03 | -1.03 | 2.660158 | -3 | 0.583408 | 0.583408 | 0.18243 | 0.11102 | 0.05386 | 0.09802 | 18.88889 | 0.03 | 0 | Neutral |
| testneg_2069 | 0 | 0 | 2.10021 | -3 | 0.382515 | -3 | 0.15526 | 0.08511 | 0.19898 | 0.12788 | 17.84483 | 0 | 0 | Neutral |
| testneg_2070 | 0.03 | 1.22 | 2.392056 | -3 | 6.16015 | 0.837125 | 0.72501 | 0.7625 | 0.103 | 0.403 | 10.41667 | 0.617978823 | 91.8 | Disease_causing |
| testneg_2071 | -0.07 | -0.7 | -0.068708 | -3 | 2.438756 | 1.937541 | 0.25077 | 0.06261 | 0.39019 | 0.18821 | 12.34649 | 0.16 | 0 | Neutral |
| testneg_2072 | 0.21 | 0.15 | 6.256857 | -3 | -2.30753 | -3 | 0.65674 | 0.10572 | 0.13118 | 0.08646 | 39.64236 | 0.16 | 0.2 | Neutral |
| testneg_2073 | 0.22 | -2.66 | 5.165273 | -3 | 0.338954 | -3 | 0.18465 | 0.11262 | 0.25422 | 0.10102 | 18.24619 | 0.092916667 | 0 | Neutral |

| | | | | | | | | | | | | | | |
|---|---|---|---|---|---|---|---|---|---|---|---|---|---|---|
| testneg_2074 | -0.01 | -0.8 | 7.383324 | -3 | 1.641419 | -3 | 0.22761 | 0.14229 | 0.11891 | 0.11186 | 11.74806 | 0.45 | 33.8 | Neutral |
| testneg_2075 | -0.03 | -0.1 | -3.346595 | -3 | 2.235719 | -3 | 0.24949 | 0.29709 | 0.12511 | 0.11897 | 19.44444 | 0.06 | 0 | Neutral |
| testneg_2076 | 0.18 | -1.05 | 3.183437 | -3 | 5.893689 | -3 | 0.20448 | 0.12788 | 0.21383 | 0.11171 | 10.07288 | 0.255173252 | 1.6 | Neutral |
| testneg_2077 | 0.03 | -1.03 | 4.859478 | -3 | -1.487172 | -1.487172 | 0.11501 | 0.13748 | 0.103 | 0.403 | 18.83333 | 0.32 | 3.6 | Neutral |
| testneg_2078 | 0.16 | -1.37 | 6.076778 | -3 | 0.327712 | -3 | 0.33818 | 0.12788 | 0.57517 | 0.10873 | 26.84685 | 0.15 | 0 | Neutral |
| testneg_2079 | -0.07 | -0.01 | 2.291931 | -3 | 0.405386 | -3 | 0.99998 | 0.86471 | 0.08476 | 0.12788 | 10 | 0.095848214 | 0 | Neutral |
| testneg_2080 | 0.08 | 0.03 | -2.363283 | -2.363283 | 5.119697 | -3 | 0.05846 | 0.12788 | 0.61834 | 0.10784 | 36.92514 | 0.119181287 | 0 | Neutral |
| testneg_2081 | 0.02 | -0.63 | 6.708054 | -3 | -1.168846 | -3 | 0.28063 | 0.40063 | 0.19898 | 0.12788 | 18.04026 | 0.376904762 | 11 | Neutral |
| testneg_2082 | -0.03 | -0.1 | 0.975747 | -3 | 10.968892 | -3 | 0.53447 | 0.10345 | 0.10183 | 0.11847 | 21.34347 | 0.35 | 6.2 | Neutral |
| testneg_2083 | 0 | 0 | 0.021671 | -3 | 6.843902 | -3 | 0.13156 | 0.10621 | 0.07652 | 0.30451 | 10.41667 | 0.398543956 | 16.4 | Neutral |
| testneg_2084 | 0 | 0 | 1.465695 | -3 | 3.446417 | -3 | 0.94822 | 0.14005 | 0.29013 | 0.11368 | 23.81085 | 0.063938176 | 0 | Neutral |
| testneg_2085 | 0.02 | 0.45 | 0.379358 | -3 | 0.406963 | -3 | 0.19898 | 0.10996 | 0.48825 | 0.10951 | 27.26565 | 0.01 | 0 | Neutral |
| testneg_2086 | 0 | 0 | -0.187456 | -0.187456 | 0.148971 | -3 | 0.19898 | 0.12788 | 0.17007 | 0.12771 | 28.23755 | 0.01 | 0 | Neutral |
| testneg_2087 | 0.18 | -1.05 | 8.297965 | -3 | 0.237568 | -3 | 0.71762 | 0.14417 | 0.98118 | 0.10229 | 9.76428 | 0.4 | 18.6 | Neutral |
| testneg_2088 | 0 | 0 | 0.393827 | -3 | 2.981794 | -3 | 0.07516 | 0.15472 | 0.17394 | 0.15823 | 10.41667 | 0.186736014 | 0 | Neutral |
| testneg_2089 | 0 | 0 | 1.145768 | -3 | 4.429067 | -3 | 0.22658 | 0.20646 | 0.19898 | 0.09824 | 15.60606 | 0.101187811 | 0 | Neutral |
| testneg_2090 | -0.06 | 0.57 | 0.709477 | -3 | 2.009531 | 0.005079 | 0.19898 | 0.12788 | 0.47322 | 0.11118 | 24.07509 | 0.02 | 0 | Neutral |
| testneg_2091 | 0.02 | -0.68 | 1.719272 | -3 | 0.72503 | -3 | 0.024 | 0.07726 | 0.56862 | 0.12788 | 29.81906 | 0.04 | 0 | Neutral |
| testneg_2092 | 0.08 | 0.03 | -0.730591 | -3 | 2.317066 | -3 | 0.52953 | 0.10241 | 0.09569 | 0.09246 | 31.59722 | 0.062884615 | 0 | Neutral |
| testneg_2093 | 0 | 0 | 2.244975 | -3 | 2.024568 | -3 | 0.92832 | 0.16306 | 0.40996 | 0.57507 | 9.84127 | 0.10792619 | 0 | Neutral |
| testneg_2094 | -0.08 | -0.03 | 6.10454 | -3 | -0.658155 | -3 | 0.56605 | 0.15893 | 0.05087 | 0.09682 | 14.69298 | 0.24 | 0.6 | Neutral |
| testneg_2095 | 0 | 0 | 1.590262 | -3 | 3.592951 | -3 | 0.44385 | 0.1311 | 0.18122 | 0.16895 | 8.97898 | 0.214234668 | 0.2 | Neutral |
| testneg_2096 | 0 | -0.01 | 0.439263 | -3 | 1.867073 | -3 | 0.87835 | 0.13522 | 0.07999 | 0.23927 | 10.83333 | 0.065046717 | 0 | Neutral |
| testneg_2097 | 0.22 | -2.66 | 3.630623 | -3 | 6.939074 | -3 | 0.29356 | 0.47581 | 0.07467 | 0.09224 | 19.44444 | 0.55 | 75 | Disease_causing |
| testneg_2098 | 0 | 0 | 1.718666 | -3 | 7.706675 | -3 | 0.07889 | 0.08835 | 0.09806 | 0.12788 | 18.33333 | 0.28175 | 2.2 | Neutral |
| testneg_2099 | 0.05 | -0.44 | 3.440427 | -3 | 3.598054 | -3 | 0.19898 | 0.12788 | 0.50275 | 0.38957 | 12.70833 | 0.508341575 | 58.4 | Disease_causing |
| testneg_2100 | -0.16 | -0.08 | 2.465346 | -3 | 1.26254 | -3 | 0.25989 | 0.11651 | 0.35189 | 0.12788 | 33.51351 | 0.053214286 | 0 | Neutral |
| testneg_2101 | 0 | 0 | -0.044046 | -3 | 2.86988 | -3 | 0.47544 | 0.17658 | 0.12987 | 0.18163 | 4.44444 | 0.156971487 | 0 | Neutral |
| testneg_2102 | 0 | 0 | 0.256955 | -3 | -1.00023 | -3 | 0.1846 | 0.08682 | 0.43158 | 0.10472 | 30.20956 | 0.01 | 0 | Neutral |
| testneg_2103 | 0 | 0 | 0.946307 | -3 | 4.711026 | -3 | 0.18081 | 0.1258 | 0.03898 | 0.09943 | 10.29424 | 0.044365079 | 0 | Neutral |
| testneg_2104 | 0.16 | -1.37 | -0.121102 | -0.121102 | -1.046088 | -3 | 0.78935 | 0.17351 | 0.93246 | 0.12788 | 6.99956 | 0.071079545 | 0 | Neutral |
| testneg_2105 | -0.03 | -1.22 | 4.490043 | -3 | -0.574737 | -3 | 0.7342 | 0.11023 | 0.56678 | 0.09955 | 19.05127 | 0.070053763 | 0 | Neutral |
| testneg_2106 | 0 | 0 | -0.233364 | -3 | 3.663929 | -3 | 0.26323 | 0.2399 | 0.25731 | 0.12662 | 20.27778 | 0.102291667 | 0 | Neutral |
| testneg_2107 | 0 | -0.32 | 3.991191 | -3 | 2.381789 | -3 | 0.07428 | 0.52467 | 0.63418 | 0.69224 | 10 | 0.6100625 | 91.8 | Disease_causing |
| testneg_2108 | 0 | 0 | 1.103204 | -3 | 3.992072 | -3 | 0.57199 | 0.11262 | 0.05767 | 0.07325 | 21.01901 | 0.09 | 0 | Neutral |
| testneg_2109 | 0 | 0 | -0.398253 | -3 | 2.135922 | -3 | 0.04668 | 0.12788 | 0.14498 | 0.33896 | 10.41667 | 0.108309991 | 0 | Neutral |
| testneg_2110 | 0.18 | -1.05 | 6.533819 | -3 | 1.668921 | -3 | 0.50303 | 0.23508 | 0.15994 | 0.11186 | 10.83333 | 0.393356481 | 19.6 | Neutral |
| testneg_2111 | 0 | -0.01 | 2.038329 | -3 | 4.276849 | -3 | 0.25159 | 0.12788 | 0.0778 | 0.0728 | 20.94356 | 0.069833333 | 0 | Neutral |
| testneg_2112 | -0.06 | -0.18 | 1.759866 | -3 | 0.072403 | -3 | 0.7249 | 0.41559 | 0.18671 | 0.12788 | 12.29167 | 0.03 | 0 | Neutral |
| testneg_2113 | 0.03 | -0.85 | 0.173309 | -3 | 0.559482 | -3 | 0.34694 | 0.20397 | 0.836 | 0.15993 | 11.25 | 0.100851513 | 0 | Neutral |
| testneg_2114 | -0.1 | -1.9 | 2.484571 | -3 | 1.106407 | -3 | 0.13575 | 0.12788 | 0.19898 | 0.1054 | 10.22898 | 0.04 | 0 | Neutral |
| testneg_2115 | 0 | 0 | 0.515441 | -3 | -0.154534 | -3 | 0.2741 | 0.10478 | 0.34946 | 0.19092 | 16.8018 | 0.020227273 | 0 | Neutral |
| testneg_2116 | -0.09 | 0.67 | 4.929625 | -3 | -0.022376 | -3 | 0.07447 | 0.23492 | 0.05817 | 0.09497 | 10.41667 | 0.17 | 0 | Neutral |
| testneg_2117 | 0.08 | 0.03 | 0.202818 | -3 | 5.393665 | 5.393665 | 0.07118 | 0.08861 | 0.03809 | 0.16044 | 19.44444 | 0.461163696 | 38.2 | Neutral |
| testneg_2118 | -0.19 | 0.25 | -2.608658 | -3 | 0.411241 | -3 | 0.19898 | 0.12788 | 0.19199 | 0.16616 | 24.6391 | 0.130805405 | 0 | Neutral |
| testneg_2119 | 0.02 | -0.68 | 1.9182 | -3 | 2.700108 | -3 | 0.43762 | 0.12788 | 0.41102 | 0.0974 | 23.04078 | 0.042868449 | 0 | Neutral |
| testneg_2120 | 0.18 | 0.16 | 5.915401 | -3 | 1.995777 | -3 | 0.16777 | 0.1749 | 0.20632 | 0.12788 | 20.89477 | 0.251895995 | 1.6 | Neutral |
| testneg_2121 | 0 | -0.01 | 0.061667 | -3 | 5.068794 | -3 | 0.13751 | 0.13274 | 0.04627 | 0.05223 | 15.39358 | 0.081594828 | 0 | Neutral |
| testneg_2122 | -0.05 | -0.29 | 0.52744 | -3 | 1.416689 | 1.416689 | 0.23278 | 0.11027 | 0.33163 | 0.26601 | 5.74956 | 0.08170566 | 0 | Neutral |
| testneg_2123 | 0 | 0 | 0.227306 | -3 | 0.084517 | -3 | 0.30412 | 0.54626 | 0.25179 | 0.09355 | 16.8018 | 0.03 | 0 | Neutral |
| testneg_2124 | 0.07 | 0.7 | 3.542059 | -3 | -0.729977 | -0.729977 | 0.10569 | 0.09059 | 0.17974 | 0.12788 | 16.59347 | 0.040134011 | 0 | Neutral |
| testneg_2125 | -0.01 | -0.8 | 2.607371 | -3 | -0.061652 | -3 | 0.53239 | 0.17509 | 0.83074 | 0.14769 | 10.83333 | 0.1 | 0 | Neutral |
| testneg_2126 | 0 | 0 | 1.006231 | -3 | 1.029613 | -3 | 0.14199 | 0.1327 | 0.2183 | 0.10789 | 15.59524 | 0.01 | 0 | Neutral |
| testneg_2127 | 0.08 | 0.03 | -1.290562 | -3 | -2.337287 | -3 | 0.19898 | 0.12895 | 0.04946 | 0.12788 | 40.39594 | 0.03 | 0 | Neutral |
| testneg_2128 | -0.01 | -0.79 | -2.624716 | -3 | 4.197845 | -3 | 0.05932 | 0.09551 | 0.58292 | 0.87725 | 5.96836 | 0.338779762 | 5 | Neutral |
| testneg_2129 | 0 | 0 | -0.223347 | -3 | 4.038181 | 4.038181 | 0.25924 | 0.19965 | 0.13815 | 0.40001 | 4.44444 | 0.539873706 | 69.2 | Disease_causing |
| testneg_2130 | 0 | 0 | -0.090459 | -0.090459 | 5.971271 | -3 | 0.15581 | 0.19351 | 0.16973 | 0.17555 | 9.55514 | 0.504037666 | 56.8 | Disease_causing |
| testneg_2131 | 0 | 0 | 0.227306 | -3 | 1.240738 | -3 | 0.30412 | 0.54626 | 0.32238 | 0.10913 | 10.41667 | 0.021096917 | 0 | Neutral |
| testneg_2132 | 0 | 0 | 1.872374 | -3 | 6.288981 | -3 | 0.28907 | 0.12788 | 0.12463 | 0.12788 | 19.51632 | 0.123765891 | 0 | Neutral |
| testneg_2133 | -0.07 | -0.38 | -1.026052 | -3 | 2.322245 | -3 | 0.16469 | 0.16042 | 0.30734 | 0.12788 | 18.88889 | 0.051687182 | 0 | Neutral |
| testneg_2134 | 0 | 0 | 0.040296 | -3 | -0.272739 | -3 | 0.25856 | 0.17179 | 0.44594 | 0.13793 | 9.0522 | 0.047648604 | 0 | Neutral |
| testneg_2135 | 0.18 | -1.05 | 5.997252 | -3 | -1.43595 | -3 | 0.68029 | 0.12788 | 0.05999 | 0.12788 | 18.6768 | 0.178611111 | 0 | Neutral |
| testneg_2136 | 0 | 0 | -0.274465 | -3 | 1.639031 | -3 | 0.04699 | 0.08149 | 0.15443 | 0.10913 | 18.23893 | 0.03 | 0 | Neutral |
| testneg_2137 | 0.16 | -1.37 | 6.571813 | 6.279292 | -1.097042 | -3 | 0.09488 | 0.10426 | 0.10703 | 0.11611 | 33.60688 | 0.56 | 78.8 | Disease_causing |
| testneg_2138 | 0 | 0 | 1.354498 | -3 | 2.148373 | -3 | 0.33713 | 0.12788 | 0.21376 | 0.12708 | 19.09347 | 0.000101525 | 0 | Neutral |
| testneg_2139 | -0.16 | -0.08 | 1.741968 | -3 | 2.70354 | -3 | 0.16057 | 0.12305 | 0.46735 | 0.10674 | 40.45114 | 0.09 | 0 | Neutral |
| testneg_2140 | 0 | 0 | 1.576631 | -3 | -1.572727 | -1.572727 | 0.15509 | 0.10665 | 0.14231 | 0.41445 | 18.6768 | 0.053523504 | 0 | Neutral |
| testneg_2141 | 0 | 0 | 0.032907 | -3 | 1.282461 | -3 | 0.04555 | 0.06327 | 0.59131 | 0.66216 | 10.26553 | 0.080286218 | 0 | Neutral |
| testneg_2142 | 0.15 | -1.06 | 2.457352 | -3 | 2.191197 | -3 | 0.13974 | 0.10225 | 0.18667 | 0.10488 | 16.8018 | 0.03 | 0 | Neutral |
| testneg_2143 | -0.07 | -0.38 | 5.958045 | -3 | 0.820444 | 0.820444 | 0.11133 | 0.12788 | 0.17736 | 0.14772 | 16.84327 | 0.34 | 9 | Neutral |
| testneg_2144 | -0.07 | -0.38 | -1.968112 | -3 | 5.810701 | -3 | 0.08779 | 0.1777 | 0.21265 | 0.1284 | 10.41667 | 0.311040936 | 2.6 | Neutral |
| testneg_2145 | 0.01 | -1.44 | -2.044558 | -3 | 1.55056 | -3 | 0.14511 | 0.12788 | 0.12857 | 0.09376 | 21.22834 | 0.04 | 0 | Neutral |
| testneg_2146 | 0.15 | -0.28 | -0.255381 | -3 | -0.111739 | -3 | 0.0917 | 0.13124 | 0.08237 | 0.09015 | 29.9024 | 0.03 | 0 | Neutral |
| testneg_2147 | 0 | 0 | 1.019099 | -3 | 1.912827 | -3 | 0.19898 | 0.12788 | 0.1203 | 0.13872 | 19.44853 | 0.050163016 | 0 | Neutral |
| testneg_2148 | -0.08 | -0.03 | 3.600262 | -3 | 1.962395 | -3 | 0.22944 | 0.09039 | 0.45399 | 0.10245 | 28.62818 | 0.05 | 0 | Neutral |
| testneg_2149 | -0.07 | -0.33 | 1.125228 | -3 | -0.614161 | -3 | 0.15808 | 0.14548 | 0.17796 | 0.09522 | 10 | 0.02 | 0 | Neutral |
| testneg_2150 | 0 | 0 | 0.032879 | -3 | 5.235923 | -3 | 0.20448 | 0.12788 | 0.35026 | 0.09817 | 23.83768 | 0.05 | 0 | Neutral |
| testneg_2151 | 0 | 0 | 0.080104 | -3 | 3.12089 | -3 | 0.73947 | 0.2058 | 0.23578 | 0.12788 | 30.26287 | 0.084876374 | 0 | Neutral |
| testneg_2152 | 0 | 0 | 0.440494 | 0.440494 | 1.384899 | 1.384899 | 0.61027 | 0.14229 | 0.56315 | 0.13761 | 13.28168 | 0.08 | 0 | Neutral |
| testneg_2153 | -0.02 | -0.13 | 5.682853 | -3 | 5.068794 | -3 | 0.31344 | 0.12788 | 0.04627 | 0.05223 | 17.21847 | 0.340541667 | 8 | Neutral |
| testneg_2154 | 0.09 | -0.3 | 0.922131 | -3 | 5.486648 | -3 | 0.05886 | 0.38242 | 0.16708 | 0.11594 | 10.83333 | 0.307893218 | 2.2 | Neutral |
| testneg_2155 | 0.08 | 0.03 | 1.355095 | -3 | 2.441441 | -3 | 0.28907 | 0.12788 | 0.08224 | 0.29179 | 12.90213 | 0.153053597 | 0 | Neutral |
| testneg_2156 | 0.04 | -0.23 | 3.570813 | -3 | -0.172518 | -3 | 0.68054 | 0.09208 | 0.19452 | 0.79779 | 10.03745 | 0.160311303 | 0 | Neutral |
| testneg_2157 | 0.02 | -0.68 | 2.77906 | -3 | 2.117525 | -3 | 0.35422 | 0.11185 | 0.37463 | 0.23021 | 28.84083 | 0.160281715 | 0 | Neutral |
| testneg_2158 | 0.21 | 0.15 | -0.762386 | -3 | 1.57381 | -3 | 0.07014 | 0.53339 | 0.13796 | 0.27869 | 19.50687 | 0.30525 | 1 | Neutral |
| testneg_2159 | 0.15 | 0.06 | -3.376225 | -3 | 1.476163 | -3 | 0.04123 | 0.06896 | 0.19898 | 0.09395 | 26.50719 | 0.050305944 | 0 | Neutral |
| testneg_2160 | -0.15 | -0.04 | 0.939929 | -3 | 5.898121 | -3 | 0.18701 | 0.12788 | 0.1296 | 0.164 | 13.51351 | 0.486761905 | 48.6 | Neutral |
| testneg_2161 | 0.02 | 0.45 | 3.640221 | 3.640221 | 1.366314 | -3 | 0.34673 | 0.24308 | 0.35051 | 0.11664 | 13.27141 | 0.46 | 38.8 | Neutral |
| testneg_2162 | 0 | 0 | 0.930011 | 0.930011 | 2.28341 | -3 | 0.12786 | 0.72066 | 0.27723 | 0.13229 | 16.84579 | 0.22639411 | 0.2 | Neutral |
| testneg_2163 | -0.06 | -0.18 | 0.491194 | 0.491194 | 2.768215 | -3 | 0.72124 | 0.09906 | 0.26928 | 0.12774 | 13.51351 | 0.041666667 | 0 | Neutral |
| testneg_2164 | 0.07 | 0.38 | 5.36374 | -3 | 0.055981 | -3 | 0.40179 | 0.12112 | 0.06547 | 0.09123 | 18.34652 | 0.11 | 0 | Neutral |
| testneg_2165 | 0 | -0.01 | 0.359803 | -3 | -0.102403 | -3 | 0.01635 | 0.12788 | 0.19355 | 0.18675 | 16.8018 | 0.08 | 0 | Neutral |
| testneg_2166 | 0 | 0 | 2.30653 | -3 | 5.23082 | -3 | 0.49106 | 0.89143 | 0.90112 | 0.20622 | 10.41667 | 0.43485005 | 31.4 | Neutral |
| testneg_2167 | -0.08 | -0.03 | -0.741718 | -3 | 1.11036 | -3 | 0.06218 | 0.10661 | 0.45399 | 0.10245 | 30.48962 | 0.03 | 0 | Neutral |
| testneg_2168 | 0 | 0 | -0.859115 | -3 | 1.630979 | -3 | 0.19898 | 0.12788 | 0.35532 | 0.24703 | 22.72602 | 0.053121212 | 0 | Neutral |
| testneg_2169 | 0.14 | 0.54 | 4.368956 | -3 | 3.22287 | -3 | 0.94229 | 0.26731 | 0.11983 | 0.24162 | 21.32913 | 0.632883929 | 95.8 | Disease_causing |
| testneg_2170 | 0.04 | -0.23 | 3.087586 | -3 | -0.133882 | -3 | 0.3828 | 0.5569 | 0.1056 | 0.12788 | 13.91667 | 0.11 | 0 | Neutral |
| testneg_2171 | -0.16 | -0.08 | 1.758868 | -3 | 0.124579 | -3 | 0.39163 | 0.2681 | 0.18537 | 0.1111 | 16.668 | 0.08 | 0 | Neutral |
| testneg_2172 | 0 | 0 | 1.864882 | -3 | -1.528492 | -3 | 0.39648 | 0.97288 | 0.12581 | 0.08118 | 10 | 0.07 | 0 | Neutral |
| testneg_2173 | 0.02 | 0.45 | 3.640221 | 3.640221 | 5.50061 | -3 | 0.34673 | 0.24308 | 0.09972 | 0.2187 | 27.95796 | 0.85 | 100 | Disease_causing |
| testneg_2174 | 0.02 | 0.45 | 4.330421 | -3 | 2.839131 | -3 | 0.10562 | 0.09329 | 0.62199 | 0.14032 | 12.46106 | 0.270665991 | 0.6 | Neutral |
| testneg_2175 | 0.04 | -0.23 | 7.332994 | -3 | 1.883305 | -3 | 0.1434 | 0.09092 | 0.54891 | 0.55746 | 32.91748 | 0.41 | 22.6 | Neutral |
| testneg_2176 | 0 | -0.01 | -0.099884 | -3 | 5.7343 | -3 | 0.0887 | 0.10047 | 0.06112 | 0.06901 | 21.49912 | 0.090250627 | 0 | Neutral |
| testneg_2177 | 0.01 | 0.8 | 1.074849 | -3 | 2.151559 | -3 | 0.06775 | 0.09511 | 0.82591 | 0.12207 | 10.625 | 0.05 | 0 | Neutral |
| testneg_2178 | -0.08 | -0.03 | 1.50021 | -3 | 2.400759 | -3 | 0.27618 | 0.11809 | 0.07872 | 0.12788 | 19.32978 | 0.02 | 0 | Neutral |
| testneg_2179 | 0.03 | 0.1 | 5.650756 | -3 | 5.397494 | -3 | 0.07973 | 0.09208 | 0.93001 | 0.11682 | 20.62016 | 0.31 | 6.4 | Neutral |
| testneg_2180 | 0 | 0 | -0.031197 | -3 | 0.603592 | -3 | 0.1017 | 0.11809 | 0.08973 | 0.0992 | 19.34809 | 0.01 | 0 | Neutral |
| testneg_2181 | 0 | 0 | 0.159372 | 0.159372 | 0.096056 | -3 | 0.05228 | 0.12788 | 0.41655 | 0.49485 | 22.03458 | 0.15 | 0 | Neutral |
| testneg_2182 | -0.03 | -0.1 | 1.609601 | -3 | 1.731383 | -3 | 0.3693 | 0.12788 | 0.1617 | 0.17928 | 10.83333 | 0.05132906 | 0 | Neutral |
| testneg_2183 | 0.11 | -0.19 | 2.575251 | -3 | 1.714629 | -3 | 0.8426 | 0.47836 | 0.03235 | 0.0948 | 17.77778 | 0.07 | 0 | Neutral |
| testneg_2184 | 0 | 0 | 0.660912 | -3 | 10.549799 | -3 | 0.33818 | 0.12788 | 0.10285 | 0.10886 | 18.88889 | 0.34 | 6.8 | Neutral |
| testneg_2185 | 0 | -0.01 | 0.71533 | -3 | 2.855535 | -3 | 0.42802 | 0.12096 | 0.46058 | 0.12903 | 22.95682 | 0.020606618 | 0 | Neutral |
| testneg_2186 | 0.14 | -0.37 | 3.878731 | -3 | 2.835668 | -3 | 0.06957 | 0.07707 | 0.72946 | 0.28603 | 9.2154 | 0.378730769 | 9.8 | Neutral |
| testneg_2187 | 0 | 0 | 1.538591 | -3 | 0.991286 | -3 | 0.64302 | 0.11262 | 0.2439 | 0.08848 | 10.6451 | 0.000412415 | 0 | Neutral |
| testneg_2188 | 0 | -0.01 | 2.059017 | -3 | 2.603226 | -3 | 0.18206 | 0.10727 | 0.1653 | 0.0977 | 13.51351 | 0.010404579 | 0 | Neutral |
| testneg_2189 | 0 | 0 | 0.342408 | -3 | 1.190149 | -3 | 0.5255 | 0.15968 | 0.16841 | 0.11143 | 22.21675 | 0.03 | 0 | Neutral |
| testneg_2190 | -0.14 | 0.37 | 1.688437 | -3 | 1.043597 | -3 | 0.15155 | 0.09923 | 0.19898 | 0.1005 | 18.05832 | 0.020467308 | 0 | Neutral |
| testneg_2191 | 0.11 | -0.19 | 4.821873 | -3 | 0.805308 | -3 | 0.78664 | 0.11725 | 0.19898 | 0.12788 | 18.33333 | 0.05 | 0 | Neutral |
| testneg_2192 | -0.07 | -0.38 | 2.097779 | -3 | -0.617826 | -3 | 0.81054 | 0.28104 | 0.06395 | 0.0759 | 13.64341 | 0.020666667 | 0 | Neutral |
| testneg_2193 | 0 | 0 | 0.297064 | -0.109042 | 1.258706 | -3 | 0.2029 | 0.12788 | 0.85345 | 0.13067 | 27.4024 | 0.020078704 | 0 | Neutral |
| testneg_2194 | -0.09 | 2.17 | 4.626824 | -3 | -0.102422 | -3 | 0.1296 | 0.164 | 0.32089 | 0.1035 | 24.03664 | 0.1645 | 0 | Neutral |
| testneg_2195 | 0 | 0 | -0.553742 | -3 | 1.464526 | -3 | 0.1424 | 0.12788 | 0.12513 | 0.12171 | 12.36538 | 0.000372294 | 0 | Neutral |

| | | | | | | | | | | | | | | |
|---|---|---|---|---|---|---|---|---|---|---|---|---|---|---|
| testneg_2196 | 0 | 0 | 0.393827 | -3 | 2.310369 | 2.310369 | 0.07516 | 0.15472 | 0.54895 | 0.15082 | 10.41667 | 0.169833333 | 0.2 | Neutral |
| testneg_2197 | -0.08 | -0.03 | 0.287286 | -3 | 1.491183 | -3 | 0.1592 | 0.12788 | 0.31043 | 0.09688 | 18.64268 | 0.01 | 0 | Neutral |
| testneg_2198 | -0.08 | -0.03 | -0.514498 | -3 | 1.673428 | 1.673428 | 0.81305 | 0.17045 | 0.32352 | 0.10243 | 13.02635 | 0.040190812 | 0 | Neutral |
| testneg_2199 | -0.03 | -0.1 | -1.42588 | -3 | 4.115851 | -3 | 0.16469 | 0.16042 | 0.07258 | 0.06248 | 16.88659 | 0.1571337 | 0 | Neutral |
| testneg_2200 | 0 | -0.01 | 1.789938 | -3 | 11.796446 | -3 | 0.33304 | 0.15588 | 0.13626 | 0.12788 | 18.59649 | 0.47 | 41.8 | Neutral |
| testneg_2201 | 0 | 0 | 0.634312 | -3 | -2.282519 | -3 | 0.07681 | 0.22865 | 0.1485 | 0.30134 | 19.31062 | 0.160083333 | 0 | Neutral |
| testneg_2202 | 0 | 0 | 1.511108 | -3 | 5.393665 | 5.393665 | 0.05873 | 0.09792 | 0.03809 | 0.16044 | 21.49819 | 0.49 | 50.6 | Disease_causing |
| testneg_2203 | -0.03 | 1.03 | 5.956083 | -3 | 6.964138 | -3 | 0.52278 | 0.41207 | 0.06395 | 0.0759 | 8.00964 | 0.709761905 | 98.6 | Disease_causing |
| testneg_2204 | 0.05 | -0.81 | 3.730075 | -3 | 1.08726 | -3 | 0.51577 | 0.19561 | 0.06667 | 0.08666 | 11.58108 | 0.07 | 0 | Neutral |
| testneg_2205 | 0 | 0 | 1.323079 | -3 | 0.90918 | -3 | 0.5103 | 0.10923 | 0.098 | 0.08791 | 26.09616 | 0.02 | 0 | Neutral |
| testneg_2206 | -0.04 | 0.89 | 4.817344 | -3 | 3.876605 | -3 | 0.44644 | 0.10813 | 0.83732 | 0.19325 | 25.02646 | 0.452571712 | 35.2 | Neutral |
| testneg_2207 | -0.01 | -0.79 | 1.556123 | -3 | 0.820354 | -3 | 0.08775 | 0.12788 | 0.21587 | 0.11701 | 13.51351 | 0.01 | 0 | Neutral |
| testneg_2208 | -0.08 | -0.03 | 3.093558 | -3 | 1.441342 | -3 | 0.28499 | 0.10637 | 0.11072 | 0.38259 | 15.63034 | 0.204067369 | 0 | Neutral |
| testneg_2209 | 0.07 | 0.01 | 0.631605 | -3 | 3.411835 | -3 | 0.14524 | 0.10162 | 0.09504 | 0.38939 | 20.8066 | 0.266499286 | 1 | Neutral |
| testneg_2210 | 0 | 0 | -0.535508 | -3 | 5.494352 | -3 | 0.01196 | 0.12788 | 0.14646 | 0.36445 | 19.44444 | 0.337130589 | 5.6 | Neutral |
| testneg_2211 | 0.04 | -0.23 | 7.332994 | -3 | 1.019613 | -3 | 0.1434 | 0.09092 | 0.08789 | 0.09565 | 29.37754 | 0.25 | 1.6 | Neutral |
| testneg_2212 | 0 | 0 | 1.66597 | -3 | -0.566651 | -3 | 0.66027 | 0.23984 | 0.38016 | 0.14503 | 18.33333 | 0.055549398 | 0 | Neutral |
| testneg_2213 | 0 | 0 | 0.586127 | -3 | 5.218039 | -3 | 0.19898 | 0.12788 | 0.07675 | 0.11464 | 27.46106 | 0.06134957 | 0 | Neutral |
| testneg_2214 | 0 | 0 | 1.436812 | -3 | 1.70493 | -3 | 0.29302 | 0.17394 | 0.72068 | 0.99841 | 6.35445 | 0.081161616 | 0 | Neutral |
| testneg_2215 | 0 | 0 | -0.535508 | -3 | 6.384552 | -3 | 0.01196 | 0.12788 | 0.05873 | 0.09792 | 11.56023 | 0.17 | 0 | Neutral |
| testneg_2216 | 0.18 | -1.05 | 3.764078 | -3 | 3.685438 | -3 | 0.15267 | 0.12788 | 0.16545 | 0.10002 | 28.04989 | 0.13 | 0 | Neutral |
| testneg_2217 | 0.01 | 0.8 | 1.842694 | -3 | 1.048529 | -3 | 0.19898 | 0.12788 | 0.54625 | 0.11856 | 11.25 | 0.02 | 0 | Neutral |
| testneg_2218 | 0.14 | -0.69 | 0.627341 | -3 | 0.506986 | -3 | 0.03991 | 0.08342 | 0.65857 | 0.26294 | 12.40248 | 0.11 | 0 | Neutral |
| testneg_2219 | -0.11 | 1.95 | 5.384186 | -3 | -0.038654 | -3 | 0.18998 | 0.26095 | 0.18206 | 0.10727 | 15.83333 | 0.263636364 | 1 | Neutral |
| testneg_2220 | 0.06 | -0.57 | 2.623327 | -3 | 0.790296 | -3 | 0.12786 | 0.72066 | 0.049 | 0.11324 | 48.08819 | 0.12 | 0 | Neutral |
| testneg_2221 | 0 | 0 | 1.067933 | -3 | 0.372849 | -3 | 0.08775 | 0.12788 | 0.10286 | 0.26819 | 18.6768 | 0.069798273 | 0 | Neutral |
| testneg_2222 | 0.2 | -1.26 | 1.812946 | -3 | 1.467602 | -3 | 0.18153 | 0.10845 | 0.29523 | 0.27623 | 12.35927 | 0.120729443 | 0 | Neutral |
| testneg_2223 | 0.23 | -1.86 | 3.479077 | -3 | 0.22446 | -3 | 0.17487 | 0.1672 | 0.32144 | 0.11774 | 10.12902 | 0.11 | 0 | Neutral |
| testneg_2224 | -0.18 | -0.16 | 2.954569 | -3 | 5.682246 | -3 | 0.58677 | 0.11579 | 0.47019 | 0.12788 | 19.11082 | 0.268208042 | 0 | Neutral |
| testneg_2225 | 0.12 | -1.16 | 4.661827 | -3 | 1.070904 | -3 | 0.13023 | 0.29692 | 0.40361 | 0.17821 | 9.96199 | 0.381035714 | 14.6 | Neutral |
| testneg_2226 | -0.04 | 0.23 | 0.396034 | -3 | 1.835259 | -3 | 0.61343 | 0.11838 | 0.14767 | 0.09559 | 20.83281 | 0.03 | 0 | Neutral |
| testneg_2227 | 0.19 | -0.27 | 2.820155 | -3 | -0.058678 | -3 | 0.06254 | 0.23561 | 0.16002 | 0.10481 | 23.48703 | 0.1 | 0 | Neutral |
| testneg_2228 | 0.18 | 0.16 | 2.268681 | -3 | 3.061286 | -3 | 0.25236 | 0.11809 | 0.07537 | 0.1399 | 14.27748 | 0.23 | 0 | Neutral |
| testneg_2229 | -0.04 | 0.23 | 1.485276 | -3 | 6.952572 | -3 | 0.39087 | 0.10381 | 0.12077 | 0.11244 | 28.659 | 0.240607143 | 1 | Neutral |
| testneg_2230 | 0.04 | -0.23 | 2.907554 | -3 | -1.290287 | -3 | 0.09356 | 0.2448 | 0.18328 | 0.12788 | 11.06295 | 0.1 | 0 | Neutral |
| testneg_2231 | 0.19 | -0.25 | 0.670876 | -3 | 0.202818 | -3 | 0.09645 | 0.13239 | 0.07118 | 0.08861 | 18.54402 | 0.03 | 0 | Neutral |
| testneg_2232 | 0.18 | -1.05 | 5.645081 | -3 | 0.275531 | -3 | 0.30739 | 0.16306 | 0.06067 | 0.07796 | 28.84829 | 0.17625 | 0 | Neutral |
| testneg_2233 | -0.15 | -0.04 | 1.067435 | -3 | 3.592336 | -3 | 0.06711 | 0.09765 | 0.03297 | 0.16627 | 12.57218 | 0.295138889 | 1 | Neutral |
| testneg_2234 | 0.22 | -2.66 | 6.87277 | -3 | -0.294681 | -3 | 0.09724 | 0.09172 | 0.4129 | 0.11172 | 37.04096 | 0.19 | 0 | Neutral |
| testneg_2235 | -0.0083217 | -0.1418881 | -0.105139 | -3 | 2.117024 | 0.932372 | 0.08489 | 0.12788 | 0.53784 | 0.5574 | 7.00579 | 0.112334805 | 0 | Neutral |
| testneg_2236 | 0.06 | 0.18 | 2.289276 | -3 | 0.789388 | 0.513139 | 0.32977 | 0.15427 | 0.6532 | 0.12788 | 13.01229 | 0.03 | 0 | Neutral |
| testneg_2237 | 0.07 | 0.7 | 5.597504 | -3 | 0.631367 | -3 | 0.10187 | 0.12339 | 0.76138 | 0.16258 | 30.01332 | 0.27 | 1.8 | Neutral |
| testneg_2238 | -0.18 | 1.05 | 2.40944 | 1.660423 | 1.877801 | -3 | 0.17075 | 0.12788 | 0.25312 | 0.12788 | 32.99221 | 0.12019434 | 0 | Neutral |
| testneg_2239 | 0 | 0 | 0.55724 | 0.55724 | 1.672333 | 1.672333 | 0.12937 | 0.23004 | 0.15338 | 0.13614 | 22.40741 | 0.189061728 | 0 | Neutral |
| testneg_2240 | 0.02 | -0.63 | -2.323867 | -3 | 0.36854 | -3 | 0.07595 | 0.11822 | 0.58292 | 0.87725 | 23.7623 | 0.111545455 | 0 | Neutral |
| testneg_2241 | 0.02 | 0.41 | 0.799457 | -3 | 5.180313 | -3 | 0.08714 | 0.08945 | 0.19898 | 0.12788 | 20.83333 | 0.082303419 | 0 | Neutral |
| testneg_2242 | 0.07 | -0.31 | 0.268697 | -3 | -1.816133 | -3 | 0.20657 | 0.12788 | 0.42366 | 0.14372 | 18.33333 | 0.04023823 | 0 | Neutral |
| testneg_2243 | 0.18 | -1.05 | 1.792345 | -3 | 1.504305 | -3 | 0.14806 | 0.09574 | 0.22233 | 0.31758 | 20 | 0.132176802 | 0 | Neutral |
| testneg_2244 | 0 | 0 | -0.262607 | -3 | 0.293949 | -3 | 0.21576 | 0.17241 | 0.16616 | 0.11073 | 20 | 0.01086663 | 0 | Neutral |
| testneg_2245 | 0.01 | 0.8 | 4.988796 | 0.968512 | 1.445664 | -3 | 0.40074 | 0.36296 | 0.14038 | 0.21177 | 26.38889 | 0.528535571 | 64.6 | Disease_causing |
| testneg_2246 | 0.14 | -0.69 | 2.950487 | -3 | 7.833352 | -3 | 0.14495 | 0.37127 | 0.08067 | 0.34914 | 57.48821 | 0.869287578 | 100 | Disease_causing |
| testneg_2247 | 0 | 0 | -0.378226 | -3 | 2.395393 | -3 | 0.99668 | 0.30319 | 0.6271 | 0.09663 | 6.35445 | 0.07 | 0 | Neutral |
| testneg_2248 | 0.18 | -1.05 | 3.656555 | -3 | 4.182656 | -3 | 0.1647 | 0.12076 | 0.12933 | 0.12788 | 21.41799 | 0.180597235 | 0 | Neutral |
| testneg_2249 | 0 | 0 | 1.731602 | -3 | 2.457626 | -3 | 0.0146 | 0.03045 | 0.33952 | 0.12788 | 19.81145 | 0.05 | 0 | Neutral |
| testneg_2250 | 0.15 | -0.82 | 2.176784 | -3 | 6.75693 | -3 | 0.27887 | 0.18663 | 0.18175 | 0.13527 | 10.83333 | 0.649295615 | 96.2 | Disease_causing |
| testneg_2251 | 0 | 0 | -0.32247 | -3 | 4.226361 | -3 | 0.35092 | 0.10596 | 0.82846 | 0.13042 | 10.83333 | 0.07 | 0 | Neutral |
| testneg_2252 | 0.03 | 0.12 | -0.432053 | -3 | 1.215719 | -3 | 0.10472 | 0.10014 | 0.56487 | 0.12788 | 18.6768 | 0.01 | 0 | Neutral |
| testneg_2253 | 0 | 0 | 0.245964 | -3 | 2.087834 | 1.600011 | 0.74099 | 0.44443 | 0.19898 | 0.12788 | 9.98688 | 0.030204082 | 0 | Neutral |
| testneg_2254 | 0 | 0 | 1.081416 | -3 | 0.47202 | -3 | 0.59345 | 0.26484 | 0.13295 | 0.12914 | 19.43262 | 0.010930736 | 0 | Neutral |
| testneg_2255 | 0.01166667 | -0.0684804 | 2.8284 | -3 | -0.445161 | -0.445161 | 0.20417 | 0.10681 | 0.09728 | 0.12788 | 48.95686 | 0.020606618 | 0 | Neutral |
| testneg_2256 | 0 | 0 | 1.932433 | -3 | 1.614723 | 1.614723 | 0.54248 | 0.12333 | 0.74944 | 0.1263 | 4.72222 | 0.02 | 0 | Neutral |
| testneg_2257 | -0.01 | -0.79 | 1.315391 | -3 | 1.096466 | -3 | 0.99417 | 0.17365 | 0.49745 | 0.1259 | 21.94444 | 0.053555008 | 0 | Neutral |
| testneg_2258 | 0.02 | -0.28 | 5.749527 | -3 | 5.070556 | -3 | 0.1214 | 0.0899 | 0.10185 | 0.12788 | 35.76924 | 0.311524927 | 5.6 | Neutral |
| testneg_2259 | 0.03 | 1.22 | 5.217351 | 2.878966 | -0.252838 | -3 | 0.28227 | 0.10211 | 0.0611 | 0.12135 | 10.41667 | 0.43 | 32.6 | Neutral |
| testneg_2260 | -0.07 | -0.38 | 2.459203 | -3 | 1.802906 | -3 | 0.23292 | 0.15913 | 0.33886 | 0.15644 | 11.25 | 0.131708586 | 0 | Neutral |
| testneg_2261 | 0 | 0 | 3.603309 | -3 | -0.133106 | -0.133106 | 0.26192 | 0.15452 | 0.07096 | 0.09965 | 13.39912 | 0.040480549 | 0 | Neutral |
| testneg_2262 | 0 | 0 | 0.560537 | -3 | 6.577474 | -3 | 0.32804 | 0.10857 | 0.08646 | 0.24855 | 27.26682 | 0.37162381 | 12.4 | Neutral |
| testneg_2263 | 0 | 0 | 0.701413 | -3 | -0.292428 | -3 | 0.1214 | 0.0899 | 0.14862 | 0.12788 | 33.49673 | 0.001003715 | 0 | Neutral |
| testneg_2264 | -0.05 | -0.29 | 2.048887 | 2.048887 | 2.475201 | -3 | 0.28128 | 0.12883 | 0.11264 | 0.11163 | 11.59396 | 0.1 | 0 | Neutral |
| testneg_2265 | 0.14 | -1.01 | 7.382501 | -3 | -1.528753 | -1.528753 | 0.60703 | 0.12378 | 0.10482 | 0.12788 | 20.66855 | 0.28225 | 3.8 | Neutral |
| testneg_2266 | 0.08 | 0.03 | -0.412359 | -3 | 1.662324 | -3 | 0.4003 | 0.10884 | 0.16777 | 0.1749 | 17.50754 | 0.075712229 | 0 | Neutral |
| testneg_2267 | 0.16 | 0.08 | 0.019411 | -3 | 1.232126 | -3 | 0.10863 | 0.098 | 0.25893 | 0.10353 | 17.73063 | 0.01 | 0 | Neutral |
| testneg_2268 | 0 | 0 | 3.979527 | -3 | 5.185173 | -3 | 0.20914 | 0.10089 | 0.06118 | 0.08724 | 10.30286 | 0.192291667 | 0 | Neutral |
| testneg_2269 | -0.03 | -0.1 | 0.325459 | -3 | 4.714948 | -3 | 0.05027 | 0.10981 | 0.20327 | 0.11174 | 7.02397 | 0.1 | 0 | Neutral |
| testneg_2270 | 0.03 | -1.03 | 3.410902 | -3 | -0.410022 | -3 | 0.20701 | 0.10345 | 0.17419 | 0.33842 | 9.33732 | 0.184603175 | 0 | Neutral |
| testneg_2271 | 0.03 | 0.1 | -1.156564 | -3 | -0.272373 | -3 | 0.54213 | 0.09044 | 0.19898 | 0.29199 | 10.22898 | 0.061404762 | 0 | Neutral |
| testneg_2272 | 0.02 | 0.41 | -2.259592 | -3 | -0.060754 | -3 | 0.06311 | 0.14802 | 0.70946 | 0.11809 | 29.5935 | 0.07 | 0 | Neutral |
| testneg_2273 | 0.01 | 0.8 | 2.486418 | -3 | 1.7309 | -3 | 0.09211 | 0.08973 | 0.21399 | 0.10708 | 18.59649 | 0.02 | 0 | Neutral |
| testneg_2274 | -0.15 | -0.04 | 6.902692 | -3 | 2.97409 | -3 | 0.13356 | 0.09963 | 0.16544 | 0.43246 | 10.80534 | 0.656504099 | 94 | Disease_causing |
| testneg_2275 | 0.14 | -1.01 | -0.270969 | -3 | 1.26259 | 1.26259 | 0.04441 | 0.13603 | 0.44722 | 0.11641 | 12.38369 | 0.06 | 0 | Neutral |
| testneg_2276 | -0.05 | -0.29 | -2.95337 | -3 | 0.478518 | -3 | 0.19898 | 0.12788 | 0.89959 | 0.55078 | 12.12979 | 0.051363941 | 0 | Neutral |
| testneg_2277 | -0.09 | 0.3 | 1.470083 | -3 | -0.493225 | -3 | 0.19969 | 0.1163 | 0.09235 | 0.18135 | 12.89568 | 0.070185759 | 0 | Neutral |
| testneg_2278 | 0.14 | -0.37 | 1.764776 | -3 | 1.627059 | -3 | 0.6975 | 0.53794 | 0.19322 | 0.12788 | 24.02248 | 0.082210784 | 0 | Neutral |
| testneg_2279 | 0.01 | -0.57 | -0.236734 | -3 | 1.284188 | -3 | 0.0832 | 0.09323 | 0.17808 | 0.12788 | 17.21847 | 0.01 | 0 | Neutral |
| testneg_2280 | 0.21 | 0.15 | 3.945475 | -3 | 1.699389 | -3 | 0.27115 | 0.14766 | 0.19898 | 0.12788 | 21.05089 | 0.080787037 | 0 | Neutral |
| testneg_2281 | 0.05 | -0.81 | 2.225911 | -3 | -1.781198 | -3 | 0.91619 | 0.10042 | 0.43897 | 0.12441 | 15.40753 | 0.020502874 | 0 | Neutral |
| testneg_2282 | 0.21 | 0.15 | 1.523309 | -3 | 0.632016 | -3 | 0.05378 | 0.12788 | 0.21289 | 0.08709 | 19.07749 | 0.050616601 | 0 | Neutral |
| testneg_2283 | 0.03 | 0.12 | 0.636994 | -3 | -0.025657 | -3.451557 | 0.1601 | 0.12546 | 0.16259 | 0.102 | 13.51351 | 0.01 | 0 | Neutral |
| testneg_2284 | -0.21 | -0.15 | 0.71504 | -3 | 5.312389 | -3 | 0.21894 | 0.09041 | 0.07293 | 0.12989 | 11.08369 | 0.18105618 | 0.2 | Neutral |
| testneg_2285 | -0.04 | 0.23 | 0.729199 | -3 | 1.940661 | -3 | 0.11373 | 0.10739 | 0.25151 | 0.12788 | 18.70817 | 0.01 | 0 | Neutral |
| testneg_2286 | 0.06 | -0.57 | 0.282279 | -3 | 0.201766 | -3 | 0.03526 | 0.06855 | 0.16388 | 0.14439 | 29.86111 | 0.1 | 0 | Neutral |
| testneg_2287 | 0 | 0 | 1.446796 | -3 | 2.048887 | -3 | 0.24528 | 0.37575 | 0.28128 | 0.12883 | 19.41289 | 0.037083333 | 0 | Neutral |
| testneg_2288 | 0.14 | -0.37 | 4.088702 | -3 | 0.159372 | 0.159372 | 0.14632 | 0.12662 | 0.05228 | 0.12788 | 16.88659 | 0.04 | 0 | Neutral |
| testneg_2289 | 0.01 | 0.8 | 0.480519 | -3 | 1.267038 | -3 | 0.08714 | 0.08945 | 0.19898 | 0.12788 | 18.33333 | 0.02 | 0 | Neutral |
| testneg_2290 | 0.03 | -1.03 | 5.957726 | -3 | 5.017223 | -3 | 0.44103 | 0.10698 | 0.06744 | 0.12788 | 13.46801 | 0.45 | 33.4 | Neutral |
| testneg_2291 | -0.16 | 0.37 | -2.275335 | -3 | 1.261852 | -3 | 0.05663 | 0.12788 | 0.14638 | 0.12788 | 29.79424 | 0.120280338 | 0 | Neutral |
| testneg_2292 | 0.07 | 0.7 | 0.08175 | -3 | 0.542661 | -3 | 0.32334 | 0.10525 | 0.32531 | 0.12351 | 10.71967 | 0.02 | 0 | Neutral |
| testneg_2293 | 0.18 | 0.16 | -2.635087 | -3 | -0.135555 | -3 | 0.19898 | 0.12788 | 0.09657 | 0.58605 | 12.30981 | 0.12 | 0 | Neutral |
| testneg_2294 | 0 | 0 | 0.844548 | -3 | 0.058387 | -3 | 0.9237 | 0.55859 | 0.57349 | 0.09764 | 4.44444 | 0.010556918 | 0 | Neutral |
| testneg_2295 | 0.06 | 0.18 | 5.333191 | -3 | 1.75357 | -3 | 0.19898 | 0.12788 | 0.06736 | 0.06344 | 20.31976 | 0.12 | 0 | Neutral |
| testneg_2296 | 0.04 | -0.23 | 2.953578 | -3 | 2.42661 | -3 | 0.33957 | 0.12788 | 0.17226 | 0.13001 | 13.74564 | 0.08 | 0 | Neutral |
| testneg_2297 | 0 | 0 | -1.158589 | -3 | 0.196323 | -3 | 0.16362 | 0.11879 | 0.13693 | 0.12788 | 20.36859 | 0.000336729 | 0 | Neutral |
| testneg_2298 | 0.02 | -0.28 | -3.482963 | -3 | 2.079528 | -3 | 0.19898 | 0.12788 | 0.04627 | 0.05223 | 30.28233 | 0.024166667 | 0 | Neutral |
| testneg_2299 | 0.21 | 0.15 | 0.770086 | 0.770086 | 4.193908 | -3 | 0.20527 | 0.10053 | 0.23088 | 0.12788 | 28.87729 | 0.131666667 | 0 | Neutral |
| testneg_2300 | 0.17 | 0.38 | 1.603588 | -3 | 1.416956 | -3 | 0.10496 | 0.21491 | 0.10314 | 0.08894 | 11.25 | 0.071984994 | 0 | Neutral |
| testneg_2301 | 0.07 | 0.01 | 1.388898 | -3 | 5.611956 | -3 | 0.17146 | 0.19882 | 0.14187 | 0.1353 | 10 | 0.514705413 | 60.6 | Disease_causing |
| testneg_2302 | -0.08 | -0.03 | 2.829858 | -3 | -0.303489 | -3 | 0.10895 | 0.10442 | 0.508 | 0.1281 | 10.83333 | 0.030386905 | 0 | Neutral |
| testneg_2303 | 0 | 0 | 0.393827 | -3 | 0.997782 | -3 | 0.07516 | 0.15472 | 0.48817 | 0.10601 | 18.6768 | 0.02 | 0 | Neutral |
| testneg_2304 | -0.03 | -0.1 | 1.347962 | -3 | 1.51996 | -3 | 0.0418 | 0.12788 | 0.09105 | 0.27434 | 61.07469 | 0.230054945 | 0.2 | Neutral |
| testneg_2305 | 0 | 0 | 0.269414 | -3 | 6.897399 | -3 | 0.43921 | 0.153 | 0.09426 | 0.1658 | 18.88889 | 0.542038534 | 71.2 | Disease_causing |
| testneg_2306 | 0 | 0 | -0.287761 | -3 | 1.385917 | -3 | 0.85345 | 0.13067 | 0.98641 | 0.83983 | 10.83333 | 0.07 | 0 | Neutral |
| testneg_2307 | 0 | 0 | -0.262607 | -3 | 2.947657 | -3 | 0.21576 | 0.17241 | 0.00971 | 0.0791 | 18.33333 | 0.05 | 0 | Neutral |
| testneg_2308 | 0 | -0.01 | 0.169949 | -3 | 3.061286 | -3 | 0.38445 | 0.58278 | 0.07537 | 0.1399 | 11.875 | 0.262446286 | 0.4 | Neutral |
| testneg_2309 | 0.07 | 0.01 | -2.597352 | -3 | 1.856425 | -3 | 0.72062 | 0.14321 | 0.09806 | 0.12788 | 17.21847 | 0.04 | 0 | Neutral |
| testneg_2310 | -0.03 | -1.22 | -1.294775 | -3 | 2.967342 | -3 | 0.50063 | 0.154 | 0.19898 | 0.09927 | 16.26913 | 0.104671717 | 0 | Neutral |
| testneg_2311 | 0.03 | -1.03 | 2.432661 | -3 | -0.90985 | -3 | 0.1541 | 0.1077 | 0.15617 | 0.24339 | 56.5741 | 0.140244484 | 0 | Neutral |
| testneg_2312 | 0.05 | -0.81 | 7.467882 | -3 | 3.840037 | -3 | 0.10774 | 0.13117 | 0.49684 | 0.29035 | 10.41667 | 0.76 | 99.8 | Disease_causing |
| testneg_2313 | 0 | 0 | 0.08585 | -3 | 1.681428 | -3 | 0.5471 | 0.11262 | 0.59691 | 0.11049 | 22.64794 | 0.02 | 0 | Neutral |
| testneg_2314 | 0 | 0 | 1.537848 | -3 | 0.224925 | -3 | 0.11391 | 0.12788 | 0.03676 | 0.12788 | 18.33333 | 0.000597148 | 0 | Neutral |
| testneg_2315 | 0 | 0 | 0.475838 | -3 | 0.879231 | -3 | 0.25516 | 0.12788 | 0.13698 | 0.12788 | 13.51351 | 0 | 0 | Neutral |
| testneg_2316 | 0.05 | 0.29 | 2.036657 | 2.036657 | 1.124718 | -3 | 0.91691 | 0.25567 | 0.55043 | 0.12112 | 11.75623 | 0.095677966 | 0.2 | Neutral |
| testneg_2317 | 0.04 | -0.23 | 4.017255 | -3 | -0.303923 | -3 | 0.19898 | 0.16603 | 0.04593 | 0.09315 | 18.33333 | 0.06 | 0 | Neutral |

| | | | | | | | | | | | | | | |
|---|---|---|---|---|---|---|---|---|---|---|---|---|---|---|
| testneg_2318 | 0.18 | -1.05 | 2.793735 | -3 | 3.568323 | -3 | 0.19898 | 0.12788 | 0.63798 | 0.19967 | 11.08369 | 0.397683081 | 21.8 | Neutral |
| testneg_2319 | 0 | 0 | 7.325251 | -3 | 1.104931 | -3 | 0.16601 | 0.14015 | 0.06708 | 0.09112 | 13.56809 | 0.340142857 | 10 | Neutral |
| testneg_2320 | 0.05 | -0.81 | 0.820857 | -3 | -1.295182 | -3 | 0.0887 | 0.10047 | 0.26913 | 0.1087 | 37.98501 | 0.01 | 0 | Neutral |
| testneg_2321 | -0.02 | 0.63 | 2.293151 | -3 | 2.022849 | -3 | 0.47997 | 0.10506 | 0.20708 | 0.12788 | 18.33333 | 0.0200761 | 0 | Neutral |
| testneg_2322 | 0.05 | 0.29 | 1.381522 | -3 | -0.269407 | -3 | 0.11325 | 0.17021 | 0.60079 | 0.11198 | 22.62411 | 0.04 | 0 | Neutral |
| testneg_2323 | 0 | 0 | -0.211085 | -3 | 0.185804 | -3 | 0.04543 | 0.28103 | 0.08306 | 0.12788 | 16.60714 | 0.070975524 | 0 | Neutral |
| testneg_2324 | 0.15 | -0.28 | -1.251158 | -3 | 0.329621 | 0.329621 | 0.20127 | 0.13478 | 0.4129 | 0.11172 | 30.85749 | 0.03 | 0 | Neutral |
| testneg_2325 | -0.18 | -0.16 | 0.767258 | -3 | 5.766968 | -3 | 0.09871 | 0.09286 | 0.19898 | 0.10593 | 18.05832 | 0.2 | 0 | Neutral |
| testneg_2326 | 0.14 | -0.37 | 0.531203 | -3 | 4.444591 | -3 | 0.30951 | 0.64738 | 0.22239 | 0.12885 | 11.25 | 0.223285714 | 0 | Neutral |
| testneg_2327 | -0.15 | 0.28 | 6.629133 | -3 | -0.599522 | -3 | 0.21814 | 0.13855 | 0.22225 | 0.11186 | 18.33333 | 0.27 | 2.4 | Neutral |
| testneg_2328 | 0 | 0 | 1.095963 | -3 | -1.284127 | -3 | 0.08846 | 0.06651 | 0.11096 | 0.10074 | 37.22222 | 0.01 | 0 | Neutral |
| testneg_2329 | 0 | -0.01 | 1.609618 | -3 | 0.407117 | -3 | 0.67038 | 0.1794 | 0.17728 | 0.46704 | 18.88889 | 0.080229167 | 0 | Neutral |
| testneg_2330 | 0.21 | 0.15 | 4.124651 | -3 | 0.183348 | -3 | 0.08393 | 0.1123 | 0.064 | 0.24721 | 39.38398 | 0.233703141 | 0 | Neutral |
| testneg_2331 | 0.16 | -1.37 | 2.896502 | -3 | 2.552263 | -3 | 0.18096 | 0.12788 | 0.1977 | 0.09728 | 29.61945 | 0.08 | 0 | Neutral |
| testneg_2332 | -0.02 | 0.68 | 0.381493 | 0.381493 | 2.987273 | -3 | 0.14432 | 0.16919 | 0.03502 | 0.08119 | 10.83333 | 0.112235507 | 0 | Neutral |
| testneg_2333 | 0.14 | -1.01 | 3.372659 | -3 | 0.316713 | -3 | 0.15263 | 0.12788 | 0.2755 | 0.12788 | 18.6768 | 0.02 | 0 | Neutral |
| testneg_2334 | -0.16 | -0.08 | 1.558809 | -3 | 6.268455 | -3 | 0.19898 | 0.12788 | 0.0594 | 0.16845 | 17.86874 | 0.5 | 55.2 | Disease_causing |
| testneg_2335 | -0.03 | -0.1 | 6.283472 | -3 | -2.618223 | -3 | 0.17529 | 0.09662 | 0.22179 | 0.09831 | 49.51585 | 0.163785714 | 0 | Neutral |
| testneg_2336 | 0 | -0.01 | 0.362326 | -3 | 2.432523 | -3 | 0.29417 | 0.10965 | 0.25011 | 0.12092 | 8.97898 | 0.01 | 0 | Neutral |
| testneg_2337 | -0.14 | 1.85 | 0.310587 | 0.310587 | -0.196592 | -0.196592 | 0.2132 | 0.12788 | 0.32733 | 0.16869 | 10.41667 | 0.180518398 | 0 | Neutral |
| testneg_2338 | -0.02 | 0.36 | 3.195501 | -3 | -0.603184 | -3 | 0.22899 | 0.31462 | 0.09176 | 0.07645 | 22.86321 | 0.071044408 | 0 | Neutral |
| testneg_2339 | 0.02 | 0.45 | 0.379358 | -3 | 5.330022 | -3 | 0.19898 | 0.10996 | 0.15676 | 0.12788 | 22.04795 | 0.093185606 | 0 | Neutral |
| testneg_2340 | -0.06 | -0.18 | 1.651093 | -3 | 0.179304 | -3 | 0.08332 | 0.09046 | 0.56817 | 0.10766 | 20.32051 | 0.010333333 | 0 | Neutral |
| testneg_2341 | -0.14 | 0.37 | 1.688437 | -3 | 0.958148 | -3 | 0.15155 | 0.09923 | 0.22575 | 0.37247 | 24.86268 | 0.110323899 | 0 | Neutral |
| testneg_2342 | -0.02 | 0.63 | 1.852975 | -3 | 1.599412 | 1.599412 | 0.18224 | 0.10419 | 0.13099 | 0.17453 | 18.72417 | 0.110202206 | 0 | Neutral |
| testneg_2343 | 0.1 | 1.9 | 4.284152 | -3 | -0.180006 | -3 | 0.2674 | 0.28725 | 0.09806 | 0.12788 | 10.41667 | 0.15 | 0 | Neutral |
| testneg_2344 | -0.07 | 0.31 | 0.335227 | -3 | -1.352736 | -3 | 0.04346 | 0.10785 | 0.27976 | 0.12613 | 16.8018 | 0.04 | 0 | Neutral |
| testneg_2345 | -0.0087273 | -0.0077576 | 1.035348 | -3 | 1.52981 | -3 | 0.87268 | 0.21837 | 0.34001 | 0.12788 | 19.51415 | 0.020202041 | 0 | Neutral |
| testneg_2346 | 0 | 0 | 1.315096 | -3 | -0.179376 | -3 | 0.26972 | 0.12484 | 0.99994 | 0.51636 | 5.92466 | 0.06019697 | 0 | Neutral |
| testneg_2347 | -0.07 | -0.33 | 1.183832 | -3 | 2.228748 | -3 | 0.3889 | 0.10831 | 0.27585 | 0.20571 | 12.52798 | 0.065202381 | 0 | Neutral |
| testneg_2348 | -0.16 | -0.08 | 3.003989 | -3 | -1.543536 | -3 | 0.33264 | 0.10493 | 0.06744 | 0.12788 | 20.42553 | 0.07 | 0 | Neutral |
| testneg_2349 | 0 | 0 | -0.101683 | -3 | -0.433937 | -0.433937 | 0.72657 | 0.11262 | 0.84572 | 0.78546 | 6.81965 | 0.06 | 0 | Neutral |
| testneg_2350 | 0.15 | 0.04 | 1.056 | 1.056 | 4.143027 | -3 | 0.21153 | 0.11357 | 0.11746 | 0.1471 | 18.88889 | 0.330542857 | 6.8 | Neutral |
| testneg_2351 | 0 | 0 | -0.297936 | -3 | 4.873924 | -3 | 0.23126 | 0.16717 | 0.66799 | 0.12788 | 5.97222 | 0.130869968 | 0 | Neutral |
| testneg_2352 | 0.18 | -1.05 | 3.974272 | -3 | 1.59057 | -3 | 0.6323 | 0.09928 | 0.26789 | 0.67432 | 10.41667 | 0.280175439 | 0.8 | Neutral |
| testneg_2353 | 0 | -0.01 | 7.854373 | -3 | 6.126062 | -3 | 0.4113 | 0.2303 | 0.17458 | 0.12788 | 46.47059 | 0.73 | 99.8 | Disease_causing |
| testneg_2354 | 0 | 0 | -0.577654 | -3 | 3.13959 | -3 | 0.08408 | 0.12788 | 0.10187 | 0.08499 | 18.33333 | 0.046287879 | 0 | Neutral |
| testneg_2355 | -0.03 | -0.1 | 5.381477 | -3 | 3.735674 | -3 | 0.34698 | 0.12788 | 0.13023 | 0.29692 | 5.73716 | 0.723750489 | 99.2 | Disease_causing |
| testneg_2356 | 0.05 | -0.81 | 1.20542 | -3 | 0.040537 | -3 | 0.06027 | 0.11574 | 0.45343 | 0.12014 | 10.83333 | 0.02245614 | 0 | Neutral |
| testneg_2357 | 0 | 0 | 1.031442 | -3 | 4.115885 | -3 | 0.2761 | 0.24559 | 0.09356 | 0.2448 | 12.75394 | 0.386930305 | 18.6 | Neutral |
| testneg_2358 | 0 | -0.01 | 0.71535 | -3 | 1.050349 | -3 | 0.92832 | 0.16306 | 0.19989 | 0.06568 | 12.34649 | 0.010192593 | 0 | Neutral |
| testneg_2359 | 0 | 0 | 0.889458 | 0.889458 | 0.717822 | -3 | 0.48937 | 0.22085 | 0.29044 | 0.12142 | 24.58607 | 0.040599867 | 0 | Neutral |
| testneg_2360 | -0.02 | -0.13 | 1.290429 | -1.019571 | -0.062057 | -3 | 0.19898 | 0.20319 | 0.26489 | 0.10549 | 38.98206 | 0.03 | 0 | Neutral |
| testneg_2361 | 0.16 | -0.37 | 5.42275 | -3 | 0.27631 | -3 | 0.44816 | 0.56655 | 0.19898 | 0.10591 | 10.41667 | 0.26 | 0.2 | Neutral |
| testneg_2362 | -0.02 | 0.28 | 2.201231 | -3 | 1.137849 | -3 | 0.20231 | 0.12788 | 0.15147 | 0.12788 | 23.20309 | 0.01 | 0 | Neutral |
| testneg_2363 | -0.01 | 0.57 | -2.581564 | -3 | -0.383235 | -3 | 0.19898 | 0.12788 | 0.87847 | 0.96194 | 15.63985 | 0.080666667 | 0 | Neutral |
| testneg_2364 | 0 | 0 | 1.040567 | -3 | 3.088745 | -3 | 0.10869 | 0.1503 | 0.17412 | 0.10595 | 11.19839 | 0.070347852 | 0 | Neutral |
| testneg_2365 | 0 | -0.01 | 2.663664 | -3 | 1.829958 | -3 | 0.19898 | 0.26604 | 0.47775 | 0.10103 | 22.19424 | 0.04 | 0 | Neutral |
| testneg_2366 | -0.01 | -0.8 | 4.655164 | -3 | 1.645021 | 1.645021 | 0.22899 | 0.31462 | 0.19898 | 0.14957 | 18.33333 | 0.355231407 | 11.6 | Neutral |
| testneg_2367 | 0 | 0 | -0.32829 | -3 | 3.805191 | -3 | 0.18509 | 0.12831 | 0.56307 | 0.10536 | 21.35235 | 0.07 | 0 | Neutral |
| testneg_2368 | 0 | 0 | 1.318565 | -3 | 0.245964 | -3 | 0.22658 | 0.20646 | 0.74099 | 0.44443 | 5.73716 | 0.062461362 | 0 | Neutral |
| testneg_2369 | 0 | -0.73 | 6.453651 | -3 | 1.762821 | 1.762821 | 0.07889 | 0.08835 | 0.1375 | 0.112 | 11.98582 | 0.265785714 | 1.8 | Neutral |
| testneg_2370 | -0.17 | -0.38 | 3.271662 | -3 | 1.708706 | -3 | 0.48662 | 0.23781 | 0.1019 | 0.17526 | 13.14653 | 0.339454451 | 6.4 | Neutral |
| testneg_2371 | 0 | -0.73 | 1.695441 | -3 | 2.12021 | 2.12021 | 0.15202 | 0.12722 | 0.07698 | 0.12788 | 30.48443 | 0.051172619 | 0 | Neutral |
| testneg_2372 | 0.07 | 0.7 | 3.069764 | -3 | 0.660523 | -3 | 0.25474 | 0.11262 | 0.06897 | 0.08445 | 32.98441 | 0.03 | 0 | Neutral |
| testneg_2373 | 0.14 | -0.37 | 1.075133 | 1.075133 | 4.026759 | -3 | 0.0606 | 0.39625 | 0.09616 | 0.11651 | 21.97988 | 0.28234944 | 0.8 | Neutral |
| testneg_2374 | 0 | 0 | 1.565142 | -3 | 6.835273 | -3 | 0.493 | 0.35835 | 0.08412 | 0.28307 | 19.63739 | 0.602859477 | 87.6 | Disease_causing |
| testneg_2375 | 0.07 | 0.38 | -1.416093 | -3 | 0.789411 | -3 | 0.12543 | 0.12788 | 0.13067 | 0.1005 | 23.68839 | 0.01 | 0 | Neutral |
| testneg_2376 | 0 | 0 | 1.44947 | -3 | 1.683005 | -3 | 0.18459 | 0.33541 | 0.19898 | 0.12788 | 18.52048 | 0.03 | 0 | Neutral |
| testneg_2377 | 0.01 | -1.44 | 1.53664 | -3 | 3.266081 | -3 | 0.17576 | 0.12193 | 0.09556 | 0.10109 | 24.68926 | 0.06 | 0 | Neutral |
| testneg_2378 | -0.01 | -0.8 | 1.614908 | -3 | 1.490016 | -3 | 0.71406 | 0.2012 | 0.93853 | 0.93439 | 10 | 0.13 | 0 | Neutral |
| testneg_2379 | 0 | 0 | 1.467396 | -3 | 0.379738 | -3 | 0.96995 | 0.66487 | 0.11731 | 0.06348 | 19.29699 | 0.040327957 | 0 | Neutral |
| testneg_2380 | -0.14 | -0.54 | -0.251883 | -3 | 0.243251 | -3 | 0.39258 | 0.12788 | 0.09402 | 0.12788 | 35.39101 | 0.03 | 0 | Neutral |
| testneg_2381 | -0.08 | -0.03 | 0.390383 | 0.390383 | 4.202319 | -3 | 0.08213 | 0.08913 | 0.19898 | 0.30319 | 19.09347 | 0.34 | 5.6 | Neutral |
| testneg_2382 | 0.06 | 0.18 | -1.019571 | -3 | 2.705771 | -3 | 0.19898 | 0.20319 | 0.06344 | 0.12788 | 27.51545 | 0.099603175 | 0 | Neutral |
| testneg_2383 | 0.07 | 0.38 | -2.188525 | -2.188525 | 2.031708 | -3 | 0.07282 | 0.12788 | 0.13061 | 0.21023 | 21.44363 | 0.10319463 | 0 | Neutral |
| testneg_2384 | -0.03 | -0.1 | 3.804607 | -3 | -0.303923 | -3 | 0.36694 | 0.12788 | 0.04593 | 0.09315 | 25.6804 | 0.04 | 0 | Neutral |
| testneg_2385 | -0.05 | -0.29 | 0.810623 | -3 | 1.989667 | -3 | 0.20502 | 0.10447 | 0.06517 | 0.15107 | 30.83789 | 0.080818713 | 0 | Neutral |
| testneg_2386 | -0.03 | 1.03 | 3.857181 | -3 | 0.275531 | -3 | 0.37855 | 0.47307 | 0.06067 | 0.07796 | 16.98432 | 0.13 | 0 | Neutral |
| testneg_2387 | -0.08 | -0.03 | -0.125177 | -3 | 1.150826 | -3 | 0.27308 | 0.12788 | 0.12951 | 0.08688 | 15.17332 | 0.01 | 0 | Neutral |
| testneg_2388 | 0 | 0 | 3.506817 | -3 | 1.075372 | -3 | 0.23088 | 0.21602 | 0.05716 | 0.12491 | 37.55753 | 0.07 | 0 | Neutral |
| testneg_2389 | 0 | 0 | 0.713568 | -3 | -0.154534 | -3 | 0.11072 | 0.38259 | 0.34946 | 0.19092 | 17.53943 | 0.13340513 | 0 | Neutral |
| testneg_2390 | 0.14 | -0.69 | 1.524427 | -3 | 2.091718 | -3 | 0.99317 | 0.60884 | 0.34946 | 0.19092 | 10 | 0.21 | 0 | Neutral |
| testneg_2391 | 0 | -0.01 | 0.594548 | -3 | 2.598788 | -3 | 0.21928 | 0.22085 | 0.19898 | 0.10307 | 19.22222 | 0.0425 | 0 | Neutral |
| testneg_2392 | 0 | 0 | -0.027704 | -3 | 3.364563 | -3 | 0.45299 | 0.16306 | 0.07258 | 0.06248 | 20.26596 | 0.1 | 0 | Neutral |
| testneg_2393 | 0 | 0 | -0.32829 | -3 | 3.085122 | -3 | 0.18509 | 0.12831 | 0.17974 | 0.12788 | 13.85334 | 0.03033908 | 0 | Neutral |
| testneg_2394 | 0 | 0 | 1.472855 | -3 | 1.606239 | 1.606239 | 0.56554 | 0.76978 | 0.89002 | 0.12788 | 10 | 0.05 | 0 | Neutral |
| testneg_2395 | 0 | 0 | 0.099679 | -3 | 4.037788 | -3 | 0.99448 | 0.67288 | 0.08213 | 0.08913 | 11.25 | 0.1201809 | 0 | Neutral |
| testneg_2396 | 0.19 | 0.51 | 3.140578 | -3 | 0.466937 | -3 | 0.14273 | 0.1162 | 0.07015 | 0.0889 | 12.98885 | 0.04 | 0 | Neutral |
| testneg_2397 | -0.12 | 1.14 | 2.5207 | -0.816186 | 4.985142 | -3 | 0.03788 | 0.10996 | 0.21596 | 0.08969 | 18.88889 | 0.240268574 | 0 | Neutral |
| testneg_2398 | 0 | 0 | 0.386087 | 0.386087 | 1.575308 | -3 | 0.26927 | 0.10752 | 0.20655 | 0.15317 | 11.79445 | 0.051671983 | 0 | Neutral |
| testneg_2399 | 0 | 0 | 0.930668 | -3 | 2.718335 | -3 | 0.3793 | 0.24105 | 0.10042 | 0.09296 | 19.11189 | 0.050537613 | 0 | Neutral |
| testneg_2400 | -0.08 | -0.03 | 3.657735 | -3 | -0.09322 | -3 | 0.24528 | 0.37575 | 0.14341 | 0.10134 | 20.14484 | 0.1 | 0 | Neutral |
| testneg_2401 | 0.22 | -2.66 | 2.537219 | -3 | 6.095504 | -3 | 0.0328 | 0.0691 | 0.78211 | 0.35815 | 16.84579 | 0.41 | 19.2 | Neutral |
| testneg_2402 | -0.05 | 0.81 | 2.250533 | -3 | 5.338361 | -3 | 0.14021 | 0.43531 | 0.21587 | 0.11701 | 16.8018 | 0.311148841 | 2 | Neutral |
| testneg_2403 | 0.05 | -0.81 | 5.496455 | -3 | 2.558076 | -3 | 0.17301 | 0.10048 | 0.12921 | 0.11605 | 20.27778 | 0.16996368 | 0.2 | Neutral |
| testneg_2404 | 0 | -0.01 | 0.834318 | -3 | -2.781194 | -3 | 0.31726 | 0.17249 | 0.16374 | 0.14659 | 11.00058 | 0.06 | 0 | Neutral |
| testneg_2405 | 0.07 | 0.33 | 3.297157 | -3 | 0.124643 | -3 | 0.39553 | 0.11991 | 0.10042 | 0.09296 | 31.19662 | 0.021049673 | 0 | Neutral |
| testneg_2406 | -0.03 | 0.85 | 5.601904 | -3 | -0.465282 | -3 | 0.16728 | 0.11868 | 0.42872 | 0.12788 | 21.417 | 0.15 | 0 | Neutral |
| testneg_2407 | 0.2 | -1.26 | 3.848136 | -3 | -1.975003 | -1.975003 | 0.0811 | 0.09449 | 0.08465 | 0.15428 | 10.83333 | 0.219502315 | 0 | Neutral |
| testneg_2408 | -0.03 | 1.03 | 3.06742 | -3 | 5.537026 | -3 | 0.87835 | 0.13522 | 0.1856 | 0.12788 | 13.86082 | 0.32 | 5.4 | Neutral |
| testneg_2409 | -0.06 | 0.57 | 4.072619 | -3 | 0.849086 | -3 | 0.16388 | 0.14439 | 0.30241 | 0.104 | 18.88889 | 0.061742424 | 0 | Neutral |
| testneg_2410 | 0.02 | 0.45 | 2.102442 | -3 | 1.064707 | -3 | 0.51675 | 0.11055 | 0.27637 | 0.0872 | 18.37773 | 0.010175237 | 0 | Neutral |
| testneg_2411 | 0.02 | -0.28 | 0.430481 | 0.430481 | 0.706599 | -3 | 0.31621 | 0.12788 | 0.1077 | 0.12788 | 18.33333 | 0.02 | 0 | Neutral |
| testneg_2412 | -0.09 | 0.25 | 3.307948 | -3 | 6.830695 | -3 | 0.06789 | 0.09707 | 0.53016 | 0.10373 | 13.75 | 0.380277778 | 12.6 | Neutral |
| testneg_2413 | 0 | 0 | 0.094005 | -3 | 5.262041 | -3 | 0.64302 | 0.11262 | 0.3667 | 0.2135 | 18.62567 | 0.205332418 | 0 | Neutral |
| testneg_2414 | 0 | 0 | 0.569405 | -3 | 0.080284 | -3 | 0.47544 | 0.17658 | 0.63818 | 0.12788 | 10.41667 | 0.010540936 | 0 | Neutral |
| testneg_2415 | 0 | 0 | 0.18237 | -3 | 4.759697 | -3 | 0.60251 | 0.39112 | 0.19898 | 0.10335 | 10 | 0.075521096 | 0 | Neutral |
| testneg_2416 | 0.03 | -1.03 | 0.171344 | -3 | -0.370487 | -3 | 0.27187 | 0.41985 | 0.61872 | 0.10479 | 10.83333 | 0.053395062 | 0 | Neutral |
| testneg_2417 | 0.04 | 2.01 | 2.746502 | -3 | 0.329621 | 0.329621 | 0.92237 | 0.11252 | 0.4129 | 0.11172 | 23.40643 | 0.09 | 0 | Neutral |
| testneg_2418 | 0.02 | 0.45 | 3.221373 | -3 | 1.139548 | -3 | 0.16781 | 0.12788 | 0.92262 | 0.12859 | 11.875 | 0.04012549 | 0 | Neutral |
| testneg_2419 | 0 | 0 | -0.387684 | -3 | 0.584676 | -3 | 0.25179 | 0.09355 | 0.31701 | 0.09986 | 17.21847 | 0.000277831 | 0 | Neutral |
| testneg_2420 | 0.19 | 0.51 | 1.770771 | -3 | 0.243635 | -3 | 0.71406 | 0.2012 | 0.47274 | 0.11928 | 5 | 0.07 | 0 | Neutral |
| testneg_2421 | 0.04 | -0.23 | 6.426345 | -3 | 2.798182 | 2.798182 | 0.15119 | 0.08823 | 0.31043 | 0.09688 | 18.30413 | 0.32 | 3.8 | Neutral |
| testneg_2422 | 0 | 0 | 2.055296 | -3 | -0.416655 | -0.416655 | 0.15092 | 0.09074 | 0.05453 | 0.12788 | 22.95764 | 0.02 | 0 | Neutral |
| testneg_2423 | 0 | -0.32 | -0.748689 | -3 | 6.011632 | -3 | 0.07007 | 0.12788 | 0.16968 | 0.12788 | 18.17804 | 0.14 | 0 | Neutral |
| testneg_2424 | -0.04 | -2.01 | 3.282272 | -3 | -0.010919 | -3 | 0.29356 | 0.47581 | 0.02059 | 0.066 | 29.16667 | 0.13 | 0 | Neutral |
| testneg_2425 | 0.21 | 0.15 | 1.704228 | -3 | 2.172359 | -3 | 0.27062 | 0.12378 | 0.15989 | 0.12788 | 18.6768 | 0.035 | 0 | Neutral |
| testneg_2426 | 0.18 | 0.16 | -0.109518 | -3 | 1.223596 | -3 | 0.29804 | 0.12768 | 0.24519 | 0.12788 | 22.36268 | 0.02 | 0 | Neutral |
| testneg_2427 | 0.23 | -1.86 | 0.810517 | -3 | 3.743601 | -3 | 0.08729 | 0.10515 | 0.04694 | 0.44682 | 20.98109 | 0.363149351 | 6.2 | Neutral |
| testneg_2428 | 0 | 0 | 1.856947 | -3 | 7.336513 | -3 | 0.56862 | 0.12788 | 0.45342 | 0.14389 | 20.85927 | 0.368916126 | 12.6 | Neutral |
| testneg_2429 | -0.01 | -0.8 | -0.878612 | -3 | 4.336137 | -3 | 0.11802 | 0.09304 | 0.19898 | 0.12788 | 31.83575 | 0.070588745 | 0 | Neutral |
| testneg_2430 | 0.14 | -1.01 | 2.081126 | -3 | 2.980687 | -3 | 0.2663 | 0.32806 | 0.74558 | 0.10643 | 7.06104 | 0.142047014 | 0 | Neutral |
| testneg_2431 | 0.16 | -0.37 | 2.648188 | -0.060441 | 2.97112 | -3 | 0.64701 | 0.94571 | 0.5901 | 0.19444 | 10.41667 | 0.473360507 | 43 | Neutral |
| testneg_2432 | -0.04 | -2.01 | 0.637803 | -3 | 5.687852 | -3 | 0.29357 | 0.79524 | 0.36709 | 0.19019 | 29.01158 | 0.562512481 | 79.8 | Disease_causing |
| testneg_2433 | 0.14 | -1.01 | -0.270969 | -3 | 3.619507 | -3 | 0.04441 | 0.13603 | 0.67676 | 0.12788 | 11.50811 | 0.141311712 | 0 | Neutral |
| testneg_2434 | 0 | 0 | 2.165106 | -3 | 1.372356 | -3 | 0.30257 | 0.17298 | 0.60026 | 0.35923 | 12.67869 | 0.089873434 | 0 | Neutral |
| testneg_2435 | 0.00172727 | -0.1280909 | 1.232154 | -3 | 3.446637 | -3 | 0.17768 | 0.10832 | 0.13413 | 0.12788 | 18.46051 | 0.03 | 0 | Neutral |
| testneg_2436 | -0.02 | 0.36 | 0.523312 | -3 | -0.959754 | -3 | 0.41987 | 0.67373 | 0.17222 | 0.12788 | 10 | 0.05 | 0 | Neutral |
| testneg_2437 | 0.07 | 0.7 | 5.847469 | -3 | 3.965752 | -3 | 0.28341 | 0.1005 | 0.30471 | 0.11609 | 25.37072 | 0.31 | 4.4 | Neutral |
| testneg_2438 | 0 | 0 | -0.22828 | -3 | 2.17165 | -3 | 0.31694 | 0.13475 | 0.72501 | 0.7625 | 12.29167 | 0.07 | 0 | Neutral |
| testneg_2439 | 0.02 | 0.45 | 4.020676 | -3 | 1.828834 | -3 | 0.44067 | 0.11201 | 0.32016 | 0.11638 | 12.46106 | 0.050082584 | 0 | Neutral |

| | | | | | | | | | | | | | | |
|---|---|---|---|---|---|---|---|---|---|---|---|---|---|---|
| testneg_2440 | 0.15 | -0.28 | 1.351643 | -3 | 1.345014 | -3 | 0.86306 | 0.10962 | 0.3437 | 0.5169 | 5 | 0.081078967 | 0 | Neutral |
| testneg_2441 | 0 | 0 | -0.590049 | -3 | 1.697009 | -3 | 0.50063 | 0.154 | 0.52771 | 0.10974 | 8.61621 | 0.011 | 0 | Neutral |
| testneg_2442 | 0.14 | -1.01 | 0.207897 | -3 | 1.414521 | 0.04439 | 0.20616 | 0.29531 | 0.40081 | 0.15418 | 10.83333 | 0.169353355 | 0 | Neutral |
| testneg_2443 | 0.04 | -0.23 | -0.122196 | -0.122196 | 0.525606 | -3 | 0.31961 | 0.17571 | 0.08401 | 0.16123 | 10 | 0.118757937 | 0 | Neutral |
| testneg_2444 | 0.06 | -0.57 | 0.297516 | 0.297516 | -1.29721 | -3 | 0.19294 | 0.10464 | 0.23385 | 0.12788 | 12.79101 | 0.01 | 0 | Neutral |
| testneg_2445 | 0 | 0 | -0.353049 | -3 | 1.448132 | -3 | 0.9874 | 0.15588 | 0.694 | 0.09974 | 13.9025 | 0.03 | 0 | Neutral |
| testneg_2446 | -0.19 | 0.27 | -1.643119 | -3 | 3.320732 | 3.320732 | 0.1535 | 0.12382 | 0.19898 | 0.13249 | 39.62462 | 0.35 | 6 | Neutral |
| testneg_2447 | -0.01 | 0.57 | -2.581564 | -3 | -2.188525 | -2.188525 | 0.19898 | 0.12788 | 0.07282 | 0.12788 | 18.23515 | 0.03 | 0 | Neutral |
| testneg_2448 | 0 | 0 | 1.609578 | -3 | -0.750318 | -3 | 0.19898 | 0.07094 | 0.26068 | 0.46457 | 39.23611 | 0.058238233 | 0 | Neutral |
| testneg_2449 | 0.21 | 0.15 | 4.012434 | -3 | 4.098385 | -3 | 0.28629 | 0.77234 | 0.09531 | 0.11449 | 10.41667 | 0.355619963 | 7.8 | Neutral |
| testneg_2450 | 0 | 0.73 | 5.650938 | -3 | 2.920916 | -3 | 0.16671 | 0.18311 | 0.19898 | 0.12788 | 34.53102 | 0.366045918 | 13.4 | Neutral |
| testneg_2451 | 0 | 0 | -0.725688 | -3 | 0.96317 | -3 | 0.17285 | 0.09314 | 0.20379 | 0.35591 | 4.72222 | 0.040833333 | 0 | Neutral |
| testneg_2452 | -0.02 | 0.68 | 4.387799 | -3 | 2.642734 | -3 | 0.04555 | 0.06327 | 0.19898 | 0.12788 | 30.96609 | 0.149263378 | 0 | Neutral |
| testneg_2453 | 0.22 | -2.66 | 4.829373 | -3 | 1.360454 | -3 | 0.42306 | 0.86891 | 0.64914 | 0.12481 | 4.72222 | 0.26 | 0.6 | Neutral |
| testneg_2454 | 0.07 | 0.38 | 2.241132 | -3 | -1.359951 | -3 | 0.10133 | 0.32155 | 0.47816 | 0.66526 | 10.83333 | 0.226222474 | 0 | Neutral |
| testneg_2455 | 0.14 | -1.01 | 2.672539 | -3 | 1.259487 | -3 | 0.46371 | 0.11637 | 0.31648 | 0.12788 | 23.12507 | 0.03 | 0 | Neutral |
| testneg_2456 | -0.05 | -0.29 | 0.23755 | -3 | 1.182522 | -3 | 0.55193 | 0.22384 | 0.38002 | 0.22508 | 6.20629 | 0.08735208 | 0 | Neutral |
| testneg_2457 | 0 | 0 | -0.170831 | -3 | 6.245625 | -3 | 0.07044 | 0.11679 | 0.05535 | 0.12788 | 20 | 0.129327922 | 0 | Neutral |
| testneg_2458 | 0.14 | -1.01 | 4.957111 | -3 | 0.099246 | -3 | 0.07748 | 0.11591 | 0.58124 | 0.31208 | 11.89315 | 0.260255925 | 1 | Neutral |
| testneg_2459 | 0.21 | 0.15 | 4.361375 | -3 | 5.306961 | -3 | 0.10459 | 0.12783 | 0.1201 | 0.1062 | 18.6768 | 0.257161111 | 1.2 | Neutral |
| testneg_2460 | 0 | 0 | -0.203227 | -3 | 0.061858 | -3 | 0.58827 | 0.57942 | 0.28474 | 0.12788 | 18.44266 | 0.04 | 0 | Neutral |
| testneg_2461 | -0.02 | 0.68 | 0.243834 | -3 | 1.743396 | -3 | 0.58545 | 0.14414 | 0.11897 | 0.3919 | 19.31102 | 0.14244997 | 0 | Neutral |
| testneg_2462 | -0.05 | -0.29 | 0.790139 | -3 | 0.496899 | 0.29615 | 0.26482 | 0.12788 | 0.17325 | 0.17964 | 6.29982 | 0.06 | 0 | Neutral |
| testneg_2463 | 0.03 | -0.85 | 4.652483 | -3 | 3.776689 | -3 | 0.06841 | 0.10927 | 0.17818 | 0.15728 | 18.30413 | 0.462568337 | 40.2 | Neutral |
| testneg_2464 | 0 | -0.01 | 2.038329 | -3 | 0.4795 | -3 | 0.25159 | 0.12788 | 0.0935 | 0.0819 | 30.85405 | 0.01 | 0 | Neutral |
| testneg_2465 | -0.03 | -0.1 | 1.538056 | -3 | 0.870772 | 0.870772 | 0.53172 | 0.12788 | 0.10569 | 0.09059 | 13.46801 | 0.02 | 0 | Neutral |
| testneg_2466 | 0.21 | 0.15 | 1.631249 | -3 | 4.508189 | -3 | 0.47166 | 0.13548 | 0.0435 | 0.14527 | 21.38889 | 0.39736245 | 14.4 | Neutral |
| testneg_2467 | 0 | 0 | 1.099435 | -3 | 4.311522 | -3 | 0.21383 | 0.11171 | 0.08857 | 0.83499 | 12.76087 | 0.260668581 | 1 | Neutral |
| testneg_2468 | 0 | 0 | 0.839823 | -3 | 2.454179 | -3 | 0.26997 | 0.63944 | 0.11788 | 0.12788 | 18.33333 | 0.075900538 | 0 | Neutral |
| testneg_2469 | 0.14 | -1.85 | 2.670295 | -3 | 5.969277 | -3 | 0.09807 | 0.24901 | 0.31067 | 0.35414 | 10.41667 | 0.799619918 | 100 | Disease_causing |
| testneg_2470 | 0.14 | -1.01 | 0.341374 | -3 | -0.295414 | -3 | 0.10646 | 0.1069 | 0.09741 | 0.11013 | 21.261 | 0.02 | 0 | Neutral |
| testneg_2471 | 0 | 0 | 0.302733 | -3 | -1.64259 | -3 | 0.19898 | 0.12788 | 0.19898 | 0.12788 | 38.64057 | 0.000199966 | 0 | Neutral |
| testneg_2472 | 0 | 0 | 0.894121 | -3 | 0.599729 | -3 | 0.02146 | 0.10351 | 0.38762 | 0.12788 | 22.54725 | 0.03 | 0 | Neutral |
| testneg_2473 | 0.04 | -0.23 | 4.771654 | -3 | -1.184577 | -3 | 0.6201 | 0.12788 | 0.34826 | 0.49454 | 10.41667 | 0.24 | 0.6 | Neutral |
| testneg_2474 | 0 | -0.01 | 0.981481 | -3 | 5.728641 | -3 | 0.24516 | 0.11809 | 0.27806 | 0.09713 | 20.71807 | 0.059892027 | 0 | Neutral |
| testneg_2475 | 0 | 0 | 1.179541 | -3 | 1.796297 | -3 | 0.25827 | 0.21386 | 0.10566 | 0.12788 | 18.02082 | 0.020190632 | 0 | Neutral |
| testneg_2476 | 0.21 | 0.15 | 0.770086 | -3 | 3.15847 | -3 | 0.20527 | 0.10053 | 0.19898 | 0.12788 | 17.21847 | 0.09 | 0 | Neutral |
| testneg_2477 | 0.09 | -0.3 | 6.573032 | -3 | 4.213187 | -3 | 0.48906 | 0.11262 | 0.08529 | 0.12376 | 36.4755 | 0.41 | 21.6 | Neutral |
| testneg_2478 | 0.01 | 0.79 | 3.838754 | -3 | -1.752013 | -3 | 0.18081 | 0.1258 | 0.0609 | 0.47188 | 18.88889 | 0.20875 | 0 | Neutral |
| testneg_2479 | 0 | 0 | 0.153038 | -3 | 4.115116 | -3 | 0.92659 | 0.13588 | 0.09532 | 0.35459 | 23.31464 | 0.38259974 | 7.4 | Neutral |
| testneg_2480 | 0.15 | -0.28 | 0.118932 | 0.118932 | -0.432565 | -3 | 0.19567 | 0.09606 | 0.1105 | 0.44945 | 15.37063 | 0.120912879 | 0 | Neutral |
| testneg_2481 | 0.02 | -0.63 | 2.326926 | -3 | 2.422379 | 2.422379 | 0.07558 | 0.12788 | 0.2485 | 0.10364 | 18.33333 | 0.08 | 0 | Neutral |
| testneg_2482 | 0.21 | 0.15 | 1.478329 | -3 | 3.26904 | -3 | 0.27614 | 0.12085 | 0.18554 | 0.12299 | 19.9225 | 0.081394231 | 0 | Neutral |
| testneg_2483 | 0.18 | -1.05 | 7.147075 | -3 | 1.509728 | -1.705186 | 0.25989 | 0.11651 | 0.0435 | 0.14527 | 25.55556 | 0.325535714 | 4.8 | Neutral |
| testneg_2484 | -0.17 | 1.22 | -2.338688 | -2.338688 | 0.72503 | -3 | 0.33129 | 0.09957 | 0.56862 | 0.12788 | 21.49912 | 0.12 | 0 | Neutral |
| testneg_2485 | 0 | 0 | 1.922434 | -3 | -0.745568 | -3 | 0.59342 | 0.12788 | 0.14349 | 0.27976 | 13.75154 | 0.041520053 | 0 | Neutral |
| testneg_2486 | 0 | 0 | 0.040441 | -3 | 7.532012 | -3 | 0.54777 | 0.25991 | 0.99704 | 0.17463 | 10 | 0.47369306 | 44.4 | Neutral |
| testneg_2487 | 0 | 0 | 0.619175 | 0.619175 | -1.646194 | -3 | 0.68138 | 0.20944 | 0.10362 | 0.09649 | 17.77778 | 0.04 | 0 | Neutral |
| testneg_2488 | 0 | 0 | 0.777519 | -3 | 1.326802 | -3 | 0.75071 | 0.63704 | 0.05183 | 0.18779 | 17.40099 | 0.113569171 | 0 | Neutral |
| testneg_2489 | 0 | 0 | 0.11979 | -3 | 0.964928 | -3 | 0.41647 | 0.11239 | 0.14403 | 0.30507 | 13.51351 | 0.030100943 | 0 | Neutral |
| testneg_2490 | 0.02 | -0.63 | 0.686794 | 0.665096 | 0.823839 | -3 | 0.15132 | 0.11724 | 0.13031 | 0.85678 | 33.68056 | 0.17 | 0 | Neutral |
| testneg_2491 | 0.01 | 0.8 | 6.842888 | -3 | 3.738347 | -3 | 0.19898 | 0.12788 | 0.50388 | 0.12295 | 21.80212 | 0.447831317 | 35.8 | Neutral |
| testneg_2492 | -0.07 | -0.38 | -1.183408 | -3 | 3.787163 | -3 | 0.9862 | 0.4528 | 0.07675 | 0.11464 | 16.42913 | 0.15 | 0 | Neutral |
| testneg_2493 | 0.04 | -0.23 | 2.909519 | -3 | 1.540719 | -3 | 0.2098 | 0.44672 | 0.13271 | 0.22958 | 24.52899 | 0.404774351 | 18.4 | Neutral |
| testneg_2494 | 0 | 0 | 0.889458 | 0.889458 | -1.428127 | -3 | 0.48937 | 0.22085 | 0.11564 | 0.1184 | 21.38889 | 0.05 | 0 | Neutral |
| testneg_2495 | 0 | 0 | 1.064634 | -3 | 2.100416 | -3 | 0.25731 | 0.12662 | 0.48298 | 0.1128 | 22.91417 | 0.000741834 | 0 | Neutral |
| testneg_2496 | 0.08 | 0.03 | 2.539499 | -3 | 7.435597 | -3 | 0.19898 | 0.10894 | 0.53996 | 0.14577 | 17.21847 | 0.437760638 | 30.2 | Neutral |
| testneg_2497 | -0.08 | -0.03 | 2.333739 | -3 | 5.53954 | 0.017414 | 0.11564 | 0.1184 | 0.19898 | 0.09927 | 18.94416 | 0.162792208 | 0 | Neutral |
| testneg_2498 | -0.02 | 0.68 | 1.913159 | -3 | 1.299176 | -3 | 0.91921 | 0.32331 | 0.28199 | 0.10688 | 20.83333 | 0.050934783 | 0 | Neutral |
| testneg_2499 | 0.07 | 0.38 | -2.600286 | -3 | 4.13176 | -3 | 0.03137 | 0.12788 | 0.226 | 0.28784 | 42.5731 | 0.302685185 | 2 | Neutral |
| testneg_2500 | -0.14 | 0.37 | 3.004861 | -3 | 3.074697 | -3 | 0.10486 | 0.08083 | 0.37282 | 0.07943 | 32.12262 | 0.14 | 0 | Neutral |
| testneg_2501 | 0.01 | 0.8 | 1.447055 | -3 | 0.337984 | -3 | 0.60959 | 0.76078 | 0.68004 | 0.12345 | 10.41667 | 0.082087369 | 0 | Neutral |
| testneg_2502 | 0 | 0 | 0.956772 | -3 | 5.172967 | -3 | 0.19898 | 0.94603 | 0.19898 | 0.09927 | 14.75597 | 0.158097211 | 0 | Neutral |
| testneg_2503 | 0.09 | -0.3 | -0.408308 | -3 | 1.351912 | -3 | 0.28165 | 0.15912 | 0.18584 | 0.11114 | 19.67873 | 0.03020999 | 0 | Neutral |
| testneg_2504 | -0.15 | -0.04 | -0.451132 | -3 | 2.395534 | -3 | 0.12048 | 0.2381 | 0.13967 | 0.15696 | 21.6335 | 0.35 | 6 | Neutral |
| testneg_2505 | -0.03 | 0.85 | 1.432399 | -3 | 5.736168 | -3 | 0.93284 | 0.71489 | 0.33245 | 0.45921 | 18.33333 | 0.479015351 | 47 | Neutral |
| testneg_2506 | 0 | -0.01 | 0.684698 | -3 | 6.045853 | -3 | 0.09661 | 0.09981 | 0.07467 | 0.09224 | 13.51351 | 0.1 | 0 | Neutral |
| testneg_2507 | -0.15 | 0.28 | -0.668638 | -0.668638 | -0.801037 | -3 | 0.37131 | 0.10994 | 0.2049 | 0.12678 | 18.76882 | 0.040202206 | 0 | Neutral |
| testneg_2508 | 0 | 0 | 1.264197 | -3 | -0.483476 | -3 | 0.37542 | 0.19444 | 0.16791 | 0.22163 | 12.70833 | 0.05015625 | 0 | Neutral |
| testneg_2509 | 0.02 | -0.36 | -1.684874 | -3 | 4.644088 | -3 | 0.16929 | 0.12788 | 0.07999 | 0.10413 | 14.97514 | 0.074007937 | 0 | Neutral |
| testneg_2510 | 0.07 | 0.38 | 1.305054 | -3 | 1.043597 | -3 | 0.16248 | 0.14352 | 0.19898 | 0.1005 | 13.13152 | 0.01 | 0 | Neutral |
| testneg_2511 | 0.03 | 0.1 | -0.756067 | -3 | 3.484642 | -3 | 0.58124 | 0.31208 | 0.10485 | 0.09565 | 17.21847 | 0.152369048 | 0 | Neutral |
| testneg_2512 | -0.04 | 0.23 | -1.151035 | -1.151035 | 6.702655 | -3 | 0.36225 | 0.1054 | 0.21972 | 0.12788 | 31.2074 | 0.2 | 0.2 | Neutral |
| testneg_2513 | 0.17 | 0.38 | 3.17007 | -3 | 6.245625 | -3 | 0.12377 | 0.12788 | 0.05535 | 0.12788 | 42.85714 | 0.268833333 | 1.4 | Neutral |
| testneg_2514 | -0.15 | 0.82 | 2.970884 | -3 | 3.597707 | -3 | 0.29362 | 0.22177 | 0.99543 | 0.12788 | 7.39391 | 0.394734638 | 14.4 | Neutral |
| testneg_2515 | 0 | 0 | -0.4932 | -3 | -0.771963 | -3 | 0.20379 | 0.43705 | 0.14668 | 0.17962 | 8.53137 | 0.091603175 | 0 | Neutral |
| testneg_2516 | 0.03 | 1.22 | 0.949523 | 0.949523 | 2.23696 | -3 | 0.10056 | 0.12608 | 0.68865 | 0.19414 | 20.77357 | 0.190073529 | 0 | Neutral |
| testneg_2517 | 0 | 0 | 0.844548 | -3 | 0.643368 | -3 | 0.9237 | 0.55859 | 0.95071 | 0.25454 | 4.72222 | 0.040043103 | 0 | Neutral |
| testneg_2518 | 0.05 | 0.29 | -0.981122 | -3 | -0.247306 | -3 | 0.07604 | 0.08642 | 0.35945 | 0.12025 | 21.38889 | 0.02 | 0 | Neutral |
| testneg_2519 | -0.15 | -0.04 | 0.991883 | -3 | 5.405576 | -3 | 0.26816 | 0.13347 | 0.52765 | 0.20417 | 20.0118 | 0.481940951 | 45.4 | Neutral |
| testneg_2520 | 0.02 | -0.63 | 1.42948 | -3 | -0.2352 | -3 | 0.13475 | 0.12788 | 0.19564 | 0.59275 | 17.18434 | 0.061428571 | 0 | Neutral |
| testneg_2521 | 0.19 | 0.51 | 1.770771 | -3 | 1.500407 | -3 | 0.71406 | 0.2012 | 0.19898 | 0.12788 | 20.27778 | 0.08 | 0 | Neutral |
| testneg_2522 | 0.04 | -0.23 | 7.951887 | -3 | 0.847859 | -3 | 0.82846 | 0.13042 | 0.42797 | 0.17974 | 5.97222 | 0.502003968 | 57.2 | Disease_causing |
| testneg_2523 | -0.03 | -1.22 | 4.38057 | -3 | 4.819528 | -3 | 0.5632 | 0.28442 | 0.1211 | 0.1526 | 10.41667 | 0.732037027 | 99.4 | Disease_causing |
| testneg_2524 | 0.18 | -1.05 | 7.116413 | -3 | 2.029095 | -3 | 0.19052 | 0.12788 | 0.96377 | 0.1167 | 6.94614 | 0.3 | 5 | Neutral |
| testneg_2525 | 0.02 | 0.45 | 1.074016 | -3 | 5.422894 | -3 | 0.4132 | 0.12788 | 0.36709 | 0.19019 | 59.60334 | 0.344210902 | 7 | Neutral |
| testneg_2526 | 0 | 0 | 1.080539 | -3 | 0.111854 | 0.111854 | 0.10285 | 0.11721 | 0.14462 | 0.11239 | 12.70833 | 0.01 | 0 | Neutral |
| testneg_2527 | 0 | -0.01 | 0.952222 | -3 | -0.484253 | -3 | 0.08373 | 0.09191 | 0.12836 | 0.09736 | 17.77778 | 0.01 | 0 | Neutral |
| testneg_2528 | 0 | 0 | -0.328773 | -3 | 2.179755 | -3 | 0.11797 | 0.12788 | 0.93246 | 0.12788 | 18.88889 | 0.02 | 0 | Neutral |
| testneg_2529 | 0.14 | -0.37 | 4.646741 | -3 | -1.160549 | -1.160549 | 0.53447 | 0.10345 | 0.04487 | 0.06561 | 13.51351 | 0.050417391 | 0 | Neutral |
| testneg_2530 | 0.23 | -1.86 | 5.446696 | -3 | -0.598616 | -3 | 0.12259 | 0.1041 | 0.37635 | 0.16791 | 12.35574 | 0.330555556 | 7 | Neutral |
| testneg_2531 | 0 | 0 | 1.687316 | -3 | 2.494353 | -3 | 0.26434 | 0.12788 | 0.91377 | 0.5142 | 10.22898 | 0.06 | 0 | Neutral |
| testneg_2532 | 0.14 | -0.37 | 4.837528 | -3 | -2.148868 | -3 | 0.04161 | 0.12788 | 0.16155 | 0.20638 | 19.09347 | 0.3 | 1.8 | Neutral |
| testneg_2533 | -0.05 | -0.29 | 0.368341 | -3 | -0.39724 | -3 | 0.77847 | 0.10751 | 0.09749 | 0.82924 | 21.94444 | 0.13 | 0 | Neutral |
| testneg_2534 | -0.16 | 1.37 | 0.338162 | 0.338162 | 1.926134 | -3 | 0.58827 | 0.57942 | 0.27783 | 0.62384 | 5 | 0.330257228 | 2.4 | Neutral |
| testneg_2535 | 0.03 | 0.12 | 2.733007 | -3 | 0.189843 | -3 | 0.98442 | 0.44966 | 0.15524 | 0.10607 | 14.18076 | 0.068083333 | 0 | Neutral |
| testneg_2536 | -0.03 | -1.22 | 3.683701 | -3 | 7.854373 | -3 | 0.61008 | 0.17538 | 0.4113 | 0.2303 | 33.07749 | 0.742625253 | 99 | Disease_causing |
| testneg_2537 | -0.01 | -0.79 | 0.34963 | -3 | 2.792362 | -3 | 0.17173 | 0.15363 | 0.1144 | 0.09798 | 13.51351 | 0.066159524 | 0 | Neutral |
| testneg_2538 | 0 | 0 | -0.005535 | -3 | 3.086284 | -3 | 0.98312 | 0.12788 | 0.13099 | 0.17453 | 10.09538 | 0.154869602 | 0 | Neutral |
| testneg_2539 | 0 | -0.32 | -0.157625 | -3 | -2.020063 | -3 | 0.14668 | 0.12788 | 0.12951 | 0.08688 | 13.62573 | 0.005200079 | 0 | Neutral |
| testneg_2540 | 0.18 | -1.05 | 1.792345 | -3 | 1.240568 | -3 | 0.14806 | 0.09574 | 0.63645 | 0.14596 | 20 | 0.07011988 | 0 | Neutral |
| testneg_2541 | -0.06 | 0.57 | 1.272946 | -3 | -0.262132 | -3 | 0.12977 | 0.1128 | 0.11788 | 0.12788 | 12.40248 | 0.02 | 0 | Neutral |
| testneg_2542 | -0.03 | -0.1 | 6.245625 | -3 | 1.084274 | -3 | 0.05535 | 0.12788 | 0.07606 | 0.13082 | 29.38849 | 0.3 | 2.8 | Neutral |
| testneg_2543 | 0.04 | -0.23 | 2.208687 | -3 | 5.648148 | -3 | 0.19898 | 0.12788 | 0.1533 | 0.13168 | 26.80135 | 0.264653955 | 1.8 | Neutral |
| testneg_2544 | 0 | 0 | 0.777519 | -3 | 4.326431 | -3 | 0.75071 | 0.63704 | 0.13195 | 0.57267 | 10 | 0.352690574 | 8.4 | Neutral |
| testneg_2545 | 0.18 | -1.05 | 5.727521 | -3 | -0.641871 | -3 | 0.65674 | 0.10572 | 0.19898 | 0.12788 | 51.16516 | 0.13 | 0 | Neutral |
| testneg_2546 | 0 | 0 | 1.941287 | -3 | 0.179864 | -3 | 0.24977 | 0.11262 | 0.10472 | 0.10014 | 18.72417 | 0.01 | 0 | Neutral |
| testneg_2547 | -0.03 | -0.1 | 2.364483 | -3 | 2.768711 | 2.768711 | 0.16716 | 0.10951 | 0.38324 | 0.25193 | 28.56139 | 0.282693655 | 1.2 | Neutral |
| testneg_2548 | 0.05 | 0.29 | 2.385227 | -3 | 5.703257 | -3 | 0.21265 | 0.1284 | 0.23308 | 0.77395 | 10.83333 | 0.470934045 | 41 | Neutral |
| testneg_2549 | 0 | 0 | -0.262607 | -3 | 5.144395 | -3 | 0.21576 | 0.17241 | 0.12848 | 0.65153 | 19.44444 | 0.472661863 | 44.4 | Neutral |
| testneg_2550 | 0 | 0 | 0.585907 | -3 | 0.552408 | -3 | 0.81212 | 0.14915 | 0.12485 | 0.13806 | 19.44444 | 0.08 | 0 | Neutral |
| testneg_2551 | 0 | 0 | 0.285584 | 0.285584 | 0.036613 | -3 | 0.20527 | 0.10053 | 0.20701 | 0.10345 | 22.89111 | 0.01 | 0 | Neutral |
| testneg_2552 | 0.07 | 0.38 | -1.124893 | -3 | 6.883386 | -3 | 0.07923 | 0.12788 | 0.1594 | 0.22002 | 27.38943 | 0.421833333 | 25 | Neutral |
| testneg_2553 | 0.1 | 1.9 | 0.416226 | -3 | -0.950375 | -3 | 0.32111 | 0.12788 | 0.08184 | 0.10206 | 14.79675 | 0.07 | 0 | Neutral |
| testneg_2554 | -0.02 | 0.68 | 5.284085 | -3 | 0.131816 | -3 | 0.08509 | 0.12788 | 0.05564 | 0.06933 | 18.6768 | 0.13 | 0 | Neutral |
| testneg_2555 | 0 | -0.01 | 0.545018 | -3 | 7.994956 | -3 | 0.11818 | 0.10561 | 0.16485 | 0.13045 | 16.59347 | 0.332354167 | 6.2 | Neutral |
| testneg_2556 | 0 | 0 | 2.049487 | -3 | 3.199571 | -3 | 0.54349 | 0.60885 | 0.30474 | 0.12788 | 16.8018 | 0.112577173 | 0 | Neutral |
| testneg_2557 | 0 | 0 | 1.650287 | -3 | 3.635653 | -3 | 0.32974 | 0.12245 | 0.11501 | 0.13748 | 18.11275 | 0.180628882 | 0 | Neutral |
| testneg_2558 | 0.14 | -0.69 | 4.913042 | -3 | -0.427887 | -3 | 0.28063 | 0.40063 | 0.16974 | 0.80184 | 10.83333 | 0.401115579 | 18.4 | Neutral |
| testneg_2559 | -0.1 | -1.9 | -1.487847 | -3 | -0.440657 | -3 | 0.06728 | 0.20664 | 0.30742 | 0.30362 | 24.60125 | 0.25 | 0 | Neutral |
| testneg_2560 | 0 | 0 | 2.485796 | 2.485796 | 1.295804 | -3 | 0.61011 | 0.09584 | 0.64795 | 0.21581 | 10.7989 | 0.268333333 | 4.8 | Neutral |
| testneg_2561 | 0.19 | 0.51 | -0.093394 | -0.093394 | -1.060552 | -3 | 0.34587 | 0.29822 | 0.2559 | 0.10848 | 21.63869 | 0.07 | 0 | Neutral |

| | | | | | | | | | | | | | |
|---|---|---|---|---|---|---|---|---|---|---|---|---|---|
| testneg_2562 | 0 | 0 | 1.52981 | -3 | -0.245689 | -3 | 0.34001 | 0.12788 | 0.19898 | 0.58778 | 18.9586 | 0.042129505 | 0 Neutral |
| testneg_2563 | 0.18 | 0.16 | 5.156881 | -3 | 2.135175 | -3 | 0.8169 | 0.23301 | 0.47192 | 0.11305 | 19 | 0.224502012 | 0 Neutral |
| testneg_2564 | -0.02 | 0.63 | 3.191277 | -3 | -0.272739 | -3 | 0.37074 | 0.11532 | 0.44594 | 0.13793 | 21.32936 | 0.1 | 0 Neutral |
| testneg_2565 | 0.15 | -0.28 | 0.118932 | 0.118932 | -0.586534 | -0.586534 | 0.19567 | 0.09606 | 0.09439 | 0.1884 | 27.03081 | 0.128511006 | 0 Neutral |
| testneg_2566 | 0.16 | -0.37 | 5.827202 | -3 | 0.590738 | -3 | 0.13275 | 0.12788 | 0.01438 | 0.03448 | 18.1877 | 0.1 | 0 Neutral |
| testneg_2567 | 0.2 | -1.26 | 1.388225 | 0.985845 | 0.586242 | -3 | 0.10517 | 0.09803 | 0.559 | 0.2286 | 15.7187 | 0.163095238 | 0 Neutral |
| testneg_2568 | 0 | 0 | -0.029976 | -3 | 6.382916 | -3 | 0.11731 | 0.06348 | 0.46244 | 0.56748 | 25.218 | 0.335281537 | 8.2 Neutral |
| testneg_2569 | -0.02 | 0.63 | 3.995565 | -3 | 4.448407 | -3 | 0.31773 | 0.10816 | 0.19898 | 0.12788 | 21.40097 | 0.232407037 | 0 Neutral |
| testneg_2570 | 0 | 0 | -0.553742 | -3 | -2.628134 | -3 | 0.1424 | 0.12788 | 0.26971 | 0.08509 | 37.73562 | 0.01 | 0 Neutral |
| testneg_2571 | 0.01 | 0.8 | 3.096454 | -3 | 1.816161 | -3 | 0.31824 | 0.6682 | 0.03898 | 0.09943 | 11.23846 | 0.13 | 0 Neutral |
| testneg_2572 | 0 | 0 | -0.464766 | -3 | 3.548299 | -3 | 0.62197 | 0.12788 | 0.26337 | 0.12771 | 18.19459 | 0.050788515 | 0 Neutral |
| testneg_2573 | 0 | 0 | 1.679142 | -3 | 1.939105 | 1.939105 | 0.56966 | 0.68625 | 0.31732 | 0.27963 | 29.77151 | 0.216997421 | 0.2 Neutral |
| testneg_2574 | 0.00615385 | 0.00013986 | -0.345708 | -3 | 0.432231 | -3 | 0.32677 | 0.12788 | 0.83955 | 0.10333 | 19.97081 | 0.01 | 0 Neutral |
| testneg_2575 | 0 | 0.32 | -0.420859 | -3 | -0.110087 | -3 | 0.0597 | 0.18092 | 0.82126 | 0.17937 | 10 | 0.12 | 0 Neutral |
| testneg_2576 | 0 | -0.01 | 0.169949 | -3 | 2.16064 | -3 | 0.38445 | 0.58278 | 0.16734 | 0.15176 | 10.83333 | 0.162410885 | 0 Neutral |
| testneg_2577 | 0.08 | 0.03 | -0.129461 | -3 | 0.84675 | -3 | 0.18416 | 0.13292 | 0.4358 | 0.1222 | 17.12644 | 0.020327957 | 0 Neutral |
| testneg_2578 | 0.01 | 0.8 | 3.914355 | -3 | 4.597558 | -3 | 0.4113 | 0.2303 | 0.0532 | 0.07746 | 22.05467 | 0.302 | 1.8 Neutral |
| testneg_2579 | 0.07 | 0.38 | 0.708784 | -3 | 2.663275 | -3 | 0.28156 | 0.13431 | 0.24087 | 0.27985 | 18.33333 | 0.181693452 | 0 Neutral |
| testneg_2580 | 0.09 | -2.17 | 5.494114 | -3 | 3.178384 | -3 | 0.1834 | 0.12788 | 0.07923 | 0.08035 | 39.42253 | 0.22 | 0.4 Neutral |
| testneg_2581 | 0.21 | 0.15 | 4.319063 | -3 | 0.718564 | -3 | 0.1647 | 0.12076 | 0.06873 | 0.12788 | 19.55601 | 0.050954545 | 0 Neutral |
| testneg_2582 | 0 | -0.01 | 0.130932 | -3 | 3.089818 | -3 | 0.19898 | 0.12788 | 0.09026 | 0.09447 | 29.45989 | 0.04 | 0 Neutral |
| testneg_2583 | 0 | 0 | 2.194532 | -3 | 1.704751 | -3 | 0.48075 | 0.20313 | 0.29548 | 0.10732 | 20.83333 | 0.020760034 | 0 Neutral |
| testneg_2584 | 0.03 | 0.1 | -1.565469 | -3 | 2.845752 | -3 | 0.16204 | 0.12788 | 0.06708 | 0.09112 | 20.27778 | 0.036503477 | 0 Neutral |
| testneg_2585 | -0.08 | -0.03 | -0.057252 | -3 | 0.545422 | -3 | 0.83176 | 0.23122 | 0.29425 | 0.50236 | 6.94444 | 0.096784557 | 0 Neutral |
| testneg_2586 | 0.04 | -0.23 | 0.407117 | -3 | 0.343798 | -3 | 0.17728 | 0.46704 | 0.13575 | 0.12788 | 16.84579 | 0.04 | 0 Neutral |
| testneg_2587 | -0.08 | -0.03 | 1.880384 | -3 | 5.632523 | -3 | 0.62273 | 0.10215 | 0.17764 | 0.09681 | 28.06527 | 0.12 | 0 Neutral |
| testneg_2588 | -0.01 | 1.44 | 0.390517 | -3 | 9.742117 | -3 | 0.21811 | 0.18489 | 0.22582 | 0.12788 | 19.09347 | 0.50225 | 54.6 Disease_causing |
| testneg_2589 | -0.01 | 1.44 | 0.734765 | -3 | 1.624942 | 1.624942 | 0.07204 | 0.07708 | 0.16208 | 0.10597 | 10.56442 | 0.1 | 0 Neutral |
| testneg_2590 | 0.03 | -1.03 | 7.525909 | -3 | 2.091497 | -3 | 0.10835 | 0.12788 | 0.85887 | 0.1489 | 13.37601 | 0.404541667 | 21.4 Neutral |
| testneg_2591 | 0 | 0 | 1.309085 | -3 | 0.8405 | -0.624038 | 0.64125 | 0.28792 | 0.31651 | 0.12788 | 10.57994 | 0.01 | 0 Neutral |
| testneg_2592 | 0.11 | -1.95 | 6.022895 | -3 | -0.086787 | -3 | 0.09583 | 0.09376 | 0.09281 | 0.13929 | 17.77778 | 0.250226449 | 1.2 Neutral |
| testneg_2593 | 0.02 | -0.63 | 0.686794 | 0.686794 | 4.004011 | -3 | 0.15132 | 0.11724 | 0.37736 | 0.11686 | 21.40097 | 0.11 | 0 Neutral |
| testneg_2594 | 0.03 | 0.1 | 1.701517 | -3 | -0.034419 | -0.034419 | 0.57564 | 0.12788 | 0.62203 | 0.12788 | 11.92982 | 0.010037037 | 0 Neutral |
| testneg_2595 | -0.02 | -0.45 | 5.786078 | -3 | 1.516343 | -3 | 0.42306 | 0.86891 | 0.19898 | 0.12788 | 15.59524 | 0.413287546 | 22.6 Neutral |
| testneg_2596 | 0.03 | -1.03 | 5.396431 | -0.172518 | 3.486581 | -3 | 0.19452 | 0.79779 | 0.04851 | 0.12788 | 9.74181 | 0.471189994 | 43.8 Neutral |
| testneg_2597 | 0 | 0 | 0.770471 | -3 | 3.607231 | -3 | 0.99859 | 0.12788 | 0.05079 | 0.0924 | 30.48258 | 0.11 | 0 Neutral |
| testneg_2598 | 0 | 0 | 0.78752 | -3 | 2.155222 | -3 | 0.39305 | 0.18672 | 0.09806 | 0.12788 | 11.25 | 0.02005618 | 0 Neutral |
| testneg_2599 | 0.05 | -0.81 | -0.215991 | -3 | 7.993415 | -3 | 0.21967 | 0.09467 | 0.17296 | 0.09548 | 27.51545 | 0.2945 | 2 Neutral |
| testneg_2600 | 0.07 | 0.38 | 0.540639 | -3 | 0.659043 | 0.659043 | 0.136 | 0.17348 | 0.03137 | 0.12788 | 43.95264 | 0.044166667 | 0 Neutral |
| testneg_2601 | 0.21 | 0.15 | 2.968197 | -3 | 2.197733 | -3 | 0.13278 | 0.20513 | 0.19969 | 0.1163 | 10.41667 | 0.14 | 0 Neutral |
| testneg_2602 | -0.07 | -0.01 | 0.439914 | -3 | 2.299586 | -3 | 0.09275 | 0.14911 | 0.19898 | 0.12788 | 27.26682 | 0.05 | 0 Neutral |
| testneg_2603 | 0.03 | 0.12 | 0.235146 | -3 | 0.79796 | -3 | 0.16856 | 0.19221 | 0.57466 | 0.12788 | 21.25537 | 0.030345238 | 0 Neutral |
| testneg_2604 | 0 | 0 | 0.18944 | -3 | 3.978626 | -3 | 0.15107 | 0.10457 | 0.65734 | 0.1013 | 26.22294 | 0.0504 | 0 Neutral |
| testneg_2605 | 0 | 0 | 0.82819 | 0.82819 | 0.415281 | -3 | 0.19898 | 0.12788 | 0.1472 | 0.12788 | 18.33333 | 0.03 | 0 Neutral |
| testneg_2606 | 0 | -0.01 | 0.491559 | -3 | 3.894139 | -3 | 0.64319 | 0.12788 | 0.18147 | 0.12788 | 39.33542 | 0.071524582 | 0 Neutral |
| testneg_2607 | 0 | 0 | -0.031197 | -3 | 1.768293 | -3 | 0.1017 | 0.11809 | 0.15016 | 0.08826 | 32.95898 | 0.01 | 0 Neutral |
| testneg_2608 | 0.14 | -0.37 | 5.240194 | -3 | 0.063515 | -3 | 0.19898 | 0.12788 | 0.11773 | 0.48758 | 19.21453 | 0.34025 | 5.8 Neutral |
| testneg_2609 | 0.16 | 0.08 | 1.309209 | -3 | 2.10021 | -3 | 0.84588 | 0.14896 | 0.15526 | 0.08511 | 18.33333 | 0.059742424 | 0 Neutral |
| testneg_2610 | 0.14 | -1.01 | 2.126776 | -3 | 0.127377 | -3 | 0.9541 | 0.96732 | 0.46791 | 0.7316 | 5.349 | 0.220534625 | 0 Neutral |
| testneg_2611 | 0 | 0 | -0.784713 | -0.784713 | 3.89714 | -3 | 0.54899 | 0.31462 | 0.33022 | 0.13818 | 13.29947 | 0.310553988 | 4.2 Neutral |
| testneg_2612 | 0.01254545 | -0.088 | 7.130316 | -3 | 1.196757 | -3 | 0.19898 | 0.12895 | 0.55315 | 0.09195 | 13.51351 | 0.319223485 | 5.6 Neutral |
| testneg_2613 | 0 | 0 | 0.402508 | 0.402508 | -0.913672 | -3 | 0.29794 | 0.11106 | 0.65897 | 0.67602 | 11.875 | 0.07 | 0 Neutral |
| testneg_2614 | -0.01 | -0.79 | 0.026216 | -3 | 1.275987 | 0.217286 | 0.94674 | 0.12413 | 0.10383 | 0.08853 | 13.11828 | 0.04 | 0 Neutral |
| testneg_2615 | 0.05 | -0.81 | 3.374642 | -3 | 6.218059 | -3 | 0.33264 | 0.10493 | 0.2559 | 0.10848 | 23.73089 | 0.21 | 0.2 Neutral |
| testneg_2616 | 0 | 0 | 1.604498 | -3 | 5.567099 | -3 | 0.6992 | 0.1804 | 0.05732 | 0.1098 | 30.70461 | 0.121223291 | 0 Neutral |
| testneg_2617 | 0.15 | -0.28 | 1.144976 | -3 | 1.104931 | 1.104931 | 0.53592 | 0.11293 | 0.06708 | 0.09112 | 20.83333 | 0.030480549 | 0 Neutral |
| testneg_2618 | -0.08 | -0.03 | 1.624358 | -3 | 1.59863 | 1.59863 | 0.16248 | 0.14352 | 0.23401 | 0.12374 | 10 | 0.020645833 | 0 Neutral |
| testneg_2619 | 0 | 0 | 0.034609 | -3 | 0.037859 | -3 | 0.42033 | 0.17821 | 0.19391 | 0.08567 | 16.88889 | 0.010472371 | 0 Neutral |
| testneg_2620 | 0 | -0.01 | 0.238703 | -3 | 0.635023 | -3 | 0.0887 | 0.10047 | 0.12136 | 0.27707 | 59.01488 | 0.17 | 0 Neutral |
| testneg_2621 | 0 | 0 | 1.649085 | -3 | 5.161345 | 0.04439 | 0.15822 | 0.12788 | 0.40081 | 0.15418 | 15.46432 | 0.224703256 | 0.8 Neutral |
| testneg_2622 | -0.03 | -0.1 | 0.271296 | -3 | 4.404174 | -3 | 0.0978 | 0.26338 | 0.09828 | 0.07733 | 17.21847 | 0.19 | 0 Neutral |
| testneg_2623 | 0.16 | 0.08 | 4.995915 | -3 | 0.754237 | -3 | 0.76138 | 0.16258 | 0.13118 | 0.08646 | 13.51351 | 0.11 | 0 Neutral |
| testneg_2624 | 0 | 0 | 1.568837 | -3 | 2.028242 | -3 | 0.10756 | 0.0892 | 0.27058 | 0.13363 | 9.75526 | 0.050417391 | 0 Neutral |
| testneg_2625 | 0 | 0 | 1.078217 | -3 | 1.88634 | -3 | 0.9327 | 0.12788 | 0.06395 | 0.0759 | 11.74281 | 0.01 | 0 Neutral |
| testneg_2626 | 0.01 | -1.44 | 6.204767 | -3 | 3.867182 | -3 | 0.28532 | 0.10663 | 0.16777 | 0.1749 | 16.83616 | 0.67 | 97 Disease_causing |
| testneg_2627 | 0.04 | -0.23 | 7.895169 | -3 | 5.464524 | -3 | 0.30257 | 0.17298 | 0.31507 | 0.61057 | 12.70833 | 0.911962121 | 100 Disease_causing |
| testneg_2628 | -0.03 | -0.12 | 3.715248 | -3 | -1.160549 | -1.160549 | 0.83067 | 0.10583 | 0.04487 | 0.06561 | 13.14653 | 0.04 | 0 Neutral |
| testneg_2629 | 0 | 0 | 0.990948 | -3 | 1.511177 | -3 | 0.31859 | 0.11262 | 0.9193 | 0.14584 | 11.46341 | 0.010234505 | 0 Neutral |
| testneg_2630 | 0.23 | -1.86 | 4.93441 | -3 | 0.565031 | -3 | 0.11564 | 0.09588 | 0.19898 | 0.12788 | 30.74611 | 0.080325397 | 0 Neutral |
| testneg_2631 | -0.07 | -0.38 | -2.304927 | -2.304927 | 1.568815 | -3 | 0.02665 | 0.05082 | 0.14379 | 0.12788 | 12.29167 | 0.050137399 | 0 Neutral |
| testneg_2632 | 0.08 | 0.03 | -0.196592 | -0.196592 | 3.997628 | -3 | 0.32733 | 0.16869 | 0.66799 | 0.12788 | 11.25 | 0.207409824 | 0 Neutral |
| testneg_2633 | 0 | 0 | 1.887991 | -3 | 5.471755 | -3 | 0.18869 | 0.23984 | 0.1784 | 0.12788 | 12.29167 | 0.21167803 | 0.4 Neutral |
| testneg_2634 | -0.15 | 0.28 | 1.579861 | -3 | 4.3659 | -3 | 0.64566 | 0.22552 | 0.12453 | 0.17736 | 10.41667 | 0.486176718 | 48.4 Neutral |
| testneg_2635 | -0.02 | 0.68 | 0.791221 | -3 | 3.635933 | -3 | 0.35422 | 0.11185 | 0.21265 | 0.1284 | 30.22972 | 0.070431561 | 0 Neutral |
| testneg_2636 | 0 | -0.01 | -0.496432 | -3 | 2.71072 | -3 | 0.06247 | 0.14184 | 0.09719 | 0.09576 | 10.85218 | 0.06 | 0 Neutral |
| testneg_2637 | -0.06 | 0.57 | 5.1687 | -3 | 5.17674 | -3 | 0.1909 | 0.14279 | 0.07994 | 0.1101 | 17.94606 | 0.442752874 | 31.2 Neutral |
| testneg_2638 | 0.02 | -0.68 | 4.742104 | -3 | 2.789761 | -3 | 0.21623 | 0.42074 | 0.09099 | 0.2051 | 16.8279 | 0.638923594 | 94.4 Disease_causing |
| testneg_2639 | 0 | 0 | -0.259811 | -3 | 0.516563 | -3 | 0.70964 | 0.12663 | 0.49083 | 0.14252 | 20.95603 | 0.051044735 | 0 Neutral |
| testneg_2640 | 0 | 0 | 0.237747 | -3 | 0.91144 | -3 | 0.11261 | 0.14664 | 0.28817 | 0.10531 | 13.2927 | 0.01 | 0 Neutral |
| testneg_2641 | 0.14 | -0.37 | 6.556357 | -3 | 6.347701 | -3 | 0.12465 | 0.09534 | 0.20379 | 0.35591 | 10.41667 | 0.827544643 | 100 Disease_causing |
| testneg_2642 | 0.19 | -0.27 | 4.230056 | -3 | 2.478986 | -3 | 0.08489 | 0.12788 | 0.13483 | 0.12788 | 17.56226 | 0.09066394 | 0 Neutral |
| testneg_2643 | 0 | 0 | 2.047211 | -3 | 4.231687 | -3 | 0.16823 | 0.10796 | 0.09616 | 0.11651 | 22.54161 | 0.06 | 0 Neutral |
| testneg_2644 | 0 | 0.32 | 2.196162 | -3 | 4.448384 | -3 | 0.12977 | 0.1128 | 0.03646 | 0.09863 | 12.9913 | 0.081787879 | 0 Neutral |
| testneg_2645 | 0 | 0 | 1.433534 | -3 | 3.472891 | -3 | 0.30275 | 0.12788 | 0.10168 | 0.12788 | 21.84397 | 0.053916667 | 0 Neutral |
| testneg_2646 | -0.06 | -0.18 | 5.244235 | -3 | 1.771608 | -3 | 0.96842 | 0.23432 | 0.47887 | 0.09483 | 11.54762 | 0.24 | 0.4 Neutral |
| testneg_2647 | 0.03 | -1.03 | 3.635917 | -3 | 3.445512 | -3 | 0.75978 | 0.11423 | 0.38126 | 0.27764 | 18.75947 | 0.477264744 | 45.2 Neutral |
| testneg_2648 | -0.02 | -0.45 | -1.582634 | -3 | -0.014257 | -3 | 0.05663 | 0.12788 | 0.69666 | 0.11055 | 31.31994 | 0.05 | 0 Neutral |
| testneg_2649 | 0.09 | -0.3 | 7.106089 | -3 | -1.198268 | -3 | 0.14484 | 0.14928 | 0.06187 | 0.36607 | 10 | 0.584 | 84.8 Disease_causing |
| testneg_2650 | 0.11 | -0.19 | -0.373191 | -0.775943 | 3.872922 | -3 | 0.13696 | 0.09566 | 0.1675 | 0.09413 | 21.10568 | 0.08 | 0 Neutral |
| testneg_2651 | 0.04 | -0.23 | 1.084734 | -3 | 0.774689 | 0.774689 | 0.1304 | 0.12788 | 0.24219 | 0.17458 | 12.74476 | 0.050792308 | 0 Neutral |
| testneg_2652 | -0.0029762 | 0.28392857 | 2.182722 | -3 | 4.092031 | -3 | 0.26913 | 0.1087 | 0.50134 | 0.15488 | 51.9464 | 0.29011797 | 1 Neutral |
| testneg_2653 | 0.18 | -1.05 | 7.069037 | -3 | 2.237466 | -3 | 0.08631 | 0.10194 | 0.28085 | 0.13803 | 16.8018 | 0.329599099 | 7.6 Neutral |
| testneg_2654 | 0.02 | 0.41 | 0.799457 | -3 | 2.908555 | -3 | 0.08714 | 0.08945 | 0.14175 | 0.12788 | 19.44444 | 0.04 | 0 Neutral |
| testneg_2655 | 0.03 | -1.03 | 6.011522 | -3 | 2.212242 | -3 | 0.84468 | 0.55945 | 0.26434 | 0.12788 | 20.3131 | 0.466775362 | 41.6 Neutral |
| testneg_2656 | 0.01 | 0.79 | 0.420754 | -3 | 5.365478 | -3 | 0.54113 | 0.11017 | 0.46837 | 0.14896 | 19.44444 | 0.2852 | 1 Neutral |
| testneg_2657 | 0.14 | -1.01 | 2.897476 | -3 | 1.489631 | -3 | 0.15958 | 0.41628 | 0.08579 | 0.6068 | 10.83333 | 0.385427451 | 11.8 Neutral |
| testneg_2658 | 0 | 0 | 2.299451 | -3 | 6.048042 | -3 | 0.24832 | 0.84371 | 0.06958 | 0.16086 | 4.44444 | 0.551040598 | 73.8 Disease_causing |
| testneg_2659 | -0.03 | -0.1 | 0.212913 | -3 | 3.544558 | -3 | 0.11391 | 0.19434 | 0.22896 | 0.24839 | 10.22898 | 0.410669908 | 26.2 Neutral |
| testneg_2660 | 0.03 | -0.85 | 6.267746 | -3 | 4.578145 | -3 | 0.09432 | 0.15749 | 0.1675 | 0.09413 | 20.8805 | 0.4953125 | 53.6 Disease_causing |
| testneg_2661 | 0.01 | 0.8 | 7.536392 | -3 | 3.527525 | -3 | 0.3954 | 0.17605 | 0.22443 | 0.10657 | 10.41667 | 0.708930011 | 98.8 Disease_causing |
| testneg_2662 | 0.02 | 0.45 | 1.21068 | -3 | -0.105078 | -3 | 0.95247 | 0.11285 | 0.0501 | 0.12788 | 22.89355 | 0.030205134 | 0 Neutral |
| testneg_2663 | 0 | 0 | 0.82819 | 0.82819 | 2.147578 | -3 | 0.19898 | 0.12788 | 0.47335 | 0.12788 | 4.72222 | 0.04 | 0 Neutral |
| testneg_2664 | -0.02 | -0.45 | 5.342396 | -3 | 0.823839 | -3 | 0.22169 | 0.11262 | 0.13031 | 0.85678 | 23.9403 | 0.35 | 7.2 Neutral |
| testneg_2665 | 0 | 0 | 0.343354 | -3 | 2.414298 | -3 | 0.10241 | 0.08165 | 0.09426 | 0.1658 | 20 | 0.117746823 | 0 Neutral |
| testneg_2666 | -0.02 | -0.41 | 0.433545 | -3 | -0.347591 | -3 | 0.06996 | 0.35923 | 0.32094 | 0.36254 | 10.83333 | 0.233060606 | 0 Neutral |
| testneg_2667 | 0.14 | -0.37 | 0.198379 | -3 | 2.996324 | -3 | 0.03991 | 0.08342 | 0.2559 | 0.10848 | 20.52758 | 0.07 | 0 Neutral |
| testneg_2668 | 0.02 | 0.45 | 5.906217 | -3 | 0.990803 | -3 | 0.72163 | 0.10596 | 0.10822 | 0.10942 | 15.21341 | 0.188504274 | 0.4 Neutral |
| testneg_2669 | 0 | 0 | 1.397753 | -3 | 4.086754 | 4.086754 | 0.43918 | 0.46357 | 0.07275 | 0.12788 | 26.3228 | 0.336138736 | 3.4 Neutral |
| testneg_2670 | 0.14 | -1.01 | -0.384327 | -3 | 1.962423 | -3 | 0.18507 | 0.12788 | 0.13961 | 0.09565 | 11.875 | 0.02 | 0 Neutral |
| testneg_2671 | -0.06 | 0.57 | 0.313057 | -3 | 5.043324 | -3 | 0.38445 | 0.58278 | 0.3405 | 0.1816 | 10.41667 | 0.52369371 | 60.4 Disease_causing |
| testneg_2672 | 0.21 | 0.15 | 0.824266 | -3 | -0.377332 | -3 | 0.113 | 0.12788 | 0.58587 | 0.09866 | 19.53379 | 0.0275 | 0 Neutral |
| testneg_2673 | 0 | 0 | 1.975201 | -3 | 5.422894 | -3 | 0.34495 | 0.16861 | 0.36709 | 0.19019 | 25.83333 | 0.493163467 | 52.8 Disease_causing |
| testneg_2674 | -0.02 | -0.45 | 4.576491 | -3 | 5.70179 | -3 | 0.09488 | 0.10426 | 0.37909 | 0.4568 | 18.33333 | 0.695858877 | 98.2 Disease_causing |
| testneg_2675 | 0 | 0 | 0.708831 | -3 | -1.781198 | -3 | 0.40803 | 0.47186 | 0.43897 | 0.12441 | 12.16924 | 0.02 | 0 Neutral |
| testneg_2676 | -0.04 | 0.23 | -0.274997 | -3 | 6.246684 | -3 | 0.37501 | 0.0997 | 0.14485 | 0.16945 | 26.9055 | 0.387310529 | 13.4 Neutral |
| testneg_2677 | -0.02 | -0.45 | 1.768293 | -3 | 2.980314 | -3 | 0.15016 | 0.08826 | 0.19898 | 0.535 | 18.33333 | 0.170007684 | 0 Neutral |
| testneg_2678 | 0.05 | -0.81 | 4.217475 | -3 | 2.830718 | -3 | 0.1551 | 0.09276 | 0.11229 | 0.08897 | 19.33702 | 0.100445309 | 0 Neutral |
| testneg_2679 | -0.08 | -0.03 | 0.399863 | -3 | 1.871771 | 1.091668 | 0.2663 | 0.32806 | 0.86373 | 0.11969 | 14.23074 | 0.060313397 | 0 Neutral |
| testneg_2680 | 0 | 0 | 0.917232 | -3 | -0.355202 | -3 | 0.25556 | 0.09342 | 0.21802 | 0.10187 | 16.91148 | 0.00021391 | 0 Neutral |
| testneg_2681 | 0.18 | -1.05 | 7.032758 | -3 | -1.759585 | -3 | 0.07428 | 0.07645 | 0.17764 | 0.09681 | 19.28063 | 0.2 | 0.2 Neutral |
| testneg_2682 | 0.21 | 0.15 | 6.222747 | -3 | 2.208499 | -3 | 0.1602 | 0.11262 | 0.29372 | 0.07609 | 23.81803 | 0.151339286 | 0 Neutral |
| testneg_2683 | 0.15 | -1.06 | 5.926391 | -3 | -1.814506 | -3 | 0.84682 | 0.35105 | 0.05999 | 0.12788 | 12.31231 | 0.2875 | 1.6 Neutral |

| | | | | | | | | | | | | | | |
|---|---|---|---|---|---|---|---|---|---|---|---|---|---|---|
| testneg_2684 | 0.08 | 0.03 | -2.199413 | -3 | 0.2182 | -3 | 0.10604 | 0.10785 | 0.19898 | 0.12788 | 42.41529 | 0.02 | 0 | Neutral |
| testneg_2685 | -0.07 | 0.31 | 0.335227 | -3 | 3.034427 | -3 | 0.04346 | 0.10785 | 0.3298 | 0.21875 | 18.37823 | 0.224576535 | 0.2 | Neutral |
| testneg_2686 | 0.04 | -0.23 | 1.294129 | -3 | 0.272659 | 0.272659 | 0.14498 | 0.33896 | 0.17093 | 0.23281 | 11.875 | 0.186280828 | 0 | Neutral |
| testneg_2687 | -0.06 | -0.18 | 0.975671 | -3 | 2.022528 | -3 | 0.29739 | 0.10893 | 0.57953 | 0.15513 | 14.48696 | 0.05 | 0 | Neutral |
| testneg_2688 | -0.15 | -0.04 | 5.828451 | -3 | 0.086807 | -3 | 0.98312 | 0.12788 | 0.30471 | 0.11609 | 11.51319 | 0.22 | 0 | Neutral |
| testneg_2689 | 0.03 | -1.03 | 2.707374 | -3 | 1.096466 | -3 | 0.33127 | 0.49952 | 0.49745 | 0.1259 | 20.95238 | 0.109821429 | 0 | Neutral |
| testneg_2690 | 0.2 | -1.26 | 3.950466 | -3 | 0.728741 | -3 | 0.12886 | 0.12788 | 0.79857 | 0.12788 | 26.80135 | 0.07 | 0 | Neutral |
| testneg_2691 | 0.1 | 1.9 | 0.416226 | -3 | 0.782368 | -3 | 0.32111 | 0.12788 | 0.31651 | 0.12788 | 13.02635 | 0.08 | 0 | Neutral |
| testneg_2692 | 0 | 0 | 0.87666 | -3 | -0.859115 | -3 | 0.1192 | 0.1508 | 0.19898 | 0.12788 | 29.80592 | 0.02 | 0 | Neutral |
| testneg_2693 | -0.05 | -0.29 | -0.062714 | -3 | 1.420779 | -3 | 0.05323 | 0.32175 | 0.14386 | 0.12788 | 37.5 | 0.126544118 | 0 | Neutral |
| testneg_2694 | 0 | 0 | 0.730301 | -3 | 4.893835 | -3 | 0.40612 | 0.45135 | 0.0778 | 0.0728 | 10.83333 | 0.105775577 | 0 | Neutral |
| testneg_2695 | -0.0069091 | 0.03145455 | 9.808835 | -3 | -1.110257 | -3 | 0.41193 | 0.10133 | 0.15217 | 0.12544 | 10 | 0.405084065 | 17 | Neutral |
| testneg_2696 | 0 | 0 | 0.818736 | -3 | 0.655162 | -3 | 0.28907 | 0.12788 | 0.26756 | 0.15588 | 21.2644 | 0.051090894 | 0 | Neutral |
| testneg_2697 | 0.21 | 0.15 | 1.305251 | -3 | 7.340698 | -3 | 0.06304 | 0.12788 | 0.57188 | 0.34038 | 18.30413 | 0.491648267 | 52.2 | Disease_causing |
| testneg_2698 | 0 | 0 | -0.553742 | -3 | -0.55851 | -3 | 0.1424 | 0.12788 | 0.06874 | 0.12788 | 17.94606 | 0.01 | 0 | Neutral |
| testneg_2699 | 0 | -0.32 | 0.941916 | -3 | 4.834981 | 1.759764 | 0.07963 | 0.23713 | 0.11043 | 0.12788 | 20.27778 | 0.26125 | 0.2 | Neutral |
| testneg_2700 | 0.07 | 0.7 | 4.681701 | -3 | -0.178479 | -3 | 0.10571 | 0.12788 | 0.25893 | 0.10353 | 35.84347 | 0.040158521 | 0 | Neutral |
| testneg_2701 | -0.03 | -0.1 | 0.212913 | -3 | 5.715689 | -3 | 0.11391 | 0.19434 | 0.19898 | 0.12788 | 19.93999 | 0.262081524 | 0.4 | Neutral |
| testneg_2702 | 0.18 | 0.16 | 0.843463 | -3 | -1.457001 | -3 | 0.98312 | 0.12788 | 0.25119 | 0.13626 | 9.1693 | 0.100280303 | 0 | Neutral |
| testneg_2703 | 0.17 | 0.38 | 5.910984 | -3 | 2.431872 | -3 | 0.9355 | 0.40418 | 0.1664 | 0.10113 | 12.29167 | 0.41 | 19 | Neutral |
| testneg_2704 | 0 | 0 | 1.609578 | -3 | 0.839368 | 0.839368 | 0.19898 | 0.07094 | 0.18785 | 0.14723 | 45.19751 | 0.05 | 0 | Neutral |
| testneg_2705 | 0 | 0 | 1.683026 | -3 | 0.791681 | -3 | 0.84106 | 0.63405 | 0.15689 | 0.28307 | 4.72222 | 0.060588235 | 0 | Neutral |
| testneg_2706 | -0.16 | -0.08 | -1.975003 | -1.975003 | 5.102298 | -3 | 0.08465 | 0.15428 | 0.10213 | 0.17266 | 12.29167 | 0.598099206 | 87.4 | Disease_causing |
| testneg_2707 | 0 | -0.32 | -0.680064 | -3 | 0.401809 | -3 | 0.19898 | 0.10512 | 0.99531 | 0.86203 | 11.25 | 0.11 | 0 | Neutral |
| testneg_2708 | 0 | 0 | 0.036455 | -3 | 2.17123 | -3 | 0.15041 | 0.43468 | 0.18597 | 0.12788 | 10.83333 | 0.041471297 | 0 | Neutral |
| testneg_2709 | -0.09 | 0.67 | -0.04628 | -3 | 0.282279 | -3 | 0.19898 | 0.12788 | 0.03526 | 0.06855 | 23.48674 | 0.03 | 0 | Neutral |
| testneg_2710 | -0.21 | -0.15 | -2.734447 | -3 | 1.170813 | -3 | 0.48023 | 0.12788 | 0.63818 | 0.12788 | 20.39683 | 0.07 | 0 | Neutral |
| testneg_2711 | -0.03 | 1.03 | 5.710421 | -3 | 5.643105 | -3 | 0.43736 | 0.10017 | 0.08775 | 0.12788 | 16.88659 | 0.529422414 | 65.2 | Disease_causing |
| testneg_2712 | 0.14 | -0.37 | 0.752578 | -3 | -0.37668 | -3 | 0.19898 | 0.12788 | 0.06187 | 0.36607 | 45.13889 | 0.135833333 | 0 | Neutral |
| testneg_2713 | 0.04 | 2.01 | -0.873402 | -3 | 1.045902 | -3 | 0.10472 | 0.10014 | 0.31339 | 0.10191 | 31.84685 | 0.08 | 0 | Neutral |
| testneg_2714 | 0 | 0 | -0.179321 | -3 | -1.745 | -3 | 0.14101 | 0.14318 | 0.1743 | 0.12788 | 10 | 0.01 | 0 | Neutral |
| testneg_2715 | 0.06 | 0.18 | -1.363655 | -3 | 1.610282 | -3 | 0.13203 | 0.12092 | 0.2663 | 0.32806 | 13.81407 | 0.06 | 0 | Neutral |
| testneg_2716 | 0.09 | -2.17 | 0.870138 | -3 | 2.628269 | -3 | 0.26961 | 0.09979 | 0.18584 | 0.11114 | 21.76143 | 0.06017042 | 0 | Neutral |
| testneg_2717 | -0.01 | 0.57 | 5.88281 | -3 | -2.609079 | -3 | 0.17322 | 0.10311 | 0.19898 | 0.09533 | 35.38622 | 0.156904762 | 0 | Neutral |
| testneg_2718 | -0.07 | -0.38 | 3.896611 | -3 | -1.160549 | -1.160549 | 0.57349 | 0.09764 | 0.04487 | 0.06561 | 16.8018 | 0.05 | 0 | Neutral |
| testneg_2719 | 0.14 | -0.69 | 2.199376 | 2.199376 | 1.021238 | -3 | 0.08489 | 0.12788 | 0.12042 | 0.20759 | 9.1046 | 0.26 | 4.2 | Neutral |
| testneg_2720 | -0.22 | 2.66 | 0.984298 | -3 | 2.310369 | -3 | 0.34406 | 0.09894 | 0.54895 | 0.15082 | 11.875 | 0.260206207 | 0.4 | Neutral |
| testneg_2721 | -0.03 | -0.1 | -0.743536 | -3 | -0.806542 | -3 | 0.13708 | 0.81545 | 0.40116 | 0.43982 | 14.89385 | 0.24 | 0 | Neutral |
| testneg_2722 | 0 | 0 | -0.489156 | -3 | 1.435364 | -3 | 0.05887 | 0.10046 | 0.20357 | 0.12788 | 29.21174 | 0.02 | 0 | Neutral |
| testneg_2723 | 0.03 | 0.1 | 4.629819 | -3 | 0.86962 | -3 | 0.24274 | 0.13034 | 0.15989 | 0.12788 | 13.51351 | 0.041055556 | 0 | Neutral |
| testneg_2724 | -0.04 | 0.89 | 1.989667 | -3 | 2.212596 | -3 | 0.06517 | 0.15107 | 0.17226 | 0.13001 | 20.89414 | 0.114166667 | 0 | Neutral |
| testneg_2725 | -0.18 | -0.16 | 2.72063 | 2.72063 | 5.405576 | -3 | 0.65141 | 0.18046 | 0.52765 | 0.20417 | 11.13761 | 0.746745951 | 99 | Disease_causing |
| testneg_2726 | 0.14 | -0.37 | 2.61912 | -3 | -0.30377 | -3 | 0.30257 | 0.17298 | 0.19416 | 0.0901 | 21.08109 | 0.04 | 0 | Neutral |
| testneg_2727 | 0 | 0 | -0.566651 | -3 | 2.196766 | -3 | 0.38016 | 0.14503 | 0.33122 | 0.12788 | 10 | 0.01 | 0 | Neutral |
| testneg_2728 | -0.08 | -0.03 | 0.919374 | -3 | 6.034304 | -3 | 0.86851 | 0.17715 | 0.16057 | 0.12305 | 16.66667 | 0.201469814 | 0 | Neutral |
| testneg_2729 | -0.07 | -0.38 | 1.812519 | -3 | 2.644462 | -3 | 0.13767 | 0.11503 | 0.10803 | 0.14652 | 4.72222 | 0.113413165 | 0 | Neutral |
| testneg_2730 | -0.08 | -0.03 | 3.582832 | -3 | -2.354628 | -3 | 0.72124 | 0.11262 | 0.06547 | 0.09123 | 8.97898 | 0.04 | 0 | Neutral |
| testneg_2731 | 0.07 | 0.38 | 0.542406 | -3 | -0.466806 | -3 | 0.96341 | 0.18404 | 0.22896 | 0.24839 | 10.83333 | 0.108863155 | 0 | Neutral |
| testneg_2732 | 0.06 | -1.29 | 6.321709 | -3 | 1.940809 | -3 | 0.96879 | 0.38765 | 0.23278 | 0.11027 | 11.62434 | 0.42 | 27.8 | Neutral |
| testneg_2733 | -0.04 | 0.23 | 2.344511 | -3 | 7.238335 | -3 | 0.42585 | 0.18081 | 0.12508 | 0.231 | 12.38369 | 0.71191282 | 99.6 | Disease_causing |
| testneg_2734 | 0.03 | 0.1 | 5.285192 | -3 | 0.275543 | 0.275543 | 0.13325 | 0.12788 | 0.4738 | 0.14744 | 27.51545 | 0.21 | 0.6 | Neutral |
| testneg_2735 | 0 | 0 | -0.060097 | -3 | -1.187437 | -3 | 0.03651 | 0.06141 | 0.44479 | 0.20004 | 20.83333 | 0.11 | 0 | Neutral |
| testneg_2736 | 0 | 0 | 2.010403 | -3 | 5.968963 | -3 | 0.2543 | 0.15588 | 0.33256 | 0.14909 | 10.41667 | 0.439196698 | 32.8 | Neutral |
| testneg_2737 | -0.16 | 0.37 | 5.840704 | -3 | 1.351912 | -3 | 0.37909 | 0.4568 | 0.18584 | 0.11114 | 13.01295 | 0.38 | 13.6 | Neutral |
| testneg_2738 | 0 | 0 | 0.300973 | -3 | 1.222548 | 1.222548 | 0.1589 | 0.17874 | 0.06705 | 0.12788 | 33.37997 | 0.04 | 0 | Neutral |
| testneg_2739 | -0.03 | -0.1 | 1.759623 | -3 | 2.272008 | -3 | 0.16475 | 0.26069 | 0.23913 | 0.12788 | 11.65872 | 0.055076923 | 0 | Neutral |
| testneg_2740 | 0.14 | -1.01 | 3.908744 | -3 | 0.255551 | 0.255551 | 0.47379 | 0.18145 | 0.03553 | 0.12788 | 35.46155 | 0.09 | 0 | Neutral |
| testneg_2741 | 0.17 | 0.38 | 3.218072 | -3 | 1.247019 | -3 | 0.9637 | 0.12788 | 0.88456 | 0.11674 | 10 | 0.08 | 0 | Neutral |
| testneg_2742 | 0.08 | 0.03 | 1.617731 | -3 | 0.64021 | -3 | 0.87038 | 0.43956 | 0.18597 | 0.12788 | 7.17593 | 0.050270869 | 0 | Neutral |
| testneg_2743 | 0 | 0 | 0.441601 | -3 | -0.422206 | -3 | 0.04032 | 0.12788 | 0.1231 | 0.3662 | 91.47641 | 0.258467262 | 0.2 | Neutral |
| testneg_2744 | 0 | 0 | 0.438443 | 0.438443 | 0.977001 | -3 | 0.36857 | 0.36135 | 0.42748 | 0.10231 | 12.10459 | 0.022546296 | 0 | Neutral |
| testneg_2745 | 0.05 | -0.81 | 0.48163 | -3 | 1.048529 | -3 | 0.03008 | 0.12788 | 0.54625 | 0.11856 | 17.21847 | 0.04 | 0 | Neutral |
| testneg_2746 | -0.18 | -0.16 | -1.600047 | -3 | 5.33097 | -3 | 0.47217 | 0.15544 | 0.08814 | 0.42357 | 6.57332 | 0.49287444 | 52.2 | Disease_causing |
| testneg_2747 | 0.18 | -1.05 | 2.686668 | -3 | 2.669855 | -3 | 0.10517 | 0.09803 | 0.80849 | 0.21975 | 10 | 0.233021978 | 0.4 | Neutral |
| testneg_2748 | 0 | 0 | -0.68826 | -0.68826 | -2.112238 | -3 | 0.32076 | 0.14229 | 0.39576 | 0.10761 | 16.8018 | 0.01 | 0 | Neutral |
| testneg_2749 | 0.03 | 0.1 | -1.566121 | -1.566121 | 0.641246 | -3 | 0.12003 | 0.12788 | 0.21118 | 0.10074 | 37.26752 | 0.01 | 0 | Neutral |
| testneg_2750 | -0.06 | -0.18 | 1.556626 | -3 | 2.299451 | -3 | 0.46919 | 0.60081 | 0.24832 | 0.84371 | 12.29167 | 0.268047619 | 0.8 | Neutral |
| testneg_2751 | 0 | 0 | 0.889458 | 0.889458 | -1.010002 | -3 | 0.48937 | 0.22085 | 0.07408 | 0.12788 | 27.2619 | 0.05 | 0 | Neutral |
| testneg_2752 | 0 | 0 | -0.583572 | -0.583572 | 1.821777 | -3 | 0.04555 | 0.06327 | 0.11187 | 0.12788 | 16.8018 | 0.04 | 0 | Neutral |
| testneg_2753 | 0 | 0 | 0.889458 | 0.889458 | 5.070123 | -3 | 0.48937 | 0.22085 | 0.34946 | 0.19092 | 11.875 | 0.448453535 | 35.6 | Neutral |
| testneg_2754 | -0.14 | 1.01 | 3.981462 | -3 | -0.237972 | -3 | 0.4113 | 0.2303 | 0.21972 | 0.12788 | 32.73994 | 0.15 | 0 | Neutral |
| testneg_2755 | 0 | 0 | -0.156459 | -3 | 2.761391 | -3 | 0.10774 | 0.13117 | 0.31859 | 0.11262 | 12.29167 | 0.03 | 0 | Neutral |
| testneg_2756 | 0 | -0.32 | 2.97409 | -3 | 3.396513 | -3 | 0.16544 | 0.43246 | 0.17782 | 0.1217 | 13.7042 | 0.239934211 | 0.2 | Neutral |
| testneg_2757 | -0.08 | -0.03 | 2.342086 | -3 | 3.482726 | -3 | 0.5901 | 0.19444 | 0.05999 | 0.12788 | 17.21847 | 0.130052632 | 0 | Neutral |
| testneg_2758 | 0.02 | 0.45 | 6.259081 | -3 | -0.78887 | -3 | 0.89972 | 0.10766 | 0.47188 | 0.10037 | 10.30222 | 0.23 | 0.4 | Neutral |
| testneg_2759 | -0.08 | -0.03 | 4.604598 | -3 | 7.263562 | -3 | 0.09767 | 0.39937 | 0.1725 | 0.09643 | 20.23352 | 0.63 | 94.4 | Disease_causing |
| testneg_2760 | 0.07 | -0.31 | 1.726579 | -3 | 2.083556 | -3 | 0.25421 | 0.16235 | 0.26482 | 0.12788 | 29.16667 | 0.04 | 0 | Neutral |
| testneg_2761 | 0.21 | 0.15 | 1.704228 | -3 | 0.055332 | -3 | 0.27062 | 0.12378 | 0.12051 | 0.08503 | 18.6768 | 0.03 | 0 | Neutral |
| testneg_2762 | -0.000303 | 0.00015152 | -0.119638 | -3 | 5.652146 | -3 | 0.16379 | 0.31462 | 0.09951 | 0.14299 | 10.08603 | 0.493824726 | 51.4 | Disease_causing |
| testneg_2763 | -0.15 | 0.28 | 2.650636 | -3 | 5.028164 | -3 | 0.82595 | 0.52905 | 0.29209 | 0.09791 | 10.00351 | 0.327583333 | 3.4 | Neutral |
| testneg_2764 | 0 | 0 | 1.252608 | -3 | 5.494664 | -3 | 0.28334 | 0.10368 | 0.19898 | 0.12788 | 19.83449 | 0.06 | 0 | Neutral |
| testneg_2765 | 0 | 0 | 1.569245 | -3 | -1.528492 | -3 | 0.99859 | 0.12788 | 0.12581 | 0.08118 | 10 | 0.04 | 0 | Neutral |
| testneg_2766 | -0.03 | -0.1 | 1.318946 | -3 | 1.528824 | -3 | 0.85508 | 0.12634 | 0.05216 | 0.23851 | 13.3297 | 0.069922222 | 0 | Neutral |
| testneg_2767 | 0.16 | -1.37 | 6.076778 | -3 | 0.854295 | -3 | 0.33818 | 0.12788 | 0.71844 | 0.20204 | 10.41667 | 0.39093295 | 17.8 | Neutral |
| testneg_2768 | 0 | -0.01 | 0.981481 | -3 | 6.119768 | -3 | 0.24516 | 0.11809 | 0.19898 | 0.12788 | 18.63562 | 0.073706897 | 0 | Neutral |
| testneg_2769 | -0.07 | -0.38 | 1.811735 | -3 | 1.885526 | -3 | 0.15767 | 0.12788 | 0.11233 | 0.28224 | 19.09347 | 0.092956349 | 0 | Neutral |
| testneg_2770 | -0.08 | -0.03 | -1.031651 | -3 | -2.565987 | -3 | 0.90767 | 0.12788 | 0.06072 | 0.09873 | 12.70833 | 0.03 | 0 | Neutral |
| testneg_2771 | 0.14 | -0.37 | 6.930926 | -3 | 0.208107 | -3 | 0.41743 | 0.78545 | 0.10863 | 0.098 | 16.8018 | 0.39 | 14.4 | Neutral |
| testneg_2772 | 0.21 | 0.15 | 2.322245 | -3 | 0.116245 | -3 | 0.30734 | 0.12788 | 0.8238 | 0.38836 | 20.27778 | 0.12 | 0 | Neutral |
| testneg_2773 | 0 | 0 | 1.826458 | 1.826458 | 1.457223 | -3 | 0.46401 | 0.24608 | 0.06845 | 0.13843 | 11.875 | 0.17 | 0 | Neutral |
| testneg_2774 | 0.05 | 0.29 | -1.763617 | -3 | 6.415747 | -3 | 0.19433 | 0.12788 | 0.36852 | 0.0903 | 50.51283 | 0.150162833 | 0 | Neutral |
| testneg_2775 | 0.03 | 0.12 | 0.235146 | -3 | 7.401156 | -3 | 0.16856 | 0.19221 | 0.1311 | 0.12667 | 30.89369 | 0.42 | 26.2 | Neutral |
| testneg_2776 | 0.21 | 0.15 | 1.631249 | -3 | 1.579861 | -3 | 0.47166 | 0.13548 | 0.64566 | 0.22552 | 20.83333 | 0.126532706 | 0 | Neutral |
| testneg_2777 | -0.02 | 0.68 | 1.8794 | -3 | -0.074833 | -3 | 0.54711 | 0.11809 | 0.65981 | 0.47389 | 10.72072 | 0.060542788 | 0 | Neutral |
| testneg_2778 | -0.14 | 1.85 | -0.749376 | -0.749376 | 3.628039 | 3.628039 | 0.51437 | 0.17173 | 0.055 | 0.20858 | 10.83333 | 0.553507326 | 75.8 | Disease_causing |
| testneg_2779 | 0.04 | -0.23 | 1.711588 | -3 | 1.495922 | -3 | 0.44615 | 0.09968 | 0.1316 | 0.28283 | 10.83333 | 0.05442316 | 0 | Neutral |
| testneg_2780 | 0 | -0.01 | -0.052748 | -3 | 5.105753 | 1.261526 | 0.91139 | 0.65138 | 0.25637 | 0.37408 | 23.51844 | 0.409169109 | 18.2 | Neutral |
| testneg_2781 | 0 | 0 | 0.494101 | -3 | 4.155257 | -3 | 0.28038 | 0.11814 | 0.55193 | 0.22384 | 14.97692 | 0.171062315 | 0 | Neutral |
| testneg_2782 | -0.16 | -0.08 | 2.86141 | -3 | 0.05288 | -3 | 0.06517 | 0.15107 | 0.15025 | 0.4872 | 21.42525 | 0.322256211 | 2 | Neutral |
| testneg_2783 | 0 | 0 | 1.302665 | -3 | -0.6503 | -3 | 0.71803 | 0.95763 | 0.13994 | 0.11181 | 10.41667 | 0.06 | 0 | Neutral |
| testneg_2784 | 0 | 0 | 0.585709 | -3 | 5.711744 | -3 | 0.77644 | 0.09695 | 0.22874 | 0.13202 | 17.78631 | 0.181440126 | 0 | Neutral |
| testneg_2785 | 0.07 | -0.31 | 4.193501 | -3 | 0.503836 | -3 | 0.20737 | 0.11262 | 0.19898 | 0.07875 | 43.56906 | 0.02 | 0 | Neutral |
| testneg_2786 | -0.08 | -0.03 | 1.562114 | -3 | 5.671166 | -3 | 0.78356 | 0.13377 | 0.08982 | 0.15279 | 21.54146 | 0.406698718 | 16.2 | Neutral |
| testneg_2787 | 0.05 | -0.81 | 4.053211 | -3 | 3.273184 | -3 | 0.58677 | 0.11579 | 0.83114 | 0.60631 | 10.43529 | 0.416302874 | 21.6 | Neutral |
| testneg_2788 | 0 | 0 | 0.368677 | -3 | -1.873728 | -1.873728 | 0.02518 | 0.07165 | 0.19898 | 0.12788 | 23.91414 | 0.03 | 0 | Neutral |
| testneg_2789 | -0.06 | -0.18 | 2.055896 | -3 | 6.490817 | -3 | 0.15509 | 0.10665 | 0.12446 | 0.08908 | 16.01246 | 0.197234848 | 0.2 | Neutral |
| testneg_2790 | 0.16 | -1.37 | -0.121102 | -0.121102 | 0.19932 | -3 | 0.78935 | 0.17351 | 0.19898 | 0.09927 | 10.22898 | 0.06 | 0 | Neutral |
| testneg_2791 | 0.07 | 0.38 | 1.022289 | -3 | 1.143955 | -3 | 0.18281 | 0.12788 | 0.62645 | 0.12788 | 13.46801 | 0.01 | 0 | Neutral |
| testneg_2792 | 0 | 0 | 0.556564 | -3 | -0.294826 | -3 | 0.18834 | 0.12788 | 0.63818 | 0.12788 | 10.56765 | 0.000240999 | 0 | Neutral |
| testneg_2793 | 0 | 0 | 1.27406 | -3 | 1.957496 | -3 | 0.37855 | 0.47307 | 0.06395 | 0.0759 | 12.52072 | 0.02 | 0 | Neutral |
| testneg_2794 | 0 | 0 | 0.043071 | -3 | 4.969986 | -3 | 0.23027 | 0.37171 | 0.49831 | 0.29427 | 36.95035 | 0.461365916 | 41.4 | Neutral |
| testneg_2795 | -0.04 | -2.01 | 2.038814 | -3 | 3.976336 | -3 | 0.24524 | 0.10803 | 0.13185 | 0.51629 | 12.77955 | 0.380940146 | 12.8 | Neutral |
| testneg_2796 | -0.03 | -0.1 | 3.804607 | -3 | 0.350329 | -3 | 0.36694 | 0.12788 | 0.19898 | 0.12788 | 25.6804 | 0.03 | 0 | Neutral |
| testneg_2797 | 0 | 0 | 1.870488 | -3 | 3.085995 | -1.606767 | 0.3548 | 0.10717 | 0.09471 | 0.12788 | 20.48471 | 0.038175156 | 0 | Neutral |
| testneg_2798 | 0 | 0 | 0.844548 | -3 | 0.041481 | -0.17228 | 0.9237 | 0.55859 | 0.47147 | 0.09503 | 11.41859 | 0.03 | 0 | Neutral |
| testneg_2799 | 0 | 0 | 0.600565 | -3 | 4.710858 | -3 | 0.99564 | 0.25794 | 0.58985 | 0.10504 | 11.77783 | 0.101371212 | 0 | Neutral |
| testneg_2800 | 0 | 0 | -0.095889 | -3 | 2.111463 | -0.354552 | 0.14643 | 0.12788 | 0.09815 | 0.12788 | 24.56929 | 0.02 | 0 | Neutral |
| testneg_2801 | 0 | 0 | 0.332849 | -3 | 1.461941 | -3 | 0.99595 | 0.23021 | 0.49276 | 0.10648 | 17.3494 | 0.04 | 0 | Neutral |
| testneg_2802 | -0.18 | -0.16 | 2.074931 | -3 | -0.808449 | -0.808449 | 0.18488 | 0.53379 | 0.13258 | 0.9258 | 10 | 0.28442055 | 0.6 | Neutral |
| testneg_2803 | 0 | -0.01 | -0.052748 | -3 | 3.640451 | -3 | 0.91139 | 0.65138 | 0.19898 | 0.12788 | 26.38889 | 0.13 | 0 | Neutral |
| testneg_2804 | 0.07 | 0.7 | 5.309515 | -3 | -1.14689 | -3 | 0.53292 | 0.32174 | 0.14704 | 0.12788 | 17.26246 | 0.218969551 | 0 | Neutral |
| testneg_2805 | 0.04 | 2.01 | 5.43333 | -3 | 2.421448 | -3 | 0.15704 | 0.72174 | 0.48075 | 0.20313 | 10 | 0.722303385 | 99.6 | Disease_causing |

| | | | | | | | | | | | | | | |
|---|---|---|---|---|---|---|---|---|---|---|---|---|---|---|
| testneg_2806 | 0.12 | -1.16 | 4.96652 | -3 | 4.641576 | -3 | 0.24287 | 0.11626 | 0.10385 | 0.09775 | 18.88889 | 0.251522436 | 0.4 | Neutral |
| testneg_2807 | 0.05 | -0.81 | -0.646746 | -3 | 0.281453 | -3 | 0.12073 | 0.10969 | 0.16052 | 0.12788 | 16.42913 | 0.01 | 0 | Neutral |
| testneg_2808 | -0.01 | -0.8 | 7.657206 | -3 | -1.690706 | -3 | 0.19898 | 0.12788 | 0.08779 | 0.1777 | 10.41667 | 0.51 | 58.8 | Disease_causing |
| testneg_2809 | 0.21 | 0.15 | 2.322245 | -3 | 4.093803 | -3 | 0.30734 | 0.12788 | 0.35234 | 0.1303 | 10.36687 | 0.15 | 0 | Neutral |
| testneg_2810 | 0.09 | -0.3 | 5.832895 | -3 | -0.850513 | -3 | 0.15524 | 0.10607 | 0.06982 | 0.10226 | 20.60663 | 0.11 | 0 | Neutral |
| testneg_2811 | 0.03 | 0.12 | 1.304269 | -3 | 1.430763 | -3 | 0.45029 | 0.12788 | 0.2578 | 0.09796 | 12.34649 | 0.01 | 0 | Neutral |
| testneg_2812 | 0 | 0 | 1.247588 | -3 | 3.006809 | -3 | 0.12835 | 0.15143 | 0.84365 | 0.12788 | 10.26467 | 0.061592262 | 0 | Neutral |
| testneg_2813 | -0.15 | -0.04 | 1.977578 | -3 | 5.235925 | -3 | 0.06877 | 0.17777 | 0.50161 | 0.12788 | 20.71976 | 0.388333333 | 10.4 | Neutral |
| testneg_2814 | -0.07 | -0.7 | 3.792287 | -3 | 4.061878 | -3 | 0.07864 | 0.12788 | 0.13322 | 0.11297 | 34.45035 | 0.151398771 | 0 | Neutral |
| testneg_2815 | -0.02 | -0.13 | 3.206825 | -3 | -1.097042 | -3 | 0.55197 | 0.11305 | 0.10703 | 0.11611 | 33.30895 | 0.031811975 | 0 | Neutral |
| testneg_2816 | 0.2 | -1.26 | 1.286107 | -3 | 4.793875 | -3 | 0.38207 | 0.35052 | 0.19898 | 0.20319 | 10.83333 | 0.581393281 | 82.6 | Disease_causing |
| testneg_2817 | -0.09 | 0.3 | -0.208904 | -3 | 6.470932 | -3 | 0.40398 | 0.12788 | 0.16662 | 0.11306 | 39.41832 | 0.215727273 | 0.2 | Neutral |
| testneg_2818 | 0 | 0 | 0.5524 | -3 | -0.111739 | -3 | 0.44644 | 0.10813 | 0.08237 | 0.09015 | 21.42913 | 0.010529264 | 0 | Neutral |
| testneg_2819 | 0.14 | -1.01 | 6.136401 | -3 | 3.265474 | -3 | 0.3238 | 0.18333 | 0.17458 | 0.12788 | 31.0866 | 0.44 | 31.6 | Neutral |
| testneg_2820 | 0.07 | 0.7 | 4.605651 | -3 | 0.125673 | -3 | 0.10719 | 0.08618 | 0.04912 | 0.08574 | 29.9024 | 0.06 | 0 | Neutral |
| testneg_2821 | -0.14 | 1.85 | 0.866269 | -3 | 1.932311 | 1.932311 | 0.30876 | 0.28825 | 0.97673 | 0.22976 | 10.41667 | 0.291678161 | 0 | Neutral |
| testneg_2822 | 0.1 | 1.9 | 6.154974 | -3 | 7.397319 | -3 | 0.04515 | 0.14318 | 0.19355 | 0.18675 | 13.32524 | 0.822222222 | 100 | Disease_causing |
| testneg_2823 | 0.02 | -0.68 | 0.188403 | -3 | 6.264472 | -3 | 0.39111 | 0.15887 | 0.71276 | 0.88182 | 16.8018 | 0.568119959 | 78.4 | Disease_causing |
| testneg_2824 | -0.18 | -0.16 | -1.333487 | -3 | 0.986854 | -3 | 0.07598 | 0.08456 | 0.13468 | 0.09235 | 57.98926 | 0.1 | 0 | Neutral |
| testneg_2825 | 0.04 | -0.23 | 2.906184 | -3 | 0.909747 | -3 | 0.32188 | 0.08974 | 0.19898 | 0.12788 | 29.10257 | 0.02 | 0 | Neutral |
| testneg_2826 | 0 | 0 | -0.726772 | -3 | 4.226361 | -3 | 0.1072 | 0.63446 | 0.82846 | 0.13042 | 18.6768 | 0.26196484 | 0.8 | Neutral |
| testneg_2827 | 0.21 | 0.15 | -0.762386 | -3 | 1.636511 | -3 | 0.07014 | 0.53339 | 0.16355 | 0.12788 | 16.8018 | 0.111774892 | 0 | Neutral |
| testneg_2828 | -0.08 | -0.03 | 3.600262 | -3 | 6.276709 | -3 | 0.22944 | 0.09039 | 0.20379 | 0.35591 | 21.94444 | 0.569422863 | 80.6 | Disease_causing |
| testneg_2829 | -0.06 | -0.18 | 2.864843 | -3 | 1.749206 | -3 | 0.63363 | 0.14795 | 0.03651 | 0.06141 | 13.69873 | 0.05 | 0 | Neutral |
| testneg_2830 | -0.18 | 1.05 | 1.550311 | -3 | 0.66227 | -3 | 0.17443 | 0.19181 | 0.23141 | 0.12788 | 26.34995 | 0.13 | 0 | Neutral |
| testneg_2831 | 0 | 0 | -0.031197 | -3 | 1.890706 | -3 | 0.1017 | 0.11809 | 0.48688 | 0.13071 | 19.89856 | 0.02 | 0 | Neutral |
| testneg_2832 | 0 | -0.01 | 5.907423 | -3 | -1.589955 | -3 | 0.00451 | 0.2022 | 0.05732 | 0.1098 | 59.35071 | 0.27 | 1.2 | Neutral |
| testneg_2833 | 0.21 | 0.15 | 5.170625 | -3 | 5.018695 | -3 | 0.10153 | 0.13343 | 0.19301 | 0.12955 | 20.5518 | 0.383754579 | 17 | Neutral |
| testneg_2834 | -0.04 | -2.01 | 3.157424 | -3 | -0.099626 | -0.099626 | 0.09067 | 0.10671 | 0.11281 | 0.0958 | 13.02635 | 0.062777778 | 0 | Neutral |
| testneg_2835 | -0.23 | 1.86 | 5.658336 | -3 | -0.657715 | -3 | 0.45014 | 0.12788 | 0.07684 | 0.09138 | 16.8018 | 0.23 | 0.4 | Neutral |
| testneg_2836 | 0 | 0 | 0.365683 | -3 | 2.817469 | -3 | 0.31648 | 0.12788 | 0.84491 | 0.30945 | 19.87691 | 0.09 | 0 | Neutral |
| testneg_2837 | 0.03 | -1.03 | 4.715557 | -3 | 5.765818 | -3 | 0.04243 | 0.22343 | 0.10835 | 0.12788 | 18.33333 | 0.489230072 | 50 | Neutral |
| testneg_2838 | 0 | -0.01 | 0.632196 | -3 | 2.058739 | -3 | 0.0984 | 0.1489 | 0.19898 | 0.12788 | 18.0784 | 0.03 | 0 | Neutral |
| testneg_2839 | 0.03 | -1.03 | 2.660158 | -3 | 2.058438 | -3 | 0.18243 | 0.11102 | 0.0749 | 0.12788 | 17.94606 | 0.021602564 | 0 | Neutral |
| testneg_2840 | 0.01 | 0.79 | 2.557398 | -3 | 4.140289 | -3 | 0.961 | 0.25991 | 0.0922 | 0.07873 | 19.82027 | 0.190731091 | 0 | Neutral |
| testneg_2841 | 0 | 0 | -0.535508 | -3 | 0.878873 | -3 | 0.01196 | 0.12788 | 0.62197 | 0.12788 | 18.09942 | 0.03 | 0 | Neutral |
| testneg_2842 | 0 | 0 | 1.115221 | -3 | 3.296898 | -3 | 0.64188 | 0.09494 | 0.13559 | 0.1946 | 18.6768 | 0.17627876 | 0 | Neutral |
| testneg_2843 | -0.03 | -0.12 | -1.873728 | -1.873728 | 2.252712 | -3 | 0.19898 | 0.12788 | 0.47789 | 0.10803 | 22.77778 | 0.02 | 0 | Neutral |
| testneg_2844 | -0.08 | -0.03 | 2.82043 | -3 | 1.846205 | -3 | 0.09346 | 0.12788 | 0.23078 | 0.10588 | 21.80782 | 0.021388889 | 0 | Neutral |
| testneg_2845 | 0 | 0 | -0.566651 | -3 | -0.593437 | -3 | 0.38016 | 0.14503 | 0.18264 | 0.10206 | 18.6768 | 0.01 | 0 | Neutral |
| testneg_2846 | 0 | 0 | 5.510751 | -3 | 1.175658 | -3 | 0.35428 | 0.12971 | 0.10285 | 0.10886 | 17.90495 | 0.175 | 0 | Neutral |
| testneg_2847 | 0 | -0.01 | 0.71533 | -3 | 2.273331 | -3 | 0.42802 | 0.12096 | 0.11237 | 0.08916 | 27.84878 | 0.01026905 | 0 | Neutral |
| testneg_2848 | -0.14 | 1.01 | 0.576229 | -3 | -0.369883 | -3 | 0.18022 | 0.12788 | 0.37909 | 0.4568 | 26.95989 | 0.119726515 | 0 | Neutral |
| testneg_2849 | 0.07 | 0.33 | 1.275096 | -3 | 1.8794 | -3 | 0.1592 | 0.12788 | 0.54711 | 0.11809 | 18.76159 | 0.01 | 0 | Neutral |
| testneg_2850 | -0.08 | -0.03 | 5.054259 | -3 | 5.840704 | -3 | 0.29478 | 0.12788 | 0.37909 | 0.4568 | 12.11131 | 0.74024025 | 99 | Disease_causing |
| testneg_2851 | -0.14 | 1.01 | 5.407194 | -3 | 4.1206 | -3 | 0.03631 | 0.80564 | 0.66693 | 0.0986 | 12.29167 | 0.63 | 94.8 | Disease_causing |
| testneg_2852 | -0.04 | 0.23 | 0.6838 | 0.6838 | 3.192398 | -3 | 0.67327 | 0.23543 | 0.19898 | 0.12788 | 26.04694 | 0.158273973 | 0 | Neutral |
| testneg_2853 | 0 | 0 | 2.041671 | -3 | 0.822036 | -3 | 0.37056 | 0.12309 | 0.13575 | 0.12788 | 13.19331 | 0 | 0 | Neutral |
| testneg_2854 | 0.05 | -0.81 | 3.217486 | -3 | -0.641545 | -3 | 0.74699 | 0.12286 | 0.09749 | 0.82924 | 32.29167 | 0.189666667 | 0 | Neutral |
| testneg_2855 | 0.02 | -0.28 | -1.20584 | -3 | 0.94842 | -3 | 0.53216 | 0.57092 | 0.16121 | 0.10081 | 42.85462 | 0.080278638 | 0 | Neutral |
| testneg_2856 | -0.03 | -0.1 | 0.461043 | -3 | 7.363899 | -3 | 0.62864 | 0.11103 | 0.0669 | 0.10997 | 29.95254 | 0.25 | 1 | Neutral |
| testneg_2857 | -0.04 | 0.23 | 1.375409 | -3 | 4.228814 | -3 | 0.10285 | 0.11721 | 0.07675 | 0.11464 | 18.27736 | 0.081376832 | 0 | Neutral |
| testneg_2858 | -0.03 | -0.1 | 2.935592 | -3 | -0.048251 | -0.048251 | 0.1909 | 0.14279 | 0.15646 | 0.12788 | 16.48772 | 0.04 | 0 | Neutral |
| testneg_2859 | -0.16 | -0.08 | 1.251399 | -3 | -0.681767 | -3 | 0.20329 | 0.10718 | 0.226 | 0.28784 | 12.34649 | 0.102916667 | 0 | Neutral |
| testneg_2860 | 0 | 0 | -0.491928 | -3 | 0.915016 | -3 | 0.1056 | 0.12788 | 0.13615 | 0.12788 | 13.72212 | 0 | 0 | Neutral |
| testneg_2861 | 0.2 | -1.26 | 3.642099 | -3 | 1.473246 | -3 | 0.17768 | 0.10832 | 0.29013 | 0.11368 | 31.44713 | 0.06 | 0 | Neutral |
| testneg_2862 | 0.05 | -0.81 | 0.099811 | -3 | 1.690495 | -3 | 0.2877 | 0.18291 | 0.11891 | 0.11186 | 17.28802 | 0.03 | 0 | Neutral |
| testneg_2863 | 0.04 | 2.01 | 5.095532 | -3 | 0.152832 | -3 | 0.01573 | 0.10031 | 0.04829 | 0.09848 | 30.49816 | 0.14 | 0 | Neutral |
| testneg_2864 | 0 | 0 | -0.157172 | -3 | 3.261846 | -3 | 0.39344 | 0.12788 | 0.06474 | 0.23607 | 11.25 | 0.170565926 | 0 | Neutral |
| testneg_2865 | 0.01666667 | -0.1875 | 0.933869 | -3 | 0.30563 | -3 | 0.9862 | 0.4528 | 0.1417 | 0.1398 | 12.33727 | 0.11 | 0 | Neutral |
| testneg_2866 | 0 | 0 | 1.383368 | -3 | 3.010558 | -3 | 0.19898 | 0.13057 | 0.1331 | 0.12788 | 19.97695 | 0.030228833 | 0 | Neutral |
| testneg_2867 | 0.05 | 0.29 | 2.297641 | -3 | 2.367948 | -3 | 0.80629 | 0.12788 | 0.85781 | 0.22949 | 12.29167 | 0.11 | 0 | Neutral |
| testneg_2868 | 0.02 | -0.63 | 0.686794 | 0.665096 | 3.233126 | -3 | 0.15132 | 0.11724 | 0.66422 | 0.99377 | 10.83333 | 0.284484127 | 2.2 | Neutral |
| testneg_2869 | 0.03 | 0.1 | 4.072509 | -3 | 0.433941 | 0.433941 | 0.19898 | 0.12788 | 0.30475 | 0.21064 | 15.66142 | 0.179676277 | 0 | Neutral |
| testneg_2870 | 0 | 0 | 0.919505 | -3 | 2.65604 | -3 | 0.19898 | 0.12788 | 0.25171 | 0.13527 | 13.51351 | 0.080689186 | 0 | Neutral |
| testneg_2871 | -0.06 | -0.18 | 4.409326 | -3 | 1.836976 | -3 | 0.62708 | 0.24932 | 0.84491 | 0.30945 | 5.35211 | 0.361795303 | 9.2 | Neutral |
| testneg_2872 | 0.09 | -0.3 | 4.414734 | -3 | 0.953612 | 0.410924 | 0.05657 | 0.14812 | 0.18431 | 0.13863 | 12.77241 | 0.239349206 | 0.2 | Neutral |
| testneg_2873 | -0.19 | 0.27 | 1.951681 | -3 | 1.13018 | -3 | 0.1875 | 0.10316 | 0.50505 | 0.11563 | 34.48179 | 0.06 | 0 | Neutral |
| testneg_2874 | 0 | 0 | 0.346053 | -3 | 2.971017 | 2.971017 | 0.3047 | 0.11929 | 0.15581 | 0.10782 | 29.724 | 0.100428779 | 0 | Neutral |
| testneg_2875 | 0 | 0 | -0.843404 | -3 | 1.853291 | -3 | 0.11587 | 0.12001 | 0.13468 | 0.09235 | 28.38173 | 0.01 | 0 | Neutral |
| testneg_2876 | 0.02 | -0.63 | 2.466761 | -3 | 1.65767 | 1.65767 | 0.0538 | 0.23912 | 0.89992 | 0.10231 | 13.75 | 0.13 | 0 | Neutral |
| testneg_2877 | 0 | 0 | -0.784713 | -0.784713 | 2.598387 | -3 | 0.54899 | 0.31462 | 0.24331 | 0.11911 | 10.68487 | 0.049554992 | 0 | Neutral |
| testneg_2878 | 0 | -0.01 | 0.741645 | -3 | 2.986041 | -3 | 0.04606 | 0.20522 | 0.05183 | 0.18779 | 27.64576 | 0.328812392 | 5 | Neutral |
| testneg_2879 | 0 | 0 | 0.760575 | -3 | -0.071864 | -3 | 0.1878 | 0.08946 | 0.97005 | 0.31407 | 13.93265 | 0.03 | 0 | Neutral |
| testneg_2880 | -0.06 | -0.18 | 0.491194 | 0.491194 | 2.459196 | -3 | 0.72124 | 0.09906 | 0.63484 | 0.16728 | 17.11963 | 0.13 | 0 | Neutral |
| testneg_2881 | 0 | -0.01 | 0.242595 | -3 | 0.238703 | -3 | 0.11358 | 0.12788 | 0.0887 | 0.10047 | 26.88873 | 0.01 | 0 | Neutral |
| testneg_2882 | 0 | 0 | 0.494384 | -3 | 5.579113 | -3 | 0.20604 | 0.10821 | 0.94859 | 0.75045 | 8.97898 | 0.302762594 | 3.2 | Neutral |
| testneg_2883 | 0.14 | -1.01 | 2.621366 | -3 | 5.308589 | -3 | 0.81899 | 0.12788 | 0.10168 | 0.12788 | 12.32809 | 0.181416667 | 0 | Neutral |
| testneg_2884 | 0.09 | -0.3 | 4.733195 | -3 | 1.453519 | -3 | 0.161 | 0.11597 | 0.15147 | 0.12788 | 17.32838 | 0.037124459 | 0 | Neutral |
| testneg_2885 | 0 | -0.01 | 2.432573 | -3 | 1.150515 | -3 | 0.32804 | 0.10857 | 0.12408 | 0.08828 | 18.88889 | 0.000580808 | 0 | Neutral |
| testneg_2886 | 0.06 | -0.57 | 5.777528 | -3 | 0.196098 | -3 | 0.17672 | 0.14229 | 0.12682 | 0.11357 | 18.98474 | 0.163632353 | 0 | Neutral |
| testneg_2887 | 0 | 0 | 1.609578 | -3 | 0.853858 | -3 | 0.19898 | 0.07094 | 0.35513 | 0.21966 | 27.5 | 0.04 | 0 | Neutral |
| testneg_2888 | 0.22 | -2.66 | 3.810217 | -3 | 0.448323 | -3 | 0.10562 | 0.09329 | 0.19898 | 0.08486 | 16.42913 | 0.08 | 0 | Neutral |
| testneg_2889 | 0 | 0 | -0.014147 | -3 | -2.277439 | -3 | 0.14101 | 0.14318 | 0.17451 | 0.11227 | 17.77778 | 0.01 | 0 | Neutral |
| testneg_2890 | 0.12 | -1.14 | 1.65767 | 1.65767 | 1.304689 | -3 | 0.89992 | 0.10231 | 0.06134 | 0.08207 | 10.22898 | 0.071725 | 0 | Neutral |
| testneg_2891 | 0.14 | -1.01 | 2.297111 | -3 | 6.254737 | -3 | 0.39111 | 0.15887 | 0.39225 | 0.10625 | 17.21847 | 0.260436364 | 0.6 | Neutral |
| testneg_2892 | 0 | 0 | 0.374605 | -3 | 2.264893 | -3 | 0.45112 | 0.12083 | 0.07204 | 0.07708 | 10.18172 | 0.01 | 0 | Neutral |
| testneg_2893 | 0 | -0.32 | -0.334946 | -3 | 1.013474 | -3 | 0.10895 | 0.11082 | 0.19898 | 0.10778 | 25.94766 | 0.01 | 0 | Neutral |
| testneg_2894 | -0.03 | -0.1 | 3.350172 | -3 | 1.485335 | -3 | 0.13331 | 0.13638 | 0.19894 | 0.27411 | 10.83333 | 0.250772412 | 1 | Neutral |
| testneg_2895 | -0.01 | 1.44 | 4.608667 | -3 | -1.218152 | -3 | 0.97273 | 0.27711 | 0.19898 | 0.12788 | 16.52897 | 0.15 | 0 | Neutral |
| testneg_2896 | 0 | 0 | 1.4365 | -3 | -0.343322 | -3 | 0.68029 | 0.12788 | 0.06541 | 0.12012 | 11.25 | 0.01 | 0 | Neutral |
| testneg_2897 | -0.08 | -0.03 | 6.667592 | -3 | 5.593394 | -3 | 0.35475 | 0.12788 | 0.25674 | 0.83834 | 10.41667 | 0.79 | 99.6 | Disease_causing |
| testneg_2898 | 0 | 0 | 0.082683 | -3 | -0.605051 | -3 | 0.19898 | 0.12788 | 0.48144 | 0.46642 | 15.89722 | 0.031338936 | 0 | Neutral |
| testneg_2899 | 0 | -0.01 | 0.242595 | -3 | 0.476024 | -3 | 0.11358 | 0.12788 | 0.24832 | 0.21196 | 14.45887 | 0.020350347 | 0 | Neutral |
| testneg_2900 | 0 | 0 | -0.072696 | -3 | 1.849294 | -3 | 0.30326 | 0.21562 | 0.11042 | 0.09881 | 20.42553 | 0.029285714 | 0 | Neutral |
| testneg_2901 | 0 | 0 | 2.437503 | -3 | 5.464083 | -3 | 0.11337 | 0.09363 | 0.19898 | 0.12788 | 18.33333 | 0.1 | 0 | Neutral |
| testneg_2902 | 0.00454545 | -0.0153636 | 2.150952 | -3 | 5.393603 | -3 | 0.19898 | 0.12788 | 0.19898 | 0.09584 | 18.0638 | 0.080709813 | 0 | Neutral |
| testneg_2903 | 0 | 0 | 4.038553 | -3 | 1.360191 | -3 | 0.54909 | 0.85056 | 0.10485 | 0.09565 | 19.09347 | 0.15 | 0 | Neutral |
| testneg_2904 | 0.21 | 0.15 | 4.652896 | -3 | 1.015573 | -3 | 0.21458 | 0.10398 | 0.16974 | 0.80184 | 30.14905 | 0.25 | 0.2 | Neutral |
| testneg_2905 | 0 | 0 | 0.42734 | -3 | 2.884683 | -3 | 0.19883 | 0.09434 | 0.11957 | 0.08975 | 16.37968 | 0.02 | 0 | Neutral |
| testneg_2906 | 0.02 | -0.28 | -0.483554 | -3 | -1.743651 | -3 | 0.19898 | 0.56316 | 0.38072 | 0.15993 | 21.38889 | 0.221236602 | 0 | Neutral |
| testneg_2907 | 0 | 0 | 0.453556 | -3 | 4.86235 | -3 | 0.28191 | 0.20693 | 0.12547 | 0.09267 | 21.53005 | 0.135599262 | 0 | Neutral |
| testneg_2908 | 0 | 0 | 1.335711 | -3 | 3.266081 | -3 | 0.2741 | 0.10478 | 0.09556 | 0.10109 | 24.68926 | 0.035941302 | 0 | Neutral |
| testneg_2909 | 0.07 | 0.38 | -2.352243 | -2.352243 | 2.696847 | -3 | 0.04568 | 0.08746 | 0.46037 | 0.1343 | 34.02305 | 0.18 | 0 | Neutral |
| testneg_2910 | 0 | 0 | 0.194425 | -3 | 1.820749 | -3 | 0.19898 | 0.10106 | 0.18559 | 0.09173 | 30.0251 | 0.000521328 | 0 | Neutral |
| testneg_2911 | -0.05 | -0.29 | -0.242457 | -3 | 2.84362 | -3 | 0.48662 | 0.23781 | 0.2903 | 0.11176 | 17.21847 | 0.072613636 | 0 | Neutral |
| testneg_2912 | 0 | 0 | 1.445184 | 1.445184 | 1.941255 | -3 | 0.33469 | 0.17432 | 0.35401 | 0.08438 | 9.46862 | 0.03 | 0 | Neutral |
| testneg_2913 | 0.19 | -0.25 | 3.925425 | -3 | 1.298472 | -3 | 0.24239 | 0.111 | 0.75415 | 0.29199 | 11.34055 | 0.206414141 | 0.2 | Neutral |
| testneg_2914 | 0 | 0 | 1.545946 | -3 | 0.439979 | -3 | 0.03844 | 0.12788 | 0.0811 | 0.09449 | 32.35237 | 0.03 | 0 | Neutral |
| testneg_2915 | -0.02 | 0.68 | 3.271047 | -3 | 3.146426 | -3 | 0.07684 | 0.09138 | 0.19898 | 0.12788 | 13.3074 | 0.150090909 | 0 | Neutral |
| testneg_2916 | 0.04 | -0.89 | 6.423333 | -3 | -2.069741 | -2.069741 | 0.16962 | 0.12788 | 0.16259 | 0.102 | 20.83333 | 0.18 | 0.2 | Neutral |
| testneg_2917 | -0.14 | 1.01 | 1.907441 | 1.907441 | 5.288301 | -3 | 0.20708 | 0.12788 | 0.56817 | 0.10766 | 31.53846 | 0.279605263 | 1 | Neutral |
| testneg_2918 | -0.03 | -1.22 | 2.648676 | -3 | 2.640446 | -3 | 0.30684 | 0.12788 | 0.26789 | 0.67432 | 21.10202 | 0.260577323 | 0.6 | Neutral |
| testneg_2919 | -0.21 | -0.15 | 3.777303 | -3 | 6.246312 | -3 | 0.44526 | 0.09404 | 0.87611 | 0.16759 | 28.95148 | 0.479807692 | 46.2 | Neutral |
| testneg_2920 | -0.07 | -0.38 | 1.785837 | -3 | 2.927374 | -3 | 0.22564 | 0.12009 | 0.78161 | 0.14363 | 10.41667 | 0.095928708 | 0 | Neutral |
| testneg_2921 | 0.18 | -1.05 | 7.749151 | -3 | 4.791156 | -3 | 0.11316 | 0.12788 | 0.17935 | 0.36953 | 12.31231 | 0.757858226 | 99.6 | Disease_causing |
| testneg_2922 | 0.21 | 0.15 | -0.762386 | -3 | 5.144919 | -3 | 0.07014 | 0.53339 | 0.32531 | 0.12351 | 16.8018 | 0.287693627 | 1 | Neutral |
| testneg_2923 | 0.22 | -2.66 | 6.97653 | 0.747661 | -0.923371 | -3 | 0.16204 | 0.12788 | 0.07611 | 0.11551 | 10.41667 | 0.32 | 4.2 | Neutral |
| testneg_2924 | 0.04 | -0.23 | 7.683159 | 7.683159 | 1.445664 | -3 | 0.89972 | 0.10766 | 0.14038 | 0.21177 | 10.87254 | 0.82 | 100 | Disease_causing |
| testneg_2925 | 0.09 | -0.25 | 1.849048 | -3 | 5.262564 | -3 | 0.15911 | 0.11022 | 0.21755 | 0.23824 | 30.39547 | 0.341985536 | 9 | Neutral |
| testneg_2926 | -0.2 | 1.26 | -0.70557 | -0.70557 | 1.74381 | -3 | 0.29008 | 0.19268 | 0.67644 | 0.11323 | 22.11633 | 0.180263158 | 0 | Neutral |
| testneg_2927 | -0.02 | 0.36 | -0.710033 | -3 | 3.08734 | -3 | 0.12689 | 0.13376 | 0.66693 | 0.0986 | 10.18363 | 0.08 | 0 | Neutral |

| | | | | | | | | | | | | | | |
|---|---|---|---|---|---|---|---|---|---|---|---|---|---|---|
| testneg_2928 | 0 | 0 | -0.39722 | -3 | 2.422379 | 2.422379 | 0.19343 | 0.11347 | 0.2485 | 0.10364 | 10.41667 | 0.06 | 0 | Neutral |
| testneg_2929 | 0.08 | 0.03 | 2.365497 | -3 | 1.649085 | -3 | 0.61491 | 0.11954 | 0.15822 | 0.12788 | 8.57881 | 0.03 | 0 | Neutral |
| testneg_2930 | -0.07 | -0.01 | 0.439914 | -3 | 7.431299 | -3 | 0.09275 | 0.14911 | 0.10371 | 0.08655 | 26.34995 | 0.429848485 | 25.8 | Neutral |
| testneg_2931 | 0 | 0 | 1.381745 | 0.782982 | -0.844526 | -3 | 0.18485 | 0.40044 | 0.28128 | 0.12883 | 12.29167 | 0.04 | 0 | Neutral |
| testneg_2932 | 0.004 | -0.4576364 | 7.731489 | -3 | 4.370033 | 4.08472 | 0.03378 | 0.12788 | 0.10213 | 0.22954 | 28.53489 | 0.708108527 | 98.4 | Disease_causing |
| testneg_2933 | 0 | 0 | 0.755726 | -3 | 3.002533 | 3.002533 | 0.08454 | 0.22908 | 0.14304 | 0.10715 | 35.41667 | 0.16 | 0 | Neutral |
| testneg_2934 | -0.07 | -0.38 | 3.673027 | -3 | 0.018659 | -3 | 0.19989 | 0.16777 | 0.38942 | 0.345 | 23.63636 | 0.301435685 | 2.4 | Neutral |
| testneg_2935 | -0.03 | -0.1 | 3.824071 | -3 | 3.263002 | -3 | 0.45029 | 0.12788 | 0.33992 | 0.15484 | 22.87765 | 0.357901662 | 10 | Neutral |
| testneg_2936 | 0 | -0.32 | -0.324056 | -3 | 5.265959 | -3 | 0.08999 | 0.0989 | 0.17231 | 0.1358 | 18.88889 | 0.250756548 | 0.4 | Neutral |
| testneg_2937 | 0.04 | -0.23 | 3.446417 | -3 | 0.224881 | -3 | 0.29013 | 0.11368 | 0.1653 | 0.0977 | 26.34564 | 0.010299567 | 0 | Neutral |
| testneg_2938 | 0.08 | 0.03 | 1.175993 | -3 | 0.318608 | -3 | 0.06781 | 0.09009 | 0.11373 | 0.10739 | 10.22898 | 0.02 | 0 | Neutral |
| testneg_2939 | 0.07 | -0.31 | 3.062683 | -3 | 2.307137 | -3 | 0.18507 | 0.12788 | 0.91587 | 0.10818 | 20.19804 | 0.057318182 | 0 | Neutral |
| testneg_2940 | 0.09 | -0.3 | 1.802254 | -3 | -0.867132 | -3 | 0.3793 | 0.24105 | 0.08353 | 0.21479 | 11.25 | 0.147122253 | 0 | Neutral |
| testneg_2941 | 0.04 | -0.23 | -0.126343 | -3 | -0.448326 | -3 | 0.11359 | 0.08631 | 0.03466 | 0.19518 | 24.07518 | 0.09 | 0 | Neutral |
| testneg_2942 | -0.07 | -0.38 | 3.920732 | -3 | 7.073967 | -3 | 0.0811 | 0.09449 | 0.11873 | 0.10499 | 32.40682 | 0.351547619 | 10.4 | Neutral |
| testneg_2943 | 0.18 | -1.05 | 6.546148 | -3 | 1.448132 | -3 | 0.59864 | 0.14106 | 0.694 | 0.09974 | 19.14601 | 0.312286765 | 4.2 | Neutral |
| testneg_2944 | -0.03 | -0.1 | 1.158025 | -3 | 7.755149 | -3 | 0.23968 | 0.12788 | 0.84365 | 0.12788 | 10.83333 | 0.317380952 | 4.8 | Neutral |
| testneg_2945 | 0 | -0.01 | 7.813348 | -3 | 0.952714 | -3 | 0.08836 | 0.0911 | 0.19979 | 0.09328 | 13.51351 | 0.32 | 5.8 | Neutral |
| testneg_2946 | 0.12 | -1.14 | 1.65767 | 1.65767 | -0.09216 | -3 | 0.89992 | 0.10231 | 0.39932 | 0.10693 | 10.11674 | 0.08 | 0 | Neutral |
| testneg_2947 | 0.22 | -2.66 | 6.776625 | -3 | 3.744082 | -3 | 0.17039 | 0.12788 | 0.94859 | 0.75045 | 8.84474 | 0.589722222 | 80.4 | Disease_causing |
| testneg_2948 | 0 | 0 | 0.372192 | -3 | 3.93025 | -3 | 0.53654 | 0.22867 | 0.19898 | 0.08486 | 18.33333 | 0.1025 | 0 | Neutral |
| testneg_2949 | 0.06 | -0.57 | 1.240379 | -3 | 1.742395 | -3 | 0.64591 | 0.38907 | 0.11055 | 0.19829 | 14.14796 | 0.150655453 | 0 | Neutral |
| testneg_2950 | -0.21 | -0.15 | -0.114657 | -3 | 1.932858 | -3 | 0.93228 | 0.1905 | 0.63645 | 0.14596 | 11.25 | 0.159960317 | 0 | Neutral |
| testneg_2951 | -0.08 | -0.03 | 0.201228 | -3 | 0.487522 | -3 | 0.19898 | 0.12788 | 0.16462 | 0.18467 | 20.27778 | 0.070346079 | 0 | Neutral |
| testneg_2952 | 0.02 | 0.45 | 3.640221 | 3.640221 | 2.624257 | -3 | 0.34673 | 0.24308 | 0.19043 | 0.07855 | 13.51351 | 0.47 | 43.8 | Neutral |
| testneg_2953 | 0.23 | -1.04 | 3.910876 | -3 | -1.648545 | -3 | 0.24287 | 0.11626 | 0.79857 | 0.12788 | 18.88889 | 0.07 | 0 | Neutral |
| testneg_2954 | 0.02 | -0.63 | 2.139035 | -3 | -0.113262 | -3 | 0.85829 | 0.4292 | 0.56154 | 0.0939 | 5.69444 | 0.03 | 0 | Neutral |
| testneg_2955 | 0 | -0.01 | 2.663664 | -3 | -3.156107 | -3 | 0.19898 | 0.26604 | 0.31646 | 0.09844 | 13.19331 | 0.030174031 | 0 | Neutral |
| testneg_2956 | -0.09 | 1.07 | 6.083672 | -3 | 1.729486 | -3 | 0.71697 | 0.19162 | 0.26068 | 0.46457 | 12.29167 | 0.5955 | 82.8 | Disease_causing |
| testneg_2957 | -0.01 | -0.79 | 3.736149 | -3 | -0.009242 | -3 | 0.4404 | 0.38303 | 0.11945 | 0.10135 | 22.79111 | 0.12 | 0 | Neutral |
| testneg_2958 | -0.18 | -0.16 | -0.566382 | -3 | 1.290195 | 1.290195 | 0.23152 | 0.09962 | 0.10761 | 0.12246 | 21.25537 | 0.06 | 0 | Neutral |
| testneg_2959 | 0.03 | 1.22 | 5.545455 | -3 | 3.557638 | -3 | 0.10571 | 0.12788 | 0.15867 | 0.09641 | 24.39852 | 0.28 | 1.6 | Neutral |
| testneg_2960 | 0 | 0 | 0.132481 | -3 | 5.047397 | 0.726355 | 0.33026 | 0.30576 | 0.09439 | 0.1884 | 12.70833 | 0.419067318 | 25.4 | Neutral |
| testneg_2961 | 0 | 0 | -0.484253 | -3 | -0.355328 | -3 | 0.12836 | 0.09736 | 0.26789 | 0.67432 | 10.83333 | 0.042081597 | 0 | Neutral |
| testneg_2962 | 0 | 0 | 0.287181 | -3 | -0.241891 | -3 | 0.93834 | 0.11088 | 0.13873 | 0.12788 | 23.35718 | 0.02 | 0 | Neutral |
| testneg_2963 | 0 | 0 | 1.426245 | 1.320713 | 5.135383 | -3 | 0.12643 | 0.12266 | 0.12682 | 0.11357 | 12.96607 | 0.111826331 | 0 | Neutral |
| testneg_2964 | 0.07 | 0.01 | 0.631605 | -3 | 1.712042 | -3 | 0.14524 | 0.10162 | 0.31004 | 0.09727 | 18.88889 | 0.001143791 | 0 | Neutral |
| testneg_2965 | 0.14 | -0.37 | 3.287017 | -3 | 0.496531 | -3 | 0.1594 | 0.22002 | 0.82148 | 0.14847 | 16.8018 | 0.190034965 | 0 | Neutral |
| testneg_2966 | 0.14 | -0.69 | 1.524427 | -3 | -0.483923 | -3 | 0.99317 | 0.60884 | 0.23574 | 0.12788 | 10.22898 | 0.07 | 0 | Neutral |
| testneg_2967 | 0 | -0.01 | 0.217304 | -2.734447 | -0.362155 | -3 | 0.48023 | 0.12788 | 0.19898 | 0.06796 | 19.84127 | 0.00050817 | 0 | Neutral |
| testneg_2968 | 0.18 | -1.05 | 7.345624 | -3 | 5.35839 | -3 | 0.06366 | 0.12788 | 0.93035 | 0.31722 | 5.74956 | 0.71 | 99.2 | Disease_causing |
| testneg_2969 | 0 | 0 | -0.416365 | -3 | 6.394805 | 6.394805 | 0.06793 | 0.12788 | 0.10285 | 0.10186 | 20 | 0.44 | 28.2 | Neutral |
| testneg_2970 | 0 | -0.32 | 1.773334 | -3 | 4.539303 | -3 | 0.30311 | 0.11121 | 0.62808 | 0.25222 | 10.7989 | 0.260539414 | 1.6 | Neutral |
| testneg_2971 | 0 | 0 | 1.108894 | -3 | 0.553738 | -3 | 0.30942 | 0.13105 | 0.11827 | 0.23034 | 10.41667 | 0.039628064 | 0 | Neutral |
| testneg_2972 | 0 | 0 | -0.566651 | -3 | 0.47696 | -3 | 0.38016 | 0.14503 | 0.19898 | 0.25991 | 9.67871 | 0.031884821 | 0 | Neutral |
| testneg_2973 | 0.04 | -1.36 | 2.629126 | -3 | 1.457871 | -3 | 0.0984 | 0.1489 | 0.08837 | 0.09327 | 19.44444 | 0.079321429 | 0 | Neutral |
| testneg_2974 | 0.18 | -1.05 | 5.200236 | -3 | 2.902146 | -3 | 0.59871 | 0.34719 | 0.14941 | 0.0838 | 26.66667 | 0.320333333 | 0.6 | Neutral |
| testneg_2975 | 0 | 0 | 2.317085 | -3 | 1.867073 | -3 | 0.35412 | 0.10819 | 0.07999 | 0.23927 | 10.41667 | 0.06 | 0 | Neutral |
| testneg_2976 | 0 | 0 | 0.687946 | -3 | 4.998465 | 4.998465 | 0.09269 | 0.12788 | 0.11233 | 0.28224 | 4.72222 | 0.466257813 | 42.4 | Neutral |
| testneg_2977 | 0.01 | 0.8 | 4.81138 | -3 | 5.035077 | -3 | 0.284 | 0.15339 | 0.34109 | 0.2167 | 12.40248 | 0.770087719 | 99.6 | Disease_causing |
| testneg_2978 | 0 | 0 | 1.28208 | -3 | 1.125228 | -3 | 0.38375 | 0.09256 | 0.15808 | 0.14548 | 20.83333 | 0.040975415 | 0 | Neutral |
| testneg_2979 | -0.07 | -0.38 | 2.626972 | -3 | 2.296233 | -3 | 0.47775 | 0.10103 | 0.12508 | 0.231 | 20.55556 | 0.173133882 | 0 | Neutral |
| testneg_2980 | 0.14 | -0.69 | 4.135613 | -3 | 3.353608 | -3 | 0.62172 | 0.18135 | 0.19821 | 0.20954 | 18.88889 | 0.689748111 | 97.6 | Disease_causing |
| testneg_2981 | 0.03 | 0.1 | -1.156564 | -3 | 5.704027 | -3 | 0.54213 | 0.09044 | 0.09504 | 0.38939 | 22.63149 | 0.353806217 | 6.2 | Neutral |
| testneg_2982 | 0.00872727 | -0.0578182 | 8.801009 | 1.79061 | 0.617218 | -3 | 0.12112 | 0.11353 | 0.10458 | 0.10499 | 33.48036 | 0.42875 | 24.8 | Neutral |
| testneg_2983 | 0 | 0 | 0.588209 | -3 | 1.6835 | 1.091417 | 0.07431 | 0.11513 | 0.06617 | 0.12788 | 30.40992 | 0.010611111 | 0 | Neutral |
| testneg_2984 | 0.03 | 0.12 | 2.135113 | -3 | -0.608116 | -3 | 0.40391 | 0.13431 | 0.0925 | 0.08222 | 10.41667 | 0.010555556 | 0 | Neutral |
| testneg_2985 | 0.18 | -1.05 | 6.85485 | -3 | 0.616227 | -3 | 0.1413 | 0.74044 | 0.02971 | 0.33983 | 20.29919 | 0.58 | 85 | Disease_causing |
| testneg_2986 | 0 | -0.01 | 1.006749 | -3 | 0.728537 | -3 | 0.7003 | 0.12788 | 0.44541 | 0.13979 | 10.9543 | 0.025725 | 0 | Neutral |
| testneg_2987 | 0 | 0.73 | 1.370152 | -3 | -0.457516 | -3 | 0.10486 | 0.08083 | 0.01635 | 0.12788 | 13.51351 | 0.02 | 0 | Neutral |
| testneg_2988 | -0.18 | -0.16 | 2.72063 | 2.72063 | 5.095304 | -3 | 0.65141 | 0.18046 | 0.18559 | 0.13755 | 5.69444 | 0.71617029 | 97.2 | Disease_causing |
| testneg_2989 | 0.07 | 0.7 | 3.89415 | -3 | 5.544421 | -0.144535 | 0.16671 | 0.18311 | 0.39932 | 0.10693 | 18.92511 | 0.394883769 | 17.8 | Neutral |
| testneg_2990 | 0.05 | -0.81 | -0.765143 | -0.765143 | 6.195353 | -3 | 0.31627 | 0.23288 | 0.28532 | 0.10663 | 10.22898 | 0.23027027 | 0.4 | Neutral |
| testneg_2991 | 0.14 | -0.37 | 0.334634 | -3 | 0.278757 | -3 | 0.18925 | 0.32479 | 0.47816 | 0.66526 | 10 | 0.191039427 | 0 | Neutral |
| testneg_2992 | 0.18 | -1.05 | 7.618716 | -3 | 1.531136 | -3 | 0.38445 | 0.58278 | 0.85887 | 0.1489 | 10 | 0.550272947 | 75.6 | Disease_causing |
| testneg_2993 | 0 | -0.01 | 0.637461 | -3 | 1.485335 | 1.062107 | 0.15556 | 0.12841 | 0.19894 | 0.27411 | 13.36964 | 0.050568733 | 0 | Neutral |
| testneg_2994 | 0 | -0.32 | 2.333735 | -3 | 3.118033 | -3 | 0.08224 | 0.29179 | 0.19721 | 0.12485 | 10 | 0.121296541 | 0 | Neutral |
| testneg_2995 | 0 | -0.32 | -1.725964 | -3 | 0.71227 | -3 | 0.13203 | 0.12092 | 0.86437 | 0.69972 | 10.83333 | 0.058541667 | 0 | Neutral |
| testneg_2996 | 0 | 0 | 1.309085 | -3 | 8.109614 | -3 | 0.64125 | 0.28792 | 0.25896 | 0.45481 | 11.55672 | 0.516923077 | 61.6 | Disease_causing |
| testneg_2997 | 0.11 | -0.19 | 3.47162 | -3 | -1.246222 | -3 | 0.60703 | 0.12378 | 0.80264 | 0.11567 | 18.79036 | 0.04 | 0 | Neutral |
| testneg_2998 | 0 | 0 | -0.067945 | -3 | -0.518868 | -3 | 0.02844 | 0.12788 | 0.08386 | 0.12788 | 17.21847 | 0.03 | 0 | Neutral |
| testneg_2999 | 0 | 0 | 0.300937 | -3 | 0.285978 | -3 | 0.46318 | 0.14229 | 0.19898 | 0.09824 | 17.12264 | 0.01 | 0 | Neutral |
| testneg_3000 | 0.21 | 0.15 | 0.783216 | -3 | 5.884055 | -3 | 0.33026 | 0.30576 | 0.11042 | 0.09881 | 21.53664 | 0.27075 | 0.6 | Neutral |
| testneg_3001 | -0.08 | -0.03 | 1.513879 | 0.606649 | 1.01759 | -3 | 0.62199 | 0.14032 | 0.0406 | 0.12788 | 21.41414 | 0.05 | 0 | Neutral |
| testneg_3002 | 0 | 0 | 1.445184 | 1.445184 | 4.311522 | -3 | 0.33469 | 0.17432 | 0.08857 | 0.83499 | 10.41667 | 0.454305785 | 34.8 | Neutral |
| testneg_3003 | 0.03 | -1.03 | 0.221 | -3 | 6.526232 | -3 | 0.37637 | 0.12788 | 0.2016 | 0.13956 | 5 | 0.372511905 | 12.6 | Neutral |
| testneg_3004 | -0.07 | -0.38 | -1.803731 | -3 | 0.6695 | -3 | 0.07963 | 0.23713 | 0.06517 | 0.15107 | 20.83333 | 0.18678714 | 0 | Neutral |
| testneg_3005 | -0.05 | 0.81 | 4.665144 | -3 | -3.203793 | -3 | 0.30122 | 0.09165 | 0.57349 | 0.09764 | 5.69444 | 0.1 | 0 | Neutral |
| testneg_3006 | 0 | 0 | -0.179197 | -3 | -0.116624 | -3 | 0.86835 | 0.1513 | 0.16462 | 0.18467 | 10.41667 | 0.041721003 | 0 | Neutral |
| testneg_3007 | 0 | 0 | 0.484587 | -3 | -2.091153 | -3 | 0.30056 | 0.12272 | 0.69696 | 0.10507 | 28.50172 | 0.01 | 0 | Neutral |
| testneg_3008 | 0.09 | -0.3 | 4.263993 | -3 | 1.069398 | -3 | 0.25764 | 0.53109 | 0.06431 | 0.12788 | 13.08333 | 0.127057387 | 0 | Neutral |
| testneg_3009 | -0.05 | 0.81 | 0.534593 | -3 | 0.162648 | -3 | 0.44526 | 0.09404 | 0.38002 | 0.22508 | 22.25519 | 0.071242781 | 0 | Neutral |
| testneg_3010 | 0 | 0 | 1.200491 | -3 | -0.718609 | -3 | 0.34198 | 0.1113 | 0.18371 | 0.10748 | 18.452 | 0 | 0 | Neutral |
| testneg_3011 | 0 | 0 | 2.013802 | -3 | 0.581883 | -3 | 0.22239 | 0.12885 | 0.32144 | 0.11774 | 18.88889 | 0.000148717 | 0 | Neutral |
| testneg_3012 | 0.05 | -0.81 | 3.12205 | -3 | 1.327761 | 1.327761 | 0.34633 | 0.47228 | 0.19898 | 0.19048 | 16.8018 | 0.366673195 | 10.8 | Neutral |
| testneg_3013 | 0.04 | -0.23 | 0.982196 | -3 | 0.192939 | -3 | 0.19898 | 0.09322 | 0.17725 | 0.12788 | 32.69082 | 0.01 | 0 | Neutral |
| testneg_3014 | -0.03 | -0.1 | 6.036832 | -3 | 2.177624 | -3 | 0.16826 | 0.11262 | 0.19898 | 0.12788 | 19.43262 | 0.220916259 | 0.4 | Neutral |
| testneg_3015 | 0.21 | 0.15 | 5.918263 | -3 | 1.613091 | 1.613091 | 0.25421 | 0.16235 | 0.53922 | 0.12788 | 28.47222 | 0.244166667 | 0 | Neutral |
| testneg_3016 | 0.22 | -2.66 | 3.002533 | -3 | 2.655425 | -3 | 0.14304 | 0.10715 | 0.19883 | 0.09434 | 41.32166 | 0.12 | 0 | Neutral |
| testneg_3017 | 0.01 | 0.8 | 0.038109 | -3 | -1.898366 | -1.898366 | 0.12497 | 0.09805 | 0.11983 | 0.24162 | 33.08485 | 0.086730769 | 0 | Neutral |
| testneg_3018 | -0.002 | -0.037 | 9.170362 | -3 | 3.850429 | 1.634389 | 0.1144 | 0.12788 | 0.23939 | 0.25284 | 22.95229 | 0.694222222 | 98.6 | Disease_causing |
| testneg_3019 | -0.17 | -0.38 | 1.89302 | -3 | -0.237647 | -3 | 0.16475 | 0.26069 | 0.16116 | 0.08314 | 15.94021 | 0.08 | 0 | Neutral |
| testneg_3020 | 0.04 | -0.23 | 4.657768 | -3 | -0.089221 | -3 | 0.22645 | 0.1704 | 0.10277 | 0.12788 | 13.05427 | 0.071064068 | 0 | Neutral |
| testneg_3021 | 0 | 0 | 0.645288 | -3 | 0.559242 | -3 | 0.72657 | 0.11262 | 0.87803 | 0.12788 | 13.86082 | 0.01 | 0 | Neutral |
| testneg_3022 | -0.22 | 2.66 | 5.679638 | -3 | 11.578151 | -3 | 0.61834 | 0.10784 | 0.16485 | 0.13045 | 35.2867 | 0.63 | 93.2 | Disease_causing |
| testneg_3023 | 0 | 0 | 1.825331 | -3 | 4.004011 | -3 | 0.13029 | 0.12788 | 0.37736 | 0.11686 | 8.97898 | 0.050836247 | 0 | Neutral |
| testneg_3024 | 0.04 | -0.23 | 1.620184 | -3 | 0.619175 | 0.619175 | 0.20502 | 0.10447 | 0.68138 | 0.20944 | 29.72678 | 0.07 | 0 | Neutral |
| testneg_3025 | 0 | 0 | 1.39998 | -3 | -2.137796 | -3 | 0.161 | 0.11597 | 0.13698 | 0.12788 | 13.51351 | 0 | 0 | Neutral |
| testneg_3026 | 0 | 0 | 0.532295 | -3 | 6.623383 | -3 | 0.755 | 0.12788 | 0.29172 | 0.08143 | 19.09347 | 0.112094643 | 0 | Neutral |
| testneg_3027 | 0.04 | -1.61 | 4.284903 | -3 | 7.338844 | -3 | 0.3828 | 0.5569 | 0.13433 | 0.09578 | 19.90385 | 0.589272727 | 84.2 | Disease_causing |
| testneg_3028 | 0.01545455 | -0.092 | 6.776197 | -3 | 1.186176 | -3 | 0.7219 | 0.41509 | 0.19898 | 0.12788 | 10.93254 | 0.445871212 | 32.2 | Neutral |
| testneg_3029 | 0.01 | 0.8 | 4.81138 | -3 | -0.676178 | -3 | 0.284 | 0.15339 | 0.19898 | 0.08855 | 11.98582 | 0.08 | 0 | Neutral |
| testneg_3030 | 0.02 | -0.63 | 1.006549 | -3 | 3.876605 | -3 | 0.12591 | 0.09845 | 0.83732 | 0.19325 | 27.4024 | 0.23441881 | 0 | Neutral |
| testneg_3031 | 0 | 0 | 0.602803 | -3 | 5.00838 | -3 | 0.12259 | 0.1041 | 0.5043 | 0.3225 | 12.35574 | 0.232489274 | 0.6 | Neutral |
| testneg_3032 | -0.05 | -0.29 | -0.803439 | -3 | 1.014441 | -3 | 0.50276 | 0.09638 | 0.1784 | 0.12788 | 15.97222 | 0.010557843 | 0 | Neutral |
| testneg_3033 | 0.21 | 0.15 | 1.704228 | -3 | -1.047288 | -3 | 0.27062 | 0.12378 | 0.52005 | 0.12788 | 23.40643 | 0.022916667 | 0 | Neutral |
| testneg_3034 | -0.04 | 0.23 | 0.6838 | 0.6838 | 4.373261 | -3 | 0.67327 | 0.23543 | 0.29356 | 0.47581 | 18.88889 | 0.50016353 | 53.2 | Disease_causing |
| testneg_3035 | 0.17 | 0.38 | 3.132136 | -3 | 1.139058 | -3 | 0.0105 | 0.12788 | 0.27887 | 0.18663 | 20.11942 | 0.227857143 | 0.2 | Neutral |
| testneg_3036 | -0.09 | 0.3 | 4.962635 | -3 | -1.124893 | -3 | 0.84596 | 0.10633 | 0.07923 | 0.12788 | 23.78988 | 0.1 | 0 | Neutral |
| testneg_3037 | 0 | 0 | 1.445184 | 1.445184 | 5.938598 | -3 | 0.33469 | 0.17432 | 0.10835 | 0.12788 | 18.33333 | 0.223565005 | 0 | Neutral |
| testneg_3038 | 0 | 0 | 1.941287 | -3 | 3.82965 | -3 | 0.24977 | 0.11262 | 0.11957 | 0.08975 | 12.20857 | 0.048841392 | 0 | Neutral |
| testneg_3039 | 0.04 | 2.01 | 4.400914 | -3 | 6.097788 | -3 | 0.10161 | 0.13484 | 0.22944 | 0.09039 | 20.83333 | 0.350607143 | 9.6 | Neutral |
| testneg_3040 | 0 | 0.32 | 3.268811 | -3 | 1.942918 | -3 | 0.17141 | 0.1104 | 0.11187 | 0.10744 | 28.10041 | 0.03 | 0 | Neutral |
| testneg_3041 | 0 | 0 | 0.306558 | -3 | 3.465539 | 1.154108 | 0.08999 | 0.0989 | 0.13139 | 0.11068 | 11.25 | 0.045864719 | 0 | Neutral |
| testneg_3042 | 0.18 | -1.05 | 2.686668 | -3 | 1.186074 | -3 | 0.10517 | 0.09803 | 0.15341 | 0.12272 | 10 | 0.03 | 0 | Neutral |
| testneg_3043 | 0 | 0 | 0.103447 | 0.103447 | 7.50868 | -3 | 0.15813 | 0.61197 | 0.40283 | 0.34309 | 18.71842 | 0.680138889 | 96 | Disease_causing |
| testneg_3044 | -0.08 | -0.03 | 0.20058 | -3 | 0.088378 | -3 | 0.1897 | 0.12459 | 0.66693 | 0.0986 | 9.862 | 0.010200163 | 0 | Neutral |
| testneg_3045 | 0.02 | -0.63 | 0.269085 | -3 | 2.521061 | -3 | 0.09281 | 0.13929 | 0.19898 | 0.11691 | 12.07444 | 0.031009615 | 0 | Neutral |
| testneg_3046 | -0.18 | -0.16 | 3.786992 | -3 | 5.524048 | -3 | 0.22872 | 0.15588 | 0.6185 | 0.09316 | 18.15261 | 0.385821078 | 14 | Neutral |
| testneg_3047 | 0.07 | -0.31 | 2.407105 | -3 | 13.409328 | 3.422775 | 0.21095 | 0.10574 | 0.24966 | 0.12788 | 18.57511 | 0.49 | 50.6 | Disease_causing |
| testneg_3048 | 0.04 | 2.01 | 6.434147 | -3 | 1.887787 | -3 | 0.13275 | 0.12788 | 0.51398 | 0.38821 | 4.64941 | 0.508333333 | 57.2 | Disease_causing |
| testneg_3049 | 0 | 0 | 2.33001 | -3 | 0.083768 | -3 | 0.71463 | 0.29732 | 0.08041 | 0.21802 | 17.47657 | 0.099099747 | 0 | Neutral |

| | | | | | | | | | | | | | | |
|---|---|---|---|---|---|---|---|---|---|---|---|---|---|---|
| testneg_3050 | -0.08 | -0.03 | 3.256149 | -3 | 0.664179 | -3 | 0.53313 | 0.14072 | 0.2124 | 0.17288 | 31.0441 | 0.18036715 | 0 | Neutral |
| testneg_3051 | -0.03 | -0.1 | 4.076957 | 4.076957 | -1.599196 | -3 | 0.38324 | 0.25193 | 0.37711 | 0.12788 | 10.22898 | 0.488257576 | 49.8 | Neutral |
| testneg_3052 | -0.03 | 0.85 | -0.309243 | -3 | 4.281142 | -3 | 0.03137 | 0.12788 | 0.46244 | 0.56748 | 28.54167 | 0.348345142 | 7 | Neutral |
| testneg_3053 | 0.02 | -0.68 | 1.430763 | -3 | 3.494755 | -3 | 0.2578 | 0.09796 | 0.21201 | 0.20962 | 13.89684 | 0.194181469 | 0 | Neutral |
| testneg_3054 | 0 | 0 | 1.283931 | -3 | 0.34432 | -3 | 0.03499 | 0.12971 | 0.52957 | 0.18687 | 29.16667 | 0.101142241 | 0 | Neutral |
| testneg_3055 | 0 | 0 | 0.058546 | -3 | -1.295182 | -3 | 0.18998 | 0.26095 | 0.26913 | 0.1087 | 12.29167 | 0.01 | 0 | Neutral |
| testneg_3056 | -0.07 | -0.38 | -1.066068 | -3 | 2.75746 | -3 | 0.04515 | 0.14318 | 0.16053 | 0.15049 | 20.90842 | 0.238027778 | 0.4 | Neutral |
| testneg_3057 | 0.18 | -1.05 | 4.095375 | -3 | 0.774689 | 0.774689 | 0.1589 | 0.10761 | 0.24219 | 0.17458 | 13.81407 | 0.205445887 | 0 | Neutral |
| testneg_3058 | -0.08 | -0.03 | 4.08127 | -3 | 2.555217 | -3 | 0.16073 | 0.11262 | 0.29672 | 0.10485 | 19.51014 | 0.074948331 | 0 | Neutral |
| testneg_3059 | -0.06 | 0.57 | 1.295804 | -3 | 0.22996 | -3 | 0.64795 | 0.21581 | 0.0693 | 0.09293 | 10.41667 | 0.04 | 0 | Neutral |
| testneg_3060 | 0.03 | 1.22 | 6.224476 | -3 | -0.111883 | -0.111883 | 0.7074 | 0.12788 | 0.06313 | 0.12788 | 15.61022 | 0.213928571 | 0.4 | Neutral |
| testneg_3061 | 0 | 0 | -1.743199 | -3 | 1.802906 | -3 | 0.12424 | 0.09575 | 0.33886 | 0.15644 | 20 | 0.05 | 0 | Neutral |
| testneg_3062 | 0.05 | 0.29 | 2.040423 | -3 | 4.492963 | -3 | 0.835 | 0.13588 | 0.54648 | 0.19042 | 5.60741 | 0.326531664 | 3.8 | Neutral |
| testneg_3063 | -0.17 | -0.38 | 1.808824 | 1.808824 | -1.110728 | -3 | 0.2445 | 0.53069 | 0.28817 | 0.10531 | 13.87828 | 0.14 | 0 | Neutral |
| testneg_3064 | 0.07 | 0.7 | 3.310081 | -3 | 5.493427 | -3 | 0.92776 | 0.09677 | 0.14 | 0.12151 | 17.31406 | 0.229677419 | 0.4 | Neutral |
| testneg_3065 | 0.21 | 0.15 | 2.442902 | -3 | 3.238247 | 3.238247 | 0.19898 | 0.12788 | 0.19898 | 0.12788 | 39.50869 | 0.2 | 0.2 | Neutral |
| testneg_3066 | -0.02 | 0.63 | 5.78606 | -3 | -0.433367 | -3 | 0.89972 | 0.10766 | 0.13468 | 0.09235 | 41.54512 | 0.19 | 0.2 | Neutral |
| testneg_3067 | -0.14 | 0.37 | 2.017569 | -3 | 5.321508 | -3 | 0.69563 | 0.12788 | 0.06215 | 0.08575 | 10.83333 | 0.19 | 0 | Neutral |
| testneg_3068 | 0 | 0 | 0.376038 | -3 | 2.478871 | -3 | 0.18485 | 0.40044 | 0.04188 | 0.12788 | 22.85054 | 0.075605842 | 0 | Neutral |
| testneg_3069 | 0.09 | -0.3 | 1.682255 | -0.005662 | -0.887241 | -3 | 0.1664 | 0.10113 | 0.19898 | 0.12788 | 29.34685 | 0.01 | 0 | Neutral |
| testneg_3070 | 0 | 0 | -0.68826 | -3 | -1.258796 | -1.258796 | 0.32076 | 0.14229 | 0.7055 | 0.11261 | 20.8447 | 0.020121969 | 0 | Neutral |
| testneg_3071 | 0 | 0 | 0.795894 | -3 | 6.530718 | -3 | 0.99448 | 0.67288 | 0.09288 | 0.16617 | 16.8018 | 0.449065762 | 34.6 | Neutral |
| testneg_3072 | -0.06 | -0.18 | 1.657765 | -3 | 2.944773 | -3 | 0.31923 | 0.17492 | 0.42666 | 0.12788 | 10.21277 | 0.061626576 | 0 | Neutral |
| testneg_3073 | 0.01 | 0.79 | 0.470746 | -3 | 1.005948 | -3 | 0.99543 | 0.12788 | 0.12176 | 0.09356 | 13.15918 | 0.05 | 0 | Neutral |
| testneg_3074 | -0.03 | -0.1 | 1.738251 | -3 | 1.340083 | -3 | 0.17303 | 0.18733 | 0.28227 | 0.10211 | 11.875 | 0.03 | 0 | Neutral |
| testneg_3075 | 0.14 | -1.01 | 3.621071 | 2.625488 | 0.66227 | -3 | 0.87803 | 0.12788 | 0.23141 | 0.12788 | 12.46106 | 0.2825 | 4.4 | Neutral |
| testneg_3076 | 0 | 0 | 1.095746 | -3 | -0.720759 | -0.720759 | 0.24274 | 0.13034 | 0.17758 | 0.15983 | 13.02635 | 0.060253968 | 0 | Neutral |
| testneg_3077 | 0.04 | -0.23 | -0.122196 | -0.122196 | 0.386087 | 0.386087 | 0.31961 | 0.17571 | 0.26927 | 0.10752 | 14.95396 | 0.020159568 | 0 | Neutral |
| testneg_3078 | 0 | -0.32 | 1.002173 | -3 | 0.359803 | 0.359803 | 0.27308 | 0.12788 | 0.01635 | 0.12788 | 19.14829 | 0.010229915 | 0 | Neutral |
| testneg_3079 | 0 | 0 | -0.034419 | -0.034419 | 2.411098 | -3 | 0.62203 | 0.12788 | 0.2521 | 0.1784 | 23.24165 | 0.147996032 | 0 | Neutral |
| testneg_3080 | 0.15 | 0.04 | -1.177878 | -3 | 0.997782 | -3 | 0.19301 | 0.12955 | 0.48817 | 0.10601 | 20.5518 | 0.02 | 0 | Neutral |
| testneg_3081 | 0.07 | 0.38 | -0.473879 | -3 | 3.235571 | 3.235571 | 0.0695 | 0.12788 | 0.4069 | 0.3841 | 26.34995 | 0.361847813 | 8.4 | Neutral |
| testneg_3082 | 0.00055944 | 0.00363636 | 1.360315 | -3 | 2.311448 | -3 | 0.29482 | 0.15726 | 0.1231 | 0.3662 | 70.67478 | 0.31498637 | 1 | Neutral |
| testneg_3083 | 0.18 | -1.05 | 3.535138 | -3 | 3.522487 | -3 | 0.7062 | 0.83002 | 0.2755 | 0.12788 | 10.22898 | 0.352705687 | 9.6 | Neutral |
| testneg_3084 | 0 | 0 | 4.336736 | -3 | -0.368497 | -3 | 0.27275 | 0.11752 | 0.12832 | 0.07113 | 29.86111 | 0.021259615 | 0 | Neutral |
| testneg_3085 | 0 | 0 | 5.435713 | -3 | 2.403686 | 2.403686 | 0.24026 | 0.12788 | 0.48574 | 0.20473 | 13.51351 | 0.549843003 | 72.8 | Disease_causing |
| testneg_3086 | 0 | 0 | 0.383044 | -3 | 4.744051 | -3 | 0.63819 | 0.32258 | 0.19052 | 0.12788 | 13.7295 | 0.090936553 | 0 | Neutral |
| testneg_3087 | 0.14 | -1.01 | 2.399127 | -3 | 0.903176 | -3 | 0.22255 | 0.10057 | 0.10835 | 0.12788 | 11.87076 | 0.01 | 0 | Neutral |
| testneg_3088 | 0.21 | 0.15 | -1.085568 | -3 | -0.115386 | -3 | 0.22901 | 0.12788 | 0.15016 | 0.08826 | 20.98109 | 0.03 | 0 | Neutral |
| testneg_3089 | 0.14 | -0.69 | 4.331704 | -3 | 0.429708 | -3 | 0.97504 | 0.18545 | 0.19898 | 0.12788 | 20.27778 | 0.099468326 | 0 | Neutral |
| testneg_3090 | 0.03 | 0.12 | 0.565832 | -3 | 3.740739 | -3 | 0.1105 | 0.44945 | 0.26337 | 0.12771 | 10.83333 | 0.200230399 | 0 | Neutral |
| testneg_3091 | 0.14 | -0.37 | 6.395459 | -3 | -0.70331 | -3 | 0.2016 | 0.13956 | 0.47187 | 0.14227 | 30.014 | 0.310384058 | 6 | Neutral |
| testneg_3092 | 0 | -0.01 | 6.857724 | -3 | 1.69701 | -3 | 0.18224 | 0.10419 | 0.27197 | 0.12346 | 18.33333 | 0.259566666 | 2.6 | Neutral |
| testneg_3093 | 0.07 | 0.38 | 1.47114 | -3 | 0.046701 | 0.046701 | 0.65564 | 0.11468 | 0.06611 | 0.08834 | 22.77778 | 0.03 | 0 | Neutral |
| testneg_3094 | -0.01 | -0.8 | 4.805827 | -3 | 0.647256 | 0.647256 | 0.3693 | 0.12788 | 0.68436 | 0.10157 | 10.41667 | 0.08 | 0 | Neutral |
| testneg_3095 | 0 | 0 | -0.064795 | -3 | 5.243459 | -3 | 0.4566 | 0.14201 | 0.20918 | 0.10825 | 18.88889 | 0.081214547 | 0 | Neutral |
| testneg_3096 | 0.02 | 0.41 | 6.514964 | -3 | 5.969277 | -3 | 0.10895 | 0.10442 | 0.31067 | 0.35414 | 5.73716 | 0.830803396 | 100 | Disease_causing |
| testneg_3097 | 0.03 | 0.12 | 5.43953 | -3 | 0.025863 | -3 | 0.43707 | 0.10923 | 0.21755 | 0.23824 | 56.87727 | 0.32 | 4.4 | Neutral |
| testneg_3098 | 0 | 0 | 5.245361 | -3 | -0.614161 | -3 | 0.05216 | 0.23851 | 0.17796 | 0.09522 | 18.88889 | 0.23 | 0 | Neutral |
| testneg_3099 | -0.07 | -0.33 | 0.406963 | -3 | 5.007683 | -3 | 0.48825 | 0.10951 | 0.07415 | 0.12788 | 27.81922 | 0.111325443 | 0 | Neutral |
| testneg_3100 | -0.08 | -0.03 | -0.057252 | -3 | 6.001623 | -3 | 0.83176 | 0.23122 | 0.19898 | 0.12788 | 11.9773 | 0.218863636 | 0.2 | Neutral |
| testneg_3101 | 0.04 | 2.01 | -3.135825 | -3 | 2.210978 | -3 | 0.35422 | 0.11185 | 0.90308 | 0.28373 | 28.28527 | 0.172193342 | 0 | Neutral |
| testneg_3102 | 0.02 | 0.13 | 4.399727 | -3 | 0.099305 | -3 | 0.11373 | 0.10739 | 0.80889 | 0.40016 | 10.83333 | 0.17 | 0 | Neutral |
| testneg_3103 | 0 | 0 | 1.119143 | -3 | -1.327597 | -3 | 0.13482 | 0.12788 | 0.16155 | 0.20638 | 19.09347 | 0.04 | 0 | Neutral |
| testneg_3104 | 0.16 | -1.37 | 5.382361 | -3 | 0.307836 | -3 | 0.19898 | 0.12788 | 0.88976 | 0.22648 | 24.44444 | 0.29 | 2.6 | Neutral |
| testneg_3105 | -0.01 | -0.79 | 2.573684 | -3 | 2.20128 | -3 | 0.77196 | 0.14089 | 0.19898 | 0.12788 | 15.7977 | 0.04 | 0 | Neutral |
| testneg_3106 | -0.01 | 0.57 | 2.185335 | -3 | 4.028512 | -3 | 0.19898 | 0.12788 | 0.14954 | 0.12788 | 29.21954 | 0.10216129 | 0 | Neutral |
| testneg_3107 | 0 | 0 | -0.491414 | -3 | 3.498501 | -3 | 0.61027 | 0.14229 | 0.80314 | 0.42344 | 23.90474 | 0.264542749 | 0.4 | Neutral |
| testneg_3108 | 0.00233333 | -0.0513333 | 3.565578 | -3 | -0.334799 | -3 | 0.99689 | 0.28104 | 0.49665 | 0.20425 | 10 | 0.250319795 | 0.8 | Neutral |
| testneg_3109 | 0 | 0 | 2.022849 | 2.022849 | 5.544421 | -3 | 0.20708 | 0.12788 | 0.39932 | 0.10693 | 27.17793 | 0.190798368 | 0 | Neutral |
| testneg_3110 | 0.04 | -0.23 | 5.518112 | -3 | 1.247019 | -3 | 0.32977 | 0.15427 | 0.88456 | 0.11674 | 16.07589 | 0.19 | 0 | Neutral |
| testneg_3111 | -0.07 | -0.38 | -0.094721 | -3 | 2.058739 | -3 | 0.16778 | 0.82322 | 0.19898 | 0.12788 | 18.6768 | 0.1 | 0 | Neutral |
| testneg_3112 | 0.02 | 0.45 | 1.21068 | -3 | 0.634119 | -3 | 0.95247 | 0.11285 | 0.41647 | 0.11239 | 19.51014 | 0.02 | 0 | Neutral |
| testneg_3113 | 0.1 | 1.9 | 1.410622 | -3 | 5.522243 | -3 | 0.44479 | 0.20004 | 0.12855 | 0.11656 | 19.44444 | 0.303170996 | 1 | Neutral |
| testneg_3114 | 0 | 0 | -0.048245 | -3 | 0.101783 | -3 | 0.15055 | 0.22097 | 0.06196 | 0.10881 | 20.83333 | 0.031026316 | 0 | Neutral |
| testneg_3115 | 0.11 | -1.95 | 4.99961 | -3 | 5.997238 | -3 | 0.19452 | 0.79779 | 0.44479 | 0.20004 | 10.17293 | 0.892009832 | 100 | Disease_causing |
| testneg_3116 | -0.03 | -1.22 | 4.490043 | -3 | 6.264472 | -3 | 0.7342 | 0.11023 | 0.71276 | 0.88182 | 14.09268 | 0.607772043 | 88.6 | Disease_causing |
| testneg_3117 | 0 | -0.01 | 7.652567 | 7.652567 | 4.839828 | -3 | 0.73253 | 0.09747 | 0.05095 | 0.08353 | 13.51351 | 0.69 | 98.8 | Disease_causing |
| testneg_3118 | 0.03 | 0.12 | -0.231636 | -0.231636 | 0.367203 | -3 | 0.99765 | 0.95064 | 0.24832 | 0.21196 | 5.74956 | 0.152542437 | 0 | Neutral |
| testneg_3119 | 0 | 0 | 0.897808 | 0.897808 | 0.888376 | -3 | 0.17155 | 0.18585 | 0.21908 | 0.12788 | 10 | 0.03 | 0 | Neutral |
| testneg_3120 | 0.22 | -2.66 | 6.342119 | -3 | 1.27985 | -3 | 0.76186 | 0.10052 | 0.09334 | 0.10027 | 17.62359 | 0.23 | 1 | Neutral |
| testneg_3121 | 0.22 | -2.66 | 3.257494 | -3 | 6.40665 | -3 | 0.09356 | 0.2448 | 0.29828 | 0.12788 | 14.30429 | 0.492678571 | 52.2 | Disease_causing |
| testneg_3122 | -0.06 | 0.57 | 0.172291 | -3 | 0.6576 | -3 | 0.0138 | 0.06769 | 0.12147 | 0.1486 | 20.42553 | 0.132992748 | 0 | Neutral |
| testneg_3123 | -0.19 | 0.25 | -1.729462 | -3 | 0.335227 | -3 | 0.16155 | 0.20638 | 0.04346 | 0.10785 | 18.33333 | 0.1 | 0 | Neutral |
| testneg_3124 | 0.21 | 0.15 | 1.623618 | -3 | -1.140892 | -3 | 0.10083 | 0.17511 | 0.45343 | 0.12014 | 18.88889 | 0.06 | 0 | Neutral |
| testneg_3125 | 0.19 | 0.51 | 5.002476 | -3 | 5.03044 | -3 | 0.22162 | 0.09445 | 0.10561 | 0.12788 | 21.14842 | 0.333496377 | 7.2 | Neutral |
| testneg_3126 | 0.03 | -1.03 | -0.674759 | -0.674759 | 6.273159 | -3 | 0.02782 | 0.06218 | 0.39224 | 0.09664 | 12.70833 | 0.151200717 | 0 | Neutral |
| testneg_3127 | -0.07 | -0.01 | 0.450036 | -1.290749 | 2.341088 | -3 | 0.44479 | 0.20004 | 0.05564 | 0.06933 | 10.83333 | 0.06 | 0 | Neutral |
| testneg_3128 | -0.18 | -0.16 | 6.127805 | -3 | 0.218023 | -3 | 0.67549 | 0.1273 | 0.19898 | 0.12788 | 9.90291 | 0.233166667 | 0.8 | Neutral |
| testneg_3129 | 0 | 0 | 0.80714 | -3 | 1.952213 | -3 | 0.19898 | 0.12788 | 0.49504 | 0.08191 | 15.66882 | 0 | 0 | Neutral |
| testneg_3130 | 0 | 0 | 0.451796 | -3 | 1.437183 | -3 | 0.19898 | 0.12788 | 0.19898 | 0.12788 | 18.6768 | 0 | 0 | Neutral |
| testneg_3131 | 0 | 0 | -0.342253 | -3 | 2.325263 | -3 | 0.63363 | 0.14795 | 0.75315 | 0.12659 | 9.84463 | 0.03 | 0 | Neutral |
| testneg_3132 | -0.05 | -0.29 | 4.656948 | -3 | 0.03956 | -3 | 0.91587 | 0.10818 | 0.07961 | 0.11474 | 22.30665 | 0.06 | 0 | Neutral |
| testneg_3133 | 0 | 0 | 0.436295 | -3 | 5.907552 | -3 | 0.32677 | 0.12788 | 0.10438 | 0.20558 | 23.33333 | 0.389837766 | 13.2 | Neutral |
| testneg_3134 | 0 | 0 | 2.403686 | 2.403686 | 3.046218 | -3 | 0.48574 | 0.20473 | 0.16547 | 0.16092 | 16.13039 | 0.450506417 | 35 | Neutral |
| testneg_3135 | 0 | 0 | 0.535559 | -3 | 4.06036 | -3 | 0.34931 | 0.11906 | 0.06187 | 0.36607 | 10.41667 | 0.230103898 | 0.2 | Neutral |
| testneg_3136 | 0 | 0 | 0.47864 | -3 | 1.390942 | 1.390942 | 0.17597 | 0.12788 | 0.10751 | 0.08512 | 24.93393 | 0.02 | 0 | Neutral |
| testneg_3137 | 0.06 | -0.57 | 3.156075 | -3 | 0.219522 | -3 | 0.44164 | 0.13588 | 0.19898 | 0.09533 | 27.91246 | 0.0401547 | 0 | Neutral |
| testneg_3138 | 0 | 0 | 2.402629 | -3 | 0.60982 | 0.60982 | 0.19898 | 0.12788 | 0.23844 | 0.27772 | 23.51497 | 0.088724771 | 0 | Neutral |
| testneg_3139 | 0 | -0.01 | 0.558245 | -3 | -0.044091 | -3 | 0.19898 | 0.10362 | 0.07534 | 0.09292 | 11.56063 | 0.000110501 | 0 | Neutral |
| testneg_3140 | 0 | -0.32 | -1.488205 | -3 | 1.101796 | -3 | 0.08878 | 0.0974 | 0.01642 | 0.12788 | 18.44519 | 0.017048246 | 0 | Neutral |
| testneg_3141 | 0 | 0 | 0.080104 | -3 | 0.623338 | -3 | 0.73947 | 0.2058 | 0.09707 | 0.12788 | 41.77216 | 0.04 | 0 | Neutral |
| testneg_3142 | 0 | 0 | 1.079876 | -3 | 0.010179 | 0.010179 | 0.97252 | 0.8859 | 0.19898 | 0.12788 | 10.22898 | 0.070867254 | 0 | Neutral |
| testneg_3143 | -0.09 | 0.25 | 6.437271 | -3 | 7.490616 | 1.258706 | 0.1336 | 0.15669 | 0.85345 | 0.13067 | 19.44444 | 0.757811382 | 100 | Disease_causing |
| testneg_3144 | 0 | 0 | 1.269835 | -3 | -0.363052 | -3 | 0.94662 | 0.11262 | 0.3437 | 0.5169 | 4.72222 | 0.030495129 | 0 | Neutral |
| testneg_3145 | -0.07 | -0.38 | 2.459203 | -3 | 0.096543 | -3 | 0.23292 | 0.15913 | 0.28326 | 0.13944 | 10.83333 | 0.100296425 | 0 | Neutral |
| testneg_3146 | 0 | 0 | 0.902021 | -3 | -0.112406 | -3 | 0.37499 | 0.17803 | 0.94807 | 0.14977 | 4.44444 | 0.024098235 | 0 | Neutral |
| testneg_3147 | 0 | 0 | 1.967428 | -3 | 1.925465 | -3 | 0.42797 | 0.17974 | 0.21967 | 0.09467 | 16.88659 | 0.02 | 0 | Neutral |
| testneg_3148 | -0.01 | 1.44 | -1.970817 | -3 | 5.028164 | -3 | 0.2058 | 0.28174 | 0.29209 | 0.09791 | 10.36687 | 0.273179253 | 0.4 | Neutral |
| testneg_3149 | 0 | 0 | 0.380797 | -3 | 1.797648 | -3 | 0.61011 | 0.09584 | 0.30275 | 0.12788 | 15.61725 | 0.01 | 0 | Neutral |
| testneg_3150 | -0.02 | 0.63 | 4.67013 | -3 | -3.287817 | -3 | 0.10004 | 0.12788 | 0.27605 | 0.21595 | 9.66667 | 0.255550239 | 0.8 | Neutral |
| testneg_3151 | 0.18 | -1.05 | 3.225783 | -3 | 1.787025 | -3 | 0.75978 | 0.11423 | 0.58292 | 0.87725 | 11.9356 | 0.238374945 | 0.2 | Neutral |
| testneg_3152 | -0.08 | -0.03 | 3.408278 | -3 | 1.10156 | 1.10156 | 0.10719 | 0.08618 | 0.16708 | 0.11594 | 19.10253 | 0.040889328 | 0 | Neutral |
| testneg_3153 | -0.07 | -0.01 | 2.291931 | -3 | 4.387225 | -3 | 0.99998 | 0.86471 | 0.16689 | 0.11395 | 8.97898 | 0.220339862 | 0 | Neutral |
| testneg_3154 | 0.07 | 0.38 | 1.522905 | -3 | 1.806186 | 1.806186 | 0.03656 | 0.07074 | 0.19021 | 0.09379 | 27.98024 | 0.06 | 0 | Neutral |
| testneg_3155 | -0.07 | -0.38 | -2.304927 | -2.304927 | 2.795001 | -3 | 0.02665 | 0.05082 | 0.16973 | 0.17555 | 15.93508 | 0.17005618 | 0 | Neutral |
| testneg_3156 | 0 | 0 | 0.229207 | -3 | -2.304927 | -2.304927 | 0.10917 | 0.12788 | 0.02665 | 0.05082 | 34.72222 | 0.01 | 0 | Neutral |
| testneg_3157 | 0.03 | 1.22 | -1.799087 | -3 | -0.074833 | -3 | 0.11391 | 0.19434 | 0.65981 | 0.47389 | 15.86912 | 0.2 | 0 | Neutral |
| testneg_3158 | 0 | 0 | 0.123542 | -3 | -0.1782 | -3 | 0.99915 | 0.96333 | 0.19898 | 0.15276 | 12.29167 | 0.1695 | 0 | Neutral |
| testneg_3159 | -0.15 | -0.04 | 1.448116 | -3 | 1.290195 | 1.290195 | 0.19898 | 0.12788 | 0.10761 | 0.12246 | 20 | 0.05 | 0 | Neutral |
| testneg_3160 | 0.08 | 0.03 | 2.442833 | -3 | 0.931031 | -3 | 0.07701 | 0.19824 | 0.23939 | 0.25284 | 19.09347 | 0.190952381 | 0 | Neutral |
| testneg_3161 | 0 | 0 | 0.419995 | -3 | -0.828875 | -3 | 0.7174 | 0.15815 | 0.17039 | 0.12788 | 12.19355 | 0.01040416 | 0 | Neutral |
| testneg_3162 | 0 | 0 | 1.479946 | -3 | 4.653083 | 4.653083 | 0.56307 | 0.10536 | 0.1174 | 0.12788 | 21.22748 | 0.307847222 | 2.2 | Neutral |
| testneg_3163 | 0 | -0.01 | 0.868715 | -3 | 4.08472 | -3 | 0.95319 | 0.20297 | 0.10213 | 0.22954 | 10.20833 | 0.314041161 | 3.6 | Neutral |
| testneg_3164 | -0.07 | -0.01 | 0.439914 | -3 | 1.324962 | -3 | 0.09275 | 0.14911 | 0.38184 | 0.10478 | 18.33333 | 0.04 | 0 | Neutral |
| testneg_3165 | 0.14 | -0.37 | 3.020834 | -3 | 4.727776 | -3 | 0.19894 | 0.27411 | 0.16474 | 0.41578 | 10.41667 | 0.850491883 | 100 | Disease_causing |
| testneg_3166 | -0.01 | 0.57 | -2.581564 | -3 | -0.369846 | -3 | 0.19898 | 0.12788 | 0.33467 | 0.12788 | 23.68839 | 0.02 | 0 | Neutral |
| testneg_3167 | 0 | 0 | 4.103101 | -3 | 3.746168 | 2.410566 | 0.20011 | 0.27949 | 0.34412 | 0.08672 | 20.27778 | 0.270814394 | 0.2 | Neutral |
| testneg_3168 | 0 | 0 | 0.135171 | -3 | 1.385917 | -3 | 0.08143 | 0.44857 | 0.98641 | 0.83983 | 10.41667 | 0.17 | 0 | Neutral |
| testneg_3169 | 0 | 0 | -0.19086 | -3 | 4.003489 | -3 | 0.15958 | 0.41628 | 0.48504 | 0.12788 | 10.41667 | 0.138605676 | 0 | Neutral |
| testneg_3170 | -0.04 | 1.61 | 12.595297 | -3 | -0.214042 | -3 | 0.3797 | 0.1257 | 0.12836 | 0.09736 | 18.33333 | 0.413809524 | 19.8 | Neutral |
| testneg_3171 | 0.02 | -0.63 | 0.544663 | -3 | 0.771752 | -3 | 0.09767 | 0.39937 | 0.13702 | 0.12788 | 11.25 | 0.06 | 0 | Neutral |

| | | | | | | | | | | | | | | |
|---|---|---|---|---|---|---|---|---|---|---|---|---|---|---|
| testneg_3172 | 0 | 0 | 1.147853 | -3 | 1.299375 | -3 | 0.1481 | 0.1117 | 0.213 | 0.09535 | 19.18549 | 0.000192379 | 0 | Neutral |
| testneg_3173 | 0.17 | 0.38 | 1.422472 | -3 | 3.42834 | -3 | 0.39774 | 0.10813 | 0.09473 | 0.08327 | 41.69407 | 0.09 | 0 | Neutral |
| testneg_3174 | -0.07 | -0.01 | -1.197673 | -3 | 1.365197 | -3 | 0.38445 | 0.58278 | 0.27246 | 0.12788 | 10.41667 | 0.05 | 0 | Neutral |
| testneg_3175 | 0.19 | -0.27 | 2.303838 | -3 | 1.389081 | -3 | 0.17677 | 0.67006 | 0.42214 | 0.21008 | 10.41667 | 0.240185376 | 0 | Neutral |
| testneg_3176 | 0.05 | -0.81 | 2.692724 | -3 | -1.989097 | -3 | 0.52765 | 0.20417 | 0.11094 | 0.12788 | 18.031 | 0.058055556 | 0 | Neutral |
| testneg_3177 | -0.18 | 1.05 | -2.300224 | -3 | -1.379881 | -3 | 0.04792 | 0.12788 | 0.18106 | 0.12788 | 35.09447 | 0.15875 | 0 | Neutral |
| testneg_3178 | 0 | -0.01 | 1.125724 | -3 | -2.887311 | -3 | 0.6185 | 0.09316 | 0.6532 | 0.12788 | 20.70117 | 0.020037037 | 0 | Neutral |
| testneg_3179 | 0.03 | -1.03 | 2.174777 | -3 | 1.205908 | -3 | 0.32458 | 0.1016 | 0.32333 | 0.23356 | 14.19394 | 0.06035401 | 0 | Neutral |
| testneg_3180 | -0.05 | -0.29 | -1.010566 | -3 | -1.060552 | -3 | 0.39192 | 0.09135 | 0.2559 | 0.10848 | 21.63869 | 0.02 | 0 | Neutral |
| testneg_3181 | -0.02 | 0.63 | 2.435299 | -3 | 1.128179 | -3 | 0.72657 | 0.11262 | 0.31062 | 0.1222 | 10.30286 | 0.021 | 0 | Neutral |
| testneg_3182 | 0 | 0 | 0.374751 | -3 | 6.187765 | -3 | 0.18233 | 0.14781 | 0.03499 | 0.12971 | 29.86111 | 0.170154219 | 0 | Neutral |
| testneg_3183 | -0.07 | -0.38 | 1.180617 | -3 | 0.097083 | -3 | 0.1516 | 0.10704 | 0.12943 | 0.14391 | 17.94606 | 0.049270833 | 0 | Neutral |
| testneg_3184 | -0.02 | 0.63 | 2.813311 | -3 | -0.534952 | -3 | 0.10809 | 0.09057 | 0.12203 | 0.13515 | 20.31414 | 0.104606346 | 0 | Neutral |
| testneg_3185 | -0.04 | 0.23 | 3.260847 | -3 | 0.034534 | 0.034534 | 0.27605 | 0.12788 | 0.62381 | 0.9477 | 14.59142 | 0.160994283 | 0 | Neutral |
| testneg_3186 | -0.08 | -0.03 | 4.744051 | -3 | 0.985586 | -3 | 0.19052 | 0.12788 | 0.12077 | 0.11244 | 21.8764 | 0.040254183 | 0 | Neutral |
| testneg_3187 | 0.05 | 0.29 | 2.698795 | -3 | 4.046865 | -3 | 0.17806 | 0.12788 | 0.42666 | 0.12788 | 17.58947 | 0.117340959 | 0 | Neutral |
| testneg_3188 | 0.19 | 0.51 | 4.599202 | -3 | 4.801067 | -3 | 0.34295 | 0.12788 | 0.5103 | 0.10923 | 17.41935 | 0.264397993 | 0.8 | Neutral |
| testneg_3189 | 0 | 0 | 0.203322 | -3 | -0.339562 | -3 | 0.25312 | 0.12788 | 0.17793 | 0.10516 | 29.84187 | 0.01 | 0 | Neutral |
| testneg_3190 | 0.02 | 0.45 | 1.21068 | -3 | 0.441601 | -3 | 0.95247 | 0.11285 | 0.04032 | 0.12788 | 18.33333 | 0.026535714 | 0 | Neutral |
| testneg_3191 | -0.02 | 0.36 | 4.223829 | -3 | 0.18237 | -3 | 0.91921 | 0.32331 | 0.60251 | 0.39112 | 10.41667 | 0.327259259 | 5.2 | Neutral |
| testneg_3192 | 0.01 | 0.8 | 2.99687 | -3 | 2.489427 | -3 | 0.08393 | 0.1123 | 0.72068 | 0.99841 | 9.85525 | 0.27 | 0.2 | Neutral |
| testneg_3193 | 0.08 | 0.03 | -0.240285 | -3 | 1.181824 | -3 | 0.35082 | 0.12788 | 0.55384 | 0.15273 | 18.88889 | 0.063333333 | 0 | Neutral |
| testneg_3194 | 0 | 0 | 1.537848 | -3 | 2.001997 | -3 | 0.11391 | 0.12788 | 0.9193 | 0.14584 | 5.69444 | 0.020263889 | 0 | Neutral |
| testneg_3195 | 0.03 | -1.03 | -0.104883 | -3 | 5.901532 | -3 | 0.16091 | 0.90971 | 0.09442 | 0.1153 | 13.51351 | 0.27917381 | 0.8 | Neutral |
| testneg_3196 | 0 | 0 | 1.66597 | -3 | -2.363283 | -3 | 0.66027 | 0.23984 | 0.05846 | 0.12788 | 18.37615 | 0.02 | 0 | Neutral |
| testneg_3197 | 0 | 0 | 0.191318 | -3 | 1.23956 | -3 | 0.96113 | 0.24974 | 0.33122 | 0.12788 | 10.83333 | 0.014904762 | 0 | Neutral |
| testneg_3198 | 0 | 0 | 0.991757 | -3 | 2.71918 | -3 | 0.19896 | 0.11604 | 0.19898 | 0.12788 | 20.21801 | 0.01 | 0 | Neutral |
| testneg_3199 | 0.03 | 0.12 | 3.086284 | 0.262307 | 6.136401 | -3 | 0.13099 | 0.17453 | 0.3238 | 0.18333 | 10.41667 | 0.821396964 | 100 | Disease_causing |
| testneg_3200 | -0.16 | 1.37 | 1.198668 | -3 | 7.874757 | -3 | 0.03942 | 0.12788 | 0.12965 | 0.75487 | 16.75054 | 0.58 | 88.8 | Disease_causing |
| testneg_3201 | 0.07 | -0.31 | 2.407105 | -3 | 2.747694 | -3 | 0.21095 | 0.10574 | 0.11042 | 0.09881 | 34.21247 | 0.03 | 0 | Neutral |
| testneg_3202 | 0.05 | -0.81 | 3.217486 | -3 | 1.628444 | -3 | 0.74699 | 0.12286 | 0.19821 | 0.20954 | 18.88889 | 0.16625843 | 0 | Neutral |
| testneg_3203 | 0 | -0.32 | -0.477568 | -3 | 2.60964 | -3 | 0.53216 | 0.57092 | 0.7062 | 0.83002 | 10 | 0.272912776 | 2 | Neutral |
| testneg_3204 | 0 | 0 | 2.147643 | -3 | 2.972484 | -3 | 0.9149 | 0.27793 | 0.16158 | 0.3836 | 18.09057 | 0.262870879 | 0.8 | Neutral |
| testneg_3205 | 0.03 | -1.03 | 1.837062 | -3 | 7.159958 | -3 | 0.59066 | 0.10874 | 0.10803 | 0.14652 | 11.25 | 0.442190476 | 33.4 | Neutral |
| testneg_3206 | 0.07 | 0.33 | 3.636293 | -3 | -0.702862 | -3 | 0.17973 | 0.12278 | 0.07275 | 0.12788 | 18.62062 | 0.03 | 0 | Neutral |
| testneg_3207 | 0 | 0 | 1.25978 | 1.25978 | 5.295382 | -3 | 0.60251 | 0.39112 | 0.19898 | 0.09584 | 23.40482 | 0.163818543 | 0 | Neutral |
| testneg_3208 | 0.07 | 0.38 | 0.034534 | 0.034534 | 4.563044 | -3 | 0.62381 | 0.9477 | 0.01952 | 0.08329 | 17.77778 | 0.200833333 | 0 | Neutral |
| testneg_3209 | -0.22 | 2.66 | -1.189105 | -3 | 0.860716 | -3 | 0.25561 | 0.11004 | 0.57729 | 0.10831 | 8.97898 | 0.17 | 0 | Neutral |
| testneg_3210 | -0.07 | -0.01 | -0.23732 | -0.23732 | 1.641098 | -3 | 0.07473 | 0.51812 | 0.13906 | 0.09925 | 10.28409 | 0.060615106 | 0 | Neutral |
| testneg_3211 | -0.16 | 1.37 | 5.815338 | -3 | 1.538088 | -3 | 0.05932 | 0.09551 | 0.24736 | 0.12777 | 19.05506 | 0.27 | 1.2 | Neutral |
| testneg_3212 | 0 | 0 | 1.676782 | -3 | -0.833638 | -0.833638 | 0.49504 | 0.08191 | 0.51282 | 0.12075 | 11.875 | 0.010124806 | 0 | Neutral |
| testneg_3213 | -0.02 | -0.45 | 0.359749 | -3 | 0.355585 | -3 | 0.08954 | 0.63401 | 0.13195 | 0.57267 | 10 | 0.231214286 | 0 | Neutral |
| testneg_3214 | 0 | 0 | 2.105667 | -3 | 5.753514 | -3 | 0.17855 | 0.11262 | 0.22179 | 0.09831 | 18.33333 | 0.08 | 0 | Neutral |
| testneg_3215 | -0.02 | 0.63 | 2.813311 | -3 | 4.842975 | -3 | 0.10809 | 0.09057 | 0.23975 | 0.13861 | 20.86969 | 0.337129205 | 7 | Neutral |
| testneg_3216 | 0.04 | -0.23 | -0.122196 | -0.122196 | -1.755229 | -3 | 0.31961 | 0.17571 | 0.05216 | 0.23851 | 18.88889 | 0.131129149 | 0 | Neutral |
| testneg_3217 | -0.02 | 0.63 | 2.082622 | -3 | 1.311902 | -3 | 0.14335 | 0.14499 | 0.10337 | 0.12788 | 13.51351 | 0.03 | 0 | Neutral |
| testneg_3218 | 0 | 0 | 0.966502 | -3 | 3.036045 | -3 | 0.09508 | 0.09834 | 0.16098 | 0.12788 | 25.08737 | 0.03 | 0 | Neutral |
| testneg_3219 | 0 | 0 | 0.675629 | -3 | 3.157424 | -3 | 0.29145 | 0.11884 | 0.09067 | 0.10671 | 29.02362 | 0.04 | 0 | Neutral |
| testneg_3220 | -0.06 | -0.18 | 2.473481 | -3 | 8.177631 | -3 | 0.23991 | 0.1515 | 0.11522 | 0.11925 | 17.33333 | 0.489926053 | 50.6 | Disease_causing |
| testneg_3221 | 0 | 0 | -0.859115 | -3 | 1.731383 | -3 | 0.19898 | 0.12788 | 0.1617 | 0.17928 | 22.50441 | 0.07 | 0 | Neutral |
| testneg_3222 | 0.05 | -0.44 | 2.978619 | -3 | 0.467995 | -3 | 0.28165 | 0.15912 | 0.11237 | 0.08916 | 23.68839 | 0.054033734 | 0 | Neutral |
| testneg_3223 | 0.18 | -1.05 | 3.992521 | -3 | 2.136098 | -3 | 0.19898 | 0.12788 | 0.13374 | 0.12788 | 26.80135 | 0.07 | 0 | Neutral |
| testneg_3224 | 0.14 | -1.01 | 2.75054 | -3 | 0.128665 | -3 | 0.13691 | 0.11058 | 0.26812 | 0.30371 | 13.43868 | 0.125 | 0 | Neutral |
| testneg_3225 | 0.18 | -1.05 | 0.192071 | -3 | 0.10268 | -3 | 0.18312 | 0.12788 | 0.34193 | 0.12788 | 13.51351 | 0.02 | 0 | Neutral |
| testneg_3226 | 0 | -0.01 | 0.231051 | -3 | 1.917158 | 1.917158 | 0.44164 | 0.13588 | 0.11618 | 0.20784 | 13.67601 | 0.11117241 | 0 | Neutral |
| testneg_3227 | 0.01 | 0.79 | 0.585649 | 0.585649 | 5.257312 | -3 | 0.07185 | 0.68732 | 0.32897 | 0.10658 | 12.46106 | 0.303795855 | 1.4 | Neutral |
| testneg_3228 | -0.17 | -0.38 | 0.128896 | -3 | 5.635253 | -3 | 0.67331 | 0.10118 | 0.19749 | 0.11482 | 30.55556 | 0.16 | 0.2 | Neutral |
| testneg_3229 | -0.03 | -0.1 | 4.70293 | -3 | 2.524962 | -3 | 0.1878 | 0.08946 | 0.36324 | 0.15729 | 13.51351 | 0.310941729 | 4 | Neutral |
| testneg_3230 | 0 | 0 | -0.480488 | -3 | 5.434272 | -3 | 0.11797 | 0.12788 | 0.18925 | 0.32479 | 12.29167 | 0.288667134 | 1.6 | Neutral |
| testneg_3231 | -0.01 | -0.79 | 1.377684 | -3 | 0.076953 | -3 | 0.09406 | 0.19858 | 0.10187 | 0.12339 | 16.98432 | 0.05 | 0 | Neutral |
| testneg_3232 | -0.03 | -0.12 | -0.363694 | -3 | 0.330245 | -3 | 0.07098 | 0.39918 | 0.54248 | 0.12333 | 4.72222 | 0.060982143 | 0 | Neutral |
| testneg_3233 | 0 | 0 | 1.800083 | -3 | 1.222311 | -3 | 0.61211 | 0.11059 | 0.1254 | 0.08697 | 12.35574 | 0.01 | 0 | Neutral |
| testneg_3234 | 0.22 | -2.66 | 6.678689 | -3 | 4.83533 | -3 | 0.20765 | 0.27124 | 0.23975 | 0.13861 | 18.88889 | 0.79 | 100 | Disease_causing |
| testneg_3235 | 0.19 | -0.27 | 6.722995 | -3 | -1.817551 | -3 | 0.5132 | 0.56457 | 0.44541 | 0.13979 | 21.94444 | 0.464857143 | 39 | Neutral |
| testneg_3236 | -0.14 | 1.85 | -1.596835 | -3 | -1.441355 | -3 | 0.17443 | 0.19181 | 0.06729 | 0.17051 | 18.88889 | 0.253095238 | 0.2 | Neutral |
| testneg_3237 | 0 | 0 | 0.365268 | -3 | 1.383004 | -3 | 0.53627 | 0.12788 | 0.45342 | 0.14389 | 14.47674 | 0.030478022 | 0 | Neutral |
| testneg_3238 | 0.2 | -1.26 | 5.986437 | -3 | -1.899809 | -3 | 0.12325 | 0.09383 | 0.10183 | 0.11847 | 23.83333 | 0.13 | 0 | Neutral |
| testneg_3239 | -0.15 | -0.04 | 0.939929 | -3 | 1.036111 | -3 | 0.18701 | 0.12788 | 0.18869 | 0.23984 | 17.21847 | 0.088722222 | 0 | Neutral |
| testneg_3240 | 0 | -0.01 | 0.639661 | -3 | 6.281029 | -3 | 0.16263 | 0.53493 | 0.3962 | 0.10523 | 23.02131 | 0.238724042 | 0.4 | Neutral |
| testneg_3241 | 0.05 | 0.29 | 1.902852 | -3 | 4.154435 | -3 | 0.56966 | 0.68625 | 0.90112 | 0.20622 | 10.83333 | 0.455976046 | 34.8 | Neutral |
| testneg_3242 | 0 | 0 | 0.256955 | -3 | 0.594035 | -3 | 0.1846 | 0.08682 | 0.07347 | 0.08817 | 13.51351 | 0.01 | 0 | Neutral |
| testneg_3243 | 0.07 | 0.38 | -3.00145 | -3 | 0.809929 | -0.228053 | 0.22645 | 0.1704 | 0.11479 | 0.12788 | 17.78322 | 0.040108708 | 0 | Neutral |
| testneg_3244 | -0.06 | 0.57 | -0.000729 | -3 | 4.209007 | -3 | 0.35376 | 0.09478 | 0.10724 | 0.49624 | 12.29167 | 0.320213868 | 3 | Neutral |
| testneg_3245 | 0.04 | -0.23 | 3.709266 | -3 | 1.441342 | -3 | 0.08067 | 0.34914 | 0.11072 | 0.38259 | 20.83333 | 0.526610644 | 65.8 | Disease_causing |
| testneg_3246 | 0.09 | -1.07 | 2.513197 | -3 | 1.65767 | 1.65767 | 0.31732 | 0.27963 | 0.89992 | 0.10231 | 39.13978 | 0.1 | 0 | Neutral |
| testneg_3247 | 0 | 0.32 | 0.284017 | -3 | -2.027863 | -3 | 0.3025 | 0.2499 | 0.05564 | 0.06933 | 10 | 0.021180556 | 0 | Neutral |
| testneg_3248 | -0.03 | -0.1 | 0.642895 | -3 | 2.837786 | -3 | 0.26971 | 0.08509 | 0.7823 | 0.5949 | 26.83742 | 0.131698825 | 0 | Neutral |
| testneg_3249 | 0 | 0 | 0.006114 | -1.917463 | -0.142748 | -3 | 0.15848 | 0.12754 | 0.04346 | 0.10785 | 10.41667 | 0.003194444 | 0 | Neutral |
| testneg_3250 | 0.02 | 0.45 | 2.330575 | -3 | -1.728128 | -3 | 0.63233 | 0.16248 | 0.37635 | 0.16791 | 11.64954 | 0.099958695 | 0 | Neutral |
| testneg_3251 | 0 | 0 | 2.123947 | -3 | 1.545826 | -3 | 0.12867 | 0.11159 | 0.51137 | 0.19235 | 12.29167 | 0.0325 | 0 | Neutral |
| testneg_3252 | 0.03 | 1.22 | 5.062556 | -3 | 5.381477 | -3 | 0.08373 | 0.09191 | 0.34698 | 0.12788 | 17.77778 | 0.355735294 | 11 | Neutral |
| testneg_3253 | 0.03 | -1.03 | 2.432661 | -3 | -1.726521 | -3 | 0.1541 | 0.1077 | 0.03502 | 0.08119 | 20.98109 | 0.020376526 | 0 | Neutral |
| testneg_3254 | 0 | 0 | 0.517865 | -3 | 1.765069 | -3 | 0.76942 | 0.13267 | 0.09099 | 0.2051 | 12.71889 | 0.068882479 | 0 | Neutral |
| testneg_3255 | -0.08 | -0.03 | -1.458126 | -3 | 3.197244 | -3 | 0.05904 | 0.09282 | 0.09611 | 0.07772 | 30.54981 | 0.1 | 0 | Neutral |
| testneg_3256 | 0 | 0 | -0.100514 | -3 | 1.401789 | -3 | 0.16469 | 0.16042 | 0.13061 | 0.21023 | 19.44444 | 0.103181189 | 0 | Neutral |
| testneg_3257 | 0.01 | 0.8 | -0.214756 | -3 | 1.221756 | -3 | 0.07428 | 0.07645 | 0.29268 | 0.16012 | 27.506 | 0.090083333 | 0 | Neutral |
| testneg_3258 | 0.14 | -0.69 | 3.693454 | -3 | 0.57936 | -3 | 0.24144 | 0.12122 | 0.28269 | 0.12505 | 40.00719 | 0.023157596 | 0 | Neutral |
| testneg_3259 | 0.09 | -0.3 | 0.771752 | -3 | 1.925771 | -3 | 0.13702 | 0.12788 | 0.33418 | 0.09979 | 14.3109 | 0.01 | 0 | Neutral |
| testneg_3260 | -0.16 | 1.37 | 6.154007 | -3 | 3.494072 | -3 | 0.33957 | 0.12788 | 0.2118 | 0.2009 | 18.33333 | 0.670962629 | 97.4 | Disease_causing |
| testneg_3261 | -0.02 | -0.45 | 4.5747 | -3 | 0.061667 | -3 | 0.18512 | 0.12788 | 0.13751 | 0.13274 | 12.29692 | 0.119466627 | 0 | Neutral |
| testneg_3262 | 0.16 | -0.37 | 4.774952 | -3 | 1.183232 | -3 | 0.10472 | 0.10014 | 0.19898 | 0.12788 | 18.70486 | 0.04 | 0 | Neutral |
| testneg_3263 | 0.02 | 0.45 | 1.21068 | -3 | 0.485976 | -3 | 0.95247 | 0.11285 | 0.89661 | 0.17382 | 12.37434 | 0.04 | 0 | Neutral |
| testneg_3264 | 0.02 | -0.36 | 2.56113 | -3 | 4.341836 | -3 | 0.06304 | 0.12788 | 0.10187 | 0.08499 | 26.34995 | 0.105941302 | 0 | Neutral |
| testneg_3265 | 0.03 | -0.85 | 2.749126 | -3 | 5.993267 | -3 | 0.31924 | 0.11145 | 0.18866 | 0.12788 | 34.61977 | 0.18 | 0 | Neutral |
| testneg_3266 | 0 | 0 | 2.244975 | -3 | 2.801416 | -3 | 0.92832 | 0.16306 | 0.75978 | 0.11423 | 5.74956 | 0.052110354 | 0 | Neutral |
| testneg_3267 | -0.03 | -0.1 | -0.432565 | -3 | 1.90628 | -3 | 0.1105 | 0.44945 | 0.59342 | 0.12788 | 10.41667 | 0.08 | 0 | Neutral |
| testneg_3268 | 0 | 0 | -0.040357 | -3 | 0.549926 | -3 | 0.01584 | 0.07827 | 0.98255 | 0.98129 | 6.02734 | 0.11 | 0 | Neutral |
| testneg_3269 | -0.07 | -0.38 | 1.051531 | -3 | 5.573938 | -3 | 0.16741 | 0.10762 | 0.19898 | 0.12788 | 13.75 | 0.107906566 | 0 | Neutral |
| testneg_3270 | 0 | 0 | -0.553742 | -3 | 1.384174 | -3 | 0.1424 | 0.12788 | 0.16345 | 0.11716 | 13.02635 | 9.35E-05 | 0 | Neutral |
| testneg_3271 | 0 | 0 | 0.228619 | -3 | 1.499426 | -3 | 0.26451 | 0.09729 | 0.46791 | 0.7316 | 11.0708 | 0.05 | 0 | Neutral |
| testneg_3272 | 0.01 | -0.57 | 4.493837 | -3 | 0.475124 | -3 | 0.19898 | 0.14645 | 0.17677 | 0.67006 | 15.2381 | 0.248268433 | 0 | Neutral |
| testneg_3273 | -0.18 | -0.16 | 0.910161 | -3 | 4.228466 | 1.640541 | 0.93468 | 0.12788 | 0.48688 | 0.13071 | 22.47657 | 0.219714286 | 0 | Neutral |
| testneg_3274 | -0.05 | 0.81 | 3.271518 | -3 | 1.792362 | -3 | 0.1214 | 0.0899 | 0.23427 | 0.10875 | 11.25 | 0.06 | 0 | Neutral |
| testneg_3275 | 0.04 | -0.23 | 7.284294 | -3 | 2.522105 | -3 | 0.30684 | 0.12788 | 0.24327 | 0.12192 | 12.74854 | 0.411217532 | 23.2 | Neutral |
| testneg_3276 | 0.07 | 0.38 | -2.164532 | -3 | 0.817737 | -3 | 0.10562 | 0.09329 | 0.15581 | 0.10782 | 37.50183 | 0.02 | 0 | Neutral |
| testneg_3277 | 0.01 | 0.8 | 1.360649 | -3 | 7.158496 | -3 | 0.13702 | 0.12788 | 0.16926 | 0.11871 | 12.84583 | 0.29 | 2.2 | Neutral |
| testneg_3278 | 0.14 | -0.37 | 7.061469 | -3 | 3.002533 | 3.002533 | 0.32335 | 0.10604 | 0.14304 | 0.10715 | 42.23911 | 0.43 | 26.4 | Neutral |
| testneg_3279 | 0 | 0 | 0.040441 | -3 | 2.462213 | -3 | 0.54777 | 0.25991 | 0.32016 | 0.11638 | 10 | 0.031315943 | 0 | Neutral |
| testneg_3280 | -0.07 | -0.38 | -1.73147 | -3 | 1.572604 | -3 | 0.09749 | 0.82924 | 0.21967 | 0.09467 | 43.07487 | 0.130126663 | 0 | Neutral |
| testneg_3281 | -0.07 | -0.01 | 2.75027 | -3 | 0.235742 | -3 | 0.50034 | 0.07527 | 0.3718 | 0.2248 | 29.1283 | 0.092568493 | 0 | Neutral |
| testneg_3282 | 0 | -0.01 | 0.627226 | -3 | 0.304852 | -3 | 0.29004 | 0.45465 | 0.24 | 0.2063 | 5 | 0.068304671 | 0 | Neutral |
| testneg_3283 | 0.07 | 0.01 | 2.719238 | -3 | 6.337267 | 4.003489 | 0.47775 | 0.10103 | 0.48504 | 0.12788 | 18.88889 | 0.38 | 12 | Neutral |
| testneg_3284 | 0.04 | 2.01 | -0.573377 | -3 | 6.762042 | -3 | 0.42366 | 0.14372 | 0.09445 | 0.12788 | 21.38889 | 0.328151261 | 4.8 | Neutral |
| testneg_3285 | 0 | 0 | 0.494101 | -3 | 3.478553 | -3 | 0.28038 | 0.11814 | 0.07961 | 0.11474 | 38.99952 | 0.041770833 | 0 | Neutral |
| testneg_3286 | 0 | 0 | -0.02198 | -0.02198 | 5.153522 | -3 | 0.15509 | 0.1326 | 0.1205 | 0.23508 | 18.88889 | 0.397137443 | 18.4 | Neutral |
| testneg_3287 | 0 | 0 | -0.734901 | -3 | 3.869278 | -3 | 0.64721 | 0.8758 | 0.12161 | 0.24178 | 10 | 0.351973506 | 6.4 | Neutral |
| testneg_3288 | 0 | 0 | 0.310149 | 0.310149 | 1.22522 | -3 | 0.0626 | 0.5373 | 0.7062 | 0.83002 | 12.79101 | 0.248779146 | 0.2 | Neutral |
| testneg_3289 | -0.0105455 | 0.17672727 | 11.548039 | -3 | 2.884479 | -3 | 0.19898 | 0.0989 | 0.41383 | 0.119 | 16.3887 | 0.50575 | 58.6 | Disease_causing |
| testneg_3290 | -0.14 | 1.85 | 0.923069 | -3 | -1.178484 | -1.178484 | 0.06711 | 0.09765 | 0.13669 | 0.09263 | 17.57218 | 0.11 | 0 | Neutral |
| testneg_3291 | 0 | 0 | -0.505055 | -3 | 4.799959 | -3 | 0.99859 | 0.12788 | 0.32531 | 0.12351 | 16.33387 | 0.080490431 | 0 | Neutral |
| testneg_3292 | -0.02 | 0.63 | 0.792282 | -3 | 1.039407 | -3 | 0.75008 | 0.70645 | 0.28201 | 0.33127 | 10.31261 | 0.135385555 | 0 | Neutral |
| testneg_3293 | -0.05 | -0.29 | -1.851788 | -3 | 2.531153 | -3 | 0.78741 | 0.22085 | 0.36833 | 0.10433 | 6.99956 | 0.060469208 | 0 | Neutral |

| | | | | | | | | | | | | | | |
|---|---|---|---|---|---|---|---|---|---|---|---|---|---|---|
| testneg_3294 | 0.21 | 0.15 | 5.939999 | -3 | 0.29779 | -3 | 0.99531 | 0.86203 | 0.19898 | 0.20319 | 18.76159 | 0.51 | 58 | Disease_causing |
| testneg_3295 | 0 | 0 | 0.828595 | -3 | 0.144628 | -3 | 0.93284 | 0.71489 | 0.36417 | 0.21506 | 10.41667 | 0.055658519 | 0 | Neutral |
| testneg_3296 | 0 | 0.73 | 4.722814 | -3 | 0.714207 | -3 | 0.03008 | 0.12788 | 0.04346 | 0.10785 | 16.8018 | 0.1 | 0 | Neutral |
| testneg_3297 | -0.03 | -0.1 | -0.76618 | -3 | 3.635653 | -3 | 0.1503 | 0.12788 | 0.11501 | 0.13748 | 20 | 0.206268018 | 0 | Neutral |
| testneg_3298 | 0.07 | 0.7 | 4.605651 | -3 | 3.959597 | -3 | 0.10719 | 0.08618 | 0.55618 | 0.11352 | 17.13595 | 0.21 | 0 | Neutral |
| testneg_3299 | -0.09 | 0.25 | 2.350679 | -3 | 4.228466 | -3 | 0.26264 | 0.13812 | 0.48688 | 0.13071 | 10.30286 | 0.239097925 | 0.6 | Neutral |
| testneg_3300 | -0.01 | -0.8 | 1.064707 | -3 | 2.18915 | -3 | 0.27637 | 0.0872 | 0.06187 | 0.36607 | 13.75 | 0.11 | 0 | Neutral |
| testneg_3301 | 0.21 | 0.15 | 3.345724 | -3 | 0.072482 | -3 | 0.30275 | 0.12788 | 0.07394 | 0.09408 | 9.78018 | 0.04 | 0 | Neutral |
| testneg_3302 | -0.02 | 0.63 | 2.082622 | -3 | 2.980687 | -3 | 0.14335 | 0.14499 | 0.74558 | 0.10643 | 10.30286 | 0.091368056 | 0 | Neutral |
| testneg_3303 | -0.07 | -0.38 | 1.102328 | -3 | 1.099551 | -3 | 0.74271 | 0.10951 | 0.82714 | 0.7562 | 20.98109 | 0.09 | 0 | Neutral |
| testneg_3304 | 0.03 | -1.03 | 4.859478 | -3 | 0.201766 | -3 | 0.11501 | 0.13748 | 0.16388 | 0.14439 | 23.33333 | 0.17421526 | 0 | Neutral |
| testneg_3305 | 0 | 0 | 1.332007 | -3 | 0.097678 | -3 | 0.18785 | 0.13588 | 0.17451 | 0.11227 | 10 | 0.000385185 | 0 | Neutral |
| testneg_3306 | 0.05 | -0.81 | 3.217486 | -3 | 5.703258 | -3 | 0.74699 | 0.12286 | 0.60812 | 0.19286 | 20.55556 | 0.55279233 | 71.6 | Disease_causing |
| testneg_3307 | -0.15 | -0.04 | 0.991883 | -3 | 4.435214 | -3 | 0.26816 | 0.13347 | 0.84431 | 0.65315 | 11.90227 | 0.390237428 | 15.4 | Neutral |
| testneg_3308 | 0.00072727 | -0.518 | 4.098385 | -3 | 0.504888 | -3 | 0.09531 | 0.11449 | 0.06476 | 0.07491 | 15.51622 | 0.03 | 0 | Neutral |
| testneg_3309 | 0 | 0 | -0.076686 | -3 | 1.411524 | 0.674816 | 0.84596 | 0.10633 | 0.49831 | 0.29427 | 35.56147 | 0.092768519 | 0 | Neutral |
| testneg_3310 | 0.11 | -1.95 | 5.597055 | -3 | 0.329621 | 0.329621 | 0.53496 | 0.13586 | 0.4129 | 0.11172 | 10.83333 | 0.209695815 | 0 | Neutral |
| testneg_3311 | -0.05 | -0.29 | -2.95337 | -3 | 5.135383 | -3 | 0.19898 | 0.12788 | 0.12682 | 0.11357 | 26.92151 | 0.090636364 | 0 | Neutral |
| testneg_3312 | 0 | 0 | 1.66597 | -3 | -0.811454 | -0.811454 | 0.66027 | 0.23984 | 0.72912 | 0.18218 | 18.33333 | 0.060408333 | 0 | Neutral |
| testneg_3313 | 0.22 | -2.66 | 2.115161 | -3 | -0.01697 | -3 | 0.34859 | 0.09799 | 0.8376 | 0.24333 | 5.75579 | 0.120235588 | 0 | Neutral |
| testneg_3314 | 0.09 | -0.3 | -0.247306 | -3 | 1.324195 | -3 | 0.35945 | 0.12025 | 0.07467 | 0.09224 | 20 | 0.02 | 0 | Neutral |
| testneg_3315 | 0.07 | 0.7 | 6.445404 | 2.913038 | 0.75187 | -3 | 0.60251 | 0.39112 | 0.19898 | 0.535 | 4.72222 | 0.81 | 100 | Disease_causing |
| testneg_3316 | 0 | 0 | 0.319815 | -3 | -0.888381 | -3 | 0.32804 | 0.10857 | 0.00487 | 0.0143 | 10.83333 | 0.01 | 0 | Neutral |
| testneg_3317 | -0.02 | -0.45 | -1.069153 | -3 | 0.47202 | -3 | 0.19668 | 0.16042 | 0.13295 | 0.12914 | 13.46801 | 0.03 | 0 | Neutral |
| testneg_3318 | 0.03 | 0.12 | 4.035621 | -3 | -1.989102 | -3 | 0.04615 | 0.12788 | 0.1798 | 0.0954 | 12.19153 | 0.08 | 0 | Neutral |
| testneg_3319 | 0.04 | 2.01 | -3.135825 | -3 | 1.08174 | -3 | 0.35422 | 0.11185 | 0.92582 | 0.12952 | 36.57431 | 0.1 | 0 | Neutral |
| testneg_3320 | 0.21 | 0.15 | 3.926028 | -3 | 2.99187 | -3 | 0.9516 | 0.12788 | 0.05996 | 0.12788 | 45.19818 | 0.170208333 | 0 | Neutral |
| testneg_3321 | 0 | 0 | -0.211085 | -3 | 0.400863 | -3 | 0.04543 | 0.28103 | 0.08982 | 0.15279 | 10 | 0.171163333 | 0 | Neutral |
| testneg_3322 | 0.02 | -0.63 | 2.525878 | -3 | 3.835285 | -3 | 0.19898 | 0.08275 | 0.08265 | 0.12788 | 16.80383 | 0.08 | 0 | Neutral |
| testneg_3323 | 0 | 0 | 1.636389 | -3 | 0.977001 | -3 | 0.10863 | 0.098 | 0.42748 | 0.10231 | 17.44198 | 0 | 0 | Neutral |
| testneg_3324 | 0.07 | 0.33 | 1.275096 | -3 | 5.093093 | -3 | 0.1592 | 0.12788 | 0.14 | 0.12151 | 29.30135 | 0.07 | 0 | Neutral |
| testneg_3325 | 0 | 0 | 0.543131 | -3 | -0.006272 | -0.006272 | 0.1331 | 0.12788 | 0.07335 | 0.12788 | 28.49046 | 0.01022739 | 0 | Neutral |
| testneg_3326 | 0.05 | -0.81 | 6.132257 | -3 | 1.940661 | -3 | 0.19479 | 0.12788 | 0.25151 | 0.12788 | 19.44313 | 0.2 | 0.4 | Neutral |
| testneg_3327 | 0.07 | 0.38 | -1.688101 | -3 | 0.049287 | -3 | 0.65012 | 0.21208 | 0.42919 | 0.1092 | 10.23791 | 0.05 | 0 | Neutral |
| testneg_3328 | 0.05 | 0.29 | 0.721552 | -3 | 1.041001 | -3 | 0.19898 | 0.09822 | 0.36858 | 0.09681 | 19.09347 | 0.003741497 | 0 | Neutral |
| testneg_3329 | 0.01 | 0.8 | 5.875511 | -3 | 1.199974 | -3 | 0.19898 | 0.12788 | 0.21884 | 0.16743 | 24.70073 | 0.42 | 22.6 | Neutral |
| testneg_3330 | 0.18 | -1.05 | 6.005802 | -3 | 1.635034 | -3 | 0.70627 | 0.11276 | 0.55618 | 0.11352 | 25.86874 | 0.191465086 | 0 | Neutral |
| testneg_3331 | 0.01 | 0.79 | 3.066263 | -3 | 1.624358 | -3 | 0.17487 | 0.1672 | 0.16248 | 0.14352 | 17.06927 | 0.183854167 | 0 | Neutral |
| testneg_3332 | 0.02 | -0.63 | 2.525878 | -3 | 2.768711 | 2.768711 | 0.19898 | 0.08275 | 0.38324 | 0.25193 | 10.72411 | 0.273416667 | 0.8 | Neutral |
| testneg_3333 | -0.0081119 | -0.0026573 | 1.451979 | -3 | -0.671472 | -3 | 0.26727 | 0.12788 | 0.08878 | 0.0974 | 24.15346 | 0.01 | 0 | Neutral |
| testneg_3334 | 0.22 | -2.66 | 2.631407 | 2.631407 | 0.666834 | -3 | 0.44541 | 0.13979 | 0.25625 | 0.66481 | 21.94444 | 0.470320513 | 45.2 | Neutral |
| testneg_3335 | 0.08 | -1.2 | 1.68139 | -3 | 1.060676 | -3 | 0.4094 | 0.1944 | 0.18122 | 0.16895 | 16.9194 | 0.140408333 | 0 | Neutral |
| testneg_3336 | 0.05 | 0.29 | 2.702357 | -3 | -0.133882 | -3 | 0.94674 | 0.12413 | 0.1056 | 0.12788 | 11.13095 | 0.03 | 0 | Neutral |
| testneg_3337 | 0.05 | -0.81 | 5.915153 | -3 | 2.820155 | -3 | 0.30951 | 0.12788 | 0.06254 | 0.23561 | 20.40527 | 0.563691134 | 79 | Disease_causing |
| testneg_3338 | 0 | 0 | 0.173288 | -3 | -0.277982 | -3 | 0.61211 | 0.11059 | 0.04869 | 0.05922 | 14.73943 | 0.01 | 0 | Neutral |
| testneg_3339 | 0.18 | -1.05 | 5.390189 | -3 | 2.754446 | -3 | 0.15041 | 0.43468 | 0.63798 | 0.19967 | 10.83333 | 0.766355769 | 100 | Disease_causing |
| testneg_3340 | 0.03 | 0.12 | 2.051799 | -3 | 0.778269 | -3 | 0.14624 | 0.10369 | 0.10176 | 0.12788 | 16.84579 | 0.01 | 0 | Neutral |
| testneg_3341 | 0 | -0.01 | 3.05843 | 3.05843 | 8.568693 | -3 | 0.74136 | 0.1049 | 0.1834 | 0.12788 | 37.29257 | 0.53 | 68 | Disease_causing |
| testneg_3342 | 0 | -0.01 | 4.102872 | -3 | 0.160465 | -3 | 0.37463 | 0.23021 | 0.05153 | 0.12788 | 19.44444 | 0.075 | 0 | Neutral |
| testneg_3343 | 0.22 | -2.66 | 0.385189 | 0.385189 | 1.770854 | -3 | 0.11533 | 0.12788 | 0.07558 | 0.12788 | 18.88889 | 0.072361111 | 0 | Neutral |
| testneg_3344 | -0.07 | -0.38 | 1.894127 | -3 | -1.653555 | -1.653555 | 0.62004 | 0.10842 | 0.14588 | 0.12156 | 13.08333 | 0.010103799 | 0 | Neutral |
| testneg_3345 | 0.03 | -0.85 | 4.652483 | -3 | 0.447846 | -3 | 0.06841 | 0.10927 | 0.69563 | 0.12788 | 10.83333 | 0.07 | 0 | Neutral |
| testneg_3346 | 0 | 0 | 1.302665 | -3 | 3.231366 | 3.231366 | 0.71803 | 0.95763 | 0.37237 | 0.36829 | 5.71262 | 0.368406446 | 12.2 | Neutral |
| testneg_3347 | -0.14 | 1.01 | 6.791408 | -3 | 1.330976 | -3 | 0.79692 | 0.51979 | 0.25011 | 0.12092 | 5.69444 | 0.492532051 | 53.2 | Disease_causing |
| testneg_3348 | 0 | 0 | -0.134744 | -3 | 0.29558 | -3 | 0.1192 | 0.10391 | 0.87806 | 0.38966 | 10.11674 | 0.03 | 0 | Neutral |
| testneg_3349 | 0 | -0.01 | 3.197244 | -3 | 6.981338 | -3 | 0.09611 | 0.07772 | 0.52278 | 0.41207 | 17.5501 | 0.523030372 | 65.4 | Disease_causing |
| testneg_3350 | 0.04 | -0.23 | 1.023998 | -3 | 3.809259 | 3.809259 | 0.52005 | 0.12788 | 0.42306 | 0.86891 | 11.92982 | 0.4445 | 32.8 | Neutral |
| testneg_3351 | 0 | -0.32 | 5.730226 | -3 | 0.594827 | -3 | 0.13704 | 0.10054 | 0.28474 | 0.12788 | 27.64576 | 0.11 | 0 | Neutral |
| testneg_3352 | 0.18 | -1.05 | 4.102055 | -3 | 0.219231 | -3 | 0.24823 | 0.12788 | 0.68423 | 0.08163 | 13.51351 | 0.050102041 | 0 | Neutral |
| testneg_3353 | -0.05 | -0.29 | -0.079939 | -3 | -0.086754 | -3 | 0.50461 | 0.11034 | 0.07872 | 0.12788 | 13.37927 | 0.02 | 0 | Neutral |
| testneg_3354 | 0 | 0 | 1.317673 | -3 | 0.917232 | -3 | 0.45415 | 0.12971 | 0.25556 | 0.09342 | 16.63097 | 0.000521779 | 0 | Neutral |
| testneg_3355 | -0.07 | -0.38 | -1.026052 | -3 | 4.202013 | -3 | 0.16469 | 0.16042 | 0.03505 | 0.0907 | 18.6768 | 0.172134528 | 0 | Neutral |
| testneg_3356 | 0 | 0 | -0.338929 | -3 | -0.905147 | -3 | 0.6323 | 0.09928 | 0.13169 | 0.06895 | 10.41667 | 0.01 | 0 | Neutral |
| testneg_3357 | 0.04 | -0.23 | 1.048418 | -3 | 7.951887 | -3 | 0.27197 | 0.12346 | 0.82846 | 0.13042 | 10.83333 | 0.32 | 4.6 | Neutral |
| testneg_3358 | 0 | 0 | 0.340894 | -3 | 0.699547 | -3 | 0.07828 | 0.76259 | 0.25637 | 0.37408 | 13.41439 | 0.195485302 | 0 | Neutral |
| testneg_3359 | -0.14 | 0.37 | -0.030786 | -3 | 0.976924 | -3 | 0.56416 | 0.12788 | 0.07427 | 0.09451 | 10.52365 | 0.05 | 0 | Neutral |
| testneg_3360 | 0 | 0 | -0.137186 | -3 | 3.445512 | -3 | 0.38002 | 0.22508 | 0.38126 | 0.27764 | 21.43745 | 0.342365279 | 7.2 | Neutral |
| testneg_3361 | 0 | 0 | 1.326221 | -3 | -0.139258 | -3 | 0.25077 | 0.06261 | 0.31025 | 0.09491 | 17.70469 | 0.000218743 | 0 | Neutral |
| testneg_3362 | 0 | 0 | 2.30653 | -3 | 3.218364 | -3 | 0.49106 | 0.89143 | 0.39305 | 0.18672 | 4.72222 | 0.334717419 | 6 | Neutral |
| testneg_3363 | 0 | 0 | 1.687316 | -3 | 5.12683 | -3 | 0.26434 | 0.12788 | 0.70616 | 0.09323 | 13.08587 | 0.056519397 | 0 | Neutral |
| testneg_3364 | -0.01 | -0.8 | 0.369076 | -3 | 3.063227 | -3 | 0.11703 | 0.09364 | 0.05153 | 0.12788 | 18.6768 | 0.06 | 0 | Neutral |
| testneg_3365 | 0.02 | 0.45 | 5.641127 | -3 | 0.666834 | -3 | 0.14458 | 0.14062 | 0.25625 | 0.66481 | 13.51351 | 0.407331068 | 21.4 | Neutral |
| testneg_3366 | 0 | 0 | -0.480488 | -3 | -1.803416 | -3 | 0.11797 | 0.12788 | 0.62381 | 0.12788 | 20.39295 | 0.01 | 0 | Neutral |
| testneg_3367 | -0.07 | -0.38 | 3.545912 | -3 | 1.603588 | -3 | 0.21811 | 0.18489 | 0.10496 | 0.21491 | 18.88889 | 0.325509246 | 6.4 | Neutral |
| testneg_3368 | -0.08 | -0.03 | 7.833352 | -3 | -1.039289 | -1.039289 | 0.08067 | 0.34914 | 0.08538 | 0.08065 | 32.29167 | 0.409375 | 19.2 | Neutral |
| testneg_3369 | -0.03 | -0.1 | 0.968512 | -3 | 0.861662 | -1.394924 | 0.40074 | 0.36296 | 0.04831 | 0.12788 | 28.87297 | 0.06 | 0 | Neutral |
| testneg_3370 | -0.01 | -0.79 | 0.34963 | -3 | 6.654254 | -3 | 0.17173 | 0.15363 | 0.24906 | 0.11684 | 21.1411 | 0.29 | 2.6 | Neutral |
| testneg_3371 | -0.02 | 0.36 | 3.783871 | -3 | 0.155941 | -3 | 0.33923 | 0.12032 | 0.48817 | 0.10601 | 19.09347 | 0.030465368 | 0 | Neutral |
| testneg_3372 | -0.02 | -0.45 | 5.92982 | -3 | 3.222724 | -3 | 0.99948 | 0.5595 | 0.08183 | 0.11603 | 5.97222 | 0.485181818 | 49.8 | Neutral |
| testneg_3373 | 0.09 | -0.3 | 4.724251 | -3 | 1.67135 | -3 | 0.23859 | 0.1202 | 0.64125 | 0.28792 | 13.62722 | 0.250321429 | 0.8 | Neutral |
| testneg_3374 | -0.18 | 1.05 | 2.778725 | 2.778725 | 0.994999 | -3 | 0.33528 | 0.29463 | 0.51282 | 0.12075 | 12.70833 | 0.39 | 18 | Neutral |
| testneg_3375 | 0.14 | -1.01 | 6.804639 | -3 | 3.440718 | -3 | 0.07577 | 0.09128 | 0.24011 | 0.28559 | 18.88889 | 0.592179781 | 85 | Disease_causing |
| testneg_3376 | -0.02 | -0.45 | 3.092567 | -3 | 2.100416 | -3 | 0.25473 | 0.07863 | 0.48298 | 0.1128 | 13.51351 | 0.046736111 | 0 | Neutral |
| testneg_3377 | -0.18 | -0.16 | 3.778693 | -3 | 4.785647 | -3 | 0.33467 | 0.12788 | 0.15581 | 0.10782 | 36.16733 | 0.203463076 | 0 | Neutral |
| testneg_3378 | -0.01 | -0.79 | -0.811454 | -0.811454 | 0.827033 | -3 | 0.72912 | 0.18218 | 0.38934 | 0.15403 | 10.41667 | 0.0796875 | 0 | Neutral |
| testneg_3379 | -0.07 | -0.33 | -1.772514 | -3 | -0.669593 | -3 | 0.46735 | 0.10674 | 0.14862 | 0.12788 | 48.9049 | 0.02 | 0 | Neutral |
| testneg_3380 | -0.07 | -0.01 | -1.980002 | -3 | 3.480317 | -3 | 0.12176 | 0.09356 | 0.16262 | 0.15786 | 18.92511 | 0.210453623 | 0 | Neutral |
| testneg_3381 | -0.19 | 0.27 | 4.95955 | -3 | 1.770771 | -3 | 0.06247 | 0.14184 | 0.71406 | 0.2012 | 4.44444 | 0.380406142 | 10.4 | Neutral |
| testneg_3382 | 0.09 | -0.67 | 1.431731 | -3 | 2.531483 | -3 | 0.68228 | 0.09762 | 0.47462 | 0.10826 | 20.94356 | 0.05 | 0 | Neutral |
| testneg_3383 | 0.09 | -2.17 | 3.628039 | 3.628039 | -0.151164 | -0.151164 | 0.055 | 0.20858 | 0.65708 | 0.0936 | 19.11866 | 0.49 | 51.2 | Disease_causing |
| testneg_3384 | 0 | 0 | 1.134819 | -3 | 3.650561 | -3 | 0.89357 | 0.22348 | 0.0961 | 0.21256 | 17.21847 | 0.290803846 | 1.4 | Neutral |
| testneg_3385 | 0 | 0 | 0.018971 | -3 | 1.094334 | -3 | 0.17677 | 0.67006 | 0.21894 | 0.09041 | 26.96078 | 0.054214286 | 0 | Neutral |
| testneg_3386 | 0 | 0 | 1.568529 | -3 | 1.762821 | 1.762821 | 0.76942 | 0.13267 | 0.1375 | 0.112 | 32.29167 | 0.04 | 0 | Neutral |
| testneg_3387 | 0.07 | 0.33 | 0.385624 | -3 | 1.632718 | -3 | 0.49012 | 0.12138 | 0.19898 | 0.12788 | 42.96227 | 0.02 | 0 | Neutral |
| testneg_3388 | -0.18 | 1.05 | 3.759941 | -3 | -1.020022 | -3 | 0.46058 | 0.12903 | 0.61867 | 0.15658 | 30.00722 | 0.2 | 0 | Neutral |
| testneg_3389 | 0 | 0 | 0.785748 | -3 | 2.115914 | -3 | 0.52388 | 0.25831 | 0.9807 | 0.2062 | 11.25 | 0.094126862 | 0 | Neutral |
| testneg_3390 | -0.01 | 0.57 | -3.231094 | -3 | 0.865658 | -3 | 0.05323 | 0.32175 | 0.64108 | 0.51829 | 12.43458 | 0.235 | 0 | Neutral |
| testneg_3391 | 0 | 0 | -0.78698 | -3 | 5.495903 | -3 | 0.39193 | 0.11262 | 0.69426 | 0.13242 | 10 | 0.193031788 | 0 | Neutral |
| testneg_3392 | 0 | 0 | 1.474935 | -3 | 0.364762 | -3 | 0.96651 | 0.28104 | 0.85075 | 0.16646 | 5.88477 | 0.020928521 | 0 | Neutral |
| testneg_3393 | 0 | -0.01 | 0.834318 | -3 | 4.892896 | -3 | 0.31726 | 0.17249 | 0.19898 | 0.0989 | 16.3887 | 0.087907066 | 0 | Neutral |
| testneg_3394 | 0.11 | -1.95 | 3.52493 | -3 | 1.222548 | 1.222548 | 0.04916 | 0.14944 | 0.06705 | 0.12788 | 25.55193 | 0.17 | 0 | Neutral |
| testneg_3395 | 0 | 0 | 0.09433 | -3 | -1.532457 | -3 | 0.18925 | 0.32479 | 0.53784 | 0.5574 | 5.75579 | 0.100313679 | 0 | Neutral |
| testneg_3396 | 0.00055944 | 0.00363636 | 1.360315 | -3 | 0.100154 | -3 | 0.29482 | 0.15726 | 0.21122 | 0.12686 | 10.41667 | 0.01 | 0 | Neutral |
| testneg_3397 | 0.07 | 0.38 | 2.359524 | -3 | 0.689808 | -3 | 0.33957 | 0.12788 | 0.16404 | 0.4047 | 11.25 | 0.083330894 | 0 | Neutral |
| testneg_3398 | -0.02 | 0.63 | -0.628934 | -3 | 3.901083 | -3 | 0.17576 | 0.12193 | 0.52118 | 0.36073 | 9.75077 | 0.276169623 | 1.4 | Neutral |
| testneg_3399 | 0 | 0 | 0.727897 | -3 | 1.762547 | -3 | 0.34209 | 0.09515 | 0.28474 | 0.12788 | 18.52533 | 0.000107577 | 0 | Neutral |
| testneg_3400 | 0.21 | 0.15 | 5.338361 | -3 | 7.556384 | 1.367495 | 0.21587 | 0.11701 | 0.17843 | 0.15468 | 10.11292 | 0.808303571 | 100 | Disease_causing |
| testneg_3401 | 0 | 0 | 1.445184 | 1.445184 | -0.410022 | -3 | 0.33469 | 0.17432 | 0.17419 | 0.33842 | 4.72222 | 0.120862472 | 0 | Neutral |
| testneg_3402 | -0.06 | -0.18 | -0.580228 | -3 | -0.031518 | -3 | 0.13331 | 0.09729 | 0.11281 | 0.0958 | 17.21847 | 0.01 | 0 | Neutral |
| testneg_3403 | 0.18 | -1.05 | 2.686668 | -3 | 5.390365 | -3 | 0.10517 | 0.09803 | 0.09749 | 0.82924 | 23.33333 | 0.497083333 | 53 | Disease_causing |
| testneg_3404 | -0.08 | -0.03 | 2.090774 | -3 | 3.705834 | -3 | 0.35421 | 0.10694 | 0.16781 | 0.12788 | 19.26372 | 0.070283242 | 0 | Neutral |
| testneg_3405 | -0.05 | 0.44 | -1.084107 | -3 | 2.171997 | -3 | 0.05378 | 0.12788 | 0.20518 | 0.12378 | 20.37554 | 0.06 | 0 | Neutral |
| testneg_3406 | 0.01 | 0.79 | -0.188304 | -3 | 3.179644 | -3 | 0.16418 | 0.12788 | 0.12871 | 0.29252 | 10.41667 | 0.210093434 | 0 | Neutral |
| testneg_3407 | 0 | 0 | -0.51617 | -3 | 0.433941 | 0.433941 | 0.19898 | 0.12788 | 0.30475 | 0.21064 | 5.97222 | 0.04 | 0 | Neutral |
| testneg_3408 | 0.19 | 0.51 | 6.158659 | -3 | 1.210357 | -3 | 0.22258 | 0.10471 | 0.13669 | 0.09263 | 32.06474 | 0.154673203 | 0.2 | Neutral |
| testneg_3409 | 0.01 | 0.8 | 5.791189 | -3 | 2.454179 | -3 | 0.09918 | 0.11061 | 0.11788 | 0.12788 | 11.58727 | 0.263138728 | 2.4 | Neutral |
| testneg_3410 | 0.15 | 0.06 | -3.376225 | -3 | -3.186924 | -3 | 0.04123 | 0.06896 | 0.19898 | 0.09822 | 19.09347 | 0.05 | 0 | Neutral |
| testneg_3411 | 0.06 | 0.18 | -0.872395 | -3 | 4.099778 | 4.099778 | 0.03631 | 0.80564 | 0.06092 | 0.07159 | 10.41667 | 0.4 | 19.4 | Neutral |
| testneg_3412 | -0.08 | -0.03 | 0.950149 | -3 | 0.552815 | 0.552815 | 0.31318 | 0.11262 | 0.20417 | 0.10681 | 18.33333 | 0.01 | 0 | Neutral |
| testneg_3413 | 0.04 | -0.23 | 0.627475 | -3 | -0.948009 | -3 | 0.30951 | 0.12788 | 0.10379 | 0.34799 | 19.14729 | 0.102330317 | 0 | Neutral |
| testneg_3414 | 0 | -0.01 | 0.967447 | -3 | 5.162562 | -3 | 0.22658 | 0.20646 | 0.16299 | 0.10454 | 11.23846 | 0.134467965 | 0 | Neutral |
| testneg_3415 | 0.08 | -1.2 | -0.298935 | -3 | 3.364563 | -3 | 0.78462 | 0.12258 | 0.07258 | 0.06248 | 13.46801 | 0.101752646 | 0 | Neutral |

| | | | | | | | | | | | | | | |
|---|---|---|---|---|---|---|---|---|---|---|---|---|---|---|
| testneg_3416 | 0 | 0 | -0.179197 | -3 | 0.878873 | -3 | 0.86835 | 0.1513 | 0.62197 | 0.12788 | 9.89595 | 0.01 | 0 | Neutral |
| testneg_3417 | 0.21 | 0.15 | 5.309693 | -3 | 5.126283 | -3 | 0.09951 | 0.14299 | 0.68423 | 0.08163 | 18.21104 | 0.38 | 16.8 | Neutral |
| testneg_3418 | 0 | 0 | 1.108803 | -3 | -1.110257 | -3 | 0.85092 | 0.58871 | 0.15217 | 0.12544 | 6.94444 | 0.010823562 | 0 | Neutral |
| testneg_3419 | 0.08 | 0.03 | 5.535707 | -3 | 0.980675 | -3 | 0.1541 | 0.1077 | 0.35137 | 0.09161 | 28.4414 | 0.09 | 0 | Neutral |
| testneg_3420 | 0 | 0 | 0.728127 | -3 | 6.462116 | -3 | 0.07681 | 0.22865 | 0.06517 | 0.15107 | 19.31062 | 0.576691392 | 80.6 | Disease_causing |
| testneg_3421 | -0.15 | -0.04 | 2.846922 | -3 | 0.876243 | -3 | 0.18081 | 0.1258 | 0.694 | 0.09974 | 14.5275 | 0.06 | 0 | Neutral |
| testneg_3422 | 0 | 0 | 0.116062 | -3 | -0.765143 | -0.765143 | 0.4811 | 0.13398 | 0.31627 | 0.23288 | 10.83333 | 0.043316667 | 0 | Neutral |
| testneg_3423 | -0.02 | 0.28 | 2.201231 | -3 | 1.902977 | 1.902977 | 0.20231 | 0.12788 | 0.18504 | 0.0822 | 12.94465 | 0.04 | 0 | Neutral |
| testneg_3424 | 0.02 | -0.63 | -2.323867 | -3 | 2.259498 | -3 | 0.07595 | 0.11822 | 0.07175 | 0.29293 | 21.25537 | 0.135 | 0 | Neutral |
| testneg_3425 | 0 | 0 | -0.041068 | -0.190389 | 1.259325 | -3 | 0.19898 | 0.07875 | 0.16394 | 0.08512 | 18.23575 | 0.01 | 0 | Neutral |
| testneg_3426 | -0.14 | -0.54 | -0.049099 | -3 | -0.13661 | -3 | 0.16734 | 0.15176 | 0.23624 | 0.12788 | 22.40741 | 0.05 | 0 | Neutral |
| testneg_3427 | 0 | -0.32 | -1.141522 | -3 | 4.231518 | -3 | 0.09555 | 0.09639 | 0.13669 | 0.09263 | 22.31846 | 0.073080139 | 0 | Neutral |
| testneg_3428 | 0 | 0 | 1.650794 | -3 | 0.120012 | 0.120012 | 0.14643 | 0.12788 | 0.20491 | 0.22087 | 10.83333 | 0.040714286 | 0 | Neutral |
| testneg_3429 | 0 | -0.01 | 2.241825 | -3 | -0.151164 | -0.151164 | 0.46318 | 0.14229 | 0.65708 | 0.0936 | 19.34618 | 0.02 | 0 | Neutral |
| testneg_3430 | 0.09 | -0.3 | 0.037077 | -3 | -0.624738 | -3 | 0.09004 | 0.07829 | 0.0819 | 0.1823 | 19.85748 | 0.091244204 | 0 | Neutral |
| testneg_3431 | 0.11 | -0.19 | 2.886405 | -3 | 5.401127 | -3 | 0.62966 | 0.14896 | 0.67796 | 0.10874 | 19.44444 | 0.28 | 2.6 | Neutral |
| testneg_3432 | 0.17 | 0.38 | 5.890584 | -3 | 4.19568 | -3 | 0.10349 | 0.12788 | 0.35513 | 0.21966 | 20.83333 | 0.654961837 | 94.4 | Disease_causing |
| testneg_3433 | -0.06 | -0.18 | 2.785604 | -3 | 0.395766 | -3 | 0.10021 | 0.1105 | 0.98578 | 0.91075 | 8.13938 | 0.1396875 | 0 | Neutral |
| testneg_3434 | 0.04 | -0.23 | 7.185218 | -3 | 1.420291 | -3 | 0.29357 | 0.79524 | 0.19898 | 0.12788 | 11.25 | 0.539714286 | 71.8 | Disease_causing |
| testneg_3435 | 0 | 0 | 0.556564 | -3 | 8.245176 | -3 | 0.18834 | 0.12788 | 0.17969 | 0.11473 | 21.20954 | 0.311185185 | 3.8 | Neutral |
| testneg_3436 | 0.03 | -1.03 | 2.930805 | -3 | 4.939659 | -3 | 0.30942 | 0.13105 | 0.13563 | 0.12378 | 6.43275 | 0.268938087 | 1.8 | Neutral |
| testneg_3437 | 0 | 0 | 0.013394 | -3 | 0.753595 | -3 | 0.12582 | 0.12788 | 0.1273 | 0.12788 | 17.21847 | 0 | 0 | Neutral |
| testneg_3438 | 0.06 | 0.18 | 3.163183 | -3 | 3.112739 | -3 | 0.61343 | 0.11838 | 0.25448 | 0.15814 | 11.98582 | 0.343492252 | 7.6 | Neutral |
| testneg_3439 | 0.16 | 0.08 | 3.167319 | -3 | -0.038552 | -3 | 0.25195 | 0.09584 | 0.0749 | 0.12788 | 9.79693 | 0.04 | 0 | Neutral |
| testneg_3440 | -0.04 | 0.23 | -0.341225 | -3 | 4.003216 | -3 | 0.11329 | 0.12116 | 0.18554 | 0.12299 | 25.47756 | 0.090372414 | 0 | Neutral |
| testneg_3441 | -0.08 | 1.2 | 2.487097 | -3 | 3.854582 | -3 | 0.28485 | 0.12788 | 0.44953 | 0.14403 | 19.90385 | 0.32652381 | 2.4 | Neutral |
| testneg_3442 | -0.03 | -0.1 | 2.059294 | -3 | 6.517429 | -3 | 0.66711 | 0.31094 | 0.29672 | 0.10485 | 33.56481 | 0.262123016 | 1.2 | Neutral |
| testneg_3443 | 0 | -0.01 | 2.053357 | -3 | 5.29955 | -3 | 0.22761 | 0.14229 | 0.36833 | 0.10433 | 9.4445 | 0.1 | 0 | Neutral |
| testneg_3444 | 0.06 | 0.18 | 2.716241 | -3 | -0.009891 | -3 | 0.12591 | 0.09845 | 0.06366 | 0.12788 | 20.68008 | 0.010241841 | 0 | Neutral |
| testneg_3445 | 0 | 0 | -0.22016 | -0.22016 | 0.063598 | -3 | 0.17911 | 0.58893 | 0.1201 | 0.1062 | 32.84553 | 0.073035714 | 0 | Neutral |
| testneg_3446 | -0.07 | -0.33 | 0.985586 | -3 | 2.037511 | -3 | 0.12077 | 0.11244 | 0.71406 | 0.2012 | 18.16092 | 0.075551379 | 0 | Neutral |
| testneg_3447 | 0.18 | -1.05 | 2.368719 | -3 | 6.624966 | -3 | 0.30209 | 0.12788 | 0.08837 | 0.10939 | 19.44444 | 0.24 | 0.4 | Neutral |
| testneg_3448 | -0.02 | -0.45 | 1.825007 | -3 | 0.120012 | -3 | 0.96801 | 0.20777 | 0.20491 | 0.22087 | 10 | 0.091105515 | 0 | Neutral |
| testneg_3449 | 0 | 0 | 1.716798 | -3 | 4.607321 | -3 | 0.70349 | 0.12788 | 0.15406 | 0.1454 | 12.29167 | 0.184615801 | 0 | Neutral |
| testneg_3450 | -0.06 | -0.18 | 4.193908 | -3 | 0.176305 | -3 | 0.23088 | 0.12788 | 0.35513 | 0.21966 | 12.46937 | 0.176561318 | 0 | Neutral |
| testneg_3451 | 0 | -0.01 | 0.389904 | -3 | 3.446463 | 3.446463 | 0.95998 | 0.11262 | 0.1077 | 0.12788 | 18.33333 | 0.184166667 | 0 | Neutral |
| testneg_3452 | 0.14 | -0.37 | 0.198379 | -3 | 0.341911 | -3 | 0.03991 | 0.08342 | 0.99994 | 0.51636 | 10 | 0.14 | 0 | Neutral |
| testneg_3453 | 0.14 | -1.01 | -0.035874 | -3 | 3.095581 | -3 | 0.19898 | 0.12788 | 0.41269 | 0.19135 | 12.70833 | 0.206527778 | 0 | Neutral |
| testneg_3454 | 0.01 | 0.79 | 0.585649 | 0.585649 | -0.056564 | -3 | 0.07185 | 0.68732 | 0.01642 | 0.12788 | 10.38652 | 0.11 | 0 | Neutral |
| testneg_3455 | 0 | 0 | 1.828554 | -3 | 5.777528 | -3 | 0.04208 | 0.12788 | 0.17672 | 0.14229 | 10.41667 | 0.323875 | 5.2 | Neutral |
| testneg_3456 | -0.15 | 1.06 | 0.741676 | -3 | 0.432868 | -3 | 0.07701 | 0.19824 | 0.37292 | 0.11979 | 19.01008 | 0.16 | 0 | Neutral |
| testneg_3457 | 0 | 0 | 0.507327 | -3 | 0.675769 | -3 | 0.11172 | 0.10034 | 0.08723 | 0.09503 | 35.34347 | 0.01 | 0 | Neutral |
| testneg_3458 | 0 | 0 | -0.739513 | -3 | 5.162014 | -3 | 0.39305 | 0.18672 | 0.12642 | 0.11218 | 10.41667 | 0.11478655 | 0 | Neutral |
| testneg_3459 | 0 | 0 | -0.090459 | -0.090459 | 1.548425 | -3 | 0.15581 | 0.19351 | 0.13169 | 0.06895 | 11.66667 | 0.02 | 0 | Neutral |
| testneg_3460 | 0.06 | -0.57 | 6.157422 | -3 | 4.228814 | -3 | 0.19898 | 0.12788 | 0.07675 | 0.11464 | 18.41346 | 0.36 | 12.6 | Neutral |
| testneg_3461 | 0 | -0.01 | 0.895414 | -3 | 0.094994 | -3 | 0.38527 | 0.24812 | 0.56904 | 0.14839 | 10.41667 | 0.041252451 | 0 | Neutral |
| testneg_3462 | 0 | 0 | 0.374605 | -3 | 2.97112 | -3 | 0.45112 | 0.12083 | 0.5901 | 0.19444 | 10.83333 | 0.08 | 0 | Neutral |
| testneg_3463 | 0.05 | 0.29 | -0.618647 | -3 | 3.067866 | -3 | 0.14705 | 0.08702 | 0.04831 | 0.12788 | 24.04872 | 0.07 | 0 | Neutral |
| testneg_3464 | 0 | 0 | 0.976924 | -3 | 4.444691 | -3 | 0.07427 | 0.09451 | 0.32022 | 0.09529 | 12.00474 | 0.044895811 | 0 | Neutral |
| testneg_3465 | 0.12 | -1.14 | 3.586622 | -3 | 3.784977 | -3 | 0.25555 | 0.12094 | 0.07875 | 0.11082 | 17.63514 | 0.131168831 | 0 | Neutral |
| testneg_3466 | 0.03 | -1.03 | 3.292745 | -3 | -0.095386 | -3 | 0.24247 | 0.19222 | 0.12642 | 0.11218 | 11.25 | 0.06 | 0 | Neutral |
| testneg_3467 | -0.02 | 0.68 | 1.148633 | -3 | 0.603592 | -3 | 0.54724 | 0.12788 | 0.08973 | 0.0992 | 16.01246 | 0.03 | 0 | Neutral |
| testneg_3468 | 0 | -0.01 | 0.746602 | -3 | 1.156713 | 1.156713 | 0.20304 | 0.31462 | 0.36458 | 0.16397 | 18.33333 | 0.121463465 | 0 | Neutral |
| testneg_3469 | 0.21 | 0.15 | 1.091899 | -3 | 0.911965 | -0.352465 | 0.25077 | 0.06261 | 0.28685 | 0.1118 | 5.47619 | 0.03 | 0 | Neutral |
| testneg_3470 | 0 | -0.01 | -0.16546 | -3 | 6.715495 | -3 | 0.39399 | 0.12788 | 0.17672 | 0.14229 | 23.65801 | 0.35239411 | 7.6 | Neutral |
| testneg_3471 | 0 | 0 | -0.274465 | -3 | 1.417365 | 1.417365 | 0.04699 | 0.08149 | 0.61184 | 0.55597 | 11.25 | 0.110428977 | 0 | Neutral |
| testneg_3472 | 0 | 0 | 0.991757 | -3 | 5.950905 | -3 | 0.19896 | 0.11604 | 0.19898 | 0.12788 | 14.38008 | 0.07 | 0 | Neutral |
| testneg_3473 | 0.04 | -0.23 | 1.129401 | -3 | 3.59862 | -3 | 0.17301 | 0.14089 | 0.2016 | 0.13956 | 19.44444 | 0.231164877 | 0.2 | Neutral |
| testneg_3474 | 0.21 | 0.15 | 4.83533 | -3 | 6.36911 | -3 | 0.23975 | 0.13861 | 0.17412 | 0.10595 | 18.31857 | 0.463888889 | 41.2 | Neutral |
| testneg_3475 | 0 | 0 | 0.612285 | -3 | 6.625628 | -3 | 0.16078 | 0.39498 | 0.84031 | 0.16749 | 5.69444 | 0.524696928 | 60 | Disease_causing |
| testneg_3476 | 0.08 | 0.03 | -0.020273 | -3 | -1.316616 | -1.316616 | 0.27843 | 0.54325 | 0.07347 | 0.08817 | 18.6768 | 0.083714386 | 0 | Neutral |
| testneg_3477 | 0 | 0 | 0.19536 | -3 | 1.45889 | -3 | 0.25009 | 0.11017 | 0.30209 | 0.14229 | 21.68094 | 0.04 | 0 | Neutral |
| testneg_3478 | -0.18 | 1.05 | 5.508045 | -3 | 4.729514 | -3 | 0.26884 | 0.64999 | 0.19898 | 0.0989 | 31.62471 | 0.56 | 77.4 | Disease_causing |
| testneg_3479 | -0.15 | 0.28 | 0.368178 | -3 | 3.807457 | 1.576651 | 0.13458 | 0.17634 | 0.12513 | 0.12171 | 20.39683 | 0.276241758 | 1.2 | Neutral |
| testneg_3480 | 0.19 | -0.27 | -0.330967 | -3 | 4.129028 | -3 | 0.20657 | 0.12788 | 0.03505 | 0.0907 | 10.28409 | 0.08489899 | 0 | Neutral |
| testneg_3481 | -0.09 | 0.3 | 2.309209 | -3 | 5.008787 | -3 | 0.04863 | 0.12149 | 0.39111 | 0.15887 | 19.09347 | 0.44608039 | 33.2 | Neutral |
| testneg_3482 | -0.06 | -0.18 | 0.491194 | 0.491194 | 1.988072 | -3 | 0.72124 | 0.09906 | 0.35843 | 0.15758 | 7.77778 | 0.081344894 | 0 | Neutral |
| testneg_3483 | -0.21 | -0.15 | -2.232039 | -3 | -0.539741 | -3 | 0.42836 | 0.12788 | 0.10895 | 0.11082 | 20.92347 | 0.06 | 0 | Neutral |
| testneg_3484 | 0.07 | 0.38 | 1.305054 | -3 | 1.310598 | -3 | 0.16248 | 0.14352 | 0.59302 | 0.10897 | 13.37808 | 0.01 | 0 | Neutral |
| testneg_3485 | 0 | 0 | -0.113262 | -3 | 0.627649 | 0.347722 | 0.56154 | 0.0939 | 0.05564 | 0.06933 | 4.44444 | 0.03 | 0 | Neutral |
| testneg_3486 | 0.14 | -1.01 | 3.030637 | -3 | 11.240696 | -3 | 0.26482 | 0.12788 | 0.10241 | 0.08165 | 11.23271 | 0.449027778 | 33.6 | Neutral |
| testneg_3487 | 0 | 0 | 0.724196 | -3 | -0.2352 | -3 | 0.11337 | 0.09363 | 0.19564 | 0.59275 | 10.41667 | 0.04 | 0 | Neutral |
| testneg_3488 | -0.07 | -0.38 | 4.209007 | -3 | 2.084403 | -3 | 0.10724 | 0.49624 | 0.24832 | 0.21196 | 10.41667 | 0.564803571 | 78.4 | Disease_causing |
| testneg_3489 | 0 | 0 | 0.457876 | -3 | 1.959546 | -3 | 0.10379 | 0.34799 | 0.04983 | 0.08759 | 10 | 0.036666667 | 0 | Neutral |
| testneg_3490 | 0 | 0 | 0.085674 | -3 | 4.350853 | -3 | 0.57975 | 0.1248 | 0.12161 | 0.24178 | 26.34995 | 0.296301754 | 1.2 | Neutral |
| testneg_3491 | 0.01 | 0.8 | -0.725111 | -0.725111 | 4.53569 | -3 | 0.05116 | 0.08996 | 0.13559 | 0.1946 | 13.51351 | 0.294391088 | 1.8 | Neutral |
| testneg_3492 | 0 | 0 | 0.453556 | -3 | 3.913294 | -3 | 0.28191 | 0.20693 | 0.94531 | 0.91608 | 5.75579 | 0.401412671 | 21.8 | Neutral |
| testneg_3493 | 0 | 0.32 | -0.420859 | -3 | -2.800363 | -3 | 0.0597 | 0.18092 | 0.44103 | 0.10698 | 24.05425 | 0.063627874 | 0 | Neutral |
| testneg_3494 | 0.08 | -1.2 | 2.008385 | -3 | 0.439346 | -3 | 0.07275 | 0.12788 | 0.06072 | 0.09873 | 24.41367 | 0.03 | 0 | Neutral |
| testneg_3495 | 0 | 0 | 0.254114 | -3 | 2.52259 | -3 | 0.19002 | 0.12138 | 0.5127 | 0.12263 | 13.125 | 0.01 | 0 | Neutral |
| testneg_3496 | 0 | 0.73 | 4.722814 | -3 | 1.415933 | -3 | 0.03008 | 0.12788 | 0.26767 | 0.25506 | 17.21847 | 0.30021401 | 4.2 | Neutral |
| testneg_3497 | 0.18 | -1.05 | 4.454666 | -3 | 3.557638 | -3 | 0.09462 | 0.11807 | 0.15867 | 0.09641 | 23.81804 | 0.131428571 | 0 | Neutral |
| testneg_3498 | 0 | 0 | 1.003734 | -3 | 2.428041 | -3 | 0.11325 | 0.17021 | 0.07606 | 0.13082 | 18.62854 | 0.1 | 0 | Neutral |
| testneg_3499 | 0.16 | 0.08 | 0.076953 | -3 | 1.164204 | -3 | 0.10187 | 0.12339 | 0.19898 | 0.12788 | 28.47222 | 0.01 | 0 | Neutral |
| testneg_3500 | 0 | 0 | 0.014187 | -3 | 5.007683 | -3 | 0.99998 | 0.9588 | 0.07415 | 0.12788 | 16.69775 | 0.180323427 | 0 | Neutral |
| testneg_3501 | 0 | 0 | 0.09433 | -3 | 0.728537 | -3 | 0.18925 | 0.32479 | 0.08521 | 0.11239 | 7.00579 | 0.01037037 | 0 | Neutral |
| testneg_3502 | 0 | 0 | 1.568756 | -3 | 6.206238 | 6.041443 | 0.9516 | 0.12788 | 0.10703 | 0.11611 | 23.22141 | 0.380384615 | 7.4 | Neutral |
| testneg_3503 | -0.01 | -0.8 | 5.486648 | -3 | -0.275748 | -3 | 0.16708 | 0.11594 | 0.13779 | 0.0945 | 21.3173 | 0.12 | 0 | Neutral |
| testneg_3504 | -0.0356364 | 0.22672727 | -1.616402 | -3 | 2.209838 | -3 | 0.07837 | 0.12788 | 0.11261 | 0.13673 | 18.94049 | 0.1 | 0 | Neutral |
| testneg_3505 | 0 | 0 | 0.599522 | -3 | 0.299382 | -3 | 0.63363 | 0.14795 | 0.75728 | 0.34838 | 6.00877 | 0.030877897 | 0 | Neutral |
| testneg_3506 | 0 | 0 | -0.22828 | -3 | 1.616146 | -3 | 0.31694 | 0.13475 | 0.41953 | 0.44852 | 21.53664 | 0.092494939 | 0 | Neutral |
| testneg_3507 | 0 | 0 | 1.362887 | -3 | -1.56406 | -3 | 0.38527 | 0.24812 | 0.13796 | 0.27869 | 10.22898 | 0.071496354 | 0 | Neutral |
| testneg_3508 | -0.08 | -0.03 | 5.412199 | -3 | -0.057969 | -3 | 0.18165 | 0.49827 | 0.68233 | 0.58998 | 4.44444 | 0.567238095 | 76.4 | Disease_causing |
| testneg_3509 | 0.03 | -0.85 | 0.173309 | -3 | -1.171244 | -1.171244 | 0.34694 | 0.20397 | 0.19898 | 0.12788 | 18.73163 | 0.04 | 0 | Neutral |
| testneg_3510 | -0.06 | 1.29 | 4.344772 | -3 | 4.032469 | -3 | 0.34404 | 0.13209 | 0.06895 | 0.06354 | 20.83333 | 0.270557276 | 0.4 | Neutral |
| testneg_3511 | 0 | 0 | 1.505786 | -3 | -0.437316 | -3 | 0.19898 | 0.10591 | 0.20231 | 0.12788 | 9.6023 | 0 | 0 | Neutral |
| testneg_3512 | -0.01 | 0.57 | 0.083768 | -3 | 0.579303 | -3 | 0.08041 | 0.21802 | 0.2852 | 0.12788 | 4.44444 | 0.05 | 0 | Neutral |
| testneg_3513 | -0.06 | -0.18 | 1.651093 | -3 | 1.940661 | -3 | 0.08332 | 0.09046 | 0.25151 | 0.12788 | 13.11539 | 0.01 | 0 | Neutral |
| testneg_3514 | 0.02 | -0.63 | 3.47892 | -3 | 5.495658 | -3 | 0.18423 | 0.10808 | 0.15294 | 0.12788 | 18.8582 | 0.239886364 | 0.6 | Neutral |
| testneg_3515 | 0 | -0.01 | 0.639661 | -3 | 0.62743 | -3 | 0.16263 | 0.53493 | 0.08878 | 0.0974 | 29.52203 | 0.05 | 0 | Neutral |
| testneg_3516 | -0.05 | -0.29 | 0.116519 | 0.116519 | -0.457394 | -3 | 0.2207 | 0.09998 | 0.19898 | 0.46867 | 12.47902 | 0.08 | 0 | Neutral |
| testneg_3517 | 0 | 0.32 | 3.274084 | 0.450036 | 0.330554 | -3 | 0.44479 | 0.20004 | 0.06431 | 0.12788 | 21.88889 | 0.109571268 | 0 | Neutral |
| testneg_3518 | 0.18 | -1.05 | 3.050835 | -3 | 1.484366 | -3 | 0.49512 | 0.08696 | 0.50592 | 0.0952 | 13.40441 | 0.07 | 0 | Neutral |
| testneg_3519 | 0.14 | -1.01 | 2.126776 | -3 | 0.361904 | -1.390553 | 0.9541 | 0.96732 | 0.08789 | 0.6361 | 6.0461 | 0.249722222 | 0 | Neutral |
| testneg_3520 | -0.04 | 0.23 | 0.190526 | -3 | 2.413901 | -3 | 0.1834 | 0.12788 | 0.1228 | 0.10482 | 33.3703 | 0.03 | 0 | Neutral |
| testneg_3521 | -0.14 | 0.69 | 2.798182 | 2.798182 | 1.967428 | -3 | 0.31043 | 0.09688 | 0.42797 | 0.17974 | 6.87637 | 0.521048799 | 62 | Disease_causing |
| testneg_3522 | 0 | 0 | 0.412565 | -3 | 1.28208 | -3 | 0.37499 | 0.17803 | 0.38375 | 0.09256 | 13.80482 | 0.01 | 0 | Neutral |
| testneg_3523 | -0.02 | -0.41 | 0.579456 | 0.579456 | 4.492963 | -3 | 0.47678 | 0.38085 | 0.54648 | 0.19042 | 9.80134 | 0.514905348 | 59.8 | Disease_causing |
| testneg_3524 | 0 | 0 | 0.173288 | -3 | 1.907441 | 1.907441 | 0.61211 | 0.11059 | 0.20708 | 0.12788 | 20.91877 | 0.04 | 0 | Neutral |
| testneg_3525 | 0.01 | 0.79 | 3.986254 | -3 | 1.242354 | -3 | 0.16797 | 0.12788 | 0.66381 | 0.13039 | 26.34995 | 0.080901261 | 0 | Neutral |
| testneg_3526 | 0 | -0.73 | 2.726423 | -3 | 0.837358 | -3 | 0.14273 | 0.1162 | 0.8433 | 0.13128 | 10.82179 | 0.05 | 0 | Neutral |
| testneg_3527 | 0 | 0 | 0.29846 | -3 | 1.098308 | -3 | 0.07543 | 0.10211 | 0.09346 | 0.12788 | 28.13493 | 0.01 | 0 | Neutral |
| testneg_3528 | 0.07 | 0.01 | 1.001127 | -3 | -0.235499 | -3 | 0.14 | 0.12151 | 0.1203 | 0.13872 | 12.29167 | 0.041666667 | 0 | Neutral |
| testneg_3529 | 0 | 0 | 1.432106 | -3 | 5.588206 | -3 | 0.11325 | 0.17021 | 0.16662 | 0.11306 | 29.03465 | 0.177261905 | 0 | Neutral |
| testneg_3530 | 0.18 | 0.16 | 5.112733 | -3 | -2.351627 | -3 | 0.03676 | 0.12788 | 0.16646 | 0.16629 | 36.69077 | 0.307916667 | 2.2 | Neutral |
| testneg_3531 | -0.07 | -0.38 | -0.763372 | -3 | 5.035077 | -3 | 0.19979 | 0.09328 | 0.34109 | 0.2167 | 17.21847 | 0.282115801 | 1.2 | Neutral |
| testneg_3532 | -0.03 | -0.1 | 1.080231 | -3 | 2.304681 | -3 | 0.05996 | 0.12788 | 0.79983 | 0.1105 | 43.86228 | 0.040345238 | 0 | Neutral |
| testneg_3533 | 0.18 | 0.16 | -1.950188 | -3 | 1.102485 | -3 | 0.09273 | 0.08316 | 0.25731 | 0.12662 | 12.50388 | 0.03 | 0 | Neutral |
| testneg_3534 | 0.01 | -0.57 | 6.366826 | -3 | 5.362959 | -3 | 0.06781 | 0.09009 | 0.31507 | 0.61057 | 9.70275 | 0.685690162 | 95.4 | Disease_causing |
| testneg_3535 | 0.03 | 0.12 | 2.332457 | -3 | 1.828554 | -3 | 0.358 | 0.11262 | 0.04208 | 0.12788 | 20.27778 | 0.010801282 | 0 | Neutral |
| testneg_3536 | 0 | 0 | -0.35638 | -3 | 1.902068 | -3 | 0.18633 | 0.14783 | 0.44382 | 0.33284 | 12.29167 | 0.065619383 | 0 | Neutral |
| testneg_3537 | 0 | 0 | 1.157752 | -3 | -0.4837 | -3 | 0.10569 | 0.09059 | 0.23276 | 0.31401 | 27.4024 | 0.06 | 0 | Neutral |

| | | | | | | | | | | | | | | |
|---|---|---|---|---|---|---|---|---|---|---|---|---|---|---|
| testneg_3538 | -0.17 | 1.22 | -2.338688 | -2.338688 | 0.22996 | -3 | 0.33129 | 0.09957 | 0.0693 | 0.09293 | 18.88889 | 0.1 | 0 | Neutral |
| testneg_3539 | 0 | 0 | 0.391204 | -3 | 1.887787 | -3 | 0.19898 | 0.13162 | 0.51398 | 0.38821 | 11.28501 | 0.051825397 | 0 | Neutral |
| testneg_3540 | 0.05 | -0.81 | 3.383188 | -3 | 1.078539 | -3 | 0.29356 | 0.47581 | 0.27077 | 0.10337 | 29.16667 | 0.11167003 | 0 | Neutral |
| testneg_3541 | -0.06 | 0.57 | 1.26369 | -3 | 3.530544 | -3 | 0.19898 | 0.12788 | 0.119 | 0.0978 | 34.44629 | 0.07 | 0 | Neutral |
| testneg_3542 | 0 | -0.32 | -2.498179 | -3 | 2.103651 | -3 | 0.50682 | 0.12788 | 0.13521 | 0.10126 | 16.43865 | 0.03 | 0 | Neutral |
| testneg_3543 | 0.02 | 0.41 | 3.953483 | -3 | 2.061627 | -3 | 0.97273 | 0.27711 | 0.13271 | 0.22958 | 13.18841 | 0.35884316 | 7.6 | Neutral |
| testneg_3544 | -0.22 | 2.66 | 0.984298 | -3 | 0.552009 | -3 | 0.34406 | 0.09894 | 0.53172 | 0.12788 | 13.51351 | 0.16 | 0 | Neutral |
| testneg_3545 | 0.01 | 0.8 | -0.516038 | -0.516038 | 4.652089 | -3 | 0.19898 | 0.12788 | 0.23005 | 0.37216 | 25.86756 | 0.327595727 | 4.4 | Neutral |
| testneg_3546 | 0.07 | 0.7 | 2.976868 | -3 | 0.942734 | -3 | 0.19898 | 0.12788 | 0.78537 | 0.33817 | 15.03769 | 0.145136986 | 0 | Neutral |
| testneg_3547 | 0.09 | -0.3 | 0.094994 | -3 | 0.343309 | -3 | 0.56904 | 0.14839 | 0.36857 | 0.36135 | 10.83333 | 0.135892857 | 0 | Neutral |
| testneg_3548 | -0.07 | -0.38 | 2.173191 | -3 | 0.089405 | -3 | 0.18233 | 0.14781 | 0.06301 | 0.32138 | 18.88889 | 0.15 | 0 | Neutral |
| testneg_3549 | 0 | 0 | 1.941287 | -3 | 2.493168 | -3 | 0.24977 | 0.11262 | 0.14897 | 0.19122 | 12.58479 | 0.090979167 | 0 | Neutral |
| testneg_3550 | 0.03 | 0.1 | -0.948009 | -3 | 2.531224 | -3 | 0.10379 | 0.34799 | 0.13061 | 0.21023 | 20.84487 | 0.339594329 | 4.2 | Neutral |
| testneg_3551 | -0.02 | 0.63 | 3.995565 | -3 | 2.433198 | -3 | 0.31773 | 0.10816 | 0.11657 | 0.21661 | 19.65091 | 0.349893875 | 6.8 | Neutral |
| testneg_3552 | 0 | 0 | 0.447219 | -3 | 3.05843 | 3.05843 | 0.13884 | 0.12788 | 0.74136 | 0.1049 | 19.09561 | 0.122479417 | 0 | Neutral |
| testneg_3553 | 0 | 0 | 1.193153 | -3 | 4.31851 | -3 | 0.12259 | 0.1041 | 0.04694 | 0.44682 | 14.95654 | 0.23316619 | 0.8 | Neutral |
| testneg_3554 | -0.15 | 0.82 | 2.976115 | -3 | 6.24493 | -3 | 0.40465 | 0.12971 | 0.19301 | 0.10828 | 11.83502 | 0.390094086 | 15.4 | Neutral |
| testneg_3555 | 0.17 | 0.38 | -1.528753 | -1.528753 | -1.110257 | -3 | 0.10482 | 0.12788 | 0.15217 | 0.12544 | 12.70833 | 0.020481859 | 0 | Neutral |
| testneg_3556 | 0.04 | 2.01 | 3.380302 | -3 | 5.739294 | -3 | 0.18281 | 0.12788 | 0.07701 | 0.19824 | 23.68839 | 0.560905158 | 74.8 | Disease_causing |
| testneg_3557 | 0 | 0 | 0.701413 | -3 | 4.026354 | -3 | 0.1214 | 0.0899 | 0.25422 | 0.10102 | 18.33333 | 0.04 | 0 | Neutral |
| testneg_3558 | 0.09 | -0.3 | 6.132176 | -3 | 6.319089 | -3 | 0.51577 | 0.19561 | 0.19898 | 0.12788 | 21.01658 | 0.680045872 | 97.2 | Disease_causing |
| testneg_3559 | 0 | 0 | -0.032453 | -3 | 0.195289 | -3 | 0.31664 | 0.14181 | 0.09972 | 0.2187 | 20 | 0.111654657 | 0 | Neutral |
| testneg_3560 | 0 | 0 | 0.36941 | -3 | 0.60982 | 0.60982 | 0.15406 | 0.1454 | 0.23844 | 0.27772 | 10.26553 | 0.059371817 | 0 | Neutral |
| testneg_3561 | 0 | 0.32 | 3.274084 | -3 | 0.335227 | -3 | 0.44479 | 0.20004 | 0.04346 | 0.10785 | 10.83333 | 0.061396104 | 0 | Neutral |
| testneg_3562 | 0.00281818 | -0.0195455 | 2.555582 | -3 | 2.46715 | -3 | 0.65012 | 0.21208 | 0.20951 | 0.16278 | 10.41667 | 0.259063417 | 0.8 | Neutral |
| testneg_3563 | 0.07 | 0.38 | 0.991286 | -3 | 2.706663 | -3 | 0.2439 | 0.08848 | 0.09555 | 0.09639 | 15.34131 | 0.04 | 0 | Neutral |
| testneg_3564 | 0 | 0 | 1.332007 | -3 | 3.203428 | -3 | 0.18785 | 0.13588 | 0.35325 | 0.12614 | 12.37434 | 0.03547027 | 0 | Neutral |
| testneg_3565 | 0 | 0 | 0.577899 | -3 | 1.617668 | -3 | 0.54231 | 0.15254 | 0.09273 | 0.08316 | 18.48597 | 0.02 | 0 | Neutral |
| testneg_3566 | -0.03 | -1.22 | 2.92942 | -3 | 0.774415 | -3 | 0.15107 | 0.10457 | 0.90112 | 0.20622 | 10.83333 | 0.138856209 | 0 | Neutral |
| testneg_3567 | 0.14 | -0.37 | 4.619073 | -3 | -0.165659 | -3 | 0.16125 | 0.08406 | 0.19698 | 0.1459 | 7.20975 | 0.10447002 | 0 | Neutral |
| testneg_3568 | -0.01 | 1.44 | -0.27139 | -3 | 2.395534 | -3 | 0.37637 | 0.12788 | 0.13967 | 0.15696 | 21.46862 | 0.219379316 | 0 | Neutral |
| testneg_3569 | -0.06 | 0.57 | -0.421956 | -3 | 7.20297 | 2.266625 | 0.2049 | 0.12678 | 0.19898 | 0.12788 | 19.09347 | 0.34 | 6.4 | Neutral |
| testneg_3570 | -0.05 | -0.29 | -1.851788 | -3 | 5.284085 | -3 | 0.78741 | 0.22085 | 0.08509 | 0.12788 | 10.22898 | 0.181216358 | 0 | Neutral |
| testneg_3571 | -0.03 | -0.1 | 3.804607 | -3 | 0.696647 | -3 | 0.36694 | 0.12788 | 0.25731 | 0.12662 | 27.06929 | 0.03 | 0 | Neutral |
| testneg_3572 | 0 | 0 | 1.554241 | -3 | -0.843404 | -3 | 0.35412 | 0.10819 | 0.11587 | 0.12001 | 11.25 | 0 | 0 | Neutral |
| testneg_3573 | 0 | 0 | 1.263979 | -3 | 4.142037 | -3 | 0.14901 | 0.12788 | 0.95535 | 0.12788 | 11.25 | 0.061590599 | 0 | Neutral |
| testneg_3574 | 0 | 0 | -0.137475 | -3 | 1.344878 | -3 | 0.19898 | 0.12788 | 0.05444 | 0.12788 | 27.46106 | 0.013095238 | 0 | Neutral |
| testneg_3575 | 0.08 | 0.03 | -0.765287 | -3 | 0.000331 | -3 | 0.21002 | 0.12788 | 0.14432 | 0.22579 | 16.89313 | 0.061417815 | 0 | Neutral |
| testneg_3576 | 0 | 0 | 0.559436 | -3 | 1.418339 | -3 | 0.77944 | 0.1858 | 0.24185 | 0.12788 | 11.25 | 0.01 | 0 | Neutral |
| testneg_3577 | 0 | 0 | 1.108803 | -3 | 1.467396 | -3 | 0.85092 | 0.58871 | 0.96995 | 0.66487 | 5 | 0.061894598 | 0 | Neutral |
| testneg_3578 | 0 | -0.01 | 0.39182 | -3 | -0.195051 | -3 | 0.09432 | 0.15749 | 0.36417 | 0.21506 | 5.69444 | 0.06496408 | 0 | Neutral |
| testneg_3579 | 0 | 0 | 1.008465 | -3 | 0.628811 | 0.628811 | 0.45299 | 0.16306 | 0.07874 | 0.12788 | 22.87765 | 0.031480735 | 0 | Neutral |
| testneg_3580 | 0 | 0 | 0.708402 | -3 | 1.75958 | 1.75958 | 0.20765 | 0.27124 | 0.65897 | 0.67602 | 10.41667 | 0.138718644 | 0 | Neutral |
| testneg_3581 | 0 | 0 | 1.095746 | -3 | 3.25138 | -3 | 0.24274 | 0.13034 | 0.1533 | 0.13168 | 17.21847 | 0.13401946 | 0 | Neutral |
| testneg_3582 | -0.03 | -0.1 | 6.638252 | -3 | -0.745211 | -3 | 0.24452 | 0.21203 | 0.12832 | 0.07113 | 29.16667 | 0.31 | 3.6 | Neutral |
| testneg_3583 | 0 | 0 | 0.740665 | -3 | 2.018667 | -3 | 0.23278 | 0.11027 | 0.39344 | 0.12788 | 12.79101 | 0 | 0 | Neutral |
| testneg_3584 | 0 | 0 | 1.241234 | -3 | -0.338929 | -3 | 0.6258 | 0.16461 | 0.6323 | 0.09928 | 10.83333 | 0.02 | 0 | Neutral |
| testneg_3585 | 0 | -0.01 | 1.806927 | -3 | 0.64288 | -3 | 0.17973 | 0.12278 | 0.57188 | 0.34038 | 6.94444 | 0.03 | 0 | Neutral |
| testneg_3586 | 0.04 | -0.23 | 8.131269 | -3 | 5.315177 | 5.315177 | 0.59342 | 0.12788 | 0.12091 | 0.13585 | 10.41667 | 0.814974311 | 100 | Disease_causing |
| testneg_3587 | 0.02 | 0.41 | 3.953483 | -3 | 5.555682 | -3 | 0.97273 | 0.27711 | 0.54147 | 0.20818 | 13.39912 | 0.730491947 | 99.4 | Disease_causing |
| testneg_3588 | -0.03 | -0.1 | 1.750131 | -3 | 3.50895 | -3 | 0.85075 | 0.16646 | 0.28165 | 0.15912 | 10.22898 | 0.292234147 | 2.6 | Neutral |
| testneg_3589 | 0.09 | -0.3 | 2.550986 | 1.466236 | 1.06169 | -3 | 0.08408 | 0.12788 | 0.34673 | 0.24308 | 19.09347 | 0.199511218 | 0 | Neutral |
| testneg_3590 | 0 | 0 | 0.116245 | -3 | -1.118982 | -3 | 0.8238 | 0.38836 | 0.32144 | 0.11774 | 10.83333 | 0.02 | 0 | Neutral |
| testneg_3591 | 0 | 0 | 0.258619 | -3 | 0.603046 | -3 | 0.30823 | 0.17188 | 0.10249 | 0.1399 | 10.41667 | 0.079202781 | 0 | Neutral |
| testneg_3592 | 0 | 0 | 0.977938 | -3 | -0.347591 | -3 | 0.47975 | 0.18568 | 0.32094 | 0.36254 | 10.83333 | 0.075384615 | 0 | Neutral |
| testneg_3593 | -0.03 | -0.1 | 5.488834 | -3 | -0.02246 | -3 | 0.38115 | 0.21061 | 0.19898 | 0.12788 | 18.88889 | 0.26665 | 1.4 | Neutral |
| testneg_3594 | 0 | 0 | 1.825331 | -3 | 1.524427 | -3 | 0.13029 | 0.12788 | 0.99317 | 0.60884 | 4.72222 | 0.060119868 | 0 | Neutral |
| testneg_3595 | -0.06 | -0.18 | 2.775182 | -3 | 3.587651 | -3 | 0.59342 | 0.12788 | 0.13615 | 0.12788 | 11.60901 | 0.099548402 | 0 | Neutral |
| testneg_3596 | 0 | 0 | 0.053978 | -3 | 0.944356 | -3 | 0.1289 | 0.14878 | 0.32532 | 0.09651 | 11.875 | 0.02 | 0 | Neutral |
| testneg_3597 | 0 | 0 | 2.007328 | -3 | 7.107534 | -3 | 0.26264 | 0.13812 | 0.11187 | 0.12788 | 4.72222 | 0.337182589 | 6.4 | Neutral |
| testneg_3598 | 0 | 0 | 2.147911 | -3 | -1.561749 | -3 | 0.39087 | 0.10381 | 0.76186 | 0.10052 | 17.3494 | 0.00225 | 0 | Neutral |
| testneg_3599 | -0.08 | -0.03 | 4.471379 | -3 | 0.318861 | -3 | 0.17404 | 0.12788 | 0.1617 | 0.17928 | 19.8374 | 0.212940171 | 0.2 | Neutral |
| testneg_3600 | -0.01 | -0.8 | 2.675128 | -3 | 0.332327 | -3 | 0.19898 | 0.12788 | 0.83955 | 0.10333 | 22.64938 | 0.02 | 0 | Neutral |
| testneg_3601 | 0.02 | -0.63 | -0.055258 | -3 | -0.758692 | -3 | 0.16139 | 0.09116 | 0.63553 | 0.26358 | 20.27778 | 0.060374408 | 0 | Neutral |
| testneg_3602 | 0 | 0 | -0.22828 | -3 | 6.535837 | -3 | 0.31694 | 0.13475 | 0.31434 | 0.11315 | 18.88889 | 0.145610606 | 0 | Neutral |
| testneg_3603 | 0.07 | 0.38 | 0.329293 | -3 | 4.715095 | -3 | 0.23809 | 0.18937 | 0.17016 | 0.12748 | 20.4583 | 0.197666667 | 0 | Neutral |
| testneg_3604 | -0.17 | -0.38 | -0.695669 | -3 | 0.71132 | -3 | 0.17818 | 0.15728 | 0.4722 | 0.13321 | 28.84995 | 0.161658508 | 0 | Neutral |
| testneg_3605 | 0 | 0 | 0.654716 | -3 | 5.582498 | -3 | 0.76942 | 0.13267 | 0.85829 | 0.4292 | 20.83333 | 0.297767262 | 1.8 | Neutral |
| testneg_3606 | 0 | 0 | 1.528973 | -3 | 4.515915 | -3 | 0.50403 | 0.21568 | 0.49644 | 0.10958 | 27.31171 | 0.114 | 0 | Neutral |
| testneg_3607 | 0 | 0 | 0.720647 | -3 | -0.761124 | -3 | 0.161 | 0.11597 | 0.08465 | 0.15428 | 18.6768 | 0.056891026 | 0 | Neutral |
| testneg_3608 | 0.07 | 0.38 | 3.807502 | 3.807502 | 0.39745 | -3 | 0.34673 | 0.24308 | 0.11404 | 0.12788 | 10.29033 | 0.47 | 42.6 | Neutral |
| testneg_3609 | 0 | 0 | 1.229821 | -3 | 1.962395 | -3 | 0.46091 | 0.13852 | 0.45399 | 0.10245 | 28.29365 | 0.02 | 0 | Neutral |
| testneg_3610 | 0 | 0 | 3.506817 | -3 | 0.520281 | -3 | 0.23088 | 0.21602 | 0.1336 | 0.15669 | 12.22222 | 0.218397432 | 0 | Neutral |
| testneg_3611 | 0.08 | 0.03 | 0.172519 | -3 | 5.495658 | -3 | 0.85887 | 0.1489 | 0.15294 | 0.12788 | 10 | 0.174665452 | 0 | Neutral |
| testneg_3612 | -0.07 | -0.38 | 1.276657 | -3 | 2.128864 | -3 | 0.98565 | 0.23629 | 0.20655 | 0.15317 | 14.62121 | 0.142052759 | 0 | Neutral |
| testneg_3613 | 0.14 | -0.69 | 3.257071 | -3 | 0.018319 | -3 | 0.59864 | 0.14106 | 0.3828 | 0.5569 | 13.57684 | 0.233892643 | 0.2 | Neutral |
| testneg_3614 | 0 | 0 | 1.941287 | -3 | 1.107552 | -3 | 0.24977 | 0.11262 | 0.11687 | 0.09069 | 24.66933 | 0.01 | 0 | Neutral |
| testneg_3615 | -0.03 | -0.1 | 7.121043 | -3 | 4.819528 | 4.819528 | 0.35945 | 0.12025 | 0.1211 | 0.1526 | 21.94444 | 0.759086066 | 100 | Disease_causing |
| testneg_3616 | 0 | 0 | 1.075324 | 1.075324 | 0.097083 | -3 | 0.49934 | 0.11069 | 0.12943 | 0.14391 | 13.98631 | 0.08 | 0 | Neutral |
| testneg_3617 | -0.07 | 0.31 | -0.564575 | -3 | 1.750565 | -3 | 0.68029 | 0.12788 | 0.19898 | 0.12788 | 12.79101 | 0.02 | 0 | Neutral |
| testneg_3618 | 0.09 | -0.25 | 1.250879 | -3 | 1.664239 | -3 | 0.17325 | 0.17964 | 0.08814 | 0.42357 | 14.35071 | 0.168827506 | 0 | Neutral |
| testneg_3619 | 0 | 0 | 0.616139 | -3 | 3.348627 | -3 | 0.47997 | 0.10506 | 0.04808 | 0.10018 | 16.47059 | 0.05 | 0 | Neutral |
| testneg_3620 | 0.14 | -1.01 | 2.428056 | -3 | 2.032988 | -3 | 0.08684 | 0.08917 | 0.19087 | 0.44337 | 12.3206 | 0.138666667 | 0 | Neutral |
| testneg_3621 | 0.04454545 | -0.2783636 | 9.766075 | -3 | 1.800429 | 1.568815 | 0.10021 | 0.1105 | 0.14379 | 0.12788 | 15.95907 | 0.350339862 | 10.8 | Neutral |
| testneg_3622 | -0.01 | -0.79 | 1.459804 | -3 | 1.446802 | -3 | 0.84106 | 0.63405 | 0.21115 | 0.17723 | 9.04816 | 0.106771529 | 0 | Neutral |
| testneg_3623 | 0.08 | 0.03 | 0.628497 | -3 | 1.800429 | 1.568815 | 0.39344 | 0.12788 | 0.14379 | 0.12788 | 12.01754 | 0.04 | 0 | Neutral |
| testneg_3624 | 0.02 | 0.45 | 4.160232 | -3 | 1.894135 | -3 | 0.14484 | 0.14928 | 0.75606 | 0.18509 | 17.77778 | 0.25109127 | 0.4 | Neutral |
| testneg_3625 | -0.04 | 0.23 | 2.272008 | -3 | -0.05199 | -3 | 0.23913 | 0.12788 | 0.13413 | 0.12788 | 17.21847 | 0.01 | 0 | Neutral |
| testneg_3626 | 0 | 0 | -0.211085 | -3 | 2.928857 | 2.928857 | 0.04543 | 0.28103 | 0.11534 | 0.12281 | 18.88889 | 0.192403846 | 0 | Neutral |
| testneg_3627 | 0 | 0 | -0.346224 | -3 | -3.299404 | -3 | 0.14632 | 0.12662 | 0.20849 | 0.12788 | 20.27778 | 0.01 | 0 | Neutral |
| testneg_3628 | 0.06 | -0.57 | -0.383439 | -3 | 2.328483 | -3 | 0.23152 | 0.09962 | 0.1577 | 0.2732 | 33.51366 | 0.120916667 | 0 | Neutral |
| testneg_3629 | 0 | 0 | 0.191318 | -3 | 5.0589 | -3 | 0.96113 | 0.24974 | 0.09658 | 0.11286 | 23.4681 | 0.115788177 | 0 | Neutral |
| testneg_3630 | 0.18 | 0.16 | -2.635087 | -3 | 7.038327 | -3 | 0.19898 | 0.12788 | 0.30367 | 0.41964 | 20.30197 | 0.45127005 | 37.6 | Neutral |
| testneg_3631 | 0.04 | -0.23 | 3.764855 | -3 | 6.71239 | -3 | 0.44067 | 0.11201 | 0.68568 | 0.65219 | 12.46106 | 0.661999008 | 94.6 | Disease_causing |
| testneg_3632 | 0 | 0 | 2.171631 | -3 | 3.980559 | -3 | 0.94859 | 0.75045 | 0.06723 | 0.10296 | 4.72222 | 0.127105311 | 0 | Neutral |
| testneg_3633 | 0 | 0 | 3.506817 | -3 | 2.191878 | -3 | 0.23088 | 0.21602 | 0.18127 | 0.12788 | 15.59524 | 0.101833333 | 0 | Neutral |
| testneg_3634 | 0.04 | -0.23 | 4.771654 | -3 | -2.240872 | -3 | 0.6201 | 0.12788 | 0.14175 | 0.12788 | 11.25 | 0.053623659 | 0 | Neutral |
| testneg_3635 | -0.07 | -0.38 | 3.645998 | -3 | 2.682586 | -3 | 0.19898 | 0.12788 | 0.10383 | 0.08853 | 24.99104 | 0.070819918 | 0 | Neutral |
| testneg_3636 | -0.03 | -0.12 | 1.79191 | -3 | -0.227871 | -3 | 0.19364 | 0.14027 | 0.44654 | 0.10907 | 33.77193 | 0.02 | 0 | Neutral |
| testneg_3637 | -0.21 | -0.15 | 0.197741 | -3 | 4.202319 | -3 | 0.31923 | 0.17492 | 0.19898 | 0.30319 | 9.18689 | 0.490400924 | 50.6 | Disease_causing |
| testneg_3638 | -0.01 | -0.79 | -0.151787 | -3 | 7.291769 | -3 | 0.85984 | 0.40044 | 0.7351 | 0.16019 | 6.42207 | 0.488470509 | 50 | Neutral |
| testneg_3639 | -0.16 | 0.37 | 5.039547 | -3 | 1.437875 | -3 | 0.42512 | 0.29333 | 0.16636 | 0.12384 | 21.25537 | 0.235 | 0.4 | Neutral |
| testneg_3640 | -0.0110909 | 0.08709091 | 2.966005 | -3 | 4.617349 | -3 | 0.26192 | 0.15452 | 0.16436 | 0.19109 | 26.9055 | 0.628469557 | 93.2 | Disease_causing |
| testneg_3641 | 0 | 0 | 6.139143 | -3 | -0.21059 | -3 | 0.26372 | 0.12788 | 0.03443 | 0.1032 | 10.41667 | 0.19 | 0.4 | Neutral |
| testneg_3642 | -0.08 | -0.03 | 4.455854 | -3 | 2.091915 | -3 | 0.10822 | 0.10942 | 0.98238 | 0.10873 | 5.89559 | 0.093333333 | 0 | Neutral |
| testneg_3643 | 0 | 0 | 1.474437 | -3 | 0.822036 | -3 | 0.54648 | 0.19042 | 0.13575 | 0.12788 | 6.57332 | 0.003389356 | 0 | Neutral |
| testneg_3644 | 0 | 0 | 0.237368 | -3 | 3.064064 | -3 | 0.1602 | 0.11262 | 0.5481 | 0.10207 | 28.52352 | 0.03 | 0 | Neutral |
| testneg_3645 | 0 | 0 | 0.870324 | -3 | -0.771963 | -3 | 0.16355 | 0.12788 | 0.14668 | 0.17962 | 22.93219 | 0.052746246 | 0 | Neutral |
| testneg_3646 | 0.03 | 0.12 | 1.451994 | -3 | -0.513734 | -3 | 0.07828 | 0.76259 | 0.256 | 0.20036 | 10.41667 | 0.180550725 | 0 | Neutral |
| testneg_3647 | 0 | 0 | 0.719031 | -3 | 1.645732 | -3 | 0.9924 | 0.37896 | 0.09997 | 0.09515 | 12.43093 | 0.032098391 | 0 | Neutral |
| testneg_3648 | -0.21 | -0.15 | 3.884876 | -3 | -0.392743 | -3 | 0.29004 | 0.45465 | 0.16275 | 0.12788 | 10.41667 | 0.151833333 | 0 | Neutral |
| testneg_3649 | -0.05 | 0.81 | 1.849303 | -3 | 2.022528 | -3 | 0.59947 | 0.10764 | 0.57953 | 0.15513 | 34.05449 | 0.09 | 0 | Neutral |
| testneg_3650 | 0 | 0 | -0.22828 | -3 | 0.985143 | -3 | 0.31694 | 0.13475 | 0.10042 | 0.09296 | 18.55633 | 0.02 | 0 | Neutral |
| testneg_3651 | -0.01 | -0.8 | 4.869565 | -3 | 2.445411 | -3 | 0.0893 | 0.11207 | 0.22179 | 0.09831 | 18.6768 | 0.1 | 0 | Neutral |
| testneg_3652 | 0.04 | -0.23 | 0.21324 | -3 | 2.475186 | -3 | 0.46664 | 0.09008 | 0.9581 | 0.12788 | 18.88889 | 0.04 | 0 | Neutral |
| testneg_3653 | 0.03 | -1.03 | 3.292745 | -3 | 2.477566 | -3 | 0.24247 | 0.19222 | 0.29451 | 0.13582 | 18.6768 | 0.357993837 | 12.8 | Neutral |
| testneg_3654 | 0.03 | 0.12 | 4.361468 | -3 | 0.188506 | -3 | 0.39015 | 0.23372 | 0.54826 | 0.24974 | 10.43529 | 0.291481331 | 2.6 | Neutral |
| testneg_3655 | -0.05 | -0.29 | 5.464524 | -3 | 3.029702 | -3 | 0.31507 | 0.61057 | 0.38942 | 0.345 | 9.70275 | 0.877083333 | 100 | Disease_causing |
| testneg_3656 | -0.03 | -0.1 | 2.928572 | -3 | 0.839935 | -3 | 0.60285 | 0.15179 | 0.17412 | 0.10595 | 4.99231 | 0.050594207 | 0 | Neutral |
| testneg_3657 | -0.01 | 0.57 | 2.185335 | -3 | -0.164302 | -3 | 0.19898 | 0.12788 | 0.35475 | 0.12788 | 19.66245 | 0.010559468 | 0 | Neutral |
| testneg_3658 | 0.14 | -0.37 | 7.061469 | -3 | 2.605779 | -3 | 0.32335 | 0.10604 | 0.22817 | 0.10634 | 17.21847 | 0.34 | 8 | Neutral |
| testneg_3659 | 0.07 | 0.38 | -2.148868 | -3 | 0.441575 | -3 | 0.16155 | 0.20638 | 0.09493 | 0.104 | 29.88846 | 0.060232684 | 0 | Neutral |

| | | | | | | | | | | | | | | |
|---|---|---|---|---|---|---|---|---|---|---|---|---|---|---|
| testneg_3660 | -0.23 | 1.04 | -1.164424 | -3 | 0.113253 | -3 | 0.12073 | 0.10969 | 0.38072 | 0.15993 | 18.33333 | 0.160282738 | 0 | Neutral |
| testneg_3661 | 0.14 | -1.01 | 0.94469 | -3 | -0.392377 | -3 | 0.08629 | 0.12788 | 0.24087 | 0.27985 | 10 | 0.08 | 0 | Neutral |
| testneg_3662 | 0 | 0 | 0.198253 | -3 | 1.821777 | -3 | 0.05535 | 0.12788 | 0.11187 | 0.12788 | 18.88889 | 0.04 | 0 | Neutral |
| testneg_3663 | 0 | 0 | 0.007192 | -3 | 2.020327 | -3 | 0.13011 | 0.14157 | 0.13177 | 0.12788 | 46.00189 | 0.02 | 0 | Neutral |
| testneg_3664 | 0.08 | 0.03 | -0.144427 | -3 | 0.565761 | -3 | 0.13575 | 0.12788 | 0.10383 | 0.08853 | 19.92008 | 0.02 | 0 | Neutral |
| testneg_3665 | -0.01 | -0.8 | 4.697901 | -3 | -1.743937 | -3 | 0.11791 | 0.39607 | 0.25643 | 0.12788 | 22.85054 | 0.161125 | 0 | Neutral |
| testneg_3666 | 0.05 | 0.29 | 1.745411 | -3 | 1.910339 | -3 | 0.16835 | 0.09282 | 0.10161 | 0.04472 | 20.90186 | 0.02 | 0 | Neutral |
| testneg_3667 | 0 | 0 | 0.332849 | 0.332849 | 0.726274 | -3 | 0.99595 | 0.23021 | 0.10228 | 0.22182 | 22.1048 | 0.17 | 0 | Neutral |
| testneg_3668 | 0.01 | 0.8 | 2.76764 | -3 | 4.382959 | -3 | 0.22795 | 0.12788 | 0.86513 | 0.31355 | 56.34955 | 0.360961839 | 7.8 | Neutral |
| testneg_3669 | 0 | 0 | 0.008676 | -3 | 0.980675 | -3 | 0.41987 | 0.67373 | 0.35137 | 0.09161 | 16.0119 | 0.03 | 0 | Neutral |
| testneg_3670 | -0.1 | -1.9 | 4.905835 | -3 | 1.247019 | -3 | 0.06708 | 0.09234 | 0.88456 | 0.11674 | 10 | 0.13 | 0 | Neutral |
| testneg_3671 | 0 | 0 | 5.995443 | -3 | -1.452684 | -3 | 0.19898 | 0.14645 | 0.24219 | 0.17458 | 15.39358 | 0.47 | 43 | Neutral |
| testneg_3672 | 0.03 | -1.03 | 2.514378 | -3 | -0.221228 | -3 | 0.3793 | 0.24105 | 0.11153 | 0.10767 | 11.25 | 0.041714466 | 0 | Neutral |
| testneg_3673 | 0.02 | 0.45 | 2.330575 | -3 | 3.817144 | -3 | 0.63233 | 0.16248 | 0.12321 | 0.08084 | 18.28557 | 0.150086207 | 0 | Neutral |
| testneg_3674 | -0.05 | -0.29 | 2.605779 | -3 | 1.350455 | -3 | 0.22817 | 0.10634 | 0.38695 | 0.12579 | 12.29167 | 0.01 | 0 | Neutral |
| testneg_3675 | 0.22 | -2.66 | 6.183054 | -3 | 3.527598 | -3 | 0.11791 | 0.39607 | 0.04615 | 0.12788 | 20.30197 | 0.459615385 | 37.8 | Neutral |
| testneg_3676 | -0.01 | -0.79 | 1.315391 | -3 | 4.247448 | 4.247448 | 0.99417 | 0.17365 | 0.44541 | 0.13979 | 12.70833 | 0.47689652 | 45 | Neutral |
| testneg_3677 | 0 | 0 | 1.827547 | -3 | 0.738714 | -3 | 0.35584 | 0.10874 | 0.17158 | 0.1149 | 13.51351 | 0 | 0 | Neutral |
| testneg_3678 | 0.17 | 0.38 | 5.35839 | -3 | 0.222999 | -3 | 0.93035 | 0.31722 | 0.1201 | 0.1062 | 6.06506 | 0.21 | 0.2 | Neutral |
| testneg_3679 | -0.03 | -0.1 | -0.677314 | -3 | -2.609079 | -3 | 0.3783 | 0.08224 | 0.19898 | 0.09533 | 27.3569 | 0.02 | 0 | Neutral |
| testneg_3680 | 0.04 | -0.23 | 2.504423 | -3 | 2.931463 | -3 | 0.14482 | 0.08592 | 0.27585 | 0.20571 | 19.66924 | 0.170086207 | 0 | Neutral |
| testneg_3681 | -0.02 | -0.41 | 0.579456 | 0.579456 | 4.58678 | -3 | 0.47678 | 0.38085 | 0.06112 | 0.06901 | 29.81459 | 0.183348666 | 0 | Neutral |
| testneg_3682 | 0.15 | 0.06 | 2.256555 | -3 | -0.77291 | -3 | 0.13468 | 0.09235 | 0.19898 | 0.12788 | 55.5475 | 0.04 | 0 | Neutral |
| testneg_3683 | 0 | 0 | -0.172145 | -0.346224 | 5.502802 | -3 | 0.14632 | 0.12662 | 0.05291 | 0.08633 | 8.97898 | 0.102083457 | 0 | Neutral |
| testneg_3684 | 0.18 | 0.16 | 4.65145 | -3 | -0.022585 | -3 | 0.19898 | 0.58778 | 0.2543 | 0.12929 | 10 | 0.143666667 | 0 | Neutral |
| testneg_3685 | 0.07 | 0.7 | -1.327597 | -3 | -2.237532 | -3 | 0.16155 | 0.20638 | 0.42836 | 0.12788 | 18.88889 | 0.061055556 | 0 | Neutral |
| testneg_3686 | -0.06 | -0.18 | 0.944356 | -3 | 5.187167 | -3 | 0.32532 | 0.09651 | 0.19898 | 0.12788 | 10.41667 | 0.088792797 | 0 | Neutral |
| testneg_3687 | 0 | 0 | 0.413917 | -3 | 5.840966 | -3 | 0.0832 | 0.09323 | 0.42235 | 0.12474 | 13.51351 | 0.07 | 0 | Neutral |
| testneg_3688 | 0.21 | 0.15 | 3.260141 | -3 | 0.249606 | -3 | 0.54792 | 0.54626 | 0.7062 | 0.83002 | 4.72222 | 0.325 | 3.4 | Neutral |
| testneg_3689 | 0 | 0 | 0.616227 | -3 | 3.022742 | -3 | 0.02971 | 0.33983 | 0.08975 | 0.07399 | 44.47407 | 0.14 | 0 | Neutral |
| testneg_3690 | -0.04 | 0.23 | 1.995788 | -3 | 6.090617 | -3 | 0.19043 | 0.07855 | 0.10185 | 0.12788 | 37.71421 | 0.150685446 | 0 | Neutral |
| testneg_3691 | 0.02 | 0.13 | 2.387064 | -3 | 1.209764 | -3 | 0.08186 | 0.11813 | 0.16636 | 0.12384 | 21.38889 | 0.010037037 | 0 | Neutral |
| testneg_3692 | -0.02 | 0.63 | 2.937875 | -3 | 1.064707 | -3 | 0.83098 | 0.57486 | 0.27637 | 0.0872 | 6.94444 | 0.11 | 0 | Neutral |
| testneg_3693 | 0.18 | -1.05 | 4.055237 | -3 | 3.483266 | -3 | 0.26961 | 0.09979 | 0.25856 | 0.17179 | 13.94349 | 0.54724194 | 69.6 | Disease_causing |
| testneg_3694 | -0.07 | -0.7 | -0.548079 | -3 | 2.399001 | -3 | 0.19898 | 0.12788 | 0.15562 | 0.10384 | 13.7487 | 0.020155844 | 0 | Neutral |
| testneg_3695 | 0.03 | -1.03 | 1.493352 | -3 | -2.290249 | -3 | 0.17364 | 0.18111 | 0.14862 | 0.12788 | 33.49673 | 0.040267857 | 0 | Neutral |
| testneg_3696 | 0 | 0 | 0.861365 | -3 | 1.730022 | -3 | 0.37953 | 0.11262 | 0.0472 | 0.12788 | 17.56711 | 0.000597148 | 0 | Neutral |
| testneg_3697 | 0.21 | 0.15 | 1.263794 | -0.859579 | 2.470276 | -3 | 0.19898 | 0.12895 | 0.06841 | 0.12788 | 24.47867 | 0.06 | 0 | Neutral |
| testneg_3698 | 0.2 | -1.26 | -1.220105 | -1.220105 | 4.076957 | -3 | 0.02971 | 0.33983 | 0.38324 | 0.25193 | 30.95448 | 0.58476704 | 86.6 | Disease_causing |
| testneg_3699 | 0 | 0 | 1.028069 | -3 | 2.76764 | -3 | 0.2771 | 0.14453 | 0.22795 | 0.12788 | 56.374 | 0.055453934 | 0 | Neutral |
| testneg_3700 | 0.05 | 0.29 | 3.039015 | -3 | 7.979277 | -3 | 0.204 | 0.15424 | 0.27004 | 0.17052 | 5.97222 | 0.795265256 | 100 | Disease_causing |
| testneg_3701 | 0.04 | -0.23 | 4.231687 | -3 | -0.28574 | -3 | 0.09616 | 0.11651 | 0.29455 | 0.12388 | 21.98605 | 0.03 | 0 | Neutral |
| testneg_3702 | 0 | 0 | 1.241234 | -3 | 3.470123 | -3 | 0.6258 | 0.16461 | 0.32532 | 0.09651 | 11.25 | 0.073858466 | 0 | Neutral |
| testneg_3703 | 0.14 | -0.37 | -0.136533 | -3 | 0.469669 | -0.133809 | 0.05932 | 0.09551 | 0.09556 | 0.10109 | 29.35429 | 0.0325 | 0 | Neutral |
| testneg_3704 | -0.03 | -0.12 | 3.470123 | -3 | 0.885869 | 0.885869 | 0.32532 | 0.09651 | 0.12003 | 0.12788 | 39.60168 | 0.04 | 0 | Neutral |
| testneg_3705 | -0.06 | -0.18 | 0.867502 | -3 | 0.829687 | -3 | 0.71511 | 0.34803 | 0.06841 | 0.12788 | 23.88889 | 0.045931373 | 0 | Neutral |
| testneg_3706 | 0 | 0 | 0.849647 | -3 | 0.56175 | -3 | 0.96651 | 0.28104 | 0.18246 | 0.12791 | 13.75 | 0.01 | 0 | Neutral |
| testneg_3707 | -0.01 | -0.79 | -3.816017 | -3 | 0.016238 | -3 | 0.06095 | 0.08179 | 0.58985 | 0.10504 | 13.80482 | 0.04 | 0 | Neutral |
| testneg_3708 | 0.04 | -0.23 | 4.461467 | -3 | 12.061237 | -3 | 0.20547 | 0.12178 | 0.19898 | 0.12788 | 16.88406 | 0.52 | 62.8 | Disease_causing |
| testneg_3709 | 0 | -0.01 | 0.125306 | -3 | -0.053373 | -3 | 0.54248 | 0.12333 | 0.06562 | 0.12788 | 18.61111 | 0.02 | 0 | Neutral |
| testneg_3710 | 0.03 | 1.22 | -0.130411 | -3 | 1.619805 | -3 | 0.96995 | 0.66487 | 0.19898 | 0.12788 | 17.28741 | 0.12 | 0 | Neutral |
| testneg_3711 | 0 | 0 | 1.434564 | -3 | 0.736335 | -3 | 0.29284 | 0.11262 | 0.98641 | 0.83983 | 4.44444 | 0.05 | 0 | Neutral |
| testneg_3712 | 0.18 | -1.05 | 7.271223 | 3.213612 | 1.795415 | -3 | 0.26323 | 0.2399 | 0.65717 | 0.16156 | 17.77778 | 0.817796296 | 100 | Disease_causing |
| testneg_3713 | -0.0022727 | -0.0332576 | 2.427467 | -3 | 1.614723 | 1.614723 | 0.43866 | 0.12121 | 0.74944 | 0.1263 | 11.25 | 0.03 | 0 | Neutral |
| testneg_3714 | -0.01 | -0.8 | 1.064707 | -3 | -0.279308 | -3 | 0.27637 | 0.0872 | 0.12885 | 0.61089 | 12.29167 | 0.070589248 | 0 | Neutral |
| testneg_3715 | 0.08 | 0.03 | -1.571869 | -3 | 0.312568 | -3 | 0.54489 | 0.09921 | 0.18264 | 0.10206 | 17.97975 | 0.04 | 0 | Neutral |
| testneg_3716 | 0.03 | -1.03 | 1.120342 | -3 | 0.857941 | -3 | 0.54015 | 0.13176 | 0.06939 | 0.08981 | 20 | 0.04 | 0 | Neutral |
| testneg_3717 | -0.07 | -0.33 | -0.49005 | -3 | 3.327226 | -3 | 0.27308 | 0.12788 | 0.22415 | 0.70109 | 15.5349 | 0.205331197 | 0.2 | Neutral |
| testneg_3718 | 0.06 | -0.57 | 3.970844 | -3 | -2.320937 | -3 | 0.13482 | 0.12788 | 0.12051 | 0.08503 | 13.51351 | 0.03 | 0 | Neutral |
| testneg_3719 | 0.01 | 0.8 | -0.725111 | -0.725111 | -0.408308 | -3 | 0.05116 | 0.08996 | 0.28165 | 0.15912 | 13.51351 | 0.112 | 0 | Neutral |
| testneg_3720 | 0.14 | -1.01 | 2.672539 | 2.672539 | 0.660523 | -3 | 0.46371 | 0.11637 | 0.06897 | 0.08445 | 24.65225 | 0.25 | 1.8 | Neutral |
| testneg_3721 | 0 | 0 | -0.076686 | -3 | 2.907554 | -3 | 0.84596 | 0.10633 | 0.09356 | 0.2448 | 11.25 | 0.127109556 | 0 | Neutral |
| testneg_3722 | -0.05 | -0.29 | -0.915676 | -3 | 2.671237 | -3 | 0.05767 | 0.07325 | 0.36852 | 0.0903 | 40.54161 | 0.06 | 0 | Neutral |
| testneg_3723 | 0.14 | -0.69 | 1.091597 | -0.390123 | 0.567562 | -3 | 0.09657 | 0.58605 | 0.17046 | 0.21736 | 6.03357 | 0.22 | 0 | Neutral |
| testneg_3724 | 0.15 | -0.28 | 1.144976 | -3 | -0.749376 | -0.749376 | 0.53592 | 0.11293 | 0.51437 | 0.17173 | 17.4528 | 0.07 | 0 | Neutral |
| testneg_3725 | 0.21 | 0.15 | 2.883248 | -3 | 4.078873 | -3 | 0.12957 | 0.12788 | 0.15364 | 0.08789 | 20.35088 | 0.134871212 | 0 | Neutral |
| testneg_3726 | -0.07 | -0.38 | 0.868703 | 0.868157 | 0.500506 | -3 | 0.06526 | 0.09018 | 0.68233 | 0.58998 | 10.83333 | 0.150405479 | 0 | Neutral |
| testneg_3727 | 0.14 | -1.01 | -0.384327 | -3 | -0.739513 | -3 | 0.18507 | 0.12788 | 0.39305 | 0.18672 | 10 | 0.06 | 0 | Neutral |
| testneg_3728 | 0 | 0 | -0.115728 | -3 | 3.103955 | -3 | 0.0922 | 0.07873 | 0.06667 | 0.08666 | 20.15479 | 0.05 | 0 | Neutral |
| testneg_3729 | 0.18 | 0.16 | 0.416167 | -3 | -0.750407 | -3 | 0.1224 | 0.11899 | 0.07631 | 0.12788 | 30.42735 | 0.02 | 0 | Neutral |
| testneg_3730 | -0.19 | 0.27 | -1.643119 | -3 | 4.571774 | -1.24052 | 0.1535 | 0.12382 | 0.22582 | 0.12788 | 39.62462 | 0.182 | 0 | Neutral |
| testneg_3731 | 0 | 0 | 2.634633 | -3 | -1.013068 | -3 | 0.40491 | 0.17596 | 0.40076 | 0.11341 | 4.72222 | 0.02174359 | 0 | Neutral |
| testneg_3732 | -0.14 | -0.54 | -0.251883 | -3 | 0.037077 | -3 | 0.39258 | 0.12788 | 0.09004 | 0.07829 | 20.82922 | 0.03 | 0 | Neutral |
| testneg_3733 | 0.04 | -0.23 | 4.017255 | -3 | -1.728523 | -3 | 0.19898 | 0.16603 | 0.25884 | 0.12788 | 11.74429 | 0.07 | 0 | Neutral |
| testneg_3734 | 0 | 0 | 0.122273 | -3 | -0.759298 | -3 | 0.0893 | 0.11207 | 0.06196 | 0.10881 | 13.02635 | 0.000217494 | 0 | Neutral |
| testneg_3735 | -0.02 | -0.45 | 2.696002 | -3 | 0.252532 | -3 | 0.3922 | 0.65186 | 0.06215 | 0.08575 | 10.83333 | 0.07 | 0 | Neutral |
| testneg_3736 | -0.02 | 0.36 | 2.643915 | -3 | 0.419787 | -3 | 0.29548 | 0.10732 | 0.32089 | 0.1035 | 10.88338 | 0.02 | 0 | Neutral |
| testneg_3737 | 0 | 0 | 1.579231 | -3 | -0.110263 | -3 | 0.72068 | 0.99841 | 0.38207 | 0.35052 | 5.74956 | 0.14 | 0 | Neutral |
| testneg_3738 | 0.03 | -1.03 | 4.741766 | -3 | 1.636161 | 1.636161 | 0.38719 | 0.15548 | 0.21153 | 0.11357 | 13.2432 | 0.11 | 0 | Neutral |
| testneg_3739 | 0.03 | 1.22 | 1.790281 | -3 | -0.351099 | -3 | 0.02059 | 0.066 | 0.19898 | 0.12788 | 26.37478 | 0.08 | 0 | Neutral |
| testneg_3740 | 0.05 | -0.81 | -0.021519 | -3 | 7.771345 | -3 | 0.30096 | 0.17203 | 0.32406 | 0.28376 | 16.39507 | 0.649228007 | 95.2 | Disease_causing |
| testneg_3741 | 0 | 0 | 1.309085 | -3 | 1.636782 | -3 | 0.64125 | 0.28792 | 0.15107 | 0.10457 | 13.13218 | 0.01 | 0 | Neutral |
| testneg_3742 | 0.07 | 0.7 | 0.131981 | -3 | 1.509728 | -3 | 0.69254 | 0.10021 | 0.0435 | 0.14527 | 21.94444 | 0.11 | 0 | Neutral |
| testneg_3743 | 0 | 0 | 1.221853 | 1.221853 | -0.518868 | -3 | 0.92659 | 0.13588 | 0.08386 | 0.12788 | 26.34995 | 0.05 | 0 | Neutral |
| testneg_3744 | 0.06 | -0.57 | 0.297516 | 0.297516 | 1.431446 | -3 | 0.19294 | 0.10464 | 0.23393 | 0.12567 | 13.94349 | 0.01 | 0 | Neutral |
| testneg_3745 | -0.03 | -0.12 | 1.183797 | -3 | -0.187467 | -0.187467 | 0.99765 | 0.95064 | 0.12314 | 0.11709 | 4.44444 | 0.110214318 | 0 | Neutral |
| testneg_3746 | 0 | 0 | 0.844548 | -3 | 0.836822 | -3 | 0.9237 | 0.55859 | 0.04866 | 0.05972 | 10 | 0.02 | 0 | Neutral |
| testneg_3747 | 0.03 | 0.12 | -0.432053 | -3 | 4.795372 | -3 | 0.10472 | 0.10014 | 0.02518 | 0.07165 | 21.41414 | 0.08 | 0 | Neutral |
| testneg_3748 | 0 | 0 | 0.71351 | -3 | -0.000915 | -3 | 0.29302 | 0.17394 | 0.10349 | 0.12788 | 24.0978 | 0.02035401 | 0 | Neutral |
| testneg_3749 | 0 | -0.01 | 0.306573 | -3 | 0.630495 | 0.630495 | 0.03898 | 0.09943 | 0.14419 | 0.1668 | 9.52684 | 0.1 | 0 | Neutral |
| testneg_3750 | 0.02 | -0.63 | 1.995171 | -3 | 1.30214 | -3 | 0.91377 | 0.5142 | 0.19898 | 0.12788 | 17.18434 | 0.05 | 0 | Neutral |
| testneg_3751 | 0.05 | 0.29 | 0.113571 | -3 | -3.291659 | -3 | 0.02844 | 0.12788 | 0.11957 | 0.08975 | 16.93015 | 0.04 | 0 | Neutral |
| testneg_3752 | 0 | 0 | 1.56019 | -3 | 5.471755 | -3 | 0.25487 | 0.13098 | 0.1784 | 0.12788 | 14.16667 | 0.098235663 | 0 | Neutral |
| testneg_3753 | -0.01 | -0.79 | 5.611956 | -3 | 0.369379 | -3 | 0.14187 | 0.1353 | 0.16388 | 0.14439 | 10.83333 | 0.316088672 | 6.6 | Neutral |
| testneg_3754 | 0.07 | 0.38 | 3.807502 | 3.807502 | 0.188594 | -3 | 0.34673 | 0.24308 | 0.35499 | 0.12052 | 41.54416 | 0.44 | 32.2 | Neutral |
| testneg_3755 | -0.05 | -0.29 | -0.038654 | -3 | 1.267807 | -3 | 0.18206 | 0.10727 | 0.18706 | 0.10259 | 22.64794 | 0.01 | 0 | Neutral |
| testneg_3756 | 0 | -0.01 | 0.834318 | -3 | 3.6541 | -3 | 0.31726 | 0.17249 | 0.19898 | 0.37148 | 6.98099 | 0.294110148 | 3.8 | Neutral |
| testneg_3757 | 0.04 | -0.23 | 2.208687 | -3 | 0.14995 | -3 | 0.19898 | 0.12788 | 0.1856 | 0.12788 | 18.8724 | 0 | 0 | Neutral |
| testneg_3758 | -0.08 | -0.03 | 2.585384 | -2.286968 | 1.049997 | -3 | 0.17039 | 0.12788 | 0.75415 | 0.29199 | 23.92206 | 0.090048416 | 0 | Neutral |
| testneg_3759 | 0.15 | -0.28 | 1.271267 | -3 | 2.71072 | -3 | 0.3221 | 0.3367 | 0.09719 | 0.09576 | 20.83333 | 0.12 | 0 | Neutral |
| testneg_3760 | -0.02 | -0.41 | -0.005856 | -3 | 0.659218 | -3 | 0.31124 | 0.08875 | 0.44429 | 0.20468 | 5 | 0.042025645 | 0 | Neutral |
| testneg_3761 | -0.01 | -0.79 | 0.771742 | -3 | 2.370092 | -3 | 0.18633 | 0.14783 | 0.59066 | 0.10874 | 10.09538 | 0.05 | 0 | Neutral |
| testneg_3762 | -0.03 | -0.1 | 2.079972 | 2.079972 | 4.639274 | -3 | 0.25549 | 0.12378 | 0.85508 | 0.12634 | 22.34127 | 0.196744514 | 0 | Neutral |
| testneg_3763 | 0 | 0 | 0.338533 | -3 | 5.176715 | -3 | 0.09618 | 0.10294 | 0.206 | 0.14631 | 19.58583 | 0.208467248 | 0 | Neutral |
| testneg_3764 | 0.22 | -2.66 | 3.422072 | -3 | 5.180313 | -3 | 0.20431 | 0.17304 | 0.19898 | 0.12788 | 22.84781 | 0.360767588 | 13.8 | Neutral |
| testneg_3765 | 0.14 | -1.01 | 7.382501 | -3 | 3.349863 | -3 | 0.60703 | 0.12378 | 0.1472 | 0.12788 | 18.88889 | 0.495416667 | 51.8 | Disease_causing |
| testneg_3766 | 0.09 | -2.17 | 5.760643 | -3 | 0.050268 | -3 | 0.85252 | 0.83576 | 0.51934 | 0.12046 | 11.25 | 0.37 | 9.4 | Neutral |
| testneg_3767 | 0.12 | -1.14 | -0.006272 | -0.006272 | 3.299396 | -3 | 0.07335 | 0.12788 | 0.12547 | 0.09267 | 13.51351 | 0.076666667 | 0 | Neutral |
| testneg_3768 | 0.21 | 0.15 | 0.153476 | -3 | 2.733966 | -3 | 0.05978 | 0.10176 | 0.34205 | 0.11195 | 37.72358 | 0.07 | 0 | Neutral |
| testneg_3769 | 0.07 | 0.7 | 5.311585 | -3 | -0.108631 | -3 | 0.24026 | 0.12788 | 0.11055 | 0.19829 | 11.74429 | 0.333541667 | 5.4 | Neutral |
| testneg_3770 | 0.03 | -1.03 | 5.917112 | -3 | 0.866411 | -3 | 0.04863 | 0.12149 | 0.13011 | 0.14157 | 16.84579 | 0.295 | 2 | Neutral |
| testneg_3771 | -0.15 | -0.04 | 3.653806 | -3 | 6.365761 | -3 | 0.10371 | 0.08655 | 0.21348 | 0.10616 | 18.6768 | 0.3 | 1.8 | Neutral |
| testneg_3772 | 0.21 | 0.15 | 4.637191 | -3 | -0.036185 | -3 | 0.9355 | 0.40418 | 0.54792 | 0.54626 | 5.97222 | 0.364847661 | 10.4 | Neutral |
| testneg_3773 | -0.08 | -0.03 | 3.376108 | -3 | -1.08749 | -3 | 0.16474 | 0.41578 | 0.08265 | 0.12788 | 17.01711 | 0.100037037 | 0 | Neutral |
| testneg_3774 | -0.02 | 0.36 | 4.584749 | -3 | -0.187363 | -3 | 0.86306 | 0.10962 | 0.90649 | 0.3543 | 5 | 0.201527778 | 0 | Neutral |
| testneg_3775 | -0.08 | -0.03 | 1.580417 | -3 | 5.916123 | -3 | 0.29417 | 0.10965 | 0.19898 | 0.11773 | 21.84591 | 0.112083333 | 0 | Neutral |
| testneg_3776 | 0.04 | -0.23 | 1.5079 | -3 | 0.827167 | -3 | 0.34694 | 0.20397 | 0.07211 | 0.12934 | 16.42857 | 0.020714286 | 0 | Neutral |
| testneg_3777 | -0.05 | 0.81 | 2.250533 | -3 | 0.634039 | -3 | 0.14021 | 0.43531 | 0.08337 | 0.18096 | 27.64576 | 0.240235769 | 0.2 | Neutral |
| testneg_3778 | 0.15 | -0.28 | 1.374995 | -3 | 1.995788 | -3 | 0.26727 | 0.12788 | 0.19043 | 0.07855 | 20.72728 | 0.01 | 0 | Neutral |
| testneg_3779 | 0 | 0 | 0.082683 | -3 | 4.056275 | -3 | 0.19898 | 0.12788 | 0.1324 | 0.09503 | 27.812 | 0.050608883 | 0 | Neutral |
| testneg_3780 | -0.08 | -0.03 | -0.949463 | -3 | 2.691039 | -3 | 0.09018 | 0.16175 | 0.15056 | 0.40109 | 18.55633 | 0.298350392 | 2.6 | Neutral |
| testneg_3781 | 0.05 | -0.81 | 0.331709 | 0.331709 | 3.968663 | -3 | 0.16534 | 0.26726 | 0.16621 | 0.49845 | 10.74877 | 0.573233417 | 79.4 | Disease_causing |

| | | | | | | | | | | | | | | |
|---|---|---|---|---|---|---|---|---|---|---|---|---|---|---|
| testneg_3782 | -0.01 | 0.57 | 0.995275 | -3 | 1.890706 | -3 | 0.05886 | 0.38242 | 0.48688 | 0.13071 | 9.21981 | 0.161602564 | 0 | Neutral |
| testneg_3783 | -0.06 | -0.18 | 0.622743 | -3 | 1.653519 | -3 | 0.18979 | 0.07948 | 0.11042 | 0.09881 | 21.53664 | 0.010526316 | 0 | Neutral |
| testneg_3784 | 0.05 | 0.29 | 1.89239 | -3 | 0.118367 | -3 | 0.11791 | 0.39607 | 0.1472 | 0.12788 | 14.18481 | 0.057008929 | 0 | Neutral |
| testneg_3785 | 0 | 0 | 0.346053 | -3 | 0.052669 | -3 | 0.3047 | 0.11929 | 0.10781 | 0.09901 | 29.9024 | 0.01 | 0 | Neutral |
| testneg_3786 | 0.09 | -0.3 | 1.436098 | -3 | 0.318608 | -3 | 0.45628 | 0.15819 | 0.11373 | 0.10739 | 14.16667 | 0.03 | 0 | Neutral |
| testneg_3787 | -0.19 | -0.51 | 0.918079 | 0.918079 | 1.117264 | -3 | 0.83039 | 0.1727 | 0.23968 | 0.12788 | 10.41667 | 0.09 | 0 | Neutral |
| testneg_3788 | -0.03 | -0.12 | -2.124954 | -3 | 0.967941 | -3 | 0.34153 | 0.08855 | 0.08365 | 0.34122 | 11.25 | 0.08 | 0 | Neutral |
| testneg_3789 | -0.04 | -2.01 | 6.015793 | -3 | 3.595391 | -3 | 0.10236 | 0.09622 | 0.09288 | 0.16617 | 23.70087 | 0.595424706 | 89.2 | Disease_causing |
| testneg_3790 | 0 | 0 | -0.044681 | -3 | 1.305769 | -3 | 0.31584 | 0.11996 | 0.16468 | 0.11275 | 5.4902 | 0.000644002 | 0 | Neutral |
| testneg_3791 | 0 | 0 | 5.987437 | -3 | -0.667383 | -3 | 0.05535 | 0.12788 | 0.19898 | 0.08275 | 29.66438 | 0.16 | 0.2 | Neutral |
| testneg_3792 | 0.04 | -0.23 | 2.379002 | -3 | 5.076843 | 5.076843 | 0.25236 | 0.11809 | 0.45281 | 0.21621 | 7.07967 | 0.564660241 | 79.4 | Disease_causing |
| testneg_3793 | -0.02 | -0.45 | -0.263596 | -3 | 1.682966 | -3 | 0.19898 | 0.12788 | 0.08814 | 0.42357 | 19.18549 | 0.102206478 | 0 | Neutral |
| testneg_3794 | 0.07 | 0.38 | 2.307955 | -3 | 2.310369 | -3 | 0.22872 | 0.15588 | 0.54895 | 0.15082 | 10 | 0.158645513 | 0 | Neutral |
| testneg_3795 | -0.19 | 0.27 | -1.643119 | -3 | -0.078944 | -3 | 0.1535 | 0.12382 | 0.13255 | 0.0962 | 33.61111 | 0.060169102 | 0 | Neutral |
| testneg_3796 | 0.04 | -0.23 | 0.690974 | -3 | 4.263298 | -3 | 0.37065 | 0.12788 | 0.18328 | 0.12788 | 19.87208 | 0.06 | 0 | Neutral |
| testneg_3797 | 0 | -0.01 | 1.790205 | -3 | 0.590072 | -3 | 0.11796 | 0.12788 | 0.0819 | 0.1823 | 22.82634 | 0.08 | 0 | Neutral |
| testneg_3798 | 0.21 | 0.15 | 2.276796 | -3 | 0.081615 | 0.081615 | 0.06939 | 0.08981 | 0.14491 | 0.10508 | 31.25 | 0.04 | 0 | Neutral |
| testneg_3799 | 0 | 0 | -0.471382 | -0.471382 | 0.403196 | -3 | 0.21933 | 0.18619 | 0.29965 | 0.26486 | 10.41667 | 0.065588235 | 0 | Neutral |
| testneg_3800 | 0 | 0 | 1.605285 | -3 | 2.177568 | -3 | 0.5068 | 0.12276 | 0.08876 | 0.09037 | 10.83333 | 0.012222222 | 0 | Neutral |
| testneg_3801 | 0 | 0 | 0.094365 | -3 | 0.707491 | -3 | 0.55179 | 0.29199 | 0.43762 | 0.12788 | 11.25 | 0.010194193 | 0 | Neutral |
| testneg_3802 | -0.06 | -0.18 | 3.046495 | -3 | 0.328138 | -3 | 0.76924 | 0.15588 | 0.50592 | 0.0952 | 21.30134 | 0.06 | 0 | Neutral |
| testneg_3803 | 0.09 | -0.25 | 2.60676 | -3 | 7.197642 | -3 | 0.09439 | 0.07415 | 0.15708 | 0.12788 | 17.21847 | 0.291428571 | 2.4 | Neutral |
| testneg_3804 | -0.03 | -0.1 | 2.765088 | -3 | 2.016143 | -3 | 0.85092 | 0.58871 | 0.18373 | 0.12788 | 18.57759 | 0.12 | 0 | Neutral |
| testneg_3805 | -0.03 | 1.03 | 2.97112 | -3 | 0.84675 | -3 | 0.5901 | 0.19444 | 0.4358 | 0.1222 | 19.32823 | 0.103416667 | 0 | Neutral |
| testneg_3806 | -0.21 | -0.15 | -0.83994 | -3 | 2.402629 | -3 | 0.95319 | 0.20297 | 0.19898 | 0.12788 | 21.48002 | 0.130060241 | 0 | Neutral |
| testneg_3807 | 0 | 0 | 6.373062 | -3 | 0.576122 | -3 | 0.36793 | 0.15588 | 0.11827 | 0.23034 | 10.41667 | 0.506561765 | 57.8 | Disease_causing |
| testneg_3808 | 0.09 | -0.3 | 4.733195 | -3 | 4.161435 | -3 | 0.161 | 0.11597 | 0.74271 | 0.10951 | 19.51932 | 0.182450192 | 0 | Neutral |
| testneg_3809 | 0.2 | -1.26 | 1.061885 | -3 | 1.65704 | -3 | 0.17725 | 0.12788 | 0.13523 | 0.12788 | 23.68839 | 0.030112554 | 0 | Neutral |
| testneg_3810 | 0 | 0 | -1.231395 | -3 | 0.431179 | -3 | 0.46757 | 0.14122 | 0.47751 | 0.2062 | 20.27778 | 0.07 | 0 | Neutral |
| testneg_3811 | 0 | 0 | 0.205888 | -3 | 3.680141 | -3 | 0.08922 | 0.10539 | 0.01642 | 0.12788 | 7.10199 | 0.041671569 | 0 | Neutral |
| testneg_3812 | 0 | -0.01 | 2.002215 | -3 | 6.066376 | -3 | 0.95682 | 0.22744 | 0.15341 | 0.12272 | 9.82124 | 0.165229115 | 0 | Neutral |
| testneg_3813 | 0.14 | -0.37 | 1.599249 | -3 | -0.235499 | -3 | 0.08714 | 0.08945 | 0.1203 | 0.13872 | 18.33333 | 0.06 | 0 | Neutral |
| testneg_3814 | 0.06 | -0.57 | 1.985669 | -3 | 3.640704 | -3 | 0.27308 | 0.12788 | 0.4069 | 0.3841 | 12.76316 | 0.310155992 | 2.2 | Neutral |
| testneg_3815 | 0 | 0 | -0.030607 | -3 | 2.698795 | -3 | 0.17146 | 0.19882 | 0.17806 | 0.12788 | 19.37853 | 0.054880952 | 0 | Neutral |
| testneg_3816 | 0.04 | -0.89 | -0.197119 | -3 | 2.171112 | -3 | 0.28191 | 0.20693 | 0.86513 | 0.31355 | 11.875 | 0.191883478 | 0 | Neutral |
| testneg_3817 | 0.00109091 | -0.0100909 | 1.971045 | -3 | 1.001127 | -3 | 0.14457 | 0.11614 | 0.14 | 0.12151 | 20.11599 | 0.000117712 | 0 | Neutral |
| testneg_3818 | -0.07 | -0.38 | -1.415783 | -3 | 1.51257 | -3 | 0.13278 | 0.20513 | 0.28066 | 0.18241 | 19.11104 | 0.164126192 | 0 | Neutral |
| testneg_3819 | -0.18 | -0.16 | 0.369791 | -3 | 1.999648 | 1.999648 | 0.66227 | 0.11002 | 0.12626 | 0.10279 | 23.64499 | 0.09 | 0 | Neutral |
| testneg_3820 | 0 | 0 | 0.329118 | -3 | 4.623974 | -3 | 0.51707 | 0.45685 | 0.09866 | 0.104 | 21.2926 | 0.149632937 | 0 | Neutral |
| testneg_3821 | -0.04 | 0.23 | 0.98325 | -3 | -0.194401 | -0.194401 | 0.16054 | 0.11683 | 0.06749 | 0.1776 | 5.70654 | 0.07736746 | 0 | Neutral |
| testneg_3822 | 0 | 0 | 1.38626 | -3 | 6.502111 | -3 | 0.50248 | 0.10939 | 0.70946 | 0.11809 | 10.83333 | 0.121651357 | 0 | Neutral |
| testneg_3823 | -0.04 | 0.89 | 1.989667 | -3 | 3.901083 | -3 | 0.06517 | 0.15107 | 0.52118 | 0.36073 | 18.33333 | 0.560711796 | 79 | Disease_causing |
| testneg_3824 | -0.03 | -0.1 | 0.545839 | -3 | 1.555382 | -3 | 0.58545 | 0.14414 | 0.80134 | 0.08685 | 18.47769 | 0.020126633 | 0 | Neutral |
| testneg_3825 | 0.07 | 0.38 | -0.283022 | -3 | 1.30641 | -3 | 0.32111 | 0.12788 | 0.19898 | 0.12788 | 14.79675 | 0.005 | 0 | Neutral |
| testneg_3826 | 0.02 | -0.63 | 5.700045 | -3 | -1.300607 | -3 | 0.12682 | 0.11357 | 0.37424 | 0.12788 | 17.25184 | 0.11 | 0 | Neutral |
| testneg_3827 | 0 | 0 | 2.434662 | -3 | 1.217695 | -3 | 0.19898 | 0.12788 | 0.27629 | 0.11166 | 10.41667 | 0.00013516 | 0 | Neutral |
| testneg_3828 | -0.03 | 0.85 | -0.534952 | -3 | 5.969277 | -3 | 0.12203 | 0.13515 | 0.31067 | 0.35414 | 10.41667 | 0.440595238 | 32.4 | Neutral |
| testneg_3829 | 0 | -0.01 | 1.221125 | -3 | 0.200393 | 0.200393 | 0.36793 | 0.15588 | 0.19898 | 0.08714 | 18.33333 | 0.01780303 | 0 | Neutral |
| testneg_3830 | 0 | 0 | 0.353546 | -3 | 5.8834 | -3 | 0.28191 | 0.20693 | 0.20112 | 0.17491 | 10.41667 | 0.452857019 | 41 | Neutral |
| testneg_3831 | 0.14 | -1.01 | 4.833027 | -3 | -1.390553 | -3 | 0.1004 | 0.11409 | 0.08789 | 0.6361 | 14.33231 | 0.250640186 | 0.4 | Neutral |
| testneg_3832 | 0.09 | -0.3 | -1.322992 | -3 | -0.60097 | -3 | 0.04863 | 0.12149 | 0.18153 | 0.10845 | 12.29167 | 0.04 | 0 | Neutral |
| testneg_3833 | 0.21 | 0.15 | 3.65813 | -3 | 1.886693 | -3 | 0.1205 | 0.23508 | 0.8433 | 0.13128 | 13.19613 | 0.209720635 | 0 | Neutral |
| testneg_3834 | 0.03 | 0.12 | 2.511997 | -3 | 5.39414 | -3 | 0.99765 | 0.95064 | 0.50034 | 0.07527 | 9.23329 | 0.243333333 | 0 | Neutral |
| testneg_3835 | 0 | 0 | 1.708897 | -3 | -0.614942 | -3 | 0.5202 | 0.16306 | 0.4129 | 0.11172 | 28.47222 | 0.020571895 | 0 | Neutral |
| testneg_3836 | 0 | 0 | 0.275842 | -3 | 2.708722 | -3 | 0.14462 | 0.11239 | 0.17412 | 0.10595 | 6.9869 | 0.010458007 | 0 | Neutral |
| testneg_3837 | 0.03 | 0.12 | 2.981794 | -3 | 0.284017 | -3 | 0.17394 | 0.15823 | 0.3025 | 0.2499 | 10.83333 | 0.21648539 | 0.2 | Neutral |
| testneg_3838 | 0.08 | 0.03 | 0.136092 | -3 | -0.384446 | -3 | 0.16507 | 0.10992 | 0.36829 | 0.11457 | 12.70833 | 0.0100761 | 0 | Neutral |
| testneg_3839 | -0.07 | -0.38 | -0.136973 | -3 | 3.006809 | -3 | 0.21843 | 0.85214 | 0.84365 | 0.12788 | 4.72222 | 0.130120482 | 0 | Neutral |
| testneg_3840 | 0.04 | 2.01 | 1.269375 | -3 | 1.627261 | -3 | 0.54489 | 0.09921 | 0.09439 | 0.07415 | 17.97975 | 0.08 | 0 | Neutral |
| testneg_3841 | 0.08 | 0.03 | 4.419543 | -3 | 4.426423 | -3 | 0.45029 | 0.12788 | 0.08041 | 0.21802 | 18.33333 | 0.570887058 | 76.8 | Disease_causing |
| testneg_3842 | 0.06 | -0.57 | 1.240379 | -3 | 5.036141 | -3 | 0.64591 | 0.38907 | 0.04787 | 0.12788 | 9.89595 | 0.183552579 | 0 | Neutral |
| testneg_3843 | -0.11 | 1.95 | 0.156645 | -3 | 5.882463 | -3 | 0.11802 | 0.09304 | 0.06099 | 0.12788 | 13.80454 | 0.25 | 0.2 | Neutral |
| testneg_3844 | 0.09 | -0.25 | 1.672239 | -3 | -0.942639 | -3 | 0.25989 | 0.11651 | 0.05996 | 0.12788 | 68.71674 | 0.08 | 0 | Neutral |
| testneg_3845 | 0.14 | -0.69 | 1.111818 | -3 | 6.282616 | -3 | 0.08775 | 0.12788 | 0.38375 | 0.09256 | 13.46801 | 0.15 | 0 | Neutral |
| testneg_3846 | 0 | 0 | 0.285584 | 0.285584 | 16.024484 | -3 | 0.20527 | 0.10053 | 0.16393 | 0.08897 | 25.14162 | 0.33205 | 3.6 | Neutral |
| testneg_3847 | 0 | 0 | 1.431242 | -3 | -0.14171 | -3 | 0.83074 | 0.14769 | 0.10782 | 0.14795 | 20 | 0.080050505 | 0 | Neutral |
| testneg_3848 | 0 | 0 | 1.390544 | -3 | -1.316616 | -1.316616 | 0.30257 | 0.17298 | 0.07347 | 0.08817 | 23.6184 | 0.02 | 0 | Neutral |
| testneg_3849 | -0.18 | -0.16 | 2.830718 | -3 | 6.431621 | 6.431621 | 0.11229 | 0.08897 | 0.26676 | 0.76568 | 29.21456 | 0.599307018 | 88 | Disease_causing |
| testneg_3850 | -0.01 | -0.79 | 0.117628 | -3 | 5.774454 | -3 | 0.11134 | 0.13818 | 0.10921 | 0.34026 | 28.85698 | 0.470841919 | 44 | Neutral |
| testneg_3851 | -0.08 | -0.03 | 1.395184 | -3 | 4.563044 | -3 | 0.11335 | 0.14225 | 0.01952 | 0.08329 | 18.33333 | 0.14 | 0 | Neutral |
| testneg_3852 | -0.09 | 0.3 | -0.908567 | -3 | 7.13751 | -3 | 0.39774 | 0.10813 | 0.2852 | 0.12788 | 16.43865 | 0.29 | 3.4 | Neutral |
| testneg_3853 | 0 | 0 | -0.005535 | -3 | 2.8284 | 0.552815 | 0.98312 | 0.12788 | 0.20417 | 0.10681 | 18.13517 | 0.040113085 | 0 | Neutral |
| testneg_3854 | 0.09 | -0.3 | -0.129706 | -3 | 4.046865 | -3 | 0.53996 | 0.14577 | 0.42666 | 0.12788 | 10.62943 | 0.151547619 | 0 | Neutral |
| testneg_3855 | 0.02 | 0.45 | 1.88229 | -3 | 2.495623 | -3 | 0.64922 | 0.10176 | 0.19898 | 0.10106 | 13.51351 | 0.04 | 0 | Neutral |
| testneg_3856 | 0.14 | -1.01 | 0.820712 | -3 | -0.155218 | -3 | 0.42755 | 0.09484 | 0.06939 | 0.08981 | 20 | 0.02 | 0 | Neutral |
| testneg_3857 | 0 | 0 | 0.796034 | -3 | 4.142425 | -3 | 0.99915 | 0.96333 | 0.25447 | 0.12788 | 16.85532 | 0.18 | 0 | Neutral |
| testneg_3858 | 0.05 | 0.29 | -1.828653 | -3 | 0.990803 | -3 | 0.4097 | 0.36733 | 0.10822 | 0.10942 | 15.21341 | 0.05016902 | 0 | Neutral |
| testneg_3859 | 0 | -0.32 | -2.498179 | -3 | 3.286987 | -3 | 0.50682 | 0.12788 | 0.19898 | 0.12788 | 13.02932 | 0.060652254 | 0 | Neutral |
| testneg_3860 | 0.07 | -0.31 | 0.202903 | -3 | 4.06036 | -3 | 0.06711 | 0.09765 | 0.06187 | 0.36607 | 12.57218 | 0.300125 | 2 | Neutral |
| testneg_3861 | 0 | 0 | 0.096068 | -3 | 1.417365 | 1.417365 | 0.30329 | 0.14582 | 0.61184 | 0.55597 | 18.33333 | 0.090445736 | 0 | Neutral |
| testneg_3862 | 0 | 0 | 0.488023 | -3 | 0.072482 | -3 | 0.85508 | 0.12634 | 0.07394 | 0.09408 | 15.37063 | 0.01 | 0 | Neutral |
| testneg_3863 | -0.15 | -0.04 | 1.117264 | -3 | 0.583408 | 0.583408 | 0.23968 | 0.12788 | 0.05386 | 0.09802 | 20 | 0.040236111 | 0 | Neutral |
| testneg_3864 | -0.01 | 0.57 | -2.581564 | -3 | 1.600017 | -3 | 0.19898 | 0.12788 | 0.07994 | 0.1101 | 24.0281 | 0.03 | 0 | Neutral |
| testneg_3865 | -0.07 | -0.38 | 0.72503 | -3 | -0.061002 | -0.061002 | 0.56862 | 0.12788 | 0.35561 | 0.08845 | 49.88273 | 0.020037037 | 0 | Neutral |
| testneg_3866 | -0.18 | -0.16 | 1.547948 | -3 | 5.728641 | -1.024719 | 0.9924 | 0.37896 | 0.27806 | 0.09713 | 9.67871 | 0.24 | 0.4 | Neutral |
| testneg_3867 | 0 | 0 | -0.031197 | -3 | 3.446417 | 3.446417 | 0.1017 | 0.11809 | 0.29013 | 0.11368 | 34.76461 | 0.17 | 0 | Neutral |
| testneg_3868 | 0 | 0 | -0.342648 | -3 | -1.457604 | -3 | 0.71463 | 0.29732 | 0.11746 | 0.1471 | 21.38889 | 0.125236051 | 0 | Neutral |
| testneg_3869 | 0.04 | -0.23 | 3.591512 | -3 | -0.423572 | -3 | 0.99765 | 0.95064 | 0.54489 | 0.09921 | 6.55362 | 0.16 | 0 | Neutral |
| testneg_3870 | 0.03 | -1.03 | -0.150684 | -3 | 0.200864 | -3 | 0.18866 | 0.12788 | 0.36257 | 0.26115 | 26.10625 | 0.08 | 0 | Neutral |
| testneg_3871 | 0.01 | -1.44 | 2.788395 | -3 | 2.472827 | -3 | 0.23567 | 0.12788 | 0.50134 | 0.15488 | 51.7632 | 0.219867284 | 0.2 | Neutral |
| testneg_3872 | 0.01 | 0.79 | 1.378098 | -3 | 6.199426 | -3 | 0.11501 | 0.13748 | 0.18866 | 0.12788 | 38.18837 | 0.19035119 | 0 | Neutral |
| testneg_3873 | 0.14 | -1.85 | 5.628007 | -3 | 2.440516 | -3 | 0.52836 | 0.12785 | 0.92776 | 0.09677 | 13.28792 | 0.239272727 | 0.2 | Neutral |
| testneg_3874 | 0.07 | 0.01 | -0.676534 | -0.676534 | -0.473257 | -3 | 0.16262 | 0.15786 | 0.46142 | 0.11057 | 19.10645 | 0.021511299 | 0 | Neutral |
| testneg_3875 | 0.08 | 0.03 | -0.196592 | -0.196592 | 5.931499 | -3 | 0.32733 | 0.16869 | 0.11019 | 0.08132 | 19.44444 | 0.220497543 | 0 | Neutral |
| testneg_3876 | 0.02 | -0.28 | 0.69881 | -3 | 4.507329 | -3 | 0.20692 | 0.70649 | 0.47789 | 0.10803 | 4.44444 | 0.171590742 | 0 | Neutral |
| testneg_3877 | 0 | -0.01 | 7.854373 | -3 | 0.390383 | -3 | 0.4113 | 0.2303 | 0.08213 | 0.08913 | 31.25 | 0.4 | 16.6 | Neutral |
| testneg_3878 | 0.09 | -0.3 | 1.790743 | -3 | 5.393665 | 5.393665 | 0.7823 | 0.5949 | 0.03809 | 0.16044 | 10.41667 | 0.582388889 | 86.6 | Disease_causing |
| testneg_3879 | 0.15 | -0.82 | 2.442187 | -3 | 5.704027 | -3 | 0.07875 | 0.11082 | 0.09504 | 0.38939 | 22.63149 | 0.475571429 | 44 | Neutral |
| testneg_3880 | 0.17 | 0.38 | 3.132136 | -3 | 0.347716 | -3 | 0.0105 | 0.12788 | 0.0634 | 0.0824 | 13.26939 | 0.07 | 0 | Neutral |
| testneg_3881 | 0.06 | -0.57 | 2.475201 | -3 | 3.850429 | -3 | 0.11264 | 0.11163 | 0.23939 | 0.25284 | 11.64938 | 0.338745098 | 7.2 | Neutral |
| testneg_3882 | 0 | 0 | 0.900502 | -0.566651 | 1.225058 | -3 | 0.38016 | 0.14503 | 0.30209 | 0.14229 | 19.34591 | 0.05 | 0 | Neutral |
| testneg_3883 | 0 | 0 | 1.898828 | -3 | 0.322383 | -3 | 0.38429 | 0.11956 | 0.50034 | 0.07527 | 29.274 | 0.000555556 | 0 | Neutral |
| testneg_3884 | 0.07 | 0.38 | -1.867166 | -3 | 1.224579 | -3 | 0.92235 | 0.62242 | 0.19898 | 0.12788 | 10.8523 | 0.07 | 0 | Neutral |
| testneg_3885 | 0 | 0.32 | 2.336378 | -3 | 1.193729 | -3 | 0.75978 | 0.11423 | 0.85345 | 0.13067 | 20.042 | 0.03 | 0 | Neutral |
| testneg_3886 | 0.15 | -0.28 | 1.45686 | -3 | 1.1308 | 1.1308 | 0.38552 | 0.14172 | 0.19898 | 0.11185 | 35.49268 | 0.021666667 | 0 | Neutral |
| testneg_3887 | -0.18 | -0.16 | 3.027444 | -3 | 5.108849 | -3 | 0.08475 | 0.19889 | 0.49064 | 0.12294 | 21.53664 | 0.423198413 | 25 | Neutral |
| testneg_3888 | 0.04 | -0.23 | 2.281524 | -3 | 0.922605 | -3 | 0.10893 | 0.12072 | 0.2152 | 0.12788 | 28.47222 | 0.01 | 0 | Neutral |
| testneg_3889 | 0.21 | 0.15 | 2.141778 | -3 | 0.582961 | -3 | 0.19898 | 0.21177 | 0.32335 | 0.10604 | 19.8668 | 0.060513158 | 0 | Neutral |
| testneg_3890 | -0.07 | -0.38 | 2.948419 | -3 | 2.51331 | -3 | 0.04863 | 0.12149 | 0.14601 | 0.12788 | 10.83333 | 0.11 | 0 | Neutral |
| testneg_3891 | 0 | 0 | 1.764306 | -3 | 1.072249 | -3 | 0.14458 | 0.14062 | 0.0895 | 0.12788 | 23.6906 | 0.010769231 | 0 | Neutral |
| testneg_3892 | 0 | 0 | 0.796034 | -3 | 6.6361 | -3 | 0.99915 | 0.96333 | 0.26618 | 0.2057 | 4.72222 | 0.391961975 | 21.8 | Neutral |
| testneg_3893 | 0 | 0 | -0.39694 | -3 | 0.333581 | -3 | 0.21807 | 0.12788 | 0.78515 | 0.35093 | 18.88889 | 0.035249475 | 0 | Neutral |
| testneg_3894 | 0.00415385 | -0.0140769 | 2.117525 | -3 | -1.397029 | -3 | 0.37463 | 0.23021 | 0.52932 | 0.65749 | 10.41667 | 0.110889328 | 0 | Neutral |
| testneg_3895 | -0.08 | -0.03 | -0.761635 | -3 | 2.377826 | -3 | 0.04866 | 0.05972 | 0.96377 | 0.1167 | 10 | 0.082636905 | 0 | Neutral |
| testneg_3896 | -0.005 | -0.015 | 1.866374 | -3 | 0.890109 | -3 | 0.22034 | 0.12788 | 0.52771 | 0.10974 | 12.27213 | 0.000871212 | 0 | Neutral |
| testneg_3897 | 0 | 0 | 0.864288 | -3 | 6.203124 | -3 | 0.21014 | 0.12378 | 0.10581 | 0.11631 | 18.88889 | 0.09121127 | 0 | Neutral |
| testneg_3898 | -0.08 | -0.03 | 1.155408 | -3 | 0.354179 | 0.354179 | 0.04161 | 0.12788 | 0.1462 | 0.18896 | 18.71322 | 0.14 | 0 | Neutral |
| testneg_3899 | -0.02 | 0.68 | 0.349669 | -3 | 1.192946 | -3 | 0.19898 | 0.12788 | 0.13061 | 0.21023 | 16.06843 | 0.061479752 | 0 | Neutral |
| testneg_3900 | 0 | 0 | 0.435323 | -3 | 2.081303 | -3 | 0.20527 | 0.10053 | 0.1336 | 0.15669 | 27.4024 | 0.072212632 | 0 | Neutral |
| testneg_3901 | 0 | 0 | 1.669413 | -3 | 0.348638 | -3 | 0.7597 | 0.12788 | 0.19898 | 0.12788 | 14.12935 | 0 | 0 | Neutral |
| testneg_3902 | -0.02 | 0.36 | 5.54728 | -3 | 7.159958 | 4.517994 | 0.14862 | 0.12788 | 0.10803 | 0.14652 | 22.20588 | 0.8 | 100 | Disease_causing |
| testneg_3903 | 0 | 0 | 0.212473 | -3 | -0.754995 | -3 | 0.61098 | 0.11179 | 0.85482 | 0.1072 | 31.06175 | 0.02 | 0 | Neutral |

| | | | | | | | | | | | | | | |
|---|---|---|---|---|---|---|---|---|---|---|---|---|---|---|
| testneg_3904 | 0 | 0 | 1.414183 | -3 | 1.268416 | -3 | 0.55618 | 0.11352 | 0.43977 | 0.17284 | 17.13595 | 0.03018976 | 0 | Neutral |
| testneg_3905 | 0 | 0 | 0.256955 | -3 | 2.840836 | -3 | 0.1846 | 0.08682 | 0.0669 | 0.10997 | 32.10521 | 0.03 | 0 | Neutral |
| testneg_3906 | 0 | 0 | 0.383044 | -3 | 0.312568 | -3 | 0.63819 | 0.32258 | 0.18264 | 0.10206 | 17.95923 | 0.012563636 | 0 | Neutral |
| testneg_3907 | 0 | 0 | -0.330275 | -0.330275 | 1.651197 | -3 | 0.10462 | 0.1278 | 0.76918 | 0.13588 | 12.30072 | 0.04 | 0 | Neutral |
| testneg_3908 | 0.08 | 0.03 | -2.363283 | -3 | 1.796297 | -3 | 0.05846 | 0.12788 | 0.10566 | 0.12788 | 18.02082 | 0.041233522 | 0 | Neutral |
| testneg_3909 | 0 | 0 | -0.35638 | -3 | 4.486783 | -3 | 0.18633 | 0.14783 | 0.19898 | 0.12788 | 17.77778 | 0.091180556 | 0 | Neutral |
| testneg_3910 | 0 | 0 | 0.346053 | -3 | 4.017255 | -3 | 0.3047 | 0.11929 | 0.19898 | 0.16603 | 10.83333 | 0.209339203 | 0 | Neutral |
| testneg_3911 | -0.11 | 1.95 | 0.156645 | -3 | 4.133728 | -3 | 0.11802 | 0.09304 | 0.13794 | 0.1141 | 18.70407 | 0.21 | 0 | Neutral |
| testneg_3912 | 0.02 | -0.36 | 6.091135 | -3 | 3.619471 | -3 | 0.13669 | 0.09263 | 0.33026 | 0.30576 | 21.7629 | 0.608615385 | 86.2 | Disease_causing |
| testneg_3913 | 0.16 | -0.37 | 3.256775 | -3 | 2.891351 | -3 | 0.16309 | 0.12788 | 0.93246 | 0.12788 | 18.84426 | 0.1 | 0 | Neutral |
| testneg_3914 | -0.07 | -0.38 | 0.312194 | -3 | 1.540301 | -3 | 0.60251 | 0.39112 | 0.15676 | 0.12788 | 15.59524 | 0.04 | 0 | Neutral |
| testneg_3915 | 0.03 | 0.12 | -1.487172 | -1.487172 | 0.641246 | -3 | 0.103 | 0.403 | 0.21118 | 0.10074 | 16.8018 | 0.05 | 0 | Neutral |
| testneg_3916 | 0.06 | -0.57 | 3.056858 | -3 | 0.203322 | -3 | 0.95535 | 0.12788 | 0.25312 | 0.12788 | 19.42906 | 0.04 | 0 | Neutral |
| testneg_3917 | 0.02 | -0.36 | 3.235017 | -3 | 4.425066 | -3 | 0.30329 | 0.14582 | 0.16355 | 0.20781 | 11.25 | 0.651738777 | 92.6 | Disease_causing |
| testneg_3918 | 0.06 | -0.57 | 5.361725 | -3 | 1.849259 | -3 | 0.06708 | 0.09112 | 0.41261 | 0.12096 | 29.20811 | 0.12 | 0 | Neutral |
| testneg_3919 | 0 | 0 | 1.118558 | -3 | 0.176615 | -3 | 0.06095 | 0.08179 | 0.11114 | 0.09582 | 10.22898 | 0.02 | 0 | Neutral |
| testneg_3920 | 0 | 0 | 1.756005 | -3 | 3.285131 | -3 | 0.22155 | 0.1062 | 0.65981 | 0.47389 | 11.41859 | 0.128160714 | 0 | Neutral |
| testneg_3921 | -0.01 | 0.57 | 2.71072 | -3 | 5.791651 | 4.805369 | 0.09719 | 0.09576 | 0.22415 | 0.70109 | 23.16522 | 0.590291667 | 86.6 | Disease_causing |
| testneg_3922 | 0 | -0.32 | 3.770006 | -3 | 4.759697 | -3 | 0.19898 | 0.11167 | 0.19898 | 0.10335 | 17.77778 | 0.162026645 | 0 | Neutral |
| testneg_3923 | 0.02 | 0.45 | 0.043893 | -3 | 3.1183 | -3 | 0.75906 | 0.09702 | 0.12921 | 0.11605 | 11.30228 | 0.09 | 0 | Neutral |
| testneg_3924 | 0.02 | 0.13 | -0.076183 | -3 | 0.598202 | -3 | 0.16418 | 0.12788 | 0.0959 | 0.12788 | 35.02896 | 0.010690476 | 0 | Neutral |
| testneg_3925 | -0.05 | -0.29 | 1.981611 | -3 | 3.162135 | -3 | 0.24977 | 0.11262 | 0.16524 | 0.12788 | 10.70979 | 0.050133826 | 0 | Neutral |
| testneg_3926 | 0 | 0 | 0.402508 | 0.402508 | -0.763372 | -3 | 0.29794 | 0.11106 | 0.19979 | 0.09328 | 18.68586 | 0.01 | 0 | Neutral |
| testneg_3927 | 0 | 0 | 1.538152 | -3 | 1.092053 | -3 | 0.03109 | 0.10705 | 0.08837 | 0.10939 | 18.89211 | 0.03 | 0 | Neutral |
| testneg_3928 | 0.05 | -0.81 | 0.331709 | 0.331709 | 0.615391 | -3 | 0.16534 | 0.26726 | 0.19898 | 0.19444 | 19.09347 | 0.1734588 | 0 | Neutral |
| testneg_3929 | 0.21 | 0.15 | 6.611007 | -3 | -2.240872 | -3 | 0.40897 | 0.13253 | 0.14175 | 0.12788 | 20.12164 | 0.233205128 | 0.8 | Neutral |
| testneg_3930 | -0.21 | -0.15 | 1.078479 | -3 | 5.428919 | -3 | 0.22817 | 0.10634 | 0.33418 | 0.09979 | 21.5812 | 0.16 | 0 | Neutral |
| testneg_3931 | 0.09 | -0.25 | 1.704591 | -3 | 4.841698 | -3 | 0.56307 | 0.10536 | 0.29356 | 0.46298 | 12.20655 | 0.300233333 | 3 | Neutral |
| testneg_3932 | 0.04 | -0.23 | 3.31131 | -3 | 0.665575 | -3 | 0.19898 | 0.15588 | 0.11613 | 0.1109 | 18.452 | 0.04 | 0 | Neutral |
| testneg_3933 | 0.01 | 0.8 | 6.526897 | -3 | 8.001358 | -3 | 0.16692 | 0.09243 | 0.10895 | 0.11082 | 30.10558 | 0.598712121 | 86.2 | Disease_causing |
| testneg_3934 | 0 | 0 | 0.889458 | 0.889458 | 1.910339 | -3 | 0.48937 | 0.22085 | 0.10161 | 0.04472 | 20.53369 | 0.050480549 | 0 | Neutral |
| testneg_3935 | 0 | 0 | 1.861867 | -3 | -0.221228 | -3 | 0.32963 | 0.1035 | 0.11153 | 0.10767 | 22.38095 | 0.000947682 | 0 | Neutral |
| testneg_3936 | 0 | -0.01 | 1.828167 | -3 | 1.420601 | -3 | 0.25474 | 0.11262 | 0.61332 | 0.1004 | 10.22898 | 0 | 0 | Neutral |
| testneg_3937 | 0 | 0 | 0.493256 | -3 | -0.133106 | -0.133106 | 0.13126 | 0.11628 | 0.07096 | 0.09965 | 19.33934 | 0.01 | 0 | Neutral |
| testneg_3938 | 0.05 | -0.81 | 4.970627 | -3 | 2.896291 | -3 | 0.31648 | 0.12788 | 0.19898 | 0.1092 | 30.4296 | 0.129348901 | 0 | Neutral |
| testneg_3939 | 0 | 0 | 0.181115 | -3 | 6.206483 | -3 | 0.19898 | 0.12788 | 0.18861 | 0.10842 | 15.01525 | 0.073185606 | 0 | Neutral |
| testneg_3940 | 0 | 0 | 2.044018 | -3 | 6.630613 | 1.849259 | 0.63431 | 0.09837 | 0.41261 | 0.12096 | 13.51351 | 0.19005618 | 0.2 | Neutral |
| testneg_3941 | 0.14 | -0.69 | 5.324969 | -3 | 0.551092 | -3 | 0.835 | 0.13588 | 0.10895 | 0.10442 | 10 | 0.167321429 | 0 | Neutral |
| testneg_3942 | 0 | 0 | 0.824678 | -3 | 1.294927 | -3 | 0.6854 | 0.12563 | 0.35486 | 0.11584 | 9.13729 | 0 | 0 | Neutral |
| testneg_3943 | 0.09 | -0.3 | -0.335332 | -3 | 0.489857 | -3 | 0.50388 | 0.12295 | 0.36257 | 0.26115 | 17.60392 | 0.071016231 | 0 | Neutral |
| testneg_3944 | -0.08 | -0.03 | 3.810101 | -3 | 5.305351 | -3 | 0.19898 | 0.12788 | 0.12933 | 0.12788 | 21.38889 | 0.240519481 | 0.6 | Neutral |
| testneg_3945 | 0 | 0 | -0.070594 | -3 | 0.467995 | -3 | 0.17529 | 0.09662 | 0.11237 | 0.08916 | 60.03868 | 0.07 | 0 | Neutral |
| testneg_3946 | 0.06 | -0.57 | 1.369525 | -3 | 4.894593 | -3 | 0.17285 | 0.09314 | 0.13743 | 0.12788 | 10.83333 | 0.08 | 0 | Neutral |
| testneg_3947 | 0.04 | -0.23 | 0.007971 | -3 | 3.437388 | -3 | 0.12591 | 0.09845 | 0.22645 | 0.1704 | 17.917 | 0.192487234 | 0 | Neutral |
| testneg_3948 | 0 | 0 | 0.27864 | -3 | 0.970927 | -3 | 0.17737 | 0.54971 | 0.63497 | 0.23021 | 14.19394 | 0.111063435 | 0 | Neutral |
| testneg_3949 | 0 | 0 | 2.918397 | -3 | -0.244708 | -3 | 0.63378 | 0.09451 | 0.06547 | 0.09123 | 17.21847 | 0.04 | 0 | Neutral |
| testneg_3950 | 0 | 0 | 1.66597 | -3 | 3.506817 | -3 | 0.66027 | 0.23984 | 0.23088 | 0.21602 | 10.83333 | 0.274672392 | 1.4 | Neutral |
| testneg_3951 | 0 | -0.01 | 1.028355 | -3 | -1.487847 | -3 | 0.87835 | 0.13522 | 0.06728 | 0.20664 | 24.83516 | 0.115582251 | 0 | Neutral |
| testneg_3952 | 0 | 0 | 0.287181 | -3 | 0.870138 | -3 | 0.93834 | 0.11088 | 0.26961 | 0.09979 | 23.60378 | 0.015197802 | 0 | Neutral |
| testneg_3953 | 0 | -0.01 | 0.729709 | -3 | 2.23696 | -3 | 0.40557 | 0.11865 | 0.68865 | 0.19414 | 29.16667 | 0.06 | 0 | Neutral |
| testneg_3954 | 0 | 0 | 1.630014 | -3 | -0.196592 | -0.196592 | 0.1109 | 0.17295 | 0.32733 | 0.16869 | 22.00588 | 0.121427989 | 0 | Neutral |
| testneg_3955 | 0 | 0 | 0.372849 | -3 | 1.406345 | -3 | 0.10286 | 0.26819 | 0.7062 | 0.83002 | 10 | 0.134796732 | 0 | Neutral |
| testneg_3956 | 0.05 | -0.81 | 2.508019 | -3 | -0.415438 | -3 | 0.51319 | 0.12971 | 0.19898 | 0.12788 | 31.82796 | 0.02 | 0 | Neutral |
| testneg_3957 | 0.18 | -1.05 | -0.040554 | -3 | -0.03313 | -3 | 0.21848 | 0.11049 | 0.07537 | 0.1399 | 21.38889 | 0.1 | 0 | Neutral |
| testneg_3958 | 0.11 | -1.95 | 6.022895 | -3 | 0.268278 | -3 | 0.09583 | 0.09376 | 0.09669 | 0.12526 | 18.88889 | 0.141414474 | 0 | Neutral |
| testneg_3959 | 0 | 0 | -0.674729 | -3 | 2.51331 | -3 | 0.24765 | 0.32858 | 0.14601 | 0.12788 | 10.83333 | 0.060082645 | 0 | Neutral |
| testneg_3960 | 0 | 0 | -0.055895 | -3 | 2.569635 | -3 | 0.31694 | 0.13475 | 0.3962 | 0.10523 | 17.77778 | 0.020072002 | 0 | Neutral |
| testneg_3961 | -0.06 | 0.57 | 1.042833 | -3 | 0.260712 | -3 | 0.19898 | 0.12788 | 0.17643 | 0.10767 | 12.40248 | 0.010247606 | 0 | Neutral |
| testneg_3962 | 0.08 | 0.03 | -2.363283 | -3 | 6.448066 | -3 | 0.05846 | 0.12788 | 0.40707 | 0.37988 | 18.71322 | 0.414002243 | 21.6 | Neutral |
| testneg_3963 | 0.07 | -0.31 | 4.157651 | -3 | 0.079306 | -3 | 0.19898 | 0.12788 | 0.19898 | 0.12788 | 13.51351 | 0.02 | 0 | Neutral |
| testneg_3964 | 0 | 0 | 0.342408 | 0.342408 | 2.686668 | -3 | 0.5255 | 0.15968 | 0.10517 | 0.09803 | 16.07936 | 0.060469872 | 0 | Neutral |
| testneg_3965 | 0 | -0.32 | -0.590938 | -3 | 5.258974 | -3 | 0.06513 | 0.12353 | 0.19898 | 0.28128 | 21.38889 | 0.33 | 7 | Neutral |
| testneg_3966 | 0 | 0 | 1.243121 | -3 | 0.513838 | -3 | 0.15958 | 0.41628 | 0.68004 | 0.12345 | 6.00877 | 0.020333333 | 0 | Neutral |
| testneg_3967 | 0 | 0 | -0.2352 | -3 | 2.28293 | -3 | 0.19564 | 0.59275 | 0.08332 | 0.09046 | 11.25 | 0.06 | 0 | Neutral |
| testneg_3968 | -0.02 | -0.41 | 0.433545 | -3 | 1.9552 | -3 | 0.06996 | 0.35923 | 0.19237 | 0.53191 | 10 | 0.25 | 0 | Neutral |
| testneg_3969 | 0 | 0 | 0.572592 | -3 | 1.824706 | -3 | 0.68918 | 0.16819 | 0.90955 | 0.10096 | 17.77778 | 0.02 | 0 | Neutral |
| testneg_3970 | 0.00238095 | -0.0190476 | 1.975867 | -3 | -0.940094 | -3 | 0.19898 | 0.12788 | 0.03646 | 0.09863 | 29.26645 | 0.01 | 0 | Neutral |
| testneg_3971 | 0.07 | -0.31 | 2.84362 | -3 | 0.076953 | 0.076953 | 0.2903 | 0.11176 | 0.10187 | 0.12339 | 29.16667 | 0.03 | 0 | Neutral |
| testneg_3972 | -0.22 | 2.66 | 0.984298 | -3 | -0.194401 | -0.194401 | 0.34406 | 0.09894 | 0.06749 | 0.1776 | 12.29167 | 0.2 | 0 | Neutral |
| testneg_3973 | 0 | 0.32 | 3.268811 | -3 | 2.178935 | -3 | 0.17141 | 0.1104 | 0.77847 | 0.10751 | 11.7735 | 0.05 | 0 | Neutral |
| testneg_3974 | 0 | -0.01 | 2.125818 | -3 | 2.28341 | -3 | 0.15713 | 0.3507 | 0.27723 | 0.13229 | 26.69988 | 0.195 | 0 | Neutral |
| testneg_3975 | 0 | 0 | 0.496899 | -3 | 0.65848 | -3 | 0.17325 | 0.17964 | 0.12949 | 0.44208 | 10.41667 | 0.08663257 | 0 | Neutral |
| testneg_3976 | -0.03 | -0.1 | 2.078666 | -3 | 4.326431 | -3 | 0.17576 | 0.12193 | 0.13195 | 0.57267 | 8.97898 | 0.380335783 | 12.8 | Neutral |
| testneg_3977 | 0 | 0 | -0.475685 | -3 | 4.456029 | -3 | 0.30942 | 0.13105 | 0.19898 | 0.09533 | 27.3569 | 0.075471612 | 0 | Neutral |
| testneg_3978 | -0.02 | 0.63 | -1.996459 | -3 | 5.144204 | -3 | 0.04988 | 0.25503 | 0.11261 | 0.13673 | 20.76828 | 0.557834162 | 78 | Disease_causing |
| testneg_3979 | 0.2 | -1.26 | 2.120173 | -3 | 0.298529 | -3 | 0.14477 | 0.18932 | 0.26264 | 0.13812 | 13.60087 | 0.14123511 | 0 | Neutral |
| testneg_3980 | 0.06 | 0.18 | -1.153792 | -3 | 0.77784 | -3 | 0.65284 | 0.2592 | 0.15685 | 0.17839 | 10.31261 | 0.101225379 | 0 | Neutral |
| testneg_3981 | 0 | 0 | -0.163107 | -3 | 1.266152 | -3 | 0.21289 | 0.08709 | 0.1846 | 0.375 | 12.38369 | 0.048250733 | 0 | Neutral |
| testneg_3982 | 0.07 | 0.01 | -1.422633 | -3 | 0.166429 | -3 | 0.07598 | 0.08456 | 0.81659 | 0.10424 | 9.87205 | 0.02 | 0 | Neutral |
| testneg_3983 | -0.03 | -0.1 | 2.112536 | -3 | -0.165443 | -3 | 0.11873 | 0.10499 | 0.05221 | 0.12788 | 32.0954 | 0.02 | 0 | Neutral |
| testneg_3984 | 0 | 0 | 0.992597 | -3 | -0.771216 | -3 | 0.30122 | 0.09165 | 0.37986 | 0.14229 | 15.51047 | 0.020296703 | 0 | Neutral |
| testneg_3985 | -0.07 | -0.7 | 2.642734 | -3 | 7.04333 | -3 | 0.19898 | 0.12788 | 0.17643 | 0.10767 | 23.24968 | 0.314510135 | 3.4 | Neutral |
| testneg_3986 | 0.05 | -0.81 | 3.217486 | -3 | 0.713568 | -3 | 0.74699 | 0.12286 | 0.11072 | 0.38259 | 12.29167 | 0.184197012 | 0 | Neutral |
| testneg_3987 | -0.17 | -0.38 | 2.753905 | -3 | -0.593094 | -3 | 0.05904 | 0.09282 | 0.19898 | 0.12788 | 57.64633 | 0.10041958 | 0 | Neutral |
| testneg_3988 | 0 | 0 | 0.760575 | -3 | 0.96317 | -3 | 0.1878 | 0.08946 | 0.20379 | 0.35591 | 14.43172 | 0.038477564 | 0 | Neutral |
| testneg_3989 | 0 | 0 | 0.864006 | -3 | -0.696429 | -3 | 0.19898 | 0.12788 | 0.26476 | 0.08217 | 37.10735 | 0 | 0 | Neutral |
| testneg_3990 | 0.07 | 0.38 | 0.698423 | -3 | 1.015573 | -3 | 0.64152 | 0.15239 | 0.16974 | 0.80184 | 21.95402 | 0.163491186 | 0 | Neutral |
| testneg_3991 | 0 | 0 | -0.409329 | -3 | 2.876648 | -3 | 0.95993 | 0.15588 | 0.06376 | 0.0935 | 22.44024 | 0.07625 | 0 | Neutral |
| testneg_3992 | 0 | -0.01 | -0.3595 | -3 | -1.543536 | -3 | 0.19898 | 0.12788 | 0.06744 | 0.12788 | 24.44444 | 0.010540936 | 0 | Neutral |
| testneg_3993 | 0 | 0 | 0.232243 | -3 | -0.103471 | -3 | 0.49745 | 0.1259 | 0.0669 | 0.10997 | 19.44444 | 0.014155797 | 0 | Neutral |
| testneg_3994 | 0 | -0.01 | 1.025435 | -3 | 3.034495 | -3 | 0.73731 | 0.26107 | 0.51934 | 0.12046 | 10.83333 | 0.057929293 | 0 | Neutral |
| testneg_3995 | -0.01 | -0.79 | 1.966701 | -3 | -0.19988 | -3 | 0.56285 | 0.18619 | 0.45399 | 0.10245 | 26.34921 | 0.05 | 0 | Neutral |
| testneg_3996 | 0.01 | -0.57 | 1.940863 | -3 | 3.844779 | -3 | 0.24038 | 0.15296 | 0.04063 | 0.05848 | 27.26496 | 0.120833333 | 0 | Neutral |
| testneg_3997 | 0 | 0 | 0.760575 | -3 | -0.603839 | -3 | 0.1878 | 0.08946 | 0.16485 | 0.13045 | 13.51351 | 0.01 | 0 | Neutral |
| testneg_3998 | 0.07 | 0.38 | 1.79061 | -3 | 4.202319 | -3 | 0.12112 | 0.11353 | 0.19898 | 0.30319 | 17.47964 | 0.297261197 | 3.2 | Neutral |
| testneg_3999 | 0 | 0 | 1.178544 | -3 | 4.020687 | -3 | 0.19898 | 0.12788 | 0.34001 | 0.12788 | 43.4379 | 0.040386905 | 0 | Neutral |
| testneg_4000 | -0.03 | -0.1 | 3.350172 | -3 | 3.29285 | -3 | 0.13331 | 0.13638 | 0.42214 | 0.21008 | 18.33333 | 0.471992807 | 43.8 | Neutral |
| testneg_4001 | 0.22 | -2.66 | 2.864158 | -3 | 2.058438 | -3 | 0.61718 | 0.1648 | 0.0749 | 0.12788 | 8.49181 | 0.149333333 | 0 | Neutral |
| testneg_4002 | 0 | 0 | 1.569299 | -3 | 4.70798 | -3 | 0.34153 | 0.08855 | 0.17527 | 0.13941 | 19.22222 | 0.180966792 | 0 | Neutral |
| testneg_4003 | 0 | -0.01 | 1.312464 | -3 | -0.971742 | -0.971742 | 0.22489 | 0.16646 | 0.17222 | 0.12788 | 18.33333 | 0.012426471 | 0 | Neutral |
| testneg_4004 | -0.07 | -0.38 | 3.412931 | -3 | 4.406142 | -3 | 0.18964 | 0.34036 | 0.14735 | 0.12788 | 10.41667 | 0.386849211 | 17.8 | Neutral |
| testneg_4005 | 0.08 | 0.03 | -0.301086 | -3 | 1.811846 | -3 | 0.1192 | 0.1508 | 0.1287 | 0.10423 | 18.88889 | 0.044329309 | 0 | Neutral |
| testneg_4006 | 0.23 | -1.04 | 6.202748 | -3 | -0.225897 | -3 | 0.4172 | 0.11643 | 0.46037 | 0.1343 | 16.8018 | 0.22 | 0.4 | Neutral |
| testneg_4007 | 0 | 0 | -0.027704 | -3 | 3.999282 | -3 | 0.45299 | 0.16306 | 0.07195 | 0.10356 | 12.29167 | 0.114880104 | 0 | Neutral |
| testneg_4008 | -0.03 | -0.1 | 2.059294 | -3 | 1.087899 | -3 | 0.66711 | 0.31094 | 0.19898 | 0.0522 | 20.89902 | 0.030791925 | 0 | Neutral |
| testneg_4009 | -0.03 | -0.1 | 0.539897 | -3 | 0.977938 | -3 | 0.09866 | 0.104 | 0.47975 | 0.18568 | 16.87457 | 0.060393288 | 0 | Neutral |
| testneg_4010 | 0.08 | 0.03 | 1.795418 | -3 | 2.455519 | -3 | 0.11055 | 0.19829 | 0.04668 | 0.12788 | 20.93158 | 0.11 | 0 | Neutral |
| testneg_4011 | 0.21 | 0.15 | 2.141778 | -3 | 2.614089 | -3 | 0.19898 | 0.21177 | 0.16052 | 0.12788 | 23.27951 | 0.145257576 | 0 | Neutral |
| testneg_4012 | -0.12 | 1.14 | 4.710069 | -3 | 1.214807 | -3 | 0.50461 | 0.11034 | 0.56204 | 0.10762 | 5.94234 | 0.139767611 | 0 | Neutral |
| testneg_4013 | 0 | 0 | 0.302733 | -3 | 5.650938 | -3 | 0.19898 | 0.12788 | 0.16671 | 0.18311 | 28.05556 | 0.327974258 | 6.2 | Neutral |
| testneg_4014 | 0.03 | -1.03 | 6.512471 | -3 | 2.23696 | -3 | 0.35584 | 0.10874 | 0.68865 | 0.19414 | 25 | 0.46909707 | 41.6 | Neutral |
| testneg_4015 | 0 | 0 | -0.258914 | -3 | 5.438293 | -3 | 0.15623 | 0.1447 | 0.25029 | 0.12267 | 17.40099 | 0.104002165 | 0 | Neutral |
| testneg_4016 | 0 | 0 | 1.675065 | -3 | 1.003508 | -3 | 0.17285 | 0.09314 | 0.20431 | 0.17304 | 13.38585 | 0.020465368 | 0 | Neutral |
| testneg_4017 | -0.05 | -0.29 | 2.477566 | -3 | -0.586534 | -0.586534 | 0.29451 | 0.13582 | 0.09439 | 0.1884 | 22.62502 | 0.13198539 | 0 | Neutral |
| testneg_4018 | 0.01 | -1.44 | 1.53664 | -3 | 7.140042 | -3 | 0.17576 | 0.12193 | 0.04142 | 0.09564 | 18.6768 | 0.27 | 1.6 | Neutral |
| testneg_4019 | 0.22 | -2.66 | 6.839727 | -3 | -0.586062 | -3 | 0.76186 | 0.10052 | 0.04032 | 0.12788 | 20.83333 | 0.23 | 0.4 | Neutral |
| testneg_4020 | 0 | 0 | 1.512957 | -3 | 3.544108 | -3 | 0.29548 | 0.10732 | 0.13974 | 0.10225 | 19.81145 | 0.026547619 | 0 | Neutral |
| testneg_4021 | 0 | -0.01 | 1.376749 | -3 | 5.604435 | -3 | 0.46242 | 0.39055 | 0.10261 | 0.0933 | 11.08333 | 0.157534119 | 0 | Neutral |
| testneg_4022 | 0.21 | 0.15 | 3.452327 | -3 | 5.124676 | -3 | 0.08047 | 0.06746 | 0.26068 | 0.46457 | 32.9221 | 0.4625 | 39 | Neutral |
| testneg_4023 | 0.21 | 0.15 | -0.029006 | -3 | 1.413393 | -3 | 0.13994 | 0.11181 | 0.2293 | 0.12788 | 19.56386 | 0.020732323 | 0 | Neutral |
| testneg_4024 | 0 | 0 | 0.007189 | -3 | 3.89714 | -3 | 0.41647 | 0.11239 | 0.33022 | 0.13818 | 17.86874 | 0.178219975 | 0 | Neutral |
| testneg_4025 | 0.15 | -0.28 | 1.45686 | -3 | 1.340722 | -3 | 0.38552 | 0.14172 | 0.35421 | 0.10694 | 8.97898 | 0.011645022 | 0 | Neutral |

| | | | | | | | | | | | | | | |
|---|---|---|---|---|---|---|---|---|---|---|---|---|---|---|
| testneg_4026 | 0.09 | -0.67 | 4.592967 | -3 | 3.497813 | 3.497813 | 0.63484 | 0.16728 | 0.05873 | 0.09792 | 18.17595 | 0.358046133 | 7.2 | Neutral |
| testneg_4027 | -0.07 | -0.01 | 2.087185 | -3 | 2.054988 | 1.699603 | 0.06392 | 0.12788 | 0.15406 | 0.1454 | 13.51351 | 0.107929825 | 0 | Neutral |
| testneg_4028 | 0 | 0 | 0.739398 | -3 | 4.344355 | -3 | 0.48906 | 0.11262 | 0.74919 | 0.85508 | 31.20854 | 0.255111111 | 0.6 | Neutral |
| testneg_4029 | 0.1 | 1.9 | 3.279128 | -3 | 4.585021 | -3 | 0.1105 | 0.44945 | 0.44774 | 0.15015 | 7.61555 | 0.773587118 | 100 | Disease_causing |
| testneg_4030 | 0 | 0 | 0.205123 | -3 | 4.028152 | -3 | 0.12073 | 0.10969 | 0.19898 | 0.12788 | 10 | 0.04 | 0 | Neutral |
| testneg_4031 | 0.05 | -0.81 | 5.140271 | -3 | 5.567762 | 5.567762 | 0.09542 | 0.14806 | 0.74641 | 0.66512 | 10.41667 | 0.828708333 | 100 | Disease_causing |
| testneg_4032 | 0 | 0 | 1.262077 | -3 | 4.617328 | -3 | 0.11351 | 0.10667 | 0.19898 | 0.12788 | 18.71981 | 0.03811553 | 0 | Neutral |
| testneg_4033 | 0 | 0 | -0.859115 | -3 | 1.672333 | 1.672333 | 0.19898 | 0.12788 | 0.15338 | 0.13614 | 30.82301 | 0.087852564 | 0 | Neutral |
| testneg_4034 | 0.16 | -1.37 | 6.076778 | -3 | 0.249822 | -3 | 0.33818 | 0.12788 | 0.40021 | 0.21702 | 10.83333 | 0.424833333 | 28.6 | Neutral |
| testneg_4035 | 0.18 | -1.05 | 2.305068 | -3 | 1.056502 | -3 | 0.19898 | 0.12788 | 0.08686 | 0.0953 | 51.10614 | 0.04 | 0 | Neutral |
| testneg_4036 | 0.01 | 0.8 | 4.146576 | -3 | 4.828386 | -3 | 0.18488 | 0.53379 | 0.33467 | 0.12788 | 19.34552 | 0.499222222 | 54.6 | Disease_causing |
| testneg_4037 | 0 | 0 | 0.310102 | -3 | 1.349086 | -3 | 0.12576 | 0.08486 | 0.17199 | 0.10586 | 10.51205 | 0.01 | 0 | Neutral |
| testneg_4038 | 0.09 | -0.67 | 4.020687 | -3 | 4.134413 | -3 | 0.34001 | 0.12788 | 0.10781 | 0.09901 | 29.98419 | 0.15003367 | 0 | Neutral |
| testneg_4039 | 0 | 0 | 0.318608 | -3 | 0.725803 | -3 | 0.11373 | 0.10739 | 0.19898 | 0.12788 | 11.875 | 0 | 0 | Neutral |
| testneg_4040 | 0 | 0 | 0.547246 | -3 | -0.039157 | -3 | 0.81169 | 0.11096 | 0.04606 | 0.20522 | 18.33333 | 0.06 | 0 | Neutral |
| testneg_4041 | -0.0025714 | -0.1897143 | 2.053049 | -3 | 0.797383 | -3 | 0.26586 | 0.10825 | 0.10314 | 0.08894 | 11.25 | 0.01 | 0 | Neutral |
| testneg_4042 | 0 | 0 | -0.50303 | -3 | 2.351073 | -3 | 0.15192 | 0.14302 | 0.16957 | 0.12788 | 10.22898 | 0.020575658 | 0 | Neutral |
| testneg_4043 | 0.02 | -0.63 | 5.557447 | -3 | 0.125659 | -3 | 0.34633 | 0.47228 | 0.21289 | 0.1079 | 16.35932 | 0.242420635 | 0.8 | Neutral |
| testneg_4044 | -0.03 | -0.1 | 6.666841 | -3 | -1.30762 | -3 | 0.1434 | 0.09092 | 0.4676 | 0.11257 | 55.15466 | 0.209861111 | 0 | Neutral |
| testneg_4045 | 0.08 | -1.2 | 1.056242 | -3 | 0.24062 | 0.24062 | 0.19832 | 0.19622 | 0.18351 | 0.11213 | 15.29221 | 0.05 | 0 | Neutral |
| testneg_4046 | 0 | 0 | 0.525606 | -3 | 5.393767 | 1.869406 | 0.08401 | 0.16123 | 0.0882 | 0.12788 | 14.16667 | 0.227185897 | 0.4 | Neutral |
| testneg_4047 | 0.02 | 0.13 | 5.712534 | -3 | 1.335647 | -3 | 0.24026 | 0.12788 | 0.54876 | 0.22335 | 8.97898 | 0.441434779 | 31.8 | Neutral |
| testneg_4048 | -0.08 | -0.03 | -0.451006 | -3 | -1.186234 | -3 | 0.3425 | 0.11597 | 0.35843 | 0.15758 | 11.875 | 0.050232684 | 0 | Neutral |
| testneg_4049 | -0.19 | 0.27 | -1.643119 | -3 | -0.800576 | -3 | 0.1535 | 0.12382 | 0.05245 | 0.13458 | 40.76917 | 0.15 | 0 | Neutral |
| testneg_4050 | -0.04 | 1.61 | 12.292649 | -3 | 0.513578 | 0.513578 | 0.1247 | 0.10331 | 0.19898 | 0.07409 | 42.7598 | 0.34 | 4.6 | Neutral |
| testneg_4051 | -0.15 | -0.04 | 4.2907 | 0.368341 | 1.864364 | -3 | 0.77847 | 0.10751 | 0.12943 | 0.14391 | 11.25 | 0.23 | 0.2 | Neutral |
| testneg_4052 | -0.14 | 1.85 | 0.310587 | 0.310587 | 0.258063 | -3 | 0.2132 | 0.12788 | 0.99941 | 0.50288 | 11.59277 | 0.21 | 0 | Neutral |
| testneg_4053 | -0.19 | -0.51 | 3.374283 | -3 | -0.01248 | -3 | 0.19898 | 0.28128 | 0.61231 | 0.13362 | 19.44444 | 0.250178571 | 0.2 | Neutral |
| testneg_4054 | 0 | 0 | 2.493306 | -3 | 0.395779 | -3 | 0.23506 | 0.24555 | 0.46723 | 0.08832 | 10.25326 | 0.011368081 | 0 | Neutral |
| testneg_4055 | 0.03 | -1.03 | 1.559704 | -3 | 1.557286 | -3 | 0.0138 | 0.06769 | 0.27705 | 0.22346 | 18.33333 | 0.11 | 0 | Neutral |
| testneg_4056 | 0.09 | -0.25 | 1.300233 | -3 | 1.245581 | -1.330911 | 0.13119 | 0.12788 | 0.60793 | 0.18264 | 26.80135 | 0.07 | 0 | Neutral |
| testneg_4057 | 0.07 | 0.7 | 0.208899 | 0.208899 | -2.721335 | -3 | 0.64608 | 0.35524 | 0.19898 | 0.09927 | 15.97286 | 0.06 | 0 | Neutral |
| testneg_4058 | -0.06 | -0.18 | 4.235243 | -3 | 4.720694 | -3 | 0.14806 | 0.09574 | 0.12453 | 0.17736 | 18.88889 | 0.61010582 | 87 | Disease_causing |
| testneg_4059 | 0.07 | 0.33 | 2.134433 | -3 | 1.217986 | -3 | 0.27887 | 0.19826 | 0.19898 | 0.12788 | 29.96713 | 0.04 | 0 | Neutral |
| testneg_4060 | 0.05 | 0.29 | -1.828653 | -3 | 4.444691 | -3 | 0.4097 | 0.36733 | 0.32022 | 0.09529 | 10 | 0.181676966 | 0 | Neutral |
| testneg_4061 | -0.14 | -0.54 | 3.948867 | -3 | 6.051826 | -3 | 0.21236 | 0.12788 | 0.2559 | 0.10848 | 21.63869 | 0.2819875 | 2.4 | Neutral |
| testneg_4062 | 0.04 | -0.23 | -1.047288 | -3 | 0.272659 | 0.272659 | 0.52005 | 0.12788 | 0.17093 | 0.23281 | 22.71196 | 0.099855418 | 0 | Neutral |
| testneg_4063 | 0.15 | 0.06 | 4.843192 | -3 | 1.940661 | -3 | 0.25758 | 0.2262 | 0.25151 | 0.12788 | 19.26372 | 0.153689353 | 0 | Neutral |
| testneg_4064 | 0.03 | -0.85 | 1.181282 | -3 | 1.507517 | -3 | 0.113 | 0.12788 | 0.19898 | 0.10482 | 71.0688 | 0.08 | 0 | Neutral |
| testneg_4065 | 0 | 0 | 0.178815 | -3 | -1.141522 | -3 | 0.5471 | 0.11262 | 0.09555 | 0.09639 | 19.09347 | 0.02 | 0 | Neutral |
| testneg_4066 | 0 | 0 | 0.777519 | -3 | 0.602917 | -3 | 0.75071 | 0.63704 | 0.16125 | 0.08406 | 14.14796 | 0.02 | 0 | Neutral |
| testneg_4067 | 0.17 | 0.38 | 4.475922 | -3 | 1.792139 | 1.792139 | 0.19898 | 0.20319 | 0.1109 | 0.12788 | 23.68839 | 0.147718254 | 0 | Neutral |
| testneg_4068 | -0.01 | -0.8 | 6.19532 | -3 | -0.416655 | -0.416655 | 0.36257 | 0.26115 | 0.05453 | 0.12788 | 19.07749 | 0.281269231 | 1.4 | Neutral |
| testneg_4069 | 0 | -0.32 | 3.805191 | -3 | -0.37295 | -3 | 0.56307 | 0.10536 | 0.07237 | 0.12788 | 20.37918 | 0.04 | 0 | Neutral |
| testneg_4070 | 0.03 | -1.03 | 6.867053 | -3 | -0.174435 | -3 | 0.08878 | 0.0974 | 0.31542 | 0.26739 | 14.45887 | 0.41 | 23.4 | Neutral |
| testneg_4071 | 0 | 0 | -0.034875 | -3 | 0.099246 | -3 | 0.2547 | 0.07589 | 0.58124 | 0.31208 | 10 | 0.031728666 | 0 | Neutral |
| testneg_4072 | -0.0138182 | -0.2629091 | 4.820901 | -3 | 0.982788 | -3 | 0.06358 | 0.5364 | 0.1481 | 0.1117 | 18.69495 | 0.200857143 | 0 | Neutral |
| testneg_4073 | 0.01 | -1.44 | 0.634175 | -3 | 2.025379 | -3 | 0.37637 | 0.12788 | 0.13374 | 0.12788 | 10 | 0.03 | 0 | Neutral |
| testneg_4074 | 0.1 | 1.9 | -1.532659 | -3 | 1.36361 | -3 | 0.39258 | 0.12788 | 0.74881 | 0.12788 | 21.25537 | 0.1 | 0 | Neutral |
| testneg_4075 | 0.01 | 0.79 | 0.696048 | -3 | -1.087729 | -3 | 0.25025 | 0.18924 | 0.03563 | 0.10455 | 18.33333 | 0.031579365 | 0 | Neutral |
| testneg_4076 | -0.02 | 0.36 | 1.388896 | -3 | 0.043004 | -3 | 0.40398 | 0.12788 | 0.3012 | 0.10317 | 19.30894 | 0.01 | 0 | Neutral |
| testneg_4077 | 0.07 | 0.38 | -3.00145 | -3 | 0.437329 | -3 | 0.22645 | 0.1704 | 0.19898 | 0.12788 | 17.917 | 0.03089738 | 0 | Neutral |
| testneg_4078 | 0 | -0.01 | 1.245581 | -3 | 0.641608 | -3 | 0.60793 | 0.18264 | 0.32458 | 0.1016 | 12.31894 | 0.01 | 0 | Neutral |
| testneg_4079 | 0.04 | -0.23 | 5.690551 | -3 | 0.014287 | -3 | 0.33264 | 0.10493 | 0.30103 | 0.1462 | 20.98109 | 0.222183333 | 0.8 | Neutral |
| testneg_4080 | -0.07 | -0.38 | 2.4232 | -3 | 4.454666 | -3 | 0.38002 | 0.22508 | 0.09462 | 0.11807 | 19.65901 | 0.204583333 | 0 | Neutral |
| testneg_4081 | -0.07 | -0.01 | 0.141736 | -3 | 0.941458 | -3 | 0.0811 | 0.09449 | 0.16424 | 0.11262 | 27.4024 | 0.01 | 0 | Neutral |
| testneg_4082 | 0 | 0 | 0.19536 | -3 | 1.729635 | 1.729635 | 0.25009 | 0.11017 | 0.57588 | 0.09682 | 11.05428 | 0.02 | 0 | Neutral |
| testneg_4083 | 0.02 | 0.45 | -0.887241 | -3 | -0.024667 | -3 | 0.19898 | 0.12788 | 0.59599 | 0.12477 | 17.21847 | 0.01 | 0 | Neutral |
| testneg_4084 | 0 | 0 | 5.586363 | -3 | -2.290249 | -3 | 0.65909 | 0.31908 | 0.14862 | 0.12788 | 21.78922 | 0.24842146 | 0.4 | Neutral |
| testneg_4085 | 0 | -0.01 | 1.125724 | -3 | 3.612387 | -3 | 0.6185 | 0.09316 | 0.17075 | 0.12788 | 20.53807 | 0.07 | 0 | Neutral |
| testneg_4086 | 0.03 | -1.03 | 5.957726 | -3 | -1.803416 | -3 | 0.44103 | 0.10698 | 0.62381 | 0.12788 | 35.59312 | 0.17 | 0 | Neutral |
| testneg_4087 | 0 | 0 | 1.150515 | -3 | -1.00023 | -3 | 0.12408 | 0.08828 | 0.43158 | 0.10472 | 10.83333 | 0.01 | 0 | Neutral |
| testneg_4088 | 0.05 | -0.81 | 1.982936 | -3 | 1.492764 | -3 | 0.07044 | 0.11679 | 0.10822 | 0.10942 | 21.79117 | 0.010099635 | 0 | Neutral |
| testneg_4089 | 0 | 0 | 0.452231 | -3 | 4.813885 | -3 | 0.46691 | 0.07886 | 0.08237 | 0.09015 | 19.68594 | 0.063629227 | 0 | Neutral |
| testneg_4090 | -0.1 | -1.9 | 1.709952 | -3 | 3.580866 | 3.580866 | 0.4172 | 0.11643 | 0.3012 | 0.10317 | 18.75339 | 0.233333333 | 0.4 | Neutral |
| testneg_4091 | -0.0034545 | 0.00218182 | 9.919363 | -3 | 4.025857 | -3 | 0.11351 | 0.10667 | 0.50275 | 0.38957 | 12.15583 | 0.721667565 | 99.2 | Disease_causing |
| testneg_4092 | 0 | -0.32 | 2.219571 | -3 | 0.941706 | -3 | 0.25827 | 0.21386 | 0.15294 | 0.12788 | 17.21847 | 0.021278711 | 0 | Neutral |
| testneg_4093 | 0 | 0 | 0.628299 | -3 | 5.833165 | -3 | 0.40612 | 0.45135 | 0.25179 | 0.09355 | 10.22898 | 0.13491958 | 0 | Neutral |
| testneg_4094 | 0 | 0 | 0.002341 | -3 | -0.253658 | -3 | 0.10799 | 0.10754 | 0.07293 | 0.12989 | 10.28409 | 0.000801282 | 0 | Neutral |
| testneg_4095 | 0 | 0 | -0.011496 | -3 | -0.757789 | -3 | 0.01409 | 0.11307 | 0.40491 | 0.17596 | 19.44444 | 0.096533966 | 0 | Neutral |
| testneg_4096 | 0 | 0 | 1.641098 | -3 | -2.464297 | -2.464297 | 0.13906 | 0.09925 | 0.11524 | 0.12788 | 17.21847 | 0 | 0 | Neutral |
| testneg_4097 | 0 | 0 | 2.988133 | -3 | 4.311484 | -3 | 0.99338 | 0.14229 | 0.29525 | 0.12788 | 16.875 | 0.209546784 | 0.2 | Neutral |
| testneg_4098 | 0.18 | -1.05 | 3.859718 | -3 | 0.728127 | -3 | 0.19898 | 0.12788 | 0.07681 | 0.22865 | 18.13538 | 0.207369565 | 0 | Neutral |
| testneg_4099 | 0 | 0 | 0.260929 | -3 | 4.565186 | -3 | 0.53654 | 0.22867 | 0.31744 | 0.10986 | 18.6768 | 0.092970779 | 0 | Neutral |
| testneg_4100 | -0.22 | 2.66 | 1.021201 | 1.021201 | 3.329851 | -3 | 0.15785 | 0.35538 | 0.5184 | 0.11889 | 6.658 | 0.38 | 6.8 | Neutral |
| testneg_4101 | 0 | 0 | 1.264197 | -3 | 3.350172 | -3 | 0.37542 | 0.19444 | 0.13331 | 0.13638 | 12.70833 | 0.268073051 | 1.4 | Neutral |
| testneg_4102 | 0.022 | 0.01145455 | 0.7799 | -3 | -0.887241 | -3 | 0.60285 | 0.15179 | 0.19898 | 0.12788 | 16.8018 | 0.017083612 | 0 | Neutral |
| testneg_4103 | -0.03 | -0.1 | 1.347962 | -3 | 2.171434 | -3 | 0.0418 | 0.12788 | 0.49425 | 0.25538 | 16.8018 | 0.150304862 | 0 | Neutral |
| testneg_4104 | 0.17 | 0.38 | 0.917643 | -3 | 2.261013 | -1.624227 | 0.37074 | 0.10627 | 0.19237 | 0.53191 | 11.6958 | 0.15 | 0 | Neutral |
| testneg_4105 | -0.03 | -0.1 | 3.202429 | -3 | 1.513487 | -3 | 0.39964 | 0.22293 | 0.70872 | 0.09884 | 22.50346 | 0.100568182 | 0 | Neutral |
| testneg_4106 | 0.15 | 0.06 | -0.025439 | -3 | 4.752866 | -3 | 0.21355 | 0.12788 | 0.75728 | 0.34838 | 19.88534 | 0.288048246 | 2 | Neutral |
| testneg_4107 | -0.15 | 1.06 | 2.902826 | -3 | 4.189674 | -3 | 0.21596 | 0.08969 | 0.15092 | 0.09074 | 20.8819 | 0.25 | 0 | Neutral |
| testneg_4108 | 0.03 | 1.22 | 1.359632 | -3 | 1.586397 | 1.586397 | 0.38171 | 0.15977 | 0.65284 | 0.2592 | 12.00275 | 0.156761905 | 0 | Neutral |
| testneg_4109 | 0.14 | -1.01 | -0.60097 | -3 | 1.971338 | -3 | 0.18153 | 0.10845 | 0.09682 | 0.09596 | 16.16034 | 0.02 | 0 | Neutral |
| testneg_4110 | -0.07 | -0.01 | 3.213612 | 1.535215 | -0.09441 | -3 | 0.26323 | 0.2399 | 0.32188 | 0.08974 | 11.875 | 0.126166667 | 0 | Neutral |
| testneg_4111 | 0.14 | -0.37 | 0.334634 | -3 | -0.187363 | -3 | 0.18925 | 0.32479 | 0.90649 | 0.3543 | 5 | 0.17 | 0 | Neutral |
| testneg_4112 | 0 | 0 | -0.363052 | -3 | 4.563044 | -3 | 0.3437 | 0.5169 | 0.01952 | 0.08329 | 10.41667 | 0.103139148 | 0 | Neutral |
| testneg_4113 | 0.01 | -0.57 | 0.379634 | -3 | 0.752523 | -3 | 0.87611 | 0.16759 | 0.65857 | 0.26294 | 10.88192 | 0.076333333 | 0 | Neutral |
| testneg_4114 | 0 | 0 | 0.580159 | -3 | 1.786946 | -3 | 0.35908 | 0.18099 | 0.20577 | 0.12653 | 11.25 | 0.01029439 | 0 | Neutral |
| testneg_4115 | 0 | 0 | 0.726872 | -3 | 1.105669 | -3 | 0.04988 | 0.25503 | 0.44385 | 0.1311 | 5 | 0.111302521 | 0 | Neutral |
| testneg_4116 | -0.03 | -0.1 | 0.461043 | -3 | -1.367259 | -3 | 0.62864 | 0.11103 | 0.06099 | 0.12788 | 22.33173 | 0.03 | 0 | Neutral |
| testneg_4117 | 0.22 | -2.66 | 5.935147 | -3 | 0.345417 | -3 | 0.19898 | 0.10269 | 0.07996 | 0.08354 | 44.4235 | 0.13 | 0 | Neutral |
| testneg_4118 | 0.14 | -0.37 | 2.030208 | -3 | 3.118861 | -3 | 0.06957 | 0.07707 | 0.37275 | 0.16262 | 18.66082 | 0.236214213 | 0 | Neutral |
| testneg_4119 | 0 | 0 | 0.699729 | -0.925939 | 1.432015 | -3 | 0.39192 | 0.09135 | 0.64108 | 0.51829 | 10.55958 | 0.031374365 | 0 | Neutral |
| testneg_4120 | 0.04 | -0.23 | 7.35694 | -3 | 1.756423 | -3 | 0.32669 | 0.10659 | 0.2521 | 0.1784 | 12.70833 | 0.487954545 | 49.4 | Neutral |
| testneg_4121 | 0 | 0 | 0.40958 | -3 | 1.668583 | -3 | 0.06728 | 0.20664 | 0.04869 | 0.05922 | 14.28427 | 0.037574337 | 0 | Neutral |
| testneg_4122 | 0.07 | 0.38 | 0.972199 | 0.972199 | 2.83359 | -3 | 0.42751 | 0.10919 | 0.19898 | 0.10512 | 18.88889 | 0.068759804 | 0 | Neutral |
| testneg_4123 | 0 | 0 | 0.550269 | -3 | 3.551618 | -3 | 0.22169 | 0.11262 | 0.10249 | 0.1399 | 19.90556 | 0.20495216 | 0.4 | Neutral |
| testneg_4124 | -0.07 | -0.38 | 1.102328 | -3 | -0.691406 | -3 | 0.74271 | 0.10951 | 0.04046 | 0.06599 | 20.98109 | 0.020571895 | 0 | Neutral |
| testneg_4125 | 0.07 | 0.01 | 1.001127 | -3 | 1.897319 | -3 | 0.14 | 0.12151 | 0.39225 | 0.10625 | 12.29167 | 0.000470265 | 0 | Neutral |
| testneg_4126 | -0.2 | 1.26 | -0.70557 | -3 | -1.158834 | -3 | 0.29008 | 0.19268 | 0.12739 | 0.12788 | 10.30286 | 0.15 | 0 | Neutral |
| testneg_4127 | 0 | 0 | 0.546525 | -3 | 3.161955 | -3 | 0.12545 | 0.08963 | 0.11229 | 0.08897 | 26.04838 | 0.05 | 0 | Neutral |
| testneg_4128 | 0.14 | -1.01 | 0.772419 | -3 | 4.361375 | -3 | 0.83936 | 0.42313 | 0.10459 | 0.12783 | 18.6768 | 0.220331028 | 0 | Neutral |
| testneg_4129 | -0.01 | 0.57 | 3.674344 | -3 | 7.386822 | -3 | 0.13565 | 0.46572 | 0.20249 | 0.10327 | 10.83333 | 0.63 | 91.2 | Disease_causing |
| testneg_4130 | -0.05 | 0.81 | 1.623152 | 1.623152 | -0.042671 | -3 | 0.19898 | 0.12788 | 0.20951 | 0.16278 | 18.33333 | 0.148934115 | 0 | Neutral |
| testneg_4131 | 0.07 | -0.31 | 1.218458 | -3 | -0.684283 | -3 | 0.13119 | 0.12788 | 0.2098 | 0.44672 | 29.45989 | 0.089666667 | 0 | Neutral |
| testneg_4132 | -0.06 | -0.18 | 3.350472 | -0.969128 | -0.791573 | -3 | 0.09577 | 0.2436 | 0.12886 | 0.12788 | 27.23767 | 0.103143939 | 0 | Neutral |
| testneg_4133 | 0 | 0 | 0.284143 | -3 | 0.651919 | -3 | 0.71762 | 0.14417 | 0.15025 | 0.4872 | 18.70407 | 0.070335097 | 0 | Neutral |
| testneg_4134 | 0 | 0 | 1.565142 | -3 | 1.022212 | 0.433904 | 0.493 | 0.35835 | 0.09093 | 0.28643 | 12.70833 | 0.15037037 | 0 | Neutral |
| testneg_4135 | 0.2 | -1.26 | 4.671744 | -3 | 6.433557 | -3 | 0.29302 | 0.17394 | 0.16621 | 0.49845 | 13.94832 | 0.825249098 | 99.8 | Disease_causing |
| testneg_4136 | 0 | 0 | 4.240578 | -3 | 1.542868 | -3 | 0.17903 | 0.37232 | 0.68233 | 0.58998 | 4.44444 | 0.382916451 | 14.8 | Neutral |
| testneg_4137 | 0 | -0.32 | 4.577438 | -3 | 2.457626 | -3 | 0.56966 | 0.68625 | 0.33952 | 0.12788 | 18.6768 | 0.253988095 | 0.4 | Neutral |
| testneg_4138 | 0.03 | 0.1 | 0.500506 | -3 | 1.021201 | 1.021201 | 0.68233 | 0.58998 | 0.15785 | 0.35538 | 4.72222 | 0.17410776 | 0 | Neutral |
| testneg_4139 | -0.03 | -0.1 | 5.381477 | -3 | 4.935141 | -3 | 0.34698 | 0.12788 | 0.10383 | 0.08853 | 21.93548 | 0.348535714 | 8.4 | Neutral |
| testneg_4140 | 0.08 | 0.03 | -0.130309 | -3 | 4.54534 | -3 | 0.07049 | 0.12788 | 0.06713 | 0.11112 | 29.93981 | 0.09 | 0 | Neutral |
| testneg_4141 | -0.02 | -0.45 | 6.344462 | -3 | 0.985076 | -3 | 0.29624 | 0.1067 | 0.82131 | 0.12788 | 33.64168 | 0.18 | 0.2 | Neutral |
| testneg_4142 | 0.03 | 0.12 | 3.392117 | -3 | 1.565142 | -0.038604 | 0.2231 | 0.22973 | 0.493 | 0.35835 | 5 | 0.378779327 | 13.6 | Neutral |
| testneg_4143 | 0 | 0 | 0.250946 | -3 | 0.297691 | -3 | 0.39774 | 0.10813 | 0.07275 | 0.09562 | 12.78425 | 0.000860016 | 0 | Neutral |
| testneg_4144 | 0.01 | 0.8 | 2.2955 | -3 | -0.678462 | -3 | 0.36566 | 0.15312 | 0.67216 | 0.17808 | 10.62943 | 0.08 | 0 | Neutral |
| testneg_4145 | -0.02 | 0.68 | 1.730125 | -3 | 0.682078 | 0.504046 | 0.10671 | 0.10867 | 0.30471 | 0.11609 | 19.79537 | 0.010095238 | 0 | Neutral |
| testneg_4146 | 0.05 | 0.29 | 3.391965 | -3 | -0.418378 | -3 | 0.19898 | 0.12788 | 0.31744 | 0.10986 | 18.6768 | 0.015 | 0 | Neutral |
| testneg_4147 | 0 | 0 | 0.310149 | -3 | -0.40166 | -3 | 0.0626 | 0.5373 | 0.08683 | 0.26961 | 31.04551 | 0.276042857 | 0.2 | Neutral |

| | | | | | | | | | | | | | | |
|---|---|---|---|---|---|---|---|---|---|---|---|---|---|---|
| testneg_4148 | 0 | -0.01 | 1.616889 | -3 | 1.019099 | -3 | 0.75071 | 0.63704 | 0.19898 | 0.12788 | 13.89558 | 0.02 | 0 | Neutral |
| testneg_4149 | -0.03 | 1.03 | 2.996727 | -3 | 2.61677 | 0.587716 | 0.19898 | 0.17518 | 0.60285 | 0.15179 | 18.88889 | 0.405735608 | 23.4 | Neutral |
| testneg_4150 | 0 | -0.01 | 2.324316 | -3 | 8.521994 | -3 | 0.13767 | 0.11503 | 0.07611 | 0.11551 | 10 | 0.36 | 8 | Neutral |
| testneg_4151 | 0 | 0 | 0.196897 | -3 | 1.613942 | -3 | 0.74699 | 0.12286 | 0.34399 | 0.08193 | 35.83482 | 0.010128395 | 0 | Neutral |
| testneg_4152 | 0.14 | -0.37 | 6.149616 | -3 | -0.885978 | -3 | 0.51037 | 0.10999 | 0.14007 | 0.10647 | 29.16667 | 0.15 | 0 | Neutral |
| testneg_4153 | 0.03 | -1.03 | 2.763749 | -3 | 3.903693 | -3 | 0.13203 | 0.12092 | 0.14231 | 0.41445 | 18.6768 | 0.449573144 | 33.6 | Neutral |
| testneg_4154 | 0 | 0 | 1.626934 | -3 | 0.833216 | -3 | 0.20942 | 0.14229 | 0.20127 | 0.13478 | 8.42931 | 0.040346476 | 0 | Neutral |
| testneg_4155 | 0 | -0.01 | -0.302033 | -3 | 5.706361 | -3 | 0.19898 | 0.12788 | 0.51852 | 0.11202 | 11.99102 | 0.060152089 | 0 | Neutral |
| testneg_4156 | 0 | 0 | 1.118558 | -3 | 1.133629 | -3 | 0.06095 | 0.08179 | 0.19898 | 0.12788 | 12.91318 | 0.01 | 0 | Neutral |
| testneg_4157 | 0.06 | -0.57 | 1.64952 | -3 | 1.326868 | -3 | 0.06513 | 0.12353 | 0.40264 | 0.12788 | 45.41667 | 0.02 | 0 | Neutral |
| testneg_4158 | 0 | -0.01 | 1.149331 | -3 | 0.185804 | -3 | 0.15582 | 0.14216 | 0.08306 | 0.12788 | 13.51351 | 0.01 | 0 | Neutral |
| testneg_4159 | 0 | -0.32 | -1.178577 | -3 | 1.488553 | -3 | 0.19898 | 0.12788 | 0.35584 | 0.10874 | 18.69495 | 0.01 | 0 | Neutral |
| testneg_4160 | 0 | 0 | 1.687316 | -3 | 3.808656 | -3 | 0.26434 | 0.12788 | 0.15152 | 0.12645 | 17.21847 | 0.04 | 0 | Neutral |
| testneg_4161 | -0.21 | -0.15 | -0.569149 | -3 | 1.572604 | -3 | 0.27528 | 0.12788 | 0.21967 | 0.09467 | 18.77973 | 0.05 | 0 | Neutral |
| testneg_4162 | 0 | 0 | 0.619175 | 0.619175 | 0.780526 | -3 | 0.68138 | 0.20944 | 0.50161 | 0.12788 | 11.16015 | 0.033779762 | 0 | Neutral |
| testneg_4163 | 0 | 0 | 1.797648 | -3 | 1.315669 | -3 | 0.30275 | 0.12788 | 0.13331 | 0.09729 | 13.46631 | 0 | 0 | Neutral |
| testneg_4164 | 0.14 | -0.37 | 6.333404 | -3 | -1.335936 | -3 | 0.14477 | 0.18932 | 0.13118 | 0.08646 | 18.41686 | 0.212768199 | 0.4 | Neutral |
| testneg_4165 | 0 | 0 | 0.079919 | -3 | 2.16064 | -3 | 0.50063 | 0.154 | 0.16734 | 0.15176 | 19.44444 | 0.1144916 | 0 | Neutral |
| testneg_4166 | 0.17 | 0.38 | 2.059872 | -3 | -0.383439 | -3 | 0.605 | 0.10748 | 0.23152 | 0.09962 | 12.46106 | 0.03 | 0 | Neutral |
| testneg_4167 | -0.07 | -0.38 | 1.785837 | -3 | 1.706302 | -3 | 0.22564 | 0.12009 | 0.17639 | 0.12919 | 11.25 | 0.01 | 0 | Neutral |
| testneg_4168 | 0.21 | 0.15 | 1.098308 | -3 | 3.074697 | -3 | 0.09346 | 0.12788 | 0.37282 | 0.07943 | 21.87607 | 0.070159508 | 0 | Neutral |
| testneg_4169 | 0.07 | 0.01 | 5.328732 | -3 | 4.714948 | -3 | 0.64922 | 0.10176 | 0.20327 | 0.11174 | 30.61883 | 0.277098981 | 1.4 | Neutral |
| testneg_4170 | -0.07 | -0.7 | 2.840988 | -3 | -1.521094 | -3 | 0.1014 | 0.12788 | 0.29672 | 0.10485 | 25.10109 | 0.03 | 0 | Neutral |
| testneg_4171 | 0.07 | 0.38 | -2.0307 | -3 | -0.171668 | -3 | 0.06848 | 0.09963 | 0.45029 | 0.12788 | 27.4024 | 0.02 | 0 | Neutral |
| testneg_4172 | -0.09 | 0.67 | 0.082232 | -3 | 0.345315 | -3 | 0.08475 | 0.19889 | 0.09964 | 0.10384 | 13.02635 | 0.07 | 0 | Neutral |
| testneg_4173 | -0.17 | -0.38 | -0.92068 | -3 | 0.24062 | -3 | 0.28038 | 0.11814 | 0.18351 | 0.11213 | 28.79924 | 0.04 | 0 | Neutral |
| testneg_4174 | 0 | 0 | 1.781646 | -3 | 0.313043 | -3 | 0.17258 | 0.12788 | 0.18706 | 0.10259 | 12.50976 | 0.000198039 | 0 | Neutral |
| testneg_4175 | -0.1 | -1.9 | 3.35171 | -3 | 1.156713 | 1.156713 | 0.20112 | 0.17491 | 0.36458 | 0.16397 | 11.25 | 0.293333333 | 1.8 | Neutral |
| testneg_4176 | -0.02 | 0.63 | 2.528853 | -3 | 5.678943 | -3 | 0.39225 | 0.10625 | 0.37909 | 0.4568 | 18.33333 | 0.433358379 | 29.8 | Neutral |
| testneg_4177 | 0 | 0 | -0.104866 | -3 | 1.565943 | -3 | 0.2698 | 0.18815 | 0.17796 | 0.09522 | 18.33333 | 0.02 | 0 | Neutral |
| testneg_4178 | 0 | 0 | 0.11336 | -3 | 7.947701 | -3 | 0.26192 | 0.15452 | 0.10183 | 0.11847 | 22.44444 | 0.401361111 | 19.2 | Neutral |
| testneg_4179 | 0.05 | -0.81 | -0.765143 | -0.765143 | 0.905656 | -3 | 0.31627 | 0.23288 | 0.30585 | 0.10254 | 13.29632 | 0.0305 | 0 | Neutral |
| testneg_4180 | 0.17 | 0.38 | 2.7281 | -3 | -0.811454 | -0.811454 | 0.33163 | 0.26601 | 0.72912 | 0.18218 | 5 | 0.199006944 | 0 | Neutral |
| testneg_4181 | -0.02 | 0.63 | 5.183006 | -3 | 4.651185 | -3 | 0.04208 | 0.12788 | 0.18373 | 0.12788 | 42.5 | 0.328642587 | 6 | Neutral |
| testneg_4182 | 0 | -0.01 | -1.285414 | -3 | -0.674759 | -0.674759 | 0.0693 | 0.09293 | 0.02782 | 0.06218 | 20.83333 | 0.03 | 0 | Neutral |
| testneg_4183 | -0.23 | 1.86 | 2.603661 | -3 | -0.071864 | -3 | 0.06708 | 0.09234 | 0.97005 | 0.31407 | 4.44444 | 0.22 | 0 | Neutral |
| testneg_4184 | 0.16 | -1.37 | -0.121102 | -0.121102 | 0.612491 | -3 | 0.78935 | 0.17351 | 0.52768 | 0.12788 | 9.862 | 0.06 | 0 | Neutral |
| testneg_4185 | 0 | 0 | 0.121522 | -3 | 1.906984 | -3 | 0.25421 | 0.16235 | 0.10385 | 0.09775 | 29.16667 | 0.03 | 0 | Neutral |
| testneg_4186 | 0.16 | -0.37 | 4.456884 | -3 | 1.707083 | -3 | 0.38115 | 0.21061 | 0.90296 | 0.28521 | 18.33333 | 0.37 | 11.6 | Neutral |
| testneg_4187 | 0.14 | -0.69 | 2.950487 | -3 | -3.056434 | -3 | 0.14495 | 0.37127 | 0.17786 | 0.15314 | 40.14476 | 0.31175 | 2.2 | Neutral |
| testneg_4188 | 0.08 | 0.03 | -1.16707 | -3 | 1.861867 | 1.861867 | 0.18416 | 0.13292 | 0.32963 | 0.1035 | 6.19048 | 0.05 | 0 | Neutral |
| testneg_4189 | 0 | -0.01 | 2.227487 | -3 | 4.971354 | -3 | 0.85984 | 0.40044 | 0.17258 | 0.12788 | 9.12252 | 0.156548696 | 0 | Neutral |
| testneg_4190 | 0 | 0 | -0.471382 | -3 | 5.879594 | -3 | 0.21933 | 0.18619 | 0.19898 | 0.12788 | 12.38369 | 0.157652881 | 0 | Neutral |
| testneg_4191 | -0.16 | 1.37 | 2.581597 | 2.581597 | -0.260345 | -3 | 0.14021 | 0.43531 | 0.08499 | 0.08678 | 10.41667 | 0.3403125 | 9.2 | Neutral |
| testneg_4192 | -0.19 | 0.27 | 1.951681 | -3 | 3.537755 | -3 | 0.1875 | 0.10316 | 0.06922 | 0.07026 | 40.503 | 0.120821612 | 0 | Neutral |
| testneg_4193 | 0.01 | 0.79 | 3.426239 | -3 | 3.328368 | -3 | 0.32669 | 0.10659 | 0.14731 | 0.12788 | 18.76159 | 0.137531616 | 0 | Neutral |
| testneg_4194 | 0.02 | -0.63 | 2.466761 | -3 | 0.465329 | 0.465329 | 0.0538 | 0.23912 | 0.11131 | 0.68435 | 18.88889 | 0.338095238 | 3.4 | Neutral |
| testneg_4195 | 0 | 0 | 0.71683 | -3 | 0.186761 | -3 | 0.42583 | 0.10901 | 0.19898 | 0.12788 | 23.23037 | 0.00026905 | 0 | Neutral |
| testneg_4196 | -0.08 | -0.03 | 5.981807 | -3 | 0.835262 | -3 | 0.25827 | 0.21386 | 0.13195 | 0.57267 | 8.97898 | 0.608759615 | 85.4 | Disease_causing |
| testneg_4197 | -0.06 | -0.18 | 0.96728 | -3 | -1.094626 | -3 | 0.11114 | 0.09582 | 0.3783 | 0.08224 | 17.21847 | 0.01 | 0 | Neutral |
| testneg_4198 | 0 | 0 | 0.674234 | -3 | -1.905689 | -3 | 0.08465 | 0.1932 | 0.19898 | 0.06726 | 45 | 0.03 | 0 | Neutral |
| testneg_4199 | 0 | -0.01 | 1.886678 | -3 | 0.10287 | -3 | 0.11187 | 0.10744 | 0.15443 | 0.10913 | 13.51351 | 0 | 0 | Neutral |
| testneg_4200 | 0 | 0 | 0.275913 | -3 | 0.631528 | -3 | 0.61027 | 0.14229 | 0.11688 | 0.13208 | 22.35492 | 0.096602035 | 0 | Neutral |
| testneg_4201 | 0.04 | 2.01 | -1.062451 | -1.062451 | -0.720759 | -3 | 0.03072 | 0.27473 | 0.17758 | 0.15983 | 16.54706 | 0.31 | 1.4 | Neutral |
| testneg_4202 | 0 | -0.32 | -1.178577 | -3 | -1.859495 | -3 | 0.19898 | 0.12788 | 0.1214 | 0.0899 | 15.91585 | 0.01 | 0 | Neutral |
| testneg_4203 | -0.02 | 0.68 | 5.845448 | -3 | 1.1308 | 1.1308 | 0.06926 | 0.08399 | 0.19898 | 0.11185 | 50.67468 | 0.173611111 | 0 | Neutral |
| testneg_4204 | -0.08 | -0.03 | 3.600262 | -3 | -2.154098 | -3 | 0.22944 | 0.09039 | 0.29172 | 0.08143 | 19.09347 | 0.030161395 | 0 | Neutral |
| testneg_4205 | 0 | 0 | 0.839823 | -3 | 3.715243 | -3 | 0.26997 | 0.63944 | 0.25516 | 0.12788 | 27.3779 | 0.13 | 0 | Neutral |
| testneg_4206 | 0 | 0 | 0.159372 | 0.159372 | 3.355037 | -3 | 0.05228 | 0.12788 | 0.17443 | 0.19181 | 22.03458 | 0.26460748 | 0.2 | Neutral |
| testneg_4207 | 0.14 | -0.37 | 3.05134 | -3 | 2.43732 | -3 | 0.20692 | 0.70649 | 0.66422 | 0.99377 | 4.44444 | 0.571583333 | 81.2 | Disease_causing |
| testneg_4208 | 0 | -0.32 | -0.849297 | -0.849297 | 1.447389 | 1.447389 | 0.23691 | 0.1015 | 0.14419 | 0.1668 | 26.07351 | 0.12 | 0 | Neutral |
| testneg_4209 | 0.22 | -2.66 | 2.631407 | 2.631407 | 1.534492 | -3 | 0.44541 | 0.13979 | 0.10183 | 0.11847 | 22.98461 | 0.29 | 3.2 | Neutral |
| testneg_4210 | 0.02 | 0.13 | 4.422139 | -3 | 4.806063 | -3 | 0.04699 | 0.08149 | 0.22239 | 0.12885 | 11.25 | 0.240675748 | 1.2 | Neutral |
| testneg_4211 | 0 | 0 | -0.077498 | -3 | 3.587651 | -3 | 0.71463 | 0.29732 | 0.13615 | 0.12788 | 10.96644 | 0.09097619 | 0 | Neutral |
| testneg_4212 | 0 | 0 | 0.329118 | -3 | 1.681428 | -3 | 0.51707 | 0.45685 | 0.59691 | 0.11049 | 19.37199 | 0.03 | 0 | Neutral |
| testneg_4213 | 0.09 | -0.3 | 3.901567 | -3 | 5.255136 | -3 | 0.29044 | 0.12142 | 0.38911 | 0.12779 | 18.543 | 0.237470238 | 1.2 | Neutral |
| testneg_4214 | 0.07 | 0.7 | 3.173658 | -3 | 7.01315 | -3 | 0.95857 | 0.12137 | 0.09902 | 0.12788 | 18.33333 | 0.38 | 15 | Neutral |
| testneg_4215 | 0 | 0 | 2.050076 | -3 | -1.457001 | -3 | 0.54049 | 0.33876 | 0.25119 | 0.13626 | 9.57763 | 0.081492063 | 0 | Neutral |
| testneg_4216 | 0 | 0 | -0.061795 | -3 | -0.126067 | -3 | 0.34698 | 0.12788 | 0.19898 | 0.09171 | 22.67913 | 0.01 | 0 | Neutral |
| testneg_4217 | 0 | 0 | -0.174072 | -3 | -0.024667 | -3 | 0.97678 | 0.09885 | 0.59599 | 0.12477 | 10 | 0.01 | 0 | Neutral |
| testneg_4218 | 0.03 | 0.12 | -0.825882 | -3 | 1.308115 | -3 | 0.3221 | 0.3367 | 0.55618 | 0.11352 | 11.99492 | 0.05 | 0 | Neutral |
| testneg_4219 | 0 | -0.01 | 0.679648 | -3 | 2.169051 | -3 | 0.70964 | 0.12663 | 0.0676 | 0.12788 | 19.97695 | 0.013809524 | 0 | Neutral |
| testneg_4220 | 0.14 | -0.37 | 5.734512 | -3 | -0.470426 | -3 | 0.18498 | 0.12788 | 0.08982 | 0.15279 | 16.42913 | 0.33 | 6.6 | Neutral |
| testneg_4221 | 0.01 | -1.44 | -2.044558 | -3 | 3.439292 | -3 | 0.14511 | 0.12788 | 0.28162 | 0.12788 | 19.03093 | 0.061194296 | 0 | Neutral |
| testneg_4222 | 0 | -0.01 | 0.183879 | -3 | 0.67331 | -3 | 0.19898 | 0.12788 | 0.13535 | 0.13562 | 26.34175 | 0.06 | 0 | Neutral |
| testneg_4223 | 0 | 0 | 1.550971 | -3 | 4.479754 | -3 | 0.73311 | 0.10866 | 0.19898 | 0.12788 | 10.22898 | 0.030084631 | 0 | Neutral |
| testneg_4224 | 0 | 0 | 0.339713 | -3 | 3.567601 | -3 | 0.30739 | 0.16306 | 0.04655 | 0.10225 | 19.99092 | 0.080871212 | 0 | Neutral |
| testneg_4225 | 0.07 | 0.7 | 5.438117 | -3 | -0.020693 | -3 | 0.12463 | 0.12788 | 0.80034 | 0.10518 | 17.64787 | 0.11 | 0 | Neutral |
| testneg_4226 | 0.07 | 0.7 | 1.763064 | -3 | 2.457352 | -3 | 0.11873 | 0.10499 | 0.13974 | 0.10225 | 32.82936 | 0.03 | 0 | Neutral |
| testneg_4227 | 0.21 | 0.15 | 6.232965 | -3 | 3.200417 | -3 | 0.90811 | 0.17448 | 0.51137 | 0.19235 | 10.41667 | 0.730219877 | 100 | Disease_causing |
| testneg_4228 | 0 | 0 | -0.100514 | -3 | 0.418262 | -3 | 0.16469 | 0.16042 | 0.69007 | 0.09893 | 12.29167 | 0.020313188 | 0 | Neutral |
| testneg_4229 | -0.08 | -0.03 | 2.428041 | -3 | 0.442945 | -3 | 0.07606 | 0.13082 | 0.19499 | 0.12111 | 17.09579 | 0.02 | 0 | Neutral |
| testneg_4230 | 0 | 0 | 0.189829 | -3 | 0.009894 | -3 | 0.20219 | 0.10986 | 0.07996 | 0.08354 | 22.77196 | 0.01 | 0 | Neutral |
| testneg_4231 | 0 | 0 | 0.419264 | -3 | 2.437047 | -3 | 0.93284 | 0.71489 | 0.10395 | 0.08737 | 18.33333 | 0.07 | 0 | Neutral |
| testneg_4232 | -0.04 | 0.89 | 3.843462 | -3 | -0.187363 | -3 | 0.19898 | 0.12788 | 0.90649 | 0.3543 | 27.52646 | 0.17114565 | 0 | Neutral |
| testneg_4233 | -0.02 | 0.63 | 3.191277 | 3.191277 | 0.282797 | -3 | 0.37074 | 0.11532 | 0.20448 | 0.12788 | 21.85988 | 0.381666667 | 14.4 | Neutral |
| testneg_4234 | 0 | 0 | 1.902798 | -3 | 1.018572 | 1.018572 | 0.09259 | 0.11165 | 0.37094 | 0.10762 | 18.33333 | 0.01 | 0 | Neutral |
| testneg_4235 | 0 | -0.01 | -0.496432 | -3 | 0.372479 | -3 | 0.06247 | 0.14184 | 0.19102 | 0.12788 | 8.97898 | 0.02 | 0 | Neutral |
| testneg_4236 | 0 | 0 | 0.329118 | -3 | 4.76387 | -3 | 0.51707 | 0.45685 | 0.37109 | 0.12788 | 13.97697 | 0.112694376 | 0 | Neutral |
| testneg_4237 | -0.18 | -0.16 | -1.333487 | -3 | -0.303489 | -3 | 0.07598 | 0.08456 | 0.508 | 0.1281 | 12.32779 | 0.08 | 0 | Neutral |
| testneg_4238 | 0.04 | -0.23 | 7.386316 | -3 | 5.060627 | -3 | 0.14785 | 0.09084 | 0.11873 | 0.10499 | 32.82936 | 0.44 | 32.8 | Neutral |
| testneg_4239 | 0.03 | 0.12 | 1.294111 | -3 | -1.167722 | -3 | 0.22575 | 0.37247 | 0.84431 | 0.65315 | 10 | 0.140915532 | 0 | Neutral |
| testneg_4240 | 0.07 | 0.38 | 0.238195 | -3 | 6.216009 | -3 | 0.38749 | 0.1076 | 0.19898 | 0.12788 | 20.93537 | 0.12009434 | 0 | Neutral |
| testneg_4241 | -0.05 | 0.81 | 4.111365 | -3 | -0.154534 | -3 | 0.1462 | 0.18896 | 0.34946 | 0.19092 | 10 | 0.290092783 | 1.6 | Neutral |
| testneg_4242 | 0 | -0.01 | -0.190389 | -3 | 0.382515 | -3 | 0.19898 | 0.07875 | 0.19898 | 0.12788 | 25.00392 | 0.01 | 0 | Neutral |
| testneg_4243 | 0 | 0 | 1.002333 | -3 | 4.092079 | -3 | 0.13918 | 0.12788 | 0.21033 | 0.35953 | 15.32509 | 0.231251161 | 0.4 | Neutral |
| testneg_4244 | 0 | 0 | 0.361863 | -3 | 1.586053 | -3 | 0.04305 | 0.06354 | 0.30429 | 0.29938 | 16.8018 | 0.09 | 0 | Neutral |
| testneg_4245 | 0 | 0 | 0.694077 | -3 | 7.320532 | -3 | 0.21235 | 0.1449 | 0.19898 | 0.12788 | 18.76882 | 0.302083333 | 4 | Neutral |
| testneg_4246 | 0.07 | 0.38 | 2.92632 | -3 | 1.446803 | -3 | 0.47442 | 0.11873 | 0.143 | 0.11236 | 17.63341 | 0.03 | 0 | Neutral |
| testneg_4247 | 0 | -0.01 | 1.129155 | -3 | 3.592951 | -3 | 0.16835 | 0.09282 | 0.18122 | 0.16895 | 27.75747 | 0.188299108 | 0 | Neutral |
| testneg_4248 | 0 | 0 | 1.196382 | -3 | -0.433937 | -3 | 0.11818 | 0.10561 | 0.84572 | 0.78546 | 10.83333 | 0.054642857 | 0 | Neutral |
| testneg_4249 | -0.07 | -0.38 | 6.900185 | -3 | 4.025044 | -3 | 0.82269 | 0.15732 | 0.18512 | 0.12788 | 37.64126 | 0.589529412 | 82.6 | Disease_causing |
| testneg_4250 | -0.03 | -0.12 | -0.387026 | -3 | 7.455807 | -3 | 0.04831 | 0.12788 | 0.13961 | 0.09565 | 22.23291 | 0.33 | 5 | Neutral |
| testneg_4251 | 0.05 | 0.29 | 2.83636 | -3 | 4.95996 | -3 | 0.19002 | 0.12138 | 0.05453 | 0.12788 | 17.71032 | 0.156311872 | 0.2 | Neutral |
| testneg_4252 | 0 | 0 | 1.785947 | -3 | 1.730485 | -3 | 0.37986 | 0.14229 | 0.30275 | 0.12788 | 8.80077 | 0.000201927 | 0 | Neutral |
| testneg_4253 | 0.17 | 0.38 | 1.603588 | -3 | 1.290195 | 1.290195 | 0.10496 | 0.21491 | 0.10761 | 0.12246 | 18.88889 | 0.08055393 | 0 | Neutral |
| testneg_4254 | 0 | 0 | -0.353049 | -3 | 4.202319 | -3 | 0.9874 | 0.15588 | 0.19898 | 0.30319 | 15 | 0.270518828 | 0.6 | Neutral |
| testneg_4255 | 0.02490909 | -0.3747273 | 11.779065 | -0.076898 | 0.167824 | -3 | 0.07973 | 0.09208 | 0.11264 | 0.11163 | 32.68623 | 0.33 | 4.2 | Neutral |
| testneg_4256 | 0.04 | -0.23 | -0.448238 | -3 | 1.388133 | -3 | 0.1592 | 0.12788 | 0.35142 | 0.18303 | 27.94286 | 0.06 | 0 | Neutral |
| testneg_4257 | 0.09 | -0.25 | -0.154606 | -3 | 3.699144 | -3 | 0.12886 | 0.12788 | 0.10782 | 0.14795 | 18.88212 | 0.254512175 | 0.4 | Neutral |
| testneg_4258 | 0 | 0 | 2.044018 | -3 | 3.808656 | 2.705972 | 0.63431 | 0.09837 | 0.15152 | 0.12645 | 11.92982 | 0.14 | 0 | Neutral |
| testneg_4259 | 0.04 | -0.23 | 3.629239 | -3 | 2.439845 | -3 | 0.30042 | 0.13517 | 0.09964 | 0.10384 | 13.08587 | 0.08 | 0 | Neutral |
| testneg_4260 | 0 | 0 | 0.036223 | -3 | 5.351052 | -3 | 0.9355 | 0.40418 | 0.87877 | 0.60823 | 5 | 0.306883849 | 5 | Neutral |
| testneg_4261 | 0.02 | -0.63 | 6.188719 | -3 | 5.698111 | -3 | 0.25474 | 0.11262 | 0.22645 | 0.1704 | 17.917 | 0.789232955 | 99.8 | Disease_causing |
| testneg_4262 | -0.03 | -0.12 | 3.715248 | -3 | 3.471969 | -3 | 0.83067 | 0.10583 | 0.99859 | 0.12788 | 19.40613 | 0.159545455 | 0 | Neutral |
| testneg_4263 | 0 | -0.73 | 6.453651 | -3 | -0.445161 | -0.445161 | 0.07889 | 0.08835 | 0.09728 | 0.12788 | 36.66548 | 0.18 | 0 | Neutral |
| testneg_4264 | 0.11 | -0.19 | 2.716468 | -3 | 0.10268 | -3 | 0.28066 | 0.18241 | 0.34193 | 0.12788 | 14.85392 | 0.04 | 0 | Neutral |
| testneg_4265 | 0 | 0 | 0.726154 | -3 | 0.212443 | -3 | 0.39964 | 0.22293 | 0.07701 | 0.09936 | 18.69073 | 0.013684524 | 0 | Neutral |
| testneg_4266 | 0.14 | -0.69 | 7.253468 | -3 | 1.464574 | 1.464574 | 0.48936 | 0.10963 | 0.60603 | 0.12788 | 16.06537 | 0.31 | 4.6 | Neutral |
| testneg_4267 | -0.06 | -0.18 | 4.281441 | -3 | 6.811258 | -3 | 0.47775 | 0.10103 | 0.12269 | 0.08105 | 24.03513 | 0.35 | 8.6 | Neutral |
| testneg_4268 | -0.03 | -0.12 | 1.183797 | -3 | -0.388631 | -3 | 0.99765 | 0.95064 | 0.19227 | 0.1096 | 12.1131 | 0.11 | 0 | Neutral |
| testneg_4269 | 0 | 0 | 1.289271 | -3 | 3.715248 | -3 | 0.19898 | 0.10894 | 0.83067 | 0.10583 | 13.14653 | 0.035211039 | 0 | Neutral |

| | | | | | | | | | | | | | | |
|---|---|---|---|---|---|---|---|---|---|---|---|---|---|---|
| testneg_4270 | 0.03 | -1.03 | 4.326488 | -3 | 3.036954 | -3 | 0.10604 | 0.10785 | 0.36541 | 0.27749 | 19.09347 | 0.491869104 | 50.8 | Disease_causing |
| testneg_4271 | -0.18 | -0.16 | 1.361743 | -3 | 1.105669 | -3 | 0.19898 | 0.12788 | 0.44385 | 0.1311 | 10.41667 | 0.08 | 0 | Neutral |
| testneg_4272 | -0.05 | -0.29 | 0.483862 | -3 | 6.154021 | -3 | 0.21551 | 0.12788 | 0.13669 | 0.09263 | 21.68901 | 0.130086207 | 0 | Neutral |
| testneg_4273 | 0 | 0 | 0.356924 | -3 | 1.954724 | -3 | 0.3889 | 0.10831 | 0.49831 | 0.29427 | 37.59997 | 0.06 | 0 | Neutral |
| testneg_4274 | -0.02 | 0.63 | 0.385855 | -3 | 6.562934 | -3 | 0.10153 | 0.13343 | 0.12965 | 0.75487 | 18.6768 | 0.547678571 | 70.6 | Disease_causing |
| testneg_4275 | 0.01 | -1.3 | 3.151927 | -3 | 4.694696 | -3 | 0.19898 | 0.08029 | 0.13169 | 0.06895 | 17.21847 | 0.123431373 | 0 | Neutral |
| testneg_4276 | 0 | 0 | 0.754525 | -3 | 0.27272 | -3 | 0.80629 | 0.12788 | 0.02967 | 0.06605 | 9.21523 | 0.005053763 | 0 | Neutral |
| testneg_4277 | 0 | 0 | 0.826657 | -3 | 2.653525 | -3 | 0.10385 | 0.09775 | 0.20546 | 0.10734 | 18.6768 | 0.01 | 0 | Neutral |
| testneg_4278 | 0 | 0 | 0.094005 | -3 | 6.609854 | -3 | 0.64302 | 0.11262 | 0.18096 | 0.12788 | 29.61945 | 0.127541667 | 0 | Neutral |
| testneg_4279 | 0 | 0 | 4.136563 | -3 | 2.174777 | -3 | 0.99689 | 0.28104 | 0.32458 | 0.1016 | 13.80454 | 0.155287879 | 0 | Neutral |
| testneg_4280 | -0.03 | -0.1 | 6.245625 | -3 | 0.819832 | -3 | 0.05535 | 0.12788 | 0.19898 | 0.12788 | 54.00594 | 0.23 | 0.2 | Neutral |
| testneg_4281 | 0.07 | 0.38 | -1.623395 | -3 | 0.008926 | -3 | 0.10799 | 0.10754 | 0.12455 | 0.0998 | 13.51351 | 0.01 | 0 | Neutral |
| testneg_4282 | -0.03 | -0.1 | 4.076957 | 4.076957 | 1.641098 | -3 | 0.38324 | 0.25193 | 0.13906 | 0.09925 | 10.22898 | 0.49 | 50.8 | Disease_causing |
| testneg_4283 | 0 | 0 | 0.034609 | 0.034609 | 3.206814 | -3 | 0.42033 | 0.17821 | 0.7022 | 0.35308 | 10.41667 | 0.224022607 | 0.2 | Neutral |
| testneg_4284 | 0.07 | -0.31 | 3.062683 | -3 | 5.874971 | -3 | 0.18507 | 0.12788 | 0.16393 | 0.08897 | 41.67748 | 0.183644987 | 0 | Neutral |
| testneg_4285 | 0 | 0 | -0.505055 | -3 | 2.347933 | -3 | 0.99859 | 0.12788 | 0.43158 | 0.10472 | 4.44444 | 0.050296474 | 0 | Neutral |
| testneg_4286 | -0.11 | 0.19 | 1.182242 | -3 | 4.697701 | 4.697701 | 0.36486 | 0.14268 | 0.1415 | 0.15572 | 9.6262 | 0.536 | 69.4 | Disease_causing |
| testneg_4287 | 0 | 0 | 0.844548 | -3 | 2.418801 | -3 | 0.9237 | 0.55859 | 0.05318 | 0.12788 | 21.71558 | 0.081586538 | 0 | Neutral |
| testneg_4288 | -0.15 | 0.28 | -2.193108 | -3 | 4.889881 | -3 | 0.06996 | 0.35923 | 0.31434 | 0.11315 | 18.33333 | 0.36 | 5.4 | Neutral |
| testneg_4289 | 0.01 | 0.8 | -0.214756 | -3 | 5.829822 | -3 | 0.07428 | 0.07645 | 0.06895 | 0.06354 | 24.85825 | 0.121814286 | 0 | Neutral |
| testneg_4290 | 0.03 | 0.1 | 2.481102 | -3 | -0.183041 | -3 | 0.25804 | 0.11876 | 0.09323 | 0.16732 | 50.65476 | 0.123333333 | 0 | Neutral |
| testneg_4291 | 0.2 | -1.26 | 3.078755 | 1.112437 | -0.816744 | -3 | 0.62172 | 0.18135 | 0.05087 | 0.09682 | 12.29167 | 0.159583333 | 0 | Neutral |
| testneg_4292 | 0.06 | -0.57 | 1.87579 | -3 | 0.690746 | -3 | 0.16965 | 0.25991 | 0.19898 | 0.12788 | 20.86969 | 0.031 | 0 | Neutral |
| testneg_4293 | 0 | 0 | 1.107432 | -3 | 0.290985 | -3 | 0.5726 | 0.1165 | 0.20986 | 0.10633 | 17.94606 | 0.000465368 | 0 | Neutral |
| testneg_4294 | 0 | 0 | 0.371766 | 0.371766 | 0.188403 | -3 | 0.3095 | 0.13502 | 0.39111 | 0.15887 | 13.27141 | 0.070228639 | 0 | Neutral |
| testneg_4295 | 0.05 | -0.81 | 6.066721 | -3 | 1.01961 | -3 | 0.15041 | 0.43468 | 0.53016 | 0.10373 | 18.33333 | 0.2905 | 2 | Neutral |
| testneg_4296 | 0 | 0 | 2.491045 | -3 | -0.406057 | -0.406057 | 0.55193 | 0.22384 | 0.7413 | 0.18974 | 10.41667 | 0.070259136 | 0 | Neutral |
| testneg_4297 | 0 | -0.01 | 2.663664 | -3 | 0.859718 | -3 | 0.19898 | 0.26604 | 0.37736 | 0.11686 | 31.61232 | 0.030477273 | 0 | Neutral |
| testneg_4298 | 0.00818182 | -0.38 | 5.643105 | -1.684975 | 5.399087 | -3 | 0.08775 | 0.12788 | 0.1594 | 0.22002 | 23.68839 | 0.720049127 | 98.6 | Disease_causing |
| testneg_4299 | 0 | 0 | -0.181123 | -3 | 2.009531 | -3 | 0.12867 | 0.11159 | 0.47322 | 0.11118 | 13.60057 | 8.07E-05 | 0 | Neutral |
| testneg_4300 | 0.03 | 0.12 | 0.199359 | 0.199359 | 5.734507 | -3 | 0.07626 | 0.12788 | 0.06731 | 0.2149 | 29.45604 | 0.388481191 | 15 | Neutral |
| testneg_4301 | 0 | -0.32 | 0.984662 | -3 | 0.829195 | -3 | 0.32022 | 0.09529 | 0.68423 | 0.08163 | 16.73378 | 0.01 | 0 | Neutral |
| testneg_4302 | 0 | -0.32 | 6.416028 | -3 | 1.885547 | -3 | 0.09306 | 0.09084 | 0.15992 | 0.12788 | 13.23087 | 0.221827957 | 1.2 | Neutral |
| testneg_4303 | -0.07 | -0.33 | -2.373259 | -2.373259 | 1.53664 | -3 | 0.04525 | 0.12788 | 0.17576 | 0.12193 | 19.09347 | 0.050432417 | 0 | Neutral |
| testneg_4304 | -0.03 | -0.1 | 2.93409 | -3 | -2.609079 | -3 | 0.09531 | 0.11449 | 0.19898 | 0.09533 | 27.3569 | 0.04 | 0 | Neutral |
| testneg_4305 | 0.14 | -1.01 | 0.341374 | -3 | -0.825189 | -0.940094 | 0.10646 | 0.1069 | 0.03646 | 0.09863 | 20.00508 | 0.02 | 0 | Neutral |
| testneg_4306 | 0 | 0 | 1.164364 | -3 | 0.359737 | -3 | 0.19898 | 0.12788 | 0.21355 | 0.12788 | 14.2272 | 0 | 0 | Neutral |
| testneg_4307 | 0.04 | -0.23 | 4.599131 | -3 | 8.23651 | -3 | 0.21848 | 0.11049 | 0.07071 | 0.12788 | 18.88889 | 0.53 | 65.6 | Disease_causing |
| testneg_4308 | 0 | -0.01 | 2.539007 | -3 | 1.832674 | -3 | 0.47751 | 0.2062 | 0.1589 | 0.10761 | 11.93907 | 0.030116673 | 0 | Neutral |
| testneg_4309 | 0 | 0 | 1.599689 | -3 | 0.601183 | -3 | 0.17057 | 0.12788 | 0.17639 | 0.12919 | 13.00459 | 0 | 0 | Neutral |
| testneg_4310 | 0 | 0 | 1.441342 | -0.3269 | 4.176231 | -3 | 0.11072 | 0.38259 | 0.55315 | 0.09195 | 16.8018 | 0.1721 | 0 | Neutral |
| testneg_4311 | 0.22 | -2.66 | 5.950905 | -3 | -0.929745 | -3 | 0.19898 | 0.12788 | 0.09284 | 0.26053 | 5.55556 | 0.44 | 32.8 | Neutral |
| testneg_4312 | -0.02 | -0.45 | 1.35797 | -3 | 5.543296 | -3 | 0.51994 | 0.37283 | 0.09642 | 0.09812 | 20 | 0.201125823 | 0 | Neutral |
| testneg_4313 | 0.22 | -2.66 | 5.564432 | -3 | 0.442945 | -3 | 0.32897 | 0.10658 | 0.19499 | 0.12111 | 22.32495 | 0.121805556 | 0 | Neutral |
| testneg_4314 | 0.15 | -0.28 | -0.703388 | -3 | 1.720879 | -3 | 0.44526 | 0.09404 | 0.17853 | 0.32085 | 19.28706 | 0.100834807 | 0 | Neutral |
| testneg_4315 | 0.07 | 0.38 | 0.035747 | -3 | 2.944639 | -3 | 0.38873 | 0.11686 | 0.47335 | 0.12788 | 11.25 | 0.048606918 | 0 | Neutral |
| testneg_4316 | -0.07 | -0.38 | 0.610011 | -3 | 4.361375 | -3 | 0.52278 | 0.41207 | 0.10459 | 0.12783 | 17.5501 | 0.20269697 | 0 | Neutral |
| testneg_4317 | 0.03 | -1.03 | 3.488962 | -3 | 1.464574 | 1.464574 | 0.59245 | 0.12788 | 0.60603 | 0.12788 | 13.40128 | 0.07 | 0 | Neutral |
| testneg_4318 | -0.02 | -0.45 | -0.208207 | -3 | -1.367472 | -3 | 0.07219 | 0.12788 | 0.049 | 0.11324 | 43.93658 | 0.02 | 0 | Neutral |
| testneg_4319 | 0 | -0.01 | 0.627226 | -3 | 4.773876 | -3 | 0.29004 | 0.45465 | 0.25453 | 0.1177 | 13.50732 | 0.097971429 | 0 | Neutral |
| testneg_4320 | 0.2 | -1.26 | 2.143533 | -3 | 7.168928 | -3 | 0.40897 | 0.13253 | 0.21095 | 0.10574 | 23.48109 | 0.32 | 4.6 | Neutral |
| testneg_4321 | -0.06 | 0.57 | -1.48217 | -3 | 1.651093 | 1.651093 | 0.1621 | 0.1641 | 0.08332 | 0.09046 | 20.83333 | 0.070713349 | 0 | Neutral |
| testneg_4322 | 0.05 | -0.81 | 6.767868 | -3 | 5.957971 | -3 | 0.50224 | 0.4059 | 0.20701 | 0.10345 | 15.09363 | 0.731583333 | 99.6 | Disease_causing |
| testneg_4323 | 0 | 0 | 0.432513 | -3 | 0.729199 | -3 | 0.99595 | 0.23021 | 0.11373 | 0.10739 | 10.23129 | 0.03 | 0 | Neutral |
| testneg_4324 | 0 | 0 | -0.415284 | -3 | 0.431201 | 0.431201 | 0.11301 | 0.10716 | 0.11019 | 0.08132 | 16.88725 | 0.01 | 0 | Neutral |
| testneg_4325 | 0 | 0 | 1.584574 | 1.584574 | -0.123577 | -3 | 0.29362 | 0.22177 | 0.19668 | 0.16042 | 10.87198 | 0.145185185 | 0 | Neutral |
| testneg_4326 | 0 | 0 | -0.262879 | -0.262879 | 7.433401 | -3 | 0.61011 | 0.09584 | 0.7219 | 0.41509 | 20.08636 | 0.339125 | 9.8 | Neutral |
| testneg_4327 | 0.04 | -0.23 | 1.879054 | -3 | 3.446417 | 3.446417 | 0.505 | 0.12788 | 0.29013 | 0.11368 | 21.51158 | 0.2 | 0 | Neutral |
| testneg_4328 | 0.18 | -1.05 | 2.597662 | -3 | 0.944979 | -3 | 0.06027 | 0.11574 | 0.1856 | 0.12788 | 7.01832 | 0.04 | 0 | Neutral |
| testneg_4329 | 0.04 | -0.23 | 2.563054 | -3 | 3.557638 | -3 | 0.81899 | 0.12788 | 0.15867 | 0.09641 | 17.21847 | 0.064476469 | 0 | Neutral |
| testneg_4330 | -0.1 | -1.9 | 2.306738 | -3 | 5.898121 | -3 | 0.16091 | 0.11571 | 0.1296 | 0.164 | 29.65357 | 0.463042335 | 40.2 | Neutral |
| testneg_4331 | 0.02 | 0.45 | 1.791247 | -3 | 1.594672 | -3 | 0.39193 | 0.11262 | 0.3298 | 0.21875 | 12.81947 | 0.039313492 | 0 | Neutral |
| testneg_4332 | 0 | 0 | -0.032453 | -3 | 2.290803 | -3 | 0.31664 | 0.14181 | 0.02329 | 0.07615 | 27.95796 | 0.037202381 | 0 | Neutral |
| testneg_4333 | -0.07 | -0.38 | 1.953114 | -3 | 0.256741 | -3 | 0.96879 | 0.38765 | 0.3012 | 0.10317 | 10.73171 | 0.039083333 | 0 | Neutral |
| testneg_4334 | 0.15 | -0.28 | 0.937469 | -3 | 4.454666 | -3 | 0.58677 | 0.11579 | 0.09462 | 0.11807 | 28.24388 | 0.1 | 0 | Neutral |
| testneg_4335 | 0.06 | -0.57 | 0.297516 | 0.297516 | 1.416458 | -3 | 0.19294 | 0.10464 | 0.10738 | 0.0974 | 32.67285 | 0.02 | 0 | Neutral |
| testneg_4336 | 0.12 | -1.16 | 1.957496 | -3 | 1.474779 | 1.474779 | 0.06395 | 0.0759 | 0.0882 | 0.12788 | 23.74677 | 0.05 | 0 | Neutral |
| testneg_4337 | -0.02 | 0.63 | 2.174684 | -3 | -0.526282 | -3 | 0.49246 | 0.11033 | 0.16663 | 0.12788 | 20.83333 | 0.03 | 0 | Neutral |
| testneg_4338 | 0.14 | -1.01 | 3.139463 | -3 | 4.116041 | -3 | 0.1228 | 0.10482 | 0.68423 | 0.08163 | 13.40128 | 0.159871795 | 0 | Neutral |
| testneg_4339 | -0.01 | 0.57 | 1.661207 | -3 | 6.368332 | -3 | 0.13555 | 0.16663 | 0.09275 | 0.14911 | 4.44444 | 0.577836371 | 83.2 | Disease_causing |
| testneg_4340 | 0.14 | -0.69 | 1.423539 | -3 | -0.867132 | -3 | 0.19898 | 0.12788 | 0.08353 | 0.21479 | 10 | 0.068540545 | 0 | Neutral |
| testneg_4341 | 0 | -0.01 | 2.171434 | -3 | 0.527682 | -3 | 0.49425 | 0.25538 | 0.10803 | 0.14652 | 10.41667 | 0.089948797 | 0 | Neutral |
| testneg_4342 | 0.14 | -0.37 | 1.075133 | -3 | -0.142939 | -3 | 0.0606 | 0.39625 | 0.11254 | 0.19973 | 20.13833 | 0.31 | 1.2 | Neutral |
| testneg_4343 | -0.07 | -0.38 | 0.816148 | -3 | 0.452231 | -3 | 0.26347 | 0.16383 | 0.46691 | 0.07886 | 22.77708 | 0.03 | 0 | Neutral |
| testneg_4344 | 0.01 | 0.8 | 5.808678 | -3 | 5.679638 | -3 | 0.11584 | 0.19704 | 0.61834 | 0.10784 | 20.66855 | 0.585570666 | 84 | Disease_causing |
| testneg_4345 | -0.08 | -0.03 | 0.630495 | 0.630495 | 0.842664 | -3 | 0.14419 | 0.1668 | 0.47187 | 0.14227 | 18.13683 | 0.15 | 0 | Neutral |
| testneg_4346 | 0 | 0 | 1.600017 | -3 | -0.0507 | -3 | 0.07994 | 0.1101 | 0.02967 | 0.06605 | 27.31935 | 0.01 | 0 | Neutral |
| testneg_4347 | 0.09 | -0.3 | -0.706782 | -3 | 1.031442 | -3 | 0.18047 | 0.2356 | 0.2761 | 0.24559 | 18.33333 | 0.151352969 | 0 | Neutral |
| testneg_4348 | 0.04 | -0.23 | 1.5079 | -3 | 0.082365 | -3 | 0.34694 | 0.20397 | 0.10011 | 0.08014 | 12.34649 | 0.021356112 | 0 | Neutral |
| testneg_4349 | -0.0190909 | 0.62818182 | 7.974744 | -3 | -0.211546 | -3 | 0.087 | 0.12788 | 0.2442 | 0.11401 | 22.89111 | 0.34 | 6.4 | Neutral |
| testneg_4350 | -0.09 | 0.3 | 0.216563 | -3 | 0.367308 | 0.367308 | 0.90296 | 0.28521 | 0.20047 | 0.11827 | 18.6791 | 0.041715686 | 0 | Neutral |
| testneg_4351 | -0.01 | -0.79 | 2.489008 | -3 | 0.772419 | -3 | 0.09416 | 0.17821 | 0.83936 | 0.42313 | 21.49912 | 0.195887446 | 0 | Neutral |
| testneg_4352 | -0.02 | 0.36 | 2.804789 | -3 | 0.452441 | 0.452441 | 0.53313 | 0.14072 | 0.48437 | 0.09986 | 21.2963 | 0.042426471 | 0 | Neutral |
| testneg_4353 | 0.03 | 0.12 | 4.738569 | -3 | 5.378158 | -3 | 0.04161 | 0.12788 | 0.4334 | 0.09454 | 19.06745 | 0.26 | 1.2 | Neutral |
| testneg_4354 | -0.04 | 0.23 | 1.474779 | 1.474779 | -0.614161 | -3 | 0.0882 | 0.12788 | 0.17796 | 0.09522 | 20 | 0.04 | 0 | Neutral |
| testneg_4355 | -0.06 | -0.18 | 4.235243 | -3 | 4.075251 | -3 | 0.14806 | 0.09574 | 0.13001 | 0.13176 | 31.25 | 0.323464696 | 4.8 | Neutral |
| testneg_4356 | 0 | -0.01 | 0.545018 | -3 | 0.096517 | -3 | 0.11818 | 0.10561 | 0.04946 | 0.12788 | 26.48342 | 0.01 | 0 | Neutral |
| testneg_4357 | 0.08 | -1.2 | -1.685253 | -3 | 5.011586 | -3 | 0.16091 | 0.90971 | 0.75008 | 0.70645 | 12.29167 | 0.571158897 | 84.2 | Disease_causing |
| testneg_4358 | 0.2 | -1.26 | 3.801929 | -3 | 1.192651 | -3 | 0.16974 | 0.80184 | 0.65952 | 0.49708 | 10 | 0.402995238 | 18.4 | Neutral |
| testneg_4359 | -0.05 | -1.52 | 1.347937 | -3 | -0.410022 | -3 | 0.9827 | 0.16837 | 0.17419 | 0.33842 | 10.81604 | 0.14 | 0 | Neutral |
| testneg_4360 | 0 | 0 | 1.376454 | -3 | -1.304747 | -2.106364 | 0.1679 | 0.24232 | 0.10383 | 0.08853 | 19.92008 | 0.021666667 | 0 | Neutral |
| testneg_4361 | 0 | 0 | 0.27864 | -3 | 2.28293 | -3 | 0.17737 | 0.54971 | 0.08332 | 0.09046 | 6.95654 | 0.040769231 | 0 | Neutral |
| testneg_4362 | -0.17 | -0.38 | -1.129477 | -3 | 3.564366 | -3 | 0.11006 | 0.14352 | 0.59864 | 0.14106 | 19.96197 | 0.329736842 | 4 | Neutral |
| testneg_4363 | 0 | 0 | 3.345748 | -3 | 1.499766 | -3 | 0.12786 | 0.72066 | 0.08836 | 0.0911 | 19.45437 | 0.135 | 0 | Neutral |
| testneg_4364 | 0.21 | 0.15 | 3.070963 | -3 | -0.826901 | -1.178166 | 0.11791 | 0.39607 | 0.21289 | 0.1079 | 25.62665 | 0.13 | 0 | Neutral |
| testneg_4365 | -0.02 | 0.68 | 3.51321 | -3 | 1.411328 | -3 | 0.07887 | 0.07487 | 0.13143 | 0.90853 | 28.52371 | 0.21 | 0 | Neutral |
| testneg_4366 | 0.08 | 0.03 | 1.355095 | -3 | 0.468445 | -3 | 0.28907 | 0.12788 | 0.37635 | 0.16791 | 13.3188 | 0.05 | 0 | Neutral |
| testneg_4367 | -0.14 | 1.01 | 4.888363 | -3 | 2.835051 | -3 | 0.87268 | 0.21837 | 0.20502 | 0.12788 | 18.88889 | 0.337610579 | 6.6 | Neutral |
| testneg_4368 | 0.03 | -1.03 | 7.475712 | -3 | -1.942007 | -3 | 0.26971 | 0.08509 | 0.064 | 0.24721 | 55.97902 | 0.4 | 15.4 | Neutral |
| testneg_4369 | 0.07 | 0.7 | 5.466543 | -3 | 1.497642 | -3 | 0.68729 | 0.10641 | 0.89959 | 0.55078 | 15.04363 | 0.308161765 | 3 | Neutral |
| testneg_4370 | 0 | 0 | -0.397788 | -3 | 8.064683 | -3 | 0.15911 | 0.11022 | 0.227 | 0.14389 | 29.33031 | 0.386041667 | 14.8 | Neutral |
| testneg_4371 | 0 | 0 | 1.897767 | 1.897767 | 2.293151 | -3 | 0.38911 | 0.12779 | 0.47997 | 0.10506 | 10.41667 | 0.07 | 0 | Neutral |
| testneg_4372 | 0.03 | 1.22 | 2.753966 | 2.753966 | 0.571926 | -3 | 0.1551 | 0.09276 | 0.15406 | 0.1454 | 30.85006 | 0.42 | 29.2 | Neutral |
| testneg_4373 | 0 | 0 | 1.649381 | -3 | 2.355503 | -3 | 0.22686 | 0.24974 | 0.6513 | 0.15254 | 19.44444 | 0.16424359 | 0 | Neutral |
| testneg_4374 | 0 | 0 | 1.74381 | -3 | 1.809571 | -3 | 0.67644 | 0.11323 | 0.25151 | 0.12788 | 20.97303 | 0.01 | 0 | Neutral |
| testneg_4375 | 0 | 0 | -0.513734 | -3 | -1.516438 | -3 | 0.256 | 0.20036 | 0.04046 | 0.06599 | 14.16667 | 0.02 | 0 | Neutral |
| testneg_4376 | -0.02 | 0.36 | 2.054266 | -3 | 0.2182 | -3 | 0.16777 | 0.1749 | 0.19898 | 0.12788 | 35.71709 | 0.03 | 0 | Neutral |
| testneg_4377 | 0 | -0.01 | 1.609618 | -3 | 6.36911 | -3 | 0.67038 | 0.1794 | 0.17412 | 0.10595 | 12.71494 | 0.173624121 | 0.2 | Neutral |
| testneg_4378 | 0.14 | -0.37 | 1.599249 | -3 | 2.756541 | -3 | 0.08714 | 0.08945 | 0.25236 | 0.11809 | 14.27748 | 0.030158107 | 0 | Neutral |
| testneg_4379 | 0.01 | 0.79 | 5.981726 | -3 | 3.474152 | 3.474152 | 0.28156 | 0.11579 | 0.12642 | 0.11218 | 16.0119 | 0.446031891 | 32.4 | Neutral |
| testneg_4380 | 0 | -0.01 | 7.512937 | 7.512937 | 1.56241 | -3 | 0.56315 | 0.13761 | 0.08386 | 0.12788 | 10.41667 | 0.742666667 | 99.6 | Disease_causing |
| testneg_4381 | 0.05 | -0.81 | 1.982936 | -3 | 0.79796 | -3 | 0.07044 | 0.11679 | 0.57466 | 0.12788 | 10.41667 | 0.01 | 0 | Neutral |
| testneg_4382 | 0 | 0 | 0.651919 | -3 | 1.019702 | -3 | 0.15025 | 0.4872 | 0.98238 | 0.10873 | 13.84134 | 0.04 | 0 | Neutral |
| testneg_4383 | 0.14 | -0.37 | 6.930926 | -3 | 0.82819 | 0.82819 | 0.41743 | 0.78545 | 0.19898 | 0.12788 | 10 | 0.42 | 24.4 | Neutral |
| testneg_4384 | 0 | 0 | 1.252726 | -3 | 0.162648 | -3 | 0.54231 | 0.15254 | 0.38002 | 0.22508 | 18.28933 | 0.050475718 | 0 | Neutral |
| testneg_4385 | 0 | 0 | 1.144493 | -3 | 2.76764 | -3 | 0.13794 | 0.40936 | 0.22795 | 0.12788 | 54.9243 | 0.110690476 | 0 | Neutral |
| testneg_4386 | 0 | -0.32 | 0.457146 | -3 | 0.202863 | -3 | 0.30951 | 0.12788 | 0.47451 | 0.3285 | 11.02713 | 0.04215091 | 0 | Neutral |
| testneg_4387 | 0 | 0 | 1.333395 | -3 | 2.243237 | -3 | 0.12259 | 0.1041 | 0.43977 | 0.17284 | 10.2717 | 0.04 | 0 | Neutral |
| testneg_4388 | -0.07 | -0.7 | 2.924343 | -1.464692 | 1.266295 | -3 | 0.49629 | 0.11989 | 0.32022 | 0.40347 | 19.11765 | 0.175447904 | 0 | Neutral |
| testneg_4389 | -0.1 | -1.9 | 1.840916 | -3 | 0.196354 | -3 | 0.17226 | 0.13001 | 0.18067 | 0.14759 | 12.08378 | 0.09 | 0 | Neutral |
| testneg_4390 | 0 | -0.32 | 1.05419 | -3 | 7.568352 | -3 | 0.45152 | 0.34457 | 0.14021 | 0.43531 | 18.33333 | 0.735875125 | 99.8 | Disease_causing |
| testneg_4391 | 0.03 | -1.03 | 5.25972 | -3 | 2.767554 | -3 | 0.01409 | 0.11307 | 0.0435 | 0.14527 | 11.25 | 0.3625 | 11 | Neutral |

| | | | | | | | | | | | | | | |
|---|---|---|---|---|---|---|---|---|---|---|---|---|---|---|
| testneg_4392 | 0.21 | 0.15 | 1.631249 | -3 | 0.852085 | -3 | 0.47166 | 0.13548 | 0.56554 | 0.76978 | 20.27778 | 0.15 | 0 | Neutral |
| testneg_4393 | 0 | 0 | 1.895788 | -3 | -0.230595 | -3 | 0.04863 | 0.12149 | 0.31817 | 0.17348 | 30.16667 | 0.114940476 | 0 | Neutral |
| testneg_4394 | -0.07 | -0.01 | -1.696655 | -3 | 5.391933 | -3 | 0.03526 | 0.06855 | 0.05716 | 0.12491 | 53.89135 | 0.141236111 | 0 | Neutral |
| testneg_4395 | -0.06 | -0.18 | 2.786985 | -3 | 2.264613 | -3 | 0.23292 | 0.15913 | 0.08408 | 0.12788 | 18.88889 | 0.068357143 | 0 | Neutral |
| testneg_4396 | 0.22 | -2.66 | 4.82427 | -3 | 3.165611 | 2.941559 | 0.09323 | 0.16732 | 0.30474 | 0.12788 | 44.89949 | 0.36 | 7.8 | Neutral |
| testneg_4397 | -0.1 | -1.9 | 2.551866 | -3 | 2.39641 | -3 | 0.32677 | 0.12788 | 0.25059 | 0.42498 | 11.875 | 0.23 | 0 | Neutral |
| testneg_4398 | 0 | -0.01 | 0.328189 | -3 | 1.011117 | -3 | 0.33957 | 0.12788 | 0.15524 | 0.10607 | 21.16317 | 0.01 | 0 | Neutral |
| testneg_4399 | 0.09 | -0.25 | -0.159404 | -3 | 0.523037 | -3 | 0.95319 | 0.20297 | 0.0608 | 0.09168 | 10.41667 | 0.05 | 0 | Neutral |
| testneg_4400 | 0 | 0 | 1.183859 | -3 | 0.47877 | -3 | 0.72501 | 0.7625 | 0.56204 | 0.10762 | 10.41667 | 0.03 | 0 | Neutral |
| testneg_4401 | 0.09 | -0.3 | 6.541257 | -3 | 1.864851 | -3 | 0.99531 | 0.86203 | 0.0773 | 0.14062 | 5.074 | 0.56625 | 79.8 | Disease_causing |
| testneg_4402 | 0.14 | -0.37 | 1.075133 | 1.075133 | 0.751387 | -3 | 0.0606 | 0.39625 | 0.19898 | 0.535 | 10.64018 | 0.330570487 | 3.4 | Neutral |
| testneg_4403 | 0.01 | 0.79 | 5.588206 | -0.001756 | 3.827062 | -3 | 0.16662 | 0.11306 | 0.73985 | 0.13844 | 26.53465 | 0.416571429 | 24.4 | Neutral |
| testneg_4404 | 0.17 | 0.38 | 5.080349 | -3 | 0.052319 | -3 | 0.20219 | 0.10986 | 0.15677 | 0.18537 | 13.35742 | 0.230872409 | 0.4 | Neutral |
| testneg_4405 | -0.05 | -0.29 | -2.985743 | -3 | 5.596731 | -3 | 0.55367 | 0.57993 | 0.19898 | 0.12788 | 11.875 | 0.23 | 0 | Neutral |
| testneg_4406 | 0.02 | 0.41 | 2.511378 | -3 | -1.543442 | -3 | 0.05886 | 0.38242 | 0.02759 | 0.12458 | 9.21981 | 0.1 | 0 | Neutral |
| testneg_4407 | 0 | 0 | -0.100514 | -3 | 3.390226 | -3 | 0.16469 | 0.16042 | 0.40403 | 0.12788 | 17.94606 | 0.080909091 | 0 | Neutral |
| testneg_4408 | 0 | 0 | 1.362887 | -3 | -0.55255 | -3 | 0.38527 | 0.24812 | 0.06793 | 0.12788 | 13.60998 | 0.01 | 0 | Neutral |
| testneg_4409 | 0 | 0 | -0.4932 | -3 | 0.867971 | -3 | 0.20379 | 0.43705 | 0.24966 | 0.12788 | 17.11678 | 0.02 | 0 | Neutral |
| testneg_4410 | 0.07 | 0.01 | 3.022242 | -3 | -2.160669 | -3 | 0.53627 | 0.12788 | 0.12051 | 0.08503 | 20.96973 | 0.04 | 0 | Neutral |
| testneg_4411 | 0 | -0.01 | 4.857685 | -3 | 1.528609 | -3 | 0.08714 | 0.08945 | 0.20249 | 0.10327 | 21.62577 | 0.06 | 0 | Neutral |
| testneg_4412 | -0.14 | 0.37 | -0.694905 | -3 | 0.371652 | -3 | 0.4132 | 0.12788 | 0.05087 | 0.09682 | 34.73889 | 0.04 | 0 | Neutral |
| testneg_4413 | 0 | 0 | 0.279445 | -3 | 4.895649 | -3 | 0.25179 | 0.09355 | 0.65952 | 0.49708 | 18.34652 | 0.208455914 | 0 | Neutral |
| testneg_4414 | 0.07 | -0.31 | 2.496725 | -3 | 1.951681 | -3 | 0.92237 | 0.11252 | 0.1875 | 0.10316 | 21.38889 | 0.020384058 | 0 | Neutral |
| testneg_4415 | 0.02 | 0.41 | 1.724361 | -3 | 5.283415 | -3 | 0.08454 | 0.22908 | 0.20513 | 0.09801 | 12.43093 | 0.23 | 0.2 | Neutral |
| testneg_4416 | 0 | 0 | 1.635258 | -3 | 0.390383 | -3 | 0.07701 | 0.19824 | 0.08213 | 0.08913 | 19.09347 | 0.02125 | 0 | Neutral |
| testneg_4417 | 0 | 0 | 0.029606 | -3 | -0.009242 | -3 | 0.14635 | 0.13588 | 0.11945 | 0.10135 | 24.33469 | 0.015 | 0 | Neutral |
| testneg_4418 | 0.07 | 0.33 | 3.81823 | -3 | 2.257698 | -3 | 0.1755 | 0.10455 | 0.30869 | 0.11948 | 23.09717 | 0.05 | 0 | Neutral |
| testneg_4419 | 0.02 | 0.45 | -0.784242 | -3 | 2.569635 | -3 | 0.18153 | 0.10845 | 0.3962 | 0.10523 | 12.77593 | 0.03 | 0 | Neutral |
| testneg_4420 | 0.03 | 1.22 | 2.219737 | -3 | 3.066776 | -3 | 0.72124 | 0.11262 | 0.2877 | 0.18291 | 11.875 | 0.226327163 | 0 | Neutral |
| testneg_4421 | 0.21 | 0.15 | 4.118971 | -3 | 0.05475 | -3 | 0.126 | 0.14076 | 0.13468 | 0.09235 | 41.03271 | 0.06 | 0 | Neutral |
| testneg_4422 | -0.02 | 0.68 | 0.349669 | -3 | 4.89523 | -3 | 0.19898 | 0.12788 | 0.12855 | 0.11656 | 11.30424 | 0.10022807 | 0 | Neutral |
| testneg_4423 | 0.21 | 0.15 | 3.072922 | -3 | -0.323717 | -3 | 0.17301 | 0.14089 | 0.23691 | 0.1015 | 25 | 0.05 | 0 | Neutral |
| testneg_4424 | 0 | 0 | -0.480488 | -3 | 3.063227 | -3 | 0.11797 | 0.12788 | 0.05153 | 0.12788 | 18.88889 | 0.04 | 0 | Neutral |
| testneg_4425 | -0.03 | -0.1 | 0.483877 | -0.330967 | 0.921828 | -3 | 0.20657 | 0.12788 | 0.31646 | 0.09844 | 10.41667 | 0.01 | 0 | Neutral |
| testneg_4426 | 0.04 | -0.23 | 6.936214 | -3 | 0.766889 | -3 | 0.43918 | 0.46357 | 0.13356 | 0.12788 | 13.43868 | 0.406321429 | 20 | Neutral |
| testneg_4427 | -0.08 | -0.03 | 1.629083 | -3 | 2.322079 | -3 | 0.25764 | 0.53109 | 0.93246 | 0.12788 | 10.41667 | 0.090402576 | 0 | Neutral |
| testneg_4428 | 0.21 | 0.15 | 8.17835 | -3 | -2.373259 | -3 | 0.19948 | 0.12788 | 0.04525 | 0.12788 | 18.30413 | 0.31009434 | 3.8 | Neutral |
| testneg_4429 | 0 | 0 | 0.185804 | -3 | -0.389142 | -3 | 0.08306 | 0.12788 | 0.82148 | 0.14847 | 14.97692 | 0.022736894 | 0 | Neutral |
| testneg_4430 | 0 | 0 | 0.664346 | 0.664346 | 5.390365 | 5.390365 | 0.22169 | 0.11262 | 0.09749 | 0.82924 | 36.75849 | 0.565595238 | 79 | Disease_causing |
| testneg_4431 | 0.21 | 0.15 | 3.945475 | -3 | 0.974808 | 0.974808 | 0.27115 | 0.14766 | 0.32992 | 0.33227 | 17.99708 | 0.317313984 | 2.2 | Neutral |
| testneg_4432 | 0.22 | -2.66 | 5.950905 | -3 | 5.464083 | -3 | 0.19898 | 0.12788 | 0.19898 | 0.12788 | 18.61111 | 0.443858036 | 36.4 | Neutral |
| testneg_4433 | 0 | -0.01 | 2.002215 | -3 | 1.630555 | -3 | 0.95682 | 0.22744 | 0.17526 | 0.12632 | 16.2412 | 0.018976939 | 0 | Neutral |
| testneg_4434 | 0 | -0.01 | 4.373107 | -3 | 0.693592 | -3 | 0.17173 | 0.15363 | 0.14378 | 0.41915 | 19.64993 | 0.272452943 | 0.8 | Neutral |
| testneg_4435 | -0.03 | -0.1 | 0.619249 | -3 | -0.745568 | -3 | 0.21236 | 0.12788 | 0.1475 | 0.4323 | 18.88889 | 0.077733197 | 0 | Neutral |
| testneg_4436 | 0 | 0 | 2.323551 | -3 | 2.070789 | -3 | 0.76693 | 0.19444 | 0.91139 | 0.65138 | 11.0082 | 0.105695865 | 0 | Neutral |
| testneg_4437 | 0.02 | -0.63 | 1.133629 | -3 | 0.068383 | -3 | 0.19898 | 0.12788 | 0.15822 | 0.12788 | 17.75599 | 0.000229915 | 0 | Neutral |
| testneg_4438 | -0.07 | -0.38 | 0.610011 | -3 | 5.999687 | -3 | 0.52278 | 0.41207 | 0.46242 | 0.39055 | 18.33333 | 0.617809982 | 91.2 | Disease_causing |
| testneg_4439 | 0 | 0 | 0.068421 | -3 | 4.039717 | -3 | 0.15016 | 0.08826 | 0.20546 | 0.10734 | 29.9024 | 0.04 | 0 | Neutral |
| testneg_4440 | 0.08 | 0.03 | -1.351027 | -3 | -1.924352 | -3 | 0.28156 | 0.13431 | 0.02396 | 0.12703 | 4.72222 | 0.038055556 | 0 | Neutral |
| testneg_4441 | -0.02 | 0.63 | 5.119073 | -3 | 3.233126 | -3 | 0.29679 | 0.12788 | 0.66422 | 0.99377 | 5.97826 | 0.499673801 | 54.8 | Disease_causing |
| testneg_4442 | 0.08 | 0.03 | 2.365497 | -3 | 3.90237 | -3 | 0.61491 | 0.11954 | 0.11564 | 0.1184 | 12.87958 | 0.11 | 0 | Neutral |
| testneg_4443 | 0.22 | -2.66 | 3.002533 | 3.002533 | -0.133106 | -0.133106 | 0.14304 | 0.10715 | 0.07096 | 0.09965 | 35.87354 | 0.39 | 17 | Neutral |
| testneg_4444 | -0.08 | -0.03 | 4.781986 | -3 | 0.73392 | -1.359951 | 0.47997 | 0.10506 | 0.47816 | 0.66526 | 17.77778 | 0.212162335 | 0 | Neutral |
| testneg_4445 | -0.03 | -0.1 | 4.536943 | -3 | 0.471907 | -3 | 0.08982 | 0.15279 | 0.53424 | 0.08484 | 8.80117 | 0.11 | 0 | Neutral |
| testneg_4446 | 0.07 | 0.38 | 1.946844 | -3 | 1.56241 | -3 | 0.52457 | 0.38957 | 0.08386 | 0.12788 | 20.83333 | 0.075 | 0 | Neutral |
| testneg_4447 | 0 | -0.01 | 0.961397 | -3 | 4.304984 | -3 | 0.08373 | 0.09191 | 0.19898 | 0.14645 | 15.37063 | 0.157239386 | 0 | Neutral |
| testneg_4448 | 0.06 | 0.18 | 2.459196 | -3 | 0.413639 | -3 | 0.63484 | 0.16728 | 0.09402 | 0.12788 | 20.11299 | 0.032825758 | 0 | Neutral |
| testneg_4449 | -0.03 | -1.22 | 2.147571 | -3 | 0.740721 | -3 | 0.1741 | 0.1631 | 0.05716 | 0.12491 | 56.32191 | 0.09 | 0 | Neutral |
| testneg_4450 | 0 | 0 | -0.262607 | -3 | 1.833275 | -0.375499 | 0.21576 | 0.17241 | 0.40981 | 0.21807 | 29.86111 | 0.112401676 | 0 | Neutral |
| testneg_4451 | 0.14 | -0.37 | 5.499025 | -3 | 1.050302 | -3 | 0.19898 | 0.10269 | 0.21972 | 0.12788 | 21.29614 | 0.101333333 | 0 | Neutral |
| testneg_4452 | 0 | 0 | 1.609578 | -3 | 2.581511 | 2.581511 | 0.19898 | 0.07094 | 0.18305 | 0.12743 | 44.83711 | 0.070161395 | 0 | Neutral |
| testneg_4453 | 0.07 | -0.31 | 3.064784 | -3 | 5.559026 | -3 | 0.86835 | 0.1513 | 0.16709 | 0.08659 | 10.41667 | 0.273013889 | 2 | Neutral |
| testneg_4454 | 0 | -0.01 | 3.05843 | 3.05843 | 3.841536 | -3 | 0.74136 | 0.1049 | 0.08002 | 0.08792 | 31.25 | 0.373787879 | 7 | Neutral |
| testneg_4455 | -0.02 | 0.68 | 5.990951 | -3 | 4.945318 | -3 | 0.09542 | 0.14806 | 0.29958 | 0.11203 | 18.33333 | 0.54425075 | 70.8 | Disease_causing |
| testneg_4456 | 0 | -0.01 | 0.558245 | -3 | 3.475784 | -3 | 0.19898 | 0.10362 | 0.13374 | 0.12788 | 11.56063 | 0.028895449 | 0 | Neutral |
| testneg_4457 | 0.14 | -0.37 | 4.795372 | -3 | 0.545055 | -3 | 0.02518 | 0.07165 | 0.17935 | 0.36953 | 17.72727 | 0.270513158 | 1.2 | Neutral |
| testneg_4458 | -0.08 | -0.03 | 0.950149 | -3 | 0.955677 | 0.955677 | 0.31318 | 0.11262 | 0.10034 | 0.10906 | 11.06549 | 0.01036039 | 0 | Neutral |
| testneg_4459 | 0 | -0.01 | 1.715756 | -3 | -0.001756 | -3 | 0.32804 | 0.10857 | 0.16662 | 0.11306 | 39.41832 | 0.000235327 | 0 | Neutral |
| testneg_4460 | 0 | 0 | 0.068014 | 0.068014 | 1.811443 | -3 | 0.20869 | 0.13493 | 0.58063 | 0.12788 | 19.57265 | 0.01 | 0 | Neutral |
| testneg_4461 | 0.21 | 0.15 | 2.584632 | -3 | -0.972856 | -3 | 0.31062 | 0.1222 | 0.19898 | 0.12788 | 30.9756 | 0.022916667 | 0 | Neutral |
| testneg_4462 | -0.03 | -0.1 | 6.211821 | -3 | 0.140432 | -3 | 0.1215 | 0.12788 | 0.88976 | 0.22648 | 5 | 0.38 | 19.4 | Neutral |
| testneg_4463 | -0.03 | -0.12 | 5.525606 | -3 | 0.501771 | -3 | 0.0146 | 0.03045 | 0.62203 | 0.15583 | 14.88464 | 0.35 | 6.8 | Neutral |
| testneg_4464 | 0.18 | -1.05 | 1.855958 | -3 | 1.255802 | -3 | 0.18979 | 0.07948 | 0.20546 | 0.10734 | 17.21847 | 0.03 | 0 | Neutral |
| testneg_4465 | -0.01 | 1.44 | 1.639844 | -3 | 3.875305 | -3 | 0.20219 | 0.10986 | 0.45011 | 0.1751 | 12.77241 | 0.317263142 | 2.2 | Neutral |
| testneg_4466 | 0 | 0 | 3.692426 | -3 | -0.070448 | -3 | 0.19668 | 0.16042 | 0.47997 | 0.10506 | 16.8018 | 0.05015 | 0 | Neutral |
| testneg_4467 | 0.14 | -1.01 | 1.865393 | -3 | 1.169097 | -3 | 0.19898 | 0.10482 | 0.10011 | 0.08014 | 58.75432 | 0.06 | 0 | Neutral |
| testneg_4468 | 0.03 | -1.03 | 1.019521 | -3 | 0.444659 | -3 | 0.67216 | 0.17808 | 0.67393 | 0.12788 | 18.33333 | 0.040875959 | 0 | Neutral |
| testneg_4469 | 0 | 0 | 0.573225 | -3 | 0.780279 | -3 | 0.73947 | 0.2058 | 0.30521 | 0.36067 | 46.74747 | 0.141554487 | 0 | Neutral |
| testneg_4470 | -0.07 | -0.38 | 1.359475 | -3 | 0.040505 | -3 | 0.47975 | 0.18568 | 0.16926 | 0.11871 | 12.75145 | 0.022504181 | 0 | Neutral |
| testneg_4471 | 0 | 0 | 0.389489 | -3 | 6.815776 | -3 | 0.11858 | 0.11551 | 0.19832 | 0.19622 | 22.93866 | 0.402774725 | 18.2 | Neutral |
| testneg_4472 | 0.05 | 0.29 | -0.72639 | -3 | 0.993581 | -3 | 0.19898 | 0.12788 | 0.04508 | 0.12788 | 21.46199 | 0.02 | 0 | Neutral |
| testneg_4473 | -0.03 | -0.1 | -1.468911 | -3 | 5.099439 | -3 | 0.33057 | 0.10549 | 0.0773 | 0.14062 | 12.13337 | 0.281055556 | 1.4 | Neutral |
| testneg_4474 | -0.01 | 1.44 | 4.351718 | -3 | 2.184344 | -3 | 0.16475 | 0.26069 | 0.05472 | 0.12788 | 34.26471 | 0.201047619 | 0 | Neutral |
| testneg_4475 | 0.03 | 0.1 | -1.628421 | -1.628421 | 1.035434 | -3 | 0.54441 | 0.08891 | 0.4558 | 0.9277 | 14.16667 | 0.09 | 0 | Neutral |
| testneg_4476 | 0.21 | 0.15 | 2.654976 | -3 | -0.711743 | -3 | 0.16345 | 0.11716 | 0.43977 | 0.17284 | 13.51351 | 0.11 | 0 | Neutral |
| testneg_4477 | 0.15 | -0.28 | 5.663657 | -3 | -0.823621 | -3 | 0.21989 | 0.10981 | 0.12453 | 0.17736 | 13.13218 | 0.350494118 | 11 | Neutral |
| testneg_4478 | 0.07 | 0.38 | 1.695559 | -3 | 2.513826 | -3 | 0.31773 | 0.10816 | 0.3414 | 0.13575 | 30.24556 | 0.110575658 | 0 | Neutral |
| testneg_4479 | 0.01 | 0.79 | -0.188304 | -3 | 1.228944 | -3 | 0.16418 | 0.12788 | 0.48572 | 0.13977 | 10.83333 | 0.05 | 0 | Neutral |
| testneg_4480 | 0.14 | -0.37 | 3.499396 | -3 | 0.196475 | -3 | 0.69249 | 0.12788 | 0.15562 | 0.10384 | 13.08587 | 0.03 | 0 | Neutral |
| testneg_4481 | 0.14 | -1.01 | 1.084274 | -3 | -0.763816 | -3 | 0.07606 | 0.13082 | 0.12403 | 0.12788 | 19.10253 | 0.02 | 0 | Neutral |
| testneg_4482 | 0.18 | -1.05 | 3.225783 | -3 | -0.811455 | -0.811455 | 0.75978 | 0.11423 | 0.10638 | 0.66554 | 20.37591 | 0.240135387 | 0 | Neutral |
| testneg_4483 | 0.01 | -1.3 | 5.149724 | -3 | 1.92122 | -3 | 0.29679 | 0.12788 | 0.16728 | 0.11868 | 10 | 0.124608543 | 0 | Neutral |
| testneg_4484 | -0.04 | 0.23 | -0.274997 | -3 | 2.632198 | -3 | 0.37501 | 0.0997 | 0.23939 | 0.25284 | 11.25 | 0.120912698 | 0 | Neutral |
| testneg_4485 | 0.22 | -2.66 | 7.430811 | -3 | 0.561 | -3 | 0.11359 | 0.08631 | 0.09184 | 0.38254 | 19.09347 | 0.381277778 | 14 | Neutral |
| testneg_4486 | 0 | 0 | 1.599689 | -3 | -0.337044 | -3 | 0.17057 | 0.12788 | 0.02396 | 0.12703 | 10.83333 | 0.0025 | 0 | Neutral |
| testneg_4487 | 0 | 0 | -0.48561 | -3 | 6.776197 | -3 | 0.09432 | 0.15749 | 0.7219 | 0.41509 | 11.82131 | 0.543971464 | 70.2 | Disease_causing |
| testneg_4488 | 0 | 0 | 0.308908 | -3 | -0.826024 | -3 | 0.39305 | 0.18672 | 0.33057 | 0.10549 | 12.13337 | 0.01 | 0 | Neutral |
| testneg_4489 | -0.17 | -0.38 | 1.808824 | 1.808824 | 3.782185 | -3 | 0.2445 | 0.53069 | 0.83732 | 0.19325 | 10.41667 | 0.546153341 | 73.2 | Disease_causing |
| testneg_4490 | -0.07 | -0.38 | 1.420059 | -3 | -1.375656 | -3 | 0.38115 | 0.21061 | 0.04192 | 0.06306 | 18.73163 | 0.031705997 | 0 | Neutral |
| testneg_4491 | 0 | -0.32 | -0.118786 | -3 | 0.509747 | -3 | 0.19749 | 0.11482 | 0.19898 | 0.08486 | 29.16667 | 0.005649254 | 0 | Neutral |
| testneg_4492 | 0.05 | -0.81 | 5.114262 | -3 | -3.330542 | -3 | 0.18281 | 0.12788 | 0.02723 | 0.04872 | 21.5133 | 0.08 | 0 | Neutral |
| testneg_4493 | 0 | 0 | 0.305495 | -3 | 1.967314 | -3 | 0.98255 | 0.98129 | 0.04946 | 0.12788 | 26.7612 | 0.12 | 0 | Neutral |
| testneg_4494 | 0 | 0 | 1.000919 | -3 | -0.005339 | -3 | 0.16962 | 0.12788 | 0.47192 | 0.11305 | 17.33333 | 0 | 0 | Neutral |
| testneg_4495 | 0.07 | 0.7 | 6.611537 | -3 | 1.034081 | -3 | 0.32201 | 0.10713 | 0.13398 | 0.12788 | 16.8018 | 0.229347527 | 1.8 | Neutral |
| testneg_4496 | 0 | -0.01 | 0.074894 | -3 | 2.510161 | -3 | 0.34225 | 0.12788 | 0.2168 | 0.11297 | 21.53664 | 0.012591099 | 0 | Neutral |
| testneg_4497 | 0 | -0.01 | 0.93478 | -3 | 6.317626 | 4.538431 | 0.57239 | 0.16622 | 0.10383 | 0.08853 | 35.89889 | 0.4 | 18 | Neutral |
| testneg_4498 | 0 | 0 | 1.182206 | -3 | 4.520756 | -3 | 0.93853 | 0.93439 | 0.15267 | 0.12788 | 9.02494 | 0.128535354 | 0 | Neutral |
| testneg_4499 | 0.22 | -2.66 | 6.157395 | -3 | 0.916182 | -3 | 0.07996 | 0.08354 | 0.65857 | 0.26294 | 22.72479 | 0.33 | 5.8 | Neutral |
| testneg_4500 | -0.08 | -0.03 | 1.755955 | -3 | 5.845998 | 0.13957 | 0.9832 | 0.39904 | 0.1231 | 0.3662 | 74.67892 | 0.560061198 | 79.6 | Disease_causing |
| testneg_4501 | 0.03 | -1.03 | 3.976603 | -3 | 0.870165 | 0.667764 | 0.64125 | 0.28792 | 0.10511 | 0.12788 | 13.13218 | 0.090438127 | 0 | Neutral |
| testneg_4502 | 0.07 | 0.01 | 0.395033 | -3 | 0.36854 | -3 | 0.15202 | 0.12722 | 0.58292 | 0.87725 | 37.08756 | 0.09 | 0 | Neutral |
| testneg_4503 | 0 | 0 | -0.100514 | -3 | -0.811455 | -0.811455 | 0.16469 | 0.16042 | 0.10638 | 0.66554 | 11.875 | 0.129960311 | 0 | Neutral |
| testneg_4504 | 0.06 | -0.57 | 0.132908 | -3 | 4.260553 | -1.064743 | 0.12643 | 0.12266 | 0.13624 | 0.09458 | 10 | 0.060477273 | 0 | Neutral |
| testneg_4505 | 0 | 0 | 0.347529 | -3 | 2.698795 | -3 | 0.22443 | 0.10657 | 0.17806 | 0.12788 | 27.3569 | 0.011904663 | 0 | Neutral |
| testneg_4506 | -0.08 | -0.03 | 1.155408 | -3 | 0.101783 | -3 | 0.04161 | 0.12788 | 0.06196 | 0.10881 | 13.02635 | 0.04 | 0 | Neutral |
| testneg_4507 | 0 | 0 | 0.917232 | -3 | 0.413122 | -3 | 0.25556 | 0.09342 | 0.32531 | 0.12351 | 21.1411 | 0.01 | 0 | Neutral |
| testneg_4508 | 0 | -0.01 | 0.614361 | -3 | 3.067866 | -3 | 0.95998 | 0.11262 | 0.04831 | 0.12788 | 20.40495 | 0.061428571 | 0 | Neutral |
| testneg_4509 | 0.01 | -1.44 | 3.309654 | -3 | 0.939511 | -3 | 0.09645 | 0.13239 | 0.20379 | 0.35591 | 17.98406 | 0.267674428 | 0.8 | Neutral |
| testneg_4510 | 0 | 0 | 0.013394 | -3 | -1.165102 | -1.165102 | 0.12582 | 0.12788 | 0.37237 | 0.36829 | 16.8018 | 0.071386555 | 0 | Neutral |
| testneg_4511 | 0 | 0 | 0.307719 | -3 | 1.808824 | 1.808824 | 0.16091 | 0.11571 | 0.2445 | 0.53069 | 10.52098 | 0.080576923 | 0 | Neutral |
| testneg_4512 | 0 | 0 | 0.990948 | -3 | 2.987486 | 1.319914 | 0.31859 | 0.11262 | 0.08896 | 0.12788 | 20.83333 | 0.04 | 0 | Neutral |
| testneg_4513 | 0.00055944 | 0.00363636 | 1.360315 | -3 | 3.112733 | -3 | 0.29482 | 0.15726 | 0.11773 | 0.48758 | 10.41667 | 0.223705275 | 0.2 | Neutral |

| | | | | | | | | | | | | | | |
|---|---|---|---|---|---|---|---|---|---|---|---|---|---|---|
| testneg_4514 | 0.06 | -0.57 | 2.764203 | -3 | 2.300256 | 2.300256 | 0.68228 | 0.09762 | 0.14996 | 0.09121 | 16.25133 | 0.09 | 0 | Neutral |
| testneg_4515 | 0.21 | 0.15 | 7.586474 | -3 | 4.228466 | -3 | 0.13961 | 0.09565 | 0.48688 | 0.13071 | 11.74429 | 0.55 | 74.2 | Disease_causing |
| testneg_4516 | 0.04 | 2.01 | -0.707182 | -3 | 2.622551 | -3 | 0.06789 | 0.09707 | 0.11849 | 0.20101 | 14.16667 | 0.233766168 | 0 | Neutral |
| testneg_4517 | 0 | 0 | 0.039637 | -3 | -0.247762 | -3 | 0.20304 | 0.31462 | 0.44654 | 0.10907 | 18.88889 | 0.011116087 | 0 | Neutral |
| testneg_4518 | 0.19 | 0.51 | -0.093394 | -0.093394 | 1.678575 | -3 | 0.34587 | 0.29822 | 0.5958 | 0.0929 | 20 | 0.08 | 0 | Neutral |
| testneg_4519 | 0.16 | 0.08 | 4.384791 | -3 | 2.178269 | -3 | 0.13244 | 0.14875 | 0.2521 | 0.1784 | 21.51709 | 0.387138502 | 14.8 | Neutral |
| testneg_4520 | -0.04 | -2.01 | 3.277724 | -3 | 5.392275 | -3 | 0.10438 | 0.20558 | 0.10385 | 0.09775 | 12.65469 | 0.379444444 | 12.2 | Neutral |
| testneg_4521 | 0 | 0 | 1.040567 | -3 | 4.020236 | -3 | 0.10869 | 0.1503 | 0.1109 | 0.12788 | 17.21847 | 0.13 | 0 | Neutral |
| testneg_4522 | 0.07 | 0.38 | 1.350227 | -3 | 4.489998 | -3 | 0.20765 | 0.27124 | 0.03378 | 0.07001 | 18.33333 | 0.16 | 0 | Neutral |
| testneg_4523 | 0.16 | 0.08 | -0.169987 | -3 | 1.483649 | 1.483649 | 0.99445 | 0.10396 | 0.47678 | 0.38085 | 29.41849 | 0.182232143 | 0 | Neutral |
| testneg_4524 | 0 | 0 | 0.024952 | -3 | 1.603037 | -3 | 0.19423 | 0.14002 | 0.09522 | 0.20323 | 18.37615 | 0.082498688 | 0 | Neutral |
| testneg_4525 | 0.17 | 0.38 | 2.311167 | 2.311167 | 0.14373 | -3 | 0.05413 | 0.05691 | 0.17728 | 0.46704 | 18.88889 | 0.32 | 7.6 | Neutral |
| testneg_4526 | 0 | 0 | 0.057075 | -3 | 1.691672 | -3 | 0.97504 | 0.18545 | 0.30789 | 0.15135 | 10.83333 | 0.051679228 | 0 | Neutral |
| testneg_4527 | 0.05 | -0.81 | 3.217486 | -3 | 0.292982 | -3 | 0.74699 | 0.12286 | 0.15677 | 0.18537 | 17.21847 | 0.122892857 | 0 | Neutral |
| testneg_4528 | -0.08 | -0.03 | 2.280979 | -3 | -1.013068 | -1.27012 | 0.01704 | 0.05734 | 0.40076 | 0.11341 | 12.75441 | 0.05 | 0 | Neutral |
| testneg_4529 | 0 | 0 | 1.052989 | -3 | 0.38416 | -1.239866 | 0.9874 | 0.15588 | 0.12173 | 0.12788 | 16.42913 | 0.02 | 0 | Neutral |
| testneg_4530 | -0.07 | -0.38 | 2.173191 | -3 | 0.791297 | -3 | 0.18233 | 0.14781 | 0.50276 | 0.09638 | 21.71775 | 0.03 | 0 | Neutral |
| testneg_4531 | -0.21 | -0.15 | -2.734447 | -3 | 1.125734 | -3 | 0.48023 | 0.12788 | 0.12576 | 0.08486 | 10.80967 | 0.07 | 0 | Neutral |
| testneg_4532 | -0.02 | -0.45 | 3.127301 | -3 | 0.020099 | -3 | 0.1621 | 0.1641 | 0.12203 | 0.13515 | 18.33333 | 0.172595995 | 0 | Neutral |
| testneg_4533 | 0.02 | -0.63 | 2.125245 | -3 | 1.514563 | -3 | 0.99338 | 0.14229 | 0.60152 | 0.12166 | 5.57306 | 0.04 | 0 | Neutral |
| testneg_4534 | 0 | 0 | 1.339916 | -3 | 1.020872 | -3 | 0.19898 | 0.09171 | 0.31434 | 0.11315 | 23.79024 | 0.000223471 | 0 | Neutral |
| testneg_4535 | 0 | -0.01 | 1.961035 | -3 | 0.945866 | -3 | 0.55105 | 0.43208 | 0.23537 | 0.33997 | 10 | 0.093342105 | 0 | Neutral |
| testneg_4536 | 0.14 | -0.69 | 1.692426 | -3 | 0.372849 | -3 | 0.09306 | 0.09084 | 0.10286 | 0.26819 | 21.47432 | 0.11 | 0 | Neutral |
| testneg_4537 | -0.21 | -0.15 | 2.61677 | -3 | 0.911948 | -3 | 0.60285 | 0.15179 | 0.19898 | 0.12788 | 16.8018 | 0.07 | 0 | Neutral |
| testneg_4538 | 0.14 | -0.37 | 4.232017 | -3 | -2.565987 | -3 | 0.5726 | 0.1165 | 0.06072 | 0.09873 | 10.41667 | 0.05 | 0 | Neutral |
| testneg_4539 | 0 | 0 | 0.77406 | -3 | 1.339264 | -3 | 0.1192 | 0.1508 | 0.3012 | 0.10317 | 18.75339 | 0.02 | 0 | Neutral |
| testneg_4540 | 0 | 0 | 0.307719 | -3 | -0.493952 | -3 | 0.16091 | 0.11571 | 0.05444 | 0.12788 | 12.46106 | 0.000241934 | 0 | Neutral |
| testneg_4541 | 0.07 | 0.38 | 1.044673 | -3 | 1.474779 | 1.474779 | 0.73808 | 0.15588 | 0.0882 | 0.12788 | 32.36768 | 0.05 | 0 | Neutral |
| testneg_4542 | 0 | 0 | 0.510259 | -3 | -0.784242 | -3 | 0.08186 | 0.11813 | 0.18153 | 0.10845 | 20.83333 | 0.00094494 | 0 | Neutral |
| testneg_4543 | 0 | 0 | 0.770471 | -3 | 1.885526 | -3 | 0.99859 | 0.12788 | 0.11233 | 0.28224 | 10 | 0.070457973 | 0 | Neutral |
| testneg_4544 | -0.18 | -0.16 | 0.60982 | 0.60982 | 2.795001 | -3 | 0.23844 | 0.27772 | 0.16973 | 0.17555 | 10.26553 | 0.365712074 | 10.8 | Neutral |
| testneg_4545 | 0 | -0.01 | 2.171434 | -3 | -0.030255 | -1.434272 | 0.49425 | 0.25538 | 0.143 | 0.11236 | 14.95396 | 0.020034965 | 0 | Neutral |
| testneg_4546 | 0 | 0 | 1.059479 | -3 | 5.243885 | -3 | 0.38002 | 0.22508 | 0.13693 | 0.12788 | 10.60658 | 0.137049603 | 0 | Neutral |
| testneg_4547 | 0.14 | -0.37 | 2.102067 | -3 | 7.150139 | -3 | 0.95557 | 0.38833 | 0.13751 | 0.13274 | 14.44144 | 0.58 | 84 | Disease_causing |
| testneg_4548 | 0 | 0 | 0.57285 | -3 | 0.170454 | -3 | 0.20501 | 0.6166 | 0.10803 | 0.14652 | 10.83333 | 0.141480661 | 0 | Neutral |
| testneg_4549 | 0.21 | 0.15 | 4.319063 | -3 | -1.595702 | -3 | 0.1647 | 0.12076 | 0.82131 | 0.12788 | 26.41534 | 0.062788462 | 0 | Neutral |
| testneg_4550 | 0.01 | -0.57 | 2.700261 | -3 | 0.435447 | -3 | 0.76957 | 0.13588 | 0.21941 | 0.46348 | 5 | 0.120318296 | 0 | Neutral |
| testneg_4551 | 0.18 | -1.05 | 6.524735 | -3 | -0.59601 | -3 | 0.20116 | 0.10008 | 0.09569 | 0.09246 | 31.5924 | 0.17 | 0.2 | Neutral |
| testneg_4552 | 0 | 0 | 0.454724 | -3 | -0.365462 | -3 | 0.33022 | 0.13818 | 0.19898 | 0.12788 | 18.4511 | 0.000188746 | 0 | Neutral |
| testneg_4553 | -0.19 | -0.51 | 1.521336 | -3 | 0.101783 | -3 | 0.39553 | 0.11991 | 0.06196 | 0.10881 | 23.35749 | 0.04 | 0 | Neutral |
| testneg_4554 | 0 | 0 | 0.730138 | -3 | 0.372325 | -3 | 0.16952 | 0.10613 | 0.62197 | 0.12788 | 15.16951 | 0 | 0 | Neutral |
| testneg_4555 | 0.01 | 0.79 | 0.470746 | 0.470746 | 4.048415 | -3 | 0.99543 | 0.12788 | 0.19898 | 0.12788 | 13.27141 | 0.144506033 | 0 | Neutral |
| testneg_4556 | 0 | 0 | 0.590474 | -3 | 8.786409 | -3 | 0.14484 | 0.14928 | 0.19898 | 0.12788 | 30.80058 | 0.42 | 24.8 | Neutral |
| testneg_4557 | -0.02 | 0.36 | 6.611139 | -3 | 5.338598 | -3 | 0.36324 | 0.15729 | 0.22179 | 0.09831 | 18.6768 | 0.665232498 | 96.6 | Disease_causing |
| testneg_4558 | 0 | 0 | 1.269233 | -3 | -0.468464 | -3 | 0.18998 | 0.26095 | 0.19898 | 0.12788 | 17.55703 | 0.01 | 0 | Neutral |
| testneg_4559 | -0.15 | -0.04 | 2.846922 | -3 | -3.00145 | -3 | 0.18081 | 0.1258 | 0.22645 | 0.1704 | 12.49191 | 0.149982215 | 0 | Neutral |
| testneg_4560 | 0.07 | 0.38 | 2.239242 | -3 | 4.293539 | -3 | 0.13746 | 0.12788 | 0.11891 | 0.11186 | 17.28802 | 0.077106921 | 0 | Neutral |
| testneg_4561 | 0 | 0 | 2.566346 | -3 | 5.185419 | -3 | 0.19898 | 0.12788 | 0.17093 | 0.23281 | 17.39425 | 0.370510722 | 14.2 | Neutral |
| testneg_4562 | 0.01 | -1.44 | 1.092456 | -3 | -0.71253 | -0.71253 | 0.90296 | 0.28521 | 0.03563 | 0.10455 | 11.875 | 0.050690476 | 0 | Neutral |
| testneg_4563 | 0 | -0.32 | -0.415118 | -3 | 7.089732 | -3 | 0.1144 | 0.12788 | 0.10168 | 0.12788 | 24.64158 | 0.283888889 | 1.8 | Neutral |
| testneg_4564 | 0 | 0 | 0.004869 | -3 | 0.699547 | -3 | 0.16134 | 0.1847 | 0.25637 | 0.37408 | 11.93647 | 0.081349707 | 0 | Neutral |
| testneg_4565 | 0.14 | -1.01 | 3.372659 | -3 | 0.010391 | -3 | 0.15263 | 0.12788 | 0.03558 | 0.05202 | 27.3569 | 0.03 | 0 | Neutral |
| testneg_4566 | 0.04 | -0.23 | 4.939659 | -3 | 1.943488 | -3 | 0.13563 | 0.12378 | 0.25011 | 0.12092 | 4.44444 | 0.067718468 | 0 | Neutral |
| testneg_4567 | 0.04 | -0.23 | 1.34082 | -3 | 1.311634 | -3 | 0.20116 | 0.10008 | 0.93853 | 0.93439 | 10.22898 | 0.070296425 | 0 | Neutral |
| testneg_4568 | 0 | -0.01 | 1.955355 | -3 | -0.531475 | -3 | 0.33795 | 0.17455 | 0.08686 | 0.0953 | 16.85532 | 0.010606618 | 0 | Neutral |
| testneg_4569 | 0.17 | 0.38 | 2.7281 | -3 | 8.044918 | -3 | 0.33163 | 0.26601 | 0.06301 | 0.32138 | 10.83333 | 0.76702957 | 100 | Disease_causing |
| testneg_4570 | 0 | 0 | 0.571304 | -3 | 1.330341 | -3 | 0.99448 | 0.67288 | 0.469 | 0.25588 | 4.44444 | 0.061855459 | 0 | Neutral |
| testneg_4571 | 0 | 0 | -0.061652 | -3 | 1.258927 | -3 | 0.83074 | 0.14769 | 0.08903 | 0.12788 | 27.1735 | 0.040203862 | 0 | Neutral |
| testneg_4572 | 0.11 | -0.19 | 1.233798 | -3 | 1.790281 | -3 | 0.64701 | 0.94571 | 0.02059 | 0.066 | 18.33333 | 0.10325 | 0 | Neutral |
| testneg_4573 | 0 | 0 | 1.016702 | -3 | 2.303477 | -3 | 0.09306 | 0.09084 | 0.53216 | 0.84775 | 13.97895 | 0.08975 | 0 | Neutral |
| testneg_4574 | 0 | 0 | 1.235917 | -3 | 0.634312 | -3 | 0.07543 | 0.10211 | 0.07681 | 0.22865 | 18.43039 | 0.050154431 | 0 | Neutral |
| testneg_4575 | -0.05 | -0.29 | -1.641768 | -3 | 1.418002 | -3 | 0.31648 | 0.12788 | 0.19898 | 0.12788 | 43.99077 | 0.01 | 0 | Neutral |
| testneg_4576 | 0.14 | -0.37 | 2.01573 | -3 | 1.673881 | -3 | 0.32531 | 0.12351 | 0.07923 | 0.08035 | 13.51351 | 0.01 | 0 | Neutral |
| testneg_4577 | 0.07 | -0.31 | 3.064784 | -3 | 0.271251 | -3 | 0.86835 | 0.1513 | 0.42836 | 0.12788 | 10 | 0.060103652 | 0 | Neutral |
| testneg_4578 | 0 | 0 | 1.470632 | -3 | 6.776625 | -3 | 0.67549 | 0.1273 | 0.17039 | 0.12788 | 20.29691 | 0.170298691 | 0.2 | Neutral |
| testneg_4579 | -0.07 | -0.38 | 1.367149 | -3 | 0.076953 | -3 | 0.06877 | 0.17777 | 0.10187 | 0.12339 | 27.4024 | 0.06 | 0 | Neutral |
| testneg_4580 | 0.15 | -0.28 | 3.27749 | -3 | 7.492522 | -3 | 0.11329 | 0.12116 | 0.13433 | 0.09578 | 35.95985 | 0.38 | 16.2 | Neutral |
| testneg_4581 | -0.09 | 2.17 | 4.614663 | -3 | 1.450167 | -3 | 0.20604 | 0.10821 | 0.19898 | 0.12788 | 25.73432 | 0.11 | 0 | Neutral |
| testneg_4582 | 0 | -0.01 | 1.961035 | -3 | 0.887782 | -3 | 0.55105 | 0.43208 | 0.22155 | 0.1062 | 11.92204 | 0.010534188 | 0 | Neutral |
| testneg_4583 | 0 | 0 | 0.473553 | -3 | 1.131275 | -3 | 0.31062 | 0.1222 | 0.0931 | 0.12788 | 30.15677 | 0.01 | 0 | Neutral |
| testneg_4584 | 0.09 | -0.3 | -0.335332 | -3 | 1.211799 | -3 | 0.50388 | 0.12295 | 0.39932 | 0.10693 | 13.2019 | 0.02 | 0 | Neutral |
| testneg_4585 | 0.01 | -1.44 | 2.788395 | -3 | 0.726148 | -3 | 0.23567 | 0.12788 | 0.10782 | 0.14795 | 21.60834 | 0.12 | 0 | Neutral |
| testneg_4586 | 0 | 0.32 | 3.268811 | -3 | 1.953142 | -3 | 0.17141 | 0.1104 | 0.47483 | 0.12788 | 28.10041 | 0.03035625 | 0 | Neutral |
| testneg_4587 | -0.14 | 0.37 | 2.327743 | -3 | 0.086889 | -3 | 0.0538 | 0.0823 | 0.66693 | 0.0986 | 13.75 | 0.100040984 | 0 | Neutral |
| testneg_4588 | 0.09 | -0.3 | 6.630335 | -3 | -1.000185 | -3 | 0.9193 | 0.14584 | 0.19024 | 0.1132 | 11.23846 | 0.34 | 7.2 | Neutral |
| testneg_4589 | 0 | 0 | 0.213657 | -3 | 1.478329 | -3 | 0.3769 | 0.12788 | 0.27614 | 0.12085 | 17.47657 | 0 | 0 | Neutral |
| testneg_4590 | 0.09 | -0.3 | 6.180112 | -3 | -1.259442 | -3 | 0.38171 | 0.15977 | 0.24185 | 0.12788 | 22.61299 | 0.241010101 | 0.6 | Neutral |
| testneg_4591 | 0 | 0 | 1.966897 | -3 | 0.300343 | -3 | 0.39262 | 0.12788 | 0.04032 | 0.12788 | 10.83333 | 0.01 | 0 | Neutral |
| testneg_4592 | -0.04 | 1.61 | 6.77188 | -3 | 4.605651 | -3 | 0.19898 | 0.13162 | 0.10719 | 0.08618 | 19.09347 | 0.520138889 | 64.8 | Disease_causing |
| testneg_4593 | -0.15 | 1.06 | 6.682345 | -3 | -0.285825 | -3 | 0.08268 | 0.09939 | 0.15016 | 0.08826 | 19.44444 | 0.26 | 1.6 | Neutral |
| testneg_4594 | 0.01 | 0.8 | 2.989973 | -3 | 1.010214 | -3 | 0.24 | 0.2063 | 0.13141 | 0.1123 | 9.91553 | 0.08 | 0 | Neutral |
| testneg_4595 | 0.01 | 0.8 | 2.60181 | -3 | 0.720142 | -3 | 0.45014 | 0.12788 | 0.28474 | 0.12788 | 36.15928 | 0.02 | 0 | Neutral |
| testneg_4596 | -0.01 | -0.79 | 1.33295 | -3 | 6.138574 | -3 | 0.23088 | 0.12788 | 0.13559 | 0.1946 | 30.13935 | 0.410514383 | 20.6 | Neutral |
| testneg_4597 | 0.02 | 0.45 | -1.0834 | -3 | 3.566467 | -3 | 0.14248 | 0.12788 | 0.17548 | 0.11384 | 23.68839 | 0.060983871 | 0 | Neutral |
| testneg_4598 | 0 | 0 | 0.028094 | -3 | 0.128042 | -3 | 0.46712 | 0.93559 | 0.13702 | 0.12788 | 10.3336 | 0.08 | 0 | Neutral |
| testneg_4599 | 0.03 | -1.03 | 1.493352 | -3 | 4.428774 | -3 | 0.17364 | 0.18111 | 0.07175 | 0.29293 | 11.92061 | 0.496570902 | 53.6 | Disease_causing |
| testneg_4600 | -0.02 | -0.45 | 6.418252 | -3 | 7.917194 | -3 | 0.43314 | 0.67252 | 0.39019 | 0.18821 | 10.41667 | 0.952615863 | 100 | Disease_causing |
| testneg_4601 | -0.07 | -0.38 | 2.16463 | -3 | 0.390383 | -3 | 0.13555 | 0.16663 | 0.08213 | 0.08913 | 10.41667 | 0.04 | 0 | Neutral |
| testneg_4602 | -0.15 | -0.04 | 5.146748 | -3 | 2.179755 | -3 | 0.16778 | 0.82322 | 0.93246 | 0.12788 | 10.41667 | 0.349436813 | 4.6 | Neutral |
| testneg_4603 | 0 | 0 | -0.68826 | -3 | 2.859591 | -3 | 0.32076 | 0.14229 | 0.1576 | 0.12788 | 10.41667 | 0.03 | 0 | Neutral |
| testneg_4604 | 0.07 | 0.38 | -1.041977 | -3 | -0.01248 | -3 | 0.49665 | 0.20425 | 0.61231 | 0.13362 | 18.88889 | 0.162467949 | 0 | Neutral |
| testneg_4605 | -0.02 | 0.63 | 2.115956 | -3 | 5.183855 | -3 | 0.47443 | 0.11062 | 0.12853 | 0.08565 | 27.13392 | 0.120076923 | 0 | Neutral |
| testneg_4606 | -0.08 | -0.03 | -0.165646 | -3 | 1.677581 | -3 | 0.69426 | 0.13242 | 0.74919 | 0.85508 | 10 | 0.093571548 | 0 | Neutral |
| testneg_4607 | 0 | 0 | 0.506959 | -3 | 2.840836 | -3 | 0.84236 | 0.14093 | 0.0669 | 0.10997 | 17.77778 | 0.04 | 0 | Neutral |
| testneg_4608 | 0.01 | -1.44 | 2.064834 | -3 | 4.161059 | -3 | 0.99417 | 0.17365 | 0.22225 | 0.11186 | 18.88889 | 0.180368828 | 0 | Neutral |
| testneg_4609 | 0 | 0 | 0.101787 | -3 | 2.956317 | -3 | 0.13782 | 0.13794 | 0.9637 | 0.12788 | 16.8018 | 0.031193853 | 0 | Neutral |
| testneg_4610 | 0 | 0 | 4.54534 | -3 | 2.920916 | -3 | 0.06713 | 0.11112 | 0.19898 | 0.12788 | 43.84582 | 0.095810185 | 0 | Neutral |
| testneg_4611 | 0 | 0 | 1.169905 | -3 | 5.282618 | -3 | 0.51398 | 0.38821 | 0.4676 | 0.11257 | 28.03426 | 0.132158258 | 0 | Neutral |
| testneg_4612 | -0.05 | 0.81 | 4.111365 | -3 | 6.6361 | -3 | 0.1462 | 0.18896 | 0.26618 | 0.2057 | 12.29167 | 0.89 | 100 | Disease_causing |
| testneg_4613 | 0 | 0 | 1.012812 | -3 | 2.051799 | -3 | 0.53313 | 0.14072 | 0.14624 | 0.10369 | 10.83333 | 0.02 | 0 | Neutral |
| testneg_4614 | 0.04 | -0.23 | 1.635811 | -3 | 0.490527 | -3 | 0.11187 | 0.10744 | 0.11337 | 0.09363 | 16.8018 | 0.000108708 | 0 | Neutral |
| testneg_4615 | 0 | 0 | 0.583408 | 0.583408 | 0.452004 | -3 | 0.05386 | 0.09802 | 0.09176 | 0.07645 | 32.26002 | 0.06 | 0 | Neutral |
| testneg_4616 | 0.09 | -0.3 | 2.385019 | -3 | 1.262326 | -1.335936 | 0.19898 | 0.12788 | 0.13118 | 0.08646 | 13.51351 | 0.01 | 0 | Neutral |
| testneg_4617 | -0.07 | -0.38 | 2.626972 | -3 | 3.776689 | -3 | 0.47775 | 0.10103 | 0.17818 | 0.15728 | 28.01661 | 0.333453777 | 5.2 | Neutral |
| testneg_4618 | 0.03 | 1.22 | 1.399794 | -3 | -0.434903 | -3 | 0.49425 | 0.25538 | 0.40981 | 0.21807 | 17.77778 | 0.166988748 | 0 | Neutral |
| testneg_4619 | -0.01 | -0.79 | 2.794319 | -3 | 2.71072 | -3 | 0.60812 | 0.19286 | 0.09719 | 0.09576 | 22.33207 | 0.133005952 | 0 | Neutral |
| testneg_4620 | -0.19 | 0.27 | 2.726709 | -3 | 2.338836 | -3 | 0.05904 | 0.09282 | 0.10011 | 0.08014 | 32.0935 | 0.13 | 0 | Neutral |
| testneg_4621 | 0.07 | 0.38 | 0.140559 | -3 | -0.206099 | -3 | 0.68029 | 0.12788 | 0.17639 | 0.12919 | 10.83333 | 0.020741758 | 0 | Neutral |
| testneg_4622 | 0 | 0 | 2.030989 | -3 | 2.446207 | -3 | 0.26264 | 0.13812 | 0.16781 | 0.12788 | 6.00263 | 0.02 | 0 | Neutral |
| testneg_4623 | 0.08 | 0.03 | -1.691212 | -1.691212 | 1.999625 | -3 | 0.13985 | 0.08349 | 0.58587 | 0.09866 | 18.81641 | 0.020198195 | 0 | Neutral |
| testneg_4624 | -0.02 | 0.68 | 1.8794 | -3 | 1.956887 | -3 | 0.54711 | 0.11809 | 0.1087 | 0.08523 | 23.34541 | 0.04 | 0 | Neutral |
| testneg_4625 | 0 | 0 | 1.145768 | -3 | 2.360629 | -3 | 0.22658 | 0.20646 | 0.10566 | 0.12788 | 11.11559 | 0.050075188 | 0 | Neutral |
| testneg_4626 | -0.08 | -0.03 | 0.750291 | -3 | 1.129475 | -3 | 0.09275 | 0.14911 | 0.27887 | 0.19826 | 17.77778 | 0.13 | 0 | Neutral |
| testneg_4627 | 0 | 0 | 0.462587 | -3 | 2.423449 | -3 | 0.26014 | 0.12378 | 0.30837 | 0.19754 | 22.05417 | 0.093222611 | 0 | Neutral |
| testneg_4628 | -0.02 | 0.63 | 2.928857 | -3 | 0.27289 | -3 | 0.11534 | 0.12281 | 0.14668 | 0.25103 | 17.31419 | 0.154824341 | 0 | Neutral |
| testneg_4629 | 0 | 0 | 1.004251 | -3 | 1.463131 | -3 | 0.56285 | 0.18619 | 0.16709 | 0.08659 | 12.40248 | 0.011892334 | 0 | Neutral |
| testneg_4630 | 0.01 | 0.8 | 1.074849 | -3 | 3.036045 | -3 | 0.06775 | 0.09511 | 0.16098 | 0.12788 | 23.21237 | 0.07 | 0 | Neutral |
| testneg_4631 | 0.03 | -1.03 | 3.626928 | -3 | -0.725111 | -0.725111 | 0.44086 | 0.173 | 0.05116 | 0.08996 | 10.25938 | 0.08 | 0 | Neutral |
| testneg_4632 | 0.02 | 0.45 | -1.097382 | -3 | 1.123147 | -3 | 0.17471 | 0.12788 | 0.59947 | 0.14339 | 13.12327 | 0.031040541 | 0 | Neutral |
| testneg_4633 | -0.15 | -0.06 | 6.682019 | -3 | 2.835811 | -3 | 0.23005 | 0.37216 | 0.0406 | 0.12788 | 10.83333 | 0.56 | 78.4 | Disease_causing |
| testneg_4634 | -0.03 | -0.1 | 0.212913 | -3 | -1.179093 | -3 | 0.11391 | 0.19434 | 0.32948 | 0.13624 | 18.6768 | 0.1312511 | 0 | Neutral |
| testneg_4635 | -0.07 | -0.38 | 1.422631 | -3 | 4.488254 | -3 | 0.09024 | 0.08559 | 0.19898 | 0.08855 | 10.41667 | 0.080886106 | 0 | Neutral |

| | | | | | | | | | | | | | | |
|---|---|---|---|---|---|---|---|---|---|---|---|---|---|---|
| testneg_4636 | -0.03 | -0.1 | 0.968512 | -3 | 3.59862 | -3 | 0.40074 | 0.36296 | 0.2016 | 0.13956 | 27.73348 | 0.377794324 | 12.4 | Neutral |
| testneg_4637 | -0.03 | -0.12 | 0.38962 | -3 | -0.831527 | -3 | 0.49745 | 0.1103 | 0.27783 | 0.62384 | 18.6768 | 0.070669643 | 0 | Neutral |
| testneg_4638 | 0.22 | -2.66 | 7.366549 | -3 | 0.618993 | 0.618993 | 0.143 | 0.11236 | 0.04207 | 0.12788 | 23.14403 | 0.24 | 1 | Neutral |
| testneg_4639 | 0 | -0.01 | 1.245581 | -3 | 0.950521 | -3 | 0.60793 | 0.18264 | 0.2734 | 0.2578 | 10.41667 | 0.040757331 | 0 | Neutral |
| testneg_4640 | -0.07 | -0.38 | 1.400536 | -3 | 11.739453 | -3 | 0.89972 | 0.10766 | 0.18328 | 0.12788 | 18.13386 | 0.34 | 5.2 | Neutral |
| testneg_4641 | 0.07 | 0.7 | 5.788681 | -3 | 1.602273 | -3 | 0.81212 | 0.14915 | 0.08531 | 0.12788 | 13.04617 | 0.276680556 | 3.2 | Neutral |
| testneg_4642 | 0.1 | 1.9 | 2.756314 | -3 | 3.546589 | -3 | 0.29044 | 0.12142 | 0.37109 | 0.12788 | 13.14653 | 0.17 | 0 | Neutral |
| testneg_4643 | -0.04 | 0.23 | 2.526107 | -3 | 0.082683 | -3 | 0.64721 | 0.8758 | 0.19898 | 0.12788 | 11.59278 | 0.080258291 | 0 | Neutral |
| testneg_4644 | 0.04 | -0.23 | -0.057579 | -3 | 0.564419 | -3 | 0.05476 | 0.12788 | 0.04851 | 0.12788 | 28.30274 | 0.06 | 0 | Neutral |
| testneg_4645 | -0.06 | -0.18 | 2.204221 | -3 | 5.583477 | -3 | 0.26916 | 0.13285 | 0.05754 | 0.10881 | 10.22898 | 0.17 | 0 | Neutral |
| testneg_4646 | 0 | 0 | 0.767988 | -3 | 3.188044 | -3 | 0.16361 | 0.13388 | 0.04829 | 0.09848 | 13.51351 | 0.050196078 | 0 | Neutral |
| testneg_4647 | 0 | 0 | 2.10021 | -3 | 3.804607 | -3 | 0.15526 | 0.08511 | 0.36694 | 0.12788 | 28.95241 | 0.051991205 | 0 | Neutral |
| testneg_4648 | 0.03 | 1.22 | 2.244475 | -3 | -0.195771 | -3 | 0.34406 | 0.09894 | 0.26291 | 0.10317 | 11.875 | 0.04 | 0 | Neutral |
| testneg_4649 | 0.18 | -1.05 | 2.305068 | -3 | 0.418626 | 0.418626 | 0.19898 | 0.12788 | 0.53648 | 0.1104 | 33.06212 | 0.03 | 0 | Neutral |
| testneg_4650 | 0.23 | -1.04 | 2.712609 | -3 | 2.066006 | -3 | 0.03563 | 0.10455 | 0.16321 | 0.09621 | 18.33333 | 0.08 | 0 | Neutral |
| testneg_4651 | 0 | 0 | 0.772627 | -3 | -0.004368 | -3 | 0.19898 | 0.12788 | 0.1977 | 0.09728 | 20.00133 | 0 | 0 | Neutral |
| testneg_4652 | 0.01 | 0.79 | -2.720928 | -3 | 0.944356 | -3 | 0.55367 | 0.57993 | 0.32532 | 0.09651 | 10.83333 | 0.076364391 | 0 | Neutral |
| testneg_4653 | -0.04 | 1.36 | 4.511504 | -3 | 1.541736 | -3 | 0.78741 | 0.22085 | 0.21978 | 0.16522 | 18.88889 | 0.352279567 | 7 | Neutral |
| testneg_4654 | 0.19 | -0.25 | 2.546301 | -3 | 6.230771 | -3 | 0.12689 | 0.13376 | 0.10723 | 0.33098 | 14.90898 | 0.600166203 | 88.2 | Disease_causing |
| testneg_4655 | 0 | 0 | 1.372857 | -3 | 3.159116 | -3 | 0.14378 | 0.41915 | 0.54891 | 0.55746 | 11.96429 | 0.332208965 | 7.8 | Neutral |
| testneg_4656 | 0.23 | -1.04 | 7.128483 | -3 | -0.144702 | -3 | 0.25989 | 0.11651 | 0.28162 | 0.12788 | 33.51351 | 0.2 | 0.4 | Neutral |
| testneg_4657 | 0.06 | -0.57 | 3.795863 | -3 | 3.013004 | -3 | 0.68055 | 0.12788 | 0.15537 | 0.19705 | 18.28996 | 0.423351449 | 25.4 | Neutral |
| testneg_4658 | 0.14 | -0.37 | 4.514811 | -3 | 1.999648 | 1.999648 | 0.48662 | 0.23781 | 0.12626 | 0.10279 | 20.78554 | 0.16625 | 0 | Neutral |
| testneg_4659 | -0.07 | -0.33 | 1.58358 | 1.22676 | -0.388631 | -3 | 0.4069 | 0.3841 | 0.19227 | 0.1096 | 12.84759 | 0.05 | 0 | Neutral |
| testneg_4660 | 0 | 0 | 1.315096 | -3 | 1.272902 | -3 | 0.26972 | 0.12484 | 0.19898 | 0.12788 | 29.70767 | 0 | 0 | Neutral |
| testneg_4661 | 0.09 | -0.3 | 3.79415 | -3 | 6.561232 | 5.384088 | 0.26434 | 0.12788 | 0.15016 | 0.08826 | 29.9024 | 0.49 | 51.6 | Disease_causing |
| testneg_4662 | 0 | 0 | 1.031973 | -3 | 2.352957 | -3 | 0.18256 | 0.13499 | 0.83067 | 0.10583 | 19.96169 | 0.020152984 | 0 | Neutral |
| testneg_4663 | 0 | 0 | -0.153757 | -3 | 1.104931 | -3 | 0.23027 | 0.37171 | 0.06708 | 0.09112 | 20.83333 | 0.04 | 0 | Neutral |
| testneg_4664 | 0.03 | 0.1 | 1.374967 | -3 | 5.634496 | -3 | 0.26997 | 0.63944 | 0.13331 | 0.13638 | 20 | 0.568870884 | 81.2 | Disease_causing |
| testneg_4665 | -0.02 | -0.45 | 3.615951 | -3 | -0.018797 | -3 | 0.73808 | 0.15588 | 0.11891 | 0.11186 | 30.61151 | 0.06 | 0 | Neutral |
| testneg_4666 | -0.07 | -0.38 | 3.896611 | -3 | 2.130388 | -3 | 0.57349 | 0.09764 | 0.19898 | 0.12788 | 14.12935 | 0.060473326 | 0 | Neutral |
| testneg_4667 | 0.16 | 0.08 | 4.242507 | -3 | 5.851562 | -3 | 0.11329 | 0.12116 | 0.25485 | 0.12788 | 20.72912 | 0.286658473 | 3.8 | Neutral |
| testneg_4668 | 0 | 0 | -0.557277 | -3 | 1.882474 | -3 | 0.559 | 0.2286 | 0.45354 | 0.09957 | 11.47365 | 0.02 | 0 | Neutral |
| testneg_4669 | 0 | 0 | -0.284617 | -3 | -2.985743 | -3 | 0.6975 | 0.53794 | 0.55367 | 0.57993 | 4.44444 | 0.073333333 | 0 | Neutral |
| testneg_4670 | 0.03 | 1.22 | 0.647985 | -3 | 4.257584 | -3 | 0.21728 | 0.1132 | 0.05005 | 0.07051 | 28.47222 | 0.114428419 | 0 | Neutral |
| testneg_4671 | 0.01 | -1.44 | 6.204767 | -3 | 1.334738 | -3 | 0.28532 | 0.10663 | 0.98238 | 0.10873 | 10.2717 | 0.21 | 0.6 | Neutral |
| testneg_4672 | 0 | 0 | 0.018114 | -3 | 3.355037 | -3 | 0.21884 | 0.16743 | 0.17443 | 0.19181 | 10.41667 | 0.249793317 | 0.8 | Neutral |
| testneg_4673 | 0.14 | -1.01 | 6.561184 | -3 | -0.454393 | -1.150939 | 0.46056 | 0.12785 | 0.03443 | 0.1032 | 11.25 | 0.236388889 | 1.2 | Neutral |
| testneg_4674 | -0.08 | -0.03 | -0.125177 | -3 | 0.514719 | -3 | 0.27308 | 0.12788 | 0.21331 | 0.09513 | 14.16667 | 0.01 | 0 | Neutral |
| testneg_4675 | 0 | 0 | 1.966897 | -3 | -0.687336 | -3 | 0.39262 | 0.12788 | 0.09997 | 0.09515 | 10.83333 | 0.01 | 0 | Neutral |
| testneg_4676 | 0.01 | 0.8 | 2.118637 | -3 | 0.626243 | 0.626243 | 0.19898 | 0.12788 | 0.34412 | 0.08672 | 23.57096 | 0.010676706 | 0 | Neutral |
| testneg_4677 | -0.08 | -0.03 | 1.155408 | -3 | 1.357664 | 1.357664 | 0.04161 | 0.12788 | 0.1798 | 0.0954 | 18.74088 | 0.06 | 0 | Neutral |
| testneg_4678 | -0.04 | -2.01 | 3.858825 | -3 | -0.727739 | -3 | 0.51398 | 0.38821 | 0.19898 | 0.08223 | 16.55371 | 0.12 | 0 | Neutral |
| testneg_4679 | -0.15 | -0.06 | 2.266671 | -3 | 2.854385 | -3 | 0.09058 | 0.50106 | 0.2218 | 0.11533 | 25.87147 | 0.212047009 | 0 | Neutral |
| testneg_4680 | 0 | 0 | 1.887682 | -3 | 1.056 | 1.056 | 0.18282 | 0.12022 | 0.21153 | 0.11357 | 10.83333 | 0.01 | 0 | Neutral |
| testneg_4681 | 0.15 | 0.04 | 0.346819 | -3 | 3.835265 | -3 | 0.76942 | 0.13267 | 0.76186 | 0.10052 | 11.875 | 0.14 | 0 | Neutral |
| testneg_4682 | 0.15 | -0.82 | 2.20332 | -3 | 0.388086 | -3 | 0.09259 | 0.11165 | 0.15432 | 0.095 | 17.77778 | 0.01 | 0 | Neutral |
| testneg_4683 | 0.01018182 | 0.08981818 | 10.879439 | -3 | 0.332188 | -3 | 0.02146 | 0.10351 | 0.13974 | 0.10225 | 18.5792 | 0.34 | 6.2 | Neutral |
| testneg_4684 | 0.21 | 0.15 | 2.804277 | -3 | 1.922434 | -3 | 0.02124 | 0.05186 | 0.59342 | 0.12788 | 21.52409 | 0.0755 | 0 | Neutral |
| testneg_4685 | 0.02 | -0.68 | -0.505763 | -3 | 0.436559 | -3 | 0.11585 | 0.12788 | 0.14638 | 0.12788 | 19.5422 | 0.01 | 0 | Neutral |
| testneg_4686 | -0.01 | -0.79 | 1.315391 | -3 | 8.23651 | -3 | 0.99417 | 0.17365 | 0.07071 | 0.12788 | 18.33333 | 0.399615385 | 19.6 | Neutral |
| testneg_4687 | -0.04 | 0.23 | 2.067026 | -3 | -1.030525 | -3 | 0.19898 | 0.40608 | 0.07887 | 0.07487 | 18.15705 | 0.05 | 0 | Neutral |
| testneg_4688 | 0 | 0 | 0.356924 | -3 | 4.492963 | -3 | 0.3889 | 0.10831 | 0.54648 | 0.19042 | 7.10199 | 0.156267813 | 0 | Neutral |
| testneg_4689 | -0.06 | 0.57 | 1.26369 | -3 | 0.94538 | -3 | 0.19898 | 0.12788 | 0.0328 | 0.0691 | 30.93079 | 0.02 | 0 | Neutral |
| testneg_4690 | 0 | 0 | 1.048155 | -3 | 5.723508 | -3 | 0.42523 | 0.13467 | 0.10893 | 0.12072 | 18.33333 | 0.109444444 | 0 | Neutral |
| testneg_4691 | 0.14 | -0.37 | 1.800547 | -3 | 4.959501 | -3 | 0.12871 | 0.29252 | 0.14301 | 0.12788 | 17.21847 | 0.276607143 | 1 | Neutral |
| testneg_4692 | -0.01 | -0.8 | 7.657206 | -3 | 0.259095 | -3 | 0.19898 | 0.12788 | 0.99994 | 0.51636 | 10.41667 | 0.404472222 | 25 | Neutral |
| testneg_4693 | -0.04 | 0.23 | -2.175398 | -3 | 6.071016 | -3 | 0.10228 | 0.22182 | 0.24791 | 0.09675 | 21.40097 | 0.312142857 | 2.2 | Neutral |
| testneg_4694 | -0.01 | 0.57 | 0.47202 | 0.47202 | 1.27985 | -3 | 0.13295 | 0.12914 | 0.09334 | 0.10027 | 23.68839 | 0.03 | 0 | Neutral |
| testneg_4695 | 0 | 0 | 0.059973 | -3 | 0.567562 | -3 | 0.21095 | 0.10574 | 0.17046 | 0.21736 | 15.89191 | 0.020708658 | 0 | Neutral |
| testneg_4696 | 0 | -0.01 | 0.558245 | 0.558245 | 2.027702 | -3 | 0.19898 | 0.10362 | 0.55155 | 0.10709 | 20.74751 | 0.02 | 0 | Neutral |
| testneg_4697 | 0 | 0 | 0.477103 | -3 | 4.845098 | -3 | 0.64072 | 0.12788 | 0.09093 | 0.28643 | 35.83104 | 0.298191129 | 1.2 | Neutral |
| testneg_4698 | -0.03 | -0.1 | 6.036832 | -3 | -0.505763 | -3 | 0.16826 | 0.11262 | 0.11585 | 0.12788 | 19.43262 | 0.17 | 0.2 | Neutral |
| testneg_4699 | 0.09 | -0.3 | 6.596925 | -3 | 6.085767 | -3 | 0.19898 | 0.12788 | 0.19898 | 0.12788 | 10.30706 | 0.641666667 | 89.2 | Disease_causing |
| testneg_4700 | 0.03 | -1.03 | 4.715557 | -3 | 1.222548 | 1.222548 | 0.04243 | 0.22343 | 0.06705 | 0.12788 | 23.77111 | 0.180316537 | 0 | Neutral |
| testneg_4701 | 0 | 0 | -0.456898 | -3 | 2.231851 | -3 | 0.06397 | 0.12788 | 0.13521 | 0.10126 | 33.95189 | 0.02 | 0 | Neutral |
| testneg_4702 | 0.18 | -1.05 | 4.095375 | -3 | 5.694121 | -3 | 0.1589 | 0.10761 | 0.11042 | 0.09881 | 23.01096 | 0.226666667 | 0.6 | Neutral |
| testneg_4703 | -0.08 | -0.03 | 0.083519 | -3 | 0.500506 | -3 | 0.18488 | 0.53379 | 0.68233 | 0.58998 | 4.72222 | 0.1609375 | 0 | Neutral |
| testneg_4704 | -0.15 | 0.28 | -2.193108 | -3 | 4.83533 | -3 | 0.06996 | 0.35923 | 0.23975 | 0.13861 | 10.83333 | 0.610271046 | 90.6 | Disease_causing |
| testneg_4705 | 0.05 | 0.29 | 0.42015 | -3 | 2.667661 | -3 | 0.02844 | 0.12788 | 0.10498 | 0.08421 | 17.21847 | 0.055560606 | 0 | Neutral |
| testneg_4706 | 0.22 | -2.66 | 2.450036 | -3 | 3.907702 | -3 | 0.09997 | 0.1797 | 0.14498 | 0.33896 | 12.6082 | 0.633569843 | 95.2 | Disease_causing |
| testneg_4707 | -0.01 | 0.57 | 2.71072 | -3 | 5.244539 | -3 | 0.09719 | 0.09576 | 0.21289 | 0.1079 | 17.85763 | 0.153333333 | 0 | Neutral |
| testneg_4708 | 0.22 | -2.66 | 5.821758 | -3 | 1.747261 | 1.747261 | 0.11359 | 0.08631 | 0.17736 | 0.14772 | 20.5518 | 0.289337607 | 2 | Neutral |
| testneg_4709 | 0 | 0 | 1.012321 | -3 | -0.381312 | -2.299248 | 0.21805 | 0.12777 | 0.12173 | 0.12788 | 23.09941 | 0.0005209 | 0 | Neutral |
| testneg_4710 | 0 | 0 | 0.197366 | -3 | 1.795813 | -3 | 0.95978 | 0.09493 | 0.28199 | 0.10688 | 22.80197 | 0.020111125 | 0 | Neutral |
| testneg_4711 | 0 | 0 | 0.996313 | -3 | 2.191197 | -3 | 0.25744 | 0.2162 | 0.18667 | 0.10488 | 15 | 0.02 | 0 | Neutral |
| testneg_4712 | 0.06 | -0.57 | 0.237935 | -3 | 0.727897 | -3 | 0.16362 | 0.11879 | 0.34209 | 0.09515 | 20.30197 | 0.004217687 | 0 | Neutral |
| testneg_4713 | 0.01 | 0.79 | -2.550688 | -3 | -0.462963 | -0.462963 | 0.19898 | 0.12788 | 0.1946 | 0.14591 | 20.5518 | 0.06 | 0 | Neutral |
| testneg_4714 | -0.07 | -0.38 | 3.331643 | 3.331643 | 1.606239 | -3 | 0.26337 | 0.12771 | 0.89002 | 0.12788 | 10.41667 | 0.43 | 23.8 | Neutral |
| testneg_4715 | 0.18 | -1.05 | 5.492254 | -3 | -0.102803 | -3 | 0.28063 | 0.40063 | 0.1134 | 0.12788 | 29.16667 | 0.25 | 0 | Neutral |
| testneg_4716 | 0.16 | -0.37 | -2.713571 | -2.713571 | 0.277234 | -3 | 0.07506 | 0.09112 | 0.18559 | 0.09173 | 35.47861 | 0.03 | 0 | Neutral |
| testneg_4717 | -0.17 | 1.22 | -2.338688 | -2.338688 | 0.120012 | -3 | 0.33129 | 0.09957 | 0.20491 | 0.22087 | 18.88889 | 0.15 | 0 | Neutral |
| testneg_4718 | 0.07 | 0.01 | 0.631605 | -3 | 3.635979 | -3 | 0.14524 | 0.10162 | 0.35779 | 0.12788 | 29.86111 | 0.05 | 0 | Neutral |
| testneg_4719 | -0.06 | 0.57 | 1.042833 | -3 | 1.036843 | -3 | 0.19898 | 0.12788 | 0.18102 | 0.6537 | 18.88889 | 0.079311236 | 0 | Neutral |
| testneg_4720 | 0.14 | -0.37 | 3.348627 | -3 | -0.211546 | -3 | 0.04808 | 0.10018 | 0.2442 | 0.11401 | 13.2927 | 0.05 | 0 | Neutral |
| testneg_4721 | 0 | 0 | -0.011643 | -3 | 5.835814 | -3 | 0.30823 | 0.17188 | 0.505 | 0.12788 | 19.44444 | 0.141526089 | 0 | Neutral |
| testneg_4722 | 0.19 | -0.25 | 3.963694 | -3 | 4.253999 | -3 | 0.42512 | 0.29333 | 0.09807 | 0.24901 | 12.19153 | 0.785973939 | 99.8 | Disease_causing |
| testneg_4723 | 0.04 | -0.23 | -1.047288 | -3 | -0.431527 | -3 | 0.52005 | 0.12788 | 0.19772 | 0.09261 | 19.23479 | 0.010392308 | 0 | Neutral |
| testneg_4724 | -0.01 | -0.79 | 0.452127 | -3 | 1.028858 | -3 | 0.24791 | 0.09675 | 0.16973 | 0.17555 | 10.12505 | 0.05 | 0 | Neutral |
| testneg_4725 | 0 | 0 | 1.550971 | -3 | -0.437663 | -3 | 0.73311 | 0.10866 | 0.34946 | 0.19092 | 10.41667 | 0.011297953 | 0 | Neutral |
| testneg_4726 | 0.01 | 0.79 | 0.259266 | -3 | 0.214112 | -3 | 0.32804 | 0.10857 | 0.17301 | 0.14089 | 11.25 | 0.050311594 | 0 | Neutral |
| testneg_4727 | 0.03 | -0.85 | -0.678462 | -3 | 0.763947 | -3 | 0.67216 | 0.17808 | 0.7413 | 0.18974 | 11.25 | 0.071606206 | 0 | Neutral |
| testneg_4728 | 0.04 | 2.01 | 5.43333 | -3 | -0.335117 | -3 | 0.15704 | 0.72174 | 0.49278 | 0.09047 | 11.92061 | 0.290814176 | 0.4 | Neutral |
| testneg_4729 | 0 | 0 | 0.563638 | -3 | 2.136098 | -3 | 0.99998 | 0.9588 | 0.13374 | 0.12788 | 10.41667 | 0.1107125 | 0 | Neutral |
| testneg_4730 | -0.07 | -0.38 | -0.872044 | -3 | 3.861717 | -3 | 0.04792 | 0.12788 | 0.58292 | 0.87725 | 17.96787 | 0.357404762 | 4.2 | Neutral |
| testneg_4731 | -0.04 | -2.01 | 2.099311 | -3 | 2.003603 | -3 | 0.98565 | 0.23629 | 0.24544 | 0.0894 | 20.36321 | 0.09 | 0 | Neutral |
| testneg_4732 | -0.07 | -0.33 | 0.406963 | -3 | 4.492087 | -3 | 0.48825 | 0.10951 | 0.37424 | 0.12788 | 18.88889 | 0.08 | 0 | Neutral |
| testneg_4733 | 0 | -0.01 | 2.053357 | -3 | 0.823242 | -3 | 0.22761 | 0.14229 | 0.10423 | 0.12788 | 18.13813 | 0.01 | 0 | Neutral |
| testneg_4734 | -0.18 | 1.05 | 1.550311 | -3 | 0.071635 | -3 | 0.17443 | 0.19181 | 0.23939 | 0.25284 | 18.33333 | 0.201833784 | 0 | Neutral |
| testneg_4735 | 0 | -0.01 | 0.664208 | -3 | 1.11197 | -3 | 0.20673 | 0.13236 | 0.12231 | 0.23155 | 10.30286 | 0.047789492 | 0 | Neutral |
| testneg_4736 | 0.07 | 0.38 | 0.878803 | -3 | 3.68445 | 2.010216 | 0.07837 | 0.12788 | 0.4069 | 0.3841 | 18.50844 | 0.352748237 | 7 | Neutral |
| testneg_4737 | 0 | 0 | 2.013802 | -3 | 1.655297 | -3 | 0.22239 | 0.12885 | 0.09515 | 0.07784 | 12.39396 | 0.000801282 | 0 | Neutral |
| testneg_4738 | 0.06 | -0.57 | 5.346373 | -3 | 0.024873 | -3 | 0.30669 | 0.11489 | 0.16507 | 0.10992 | 13.74795 | 0.08 | 0 | Neutral |
| testneg_4739 | 0.08 | 0.03 | 1.182833 | -3 | 4.013472 | -3 | 0.89661 | 0.17382 | 0.04983 | 0.08759 | 10.41667 | 0.149113585 | 0 | Neutral |
| testneg_4740 | 0.22 | -2.66 | 4.250235 | -3 | 4.388483 | -3 | 0.60412 | 0.13303 | 0.17264 | 0.09426 | 17.77778 | 0.290496032 | 0.8 | Neutral |
| testneg_4741 | 0 | 0 | -0.381881 | -1.200815 | -0.01248 | -3 | 0.28817 | 0.10531 | 0.61231 | 0.13362 | 23.04755 | 0.06 | 0 | Neutral |
| testneg_4742 | 0.21 | 0.15 | 4.012434 | -3 | 2.022528 | -3 | 0.28629 | 0.77234 | 0.57953 | 0.15513 | 13.60507 | 0.426335604 | 28.4 | Neutral |
| testneg_4743 | 0 | 0 | 2.049487 | -3 | -0.05178 | -3 | 0.54349 | 0.60885 | 0.19898 | 0.12788 | 16.8018 | 0.030247253 | 0 | Neutral |
| testneg_4744 | 0 | 0 | -0.038389 | -3 | 2.263609 | -3 | 0.16326 | 0.09256 | 0.4811 | 0.13398 | 18.6768 | 0.070178628 | 0 | Neutral |
| testneg_4745 | -0.07 | -0.38 | 1.941375 | -3 | 2.36406 | -3 | 0.23962 | 0.10029 | 0.10395 | 0.08737 | 13.02635 | 0.03 | 0 | Neutral |
| testneg_4746 | -0.03 | 1.03 | 1.03165 | -3 | 3.929013 | -3 | 0.17873 | 0.12378 | 0.03558 | 0.05202 | 23.82599 | 0.111162769 | 0 | Neutral |
| testneg_4747 | 0 | 0 | 2.064873 | 1.683039 | 0.488014 | -3 | 0.65564 | 0.11468 | 0.62339 | 0.19592 | 10.83333 | 0.1 | 0 | Neutral |
| testneg_4748 | -0.05 | 0.81 | -0.02606 | -3 | 0.557312 | -3 | 0.9327 | 0.12788 | 0.23308 | 0.77395 | 9.99261 | 0.090043103 | 0 | Neutral |
| testneg_4749 | 0.1 | 1.9 | 2.756314 | -3 | 2.410627 | -3 | 0.29044 | 0.12142 | 0.07275 | 0.12788 | 28.72196 | 0.1 | 0 | Neutral |
| testneg_4750 | -0.02 | -0.41 | 0.433545 | -3 | -0.673818 | -3 | 0.06996 | 0.35923 | 0.76138 | 0.16258 | 13.80486 | 0.19 | 0 | Neutral |
| testneg_4751 | 0.14 | -0.37 | -0.303574 | -3 | -1.030525 | -3 | 0.04346 | 0.10785 | 0.07887 | 0.07487 | 26.15385 | 0.048333333 | 0 | Neutral |
| testneg_4752 | 0 | 0 | 0.758635 | -3 | 2.144749 | -3 | 0.19898 | 0.09822 | 0.07071 | 0.11499 | 19.09347 | 0.000709256 | 0 | Neutral |
| testneg_4753 | 0.21 | 0.15 | 4.012434 | -3 | 2.171434 | -3 | 0.28629 | 0.77234 | 0.49425 | 0.25538 | 4.72222 | 0.564754386 | 78.4 | Disease_causing |
| testneg_4754 | -0.07 | -0.33 | 1.125228 | -3 | 4.792565 | -3 | 0.15808 | 0.14548 | 0.15432 | 0.095 | 10 | 0.139078431 | 0 | Neutral |
| testneg_4755 | -0.01 | -0.79 | 2.218156 | -3 | 3.017549 | -3 | 0.37327 | 0.2201 | 0.23385 | 0.12788 | 10 | 0.109928161 | 0 | Neutral |
| testneg_4756 | 0 | 0 | 1.930844 | -3 | -0.408308 | -3 | 0.74136 | 0.1049 | 0.28165 | 0.15912 | 18.39615 | 0.05 | 0 | Neutral |
| testneg_4757 | -0.16 | -0.08 | 1.042194 | -3 | 0.840944 | -3 | 0.19898 | 0.12788 | 0.03505 | 0.0907 | 23.68839 | 0.05 | 0 | Neutral |

| | | | | | | | | | | | | | | |
|---|---|---|---|---|---|---|---|---|---|---|---|---|---|---|
| testneg_4758 | 0 | 0 | 1.038487 | -3 | 2.043452 | -3 | 0.72124 | 0.11262 | 0.13011 | 0.14157 | 8.97898 | 0.033838235 | 0 | Neutral |
| testneg_4759 | 0 | 0 | -0.674729 | -3 | 1.228976 | -1.950188 | 0.24765 | 0.32858 | 0.09273 | 0.08316 | 18.6768 | 0.03 | 0 | Neutral |
| testneg_4760 | 0.08 | 0.03 | 0.23302 | -3 | 4.283158 | -3 | 0.1653 | 0.0977 | 0.08529 | 0.12376 | 18.03571 | 0.071666667 | 0 | Neutral |
| testneg_4761 | -0.12 | 1.16 | 6.464782 | -3 | 6.279647 | -3 | 0.98833 | 0.48581 | 0.15856 | 0.10994 | 12.29167 | 0.77 | 99.8 | Disease_causing |
| testneg_4762 | 0 | 0 | -0.136916 | -3 | 2.403577 | -3 | 0.04853 | 0.4186 | 0.99417 | 0.17365 | 19.09347 | 0.321838054 | 2.8 | Neutral |
| testneg_4763 | 0.18 | -1.05 | 7.032758 | -3 | 3.568323 | -3 | 0.07428 | 0.07645 | 0.63798 | 0.19967 | 24.14414 | 0.5 | 54.6 | Disease_causing |
| testneg_4764 | 0 | -0.32 | 2.305189 | -3 | 1.361303 | -3 | 0.505 | 0.12788 | 0.12855 | 0.11656 | 19.16667 | 0.010443501 | 0 | Neutral |
| testneg_4765 | 0 | 0 | -0.22828 | -3 | 3.422566 | -3 | 0.31694 | 0.13475 | 0.16462 | 0.18467 | 6.2037 | 0.203741901 | 0.4 | Neutral |
| testneg_4766 | -0.16 | -0.08 | 2.98751 | -3 | 2.212728 | -3 | 0.18022 | 0.12788 | 0.06541 | 0.12012 | 30.83804 | 0.09 | 0 | Neutral |
| testneg_4767 | 0.04 | -0.23 | 2.30123 | -3 | 3.045525 | -3 | 0.23152 | 0.09962 | 0.64775 | 0.54233 | 12.6082 | 0.225513309 | 0.6 | Neutral |
| testneg_4768 | 0 | 0 | 1.683026 | 1.683026 | 2.998118 | -3 | 0.84106 | 0.63405 | 0.143 | 0.11236 | 9.01526 | 0.082292243 | 0 | Neutral |
| testneg_4769 | 0.07 | 0.38 | 0.34268 | -3 | 0.201766 | -3 | 0.22794 | 0.10688 | 0.16388 | 0.14439 | 36.11111 | 0.053938618 | 0 | Neutral |
| testneg_4770 | 0.07 | 0.38 | -2.242853 | -3 | 2.729204 | -3 | 0.06358 | 0.5364 | 0.62339 | 0.19592 | 19.44444 | 0.404102564 | 15.2 | Neutral |
| testneg_4771 | 0.07 | 0.7 | 5.661416 | -3 | 1.986841 | 1.986841 | 0.9352 | 0.14229 | 0.19898 | 0.12788 | 17.28868 | 0.289048826 | 2.2 | Neutral |
| testneg_4772 | 0 | 0 | -0.401618 | -3 | 5.108216 | -3 | 0.59853 | 0.75237 | 0.07912 | 0.09055 | 12.7356 | 0.120833333 | 0 | Neutral |
| testneg_4773 | -0.14 | 0.37 | 2.573998 | -3 | 3.538232 | -3 | 0.12073 | 0.10969 | 0.10317 | 0.14954 | 18.33333 | 0.31 | 3.2 | Neutral |
| testneg_4774 | 0 | 0 | -0.001679 | -3 | 1.987946 | -3 | 0.70964 | 0.12663 | 0.16662 | 0.11306 | 29.59021 | 0.02 | 0 | Neutral |
| testneg_4775 | -0.19 | 0.27 | 1.951681 | -3 | 6.886079 | -3 | 0.1875 | 0.10316 | 0.09522 | 0.20323 | 33.69565 | 0.520714286 | 62.8 | Disease_causing |
| testneg_4776 | -0.08 | -0.03 | 5.10743 | -3 | 6.267382 | -3 | 0.13067 | 0.1005 | 0.28211 | 0.25332 | 26.84685 | 0.709761905 | 99 | Disease_causing |
| testneg_4777 | 0 | 0 | -0.262607 | -3 | 0.823665 | -3 | 0.21576 | 0.17241 | 0.27915 | 0.27706 | 10.83333 | 0.056389752 | 0 | Neutral |
| testneg_4778 | 0.01 | 0.79 | 0.585649 | 0.585649 | 0.063598 | -3 | 0.07185 | 0.68732 | 0.1201 | 0.1062 | 18.33333 | 0.113205128 | 0 | Neutral |
| testneg_4779 | 0 | 0 | 1.26254 | -3 | 4.845396 | -3 | 0.35189 | 0.12788 | 0.17231 | 0.1358 | 19.17614 | 0.224286875 | 0 | Neutral |
| testneg_4780 | 0 | -0.01 | 1.202769 | -3 | -0.307406 | -3 | 0.20231 | 0.12788 | 0.62858 | 0.16645 | 23.09286 | 0.05 | 0 | Neutral |
| testneg_4781 | 0.02 | -0.63 | 3.104093 | -3 | 1.421274 | -3 | 0.65564 | 0.11468 | 0.49745 | 0.1259 | 20 | 0.04 | 0 | Neutral |
| testneg_4782 | 0 | 0 | -0.5391 | -3 | 2.157433 | -3 | 0.16418 | 0.12788 | 0.84318 | 0.11313 | 16.8018 | 0.01 | 0 | Neutral |
| testneg_4783 | -0.07 | -0.38 | 1.102328 | -3 | 1.820578 | -3 | 0.74271 | 0.10951 | 0.31507 | 0.61057 | 12.40248 | 0.060366667 | 0 | Neutral |
| testneg_4784 | 0.19 | -0.27 | 6.763226 | -3 | 3.146117 | -3 | 0.63818 | 0.12788 | 0.93246 | 0.12788 | 11.25 | 0.418824405 | 22.6 | Neutral |
| testneg_4785 | 0 | 0 | 1.686834 | -3 | 5.826992 | -3 | 0.29828 | 0.12788 | 0.20084 | 0.0851 | 17.36842 | 0.06 | 0 | Neutral |
| testneg_4786 | 0.14 | -1.01 | 0.745507 | -3 | -0.360768 | -0.360768 | 0.82126 | 0.17937 | 0.47147 | 0.09503 | 19.66924 | 0.050606618 | 0 | Neutral |
| testneg_4787 | 0 | 0 | 1.992408 | -3 | 4.45544 | -3 | 0.19898 | 0.25991 | 0.16692 | 0.09243 | 19.5422 | 0.151904762 | 0 | Neutral |
| testneg_4788 | -0.07 | -0.38 | 2.555278 | -3 | 0.769221 | -3 | 0.21814 | 0.13855 | 0.07297 | 0.12788 | 21.79342 | 0.020315558 | 0 | Neutral |
| testneg_4789 | 0.04 | -0.89 | -0.197119 | -3 | 1.666522 | -3 | 0.28191 | 0.20693 | 0.19883 | 0.09434 | 16.25133 | 0.04125 | 0 | Neutral |
| testneg_4790 | 0 | 0 | 0.944979 | -3 | 1.326802 | -3 | 0.32022 | 0.09529 | 0.05183 | 0.18779 | 28.78281 | 0.080804732 | 0 | Neutral |
| testneg_4791 | 0.03 | 0.12 | 2.628269 | -3 | 1.988228 | -3 | 0.18584 | 0.11114 | 0.18504 | 0.0822 | 19.66924 | 0.010168852 | 0 | Neutral |
| testneg_4792 | 0 | 0 | 0.628682 | -3 | 0.425569 | -3 | 0.16404 | 0.09378 | 0.12879 | 0.12788 | 27.0922 | 0.01 | 0 | Neutral |
| testneg_4793 | 0.04 | -0.23 | 6.319089 | -3 | 2.176387 | -3 | 0.19898 | 0.12788 | 0.72068 | 0.99841 | 14.37767 | 0.46587037 | 42.8 | Neutral |
| testneg_4794 | 0 | 0 | 3.466764 | -3 | 10.298141 | -3 | 0.34205 | 0.11195 | 0.14704 | 0.12788 | 19.09347 | 0.46 | 39.2 | Neutral |
| testneg_4795 | 0 | 0 | 4.43194 | -3 | 3.760866 | -3 | 0.36829 | 0.11457 | 0.75033 | 0.64204 | 10.41667 | 0.436655023 | 29.4 | Neutral |
| testneg_4796 | 0.03 | 0.1 | -1.156564 | -3 | -0.285087 | -3 | 0.54213 | 0.09044 | 0.09522 | 0.20323 | 13.51351 | 0.084769841 | 0 | Neutral |
| testneg_4797 | -0.16 | -0.08 | 1.555057 | -3 | 1.986323 | -3 | 0.06187 | 0.36607 | 0.19898 | 0.12788 | 10 | 0.2 | 0 | Neutral |
| testneg_4798 | 0.07 | 0.7 | 2.703607 | -3 | 1.189051 | -3 | 0.11585 | 0.12788 | 0.19227 | 0.1096 | 27.27735 | 0.02 | 0 | Neutral |
| testneg_4799 | -0.07 | -0.38 | 1.30006 | -3 | 3.006651 | -3 | 0.13704 | 0.10054 | 0.03441 | 0.10663 | 18.6768 | 0.04 | 0 | Neutral |
| testneg_4800 | 0.05 | -0.81 | 1.982936 | -3 | 2.729304 | -3 | 0.07044 | 0.11679 | 0.0917 | 0.13124 | 29.86111 | 0.082930162 | 0 | Neutral |
| testneg_4801 | 0 | 0.73 | 5.650938 | -3 | 0.309305 | -3 | 0.16671 | 0.18311 | 0.16973 | 0.17555 | 14.47674 | 0.47 | 46.6 | Neutral |
| testneg_4802 | -0.15 | -0.04 | 0.939929 | -3 | 2.900481 | -3 | 0.18701 | 0.12788 | 0.33992 | 0.15484 | 20.21004 | 0.187398649 | 0 | Neutral |
| testneg_4803 | -0.07 | -0.01 | -0.919759 | -3 | -2.200892 | -3 | 0.24791 | 0.09675 | 0.17818 | 0.15728 | 19.76246 | 0.07 | 0 | Neutral |
| testneg_4804 | 0 | 0 | -0.272804 | -0.272804 | 2.158998 | -3 | 0.17839 | 0.12009 | 0.81659 | 0.10424 | 17.68062 | 0.01 | 0 | Neutral |
| testneg_4805 | -0.03 | -0.1 | 1.285186 | 1.285186 | -1.970817 | -3 | 0.35957 | 0.11476 | 0.2058 | 0.28174 | 12.4985 | 0.111792082 | 0 | Neutral |
| testneg_4806 | 0.15 | 0.04 | -0.646831 | -3 | 1.559025 | -3 | 0.20849 | 0.12788 | 0.19898 | 0.12788 | 29.34685 | 0.02 | 0 | Neutral |
| testneg_4807 | -0.02 | 0.36 | 0.523312 | -3 | -1.024972 | -3 | 0.41987 | 0.67373 | 0.23955 | 0.97212 | 4.72222 | 0.171133381 | 0 | Neutral |
| testneg_4808 | 0 | 0 | 1.492291 | -3 | 0.867971 | -3 | 0.11791 | 0.39607 | 0.24966 | 0.12788 | 18.59326 | 0.030409699 | 0 | Neutral |
| testneg_4809 | 0 | 0 | 0.194425 | -3 | 0.567562 | -3 | 0.19898 | 0.10106 | 0.17046 | 0.21736 | 30.0251 | 0.05 | 0 | Neutral |
| testneg_4810 | 0.14 | -0.69 | 1.36796 | -3 | 4.858929 | 2.0376 | 0.38719 | 0.15548 | 0.82148 | 0.14847 | 17.77778 | 0.409544512 | 19.8 | Neutral |
| testneg_4811 | 0.03 | 1.22 | 2.69903 | -3 | -0.729977 | -0.729977 | 0.1192 | 0.10391 | 0.17974 | 0.12788 | 16.42511 | 0.05 | 0 | Neutral |
| testneg_4812 | 0.14 | -1.01 | 3.57776 | -3 | -0.056225 | -3 | 0.63431 | 0.09837 | 0.08059 | 0.09576 | 37.80687 | 0.05 | 0 | Neutral |
| testneg_4813 | 0 | 0 | 1.548509 | -3 | -1.164458 | -3 | 0.06711 | 0.09765 | 0.08183 | 0.11603 | 7.40923 | 0.000926037 | 0 | Neutral |
| testneg_4814 | 0 | -0.01 | -1.288398 | -3 | -1.624227 | -1.624227 | 0.19898 | 0.12788 | 0.19237 | 0.53191 | 19.09347 | 0.060527778 | 0 | Neutral |
| testneg_4815 | -0.1 | -1.9 | 2.134495 | 1.517891 | 4.105708 | -3 | 0.42755 | 0.09484 | 0.33245 | 0.45921 | 11.25 | 0.442418762 | 33.4 | Neutral |
| testneg_4816 | 0 | 0 | 0.596959 | -3 | 1.053971 | -3 | 0.18925 | 0.32479 | 0.19898 | 0.12788 | 28.51351 | 0.03 | 0 | Neutral |
| testneg_4817 | 0.07 | 0.38 | 0.034534 | 0.034534 | 4.28255 | -3 | 0.62381 | 0.9477 | 0.45075 | 0.12232 | 12.40248 | 0.22 | 0 | Neutral |
| testneg_4818 | 0 | 0 | 1.190149 | -3 | 1.772012 | -3 | 0.16841 | 0.11143 | 0.26731 | 0.16415 | 21.49575 | 0.050877416 | 0 | Neutral |
| testneg_4819 | -0.03 | -0.1 | 0.545839 | -3 | -0.40895 | -3 | 0.58545 | 0.14414 | 0.15646 | 0.12788 | 18.89436 | 0.02 | 0 | Neutral |
| testneg_4820 | -0.15 | -0.04 | 3.653806 | -3 | -0.470915 | -3 | 0.10371 | 0.08655 | 0.42836 | 0.12788 | 16.8018 | 0.08 | 0 | Neutral |
| testneg_4821 | 0.07 | 0.33 | 1.750479 | -3 | 2.562908 | 0.984611 | 0.15767 | 0.12788 | 0.82148 | 0.14847 | 16.8018 | 0.090210859 | 0 | Neutral |
| testneg_4822 | -0.02 | -0.45 | 5.92982 | -3 | 7.007099 | -3 | 0.99948 | 0.5595 | 0.52836 | 0.12785 | 7.66785 | 0.8 | 99.6 | Disease_causing |
| testneg_4823 | 0 | -0.01 | 1.723095 | 1.723095 | 4.479766 | -3 | 0.91293 | 0.12788 | 0.1294 | 0.10964 | 12.12667 | 0.11 | 0 | Neutral |
| testneg_4824 | -0.0043357 | 0.02363636 | -0.760295 | -3 | 0.07066 | -3 | 0.33082 | 0.12788 | 0.08903 | 0.12788 | 23.13568 | 0.020755656 | 0 | Neutral |
| testneg_4825 | 0.2 | -1.26 | 5.986437 | -3 | 1.624942 | 1.624942 | 0.12325 | 0.09383 | 0.16208 | 0.10597 | 20.83333 | 0.15984375 | 0 | Neutral |
| testneg_4826 | 0.02 | -0.68 | 1.270761 | -3 | -1.873728 | -1.873728 | 0.24144 | 0.12122 | 0.19898 | 0.12788 | 39.15938 | 0.01 | 0 | Neutral |
| testneg_4827 | 0.17 | 0.38 | 3.129469 | -3 | 1.782334 | -3 | 0.17075 | 0.12788 | 0.79857 | 0.12788 | 21.27435 | 0.05 | 0 | Neutral |
| testneg_4828 | 0 | 0 | 0.159372 | 0.159372 | 6.416133 | -3 | 0.05228 | 0.12788 | 0.18499 | 0.10583 | 32.40433 | 0.161904762 | 0 | Neutral |
| testneg_4829 | 0.09125 | -0.2675 | 10.838582 | -3 | 1.344374 | -3 | 0.29478 | 0.12788 | 0.34109 | 0.2167 | 12.33727 | 0.548805704 | 73.8 | Disease_causing |
| testneg_4830 | -0.06 | -0.18 | 4.170859 | -3 | 2.798182 | 2.798182 | 0.09998 | 0.09022 | 0.31043 | 0.09688 | 17.01711 | 0.19 | 0 | Neutral |
| testneg_4831 | 0 | 0 | 1.39998 | -3 | -0.186131 | -3 | 0.161 | 0.11597 | 0.11802 | 0.09304 | 18.70407 | 0 | 0 | Neutral |
| testneg_4832 | 0 | 0 | 0.389761 | -3 | 4.866108 | -3 | 0.73601 | 0.26791 | 0.19898 | 0.12788 | 40.2439 | 0.107557124 | 0 | Neutral |
| testneg_4833 | 0 | 0 | 0.988618 | -3 | 0.659218 | -3 | 0.15019 | 0.10342 | 0.44429 | 0.20468 | 12.81915 | 0.02 | 0 | Neutral |
| testneg_4834 | 0 | 0 | 1.252902 | -3 | 0.259883 | -3 | 0.6957 | 0.38769 | 0.56204 | 0.10762 | 10 | 0.01 | 0 | Neutral |
| testneg_4835 | 0 | 0 | 0.089128 | -3 | 1.960992 | -3 | 0.34495 | 0.16861 | 0.15486 | 0.28768 | 10.83333 | 0.073589912 | 0 | Neutral |
| testneg_4836 | 0 | 0 | 0.785121 | -3 | 2.436626 | -3 | 0.10004 | 0.12788 | 0.26181 | 0.12875 | 15.77966 | 0.01 | 0 | Neutral |
| testneg_4837 | 0 | 0 | 0.470493 | -3 | -2.618223 | -3 | 0.42802 | 0.12096 | 0.22179 | 0.09831 | 19.55556 | 0.01 | 0 | Neutral |
| testneg_4838 | 0 | 0 | -0.02198 | -0.02198 | 4.395754 | -3 | 0.15509 | 0.1326 | 0.23879 | 0.12788 | 10.62943 | 0.085810606 | 0 | Neutral |
| testneg_4839 | 0.15 | -0.28 | -0.431211 | -3 | 4.539303 | -0.465307 | 0.36676 | 0.37749 | 0.62808 | 0.25222 | 10 | 0.498004128 | 54.4 | Disease_causing |
| testneg_4840 | 0 | 0 | 1.495836 | -3 | 7.812758 | -3 | 0.24977 | 0.11262 | 0.36486 | 0.14268 | 17.5116 | 0.381951659 | 13.8 | Neutral |
| testneg_4841 | 0 | 0 | 0.984324 | -3 | 2.323036 | -3 | 0.21998 | 0.20297 | 0.65141 | 0.18046 | 10.41667 | 0.119071787 | 0 | Neutral |
| testneg_4842 | 0.14 | -1.01 | 6.843378 | -3 | 2.723672 | -3 | 0.23624 | 0.12788 | 0.13743 | 0.12788 | 12.22222 | 0.41 | 22.6 | Neutral |
| testneg_4843 | -0.02 | -0.13 | 1.290429 | -1.019571 | 0.330554 | -3 | 0.19898 | 0.20319 | 0.06431 | 0.12788 | 30.29126 | 0.030131868 | 0 | Neutral |
| testneg_4844 | -0.03 | -0.1 | 0.907219 | -3 | 0.637213 | -3 | 0.19898 | 0.12788 | 0.19898 | 0.535 | 6.94444 | 0.053265504 | 0 | Neutral |
| testneg_4845 | 0.03 | 1.22 | 2.297873 | -3 | 0.34022 | -3 | 0.31168 | 0.12788 | 0.19898 | 0.12788 | 14.54964 | 0.04 | 0 | Neutral |
| testneg_4846 | 0 | 0 | 0.902021 | -3 | -0.05829 | -3 | 0.37499 | 0.17803 | 0.38115 | 0.21061 | 11.96702 | 0.040450297 | 0 | Neutral |
| testneg_4847 | 0.01 | 0.8 | 3.671807 | -3 | 0.129782 | -3 | 0.17222 | 0.12788 | 0.0882 | 0.12788 | 18.88889 | 0.04 | 0 | Neutral |
| testneg_4848 | 0 | 0 | 0.628811 | -3 | 1.592614 | -3 | 0.07874 | 0.12788 | 0.37637 | 0.12788 | 14.24157 | 0 | 0 | Neutral |
| testneg_4849 | 0.05 | 0.29 | 2.698795 | -3 | 0.877689 | -3 | 0.17806 | 0.12788 | 0.09515 | 0.07784 | 23.68839 | 0.010281746 | 0 | Neutral |
| testneg_4850 | 0.06 | -0.57 | 2.475201 | -3 | 1.749597 | -3 | 0.11264 | 0.11163 | 0.90112 | 0.20622 | 11.23271 | 0.061402585 | 0 | Neutral |
| testneg_4851 | 0.17 | 0.38 | 2.280202 | -0.725688 | 7.530767 | -3 | 0.17285 | 0.09314 | 0.1897 | 0.12459 | 15.55329 | 0.32 | 3.4 | Neutral |
| testneg_4852 | 0.17 | -1.22 | 4.581463 | -3 | 1.512216 | 1.241234 | 0.16741 | 0.10762 | 0.6258 | 0.16461 | 12.07317 | 0.250229034 | 0.2 | Neutral |
| testneg_4853 | -0.05 | -0.29 | -1.760208 | -3 | 4.969986 | -3 | 0.14632 | 0.12662 | 0.49831 | 0.29427 | 34.45035 | 0.328150585 | 4.2 | Neutral |
| testneg_4854 | 0.17 | 0.38 | 1.490909 | -3 | 0.538945 | -3 | 0.90259 | 0.31502 | 0.11746 | 0.1471 | 12.29167 | 0.164627578 | 0 | Neutral |
| testneg_4855 | -0.18 | 1.05 | 3.652922 | -3 | 0.22112 | -3 | 0.39344 | 0.12788 | 0.75742 | 0.16306 | 12.70833 | 0.178558693 | 0 | Neutral |
| testneg_4856 | 0.01 | 0.8 | 2.947434 | -3 | 6.230182 | -3 | 0.27915 | 0.27706 | 0.08268 | 0.09939 | 12.19407 | 0.401395833 | 19.4 | Neutral |
| testneg_4857 | 0 | -0.01 | 1.609618 | -3 | 0.892974 | 0.892974 | 0.67038 | 0.1794 | 0.06986 | 0.12788 | 14.38506 | 0.02 | 0 | Neutral |
| testneg_4858 | 0 | 0 | 1.73928 | -3 | 4.431523 | -3 | 0.17643 | 0.10767 | 0.14007 | 0.10647 | 18.88889 | 0.031424347 | 0 | Neutral |
| testneg_4859 | 0 | 0 | 0.889458 | 0.889458 | 2.076532 | -3 | 0.48937 | 0.22085 | 0.08775 | 0.12788 | 31.80135 | 0.06 | 0 | Neutral |
| testneg_4860 | 0.05 | -0.81 | -0.842653 | -3 | 2.67688 | -3 | 0.06708 | 0.09234 | 0.03391 | 0.09745 | 21.30081 | 0.05 | 0 | Neutral |
| testneg_4861 | 0.09 | -2.17 | 3.628039 | -3 | 4.093803 | -3 | 0.055 | 0.20858 | 0.35234 | 0.1303 | 13.95683 | 0.477262766 | 44.2 | Neutral |
| testneg_4862 | -0.14 | 0.37 | 1.291415 | -3 | 11.548039 | -3 | 0.96801 | 0.20777 | 0.19898 | 0.0989 | 22.41987 | 0.46 | 35 | Neutral |
| testneg_4863 | 0 | -0.01 | 0.912506 | -3 | 0.958989 | -3 | 0.38429 | 0.11956 | 0.39436 | 0.16446 | 18.6768 | 0.03 | 0 | Neutral |
| testneg_4864 | -0.06 | 0.57 | 1.112541 | -3 | 4.639274 | -3 | 0.07982 | 0.12788 | 0.85508 | 0.12634 | 39.30712 | 0.13125 | 0 | Neutral |
| testneg_4865 | -0.03 | 1.03 | -1.584877 | -3 | 4.038181 | 4.038181 | 0.01409 | 0.11307 | 0.13815 | 0.40001 | 10.41667 | 0.461153846 | 40.4 | Neutral |
| testneg_4866 | -0.15 | -0.04 | 3.431501 | -3 | 0.144694 | -3 | 0.1289 | 0.14878 | 0.12048 | 0.2381 | 11.875 | 0.265787393 | 0.6 | Neutral |
| testneg_4867 | -0.07 | -0.38 | 0.37014 | -3 | 0.839368 | 0.839368 | 0.12048 | 0.2381 | 0.18785 | 0.14723 | 22.5469 | 0.18 | 0 | Neutral |
| testneg_4868 | 0.14 | -0.37 | 5.734512 | -3 | -0.06042 | -3 | 0.18498 | 0.12788 | 0.0961 | 0.21256 | 13.51351 | 0.388338624 | 17.8 | Neutral |
| testneg_4869 | 0 | 0 | 0.969725 | -3 | -1.448973 | -3 | 0.68029 | 0.12788 | 0.6975 | 0.53794 | 4.72222 | 0.030375439 | 0 | Neutral |
| testneg_4870 | 0 | 0 | 0.734956 | -3 | -0.298555 | -0.298555 | 0.14954 | 0.12788 | 0.10314 | 0.12173 | 20.35088 | 0.01 | 0 | Neutral |
| testneg_4871 | -0.05 | -0.29 | 1.376936 | -3 | -1.745 | -3 | 0.89992 | 0.10231 | 0.1743 | 0.12788 | 4.44444 | 0.01 | 0 | Neutral |
| testneg_4872 | 0 | 0 | 0.44344 | -3 | 6.151216 | -3 | 0.69666 | 0.11055 | 0.34324 | 0.085 | 18.75947 | 0.06225 | 0 | Neutral |
| testneg_4873 | -0.02 | -0.45 | 2.195922 | -3 | 2.273079 | -3 | 0.99704 | 0.17463 | 0.09284 | 0.10978 | 4.72222 | 0.07 | 0 | Neutral |
| testneg_4874 | -0.16 | 1.37 | 4.353927 | -3 | -0.697584 | -3 | 0.03072 | 0.27473 | 0.19898 | 0.12788 | 27.40684 | 0.29 | 0 | Neutral |
| testneg_4875 | -0.19 | 0.25 | -2.608658 | -3 | 0.339282 | -3 | 0.19898 | 0.12788 | 0.2547 | 0.07589 | 24.8155 | 0.060151803 | 0 | Neutral |
| testneg_4876 | 0 | 0 | 0.080104 | -3 | 1.360315 | -3 | 0.73947 | 0.2058 | 0.29482 | 0.15726 | 30.95732 | 0.11 | 0 | Neutral |
| testneg_4877 | 0.16 | -1.37 | 7.201556 | -3 | 1.87465 | 1.87465 | 0.93942 | 0.23984 | 0.16616 | 0.11073 | 13.90883 | 0.47 | 43.4 | Neutral |
| testneg_4878 | 0 | 0 | 1.38626 | -3 | 7.097934 | -3 | 0.50248 | 0.10939 | 0.2185 | 0.20593 | 10.41667 | 0.374543024 | 15.6 | Neutral |
| testneg_4879 | 0 | 0 | 0.244145 | -3 | 7.731489 | -3 | 0.9662 | 0.28104 | 0.03378 | 0.12788 | 27.22222 | 0.3344375 | 7.8 | Neutral |

| | | | | | | | | | | | | | | |
|---|---|---|---|---|---|---|---|---|---|---|---|---|---|---|
| testneg_4880 | -0.05 | -0.29 | 4.882717 | -3 | -0.416655 | -0.416655 | 0.62339 | 0.19592 | 0.05453 | 0.12788 | 20.20694 | 0.109172204 | 0 | Neutral |
| testneg_4881 | 0.08 | -1.2 | -0.35214 | -3 | 12.519035 | -3 | 0.11134 | 0.13818 | 0.03646 | 0.09863 | 34.29745 | 0.4 | 16.4 | Neutral |
| testneg_4882 | -0.03 | -0.1 | 3.084101 | -3 | 5.496455 | -3 | 0.39015 | 0.23372 | 0.17301 | 0.10048 | 8.73053 | 0.356814139 | 11.4 | Neutral |
| testneg_4883 | -0.15 | 0.28 | 3.882157 | -3 | 2.023546 | -3 | 0.15842 | 0.1294 | 0.33528 | 0.29463 | 16.81174 | 0.308502019 | 3 | Neutral |
| testneg_4884 | 0 | -0.01 | 1.902172 | -3 | 0.586954 | -3 | 0.23968 | 0.12788 | 0.4118 | 0.1154 | 21.38889 | 0.01 | 0 | Neutral |
| testneg_4885 | 0 | 0 | 1.316654 | -3 | -0.23732 | -0.23732 | 0.16613 | 0.12378 | 0.07473 | 0.51812 | 10.28409 | 0.060785516 | 0 | Neutral |
| testneg_4886 | -0.02 | 0.68 | 0.03082 | -3 | 1.753334 | -3 | 0.23385 | 0.12788 | 0.19898 | 0.12788 | 5.76832 | 0.010374126 | 0 | Neutral |
| testneg_4887 | 0.07 | -0.31 | 0.469669 | -3 | 1.986841 | 1.986841 | 0.09556 | 0.10109 | 0.19898 | 0.12788 | 26.70284 | 0.04 | 0 | Neutral |
| testneg_4888 | -0.03 | -0.1 | -0.76618 | -3 | 3.251074 | -3 | 0.1503 | 0.12788 | 0.20632 | 0.12788 | 22.27594 | 0.040298574 | 0 | Neutral |
| testneg_4889 | -0.17 | -0.38 | 0.481912 | -3 | 0.451974 | -3 | 0.22155 | 0.1062 | 0.42306 | 0.86891 | 10 | 0.12 | 0 | Neutral |
| testneg_4890 | 0.16 | -0.37 | -2.713571 | -2.713571 | 0.406982 | -3 | 0.07506 | 0.09112 | 0.13255 | 0.0962 | 28.5773 | 0.03 | 0 | Neutral |
| testneg_4891 | 0.14 | -0.37 | 5.019921 | -3 | 2.354743 | -3 | 0.41953 | 0.44852 | 0.13322 | 0.11297 | 37.92553 | 0.265995693 | 0 | Neutral |
| testneg_4892 | 0 | 0 | 0.528019 | -3 | 0.9704 | -3 | 0.42621 | 0.49093 | 0.06092 | 0.07159 | 12.29167 | 0.02 | 0 | Neutral |
| testneg_4893 | -0.08 | -0.03 | 1.562114 | -3 | 1.043597 | -3 | 0.78356 | 0.13377 | 0.19898 | 0.1005 | 22.30912 | 0.020151723 | 0 | Neutral |
| testneg_4894 | 0.02 | -0.36 | 1.712692 | -3 | 1.5447 | -3 | 0.06848 | 0.09963 | 0.08932 | 0.32882 | 16.84579 | 0.092546032 | 0 | Neutral |
| testneg_4895 | 0 | 0 | 1.560019 | -3 | 0.395033 | -3 | 0.19898 | 0.13495 | 0.15202 | 0.12722 | 25.55762 | 0.01 | 0 | Neutral |
| testneg_4896 | 0.16 | -0.37 | 0.701088 | -3 | 1.202078 | -3 | 0.99538 | 0.40044 | 0.16286 | 0.15786 | 10.87424 | 0.170729167 | 0 | Neutral |
| testneg_4897 | 0 | 0 | 0.387274 | -3 | 2.044018 | -3 | 0.35412 | 0.10819 | 0.63431 | 0.09837 | 13.281 | 0.000243902 | 0 | Neutral |
| testneg_4898 | -0.07 | 0.31 | -0.329074 | -3 | 12.520387 | 7.163551 | 0.04916 | 0.14944 | 0.22582 | 0.12788 | 10.35388 | 0.76 | 100 | Disease_causing |
| testneg_4899 | 0.05 | -0.81 | 6.767868 | -3 | 0.221712 | -3 | 0.50224 | 0.4059 | 0.54648 | 0.19042 | 10 | 0.559258242 | 73.8 | Disease_causing |
| testneg_4900 | 0 | -0.01 | 6.151767 | -3 | 5.639831 | -3 | 0.08002 | 0.08792 | 0.46142 | 0.11057 | 32.21639 | 0.32 | 7.4 | Neutral |
| testneg_4901 | 0.14 | -0.37 | 0.198379 | -3 | 0.643368 | -3 | 0.03991 | 0.08342 | 0.95071 | 0.25454 | 4.72222 | 0.1 | 0 | Neutral |
| testneg_4902 | 0 | 0 | -0.695904 | -3 | 0.451382 | -3 | 0.19364 | 0.14027 | 0.21774 | 0.10131 | 25.38961 | 0.010149387 | 0 | Neutral |
| testneg_4903 | -0.18 | 1.05 | 3.812572 | -3 | -0.153757 | -3 | 0.59853 | 0.75237 | 0.23027 | 0.37171 | 10 | 0.371944444 | 11 | Neutral |
| testneg_4904 | 0.07 | 0.01 | 2.699312 | -3 | 2.364247 | -3 | 0.17843 | 0.15468 | 0.09532 | 0.35459 | 18.21835 | 0.316832751 | 3.6 | Neutral |
| testneg_4905 | 0 | 0 | 0.110295 | -3 | -0.330275 | -0.330275 | 0.36709 | 0.19019 | 0.10462 | 0.1278 | 31.87292 | 0.030526681 | 0 | Neutral |
| testneg_4906 | 0.03 | 0.12 | 2.981794 | -3 | 0.85666 | -3 | 0.17394 | 0.15823 | 0.10278 | 0.08667 | 29.16667 | 0.06 | 0 | Neutral |
| testneg_4907 | 0 | 0 | -0.801037 | -3 | -0.01145 | -0.01145 | 0.2049 | 0.12678 | 0.78462 | 0.12258 | 18.6768 | 0.01 | 0 | Neutral |
| testneg_4908 | 0 | -0.01 | 0.93478 | -3 | 1.407842 | -3 | 0.57239 | 0.16622 | 0.63418 | 0.69224 | 14.22149 | 0.051452409 | 0 | Neutral |
| testneg_4909 | -0.15 | -0.04 | 2.20379 | -3 | 4.660061 | -3 | 0.60703 | 0.12378 | 0.1897 | 0.12459 | 18.16845 | 0.156998106 | 0 | Neutral |
| testneg_4910 | 0.2 | -1.26 | -1.784808 | -3 | 6.142434 | 5.730622 | 0.11133 | 0.12788 | 0.18597 | 0.12788 | 19.66924 | 0.42125 | 21 | Neutral |
| testneg_4911 | -0.01 | -0.79 | 1.796129 | -3 | 6.385153 | -3 | 0.16052 | 0.12788 | 0.21989 | 0.10981 | 27.32556 | 0.149666667 | 0 | Neutral |
| testneg_4912 | 0 | 0 | 0.394809 | -3 | 0.999083 | -3 | 0.04393 | 0.12788 | 0.84262 | 0.10997 | 13.2927 | 0.030175557 | 0 | Neutral |
| testneg_4913 | 0.14 | -1.01 | 1.433381 | -3 | 6.808241 | -3 | 0.12543 | 0.12788 | 0.14904 | 0.13261 | 18.76159 | 0.381477352 | 14.4 | Neutral |
| testneg_4914 | 0.18 | 0.16 | 0.556181 | 0.556181 | 1.619888 | -3 | 0.32406 | 0.28376 | 0.19477 | 0.12788 | 8.97898 | 0.08 | 0 | Neutral |
| testneg_4915 | -0.09 | 0.25 | 5.966239 | 1.957153 | 1.562114 | -3 | 0.89972 | 0.10766 | 0.78356 | 0.13377 | 9.14777 | 0.438 | 30.8 | Neutral |
| testneg_4916 | 0 | 0 | -0.801037 | -3 | 6.420066 | -3 | 0.2049 | 0.12678 | 0.13433 | 0.09578 | 26.53988 | 0.13 | 0 | Neutral |
| testneg_4917 | 0.04 | -0.23 | 5.836424 | 5.836424 | 4.741413 | -3 | 0.1503 | 0.12788 | 0.21115 | 0.17723 | 25.2451 | 0.838968254 | 100 | Disease_causing |
| testneg_4918 | 0 | 0 | 0.036223 | -3 | -0.564575 | -3 | 0.9355 | 0.40418 | 0.68029 | 0.12788 | 5.97222 | 0.014840217 | 0 | Neutral |
| testneg_4919 | -0.16 | -0.08 | 3.747066 | -3 | 2.673981 | -3 | 0.1535 | 0.12382 | 0.48936 | 0.10963 | 35.32468 | 0.133392857 | 0 | Neutral |
| testneg_4920 | 0.03 | -1.03 | 4.715557 | -3 | 3.506817 | -3 | 0.04243 | 0.22343 | 0.23088 | 0.21602 | 11.92982 | 0.729805696 | 98.8 | Disease_causing |
| testneg_4921 | 0 | 0 | 0.440494 | 0.440494 | -0.072198 | -0.072198 | 0.61027 | 0.14229 | 0.1025 | 0.3268 | 19.66924 | 0.180182084 | 0 | Neutral |
| testneg_4922 | 0.14 | -1.85 | -1.690089 | -3 | 0.010179 | -3 | 0.20918 | 0.10825 | 0.19898 | 0.12788 | 27.95796 | 0.04 | 0 | Neutral |
| testneg_4923 | 0.21 | 0.15 | 3.070963 | -3 | 1.238027 | -3 | 0.11791 | 0.39607 | 0.35499 | 0.12052 | 45.31794 | 0.16 | 0 | Neutral |
| testneg_4924 | 0.1 | 1.9 | 6.154974 | -3 | 6.010756 | -3 | 0.04515 | 0.14318 | 0.92262 | 0.12859 | 10.83333 | 0.611308675 | 91.8 | Disease_causing |
| testneg_4925 | -0.01 | -0.8 | -0.439118 | -3 | 1.52981 | -3 | 0.9149 | 0.27793 | 0.34001 | 0.12788 | 12.34395 | 0.04 | 0 | Neutral |
| testneg_4926 | 0 | 0 | 7.325251 | -3 | 3.488071 | -3 | 0.16601 | 0.14015 | 0.1485 | 0.30134 | 24.20397 | 0.705353641 | 98.8 | Disease_causing |
| testneg_4927 | -0.08 | -0.03 | 2.768711 | 2.768711 | 0.385855 | -3 | 0.38324 | 0.25193 | 0.10153 | 0.13343 | 10.22898 | 0.46 | 42.6 | Neutral |
| testneg_4928 | 0.03 | 0.12 | 3.392117 | -3 | 7.064425 | -3 | 0.2231 | 0.22973 | 0.15856 | 0.10994 | 18.33333 | 0.555071429 | 73.4 | Disease_causing |
| testneg_4929 | 0.14 | -1.01 | 0.599729 | 0.599729 | 0.162506 | -3 | 0.38762 | 0.12788 | 0.06112 | 0.06901 | 34.50745 | 0.04 | 0 | Neutral |
| testneg_4930 | 0.05 | -0.81 | 2.983048 | -3 | 2.686841 | -3 | 0.32963 | 0.1035 | 0.06845 | 0.13843 | 14.17968 | 0.206762436 | 0.4 | Neutral |
| testneg_4931 | 0 | 0 | 0.018301 | -3 | 1.430967 | -3 | 0.0611 | 0.12135 | 0.2521 | 0.1784 | 28.47222 | 0.080216291 | 0 | Neutral |
| testneg_4932 | -0.02 | 0.36 | 2.178935 | -3 | 5.33097 | -3 | 0.77847 | 0.10751 | 0.08814 | 0.42357 | 11.25 | 0.358958491 | 7.6 | Neutral |
| testneg_4933 | 0 | 0 | 0.141681 | -3 | 2.354743 | -3 | 0.11458 | 0.16496 | 0.13322 | 0.11297 | 22.86114 | 0.050886093 | 0 | Neutral |
| testneg_4934 | 0 | 0 | 1.362887 | -3 | 6.678015 | -3 | 0.38527 | 0.24812 | 0.19898 | 0.12788 | 10.3336 | 0.267690017 | 3 | Neutral |
| testneg_4935 | 0.02 | -0.68 | 1.222356 | -3 | 0.131996 | -3 | 0.17631 | 0.14648 | 0.19821 | 0.20954 | 21.45033 | 0.091055556 | 0 | Neutral |
| testneg_4936 | 0.14 | -0.37 | 7.061469 | -3 | -1.487172 | -1.487172 | 0.32335 | 0.10604 | 0.103 | 0.403 | 16.8018 | 0.434 | 31.6 | Neutral |
| testneg_4937 | -0.04 | 1.36 | -1.284127 | -3 | 1.285876 | -1.93808 | 0.11096 | 0.10074 | 0.07552 | 0.12788 | 45.18018 | 0.08 | 0 | Neutral |
| testneg_4938 | 0.21 | 0.15 | 1.418708 | -3 | 1.448132 | -3 | 0.6532 | 0.12788 | 0.694 | 0.09974 | 18.96658 | 0.04 | 0 | Neutral |
| testneg_4939 | 0 | 0 | -0.115728 | -0.115728 | 0.362935 | -3 | 0.0922 | 0.07873 | 0.12488 | 0.12788 | 42.23154 | 0.01 | 0 | Neutral |
| testneg_4940 | 0 | -0.01 | 2.237213 | -3 | 3.95483 | -3 | 0.31244 | 0.4615 | 0.27605 | 0.21595 | 4.44444 | 0.467607089 | 41.8 | Neutral |
| testneg_4941 | -0.08 | -0.03 | 6.695303 | -3 | 4.099778 | -3 | 0.09871 | 0.09286 | 0.06092 | 0.07159 | 17.21847 | 0.365 | 10 | Neutral |
| testneg_4942 | 0.15 | 0.06 | 0.364762 | -3 | 1.792563 | -3 | 0.85075 | 0.16646 | 0.29739 | 0.10893 | 6.94444 | 0.047083333 | 0 | Neutral |
| testneg_4943 | 0.14 | -1.01 | 3.635519 | -3 | 5.113079 | -3 | 0.07995 | 0.12788 | 0.9581 | 0.12788 | 14.14216 | 0.250889328 | 0.8 | Neutral |
| testneg_4944 | 0 | 0 | 0.200125 | -3 | -0.01928 | -3 | 0.13863 | 0.2579 | 0.09445 | 0.12788 | 12.70833 | 0.02 | 0 | Neutral |
| testneg_4945 | -0.01 | -0.8 | 6.193509 | -3 | 1.663993 | -3 | 0.31817 | 0.17348 | 0.13997 | 0.19598 | 31.38802 | 0.58 | 80.6 | Disease_causing |
| testneg_4946 | -0.22 | 2.66 | 1.021201 | 1.021201 | 1.435364 | -3 | 0.15785 | 0.35538 | 0.20357 | 0.12788 | 16.42857 | 0.24 | 0 | Neutral |
| testneg_4947 | -0.07 | -0.38 | 2.568127 | -3 | 7.68794 | -3 | 0.65141 | 0.18046 | 0.19898 | 0.12788 | 10.41667 | 0.467823816 | 43.4 | Neutral |
| testneg_4948 | 0 | 0 | 1.224946 | -3 | 2.266693 | -3 | 0.82126 | 0.17937 | 0.08401 | 0.16123 | 10.41667 | 0.10674092 | 0 | Neutral |
| testneg_4949 | 0.03 | 0.12 | 3.561858 | -3 | 0.433941 | 0.433941 | 0.7342 | 0.11023 | 0.30475 | 0.21064 | 9.88628 | 0.138743687 | 0 | Neutral |
| testneg_4950 | 0 | 0 | 0.094005 | -3 | -0.009875 | -3 | 0.64302 | 0.11262 | 0.25236 | 0.11809 | 9.93588 | 0 | 0 | Neutral |
| testneg_4951 | 0 | 0 | 0.068014 | 0.068014 | 0.476024 | -3 | 0.20869 | 0.13493 | 0.24832 | 0.21196 | 11.34615 | 0.050554422 | 0 | Neutral |
| testneg_4952 | -0.02 | -0.45 | -0.197057 | -3 | 1.377684 | -3 | 0.16797 | 0.12788 | 0.09406 | 0.19858 | 26.9055 | 0.1001973 | 0 | Neutral |
| testneg_4953 | 0 | 0 | -0.961702 | -0.961702 | 0.442945 | -3 | 0.46979 | 0.1705 | 0.19499 | 0.12111 | 12.70833 | 0.02 | 0 | Neutral |
| testneg_4954 | 0.16 | -1.37 | 3.320732 | 3.320732 | 0.596978 | -3 | 0.19898 | 0.13249 | 0.05095 | 0.08353 | 13.51351 | 0.42 | 27.8 | Neutral |
| testneg_4955 | 0.17 | 0.38 | 3.17007 | -3 | 1.574468 | -3 | 0.12377 | 0.12788 | 0.12121 | 0.4142 | 20.28172 | 0.254996951 | 0.2 | Neutral |
| testneg_4956 | 0.07 | 0.38 | 1.715954 | -3 | -0.750318 | -3 | 0.13126 | 0.11628 | 0.26068 | 0.46457 | 10.41667 | 0.055625 | 0 | Neutral |
| testneg_4957 | 0 | 0 | 0.260427 | -3 | 2.395534 | -3 | 0.08465 | 0.1932 | 0.13967 | 0.15696 | 22.02418 | 0.215515478 | 0 | Neutral |
| testneg_4958 | -0.1 | -1.9 | 1.643871 | -3 | 3.441233 | -3 | 0.19898 | 0.12788 | 0.22046 | 0.12788 | 18.73032 | 0.08 | 0 | Neutral |
| testneg_4959 | 0 | 0 | -0.468798 | -3 | 6.153381 | -3 | 0.03631 | 0.80564 | 0.08489 | 0.12788 | 12.38369 | 0.29 | 1 | Neutral |
| testneg_4960 | 0 | 0 | 1.369699 | -3 | 0.548908 | -3 | 0.50063 | 0.154 | 0.07258 | 0.06248 | 21.26172 | 0.03 | 0 | Neutral |
| testneg_4961 | -0.15 | 0.28 | 0.075246 | -3 | 2.747402 | 2.747402 | 0.19898 | 0.12788 | 0.11134 | 0.13818 | 37.68848 | 0.281102273 | 1 | Neutral |
| testneg_4962 | -0.1 | -1.9 | 1.624194 | -3 | 2.156166 | -3 | 0.26181 | 0.12875 | 0.13185 | 0.51629 | 21.43745 | 0.176137019 | 0 | Neutral |
| testneg_4963 | 0 | 0 | 1.243151 | -3 | 1.567144 | -3 | 0.91377 | 0.5142 | 0.11043 | 0.12788 | 11.93907 | 0.02 | 0 | Neutral |
| testneg_4964 | 0.12 | -1.16 | 3.511197 | -3 | -0.654233 | -3 | 0.19087 | 0.44337 | 0.04795 | 0.07008 | 12.3206 | 0.1 | 0 | Neutral |
| testneg_4965 | 0 | 0 | -0.049164 | -0.049164 | 2.013802 | -3 | 0.12322 | 0.1246 | 0.22239 | 0.12885 | 21.38889 | 0.01 | 0 | Neutral |
| testneg_4966 | -0.03 | -0.12 | -0.214687 | -3 | 6.179448 | -3 | 0.19668 | 0.16042 | 0.17296 | 0.09548 | 17.21847 | 0.258088034 | 0.8 | Neutral |
| testneg_4967 | 0.02 | -0.36 | 2.305765 | -3 | 0.038831 | -3 | 0.11055 | 0.1127 | 0.02498 | 0.08483 | 14.88684 | 0.01 | 0 | Neutral |
| testneg_4968 | 0.15 | 0.06 | 0.572808 | -3 | 0.459406 | -3 | 0.12036 | 0.0873 | 0.03981 | 0.09432 | 20.57907 | 0.03 | 0 | Neutral |
| testneg_4969 | 0.08 | 0.03 | -0.144427 | -3 | 0.723398 | 0.723398 | 0.13575 | 0.12788 | 0.19898 | 0.14957 | 27.4024 | 0.080933916 | 0 | Neutral |
| testneg_4970 | 0 | 0 | 0.240863 | -3 | -0.360768 | -0.360768 | 0.22816 | 0.10878 | 0.47147 | 0.09503 | 19.66924 | 0.01 | 0 | Neutral |
| testneg_4971 | 0.03 | -1.03 | 3.203334 | -3 | 4.775788 | -3 | 0.12633 | 0.19996 | 0.56204 | 0.10762 | 18.46847 | 0.339677419 | 6.2 | Neutral |
| testneg_4972 | -0.19 | 0.25 | 4.16391 | -3 | 7.749981 | -3 | 0.46058 | 0.12903 | 0.10278 | 0.08667 | 35.12807 | 0.513333333 | 63 | Disease_causing |
| testneg_4973 | 0.04 | -0.23 | 1.605283 | -3 | 0.295108 | -3 | 0.7257 | 0.64394 | 0.09527 | 0.12788 | 18.88889 | 0.053666667 | 0 | Neutral |
| testneg_4974 | 0 | 0 | 4.656753 | -3 | -0.133106 | -0.133106 | 0.52388 | 0.25831 | 0.07096 | 0.09965 | 11.57454 | 0.073611111 | 0 | Neutral |
| testneg_4975 | 0 | 0 | 0.884014 | -3 | 1.73928 | -3 | 0.16125 | 0.08406 | 0.17643 | 0.10767 | 14.96975 | 0.000217494 | 0 | Neutral |
| testneg_4976 | 0.03 | 1.22 | -0.307704 | -3 | 0.875146 | -3 | 0.12497 | 0.09805 | 0.09284 | 0.10978 | 20.98109 | 0.05 | 0 | Neutral |
| testneg_4977 | 0.03 | -1.03 | 2.174777 | -3 | 4.361375 | -3 | 0.32458 | 0.1016 | 0.10459 | 0.12783 | 18.3314 | 0.097300725 | 0 | Neutral |
| testneg_4978 | -0.08 | -0.03 | -0.385913 | -3 | 7.490616 | 1.258706 | 0.0752 | 0.74467 | 0.85345 | 0.13067 | 26.54712 | 0.545 | 75 | Disease_causing |
| testneg_4979 | 0.07 | 0.33 | 0.160465 | -3 | 2.247345 | -3 | 0.05153 | 0.12788 | 0.09334 | 0.10027 | 18.6768 | 0.05 | 0 | Neutral |
| testneg_4980 | -0.01 | -0.8 | 7.657206 | -3 | 4.785037 | -3 | 0.19898 | 0.12788 | 0.31025 | 0.09491 | 9.30313 | 0.608333333 | 89.2 | Disease_causing |
| testneg_4981 | 0.04 | -0.23 | -0.122196 | -0.122196 | 0.116245 | -3 | 0.31961 | 0.17571 | 0.8238 | 0.38836 | 10.83333 | 0.10485479 | 0 | Neutral |
| testneg_4982 | -0.06 | 0.57 | 5.853414 | -3 | 4.136929 | -3 | 0.19832 | 0.19622 | 0.19898 | 0.12788 | 29.21475 | 0.53 | 67.4 | Disease_causing |
| testneg_4983 | 0.21 | 0.15 | 4.18135 | -3 | 5.118069 | -3 | 0.19898 | 0.12788 | 0.33818 | 0.12788 | 10.41667 | 0.30421952 | 6 | Neutral |
| testneg_4984 | -0.07 | -0.38 | 4.718197 | 1.824706 | 0.204096 | -3 | 0.90955 | 0.10096 | 0.08531 | 0.12788 | 18.88889 | 0.16 | 0 | Neutral |
| testneg_4985 | 0 | 0 | 0.383044 | -3 | 0.975671 | -3 | 0.63819 | 0.32258 | 0.29739 | 0.10893 | 13.80963 | 0.01 | 0 | Neutral |
| testneg_4986 | 0 | 0 | -0.203664 | -3 | 5.503167 | -3 | 0.59321 | 0.59981 | 0.13961 | 0.09565 | 15.38642 | 0.141225427 | 0 | Neutral |
| testneg_4987 | 0.18 | -1.05 | 2.222067 | 1.525463 | 0.106535 | -3 | 0.09462 | 0.11807 | 0.44429 | 0.20468 | 25.74388 | 0.2 | 0 | Neutral |
| testneg_4988 | 0 | 0 | -0.19086 | -3 | 1.606239 | 1.606239 | 0.15958 | 0.41628 | 0.89002 | 0.12788 | 10.41667 | 0.040839161 | 0 | Neutral |
| testneg_4989 | 0 | 0 | 0.50666 | -3 | 1.729486 | -3 | 0.72062 | 0.14321 | 0.26068 | 0.46457 | 26.52893 | 0.090928571 | 0 | Neutral |
| testneg_4990 | 0 | -0.01 | 0.494045 | -3 | 1.959161 | -3 | 0.1304 | 0.12788 | 0.18312 | 0.12788 | 27.4024 | 0.000217494 | 0 | Neutral |
| testneg_4991 | 0.08 | 0.03 | 2.365497 | -3 | 1.21068 | -3 | 0.61491 | 0.11954 | 0.95247 | 0.11285 | 12.70424 | 0.03 | 0 | Neutral |
| testneg_4992 | -0.09 | 0.3 | -2.556301 | -3 | 0.888014 | -3 | 0.12379 | 0.12788 | 0.47451 | 0.3285 | 11.78612 | 0.080269886 | 0 | Neutral |
| testneg_4993 | 0 | 0 | 0.347529 | -3 | 4.643128 | -3 | 0.22443 | 0.10657 | 0.13083 | 0.23231 | 10.41667 | 0.208367244 | 0.2 | Neutral |
| testneg_4994 | 0 | 0 | 1.256275 | -3 | 1.13018 | -3 | 0.54724 | 0.12788 | 0.50505 | 0.11563 | 18.6768 | 0.005 | 0 | Neutral |
| testneg_4995 | 0 | 0 | 0.900502 | -0.566651 | 0.458968 | -3 | 0.38016 | 0.14503 | 0.25485 | 0.12788 | 13.89832 | 0.000223471 | 0 | Neutral |
| testneg_4996 | -0.05 | -0.29 | 1.332595 | -3 | 2.918397 | -3 | 0.77196 | 0.14089 | 0.63378 | 0.09451 | 11.83754 | 0.05 | 0 | Neutral |
| testneg_4997 | 0.07 | 0.38 | 2.246546 | -3 | 5.845998 | -3 | 0.32022 | 0.09529 | 0.1231 | 0.3662 | 74.64973 | 0.42885101 | 26.4 | Neutral |
| testneg_4998 | -0.21 | -0.15 | -0.114657 | -3 | -0.006272 | -0.006272 | 0.93228 | 0.1905 | 0.07335 | 0.12788 | 18.50941 | 0.08 | 0 | Neutral |
| testneg_4999 | 0.03 | 0.12 | -1.817927 | -3 | 5.665903 | -3 | 0.12455 | 0.0998 | 0.13031 | 0.85678 | 24.83714 | 0.366862637 | 9 | Neutral |
| testneg_5000 | -0.07 | -0.01 | -0.474074 | -3 | -0.800576 | -0.800576 | 0.06419 | 0.35525 | 0.05245 | 0.13458 | 24.1453 | 0.23 | 0 | Neutral |
| testneg_5001 | -0.01 | -0.79 | 1.105669 | -3 | 3.146686 | -3 | 0.44385 | 0.1311 | 0.0634 | 0.0824 | 10.83333 | 0.087852308 | 0 | Neutral |

| | | | | | | | | | | | | | | |
|---|---|---|---|---|---|---|---|---|---|---|---|---|---|---|
| testneg_5002 | -0.02 | -0.45 | -0.038552 | -3 | 1.089669 | -3 | 0.0749 | 0.12788 | 0.13796 | 0.27869 | 13.02635 | 0.070288693 | 0 | Neutral |
| testneg_5003 | 0 | 0 | -0.843404 | -3 | 1.673428 | 1.673428 | 0.11587 | 0.12001 | 0.32352 | 0.10243 | 10.2717 | 0.02 | 0 | Neutral |
| testneg_5004 | -0.03 | -0.1 | 4.591842 | -3 | 0.341374 | -3 | 0.84596 | 0.10633 | 0.10646 | 0.1069 | 20.38664 | 0.06 | 0 | Neutral |
| testneg_5005 | 0.07 | 0.7 | 4.354789 | -3 | 3.653901 | 3.099225 | 0.64319 | 0.12788 | 0.4069 | 0.3841 | 20.38801 | 0.620772202 | 94.4 | Disease_causing |
| testneg_5006 | 0 | 0 | -0.68826 | -3 | 0.10268 | -3 | 0.32076 | 0.14229 | 0.34193 | 0.12788 | 16.8018 | 0.002763158 | 0 | Neutral |
| testneg_5007 | 0 | 0 | -0.489156 | -0.925497 | 1.137849 | -3 | 0.05887 | 0.10046 | 0.15147 | 0.12788 | 22.30747 | 0.02 | 0 | Neutral |
| testneg_5008 | 0.04 | -0.23 | 6.599414 | -3 | 0.789 | -3 | 0.26668 | 0.11598 | 0.82156 | 0.14621 | 11.90227 | 0.269444444 | 2.8 | Neutral |
| testneg_5009 | 0 | 0 | 1.132066 | -3 | 1.158382 | -3 | 0.99317 | 0.60884 | 0.19087 | 0.44337 | 4.72222 | 0.081092857 | 0 | Neutral |
| testneg_5010 | -0.15 | -0.04 | 1.334446 | 1.334446 | 1.114692 | -3 | 0.15514 | 0.14852 | 0.05124 | 0.07099 | 18.88889 | 0.082892308 | 0 | Neutral |
| testneg_5011 | 0.03 | -1.03 | 2.514378 | -3 | -0.448326 | -3 | 0.3793 | 0.24105 | 0.03466 | 0.19518 | 27.68948 | 0.18 | 0 | Neutral |
| testneg_5012 | 0 | -0.01 | 2.237213 | -3 | 1.5733 | -3 | 0.31244 | 0.4615 | 0.42512 | 0.29333 | 11.65512 | 0.15015 | 0 | Neutral |
| testneg_5013 | 0 | 0 | 1.718666 | -3 | 6.160821 | -3 | 0.07889 | 0.08835 | 0.32532 | 0.09651 | 10 | 0.1 | 0 | Neutral |
| testneg_5014 | -0.03 | -0.1 | 1.521545 | -3 | 3.666622 | -3 | 0.12173 | 0.12788 | 0.61834 | 0.10784 | 39.72458 | 0.052047619 | 0 | Neutral |
| testneg_5015 | 0 | 0 | 0.462587 | -3 | 5.463671 | -3 | 0.26014 | 0.12378 | 0.50224 | 0.4059 | 11.69872 | 0.274112322 | 2.8 | Neutral |
| testneg_5016 | 0 | 0 | 0.104367 | -3 | 4.71084 | -3 | 0.62966 | 0.14896 | 0.33166 | 0.11755 | 18.33333 | 0.07115411 | 0 | Neutral |
| testneg_5017 | -0.07 | -0.01 | 0.736531 | -3 | 4.937534 | -3 | 0.07431 | 0.11513 | 0.42235 | 0.12474 | 13.51351 | 0.09 | 0 | Neutral |
| testneg_5018 | 0 | -0.32 | -0.849297 | -3 | 0.419264 | -3 | 0.23691 | 0.1015 | 0.93284 | 0.71489 | 24.44444 | 0.080060241 | 0 | Neutral |
| testneg_5019 | 0 | 0 | -0.050489 | -3 | 2.104429 | -3 | 0.13395 | 0.29754 | 0.30585 | 0.10254 | 34.81886 | 0.038375 | 0 | Neutral |
| testneg_5020 | 0.06 | -0.57 | 5.215375 | -3 | 2.300256 | -3 | 0.11818 | 0.10561 | 0.14996 | 0.09121 | 16.043 | 0.101566558 | 0 | Neutral |
| testneg_5021 | 0.07 | -0.31 | 1.587823 | -3 | 2.851691 | -3 | 0.9989 | 0.8746 | 0.21623 | 0.42074 | 5 | 0.336515269 | 4.4 | Neutral |
| testneg_5022 | 0 | 0 | -0.009858 | -3 | 1.177138 | -3 | 0.31664 | 0.14181 | 0.29624 | 0.1067 | 14.599 | 0.0008947 | 0 | Neutral |
| testneg_5023 | -0.05 | -0.29 | -0.062714 | -3 | -2.464297 | -2.464297 | 0.05323 | 0.32175 | 0.11524 | 0.12788 | 18.88889 | 0.12 | 0 | Neutral |
| testneg_5024 | -0.06 | -0.18 | 0.531542 | -3 | 1.022401 | -3 | 0.3766 | 0.21565 | 0.10879 | 0.10667 | 12.32809 | 0.021447449 | 0 | Neutral |
| testneg_5025 | -0.03 | -0.1 | 0.988046 | -3 | 0.465329 | 0.465329 | 0.19898 | 0.12788 | 0.11131 | 0.68435 | 11.875 | 0.103139579 | 0 | Neutral |
| testneg_5026 | -0.0016883 | 0.02623377 | 1.539365 | -3 | 1.15789 | 1.15789 | 0.3221 | 0.3367 | 0.68439 | 0.09371 | 10.41667 | 0.06 | 0 | Neutral |
| testneg_5027 | 0 | -0.01 | 1.494238 | -3 | 2.493168 | -3 | 0.12545 | 0.08963 | 0.14897 | 0.19122 | 18.3868 | 0.07074473 | 0 | Neutral |
| testneg_5028 | 0 | 0 | 1.636782 | -3 | 2.671237 | -3 | 0.15107 | 0.10457 | 0.36852 | 0.0903 | 56.17595 | 0.04 | 0 | Neutral |
| testneg_5029 | 0 | 0 | 0.028094 | -3 | 5.344722 | -3 | 0.46712 | 0.93559 | 0.09284 | 0.10978 | 9.75627 | 0.15 | 0 | Neutral |
| testneg_5030 | -0.07 | -0.38 | -0.220725 | -3 | 4.750251 | -3 | 0.23005 | 0.37216 | 0.20577 | 0.12653 | 10.41667 | 0.1959375 | 0 | Neutral |
| testneg_5031 | 0 | 0 | 0.606984 | -3 | 1.619354 | -3 | 0.08932 | 0.12788 | 0.27789 | 0.17365 | 20.91877 | 0.05935 | 0 | Neutral |
| testneg_5032 | 0 | 0 | 0.919505 | -3 | 2.450036 | -3 | 0.19898 | 0.12788 | 0.09997 | 0.1797 | 30.32444 | 0.130884615 | 0 | Neutral |
| testneg_5033 | 0.06 | 0.18 | 2.169317 | -3 | -1.295182 | -3 | 0.17083 | 0.31043 | 0.26913 | 0.1087 | 20.27778 | 0.03027305 | 0 | Neutral |
| testneg_5034 | 0.23 | -1.86 | 4.93441 | -3 | 1.269233 | -3 | 0.11564 | 0.09588 | 0.18998 | 0.26095 | 20.59942 | 0.316978261 | 3.2 | Neutral |
| testneg_5035 | 0 | -0.01 | 0.389904 | -3 | -0.416655 | -0.416655 | 0.95998 | 0.11262 | 0.05453 | 0.12788 | 27.3569 | 0.03 | 0 | Neutral |
| testneg_5036 | 0 | 0 | 0.429192 | -3 | -0.052748 | -3 | 0.50224 | 0.4059 | 0.91139 | 0.65138 | 10.83333 | 0.09 | 0 | Neutral |
| testneg_5037 | -0.05 | -0.29 | -0.079939 | -3 | 0.544663 | -3 | 0.50461 | 0.11034 | 0.09767 | 0.39937 | 20.48674 | 0.114163919 | 0 | Neutral |
| testneg_5038 | -0.05 | -0.29 | 3.795903 | -3 | 1.866374 | -3 | 0.88914 | 0.39065 | 0.22034 | 0.12788 | 16.09933 | 0.12 | 0 | Neutral |
| testneg_5039 | 0.17 | -1.22 | 4.106182 | -3 | 0.686233 | -3 | 0.06394 | 0.24464 | 0.17969 | 0.11473 | 11.98582 | 0.13 | 0 | Neutral |
| testneg_5040 | -0.01 | -0.8 | 3.94187 | -3 | 5.851071 | -3 | 0.37542 | 0.19444 | 0.32022 | 0.09529 | 10.41667 | 0.410588235 | 23.2 | Neutral |
| testneg_5041 | 0.15 | -0.28 | 1.374995 | -3 | 2.778725 | 2.778725 | 0.26727 | 0.12788 | 0.33528 | 0.29463 | 18.70665 | 0.28 | 0.8 | Neutral |
| testneg_5042 | 0 | 0 | 0.44344 | -3 | 0.916588 | -3 | 0.69666 | 0.11055 | 0.06603 | 0.08071 | 18.75947 | 0.01 | 0 | Neutral |
| testneg_5043 | -0.1 | -1.9 | -0.298153 | -3 | 1.074849 | -3 | 0.12099 | 0.11502 | 0.06775 | 0.09511 | 16.69872 | 0.04 | 0 | Neutral |
| testneg_5044 | 0 | 0 | 0.913032 | -3 | 5.996767 | -3 | 0.55708 | 0.13366 | 0.38695 | 0.12579 | 12.22261 | 0.106694655 | 0 | Neutral |
| testneg_5045 | 0 | 0 | 1.20976 | -3 | 5.175016 | -3 | 0.30209 | 0.12788 | 0.4334 | 0.09454 | 20.92575 | 0.05 | 0 | Neutral |
| testneg_5046 | -0.06 | -0.18 | 1.759866 | -3 | 6.070989 | -3 | 0.7249 | 0.41559 | 0.14187 | 0.1353 | 4.44444 | 0.480732732 | 48.4 | Neutral |
| testneg_5047 | -0.03 | -0.1 | 1.093873 | -3 | 2.464114 | -3 | 0.8433 | 0.13128 | 0.0611 | 0.12135 | 12.27364 | 0.04 | 0 | Neutral |
| testneg_5048 | 0 | -0.01 | 0.6334 | -3 | 0.80597 | -3 | 0.83798 | 0.14433 | 0.48298 | 0.1128 | 11.74429 | 0.01 | 0 | Neutral |
| testneg_5049 | 0.03 | 1.22 | 0.358373 | -3 | 6.201602 | 0.890517 | 0.49106 | 0.89143 | 0.1294 | 0.10964 | 13.28168 | 0.34 | 3.6 | Neutral |
| testneg_5050 | 0.14 | -1.01 | 0.803109 | -3 | -2.359472 | -3 | 0.99317 | 0.60884 | 0.22234 | 0.09474 | 10.71529 | 0.09 | 0 | Neutral |
| testneg_5051 | 0.07 | -0.31 | 3.048285 | 2.103598 | 5.700045 | -3 | 0.20604 | 0.10821 | 0.12682 | 0.11357 | 23.47372 | 0.3 | 2.8 | Neutral |
| testneg_5052 | 0 | -0.32 | -0.556879 | -3 | 0.154542 | -3 | 0.07007 | 0.12788 | 0.19898 | 0.535 | 10.22898 | 0.064958333 | 0 | Neutral |
| testneg_5053 | 0 | 0 | 0.858524 | -3 | 3.731623 | -3 | 0.23078 | 0.10588 | 0.19898 | 0.58778 | 12.08183 | 0.19746601 | 0 | Neutral |
| testneg_5054 | 0 | 0 | 1.405787 | -3 | 1.673428 | 1.673428 | 0.33304 | 0.15588 | 0.32352 | 0.10243 | 16.8018 | 0.02 | 0 | Neutral |
| testneg_5055 | 0 | -0.32 | 0.769645 | -3 | 0.439979 | -3 | 0.32335 | 0.10604 | 0.0811 | 0.09449 | 27.4024 | 0.01 | 0 | Neutral |
| testneg_5056 | 0 | 0 | 1.756493 | -3 | -0.329074 | -3 | 0.19052 | 0.12788 | 0.04916 | 0.14944 | 7.01593 | 0.04022739 | 0 | Neutral |
| testneg_5057 | 0.02 | -0.63 | -2.196989 | -3 | 0.814088 | -3 | 0.02759 | 0.12458 | 0.55197 | 0.11305 | 17.21847 | 0.04 | 0 | Neutral |
| testneg_5058 | 0.02 | -0.68 | 2.729304 | -3 | 0.513838 | -3 | 0.0917 | 0.13124 | 0.68004 | 0.12345 | 21.38889 | 0.05 | 0 | Neutral |
| testneg_5059 | 0.03 | -1.03 | 3.488962 | -3 | 0.335227 | -3 | 0.59245 | 0.12788 | 0.04346 | 0.10785 | 16.8018 | 0.04 | 0 | Neutral |
| testneg_5060 | 0.17 | 0.38 | 2.901134 | -3 | 1.311902 | -3 | 0.82014 | 0.14054 | 0.10337 | 0.12788 | 14.60764 | 0.08 | 0 | Neutral |
| testneg_5061 | 0 | 0 | 1.69763 | -3 | -0.505055 | -3 | 0.9862 | 0.4528 | 0.99859 | 0.12788 | 9.79167 | 0.06 | 0 | Neutral |
| testneg_5062 | -0.08 | -0.03 | 4.038145 | -3 | 2.183335 | -3 | 0.19898 | 0.07094 | 0.13143 | 0.90853 | 42.19844 | 0.26 | 0.2 | Neutral |
| testneg_5063 | 0.07 | 0.38 | 0.197957 | -3 | -0.01248 | -3 | 0.10556 | 0.28423 | 0.61231 | 0.13362 | 18.88889 | 0.160136986 | 0 | Neutral |
| testneg_5064 | 0 | 0 | 1.576055 | -3 | 0.147975 | -0.123782 | 0.05886 | 0.38242 | 0.06898 | 0.11659 | 28.166 | 0.090261905 | 0 | Neutral |
| testneg_5065 | 0 | 0 | 0.560537 | -3 | 0.631229 | -3 | 0.32804 | 0.10857 | 0.82269 | 0.15732 | 39.53468 | 0.040449949 | 0 | Neutral |
| testneg_5066 | 0.21 | 0.15 | 1.651539 | 0.582514 | 5.849437 | -3 | 0.06218 | 0.10661 | 0.11972 | 0.09438 | 26.38705 | 0.180374269 | 0 | Neutral |
| testneg_5067 | -0.06 | -0.18 | 1.398739 | -3 | 2.104561 | -3 | 0.54113 | 0.11017 | 0.06781 | 0.09009 | 7.1517 | 0.02 | 0 | Neutral |
| testneg_5068 | -0.01 | -0.79 | 4.508894 | -3 | -0.202618 | -3 | 0.17181 | 0.10294 | 0.51167 | 0.13142 | 21.20422 | 0.080495169 | 0 | Neutral |
| testneg_5069 | -0.03 | -0.1 | 5.021764 | -3 | 2.750883 | -3 | 0.30684 | 0.12788 | 0.19898 | 0.12788 | 22.62443 | 0.161796717 | 0 | Neutral |
| testneg_5070 | 0 | 0 | 1.283931 | -3 | 6.461058 | -3 | 0.03499 | 0.12971 | 0.19898 | 0.12788 | 45.93756 | 0.17 | 0 | Neutral |
| testneg_5071 | 0 | 0 | -0.71253 | -3 | -1.137419 | -3 | 0.03563 | 0.10455 | 0.45405 | 0.10896 | 18.33333 | 0.03 | 0 | Neutral |
| testneg_5072 | -0.04 | -2.01 | 4.558544 | -3 | 2.439465 | -3 | 0.08753 | 0.09666 | 0.06552 | 0.07762 | 20.26124 | 0.110426136 | 0 | Neutral |
| testneg_5073 | 0.05 | -0.81 | 3.064064 | -3 | 2.578388 | -3 | 0.5481 | 0.10207 | 0.16361 | 0.13388 | 29.28648 | 0.220153601 | 0 | Neutral |
| testneg_5074 | -0.16 | -0.08 | 1.75357 | -3 | 1.514538 | -3 | 0.06736 | 0.06344 | 0.19898 | 0.11773 | 11.96702 | 0.08 | 0 | Neutral |
| testneg_5075 | 0 | 0 | 0.477103 | -3 | -0.416655 | -0.416655 | 0.64072 | 0.12788 | 0.05453 | 0.12788 | 13.46801 | 0.01 | 0 | Neutral |
| testneg_5076 | -0.14 | 1.85 | 0.252727 | -3 | 4.142425 | -3 | 0.13985 | 0.08349 | 0.25447 | 0.12788 | 31.2434 | 0.22 | 0 | Neutral |
| testneg_5077 | 0 | -0.01 | 2.855452 | -3 | 0.945866 | -3 | 0.19898 | 0.12788 | 0.23537 | 0.33997 | 9.79167 | 0.12 | 0 | Neutral |
| testneg_5078 | -0.0022727 | -0.0332576 | 2.427467 | -3 | 1.359146 | -3 | 0.43866 | 0.12121 | 0.22575 | 0.37247 | 18.88889 | 0.087089945 | 0 | Neutral |
| testneg_5079 | 0.05 | 0.29 | 1.372356 | -3 | 7.325251 | -3 | 0.60026 | 0.35923 | 0.16601 | 0.14015 | 23.53741 | 0.63 | 93 | Disease_causing |
| testneg_5080 | -0.02 | -0.41 | 1.480614 | -3 | 1.819817 | -3 | 0.11849 | 0.20101 | 0.19898 | 0.12788 | 18.33333 | 0.04 | 0 | Neutral |
| testneg_5081 | -0.15 | -0.04 | 1.715795 | -3 | -3.590514 | -3 | 0.39863 | 0.44944 | 0.47322 | 0.11118 | 11.63355 | 0.08 | 0 | Neutral |
| testneg_5082 | -0.05 | -0.29 | 0.110557 | -3 | -0.237401 | -3 | 0.11703 | 0.09364 | 0.07537 | 0.1399 | 13.16701 | 0.060229915 | 0 | Neutral |
| testneg_5083 | 0.07 | -0.31 | -0.565166 | -3 | 1.745907 | -3 | 0.9832 | 0.39904 | 0.12949 | 0.44208 | 10.83333 | 0.19 | 0 | Neutral |
| testneg_5084 | -0.08 | -0.03 | 4.069302 | -3 | 3.519231 | -3 | 0.40632 | 0.1094 | 0.33049 | 0.12788 | 12.33727 | 0.163202403 | 0 | Neutral |
| testneg_5085 | 0 | 0 | 4.894604 | -3 | -1.941492 | -3 | 0.21726 | 0.23214 | 0.37711 | 0.12788 | 10.22898 | 0.11 | 0 | Neutral |
| testneg_5086 | 0 | 0 | 0.15975 | -3 | 3.450149 | -3 | 0.19898 | 0.16603 | 0.5043 | 0.3225 | 12.70833 | 0.281361364 | 2.2 | Neutral |
| testneg_5087 | 0 | -0.32 | -2.076593 | -3 | 2.78748 | -3 | 0.1276 | 0.30104 | 0.17977 | 0.12788 | 40.75397 | 0.100410487 | 0 | Neutral |
| testneg_5088 | -0.01 | -0.79 | 3.399462 | -3 | -0.964624 | -3 | 0.05472 | 0.12788 | 0.13825 | 0.0985 | 33.56606 | 0.08 | 0 | Neutral |
| testneg_5089 | -0.08 | -0.03 | 1.755955 | 1.755955 | 3.287017 | -3 | 0.9832 | 0.39904 | 0.1594 | 0.22002 | 12.20874 | 0.35640796 | 9.4 | Neutral |
| testneg_5090 | 0.21 | 0.15 | 2.804277 | -3 | 3.761039 | -3 | 0.02124 | 0.05186 | 0.67263 | 0.09781 | 20.96853 | 0.15 | 0 | Neutral |
| testneg_5091 | 0.02 | -0.68 | 6.540751 | -3 | 0.372849 | -3 | 0.5901 | 0.19444 | 0.10286 | 0.26819 | 18.33333 | 0.538309124 | 66.4 | Disease_causing |
| testneg_5092 | 0.04 | 2.01 | 3.833274 | -3 | 1.153408 | -3 | 0.18022 | 0.12788 | 0.1755 | 0.10455 | 23.68839 | 0.08 | 0 | Neutral |
| testneg_5093 | 0.09 | -0.3 | 4.414734 | -3 | 3.818938 | -3 | 0.05657 | 0.14812 | 0.65674 | 0.10572 | 41.34161 | 0.259004284 | 0.2 | Neutral |
| testneg_5094 | -0.09 | 0.3 | -0.522638 | -3 | 0.582514 | -3 | 0.20604 | 0.10821 | 0.06218 | 0.10661 | 21.88855 | 0.02 | 0 | Neutral |
| testneg_5095 | 0 | 0 | 1.327873 | -3 | 2.632198 | -3 | 0.12943 | 0.14391 | 0.23939 | 0.25284 | 11.25 | 0.111218341 | 0 | Neutral |
| testneg_5096 | 0.14 | -0.37 | 0.752578 | -3 | 1.539573 | -3 | 0.19898 | 0.12788 | 0.87611 | 0.16759 | 45.46569 | 0.070301178 | 0 | Neutral |
| testneg_5097 | 0 | 0 | -1.247889 | -3 | -0.729977 | -0.729977 | 0.27577 | 0.09812 | 0.17974 | 0.12788 | 16.59347 | 0.01 | 0 | Neutral |
| testneg_5098 | 0.04 | -0.23 | 0.51832 | -3 | 1.464262 | -3 | 0.33082 | 0.12788 | 0.10042 | 0.09296 | 13.51351 | 0.01 | 0 | Neutral |
| testneg_5099 | 0 | -0.01 | 0.773594 | -3 | 4.091456 | -3 | 0.16826 | 0.11262 | 0.18578 | 0.27854 | 9.67871 | 0.191615335 | 0 | Neutral |
| testneg_5100 | 0 | 0 | 1.720206 | -3 | 0.862282 | -3 | 0.6992 | 0.1804 | 0.51167 | 0.13142 | 9.04596 | 0.032041667 | 0 | Neutral |
| testneg_5101 | 0 | 0 | 0.496132 | -3 | 1.686834 | -3 | 0.16252 | 0.11134 | 0.29828 | 0.12788 | 13.88763 | 0 | 0 | Neutral |
| testneg_5102 | -0.03 | -0.1 | 2.059294 | -3 | -0.833334 | -3 | 0.66711 | 0.31094 | 0.57454 | 0.14465 | 23.91147 | 0.120596491 | 0 | Neutral |
| testneg_5103 | 0.15 | -0.28 | -1.251158 | -3 | 6.264472 | -3 | 0.20127 | 0.13478 | 0.71276 | 0.88182 | 11.84783 | 0.465965666 | 40.6 | Neutral |
| testneg_5104 | 0.15 | 0.04 | 2.282995 | -3 | 0.714605 | -3 | 0.47217 | 0.15544 | 0.82126 | 0.17937 | 10 | 0.091194492 | 0 | Neutral |
| testneg_5105 | -0.04 | -2.01 | 1.963276 | -3 | 4.087357 | -3 | 0.21355 | 0.12788 | 0.1174 | 0.12788 | 31.76621 | 0.11 | 0 | Neutral |
| testneg_5106 | -0.03 | 1.03 | 3.557072 | -3 | -0.3595 | -3 | 0.2477 | 0.11928 | 0.19898 | 0.12788 | 15 | 0.05 | 0 | Neutral |
| testneg_5107 | 0.06 | -1.29 | 1.367341 | -3 | 0.059493 | -3 | 0.04694 | 0.44682 | 0.14486 | 0.33725 | 15.00992 | 0.319486326 | 2 | Neutral |
| testneg_5108 | 0 | 0 | 0.730301 | -3 | 2.72063 | -3 | 0.40612 | 0.45135 | 0.65141 | 0.18046 | 4.72222 | 0.151762809 | 0 | Neutral |
| testneg_5109 | 0.02 | 0.45 | -1.0834 | -3 | 0.704025 | -3 | 0.14248 | 0.12788 | 0.60026 | 0.35923 | 22.46683 | 0.08 | 0 | Neutral |
| testneg_5110 | -0.09 | 0.3 | 1.202839 | -3 | 3.399462 | -3 | 0.11144 | 0.08199 | 0.05472 | 0.12788 | 31.47833 | 0.082057692 | 0 | Neutral |
| testneg_5111 | -0.1 | -1.9 | 5.567099 | -3 | 2.941823 | -3 | 0.05732 | 0.1098 | 0.10285 | 0.10186 | 54.58333 | 0.25 | 0.4 | Neutral |
| testneg_5112 | 0.04 | 2.01 | 5.876399 | -3 | 5.638751 | -3 | 0.7823 | 0.5949 | 0.1837 | 0.27351 | 9.35743 | 0.877321429 | 99.8 | Disease_causing |
| testneg_5113 | -0.17 | -0.38 | -0.92068 | -3 | 6.72304 | -3 | 0.28038 | 0.11814 | 0.18127 | 0.12788 | 31.10621 | 0.27 | 1.6 | Neutral |
| testneg_5114 | 0 | -0.32 | 2.075891 | -3 | 4.133728 | -3 | 0.14375 | 0.12788 | 0.13794 | 0.1141 | 17.61785 | 0.063166667 | 0 | Neutral |
| testneg_5115 | 0.01 | -0.57 | -0.412994 | -3 | 3.559501 | -3 | 0.04393 | 0.12788 | 0.3834 | 0.3731 | 16.88725 | 0.270145152 | 0.8 | Neutral |
| testneg_5116 | 0 | 0 | 0.92637 | -3 | 0.082232 | -3 | 0.17631 | 0.14648 | 0.08475 | 0.19889 | 35.68737 | 0.100205451 | 0 | Neutral |
| testneg_5117 | 0 | 0 | 0.102218 | -3 | 1.61682 | -3 | 0.64072 | 0.12788 | 0.21596 | 0.08969 | 34.698 | 0.010571895 | 0 | Neutral |
| testneg_5118 | 0.05 | 0.29 | 2.040423 | -3 | -2.373259 | -3 | 0.835 | 0.13588 | 0.04525 | 0.12788 | 17.90495 | 0.03 | 0 | Neutral |
| testneg_5119 | 0 | -0.01 | 1.149331 | 1.149331 | -0.410022 | -3 | 0.15582 | 0.14216 | 0.17419 | 0.33842 | 10.83333 | 0.12015721 | 0 | Neutral |
| testneg_5120 | 0.07 | 0.33 | 2.165808 | -3 | 4.404978 | -3 | 0.19898 | 0.07312 | 0.03505 | 0.0907 | 23.68839 | 0.080322286 | 0 | Neutral |
| testneg_5121 | 0.14 | -0.37 | 3.499396 | -3 | -0.765143 | -0.765143 | 0.69249 | 0.12788 | 0.31627 | 0.23288 | 9.80134 | 0.179414492 | 0.2 | Neutral |
| testneg_5122 | 0.01 | -1.44 | 3.195028 | 1.550971 | 0.649412 | -3 | 0.73311 | 0.10866 | 0.3394 | 0.5826 | 10.22898 | 0.321213235 | 4.2 | Neutral |
| testneg_5123 | 0.04 | -0.89 | -1.056088 | -3 | 1.729145 | -3 | 0.06781 | 0.09009 | 0.13575 | 0.12788 | 12.46106 | 0.010201927 | 0 | Neutral |

| | | | | | | | | | | | | | | |
|---|---|---|---|---|---|---|---|---|---|---|---|---|---|---|
| testneg_5124 | -0.02 | 0.36 | -0.710033 | -3 | 3.459821 | -3 | 0.12689 | 0.13376 | 0.09866 | 0.104 | 20.52939 | 0.1 | 0 | Neutral |
| testneg_5125 | -0.02 | -0.45 | -0.208207 | -3 | 0.528801 | -3 | 0.07219 | 0.12788 | 0.12136 | 0.27707 | 64.56058 | 0.193333333 | 0 | Neutral |
| testneg_5126 | 0 | 0 | 0.768207 | -3 | -0.607068 | -3 | 0.30669 | 0.11489 | 0.60918 | 0.11247 | 17.81315 | 0 | 0 | Neutral |
| testneg_5127 | -0.08 | -0.03 | 0.775573 | -3 | -1.495059 | -3 | 0.13325 | 0.12788 | 0.9581 | 0.12788 | 19.43262 | 0.020559468 | 0 | Neutral |
| testneg_5128 | -0.15 | 0.28 | -0.668638 | -0.668638 | 5.582498 | -3 | 0.37131 | 0.10994 | 0.85829 | 0.4292 | 11.96702 | 0.37768125 | 10.2 | Neutral |
| testneg_5129 | -0.01 | 0.57 | 1.86572 | -3 | -2.586155 | -3 | 0.05873 | 0.09792 | 0.98641 | 0.83983 | 17.95445 | 0.141555556 | 0 | Neutral |
| testneg_5130 | -0.03 | -1.22 | 3.110562 | -3 | -0.134358 | -3 | 0.7597 | 0.12788 | 0.13139 | 0.11068 | 11.08333 | 0.05 | 0 | Neutral |
| testneg_5131 | 0.16 | -1.37 | 6.571813 | 6.571813 | 4.387695 | -3 | 0.09488 | 0.10426 | 0.18341 | 0.13901 | 21.43745 | 0.74 | 99.8 | Disease_causing |
| testneg_5132 | 0 | 0 | 0.389761 | -3 | 4.643757 | -3 | 0.73601 | 0.26791 | 0.113 | 0.12788 | 10.83333 | 0.09996614 | 0 | Neutral |
| testneg_5133 | 0.09 | -0.67 | 1.87178 | -3 | 1.509728 | -3 | 0.11337 | 0.09363 | 0.0435 | 0.14527 | 10.41667 | 0.060419492 | 0 | Neutral |
| testneg_5134 | 0.21 | 0.15 | 2.729204 | 0.707638 | 2.18889 | -3 | 0.62339 | 0.19592 | 0.0819 | 0.1823 | 20.8819 | 0.36 | 6.6 | Neutral |
| testneg_5135 | 0.03 | 1.22 | 7.358991 | -3 | -1.729462 | -3 | 0.03109 | 0.10705 | 0.16155 | 0.20638 | 18.89211 | 0.437083333 | 30.6 | Neutral |
| testneg_5136 | 0.02 | 0.13 | -0.076183 | -3 | 1.943409 | -3 | 0.16418 | 0.12788 | 0.41647 | 0.11239 | 16.8018 | 0.01 | 0 | Neutral |
| testneg_5137 | 0.21 | 0.15 | 6.752392 | -3 | 5.226404 | -3 | 0.08857 | 0.83499 | 0.30423 | 0.12788 | 16.42913 | 0.697809128 | 98.2 | Disease_causing |
| testneg_5138 | 0.08 | 0.03 | -0.765287 | -3 | 5.312389 | -3 | 0.21002 | 0.12788 | 0.07293 | 0.12989 | 10.28409 | 0.110444664 | 0 | Neutral |
| testneg_5139 | 0.03 | -1.03 | 2.18442 | -3 | 4.046641 | -3 | 0.19898 | 0.12788 | 0.25422 | 0.10102 | 28.20322 | 0.071125996 | 0 | Neutral |
| testneg_5140 | 0.05 | 0.29 | 2.702357 | -3 | -0.877392 | -3 | 0.94674 | 0.12413 | 0.1311 | 0.12667 | 16.48772 | 0.03 | 0 | Neutral |
| testneg_5141 | 0 | 0 | 0.318608 | -3 | -1.336661 | -3 | 0.11373 | 0.10739 | 0.20513 | 0.09801 | 10.22898 | 0 | 0 | Neutral |
| testneg_5142 | -0.03 | -0.1 | 2.877215 | -3 | 2.325904 | -3 | 0.20657 | 0.12788 | 0.10485 | 0.09565 | 10.28409 | 0.050185467 | 0 | Neutral |
| testneg_5143 | -0.17 | -0.38 | 0.857098 | -3 | 0.103919 | -3 | 0.65734 | 0.1013 | 0.83955 | 0.10333 | 34.51735 | 0.06 | 0 | Neutral |
| testneg_5144 | 0 | -0.32 | -0.119569 | -3 | 0.988861 | -3 | 0.19898 | 0.10362 | 0.1316 | 0.28283 | 12.81915 | 0.05 | 0 | Neutral |
| testneg_5145 | 0.04 | -0.23 | 3.599958 | -3 | 0.585649 | 0.585649 | 0.40491 | 0.17596 | 0.07185 | 0.68732 | 18.88889 | 0.270440039 | 0.8 | Neutral |
| testneg_5146 | -0.0023485 | -0.0163636 | 2.329732 | -3 | 0.492674 | -3 | 0.15526 | 0.08511 | 0.19898 | 0.12788 | 30.01545 | 0.01 | 0 | Neutral |
| testneg_5147 | -0.06 | -0.18 | 2.103681 | -3 | 1.59057 | -3 | 0.19898 | 0.13434 | 0.26789 | 0.67432 | 10.83333 | 0.09529885 | 0 | Neutral |
| testneg_5148 | -0.05 | -0.29 | 2.72051 | -3 | 1.999625 | -3 | 0.64701 | 0.94571 | 0.58587 | 0.09866 | 10.18228 | 0.12 | 0 | Neutral |
| testneg_5149 | 0.08 | 0.03 | -0.61726 | -3 | 3.308288 | -3 | 0.20673 | 0.13236 | 0.48585 | 0.66989 | 14.27748 | 0.302258054 | 1 | Neutral |
| testneg_5150 | 0 | 0 | 1.444282 | -3 | 0.024903 | -3 | 0.17181 | 0.10294 | 0.19416 | 0.0901 | 33.42719 | 0.000114319 | 0 | Neutral |
| testneg_5151 | -0.12 | 1.14 | 6.045853 | -3 | 4.440447 | -3 | 0.07467 | 0.09224 | 0.86437 | 0.69972 | 19.44444 | 0.569736842 | 79.6 | Disease_causing |
| testneg_5152 | 0.04 | -0.23 | 7.214565 | -3 | 1.941203 | -3 | 0.11329 | 0.12116 | 0.28334 | 0.10368 | 14.78889 | 0.275 | 4 | Neutral |
| testneg_5153 | 0.18 | 0.16 | 4.695825 | -3 | 0.388086 | -3 | 0.44615 | 0.09968 | 0.15432 | 0.095 | 17.77778 | 0.05039721 | 0 | Neutral |
| testneg_5154 | 0.12 | -1.16 | 5.826992 | -3 | 1.632718 | -3 | 0.20084 | 0.0851 | 0.19898 | 0.12788 | 30.93087 | 0.14 | 0 | Neutral |
| testneg_5155 | 0 | -0.01 | 0.836822 | -3 | 0.800388 | -3 | 0.04866 | 0.05972 | 0.47816 | 0.66526 | 10.83333 | 0.09 | 0 | Neutral |
| testneg_5156 | -0.16 | 1.37 | 6.341825 | -3 | 3.477533 | -3 | 0.23506 | 0.24555 | 0.87847 | 0.96194 | 17.21847 | 0.68 | 97.4 | Disease_causing |
| testneg_5157 | 0 | 0 | 1.156713 | 1.156713 | 2.823466 | -3 | 0.36458 | 0.16397 | 0.227 | 0.14389 | 19.84127 | 0.190254918 | 0 | Neutral |
| testneg_5158 | 0.05 | 0.29 | 1.6302 | -3 | -0.718699 | -0.718699 | 0.23468 | 0.10874 | 0.21122 | 0.12788 | 18.6768 | 0.000279137 | 0 | Neutral |
| testneg_5159 | -0.15 | -0.04 | 1.282809 | -3 | 3.445512 | -3 | 0.65564 | 0.11468 | 0.38126 | 0.27764 | 17.21847 | 0.240382554 | 0.2 | Neutral |
| testneg_5160 | 0 | 0 | 0.035553 | 0.035553 | 3.635653 | -3 | 0.15414 | 0.17421 | 0.11501 | 0.13748 | 14.23074 | 0.311994624 | 6.6 | Neutral |
| testneg_5161 | 0 | 0 | 1.28208 | -3 | 1.677581 | -3 | 0.38375 | 0.09256 | 0.74919 | 0.85508 | 20.27778 | 0.061837413 | 0 | Neutral |
| testneg_5162 | 0 | -0.01 | 1.116052 | -3 | 1.461115 | -3 | 0.1631 | 0.40344 | 0.22953 | 0.0919 | 13.51351 | 0.02 | 0 | Neutral |
| testneg_5163 | 0 | 0 | 1.614896 | -3 | 5.019095 | -3 | 0.10703 | 0.11611 | 0.51587 | 0.13949 | 33.28244 | 0.193480392 | 0 | Neutral |
| testneg_5164 | -0.02 | 0.68 | 0.697273 | -3 | -0.915676 | -3 | 0.18559 | 0.13755 | 0.05767 | 0.07325 | 10.78704 | 0.020382653 | 0 | Neutral |
| testneg_5165 | 0 | 0 | 1.870488 | -3 | 2.83636 | -3 | 0.3548 | 0.10717 | 0.19002 | 0.12138 | 11.18405 | 0.02 | 0 | Neutral |
| testneg_5166 | 0.16 | 0.08 | 0.972454 | -3 | 1.102874 | -3 | 0.25561 | 0.11004 | 0.14511 | 0.12788 | 19.03093 | 0.010238636 | 0 | Neutral |
| testneg_5167 | 0 | 0 | 0.097356 | -3 | 0.988765 | -3 | 0.14485 | 0.16945 | 0.62197 | 0.12788 | 26.11603 | 0.021655883 | 0 | Neutral |
| testneg_5168 | -0.07 | -0.01 | 2.614937 | -3 | 0.700131 | -3 | 0.14185 | 0.12788 | 0.46056 | 0.12785 | 21.69337 | 0.01 | 0 | Neutral |
| testneg_5169 | 0.003 | -0.1380909 | 2.674709 | -0.100569 | 0.377311 | -3 | 0.6185 | 0.09316 | 0.0961 | 0.21256 | 19.09354 | 0.120736508 | 0 | Neutral |
| testneg_5170 | 0.09 | -0.3 | 2.550986 | -3 | 1.518082 | -3 | 0.08408 | 0.12788 | 0.09978 | 0.12788 | 19.44444 | 0.02 | 0 | Neutral |
| testneg_5171 | 0.07 | 0.38 | 1.022289 | 1.022289 | 3.237068 | -3 | 0.18281 | 0.12788 | 0.19898 | 0.12788 | 27.51545 | 0.09072479 | 0 | Neutral |
| testneg_5172 | 0.22 | -2.66 | 2.537219 | -3 | 0.967538 | -3 | 0.0328 | 0.0691 | 0.21235 | 0.1449 | 25.79439 | 0.151213235 | 0 | Neutral |
| testneg_5173 | 0 | 0 | 1.218246 | -3 | 7.8284 | -3 | 0.20448 | 0.12788 | 0.30585 | 0.10254 | 19.05506 | 0.269444444 | 2.6 | Neutral |
| testneg_5174 | 0 | 0 | 1.875005 | -3 | 1.636161 | 1.636161 | 0.50562 | 0.2196 | 0.21153 | 0.11357 | 13.101 | 0.03 | 0 | Neutral |
| testneg_5175 | 0.14 | -1.01 | 2.521088 | -3 | -1.66039 | -3 | 0.10021 | 0.1105 | 0.06957 | 0.07707 | 13.28391 | 0.02 | 0 | Neutral |
| testneg_5176 | 0 | 0 | 6.373062 | 1.221125 | 2.165741 | -3 | 0.36793 | 0.15588 | 0.09978 | 0.12788 | 12.7356 | 0.431201923 | 29 | Neutral |
| testneg_5177 | -0.04 | 0.23 | -0.481003 | -3 | 1.627178 | -3 | 0.45405 | 0.10896 | 0.30925 | 0.10335 | 31.25 | 0.02 | 0 | Neutral |
| testneg_5178 | -0.03 | -0.1 | 0.447373 | -3 | 2.031708 | -3 | 0.37065 | 0.12788 | 0.13061 | 0.21023 | 21.9338 | 0.095967096 | 0 | Neutral |
| testneg_5179 | 0 | 0 | 1.329031 | -3 | 2.859373 | -0.405495 | 0.22489 | 0.16646 | 0.19721 | 0.12485 | 10.83333 | 0.05 | 0 | Neutral |
| testneg_5180 | 0.04 | -0.23 | 3.013855 | -3 | -0.232894 | -3 | 0.20152 | 0.09895 | 0.09724 | 0.09172 | 12.40647 | 0.02039881 | 0 | Neutral |
| testneg_5181 | -0.03 | 1.03 | 0.958989 | -3 | 1.232154 | -3 | 0.39436 | 0.16446 | 0.17768 | 0.10832 | 18.39641 | 0.05 | 0 | Neutral |
| testneg_5182 | 0.06 | 0.18 | 0.135134 | -3 | 3.557638 | -3 | 0.05932 | 0.09551 | 0.15867 | 0.09641 | 19.05506 | 0.070272401 | 0 | Neutral |
| testneg_5183 | 0.07 | 0.38 | -0.84789 | -3 | 5.853414 | -3 | 0.17146 | 0.19882 | 0.19832 | 0.19622 | 10.41667 | 0.539152165 | 66.8 | Disease_causing |
| testneg_5184 | 0.01 | -1.44 | 3.574592 | -3 | 2.504825 | -3 | 0.09493 | 0.104 | 0.09184 | 0.38254 | 28.93023 | 0.385 | 10.6 | Neutral |
| testneg_5185 | 0 | 0 | 1.125734 | -3 | -0.575222 | -1.457001 | 0.12576 | 0.08486 | 0.25119 | 0.13626 | 17.29012 | 0.030262626 | 0 | Neutral |
| testneg_5186 | 0.15 | -0.28 | 0.937469 | -3 | 2.765662 | -3 | 0.58677 | 0.11579 | 0.10438 | 0.20558 | 12.6495 | 0.178770842 | 0 | Neutral |
| testneg_5187 | 0 | 0 | 1.700589 | -3 | 4.157651 | -3 | 0.89357 | 0.22348 | 0.19898 | 0.12788 | 17.21847 | 0.093333333 | 0 | Neutral |
| testneg_5188 | 0 | 0 | 0.535401 | -3 | 1.366812 | -3 | 0.605 | 0.10748 | 0.19898 | 0.12788 | 17.21847 | 0.000259784 | 0 | Neutral |
| testneg_5189 | 0 | 0 | 0.371766 | -3 | 4.563916 | -3 | 0.3095 | 0.13502 | 0.27806 | 0.09713 | 21.42564 | 0.065178433 | 0 | Neutral |
| testneg_5190 | -0.14 | 0.37 | -2.312779 | -3 | 0.254524 | -3 | 0.07617 | 0.07182 | 0.38942 | 0.345 | 16.68085 | 0.11 | 0 | Neutral |
| testneg_5191 | -0.03 | -0.1 | 0.056976 | -3 | 0.212473 | -3 | 0.12048 | 0.2381 | 0.25473 | 0.07863 | 16.37126 | 0.04 | 0 | Neutral |
| testneg_5192 | 0 | 0 | -0.633568 | -3 | 1.420779 | -3 | 0.23027 | 0.37171 | 0.14386 | 0.12788 | 15.41667 | 0.024431818 | 0 | Neutral |
| testneg_5193 | 0 | 0 | -0.100514 | -3 | 11.238372 | -3 | 0.16469 | 0.16042 | 0.05657 | 0.14812 | 11.25 | 0.645491201 | 94.8 | Disease_causing |
| testneg_5194 | 0 | 0 | -0.3269 | -3 | 5.806574 | -3 | 0.11072 | 0.38259 | 0.47274 | 0.11928 | 10.41667 | 0.206839744 | 0.2 | Neutral |
| testneg_5195 | -0.09 | 1.07 | 3.025347 | -3 | 3.095722 | 3.095722 | 0.19969 | 0.1163 | 0.21967 | 0.09467 | 30.28249 | 0.25 | 0.2 | Neutral |
| testneg_5196 | 0 | 0 | 0.510259 | -3 | 3.156541 | -3 | 0.08186 | 0.11813 | 0.19237 | 0.53191 | 20 | 0.160097403 | 0 | Neutral |
| testneg_5197 | -0.01 | 0.57 | 0.47202 | -3 | -0.474175 | -3 | 0.13295 | 0.12914 | 0.3414 | 0.13575 | 36.40434 | 0.064808726 | 0 | Neutral |
| testneg_5198 | 0 | 0 | -0.090459 | -3 | 0.735095 | -3 | 0.15581 | 0.19351 | 0.90941 | 0.28104 | 5.73099 | 0.050814456 | 0 | Neutral |
| testneg_5199 | -0.01 | 0.57 | 0.554274 | -3 | 12.398706 | -3 | 0.32963 | 0.1035 | 0.14767 | 0.09559 | 23.76932 | 0.354166667 | 6.6 | Neutral |
| testneg_5200 | 0 | 0 | -0.260149 | -3 | 1.279217 | -3 | 0.16418 | 0.12788 | 0.12357 | 0.14417 | 9.79167 | 0.0345 | 0 | Neutral |
| testneg_5201 | -0.01 | -0.79 | 1.796129 | -3 | 2.335753 | -3 | 0.16052 | 0.12788 | 0.27789 | 0.17365 | 23.68839 | 0.117545635 | 0 | Neutral |
| testneg_5202 | -0.15 | -0.04 | 2.6592 | -3 | -0.586534 | -0.586534 | 0.24524 | 0.10803 | 0.09439 | 0.1884 | 27.64775 | 0.144166667 | 0 | Neutral |
| testneg_5203 | 0.09 | -0.67 | 1.718854 | -3 | -0.244679 | -3 | 0.19898 | 0.12788 | 0.16381 | 0.09851 | 12.29128 | 0.01 | 0 | Neutral |
| testneg_5204 | 0 | 0 | 1.323079 | -3 | -0.379975 | -3 | 0.5103 | 0.10923 | 0.05817 | 0.09497 | 28.22581 | 0.02 | 0 | Neutral |
| testneg_5205 | 0 | 0 | 0.730138 | -3 | -1.285414 | -3 | 0.16952 | 0.10613 | 0.0693 | 0.09293 | 24.56362 | 0.01 | 0 | Neutral |
| testneg_5206 | 0.03 | 0.1 | 5.424976 | -3 | 0.311006 | -3 | 0.14956 | 0.10585 | 0.10042 | 0.09296 | 26.78006 | 0.1 | 0 | Neutral |
| testneg_5207 | 0 | -0.32 | 3.202792 | -3 | 3.966623 | -3 | 0.28449 | 0.10981 | 0.16737 | 0.12788 | 18.88889 | 0.133439705 | 0 | Neutral |
| testneg_5208 | 0.01 | 0.79 | -0.188304 | -3 | -0.124213 | -0.124213 | 0.16418 | 0.12788 | 0.74641 | 0.66512 | 4.72222 | 0.06118006 | 0 | Neutral |
| testneg_5209 | 0.07 | 0.38 | 1.124718 | 1.124718 | 1.418556 | -3 | 0.13356 | 0.09963 | 0.23844 | 0.27772 | 10.26553 | 0.120757143 | 0 | Neutral |
| testneg_5210 | 0 | 0 | 1.367025 | -3 | -0.337044 | -3 | 0.34467 | 0.13076 | 0.02396 | 0.12703 | 26.57837 | 0.02 | 0 | Neutral |
| testneg_5211 | 0.02 | -0.63 | 5.718017 | -3 | -0.301372 | -3 | 0.11351 | 0.10667 | 0.11019 | 0.08132 | 18.71981 | 0.098333333 | 0 | Neutral |
| testneg_5212 | 0 | 0 | 0.089128 | -3 | 0.088089 | -3 | 0.34495 | 0.16861 | 0.12051 | 0.08503 | 8.97898 | 0.010275875 | 0 | Neutral |
| testneg_5213 | 0 | 0 | 0.494101 | -3 | 3.840666 | -3 | 0.28038 | 0.11814 | 0.1675 | 0.09413 | 28.73135 | 0.05072421 | 0 | Neutral |
| testneg_5214 | 0 | 0 | 1.605285 | -3 | 0.230515 | -3 | 0.5068 | 0.12276 | 0.19898 | 0.1092 | 17.21847 | 0.000259784 | 0 | Neutral |
| testneg_5215 | -0.16 | 1.37 | 5.809146 | -3 | 6.643804 | -3 | 0.11359 | 0.08631 | 0.12921 | 0.11605 | 16.76768 | 0.53 | 67.2 | Disease_causing |
| testneg_5216 | 0 | 0 | 1.010818 | -3 | -1.902903 | -3 | 0.36793 | 0.15588 | 0.1254 | 0.08697 | 10.83333 | 0.015 | 0 | Neutral |
| testneg_5217 | 0.07 | 0.7 | 4.447974 | -3 | 3.410902 | -3 | 0.04869 | 0.05922 | 0.20701 | 0.10345 | 24.21356 | 0.161793586 | 0 | Neutral |
| testneg_5218 | 0 | -0.01 | 0.912506 | -3 | 0.363314 | -3 | 0.38429 | 0.11956 | 0.10286 | 0.26819 | 18.33333 | 0.050781944 | 0 | Neutral |
| testneg_5219 | 0 | -0.01 | 0.888382 | -3 | 3.181514 | -3 | 0.04808 | 0.10018 | 0.21908 | 0.12788 | 18.34559 | 0.06 | 0 | Neutral |
| testneg_5220 | 0.02 | -0.63 | 2.887218 | -3 | 0.388086 | -3 | 0.3962 | 0.10523 | 0.15432 | 0.095 | 10.41667 | 0.03 | 0 | Neutral |
| testneg_5221 | 0 | 0 | 1.771975 | -3 | 2.582897 | -3 | 0.89357 | 0.22348 | 0.19898 | 0.12788 | 18.76225 | 0.037749204 | 0 | Neutral |
| testneg_5222 | 0 | -0.01 | 1.042018 | -3 | 2.570804 | -3 | 0.73808 | 0.15588 | 0.29357 | 0.79524 | 16.84579 | 0.145800793 | 0 | Neutral |
| testneg_5223 | -0.09 | 0.67 | 5.271692 | -3 | 1.519913 | -3 | 0.89959 | 0.55078 | 0.16436 | 0.19109 | 16.84579 | 0.55 | 68 | Disease_causing |
| testneg_5224 | 0.03 | 0.12 | 2.051799 | -3 | 3.390375 | -3 | 0.14624 | 0.10369 | 0.25462 | 0.12788 | 17.21847 | 0.039772727 | 0 | Neutral |
| testneg_5225 | -0.0010909 | 0.07645455 | 2.87279 | -3 | -0.459078 | -3 | 0.32406 | 0.28376 | 0.19214 | 0.12788 | 16.88659 | 0.060322581 | 0 | Neutral |
| testneg_5226 | -0.22 | 2.66 | 0.984298 | -3 | 6.773601 | -3 | 0.34406 | 0.09894 | 0.19898 | 0.12788 | 29.9024 | 0.310330113 | 2 | Neutral |
| testneg_5227 | 0.02 | -0.63 | 1.806186 | 1.806186 | 2.908555 | -3 | 0.19021 | 0.09379 | 0.14175 | 0.12788 | 18.87307 | 0.09 | 0 | Neutral |
| testneg_5228 | 0.18 | -1.05 | 6.384552 | -3 | 0.052319 | -3 | 0.05873 | 0.09792 | 0.15677 | 0.18537 | 13.51351 | 0.37 | 11.8 | Neutral |
| testneg_5229 | -0.1 | -1.9 | 0.774432 | -3 | 3.355302 | -3 | 0.18013 | 0.25346 | 0.21348 | 0.10616 | 18.88889 | 0.15 | 0 | Neutral |
| testneg_5230 | 0.14 | -1.01 | 2.821548 | -3 | 1.464574 | 1.464574 | 0.18578 | 0.27854 | 0.60603 | 0.12788 | 10.83333 | 0.100769231 | 0 | Neutral |
| testneg_5231 | 0.04 | -0.23 | 3.061005 | -3 | 0.84675 | -3 | 0.09269 | 0.12788 | 0.4358 | 0.1222 | 20.89286 | 0.04 | 0 | Neutral |
| testneg_5232 | 0.14 | -0.37 | 3.650869 | -3 | 0.148971 | -3 | 0.07961 | 0.11474 | 0.17007 | 0.12771 | 24.48718 | 0.03 | 0 | Neutral |
| testneg_5233 | 0 | 0 | -0.88087 | -3 | 0.659043 | -3 | 0.38135 | 0.22085 | 0.03137 | 0.12788 | 30.55416 | 0.034857143 | 0 | Neutral |
| testneg_5234 | 0.16 | 0.08 | -0.896036 | -3 | 0.981816 | -3 | 0.11956 | 0.3237 | 0.19416 | 0.0901 | 18.6768 | 0.07 | 0 | Neutral |
| testneg_5235 | -0.05 | -0.29 | -1.315358 | -3 | 1.687932 | -3 | 0.58984 | 0.15588 | 0.16299 | 0.10454 | 18.8735 | 0.03 | 0 | Neutral |
| testneg_5236 | -0.06 | 0.57 | 3.864502 | -3 | 0.84675 | -3 | 0.5471 | 0.11262 | 0.4358 | 0.1222 | 25.72409 | 0.07 | 0 | Neutral |
| testneg_5237 | 0 | 0 | 0.171876 | -3 | 0.997498 | -3 | 0.19969 | 0.1163 | 0.13271 | 0.22958 | 22.58454 | 0.05891351 | 0 | Neutral |
| testneg_5238 | 0.17 | 0.38 | 3.649039 | -3 | 3.443216 | -3 | 0.2877 | 0.18291 | 0.12871 | 0.29252 | 10.83333 | 0.712042409 | 99.4 | Disease_causing |
| testneg_5239 | 0 | 0 | 0.27864 | -3 | 4.140125 | -3 | 0.17737 | 0.54971 | 0.09184 | 0.38254 | 12.34649 | 0.47073054 | 43.4 | Neutral |
| testneg_5240 | 0.04 | -0.23 | 1.86404 | -3 | 1.699554 | -3 | 0.21843 | 0.85214 | 0.16121 | 0.10081 | 40.33208 | 0.108433333 | 0 | Neutral |
| testneg_5241 | 0.07 | 0.01 | -0.285825 | -3 | -0.024667 | -0.024667 | 0.15016 | 0.08826 | 0.59599 | 0.12477 | 18.33333 | 0.01 | 0 | Neutral |
| testneg_5242 | 0.03 | -1.03 | 2.660158 | -3 | 1.893832 | 0.295108 | 0.18243 | 0.11102 | 0.09527 | 0.12788 | 10.41667 | 0.03 | 0 | Neutral |
| testneg_5243 | 0 | 0 | 1.504538 | -3 | 1.890781 | -3 | 0.19898 | 0.12788 | 0.37424 | 0.12788 | 10.27548 | 0 | 0 | Neutral |
| testneg_5244 | -0.21 | -0.15 | 5.223796 | -3 | 0.904012 | -3 | 0.21814 | 0.13855 | 0.17529 | 0.09662 | 35.72379 | 0.15 | 0 | Neutral |
| testneg_5245 | 0 | 0 | 1.69763 | -3 | 3.046889 | -3 | 0.9862 | 0.4528 | 0.1066 | 0.12788 | 10.26467 | 0.07 | 0 | Neutral |

| | | | | | | | | | | | | | | |
|---|---|---|---|---|---|---|---|---|---|---|---|---|---|---|
| testneg_5246 | 0.14 | -1.01 | 6.115432 | -3 | 5.935147 | -3 | 0.19898 | 0.11167 | 0.19898 | 0.10269 | 26.01095 | 0.418216846 | 28.8 | Neutral |
| testneg_5247 | 0.14 | -0.37 | 4.664515 | 1.43553 | -0.036235 | -3 | 0.13565 | 0.46572 | 0.21118 | 0.10074 | 16.8018 | 0.19 | 0 | Neutral |
| testneg_5248 | 0 | 0 | 1.805224 | -3 | 1.285186 | 1.285186 | 0.29679 | 0.12788 | 0.35957 | 0.11476 | 11.58727 | 0.01 | 0 | Neutral |
| testneg_5249 | -0.04 | 0.23 | 1.397648 | -3 | 6.933448 | -3 | 0.11006 | 0.20781 | 0.1059 | 0.1265 | 19.44444 | 0.483592949 | 47.4 | Neutral |
| testneg_5250 | 0.004 | -0.4576364 | 7.731489 | -3 | -0.524108 | -3 | 0.03378 | 0.12788 | 0.08531 | 0.12788 | 27.22222 | 0.325833333 | 7 | Neutral |
| testneg_5251 | -0.15 | 0.28 | 0.27384 | -3 | 0.186761 | -3 | 0.17173 | 0.15363 | 0.19898 | 0.12788 | 23.23037 | 0.08 | 0 | Neutral |
| testneg_5252 | 0 | 0 | 1.988879 | -3 | 1.165033 | -3 | 0.46723 | 0.08832 | 0.86513 | 0.31355 | 12.32809 | 0.030357257 | 0 | Neutral |
| testneg_5253 | -0.03 | -0.1 | -0.001612 | -3 | 3.051125 | -3 | 0.34205 | 0.11195 | 0.0709 | 0.07843 | 19.09347 | 0.041055556 | 0 | Neutral |
| testneg_5254 | 0 | 0 | 0.192861 | -3 | 6.353931 | -3 | 0.85508 | 0.12634 | 0.27605 | 0.12788 | 9.78924 | 0.081972658 | 0 | Neutral |
| testneg_5255 | 0 | 0 | 1.16373 | -3 | 3.543418 | -3 | 0.28416 | 0.27805 | 0.98833 | 0.48581 | 11.25 | 0.298443643 | 4.8 | Neutral |
| testneg_5256 | 0.03 | 1.22 | 6.540942 | -0.139941 | -0.092284 | -3 | 0.12203 | 0.13515 | 0.38695 | 0.12579 | 10.83333 | 0.287083333 | 2.6 | Neutral |
| testneg_5257 | 0.04 | -0.23 | 2.151559 | -3 | 3.264791 | -3 | 0.82591 | 0.12207 | 0.11788 | 0.12788 | 10.41667 | 0.053985903 | 0 | Neutral |
| testneg_5258 | -0.07 | -0.38 | 2.406589 | -3 | 1.149701 | -3 | 0.1205 | 0.23508 | 0.31634 | 0.52685 | 10.83333 | 0.225230782 | 0.2 | Neutral |
| testneg_5259 | 0.16 | -1.37 | -0.461178 | -3 | 2.584779 | -3 | 0.04568 | 0.08746 | 0.19102 | 0.12788 | 27.22994 | 0.08 | 0 | Neutral |
| testneg_5260 | 0.03 | 1.22 | 2.753966 | 2.753966 | 1.802906 | -3 | 0.1551 | 0.09276 | 0.33886 | 0.15644 | 19.23346 | 0.4627 | 41.8 | Neutral |
| testneg_5261 | -0.02 | 0.63 | 0.385855 | -3 | 6.012126 | -3 | 0.10153 | 0.13343 | 0.48585 | 0.66989 | 19.09347 | 0.48558029 | 49.4 | Neutral |
| testneg_5262 | 0 | 0 | 0.470493 | -3 | -2.421511 | -3 | 0.42802 | 0.12096 | 0.37464 | 0.12788 | 31.02368 | 0.01 | 0 | Neutral |
| testneg_5263 | 0 | 0 | 1.406675 | -3 | 2.299454 | -3 | 0.16952 | 0.10613 | 0.19898 | 0.09316 | 22.17046 | 0.002448107 | 0 | Neutral |
| testneg_5264 | 0 | 0 | -0.200198 | -3 | -1.14689 | -3 | 0.2771 | 0.14453 | 0.14704 | 0.12788 | 16.42913 | 0.01 | 0 | Neutral |
| testneg_5265 | 0.17 | 0.38 | 2.7281 | -3 | 4.003489 | -3 | 0.33163 | 0.26601 | 0.48504 | 0.12788 | 10 | 0.309511905 | 2.4 | Neutral |
| testneg_5266 | -0.07 | -0.38 | 2.909977 | -3 | 5.076781 | -3 | 0.39087 | 0.10381 | 0.67549 | 0.1273 | 12.29102 | 0.230779082 | 0.2 | Neutral |
| testneg_5267 | 0 | 0 | 0.531358 | -3 | 1.412972 | -3 | 0.19898 | 0.13495 | 0.10565 | 0.08739 | 12.29167 | 0.01 | 0 | Neutral |
| testneg_5268 | 0.22 | -2.66 | 7.081701 | -3 | -0.816744 | -3 | 0.34826 | 0.49454 | 0.05087 | 0.09682 | 12.34649 | 0.368282051 | 9.4 | Neutral |
| testneg_5269 | 0 | 0 | 0.097586 | -3 | 0.104192 | -3 | 0.19898 | 0.13434 | 0.67644 | 0.11323 | 20.65799 | 0.012916667 | 0 | Neutral |
| testneg_5270 | 0 | 0 | 1.686834 | -3 | 1.549087 | -3 | 0.29828 | 0.12788 | 0.3025 | 0.2499 | 11.25 | 0.020170364 | 0 | Neutral |
| testneg_5271 | 0 | 0 | 0.474011 | -3 | 0.772225 | -3 | 0.40283 | 0.34309 | 0.18705 | 0.43522 | 20.27778 | 0.149575907 | 0 | Neutral |
| testneg_5272 | -0.03 | -0.12 | 1.627305 | -3 | -1.696114 | -3 | 0.08448 | 0.17914 | 0.19898 | 0.82835 | 12.79101 | 0.193611111 | 0 | Neutral |
| testneg_5273 | 0 | 0 | 1.91859 | -3 | 2.443393 | -3 | 0.53861 | 0.18604 | 0.17643 | 0.10767 | 5.73716 | 0.040208696 | 0 | Neutral |
| testneg_5274 | 0.03 | 0.12 | 0.636994 | -3 | 0.748865 | -1.39588 | 0.1601 | 0.12546 | 0.0478 | 0.12788 | 13.51351 | 0.01 | 0 | Neutral |
| testneg_5275 | 0.00281818 | -0.0195455 | 2.555582 | -3 | 6.546927 | -3 | 0.65012 | 0.21208 | 0.19898 | 0.12788 | 6.61194 | 0.309178982 | 4.2 | Neutral |
| testneg_5276 | 0.15 | -0.28 | 1.567816 | -3 | -0.151164 | -0.151164 | 0.80979 | 0.5595 | 0.65708 | 0.0936 | 19.90393 | 0.080105729 | 0 | Neutral |
| testneg_5277 | 0 | 0 | 0.916236 | -3 | 1.217842 | -3 | 0.37499 | 0.17803 | 0.15856 | 0.10994 | 10 | 0.01 | 0 | Neutral |
| testneg_5278 | 0 | 0 | 1.318565 | -3 | 2.048706 | -3 | 0.22658 | 0.20646 | 0.46664 | 0.09008 | 12.42141 | 0.021849206 | 0 | Neutral |
| testneg_5279 | 0 | 0 | 1.73925 | -3 | 0.513838 | -3 | 0.12971 | 0.11434 | 0.68004 | 0.12345 | 12.20312 | 0.01 | 0 | Neutral |
| testneg_5280 | 0.04 | -0.23 | 4.454617 | -3 | -1.162831 | -1.162831 | 0.7639 | 0.10408 | 0.13398 | 0.12788 | 18.03715 | 0.04 | 0 | Neutral |
| testneg_5281 | -0.08 | -0.03 | 0.395961 | -3 | 7.473966 | -3 | 0.23027 | 0.37171 | 0.07447 | 0.23492 | 11.875 | 0.721016973 | 98.8 | Disease_causing |
| testneg_5282 | 0.08 | 0.03 | -2.800363 | -3 | -0.825189 | -3 | 0.44103 | 0.10698 | 0.03646 | 0.09863 | 25.2565 | 0.040322581 | 0 | Neutral |
| testneg_5283 | 0.03 | -0.85 | 4.652483 | -3 | 5.312389 | -3 | 0.06841 | 0.10927 | 0.07293 | 0.12989 | 11.25 | 0.31 | 4.6 | Neutral |
| testneg_5284 | -0.09 | 0.25 | 3.716081 | -3 | 0.609952 | 0.609952 | 0.40632 | 0.1094 | 0.06224 | 0.12788 | 16.8018 | 0.045809524 | 0 | Neutral |
| testneg_5285 | 0.16 | 0.08 | 1.706402 | -3 | 0.235328 | -3 | 0.60703 | 0.12378 | 0.16547 | 0.16092 | 17.77778 | 0.099315986 | 0 | Neutral |
| testneg_5286 | 0 | 0 | 1.66597 | -3 | -1.06095 | -3 | 0.66027 | 0.23984 | 0.23393 | 0.12567 | 10.41667 | 0.01 | 0 | Neutral |
| testneg_5287 | 0.03 | 0.1 | -2.618223 | -3 | 4.96273 | -3 | 0.22179 | 0.09831 | 0.09807 | 0.24901 | 17.77778 | 0.290051282 | 1.6 | Neutral |
| testneg_5288 | 0 | 0 | 1.832674 | -3 | 1.75958 | 1.75958 | 0.1589 | 0.10761 | 0.65897 | 0.67602 | 6.98716 | 0.081614285 | 0 | Neutral |
| testneg_5289 | 0 | 0 | 0.342408 | -3 | 1.043597 | 1.043597 | 0.5255 | 0.15968 | 0.19898 | 0.1005 | 22.18259 | 0.030165978 | 0 | Neutral |
| testneg_5290 | -0.17 | -0.38 | 1.808824 | 1.808824 | -0.84394 | -3 | 0.2445 | 0.53069 | 0.54895 | 0.15082 | 4.44444 | 0.27 | 0.6 | Neutral |
| testneg_5291 | -0.14 | 1.01 | 1.855162 | 1.695225 | -1.133182 | -3 | 0.33294 | 0.16155 | 0.06431 | 0.12788 | 13.08333 | 0.11 | 0 | Neutral |
| testneg_5292 | -0.07 | -0.01 | -0.23732 | -0.23732 | 4.523205 | -3 | 0.07473 | 0.51812 | 0.18067 | 0.14759 | 4.72222 | 0.486313707 | 48.6 | Neutral |
| testneg_5293 | 0 | 0 | 2.575019 | -3 | 0.158776 | -3 | 0.28269 | 0.12505 | 0.08529 | 0.12376 | 32.88161 | 0.01 | 0 | Neutral |
| testneg_5294 | 0.02 | 0.45 | 0.982787 | -3 | 3.374283 | -3 | 0.86851 | 0.17715 | 0.19898 | 0.28128 | 10.41667 | 0.289203735 | 1.4 | Neutral |
| testneg_5295 | 0 | 0 | 0.327712 | -3 | -0.269735 | -0.269735 | 0.57517 | 0.10873 | 0.12461 | 0.1241 | 29.9024 | 0.02 | 0 | Neutral |
| testneg_5296 | -0.07 | -0.38 | 3.731276 | -3 | 2.859591 | -3 | 0.82455 | 0.2093 | 0.1576 | 0.12788 | 11.25 | 0.200271609 | 0.2 | Neutral |
| testneg_5297 | 0.16 | -0.37 | 3.271878 | 2.303409 | 0.284017 | -3 | 0.19898 | 0.12788 | 0.3025 | 0.2499 | 16.85532 | 0.4 | 25.8 | Neutral |
| testneg_5298 | 0.008 | 0.24890909 | 4.814261 | -3 | 3.445512 | -3 | 0.08775 | 0.12788 | 0.38126 | 0.27764 | 13.51351 | 0.528406481 | 63.4 | Disease_causing |
| testneg_5299 | 0 | 0 | -1.230837 | -1.230837 | 9.589916 | -3 | 0.14821 | 0.11262 | 0.19898 | 0.08111 | 12.29167 | 0.31 | 2.6 | Neutral |
| testneg_5300 | 0.01 | 0.8 | -0.451626 | -3 | 3.235104 | -3 | 0.17677 | 0.67006 | 0.37212 | 0.11323 | 20.38801 | 0.18 | 0 | Neutral |
| testneg_5301 | 0 | 0 | 0.253071 | -3 | -0.164302 | -3 | 0.0418 | 0.12788 | 0.35475 | 0.12788 | 26.84685 | 0.03 | 0 | Neutral |
| testneg_5302 | 0.14 | -0.37 | 1.830118 | -3 | 2.43417 | -3 | 0.55367 | 0.57993 | 0.10231 | 0.1 | 6.94444 | 0.113858173 | 0 | Neutral |
| testneg_5303 | 0.08 | 0.03 | 0.310184 | -3 | 0.072959 | -3 | 0.3769 | 0.12788 | 0.19861 | 0.12057 | 22.00525 | 0.02 | 0 | Neutral |
| testneg_5304 | 0.21 | 0.15 | 1.305251 | -3 | 1.447263 | -3 | 0.06304 | 0.12788 | 0.19531 | 0.18996 | 28.84995 | 0.12 | 0 | Neutral |
| testneg_5305 | -0.16 | -0.08 | 3.114446 | -3 | 4.640406 | -3 | 0.34209 | 0.09515 | 0.02658 | 0.18737 | 10.83333 | 0.52 | 60.8 | Disease_causing |
| testneg_5306 | 0 | 0 | 2.918397 | -3 | 6.588654 | -3 | 0.63378 | 0.09451 | 0.10422 | 0.0994 | 12.94141 | 0.2545 | 1.2 | Neutral |
| testneg_5307 | -0.03 | -1.22 | 2.92942 | -3 | 0.868157 | -3 | 0.15107 | 0.10457 | 0.06526 | 0.09018 | 20.98109 | 0.04 | 0 | Neutral |
| testneg_5308 | -0.0356364 | 0.22672727 | -1.616402 | -3 | 1.582649 | -3 | 0.07837 | 0.12788 | 0.0501 | 0.12788 | 20.34752 | 0.03 | 0 | Neutral |
| testneg_5309 | -0.03 | -0.12 | 1.782773 | -3 | 6.556357 | -3 | 0.42544 | 0.11809 | 0.12465 | 0.09534 | 26.34995 | 0.187291667 | 0.2 | Neutral |
| testneg_5310 | 0 | 0 | 0.760575 | -3 | -0.196592 | -0.196592 | 0.1878 | 0.08946 | 0.32733 | 0.16869 | 16.67431 | 0.04 | 0 | Neutral |
| testneg_5311 | 0.03 | 0.1 | 6.254981 | -3 | 4.838148 | -3 | 0.95993 | 0.15588 | 0.1576 | 0.12788 | 18.09499 | 0.609259921 | 87.4 | Disease_causing |
| testneg_5312 | 0 | 0 | -0.58291 | -3 | 5.036141 | -1.537769 | 0.64608 | 0.35524 | 0.04787 | 0.12788 | 10.54139 | 0.113456326 | 0 | Neutral |
| testneg_5313 | -0.02 | -0.45 | 3.546645 | -3 | 0.154673 | -3 | 0.18502 | 0.21329 | 0.04484 | 0.95426 | 10.41667 | 0.276666667 | 1 | Neutral |
| testneg_5314 | 0 | 0 | 1.900871 | -3 | 0.103447 | 0.103447 | 0.92514 | 0.19444 | 0.15813 | 0.61197 | 5 | 0.110829519 | 0 | Neutral |
| testneg_5315 | -0.08 | -0.03 | 2.768711 | 2.768711 | 4.623093 | -3 | 0.38324 | 0.25193 | 0.0774 | 0.1702 | 27.5731 | 0.741333333 | 99.4 | Disease_causing |
| testneg_5316 | -0.02 | 0.36 | 3.437388 | -3 | 1.897319 | -3 | 0.22645 | 0.1704 | 0.39225 | 0.10625 | 19.82027 | 0.080119682 | 0 | Neutral |
| testneg_5317 | 0.05 | 0.29 | -1.828653 | -3 | 2.135113 | -3 | 0.4097 | 0.36733 | 0.40391 | 0.13431 | 10 | 0.200658897 | 0 | Neutral |
| testneg_5318 | 0.09 | -0.3 | 6.559945 | -3 | 5.858753 | -3 | 0.06072 | 0.09873 | 0.12403 | 0.12788 | 10.83333 | 0.569545455 | 76.6 | Disease_causing |
| testneg_5319 | 0.05 | -0.81 | -0.842653 | -3 | 0.770086 | 0.770086 | 0.06708 | 0.09234 | 0.20527 | 0.10053 | 16.8018 | 0.02 | 0 | Neutral |
| testneg_5320 | 0 | -0.01 | 0.857165 | -3 | 0.025863 | -3 | 0.14635 | 0.13588 | 0.21755 | 0.23824 | 29.86111 | 0.060477273 | 0 | Neutral |
| testneg_5321 | -0.0016667 | -0.007197 | 3.942584 | -3 | 2.740775 | -3 | 0.961 | 0.25991 | 0.10379 | 0.34799 | 13.19331 | 0.516738854 | 61.4 | Disease_causing |
| testneg_5322 | -0.08 | -0.03 | -0.949463 | -3 | 1.26259 | 1.26259 | 0.09018 | 0.16175 | 0.44722 | 0.11641 | 13.84202 | 0.07 | 0 | Neutral |
| testneg_5323 | 0.05 | 0.29 | 1.381522 | -3 | 2.232989 | -3 | 0.11325 | 0.17021 | 0.22998 | 0.08738 | 27.40077 | 0.042457189 | 0 | Neutral |
| testneg_5324 | 0 | 0 | 1.712042 | -3 | 0.643669 | -3 | 0.31004 | 0.09727 | 0.39932 | 0.10693 | 17.05011 | 0 | 0 | Neutral |
| testneg_5325 | -0.05 | -0.29 | 3.001619 | -3 | -1.996459 | -3 | 0.16078 | 0.39498 | 0.04988 | 0.25503 | 10 | 0.346314935 | 6.2 | Neutral |
| testneg_5326 | 0 | 0 | 1.431043 | -3 | 2.393948 | 2.393948 | 0.08489 | 0.12788 | 0.70819 | 0.5975 | 11.00193 | 0.150557143 | 0 | Neutral |
| testneg_5327 | 0 | 0 | 1.052386 | -3 | 5.557447 | -3 | 0.87038 | 0.43956 | 0.34633 | 0.47228 | 9.71235 | 0.348011364 | 10.8 | Neutral |
| testneg_5328 | 0.15 | 0.04 | -0.962385 | -3 | 3.894139 | -3 | 0.60285 | 0.15179 | 0.18147 | 0.12788 | 16.47059 | 0.183024227 | 0 | Neutral |
| testneg_5329 | 0 | 0 | 0.323774 | -3 | 5.153595 | -3 | 0.60755 | 0.41186 | 0.85781 | 0.22949 | 14.16667 | 0.297000645 | 3.8 | Neutral |
| testneg_5330 | 0 | -0.01 | 0.334215 | -3 | 3.445512 | -3 | 0.33314 | 0.47999 | 0.38126 | 0.27764 | 29.14332 | 0.359604787 | 9.4 | Neutral |
| testneg_5331 | 0 | 0 | 0.955737 | -3 | 3.042989 | -3 | 0.24528 | 0.37575 | 0.35584 | 0.10874 | 22.6322 | 0.073480392 | 0 | Neutral |
| testneg_5332 | 0 | 0 | -0.68826 | -3 | 2.159193 | -3 | 0.32076 | 0.14229 | 0.45399 | 0.10245 | 16.113 | 0.010170953 | 0 | Neutral |
| testneg_5333 | 0.07 | 0.01 | 2.592582 | -3 | 0.100922 | -3 | 0.4404 | 0.38303 | 0.13061 | 0.21023 | 11.93907 | 0.187205744 | 0 | Neutral |
| testneg_5334 | -0.06 | -0.18 | 0.867502 | -3 | -0.313201 | -3 | 0.71511 | 0.34803 | 0.16524 | 0.12788 | 10 | 0.029 | 0 | Neutral |
| testneg_5335 | -0.0009091 | -0.0219091 | 2.018667 | -3 | -0.25961 | -3 | 0.39344 | 0.12788 | 0.31773 | 0.10816 | 14.34932 | 0.000580953 | 0 | Neutral |
| testneg_5336 | 0.08 | 0.03 | 0.709755 | -3 | 3.52493 | -3 | 0.45014 | 0.12788 | 0.04916 | 0.14944 | 10.35388 | 0.285604401 | 0.4 | Neutral |
| testneg_5337 | 0.21 | 0.15 | 3.237386 | -3 | 1.81436 | -3 | 0.28499 | 0.10637 | 0.20357 | 0.12788 | 12.43609 | 0.04016313 | 0 | Neutral |
| testneg_5338 | 0.04 | -0.23 | 1.86404 | -3 | -1.839743 | -3 | 0.21843 | 0.85214 | 0.12446 | 0.08908 | 16.42913 | 0.08 | 0 | Neutral |
| testneg_5339 | -0.07 | -0.38 | 1.85883 | -3 | 1.517891 | -3 | 0.25009 | 0.11017 | 0.42755 | 0.09484 | 19 | 0.01 | 0 | Neutral |
| testneg_5340 | -0.21 | -0.15 | -0.83994 | -3 | 0.272659 | 0.272659 | 0.95319 | 0.20297 | 0.17093 | 0.23281 | 9.71235 | 0.151141762 | 0 | Neutral |
| testneg_5341 | -0.17 | -0.38 | -1.21454 | -3 | 4.056427 | -3 | 0.07604 | 0.08642 | 0.32677 | 0.12788 | 20.83333 | 0.17 | 0 | Neutral |
| testneg_5342 | 0.07 | 0.38 | 3.226843 | -3 | 1.240379 | -3 | 0.85252 | 0.83576 | 0.64591 | 0.38907 | 10.41667 | 0.348016487 | 6 | Neutral |
| testneg_5343 | 0.21 | 0.15 | 4.361375 | -3 | 0.82266 | -3 | 0.10459 | 0.12783 | 0.07923 | 0.12788 | 27.4024 | 0.051833333 | 0 | Neutral |
| testneg_5344 | 0 | 0 | 1.716798 | -3 | 3.268557 | -3 | 0.70349 | 0.12788 | 0.09658 | 0.11286 | 29.86111 | 0.061511905 | 0 | Neutral |
| testneg_5345 | -0.05 | -0.29 | 1.417982 | -3 | 0.48539 | -3 | 0.21811 | 0.18489 | 0.76924 | 0.15588 | 33.40179 | 0.100351899 | 0 | Neutral |
| testneg_5346 | 0.06 | -0.57 | 0.234129 | -3 | 1.673476 | -3 | 0.25758 | 0.2262 | 0.40361 | 0.17821 | 11.25 | 0.14 | 0 | Neutral |
| testneg_5347 | 0 | 0 | 1.319843 | -3 | 0.809376 | -3 | 0.04056 | 0.55024 | 0.11522 | 0.11925 | 10.08333 | 0.1 | 0 | Neutral |
| testneg_5348 | 0.09 | -0.3 | 6.314583 | -3 | 0.450104 | -3 | 0.42583 | 0.10901 | 0.18067 | 0.14759 | 16.96497 | 0.281274831 | 3.2 | Neutral |
| testneg_5349 | -0.18 | 1.05 | 2.778725 | 2.778725 | 0.591914 | -3 | 0.33528 | 0.29463 | 0.88976 | 0.22648 | 10.625 | 0.54 | 66.2 | Disease_causing |
| testneg_5350 | 0 | 0 | 0.981764 | -3 | 5.844336 | -3 | 0.56904 | 0.14839 | 0.14214 | 0.09456 | 31.90299 | 0.13 | 0 | Neutral |
| testneg_5351 | 0.12 | -1.16 | 2.220096 | -3 | 0.792263 | -3 | 0.4003 | 0.10884 | 0.25447 | 0.12788 | 27.90854 | 0.020244048 | 0 | Neutral |
| testneg_5352 | 0.14 | -1.01 | 0.023757 | -3 | 6.211821 | -3 | 0.3221 | 0.3367 | 0.1215 | 0.12788 | 10.83333 | 0.3 | 3 | Neutral |
| testneg_5353 | 0 | 0 | 0.013394 | -3 | 5.721753 | -3 | 0.12582 | 0.12788 | 0.62339 | 0.19592 | 17.21847 | 0.28185658 | 2.4 | Neutral |
| testneg_5354 | 0.03 | 0.1 | -0.942639 | -3 | -0.713943 | -3 | 0.05996 | 0.12788 | 0.20547 | 0.12178 | 21.8759 | 0.02 | 0 | Neutral |
| testneg_5355 | -0.03 | -0.1 | 1.975689 | -3 | 3.39754 | -3 | 0.99665 | 0.12884 | 0.62203 | 0.15583 | 11.875 | 0.203537296 | 0 | Neutral |
| testneg_5356 | 0.09 | -0.3 | 5.235545 | -3 | -0.876313 | -3 | 0.09998 | 0.09022 | 0.08973 | 0.07662 | 12.95163 | 0.08 | 0 | Neutral |
| testneg_5357 | 0.06 | -0.57 | 4.429067 | -3 | -0.029956 | -3 | 0.19898 | 0.09824 | 0.06296 | 0.09823 | 16.48204 | 0.03 | 0 | Neutral |
| testneg_5358 | -0.15 | -0.04 | 1.905797 | -3 | 0.897605 | -3 | 0.13906 | 0.09925 | 0.08723 | 0.09503 | 35.34347 | 0.04 | 0 | Neutral |
| testneg_5359 | 0.14 | -1.01 | 3.493531 | -3 | 5.770752 | -3 | 0.09997 | 0.1797 | 0.26192 | 0.15452 | 22.36648 | 0.771994509 | 99.8 | Disease_causing |
| testneg_5360 | 0 | -0.32 | 4.678621 | -3 | -1.111616 | -3 | 0.86373 | 0.11969 | 0.05444 | 0.12788 | 29.94567 | 0.060353598 | 0 | Neutral |
| testneg_5361 | -0.05 | 0.81 | -0.593986 | -0.593986 | 1.55793 | -3 | 0.31732 | 0.27963 | 0.39305 | 0.18672 | 16.37545 | 0.150247059 | 0 | Neutral |
| testneg_5362 | 0 | 0 | 2.238523 | -3 | 0.953557 | -3 | 0.59947 | 0.10764 | 0.09522 | 0.20323 | 35.43176 | 0.09 | 0 | Neutral |
| testneg_5363 | 0 | 0 | 0.561287 | -3 | 0.941706 | -3 | 0.3797 | 0.1257 | 0.15294 | 0.12788 | 18.33333 | 0 | 0 | Neutral |
| testneg_5364 | 0 | 0 | 0.309568 | -3 | 0.645185 | -3 | 0.54724 | 0.12788 | 0.03898 | 0.09943 | 12.69679 | 0.010446429 | 0 | Neutral |
| testneg_5365 | 0 | 0 | 4.037148 | -3 | 1.88502 | -3 | 0.92832 | 0.16306 | 0.06744 | 0.12788 | 16.59347 | 0.08 | 0 | Neutral |
| testneg_5366 | 0.14 | -1.01 | 2.403577 | -3 | 4.659357 | -3 | 0.99417 | 0.17365 | 0.10084 | 0.12788 | 18.33333 | 0.237395833 | 0 | Neutral |
| testneg_5367 | 0 | -0.01 | 2.305644 | -3 | 0.118932 | 0.118932 | 0.47787 | 0.25991 | 0.19567 | 0.09606 | 14.95396 | 0.025833333 | 0 | Neutral |

| | | | | | | | | | | | | | | |
|---|---|---|---|---|---|---|---|---|---|---|---|---|---|---|
| testneg_5368 | 0 | 0 | 1.269701 | -3 | 0.692563 | -3 | 0.84236 | 0.14093 | 0.39648 | 0.97288 | 5.70654 | 0.06 | 0 | Neutral |
| testneg_5369 | 0 | 0 | -0.239359 | -3 | 0.253348 | -3 | 0.24287 | 0.11626 | 0.72946 | 0.28603 | 10.87198 | 0.02 | 0 | Neutral |
| testneg_5370 | 0.14 | -1.01 | 3.072449 | -3 | -1.23888 | -3 | 0.17529 | 0.09662 | 0.19898 | 0.16603 | 35.72379 | 0.17 | 0 | Neutral |
| testneg_5371 | -0.03 | -0.1 | 1.757173 | -3 | 4.020236 | -3 | 0.21726 | 0.23214 | 0.1109 | 0.12788 | 10.22898 | 0.181102273 | 0 | Neutral |
| testneg_5372 | 0 | 0 | 0.440494 | 0.440494 | 6.948148 | -3 | 0.61027 | 0.14229 | 0.35325 | 0.12614 | 11.83526 | 0.251788889 | 1.6 | Neutral |
| testneg_5373 | -0.06 | -0.18 | 2.632596 | -3 | 3.024546 | -3 | 0.9662 | 0.28104 | 0.31651 | 0.12788 | 16.07106 | 0.128205815 | 0 | Neutral |
| testneg_5374 | 0 | 0 | -0.998522 | -0.998522 | 6.754897 | -3 | 0.04574 | 0.05824 | 0.45997 | 0.15162 | 50 | 0.328663898 | 4.6 | Neutral |
| testneg_5375 | 0 | -0.01 | 0.684698 | -3 | 3.255832 | -3 | 0.09661 | 0.09981 | 0.11564 | 0.1184 | 23.96199 | 0.038619048 | 0 | Neutral |
| testneg_5376 | 0 | 0 | 0.700444 | -3 | 3.4658 | -3 | 0.31664 | 0.14181 | 0.52118 | 0.36073 | 18.88889 | 0.240965659 | 0.8 | Neutral |
| testneg_5377 | 0 | 0 | 2.221516 | -3 | 1.126259 | -3 | 0.16952 | 0.10613 | 0.56817 | 0.10766 | 37.32004 | 0.000289916 | 0 | Neutral |
| testneg_5378 | 0 | 0 | 0.390004 | -3 | 0.218838 | -3 | 0.85252 | 0.83576 | 0.13884 | 0.12788 | 5 | 0.032708333 | 0 | Neutral |
| testneg_5379 | -0.18 | 1.05 | 0.012106 | -3 | -0.180896 | -3 | 0.42755 | 0.09484 | 0.30423 | 0.12788 | 26.9055 | 0.09 | 0 | Neutral |
| testneg_5380 | 0 | 0 | 2.044018 | -3 | 0.007168 | -3 | 0.63431 | 0.09837 | 0.29356 | 0.47581 | 20.76356 | 0.07 | 0 | Neutral |
| testneg_5381 | -0.07 | -0.38 | 2.003914 | -3 | 2.120746 | -3 | 0.14495 | 0.37127 | 0.13369 | 0.09791 | 38.49057 | 0.08 | 0 | Neutral |
| testneg_5382 | -0.05 | -0.29 | 1.124718 | -3 | 0.196897 | -3 | 0.55043 | 0.12112 | 0.74699 | 0.12286 | 10.41667 | 0.01 | 0 | Neutral |
| testneg_5383 | -0.08 | -0.03 | 7.833352 | -3 | 3.557395 | 1.829508 | 0.08067 | 0.34914 | 0.64775 | 0.54233 | 20.83333 | 0.839384164 | 100 | Disease_causing |
| testneg_5384 | -0.05 | -0.29 | 0.235446 | -3 | 5.050499 | -3 | 0.53447 | 0.10345 | 0.12819 | 0.1229 | 13.02635 | 0.088208333 | 0 | Neutral |
| testneg_5385 | 0.14 | -0.37 | 3.758883 | -3 | -0.107298 | -3 | 0.50461 | 0.11034 | 0.09532 | 0.35459 | 13.01229 | 0.21 | 0 | Neutral |
| testneg_5386 | -0.08 | -0.03 | 5.137052 | -3 | -0.018405 | -3 | 0.36709 | 0.19019 | 0.25957 | 0.06969 | 28.39414 | 0.119857143 | 0 | Neutral |
| testneg_5387 | 0.2 | -1.26 | 2.073808 | -3 | 0.247956 | 0.247956 | 0.08683 | 0.26961 | 0.09471 | 0.12788 | 28.79339 | 0.100954545 | 0 | Neutral |
| testneg_5388 | -0.19 | -0.51 | -0.445993 | -0.445993 | 4.859478 | -3 | 0.18324 | 0.13041 | 0.11501 | 0.13748 | 19.44444 | 0.400859649 | 16 | Neutral |
| testneg_5389 | 0.08 | 0.03 | 1.214028 | -3 | 7.121043 | -3 | 0.08136 | 0.10647 | 0.35945 | 0.12025 | 23.51901 | 0.22 | 0.8 | Neutral |
| testneg_5390 | 0 | 0 | 0.232568 | -3 | 5.114974 | -3 | 0.339 | 0.13169 | 0.23962 | 0.10029 | 16.48772 | 0.06 | 0 | Neutral |
| testneg_5391 | -0.06 | -0.18 | 0.491194 | 0.491194 | 5.734507 | -3 | 0.72124 | 0.09906 | 0.06731 | 0.2149 | 25 | 0.386982257 | 11 | Neutral |
| testneg_5392 | 0 | 0 | 0.017266 | -3 | -0.474484 | -3 | 0.1228 | 0.10482 | 0.45075 | 0.12232 | 18.50844 | 0.00011236 | 0 | Neutral |
| testneg_5393 | -0.14 | 0.37 | 0.312719 | -3 | 7.325028 | -3 | 0.21728 | 0.1132 | 0.13143 | 0.90853 | 18.5792 | 0.520208333 | 63.6 | Disease_causing |
| testneg_5394 | 0 | 0 | 1.27406 | -3 | 2.015062 | -3 | 0.37855 | 0.47307 | 0.13559 | 0.1946 | 18.6768 | 0.147828063 | 0 | Neutral |
| testneg_5395 | -0.03 | 1.03 | 3.953236 | -3 | 6.964397 | -3 | 0.07219 | 0.12788 | 0.14495 | 0.37127 | 48.51693 | 0.656042037 | 98 | Disease_causing |
| testneg_5396 | 0 | 0 | 2.21496 | -3 | 6.984226 | -3 | 0.7413 | 0.18974 | 0.87877 | 0.60823 | 10.20833 | 0.473625 | 44.2 | Neutral |
| testneg_5397 | 0.21 | 0.15 | -1.218152 | -3 | 4.711466 | -3 | 0.19898 | 0.12788 | 0.10285 | 0.10886 | 27.95796 | 0.11 | 0 | Neutral |
| testneg_5398 | -0.01 | 1.44 | -0.133106 | -0.133106 | 6.986572 | -3 | 0.07096 | 0.09965 | 0.45343 | 0.12014 | 19.89489 | 0.28 | 1 | Neutral |
| testneg_5399 | 0 | -0.01 | 1.245581 | -3 | 3.391162 | -3 | 0.60793 | 0.18264 | 0.213 | 0.09535 | 18.88889 | 0.071 | 0 | Neutral |
| testneg_5400 | 0 | 0 | -0.049164 | -0.049164 | 4.0939 | -3 | 0.12322 | 0.1246 | 0.18597 | 0.12788 | 20.83333 | 0.070851746 | 0 | Neutral |
| testneg_5401 | 0.22 | -2.66 | 7.446784 | -3 | 4.11462 | -3 | 0.32532 | 0.09651 | 0.54891 | 0.55746 | 5 | 0.710835498 | 98.2 | Disease_causing |
| testneg_5402 | 0 | 0 | 0.532295 | -3 | 8.272947 | 8.272947 | 0.755 | 0.12788 | 0.16374 | 0.14659 | 20 | 0.544429434 | 72.2 | Disease_causing |
| testneg_5403 | 0.04 | -1.36 | 4.588896 | -3 | 1.605285 | -3 | 0.19898 | 0.10512 | 0.5068 | 0.12276 | 18.33333 | 0.06 | 0 | Neutral |
| testneg_5404 | 0 | 0 | 1.641098 | -3 | 3.157424 | -3 | 0.13906 | 0.09925 | 0.09067 | 0.10671 | 13.02635 | 0.03136646 | 0 | Neutral |
| testneg_5405 | 0.21 | 0.15 | 0.153476 | -3 | 0.839368 | 0.839368 | 0.05978 | 0.10176 | 0.18785 | 0.14723 | 44.04381 | 0.12 | 0 | Neutral |
| testneg_5406 | 0 | 0 | 0.634119 | -3 | -0.371298 | -3 | 0.41647 | 0.11239 | 0.06647 | 0.09129 | 18.6768 | 0.000526316 | 0 | Neutral |
| testneg_5407 | 0.03 | 0.12 | 3.408834 | -3 | 5.075013 | -3 | 0.19479 | 0.12788 | 0.47019 | 0.12788 | 19.10253 | 0.204152543 | 0.2 | Neutral |
| testneg_5408 | 0 | 0 | 1.343398 | -3 | 1.121308 | -3 | 0.06926 | 0.08399 | 0.33418 | 0.09979 | 29.47249 | 0.0075 | 0 | Neutral |
| testneg_5409 | 0 | -0.01 | -1.285414 | -3 | 5.055215 | -3 | 0.0693 | 0.09293 | 0.5958 | 0.0929 | 18.33333 | 0.072142857 | 0 | Neutral |
| testneg_5410 | 0 | 0 | -0.88087 | -3 | -0.330275 | -0.330275 | 0.38135 | 0.22085 | 0.10462 | 0.1278 | 12.65469 | 0.03 | 0 | Neutral |
| testneg_5411 | 0 | 0 | 0.0974 | -3 | 6.860523 | -3 | 0.83219 | 0.29765 | 0.10084 | 0.12788 | 10 | 0.25 | 1.8 | Neutral |
| testneg_5412 | 0.04 | -0.23 | 4.063675 | -3 | 0.250187 | -3 | 0.42544 | 0.11809 | 0.07887 | 0.07487 | 24.95885 | 0.03 | 0 | Neutral |
| testneg_5413 | 0 | 0 | 0.335364 | -3 | 5.603216 | -3 | 0.87839 | 0.33588 | 0.06112 | 0.06901 | 11.25 | 0.101464286 | 0 | Neutral |
| testneg_5414 | 0.22 | -2.66 | 6.675004 | -3 | 6.734048 | -3 | 0.92514 | 0.19444 | 0.19898 | 0.48545 | 4.72222 | 0.888901198 | 100 | Disease_causing |
| testneg_5415 | 0.21 | 0.15 | 3.763978 | -3 | 3.002533 | 3.002533 | 0.43736 | 0.10017 | 0.14304 | 0.10715 | 30.90278 | 0.255142857 | 0.2 | Neutral |
| testneg_5416 | 0 | 0 | -0.5391 | -3 | 0.905157 | -3 | 0.16418 | 0.12788 | 0.1019 | 0.17526 | 8.63248 | 0.0550625 | 0 | Neutral |
| testneg_5417 | 0.08 | -1.2 | 6.708423 | -3 | 4.149716 | -3 | 0.26291 | 0.10317 | 0.19322 | 0.12788 | 17.69595 | 0.42952381 | 29.2 | Neutral |
| testneg_5418 | 0 | -0.01 | 0.306573 | -3 | 4.692647 | -3 | 0.03898 | 0.09943 | 0.19898 | 0.12788 | 6.24231 | 0.07 | 0 | Neutral |
| testneg_5419 | 0 | 0 | 0.44595 | -3 | 2.022849 | 2.022849 | 0.15019 | 0.10342 | 0.20708 | 0.12788 | 18.88889 | 0.03 | 0 | Neutral |
| testneg_5420 | 0.09 | -0.3 | 6.461271 | -3 | 3.445512 | -3 | 0.51622 | 0.11809 | 0.38126 | 0.27764 | 34.65752 | 0.626324074 | 91 | Disease_causing |
| testneg_5421 | -0.05 | -0.29 | -1.851788 | -3 | 4.168346 | -3 | 0.78741 | 0.22085 | 0.30042 | 0.13517 | 9.80134 | 0.356640375 | 6.8 | Neutral |
| testneg_5422 | 0 | 0 | 0.90039 | -3 | -1.362764 | -3 | 0.33332 | 0.12106 | 0.08876 | 0.09037 | 13.07136 | 0.000188679 | 0 | Neutral |
| testneg_5423 | 0 | 0 | 0.050569 | -3 | 5.663657 | -3 | 0.16263 | 0.53493 | 0.21989 | 0.10981 | 18.91117 | 0.175946593 | 0 | Neutral |
| testneg_5424 | -0.05 | -0.29 | 3.393473 | -3 | 6.169136 | -3 | 0.05657 | 0.14812 | 0.41655 | 0.49485 | 11.25 | 0.814643899 | 100 | Disease_causing |
| testneg_5425 | -0.02 | 0.63 | 1.253634 | -3 | 1.613091 | -3 | 0.19898 | 0.12788 | 0.53922 | 0.12788 | 10.41667 | 0.02 | 0 | Neutral |
| testneg_5426 | -0.15 | -0.04 | 2.762158 | -3 | 2.850193 | -3 | 0.19898 | 0.12788 | 0.87803 | 0.12788 | 18.88889 | 0.110069444 | 0 | Neutral |
| testneg_5427 | 0 | 0 | -0.491414 | -3 | -1.480749 | -3 | 0.61027 | 0.14229 | 0.19898 | 0.12788 | 24.60546 | 0.03 | 0 | Neutral |
| testneg_5428 | 0.14 | -0.37 | 2.773031 | 1.507693 | 5.078819 | -3 | 0.31732 | 0.27963 | 0.1228 | 0.10482 | 34.52995 | 0.33 | 4 | Neutral |
| testneg_5429 | 0.04 | -0.23 | -1.047288 | -3 | 1.205837 | -3 | 0.52005 | 0.12788 | 0.07071 | 0.11499 | 20.90643 | 0.03 | 0 | Neutral |
| testneg_5430 | 0 | 0 | 1.156713 | 1.156713 | -0.294826 | -3 | 0.36458 | 0.16397 | 0.63818 | 0.12788 | 11.25 | 0.038875 | 0 | Neutral |
| testneg_5431 | -0.02 | 0.63 | 2.174684 | -3 | 0.486603 | -3 | 0.49246 | 0.11033 | 0.82131 | 0.12788 | 21.50794 | 0.03 | 0 | Neutral |
| testneg_5432 | 0.16 | -1.37 | 5.834642 | -3 | 0.846005 | -3 | 0.47751 | 0.2062 | 0.17639 | 0.12919 | 10.41667 | 0.27 | 1.4 | Neutral |
| testneg_5433 | 0 | 0 | 1.108803 | -3 | 2.470276 | -3 | 0.85092 | 0.58871 | 0.06841 | 0.12788 | 24.44444 | 0.08525641 | 0 | Neutral |
| testneg_5434 | -0.07 | -0.01 | 5.490135 | -3 | 0.683746 | -3 | 0.91587 | 0.10818 | 0.15486 | 0.28768 | 20.48674 | 0.37345648 | 10.4 | Neutral |
| testneg_5435 | 0.21 | 0.15 | 6.222747 | -3 | 2.387342 | -3 | 0.1602 | 0.11262 | 0.07428 | 0.52467 | 18.82075 | 0.48 | 47.4 | Neutral |
| testneg_5436 | 0 | -0.32 | 3.779532 | -3 | -0.470437 | -3 | 0.11373 | 0.10739 | 0.25904 | 0.16134 | 20.27778 | 0.183928571 | 0 | Neutral |
| testneg_5437 | -0.000303 | 0.00015152 | -0.119638 | -3 | 5.438117 | -3 | 0.16379 | 0.31462 | 0.12463 | 0.12788 | 4.3985 | 0.187689017 | 0 | Neutral |
| testneg_5438 | 0 | 0 | -0.090263 | -3 | 1.430219 | -3 | 0.80629 | 0.12788 | 0.21731 | 0.11795 | 9.33333 | 0.000816667 | 0 | Neutral |
| testneg_5439 | 0 | 0 | -0.124136 | -3 | 0.923434 | -3 | 0.16248 | 0.14352 | 0.16436 | 0.19109 | 18.30413 | 0.050911616 | 0 | Neutral |
| testneg_5440 | 0 | -0.01 | 6.857724 | -3 | -0.005662 | -3 | 0.18224 | 0.10419 | 0.1664 | 0.10113 | 20.83333 | 0.210106383 | 1.2 | Neutral |
| testneg_5441 | 0.06 | -0.57 | 2.146864 | -3 | 3.809317 | -3 | 0.19898 | 0.09316 | 0.2442 | 0.11401 | 17.21847 | 0.050297059 | 0 | Neutral |
| testneg_5442 | -0.06 | 0.57 | 0.172291 | -3 | -0.345945 | -3 | 0.0138 | 0.06769 | 0.08476 | 0.12788 | 17.77778 | 0.06 | 0 | Neutral |
| testneg_5443 | 0 | 0 | 1.836976 | -3 | -2.563034 | -3 | 0.84491 | 0.30945 | 0.11043 | 0.12788 | 10.83333 | 0.018108974 | 0 | Neutral |
| testneg_5444 | 0.01 | -1.44 | 1.237874 | -3 | 2.055296 | -3 | 0.10893 | 0.12072 | 0.15092 | 0.09074 | 21.43745 | 0.03 | 0 | Neutral |
| testneg_5445 | 0.04 | -0.23 | 3.599958 | -3 | 0.751081 | -3 | 0.40491 | 0.17596 | 0.56824 | 0.10646 | 10.83333 | 0.06 | 0 | Neutral |
| testneg_5446 | 0 | 0 | 1.012321 | -3 | 2.631274 | -3 | 0.21805 | 0.12777 | 0.14458 | 0.14062 | 13.08587 | 0.075698027 | 0 | Neutral |
| testneg_5447 | 0.01 | -1.44 | 3.677821 | -3 | 3.574921 | -3 | 0.61211 | 0.11059 | 0.11501 | 0.13748 | 20.91877 | 0.353875458 | 8.2 | Neutral |
| testneg_5448 | 0 | 0 | 1.519976 | -3 | 2.12199 | -3 | 0.19898 | 0.12788 | 0.2663 | 0.32806 | 11.93907 | 0.05 | 0 | Neutral |
| testneg_5449 | -0.02 | 0.63 | 2.266693 | 2.266693 | -1.352736 | -3 | 0.08401 | 0.16123 | 0.27976 | 0.12613 | 10.22898 | 0.140306383 | 0.2 | Neutral |
| testneg_5450 | 0.14 | -1.01 | -0.221228 | -3 | 1.5447 | -3 | 0.11153 | 0.10767 | 0.08932 | 0.32882 | 10.83333 | 0.1018794 | 0 | Neutral |
| testneg_5451 | -0.08 | -0.03 | 3.726949 | -3 | 5.654876 | -3 | 0.13863 | 0.2579 | 0.82455 | 0.2093 | 12.70833 | 0.766437039 | 99.8 | Disease_causing |
| testneg_5452 | 0.05 | 0.29 | 1.745411 | -3 | 4.668758 | -3 | 0.16835 | 0.09282 | 0.06112 | 0.06901 | 20.83333 | 0.08 | 0 | Neutral |
| testneg_5453 | -0.19 | 0.25 | 1.424214 | 1.424214 | 0.618993 | 0.618993 | 0.21153 | 0.11357 | 0.04207 | 0.12788 | 22.61299 | 0.08 | 0 | Neutral |
| testneg_5454 | -0.08 | -0.03 | 6.096653 | -3 | -0.106366 | -3 | 0.07961 | 0.11474 | 0.19898 | 0.12788 | 30.42735 | 0.15 | 0 | Neutral |
| testneg_5455 | -0.01 | -0.79 | 0.119573 | -3 | 3.326043 | -3 | 0.29451 | 0.13582 | 0.2016 | 0.13956 | 18.90684 | 0.201707287 | 0.2 | Neutral |
| testneg_5456 | 0 | 0 | 0.964615 | -3 | 2.647457 | -3 | 0.64914 | 0.12481 | 0.19898 | 0.12788 | 17.73678 | 0.014102564 | 0 | Neutral |
| testneg_5457 | -0.02 | 0.36 | 3.437388 | -3 | 0.626243 | 0.626243 | 0.22645 | 0.1704 | 0.34412 | 0.08672 | 17.18626 | 0.06 | 0 | Neutral |
| testneg_5458 | 0 | 0 | 0.03503 | -3 | 4.027424 | -3 | 0.90955 | 0.10096 | 0.16926 | 0.11871 | 17.77778 | 0.052223756 | 0 | Neutral |
| testneg_5459 | 0.04 | -0.23 | 3.416183 | -3 | 1.426932 | 1.426932 | 0.44086 | 0.173 | 0.1462 | 0.18896 | 10.41667 | 0.260001149 | 1 | Neutral |
| testneg_5460 | -0.07 | -0.38 | -1.097042 | -3 | 2.454179 | -3 | 0.10703 | 0.11611 | 0.11788 | 0.12788 | 33.60688 | 0.03 | 0 | Neutral |
| testneg_5461 | 0 | 0 | 1.371909 | -3 | 7.284716 | -3 | 0.47733 | 0.08388 | 0.227 | 0.14389 | 20.20318 | 0.322019336 | 5.4 | Neutral |
| testneg_5462 | 0 | 0 | 0.382894 | -3 | -2.410599 | -3 | 0.84236 | 0.14093 | 0.59691 | 0.11049 | 15.59524 | 0.01 | 0 | Neutral |
| testneg_5463 | 0 | 0 | 0.145858 | -3 | 4.048415 | -3 | 0.16797 | 0.12788 | 0.19898 | 0.12788 | 24.68926 | 0.041367777 | 0 | Neutral |
| testneg_5464 | -0.21 | -0.15 | -0.83994 | -3 | 5.471755 | -3 | 0.95319 | 0.20297 | 0.1784 | 0.12788 | 6.96262 | 0.266018207 | 0.8 | Neutral |
| testneg_5465 | 0 | 0 | -0.387684 | -3 | 3.405014 | -3 | 0.25179 | 0.09355 | 0.32423 | 0.08104 | 22.92612 | 0.040533428 | 0 | Neutral |
| testneg_5466 | 0 | 0 | 0.488023 | -3 | 1.56158 | -3 | 0.85508 | 0.12634 | 0.0819 | 0.1823 | 13.55321 | 0.051199218 | 0 | Neutral |
| testneg_5467 | 0.18 | -1.05 | 7.334773 | -3 | 1.729145 | -3 | 0.86835 | 0.1513 | 0.13575 | 0.12788 | 12.59789 | 0.43 | 28.6 | Neutral |
| testneg_5468 | 0 | 0 | 0.850542 | 0.850542 | 1.680224 | -3 | 0.13029 | 0.12788 | 0.05444 | 0.12788 | 16.88659 | 0.03 | 0 | Neutral |
| testneg_5469 | 0 | 0 | 0.28875 | -3 | 0.17553 | -3 | 0.40179 | 0.12112 | 0.10646 | 0.1069 | 12.59057 | 0.000221163 | 0 | Neutral |
| testneg_5470 | 0.02 | 0.45 | 5.042647 | -3 | 4.54141 | -3 | 0.45927 | 0.74955 | 0.20431 | 0.17304 | 10.22898 | 0.839851838 | 99.6 | Disease_causing |
| testneg_5471 | 0.07 | 0.38 | 1.720938 | -3 | 2.103795 | -3 | 0.28856 | 0.1544 | 0.08723 | 0.09503 | 59.22687 | 0.09 | 0 | Neutral |
| testneg_5472 | 0 | 0 | 0.61227 | -3 | 4.163711 | -3 | 0.19898 | 0.12788 | 0.21122 | 0.12788 | 18.1936 | 0.031921586 | 0 | Neutral |
| testneg_5473 | 0.14 | -1.01 | 1.424096 | -3 | 0.34432 | -3 | 0.11358 | 0.12788 | 0.52957 | 0.18687 | 16.33387 | 0.06 | 0 | Neutral |
| testneg_5474 | -0.06 | -0.18 | 1.398739 | -3 | 3.894086 | -3 | 0.54113 | 0.11017 | 0.9637 | 0.12788 | 12.52798 | 0.090714286 | 0 | Neutral |
| testneg_5475 | 0 | -0.01 | 0.63597 | -3 | -0.024667 | -3 | 0.88914 | 0.39065 | 0.59599 | 0.12477 | 10 | 0.01 | 0 | Neutral |
| testneg_5476 | 0.03 | 0.12 | 1.382401 | -3 | -1.289628 | -3 | 0.82131 | 0.12788 | 0.20116 | 0.10008 | 21.16857 | 0.02 | 0 | Neutral |
| testneg_5477 | 0 | -0.01 | -1.288398 | -3 | 2.366366 | -3 | 0.19898 | 0.12788 | 0.12051 | 0.08503 | 13.51351 | 0.02 | 0 | Neutral |
| testneg_5478 | 0.03 | -1.03 | -0.13935 | -3 | 0.829502 | -3 | 0.55367 | 0.57993 | 0.08686 | 0.0953 | 9.98688 | 0.0615 | 0 | Neutral |
| testneg_5479 | -0.03 | 0.85 | 1.432399 | -3 | 3.089818 | -3 | 0.93284 | 0.71489 | 0.09026 | 0.09447 | 6.94444 | 0.120682124 | 0 | Neutral |
| testneg_5480 | 0 | 0 | 0.138788 | -3 | -1.086999 | -3 | 0.71463 | 0.29732 | 0.13126 | 0.11628 | 17.74744 | 0.010318574 | 0 | Neutral |
| testneg_5481 | 0 | 0 | -0.113262 | -3 | 0.406982 | -3 | 0.56154 | 0.0939 | 0.13255 | 0.0962 | 11.7299 | 0.02 | 0 | Neutral |
| testneg_5482 | 0.08 | 0.03 | -1.514831 | -3 | 0.967538 | -3 | 0.19898 | 0.12788 | 0.21235 | 0.1449 | 20.78544 | 0.063333333 | 0 | Neutral |
| testneg_5483 | -0.08 | -0.03 | 2.574972 | -0.330275 | 2.51125 | -3 | 0.10462 | 0.1278 | 0.31344 | 0.12788 | 10.74877 | 0.05 | 0 | Neutral |
| testneg_5484 | 0.16 | -1.37 | 2.689971 | -3 | -0.275564 | -3 | 0.85075 | 0.16646 | 0.14432 | 0.22579 | 11.96632 | 0.19080397 | 0 | Neutral |
| testneg_5485 | 0 | 0 | -0.038389 | -3 | 5.405576 | -3 | 0.16326 | 0.09256 | 0.52765 | 0.20417 | 17.60518 | 0.237303846 | 0.2 | Neutral |
| testneg_5486 | 0.21 | 0.15 | 1.912237 | -3 | 1.35891 | -3 | 0.12511 | 0.11897 | 0.08531 | 0.12788 | 21.315 | 0.03 | 0 | Neutral |
| testneg_5487 | -0.01 | -0.8 | 5.824028 | -3 | 11.548039 | -3 | 0.76957 | 0.13588 | 0.19898 | 0.0989 | 14.36878 | 0.707777778 | 99.6 | Disease_causing |
| testneg_5488 | 0.08 | 0.03 | -1.290562 | -3 | 2.815622 | -1.287967 | 0.19898 | 0.12895 | 0.46037 | 0.1343 | 25.46373 | 0.177219478 | 0 | Neutral |
| testneg_5489 | -0.07 | -0.38 | 1.400536 | -3 | 2.46612 | 1.583224 | 0.89972 | 0.10766 | 0.10314 | 0.08894 | 18.38327 | 0.05 | 0 | Neutral |

| | | | | | | | | | | | | | | |
|---|---|---|---|---|---|---|---|---|---|---|---|---|---|---|
| testneg_5490 | -0.0025714 | -0.1897143 | 2.053049 | -3 | 4.717335 | -3 | 0.26586 | 0.10825 | 0.46242 | 0.39055 | 17.77778 | 0.361548647 | 10.4 | Neutral |
| testneg_5491 | -0.08 | -0.03 | 1.50021 | -3 | -1.122085 | -1.122085 | 0.27618 | 0.11809 | 0.0774 | 0.1702 | 38.88889 | 0.109325092 | 0 | Neutral |
| testneg_5492 | 0 | 0 | 0.164848 | -3 | 2.385228 | -3 | 0.12839 | 0.09503 | 0.09093 | 0.28643 | 14.22149 | 0.090339343 | 0 | Neutral |
| testneg_5493 | 0 | 0 | 0.479788 | -3 | 0.680291 | -3 | 0.21587 | 0.11701 | 0.23574 | 0.12788 | 13.51351 | 0 | 0 | Neutral |
| testneg_5494 | 0.05 | 0.29 | -0.076665 | -3 | 5.730226 | -3 | 0.19898 | 0.12788 | 0.13704 | 0.10054 | 34.77719 | 0.1 | 0 | Neutral |
| testneg_5495 | 0 | 0 | 1.012321 | 1.012321 | 2.317295 | -3 | 0.21805 | 0.12777 | 0.99178 | 0.93124 | 16.16034 | 0.17 | 0 | Neutral |
| testneg_5496 | -0.08 | -0.03 | 7.833352 | -3 | 1.538056 | -3 | 0.08067 | 0.34914 | 0.53172 | 0.12788 | 42.23911 | 0.52 | 64.4 | Disease_causing |
| testneg_5497 | 0 | 0 | -0.489156 | -3 | 0.60982 | 0.60982 | 0.05887 | 0.10046 | 0.23844 | 0.27772 | 24.32452 | 0.11 | 0 | Neutral |
| testneg_5498 | 0 | 0 | 1.700589 | -3 | -1.162831 | -1.162831 | 0.89357 | 0.22348 | 0.13398 | 0.12788 | 17.21847 | 0.010763376 | 0 | Neutral |
| testneg_5499 | 0 | -0.32 | 0.625959 | -3 | -0.908797 | -3 | 0.27409 | 0.10785 | 0.12282 | 0.20617 | 22.36648 | 0.080457391 | 0 | Neutral |
| testneg_5500 | 0.07 | -0.31 | -0.344853 | -3 | -2.397932 | -2.397932 | 0.50276 | 0.09638 | 0.48705 | 0.09993 | 20.89902 | 0.020098124 | 0 | Neutral |
| testneg_5501 | 0.03 | 1.22 | 0.949523 | 0.949523 | 0.827033 | -3 | 0.10056 | 0.12608 | 0.38934 | 0.15403 | 12.24684 | 0.14 | 0 | Neutral |
| testneg_5502 | 0.16 | -1.37 | 6.9826 | -3 | -0.114413 | -3 | 0.11905 | 0.23747 | 0.19898 | 0.12788 | 16.8018 | 0.318097222 | 4.4 | Neutral |
| testneg_5503 | 0 | 0 | 1.568756 | -3 | -1.333378 | -3 | 0.9516 | 0.12788 | 0.0811 | 0.09449 | 19.66924 | 0.02 | 0 | Neutral |
| testneg_5504 | 0.14 | -0.69 | 1.524427 | -3 | 4.866518 | -3 | 0.99317 | 0.60884 | 0.12949 | 0.44208 | 10 | 0.51833569 | 64.4 | Disease_causing |
| testneg_5505 | -0.06 | -0.18 | -1.108542 | -1.108542 | 6.905837 | -3 | 0.90308 | 0.28373 | 0.27004 | 0.17052 | 12.70833 | 0.51810837 | 61.2 | Disease_causing |
| testneg_5506 | -0.03 | 1.03 | -1.53473 | -3 | -0.453465 | -3 | 0.03631 | 0.80564 | 0.46399 | 0.15334 | 5 | 0.289231505 | 0.2 | Neutral |
| testneg_5507 | -0.05 | -0.29 | 0.660269 | -3 | -0.124016 | -3 | 0.19898 | 0.12788 | 0.42836 | 0.12788 | 16.8018 | 0.000540936 | 0 | Neutral |
| testneg_5508 | 0.04 | -0.23 | 4.469433 | -3 | 0.897808 | 0.897808 | 0.23468 | 0.10874 | 0.17155 | 0.18585 | 19.09347 | 0.197948413 | 0 | Neutral |
| testneg_5509 | 0 | 0 | 1.981098 | -3 | 1.397791 | -3 | 0.22162 | 0.11997 | 0.70872 | 0.09884 | 14.18151 | 0.01 | 0 | Neutral |
| testneg_5510 | 0.14 | -1.01 | 1.4604 | -3 | -0.196592 | -0.196592 | 0.53888 | 0.09904 | 0.32733 | 0.16869 | 18.76882 | 0.09 | 0 | Neutral |
| testneg_5511 | 0 | 0 | 0.276668 | -3 | 6.423333 | -3 | 0.35376 | 0.09478 | 0.16962 | 0.12788 | 11.875 | 0.114286131 | 0 | Neutral |
| testneg_5512 | 0 | 0 | -0.151164 | -0.151164 | 0.60982 | 0.60982 | 0.65708 | 0.0936 | 0.23844 | 0.27772 | 14.42934 | 0.067132035 | 0 | Neutral |
| testneg_5513 | 0.14 | -0.37 | 6.441278 | -3 | 1.539579 | -3 | 0.20772 | 0.16918 | 0.18459 | 0.33541 | 21.38889 | 0.612939174 | 90.2 | Disease_causing |
| testneg_5514 | 0 | 0 | 0.845771 | -3 | 1.902977 | 1.902977 | 0.27197 | 0.12346 | 0.18504 | 0.0822 | 11.25 | 0.03 | 0 | Neutral |
| testneg_5515 | 0 | 0 | -0.481968 | -0.481968 | 0.416167 | -3 | 0.67331 | 0.10118 | 0.1224 | 0.11899 | 31.53846 | 0.020219459 | 0 | Neutral |
| testneg_5516 | 0.07 | 0.38 | 1.022289 | 1.022289 | 0.068014 | 0.068014 | 0.18281 | 0.12788 | 0.20869 | 0.13493 | 31.12379 | 0.120693665 | 0 | Neutral |
| testneg_5517 | 0 | 0 | 1.055391 | -3 | 2.241925 | -3 | 0.15689 | 0.28307 | 0.31025 | 0.09491 | 20.91877 | 0.03 | 0 | Neutral |
| testneg_5518 | -0.07 | -0.38 | 2.393948 | 2.393948 | -0.480309 | -3 | 0.70819 | 0.5975 | 0.13693 | 0.12788 | 16.60044 | 0.15 | 0.2 | Neutral |
| testneg_5519 | 0 | 0 | 0.6379 | -3 | 0.8528 | 0.8528 | 0.40557 | 0.11865 | 0.55384 | 0.15273 | 28.47222 | 0.051181099 | 0 | Neutral |
| testneg_5520 | -0.19 | 0.25 | -1.729462 | -3 | 2.23696 | -3 | 0.16155 | 0.20638 | 0.68865 | 0.19414 | 18.33333 | 0.265333333 | 0.2 | Neutral |
| testneg_5521 | 0.14 | -1.01 | 3.072449 | -3 | 2.025744 | -3 | 0.17529 | 0.09662 | 0.1215 | 0.12788 | 35.72379 | 0.05 | 0 | Neutral |
| testneg_5522 | 0.03 | 1.22 | 0.635023 | -3 | 1.023241 | -3 | 0.12136 | 0.27707 | 0.1617 | 0.17928 | 27.82626 | 0.216966126 | 0 | Neutral |
| testneg_5523 | -0.07 | -0.38 | 4.78355 | -3 | 2.02554 | -3 | 0.25516 | 0.12788 | 0.21331 | 0.09513 | 10.2717 | 0.078697044 | 0 | Neutral |
| testneg_5524 | 0.04 | -1.36 | 5.181357 | -3 | 1.38401 | -3 | 0.11956 | 0.3237 | 0.37212 | 0.11323 | 22.88801 | 0.234345238 | 0 | Neutral |
| testneg_5525 | 0 | 0 | 0.329118 | -3 | 2.507265 | -3 | 0.51707 | 0.45685 | 0.98581 | 0.27035 | 21.94444 | 0.23 | 0.2 | Neutral |
| testneg_5526 | 0 | 0 | 1.677581 | -3 | 0.345315 | -3 | 0.74919 | 0.85508 | 0.09964 | 0.10384 | 17.21847 | 0.06 | 0 | Neutral |
| testneg_5527 | 0.17 | 0.38 | 4.608358 | -3 | 3.769116 | -3 | 0.04912 | 0.08574 | 0.54891 | 0.55746 | 18.33333 | 0.497149549 | 52.2 | Disease_causing |
| testneg_5528 | 0.03 | 1.22 | 2.244475 | -3 | 2.02727 | -3 | 0.34406 | 0.09894 | 0.82269 | 0.15732 | 29.12774 | 0.101313511 | 0 | Neutral |
| testneg_5529 | 0 | 0 | -0.101683 | -3 | 5.543521 | -3 | 0.72657 | 0.11262 | 0.32181 | 0.07041 | 13.56281 | 0.06 | 0 | Neutral |
| testneg_5530 | 0.07 | 0.01 | 3.419422 | -3 | 3.258412 | -3 | 0.05323 | 0.32175 | 0.85493 | 0.30152 | 10.83333 | 0.619833333 | 92.6 | Disease_causing |
| testneg_5531 | -0.03 | -0.1 | 5.021764 | -3 | -0.270908 | -3 | 0.30684 | 0.12788 | 0.14101 | 0.14318 | 21.07466 | 0.19 | 0 | Neutral |
| testneg_5532 | 0.07 | 0.7 | 3.275701 | -3 | 7.269131 | 4.234809 | 0.19102 | 0.12788 | 0.37109 | 0.12788 | 13.51351 | 0.58 | 81.8 | Disease_causing |
| testneg_5533 | 0.21 | 0.15 | 1.616555 | -3 | 4.24996 | -3 | 0.20072 | 0.08606 | 0.85493 | 0.30152 | 18.33333 | 0.261580947 | 0.8 | Neutral |
| testneg_5534 | 0.01 | 0.8 | 6.972724 | -3 | -0.298555 | -0.298555 | 0.44953 | 0.14403 | 0.10314 | 0.12173 | 18.33333 | 0.326071429 | 7.4 | Neutral |
| testneg_5535 | -0.03 | -0.1 | 2.738949 | -3 | 1.2481 | -3 | 0.07961 | 0.11474 | 0.11281 | 0.0958 | 28.16398 | 0.023541667 | 0 | Neutral |
| testneg_5536 | 0.15 | 0.06 | 2.604577 | -3 | 2.914276 | -3 | 0.68918 | 0.16819 | 0.2698 | 0.18815 | 10.82179 | 0.377834207 | 13.8 | Neutral |
| testneg_5537 | 0.02 | 0.45 | 1.21068 | -3 | -1.549377 | -3 | 0.95247 | 0.11285 | 0.16404 | 0.4047 | 18.22258 | 0.09 | 0 | Neutral |
| testneg_5538 | 0.07 | 0.7 | 4.037659 | -3 | 0.259866 | -3 | 0.77944 | 0.1858 | 0.04063 | 0.05848 | 27.26496 | 0.081041667 | 0 | Neutral |
| testneg_5539 | 0 | 0 | 0.104266 | -3 | 1.821777 | -3 | 0.96396 | 0.7628 | 0.11187 | 0.12788 | 10.41667 | 0.04 | 0 | Neutral |
| testneg_5540 | 0 | 0 | 1.609501 | -3 | -0.448326 | -3 | 0.16404 | 0.09378 | 0.03466 | 0.19518 | 22.27338 | 0.072268018 | 0 | Neutral |
| testneg_5541 | 0.14 | -1.01 | 2.521088 | -3 | 0.930465 | -3 | 0.10021 | 0.1105 | 0.30429 | 0.29938 | 12.02381 | 0.103006133 | 0 | Neutral |
| testneg_5542 | 0.04 | -0.23 | 3.352237 | -3 | 3.375468 | -3 | 0.17672 | 0.14229 | 0.21807 | 0.12788 | 20 | 0.14763438 | 0 | Neutral |
| testneg_5543 | 0.02 | -0.28 | -1.20584 | -3 | 0.054796 | -3 | 0.53216 | 0.57092 | 0.32531 | 0.12351 | 10.22898 | 0.05 | 0 | Neutral |
| testneg_5544 | -0.09 | 0.3 | 0.156321 | -3 | 5.537244 | 5.537244 | 0.21566 | 0.15056 | 0.19898 | 0.09972 | 10.87719 | 0.52 | 65.6 | Disease_causing |
| testneg_5545 | 0.04 | -0.23 | 4.366648 | -3 | 3.635519 | -3 | 0.34279 | 0.09378 | 0.07995 | 0.12788 | 13.72549 | 0.150849005 | 0 | Neutral |
| testneg_5546 | -0.03 | -0.1 | 2.260957 | 2.260957 | 4.940548 | 4.940548 | 0.36054 | 0.12788 | 0.14628 | 0.09488 | 29.05798 | 0.44 | 29.2 | Neutral |
| testneg_5547 | 0 | -0.73 | 6.205907 | -3 | -0.124213 | -0.124213 | 0.07406 | 0.08059 | 0.74641 | 0.66512 | 20.76356 | 0.32 | 6.4 | Neutral |
| testneg_5548 | 0 | -0.01 | 0.183879 | -3 | 1.140918 | -3 | 0.19898 | 0.12788 | 0.1846 | 0.08682 | 36.54799 | 0.000425725 | 0 | Neutral |
| testneg_5549 | 0 | 0 | 2.597899 | -3 | 3.712827 | -3 | 0.07175 | 0.29293 | 0.87202 | 0.42207 | 10.41667 | 0.483588187 | 47.2 | Neutral |
| testneg_5550 | 0.21 | 0.15 | 2.317436 | -3 | 3.164016 | -3 | 0.35325 | 0.12614 | 0.10781 | 0.09901 | 27.4024 | 0.104222222 | 0 | Neutral |
| testneg_5551 | 0 | 0 | 6.373062 | -3 | 1.018818 | 1.018818 | 0.36793 | 0.15588 | 0.13398 | 0.12788 | 17.21847 | 0.28 | 3.8 | Neutral |
| testneg_5552 | 0.22 | -2.66 | 7.833649 | -3 | 1.460399 | -3 | 0.20951 | 0.16278 | 0.80314 | 0.42344 | 12.77241 | 0.507900121 | 55.6 | Disease_causing |
| testneg_5553 | 0 | 0 | 0.383044 | -3 | 0.38184 | -3 | 0.63819 | 0.32258 | 0.35421 | 0.10694 | 14.25481 | 0.01 | 0 | Neutral |
| testneg_5554 | 0 | 0 | -0.030607 | -3 | -0.598747 | -3 | 0.17146 | 0.19882 | 0.694 | 0.09974 | 14.5275 | 0.020128395 | 0 | Neutral |
| testneg_5555 | 0.05 | -0.81 | 0.993178 | -3 | 0.276186 | -3 | 0.07098 | 0.39918 | 0.52914 | 0.25278 | 11.23846 | 0.21 | 0 | Neutral |
| testneg_5556 | 0 | 0 | 1.445184 | 1.445184 | 0.429523 | -3 | 0.33469 | 0.17432 | 0.18328 | 0.12788 | 10.41667 | 0.03 | 0 | Neutral |
| testneg_5557 | -0.18 | -0.16 | 2.409244 | -3 | 0.450722 | -3 | 0.99535 | 0.38424 | 0.16078 | 0.39498 | 5.69444 | 0.213374586 | 0 | Neutral |
| testneg_5558 | -0.04 | 1.36 | 2.289419 | -3 | 2.61669 | -3 | 0.53654 | 0.22867 | 0.2578 | 0.09796 | 18.33333 | 0.164429825 | 0 | Neutral |
| testneg_5559 | -0.09 | 0.25 | 4.334907 | -3 | 2.235084 | -3 | 0.15263 | 0.12788 | 0.03942 | 0.12788 | 24.2785 | 0.08 | 0 | Neutral |
| testneg_5560 | -0.08 | -0.03 | 3.13774 | -3 | -3.629472 | -3 | 0.26758 | 0.10473 | 0.15612 | 0.11465 | 13.46801 | 0.02047619 | 0 | Neutral |
| testneg_5561 | 0.12 | -1.14 | -0.006272 | -0.006272 | 5.095304 | -3 | 0.07335 | 0.12788 | 0.18559 | 0.13755 | 16.8018 | 0.311690476 | 2.2 | Neutral |
| testneg_5562 | 0.06 | -0.57 | 5.777528 | 5.777528 | 1.255613 | -3 | 0.17672 | 0.14229 | 0.06971 | 0.09918 | 14.19394 | 0.59 | 93.4 | Disease_causing |
| testneg_5563 | -0.14 | 0.69 | 0.233598 | 0.233598 | 3.787163 | -3 | 0.25804 | 0.11876 | 0.07675 | 0.11464 | 22.02492 | 0.15 | 0 | Neutral |
| testneg_5564 | -0.15 | 0.28 | -0.668638 | -0.668638 | 3.107095 | -3 | 0.37131 | 0.10994 | 0.19898 | 0.40608 | 11.96702 | 0.250665002 | 0.2 | Neutral |
| testneg_5565 | 0 | 0 | 2.30653 | -3 | 0.760327 | -3 | 0.49106 | 0.89143 | 0.42512 | 0.29333 | 11.55405 | 0.188795348 | 0 | Neutral |
| testneg_5566 | 0 | 0 | 0.147507 | -3 | -0.929745 | -3 | 0.0959 | 0.12788 | 0.09284 | 0.26053 | 34.33451 | 0.090386386 | 0 | Neutral |
| testneg_5567 | 0.18 | 0.16 | -1.517745 | -3 | 1.432015 | -3 | 0.56605 | 0.15893 | 0.64108 | 0.51829 | 19.07944 | 0.170763636 | 0 | Neutral |
| testneg_5568 | -0.02 | -0.45 | 0.999199 | -0.001031 | 0.752432 | -3 | 0.32022 | 0.09529 | 0.19898 | 0.1347 | 12.00474 | 0.06 | 0 | Neutral |
| testneg_5569 | 0 | 0 | 0.046064 | -3 | 4.894593 | -3 | 0.2543 | 0.15588 | 0.13743 | 0.12788 | 17.77778 | 0.11 | 0 | Neutral |
| testneg_5570 | 0 | 0 | 0.205888 | -3 | 1.251399 | -3 | 0.08922 | 0.10539 | 0.20329 | 0.10718 | 12.29167 | 0 | 0 | Neutral |
| testneg_5571 | 0.09 | -0.3 | 4.185197 | -3 | 4.602859 | -3 | 0.10472 | 0.10014 | 0.5065 | 0.20697 | 4.72222 | 0.644517424 | 92 | Disease_causing |
| testneg_5572 | -0.14 | 0.37 | 5.047397 | 0.726355 | 0.19932 | -3 | 0.09439 | 0.1884 | 0.19898 | 0.09927 | 22.62502 | 0.22 | 0 | Neutral |
| testneg_5573 | 0 | 0 | 1.605285 | -3 | 3.588713 | -3 | 0.5068 | 0.12276 | 0.32606 | 0.10251 | 12.29167 | 0.040103137 | 0 | Neutral |
| testneg_5574 | 0.03 | -1.03 | 3.623517 | -3 | 3.810693 | -3 | 0.54724 | 0.12788 | 0.41643 | 0.09692 | 11.875 | 0.17 | 0 | Neutral |
| testneg_5575 | 0.02 | 0.13 | 7.041902 | -3 | 0.080104 | -3 | 0.34467 | 0.13076 | 0.73947 | 0.2058 | 56.18299 | 0.382892308 | 15 | Neutral |
| testneg_5576 | 0 | 0 | 5.435713 | -3 | 0.885869 | 0.885869 | 0.24026 | 0.12788 | 0.12003 | 0.12788 | 37.26752 | 0.139375 | 0 | Neutral |
| testneg_5577 | -0.01 | -0.8 | 1.765078 | -3 | 3.408834 | -3 | 0.36461 | 0.11015 | 0.19479 | 0.12788 | 18.78762 | 0.040842363 | 0 | Neutral |
| testneg_5578 | -0.02 | 0.63 | -0.833334 | -3 | 0.63597 | -3 | 0.57454 | 0.14465 | 0.88914 | 0.39065 | 13.29613 | 0.081146601 | 0 | Neutral |
| testneg_5579 | 0 | 0 | 1.638668 | -3 | -1.932308 | -3 | 0.14199 | 0.1327 | 0.06474 | 0.23607 | 10.41667 | 0.052581818 | 0 | Neutral |
| testneg_5580 | -0.03 | -0.1 | 4.149716 | -3 | -2.137796 | -3 | 0.19322 | 0.12788 | 0.13698 | 0.12788 | 33.73329 | 0.04 | 0 | Neutral |
| testneg_5581 | 0.14 | -0.37 | 4.610923 | -3 | -0.26416 | -3 | 0.98833 | 0.48581 | 0.45343 | 0.12014 | 11.25 | 0.140572581 | 0 | Neutral |
| testneg_5582 | -0.15 | -0.04 | 1.473246 | -3 | -0.187363 | -3 | 0.29013 | 0.11368 | 0.90649 | 0.3543 | 20.40047 | 0.091324319 | 0 | Neutral |
| testneg_5583 | 0.14 | -1.01 | 0.296748 | -3 | 1.019702 | -3 | 0.75906 | 0.09702 | 0.98238 | 0.10873 | 8.36931 | 0.03 | 0 | Neutral |
| testneg_5584 | 0 | 0 | 1.43999 | -3 | 6.433128 | -3 | 0.10723 | 0.11163 | 0.25516 | 0.12788 | 13.51351 | 0.13 | 0 | Neutral |
| testneg_5585 | 0.07 | 0.7 | 4.605651 | -3 | 1.105669 | -3 | 0.10719 | 0.08618 | 0.44385 | 0.1311 | 10.22898 | 0.1 | 0 | Neutral |
| testneg_5586 | 0 | 0 | 0.497873 | -3 | 1.70493 | -3 | 0.33049 | 0.12788 | 0.72068 | 0.99841 | 11.91142 | 0.050380747 | 0 | Neutral |
| testneg_5587 | 0.07 | 0.7 | 5.597504 | -3 | 2.565617 | -3 | 0.10187 | 0.12339 | 0.24011 | 0.28559 | 19.44444 | 0.506614148 | 56.2 | Disease_causing |
| testneg_5588 | 0.09 | -0.3 | 1.994203 | -3 | -0.209155 | -3 | 0.13565 | 0.46572 | 0.71844 | 0.20204 | 10.41667 | 0.183614609 | 0 | Neutral |
| testneg_5589 | 0 | 0 | 3.296898 | -3 | 2.195109 | -3 | 0.13559 | 0.1946 | 0.43502 | 0.14712 | 18.76882 | 0.26776902 | 0.8 | Neutral |
| testneg_5590 | 0 | 0 | -0.264755 | -3 | 3.878129 | -3 | 0.09024 | 0.08559 | 0.06744 | 0.12788 | 10.41667 | 0.065 | 0 | Neutral |
| testneg_5591 | -0.0110909 | 0.08709091 | 2.966005 | -3 | 2.019586 | -3 | 0.26192 | 0.15452 | 0.44488 | 0.31694 | 18.88889 | 0.336714286 | 4.6 | Neutral |
| testneg_5592 | 0.21 | 0.15 | 3.541975 | -3 | 0.05475 | -3 | 0.11608 | 0.09123 | 0.13468 | 0.09235 | 40.33827 | 0.05 | 0 | Neutral |
| testneg_5593 | -0.02 | -0.41 | 3.378414 | -3 | -2.990613 | -3 | 0.39724 | 0.11058 | 0.19403 | 0.09278 | 12.78306 | 0.03 | 0 | Neutral |
| testneg_5594 | -0.09 | 0.3 | -0.178395 | -3 | -0.354552 | -3 | 0.21033 | 0.35953 | 0.09815 | 0.12788 | 18.07512 | 0.06 | 0 | Neutral |
| testneg_5595 | -0.08 | -0.03 | 0.307773 | -3 | 0.661515 | -3 | 0.14956 | 0.10585 | 0.07534 | 0.09292 | 14.64539 | 0.01 | 0 | Neutral |
| testneg_5596 | -0.05 | -0.29 | 5.645383 | -3 | 0.484944 | -3 | 0.08683 | 0.26961 | 0.08896 | 0.12788 | 26.42807 | 0.28 | 1.8 | Neutral |
| testneg_5597 | 0.01 | 0.8 | 3.096454 | -3 | 0.232171 | -0.618794 | 0.31824 | 0.6682 | 0.38934 | 0.15403 | 5.97222 | 0.271807692 | 1.8 | Neutral |
| testneg_5598 | -0.08 | -0.03 | 3.415166 | -3 | 0.973671 | -3 | 0.67796 | 0.10874 | 0.47246 | 0.47221 | 12.29167 | 0.149835539 | 0 | Neutral |
| testneg_5599 | 0.03 | 0.12 | 1.943409 | -3 | 3.47892 | -3 | 0.41647 | 0.11239 | 0.18423 | 0.10808 | 27.92726 | 0.053650946 | 0 | Neutral |
| testneg_5600 | 0.04 | 2.01 | -2.511821 | -3 | 3.848975 | -3 | 0.13873 | 0.12788 | 0.10458 | 0.10499 | 43.79527 | 0.18 | 0 | Neutral |
| testneg_5601 | -0.01 | -0.79 | 2.218156 | -3 | 5.894587 | -3 | 0.37327 | 0.2201 | 0.15364 | 0.08789 | 10.41667 | 0.256083333 | 0.6 | Neutral |
| testneg_5602 | 0.21 | 0.15 | 7.652527 | -3 | -2.165884 | -3 | 0.64701 | 0.94571 | 0.12957 | 0.12788 | 13.85965 | 0.44 | 30.8 | Neutral |
| testneg_5603 | 0.04 | -0.23 | 3.352237 | -3 | 1.034225 | -3 | 0.17672 | 0.14229 | 0.23913 | 0.12788 | 19.09347 | 0.030290476 | 0 | Neutral |
| testneg_5604 | -0.06 | 0.57 | 1.042833 | -3 | 4.338577 | -3 | 0.19898 | 0.12788 | 0.19898 | 0.12788 | 20.1245 | 0.09 | 0 | Neutral |
| testneg_5605 | 0.14 | -1.01 | 0.849231 | -3 | 0.728537 | -3 | 0.08629 | 0.12788 | 0.08521 | 0.11239 | 10.41667 | 0.01 | 0 | Neutral |
| testneg_5606 | 0 | 0 | -0.470651 | -3 | 2.195633 | -3 | 0.28063 | 0.40063 | 0.06723 | 0.10296 | 19.10645 | 0.041813856 | 0 | Neutral |
| testneg_5607 | 0 | 0 | 1.428423 | -3 | 3.302134 | -3 | 0.46091 | 0.13852 | 0.36813 | 0.09414 | 21.94444 | 0.05825 | 0 | Neutral |
| testneg_5608 | 0.21 | 0.15 | 3.044041 | -3 | 2.893051 | -3 | 0.52412 | 0.21177 | 0.15562 | 0.10384 | 18.88889 | 0.226541918 | 0 | Neutral |
| testneg_5609 | 0.02 | 0.45 | 2.472827 | -3 | 2.489811 | -3 | 0.50134 | 0.15488 | 0.10554 | 0.12788 | 64.17937 | 0.14 | 0 | Neutral |
| testneg_5610 | 0.07 | 0.38 | 2.246546 | -3 | 2.060888 | -3 | 0.32022 | 0.09529 | 0.22034 | 0.12788 | 17.20371 | 0.01031754 | 0 | Neutral |
| testneg_5611 | 0.06 | 0.18 | -1.19222 | -1.220105 | 3.061106 | -3 | 0.02971 | 0.33983 | 0.36858 | 0.09681 | 31.51003 | 0.18 | 0 | Neutral |

| | | | | | | | | | | | | | | |
|---|---|---|---|---|---|---|---|---|---|---|---|---|---|---|
| testneg_5612 | -0.03 | -0.1 | 2.260957 | -3 | 0.261126 | -3 | 0.36054 | 0.12788 | 0.65048 | 0.12778 | 21.85713 | 0.010116421 | 0 | Neutral |
| testneg_5613 | -0.01 | -0.8 | 7.038327 | 7.038327 | 0.977938 | -3 | 0.30367 | 0.41964 | 0.47975 | 0.18568 | 10.83333 | 0.893214286 | 100 | Disease_causing |
| testneg_5614 | 0 | 0 | 0.584441 | -3 | 0.532295 | -3 | 0.97678 | 0.09885 | 0.755 | 0.12788 | 7.01467 | 0.01 | 0 | Neutral |
| testneg_5615 | -0.02 | -0.45 | 6.712641 | -3 | 0.211586 | 0.211586 | 0.25731 | 0.12662 | 0.1119 | 0.24875 | 11.61223 | 0.478222826 | 44 | Neutral |
| testneg_5616 | 0 | 0 | 4.603695 | -3 | 3.979487 | -3 | 0.18559 | 0.13755 | 0.13751 | 0.13274 | 15.28953 | 0.450312239 | 34.8 | Neutral |
| testneg_5617 | -0.07 | -0.38 | 5.775288 | -3 | 0.292982 | -3 | 0.16663 | 0.12788 | 0.15677 | 0.18537 | 13.51351 | 0.40683148 | 23.4 | Neutral |
| testneg_5618 | -0.0036364 | 0.11719697 | 0.781229 | 0.781229 | 0.390383 | -3 | 0.07321 | 0.12788 | 0.08213 | 0.08913 | 39.62862 | 0.04 | 0 | Neutral |
| testneg_5619 | -0.07 | -0.38 | 2.28341 | -3 | 0.192071 | -3 | 0.27723 | 0.13229 | 0.18312 | 0.12788 | 24.68926 | 0.02 | 0 | Neutral |
| testneg_5620 | -0.01 | 0.57 | 0.120012 | -3 | 5.958218 | -3 | 0.20491 | 0.22087 | 0.18559 | 0.13755 | 10.41667 | 0.530088346 | 66.8 | Disease_causing |
| testneg_5621 | 0 | 0 | 0.179322 | -3 | 0.108944 | -3 | 0.54349 | 0.60885 | 0.55315 | 0.09195 | 16.8018 | 0.031213526 | 0 | Neutral |
| testneg_5622 | 0.14 | -0.69 | 1.524427 | -3 | 4.336137 | -3 | 0.99317 | 0.60884 | 0.19898 | 0.12788 | 20.18377 | 0.242916667 | 0 | Neutral |
| testneg_5623 | 0 | 0 | 0.56241 | -3 | -0.511954 | -3 | 0.9511 | 0.54626 | 0.16716 | 0.10951 | 25.38314 | 0.05 | 0 | Neutral |
| testneg_5624 | 0.07 | -0.31 | 4.113026 | -3 | 1.656921 | -3 | 0.14021 | 0.43531 | 0.19898 | 0.12788 | 14.12935 | 0.14 | 0 | Neutral |
| testneg_5625 | 0 | 0 | 0.484587 | -3 | -0.440113 | -3 | 0.30056 | 0.12272 | 0.53592 | 0.11293 | 38.27913 | 0.000207259 | 0 | Neutral |
| testneg_5626 | -0.15 | -0.04 | 4.208477 | -3 | -0.450207 | -3 | 0.6201 | 0.12788 | 0.17083 | 0.31043 | 10.41667 | 0.227700443 | 0 | Neutral |
| testneg_5627 | 0 | 0 | 0.497371 | -3 | 3.230388 | -3 | 0.21983 | 0.10146 | 0.1215 | 0.15825 | 17.26246 | 0.170098039 | 0 | Neutral |
| testneg_5628 | 0.16 | -0.37 | 0.701088 | -3 | -1.197202 | -1.197202 | 0.99538 | 0.40044 | 0.08688 | 0.08809 | 21.82143 | 0.1 | 0 | Neutral |
| testneg_5629 | 0.16 | 0.08 | 1.309209 | -3 | 6.773601 | -3 | 0.84588 | 0.14896 | 0.19898 | 0.12788 | 29.37104 | 0.275 | 1.6 | Neutral |
| testneg_5630 | 0 | 0 | -0.040357 | -0.040357 | 2.885372 | -3 | 0.01584 | 0.07827 | 0.21355 | 0.12788 | 10.11902 | 0.05 | 0 | Neutral |
| testneg_5631 | -0.02 | 0.68 | 0.385771 | -3 | 5.902347 | -3 | 0.11077 | 0.10076 | 0.08876 | 0.09037 | 10.83333 | 0.16 | 0 | Neutral |
| testneg_5632 | 0 | 0 | -0.32829 | -3 | -0.27139 | -3 | 0.18509 | 0.12831 | 0.37637 | 0.12788 | 12.01426 | 0.000152111 | 0 | Neutral |
| testneg_5633 | 0 | 0 | -0.109042 | -3 | 0.833236 | -3 | 0.2029 | 0.12788 | 0.4172 | 0.11643 | 16.8018 | 0 | 0 | Neutral |
| testneg_5634 | 0.18 | -1.05 | 5.955324 | -3 | 0.148971 | -3 | 0.24287 | 0.11626 | 0.17007 | 0.12771 | 31.25 | 0.11 | 0 | Neutral |
| testneg_5635 | 0.18 | -1.05 | 2.810817 | 2.810817 | 0.169559 | -3 | 0.09471 | 0.12788 | 0.86166 | 0.14254 | 29.76772 | 0.42575 | 31.6 | Neutral |
| testneg_5636 | 0 | 0 | -0.259811 | -3 | 0.792315 | 0.475838 | 0.70964 | 0.12663 | 0.25516 | 0.12788 | 19.09347 | 0.01 | 0 | Neutral |
| testneg_5637 | 0 | 0 | -0.064795 | -3 | 2.424425 | -3 | 0.4566 | 0.14201 | 0.54211 | 0.12592 | 8.6173 | 0.023489583 | 0 | Neutral |
| testneg_5638 | 0.19 | -0.27 | 3.485229 | -3 | 1.021201 | 1.021201 | 0.22872 | 0.15588 | 0.15785 | 0.35538 | 10.41667 | 0.309746558 | 2.2 | Neutral |
| testneg_5639 | 0.07 | 0.01 | 2.044231 | -3 | 3.536875 | -3 | 0.19898 | 0.10894 | 0.0895 | 0.12788 | 23.6906 | 0.06 | 0 | Neutral |
| testneg_5640 | 0.05 | -0.81 | 4.322868 | -3 | 1.019613 | -3 | 0.47888 | 0.43756 | 0.08789 | 0.09565 | 17.60476 | 0.105833333 | 0 | Neutral |
| testneg_5641 | 0 | 0 | 1.527462 | -3 | 0.877291 | -3 | 0.25528 | 0.24339 | 0.86166 | 0.14254 | 12.31231 | 0.052271906 | 0 | Neutral |
| testneg_5642 | 0.09 | -0.3 | 6.191692 | -3 | -1.039289 | -1.039289 | 0.45927 | 0.74955 | 0.08538 | 0.08065 | 10.41667 | 0.3 | 1 | Neutral |
| testneg_5643 | -0.05 | -0.29 | -0.062714 | -3 | 5.756532 | -3 | 0.05323 | 0.32175 | 0.15992 | 0.12788 | 19.44444 | 0.321388889 | 2 | Neutral |
| testneg_5644 | -0.18 | -0.16 | 2.074931 | -3 | 0.233598 | 0.233598 | 0.18488 | 0.53379 | 0.25804 | 0.11876 | 20.8819 | 0.13 | 0 | Neutral |
| testneg_5645 | -0.04 | 0.23 | 6.027034 | -3 | 3.447125 | -3 | 0.38749 | 0.1076 | 0.29372 | 0.07609 | 25.89569 | 0.298221154 | 1.6 | Neutral |
| testneg_5646 | 0 | 0 | 0.789516 | -3 | 0.780247 | -3 | 0.99564 | 0.25794 | 0.26524 | 0.11154 | 13.22362 | 0.022197318 | 0 | Neutral |
| testneg_5647 | 0 | 0 | 0.146321 | -0.03734 | 1.491195 | -3 | 0.11827 | 0.23034 | 0.03391 | 0.09745 | 32.40108 | 0.04 | 0 | Neutral |
| testneg_5648 | -0.18 | -0.16 | 2.954569 | -3 | 5.882844 | -3 | 0.58677 | 0.11579 | 0.10133 | 0.32155 | 18.55527 | 0.57467803 | 80.8 | Disease_causing |
| testneg_5649 | 0.00727273 | 0.24236364 | 12.218087 | -3 | 4.028152 | -3 | 0.08932 | 0.32882 | 0.19898 | 0.12788 | 18.33333 | 0.7 | 98.4 | Disease_causing |
| testneg_5650 | 0.08 | 0.03 | -0.196592 | -0.196592 | 2.575019 | -3 | 0.32733 | 0.16869 | 0.28269 | 0.12505 | 20.95603 | 0.09026129 | 0 | Neutral |
| testneg_5651 | 0 | 0 | 1.527462 | -3 | 0.774434 | -3 | 0.25528 | 0.24339 | 0.09235 | 0.18135 | 10.83333 | 0.088212302 | 0 | Neutral |
| testneg_5652 | 0 | 0 | 2.397094 | -3 | 4.665413 | -3 | 0.22364 | 0.12788 | 0.39225 | 0.10625 | 16.84579 | 0.051267403 | 0 | Neutral |
| testneg_5653 | 0.02 | -0.28 | 0.372767 | -3 | 0.930465 | -3 | 0.08684 | 0.08917 | 0.30429 | 0.29938 | 16.8018 | 0.060563492 | 0 | Neutral |
| testneg_5654 | -0.21 | -0.15 | -0.83994 | -3 | 1.111997 | -3 | 0.95319 | 0.20297 | 0.06957 | 0.07707 | 16.67825 | 0.070801282 | 0 | Neutral |
| testneg_5655 | 0.03 | -1.03 | 3.488962 | -3 | -0.416655 | -0.416655 | 0.59245 | 0.12788 | 0.05453 | 0.12788 | 23.68839 | 0.050227273 | 0 | Neutral |
| testneg_5656 | 0 | 0 | 1.609501 | -3 | 0.725472 | -3 | 0.16404 | 0.09378 | 0.19898 | 0.06726 | 41.31944 | 9.52E-05 | 0 | Neutral |
| testneg_5657 | 0 | 0 | 0.327712 | -3 | 1.136968 | -3 | 0.57517 | 0.10873 | 0.31744 | 0.10986 | 13.51351 | 0.000235327 | 0 | Neutral |
| testneg_5658 | 0.06 | -0.57 | 2.146864 | -3 | 13.172408 | -3 | 0.19898 | 0.09316 | 0.13433 | 0.09578 | 19.90385 | 0.347644928 | 5.6 | Neutral |
| testneg_5659 | 0 | 0 | -0.680927 | -3 | 5.028164 | -3 | 0.0832 | 0.09323 | 0.29209 | 0.09791 | 13.51351 | 0.045371418 | 0 | Neutral |
| testneg_5660 | 0 | 0 | 0.194184 | -3 | 0.819832 | -3 | 0.18744 | 0.09392 | 0.19898 | 0.12788 | 37.92207 | 0.000267214 | 0 | Neutral |
| testneg_5661 | 0.04 | -0.23 | 0.007971 | -3 | 0.157379 | -3 | 0.12591 | 0.09845 | 0.45997 | 0.15162 | 27.95796 | 0.050824542 | 0 | Neutral |
| testneg_5662 | -0.19 | -0.51 | -0.025859 | -3 | 0.790255 | -3 | 0.11359 | 0.08631 | 0.25236 | 0.11809 | 10.30286 | 0.06 | 0 | Neutral |
| testneg_5663 | -0.03 | -1.22 | 4.80743 | -0.181311 | 6.95989 | -3 | 0.12513 | 0.12171 | 0.16542 | 0.54222 | 13.04237 | 0.740228758 | 99.4 | Disease_causing |
| testneg_5664 | 0 | 0 | 1.972273 | -3 | -1.826016 | -3 | 0.27187 | 0.41985 | 0.19898 | 0.12788 | 10.41667 | 0.02 | 0 | Neutral |
| testneg_5665 | -0.16 | -0.08 | -1.975003 | -1.975003 | 6.136401 | -3 | 0.08465 | 0.15428 | 0.3238 | 0.18333 | 10 | 0.645947938 | 94.8 | Disease_causing |
| testneg_5666 | 0.07 | -0.31 | 1.843162 | -3 | 5.03044 | -3 | 0.30869 | 0.11948 | 0.10561 | 0.12788 | 18.33333 | 0.0925 | 0 | Neutral |
| testneg_5667 | 0 | 0 | 2.074832 | -3 | 6.856644 | -3 | 0.17199 | 0.10586 | 0.05221 | 0.12788 | 18.95966 | 0.220135135 | 1.2 | Neutral |
| testneg_5668 | -0.15 | -0.04 | 2.269407 | -3 | 0.892974 | 0.892974 | 0.98442 | 0.44966 | 0.06986 | 0.12788 | 25.48653 | 0.100679825 | 0 | Neutral |
| testneg_5669 | 0 | 0 | 0.227306 | -3 | 0.448407 | -3 | 0.30412 | 0.54626 | 0.12231 | 0.23155 | 11.95767 | 0.139058686 | 0 | Neutral |
| testneg_5670 | 0 | 0 | 1.263979 | -3 | 0.191986 | -3 | 0.14901 | 0.12788 | 0.03235 | 0.0948 | 18.33333 | 0.000306486 | 0 | Neutral |
| testneg_5671 | -0.08 | -0.03 | -0.867132 | -3 | 1.375504 | -3 | 0.08353 | 0.21479 | 0.19898 | 0.12788 | 24.44444 | 0.063051619 | 0 | Neutral |
| testneg_5672 | 0.02 | 0.45 | 5.906217 | -3 | -2.624716 | -3 | 0.72163 | 0.10596 | 0.05932 | 0.09551 | 12.66993 | 0.2 | 0 | Neutral |
| testneg_5673 | 0 | 0 | 1.928458 | -3 | 4.83533 | -3 | 0.45152 | 0.34457 | 0.23975 | 0.13861 | 18.33333 | 0.450846996 | 35.4 | Neutral |
| testneg_5674 | 0.03 | -1.03 | 2.514378 | -3 | 0.27289 | 0.27289 | 0.3793 | 0.24105 | 0.14668 | 0.25103 | 10.83333 | 0.1775 | 0 | Neutral |
| testneg_5675 | 0.07 | 0.38 | 2.92632 | -3 | 0.034571 | -3 | 0.47442 | 0.11873 | 0.48561 | 0.09034 | 27.12306 | 0.03 | 0 | Neutral |
| testneg_5676 | -0.02 | 0.68 | 2.648194 | -3 | 2.980314 | -3 | 0.04655 | 0.10225 | 0.19898 | 0.535 | 4.72222 | 0.310794492 | 2.4 | Neutral |
| testneg_5677 | 0.09 | -0.3 | 1.940809 | -3 | 1.26259 | 1.26259 | 0.23278 | 0.11027 | 0.44722 | 0.11641 | 10.29033 | 0.02 | 0 | Neutral |
| testneg_5678 | 0.17 | 0.38 | 3.17007 | -3 | 0.71235 | -3 | 0.12377 | 0.12788 | 0.15152 | 0.12645 | 29.08136 | 0.03 | 0 | Neutral |
| testneg_5679 | 0.02 | -0.68 | 4.742104 | -3 | 0.931031 | -3 | 0.21623 | 0.42074 | 0.23939 | 0.25284 | 11.25 | 0.411416667 | 24 | Neutral |
| testneg_5680 | 0 | -0.32 | 6.71239 | -3 | 0.823839 | -3 | 0.68568 | 0.65219 | 0.13031 | 0.85678 | 23.88889 | 0.608856209 | 90.2 | Disease_causing |
| testneg_5681 | 0.1 | 1.9 | 0.416226 | -3 | 4.251821 | -3 | 0.32111 | 0.12788 | 0.05754 | 0.10881 | 10.22898 | 0.16 | 0 | Neutral |
| testneg_5682 | 0.21 | 0.15 | 6.051557 | -3 | 2.65719 | -3 | 0.13482 | 0.12788 | 0.12185 | 0.08854 | 17.21847 | 0.22 | 0.4 | Neutral |
| testneg_5683 | 0.01509091 | -0.312 | 11.127786 | -3 | -0.21059 | -3 | 0.10917 | 0.12788 | 0.03443 | 0.1032 | 32.29167 | 0.31 | 5.4 | Neutral |
| testneg_5684 | 0 | 0 | 0.74359 | -3 | 5.700045 | -3 | 0.25528 | 0.24339 | 0.12682 | 0.11357 | 15.16061 | 0.13529326 | 0 | Neutral |
| testneg_5685 | 0 | -0.01 | 0.186761 | -3 | 4.705574 | -3 | 0.19898 | 0.12788 | 0.17478 | 0.1501 | 15.80081 | 0.200870656 | 0.2 | Neutral |
| testneg_5686 | 0.07 | 0.38 | -0.983957 | -3 | 0.323865 | -3 | 0.14901 | 0.12788 | 0.63737 | 0.18081 | 12.70833 | 0.0325 | 0 | Neutral |
| testneg_5687 | 0.03 | 1.22 | 5.911847 | -3 | 0.714207 | -3 | 0.40179 | 0.12112 | 0.04346 | 0.10785 | 11.98582 | 0.19 | 0.2 | Neutral |
| testneg_5688 | 0.01666667 | 0.058 | 13.335291 | -3 | 3.602311 | -3 | 0.11359 | 0.08631 | 0.65048 | 0.12778 | 13.51351 | 0.458333333 | 36.6 | Neutral |
| testneg_5689 | 0 | 0 | 0.330109 | -0.808449 | 1.839289 | -3 | 0.13258 | 0.9258 | 0.29672 | 0.10485 | 10.22898 | 0.08 | 0 | Neutral |
| testneg_5690 | -0.15 | 0.28 | -0.410038 | -3 | 1.377333 | -3 | 0.21848 | 0.11049 | 0.06215 | 0.08575 | 29.16667 | 0.05 | 0 | Neutral |
| testneg_5691 | 0.07 | 0.38 | -1.027981 | -3 | -0.019972 | -0.019972 | 0.10133 | 0.32155 | 0.05245 | 0.13458 | 24.70085 | 0.205388889 | 0 | Neutral |
| testneg_5692 | -0.03 | -0.1 | 0.212913 | -3 | 2.414577 | -3 | 0.11391 | 0.19434 | 0.2439 | 0.08848 | 11.01208 | 0.08 | 0 | Neutral |
| testneg_5693 | 0 | -0.01 | 1.241126 | -3 | 1.223596 | -3 | 0.89959 | 0.55078 | 0.24519 | 0.12788 | 10.41667 | 0.011433848 | 0 | Neutral |
| testneg_5694 | 0.15 | -0.28 | 0.118932 | 0.118932 | 0.497371 | -3 | 0.19567 | 0.09606 | 0.21983 | 0.10146 | 12.46106 | 0.010177116 | 0 | Neutral |
| testneg_5695 | 0 | 0 | 0.307719 | -3 | 0.131182 | -3 | 0.16091 | 0.11571 | 0.44429 | 0.20468 | 16.84579 | 0.020316964 | 0 | Neutral |
| testneg_5696 | 0 | 0 | 0.178815 | -3 | 0.949523 | 0.949523 | 0.5471 | 0.11262 | 0.10056 | 0.12608 | 10.1991 | 0.02 | 0 | Neutral |
| testneg_5697 | 0.04 | -0.23 | 0.659043 | 0.659043 | 0.743493 | -3 | 0.03137 | 0.12788 | 0.11703 | 0.09364 | 49.06907 | 0.07 | 0 | Neutral |
| testneg_5698 | -0.07 | -0.38 | 2.597387 | -3 | -0.020693 | -3 | 0.08579 | 0.6068 | 0.80034 | 0.10518 | 10.22898 | 0.100717949 | 0 | Neutral |
| testneg_5699 | -0.22 | 2.66 | -1.189105 | -3 | 2.830718 | -3 | 0.25561 | 0.11004 | 0.11229 | 0.08897 | 26.04838 | 0.193095238 | 0 | Neutral |
| testneg_5700 | 0 | 0 | 1.420779 | -1.808697 | 3.705834 | -3 | 0.14386 | 0.12788 | 0.16781 | 0.12788 | 36.11111 | 0.041602564 | 0 | Neutral |
| testneg_5701 | 0.14 | -0.37 | 6.930926 | -3 | 4.995228 | 0.16841 | 0.41743 | 0.78545 | 0.50392 | 0.12788 | 10.41667 | 0.785139296 | 100 | Disease_causing |
| testneg_5702 | 0 | 0 | 0.720837 | -3 | 0.509747 | -3 | 0.51398 | 0.38821 | 0.19898 | 0.08486 | 11.23984 | 0.014642857 | 0 | Neutral |
| testneg_5703 | 0 | 0 | 2.437503 | -3 | -0.061002 | -0.061002 | 0.11337 | 0.09363 | 0.35561 | 0.08845 | 23.16963 | 0.01 | 0 | Neutral |
| testneg_5704 | 0.22 | -2.66 | 6.932503 | -3 | 6.206483 | -3 | 0.74271 | 0.10951 | 0.18861 | 0.10842 | 7.01832 | 0.610119048 | 87.8 | Disease_causing |
| testneg_5705 | -0.004965 | 0.03867133 | 1.375958 | -3 | 3.074697 | -3 | 0.0538 | 0.23912 | 0.37282 | 0.07943 | 33.76574 | 0.16126486 | 0 | Neutral |
| testneg_5706 | 0 | 0 | -0.433937 | -3 | 5.243459 | -3 | 0.84572 | 0.78546 | 0.20918 | 0.10825 | 11.25 | 0.1128738 | 0 | Neutral |
| testneg_5707 | -0.17 | -0.38 | 1.808824 | 1.808824 | 0.696041 | -3 | 0.2445 | 0.53069 | 0.78515 | 0.35093 | 4.44444 | 0.3 | 2 | Neutral |
| testneg_5708 | 0.18 | 0.16 | 2.016768 | -3 | 0.305495 | -3 | 0.29297 | 0.11049 | 0.98255 | 0.98129 | 4.72222 | 0.12 | 0 | Neutral |
| testneg_5709 | 0 | 0 | 0.111854 | -3 | 4.406394 | -3 | 0.14462 | 0.11239 | 0.07577 | 0.09128 | 20.83333 | 0.055462454 | 0 | Neutral |
| testneg_5710 | 0.14 | -1.01 | 1.674406 | -3 | 2.815622 | -3 | 0.24 | 0.2063 | 0.46037 | 0.1343 | 16.89815 | 0.316026611 | 2.6 | Neutral |
| testneg_5711 | 0.04 | -0.23 | 1.849334 | -3 | -0.454393 | -1.150939 | 0.03788 | 0.10996 | 0.03443 | 0.1032 | 10.41667 | 0.04 | 0 | Neutral |
| testneg_5712 | 0.18 | -1.05 | 4.646906 | -3 | 4.444128 | -3 | 0.09508 | 0.09834 | 0.09235 | 0.18135 | 18.33333 | 0.6 | 84.2 | Disease_causing |
| testneg_5713 | 0.05 | -0.81 | 3.064064 | -3 | -1.378191 | -3 | 0.5481 | 0.10207 | 0.19898 | 0.12788 | 11.74429 | 0.04 | 0 | Neutral |
| testneg_5714 | -0.18 | 1.05 | 4.595565 | -3 | 3.125039 | -3 | 0.19898 | 0.12788 | 0.18147 | 0.12788 | 13.2927 | 0.296524621 | 1.6 | Neutral |
| testneg_5715 | -0.07 | -0.38 | 0.704598 | -3 | 5.518112 | -3 | 0.09657 | 0.58605 | 0.32977 | 0.15427 | 9.862 | 0.650579205 | 95.6 | Disease_causing |
| testneg_5716 | 0 | 0 | -0.352016 | -3 | 0.746831 | -3 | 0.17264 | 0.09426 | 0.16542 | 0.54222 | 13.57809 | 0.04 | 0 | Neutral |
| testneg_5717 | 0.05 | -0.81 | 2.670334 | -3 | 5.124676 | -3 | 0.31906 | 0.1705 | 0.26068 | 0.46457 | 18.33333 | 0.694060987 | 98 | Disease_causing |
| testneg_5718 | 0 | 0 | -0.164302 | -3 | 1.583634 | -3 | 0.35475 | 0.12788 | 0.12642 | 0.11218 | 18.33333 | 0.00046883 | 0 | Neutral |
| testneg_5719 | 0 | -0.32 | -0.359045 | -3 | 0.456586 | 0.456586 | 0.30951 | 0.64738 | 0.62029 | 0.09082 | 10.83333 | 0.06 | 0 | Neutral |
| testneg_5720 | 0 | -0.01 | 0.36151 | -3 | -1.566121 | -1.566121 | 0.28878 | 0.16765 | 0.12003 | 0.12788 | 39.60168 | 0.021306391 | 0 | Neutral |
| testneg_5721 | 0.02 | 0.45 | 1.21068 | -3 | 1.121308 | -3 | 0.95247 | 0.11285 | 0.33418 | 0.09979 | 19.7626 | 0.02 | 0 | Neutral |
| testneg_5722 | -0.08 | -0.03 | 1.483649 | 1.483649 | 4.066081 | -3 | 0.47678 | 0.38085 | 0.73008 | 0.16585 | 9.80134 | 0.471253501 | 44.6 | Neutral |
| testneg_5723 | -0.08 | -0.03 | 0.73341 | -3 | 0.632891 | -3 | 0.54792 | 0.54626 | 0.28856 | 0.1544 | 21.82594 | 0.17052139 | 0 | Neutral |
| testneg_5724 | -0.1 | -1.9 | -1.487847 | -3 | 4.187854 | -3 | 0.06728 | 0.20664 | 0.09176 | 0.07645 | 39.02465 | 0.25225 | 0 | Neutral |
| testneg_5725 | 0.02 | -0.63 | -2.323867 | -3 | 3.381638 | 3.381638 | 0.07595 | 0.11822 | 0.07552 | 0.12788 | 30.32444 | 0.194672619 | 0 | Neutral |
| testneg_5726 | -0.08 | 1.2 | 1.189976 | -3 | -1.05412 | -3 | 0.17303 | 0.18733 | 0.24791 | 0.09675 | 6.94444 | 0.08 | 0 | Neutral |
| testneg_5727 | -0.03 | 1.03 | 2.078465 | -3 | 0.604567 | -3 | 0.99665 | 0.12884 | 0.24964 | 0.12788 | 32.85473 | 0.07 | 0 | Neutral |
| testneg_5728 | -0.03 | -0.1 | 1.820578 | -3 | 1.15789 | 1.15789 | 0.31507 | 0.61057 | 0.68439 | 0.09371 | 10 | 0.07 | 0 | Neutral |
| testneg_5729 | 0.04 | -0.23 | 2.988584 | -3 | -0.409229 | -3 | 0.8611 | 0.12788 | 0.12282 | 0.20617 | 11.25 | 0.14 | 0 | Neutral |
| testneg_5730 | 0.04 | -0.23 | 3.570813 | -3 | 1.192651 | -3 | 0.68054 | 0.09208 | 0.65952 | 0.49708 | 10.72411 | 0.162081197 | 0 | Neutral |
| testneg_5731 | -0.06 | -0.18 | 1.285578 | -3 | -0.329218 | -3 | 0.087 | 0.12788 | 0.13532 | 0.19904 | 15.5103 | 0.06176511 | 0 | Neutral |
| testneg_5732 | -0.02 | 0.63 | 2.451821 | -3 | 0.188403 | -3 | 0.25025 | 0.18924 | 0.39111 | 0.15887 | 19.13908 | 0.150552738 | 0 | Neutral |
| testneg_5733 | -0.16 | -0.08 | 3.904599 | -3 | 0.190526 | -0.494899 | 0.07427 | 0.09451 | 0.1834 | 0.12788 | 29.23973 | 0.1 | 0 | Neutral |

| | | | | | | | | | | | | | | |
|---|---|---|---|---|---|---|---|---|---|---|---|---|---|---|
| testneg_5734 | -0.01 | 0.57 | 0.464102 | -3 | -3.138593 | -3 | 0.14668 | 0.25103 | 0.06254 | 0.23561 | 12.16268 | 0.141070676 | 0 | Neutral |
| testneg_5735 | 0 | 0 | 0.58387 | -3 | 1.460591 | -3 | 0.13244 | 0.14875 | 0.53996 | 0.14577 | 19.01709 | 0.061065205 | 0 | Neutral |
| testneg_5736 | -0.09 | 0.3 | -0.355202 | -3 | 0.316188 | -3 | 0.21802 | 0.10187 | 0.36852 | 0.0903 | 55.62414 | 0.040313259 | 0 | Neutral |
| testneg_5737 | -0.03 | -0.1 | 2.058438 | -3 | -0.718699 | -3 | 0.0749 | 0.12788 | 0.21122 | 0.12788 | 17.94606 | 0.01 | 0 | Neutral |
| testneg_5738 | 0.17 | 0.38 | 3.132136 | -3 | -3.528995 | -3 | 0.0105 | 0.12788 | 0.29525 | 0.12788 | 30.35589 | 0.08 | 0 | Neutral |
| testneg_5739 | 0 | 0 | 1.777966 | -3 | 2.191142 | -3 | 0.33264 | 0.10493 | 0.39225 | 0.10625 | 20.89747 | 0.01 | 0 | Neutral |
| testneg_5740 | -0.1 | -1.9 | 1.551505 | -3 | 1.139058 | -3 | 0.05413 | 0.05691 | 0.27887 | 0.18663 | 10.83333 | 0.16 | 0 | Neutral |
| testneg_5741 | 0.07 | 0.38 | -2.352243 | -3 | 7.49852 | -3 | 0.04568 | 0.08746 | 0.3112 | 0.33703 | 18.89436 | 0.413791667 | 21.8 | Neutral |
| testneg_5742 | 0 | 0 | 0.094735 | -3 | 2.162874 | -3 | 0.024 | 0.07726 | 0.54625 | 0.11856 | 31.26016 | 0.04 | 0 | Neutral |
| testneg_5743 | 0.16 | 0.08 | -0.28306 | -3 | 1.884645 | -3 | 0.44067 | 0.11201 | 0.10285 | 0.11721 | 12.46106 | 0.03 | 0 | Neutral |
| testneg_5744 | 0 | 0 | -0.137475 | -0.137475 | 4.280436 | -3 | 0.19898 | 0.12788 | 0.40116 | 0.43982 | 21.42525 | 0.306252333 | 2.6 | Neutral |
| testneg_5745 | 0 | -0.01 | 0.571549 | -3 | 0.820444 | 0.820444 | 0.48615 | 0.14369 | 0.17736 | 0.14772 | 14.68921 | 0.059302926 | 0 | Neutral |
| testneg_5746 | 0.15 | -0.28 | 1.731846 | -3 | 1.682022 | -3 | 0.83067 | 0.10583 | 0.06895 | 0.06354 | 19.96169 | 0.020235327 | 0 | Neutral |
| testneg_5747 | 0.14 | -1.01 | 6.115432 | -3 | -1.043136 | -3 | 0.19898 | 0.11167 | 0.19898 | 0.12788 | 21.56188 | 0.128035714 | 0 | Neutral |
| testneg_5748 | 0 | 0 | 0.323374 | -3 | -0.16841 | -3 | 0.04853 | 0.4186 | 0.21972 | 0.12788 | 26.19195 | 0.11 | 0 | Neutral |
| testneg_5749 | 0.05 | -0.81 | 2.468627 | -3 | 1.493352 | -3 | 0.69249 | 0.12788 | 0.17364 | 0.18111 | 18.33184 | 0.08 | 0 | Neutral |
| testneg_5750 | -0.04 | 0.23 | -0.274997 | -3 | -0.187365 | -3 | 0.37501 | 0.0997 | 0.6612 | 0.16696 | 10.41667 | 0.041154695 | 0 | Neutral |
| testneg_5751 | 0 | -0.01 | 0.981481 | -3 | 0.864946 | -3 | 0.24516 | 0.11809 | 0.10822 | 0.10942 | 29.11853 | 0.000311872 | 0 | Neutral |
| testneg_5752 | 0 | 0 | 1.201989 | -3 | 0.383078 | -3 | 0.2231 | 0.22973 | 0.25625 | 0.66481 | 18.88889 | 0.110618412 | 0 | Neutral |
| testneg_5753 | 0.2 | -1.26 | 2.143533 | -3 | 0.807879 | -3 | 0.40897 | 0.13253 | 0.15994 | 0.11186 | 19.01053 | 0.042666667 | 0 | Neutral |
| testneg_5754 | 0.21 | 0.15 | 2.726693 | -3 | 0.870722 | -3 | 0.28227 | 0.10211 | 0.1193 | 0.5592 | 16.40693 | 0.168875 | 0 | Neutral |
| testneg_5755 | -0.07 | -0.38 | 3.319584 | -3 | 2.598788 | -3 | 0.99543 | 0.12788 | 0.19898 | 0.10307 | 35.39671 | 0.1 | 0 | Neutral |
| testneg_5756 | 0.22 | -2.66 | 6.203124 | -3 | 1.175658 | -3 | 0.10581 | 0.11631 | 0.10285 | 0.10886 | 31.53099 | 0.15 | 0 | Neutral |
| testneg_5757 | -0.07 | -0.38 | 1.749962 | -3 | 6.384418 | -3 | 0.04916 | 0.14944 | 0.84431 | 0.65315 | 10 | 0.673066436 | 96.8 | Disease_causing |
| testneg_5758 | 0 | 0 | 0.305495 | -3 | -0.398253 | -3 | 0.98255 | 0.98129 | 0.04668 | 0.12788 | 10 | 0.070687343 | 0 | Neutral |
| testneg_5759 | -0.03 | 0.85 | 5.601904 | -3 | 3.046889 | -3 | 0.16728 | 0.11868 | 0.1066 | 0.12788 | 10 | 0.365527778 | 12.2 | Neutral |
| testneg_5760 | -0.07 | -0.38 | -0.395207 | -3 | 6.598988 | -3 | 0.2098 | 0.44672 | 0.1454 | 0.10285 | 19.09347 | 0.354140997 | 9.2 | Neutral |
| testneg_5761 | 0 | 0 | -0.681556 | -3 | -0.030574 | -3 | 0.42585 | 0.18081 | 0.22972 | 0.17259 | 5.45324 | 0.040566756 | 0 | Neutral |
| testneg_5762 | -0.08 | -0.03 | 5.449749 | -3 | -0.175246 | -3 | 0.15041 | 0.43468 | 0.12969 | 0.09906 | 11.25 | 0.28 | 1 | Neutral |
| testneg_5763 | 0.14 | -0.37 | 0.898194 | -3 | -0.086008 | -3 | 0.0925 | 0.08222 | 0.86437 | 0.69972 | 10.41667 | 0.08377381 | 0 | Neutral |
| testneg_5764 | -0.2 | 1.26 | 6.594372 | -3 | 3.619507 | -3 | 0.16841 | 0.11143 | 0.67676 | 0.12788 | 20.20624 | 0.47 | 42.8 | Neutral |
| testneg_5765 | 0.17 | 0.38 | -1.859511 | -3 | 1.334446 | 1.334446 | 0.41193 | 0.10133 | 0.15514 | 0.14852 | 10.83333 | 0.106199576 | 0 | Neutral |
| testneg_5766 | 0 | 0 | 1.708897 | -3 | -0.121121 | -3 | 0.5202 | 0.16306 | 0.213 | 0.09535 | 13.22863 | 0.02 | 0 | Neutral |
| testneg_5767 | 0 | 0 | -0.43021 | -3 | -0.212616 | -3 | 0.17041 | 0.12378 | 0.2674 | 0.28725 | 12.29167 | 0.022374365 | 0 | Neutral |
| testneg_5768 | 0.01 | 0.8 | 5.151503 | -3 | 6.007104 | 6.007104 | 0.21399 | 0.10708 | 0.17764 | 0.09681 | 32.49042 | 0.56 | 80.6 | Disease_causing |
| testneg_5769 | 0.00545455 | -0.0303636 | 5.729308 | -3 | -2.768659 | -3 | 0.06793 | 0.12788 | 0.34399 | 0.08193 | 28.06373 | 0.13 | 0 | Neutral |
| testneg_5770 | -0.0665455 | 0.28327273 | 13.012965 | -3 | -1.024972 | -3 | 0.33166 | 0.11755 | 0.23955 | 0.97212 | 5 | 0.51 | 62.6 | Disease_causing |
| testneg_5771 | 0.09 | -0.3 | 5.225996 | -3 | 5.545529 | -3 | 0.34698 | 0.12788 | 0.29172 | 0.08143 | 16.42913 | 0.33 | 7.2 | Neutral |
| testneg_5772 | 0 | 0 | 1.687316 | -3 | 0.477392 | -3 | 0.26434 | 0.12788 | 0.09682 | 0.09596 | 13.08587 | 0.000117155 | 0 | Neutral |
| testneg_5773 | 0 | 0 | 0.021671 | -3 | 0.452441 | 0.452441 | 0.13156 | 0.10621 | 0.48437 | 0.09986 | 21.85185 | 0.01 | 0 | Neutral |
| testneg_5774 | 0.09 | -0.3 | 5.817553 | -3 | 4.627145 | -3 | 0.85075 | 0.16646 | 0.19898 | 0.12788 | 6.94444 | 0.61 | 88 | Disease_causing |
| testneg_5775 | 0 | 0 | 1.757755 | -3 | 6.823573 | -3 | 0.9193 | 0.14584 | 0.07406 | 0.08059 | 11.82267 | 0.22 | 0.4 | Neutral |
| testneg_5776 | 0 | 0 | 1.963971 | -3 | 1.019099 | -3 | 0.9193 | 0.14584 | 0.19898 | 0.12788 | 13.89558 | 0.01 | 0 | Neutral |
| testneg_5777 | 0 | -0.01 | 0.129221 | -3 | 1.606239 | 1.606239 | 0.13794 | 0.40936 | 0.89002 | 0.12788 | 10.41667 | 0.05 | 0 | Neutral |
| testneg_5778 | -0.11 | 0.19 | 5.824607 | -3 | 0.278768 | -3 | 0.03391 | 0.09745 | 0.1327 | 0.09885 | 33.95664 | 0.181547619 | 0 | Neutral |
| testneg_5779 | 0.03 | 1.22 | 4.311522 | -3 | 2.617333 | -3 | 0.08857 | 0.83499 | 0.13007 | 0.12788 | 17.21847 | 0.345 | 6.2 | Neutral |
| testneg_5780 | -0.09 | 0.3 | -0.522358 | -3 | 1.123147 | -3 | 0.12836 | 0.09736 | 0.59947 | 0.14339 | 5.38184 | 0.06 | 0 | Neutral |
| testneg_5781 | 0.06 | 0.18 | 5.333191 | -3 | 4.895689 | -3 | 0.19898 | 0.12788 | 0.23385 | 0.12788 | 12.52313 | 0.415829365 | 26.6 | Neutral |
| testneg_5782 | 0 | 0 | 1.365392 | -3 | 1.84534 | -3 | 0.72062 | 0.14321 | 0.40996 | 0.57507 | 22.15335 | 0.091417537 | 0 | Neutral |
| testneg_5783 | -0.09 | 0.3 | 4.377701 | -3 | 2.354402 | -3 | 0.07189 | 0.11325 | 0.0538 | 0.0823 | 10.41667 | 0.12 | 0 | Neutral |
| testneg_5784 | -0.09 | 0.25 | 5.141418 | -3 | -0.290972 | -3 | 0.99689 | 0.28104 | 0.62273 | 0.10215 | 19.37495 | 0.21 | 0 | Neutral |
| testneg_5785 | 0.2 | -1.26 | 3.203428 | -3 | 6.499399 | -3 | 0.35325 | 0.12614 | 0.3038 | 0.11278 | 17.21847 | 0.269933333 | 3 | Neutral |
| testneg_5786 | 0 | 0 | 0.984412 | -3 | 0.120705 | -3 | 0.34931 | 0.11906 | 0.08982 | 0.15279 | 10.83333 | 0.04052305 | 0 | Neutral |
| testneg_5787 | 0 | -0.32 | 3.279823 | -3 | 1.120297 | -3 | 0.07189 | 0.11325 | 0.10919 | 0.105 | 18.88889 | 0.03 | 0 | Neutral |
| testneg_5788 | 0.16 | -0.37 | -2.713571 | -2.713571 | 4.21286 | -3 | 0.07506 | 0.09112 | 0.07447 | 0.23492 | 26.34553 | 0.300264808 | 0.2 | Neutral |
| testneg_5789 | 0 | 0 | -0.043077 | -3 | -1.333487 | -3 | 0.15055 | 0.22097 | 0.07598 | 0.08456 | 19.18549 | 0.03 | 0 | Neutral |
| testneg_5790 | 0 | -0.32 | 0.489857 | -3 | 0.557964 | -3 | 0.36257 | 0.26115 | 0.10185 | 0.12788 | 35.11316 | 0.035 | 0 | Neutral |
| testneg_5791 | -0.19 | -0.51 | -1.62333 | -3 | -0.887241 | -3 | 0.09661 | 0.09981 | 0.19898 | 0.12788 | 13.51351 | 0.0675 | 0 | Neutral |
| testneg_5792 | -0.14 | 1.01 | 0.576229 | -3 | 1.414655 | -3 | 0.18022 | 0.12788 | 0.19898 | 0.12788 | 13.46801 | 0.05 | 0 | Neutral |
| testneg_5793 | 0.07 | 0.38 | 0.991286 | -3 | -0.266975 | -3 | 0.2439 | 0.08848 | 0.1294 | 0.10964 | 11.01208 | 0.02 | 0 | Neutral |
| testneg_5794 | 0 | 0 | -0.757789 | -3 | 3.107738 | -3 | 0.40491 | 0.17596 | 0.49012 | 0.12138 | 18.77814 | 0.063778721 | 0 | Neutral |
| testneg_5795 | 0.18 | 0.16 | -0.580188 | -3 | 0.868464 | -3 | 0.1675 | 0.09413 | 0.07204 | 0.07708 | 18.40376 | 0.024875 | 0 | Neutral |
| testneg_5796 | -0.0101667 | 0.00766667 | 0.837529 | -3 | 4.841652 | -3 | 0.26913 | 0.1087 | 0.57188 | 0.34038 | 14.16667 | 0.250823003 | 0.6 | Neutral |
| testneg_5797 | 0.21 | 0.15 | 1.623618 | -3 | 5.224304 | -3 | 0.10083 | 0.17511 | 0.0959 | 0.12788 | 24.13428 | 0.250904762 | 0.2 | Neutral |
| testneg_5798 | 0 | -0.01 | 0.328189 | -3 | 5.007683 | -3 | 0.33957 | 0.12788 | 0.07415 | 0.12788 | 10.65539 | 0.06 | 0 | Neutral |
| testneg_5799 | 0.03 | -1.03 | 3.635917 | -3 | 0.827982 | -3 | 0.75978 | 0.11423 | 0.12231 | 0.23155 | 20.00448 | 0.174378543 | 0 | Neutral |
| testneg_5800 | 0 | 0 | 1.125734 | -3 | 0.974808 | 0.974808 | 0.12576 | 0.08486 | 0.32992 | 0.33227 | 11.43185 | 0.056152597 | 0 | Neutral |
| testneg_5801 | 0.07 | -0.31 | 0.268697 | -3 | -0.303923 | -3 | 0.20657 | 0.12788 | 0.04593 | 0.09315 | 10 | 0.01 | 0 | Neutral |
| testneg_5802 | -0.18 | -0.16 | 6.127805 | -3 | 1.473689 | -3 | 0.67549 | 0.1273 | 0.21677 | 0.1681 | 19.45931 | 0.48 | 47 | Neutral |
| testneg_5803 | 0 | 0 | 1.24594 | -3 | 3.159116 | -3 | 0.19898 | 0.21177 | 0.54891 | 0.55746 | 12.7356 | 0.290261604 | 2.2 | Neutral |
| testneg_5804 | 0 | 0 | 1.13018 | -3 | 2.111463 | -3 | 0.50505 | 0.11563 | 0.09815 | 0.12788 | 22.89111 | 0.02 | 0 | Neutral |
| testneg_5805 | 0 | 0 | 1.329031 | -3 | 4.719495 | -3 | 0.22489 | 0.16646 | 0.24971 | 0.12971 | 19.44444 | 0.101461557 | 0 | Neutral |
| testneg_5806 | 0.04 | -0.23 | 4.018098 | 4.018098 | 3.146728 | -3 | 0.0606 | 0.39625 | 0.28856 | 0.1544 | 32.58465 | 0.808451438 | 100 | Disease_causing |
| testneg_5807 | 0 | 0 | 0.443289 | -3 | 0.246616 | -3 | 0.09972 | 0.13992 | 0.06955 | 0.1748 | 19.71131 | 0.097218254 | 0 | Neutral |
| testneg_5808 | -0.04 | 0.23 | 3.260847 | -3 | 5.107463 | -3 | 0.27605 | 0.12788 | 0.10244 | 0.10107 | 23.4127 | 0.189933783 | 0 | Neutral |
| testneg_5809 | -0.05 | 0.44 | 3.587885 | -3 | 4.819528 | -3 | 0.77644 | 0.09695 | 0.1211 | 0.1526 | 27.4024 | 0.426802399 | 27.4 | Neutral |
| testneg_5810 | 0.03 | 1.22 | 4.663109 | -3 | 4.167444 | -3 | 0.23292 | 0.15913 | 0.87806 | 0.38966 | 10.41667 | 0.657282687 | 95 | Disease_causing |
| testneg_5811 | 0 | 0 | 0.461408 | -3 | 4.193894 | -3 | 0.16797 | 0.11809 | 0.12855 | 0.23304 | 13.78523 | 0.208984761 | 0.4 | Neutral |
| testneg_5812 | 0.03 | -0.85 | 0.173309 | -3 | 1.036843 | -3 | 0.34694 | 0.20397 | 0.18102 | 0.6537 | 20.55556 | 0.162446267 | 0 | Neutral |
| testneg_5813 | 0.07 | 0.38 | 2.43609 | -3 | 1.662461 | -3 | 0.02146 | 0.10351 | 0.14646 | 0.36445 | 18.88889 | 0.15402381 | 0 | Neutral |
| testneg_5814 | -0.05 | -0.29 | -0.90037 | -3 | 1.541854 | -3 | 0.65564 | 0.11468 | 0.1065 | 0.12788 | 17.21847 | 0.015 | 0 | Neutral |
| testneg_5815 | 0.01 | 0.8 | 1.447055 | -3 | 4.53958 | -3 | 0.60959 | 0.76078 | 0.16636 | 0.12384 | 18.33333 | 0.231145048 | 0.2 | Neutral |
| testneg_5816 | 0 | 0 | 5.011748 | -3 | 2.821522 | -3 | 0.33135 | 0.73068 | 0.58994 | 0.11602 | 11.25 | 0.321487013 | 4 | Neutral |
| testneg_5817 | 0.06 | -0.57 | 3.73543 | -3 | 0.138401 | -1.616188 | 0.19898 | 0.12788 | 0.09527 | 0.12788 | 24.52719 | 0.025681818 | 0 | Neutral |
| testneg_5818 | 0.07 | 0.38 | 1.744917 | -3 | 1.872697 | -3 | 0.15581 | 0.19351 | 0.84468 | 0.55945 | 12.60802 | 0.139256107 | 0 | Neutral |
| testneg_5819 | -0.09 | 0.3 | 2.446797 | -3 | 3.832241 | 2.812197 | 0.61008 | 0.17538 | 0.213 | 0.09535 | 15.81732 | 0.257282898 | 0 | Neutral |
| testneg_5820 | 0.04 | -0.23 | 7.410019 | -3 | -0.605051 | -3 | 0.31318 | 0.11262 | 0.48144 | 0.46642 | 14.52969 | 0.408333333 | 28.2 | Neutral |
| testneg_5821 | 0.01 | -1.44 | 2.093021 | -3 | 1.668583 | -3 | 0.54724 | 0.12788 | 0.04869 | 0.05922 | 18.88889 | 0.04 | 0 | Neutral |
| testneg_5822 | 0.18 | -1.05 | 1.744357 | -3 | 5.941706 | -3 | 0.126 | 0.14076 | 0.32065 | 0.19642 | 11.25 | 0.523689535 | 62.6 | Disease_causing |
| testneg_5823 | 0 | 0 | 2.104789 | -3 | 2.5258 | -2.07002 | 0.54792 | 0.54626 | 0.48801 | 0.09118 | 12.39141 | 0.065227273 | 0 | Neutral |
| testneg_5824 | 0 | -0.01 | 2.05177 | -3 | 0.298115 | -3 | 0.09542 | 0.14806 | 0.5481 | 0.10207 | 19.58698 | 0.03 | 0 | Neutral |
| testneg_5825 | 0 | 0 | 1.56019 | -3 | -0.71253 | -3 | 0.25487 | 0.13098 | 0.03563 | 0.10455 | 18.88889 | 0.01 | 0 | Neutral |
| testneg_5826 | 0.07 | 0.33 | 3.81823 | -3 | 0.126798 | -3 | 0.1755 | 0.10455 | 0.21731 | 0.11795 | 14.70656 | 0.02 | 0 | Neutral |
| testneg_5827 | -0.08 | -0.03 | 2.557841 | -3 | -0.464766 | -3 | 0.19898 | 0.14896 | 0.62197 | 0.12788 | 11.28916 | 0.032640845 | 0 | Neutral |
| testneg_5828 | 0 | 0 | 0.663777 | -3 | 0.240038 | -3 | 0.68055 | 0.12788 | 0.19898 | 0.08111 | 17.77778 | 0.000513158 | 0 | Neutral |
| testneg_5829 | 0 | 0 | 7.325251 | -3 | 3.781746 | -3 | 0.16601 | 0.14015 | 0.52771 | 0.10974 | 26.76176 | 0.470313187 | 43.4 | Neutral |
| testneg_5830 | 0.01 | -0.57 | 4.956408 | -3 | 0.892974 | 0.892974 | 0.28341 | 0.1005 | 0.06986 | 0.12788 | 24.10613 | 0.07 | 0 | Neutral |
| testneg_5831 | 0 | 0 | 1.076963 | -3 | 0.236781 | -3 | 0.20011 | 0.27949 | 0.05216 | 0.23851 | 18.33333 | 0.1 | 0 | Neutral |
| testneg_5832 | -0.07 | -0.38 | -0.75095 | -3 | 1.720206 | -3 | 0.95076 | 0.17388 | 0.6992 | 0.1804 | 6.00263 | 0.08 | 0 | Neutral |
| testneg_5833 | -0.03 | -1.22 | 3.652963 | -3 | -0.187456 | -3 | 0.82269 | 0.15732 | 0.19898 | 0.12788 | 47.17357 | 0.09 | 0 | Neutral |
| testneg_5834 | 0 | 0 | 1.362887 | -3 | -0.907393 | -3 | 0.38527 | 0.24812 | 0.05332 | 0.12788 | 10.22898 | 0.01 | 0 | Neutral |
| testneg_5835 | 0 | 0 | 1.428854 | -3 | 3.456817 | -3 | 0.16728 | 0.11868 | 0.20329 | 0.10718 | 10.41667 | 0.02 | 0 | Neutral |
| testneg_5836 | 0.01 | 0.8 | 0.663866 | -3 | -0.299171 | -3 | 0.08283 | 0.11797 | 0.18753 | 0.09983 | 12.43421 | 0.010291685 | 0 | Neutral |
| testneg_5837 | 0.01 | 0.79 | 2.67742 | -3 | 2.341461 | -3 | 0.39553 | 0.11991 | 0.961 | 0.25991 | 21.40097 | 0.16075188 | 0 | Neutral |
| testneg_5838 | 0 | 0 | 0.889458 | 0.889458 | 0.212654 | -3 | 0.48937 | 0.22085 | 0.17173 | 0.15363 | 20.5518 | 0.151528043 | 0 | Neutral |
| testneg_5839 | 0.02 | -0.68 | 6.639247 | -3 | 1.102328 | -3 | 0.27407 | 0.1136 | 0.74271 | 0.10951 | 6.0461 | 0.246388889 | 1.4 | Neutral |
| testneg_5840 | 0.17 | 0.38 | 4.078311 | -3 | 0.241198 | -3 | 0.20951 | 0.16278 | 0.09806 | 0.12788 | 10.83333 | 0.09 | 0 | Neutral |
| testneg_5841 | 0.08 | 0.03 | -1.259442 | -3 | 1.697009 | -3 | 0.24185 | 0.12788 | 0.52771 | 0.10974 | 15.03121 | 0.02 | 0 | Neutral |
| testneg_5842 | 0.18 | 0.16 | 4.869046 | -3 | 5.312389 | -3 | 0.12957 | 0.12788 | 0.07293 | 0.12989 | 10.28409 | 0.357418277 | 11.6 | Neutral |
| testneg_5843 | 0 | 0 | 1.038487 | -3 | 2.787374 | -3 | 0.72124 | 0.11262 | 0.19898 | 0.12788 | 33.34622 | 0.03 | 0 | Neutral |
| testneg_5844 | -0.06 | -0.18 | 3.812626 | -3 | -0.044793 | -3 | 0.93942 | 0.23984 | 0.12513 | 0.12171 | 11.44725 | 0.070361502 | 0 | Neutral |
| testneg_5845 | 0.18 | -1.05 | 4.164795 | -3 | -0.763816 | -3 | 0.16091 | 0.90971 | 0.12403 | 0.12788 | 31.59722 | 0.209258242 | 0 | Neutral |
| testneg_5846 | -0.07 | -0.33 | -3.79699 | -3 | 0.083451 | -3 | 0.19898 | 0.12788 | 0.15994 | 0.11186 | 19.5422 | 0.02 | 0 | Neutral |
| testneg_5847 | 0.09 | -0.3 | 4.834383 | -3 | 0.622743 | -3 | 0.67796 | 0.10874 | 0.18979 | 0.07948 | 11.25 | 0.05 | 0 | Neutral |
| testneg_5848 | -0.07 | 0.31 | 3.270074 | -3 | -2.584648 | -3 | 0.1837 | 0.27351 | 0.0634 | 0.0824 | 11.92982 | 0.07 | 0 | Neutral |
| testneg_5849 | 0 | 0 | 2.171631 | -3 | 6.918031 | -3 | 0.94859 | 0.75045 | 0.1201 | 0.1062 | 10 | 0.31 | 4.4 | Neutral |
| testneg_5850 | 0 | 0 | 0.703801 | -3 | 0.461043 | -3 | 0.46723 | 0.08832 | 0.62864 | 0.11103 | 10.25326 | 0.01 | 0 | Neutral |
| testneg_5851 | 0.02 | -0.63 | 1.351912 | -3 | -1.177878 | -3 | 0.18584 | 0.11114 | 0.19301 | 0.12955 | 24.11368 | 0.01 | 0 | Neutral |
| testneg_5852 | -0.03 | 1.03 | 4.283597 | -3 | 0.280808 | -3 | 0.19898 | 0.12788 | 0.81659 | 0.10424 | 18.41137 | 0.09 | 0 | Neutral |
| testneg_5853 | 0 | 0 | 1.467396 | -3 | -0.009815 | -3 | 0.96995 | 0.66487 | 0.85793 | 0.80177 | 10.41667 | 0.091397475 | 0 | Neutral |
| testneg_5854 | 0.14 | -0.37 | 0.547607 | -3 | 0.02094 | -3 | 0.07872 | 0.12788 | 0.12547 | 0.09267 | 13.51351 | 0.01 | 0 | Neutral |
| testneg_5855 | -0.14 | 1.01 | 6.105315 | 0.496298 | 0.68543 | -3 | 0.20219 | 0.10986 | 0.15267 | 0.12788 | 18.90053 | 0.28 | 2 | Neutral |

| | | | | | | | | | | | | | | |
|---|---|---|---|---|---|---|---|---|---|---|---|---|---|---|
| testneg_5856 | 0.14 | -1.01 | 2.672539 | -3 | 5.375154 | -3 | 0.46371 | 0.11637 | 0.19898 | 0.30319 | 9.82424 | 0.516322281 | 61.2 | Disease_causing |
| testneg_5857 | 0 | 0 | 1.577999 | -3 | 3.016075 | -3 | 0.17301 | 0.14089 | 0.10176 | 0.12788 | 13.64754 | 0.040172619 | 0 | Neutral |
| testneg_5858 | 0 | -0.01 | 0.707396 | -3 | 1.493352 | -3 | 0.19898 | 0.26604 | 0.17364 | 0.18111 | 19.44444 | 0.113132149 | 0 | Neutral |
| testneg_5859 | 0.01 | 0.8 | 2.947434 | -3 | 5.105753 | 1.261526 | 0.27915 | 0.27706 | 0.25637 | 0.37408 | 13.20597 | 0.784884296 | 99.8 | Disease_causing |
| testneg_5860 | 0.16 | -1.37 | 3.062901 | -3 | -1.968465 | -3 | 0.03563 | 0.10455 | 0.65048 | 0.12778 | 21.79201 | 0.091 | 0 | Neutral |
| testneg_5861 | 0 | 0 | 2.064931 | 2.064931 | 1.664734 | -3 | 0.30122 | 0.09165 | 0.119 | 0.0978 | 27.06238 | 0.070279639 | 0.2 | Neutral |
| testneg_5862 | -0.06 | -0.18 | 2.299984 | -3 | -1.616402 | -3 | 0.98926 | 0.53296 | 0.07837 | 0.12788 | 16.63344 | 0.050454545 | 0 | Neutral |
| testneg_5863 | 0.05 | -0.81 | 1.379835 | -3 | 2.736132 | -3 | 0.34117 | 0.12167 | 0.1287 | 0.10423 | 12.33727 | 0.03 | 0 | Neutral |
| testneg_5864 | -0.08 | -0.03 | 3.657735 | -3 | 4.928659 | 0.284017 | 0.24528 | 0.37575 | 0.3025 | 0.2499 | 11.41859 | 0.860422727 | 100 | Disease_causing |
| testneg_5865 | 0.18 | -1.05 | 5.456932 | -3 | 2.209838 | -3 | 0.19969 | 0.1163 | 0.11261 | 0.13673 | 22.43494 | 0.277638889 | 1.4 | Neutral |
| testneg_5866 | 0.02 | -0.68 | 4.473451 | -3 | -0.67532 | -3 | 0.37463 | 0.23021 | 0.462 | 0.1289 | 12.58475 | 0.090873379 | 0 | Neutral |
| testneg_5867 | 0 | -0.01 | 1.887787 | -3 | -0.981122 | -3 | 0.51398 | 0.38821 | 0.07604 | 0.08642 | 12.70833 | 0.02 | 0 | Neutral |
| testneg_5868 | 0.03 | 0.1 | 2.193295 | -3 | 6.151341 | -3 | 0.99689 | 0.28104 | 0.47733 | 0.08388 | 10.11654 | 0.216551136 | 0 | Neutral |
| testneg_5869 | 0.14 | -1.01 | 3.070348 | -3 | 0.727619 | -3 | 0.10809 | 0.09057 | 0.57975 | 0.1248 | 28.35558 | 0.040297619 | 0 | Neutral |
| testneg_5870 | 0.14 | -0.37 | 2.767225 | -3 | 1.643466 | -3 | 0.12545 | 0.08963 | 0.19343 | 0.11347 | 27.46106 | 0.02 | 0 | Neutral |
| testneg_5871 | 0 | 0.32 | -0.420859 | -3 | 12.039115 | -3 | 0.0597 | 0.18092 | 0.20011 | 0.27949 | 10 | 0.673092949 | 97.2 | Disease_causing |
| testneg_5872 | 0.02 | 0.45 | 5.903349 | -3 | -0.025439 | -3 | 0.49745 | 0.1259 | 0.21355 | 0.12788 | 24.98564 | 0.16 | 0 | Neutral |
| testneg_5873 | -0.03 | -0.1 | 2.935592 | -3 | 4.507347 | -3 | 0.1909 | 0.14279 | 0.68569 | 0.12463 | 16.07106 | 0.255936508 | 1.6 | Neutral |
| testneg_5874 | 0.07 | 0.38 | -3.472929 | -3 | -0.504246 | -3 | 0.4097 | 0.36733 | 0.35051 | 0.11664 | 10.83333 | 0.060162907 | 0 | Neutral |
| testneg_5875 | 0 | 0 | -0.049164 | -0.049164 | 1.823528 | -3 | 0.12322 | 0.1246 | 0.17016 | 0.12748 | 34.85998 | 0.01 | 0 | Neutral |
| testneg_5876 | 0.07 | 0.38 | 1.407842 | 1.083343 | 3.377364 | -3 | 0.63418 | 0.69224 | 0.82148 | 0.14847 | 10 | 0.333613395 | 4.4 | Neutral |
| testneg_5877 | -0.07 | -0.38 | 2.459203 | -3 | 0.280422 | -3 | 0.23292 | 0.15913 | 0.67301 | 0.15304 | 10.83333 | 0.101766234 | 0 | Neutral |
| testneg_5878 | 0 | 0 | 0.346053 | -3 | 0.953557 | -3 | 0.3047 | 0.11929 | 0.09522 | 0.20323 | 21.95652 | 0.080304667 | 0 | Neutral |
| testneg_5879 | 0 | 0 | 2.397094 | -3 | 5.586363 | -3 | 0.22364 | 0.12788 | 0.65909 | 0.31908 | 16.84579 | 0.359394659 | 12.2 | Neutral |
| testneg_5880 | 0 | 0 | 1.817109 | -3 | -0.119638 | -3 | 0.51622 | 0.11809 | 0.16379 | 0.31462 | 12.30072 | 0.03594427 | 0 | Neutral |
| testneg_5881 | 0.01 | 0.8 | 0.663866 | -3 | -2.863489 | -3 | 0.08283 | 0.11797 | 0.09028 | 0.10643 | 13.51351 | 0.02 | 0 | Neutral |
| testneg_5882 | 0 | 0 | -0.100514 | -3 | 0.573548 | -3 | 0.16469 | 0.16042 | 0.99267 | 0.1069 | 10 | 0.04 | 0 | Neutral |
| testneg_5883 | 0 | 0 | -0.71253 | -3 | 1.613091 | -3 | 0.03563 | 0.10455 | 0.53922 | 0.12788 | 12.29167 | 0.03 | 0 | Neutral |
| testneg_5884 | -0.01 | -0.8 | 2.607371 | -3 | 2.379482 | -3 | 0.53239 | 0.17509 | 0.6854 | 0.12563 | 17.77778 | 0.105 | 0 | Neutral |
| testneg_5885 | 0.19 | -0.27 | -0.330967 | -3 | 0.60982 | 0.60982 | 0.20657 | 0.12788 | 0.23844 | 0.27772 | 7.03611 | 0.09 | 0 | Neutral |
| testneg_5886 | 0.04 | -0.23 | 5.823825 | -3 | 3.730704 | -3 | 0.08531 | 0.12788 | 0.37951 | 0.19435 | 18.88889 | 0.642928298 | 91.8 | Disease_causing |
| testneg_5887 | -0.03 | -1.22 | 3.001125 | -3 | 3.591293 | -3 | 0.12974 | 0.11886 | 0.16409 | 0.12788 | 4.72222 | 0.134666667 | 0 | Neutral |
| testneg_5888 | 0.21 | 0.15 | -1.422138 | -1.685253 | 5.635366 | -3 | 0.16091 | 0.90971 | 0.50403 | 0.21568 | 18.60878 | 0.569490741 | 79 | Disease_causing |
| testneg_5889 | -0.01 | 0.57 | 1.519771 | -3 | -0.674759 | -0.674759 | 0.09356 | 0.2448 | 0.02782 | 0.06218 | 12.70833 | 0.05 | 0 | Neutral |
| testneg_5890 | 0 | -0.01 | 0.306573 | -3 | 3.541975 | -3 | 0.03898 | 0.09943 | 0.11608 | 0.09123 | 12.46232 | 0.06 | 0 | Neutral |
| testneg_5891 | 0 | -0.01 | 0.866411 | -3 | 0.03296 | -3 | 0.13011 | 0.14157 | 0.31824 | 0.6682 | 15.88561 | 0.06011236 | 0 | Neutral |
| testneg_5892 | -0.08 | -0.03 | 1.518871 | -3 | -0.130873 | -3 | 0.52953 | 0.10241 | 0.08631 | 0.10194 | 17.57805 | 0.014592391 | 0 | Neutral |
| testneg_5893 | 0 | 0 | 0.475124 | -3 | -0.298537 | -3 | 0.17677 | 0.67006 | 0.04831 | 0.12788 | 12.70833 | 0.04 | 0 | Neutral |
| testneg_5894 | 0.18 | -1.05 | 3.225783 | -3 | 4.123198 | -3 | 0.75978 | 0.11423 | 0.0925 | 0.08222 | 11.95767 | 0.16 | 0 | Neutral |
| testneg_5895 | 0 | 0 | 0.184064 | -3 | 4.19568 | -3 | 0.04436 | 0.12788 | 0.35513 | 0.21966 | 20.14867 | 0.27066391 | 1.2 | Neutral |
| testneg_5896 | 0.02 | 0.45 | 0.245416 | -3 | 3.114446 | -3 | 0.10004 | 0.12788 | 0.34209 | 0.09515 | 17.33333 | 0.05 | 0 | Neutral |
| testneg_5897 | 0 | 0 | -0.680715 | -3 | 2.75027 | -3 | 0.40264 | 0.12788 | 0.50034 | 0.07527 | 29.1283 | 0.03 | 0 | Neutral |
| testneg_5898 | -0.07 | -0.38 | -1.415783 | -3 | -1.060816 | -3 | 0.13278 | 0.20513 | 0.02658 | 0.18737 | 10.71529 | 0.151601863 | 0 | Neutral |
| testneg_5899 | 0.04 | -0.23 | 2.529555 | -3 | -1.171244 | -1.171244 | 0.11359 | 0.08631 | 0.19898 | 0.12788 | 13.46801 | 0.01 | 0 | Neutral |
| testneg_5900 | 0 | 0 | 1.39998 | -3 | 6.398922 | -3 | 0.161 | 0.11597 | 0.54792 | 0.54626 | 16.42913 | 0.336935131 | 9.6 | Neutral |
| testneg_5901 | 0.03 | 0.12 | 5.580988 | -3 | 1.256981 | -3 | 0.8169 | 0.23301 | 0.15202 | 0.12722 | 34.70126 | 0.27 | 0.8 | Neutral |
| testneg_5902 | 0.04 | -0.23 | 2.777545 | -3 | 1.846205 | -3 | 0.16345 | 0.11716 | 0.23078 | 0.10588 | 10.36687 | 0.01 | 0 | Neutral |
| testneg_5903 | 0 | 0.32 | 3.828757 | -3 | 0.512897 | -3 | 0.19898 | 0.11262 | 0.52771 | 0.10974 | 12.63911 | 0.03 | 0 | Neutral |
| testneg_5904 | 0.1 | 1.9 | 6.154974 | -3 | -0.457394 | -3 | 0.04515 | 0.14318 | 0.19898 | 0.46867 | 18.33333 | 0.476127608 | 45 | Neutral |
| testneg_5905 | -0.21 | -0.15 | 1.538424 | -3 | 0.530386 | -3 | 0.13475 | 0.12788 | 0.26909 | 0.10159 | 43.87313 | 0.05 | 0 | Neutral |
| testneg_5906 | 0.07 | 0.38 | 1.136151 | -3 | 1.782913 | -3 | 0.1119 | 0.24875 | 0.1327 | 0.09885 | 16.57328 | 0.044214286 | 0 | Neutral |
| testneg_5907 | 0.07 | 0.38 | 1.47114 | 0.847539 | 1.120297 | -3 | 0.65564 | 0.11468 | 0.10919 | 0.105 | 18.88889 | 0.04 | 0 | Neutral |
| testneg_5908 | 0 | 0 | 0.910599 | -3 | 1.5447 | -3 | 0.19898 | 0.12788 | 0.08932 | 0.32882 | 17.60101 | 0.06 | 0 | Neutral |
| testneg_5909 | 0 | 0 | -0.633568 | -3 | 1.999648 | 1.999648 | 0.23027 | 0.37171 | 0.12626 | 0.10279 | 15.85874 | 0.07 | 0 | Neutral |
| testneg_5910 | 0.14 | -1.01 | 0.997182 | -3 | 4.791856 | 4.791856 | 0.0887 | 0.10047 | 0.52028 | 0.47239 | 31.73501 | 0.48225 | 48.8 | Neutral |
| testneg_5911 | -0.03 | -0.1 | 0.539897 | -3 | -0.850513 | -3 | 0.09866 | 0.104 | 0.06982 | 0.10226 | 20.59741 | 0.02 | 0 | Neutral |
| testneg_5912 | 0 | 0 | -0.172518 | -3 | 0.266467 | -3 | 0.19452 | 0.79779 | 0.46837 | 0.14896 | 10 | 0.102 | 0 | Neutral |
| testneg_5913 | -0.15 | -0.04 | -0.451132 | -3 | 3.811924 | -3 | 0.12048 | 0.2381 | 0.2218 | 0.11533 | 17.66149 | 0.25 | 0.8 | Neutral |
| testneg_5914 | 0 | -0.32 | 2.97409 | -3 | 5.575747 | -3 | 0.16544 | 0.43246 | 0.54891 | 0.55746 | 11.25 | 0.865454737 | 100 | Disease_causing |
| testneg_5915 | 0.07 | 0.38 | 1.124718 | -3 | 4.172643 | -3 | 0.13356 | 0.09963 | 0.10893 | 0.12072 | 23.81213 | 0.07 | 0 | Neutral |
| testneg_5916 | 0 | 0 | 1.599689 | -3 | 0.919273 | -3 | 0.17057 | 0.12788 | 0.09658 | 0.11286 | 32.29167 | 0.000392308 | 0 | Neutral |
| testneg_5917 | 0.01 | 0.8 | 4.332296 | -3 | 1.036843 | -3 | 0.19564 | 0.59275 | 0.18102 | 0.6537 | 11.25 | 0.383464209 | 16.4 | Neutral |
| testneg_5918 | 0.01485714 | -0.0434286 | 1.535215 | -3 | 2.212176 | -3 | 0.26323 | 0.2399 | 0.63233 | 0.16248 | 19.42161 | 0.196537467 | 0.2 | Neutral |
| testneg_5919 | -0.03 | -0.1 | 7.170492 | -3 | 5.803451 | -3 | 0.42306 | 0.86891 | 0.16692 | 0.09243 | 16.76768 | 0.743333333 | 100 | Disease_causing |
| testneg_5920 | 0.09 | -0.3 | 5.344639 | -3 | 0.292782 | -3 | 0.6992 | 0.1804 | 0.50592 | 0.0952 | 13.48802 | 0.189 | 0 | Neutral |
| testneg_5921 | -0.08 | -0.03 | 3.221915 | -3 | -1.168963 | -3 | 0.19898 | 0.10996 | 0.52768 | 0.12788 | 13.14653 | 0.03 | 0 | Neutral |
| testneg_5922 | 0.09 | -0.3 | 5.408425 | -3 | 5.583219 | -3 | 0.37855 | 0.47307 | 0.13458 | 0.17634 | 10.83333 | 0.952869694 | 100 | Disease_causing |
| testneg_5923 | -0.05 | -0.29 | 0.018835 | -3 | -0.068708 | -3 | 0.19898 | 0.12788 | 0.25077 | 0.06261 | 10.41667 | 0.000400079 | 0 | Neutral |
| testneg_5924 | 0.18 | -1.05 | 5.964672 | -3 | 2.013485 | -3 | 0.51137 | 0.19235 | 0.87803 | 0.12788 | 10 | 0.3225 | 6.2 | Neutral |
| testneg_5925 | -0.07 | -0.38 | 3.002256 | -3 | 0.865094 | -3 | 0.19301 | 0.12955 | 0.05315 | 0.12788 | 30.22182 | 0.035181818 | 0 | Neutral |
| testneg_5926 | -0.03 | -0.1 | 6.058643 | -3 | -0.228545 | -3 | 0.13994 | 0.11181 | 0.12176 | 0.09356 | 16.63344 | 0.16 | 0 | Neutral |
| testneg_5927 | 0 | 0 | -0.843404 | -3 | 2.698195 | -3 | 0.11587 | 0.12001 | 0.08143 | 0.44857 | 9.08597 | 0.111975253 | 0 | Neutral |
| testneg_5928 | 0 | -0.01 | 6.431621 | 6.431621 | 6.090617 | 6.090617 | 0.26676 | 0.76568 | 0.10185 | 0.12788 | 24.17095 | 0.83 | 100 | Disease_causing |
| testneg_5929 | -0.09 | 0.3 | 4.208754 | -3 | 1.643466 | -3 | 0.10719 | 0.08618 | 0.19343 | 0.11347 | 19.09347 | 0.065888278 | 0 | Neutral |
| testneg_5930 | 0 | 0 | 0.438443 | 0.438443 | 3.841536 | -3 | 0.36857 | 0.36135 | 0.08002 | 0.08792 | 20.55556 | 0.126803397 | 0 | Neutral |
| testneg_5931 | 0 | 0 | 0.452231 | -3 | -1.941492 | -3 | 0.46691 | 0.07886 | 0.37711 | 0.12788 | 19.68594 | 0.01 | 0 | Neutral |
| testneg_5932 | 0.07 | 0.33 | 3.446637 | -3 | 3.390375 | -3 | 0.13413 | 0.12788 | 0.25462 | 0.12788 | 17.21847 | 0.110471898 | 0 | Neutral |
| testneg_5933 | -0.03 | 1.03 | 3.953236 | -3 | 3.260847 | -3 | 0.07219 | 0.12788 | 0.27605 | 0.12788 | 26.36285 | 0.185885625 | 0 | Neutral |
| testneg_5934 | -0.17 | 1.22 | -1.282937 | -3 | 5.486648 | -3 | 0.26192 | 0.15452 | 0.16708 | 0.11594 | 11.51465 | 0.382815558 | 11 | Neutral |
| testneg_5935 | 0 | 0 | 1.365392 | -3 | -0.614161 | -3 | 0.72062 | 0.14321 | 0.17796 | 0.09522 | 17.21847 | 0.01 | 0 | Neutral |
| testneg_5936 | 0.09 | -0.3 | 3.49868 | -3 | 2.901134 | -3 | 0.18282 | 0.12022 | 0.82014 | 0.14054 | 10.83333 | 0.220705047 | 0.2 | Neutral |
| testneg_5937 | -0.15 | -0.04 | 0.179678 | -3 | 1.351643 | -3 | 0.31168 | 0.12788 | 0.86306 | 0.10962 | 11.88333 | 0.05 | 0 | Neutral |
| testneg_5938 | -0.01 | -0.8 | 3.446913 | -3 | 0.35929 | -3 | 0.32531 | 0.12351 | 0.19898 | 0.12788 | 22.6792 | 0.02 | 0 | Neutral |
| testneg_5939 | 0 | 0 | 3.170674 | -3 | 4.465847 | -3 | 0.15713 | 0.3507 | 0.34109 | 0.2167 | 19.22908 | 0.771708551 | 100 | Disease_causing |
| testneg_5940 | 0 | 0 | 0.515441 | -3 | -0.022585 | -3 | 0.2741 | 0.10478 | 0.2543 | 0.12929 | 8.97898 | 0 | 0 | Neutral |
| testneg_5941 | -0.09 | 0.67 | 0.398037 | -3 | 2.751749 | -3 | 0.20547 | 0.12178 | 0.19479 | 0.12788 | 19.20428 | 0.050298574 | 0 | Neutral |
| testneg_5942 | 0 | -0.01 | 0.229126 | -3 | 6.179859 | -3 | 0.25561 | 0.11004 | 0.07467 | 0.09224 | 13.51351 | 0.088107143 | 0 | Neutral |
| testneg_5943 | -0.02 | -0.45 | -1.184577 | -3 | 1.613942 | -3 | 0.34826 | 0.49454 | 0.34399 | 0.08193 | 25.11193 | 0.060260417 | 0 | Neutral |
| testneg_5944 | 0.01485714 | -0.0434286 | 1.535215 | -3 | 2.722211 | -0.101413 | 0.26323 | 0.2399 | 0.2306 | 0.0952 | 11.19839 | 0.072021684 | 0 | Neutral |
| testneg_5945 | -0.08 | -0.03 | 2.145761 | -3 | 4.725264 | -3 | 0.53861 | 0.18604 | 0.13869 | 0.10313 | 12.7356 | 0.187820231 | 0 | Neutral |
| testneg_5946 | 0 | 0 | 0.287181 | -3 | -0.873164 | -3.056434 | 0.93834 | 0.11088 | 0.17786 | 0.15314 | 15.60822 | 0.040261207 | 0 | Neutral |
| testneg_5947 | -0.07 | -0.38 | 4.428125 | -3 | 1.932311 | 1.932311 | 0.76693 | 0.19444 | 0.97673 | 0.22976 | 10 | 0.3275 | 4.6 | Neutral |
| testneg_5948 | 0.12 | -1.16 | 1.957496 | -3 | -0.937961 | -3 | 0.06395 | 0.0759 | 0.13523 | 0.12788 | 21.7129 | 0.030138147 | 0 | Neutral |
| testneg_5949 | 0 | 0 | -0.09322 | -3 | 7.831582 | -3 | 0.14341 | 0.10134 | 0.09462 | 0.11807 | 37.04096 | 0.2775 | 2 | Neutral |
| testneg_5950 | 0.15 | 0.06 | 3.876605 | -3 | 2.262993 | -3 | 0.83732 | 0.19325 | 0.0961 | 0.21256 | 27.4024 | 0.386337121 | 10.8 | Neutral |
| testneg_5951 | -0.03 | -0.1 | 2.735353 | -3 | 3.544558 | -3 | 0.29478 | 0.12788 | 0.22896 | 0.24839 | 7.01832 | 0.357888986 | 10.2 | Neutral |
| testneg_5952 | 0 | 0 | 0.992597 | -3 | 0.313674 | -3 | 0.30122 | 0.09165 | 0.26812 | 0.30371 | 11.875 | 0.027099441 | 0 | Neutral |
| testneg_5953 | -0.02 | 0.36 | 3.856697 | -3 | 0.046221 | -3 | 0.07015 | 0.0889 | 0.1856 | 0.12788 | 16.43299 | 0.05 | 0 | Neutral |
| testneg_5954 | 0.02 | 0.45 | 3.769936 | -3 | -0.744113 | -3 | 0.57239 | 0.16622 | 0.12547 | 0.09267 | 35.6845 | 0.07 | 0 | Neutral |
| testneg_5955 | -0.08 | -0.03 | 3.219123 | -3 | 2.804418 | -3 | 0.31924 | 0.11145 | 0.32416 | 0.09959 | 13.02635 | 0.07 | 0 | Neutral |
| testneg_5956 | -0.05 | -0.29 | 0.790139 | -3 | 7.149184 | -3 | 0.26482 | 0.12788 | 0.52278 | 0.41207 | 11.25 | 0.507613782 | 58.2 | Disease_causing |
| testneg_5957 | -0.01 | 0.57 | 3.099279 | -3 | 1.21068 | -3 | 0.46919 | 0.60081 | 0.95247 | 0.11285 | 21.38889 | 0.16 | 0 | Neutral |
| testneg_5958 | -0.07 | -0.01 | 0.237617 | -3 | 3.003095 | -3 | 0.97252 | 0.8859 | 0.09284 | 0.26053 | 10 | 0.264198969 | 0.6 | Neutral |
| testneg_5959 | 0.05 | 0.29 | -0.214978 | -3 | 1.378582 | -3 | 0.39192 | 0.09135 | 0.08184 | 0.10206 | 10.83333 | 0.03 | 0 | Neutral |
| testneg_5960 | -0.06 | -0.18 | 3.18158 | -3 | 1.499967 | 0.732076 | 0.23439 | 0.09253 | 0.25422 | 0.10102 | 26.84685 | 0.04 | 0 | Neutral |
| testneg_5961 | 0 | -0.01 | 1.609618 | -3 | 0.717822 | -3 | 0.67038 | 0.1794 | 0.29044 | 0.12142 | 23.19718 | 0.02 | 0 | Neutral |
| testneg_5962 | 0.16 | 0.08 | -0.169987 | -3 | -0.111883 | -0.111883 | 0.99445 | 0.10396 | 0.06313 | 0.12788 | 13.73522 | 0.053035993 | 0 | Neutral |
| testneg_5963 | -0.04 | 1.61 | 4.174982 | -3 | 1.594121 | -3 | 0.42544 | 0.11809 | 0.09259 | 0.11165 | 16.84579 | 0.1 | 0 | Neutral |
| testneg_5964 | -0.01 | 0.57 | -0.767729 | -3 | -0.170031 | -3 | 0.1289 | 0.14878 | 0.07049 | 0.12788 | 19.71696 | 0.04 | 0 | Neutral |
| testneg_5965 | 0 | 0 | 0.543131 | -3 | 1.996983 | 1.996983 | 0.1331 | 0.12788 | 0.09553 | 0.11637 | 22.17119 | 0.04 | 0 | Neutral |
| testneg_5966 | 0.14 | -0.69 | 2.909681 | -3 | 1.011117 | -3 | 0.62708 | 0.24932 | 0.15524 | 0.10607 | 19.8494 | 0.080086207 | 0 | Neutral |
| testneg_5967 | 0.03 | -1.03 | 2.912037 | -3 | 2.23696 | -3 | 0.27197 | 0.12346 | 0.68865 | 0.19414 | 18.44444 | 0.214857143 | 0.2 | Neutral |
| testneg_5968 | -0.18 | 1.05 | 0.012106 | -3 | 5.467754 | -3 | 0.42755 | 0.09484 | 0.05877 | 0.12788 | 18.88889 | 0.26 | 0 | Neutral |
| testneg_5969 | -0.08 | -0.03 | 5.721411 | -3 | 4.336137 | -3 | 0.19494 | 0.14451 | 0.19898 | 0.12788 | 44.05628 | 0.394915366 | 20.4 | Neutral |
| testneg_5970 | 0.07 | 0.38 | 0.906743 | -3 | -0.075228 | -3 | 0.10349 | 0.12788 | 0.12626 | 0.10279 | 28.08943 | 0.01 | 0 | Neutral |
| testneg_5971 | -0.02 | -0.45 | 2.925716 | -3 | 1.82476 | -3 | 0.13906 | 0.09925 | 0.12253 | 0.10814 | 9.60576 | 0.03 | 0 | Neutral |
| testneg_5972 | 0 | 0 | 1.402999 | -3 | 0.330924 | -3 | 0.84682 | 0.35105 | 0.09176 | 0.07645 | 22.9859 | 0.03 | 0 | Neutral |
| testneg_5973 | 0.05 | -0.81 | 3.455713 | -3 | 1.614723 | 1.614723 | 0.12175 | 0.21177 | 0.74944 | 0.1263 | 5.73099 | 0.14 | 0 | Neutral |
| testneg_5974 | 0.14 | -0.37 | 2.235719 | -3 | 3.477533 | -3 | 0.12511 | 0.11897 | 0.87847 | 0.96194 | 12.70833 | 0.291897436 | 1.6 | Neutral |
| testneg_5975 | -0.11 | 1.95 | 2.355101 | -3 | 1.511036 | -3 | 0.18312 | 0.12788 | 0.19433 | 0.12788 | 13.46801 | 0.09 | 0 | Neutral |
| testneg_5976 | -0.0003846 | 0.02730769 | 2.005247 | -3 | 2.674709 | -3 | 0.37344 | 0.11262 | 0.6185 | 0.09316 | 10.69779 | 0.040066667 | 0 | Neutral |
| testneg_5977 | 0 | -0.01 | 0.494045 | -3 | 5.967322 | -3 | 0.1304 | 0.12788 | 0.25312 | 0.12788 | 19.42906 | 0.088022911 | 0 | Neutral |

| | | | | | | | | | | | | | | |
|---|---|---|---|---|---|---|---|---|---|---|---|---|---|---|
| testneg_5978 | 0 | 0 | -0.43649 | -3 | 0.997782 | -3 | 0.71968 | 0.54266 | 0.48817 | 0.10601 | 9.74183 | 0.010909091 | 0 | Neutral |
| testneg_5979 | 0 | 0 | 1.145768 | -3 | 0.923814 | -3 | 0.22658 | 0.20646 | 0.48627 | 0.09509 | 12.00726 | 0.02 | 0 | Neutral |
| testneg_5980 | 0.01485714 | -0.0434286 | 1.535215 | -3 | -0.392743 | -3 | 0.26323 | 0.2399 | 0.16275 | 0.12788 | 11.96702 | 0.020548048 | 0 | Neutral |
| testneg_5981 | 0.04 | -0.89 | -1.056088 | -3 | 0.447242 | -3 | 0.06781 | 0.09009 | 0.14248 | 0.12788 | 23.68839 | 0.019166667 | 0 | Neutral |
| testneg_5982 | 0 | 0 | -0.039433 | -3 | -2.188525 | -2.188525 | 0.05732 | 0.1098 | 0.07282 | 0.12788 | 44.02731 | 0.03173913 | 0 | Neutral |
| testneg_5983 | 0 | 0 | 5.586363 | -3 | 12.460288 | -3 | 0.65909 | 0.31908 | 0.13559 | 0.1946 | 17.21847 | 0.875181847 | 100 | Disease_causing |
| testneg_5984 | 0.06 | -0.57 | -2.190906 | -3 | 0.17553 | -3 | 0.03137 | 0.12788 | 0.10646 | 0.1069 | 40.06096 | 0.05 | 0 | Neutral |
| testneg_5985 | 0 | 0 | -0.325825 | -3 | 3.072677 | -3 | 0.68918 | 0.16819 | 0.19898 | 0.12788 | 7.06091 | 0.055162202 | 0 | Neutral |
| testneg_5986 | 0.19 | -0.27 | 1.091905 | 1.091905 | 5.933886 | -3 | 0.10058 | 0.13882 | 0.09099 | 0.2051 | 12.99578 | 0.559766234 | 76.6 | Disease_causing |
| testneg_5987 | -0.02 | 0.63 | 3.870997 | -3 | 1.559263 | -3 | 0.11966 | 0.16464 | 0.1662 | 0.16982 | 31.48039 | 0.311595663 | 2.8 | Neutral |
| testneg_5988 | 0 | 0 | 1.73928 | -3 | 0.061667 | -3 | 0.17643 | 0.10767 | 0.13751 | 0.13274 | 15.79871 | 0.040916553 | 0 | Neutral |
| testneg_5989 | -0.01 | 1.44 | 7.230871 | -3 | -1.705636 | -3 | 0.06953 | 0.17439 | 0.11957 | 0.08975 | 12.00335 | 0.39 | 15 | Neutral |
| testneg_5990 | 0 | 0 | 1.384899 | 1.384899 | 0.922605 | -3 | 0.56315 | 0.13761 | 0.2152 | 0.12788 | 10.41667 | 0.03 | 0 | Neutral |
| testneg_5991 | 0.00109091 | -0.0100909 | 1.971045 | -3 | 5.185173 | -3 | 0.14457 | 0.11614 | 0.06118 | 0.08724 | 10.41667 | 0.08 | 0 | Neutral |
| testneg_5992 | -0.05 | -0.29 | 1.979502 | -3 | -0.674759 | -0.674759 | 0.18498 | 0.12788 | 0.02782 | 0.06218 | 19.09347 | 0.01 | 0 | Neutral |
| testneg_5993 | 0 | 0 | 0.305495 | -3 | 1.083169 | -3 | 0.98255 | 0.98129 | 0.26621 | 0.10639 | 5 | 0.06 | 0 | Neutral |
| testneg_5994 | 0.00672727 | 0.03618182 | 1.204688 | -3 | 2.342086 | -3 | 0.71956 | 0.12519 | 0.5901 | 0.19444 | 12.70833 | 0.081011656 | 0 | Neutral |
| testneg_5995 | -0.15 | -0.06 | 6.055954 | -3 | 3.439486 | -3 | 0.11316 | 0.12788 | 0.13468 | 0.09235 | 50.974 | 0.3 | 3 | Neutral |
| testneg_5996 | 0.01 | 0.8 | -0.24364 | -3 | 0.90039 | -3 | 0.07671 | 0.09226 | 0.33332 | 0.12106 | 19.79589 | 0.02 | 0 | Neutral |
| testneg_5997 | -0.16 | 0.37 | 5.840704 | -3 | 2.243237 | -3 | 0.37909 | 0.4568 | 0.43977 | 0.17284 | 11.78188 | 0.757147113 | 100 | Disease_causing |
| testneg_5998 | -0.06 | -0.18 | 2.120248 | 2.120248 | 3.498501 | -3 | 0.91619 | 0.10042 | 0.80314 | 0.42344 | 14.4789 | 0.319689477 | 5.8 | Neutral |
| testneg_5999 | 0.05 | 0.29 | -1.763617 | -3 | 5.536712 | -3 | 0.19433 | 0.12788 | 0.31344 | 0.12788 | 13.51351 | 0.11458134 | 0 | Neutral |
| testneg_6000 | 0.19 | 0.51 | 4.599202 | -3 | 1.292485 | -3 | 0.34295 | 0.12788 | 0.53784 | 0.5574 | 4.72222 | 0.287105808 | 2 | Neutral |
| testneg_6001 | -0.08 | -0.03 | -0.48043 | -1.108616 | 1.983447 | -3 | 0.27843 | 0.54325 | 0.60812 | 0.19286 | 20 | 0.228180147 | 0 | Neutral |
| testneg_6002 | -0.03 | -0.1 | 6.609854 | -3 | 4.904147 | -3 | 0.18096 | 0.12788 | 0.27605 | 0.21595 | 35.60194 | 0.708570704 | 98.6 | Disease_causing |
| testneg_6003 | -0.08 | -0.03 | 5.050499 | -3 | 0.636994 | -3 | 0.12819 | 0.1229 | 0.1601 | 0.12546 | 13.02635 | 0.070502976 | 0 | Neutral |
| testneg_6004 | -0.14 | 1.01 | 1.907441 | 1.907441 | 1.512094 | -3 | 0.20708 | 0.12788 | 0.16636 | 0.12384 | 29.16667 | 0.11 | 0 | Neutral |
| testneg_6005 | 0 | -0.01 | 1.315543 | -3 | 0.39745 | -3 | 0.28358 | 0.38765 | 0.11404 | 0.12788 | 11.96702 | 0.02 | 0 | Neutral |
| testneg_6006 | 0.21 | 0.15 | 4.730594 | -3 | 4.920647 | -3 | 0.15192 | 0.14302 | 0.09353 | 0.12788 | 10.22898 | 0.380662212 | 14.8 | Neutral |
| testneg_6007 | 0 | 0 | 0.943793 | -0.147943 | 3.106226 | -3 | 0.41089 | 0.1897 | 0.99704 | 0.17463 | 5.69444 | 0.220670857 | 0 | Neutral |
| testneg_6008 | 0.17 | 0.38 | 5.248097 | -3 | 0.861365 | -3 | 0.12497 | 0.09805 | 0.37953 | 0.11262 | 27.77778 | 0.08 | 0 | Neutral |
| testneg_6009 | 0.05 | -0.81 | 6.398064 | -3 | 0.855893 | -3 | 0.339 | 0.13169 | 0.26916 | 0.13285 | 14.52295 | 0.288445815 | 4.4 | Neutral |
| testneg_6010 | -0.03 | -0.1 | 0.212913 | -3 | 1.472855 | -3 | 0.11391 | 0.19434 | 0.56554 | 0.76978 | 16.42913 | 0.208972423 | 0 | Neutral |
| testneg_6011 | 0.16 | -1.37 | 2.791385 | -3 | 1.367341 | -3 | 0.59853 | 0.75237 | 0.04694 | 0.44682 | 15.41667 | 0.33 | 3.6 | Neutral |
| testneg_6012 | 0 | 0 | 1.828554 | -3 | 0.852085 | -3 | 0.04208 | 0.12788 | 0.56554 | 0.76978 | 10.41667 | 0.100103395 | 0 | Neutral |
| testneg_6013 | 0 | 0 | 1.132066 | -3 | 6.169752 | -3 | 0.99317 | 0.60884 | 0.08836 | 0.0911 | 11.77817 | 0.16 | 0 | Neutral |
| testneg_6014 | 0.09 | -0.3 | -0.45976 | -3 | 3.794683 | -3 | 0.81899 | 0.12788 | 0.29356 | 0.47581 | 11.25 | 0.286631919 | 0.4 | Neutral |
| testneg_6015 | 0 | 0 | -0.674729 | -3 | 6.047971 | -3 | 0.24765 | 0.32858 | 0.53157 | 0.50154 | 4.44444 | 0.490578853 | 51.2 | Disease_causing |
| testneg_6016 | 0 | 0 | 0.386655 | -3 | 6.831326 | -3 | 0.07671 | 0.09226 | 0.3405 | 0.1816 | 16.7606 | 0.346276587 | 8 | Neutral |
| testneg_6017 | 0 | 0 | 0.505914 | -3 | -0.814992 | -3 | 0.24823 | 0.12788 | 0.12195 | 0.21397 | 15 | 0.04 | 0 | Neutral |
| testneg_6018 | -0.04 | 0.89 | -1.166085 | -3 | 1.239899 | -3 | 0.54015 | 0.13176 | 0.50687 | 0.11283 | 10.83333 | 0.05 | 0 | Neutral |
| testneg_6019 | 0.14 | -1.01 | 0.29463 | 0.29463 | 1.53776 | 1.53776 | 0.18507 | 0.12788 | 0.29455 | 0.12388 | 18.88889 | 0.04 | 0 | Neutral |
| testneg_6020 | -0.09 | 0.25 | 2.611714 | -3 | 0.29779 | -3 | 0.99543 | 0.12788 | 0.19898 | 0.20319 | 23.32524 | 0.119732587 | 0 | Neutral |
| testneg_6021 | 0 | 0 | 0.280928 | -3 | 4.17674 | -3 | 0.28211 | 0.25332 | 0.0501 | 0.12788 | 21.341 | 0.13075 | 0 | Neutral |
| testneg_6022 | 0.03 | 0.1 | -0.187467 | -0.187467 | -0.59601 | -3 | 0.12314 | 0.11709 | 0.09569 | 0.09246 | 30.90278 | 0.010215677 | 0 | Neutral |
| testneg_6023 | 0.21 | 0.15 | 1.478329 | -3 | 1.371909 | -3 | 0.27614 | 0.12085 | 0.47733 | 0.08388 | 16.79041 | 0.02019638 | 0 | Neutral |
| testneg_6024 | -0.21 | -0.15 | 2.657036 | -3 | 2.886048 | -3 | 0.12408 | 0.08828 | 0.62029 | 0.09082 | 10.83333 | 0.13 | 0 | Neutral |
| testneg_6025 | 0 | -0.01 | 0.639661 | -3 | 1.599689 | -3 | 0.16263 | 0.53493 | 0.17057 | 0.12788 | 24.13242 | 0.050633404 | 0 | Neutral |
| testneg_6026 | 0 | -0.01 | 0.492211 | -3 | -0.100475 | -3 | 0.20673 | 0.13236 | 0.11261 | 0.13673 | 14.29287 | 0.060583333 | 0 | Neutral |
| testneg_6027 | -0.15 | -0.06 | -1.278679 | -3 | 0.584422 | -3 | 0.04574 | 0.05824 | 0.17458 | 0.12788 | 51.91993 | 0.1 | 0 | Neutral |
| testneg_6028 | 0 | -0.01 | 3.021892 | -3 | 1.348057 | -3 | 0.1163 | 0.20863 | 0.9807 | 0.2062 | 10.41667 | 0.244475952 | 0.2 | Neutral |
| testneg_6029 | -0.04 | 0.23 | -2.407172 | -3 | 0.851258 | -0.771274 | 0.1621 | 0.1641 | 0.09306 | 0.12788 | 18.33333 | 0.05 | 0 | Neutral |
| testneg_6030 | 0 | 0 | 0.094005 | -3 | 2.750883 | -3 | 0.64302 | 0.11262 | 0.19898 | 0.12788 | 17.1915 | 0.021055556 | 0 | Neutral |
| testneg_6031 | -0.07 | -0.38 | 2.958244 | -3 | 1.670709 | -3 | 0.93834 | 0.11088 | 0.29247 | 0.20085 | 16.88725 | 0.160411299 | 0 | Neutral |
| testneg_6032 | 0 | 0 | -0.726772 | -3 | 1.07442 | -3 | 0.1072 | 0.63446 | 0.25422 | 0.10102 | 26.84685 | 0.070126782 | 0 | Neutral |
| testneg_6033 | -0.09 | 0.25 | 4.068872 | -3 | 2.495963 | -3 | 0.19898 | 0.12788 | 0.84365 | 0.12788 | 5.74956 | 0.137597928 | 0 | Neutral |
| testneg_6034 | 0 | 0 | 1.362024 | -3 | 0.956394 | -3 | 0.28416 | 0.27805 | 0.20765 | 0.27124 | 7.01832 | 0.062020606 | 0 | Neutral |
| testneg_6035 | -0.03 | -0.1 | 3.350172 | -3 | 6.709862 | -3 | 0.13331 | 0.13638 | 0.30585 | 0.10254 | 20.1182 | 0.379285714 | 14.8 | Neutral |
| testneg_6036 | -0.17 | -0.38 | -1.100648 | -3 | 0.915604 | -3 | 0.12957 | 0.12788 | 0.16542 | 0.54222 | 10.23502 | 0.12 | 0 | Neutral |
| testneg_6037 | 0 | -0.01 | 0.960254 | -3 | 2.449752 | -3 | 0.1602 | 0.11262 | 0.06736 | 0.06344 | 27.77086 | 0.010232684 | 0 | Neutral |
| testneg_6038 | -0.15 | -0.04 | 4.218719 | -3 | 0.190884 | -3 | 0.17258 | 0.12788 | 0.07617 | 0.07182 | 16.96229 | 0.06 | 0 | Neutral |
| testneg_6039 | -0.02 | 0.63 | 2.341961 | -3 | 2.882534 | -3 | 0.22364 | 0.12788 | 0.23385 | 0.12788 | 8.97898 | 0.05 | 0 | Neutral |
| testneg_6040 | 0 | 0 | 1.988879 | -3 | 1.504538 | -3 | 0.46723 | 0.08832 | 0.19898 | 0.12788 | 12.85159 | 0.01 | 0 | Neutral |
| testneg_6041 | -0.07 | -0.7 | 0.232874 | -3 | 5.228469 | -3 | 0.14901 | 0.12788 | 0.24287 | 0.11626 | 19.44444 | 0.081715686 | 0 | Neutral |
| testneg_6042 | 0 | -0.32 | 1.235229 | -3 | 1.65442 | -3 | 0.99948 | 0.5595 | 0.10244 | 0.10107 | 10.41667 | 0.070589744 | 0 | Neutral |
| testneg_6043 | 0 | 0.32 | -1.878052 | -3 | 1.068896 | -3 | 0.21122 | 0.12788 | 0.25448 | 0.15814 | 11.875 | 0.05 | 0 | Neutral |
| testneg_6044 | 0.02 | -0.28 | 0.865525 | -3 | 0.34432 | -3 | 0.37953 | 0.11262 | 0.52957 | 0.18687 | 12.46106 | 0.021883013 | 0 | Neutral |
| testneg_6045 | -0.03 | -0.12 | 0.38962 | -3 | 4.597558 | -3 | 0.49745 | 0.1103 | 0.0532 | 0.07746 | 10.22898 | 0.08 | 0 | Neutral |
| testneg_6046 | -0.04 | 0.23 | 0.98325 | -3 | 2.63235 | -3 | 0.16054 | 0.11683 | 0.17046 | 0.21736 | 11.75869 | 0.120047619 | 0 | Neutral |
| testneg_6047 | 0 | 0 | 0.488267 | -3 | -0.342754 | -3 | 0.1516 | 0.10704 | 0.15125 | 0.08976 | 16.48772 | 0 | 0 | Neutral |
| testneg_6048 | 0.02 | -0.36 | 1.715391 | -3 | 4.649263 | -3 | 0.60959 | 0.76078 | 0.62381 | 0.12788 | 9.862 | 0.212401002 | 0 | Neutral |
| testneg_6049 | 0.02 | -0.36 | -0.01145 | -0.01145 | 2.231851 | -3 | 0.78462 | 0.12258 | 0.13521 | 0.10126 | 11.25 | 0.020455487 | 0 | Neutral |
| testneg_6050 | 0 | 0 | 0.101787 | -3 | -3.590514 | -3 | 0.13782 | 0.13794 | 0.47322 | 0.11118 | 18.3313 | 0.01 | 0 | Neutral |
| testneg_6051 | 0.02 | -0.36 | 2.305765 | -3 | 0.90039 | -3 | 0.11055 | 0.1127 | 0.33332 | 0.12106 | 10.74877 | 0.01 | 0 | Neutral |
| testneg_6052 | -0.15 | 1.06 | 0.996068 | -3 | 2.537219 | -3 | 0.30085 | 0.18374 | 0.0328 | 0.0691 | 16.42913 | 0.160098039 | 0 | Neutral |
| testneg_6053 | 0.08 | 0.03 | -2.273827 | -2.273827 | 2.300987 | -3 | 0.12839 | 0.09503 | 0.24931 | 0.78312 | 10.41667 | 0.1384938 | 0 | Neutral |
| testneg_6054 | 0.15 | -1.06 | 4.0374 | -3 | 3.59862 | -3 | 0.19898 | 0.12788 | 0.2016 | 0.13956 | 29.54801 | 0.346491758 | 8.4 | Neutral |
| testneg_6055 | 0.21 | 0.15 | 0.770454 | -3 | 2.859591 | -3 | 0.04208 | 0.12788 | 0.1576 | 0.12788 | 10.83333 | 0.1 | 0 | Neutral |
| testneg_6056 | -0.01 | -0.79 | 4.197517 | -3 | 2.623579 | -3 | 0.16091 | 0.90971 | 0.59812 | 0.19387 | 14.23074 | 0.617476564 | 89.4 | Disease_causing |
| testneg_6057 | 0.07 | 0.38 | 0.034534 | 0.034534 | 3.329851 | -3 | 0.62381 | 0.9477 | 0.5184 | 0.11889 | 11.875 | 0.202184517 | 0 | Neutral |
| testneg_6058 | 0 | 0 | 0.691862 | -3 | 5.724953 | -3 | 0.7219 | 0.41509 | 0.09515 | 0.07784 | 22.4337 | 0.138083592 | 0 | Neutral |
| testneg_6059 | 0 | 0 | 1.144493 | -3 | 6.221201 | -3 | 0.13794 | 0.40936 | 0.22582 | 0.12788 | 19.09347 | 0.246839744 | 0.8 | Neutral |
| testneg_6060 | 0 | 0 | 0.347529 | -3 | 1.685192 | -3 | 0.22443 | 0.10657 | 0.07211 | 0.12934 | 16.0119 | 0 | 0 | Neutral |
| testneg_6061 | -0.08 | -0.03 | -0.125177 | -3 | -0.490791 | -3 | 0.27308 | 0.12788 | 0.06953 | 0.17439 | 14.22149 | 0.075555556 | 0 | Neutral |
| testneg_6062 | 0 | -0.32 | 2.219571 | 1.179541 | 2.93409 | -3 | 0.25827 | 0.21386 | 0.09531 | 0.11449 | 25.79439 | 0.101155081 | 0 | Neutral |
| testneg_6063 | -0.07 | -0.33 | -0.046982 | -3 | 3.206486 | -3 | 0.17643 | 0.10767 | 0.77364 | 0.12676 | 17.77778 | 0.05 | 0 | Neutral |
| testneg_6064 | 0 | 0 | 0.31374 | -3 | -1.141522 | -3 | 0.12582 | 0.12788 | 0.09555 | 0.09639 | 19.09347 | 0.00035401 | 0 | Neutral |
| testneg_6065 | -0.07 | -0.33 | 1.458742 | -3 | 2.189352 | -3 | 0.40557 | 0.11865 | 0.56824 | 0.10646 | 18.88889 | 0.01 | 0 | Neutral |
| testneg_6066 | -0.03 | 1.03 | -1.941492 | -3 | -0.78092 | -3 | 0.37711 | 0.12788 | 0.14432 | 0.22579 | 16.89313 | 0.071746635 | 0 | Neutral |
| testneg_6067 | -0.02 | 0.63 | 2.928857 | 2.928857 | 2.888148 | -3 | 0.11534 | 0.12281 | 0.34324 | 0.085 | 19.13908 | 0.37 | 10.2 | Neutral |
| testneg_6068 | 0.17 | 0.38 | -0.584017 | -3 | 3.503331 | -3 | 0.10244 | 0.10107 | 0.11773 | 0.48758 | 12.70833 | 0.289578375 | 1.4 | Neutral |
| testneg_6069 | 0 | 0 | 5.430083 | -3 | 0.742024 | -3 | 0.79983 | 0.1105 | 0.67301 | 0.15304 | 17.99534 | 0.26 | 0.8 | Neutral |
| testneg_6070 | 0.05 | 0.29 | 1.340083 | -3 | 6.490439 | -3 | 0.28227 | 0.10211 | 0.02498 | 0.08483 | 18.86179 | 0.157675439 | 0 | Neutral |
| testneg_6071 | 0 | 0 | 0.901141 | -3 | 5.849437 | -3 | 0.16475 | 0.26069 | 0.11972 | 0.09438 | 16.89745 | 0.18 | 0 | Neutral |
| testneg_6072 | 0 | 0 | -0.350457 | -3 | 3.629264 | -3 | 0.16139 | 0.09116 | 0.9449 | 0.9378 | 12.70833 | 0.180464542 | 0 | Neutral |
| testneg_6073 | 0.1 | 1.9 | 4.95479 | -3 | 3.009853 | -3 | 0.12077 | 0.11244 | 0.50741 | 0.17915 | 35.38774 | 0.434142857 | 29.6 | Neutral |
| testneg_6074 | 0.07 | 0.33 | 3.241723 | -3 | 4.841698 | -3 | 0.61343 | 0.11838 | 0.29356 | 0.46298 | 14.60213 | 0.49493394 | 52.4 | Disease_causing |
| testneg_6075 | 0.04 | -0.23 | 7.27275 | -3 | 1.926886 | -3 | 0.38445 | 0.58278 | 0.18431 | 0.13863 | 10.72072 | 0.65505246 | 95.6 | Disease_causing |
| testneg_6076 | 0 | 0 | -0.364413 | -3 | -2.003913 | -3 | 0.16965 | 0.25991 | 0.19898 | 0.09822 | 18.70407 | 0.011654002 | 0 | Neutral |
| testneg_6077 | 0 | 0 | -0.272804 | -0.272804 | 3.566467 | -3 | 0.17839 | 0.12009 | 0.17548 | 0.11384 | 13.02635 | 0.033981823 | 0 | Neutral |
| testneg_6078 | 0 | -0.01 | 0.07066 | -3 | -0.086787 | -3 | 0.08903 | 0.12788 | 0.09281 | 0.13929 | 18.20887 | 0.050627451 | 0 | Neutral |
| testneg_6079 | 0 | 0 | 0.386087 | 0.386087 | 5.289856 | -3 | 0.26927 | 0.10752 | 0.06955 | 0.1748 | 12.28162 | 0.33838724 | 5 | Neutral |
| testneg_6080 | 0 | 0 | 2.934031 | -3 | 3.896305 | -3 | 0.32733 | 0.16869 | 0.82595 | 0.52905 | 10.83333 | 0.526493813 | 63.6 | Disease_causing |
| testneg_6081 | 0.21 | 0.15 | 6.593669 | -3 | 0.252727 | -3 | 0.65012 | 0.21208 | 0.13985 | 0.08349 | 6.64849 | 0.271287879 | 1.2 | Neutral |
| testneg_6082 | 0 | 0 | 1.886855 | -3 | 1.139144 | -3 | 0.32201 | 0.10713 | 0.37711 | 0.12788 | 16.8018 | 0 | 0 | Neutral |
| testneg_6083 | 0.21 | 0.15 | 4.124651 | -3 | 3.978022 | -3 | 0.08393 | 0.1123 | 0.27409 | 0.10785 | 41.64764 | 0.157146624 | 0 | Neutral |
| testneg_6084 | 0.22 | -2.66 | 7.635846 | -3 | 4.448407 | -3 | 0.19898 | 0.16603 | 0.19898 | 0.12788 | 18.33333 | 0.67 | 95 | Disease_causing |
| testneg_6085 | 0.21 | 0.15 | 0.837901 | -1.401819 | 1.744576 | -3 | 0.19898 | 0.12788 | 0.22278 | 0.85665 | 22.20266 | 0.143979167 | 0 | Neutral |
| testneg_6086 | 0.07 | -0.31 | 1.843162 | -3 | 0.882369 | -3 | 0.30869 | 0.11948 | 0.30474 | 0.12788 | 19.09347 | 0 | 0 | Neutral |
| testneg_6087 | -0.08 | -0.03 | -0.573221 | -3 | -1.462302 | -3 | 0.20604 | 0.10821 | 0.03811 | 0.05629 | 51.93219 | 0.03 | 0 | Neutral |
| testneg_6088 | 0.08 | 0.03 | -1.290562 | -3 | -0.179197 | -3 | 0.19898 | 0.12895 | 0.86835 | 0.1513 | 8.97898 | 0.045311383 | 0 | Neutral |
| testneg_6089 | -0.05 | -0.29 | 0.116519 | 0.116519 | 4.038181 | 4.038181 | 0.2207 | 0.09998 | 0.13815 | 0.40001 | 18.33333 | 0.481015773 | 47.2 | Neutral |
| testneg_6090 | -0.04 | -2.01 | 3.277724 | -3 | 1.987251 | -4.39493 | 0.10438 | 0.20558 | 0.05183 | 0.18779 | 28.98934 | 0.388433566 | 15.2 | Neutral |
| testneg_6091 | 0 | 0 | 1.630014 | -3 | 2.124586 | -3 | 0.1109 | 0.17295 | 0.53292 | 0.32174 | 14.81287 | 0.142246951 | 0 | Neutral |
| testneg_6092 | 0.09 | -0.3 | 4.733195 | -3 | -1.230463 | -3 | 0.161 | 0.11597 | 0.09964 | 0.10384 | 13.51351 | 0.030833333 | 0 | Neutral |
| testneg_6093 | -0.07 | -0.38 | 1.957086 | -3 | 2.846252 | -3 | 0.2698 | 0.18815 | 0.04912 | 0.08574 | 29.16667 | 0.07 | 0 | Neutral |
| testneg_6094 | 0 | 0 | 0.443289 | -3 | 0.144628 | -3 | 0.09972 | 0.13992 | 0.36417 | 0.21506 | 7.07344 | 0.032141851 | 0 | Neutral |
| testneg_6095 | 0.14 | -0.37 | 1.599249 | -3 | 0.266442 | -3 | 0.08714 | 0.08945 | 0.16791 | 0.22163 | 19.44444 | 0.070322581 | 0 | Neutral |
| testneg_6096 | -0.02 | 0.63 | 1.852975 | -3 | 4.982429 | -3 | 0.18224 | 0.10419 | 0.93064 | 0.10933 | 10.41667 | 0.13 | 0 | Neutral |
| testneg_6097 | 0.02 | 0.45 | 1.848663 | -3 | 1.543726 | -3 | 0.09058 | 0.50106 | 0.47188 | 0.10037 | 18.33333 | 0.08 | 0 | Neutral |
| testneg_6098 | -0.07 | -0.01 | 0.87263 | -3 | 2.566155 | -3 | 0.22258 | 0.10471 | 0.17796 | 0.09522 | 10.83333 | 0.02050817 | 0 | Neutral |
| testneg_6099 | -0.02 | 0.68 | 1.225058 | -3 | 1.258927 | -3 | 0.30209 | 0.14229 | 0.08903 | 0.12788 | 36.47496 | 0.03 | 0 | Neutral |

| | | | | | | | | | | | | | | |
|---|---|---|---|---|---|---|---|---|---|---|---|---|---|---|
| testneg_6100 | -0.14 | 0.69 | 2.798182 | 2.798182 | 3.299396 | -3 | 0.31043 | 0.09688 | 0.12547 | 0.09267 | 17.21847 | 0.35 | 8.6 | Neutral |
| testneg_6101 | 0.01 | 0.8 | 1.392976 | -1.013219 | 1.955212 | -3 | 0.0684 | 0.08803 | 0.29804 | 0.12768 | 25.77678 | 0.02 | 0 | Neutral |
| testneg_6102 | 0 | 0 | 2.408752 | -3 | 5.145096 | -3 | 0.96879 | 0.38765 | 0.48936 | 0.10963 | 8.97898 | 0.138856209 | 0 | Neutral |
| testneg_6103 | 0.2 | -1.26 | 2.073808 | -3 | 4.109699 | -3 | 0.08683 | 0.26961 | 0.19898 | 0.12788 | 19.82106 | 0.286382576 | 0.6 | Neutral |
| testneg_6104 | 0.02 | -0.63 | 5.718017 | -3 | -0.707182 | -3 | 0.11351 | 0.10667 | 0.06789 | 0.09707 | 14.16667 | 0.11 | 0 | Neutral |
| testneg_6105 | -0.04 | 0.23 | -1.151035 | -1.151035 | 3.006809 | -3 | 0.36225 | 0.1054 | 0.84365 | 0.12788 | 16.88659 | 0.06 | 0 | Neutral |
| testneg_6106 | 0 | 0 | 1.73928 | -3 | -1.232987 | -3 | 0.17643 | 0.10767 | 0.10362 | 0.12788 | 22.77778 | 0.01 | 0 | Neutral |
| testneg_6107 | 0.02 | 0.13 | 4.128054 | -3 | 5.39414 | -3 | 0.18498 | 0.12788 | 0.50034 | 0.07527 | 22.85021 | 0.2 | 0.2 | Neutral |
| testneg_6108 | 0 | 0 | 1.537848 | -3 | 4.447913 | -3 | 0.11391 | 0.12788 | 0.46339 | 0.10423 | 12.7356 | 0.041908331 | 0 | Neutral |
| testneg_6109 | 0 | 0 | -0.10912 | -3 | -0.354552 | -3 | 0.85984 | 0.40044 | 0.09815 | 0.12788 | 17.18075 | 0.021 | 0 | Neutral |
| testneg_6110 | -0.18 | -0.16 | 3.803915 | -3 | 0.405636 | -3 | 0.31773 | 0.10816 | 0.65952 | 0.49708 | 13.51598 | 0.180888889 | 0 | Neutral |
| testneg_6111 | 0 | 0 | 0.028094 | -3 | 0.425705 | -3 | 0.46712 | 0.93559 | 0.19898 | 0.12788 | 18.05084 | 0.080161086 | 0 | Neutral |
| testneg_6112 | 0.03 | -1.03 | 2.597966 | -3 | 0.334858 | -3 | 0.19898 | 0.12788 | 0.21348 | 0.10616 | 11.25 | 0.02 | 0 | Neutral |
| testneg_6113 | -0.21 | -0.15 | 2.711046 | -3 | 0.881846 | -3 | 0.12951 | 0.08688 | 0.07887 | 0.07487 | 20.98388 | 0.06 | 0 | Neutral |
| testneg_6114 | -0.07 | -0.38 | -1.968112 | -3 | 0.83713 | -3 | 0.08779 | 0.1777 | 0.06027 | 0.11574 | 5.22072 | 0.051222222 | 0 | Neutral |
| testneg_6115 | -0.07 | 0.31 | -0.329074 | -3 | 3.01188 | -3 | 0.04916 | 0.14944 | 0.13331 | 0.09729 | 12.47902 | 0.15 | 0 | Neutral |
| testneg_6116 | 0 | 0 | 0.339095 | -3 | -0.01248 | -3 | 0.71463 | 0.29732 | 0.61231 | 0.13362 | 21.43378 | 0.100216636 | 0 | Neutral |
| testneg_6117 | 0.09 | -0.3 | 2.216525 | -3 | 2.272008 | -3 | 0.29417 | 0.10965 | 0.23913 | 0.12788 | 13.51351 | 0.011090614 | 0 | Neutral |
| testneg_6118 | 0.01 | 0.8 | 1.582141 | -3 | 2.70354 | -3 | 0.34117 | 0.12167 | 0.46735 | 0.10674 | 24.26446 | 0.04 | 0 | Neutral |
| testneg_6119 | 0.14 | -0.37 | 5.734512 | -3 | -2.997709 | -3 | 0.18498 | 0.12788 | 0.04484 | 0.95426 | 16.8018 | 0.37952381 | 13.8 | Neutral |
| testneg_6120 | -0.07 | -0.38 | 0.81956 | -3 | -0.267626 | -3 | 0.50741 | 0.17915 | 0.19898 | 0.12788 | 21.84836 | 0.03012043 | 0 | Neutral |
| testneg_6121 | 0 | 0 | -0.064878 | -3 | 1.846321 | -3 | 0.99668 | 0.30319 | 0.23088 | 0.12788 | 12.46937 | 0.04035119 | 0 | Neutral |
| testneg_6122 | -0.14 | 1.85 | -0.580234 | -3 | 4.365992 | -3 | 0.19024 | 0.1132 | 0.35984 | 0.11129 | 19.45582 | 0.21 | 0 | Neutral |
| testneg_6123 | 0.09 | -0.3 | 3.48051 | -3 | -0.614161 | -3 | 0.99859 | 0.12788 | 0.17796 | 0.09522 | 10 | 0.08 | 0 | Neutral |
| testneg_6124 | 0 | 0 | 2.987273 | -3 | 3.751371 | 1.541201 | 0.03502 | 0.08119 | 0.4069 | 0.3841 | 12.29167 | 0.435477151 | 29 | Neutral |
| testneg_6125 | 0.03 | 0.12 | 3.492738 | -3 | -0.603184 | -3 | 0.46058 | 0.12903 | 0.09176 | 0.07645 | 25.55238 | 0.04 | 0 | Neutral |
| testneg_6126 | 0 | -0.32 | 1.824862 | -3 | -1.73577 | -3 | 0.44816 | 0.56655 | 0.0931 | 0.12788 | 16.14678 | 0.05 | 0 | Neutral |
| testneg_6127 | -0.08 | -0.03 | 1.259325 | -3 | -1.628421 | -3 | 0.16394 | 0.08512 | 0.54441 | 0.08891 | 12.29167 | 0.02 | 0 | Neutral |
| testneg_6128 | 0 | 0 | -0.583572 | -0.583572 | 1.690602 | -3 | 0.04555 | 0.06327 | 0.18206 | 0.10727 | 13.51351 | 0.03 | 0 | Neutral |
| testneg_6129 | 0 | -0.01 | 1.132093 | -3 | 1.826458 | -3 | 0.26372 | 0.12788 | 0.46401 | 0.24608 | 17.57511 | 0.027225393 | 0 | Neutral |
| testneg_6130 | 0.14 | -0.37 | 7.312169 | -3 | 0.38972 | -3 | 0.17597 | 0.12788 | 0.06526 | 0.12788 | 27.51545 | 0.24 | 1.4 | Neutral |
| testneg_6131 | 0.14 | -0.37 | 0.198379 | -3 | 5.308145 | 3.113154 | 0.03991 | 0.08342 | 0.10244 | 0.10107 | 10.41667 | 0.220491071 | 0 | Neutral |
| testneg_6132 | 0 | 0 | 0.205888 | -3 | 1.204698 | 0.807195 | 0.08922 | 0.10539 | 0.6513 | 0.15254 | 8.44735 | 0.035 | 0 | Neutral |
| testneg_6133 | 0.21 | 0.15 | 1.651539 | -3 | 0.148971 | -3 | 0.06218 | 0.10661 | 0.17007 | 0.12771 | 36.52074 | 0.038095238 | 0 | Neutral |
| testneg_6134 | 0 | 0 | -0.22828 | -3 | 0.026216 | -3 | 0.31694 | 0.13475 | 0.94674 | 0.12413 | 10.41667 | 0.010198039 | 0 | Neutral |
| testneg_6135 | 0.14 | -0.69 | 0.136371 | -3 | 0.084906 | -3 | 0.10895 | 0.10442 | 0.44329 | 0.12788 | 4.72222 | 0.010258564 | 0 | Neutral |
| testneg_6136 | -0.02 | -0.45 | 0.081996 | -3 | 4.623974 | -3 | 0.06526 | 0.12788 | 0.09866 | 0.104 | 32.0207 | 0.09 | 0 | Neutral |
| testneg_6137 | 0.08 | 0.03 | -1.360462 | -3 | 4.129028 | -3 | 0.28537 | 0.12788 | 0.03505 | 0.0907 | 13.02635 | 0.09 | 0 | Neutral |
| testneg_6138 | 0 | 0 | 0.287181 | -3 | 8.031124 | -3 | 0.93834 | 0.11088 | 0.18842 | 0.12378 | 10.31323 | 0.25 | 1 | Neutral |
| testneg_6139 | -0.05 | -0.29 | -3.374696 | -3 | 3.273498 | -3 | 0.14785 | 0.09084 | 0.13433 | 0.09578 | 26.53988 | 0.060199966 | 0 | Neutral |
| testneg_6140 | -0.05 | -0.29 | -1.851788 | -3 | 4.836338 | 1.030888 | 0.78741 | 0.22085 | 0.2439 | 0.08848 | 7.72754 | 0.163095238 | 0 | Neutral |
| testneg_6141 | 0 | 0 | 2.047211 | -3 | 3.284967 | -3 | 0.16823 | 0.10796 | 0.64914 | 0.12481 | 10.41667 | 0.031952381 | 0 | Neutral |
| testneg_6142 | 0 | 0 | 1.495199 | -3 | 1.140918 | -3 | 0.33049 | 0.12788 | 0.1846 | 0.08682 | 24.58419 | 0.000392308 | 0 | Neutral |
| testneg_6143 | 0.03 | -1.03 | 2.763749 | -3 | 1.26259 | 1.26259 | 0.13203 | 0.12092 | 0.44722 | 0.11641 | 13.84202 | 0.03 | 0 | Neutral |
| testneg_6144 | 0.02 | -0.63 | -2.323867 | -3 | 0.694077 | -3 | 0.07595 | 0.11822 | 0.21235 | 0.1449 | 32.81921 | 0.051037848 | 0 | Neutral |
| testneg_6145 | 0 | 0 | 1.825331 | -3 | 3.907702 | -3 | 0.13029 | 0.12788 | 0.14498 | 0.33896 | 5.69444 | 0.275623733 | 1.6 | Neutral |
| testneg_6146 | 0.01 | 0.79 | 2.67742 | -3 | 1.023002 | -3 | 0.39553 | 0.11991 | 0.09602 | 0.10726 | 13.16701 | 0.02 | 0 | Neutral |
| testneg_6147 | 0.15 | -0.82 | 0.925856 | -3 | 0.172526 | -3 | 0.08628 | 0.11498 | 0.22162 | 0.11997 | 18.45692 | 0.01 | 0 | Neutral |
| testneg_6148 | 0.08 | 0.03 | -1.907668 | -3 | 0.583408 | 0.583408 | 0.1273 | 0.12788 | 0.05386 | 0.09802 | 11.25 | 0.034166667 | 0 | Neutral |
| testneg_6149 | 0.01 | 0.8 | -1.354962 | -3 | 2.494251 | -3 | 0.18243 | 0.11102 | 0.10314 | 0.08894 | 18.88889 | 0.04 | 0 | Neutral |
| testneg_6150 | 0 | 0 | 0.974808 | 0.974808 | 1.474779 | 1.474779 | 0.32992 | 0.33227 | 0.0882 | 0.12788 | 20 | 0.0655 | 0 | Neutral |
| testneg_6151 | 0 | 0 | 0.902021 | 0.902021 | 4.962346 | -3 | 0.37499 | 0.17803 | 0.20047 | 0.11827 | 16.76768 | 0.148583333 | 0 | Neutral |
| testneg_6152 | 0.03 | 0.1 | 5.424976 | -3 | -1.96598 | -3 | 0.14956 | 0.10585 | 0.08476 | 0.12788 | 14.64539 | 0.114280303 | 0 | Neutral |
| testneg_6153 | -0.000303 | -0.0069697 | 0.145016 | -3 | 3.810168 | -3 | 0.42621 | 0.49093 | 0.16002 | 0.14644 | 4.44444 | 0.322169063 | 5.8 | Neutral |
| testneg_6154 | 0 | 0 | 1.259391 | -3 | 0.776242 | -3 | 0.27528 | 0.12788 | 0.03646 | 0.09863 | 20.02958 | 0.01 | 0 | Neutral |
| testneg_6155 | 0 | -0.01 | 0.907032 | -3 | -0.200482 | -3 | 0.1205 | 0.23508 | 0.3889 | 0.10831 | 10.08138 | 0.02 | 0 | Neutral |
| testneg_6156 | 0.04 | -1.36 | 2.710461 | 1.695441 | 0.79796 | -3 | 0.15202 | 0.12722 | 0.57466 | 0.12788 | 34.70126 | 0.09 | 0 | Neutral |
| testneg_6157 | -0.08 | -0.03 | 0.306778 | -3 | 3.533603 | -3 | 0.83955 | 0.10333 | 0.98641 | 0.83983 | 9.84165 | 0.229851804 | 0 | Neutral |
| testneg_6158 | 0 | -0.01 | 0.632891 | -3 | 1.419411 | -3 | 0.28856 | 0.1544 | 0.05564 | 0.06933 | 31.44909 | 0.02015 | 0 | Neutral |
| testneg_6159 | 0 | 0 | -0.549073 | -3 | -0.573377 | -3 | 0.38016 | 0.14503 | 0.42366 | 0.14372 | 18.88889 | 0.043355133 | 0 | Neutral |
| testneg_6160 | 0 | 0 | -0.681556 | -3 | 0.896883 | -3 | 0.42585 | 0.18081 | 0.02124 | 0.05186 | 11.9764 | 0.02 | 0 | Neutral |
| testneg_6161 | 0 | 0 | 0.159372 | 0.159372 | 1.288259 | -3 | 0.05228 | 0.12788 | 0.16286 | 0.15786 | 11.23851 | 0.120982073 | 0 | Neutral |
| testneg_6162 | -0.08 | -0.03 | 1.345409 | -3 | 4.33297 | -3 | 0.17973 | 0.12278 | 0.06331 | 0.2396 | 7.59298 | 0.310884485 | 4.2 | Neutral |
| testneg_6163 | 0 | -0.01 | 0.522695 | -3 | 0.79796 | -3 | 0.84616 | 0.1471 | 0.57466 | 0.12788 | 17.33333 | 0.01 | 0 | Neutral |
| testneg_6164 | 0 | 0 | 1.519976 | -3 | 1.318968 | -3 | 0.19898 | 0.12788 | 0.18438 | 0.13272 | 10.83333 | 0.05 | 0 | Neutral |
| testneg_6165 | 0.05 | -0.81 | 3.423319 | -3 | -1.543536 | -1.543536 | 0.16053 | 0.15049 | 0.06744 | 0.12788 | 30.30219 | 0.060954545 | 0 | Neutral |
| testneg_6166 | 0.07 | 0.38 | 2.257725 | -3 | 0.934348 | -3 | 0.47888 | 0.43756 | 0.32677 | 0.12788 | 12.6233 | 0.051240836 | 0 | Neutral |
| testneg_6167 | 0.07 | 0.7 | -0.59601 | -3 | -1.653555 | -1.653555 | 0.09569 | 0.09246 | 0.14588 | 0.12156 | 14.68921 | 0.02 | 0 | Neutral |
| testneg_6168 | -0.17 | -0.38 | 1.89302 | -3 | 2.393948 | 2.393948 | 0.16475 | 0.26069 | 0.70819 | 0.5975 | 14.01961 | 0.420261905 | 24.4 | Neutral |
| testneg_6169 | 0 | 0 | 2.575019 | -3 | 0.623338 | 0.623338 | 0.28269 | 0.12505 | 0.09707 | 0.12788 | 30.29057 | 0.018214286 | 0 | Neutral |
| testneg_6170 | 0.07 | 0.33 | 3.81823 | -3 | 1.426181 | -3 | 0.1755 | 0.10455 | 0.93001 | 0.11682 | 10.10684 | 0.05 | 0 | Neutral |
| testneg_6171 | -0.12 | 1.14 | 2.652229 | -3 | 3.398658 | -3 | 0.04393 | 0.12788 | 0.06971 | 0.09918 | 10.02635 | 0.209285714 | 0 | Neutral |
| testneg_6172 | 0.11 | -1.95 | 3.776207 | -3 | 0.101783 | -3 | 0.18765 | 0.27794 | 0.06196 | 0.10881 | 9.10647 | 0.09486236 | 0 | Neutral |
| testneg_6173 | 0.22 | -2.66 | 3.810217 | -3 | 1.562114 | -3 | 0.10562 | 0.09329 | 0.78356 | 0.13377 | 29.00252 | 0.17 | 0 | Neutral |
| testneg_6174 | 0 | 0 | -0.0353 | -3 | 6.931183 | -3 | 0.98339 | 0.18221 | 0.32423 | 0.08104 | 14.38073 | 0.239615385 | 0.8 | Neutral |
| testneg_6175 | -0.21 | -0.15 | -2.747914 | -3 | 0.588209 | -3 | 0.03651 | 0.06141 | 0.07431 | 0.11513 | 18.6768 | 0.1075 | 0 | Neutral |
| testneg_6176 | -0.06 | 0.57 | 1.042833 | -3 | 0.373592 | -3 | 0.19898 | 0.12788 | 0.02723 | 0.04872 | 18.88889 | 0.010909091 | 0 | Neutral |
| testneg_6177 | 0 | 0 | 0.362957 | -3 | -1.057731 | -1.057731 | 0.59342 | 0.12788 | 0.22901 | 0.12788 | 11.92982 | 0.01 | 0 | Neutral |
| testneg_6178 | -0.21 | -0.15 | 2.241257 | -3 | 5.375154 | -3 | 0.07014 | 0.53339 | 0.19898 | 0.30319 | 16.8018 | 0.777633929 | 100 | Disease_causing |
| testneg_6179 | -0.08 | -0.03 | 1.45727 | -3 | 4.661827 | -3 | 0.06526 | 0.12788 | 0.13023 | 0.29692 | 18.76159 | 0.369701004 | 10.8 | Neutral |
| testneg_6180 | 0.03072727 | -0.4116364 | 2.905355 | -3 | -0.660504 | -3 | 0.04305 | 0.06354 | 0.19898 | 0.12788 | 13.51351 | 0.051714286 | 0 | Neutral |
| testneg_6181 | 0.05 | 0.29 | 2.036657 | 2.036657 | 5.586363 | -3 | 0.91691 | 0.25567 | 0.65909 | 0.31908 | 10 | 0.544101356 | 69.6 | Disease_causing |
| testneg_6182 | -0.07 | -0.38 | 1.905081 | -3 | -2.173353 | -3 | 0.3038 | 0.11278 | 0.0676 | 0.12788 | 17.86874 | 0.01 | 0 | Neutral |
| testneg_6183 | -0.03 | 1.03 | 0.93405 | -3 | 5.393665 | 5.393665 | 0.05677 | 0.06758 | 0.03809 | 0.16044 | 18.33333 | 0.502833333 | 58.2 | Disease_causing |
| testneg_6184 | -0.05 | -0.29 | 0.346592 | -3 | 2.859591 | -3 | 0.29145 | 0.11884 | 0.1576 | 0.12788 | 32.41109 | 0.02 | 0 | Neutral |
| testneg_6185 | -0.07 | -0.01 | 2.087185 | -3 | -3.291659 | -3 | 0.06392 | 0.12788 | 0.11957 | 0.08975 | 16.93015 | 0.02 | 0 | Neutral |
| testneg_6186 | 0.07 | 0.38 | -0.075755 | -3 | 2.650284 | -3 | 0.19898 | 0.12788 | 0.45628 | 0.15819 | 22.9497 | 0.151519231 | 0 | Neutral |
| testneg_6187 | 0 | 0 | 0.256955 | -3 | 0.675108 | -3 | 0.1846 | 0.08682 | 0.24287 | 0.11626 | 23.13449 | 0.000960784 | 0 | Neutral |
| testneg_6188 | -0.14 | 1.85 | 3.759726 | -3 | 0.197741 | -3 | 0.48906 | 0.11262 | 0.31923 | 0.17492 | 22.93882 | 0.181195286 | 0 | Neutral |
| testneg_6189 | -0.01 | -0.79 | 1.377684 | -3 | 3.17007 | -3 | 0.09406 | 0.19858 | 0.12377 | 0.12788 | 29.63692 | 0.13 | 0 | Neutral |
| testneg_6190 | 0 | 0 | -0.314986 | -3 | 5.931702 | -3 | 0.10646 | 0.1069 | 0.0895 | 0.12788 | 28.016 | 0.103785714 | 0 | Neutral |
| testneg_6191 | 0 | 0 | 0.365268 | -3 | -0.184036 | -3 | 0.53627 | 0.12788 | 0.05999 | 0.12788 | 18.6768 | 0.012012184 | 0 | Neutral |
| testneg_6192 | -0.08 | -0.03 | 2.333739 | -3 | -0.289151 | -3 | 0.11564 | 0.1184 | 0.80314 | 0.42344 | 20 | 0.100253205 | 0 | Neutral |
| testneg_6193 | 0.05 | 0.29 | 2.437307 | -3 | -0.885978 | -3 | 0.3047 | 0.11929 | 0.14007 | 0.10647 | 18.33333 | 0.008365801 | 0 | Neutral |
| testneg_6194 | 0 | 0 | 1.495836 | -3 | 0.439914 | -3 | 0.24977 | 0.11262 | 0.09275 | 0.14911 | 18.72417 | 0.04356866 | 0 | Neutral |
| testneg_6195 | 0.03 | 0.12 | 6.140208 | -3 | 1.88229 | -3 | 0.40179 | 0.12112 | 0.64922 | 0.10176 | 28.27549 | 0.2 | 0.2 | Neutral |
| testneg_6196 | 0.01 | -0.57 | -0.236734 | -3 | -0.284048 | -3 | 0.0832 | 0.09323 | 0.16709 | 0.08659 | 17.21847 | 0.01 | 0 | Neutral |
| testneg_6197 | 0.02 | -0.36 | -0.01145 | -0.01145 | 3.752366 | -3 | 0.78462 | 0.12258 | 0.24906 | 0.11684 | 17.47024 | 0.071533082 | 0 | Neutral |
| testneg_6198 | 0.05 | -0.81 | 1.982936 | -3 | 0.973671 | 0.973671 | 0.07044 | 0.11679 | 0.47246 | 0.47221 | 5 | 0.11 | 0 | Neutral |
| testneg_6199 | -0.04 | -2.01 | 4.558544 | -3 | 4.563044 | -3 | 0.08753 | 0.09666 | 0.01952 | 0.08329 | 17.21847 | 0.191666667 | 0 | Neutral |
| testneg_6200 | -0.03 | -0.1 | 1.40668 | -3 | 0.106535 | -3 | 0.08401 | 0.16123 | 0.44429 | 0.20468 | 10 | 0.140397493 | 0 | Neutral |
| testneg_6201 | 0.04 | -0.23 | 3.962157 | -3 | 0.916182 | -3 | 0.06898 | 0.11659 | 0.65857 | 0.26294 | 53.80034 | 0.22 | 0.4 | Neutral |
| testneg_6202 | -0.04 | -2.01 | 2.571576 | -3 | -0.823621 | -0.823621 | 0.37183 | 0.10564 | 0.27382 | 0.16898 | 13.02635 | 0.117395754 | 0 | Neutral |
| testneg_6203 | 0.04 | -0.23 | 7.050129 | -3 | 1.013474 | -3 | 0.34205 | 0.11195 | 0.19898 | 0.10778 | 22.80197 | 0.239209979 | 1.6 | Neutral |
| testneg_6204 | -0.16 | 0.37 | 1.664239 | -3 | 3.546589 | -3 | 0.08814 | 0.42357 | 0.37109 | 0.12788 | 19.18549 | 0.31 | 1.8 | Neutral |
| testneg_6205 | 0.09 | -0.3 | 6.221418 | -3 | 2.398349 | -3 | 0.21811 | 0.18489 | 0.73985 | 0.13844 | 17.77778 | 0.5175 | 61 | Disease_causing |
| testneg_6206 | 0.07 | -0.31 | 4.644707 | -3 | 0.446029 | -3 | 0.27614 | 0.12085 | 0.38749 | 0.1076 | 18.00699 | 0.02 | 0 | Neutral |
| testneg_6207 | 0.01 | 0.8 | 0.179864 | -3 | -1.808697 | -3 | 0.10472 | 0.10014 | 0.14386 | 0.12788 | 25 | 0.010557276 | 0 | Neutral |
| testneg_6208 | 0.06 | 0.18 | 5.333191 | -3 | 2.638763 | -3 | 0.19898 | 0.12788 | 0.13702 | 0.12788 | 20.87531 | 0.208920455 | 0.6 | Neutral |
| testneg_6209 | 0 | 0 | 1.366001 | -3 | 0.148971 | -3 | 0.04305 | 0.06354 | 0.17007 | 0.12771 | 30.45796 | 0.03 | 0 | Neutral |
| testneg_6210 | 0 | -0.01 | 1.955355 | -3 | 1.738112 | -3 | 0.33795 | 0.17455 | 0.50921 | 0.21466 | 11.25 | 0.071141463 | 0 | Neutral |
| testneg_6211 | -0.07 | 0.31 | 0.335227 | -3 | -0.009422 | -3 | 0.04346 | 0.10785 | 0.16275 | 0.12788 | 10 | 0.044119048 | 0 | Neutral |
| testneg_6212 | 0 | 0 | 0.464444 | -3 | 2.94636 | -3 | 0.07973 | 0.09208 | 0.5958 | 0.0929 | 32.07516 | 0.030162833 | 0 | Neutral |
| testneg_6213 | 0 | 0 | 0.921828 | -3 | -0.025657 | -3.451557 | 0.31646 | 0.09844 | 0.16259 | 0.102 | 18.6768 | 0 | 0 | Neutral |
| testneg_6214 | 0 | 0 | -0.138559 | -3 | 3.925425 | -3 | 0.08684 | 0.08917 | 0.24239 | 0.111 | 12.58766 | 0.040233266 | 0 | Neutral |
| testneg_6215 | 0 | 0 | 0.573225 | -3 | 5.766968 | -3 | 0.73947 | 0.2058 | 0.19898 | 0.10593 | 45.25732 | 0.117486778 | 0 | Neutral |
| testneg_6216 | 0.03 | 1.22 | 0.411241 | -3 | 7.890764 | 0.888376 | 0.19199 | 0.16616 | 0.21908 | 0.12788 | 10 | 0.488560606 | 49.8 | Neutral |
| testneg_6217 | 0 | 0 | 0.864288 | -3 | -1.914326 | -3 | 0.21014 | 0.12378 | 0.2124 | 0.17288 | 19.83528 | 0.042988095 | 0 | Neutral |
| testneg_6218 | 0 | 0 | 1.916985 | -3 | 5.704027 | -3 | 0.84031 | 0.16749 | 0.09504 | 0.38939 | 19.89425 | 0.432118107 | 27.4 | Neutral |
| testneg_6219 | -0.04 | 0.23 | 3.260847 | -3 | 5.276013 | -3 | 0.27605 | 0.12788 | 0.21677 | 0.1681 | 17.20874 | 0.561604143 | 73.4 | Disease_causing |
| testneg_6220 | -0.03 | 1.03 | 0.055698 | -3 | 3.173495 | -3 | 0.92776 | 0.09677 | 0.10337 | 0.12788 | 11.78007 | 0.144926802 | 0 | Neutral |
| testneg_6221 | 0.03 | 0.12 | 2.051799 | -3 | 0.286223 | -3 | 0.14624 | 0.10369 | 0.27629 | 0.11166 | 6.25 | 0.000275875 | 0 | Neutral |

| | | | | | | | | | | | | | | |
|---|---|---|---|---|---|---|---|---|---|---|---|---|---|---|
| testneg_6222 | 0 | 0 | 0.755726 | -3 | 0.793606 | -3 | 0.08454 | 0.22908 | 0.75906 | 0.09702 | 13.94822 | 0.03 | 0 | Neutral |
| testneg_6223 | 0.07 | 0.38 | 0.991286 | -3 | 5.372563 | -3 | 0.2439 | 0.08848 | 0.37909 | 0.4568 | 13.04965 | 0.30175 | 2.6 | Neutral |
| testneg_6224 | 0 | 0 | -0.100586 | -3 | -0.273874 | -3 | 0.14668 | 0.25103 | 0.0501 | 0.12788 | 18.691 | 0.03 | 0 | Neutral |
| testneg_6225 | 0.02 | -0.63 | 1.995777 | -3 | 3.017506 | -3 | 0.20632 | 0.12788 | 0.14862 | 0.12788 | 34.86624 | 0.04 | 0 | Neutral |
| testneg_6226 | 0 | 0 | 0.185804 | -3 | 2.846252 | -3 | 0.08306 | 0.12788 | 0.04912 | 0.08574 | 29.9024 | 0.024935897 | 0 | Neutral |
| testneg_6227 | 0 | 0 | 1.674358 | -3 | 0.531892 | -3 | 0.53654 | 0.22867 | 0.25549 | 0.12378 | 23.96199 | 0.020270563 | 0 | Neutral |
| testneg_6228 | 0 | -0.01 | 0.792294 | -3 | 3.021591 | -3 | 0.24528 | 0.37575 | 0.19898 | 0.12788 | 18.23804 | 0.070330087 | 0 | Neutral |
| testneg_6229 | -0.08 | -0.03 | 0.950149 | -3 | -0.925939 | -3 | 0.31318 | 0.11262 | 0.39192 | 0.09135 | 10.41667 | 0.01 | 0 | Neutral |
| testneg_6230 | 0.07 | 0.01 | -0.201571 | -3 | 0.096543 | -3 | 0.09332 | 0.53331 | 0.28326 | 0.13944 | 10.41667 | 0.147348901 | 0 | Neutral |
| testneg_6231 | 0 | 0 | 1.264197 | -3 | -1.323706 | -3 | 0.37542 | 0.19444 | 0.47887 | 0.09483 | 13.94516 | 0.01 | 0 | Neutral |
| testneg_6232 | 0 | 0 | -1.231395 | -3 | 0.472565 | -3 | 0.46757 | 0.14122 | 0.13185 | 0.51629 | 18.12633 | 0.08 | 0 | Neutral |
| testneg_6233 | 0 | 0 | 1.671898 | -3 | 1.259487 | -3 | 0.36225 | 0.1054 | 0.31648 | 0.12788 | 32.67733 | 0.000138916 | 0 | Neutral |
| testneg_6234 | 0.03 | 0.12 | -0.231636 | -0.231636 | 0.636308 | -3 | 0.99765 | 0.95064 | 0.2559 | 0.10848 | 12.47902 | 0.12 | 0 | Neutral |
| testneg_6235 | 0 | 0 | 0.004869 | -3 | 2.018582 | -3 | 0.16134 | 0.1847 | 0.85759 | 0.1031 | 18.26461 | 0.021002735 | 0 | Neutral |
| testneg_6236 | 0.09 | -0.3 | -1.511687 | -3 | 2.716468 | -3 | 0.13173 | 0.12788 | 0.28066 | 0.18241 | 22.90964 | 0.153561073 | 0 | Neutral |
| testneg_6237 | -0.07 | -0.01 | -0.23732 | -0.23732 | 3.236399 | -3 | 0.07473 | 0.51812 | 0.22953 | 0.0919 | 10.28409 | 0.13866436 | 0 | Neutral |
| testneg_6238 | 0 | -0.01 | 0.93478 | -3 | 1.768293 | -3 | 0.57239 | 0.16622 | 0.15016 | 0.08826 | 31.6886 | 0.021657502 | 0 | Neutral |
| testneg_6239 | -0.15 | -0.04 | 3.431501 | -3 | 1.139144 | -3 | 0.1289 | 0.14878 | 0.37711 | 0.12788 | 19.71696 | 0.12 | 0 | Neutral |
| testneg_6240 | 0.09 | -0.3 | 7.081501 | -3 | 3.668266 | 3.668266 | 0.08067 | 0.34914 | 0.63645 | 0.14596 | 32.29167 | 0.75 | 100 | Disease_causing |
| testneg_6241 | -0.08 | -0.03 | 0.746424 | -3 | 5.788681 | -3 | 0.21014 | 0.12378 | 0.81212 | 0.14915 | 5.27778 | 0.334708402 | 6.6 | Neutral |
| testneg_6242 | 0 | 0 | 0.323458 | -3 | 7.40532 | -3 | 0.96842 | 0.23432 | 0.54891 | 0.55746 | 10 | 0.450606763 | 37.4 | Neutral |
| testneg_6243 | -0.08 | -0.03 | -0.057252 | -3 | -0.671472 | -3 | 0.83176 | 0.23122 | 0.08878 | 0.0974 | 9.61744 | 0.020400641 | 0 | Neutral |
| testneg_6244 | 0 | -0.32 | 3.623859 | -3 | 6.214762 | -3 | 0.32065 | 0.19642 | 0.50592 | 0.0952 | 11.25 | 0.4 | 18.8 | Neutral |
| testneg_6245 | -0.07 | -0.38 | 3.810722 | -3 | 5.295628 | -3 | 0.22162 | 0.09445 | 0.27382 | 0.16898 | 13.51351 | 0.59128876 | 83.4 | Disease_causing |
| testneg_6246 | 0.14 | -1.01 | -0.062457 | -3 | 1.221756 | -3 | 0.14705 | 0.08702 | 0.29268 | 0.16012 | 21.51158 | 0.097032967 | 0 | Neutral |
| testneg_6247 | -0.02 | 0.36 | 3.856697 | -3 | 1.474779 | 1.474779 | 0.07015 | 0.0889 | 0.0882 | 0.12788 | 26.7629 | 0.08 | 0 | Neutral |
| testneg_6248 | -0.08 | -0.03 | -1.275641 | -3 | 1.82476 | -3 | 0.05546 | 0.28027 | 0.12253 | 0.10814 | 10.83333 | 0.122916667 | 0 | Neutral |
| testneg_6249 | 0.05 | 0.29 | 2.698795 | -3 | 3.496827 | -3 | 0.17806 | 0.12788 | 0.22794 | 0.10688 | 23.04692 | 0.067896825 | 0 | Neutral |
| testneg_6250 | 0.14 | -1.01 | 3.043761 | -3 | 0.620815 | -3 | 0.08224 | 0.29179 | 0.51587 | 0.13949 | 10.222 | 0.248240313 | 0.2 | Neutral |
| testneg_6251 | 0 | 0 | 1.591831 | -3 | -0.373191 | -1.411613 | 0.99668 | 0.30319 | 0.13696 | 0.09566 | 20.42484 | 0.05 | 0 | Neutral |
| testneg_6252 | 0 | -0.01 | -0.40139 | -3 | 5.833407 | -3 | 0.19898 | 0.12788 | 0.49644 | 0.10958 | 10.83333 | 0.063402778 | 0 | Neutral |
| testneg_6253 | 0.09 | -0.3 | -0.489139 | -3 | 2.424425 | -3 | 0.09179 | 0.12788 | 0.54211 | 0.12592 | 10.38652 | 0.031066856 | 0 | Neutral |
| testneg_6254 | 0 | 0 | -0.328773 | -3 | 0.358974 | -3 | 0.11797 | 0.12788 | 0.27197 | 0.12346 | 18.88889 | 0.000575808 | 0 | Neutral |
| testneg_6255 | 0.21 | 0.15 | 6.448066 | -3 | 2.067928 | -3 | 0.40707 | 0.37988 | 0.96377 | 0.1167 | 10 | 0.387916667 | 14.6 | Neutral |
| testneg_6256 | 0.21 | 0.15 | 0.243701 | -3 | 7.207787 | -3 | 0.06394 | 0.24464 | 0.15263 | 0.12788 | 4.44444 | 0.48625 | 49.4 | Neutral |
| testneg_6257 | -0.23 | 1.04 | 1.436839 | -3 | 0.3368 | -3 | 0.19898 | 0.12788 | 0.08837 | 0.10939 | 27.14151 | 0.1 | 0 | Neutral |
| testneg_6258 | 0 | 0 | 0.216807 | -3 | 0.463587 | -3 | 0.16475 | 0.26069 | 0.36494 | 0.12763 | 15.83207 | 0.010037037 | 0 | Neutral |
| testneg_6259 | 0 | 0 | 1.012529 | -3 | 0.144446 | -3 | 0.53284 | 0.56595 | 0.20604 | 0.10821 | 14.09241 | 0.020144592 | 0 | Neutral |
| testneg_6260 | 0 | 0 | 0.272039 | -3 | -1.246293 | -3 | 0.09024 | 0.08559 | 0.06072 | 0.09873 | 10 | 0.01 | 0 | Neutral |
| testneg_6261 | 0 | 0 | 0.883743 | 0.883743 | 5.002476 | -3 | 0.4094 | 0.1944 | 0.22162 | 0.09445 | 20.59286 | 0.152680998 | 0 | Neutral |
| testneg_6262 | 0.01527273 | 0.118 | 11.521882 | -3 | 0.954121 | -3 | 0.11096 | 0.10074 | 0.55384 | 0.15273 | 50.69444 | 0.41 | 18.2 | Neutral |
| testneg_6263 | 0.17 | -1.22 | 4.581463 | -3 | -0.285825 | -3 | 0.16741 | 0.10762 | 0.15016 | 0.08826 | 14.19394 | 0.04 | 0 | Neutral |
| testneg_6264 | 0.03 | 1.22 | 0.635023 | -3 | -1.379881 | -3 | 0.12136 | 0.27707 | 0.18106 | 0.12788 | 63.42967 | 0.170172876 | 0 | Neutral |
| testneg_6265 | -0.03 | -1.22 | 3.683701 | -3 | 0.748878 | -3 | 0.61008 | 0.17538 | 0.93064 | 0.10933 | 21.51158 | 0.10225 | 0 | Neutral |
| testneg_6266 | 0 | 0 | 1.173845 | -3 | 0.641522 | -3 | 0.88689 | 0.41514 | 0.37109 | 0.12788 | 16.8018 | 0.010215079 | 0 | Neutral |
| testneg_6267 | 0.02 | 0.45 | 2.195573 | -3 | 3.635653 | -3 | 0.10774 | 0.13117 | 0.11501 | 0.13748 | 11.25 | 0.270806373 | 1 | Neutral |
| testneg_6268 | 0.14 | -0.37 | 2.270125 | -3 | 6.530023 | -3 | 0.54724 | 0.12788 | 0.92237 | 0.11252 | 5.69444 | 0.215666667 | 0.2 | Neutral |
| testneg_6269 | 0 | 0 | 2.047211 | -3 | 4.047317 | -3 | 0.16823 | 0.10796 | 0.90308 | 0.28373 | 12.85263 | 0.217050144 | 0.2 | Neutral |
| testneg_6270 | 0.04 | -0.23 | 1.300582 | -3 | 1.955613 | -3 | 0.12424 | 0.09575 | 0.21894 | 0.09041 | 23.68839 | 0.01 | 0 | Neutral |
| testneg_6271 | 0.05 | -0.81 | 5.210825 | -3 | 1.13018 | -3 | 0.63484 | 0.16728 | 0.50505 | 0.11563 | 21.47432 | 0.18 | 0 | Neutral |
| testneg_6272 | 0.02 | -0.28 | -3.482963 | -3 | 3.199076 | -3 | 0.19898 | 0.12788 | 0.04032 | 0.12788 | 41.36636 | 0.07 | 0 | Neutral |
| testneg_6273 | 0 | 0 | -0.139257 | -3 | 0.549871 | -3 | 0.19979 | 0.09328 | 0.21241 | 0.08684 | 13.51351 | 0.000259784 | 0 | Neutral |
| testneg_6274 | 0 | 0 | 0.028094 | -3 | 1.074985 | -3 | 0.46712 | 0.93559 | 0.74558 | 0.10643 | 11.74208 | 0.07113476 | 0 | Neutral |
| testneg_6275 | 0 | 0 | 1.154268 | 1.154268 | 1.217652 | -3 | 0.23071 | 0.09582 | 0.19898 | 0.12788 | 18.76159 | 0.020091326 | 0 | Neutral |
| testneg_6276 | 0.07 | -0.31 | -1.638874 | -3 | 2.407105 | -3 | 0.22078 | 0.12788 | 0.21095 | 0.10574 | 21.94469 | 0.02 | 0 | Neutral |
| testneg_6277 | -0.07 | -0.33 | 1.125228 | -3 | 1.554546 | -3 | 0.15808 | 0.14548 | 0.17808 | 0.12788 | 10.82179 | 0.02 | 0 | Neutral |
| testneg_6278 | -0.05 | -0.29 | 0.089098 | -3 | 0.276885 | -3 | 0.84318 | 0.11313 | 0.50392 | 0.12788 | 23.65801 | 0.011892857 | 0 | Neutral |
| testneg_6279 | 0 | -0.01 | 1.715756 | -3 | -1.200565 | -3 | 0.32804 | 0.10857 | 0.21289 | 0.1079 | 25.60245 | 0.000158224 | 0 | Neutral |
| testneg_6280 | 0 | 0 | 0.103447 | 0.103447 | -0.483554 | -3 | 0.15813 | 0.61197 | 0.19898 | 0.56316 | 12.7581 | 0.17 | 0 | Neutral |
| testneg_6281 | 0 | 0 | 1.464574 | 1.464574 | 1.15899 | -3 | 0.60603 | 0.12788 | 0.17969 | 0.11473 | 21.07794 | 0.04 | 0 | Neutral |
| testneg_6282 | 0.03 | -1.03 | 2.661704 | -3 | 0.996068 | -3 | 0.23691 | 0.1015 | 0.30085 | 0.18374 | 15 | 0.089011905 | 0 | Neutral |
| testneg_6283 | 0.09 | -0.3 | 6.124886 | -3 | 4.671506 | -3 | 0.42306 | 0.86891 | 0.15856 | 0.10994 | 10.83333 | 0.639988095 | 93.4 | Disease_causing |
| testneg_6284 | 0.03 | 0.1 | 1.701517 | -3 | 0.018114 | -3 | 0.57564 | 0.12788 | 0.21884 | 0.16743 | 10.83333 | 0.05 | 0 | Neutral |
| testneg_6285 | 0.2 | -1.26 | 1.467073 | -3 | 3.422072 | -3 | 0.50562 | 0.2196 | 0.20431 | 0.17304 | 10.22898 | 0.453394241 | 33.2 | Neutral |
| testneg_6286 | -0.02 | 0.68 | 1.8794 | -3 | 5.239596 | -3 | 0.54711 | 0.11809 | 0.16089 | 0.1034 | 14.74945 | 0.132396076 | 0 | Neutral |
| testneg_6287 | 0.03 | 0.12 | 0.235146 | -3 | 2.002183 | 1.6844 | 0.16856 | 0.19221 | 0.97972 | 0.13032 | 14.44779 | 0.080669643 | 0 | Neutral |
| testneg_6288 | 0.19 | -0.27 | 3.891483 | -3 | 2.104576 | -3 | 0.19898 | 0.10591 | 0.32669 | 0.10659 | 11.875 | 0.06 | 0 | Neutral |
| testneg_6289 | 0.07 | 0.38 | 0.160149 | -3 | 6.822565 | 1.801673 | 0.07064 | 0.11699 | 0.41953 | 0.44852 | 29.16667 | 0.453038618 | 35.4 | Neutral |
| testneg_6290 | 0.22 | -2.66 | 7.740203 | -3 | -0.729977 | -0.729977 | 0.21029 | 0.27753 | 0.17974 | 0.12788 | 10.83333 | 0.415 | 20.8 | Neutral |
| testneg_6291 | 0 | 0 | -0.364413 | -3 | 2.050194 | -3 | 0.16965 | 0.25991 | 0.1897 | 0.12459 | 18.1536 | 0.020921128 | 0 | Neutral |
| testneg_6292 | 0 | 0 | 0.062047 | -3 | -0.496432 | -0.496432 | 0.21928 | 0.22085 | 0.06247 | 0.14184 | 10 | 0.09025625 | 0 | Neutral |
| testneg_6293 | 0 | -0.01 | 0.183879 | -3 | 2.36813 | -1.507792 | 0.19898 | 0.12788 | 0.63418 | 0.69224 | 21.89731 | 0.108570991 | 0 | Neutral |
| testneg_6294 | 0.06 | -0.57 | 6.407239 | -3 | 6.433557 | -3 | 0.81838 | 0.09725 | 0.16621 | 0.49845 | 18.88889 | 0.75864899 | 99.6 | Disease_causing |
| testneg_6295 | 0 | 0 | 1.252902 | -3 | 0.768578 | 0.768578 | 0.6957 | 0.38769 | 0.26758 | 0.10473 | 12.06235 | 0.02 | 0 | Neutral |
| testneg_6296 | 0.02945455 | 0.16109091 | 12.063711 | -3 | 3.469114 | -3 | 0.12596 | 0.18924 | 0.5127 | 0.12263 | 23.125 | 0.599658565 | 85 | Disease_causing |
| testneg_6297 | 0 | 0 | 0.246572 | -3 | 3.217486 | 3.217486 | 0.2674 | 0.28725 | 0.74699 | 0.12286 | 5 | 0.172217949 | 0 | Neutral |
| testneg_6298 | 0 | 0 | 0.487926 | -3 | -0.559856 | -3 | 0.39224 | 0.09664 | 0.12642 | 0.11218 | 10.83333 | 0.01 | 0 | Neutral |
| testneg_6299 | -0.07 | -0.38 | 3.513469 | -3 | 2.667578 | -3 | 0.99445 | 0.10396 | 0.15025 | 0.4872 | 20.7079 | 0.340833333 | 4.4 | Neutral |
| testneg_6300 | -0.05 | -0.29 | -1.315358 | -3 | 6.202748 | -3 | 0.58984 | 0.15588 | 0.4172 | 0.11643 | 10.41667 | 0.210972222 | 0.4 | Neutral |
| testneg_6301 | 0 | 0 | 1.683026 | -3 | 2.935592 | -3 | 0.84106 | 0.63405 | 0.1909 | 0.14279 | 13.20261 | 0.183330601 | 0 | Neutral |
| testneg_6302 | 0.07 | -0.31 | 6.226242 | -3 | -0.857046 | -3 | 0.20942 | 0.14229 | 0.27924 | 0.12788 | 22.39325 | 0.199122174 | 0.6 | Neutral |
| testneg_6303 | -0.04 | 0.23 | 2.526107 | -3 | 0.579456 | 0.579456 | 0.64721 | 0.8758 | 0.47678 | 0.38085 | 18.85167 | 0.29 | 0.4 | Neutral |
| testneg_6304 | 0.01300699 | -0.0248951 | 2.462213 | -3 | 1.052948 | -3 | 0.32016 | 0.11638 | 0.13708 | 0.81545 | 24.67415 | 0.102261494 | 0 | Neutral |
| testneg_6305 | 0 | 0 | 1.799258 | -3 | 0.804817 | -3 | 0.204 | 0.15424 | 0.12831 | 0.10983 | 10.22898 | 0.01 | 0 | Neutral |
| testneg_6306 | 0 | 0 | 0.73062 | -3 | 0.781134 | -3 | 0.08975 | 0.07399 | 0.92237 | 0.11252 | 20.27909 | 0.01 | 0 | Neutral |
| testneg_6307 | 0.03 | 1.22 | 2.753966 | -3 | 9.146604 | -3 | 0.1551 | 0.09276 | 0.44923 | 0.12788 | 13.67384 | 0.38 | 12.2 | Neutral |
| testneg_6308 | 0 | 0 | 2.012521 | -3 | 2.429829 | -3 | 0.88689 | 0.41514 | 0.25447 | 0.12788 | 26.36265 | 0.07 | 0 | Neutral |
| testneg_6309 | -0.19 | -0.51 | -0.445993 | -0.445993 | 1.091058 | -3 | 0.18324 | 0.13041 | 0.26816 | 0.13347 | 12.7356 | 0.12 | 0 | Neutral |
| testneg_6310 | -0.02 | 0.68 | 1.730125 | -3 | 1.091905 | -3 | 0.10671 | 0.10867 | 0.10058 | 0.13882 | 18.54167 | 0.07 | 0 | Neutral |
| testneg_6311 | 0 | 0 | 1.482956 | -3 | -0.456898 | -3 | 0.2852 | 0.12788 | 0.06397 | 0.12788 | 22.54725 | 0.01 | 0 | Neutral |
| testneg_6312 | 0.04 | 2.01 | -0.707182 | -3 | 0.116245 | -3 | 0.06789 | 0.09707 | 0.8238 | 0.38836 | 7.01832 | 0.1595 | 0 | Neutral |
| testneg_6313 | 0 | 0 | 2.197785 | -3 | 2.61429 | -3 | 0.39863 | 0.44944 | 0.4719 | 0.33344 | 10 | 0.276955177 | 1.6 | Neutral |
| testneg_6314 | 0.09 | -0.3 | 1.802254 | -3 | 5.673256 | -3 | 0.3793 | 0.24105 | 0.04519 | 0.12788 | 10.41667 | 0.243134921 | 0.6 | Neutral |
| testneg_6315 | 0 | 0 | 5.501633 | -3 | 0.460152 | -3 | 0.12582 | 0.12788 | 0.21033 | 0.35953 | 10.30286 | 0.384957143 | 16.4 | Neutral |
| testneg_6316 | -0.15 | 1.06 | 4.632893 | -3 | 3.040711 | -3 | 0.09997 | 0.1797 | 0.3414 | 0.13575 | 32.90163 | 0.532448413 | 74 | Disease_causing |
| testneg_6317 | -0.1 | -1.9 | 5.783525 | -3 | 1.765133 | -3 | 0.15874 | 0.13137 | 0.11114 | 0.09582 | 11.23846 | 0.2485 | 0.4 | Neutral |
| testneg_6318 | 0 | 0 | -0.445161 | -0.445161 | -1.745 | -3 | 0.09728 | 0.12788 | 0.1743 | 0.12788 | 36.66548 | 0.01 | 0 | Neutral |
| testneg_6319 | 0 | 0 | 4.925918 | -3 | 2.352012 | -3 | 0.33135 | 0.73068 | 0.45354 | 0.09957 | 20.27778 | 0.257970588 | 0.4 | Neutral |
| testneg_6320 | 0 | 0 | 1.807098 | -3 | -1.167722 | -3 | 0.19403 | 0.09278 | 0.84431 | 0.65315 | 10.81604 | 0.040475771 | 0 | Neutral |
| testneg_6321 | -0.07 | -0.01 | 1.008528 | -3 | 1.834793 | -3 | 0.1755 | 0.10455 | 0.15767 | 0.12788 | 13.51351 | 0.00050641 | 0 | Neutral |
| testneg_6322 | -0.14 | 0.37 | 2.640854 | -3 | 0.949523 | 0.949523 | 0.20072 | 0.08606 | 0.10056 | 0.12608 | 20.21801 | 0.05 | 0 | Neutral |
| testneg_6323 | -0.01 | -0.79 | 0.652225 | -3 | 0.135853 | -3 | 0.04694 | 0.44682 | 0.10919 | 0.105 | 21.41309 | 0.14 | 0 | Neutral |
| testneg_6324 | 0 | 0 | 0.725748 | -3 | 2.449334 | -0.062464 | 0.19898 | 0.12788 | 0.22415 | 0.70109 | 13.18841 | 0.104180672 | 0 | Neutral |
| testneg_6325 | -0.07 | -0.38 | 0.37014 | -3 | 1.016479 | 1.016479 | 0.12048 | 0.2381 | 0.12649 | 0.12788 | 4.44444 | 0.051339286 | 0 | Neutral |
| testneg_6326 | -0.02 | 0.36 | 2.994942 | -3 | 0.210925 | -3 | 0.2477 | 0.11928 | 0.46386 | 0.16229 | 10.83333 | 0.147583333 | 0 | Neutral |
| testneg_6327 | 0 | -0.01 | 1.801091 | -3 | 0.6037 | -3 | 0.39459 | 0.12788 | 0.19343 | 0.11347 | 10.41667 | 7.61E-05 | 0 | Neutral |
| testneg_6328 | -0.15 | -0.04 | 2.20379 | -3 | 0.099506 | -3 | 0.60703 | 0.12378 | 0.10869 | 0.1503 | 13.51598 | 0.098327586 | 0 | Neutral |
| testneg_6329 | -0.07 | -0.38 | 6.068773 | -3 | 2.937498 | -3 | 0.08509 | 0.12788 | 0.18305 | 0.12743 | 29.34685 | 0.27 | 1.4 | Neutral |
| testneg_6330 | 0.14 | -0.37 | 4.892854 | -3 | 0.867502 | -3 | 0.126 | 0.14076 | 0.71511 | 0.34803 | 18.88889 | 0.309642857 | 3.4 | Neutral |
| testneg_6331 | 0 | 0 | 0.346053 | -3 | 1.341885 | -3 | 0.3047 | 0.11929 | 0.06134 | 0.08207 | 24.7083 | 0.01 | 0 | Neutral |
| testneg_6332 | 0.03 | 0.12 | 3.066776 | -3 | 0.132481 | -3 | 0.2877 | 0.18291 | 0.33026 | 0.30576 | 10.83333 | 0.26539826 | 0.4 | Neutral |
| testneg_6333 | 0 | 0 | 0.991757 | -3 | 0.085674 | -3 | 0.19896 | 0.11604 | 0.57975 | 0.1248 | 32.74561 | 0.00011236 | 0 | Neutral |
| testneg_6334 | 0.16 | -1.37 | 4.338658 | -3 | 6.975558 | -3 | 0.08268 | 0.09939 | 0.56605 | 0.15893 | 20.55556 | 0.647064637 | 93 | Disease_causing |
| testneg_6335 | 0 | 0 | 0.100154 | -3 | 0.730371 | 0.730371 | 0.21122 | 0.12686 | 0.11042 | 0.09881 | 20.42553 | 0.01 | 0 | Neutral |
| testneg_6336 | 0 | 0 | -0.050489 | -3 | 0.236592 | -3 | 0.13395 | 0.29754 | 0.40234 | 0.10353 | 32.44933 | 0.03 | 0 | Neutral |
| testneg_6337 | 0 | 0 | 2.10021 | -3 | -2.637354 | -3 | 0.15526 | 0.08511 | 0.2698 | 0.18815 | 20 | 0.050757143 | 0 | Neutral |
| testneg_6338 | -0.23 | 1.04 | 5.857535 | -3 | -0.352903 | -3 | 0.17532 | 0.36561 | 0.23078 | 0.10588 | 12.4985 | 0.35066129 | 7.8 | Neutral |
| testneg_6339 | 0 | 0 | -0.359121 | -3 | 0.483862 | 0.483862 | 0.18283 | 0.11376 | 0.21551 | 0.12788 | 11.92982 | 0.01 | 0 | Neutral |
| testneg_6340 | 0.04 | -0.23 | 0.982052 | -3 | 0.410412 | -3 | 0.09211 | 0.08973 | 0.74944 | 0.1263 | 5.71873 | 0.01 | 0 | Neutral |
| testneg_6341 | -0.16 | -0.08 | 6.194889 | -3 | 3.009853 | -3 | 0.13083 | 0.23231 | 0.50741 | 0.17915 | 24.50652 | 0.796315217 | 100 | Disease_causing |
| testneg_6342 | 0 | 0 | 2.148718 | -3 | 5.146888 | -3 | 0.45299 | 0.16306 | 0.25831 | 0.10988 | 12.46689 | 0.133336774 | 0 | Neutral |
| testneg_6343 | -0.05 | -0.29 | 0.537263 | -3 | 4.354155 | -3 | 0.45415 | 0.12971 | 0.10244 | 0.10107 | 10.41667 | 0.091598267 | 0 | Neutral |

| | | | | | | | | | | | | | |
|---|---|---|---|---|---|---|---|---|---|---|---|---|---|
| testneg_6344 | 0.14 | -0.69 | 2.950487 | -3 | -1.122085 | -1.122085 | 0.14495 | 0.37127 | 0.0774 | 0.1702 | 74.40252 | 0.42 | 19 Neutral |
| testneg_6345 | 0 | 0.73 | 5.650938 | -3 | 2.398979 | -3 | 0.16671 | 0.18311 | 0.93246 | 0.12788 | 18.88889 | 0.320676329 | 4.6 Neutral |
| testneg_6346 | -0.19 | -0.51 | 3.124356 | -3 | 13.172408 | -3 | 0.20557 | 0.10131 | 0.13433 | 0.09578 | 33.56834 | 0.41 | 22.8 Neutral |
| testneg_6347 | -0.07 | -0.33 | 0.71235 | -3 | 0.034571 | -3 | 0.15152 | 0.12645 | 0.48561 | 0.09034 | 17.21847 | 0.01 | 0 Neutral |
| testneg_6348 | -0.02 | -0.45 | -0.208207 | -3 | 1.559108 | -3 | 0.07219 | 0.12788 | 0.19898 | 0.12788 | 19.37853 | 0.01 | 0 Neutral |
| testneg_6349 | 0 | 0 | 1.420779 | -3 | -1.73577 | -3 | 0.14386 | 0.12788 | 0.0931 | 0.12788 | 34.69482 | 0.01 | 0 Neutral |
| testneg_6350 | 0.02 | 0.45 | 5.537244 | -3 | 0.643588 | -3 | 0.19898 | 0.09972 | 0.30471 | 0.11609 | 12.67761 | 0.126027778 | 0.2 Neutral |
| testneg_6351 | 0.07 | 0.01 | -0.38551 | -3 | 1.260535 | -0.732125 | 0.16797 | 0.12788 | 0.1072 | 0.63446 | 24.68926 | 0.120391207 | 0 Neutral |
| testneg_6352 | -0.03 | -0.1 | -0.369846 | -3 | 3.41486 | -3 | 0.33467 | 0.12788 | 0.64254 | 0.10729 | 11.0044 | 0.04375 | 0 Neutral |
| testneg_6353 | 0.23 | -1.86 | 6.394805 | -3 | 6.64321 | -3 | 0.10285 | 0.10186 | 0.21654 | 0.24468 | 18.88889 | 0.755893908 | 99.2 Disease_causing |
| testneg_6354 | -0.06 | -0.18 | -0.439044 | -3 | 0.709302 | -3 | 0.3797 | 0.1257 | 0.27585 | 0.20571 | 18.88889 | 0.042 | 0 Neutral |
| testneg_6355 | -0.0190909 | 0.62818182 | 7.974744 | -3 | 0.882388 | -0.691044 | 0.087 | 0.12788 | 0.04655 | 0.10225 | 25.12484 | 0.353863636 | 10.6 Neutral |
| testneg_6356 | 0 | 0 | 0.615335 | -3 | 3.380932 | -3 | 0.70872 | 0.09884 | 0.18559 | 0.09173 | 20.28986 | 0.052960655 | 0 Neutral |
| testneg_6357 | 0 | 0.32 | 3.268811 | -3 | 0.327712 | -3 | 0.17141 | 0.1104 | 0.57517 | 0.10873 | 28.10041 | 0.02 | 0 Neutral |
| testneg_6358 | -0.0138182 | -0.2629091 | 4.820901 | 4.820901 | -0.726772 | -3 | 0.06358 | 0.5364 | 0.1072 | 0.63446 | 18.69495 | 0.82 | 100 Disease_causing |
| testneg_6359 | -0.09 | 0.3 | 0.156321 | -3 | 3.351389 | -3 | 0.21566 | 0.15056 | 0.13749 | 0.17824 | 17.89614 | 0.338132914 | 6.6 Neutral |
| testneg_6360 | -0.17 | -0.38 | 5.84848 | -3 | 1.897319 | -3 | 0.19898 | 0.12788 | 0.39225 | 0.10625 | 10.83333 | 0.259845238 | 0.8 Neutral |
| testneg_6361 | 0.09 | -0.3 | 2.810635 | -3 | 0.441119 | -3 | 0.04484 | 0.95426 | 0.24823 | 0.12788 | 16.8018 | 0.2 | 0 Neutral |
| testneg_6362 | -0.14 | 0.37 | 2.640854 | -3 | 0.276239 | -0.774602 | 0.20072 | 0.08606 | 0.1743 | 0.12788 | 17.77778 | 0.04 | 0 Neutral |
| testneg_6363 | -0.19 | 0.27 | 1.951681 | -3 | 0.8178 | -3 | 0.1875 | 0.10316 | 0.09508 | 0.09834 | 29.86111 | 0.05 | 0 Neutral |
| testneg_6364 | -0.01 | -0.8 | 5.486648 | -3 | 1.427185 | -3 | 0.16708 | 0.11594 | 0.98238 | 0.10873 | 11.25 | 0.15 | 0 Neutral |
| testneg_6365 | -0.04 | -2.01 | 4.558544 | -3 | 1.179788 | -3 | 0.08753 | 0.09666 | 0.0571 | 0.10747 | 13.02635 | 0.09 | 0 Neutral |
| testneg_6366 | 0.02 | -0.63 | 3.39983 | -3 | 2.289294 | -3 | 0.70964 | 0.12663 | 0.65696 | 0.42609 | 18.33333 | 0.294271046 | 2 Neutral |
| testneg_6367 | -0.0036364 | 0.11719697 | 0.781229 | 0.781229 | -0.156467 | -3 | 0.07321 | 0.12788 | 0.462 | 0.1289 | 40.10477 | 0.04 | 0 Neutral |
| testneg_6368 | 0 | 0.32 | 0.150421 | -3 | -0.154606 | -3 | 0.54489 | 0.09921 | 0.12886 | 0.12788 | 28.53049 | 0.03 | 0 Neutral |
| testneg_6369 | 0 | 0 | 0.82819 | 0.82819 | 1.796297 | -3 | 0.19898 | 0.12788 | 0.10566 | 0.12788 | 12.30072 | 0.03 | 0 Neutral |
| testneg_6370 | -0.15 | -0.04 | 1.221649 | -3 | 2.51331 | -3 | 0.03676 | 0.12788 | 0.14601 | 0.12788 | 12.79101 | 0.12 | 0 Neutral |
| testneg_6371 | 0 | 0 | 0.101787 | -3 | 2.69903 | -3 | 0.13782 | 0.13794 | 0.1192 | 0.10391 | 13.40128 | 0.020470751 | 0 Neutral |
| testneg_6372 | -0.23 | 1.04 | 1.436839 | -3 | 0.452231 | -3 | 0.19898 | 0.12788 | 0.46691 | 0.07886 | 25.86756 | 0.11 | 0 Neutral |
| testneg_6373 | -0.06 | -0.18 | 1.248415 | -3 | 3.489496 | 0.268639 | 0.27585 | 0.20571 | 0.69254 | 0.10021 | 21.94444 | 0.136554154 | 0 Neutral |
| testneg_6374 | 0.07 | 0.38 | 4.556342 | -3 | 6.469729 | -3 | 0.21029 | 0.27753 | 0.17296 | 0.09548 | 12.22222 | 0.538785714 | 68 Disease_causing |
| testneg_6375 | 0 | 0 | 1.315096 | -3 | 4.88138 | -3 | 0.26972 | 0.12484 | 0.28537 | 0.12788 | 11.96702 | 0.031085206 | 0 Neutral |
| testneg_6376 | 0 | 0 | 1.626934 | -3 | -1.975003 | -1.975003 | 0.20942 | 0.14229 | 0.08465 | 0.15428 | 12.30981 | 0.064153846 | 0 Neutral |
| testneg_6377 | 0.09 | -0.25 | -0.154606 | -3 | 1.448212 | -3 | 0.12886 | 0.12788 | 0.87803 | 0.12788 | 19.95887 | 0.02 | 0 Neutral |
| testneg_6378 | 0.03 | 0.12 | 2.290907 | -3 | -0.576134 | -3 | 0.30789 | 0.15135 | 0.09058 | 0.50106 | 10.83333 | 0.118491889 | 0 Neutral |
| testneg_6379 | -0.17 | -0.38 | 1.850031 | -3 | 0.973289 | -3 | 0.25904 | 0.16134 | 0.88976 | 0.22648 | 12.29167 | 0.14311711 | 0 Neutral |
| testneg_6380 | 0 | -0.01 | 2.100546 | -3 | 2.74177 | -3 | 0.97504 | 0.18545 | 0.04935 | 0.06599 | 10.41667 | 0.051757617 | 0 Neutral |
| testneg_6381 | -0.08 | -0.03 | 5.179822 | -3 | 3.016831 | -3 | 0.05916 | 0.12788 | 0.09099 | 0.2051 | 16.79207 | 0.548053519 | 72.4 Disease_causing |
| testneg_6382 | 0 | 0 | 0.724291 | -3 | 0.338128 | -3 | 0.05005 | 0.07051 | 0.32558 | 0.32796 | 4.44444 | 0.084616279 | 0 Neutral |
| testneg_6383 | 0 | 0 | 1.26533 | -3 | 1.541854 | -3 | 0.27528 | 0.12788 | 0.1065 | 0.12788 | 17.21847 | 0.00016954 | 0 Neutral |
| testneg_6384 | 0 | -0.01 | 3.451445 | -3 | 3.813566 | -3 | 0.49064 | 0.12294 | 0.43691 | 0.12794 | 19.44444 | 0.139546569 | 0 Neutral |
| testneg_6385 | 0 | -0.32 | -0.748689 | -3 | 4.865743 | -3 | 0.07007 | 0.12788 | 0.54015 | 0.13176 | 19.09347 | 0.2 | 0 Neutral |
| testneg_6386 | 0.05 | 0.29 | -1.171244 | -1.171244 | 6.490439 | -3 | 0.19898 | 0.12788 | 0.02498 | 0.08483 | 14.88684 | 0.17 | 0 Neutral |
| testneg_6387 | 0.00916667 | -0.1625 | 0.801954 | -3 | -0.491928 | -3 | 0.09661 | 0.09981 | 0.1056 | 0.12788 | 19.64993 | 0.01 | 0 Neutral |
| testneg_6388 | -0.09 | 0.3 | 3.133137 | -3 | 3.263002 | -3 | 0.26971 | 0.08509 | 0.33992 | 0.15484 | 35.48374 | 0.298734221 | 2.6 Neutral |
| testneg_6389 | 0 | 0 | -0.68826 | -3 | 0.573777 | -3 | 0.32076 | 0.14229 | 0.82156 | 0.14621 | 5.72485 | 0.02 | 0 Neutral |
| testneg_6390 | -0.03 | 1.03 | 1.209864 | -3 | 1.975456 | -3 | 0.23005 | 0.37216 | 0.86552 | 0.33462 | 11.25 | 0.215761074 | 0 Neutral |
| testneg_6391 | 0.14 | -0.69 | 4.55113 | -3 | -1.259442 | -3 | 0.14462 | 0.11239 | 0.24185 | 0.12788 | 13.75 | 0.03 | 0 Neutral |
| testneg_6392 | 0.05 | 0.29 | 1.91968 | -3 | 0.328203 | -3 | 0.20219 | 0.10986 | 0.3414 | 0.13575 | 28.57511 | 0.05 | 0 Neutral |
| testneg_6393 | 0.01 | 0.79 | 0.585649 | 0.585649 | 1.294343 | -3 | 0.07185 | 0.68732 | 0.78161 | 0.14363 | 18.88889 | 0.234123046 | 0.2 Neutral |
| testneg_6394 | 0.08 | 0.03 | 0.041625 | -3 | 0.567743 | -3 | 0.17226 | 0.13001 | 0.19623 | 0.08363 | 18.72241 | 0.02 | 0 Neutral |
| testneg_6395 | 0 | 0 | 1.305769 | -3 | 0.57285 | -3 | 0.16468 | 0.11275 | 0.20501 | 0.6166 | 23.57794 | 0.050765107 | 0 Neutral |
| testneg_6396 | 0.21 | 0.15 | 4.401318 | -3 | 2.634651 | -3 | 0.1434 | 0.09092 | 0.11115 | 0.09814 | 23.43928 | 0.126666667 | 0 Neutral |
| testneg_6397 | 0 | 0 | 1.907124 | -3 | 2.242726 | -3 | 0.18512 | 0.12788 | 0.07275 | 0.09562 | 13.02635 | 0.01 | 0 Neutral |
| testneg_6398 | 0.15 | -0.28 | 2.534333 | -3 | 1.455667 | -3 | 0.19898 | 0.12788 | 0.14341 | 0.10134 | 22.48467 | 0.01 | 0 Neutral |
| testneg_6399 | -0.09 | 0.3 | 1.539941 | -3 | 5.268788 | -3 | 0.64152 | 0.15239 | 0.12943 | 0.14391 | 22.50958 | 0.423501452 | 23.6 Neutral |
| testneg_6400 | 0.06 | -0.57 | 0.297516 | 0.297516 | 0.413333 | -3 | 0.19294 | 0.10464 | 0.36694 | 0.12788 | 27.67377 | 0.010137089 | 0 Neutral |
| testneg_6401 | 0 | -0.32 | -0.547467 | -3 | 5.068794 | -3 | 0.38762 | 0.12788 | 0.04627 | 0.05223 | 33.95189 | 0.09 | 0 Neutral |
| testneg_6402 | 0.18 | -1.05 | 5.456932 | -3 | 1.705087 | -3 | 0.19969 | 0.1163 | 0.22225 | 0.11186 | 17.77778 | 0.105625 | 0 Neutral |
| testneg_6403 | 0.03 | 0.1 | 6.254981 | -3 | -0.076764 | -3 | 0.95993 | 0.15588 | 0.16388 | 0.14439 | 18.88889 | 0.39 | 16.6 Neutral |
| testneg_6404 | 0.06 | 0.18 | -1.330673 | -1.330673 | 1.336135 | -3 | 0.03137 | 0.12788 | 0.17935 | 0.36953 | 45.55556 | 0.142916667 | 0 Neutral |
| testneg_6405 | 0.21 | 0.15 | 1.704228 | -3 | -0.095822 | -3 | 0.27062 | 0.12378 | 0.19898 | 0.19048 | 24.60526 | 0.090714286 | 0 Neutral |
| testneg_6406 | 0.14 | -0.69 | 3.352085 | -3 | 3.923847 | -3 | 0.19898 | 0.12788 | 0.19898 | 0.12788 | 25.14162 | 0.14140873 | 0 Neutral |
| testneg_6407 | -0.02 | -0.41 | 0.579456 | 0.579456 | 2.374684 | -3 | 0.47678 | 0.38085 | 0.33314 | 0.47999 | 12.73937 | 0.306394781 | 4.6 Neutral |
| testneg_6408 | 0 | 0 | 2.995532 | -3 | -0.290801 | -3 | 0.96879 | 0.38765 | 0.06112 | 0.06901 | 10.41667 | 0.069444444 | 0 Neutral |
| testneg_6409 | -0.02 | 0.63 | 2.115956 | 2.115956 | 3.686515 | -3 | 0.47443 | 0.11062 | 0.77981 | 0.62662 | 10.41667 | 0.365737179 | 11 Neutral |
| testneg_6410 | 0 | 0 | 0.332849 | -3 | 0.740166 | 0.740166 | 0.99595 | 0.23021 | 0.18642 | 0.09748 | 18.88889 | 0.05 | 0 Neutral |
| testneg_6411 | 0.09 | -0.25 | 0.253578 | -3 | 0.368238 | -3 | 0.10496 | 0.21491 | 0.17853 | 0.32085 | 10.41667 | 0.1880625 | 0 Neutral |
| testneg_6412 | 0 | 0 | 1.219225 | -3 | 5.315177 | -3 | 0.29548 | 0.10732 | 0.12091 | 0.13585 | 22.79314 | 0.242427241 | 0.4 Neutral |
| testneg_6413 | 0 | -0.32 | -2.076593 | -3 | 3.944419 | -3 | 0.1276 | 0.30104 | 0.47187 | 0.14227 | 16.32934 | 0.432193094 | 28 Neutral |
| testneg_6414 | 0 | 0 | 0.676994 | -3 | 5.288301 | -3 | 0.09356 | 0.2448 | 0.56817 | 0.10766 | 20.32051 | 0.190069444 | 0 Neutral |
| testneg_6415 | -0.08 | -0.03 | 3.206814 | -3 | 1.716425 | -3 | 0.7022 | 0.35308 | 0.77981 | 0.62662 | 5 | 0.312744451 | 3.8 Neutral |
| testneg_6416 | 0.07 | 0.01 | 1.365447 | -3 | 3.781815 | -3 | 0.07701 | 0.19824 | 0.01704 | 0.05734 | 21.01455 | 0.160821895 | 0 Neutral |
| testneg_6417 | -0.08 | -0.03 | 0.950149 | -3 | 1.509253 | 0.659344 | 0.31318 | 0.11262 | 0.81838 | 0.09725 | 11.12432 | 0.02 | 0 Neutral |
| testneg_6418 | 0.07 | -0.31 | 0.913268 | -3 | 0.667614 | -3 | 0.21265 | 0.1284 | 0.47997 | 0.10506 | 10.83333 | 0.01 | 0 Neutral |
| testneg_6419 | 0.14 | -1.01 | 2.130896 | -3 | 0.075565 | -3 | 0.1254 | 0.08697 | 0.03788 | 0.10996 | 10 | 0.02 | 0 Neutral |
| testneg_6420 | 0 | -0.01 | -0.3595 | -3 | -2.769437 | -3 | 0.19898 | 0.12788 | 0.18882 | 0.14637 | 30.22077 | 0.050083333 | 0 Neutral |
| testneg_6421 | 0.17 | 0.38 | 5.405576 | -3 | 1.504305 | -3 | 0.52765 | 0.20417 | 0.22233 | 0.31758 | 17.8804 | 0.537791667 | 71 Disease_causing |
| testneg_6422 | 0.09 | -2.17 | 3.51087 | -3 | -1.117253 | -1.117253 | 0.46339 | 0.10423 | 0.84682 | 0.35105 | 13.86929 | 0.25 | 0.4 Neutral |
| testneg_6423 | 0.08 | 0.03 | -2.273827 | -2.273827 | -0.076898 | -3 | 0.12839 | 0.09503 | 0.07973 | 0.09208 | 20.67274 | 0.03 | 0 Neutral |
| testneg_6424 | 0 | 0 | 0.741693 | 0.741693 | 0.050569 | -3 | 0.48906 | 0.11262 | 0.16263 | 0.53493 | 31.59206 | 0.131010101 | 0 Neutral |
| testneg_6425 | -0.01 | -0.79 | 1.796129 | -3 | 9.935182 | -3 | 0.16052 | 0.12788 | 0.08393 | 0.1123 | 34.5238 | 0.340602113 | 6 Neutral |
| testneg_6426 | 0 | 0 | 0.29779 | -3 | 1.831606 | -3 | 0.19898 | 0.20319 | 0.25896 | 0.45481 | 29.8569 | 0.140444489 | 0 Neutral |
| testneg_6427 | -0.21 | -0.15 | -2.747914 | -3 | 5.29955 | -3 | 0.03651 | 0.06141 | 0.36833 | 0.10433 | 13.83267 | 0.21 | 0 Neutral |
| testneg_6428 | 0 | 0 | 0.59811 | -3 | 5.716056 | -3 | 0.26971 | 0.08509 | 0.28817 | 0.10531 | 32.25184 | 0.05 | 0 Neutral |
| testneg_6429 | 0.05 | -0.81 | 5.432396 | -3 | 1.140923 | -3 | 0.23991 | 0.1515 | 0.67263 | 0.09781 | 10.41667 | 0.21777436 | 0.2 Neutral |
| testneg_6430 | 0.18 | -1.05 | 6.283203 | -3 | 6.230728 | -3 | 0.17818 | 0.15728 | 0.35499 | 0.12052 | 52.18347 | 0.550760157 | 71.4 Disease_causing |
| testneg_6431 | 0 | 0 | 1.411982 | -3 | 5.632523 | -3 | 0.13255 | 0.0962 | 0.17764 | 0.09681 | 13.74475 | 0.07 | 0 Neutral |
| testneg_6432 | -0.08 | -0.03 | 1.582915 | -3 | 0.580159 | -3 | 0.09493 | 0.104 | 0.35908 | 0.18099 | 30.44402 | 0.074627601 | 0 Neutral |
| testneg_6433 | 0.05 | -0.81 | 4.319181 | -3 | -2.421511 | -3 | 0.29356 | 0.47581 | 0.37464 | 0.12788 | 21.50407 | 0.130086207 | 0 Neutral |
| testneg_6434 | -0.07 | -0.38 | -1.803731 | -3 | -1.022735 | -3 | 0.07963 | 0.23713 | 0.08896 | 0.12788 | 20.83333 | 0.07 | 0 Neutral |
| testneg_6435 | 0.03 | 0.1 | 0.148561 | -3 | -0.059772 | -3 | 0.2559 | 0.10848 | 0.12857 | 0.09376 | 24.60165 | 0.01 | 0 Neutral |
| testneg_6436 | 0 | 0 | 0.666829 | 0.531203 | -0.474484 | -3 | 0.30951 | 0.64738 | 0.45075 | 0.12232 | 10.83333 | 0.0445 | 0 Neutral |
| testneg_6437 | -0.08 | -0.03 | 6.665409 | -3 | 1.944832 | -3 | 0.30684 | 0.12788 | 0.69254 | 0.10021 | 34.75855 | 0.287833333 | 1.6 Neutral |
| testneg_6438 | 0.18 | -1.05 | 6.27298 | -3 | -1.13933 | -3 | 0.1602 | 0.11262 | 0.11703 | 0.09364 | 25.10109 | 0.13 | 0 Neutral |
| testneg_6439 | 0 | 0 | -0.044681 | -3 | 1.791247 | -3 | 0.31584 | 0.11996 | 0.39193 | 0.11262 | 17.77778 | 0.00016954 | 0 Neutral |
| testneg_6440 | -0.1 | -1.9 | 1.551505 | -3 | 0.2237 | -3 | 0.05413 | 0.05691 | 0.17853 | 0.32085 | 10.83333 | 0.170916667 | 0 Neutral |
| testneg_6441 | -0.07 | -0.38 | 4.438262 | -3 | 1.845066 | -3 | 0.43707 | 0.10923 | 0.56154 | 0.0939 | 39.56273 | 0.06 | 0 Neutral |
| testneg_6442 | 0 | 0 | 0.699729 | -0.925939 | 4.714948 | -3 | 0.39192 | 0.09135 | 0.20327 | 0.11174 | 20.208 | 0.049375 | 0 Neutral |
| testneg_6443 | 0 | -0.01 | 0.895061 | -3 | 2.061868 | -3 | 0.5471 | 0.11262 | 0.11006 | 0.20781 | 17.21847 | 0.061004167 | 0 Neutral |
| testneg_6444 | 0 | 0 | -0.32247 | -3 | 0.659218 | -3 | 0.35092 | 0.10596 | 0.44429 | 0.20468 | 10.83333 | 0.02 | 0 Neutral |
| testneg_6445 | -0.07 | -0.38 | 1.04036 | -3 | 2.657613 | -3 | 0.23292 | 0.15913 | 0.19898 | 0.12788 | 18.33333 | 0.051071429 | 0 Neutral |
| testneg_6446 | -0.03 | -0.1 | 4.845098 | -3 | -1.428127 | -3 | 0.09093 | 0.28643 | 0.11564 | 0.1184 | 21.38889 | 0.15 | 0 Neutral |
| testneg_6447 | 0.01 | 0.8 | 1.320426 | -3 | 3.541384 | -3 | 0.11941 | 0.10144 | 0.32188 | 0.08974 | 21.58355 | 0.061924674 | 0 Neutral |
| testneg_6448 | -0.14 | 1.85 | -0.580234 | -3 | 9.170362 | -3 | 0.19024 | 0.1132 | 0.1144 | 0.12788 | 26.29285 | 0.392916667 | 16.8 Neutral |
| testneg_6449 | 0.21 | 0.15 | 1.20537 | -3 | 2.794038 | 2.794038 | 0.08846 | 0.06651 | 0.10977 | 0.31695 | 12.29167 | 0.295645833 | 1.4 Neutral |
| testneg_6450 | 0 | 0 | 5.638394 | -3 | 1.436025 | -3 | 0.08999 | 0.0989 | 0.10026 | 0.12788 | 20 | 0.15 | 0 Neutral |
| testneg_6451 | -0.15 | -0.04 | 5.243457 | -3 | 1.066553 | -3 | 0.18096 | 0.12788 | 0.19898 | 0.12788 | 27.49747 | 0.13 | 0 Neutral |
| testneg_6452 | -0.07 | -0.38 | 0.513578 | -3 | 1.11036 | -3 | 0.19898 | 0.07409 | 0.45399 | 0.10245 | 14.49939 | 0.01 | 0 Neutral |
| testneg_6453 | 0.06 | -0.57 | 0.042224 | -3 | -1.432844 | -3 | 0.08878 | 0.0974 | 0.16286 | 0.15786 | 24.7612 | 0.083152254 | 0 Neutral |
| testneg_6454 | 0 | 0 | 1.45802 | -3 | 2.864641 | -3 | 0.47788 | 0.10817 | 0.21399 | 0.10708 | 17.18434 | 0.02 | 0 Neutral |
| testneg_6455 | 0.16 | -0.37 | 3.256775 | -3 | 1.547491 | -3 | 0.16309 | 0.12788 | 0.07071 | 0.11499 | 32.9221 | 0.03 | 0 Neutral |
| testneg_6456 | 0 | 0 | 0.855485 | -3 | 0.888884 | -3 | 0.20116 | 0.10008 | 0.10277 | 0.12788 | 10.2717 | 0.004166667 | 0 Neutral |
| testneg_6457 | 0.17 | 0.38 | -1.662533 | -1.662533 | 0.729943 | -3 | 0.19898 | 0.12788 | 0.52953 | 0.10241 | 11.83017 | 0.021459216 | 0 Neutral |
| testneg_6458 | 0 | 0 | 1.217986 | -3 | 1.752994 | -3 | 0.19898 | 0.12788 | 0.15767 | 0.12788 | 38.96233 | 8.27E-05 | 0 Neutral |
| testneg_6459 | -0.06 | -0.18 | 1.759866 | -3 | -0.416365 | -3 | 0.7249 | 0.41559 | 0.06793 | 0.12788 | 13.19331 | 0.03 | 0 Neutral |
| testneg_6460 | -0.08 | -0.03 | 1.138753 | -3 | -1.713706 | -3 | 0.94859 | 0.75045 | 0.25957 | 0.06969 | 10 | 0.04 | 0 Neutral |
| testneg_6461 | 0 | 0 | 3.03947 | -3 | 0.744084 | -3 | 0.29804 | 0.12768 | 0.1311 | 0.12667 | 13.02635 | 0.018193548 | 0 Neutral |
| testneg_6462 | 0.21 | 0.15 | 1.704228 | -3 | -0.100872 | -3 | 0.27062 | 0.12378 | 0.11891 | 0.11186 | 17.28802 | 0.020381818 | 0 Neutral |
| testneg_6463 | 0.09 | -0.67 | 4.387696 | -3 | 6.195353 | -3 | 0.11802 | 0.09304 | 0.28532 | 0.10663 | 18.70407 | 0.255576923 | 1.4 Neutral |
| testneg_6464 | 0 | 0 | 1.943537 | 1.943537 | 0.415464 | -3 | 0.19237 | 0.23658 | 0.18671 | 0.12788 | 13.05556 | 0.070268707 | 0 Neutral |
| testneg_6465 | -0.02 | -0.45 | 2.197733 | -3 | 0.881601 | -3 | 0.19969 | 0.1163 | 0.16388 | 0.14439 | 5.69444 | 0.041361139 | 0 Neutral |

| | | | | | | | | | | | | | | |
|---|---|---|---|---|---|---|---|---|---|---|---|---|---|---|
| testneg_6466 | 0.02 | -0.63 | 1.098831 | -3 | 2.981794 | -3 | 0.19898 | 0.12788 | 0.17394 | 0.15823 | 23.46973 | 0.164428024 | 0 | Neutral |
| testneg_6467 | -0.02 | 0.68 | 1.225058 | -3 | 3.533603 | -2.586155 | 0.30209 | 0.14229 | 0.98641 | 0.83983 | 12.40248 | 0.293875 | 1.6 | Neutral |
| testneg_6468 | 0 | 0 | 0.347529 | -3 | -0.289151 | -3 | 0.22443 | 0.10657 | 0.80314 | 0.42344 | 10.83333 | 0.021428571 | 0 | Neutral |
| testneg_6469 | 0.04 | 2.01 | 0.133669 | -3 | 0.641608 | -3 | 0.08353 | 0.21479 | 0.32458 | 0.1016 | 20.31414 | 0.14 | 0 | Neutral |
| testneg_6470 | 0 | 0 | -0.481968 | -3 | 0.347722 | -3 | 0.67331 | 0.10118 | 0.05564 | 0.06933 | 18.88889 | 0.02 | 0 | Neutral |
| testneg_6471 | 0 | 0 | 0.525606 | -3 | 1.249535 | -3 | 0.08401 | 0.16123 | 0.75123 | 0.20667 | 17.77778 | 0.081362332 | 0 | Neutral |
| testneg_6472 | 0.04 | -0.89 | 4.932725 | -3 | 1.38401 | -3 | 0.85075 | 0.16646 | 0.37212 | 0.11323 | 14.18282 | 0.131613191 | 0 | Neutral |
| testneg_6473 | -0.08 | -0.03 | 1.978595 | -3 | 1.507411 | -3 | 0.39305 | 0.18672 | 0.04606 | 0.20522 | 10.41667 | 0.120433373 | 0 | Neutral |
| testneg_6474 | 0.03 | 1.22 | 2.035357 | -3 | 4.376293 | -3 | 0.78537 | 0.33817 | 0.16182 | 0.11544 | 12.69679 | 0.243854384 | 0 | Neutral |
| testneg_6475 | 0.04 | -0.23 | 4.740439 | -3 | 3.099902 | -3 | 0.28156 | 0.11579 | 0.15822 | 0.12788 | 21.31045 | 0.135714286 | 0 | Neutral |
| testneg_6476 | -0.08 | -0.03 | 5.770154 | -3 | 3.883879 | -3 | 0.19883 | 0.09434 | 0.30925 | 0.10335 | 13.14653 | 0.35 | 6.6 | Neutral |
| testneg_6477 | 0 | 0 | 1.165732 | -3 | 0.374591 | -3 | 0.90955 | 0.10096 | 0.19898 | 0.12788 | 24.30556 | 0.01 | 0 | Neutral |
| testneg_6478 | -0.04 | -2.01 | 1.879788 | -3 | 0.002159 | -3 | 0.64254 | 0.10729 | 0.10231 | 0.1 | 6.98099 | 0.05 | 0 | Neutral |
| testneg_6479 | 0 | 0 | 0.93075 | -3 | 3.691952 | -3 | 0.16734 | 0.15176 | 0.04694 | 0.44682 | 15.91846 | 0.309284091 | 5 | Neutral |
| testneg_6480 | -0.0114545 | -0.0138182 | 8.391913 | -3 | 0.814088 | -3 | 0.95076 | 0.17388 | 0.55197 | 0.11305 | 4.72222 | 0.46 | 40 | Neutral |
| testneg_6481 | -0.08 | -0.03 | 3.519231 | -3 | 5.728078 | -3 | 0.33049 | 0.12788 | 0.69426 | 0.13242 | 11.92061 | 0.499590887 | 54.8 | Disease_causing |
| testneg_6482 | -0.1 | -1.9 | 3.35171 | -3 | 1.009156 | -3 | 0.20112 | 0.17491 | 0.19898 | 0.12788 | 17.21847 | 0.09 | 0 | Neutral |
| testneg_6483 | 0.18 | -1.05 | 4.092733 | -3 | 5.007683 | -3 | 0.08283 | 0.11797 | 0.07415 | 0.12788 | 13.51351 | 0.25 | 1.2 | Neutral |
| testneg_6484 | 0 | 0 | 1.156713 | 1.156713 | -1.10605 | -3 | 0.36458 | 0.16397 | 0.23401 | 0.12374 | 10.83333 | 0.032724868 | 0 | Neutral |
| testneg_6485 | 0.18 | -1.05 | 1.097809 | -3 | -0.39694 | -3 | 0.43314 | 0.67252 | 0.21807 | 0.12788 | 19.37055 | 0.11 | 0 | Neutral |
| testneg_6486 | 0.04 | 2.01 | 1.631085 | -3 | 1.122188 | -3 | 0.77364 | 0.12676 | 0.26878 | 0.09179 | 27.09021 | 0.070370879 | 0 | Neutral |
| testneg_6487 | 0 | -0.01 | 0.6576 | -3 | 1.213226 | -3 | 0.12147 | 0.1486 | 0.19898 | 0.12788 | 11.98582 | 0.02 | 0 | Neutral |
| testneg_6488 | 0 | 0 | 1.405787 | -3 | 1.92122 | -3 | 0.33304 | 0.15588 | 0.16728 | 0.11868 | 10 | 0.01 | 0 | Neutral |
| testneg_6489 | -0.03 | 1.03 | 1.508354 | -3 | 0.933869 | 0.933869 | 0.03109 | 0.10705 | 0.9862 | 0.4528 | 9.05265 | 0.14 | 0 | Neutral |
| testneg_6490 | 0 | 0 | 1.840893 | -3 | 1.120342 | -3 | 0.18465 | 0.11262 | 0.54015 | 0.13176 | 12.27191 | 0.030228833 | 0 | Neutral |
| testneg_6491 | 0 | 0 | 1.611577 | -3 | 0.740118 | -3 | 0.04483 | 0.59766 | 0.04988 | 0.25503 | 10.41667 | 0.24336609 | 0 | Neutral |
| testneg_6492 | 0 | 0 | 0.614574 | -3 | 6.066854 | -3 | 0.06526 | 0.12788 | 0.03324 | 0.12788 | 27.3569 | 0.12 | 0 | Neutral |
| testneg_6493 | -0.05 | -0.29 | 3.231366 | 3.231366 | 1.992408 | -3 | 0.37237 | 0.36829 | 0.19898 | 0.25991 | 4.44444 | 0.762007576 | 99.6 | Disease_causing |
| testneg_6494 | 0 | 0 | 0.664346 | -3 | 0.312247 | -3 | 0.22169 | 0.11262 | 0.09306 | 0.12788 | 22.46235 | 0.01 | 0 | Neutral |
| testneg_6495 | 0 | -0.01 | 1.609618 | -3 | 3.381705 | -3 | 0.67038 | 0.1794 | 0.30926 | 0.11955 | 7.06934 | 0.053873667 | 0 | Neutral |
| testneg_6496 | 0 | 0 | 1.531656 | -3 | 1.632135 | -3 | 0.37237 | 0.36829 | 0.57188 | 0.34038 | 10 | 0.100662165 | 0 | Neutral |
| testneg_6497 | -0.08 | -0.03 | -0.867132 | -3 | 0.90777 | -3 | 0.08353 | 0.21479 | 0.0961 | 0.21256 | 16.8018 | 0.18 | 0 | Neutral |
| testneg_6498 | 0.12 | -1.14 | -0.006272 | -0.006272 | 0.96317 | -3 | 0.07335 | 0.12788 | 0.20379 | 0.35591 | 18.62552 | 0.131114286 | 0 | Neutral |
| testneg_6499 | 0.04 | -0.23 | 3.352237 | -3 | 0.269965 | -3 | 0.17672 | 0.14229 | 0.26789 | 0.67432 | 10.83333 | 0.199661499 | 0 | Neutral |
| testneg_6500 | 0 | 0 | 0.121522 | -3 | 0.575728 | -3 | 0.25421 | 0.16235 | 0.32606 | 0.10251 | 29.16667 | 0.02 | 0 | Neutral |
| testneg_6501 | 0 | 0 | -0.279068 | -3 | 2.395534 | -3 | 0.26323 | 0.2399 | 0.13967 | 0.15696 | 21.46862 | 0.227803114 | 0.4 | Neutral |
| testneg_6502 | -0.17 | -0.38 | 0.128896 | 0.128896 | 0.183938 | -3 | 0.67331 | 0.10118 | 0.18882 | 0.14637 | 24.85265 | 0.110374126 | 0 | Neutral |
| testneg_6503 | 0 | 0 | -0.294826 | -3 | 4.941767 | -3 | 0.63818 | 0.12788 | 0.60171 | 0.24685 | 12.76316 | 0.177288281 | 0 | Neutral |
| testneg_6504 | 0.02 | -0.63 | -0.170973 | -3 | 5.568196 | -3 | 0.5127 | 0.12263 | 0.1485 | 0.30134 | 20 | 0.379900259 | 12.8 | Neutral |
| testneg_6505 | 0 | 0 | 1.329031 | -3 | 0.465329 | 0.465329 | 0.22489 | 0.16646 | 0.11131 | 0.68435 | 10.83333 | 0.130211032 | 0 | Neutral |
| testneg_6506 | -0.03 | -0.1 | 2.296124 | -3 | -0.133882 | -3 | 0.22258 | 0.10471 | 0.1056 | 0.12788 | 11.54762 | 0.01 | 0 | Neutral |
| testneg_6507 | 0 | -0.01 | 2.053357 | -3 | 0.199148 | -3 | 0.22761 | 0.14229 | 0.13468 | 0.09235 | 28.38173 | 0.01 | 0 | Neutral |
| testneg_6508 | 0 | 0 | 1.405787 | -3 | -0.849297 | -3 | 0.33304 | 0.15588 | 0.23691 | 0.1015 | 24.44444 | 0.010116673 | 0 | Neutral |
| testneg_6509 | 0 | 0 | 1.324436 | -0.708217 | 3.894086 | -3 | 0.11533 | 0.12788 | 0.9637 | 0.12788 | 10.41667 | 0.08 | 0 | Neutral |
| testneg_6510 | 0.03 | -1.03 | 3.637005 | 3.637005 | 5.88302 | -3 | 0.13884 | 0.12788 | 0.41655 | 0.49485 | 10.83333 | 0.858624424 | 100 | Disease_causing |
| testneg_6511 | -0.03 | -0.1 | 6.609854 | -3 | 0.284017 | -3 | 0.18096 | 0.12788 | 0.3025 | 0.2499 | 36.15749 | 0.448035714 | 35.8 | Neutral |
| testneg_6512 | 0 | 0 | 1.003734 | -3 | 2.296298 | -3 | 0.11325 | 0.17021 | 0.96475 | 0.27545 | 5.44551 | 0.12 | 0 | Neutral |
| testneg_6513 | 0.14 | -1.85 | 3.507668 | -3 | 3.238888 | -3 | 0.88689 | 0.41514 | 0.1909 | 0.14279 | 16.07106 | 0.550654701 | 74 | Disease_causing |
| testneg_6514 | 0 | 0 | 0.039637 | -3 | 0.058546 | -3 | 0.20304 | 0.31462 | 0.18998 | 0.26095 | 13.39912 | 0.090121951 | 0 | Neutral |
| testneg_6515 | 0 | -0.01 | 2.125818 | -3 | 2.729204 | 0.707638 | 0.15713 | 0.3507 | 0.62339 | 0.19592 | 19.22908 | 0.31937063 | 4.4 | Neutral |
| testneg_6516 | 0 | 0 | 0.291054 | -3 | 3.227121 | -3 | 0.28499 | 0.10637 | 0.47462 | 0.10826 | 13.51351 | 0.020205566 | 0 | Neutral |
| testneg_6517 | -0.21 | -0.15 | -0.791573 | -3 | 4.455889 | -3 | 0.12886 | 0.12788 | 0.1589 | 0.10761 | 18.82566 | 0.162097561 | 0 | Neutral |
| testneg_6518 | 0.14 | -0.69 | 2.526774 | -3 | 4.892896 | -3 | 0.16797 | 0.12788 | 0.19898 | 0.0989 | 32.29511 | 0.09 | 0 | Neutral |
| testneg_6519 | 0.14 | -0.37 | 6.03866 | -3 | 2.139704 | -3 | 0.60703 | 0.12378 | 0.43921 | 0.153 | 17.77778 | 0.428325397 | 31.4 | Neutral |
| testneg_6520 | 0 | 0 | 1.80113 | -3 | 0.885869 | 0.885869 | 0.2761 | 0.24559 | 0.12003 | 0.12788 | 40.15723 | 0.04 | 0 | Neutral |
| testneg_6521 | 0 | 0 | 0.573225 | -3 | 1.791247 | -3 | 0.73947 | 0.2058 | 0.39193 | 0.11262 | 30.26287 | 0.03 | 0 | Neutral |
| testneg_6522 | 0 | 0 | 0.046064 | -3 | 0.091226 | 0.091226 | 0.2543 | 0.15588 | 0.10625 | 0.09225 | 12.38369 | 0.03 | 0 | Neutral |
| testneg_6523 | -0.07 | -0.38 | -0.094721 | -3 | 1.03321 | -3 | 0.16778 | 0.82322 | 0.35584 | 0.10874 | 23.24074 | 0.094059524 | 0 | Neutral |
| testneg_6524 | 0.17 | 0.38 | 1.703954 | -3 | 1.418046 | -3 | 0.30823 | 0.17188 | 0.42872 | 0.12788 | 20.32051 | 0.05 | 0 | Neutral |
| testneg_6525 | -0.15 | 0.28 | 3.501223 | -3 | -0.811454 | -0.811454 | 0.46339 | 0.10423 | 0.72912 | 0.18218 | 13.88808 | 0.14 | 0 | Neutral |
| testneg_6526 | 0 | 0 | -0.214627 | -3 | 1.489062 | -3 | 0.98578 | 0.91075 | 0.7823 | 0.5949 | 4.44444 | 0.117911422 | 0 | Neutral |
| testneg_6527 | 0.21 | 0.15 | 2.276796 | -3 | 3.118861 | -3 | 0.06939 | 0.08981 | 0.37275 | 0.16262 | 11.25 | 0.27 | 0 | Neutral |
| testneg_6528 | -0.09 | 0.67 | 0.082232 | -3 | -1.883946 | -3 | 0.08475 | 0.19889 | 0.04046 | 0.06599 | 18.36272 | 0.061221266 | 0 | Neutral |
| testneg_6529 | -0.08 | -0.03 | 1.732154 | -3 | -0.121102 | -0.121102 | 0.95857 | 0.12137 | 0.78935 | 0.17351 | 11.25 | 0.05 | 0 | Neutral |
| testneg_6530 | 0 | 0 | 0.052371 | 0.052371 | 1.619888 | -3 | 0.82107 | 0.27926 | 0.19477 | 0.12788 | 9.93641 | 0.01 | 0 | Neutral |
| testneg_6531 | 0.16 | 0.08 | 4.995915 | -3 | 1.957496 | -3 | 0.76138 | 0.16258 | 0.06395 | 0.0759 | 14.7692 | 0.14 | 0 | Neutral |
| testneg_6532 | 0.03 | 0.12 | 2.204017 | -3 | 4.558319 | -3 | 0.18256 | 0.13499 | 0.19898 | 0.12788 | 22.02988 | 0.118382937 | 0 | Neutral |
| testneg_6533 | -0.04 | 0.23 | 2.344511 | -3 | 2.388689 | -3 | 0.42585 | 0.18081 | 0.18771 | 0.13279 | 5.73099 | 0.222095238 | 0.2 | Neutral |
| testneg_6534 | -0.06 | -0.18 | 0.491194 | 0.491194 | 4.257584 | -3 | 0.72124 | 0.09906 | 0.05005 | 0.07051 | 25.2381 | 0.11 | 0 | Neutral |
| testneg_6535 | -0.02 | 0.68 | 0.385771 | -3 | 8.008012 | -3 | 0.11077 | 0.10076 | 0.89178 | 0.23115 | 10.15099 | 0.42 | 26.6 | Neutral |
| testneg_6536 | 0.18 | -1.05 | 5.931475 | -3 | 0.136092 | -3 | 0.19494 | 0.14451 | 0.16507 | 0.10992 | 20.27778 | 0.158792271 | 0 | Neutral |
| testneg_6537 | 0.03 | -1.03 | 4.386219 | 4.386219 | 6.935441 | -3 | 0.03137 | 0.12788 | 0.10185 | 0.12788 | 65.58036 | 0.61 | 93.2 | Disease_causing |
| testneg_6538 | 0 | 0 | 0.024873 | -3 | 0.892974 | 0.892974 | 0.16507 | 0.10992 | 0.06986 | 0.12788 | 17.45692 | 0.01 | 0 | Neutral |
| testneg_6539 | 0 | 0 | -0.234907 | -3 | 1.687645 | -3 | 0.1947 | 0.11413 | 0.04519 | 0.12788 | 19.44444 | 0.01 | 0 | Neutral |
| testneg_6540 | 0.04 | -0.23 | -1.239866 | -3 | 2.327743 | -3 | 0.12173 | 0.12788 | 0.0538 | 0.0823 | 15 | 0.02 | 0 | Neutral |
| testneg_6541 | 0.12 | -1.16 | 2.568732 | -3 | 0.009894 | -3 | 0.1516 | 0.10704 | 0.07996 | 0.08354 | 29.70842 | 0.03 | 0 | Neutral |
| testneg_6542 | 0.02 | 0.45 | -0.031677 | -3 | -0.028446 | -3 | 0.30669 | 0.11489 | 0.40076 | 0.11341 | 11.96702 | 0.01 | 0 | Neutral |
| testneg_6543 | 0.19 | -0.27 | 4.230056 | -3 | 1.56241 | -3 | 0.08489 | 0.12788 | 0.08386 | 0.12788 | 12.38369 | 0.05 | 0 | Neutral |
| testneg_6544 | 0 | -0.01 | 7.652567 | -3 | 1.348057 | -3 | 0.73253 | 0.09747 | 0.9807 | 0.2062 | 16.61973 | 0.4 | 18.6 | Neutral |
| testneg_6545 | 0 | 0 | 0.596527 | -0.294826 | 6.491905 | -3 | 0.63818 | 0.12788 | 0.2306 | 0.0952 | 11.25 | 0.132845238 | 0 | Neutral |
| testneg_6546 | 0 | 0 | -0.064925 | -3 | 2.177624 | -3 | 0.25924 | 0.19965 | 0.19898 | 0.12788 | 18.34203 | 0.020191327 | 0 | Neutral |
| testneg_6547 | 0 | -0.01 | 0.981481 | -3 | 7.566934 | -3 | 0.24516 | 0.11809 | 0.4404 | 0.38303 | 11.83382 | 0.403447179 | 22 | Neutral |
| testneg_6548 | -0.07 | -0.33 | 1.282009 | -3 | 3.483266 | 3.483266 | 0.04853 | 0.4186 | 0.25856 | 0.17179 | 10.28409 | 0.589446839 | 85.4 | Disease_causing |
| testneg_6549 | -0.02 | -0.41 | -0.086855 | -3 | -0.483923 | -3 | 0.59827 | 0.11479 | 0.23574 | 0.12788 | 29.99249 | 0.02 | 0 | Neutral |
| testneg_6550 | -0.07 | -0.38 | 3.310217 | -3 | 0.456586 | -3 | 0.2293 | 0.12788 | 0.62029 | 0.09082 | 13.63123 | 0.020305556 | 0 | Neutral |
| testneg_6551 | 0 | 0 | 0.549871 | -3 | 4.919497 | -3 | 0.21241 | 0.08684 | 0.29523 | 0.27623 | 18.6768 | 0.196296602 | 0.2 | Neutral |
| testneg_6552 | 0 | 0 | 1.262077 | -3 | 3.318604 | -3 | 0.11351 | 0.10667 | 0.96995 | 0.66487 | 10.28986 | 0.135819444 | 0 | Neutral |
| testneg_6553 | 0 | -0.01 | 0.549483 | -3 | -0.143224 | -3 | 0.46084 | 0.88583 | 0.12176 | 0.09356 | 16.03856 | 0.080300921 | 0 | Neutral |
| testneg_6554 | 0 | 0 | 2.364158 | -3 | 0.538945 | -3 | 0.79983 | 0.1105 | 0.11746 | 0.1471 | 21.40097 | 0.063326927 | 0 | Neutral |
| testneg_6555 | -0.03 | -0.1 | 1.673611 | -3 | 4.688314 | -3 | 0.10554 | 0.12788 | 0.50392 | 0.12788 | 18.70817 | 0.090967678 | 0 | Neutral |
| testneg_6556 | 0.07 | 0.7 | -2.193757 | -3 | 2.246932 | -3 | 0.18795 | 0.09487 | 0.11134 | 0.13818 | 28.85698 | 0.105823725 | 0 | Neutral |
| testneg_6557 | 0.14 | -1.01 | -0.386986 | -3 | -0.094022 | -3 | 0.1196 | 0.12788 | 0.8256 | 0.11062 | 37.51521 | 0.02 | 0 | Neutral |
| testneg_6558 | 0 | 0 | 0.720647 | -3 | 5.915401 | -3 | 0.161 | 0.11597 | 0.16777 | 0.1749 | 26.84685 | 0.320108091 | 5 | Neutral |
| testneg_6559 | 0 | -0.01 | 0.981481 | -3 | 0.395779 | -3 | 0.24516 | 0.11809 | 0.46723 | 0.08832 | 13.74524 | 0 | 0 | Neutral |
| testneg_6560 | -0.08 | -0.03 | 2.891531 | -3 | 1.309085 | -3 | 0.20152 | 0.09895 | 0.64125 | 0.28792 | 11.0671 | 0.180433591 | 0 | Neutral |
| testneg_6561 | 0 | 0 | 2.137302 | -3 | 0.072482 | -3 | 0.73311 | 0.10866 | 0.07394 | 0.09408 | 13.15036 | 0.01 | 0 | Neutral |
| testneg_6562 | 0 | -0.32 | 5.715126 | -3 | 0.381493 | 0.381493 | 0.14199 | 0.1327 | 0.14432 | 0.16919 | 10 | 0.42 | 30.2 | Neutral |
| testneg_6563 | -0.16 | -0.08 | -1.975003 | -1.975003 | 3.183749 | -3 | 0.08465 | 0.15428 | 0.12051 | 0.08503 | 20.41417 | 0.217785948 | 0.2 | Neutral |
| testneg_6564 | 0 | 0 | 1.441246 | -3 | 5.904299 | -3 | 0.33292 | 0.17111 | 0.16716 | 0.10951 | 27.43046 | 0.146811688 | 0 | Neutral |
| testneg_6565 | 0 | 0 | 0.035553 | -3 | 1.204223 | -3 | 0.15414 | 0.17421 | 0.17977 | 0.12788 | 40.28385 | 0.020445098 | 0 | Neutral |
| testneg_6566 | 0 | 0 | 0.395562 | -3 | 2.494251 | -3 | 0.2674 | 0.28725 | 0.10314 | 0.08894 | 12.29167 | 0.040198039 | 0 | Neutral |
| testneg_6567 | 0.16 | 0.08 | 1.765317 | -3 | -0.256362 | -3 | 0.072 | 0.3038 | 0.48561 | 0.09034 | 20.02547 | 0.061433126 | 0 | Neutral |
| testneg_6568 | 0.05 | 0.29 | 1.357664 | 1.357664 | 0.410723 | 0.410723 | 0.1798 | 0.0954 | 0.19721 | 0.12485 | 10 | 0.04 | 0 | Neutral |
| testneg_6569 | 0 | 0 | 0.361863 | -3 | 6.561765 | -3 | 0.04305 | 0.06354 | 0.16394 | 0.08512 | 18.6768 | 0.145390502 | 0 | Neutral |
| testneg_6570 | 0 | 0 | 0.802919 | 0.802919 | 0.386087 | 0.386087 | 0.12042 | 0.11067 | 0.26927 | 0.10752 | 16.82896 | 0.03 | 0 | Neutral |
| testneg_6571 | 0.16 | -1.37 | 6.905837 | -3 | 2.605779 | -3 | 0.27004 | 0.17052 | 0.22817 | 0.10634 | 12.29167 | 0.49025 | 51.8 | Disease_causing |
| testneg_6572 | 0 | 0.32 | 0.66894 | -3 | -0.697584 | -1.247061 | 0.81212 | 0.14915 | 0.19898 | 0.12788 | 20.26596 | 0.020399209 | 0 | Neutral |
| testneg_6573 | 0.05 | 0.29 | 2.610797 | -3 | -0.676534 | -0.676534 | 0.00971 | 0.0791 | 0.16262 | 0.15786 | 10.41667 | 0.120575297 | 0 | Neutral |
| testneg_6574 | 0.08 | 0.03 | 0.468237 | -3 | 2.859591 | -3 | 0.32334 | 0.10525 | 0.1576 | 0.12788 | 18.33333 | 0.050690476 | 0 | Neutral |
| testneg_6575 | 0 | -0.01 | 1.116052 | -3 | 2.549873 | -3 | 0.1631 | 0.40344 | 0.04484 | 0.95426 | 12.40248 | 0.246433385 | 0 | Neutral |
| testneg_6576 | 0 | 0 | 1.078588 | -3 | 0.727897 | -3 | 0.11329 | 0.12116 | 0.34209 | 0.09515 | 15.39226 | 0 | 0 | Neutral |
| testneg_6577 | 0.07 | 0.38 | 0.034534 | 0.034534 | 0.415281 | -3 | 0.62381 | 0.9477 | 0.1472 | 0.12788 | 10.83333 | 0.101623377 | 0 | Neutral |
| testneg_6578 | -0.03 | -0.1 | 5.108849 | -3 | 1.895072 | -3 | 0.49064 | 0.12294 | 0.0773 | 0.14062 | 10.41667 | 0.237083333 | 0 | Neutral |
| testneg_6579 | 0.03 | 0.1 | -0.486948 | -3 | 4.972171 | -3 | 0.25485 | 0.12788 | 0.19721 | 0.12485 | 20.46346 | 0.068270833 | 0 | Neutral |
| testneg_6580 | -0.03 | 1.03 | -0.361103 | -3 | 5.950534 | -3 | 0.26982 | 0.12788 | 0.22278 | 0.85665 | 31.88321 | 0.40028853 | 14.4 | Neutral |
| testneg_6581 | 0.06 | -0.57 | 5.794381 | -3 | -0.711858 | -3 | 0.09734 | 0.08962 | 0.12176 | 0.09356 | 25.82919 | 0.11 | 0 | Neutral |
| testneg_6582 | -0.02 | -0.45 | 2.636151 | -3 | 0.060111 | -3 | 0.94822 | 0.14005 | 0.29098 | 0.10482 | 19.42506 | 0.030817989 | 0 | Neutral |
| testneg_6583 | 0.03 | 0.12 | 1.249184 | -3 | 0.526709 | -3 | 0.40592 | 0.22108 | 0.70616 | 0.09323 | 20.95238 | 0.04 | 0 | Neutral |
| testneg_6584 | 0 | 0 | -0.031197 | -3 | -0.186131 | -3 | 0.1017 | 0.11809 | 0.11802 | 0.09304 | 22.72536 | 0.01 | 0 | Neutral |
| testneg_6585 | -0.04 | 0.23 | 0.421557 | 0.421557 | 1.176476 | -3 | 0.82507 | 0.19615 | 0.64432 | 0.10409 | 7.25667 | 0.033875 | 0 | Neutral |
| testneg_6586 | -0.08 | -0.03 | -0.125177 | -3 | 0.808178 | -3 | 0.27308 | 0.12788 | 0.35513 | 0.21966 | 20 | 0.060284389 | 0 | Neutral |
| testneg_6587 | -0.07 | -0.38 | 3.896611 | -3 | -0.034419 | -0.034419 | 0.57349 | 0.09764 | 0.62203 | 0.12788 | 20.35088 | 0.05 | 0 | Neutral |

| | | | | | | | | | | | | | | |
|---|---|---|---|---|---|---|---|---|---|---|---|---|---|---|
| testneg_6588 | 0 | 0 | 0.938815 | -3 | 3.251074 | -3 | 0.33292 | 0.17111 | 0.20632 | 0.12788 | 14.68988 | 0.05807831 | 0 | Neutral |
| testneg_6589 | 0 | -0.01 | 1.519608 | -3 | -0.526586 | -3 | 0.18206 | 0.10727 | 0.30789 | 0.15135 | 17.21847 | 0.02 | 0 | Neutral |
| testneg_6590 | 0 | 0 | 0.191318 | -3 | 3.006651 | -3 | 0.96113 | 0.24974 | 0.03441 | 0.10663 | 19.09347 | 0.065232323 | 0 | Neutral |
| testneg_6591 | 0.19 | 0.51 | 0.623338 | 0.623338 | 1.029427 | -3 | 0.09707 | 0.12788 | 0.80314 | 0.42344 | 28.09632 | 0.17 | 0 | Neutral |
| testneg_6592 | -0.03 | -0.1 | 0.680291 | -3 | 6.598988 | -3 | 0.23574 | 0.12788 | 0.1454 | 0.10285 | 13.51351 | 0.19 | 0.2 | Neutral |
| testneg_6593 | 0.05 | 0.29 | 2.163438 | -3 | 1.161345 | -3 | 0.8611 | 0.12788 | 0.62339 | 0.19592 | 10.83333 | 0.049615385 | 0 | Neutral |
| testneg_6594 | 0.01 | -0.57 | 5.360167 | -3 | 1.074461 | -3 | 0.09918 | 0.11061 | 0.14379 | 0.12788 | 21.4497 | 0.1145 | 0 | Neutral |
| testneg_6595 | -0.14 | 1.85 | 0.310587 | 0.310587 | 2.354743 | -3 | 0.2132 | 0.12788 | 0.13322 | 0.11297 | 35.56147 | 0.130314259 | 0 | Neutral |
| testneg_6596 | 0 | 0 | 1.741871 | -3 | 1.0663 | 0.102921 | 0.12848 | 0.65153 | 0.19898 | 0.12788 | 28.07329 | 0.08 | 0 | Neutral |
| testneg_6597 | -0.02 | 0.63 | 2.928857 | -3 | 2.664087 | -3 | 0.11534 | 0.12281 | 0.13796 | 0.27869 | 19.13908 | 0.323281571 | 4.8 | Neutral |
| testneg_6598 | -0.05 | -0.29 | 1.091001 | -3 | 3.364563 | 3.364563 | 0.78253 | 0.1114 | 0.07258 | 0.06248 | 28.04038 | 0.180188552 | 0.2 | Neutral |
| testneg_6599 | 0 | 0 | 0.967941 | 0.084018 | 1.242354 | -3 | 0.08365 | 0.34122 | 0.66381 | 0.13039 | 11.25 | 0.088212094 | 0 | Neutral |
| testneg_6600 | 0.02 | -0.36 | 7.015726 | -3 | 0.430481 | -3 | 0.16841 | 0.11143 | 0.31621 | 0.12788 | 19.53147 | 0.208985163 | 0.4 | Neutral |
| testneg_6601 | -0.08 | -0.03 | -0.513148 | -3 | 0.688247 | 0.33974 | 0.06474 | 0.23607 | 0.06431 | 0.12788 | 21.33333 | 0.080246036 | 0 | Neutral |
| testneg_6602 | 0.02818182 | -0.2429091 | 9.773818 | -3 | 4.550951 | -3 | 0.29417 | 0.10965 | 0.21814 | 0.13855 | 22.46228 | 0.619836066 | 89.8 | Disease_causing |
| testneg_6603 | 0.01 | -0.57 | 1.867073 | -3 | 6.087637 | -3 | 0.07999 | 0.23927 | 0.098 | 0.08791 | 17.21847 | 0.289748641 | 1.2 | Neutral |
| testneg_6604 | -0.19 | -0.51 | 2.70164 | -3 | 2.170092 | -3 | 0.1617 | 0.17928 | 0.44479 | 0.20004 | 12.81915 | 0.332418615 | 4.4 | Neutral |
| testneg_6605 | 0 | 0 | 1.817109 | -3 | 6.271279 | -3 | 0.51622 | 0.11809 | 0.12669 | 0.86628 | 14.21141 | 0.367325521 | 10 | Neutral |
| testneg_6606 | -0.01 | 1.44 | 1.762821 | 1.762821 | 1.854685 | -3 | 0.1375 | 0.112 | 0.19898 | 0.1092 | 18.41494 | 0.1 | 0 | Neutral |
| testneg_6607 | -0.01 | -0.79 | 2.44605 | -3 | -0.823621 | -0.823621 | 0.19898 | 0.06726 | 0.27382 | 0.16898 | 44.32245 | 0.09 | 0 | Neutral |
| testneg_6608 | -0.15 | -0.04 | 4.866108 | -3 | 4.276849 | -3 | 0.19898 | 0.12788 | 0.0778 | 0.0728 | 41.61931 | 0.216142857 | 0.2 | Neutral |
| testneg_6609 | 0 | 0 | 0.781134 | -3 | -0.038043 | -3 | 0.92237 | 0.11252 | 0.06729 | 0.17051 | 12.38369 | 0.052186869 | 0 | Neutral |
| testneg_6610 | 0 | 0 | -0.032453 | -3 | 0.241962 | -3 | 0.31664 | 0.14181 | 0.09734 | 0.08962 | 13.05556 | 0.010769231 | 0 | Neutral |
| testneg_6611 | -0.03 | -0.1 | 6.945269 | -3 | 0.721192 | -3 | 0.09972 | 0.13992 | 0.0634 | 0.0824 | 11.25 | 0.317333333 | 5.2 | Neutral |
| testneg_6612 | 0.01 | 0.8 | 6.273159 | -3 | 0.472111 | -3 | 0.39224 | 0.09664 | 0.35561 | 0.08845 | 23.72519 | 0.18 | 0 | Neutral |
| testneg_6613 | -0.04 | 0.89 | 1.989667 | -3 | 2.026116 | -3 | 0.06517 | 0.15107 | 0.75123 | 0.20667 | 21.46199 | 0.193351449 | 0.2 | Neutral |
| testneg_6614 | 0 | 0 | 0.55724 | 0.55724 | 3.968678 | -3 | 0.12937 | 0.23004 | 0.10286 | 0.26819 | 20.74074 | 0.555127935 | 77.2 | Disease_causing |
| testneg_6615 | 0.04 | -0.23 | 7.514495 | -3 | 2.058438 | -3 | 0.31924 | 0.11145 | 0.0749 | 0.12788 | 13.02635 | 0.369642857 | 12.8 | Neutral |
| testneg_6616 | -0.09 | 0.25 | 3.048378 | -3 | 5.537244 | -3 | 0.32861 | 0.09592 | 0.19898 | 0.09972 | 17.05846 | 0.228326659 | 0.4 | Neutral |
| testneg_6617 | 0.05 | -0.81 | 0.668367 | -3 | -0.039157 | -3 | 0.03656 | 0.07074 | 0.04606 | 0.20522 | 26.14035 | 0.14 | 0 | Neutral |
| testneg_6618 | 0 | 0 | 0.173288 | -3 | 1.596413 | -3 | 0.61211 | 0.11059 | 0.80957 | 0.17367 | 5.73716 | 0.013546435 | 0 | Neutral |
| testneg_6619 | 0 | -0.01 | 2.305644 | -3 | -0.187365 | -3 | 0.47787 | 0.25991 | 0.6612 | 0.16696 | 6.00263 | 0.06 | 0 | Neutral |
| testneg_6620 | -0.16 | -0.08 | 6.205256 | -3 | 0.024873 | -3 | 0.12377 | 0.12788 | 0.16507 | 0.10992 | 19.89872 | 0.190047619 | 0.4 | Neutral |
| testneg_6621 | 0 | 0 | 3.603309 | -3 | -1.58983 | -3 | 0.26192 | 0.15452 | 0.21677 | 0.1681 | 11.25 | 0.193707639 | 0 | Neutral |
| testneg_6622 | -0.02 | 0.36 | 1.388896 | -3 | 2.515463 | -3 | 0.40398 | 0.12788 | 0.23723 | 0.1339 | 18.54211 | 0.133566176 | 0 | Neutral |
| testneg_6623 | 0 | 0 | 2.30653 | -3 | -0.048251 | -0.048251 | 0.49106 | 0.89143 | 0.15646 | 0.12788 | 5 | 0.08452381 | 0 | Neutral |
| testneg_6624 | 0 | 0 | -0.057969 | -3 | 1.792345 | -3 | 0.68233 | 0.58998 | 0.14806 | 0.09574 | 11.25 | 0.030909091 | 0 | Neutral |
| testneg_6625 | -0.02 | 0.68 | 1.225058 | -3 | 5.007683 | -3 | 0.30209 | 0.14229 | 0.07415 | 0.12788 | 13.08587 | 0.149236842 | 0 | Neutral |
| testneg_6626 | 0.09 | -2.17 | 6.55944 | -3 | 0.489857 | -3 | 0.25561 | 0.11004 | 0.36257 | 0.26115 | 16.84579 | 0.440972222 | 34.2 | Neutral |
| testneg_6627 | 0 | 0 | 1.481937 | -3 | 5.835416 | -3 | 0.87839 | 0.33588 | 0.86513 | 0.31355 | 10.41667 | 0.297804473 | 3 | Neutral |
| testneg_6628 | 0 | 0 | 0.291054 | -3 | 1.841158 | -3 | 0.28499 | 0.10637 | 0.17075 | 0.12788 | 13.27141 | 7.61E-05 | 0 | Neutral |
| testneg_6629 | -0.08 | -0.03 | 0.307773 | -3 | 6.519516 | -3 | 0.14956 | 0.10585 | 0.28211 | 0.25332 | 23.97163 | 0.444953947 | 31.6 | Neutral |
| testneg_6630 | 0 | 0 | 1.229821 | -3 | 6.33652 | -3 | 0.46091 | 0.13852 | 0.87077 | 0.10593 | 27.99417 | 0.14 | 0 | Neutral |
| testneg_6631 | -0.05 | -0.29 | -1.851788 | -3 | 2.75027 | -3 | 0.78741 | 0.22085 | 0.50034 | 0.07527 | 15.70402 | 0.080461294 | 0 | Neutral |
| testneg_6632 | 0.09 | -0.3 | 5.862225 | -3 | 1.534492 | -3 | 0.21587 | 0.11701 | 0.10183 | 0.11847 | 21.34347 | 0.15 | 0 | Neutral |
| testneg_6633 | 0.05 | -0.81 | 3.756031 | -3 | 3.991032 | -3 | 0.37986 | 0.14229 | 0.18608 | 0.39801 | 10.83333 | 0.677220659 | 97.8 | Disease_causing |
| testneg_6634 | -0.07 | -0.38 | 2.4776 | -3 | 1.51996 | -3 | 0.09577 | 0.2436 | 0.09105 | 0.27434 | 49.73835 | 0.288035714 | 0.2 | Neutral |
| testneg_6635 | 0 | -0.01 | 3.021892 | -3 | -0.409367 | -3 | 0.1163 | 0.20863 | 0.78211 | 0.35815 | 10.41667 | 0.251514745 | 0.2 | Neutral |
| testneg_6636 | -0.15 | 0.28 | 0.62925 | -3 | -1.259137 | -3 | 0.1215 | 0.12788 | 0.15202 | 0.12722 | 36.57683 | 0.06 | 0 | Neutral |
| testneg_6637 | 0 | 0 | 1.34425 | -3 | 0.363314 | -3 | 0.16299 | 0.10454 | 0.10286 | 0.26819 | 19.42906 | 0.063958333 | 0 | Neutral |
| testneg_6638 | -0.08 | -0.03 | 1.732154 | -3 | 2.744533 | -3 | 0.95857 | 0.12137 | 0.1834 | 0.12788 | 27.98069 | 0.041356322 | 0 | Neutral |
| testneg_6639 | 0.07 | -0.31 | 5.632523 | -3 | 3.1183 | -3 | 0.17764 | 0.09681 | 0.12921 | 0.11605 | 17.92801 | 0.24697619 | 0.8 | Neutral |
| testneg_6640 | 0.16 | -0.37 | 4.774952 | -3 | 0.710444 | -3 | 0.10472 | 0.10014 | 0.56554 | 0.76978 | 10 | 0.232192641 | 0.6 | Neutral |
| testneg_6641 | 0.03 | -0.85 | 2.681264 | -3 | -1.05361 | -3 | 0.25961 | 0.07098 | 0.4558 | 0.9277 | 18.39615 | 0.13343254 | 0 | Neutral |
| testneg_6642 | 0.14 | -0.37 | 4.181702 | -3 | 2.705745 | -3 | 0.04123 | 0.06896 | 0.15041 | 0.43468 | 11.25 | 0.413762207 | 22.6 | Neutral |
| testneg_6643 | -0.2 | 1.26 | 5.557778 | -3 | -2.304927 | -2.304927 | 0.72163 | 0.10596 | 0.02665 | 0.05082 | 11.875 | 0.21 | 0.2 | Neutral |
| testneg_6644 | -0.02 | 0.68 | 1.913159 | -3 | 0.268278 | 0.268278 | 0.91921 | 0.32331 | 0.09669 | 0.12526 | 11.25 | 0.06 | 0 | Neutral |
| testneg_6645 | 0 | -0.32 | 3.34105 | -3 | 1.276927 | -3 | 0.33923 | 0.12032 | 0.024 | 0.07726 | 45.3252 | 0.03 | 0 | Neutral |
| testneg_6646 | -0.07 | -0.33 | 1.282461 | -3 | -0.190524 | -3 | 0.59131 | 0.66216 | 0.13322 | 0.11297 | 34.45035 | 0.060325906 | 0 | Neutral |
| testneg_6647 | 0 | 0 | 0.680876 | -3 | -0.093394 | -0.093394 | 0.1602 | 0.11262 | 0.34587 | 0.29822 | 30.6499 | 0.063463152 | 0 | Neutral |
| testneg_6648 | -0.06 | -0.18 | 4.092031 | -3 | -0.894688 | -3 | 0.50134 | 0.15488 | 0.05817 | 0.09497 | 48.7718 | 0.076422414 | 0 | Neutral |
| testneg_6649 | -0.07 | -0.38 | 1.741657 | -3 | 1.745688 | -3 | 0.20513 | 0.09801 | 0.11616 | 0.12788 | 32.84612 | 0.01 | 0 | Neutral |
| testneg_6650 | 0.08 | 0.03 | -0.412359 | -3 | -0.096052 | -3 | 0.4003 | 0.10884 | 0.3095 | 0.13502 | 29.15809 | 0.08 | 0 | Neutral |
| testneg_6651 | -0.07 | -0.38 | 1.619888 | -3 | 5.495658 | -3 | 0.19477 | 0.12788 | 0.15294 | 0.12788 | 17.21847 | 0.128579235 | 0 | Neutral |
| testneg_6652 | -0.18 | 1.05 | 0.601635 | -3 | -0.206553 | -3 | 0.51675 | 0.11055 | 0.33026 | 0.30576 | 18.80346 | 0.14 | 0 | Neutral |
| testneg_6653 | -0.07 | -0.38 | 1.315709 | -3 | 7.107668 | -3 | 0.28341 | 0.1005 | 0.26731 | 0.16415 | 20.78359 | 0.478539676 | 45.4 | Neutral |
| testneg_6654 | 0.04 | -0.23 | 0.982196 | -3 | 4.745771 | -3 | 0.19898 | 0.09322 | 0.14524 | 0.10162 | 44.74474 | 0.045279221 | 0 | Neutral |
| testneg_6655 | -0.07 | -0.01 | -1.378489 | -3 | -0.275748 | -3 | 0.15341 | 0.12272 | 0.13779 | 0.0945 | 11.93907 | 0.010223806 | 0 | Neutral |
| testneg_6656 | 0 | -0.01 | 5.657974 | -3 | -0.187363 | -3 | 0.25462 | 0.12788 | 0.90649 | 0.3543 | 10.18729 | 0.323030303 | 5.8 | Neutral |
| testneg_6657 | 0 | 0 | 2.17719 | -3 | 5.034072 | -3 | 0.16091 | 0.11571 | 0.08212 | 0.12788 | 23.43525 | 0.080354374 | 0 | Neutral |
| testneg_6658 | 0 | 0 | 4.336736 | -3 | 1.910801 | -3 | 0.27275 | 0.11752 | 0.15446 | 0.08618 | 20.50765 | 0.030169528 | 0 | Neutral |
| testneg_6659 | 0.07 | 0.38 | -2.0307 | -3 | 2.477566 | -3 | 0.06848 | 0.09963 | 0.29451 | 0.13582 | 12.46106 | 0.121291667 | 0 | Neutral |
| testneg_6660 | -0.02 | -0.45 | 2.569812 | -3 | 2.12021 | 2.12021 | 0.16404 | 0.09378 | 0.07698 | 0.12788 | 12.31231 | 0.0658 | 0 | Neutral |
| testneg_6661 | 0.04 | -0.23 | 5.274307 | -3 | -1.941492 | -3 | 0.25516 | 0.12788 | 0.37711 | 0.12788 | 13.51351 | 0.1 | 0 | Neutral |
| testneg_6662 | -0.0138182 | -0.2629091 | 4.820901 | -3 | 2.624257 | -3 | 0.06358 | 0.5364 | 0.19043 | 0.07855 | 18.69495 | 0.270627451 | 0.2 | Neutral |
| testneg_6663 | 0.05 | -0.81 | 0.543748 | -3 | 1.959161 | -3 | 0.34153 | 0.08855 | 0.18312 | 0.12788 | 17.78631 | 0.01 | 0 | Neutral |
| testneg_6664 | -0.07 | -0.38 | -1.556144 | -3 | 1.270092 | -3 | 0.25831 | 0.10988 | 0.63497 | 0.23021 | 15.35348 | 0.05 | 0 | Neutral |
| testneg_6665 | 0.14 | -1.01 | 0.667614 | -3 | 5.23082 | -3 | 0.47997 | 0.10506 | 0.90112 | 0.20622 | 18.33333 | 0.31347619 | 2.6 | Neutral |
| testneg_6666 | 0.03 | 0.12 | 0.235146 | -3 | 5.213775 | -3 | 0.16856 | 0.19221 | 0.10671 | 0.10867 | 32.12477 | 0.186877289 | 0.2 | Neutral |
| testneg_6667 | 0 | 0 | -0.014147 | -3 | 2.452648 | -3 | 0.14101 | 0.14318 | 0.30926 | 0.11955 | 20.83333 | 0.028 | 0 | Neutral |
| testneg_6668 | 0.01 | 0.79 | 3.405014 | -3 | 0.982052 | -3 | 0.32423 | 0.08104 | 0.09211 | 0.08973 | 10.19035 | 0.05 | 0 | Neutral |
| testneg_6669 | 0 | -0.01 | 0.558245 | -3 | 0.435323 | -3 | 0.19898 | 0.10362 | 0.20527 | 0.10053 | 20.66262 | 0 | 0 | Neutral |
| testneg_6670 | -0.06 | -0.18 | 2.103681 | -3 | 1.272902 | -3 | 0.19898 | 0.13434 | 0.19898 | 0.12788 | 29.21475 | 0.01 | 0 | Neutral |
| testneg_6671 | 0 | 0 | -0.028324 | -3 | 3.36026 | -3 | 0.20675 | 0.13709 | 0.78253 | 0.1114 | 9.88546 | 0.043384615 | 0 | Neutral |
| testneg_6672 | 0.04 | -0.23 | 1.605283 | -3 | -1.216304 | -3 | 0.7257 | 0.64394 | 0.09669 | 0.12526 | 19.44444 | 0.061291667 | 0 | Neutral |
| testneg_6673 | -0.15 | -0.04 | 0.012434 | -3 | 3.451445 | -3 | 0.08629 | 0.12788 | 0.49064 | 0.12294 | 12.32809 | 0.130323929 | 0 | Neutral |
| testneg_6674 | -0.07 | -0.01 | -0.919759 | -3 | -0.749376 | -0.749376 | 0.24791 | 0.09675 | 0.51437 | 0.17173 | 12.30072 | 0.05 | 0 | Neutral |
| testneg_6675 | -0.02 | -0.45 | 1.91875 | -3 | 4.981525 | -3 | 0.04525 | 0.12788 | 0.17977 | 0.12788 | 38.80952 | 0.13 | 0 | Neutral |
| testneg_6676 | 0 | -0.01 | 1.599412 | -3 | 6.939074 | -3 | 0.13099 | 0.17453 | 0.07467 | 0.09224 | 13.34868 | 0.368784119 | 12 | Neutral |
| testneg_6677 | -0.02 | 0.36 | -0.000915 | -3 | 4.192605 | -3 | 0.10349 | 0.12788 | 0.08454 | 0.22908 | 20.27778 | 0.339800725 | 6.4 | Neutral |
| testneg_6678 | -0.15 | -0.04 | 4.866108 | -3 | 2.798182 | 2.798182 | 0.19898 | 0.12788 | 0.31043 | 0.09688 | 38.44879 | 0.23011236 | 0.2 | Neutral |
| testneg_6679 | 0.16 | 0.08 | 4.903998 | -3 | 1.408558 | -3 | 0.22816 | 0.10878 | 0.77981 | 0.62662 | 12.29167 | 0.22 | 0.2 | Neutral |
| testneg_6680 | 0.16 | -0.37 | 5.42275 | 1.824862 | 0.62394 | -3 | 0.44816 | 0.56655 | 0.90813 | 0.10329 | 4.72222 | 0.353289627 | 4.8 | Neutral |
| testneg_6681 | 0.18 | -1.05 | 6.296843 | -3 | 2.331142 | -3 | 0.20327 | 0.11174 | 0.46037 | 0.1343 | 26.24408 | 0.32 | 4.6 | Neutral |
| testneg_6682 | 0.16 | 0.08 | 0.019411 | -3 | -1.722922 | -3 | 0.10863 | 0.098 | 0.06704 | 0.1068 | 23.68839 | 0.02 | 0 | Neutral |
| testneg_6683 | 0.21 | 0.15 | 3.813632 | -3 | 1.075324 | 1.075324 | 0.94807 | 0.14977 | 0.49934 | 0.11069 | 5.85199 | 0.095587662 | 0 | Neutral |
| testneg_6684 | 0.04 | -0.23 | -0.852628 | -3 | 10.671823 | -3 | 0.06636 | 0.1079 | 0.13749 | 0.17824 | 16.8018 | 0.48 | 44.8 | Neutral |
| testneg_6685 | 0 | 0 | 1.075324 | 1.075324 | 4.672344 | -3 | 0.49934 | 0.11069 | 0.80034 | 0.10518 | 10.38652 | 0.11 | 0 | Neutral |
| testneg_6686 | 0.03 | 1.22 | 1.359632 | -3 | 2.019398 | -3 | 0.38171 | 0.15977 | 0.09997 | 0.09515 | 20.8819 | 0.08026289 | 0 | Neutral |
| testneg_6687 | -0.17 | 1.22 | -0.126067 | -3 | 0.493256 | -3 | 0.19898 | 0.09171 | 0.13126 | 0.11628 | 22.67913 | 0.1 | 0 | Neutral |
| testneg_6688 | 0.05 | -0.81 | 2.225911 | -3 | 6.080162 | -3 | 0.91619 | 0.10042 | 0.50448 | 0.50327 | 9.76428 | 0.394380193 | 18.8 | Neutral |
| testneg_6689 | 0.03 | 0.12 | 0.90434 | -3 | 7.311333 | -3 | 0.53424 | 0.08484 | 0.10231 | 0.1 | 12.82738 | 0.26 | 1.4 | Neutral |
| testneg_6690 | 0 | 0 | 1.61019 | -3 | 0.73341 | -3 | 0.16965 | 0.25991 | 0.54792 | 0.54626 | 12.29167 | 0.085973058 | 0 | Neutral |
| testneg_6691 | 0 | -0.32 | 0.641608 | -3 | 3.148371 | -3 | 0.32458 | 0.1016 | 0.18102 | 0.6537 | 21.42525 | 0.202073308 | 0.2 | Neutral |
| testneg_6692 | 0.03 | 1.22 | 1.399794 | -3 | 5.377935 | -3 | 0.49425 | 0.25538 | 0.52028 | 0.47239 | 10 | 0.606912333 | 89.2 | Disease_causing |
| testneg_6693 | 0 | 0 | 0.353546 | -3 | 0.303207 | -3 | 0.28191 | 0.20693 | 0.23401 | 0.12374 | 10 | 0.010944984 | 0 | Neutral |
| testneg_6694 | -0.07 | -0.38 | 5.775288 | -3 | -0.392377 | -3 | 0.16663 | 0.12788 | 0.24087 | 0.27985 | 12.70833 | 0.43361828 | 29 | Neutral |
| testneg_6695 | 0.04 | -1.61 | 6.076061 | -3 | 5.078544 | -3 | 0.28038 | 0.11814 | 0.2098 | 0.44672 | 18.30413 | 0.743429779 | 99 | Disease_causing |
| testneg_6696 | 0 | 0 | 0.606074 | -3 | 1.505821 | -3 | 0.13647 | 0.11806 | 0.0421 | 0.17669 | 12.40248 | 0.042832826 | 0 | Neutral |
| testneg_6697 | 0 | -0.01 | 2.305644 | -3 | 3.473048 | -3 | 0.47787 | 0.25991 | 0.14485 | 0.16945 | 16.42913 | 0.397707683 | 16.6 | Neutral |
| testneg_6698 | -0.09 | 0.3 | 1.202839 | -3 | 0.615335 | -3 | 0.11144 | 0.08199 | 0.70872 | 0.09884 | 18.46348 | 0.04 | 0 | Neutral |
| testneg_6699 | 0 | 0 | 0.244145 | -3 | 0.754007 | -3 | 0.9662 | 0.28104 | 0.2172 | 0.10891 | 10.67125 | 0.01020999 | 0 | Neutral |
| testneg_6700 | 0 | 0 | 0.596959 | -3 | -0.536837 | -3 | 0.18925 | 0.32479 | 0.02617 | 0.12788 | 38.96325 | 0.034431818 | 0 | Neutral |
| testneg_6701 | 0 | 0 | 0.60694 | -3 | 1.937875 | -3 | 0.07387 | 0.12788 | 0.16393 | 0.08897 | 36.39339 | 0.01 | 0 | Neutral |
| testneg_6702 | 0 | 0 | 0.57586 | -3 | 9.170362 | -3 | 0.23567 | 0.12788 | 0.1144 | 0.12788 | 34.26925 | 0.31 | 3.6 | Neutral |
| testneg_6703 | 0 | -0.32 | -2.076593 | -3 | 6.193509 | -3 | 0.1276 | 0.30104 | 0.31817 | 0.17348 | 32.38889 | 0.600458695 | 89.6 | Disease_causing |
| testneg_6704 | -0.09 | 0.3 | 0.27631 | -3 | -0.61726 | -3 | 0.19898 | 0.10591 | 0.20673 | 0.13236 | 11.98582 | 0.07 | 0 | Neutral |
| testneg_6705 | 0.15 | -0.82 | -1.286656 | -1.286656 | 2.81231 | -3 | 0.10208 | 0.08867 | 0.06647 | 0.09129 | 19.84929 | 0.053445693 | 0 | Neutral |
| testneg_6706 | 0.19 | -0.25 | 0.898775 | -3 | 1.49197 | -3 | 0.3221 | 0.3367 | 0.05005 | 0.07051 | 18.33333 | 0.07 | 0 | Neutral |
| testneg_6707 | 0.2 | -1.98 | 5.220127 | -3 | 2.667225 | -3 | 0.24247 | 0.19222 | 0.14668 | 0.17962 | 8.80915 | 0.657843826 | 94.6 | Disease_causing |
| testneg_6708 | 0 | 0 | -0.22828 | -3 | 3.21678 | -3 | 0.31694 | 0.13475 | 0.99998 | 0.9588 | 12.20915 | 0.231038955 | 0.4 | Neutral |
| testneg_6709 | -0.04 | 0.23 | 1.327969 | -3 | 1.500259 | -3 | 0.60812 | 0.19286 | 0.1787 | 0.1327 | 20.27704 | 0.120590278 | 0 | Neutral |

| | | | | | | | | | | | | | | |
|---|---|---|---|---|---|---|---|---|---|---|---|---|---|---|
| testneg_6710 | -0.02 | 0.36 | 1.311291 | -3 | 3.705834 | -3 | 0.07577 | 0.09128 | 0.16781 | 0.12788 | 12.70833 | 0.08 | 0 | Neutral |
| testneg_6711 | 0 | 0 | 0.528019 | -3 | 0.338162 | -3 | 0.42621 | 0.49093 | 0.58827 | 0.57942 | 5.69444 | 0.064287737 | 0 | Neutral |
| testneg_6712 | 0.06 | -0.57 | 0.237935 | -3 | 0.256955 | -3 | 0.16362 | 0.11879 | 0.1846 | 0.08682 | 20.56995 | 0.00059419 | 0 | Neutral |
| testneg_6713 | 0.03 | 0.12 | 2.332457 | -3 | 2.050574 | -3 | 0.358 | 0.11262 | 0.69563 | 0.12788 | 20.83333 | 0.02 | 0 | Neutral |
| testneg_6714 | -0.04 | 0.23 | 1.474779 | 1.474779 | 3.593276 | -3 | 0.0882 | 0.12788 | 0.17793 | 0.10516 | 29.86111 | 0.1 | 0 | Neutral |
| testneg_6715 | -0.03 | 1.03 | 3.441233 | -3 | 1.690495 | -3 | 0.22046 | 0.12788 | 0.11891 | 0.11186 | 17.70469 | 0.059916667 | 0 | Neutral |
| testneg_6716 | 0.07 | 0.38 | 0.034534 | 0.034534 | 1.378582 | -3 | 0.62381 | 0.9477 | 0.08184 | 0.10206 | 4.28195 | 0.09 | 0 | Neutral |
| testneg_6717 | 0 | 0 | 0.103447 | 0.103447 | 0.474638 | -3 | 0.15813 | 0.61197 | 0.09284 | 0.10978 | 16.34409 | 0.05 | 0 | Neutral |
| testneg_6718 | 0 | 0 | 0.229773 | -3 | 1.568235 | -3 | 0.13275 | 0.12788 | 0.9662 | 0.28104 | 10.30744 | 0.021666667 | 0 | Neutral |
| testneg_6719 | 0.02 | 0.41 | 2.511378 | -3 | 0.882673 | -3 | 0.05886 | 0.38242 | 0.19052 | 0.12788 | 5.88186 | 0.091675028 | 0 | Neutral |
| testneg_6720 | 0.21 | 0.15 | 5.740215 | -3 | 4.819528 | -3 | 0.18499 | 0.10583 | 0.1211 | 0.1526 | 18.33333 | 0.63 | 90.8 | Disease_causing |
| testneg_6721 | 0 | -0.32 | 0.284563 | -3 | 3.477533 | -3 | 0.39436 | 0.16446 | 0.87847 | 0.96194 | 17.21847 | 0.338654503 | 4.4 | Neutral |
| testneg_6722 | 0.14 | -0.37 | -0.136533 | -3 | 7.351796 | -3 | 0.05932 | 0.09551 | 0.1211 | 0.1526 | 21.33768 | 0.44074999 | 32.2 | Neutral |
| testneg_6723 | -0.02 | 0.63 | 0.385855 | -3 | 6.445857 | -3 | 0.10153 | 0.13343 | 0.12463 | 0.12788 | 20.32229 | 0.26 | 0.4 | Neutral |
| testneg_6724 | -0.02 | -0.45 | 2.458755 | -3 | 6.693407 | -3 | 0.11587 | 0.12001 | 0.1475 | 0.4323 | 11.25 | 0.570132807 | 81.6 | Disease_causing |
| testneg_6725 | -0.05 | -1.52 | 1.926463 | -3 | 8.821972 | 2.156053 | 0.70616 | 0.09323 | 0.05546 | 0.28027 | 21.50794 | 0.49 | 52.2 | Disease_causing |
| testneg_6726 | 0.09 | -1.07 | -0.111744 | -3 | 4.70798 | -3 | 0.22078 | 0.12788 | 0.17527 | 0.13941 | 27.69024 | 0.282320717 | 1.2 | Neutral |
| testneg_6727 | 0.07 | 0.33 | 0.32945 | -3 | -0.140634 | -3 | 0.21623 | 0.42074 | 0.19898 | 0.12788 | 10 | 0.04 | 0 | Neutral |
| testneg_6728 | 0 | 0 | 0.864288 | -3 | 4.607222 | -3 | 0.21014 | 0.12378 | 0.29268 | 0.16012 | 14.27748 | 0.230493011 | 0 | Neutral |
| testneg_6729 | 0 | 0 | 0.531067 | -3 | 2.430735 | -3 | 0.23461 | 0.11809 | 0.28537 | 0.12788 | 26.23656 | 0.01 | 0 | Neutral |
| testneg_6730 | 0.14 | -0.69 | 4.217892 | -3 | -0.071864 | -0.071864 | 0.71697 | 0.19162 | 0.97005 | 0.31407 | 4.72222 | 0.281711806 | 1.2 | Neutral |
| testneg_6731 | -0.03 | -0.12 | -0.214687 | -3 | 1.118815 | -3 | 0.19668 | 0.16042 | 0.13199 | 0.14263 | 13.02635 | 0.095676269 | 0 | Neutral |
| testneg_6732 | 0 | -0.01 | 1.594546 | -3 | 0.389489 | -3 | 0.33304 | 0.15588 | 0.11858 | 0.11551 | 22.38311 | 0.010250156 | 0 | Neutral |
| testneg_6733 | 0 | 0 | 0.921407 | -3 | 0.421557 | 0.421557 | 0.19898 | 0.14229 | 0.82507 | 0.19615 | 14.10753 | 0.04 | 0 | Neutral |
| testneg_6734 | 0.18 | -1.05 | 4.914299 | -3 | 1.1308 | 1.1308 | 0.16734 | 0.15176 | 0.19898 | 0.11185 | 37.20713 | 0.1 | 0 | Neutral |
| testneg_6735 | 0.17 | 0.38 | 0.477392 | -3 | 1.123147 | -3 | 0.09682 | 0.09596 | 0.59947 | 0.14339 | 14.47674 | 0.06 | 0 | Neutral |
| testneg_6736 | 0 | 0 | 0.202333 | -0.608116 | 1.299976 | -3 | 0.0925 | 0.08222 | 0.16309 | 0.12788 | 21.89323 | 0.01 | 0 | Neutral |
| testneg_6737 | 0.09 | -0.3 | 5.898353 | -3 | 5.122764 | -3 | 0.42919 | 0.1092 | 0.19898 | 0.12788 | 27.52646 | 0.39 | 22 | Neutral |
| testneg_6738 | 0.14 | -1.01 | 3.206305 | -3 | 0.603592 | -3 | 0.38503 | 0.11286 | 0.08973 | 0.0992 | 9.89855 | 0.03 | 0 | Neutral |
| testneg_6739 | 0.08 | 0.03 | 1.175993 | -3 | 0.08175 | -3 | 0.06781 | 0.09009 | 0.32334 | 0.10525 | 22.74576 | 0.02 | 0 | Neutral |
| testneg_6740 | -0.18 | 1.05 | 4.118674 | -3 | 4.039717 | -3 | 0.04795 | 0.07008 | 0.20546 | 0.10734 | 23.68839 | 0.317291667 | 1.2 | Neutral |
| testneg_6741 | 0.05 | -0.81 | 0.203743 | -3 | 1.455085 | -3 | 0.05323 | 0.32175 | 0.15958 | 0.41628 | 11.25 | 0.284857143 | 1.4 | Neutral |
| testneg_6742 | -0.05 | -0.29 | 1.124718 | -3 | 1.220387 | 0.985437 | 0.55043 | 0.12112 | 0.1601 | 0.12546 | 13.14653 | 0.010138824 | 0 | Neutral |
| testneg_6743 | 0 | 0 | 0.09433 | -3 | -0.063333 | -3 | 0.18925 | 0.32479 | 0.493 | 0.35835 | 10 | 0.10855096 | 0 | Neutral |
| testneg_6744 | 0 | 0 | -0.101683 | -3 | 2.97112 | -3 | 0.72657 | 0.11262 | 0.5901 | 0.19444 | 12.40248 | 0.08997619 | 0 | Neutral |
| testneg_6745 | 0 | 0 | 2.165106 | -3 | 6.375543 | -3 | 0.30257 | 0.17298 | 0.07973 | 0.09208 | 23.04399 | 0.244620446 | 0 | Neutral |
| testneg_6746 | -0.05 | -0.29 | 0.368341 | -3 | 0.892974 | 0.892974 | 0.77847 | 0.10751 | 0.06986 | 0.12788 | 15.58192 | 0.02 | 0 | Neutral |
| testneg_6747 | 0.14 | -1.01 | 3.23372 | -3 | -0.484253 | -3 | 0.73947 | 0.2058 | 0.12836 | 0.09736 | 31.65176 | 0.087460317 | 0 | Neutral |
| testneg_6748 | 0.04 | -0.23 | 3.983209 | -3 | 0.097083 | -3 | 0.11144 | 0.08199 | 0.12943 | 0.14391 | 13.80482 | 0.09211281 | 0 | Neutral |
| testneg_6749 | -0.09 | 0.25 | 5.71198 | -3 | 0.002159 | -3 | 0.3112 | 0.33703 | 0.10231 | 0.1 | 6.94444 | 0.29 | 2.4 | Neutral |
| testneg_6750 | -0.21 | -0.15 | -0.374248 | -0.833744 | 4.515915 | -3 | 0.31646 | 0.09844 | 0.49644 | 0.10958 | 18.88889 | 0.133783525 | 0 | Neutral |
| testneg_6751 | 0.04 | -0.23 | 0.007971 | -3 | 1.259454 | -3 | 0.12591 | 0.09845 | 0.16259 | 0.102 | 13.51351 | 0.000360665 | 0 | Neutral |
| testneg_6752 | 0.02 | -0.28 | 0.430481 | -3 | 2.045511 | -3 | 0.31621 | 0.12788 | 0.20222 | 0.12788 | 20.84235 | 0.01 | 0 | Neutral |
| testneg_6753 | 0.02 | -0.63 | 1.995171 | -3 | 2.229991 | 2.229991 | 0.91377 | 0.5142 | 0.0359 | 0.09574 | 6.96873 | 0.09 | 0 | Neutral |
| testneg_6754 | -0.07 | -0.33 | 1.282461 | -3 | -1.333487 | -3 | 0.59131 | 0.66216 | 0.07598 | 0.08456 | 10.26553 | 0.032 | 0 | Neutral |
| testneg_6755 | -0.14 | 0.69 | 0.437223 | -3 | 3.910634 | -3 | 0.0684 | 0.08803 | 0.15092 | 0.09074 | 23.74044 | 0.162117459 | 0 | Neutral |
| testneg_6756 | 0.21 | 0.15 | 2.956317 | -3 | 0.725748 | -3 | 0.9637 | 0.12788 | 0.19898 | 0.12788 | 9.67871 | 0.06 | 0 | Neutral |
| testneg_6757 | 0 | 0 | 0.01717 | -3 | 0.441329 | -3 | 0.19898 | 0.16603 | 0.28269 | 0.12505 | 21.51158 | 0.02 | 0 | Neutral |
| testneg_6758 | 0 | 0 | 2.403686 | 2.403686 | 2.259657 | -3 | 0.48574 | 0.20473 | 0.11657 | 0.21661 | 13.40128 | 0.338016266 | 11.4 | Neutral |
| testneg_6759 | -0.08 | -0.03 | -1.275641 | -3 | 5.213254 | -0.010367 | 0.05546 | 0.28027 | 0.62172 | 0.18135 | 14.16667 | 0.610590201 | 89.2 | Disease_causing |
| testneg_6760 | -0.09 | 2.17 | 1.212248 | -3 | 4.513115 | -3 | 0.02396 | 0.12703 | 0.03526 | 0.06855 | 13.10771 | 0.22 | 0 | Neutral |
| testneg_6761 | 0.14 | -1.01 | 2.297111 | -3 | 9.90393 | -3 | 0.39111 | 0.15887 | 0.21002 | 0.12788 | 13.51351 | 0.49 | 50.8 | Disease_causing |
| testneg_6762 | 0.04 | 2.01 | 1.330982 | -3 | 0.20814 | -3 | 0.12643 | 0.12266 | 0.19898 | 0.12788 | 10 | 0.07 | 0 | Neutral |
| testneg_6763 | 0 | 0 | 1.222706 | -3 | -1.628421 | -1.628421 | 0.35376 | 0.09478 | 0.54441 | 0.08891 | 13.75 | 0.01 | 0 | Neutral |
| testneg_6764 | 0.07 | 0.38 | -3.926089 | -3 | 4.83497 | -3 | 0.12839 | 0.09503 | 0.87077 | 0.10593 | 17.79895 | 0.1 | 0 | Neutral |
| testneg_6765 | 0.04 | -0.23 | 6.188656 | -3 | 3.824646 | -3 | 0.29624 | 0.1067 | 0.56487 | 0.12788 | 13.51351 | 0.339924242 | 6.8 | Neutral |
| testneg_6766 | -0.02 | 0.36 | -0.000915 | -3 | 4.461467 | -3 | 0.10349 | 0.12788 | 0.20547 | 0.12178 | 20.27778 | 0.081547619 | 0 | Neutral |
| testneg_6767 | -0.21 | -0.15 | 0.585797 | -3 | 3.781746 | -3 | 0.25312 | 0.12788 | 0.52771 | 0.10974 | 15.722 | 0.122375 | 0 | Neutral |
| testneg_6768 | 0 | 0 | 4.50109 | -3 | 2.519988 | -3 | 0.26731 | 0.16415 | 0.16507 | 0.10992 | 21.38889 | 0.133081816 | 0 | Neutral |
| testneg_6769 | 0.04 | -0.23 | 7.072574 | -3 | 2.091718 | 1.121663 | 0.15016 | 0.08826 | 0.34946 | 0.19092 | 18.33333 | 0.459816667 | 37 | Neutral |
| testneg_6770 | 0.04 | -0.23 | 5.836424 | -3 | 6.257897 | -3 | 0.1503 | 0.12788 | 0.14735 | 0.12788 | 12.29167 | 0.5544161 | 72.8 | Disease_causing |
| testneg_6771 | 0 | 0 | 0.137295 | -3 | 0.470367 | -3 | 0.48027 | 0.52799 | 0.23784 | 0.15573 | 10.83333 | 0.090262309 | 0 | Neutral |
| testneg_6772 | 0 | 0 | -0.071864 | -3 | 3.28103 | -3 | 0.97005 | 0.31407 | 0.27077 | 0.10337 | 11.96702 | 0.07 | 0 | Neutral |
| testneg_6773 | -0.1 | -1.9 | 2.083029 | -3 | 0.002159 | -3 | 0.16929 | 0.12788 | 0.10231 | 0.1 | 14.89247 | 0.04 | 0 | Neutral |
| testneg_6774 | 0 | 0 | -0.136163 | -3 | 0.947772 | -3 | 0.0513 | 0.33622 | 0.14668 | 0.17962 | 25.95163 | 0.259 | 0 | Neutral |
| testneg_6775 | 0 | 0 | 1.687316 | -3 | 1.582915 | -3 | 0.26434 | 0.12788 | 0.09493 | 0.104 | 36.11526 | 0.01 | 0 | Neutral |
| testneg_6776 | 0 | 0 | -0.104866 | -3 | 4.914058 | -3 | 0.2698 | 0.18815 | 0.44594 | 0.13793 | 11.25552 | 0.344436721 | 10.4 | Neutral |
| testneg_6777 | 0 | 0 | 1.003967 | -3 | 5.935147 | -3 | 0.12835 | 0.15143 | 0.19898 | 0.10269 | 27.25845 | 0.167441398 | 0 | Neutral |
| testneg_6778 | 0.03 | 0.1 | -1.608862 | -3 | 3.58877 | -3 | 0.31732 | 0.27963 | 0.37074 | 0.11532 | 50.14032 | 0.142843818 | 0 | Neutral |
| testneg_6779 | 0.04 | -1.36 | 1.784639 | -3 | 4.3331 | -3 | 0.64795 | 0.21581 | 0.19898 | 0.09201 | 10.83333 | 0.189378788 | 0 | Neutral |
| testneg_6780 | -0.04 | 0.23 | 0.6838 | 0.6838 | 6.195353 | -3 | 0.67327 | 0.23543 | 0.28532 | 0.10663 | 26.5086 | 0.232435897 | 0.2 | Neutral |
| testneg_6781 | 0.07 | 0.7 | 5.661416 | -3 | 2.53141 | 2.53141 | 0.9352 | 0.14229 | 0.1123 | 0.1568 | 21.43427 | 0.57 | 81.2 | Disease_causing |
| testneg_6782 | 0.16 | -1.37 | 1.096466 | -3 | 1.837787 | -0.695009 | 0.49745 | 0.1259 | 0.75906 | 0.09702 | 21.49496 | 0.05 | 0 | Neutral |
| testneg_6783 | 0.14 | -1.01 | 2.081126 | -3 | 0.28162 | -3 | 0.2663 | 0.32806 | 0.6271 | 0.09663 | 14.22149 | 0.051742424 | 0 | Neutral |
| testneg_6784 | 0.07 | 0.38 | -2.0307 | -3 | 1.214192 | -3 | 0.06848 | 0.09963 | 0.19898 | 0.12788 | 22.61823 | 0.02 | 0 | Neutral |
| testneg_6785 | 0 | 0 | 0.608537 | -3 | 0.353052 | -3 | 0.68029 | 0.12788 | 0.71968 | 0.54266 | 10.41667 | 0.020377493 | 0 | Neutral |
| testneg_6786 | 0.14 | -1.01 | 0.302883 | -3 | 3.228883 | -3 | 0.10486 | 0.08083 | 0.14668 | 0.17962 | 22.93219 | 0.23 | 0 | Neutral |
| testneg_6787 | 0 | 0 | 0.616227 | -3 | 0.219198 | -3 | 0.02971 | 0.33983 | 0.79983 | 0.1105 | 31.06559 | 0.1 | 0 | Neutral |
| testneg_6788 | 0.14 | -1.01 | 2.213678 | -3 | 2.679679 | -3 | 0.06736 | 0.06344 | 0.17697 | 0.12974 | 29.50364 | 0.05 | 0 | Neutral |
| testneg_6789 | 0 | -0.01 | 1.947226 | -3 | -0.539741 | -3 | 0.98926 | 0.53296 | 0.10895 | 0.11082 | 20.92347 | 0.05 | 0 | Neutral |
| testneg_6790 | 0.03 | 0.12 | 1.294111 | -3 | 3.825677 | -3 | 0.22575 | 0.37247 | 0.12832 | 0.07113 | 18.68505 | 0.148454545 | 0 | Neutral |
| testneg_6791 | 0 | 0 | -0.040357 | -3 | 4.092943 | -3 | 0.01584 | 0.07827 | 0.83421 | 0.18817 | 12.22403 | 0.190489096 | 0 | Neutral |
| testneg_6792 | 0 | -0.01 | 0.583496 | -3 | 0.463587 | -3 | 0.40663 | 0.24974 | 0.36494 | 0.12763 | 13.06326 | 0.01 | 0 | Neutral |
| testneg_6793 | 0.18 | -1.05 | 5.390189 | -3 | -1.991505 | -3 | 0.15041 | 0.43468 | 0.10498 | 0.08421 | 10.83333 | 0.22 | 0 | Neutral |
| testneg_6794 | -0.05 | -0.29 | 1.952213 | -3 | 1.096703 | 1.096703 | 0.49504 | 0.08191 | 0.66215 | 0.14538 | 10.41667 | 0.051411255 | 0 | Neutral |
| testneg_6795 | -0.02 | -0.13 | 2.75746 | -3 | 1.441342 | -3 | 0.16053 | 0.15049 | 0.11072 | 0.38259 | 20.35286 | 0.257896241 | 0.2 | Neutral |
| testneg_6796 | 0.03 | 0.12 | 1.498975 | -3 | 1.144302 | -3 | 0.27843 | 0.54325 | 0.38942 | 0.345 | 21.38889 | 0.231927609 | 0 | Neutral |
| testneg_6797 | 0 | 0 | 2.064931 | -3 | 4.470743 | -3 | 0.30122 | 0.09165 | 0.49425 | 0.25538 | 14.16667 | 0.231203345 | 0.2 | Neutral |
| testneg_6798 | -0.21 | -0.15 | 0.913651 | -3 | 0.354715 | -3 | 0.10285 | 0.11721 | 0.04615 | 0.12788 | 18.33333 | 0.06 | 0 | Neutral |
| testneg_6799 | 0 | 0 | 1.130115 | -3 | 7.519278 | -3 | 0.82846 | 0.13042 | 0.20379 | 0.35591 | 10.41667 | 0.359745739 | 12.6 | Neutral |
| testneg_6800 | 0 | 0 | -0.2352 | -3 | 3.821143 | -3 | 0.19564 | 0.59275 | 0.08265 | 0.12788 | 17.01711 | 0.144870616 | 0 | Neutral |
| testneg_6801 | 0 | 0 | 2.152857 | -3 | 0.338162 | -0.203227 | 0.339 | 0.13169 | 0.58827 | 0.57942 | 4.72222 | 0.050174505 | 0 | Neutral |
| testneg_6802 | -0.0190909 | 0.62818182 | 7.974744 | -3 | 1.283687 | -3 | 0.087 | 0.12788 | 0.62199 | 0.14032 | 28.20565 | 0.427064732 | 24.8 | Neutral |
| testneg_6803 | -0.02 | -0.45 | -1.184577 | -3 | 3.628039 | -3 | 0.34826 | 0.49454 | 0.055 | 0.20858 | 10 | 0.358262271 | 8.4 | Neutral |
| testneg_6804 | 0 | 0 | 1.582075 | -3 | 5.960756 | -3 | 0.0752 | 0.74467 | 0.2477 | 0.11928 | 16.16034 | 0.240265704 | 0.6 | Neutral |
| testneg_6805 | 0 | 0 | 1.975201 | -3 | 1.170813 | 0.596527 | 0.34495 | 0.16861 | 0.63818 | 0.12788 | 10.41667 | 0.020298507 | 0 | Neutral |
| testneg_6806 | 0 | 0 | -0.342253 | -3 | -0.008971 | -3 | 0.63363 | 0.14795 | 0.6185 | 0.09316 | 10.1325 | 0.01 | 0 | Neutral |
| testneg_6807 | 0.21 | 0.15 | 1.478329 | -3 | 4.890952 | -3 | 0.27614 | 0.12085 | 0.43977 | 0.17284 | 25.99009 | 0.32078216 | 3.4 | Neutral |
| testneg_6808 | 0.04 | -0.23 | 5.478836 | -3 | 2.628269 | -3 | 0.0709 | 0.07843 | 0.18584 | 0.11114 | 19.67873 | 0.16 | 0.2 | Neutral |
| testneg_6809 | 0.01 | 0.79 | 0.97198 | -3 | -0.809518 | -3 | 0.11072 | 0.38259 | 0.47187 | 0.14227 | 10.83333 | 0.16 | 0 | Neutral |
| testneg_6810 | -0.07 | -0.01 | -2.544457 | -3 | 3.256775 | -3 | 0.08454 | 0.22908 | 0.16309 | 0.12788 | 32.22765 | 0.149757576 | 0 | Neutral |
| testneg_6811 | -0.0025714 | -0.1897143 | 2.053049 | -3 | 1.554573 | -3 | 0.26586 | 0.10825 | 0.12446 | 0.08908 | 16.01246 | 0.01 | 0 | Neutral |
| testneg_6812 | 0 | 0 | 5.590034 | -3 | -2.628134 | -3 | 0.24528 | 0.37575 | 0.26971 | 0.08509 | 30.53523 | 0.224825397 | 0 | Neutral |
| testneg_6813 | 0 | 0 | 2.193905 | -3 | 0.631229 | -3 | 0.16952 | 0.10613 | 0.82269 | 0.15732 | 44.33109 | 0.04 | 0 | Neutral |
| testneg_6814 | 0 | 0 | 0.929868 | -3 | 2.745808 | -3 | 0.43459 | 0.12595 | 0.07604 | 0.35375 | 10.41667 | 0.109906292 | 0 | Neutral |
| testneg_6815 | -0.14 | -0.54 | -0.426346 | -3 | 4.204693 | -3 | 0.25077 | 0.06261 | 0.64432 | 0.10409 | 14.27748 | 0.110849673 | 0 | Neutral |
| testneg_6816 | 0.08 | 0.03 | 1.214028 | -3 | 1.836976 | -3 | 0.08136 | 0.10647 | 0.84491 | 0.30945 | 8.612 | 0.061507326 | 0 | Neutral |
| testneg_6817 | -0.08 | -0.03 | 6.140194 | -3 | 0.398341 | 0.398341 | 0.18785 | 0.13588 | 0.10314 | 0.12173 | 12.29167 | 0.23 | 1.2 | Neutral |
| testneg_6818 | -0.15 | 0.28 | 1.817987 | -3 | -0.320232 | -3 | 0.19898 | 0.12788 | 0.23427 | 0.10875 | 27.4024 | 0.04 | 0 | Neutral |
| testneg_6819 | 0 | 0 | 1.845348 | -3 | -0.496432 | -0.496432 | 0.95682 | 0.22744 | 0.06247 | 0.14184 | 4.32527 | 0.057198524 | 0 | Neutral |
| testneg_6820 | 0 | 0 | -0.031285 | -3 | 4.217884 | -3 | 0.64661 | 0.49283 | 0.10058 | 0.13882 | 10.13889 | 0.337296244 | 6 | Neutral |
| testneg_6821 | 0.07 | -0.31 | -1.328162 | -3 | -0.041384 | -3 | 0.33082 | 0.12788 | 0.00451 | 0.2022 | 30.0251 | 0.09672619 | 0 | Neutral |
| testneg_6822 | 0.05 | -0.44 | 2.339071 | -3 | 1.644143 | -3 | 0.08753 | 0.09666 | 0.096 | 0.09851 | 21.06216 | 0.02 | 0 | Neutral |
| testneg_6823 | 0.21 | 0.15 | -0.402147 | -3 | 5.716056 | -3 | 0.0606 | 0.39625 | 0.28817 | 0.10531 | 18.19359 | 0.284285714 | 1.4 | Neutral |
| testneg_6824 | 0.21 | 0.15 | 3.349863 | -3 | 4.245674 | -3 | 0.1472 | 0.12788 | 0.31736 | 0.11947 | 27.17793 | 0.147211134 | 0 | Neutral |
| testneg_6825 | -0.04 | 0.23 | 0.6838 | 0.6838 | 3.025435 | -3 | 0.67327 | 0.23543 | 0.25674 | 0.83834 | 12.29167 | 0.344096845 | 7.6 | Neutral |
| testneg_6826 | 0.06 | -0.57 | 4.397618 | -3 | 0.061657 | -3 | 0.16544 | 0.1313 | 0.07015 | 0.0889 | 22.87402 | 0.04 | 0 | Neutral |
| testneg_6827 | 0 | 0 | 1.156713 | 1.156713 | 6.179448 | -3 | 0.36458 | 0.16397 | 0.17296 | 0.09548 | 18.33333 | 0.222005682 | 0.4 | Neutral |
| testneg_6828 | 0 | 0 | 3.164769 | -3 | 3.730704 | -3 | 0.16929 | 0.12788 | 0.37951 | 0.19435 | 23.57686 | 0.404198625 | 20.6 | Neutral |
| testneg_6829 | 0.02 | -0.63 | 2.81231 | -3 | 4.720694 | -3 | 0.06647 | 0.09129 | 0.12453 | 0.17736 | 11.875 | 0.431140873 | 31 | Neutral |
| testneg_6830 | -0.08 | -0.03 | 0.746424 | -3 | 1.855266 | 0.108451 | 0.21014 | 0.12378 | 0.13601 | 0.19952 | 6.94444 | 0.08011236 | 0 | Neutral |
| testneg_6831 | 0 | 0 | 1.234219 | -3 | 4.714948 | -3 | 0.76942 | 0.13267 | 0.20327 | 0.11174 | 24.15004 | 0.074700079 | 0 | Neutral |

| | | | | | | | | | | | | | | |
|---|---|---|---|---|---|---|---|---|---|---|---|---|---|---|
| testneg_6832 | -0.07 | -0.01 | -1.595702 | -3 | 5.745573 | 5.745573 | 0.82131 | 0.12788 | 0.08027 | 0.12788 | 21.50794 | 0.4 | 15.2 | Neutral |
| testneg_6833 | 0 | 0.73 | 5.650938 | -3 | 3.828961 | -3 | 0.16671 | 0.18311 | 0.06723 | 0.10296 | 18.88889 | 0.448530618 | 36.6 | Neutral |
| testneg_6834 | 0 | 0 | 1.748296 | -3 | 3.866934 | -3 | 0.51278 | 0.35859 | 0.08973 | 0.0992 | 13.82654 | 0.11 | 0 | Neutral |
| testneg_6835 | -0.03 | -0.12 | 1.215025 | 1.215025 | 3.107095 | -3 | 0.09028 | 0.10643 | 0.19898 | 0.40608 | 8.97898 | 0.242154927 | 0.6 | Neutral |
| testneg_6836 | -0.19 | 0.27 | 0.137787 | -3 | 0.775151 | -3 | 0.5471 | 0.11262 | 0.1331 | 0.12788 | 28.49046 | 0.07 | 0 | Neutral |
| testneg_6837 | -0.07 | -0.38 | -0.395207 | -3 | 3.592336 | -3 | 0.2098 | 0.44672 | 0.03297 | 0.16627 | 10.83333 | 0.410205301 | 19.2 | Neutral |
| testneg_6838 | -0.07 | 0.31 | 1.864364 | -3 | 1.576682 | -3 | 0.12943 | 0.14391 | 0.10362 | 0.09649 | 10.41667 | 0.03 | 0 | Neutral |
| testneg_6839 | 0.17 | 0.38 | 4.729585 | -3 | 0.090328 | -3 | 0.29277 | 0.12788 | 0.11144 | 0.08199 | 24.86406 | 0.05 | 0 | Neutral |
| testneg_6840 | 0.02 | -0.68 | 1.871771 | 1.091668 | 1.129332 | -3 | 0.86373 | 0.11969 | 0.37094 | 0.10762 | 29.86111 | 0.04 | 0 | Neutral |
| testneg_6841 | 0 | 0 | -0.32492 | -3 | 3.320732 | 3.320732 | 0.20219 | 0.10986 | 0.19898 | 0.13249 | 13.35742 | 0.239910714 | 0.8 | Neutral |
| testneg_6842 | 0.03 | -1.03 | 5.122912 | -3 | 1.122479 | -3 | 0.44816 | 0.56655 | 0.53654 | 0.22867 | 5.27778 | 0.46125 | 41.8 | Neutral |
| testneg_6843 | 0.07 | -0.31 | 2.817506 | -3 | 0.482695 | -3 | 0.08002 | 0.08792 | 0.60918 | 0.11247 | 29.96032 | 0.020263975 | 0 | Neutral |
| testneg_6844 | 0 | 0 | 2.197785 | -3 | 2.265328 | 2.265328 | 0.39863 | 0.44944 | 0.15685 | 0.17839 | 10.41667 | 0.215310164 | 0.2 | Neutral |
| testneg_6845 | -0.01 | -0.79 | 4.248239 | -3 | 4.695176 | -3 | 0.16061 | 0.12426 | 0.14432 | 0.22579 | 22.82634 | 0.591243686 | 85.4 | Disease_causing |
| testneg_6846 | 0 | 0 | 2.549835 | -3 | 2.16064 | -3 | 0.20673 | 0.13236 | 0.16734 | 0.15176 | 12.40248 | 0.091041532 | 0 | Neutral |
| testneg_6847 | 0 | 0 | 0.383952 | -3 | 3.686515 | -3 | 0.28629 | 0.77234 | 0.77981 | 0.62662 | 4.72222 | 0.313981113 | 5.4 | Neutral |
| testneg_6848 | 0 | -0.01 | 1.660423 | -3 | -0.793282 | -3 | 0.17075 | 0.12788 | 0.05324 | 0.19446 | 20.71879 | 0.07 | 0 | Neutral |
| testneg_6849 | 0 | 0 | 0.976803 | -3 | 7.198996 | -3 | 0.50687 | 0.11283 | 0.12463 | 0.12788 | 10.27677 | 0.230917561 | 2 | Neutral |
| testneg_6850 | 0.22 | -2.66 | 6.380178 | -3 | 5.148137 | 1.021021 | 0.14457 | 0.11614 | 0.09807 | 0.24901 | 17.61599 | 0.690357847 | 94.8 | Disease_causing |
| testneg_6851 | 0.16 | -1.37 | 6.807103 | 6.807103 | 0.374595 | -3 | 0.44612 | 0.24054 | 0.30367 | 0.41964 | 1.11111 | 0.84 | 100 | Disease_causing |
| testneg_6852 | 0.02109091 | -0.0216364 | 9.189407 | -3 | 0.013725 | -3 | 0.64072 | 0.12788 | 0.19898 | 0.12788 | 26.42796 | 0.35 | 7.2 | Neutral |
| testneg_6853 | -0.01 | -0.79 | -3.816017 | -3 | 2.60964 | -3 | 0.06095 | 0.08179 | 0.7062 | 0.83002 | 6.94444 | 0.171098901 | 0 | Neutral |
| testneg_6854 | 0 | 0 | 1.743625 | -3 | -0.187363 | -3 | 0.55527 | 0.12788 | 0.90649 | 0.3543 | 5 | 0.021428571 | 0 | Neutral |
| testneg_6855 | 0 | -0.01 | 0.522695 | -3 | 5.007683 | -3 | 0.84616 | 0.1471 | 0.07415 | 0.12788 | 18.6768 | 0.099035354 | 0 | Neutral |
| testneg_6856 | 0.04 | -0.23 | 1.625356 | -3 | 6.431621 | 6.431621 | 0.5065 | 0.20697 | 0.26676 | 0.76568 | 10.83333 | 0.758752457 | 99.4 | Disease_causing |
| testneg_6857 | 0.06 | -1.29 | 3.503331 | -3 | 2.031708 | -3 | 0.11773 | 0.48758 | 0.13061 | 0.21023 | 21.38889 | 0.515944041 | 62.6 | Disease_causing |
| testneg_6858 | -0.15 | -0.04 | 3.653806 | -3 | -0.196592 | -0.196592 | 0.10371 | 0.08655 | 0.32733 | 0.16869 | 18.76882 | 0.190119048 | 0 | Neutral |
| testneg_6859 | 0.14 | -1.01 | 3.635519 | -3 | 5.011754 | -3 | 0.07995 | 0.12788 | 0.19898 | 0.12788 | 22.7451 | 0.212248665 | 0.6 | Neutral |
| testneg_6860 | 0 | 0 | -0.390566 | -0.390566 | 6.678689 | -3 | 0.08283 | 0.11797 | 0.20765 | 0.27124 | 10.30286 | 0.39797619 | 18 | Neutral |
| testneg_6861 | -0.08 | -0.03 | 1.562114 | -3 | 0.47202 | -3 | 0.78356 | 0.13377 | 0.13295 | 0.12914 | 23.35718 | 0.021517638 | 0 | Neutral |
| testneg_6862 | -0.15 | 1.06 | 1.507517 | -3 | 1.402346 | -3 | 0.19898 | 0.10482 | 0.52299 | 0.12788 | 55.83765 | 0.120197432 | 0 | Neutral |
| testneg_6863 | 0.00442308 | -0.005641 | 2.309347 | -3 | 4.464164 | -3 | 0.80629 | 0.12788 | 0.58994 | 0.11602 | 10.83333 | 0.070353598 | 0 | Neutral |
| testneg_6864 | -0.03 | -0.1 | 0.447373 | -3 | 5.891609 | -3 | 0.37065 | 0.12788 | 0.0959 | 0.12788 | 39.42275 | 0.123904762 | 0 | Neutral |
| testneg_6865 | 0.03 | -1.03 | 3.13682 | -3 | 1.30641 | -3 | 0.29828 | 0.12788 | 0.19898 | 0.12788 | 18.88889 | 0.03 | 0 | Neutral |
| testneg_6866 | 0.15 | 0.06 | 2.541576 | -3 | -0.656987 | -3 | 0.71463 | 0.29732 | 0.16321 | 0.09621 | 18.37823 | 0.06 | 0 | Neutral |
| testneg_6867 | 0.07 | 0.7 | 4.354789 | -3 | -0.360768 | -0.360768 | 0.64319 | 0.12788 | 0.47147 | 0.09503 | 33.40489 | 0.060037037 | 0 | Neutral |
| testneg_6868 | 0 | 0 | -0.02198 | -0.02198 | 2.42657 | -3 | 0.15509 | 0.1326 | 0.38115 | 0.21061 | 18.88889 | 0.130169651 | 0 | Neutral |
| testneg_6869 | 0 | 0 | 0.123596 | -3 | -0.086855 | -3 | 0.53292 | 0.32174 | 0.59827 | 0.11479 | 46.54742 | 0.030954545 | 0 | Neutral |
| testneg_6870 | 0.04 | 2.01 | 0.728618 | -3 | 4.404978 | 0.141193 | 0.10241 | 0.08165 | 0.03505 | 0.0907 | 17.21847 | 0.15 | 0 | Neutral |
| testneg_6871 | 0.01 | 0.79 | 0.457477 | -3 | 4.776572 | -3 | 0.16054 | 0.11683 | 0.16125 | 0.08406 | 7.20975 | 0.081670579 | 0 | Neutral |
| testneg_6872 | 0.14 | -0.37 | 1.075133 | -3 | -0.345708 | -3 | 0.0606 | 0.39625 | 0.32677 | 0.12788 | 19.87245 | 0.11 | 0 | Neutral |
| testneg_6873 | -0.01 | -0.79 | 0.577464 | -3 | 3.803915 | -3 | 0.06366 | 0.12788 | 0.31773 | 0.10816 | 13.7814 | 0.07 | 0 | Neutral |
| testneg_6874 | 0 | 0 | 0.938815 | -3 | 1.089287 | -3 | 0.33292 | 0.17111 | 0.1336 | 0.15669 | 5.47222 | 0.04932598 | 0 | Neutral |
| testneg_6875 | 0 | 0 | 0.844548 | -3 | 4.259881 | -3 | 0.9237 | 0.55859 | 0.38756 | 0.08351 | 6.29554 | 0.077326811 | 0 | Neutral |
| testneg_6876 | 0 | 0 | 0.515441 | -3 | 2.144749 | -3 | 0.2741 | 0.10478 | 0.07071 | 0.11499 | 19.09347 | 0.000685783 | 0 | Neutral |
| testneg_6877 | 0.02 | 0.45 | 4.020676 | -3 | 5.790455 | -3 | 0.44067 | 0.11201 | 0.10895 | 0.10442 | 17.88746 | 0.288758081 | 2.8 | Neutral |
| testneg_6878 | 0 | 0 | 1.178684 | -3 | 6.459281 | -3 | 0.12974 | 0.11886 | 0.47187 | 0.14227 | 18.45565 | 0.30025 | 4.2 | Neutral |
| testneg_6879 | 0.16 | 0.08 | -0.28306 | -3 | 1.75958 | -3 | 0.44067 | 0.11201 | 0.65897 | 0.67602 | 16.42913 | 0.090394345 | 0 | Neutral |
| testneg_6880 | -0.15 | -0.04 | -0.194468 | -3 | 5.977136 | -3 | 0.1273 | 0.12788 | 0.19898 | 0.12788 | 27.91246 | 0.22 | 0 | Neutral |
| testneg_6881 | 0.05 | -0.81 | 3.271976 | -3 | 0.133748 | -3 | 0.48662 | 0.23781 | 0.96475 | 0.27545 | 10.18141 | 0.254414812 | 0.8 | Neutral |
| testneg_6882 | 0 | 0 | 0.276668 | -3 | 0.568842 | -3 | 0.35376 | 0.09478 | 0.12857 | 0.09376 | 15.97222 | 0.001185383 | 0 | Neutral |
| testneg_6883 | -0.01 | 0.57 | 1.661207 | -3 | 0.040053 | -3 | 0.13555 | 0.16663 | 0.18246 | 0.12791 | 20.83333 | 0.04 | 0 | Neutral |
| testneg_6884 | -0.04 | 0.23 | -2.565987 | -3 | 0.307203 | -0.508098 | 0.06072 | 0.09873 | 0.07872 | 0.12788 | 17.21847 | 0.04 | 0 | Neutral |
| testneg_6885 | 0.04 | -0.23 | -0.52815 | -3 | 2.079411 | -3 | 0.16052 | 0.12788 | 0.37292 | 0.11979 | 21.5133 | 0.01 | 0 | Neutral |
| testneg_6886 | 0.02 | -0.63 | 3.558021 | -3 | -0.293928 | -3 | 0.98255 | 0.98129 | 0.103 | 0.403 | 4.44444 | 0.31 | 6 | Neutral |
| testneg_6887 | 0 | -0.01 | 0.967447 | -3 | -0.22016 | -0.22016 | 0.22658 | 0.20646 | 0.17911 | 0.58893 | 32.84553 | 0.162746154 | 0 | Neutral |
| testneg_6888 | 0.08 | 0.03 | 1.795418 | -3 | 2.710438 | -3 | 0.11055 | 0.19829 | 0.19898 | 0.10116 | 14.56463 | 0.116475764 | 0 | Neutral |
| testneg_6889 | 0.21 | 0.15 | 2.53183 | -3 | 0.881051 | -3 | 0.38207 | 0.35052 | 0.48561 | 0.09034 | 11.4406 | 0.08 | 0 | Neutral |
| testneg_6890 | 0.14 | -0.37 | 4.336233 | 4.118971 | 5.0513 | -3 | 0.126 | 0.14076 | 0.19898 | 0.12788 | 25.41667 | 0.55 | 76.4 | Disease_causing |
| testneg_6891 | 0 | -0.32 | 2.774269 | -3 | 6.070697 | -3 | 0.28817 | 0.10531 | 0.08027 | 0.12788 | 22.91169 | 0.203225806 | 0.2 | Neutral |
| testneg_6892 | 0.05 | -0.81 | 2.477154 | -3 | 2.486494 | -3 | 0.09532 | 0.35459 | 0.18142 | 0.21216 | 16.668 | 0.412013889 | 19.8 | Neutral |
| testneg_6893 | 0.07 | 0.7 | 4.037659 | -3 | 6.616703 | -3 | 0.77944 | 0.1858 | 0.13997 | 0.19598 | 6.28072 | 0.748658407 | 100 | Disease_causing |
| testneg_6894 | 0.18 | -1.05 | 6.509691 | -3 | 1.802906 | -3 | 0.26586 | 0.10825 | 0.33886 | 0.15644 | 10.83333 | 0.415955882 | 23 | Neutral |
| testneg_6895 | 0.14 | -1.01 | 6.296617 | -3 | 0.79785 | -3 | 0.54213 | 0.09044 | 0.14462 | 0.11239 | 10.24716 | 0.185833333 | 0.6 | Neutral |
| testneg_6896 | 0 | -0.32 | 0.423935 | 0.423935 | 2.513033 | -3 | 0.31736 | 0.11947 | 0.19898 | 0.12788 | 29.81171 | 0.021277174 | 0 | Neutral |
| testneg_6897 | 0 | 0 | 0.701413 | -3 | 3.063265 | -3 | 0.1214 | 0.0899 | 0.26842 | 0.09683 | 18.03534 | 0.030220081 | 0 | Neutral |
| testneg_6898 | -0.18 | -0.16 | 0.369791 | 0.369791 | 2.550881 | -3 | 0.66227 | 0.11002 | 0.19898 | 0.12788 | 24.52719 | 0.080651261 | 0 | Neutral |
| testneg_6899 | 0 | 0 | 0.39971 | -3 | 0.8653 | -3 | 0.11301 | 0.10716 | 0.06729 | 0.17051 | 28.0719 | 0.08 | 0 | Neutral |
| testneg_6900 | 0 | 0 | 1.840893 | -3 | 1.373367 | -3 | 0.18465 | 0.11262 | 0.34314 | 0.18668 | 11.85525 | 0.02367957 | 0 | Neutral |
| testneg_6901 | 0 | 0 | 0.039637 | -3 | 1.733436 | -3 | 0.20304 | 0.31462 | 0.07241 | 0.12403 | 18.88889 | 0.02 | 0 | Neutral |
| testneg_6902 | -0.06 | -0.18 | -1.695325 | -3 | 3.020834 | -3 | 0.36052 | 0.2858 | 0.19894 | 0.27411 | 5.73099 | 0.268835784 | 1.6 | Neutral |
| testneg_6903 | 0.16 | 0.08 | -0.169987 | -3 | 0.040321 | -3 | 0.99445 | 0.10396 | 0.5958 | 0.0929 | 12.77241 | 0.045 | 0 | Neutral |
| testneg_6904 | -0.06 | -0.18 | 0.622743 | -3 | 3.731145 | -3 | 0.18979 | 0.07948 | 0.18067 | 0.14759 | 11.25 | 0.198615497 | 0 | Neutral |
| testneg_6905 | 0.04 | -0.23 | 7.067026 | -3 | 3.465539 | -3 | 0.0893 | 0.11207 | 0.13139 | 0.11068 | 18.75947 | 0.371142241 | 15.6 | Neutral |
| testneg_6906 | 0.07 | 0.38 | 4.917947 | -3 | 0.070259 | -3 | 0.37463 | 0.23021 | 0.09105 | 0.27434 | 35.90179 | 0.350181818 | 7.8 | Neutral |
| testneg_6907 | 0 | 0 | 0.009894 | -3 | 5.576238 | -2.118566 | 0.07996 | 0.08354 | 0.25159 | 0.12788 | 24.17334 | 0.070367746 | 0 | Neutral |
| testneg_6908 | -0.04 | 0.89 | 6.228417 | -3 | 5.674415 | -3 | 0.25905 | 0.23728 | 0.09871 | 0.09286 | 9.93136 | 0.68 | 96.6 | Disease_causing |
| testneg_6909 | 0.22 | -2.66 | 2.537219 | -3 | 3.805214 | -3 | 0.0328 | 0.0691 | 0.3437 | 0.5169 | 16.84579 | 0.390397284 | 13.4 | Neutral |
| testneg_6910 | 0 | 0 | 0.284069 | -3 | 5.824607 | -3 | 0.1247 | 0.10331 | 0.03391 | 0.09745 | 53.1051 | 0.084194139 | 0 | Neutral |
| testneg_6911 | 0.04 | -0.23 | 3.962157 | -3 | 2.411682 | -3 | 0.06898 | 0.11659 | 0.30813 | 0.19318 | 36.46163 | 0.297839494 | 1.6 | Neutral |
| testneg_6912 | -0.05 | -0.29 | -2.067049 | -3 | -0.140634 | -3 | 0.19199 | 0.16616 | 0.19898 | 0.12788 | 10 | 0.04 | 0 | Neutral |
| testneg_6913 | -0.03 | -0.1 | 3.690501 | -3 | -0.803634 | -3 | 0.83205 | 0.40904 | 0.16392 | 0.12788 | 11.25 | 0.090871212 | 0 | Neutral |
| testneg_6914 | 0 | 0 | -0.064643 | -3 | 4.764548 | -3 | 0.12766 | 0.14626 | 0.16404 | 0.4047 | 10.41667 | 0.334617027 | 9.2 | Neutral |
| testneg_6915 | 0.18 | -1.05 | 6.822565 | -3 | 7.368118 | -3 | 0.41953 | 0.44852 | 0.34412 | 0.08672 | 34.72222 | 0.68 | 99.4 | Disease_causing |
| testneg_6916 | 0.18 | -1.05 | 3.411319 | -3 | 6.423333 | -3 | 0.1664 | 0.10113 | 0.16962 | 0.12788 | 12.29167 | 0.327647521 | 8.2 | Neutral |
| testneg_6917 | -0.01 | 1.44 | -0.27139 | -3 | 3.014014 | -3 | 0.37637 | 0.12788 | 0.88456 | 0.11674 | 10 | 0.15 | 0 | Neutral |
| testneg_6918 | 0.01 | 0.8 | 0.663866 | -3 | 4.338577 | -3 | 0.08283 | 0.11797 | 0.19898 | 0.12788 | 22.51009 | 0.09 | 0 | Neutral |
| testneg_6919 | 0.07 | -0.31 | 3.54624 | -3 | 1.479722 | -3 | 0.05476 | 0.12788 | 0.57564 | 0.12788 | 21.31912 | 0.09 | 0 | Neutral |
| testneg_6920 | 0 | 0 | 0.470493 | -3 | 4.276849 | -3 | 0.42802 | 0.12096 | 0.0778 | 0.0728 | 19.22222 | 0.050955182 | 0 | Neutral |
| testneg_6921 | 0.05 | -0.81 | 6.750746 | -3 | -0.142939 | -3 | 0.10083 | 0.17511 | 0.11254 | 0.19973 | 19.78052 | 0.567075528 | 76.6 | Disease_causing |
| testneg_6922 | -0.06 | -0.18 | 3.18158 | -3 | 1.159527 | -3 | 0.23439 | 0.09253 | 0.85092 | 0.58871 | 18.6768 | 0.14 | 0 | Neutral |
| testneg_6923 | 0.04 | -0.23 | 1.605283 | -3 | 0.282092 | -3 | 0.7257 | 0.64394 | 0.11585 | 0.12788 | 19.95887 | 0.06 | 0 | Neutral |
| testneg_6924 | 0 | -0.32 | -2.859223 | -2.859223 | 6.740904 | -3 | 0.04574 | 0.05824 | 0.28334 | 0.10368 | 30.45428 | 0.193666667 | 0.2 | Neutral |
| testneg_6925 | -0.07 | -0.38 | 0.878873 | 0.878873 | 1.646708 | -3 | 0.62197 | 0.12788 | 0.25987 | 0.31819 | 18.09942 | 0.121042781 | 0 | Neutral |
| testneg_6926 | 0 | 0 | 1.883707 | -3 | 2.330279 | -3 | 0.07427 | 0.09451 | 0.30925 | 0.10335 | 29.82954 | 0.010488095 | 0 | Neutral |
| testneg_6927 | -0.21 | -0.15 | -1.257208 | -3 | -0.206153 | -3 | 0.12951 | 0.08688 | 0.24087 | 0.27985 | 7.64035 | 0.11 | 0 | Neutral |
| testneg_6928 | 0 | 0 | 0.461408 | -3 | 0.026274 | -3 | 0.16797 | 0.11809 | 0.13985 | 0.08349 | 6.27886 | 0 | 0 | Neutral |
| testneg_6929 | 0.08 | 0.03 | -0.129461 | -3 | 0.486089 | -3 | 0.18416 | 0.13292 | 0.10613 | 0.11203 | 18.33333 | 0.03 | 0 | Neutral |
| testneg_6930 | 0.02 | -0.63 | 4.896967 | -3 | 5.947088 | -3 | 0.25516 | 0.12788 | 0.35234 | 0.1303 | 10.36687 | 0.491405655 | 51.4 | Disease_causing |
| testneg_6931 | 0 | -0.01 | 0.666493 | -3 | -0.675678 | -3 | 0.06118 | 0.08724 | 0.22179 | 0.09831 | 13.86082 | 0.01 | 0 | Neutral |
| testneg_6932 | -0.06 | -0.18 | 4.409326 | -3 | -1.073085 | -3 | 0.62708 | 0.24932 | 0.15025 | 0.4872 | 13.26377 | 0.313518223 | 3.6 | Neutral |
| testneg_6933 | 0.04 | -0.23 | 1.605283 | -3 | 5.018221 | -3 | 0.7257 | 0.64394 | 0.26909 | 0.10159 | 32.83798 | 0.18 | 0 | Neutral |
| testneg_6934 | 0.1 | 1.9 | 6.503277 | -3 | 5.111017 | -3 | 0.19898 | 0.37148 | 0.92776 | 0.09677 | 4.44444 | 0.64 | 95.2 | Disease_causing |
| testneg_6935 | 0.04 | -0.23 | 3.570813 | -3 | 1.351912 | -3 | 0.68054 | 0.09208 | 0.18584 | 0.11114 | 19.16362 | 0.04 | 0 | Neutral |
| testneg_6936 | 0.17 | 0.38 | 0.031513 | -3 | -1.214088 | -3 | 0.11608 | 0.09123 | 0.04192 | 0.06306 | 19.09347 | 0.031302521 | 0 | Neutral |
| testneg_6937 | 0.12 | -1.14 | 3.586622 | -3 | 6.026804 | -3 | 0.25555 | 0.12094 | 0.17969 | 0.11473 | 20.83333 | 0.24 | 1.8 | Neutral |
| testneg_6938 | -0.15 | 0.28 | 1.817987 | -3 | 4.336137 | -3 | 0.19898 | 0.12788 | 0.19898 | 0.12788 | 28.38695 | 0.16758788 | 0 | Neutral |
| testneg_6939 | 0 | 0 | 0.930011 | -3 | 2.090134 | -3 | 0.12786 | 0.72066 | 0.08975 | 0.07399 | 20.3247 | 0.070877826 | 0 | Neutral |
| testneg_6940 | 0 | -0.01 | 1.132093 | -3 | 6.019398 | -3 | 0.26372 | 0.12788 | 0.46244 | 0.56748 | 10 | 0.31259036 | 5.6 | Neutral |
| testneg_6941 | 0 | 0 | 1.80113 | -3 | 4.533285 | -3 | 0.2761 | 0.24559 | 0.09642 | 0.09812 | 31.25 | 0.129438185 | 0 | Neutral |
| testneg_6942 | -0.03 | -0.1 | 3.572169 | -3 | 5.289856 | -3 | 0.13565 | 0.46572 | 0.06955 | 0.1748 | 19.48718 | 0.851195477 | 100 | Disease_causing |
| testneg_6943 | 0 | 0 | 1.263979 | -3 | 5.421816 | -3 | 0.14901 | 0.12788 | 0.1254 | 0.08697 | 10.41667 | 0.07 | 0 | Neutral |
| testneg_6944 | 0 | 0 | 0.035553 | -3 | 6.634872 | -3 | 0.15414 | 0.17421 | 0.20112 | 0.17491 | 10.83333 | 0.563007762 | 74.4 | Disease_causing |
| testneg_6945 | 0 | 0 | 0.493256 | -3 | 1.311291 | -3 | 0.13126 | 0.11628 | 0.07577 | 0.09128 | 18.88889 | 8.90E-05 | 0 | Neutral |
| testneg_6946 | 0.03 | 1.22 | 2.244475 | -3 | 2.305765 | -3 | 0.34406 | 0.09894 | 0.11055 | 0.1127 | 25.69121 | 0.06 | 0 | Neutral |
| testneg_6947 | 0.2 | -1.26 | 3.203428 | -3 | 1.770822 | -3 | 0.35325 | 0.12614 | 0.06313 | 0.12788 | 13.73522 | 0.050982143 | 0 | Neutral |
| testneg_6948 | 0.14 | -1.01 | 4.453224 | -3 | 1.681428 | -3 | 0.60703 | 0.12378 | 0.59691 | 0.11049 | 26.25068 | 0.07 | 0 | Neutral |
| testneg_6949 | 0 | 0 | 1.836976 | -3 | 2.151559 | -3 | 0.84491 | 0.30945 | 0.82591 | 0.12207 | 9.79167 | 0.020195076 | 0 | Neutral |
| testneg_6950 | 0.22 | -2.66 | 0.385189 | -3 | -0.783661 | -3 | 0.11533 | 0.12788 | 0.02059 | 0.066 | 11.25 | 0.060178571 | 0 | Neutral |
| testneg_6951 | -0.03 | -0.1 | 3.134404 | -3 | 4.915048 | -3 | 0.57729 | 0.10831 | 0.1679 | 0.24232 | 8.97898 | 0.547717949 | 68 | Disease_causing |
| testneg_6952 | 0 | 0 | 1.80113 | -3 | 2.357542 | -3 | 0.2761 | 0.24559 | 0.40403 | 0.12788 | 13.02635 | 0.041526242 | 0 | Neutral |
| testneg_6953 | -0.07 | -0.38 | 3.802792 | -3 | -0.061002 | -0.061002 | 0.71844 | 0.20204 | 0.35561 | 0.08845 | 22.83164 | 0.07 | 0 | Neutral |

| | | | | | | | | | | | | | | |
|---|---|---|---|---|---|---|---|---|---|---|---|---|---|---|
| testneg_6954 | 0 | -0.32 | 2.199122 | -3 | 4.417441 | -3 | 0.46058 | 0.12903 | 0.11005 | 0.14742 | 9.51452 | 0.315088716 | 4.6 | Neutral |
| testneg_6955 | 0 | 0 | 1.184608 | -3 | 0.452666 | -3 | 0.0138 | 0.06769 | 0.1215 | 0.12788 | 18.88889 | 0.03 | 0 | Neutral |
| testneg_6956 | 0 | 0 | 1.068497 | -3 | 3.817144 | -3 | 0.51278 | 0.35859 | 0.12321 | 0.08084 | 10.24716 | 0.094229963 | 0 | Neutral |
| testneg_6957 | 0 | 0 | 0.703304 | -3 | -0.48043 | -1.108616 | 0.34859 | 0.09799 | 0.27843 | 0.54325 | 20.40047 | 0.065416667 | 0 | Neutral |
| testneg_6958 | 0 | 0 | 0.462587 | -3 | 4.811385 | -3 | 0.26014 | 0.12378 | 0.17364 | 0.18111 | 20.04274 | 0.261829566 | 1 | Neutral |
| testneg_6959 | -0.02 | 0.36 | 0.553539 | -3 | 2.60676 | -3 | 0.14511 | 0.12788 | 0.09439 | 0.07415 | 19.03093 | 0.022 | 0 | Neutral |
| testneg_6960 | 0.02 | 0.13 | 4.422139 | -3 | 1.26926 | -3 | 0.04699 | 0.08149 | 0.38934 | 0.15403 | 11.25 | 0.22 | 0.2 | Neutral |
| testneg_6961 | -0.07 | -0.38 | 2.278399 | -3 | -0.139258 | -3 | 0.26489 | 0.10549 | 0.31025 | 0.09491 | 20.37668 | 0.01 | 0 | Neutral |
| testneg_6962 | 0.04 | 2.01 | -3.135825 | -3 | 2.305189 | -3 | 0.35422 | 0.11185 | 0.505 | 0.12788 | 42.30104 | 0.11 | 0 | Neutral |
| testneg_6963 | -0.17 | -0.38 | 3.567601 | -3 | 4.415736 | -3 | 0.04655 | 0.10225 | 0.11891 | 0.11186 | 15.82969 | 0.223055556 | 0 | Neutral |
| testneg_6964 | 0.15 | 0.06 | -3.376225 | -3 | 0.83713 | -3 | 0.04123 | 0.06896 | 0.06027 | 0.11574 | 10.83333 | 0.06 | 0 | Neutral |
| testneg_6965 | -0.01 | 0.57 | 2.185335 | -3 | 4.822057 | -3 | 0.19898 | 0.12788 | 0.06187 | 0.36607 | 19.37563 | 0.401610842 | 17.4 | Neutral |
| testneg_6966 | 0.03 | -1.03 | 4.715557 | -3 | 6.242245 | -3 | 0.04243 | 0.22343 | 0.10371 | 0.08655 | 16.42913 | 0.427122153 | 27.2 | Neutral |
| testneg_6967 | 0 | 0 | 0.605243 | -3 | 0.233598 | 0.233598 | 0.40803 | 0.47186 | 0.25804 | 0.11876 | 19.04762 | 0.04 | 0 | Neutral |
| testneg_6968 | 0 | 0 | 0.431179 | -3 | 1.623152 | 1.623152 | 0.47751 | 0.2062 | 0.19898 | 0.12788 | 18.33333 | 0.032377825 | 0 | Neutral |
| testneg_6969 | 0 | -0.01 | 1.315543 | -3 | 2.598788 | -3 | 0.28358 | 0.38765 | 0.19898 | 0.10307 | 29.58333 | 0.056769231 | 0 | Neutral |
| testneg_6970 | 0.06 | 0.18 | 2.289276 | -3 | 6.349802 | -3 | 0.32977 | 0.15427 | 0.13749 | 0.17824 | 13.14653 | 0.586659674 | 84.4 | Disease_causing |
| testneg_6971 | 0 | 0 | 5.815666 | -3 | 3.731402 | -3 | 0.34109 | 0.2167 | 0.49831 | 0.29427 | 35.00591 | 0.777 | 100 | Disease_causing |
| testneg_6972 | 0.03 | 0.1 | -1.628421 | -1.628421 | 1.27566 | -3 | 0.54441 | 0.08891 | 0.18535 | 0.12788 | 13.84202 | 0.03 | 0 | Neutral |
| testneg_6973 | 0 | -0.32 | -0.849297 | -0.849297 | -1.859495 | -3 | 0.23691 | 0.1015 | 0.1214 | 0.0899 | 25.55556 | 0.013690476 | 0 | Neutral |
| testneg_6974 | 0 | 0 | 0.535559 | -0.327885 | -2.285677 | -3 | 0.34931 | 0.11906 | 0.08911 | 0.09298 | 12.76316 | 0.000615023 | 0 | Neutral |
| testneg_6975 | -0.05 | -0.29 | 5.743435 | -3 | 3.668266 | -3 | 0.02518 | 0.07165 | 0.63645 | 0.14596 | 21.9697 | 0.41 | 22.2 | Neutral |
| testneg_6976 | 0.09 | -2.17 | 5.014443 | -3 | 1.176476 | -3 | 0.52953 | 0.10241 | 0.64432 | 0.10409 | 10.82179 | 0.124642857 | 0 | Neutral |
| testneg_6977 | 0 | 0 | 0.024873 | -3 | 4.459598 | -3 | 0.16507 | 0.10992 | 0.03505 | 0.0907 | 13.02635 | 0.041308761 | 0 | Neutral |
| testneg_6978 | -0.14 | 0.37 | 0.83713 | -3 | -0.164619 | -3 | 0.06027 | 0.11574 | 0.21774 | 0.10131 | 14.45887 | 0.05 | 0 | Neutral |
| testneg_6979 | 0 | 0 | 5.139254 | -3 | 1.840603 | 1.393176 | 0.19898 | 0.10269 | 0.83421 | 0.18817 | 37.02758 | 0.243405719 | 0.4 | Neutral |
| testneg_6980 | -0.09 | 0.3 | -0.522638 | -3 | 1.232154 | -3 | 0.20604 | 0.10821 | 0.17768 | 0.10832 | 23.78988 | 0.02 | 0 | Neutral |
| testneg_6981 | 0 | 0 | 0.339713 | -3 | 1.06127 | -3 | 0.30739 | 0.16306 | 0.09724 | 0.09172 | 16.38443 | 0.010372917 | 0 | Neutral |
| testneg_6982 | -0.07 | -0.38 | 4.583643 | -3 | 1.490909 | -3 | 0.62708 | 0.24932 | 0.90259 | 0.31502 | 5 | 0.351917561 | 8.4 | Neutral |
| testneg_6983 | -0.07 | -0.7 | 2.693426 | -3 | -1.275641 | -3 | 0.1014 | 0.12788 | 0.05546 | 0.28027 | 20.3247 | 0.160076923 | 0 | Neutral |
| testneg_6984 | 0.01 | -0.57 | 2.700261 | -3 | 0.667746 | -3 | 0.76957 | 0.13588 | 0.42621 | 0.49093 | 10 | 0.130458186 | 0 | Neutral |
| testneg_6985 | -0.02 | 0.36 | -0.710033 | -3 | 1.118815 | -3 | 0.12689 | 0.13376 | 0.13199 | 0.14263 | 12.981 | 0.070365691 | 0 | Neutral |
| testneg_6986 | 0.02 | -0.36 | 2.56113 | -3 | 2.846922 | -3 | 0.06304 | 0.12788 | 0.18081 | 0.1258 | 16.42913 | 0.042047619 | 0 | Neutral |
| testneg_6987 | -0.06 | 0.57 | 1.295804 | -3 | 4.447974 | -3 | 0.64795 | 0.21581 | 0.04869 | 0.05922 | 11.25 | 0.166565657 | 0 | Neutral |
| testneg_6988 | -0.06 | -0.18 | 1.398739 | -3 | 4.84544 | -3 | 0.54113 | 0.11017 | 0.81937 | 0.22945 | 12.11131 | 0.249737531 | 0.6 | Neutral |
| testneg_6989 | -0.08 | -0.03 | 1.223047 | -3 | 3.259536 | -3 | 0.4132 | 0.12788 | 0.36494 | 0.12763 | 34.04445 | 0.040288143 | 0 | Neutral |
| testneg_6990 | -0.16 | -0.08 | 2.243022 | -3 | -0.126067 | -3 | 0.07064 | 0.11699 | 0.19898 | 0.09171 | 23.79024 | 0.07 | 0 | Neutral |
| testneg_6991 | 0.14 | -1.01 | 0.207897 | -3 | -0.194401 | -0.194401 | 0.20616 | 0.29531 | 0.06749 | 0.1776 | 5.97222 | 0.153181818 | 0 | Neutral |
| testneg_6992 | -0.09 | 0.25 | 6.243256 | -3 | 1.787025 | -3 | 0.29804 | 0.12768 | 0.58292 | 0.87725 | 22.33325 | 0.459515086 | 36 | Neutral |
| testneg_6993 | -0.03 | -0.1 | 4.235802 | -3 | 5.752104 | 5.752104 | 0.9989 | 0.8746 | 0.17322 | 0.10311 | 16.85532 | 0.575 | 86 | Disease_causing |
| testneg_6994 | 0.01 | 0.79 | -2.924887 | -3 | 2.242135 | -3 | 0.32181 | 0.07041 | 0.22162 | 0.11997 | 10.08333 | 0.04 | 0 | Neutral |
| testneg_6995 | 0.04 | -0.23 | 3.747465 | -3 | 0.913148 | -3 | 0.08932 | 0.12788 | 0.09515 | 0.07784 | 20.96973 | 0.03 | 0 | Neutral |
| testneg_6996 | 0.03 | -1.03 | 4.550951 | -3 | 2.648476 | -3 | 0.21814 | 0.13855 | 0.99971 | 0.21173 | 12.33727 | 0.400961111 | 20.4 | Neutral |
| testneg_6997 | 0 | 0 | 0.433941 | 0.433941 | 2.565617 | -3 | 0.30475 | 0.21064 | 0.24011 | 0.28559 | 10.7553 | 0.220314049 | 0 | Neutral |
| testneg_6998 | 0 | 0 | 0.035561 | -3 | 8.21249 | -3 | 0.29838 | 0.63658 | 0.65564 | 0.11468 | 10.83333 | 0.390839161 | 18.2 | Neutral |
| testneg_6999 | -0.14 | 1.85 | -0.749376 | -0.749376 | 0.941706 | -3 | 0.51437 | 0.17173 | 0.15294 | 0.12788 | 18.33333 | 0.131166667 | 0 | Neutral |
| testneg_7000 | 0 | -0.01 | 0.357318 | -3 | -0.373351 | -0.373351 | 0.16826 | 0.11262 | 0.25485 | 0.12788 | 20.46346 | 0.000109319 | 0 | Neutral |
| testneg_7001 | 0.07 | 0.7 | 5.311585 | -3 | -0.542932 | -3 | 0.24026 | 0.12788 | 0.06957 | 0.07707 | 23.68839 | 0.1 | 0 | Neutral |
| testneg_7002 | 0 | 0 | 1.349695 | -3 | 5.185161 | 5.185161 | 0.20016 | 0.20159 | 0.52118 | 0.36073 | 9.75077 | 0.581189444 | 80.8 | Disease_causing |
| testneg_7003 | 0 | 0 | 2.934031 | -3 | 2.210178 | 1.0263 | 0.32733 | 0.16869 | 0.08183 | 0.11603 | 10.41667 | 0.080758425 | 0 | Neutral |
| testneg_7004 | 0 | 0 | 2.244975 | -3 | 1.698736 | -3 | 0.92832 | 0.16306 | 0.23962 | 0.10029 | 15.86272 | 0.02 | 0 | Neutral |
| testneg_7005 | 0.08 | 0.03 | 1.454077 | -3 | -2.225286 | -3 | 0.95978 | 0.09493 | 0.85759 | 0.1031 | 28.74542 | 0.05 | 0 | Neutral |
| testneg_7006 | 0 | 0 | 1.962669 | -3 | 1.156713 | 1.156713 | 0.78286 | 0.19831 | 0.36458 | 0.16397 | 11.25 | 0.08109127 | 0 | Neutral |
| testneg_7007 | 0 | 0 | 0.300973 | -3 | 1.30641 | -3 | 0.1589 | 0.17874 | 0.19898 | 0.12788 | 10.41667 | 0.010264678 | 0 | Neutral |
| testneg_7008 | -0.08 | -0.03 | 2.742488 | -3 | 0.586242 | -3 | 0.96995 | 0.66487 | 0.559 | 0.2286 | 8.25691 | 0.196987628 | 0 | Neutral |
| testneg_7009 | 0 | 0 | -0.34106 | -3 | 2.818331 | -3 | 0.61008 | 0.17538 | 0.19898 | 0.12788 | 58.33333 | 0.104013158 | 0 | Neutral |
| testneg_7010 | 0 | 0 | 0.162648 | -3 | 0.979262 | -3 | 0.38002 | 0.22508 | 0.14038 | 0.21177 | 12.74476 | 0.07047928 | 0 | Neutral |
| testneg_7011 | 0.07 | -0.31 | 0.30441 | -3 | 3.540511 | 3.061805 | 0.28817 | 0.10531 | 0.16616 | 0.11073 | 23.9497 | 0.15 | 0 | Neutral |
| testneg_7012 | 0.05 | -0.44 | 2.913025 | -1.278679 | 0.591702 | -3 | 0.04574 | 0.05824 | 0.43283 | 0.23984 | 22.08333 | 0.2 | 0 | Neutral |
| testneg_7013 | 0.22 | -2.66 | 4.250235 | -3 | 0.128902 | -3 | 0.60412 | 0.13303 | 0.81384 | 0.12094 | 18.8735 | 0.121558704 | 0 | Neutral |
| testneg_7014 | -0.01 | -0.79 | 0.577464 | -3 | -0.120714 | -3 | 0.06366 | 0.12788 | 0.99665 | 0.12884 | 13.83267 | 0.06 | 0 | Neutral |
| testneg_7015 | 0.02 | 0.45 | 0.379358 | -3 | -0.493225 | -3 | 0.19898 | 0.10996 | 0.09235 | 0.18135 | 16.8018 | 0.060370879 | 0 | Neutral |
| testneg_7016 | -0.21 | -0.15 | 3.66939 | -3 | -0.239452 | -3 | 0.66422 | 0.99377 | 0.42899 | 0.31045 | 5.69444 | 0.381201923 | 10.4 | Neutral |
| testneg_7017 | 0.08 | 0.03 | -1.242275 | -3 | 5.262041 | -3 | 0.7597 | 0.12788 | 0.3667 | 0.2135 | 10.41667 | 0.270918651 | 0.4 | Neutral |
| testneg_7018 | -0.1 | -1.9 | 0.700131 | -3 | 1.085971 | -3 | 0.46056 | 0.12785 | 0.14941 | 0.0838 | 26.66667 | 0.05 | 0 | Neutral |
| testneg_7019 | 0.1 | 1.9 | 0.416226 | -3 | 2.81231 | -3 | 0.32111 | 0.12788 | 0.06647 | 0.09129 | 24.17344 | 0.12 | 0 | Neutral |
| testneg_7020 | -0.05 | -1.52 | 3.546575 | -3 | 0.72503 | -3 | 0.19024 | 0.1132 | 0.56862 | 0.12788 | 23.70509 | 0.059259259 | 0 | Neutral |
| testneg_7021 | 0.05 | 0.29 | 0.979063 | -3 | 0.389118 | -3 | 0.39111 | 0.15887 | 0.19898 | 0.0522 | 13.51351 | 0.02 | 0 | Neutral |
| testneg_7022 | 0.22 | -2.66 | 6.380178 | -3 | -0.195907 | -3 | 0.14457 | 0.11614 | 0.3221 | 0.3367 | 6.98099 | 0.399666667 | 23.4 | Neutral |
| testneg_7023 | 0 | 0 | 0.842285 | -3 | 2.61063 | -3 | 0.10703 | 0.11611 | 0.04627 | 0.05223 | 33.60688 | 0.02 | 0 | Neutral |
| testneg_7024 | 0.18 | -1.05 | 6.294594 | -3 | 1.15899 | -3 | 0.21033 | 0.35953 | 0.17969 | 0.11473 | 12.05846 | 0.305722188 | 5.2 | Neutral |
| testneg_7025 | 0 | 0 | 0.419995 | -3 | 3.710256 | -3 | 0.7174 | 0.15815 | 0.89661 | 0.17382 | 11.25 | 0.220729085 | 0.2 | Neutral |
| testneg_7026 | -0.03 | -0.1 | -1.42588 | -3 | 4.009105 | -3 | 0.16469 | 0.16042 | 0.55384 | 0.15273 | 17.77778 | 0.421585539 | 25.6 | Neutral |
| testneg_7027 | 0.02 | 0.45 | 3.50895 | -3 | 3.167319 | -3 | 0.28165 | 0.15912 | 0.25195 | 0.09584 | 10.28409 | 0.203542989 | 0 | Neutral |
| testneg_7028 | 0.02 | -0.36 | 7.429465 | -3 | 0.635023 | -0.861865 | 0.07098 | 0.39918 | 0.12136 | 0.27707 | 39.04612 | 0.688934394 | 99.2 | Disease_causing |
| testneg_7029 | 0 | 0 | 1.132066 | -3 | 5.297491 | -3 | 0.99317 | 0.60884 | 0.19898 | 0.12788 | 13.3074 | 0.138527056 | 0 | Neutral |
| testneg_7030 | 0.01 | 0.8 | 0.590696 | -3 | 10.754173 | -3 | 0.10809 | 0.09057 | 0.09288 | 0.16617 | 13.02635 | 0.454225772 | 34.6 | Neutral |
| testneg_7031 | -0.19 | 0.27 | 1.971396 | -3 | 5.714497 | -3 | 0.46837 | 0.14896 | 0.098 | 0.08791 | 26.84685 | 0.270719697 | 0.8 | Neutral |
| testneg_7032 | 0 | 0 | 0.916236 | -3 | 3.586622 | -3 | 0.37499 | 0.17803 | 0.25555 | 0.12094 | 10.41667 | 0.067743305 | 0 | Neutral |
| testneg_7033 | 0 | 0 | 0.684399 | -3 | -0.61032 | -3 | 0.13029 | 0.12788 | 0.09522 | 0.20323 | 8.97898 | 0.040277992 | 0 | Neutral |
| testneg_7034 | -0.07 | -0.38 | 0.490574 | -3 | 2.980687 | -3 | 0.07701 | 0.19824 | 0.74558 | 0.10643 | 10.30286 | 0.12875 | 0 | Neutral |
| testneg_7035 | -0.07 | -0.38 | 2.28341 | -3 | 3.655665 | -3 | 0.27723 | 0.13229 | 0.06048 | 0.12788 | 39.3387 | 0.1 | 0 | Neutral |
| testneg_7036 | 0 | 0 | 0.844548 | -3 | 1.464574 | 1.464574 | 0.9237 | 0.55859 | 0.60603 | 0.12788 | 10.41667 | 0.04 | 0 | Neutral |
| testneg_7037 | 0.01 | 0.79 | 0.570058 | -3 | 6.059259 | -3 | 0.08922 | 0.10539 | 0.19898 | 0.12788 | 11.22416 | 0.150401242 | 0 | Neutral |
| testneg_7038 | 0 | 0 | 0.002341 | -3 | 1.273853 | -3 | 0.10799 | 0.10754 | 0.5043 | 0.3225 | 17.21847 | 0.040983923 | 0 | Neutral |
| testneg_7039 | -0.15 | -0.04 | 0.939929 | -3 | 3.542118 | -3 | 0.18701 | 0.12788 | 0.11972 | 0.09438 | 19.67965 | 0.101547619 | 0 | Neutral |
| testneg_7040 | 0 | 0 | 1.966897 | -3 | -0.184962 | -3 | 0.39262 | 0.12788 | 0.29523 | 0.27623 | 10.41667 | 0.030068493 | 0 | Neutral |
| testneg_7041 | -0.04 | 0.23 | 0.421557 | 0.421557 | 0.381493 | 0.381493 | 0.82507 | 0.19615 | 0.14432 | 0.16919 | 12.23253 | 0.130915479 | 0 | Neutral |
| testneg_7042 | 0.07 | -0.31 | 6.107142 | -3 | -0.056564 | -3 | 0.12554 | 0.10854 | 0.01642 | 0.12788 | 12.52798 | 0.128302564 | 0 | Neutral |
| testneg_7043 | 0 | 0 | 1.624378 | -3 | -0.176245 | -0.176245 | 0.2016 | 0.13956 | 0.20849 | 0.12788 | 32.29167 | 0.01 | 0 | Neutral |
| testneg_7044 | 0.14 | -0.37 | 2.102067 | -3 | 1.06262 | -3 | 0.95557 | 0.38833 | 0.2218 | 0.11533 | 25.34535 | 0.067175595 | 0 | Neutral |
| testneg_7045 | 0.06 | -0.57 | 1.240379 | -3 | 4.249749 | -3 | 0.64591 | 0.38907 | 0.0538 | 0.23912 | 17.77778 | 0.46726512 | 41 | Neutral |
| testneg_7046 | -0.03 | -0.1 | 0.618391 | -3 | 0.782249 | -3 | 0.32022 | 0.40347 | 0.5127 | 0.12263 | 13.17982 | 0.04 | 0 | Neutral |
| testneg_7047 | 0.03 | 0.1 | 1.551412 | -3 | -0.190662 | -3 | 0.25905 | 0.23728 | 0.14341 | 0.10134 | 13.72024 | 0.029375 | 0 | Neutral |
| testneg_7048 | 0 | 0 | 1.829958 | 1.829958 | 2.459115 | -3 | 0.47775 | 0.10103 | 0.3038 | 0.11278 | 27.95796 | 0.07 | 0 | Neutral |
| testneg_7049 | -0.04 | 0.23 | -0.274997 | -3 | -0.691406 | -0.691406 | 0.37501 | 0.0997 | 0.04046 | 0.06599 | 11.25 | 0.02 | 0 | Neutral |
| testneg_7050 | -0.03 | -0.12 | 1.324962 | -3 | 0.583408 | 0.583408 | 0.38184 | 0.10478 | 0.05386 | 0.09802 | 21.94444 | 0.02 | 0 | Neutral |
| testneg_7051 | 0 | 0 | 1.005512 | -3 | 3.312865 | -3 | 0.56176 | 0.11451 | 0.16345 | 0.11716 | 20.94356 | 0.053233553 | 0 | Neutral |
| testneg_7052 | 0.09 | -0.67 | -0.874528 | -3 | 1.307704 | -3 | 0.1592 | 0.12788 | 0.64749 | 0.14763 | 36.02896 | 0.053641667 | 0 | Neutral |
| testneg_7053 | -0.07 | -0.38 | 3.479092 | -3 | 1.165033 | -3 | 0.22113 | 0.10673 | 0.86513 | 0.31355 | 21.02339 | 0.169214927 | 0 | Neutral |
| testneg_7054 | 0 | 0 | 0.068014 | 0.068014 | 6.281692 | -3 | 0.20869 | 0.13493 | 0.56013 | 0.29233 | 22.15808 | 0.414553203 | 23.4 | Neutral |
| testneg_7055 | 0 | -0.01 | -0.752438 | -3 | 1.764776 | -3 | 0.10486 | 0.08083 | 0.6975 | 0.53794 | 10.22898 | 0.04 | 0 | Neutral |
| testneg_7056 | 0 | 0 | -0.048245 | -3 | 5.193514 | -3 | 0.15055 | 0.22097 | 0.57188 | 0.34038 | 13.03942 | 0.46157197 | 40.2 | Neutral |
| testneg_7057 | 0.15 | -0.28 | -1.644002 | -3 | 1.824706 | -3 | 0.05996 | 0.12788 | 0.90955 | 0.10096 | 60.3834 | 0.09 | 0 | Neutral |
| testneg_7058 | -0.1 | -1.9 | -1.487847 | -3 | 0.994999 | 0.994999 | 0.06728 | 0.20664 | 0.51282 | 0.12075 | 28.64035 | 0.120183081 | 0 | Neutral |
| testneg_7059 | 0 | -0.01 | 0.505705 | -3 | -0.163107 | -3 | 0.73395 | 0.1705 | 0.21289 | 0.08709 | 20.22479 | 0.010459327 | 0 | Neutral |
| testneg_7060 | -0.06 | -0.18 | 5.5303 | -3 | 1.555382 | -3 | 0.07467 | 0.09224 | 0.80134 | 0.08685 | 19.44444 | 0.150045872 | 0 | Neutral |
| testneg_7061 | -0.08 | -0.03 | 1.53322 | -3 | 1.217652 | -3 | 0.78537 | 0.33817 | 0.19898 | 0.12788 | 20.13042 | 0.04 | 0 | Neutral |
| testneg_7062 | 0 | 0 | -0.161926 | -3 | 4.745256 | -0.771274 | 0.42033 | 0.17821 | 0.09306 | 0.12788 | 10 | 0.112732143 | 0 | Neutral |
| testneg_7063 | 0.18 | -1.05 | 4.901007 | -3 | 0.292982 | -3 | 0.227 | 0.14389 | 0.15677 | 0.18537 | 17.93275 | 0.279269897 | 2.2 | Neutral |
| testneg_7064 | 0 | 0 | 0.059973 | -3 | 0.415836 | -3 | 0.21095 | 0.10574 | 0.15446 | 0.08618 | 18.6768 | 0 | 0 | Neutral |
| testneg_7065 | 0 | 0 | 0.402508 | 0.402508 | -0.691406 | -3 | 0.29794 | 0.11106 | 0.04046 | 0.06599 | 12.29167 | 0.01 | 0 | Neutral |
| testneg_7066 | 0.03 | -1.03 | 2.763749 | -3 | 4.708046 | 4.708046 | 0.13203 | 0.12092 | 0.14731 | 0.12788 | 29.45989 | 0.370944444 | 8.4 | Neutral |
| testneg_7067 | 0 | -0.32 | 3.968486 | -3 | 0.892974 | 0.892974 | 0.05996 | 0.12788 | 0.06986 | 0.12788 | 65.26429 | 0.1 | 0 | Neutral |
| testneg_7068 | 0.14 | -1.01 | -0.306073 | -3 | -0.673818 | -3 | 0.54211 | 0.12592 | 0.76138 | 0.16258 | 13.87133 | 0.08 | 0 | Neutral |
| testneg_7069 | 0 | 0 | 2.036514 | -3 | 0.297691 | -3 | 0.16544 | 0.1313 | 0.07275 | 0.09562 | 21.94798 | 0.02 | 0 | Neutral |
| testneg_7070 | 0 | 0 | -0.034419 | -0.034419 | 0.236592 | -3 | 0.62203 | 0.12788 | 0.40234 | 0.10353 | 9.8924 | 0.01 | 0 | Neutral |
| testneg_7071 | 0 | 0 | 0.543131 | 0.543131 | 6.254102 | -3 | 0.1331 | 0.12788 | 0.1417 | 0.1398 | 18.86584 | 0.382062212 | 14.6 | Neutral |
| testneg_7072 | 0.06 | -0.57 | 6.491948 | -3 | 1.07442 | -3 | 0.19898 | 0.12788 | 0.25422 | 0.10102 | 11.66667 | 0.24016129 | 1.6 | Neutral |
| testneg_7073 | 0 | 0 | 0.191318 | -3 | 1.454836 | -3 | 0.96113 | 0.24974 | 0.07098 | 0.39918 | 10.41667 | 0.09172734 | 0 | Neutral |
| testneg_7074 | 0.01 | 0.8 | 3.769272 | -3 | 1.524427 | -3 | 0.32022 | 0.40347 | 0.99317 | 0.60884 | 4.44444 | 0.334273343 | 8 | Neutral |
| testneg_7075 | 0.01 | 0.8 | -0.214756 | -3 | 1.502207 | -3 | 0.07428 | 0.07645 | 0.4094 | 0.1944 | 14.89369 | 0.050566239 | 0 | Neutral |

| | | | | | | | | | | | | | | |
|---|---|---|---|---|---|---|---|---|---|---|---|---|---|---|
| testneg_7076 | 0.14 | -0.37 | 3.199571 | -3 | 1.413393 | -3 | 0.30474 | 0.12788 | 0.2293 | 0.12788 | 13.51351 | 0.02 | 0 | Neutral |
| testneg_7077 | 0 | -0.01 | 0.611606 | -3 | -0.23732 | -0.23732 | 0.9149 | 0.27793 | 0.07473 | 0.51812 | 10 | 0.100842057 | 0 | Neutral |
| testneg_7078 | -0.02 | 0.68 | 4.06223 | -3 | 0.134977 | -3 | 0.80034 | 0.10518 | 0.0925 | 0.08222 | 18.6768 | 0.061160714 | 0 | Neutral |
| testneg_7079 | 0 | 0 | -0.153757 | -3 | 3.470779 | -3 | 0.23027 | 0.37171 | 0.36694 | 0.12788 | 17.9643 | 0.1 | 0 | Neutral |
| testneg_7080 | 0.04 | 2.01 | -3.135825 | -3 | 1.739278 | -3 | 0.35422 | 0.11185 | 0.17199 | 0.10586 | 30.78527 | 0.09 | 0 | Neutral |
| testneg_7081 | 0 | 0 | -0.072696 | -0.072696 | -0.02028 | -3 | 0.30326 | 0.21562 | 0.50388 | 0.12295 | 8.7796 | 0.01033908 | 0 | Neutral |
| testneg_7082 | -0.01 | 0.57 | -2.581564 | -3 | -1.722922 | -3 | 0.19898 | 0.12788 | 0.06704 | 0.1068 | 23.68839 | 0.03 | 0 | Neutral |
| testneg_7083 | -0.06 | 0.57 | 1.295804 | -3 | 5.210516 | -3 | 0.64795 | 0.21581 | 0.13173 | 0.12788 | 13.05556 | 0.212022727 | 0.2 | Neutral |
| testneg_7084 | -0.01 | -0.8 | 2.607371 | -3 | 0.969432 | -3 | 0.53239 | 0.17509 | 0.18431 | 0.13863 | 10.72072 | 0.12005291 | 0 | Neutral |
| testneg_7085 | 0 | 0 | 0.870324 | -3 | -0.111883 | -0.111883 | 0.16355 | 0.12788 | 0.06313 | 0.12788 | 21.99535 | 0.01 | 0 | Neutral |
| testneg_7086 | 0.01 | -1.44 | 5.580166 | -3 | 0.760948 | -3 | 0.07681 | 0.22865 | 0.56416 | 0.12788 | 12.24613 | 0.28 | 1.2 | Neutral |
| testneg_7087 | 0 | 0 | -1.008848 | -3 | 1.581905 | -3 | 0.38873 | 0.11686 | 0.6271 | 0.09663 | 12.76316 | 0.001213235 | 0 | Neutral |
| testneg_7088 | 0.07 | 0.38 | 1.426694 | -3 | 1.70493 | -3 | 0.88914 | 0.39065 | 0.72068 | 0.99841 | 4.44444 | 0.160386905 | 0 | Neutral |
| testneg_7089 | 0 | 0 | 1.613322 | -3 | -0.077532 | -3 | 0.39132 | 0.11179 | 0.08333 | 0.10006 | 4.72222 | 0.000177116 | 0 | Neutral |
| testneg_7090 | -0.16 | -0.08 | 3.904599 | -3 | 5.481335 | -3 | 0.07427 | 0.09451 | 0.7482 | 0.18812 | 10.83333 | 0.592144265 | 85.8 | Disease_causing |
| testneg_7091 | 0 | 0 | 1.841032 | -3 | -0.550057 | -3 | 0.40663 | 0.24974 | 0.32416 | 0.09959 | 9.74181 | 0.01 | 0 | Neutral |
| testneg_7092 | 0 | 0 | 0.200266 | -3 | 0.76545 | -3 | 0.13447 | 0.12788 | 0.20986 | 0.10633 | 16.48772 | 0 | 0 | Neutral |
| testneg_7093 | -0.03 | 1.03 | 3.933884 | -3 | -0.272804 | -3 | 0.08684 | 0.08917 | 0.17839 | 0.12009 | 13.02635 | 0.09 | 0 | Neutral |
| testneg_7094 | 0 | 0 | -0.390566 | -3 | 6.861342 | -3 | 0.08283 | 0.11797 | 0.55384 | 0.15273 | 22.96784 | 0.35 | 8 | Neutral |
| testneg_7095 | -0.15 | -0.04 | 2.692574 | -3 | 2.12021 | 2.12021 | 0.45628 | 0.15819 | 0.07698 | 0.12788 | 19.09347 | 0.1328125 | 0 | Neutral |
| testneg_7096 | 0.07 | 0.7 | -1.532885 | -3 | 3.876605 | -3 | 0.03844 | 0.12788 | 0.83732 | 0.19325 | 33.04681 | 0.271130952 | 0 | Neutral |
| testneg_7097 | -0.1 | -1.9 | 1.709952 | -3 | -0.020693 | -3 | 0.4172 | 0.11643 | 0.80034 | 0.10518 | 16.8018 | 0.04 | 0 | Neutral |
| testneg_7098 | -0.16 | -0.08 | 3.343953 | -3 | -0.165443 | -3 | 0.18355 | 0.11262 | 0.05221 | 0.12788 | 18.89409 | 0.07 | 0 | Neutral |
| testneg_7099 | -0.05 | -0.29 | 2.605779 | -3 | -0.473056 | -3 | 0.22817 | 0.10634 | 0.15486 | 0.28768 | 13.9343 | 0.09 | 0 | Neutral |
| testneg_7100 | 0 | 0 | 0.063311 | -3 | 2.090818 | -3 | 0.17404 | 0.12788 | 0.12974 | 0.11886 | 19.8374 | 0.000421211 | 0 | Neutral |
| testneg_7101 | 0.03 | -0.85 | 2.681264 | -3 | 0.583716 | -3 | 0.25961 | 0.07098 | 0.82014 | 0.14054 | 17.43377 | 0.041833333 | 0 | Neutral |
| testneg_7102 | 0 | 0 | 2.634633 | -3 | 1.687535 | -3 | 0.40491 | 0.17596 | 0.469 | 0.25588 | 10.41667 | 0.138029879 | 0 | Neutral |
| testneg_7103 | 0.04 | -0.23 | 2.067953 | -3 | 0.278169 | -3 | 0.6323 | 0.09928 | 0.10963 | 0.12788 | 18.6768 | 0.01 | 0 | Neutral |
| testneg_7104 | -0.07 | -0.38 | -0.094721 | -3 | 2.044018 | -3 | 0.16778 | 0.82322 | 0.63431 | 0.09837 | 12.34649 | 0.093333333 | 0 | Neutral |
| testneg_7105 | 0 | 0 | 0.198837 | -3 | -0.715777 | -3 | 0.99704 | 0.17463 | 0.19898 | 0.12788 | 8.97898 | 0.030884033 | 0 | Neutral |
| testneg_7106 | -0.12 | 1.14 | 4.144132 | -3 | 2.9994 | -3 | 0.18224 | 0.10419 | 0.40403 | 0.12788 | 16.01246 | 0.239583333 | 0.6 | Neutral |
| testneg_7107 | -0.02 | 0.63 | 3.679445 | -3 | -0.833975 | -3 | 0.17075 | 0.12788 | 0.20849 | 0.12788 | 35.44691 | 0.030333211 | 0 | Neutral |
| testneg_7108 | 0 | 0 | 0.717822 | -3 | 2.266867 | -3 | 0.29044 | 0.12142 | 0.49064 | 0.12294 | 16.42913 | 0.000224719 | 0 | Neutral |
| testneg_7109 | -0.08 | -0.03 | 1.755955 | -3 | 1.436098 | -3 | 0.9832 | 0.39904 | 0.45628 | 0.15819 | 19.44444 | 0.1582156 | 0 | Neutral |
| testneg_7110 | 0 | 0 | 1.354498 | -3 | 8.005214 | -3 | 0.33713 | 0.12788 | 0.05754 | 0.10881 | 10.22898 | 0.301305556 | 3 | Neutral |
| testneg_7111 | 0 | 0 | 0.094005 | -3 | -0.657349 | -3 | 0.64302 | 0.11262 | 0.14704 | 0.12788 | 12.46106 | 0.000488095 | 0 | Neutral |
| testneg_7112 | 0 | 0 | 1.550971 | -3 | 3.318431 | -3 | 0.73311 | 0.10866 | 0.57578 | 0.35982 | 10.41667 | 0.119618714 | 0 | Neutral |
| testneg_7113 | 0.07 | 0.38 | 1.157706 | -3 | 0.282453 | -3 | 0.09951 | 0.14299 | 0.42366 | 0.14372 | 28.77147 | 0.0725 | 0 | Neutral |
| testneg_7114 | 0 | 0 | 0.58387 | -3 | 7.264005 | -3 | 0.13244 | 0.14875 | 0.1856 | 0.12788 | 12.49864 | 0.373859649 | 14 | Neutral |
| testneg_7115 | 0.03 | -1.03 | 6.929425 | -3 | 5.394292 | -3 | 0.19301 | 0.10828 | 0.2442 | 0.11401 | 8.97898 | 0.549734848 | 73 | Disease_causing |
| testneg_7116 | 0.08 | 0.03 | -1.290562 | -3 | 1.59057 | -3 | 0.19898 | 0.12895 | 0.26789 | 0.67432 | 18.88889 | 0.093333333 | 0 | Neutral |
| testneg_7117 | -0.02 | 0.63 | 3.995565 | -3 | 5.588603 | 5.588603 | 0.31773 | 0.10816 | 0.12782 | 0.08502 | 13.79256 | 0.49 | 50.6 | Disease_causing |
| testneg_7118 | 0.09 | -0.25 | 3.985725 | -3 | 0.723398 | 0.723398 | 0.72946 | 0.28603 | 0.19898 | 0.14957 | 22.11934 | 0.240573593 | 0.4 | Neutral |
| testneg_7119 | 0.16 | -0.37 | 3.291397 | -3 | 6.185375 | -3 | 0.46387 | 0.17039 | 0.27629 | 0.11166 | 17.08466 | 0.363111111 | 15.4 | Neutral |
| testneg_7120 | 0 | 0 | 1.690602 | -3 | 1.394643 | -3 | 0.18206 | 0.10727 | 0.49629 | 0.11989 | 27.68982 | 0.000281746 | 0 | Neutral |
| testneg_7121 | -0.04 | 0.23 | 4.21286 | -3 | 0.116519 | 0.116519 | 0.07447 | 0.23492 | 0.2207 | 0.09998 | 18.33333 | 0.111269841 | 0 | Neutral |
| testneg_7122 | 0 | -0.01 | 1.62909 | -3 | 1.259454 | -2.069741 | 0.25447 | 0.12788 | 0.16259 | 0.102 | 38.05441 | 0.00051914 | 0 | Neutral |
| testneg_7123 | -0.03 | 0.85 | 5.601904 | -3 | 0.513126 | -3 | 0.16728 | 0.11868 | 0.50592 | 0.0952 | 10.83333 | 0.17 | 0.2 | Neutral |
| testneg_7124 | 0 | 0 | 0.38184 | -3 | 0.068865 | -3 | 0.35421 | 0.10694 | 0.35026 | 0.09817 | 18.70817 | 0 | 0 | Neutral |
| testneg_7125 | 0 | 0 | 0.546525 | -3 | 3.810722 | -3 | 0.12545 | 0.08963 | 0.22162 | 0.09445 | 29.16503 | 0.05 | 0 | Neutral |
| testneg_7126 | -0.06 | -0.18 | -0.275748 | -3 | -2.421511 | -3 | 0.13779 | 0.0945 | 0.37464 | 0.12788 | 13.90046 | 0.01 | 0 | Neutral |
| testneg_7127 | -0.07 | -0.38 | 3.500401 | -3 | 1.913341 | -3 | 0.1196 | 0.12788 | 0.67549 | 0.1273 | 26.9055 | 0.062910053 | 0 | Neutral |
| testneg_7128 | -0.12 | 1.16 | 5.480722 | -3 | 5.57053 | -3 | 0.8611 | 0.12788 | 0.73947 | 0.2058 | 18.07234 | 0.665351674 | 96.2 | Disease_causing |
| testneg_7129 | 0 | -0.01 | 2.002215 | -3 | 1.872697 | -3 | 0.95682 | 0.22744 | 0.84468 | 0.55945 | 18.49155 | 0.094110009 | 0 | Neutral |
| testneg_7130 | 0.17 | 0.38 | 6.29487 | -3 | 1.795872 | -3 | 0.18416 | 0.13292 | 0.22155 | 0.1062 | 13.18841 | 0.22 | 1.2 | Neutral |
| testneg_7131 | -0.06 | -0.18 | 3.494072 | -3 | -1.721889 | -3 | 0.2118 | 0.2009 | 0.13615 | 0.12788 | 19.92312 | 0.064575758 | 0 | Neutral |
| testneg_7132 | -0.15 | -0.04 | 4.034471 | -3 | -0.129005 | -3 | 0.14457 | 0.11614 | 0.17226 | 0.13001 | 13.85043 | 0.080801282 | 0 | Neutral |
| testneg_7133 | 0.03 | 0.1 | 4.110265 | -3 | 2.165083 | -3 | 0.17818 | 0.15728 | 0.13141 | 0.1123 | 24.65514 | 0.1 | 0 | Neutral |
| testneg_7134 | 0.02 | 0.45 | -2.376371 | -3 | 3.399799 | -3 | 0.05886 | 0.38242 | 0.13199 | 0.14263 | 8.73265 | 0.416488466 | 22.2 | Neutral |
| testneg_7135 | -0.18 | -0.16 | 6.783337 | -3 | 4.54141 | -3 | 0.1109 | 0.17295 | 0.20431 | 0.17304 | 33.01939 | 0.778428571 | 99.8 | Disease_causing |
| testneg_7136 | 0 | -0.01 | 0.242595 | -3 | -0.491414 | -3 | 0.11358 | 0.12788 | 0.61027 | 0.14229 | 17.20608 | 0.03 | 0 | Neutral |
| testneg_7137 | 0 | 0 | -0.044681 | -2.016047 | 0.527426 | -3 | 0.31584 | 0.11996 | 0.11687 | 0.09069 | 24.83405 | 0.01 | 0 | Neutral |
| testneg_7138 | -0.08 | -0.03 | 2.557841 | -3 | -0.558848 | -0.558848 | 0.19898 | 0.14896 | 0.06611 | 0.08834 | 13.27141 | 0.03 | 0 | Neutral |
| testneg_7139 | 0.01 | 0.8 | 6.169752 | -3 | 2.242726 | -3 | 0.08836 | 0.0911 | 0.07275 | 0.09562 | 21.10735 | 0.219722222 | 0.8 | Neutral |
| testneg_7140 | 0.06 | 0.18 | 2.16294 | -3 | 1.110561 | -3 | 0.10176 | 0.12788 | 0.37109 | 0.12788 | 13.51351 | 0.01 | 0 | Neutral |
| testneg_7141 | -0.03 | -1.22 | 0.837971 | -3 | 5.448182 | -3 | 0.13356 | 0.09963 | 0.48688 | 0.13071 | 13.51351 | 0.161646628 | 0 | Neutral |
| testneg_7142 | -0.02 | 0.63 | 1.096737 | 1.096737 | 1.070904 | -3 | 0.23071 | 0.09582 | 0.40361 | 0.17821 | 13.75 | 0.110455487 | 0 | Neutral |
| testneg_7143 | 0 | 0 | 0.606984 | -3 | -1.230621 | -3 | 0.08932 | 0.12788 | 0.64254 | 0.10729 | 11.92982 | 0.01 | 0 | Neutral |
| testneg_7144 | 0.02763636 | 0.20309091 | 12.965745 | -3 | 0.467459 | -3 | 0.12596 | 0.18924 | 0.20914 | 0.10089 | 25.4782 | 0.41 | 20.2 | Neutral |
| testneg_7145 | 0 | 0 | 0.755726 | -3 | 0.363292 | -3 | 0.08454 | 0.22908 | 0.21884 | 0.16743 | 11.875 | 0.091707283 | 0 | Neutral |
| testneg_7146 | 0 | -0.32 | 4.643838 | -3 | 0.270884 | 0.270884 | 0.835 | 0.13588 | 0.24536 | 0.38555 | 9.848 | 0.272606061 | 0.4 | Neutral |
| testneg_7147 | 0.14 | -0.69 | 3.53144 | 0.866429 | 1.350549 | -3 | 0.45405 | 0.10896 | 0.06296 | 0.09823 | 18.88889 | 0.1 | 0 | Neutral |
| testneg_7148 | 0.03 | -1.03 | 5.608869 | -3 | 5.29555 | -3 | 0.17806 | 0.12788 | 0.70872 | 0.09884 | 23.68839 | 0.352302504 | 12 | Neutral |
| testneg_7149 | 0 | 0 | 0.387419 | -3 | -0.130439 | -3 | 0.46399 | 0.15334 | 0.85759 | 0.1031 | 27.00364 | 0.020375 | 0 | Neutral |
| testneg_7150 | -0.03 | -0.1 | 1.285186 | -3 | 1.156114 | -3 | 0.35957 | 0.11476 | 0.24239 | 0.111 | 15.82969 | 0.000149387 | 0 | Neutral |
| testneg_7151 | -0.07 | -0.38 | 6.853158 | -3 | 0.86526 | -3 | 0.29417 | 0.10965 | 0.64914 | 0.12481 | 12.91891 | 0.26 | 1.4 | Neutral |
| testneg_7152 | 0 | 0 | 1.237105 | -3 | 5.443984 | -3 | 0.15713 | 0.3507 | 0.28271 | 0.60265 | 25.5168 | 0.585487429 | 79.4 | Disease_causing |
| testneg_7153 | -0.03 | -0.1 | 0.337194 | -3 | 0.29338 | -3 | 0.1273 | 0.12788 | 0.30367 | 0.41964 | 10.41667 | 0.06124359 | 0 | Neutral |
| testneg_7154 | -0.04 | 0.23 | -1.000328 | -3 | 2.724287 | -3 | 0.07837 | 0.12788 | 0.16361 | 0.13388 | 13.40128 | 0.153240863 | 0 | Neutral |
| testneg_7155 | 0.06 | -0.57 | 0.282279 | -3 | 0.734765 | -3 | 0.03526 | 0.06855 | 0.07204 | 0.07708 | 17.8869 | 0.04 | 0 | Neutral |
| testneg_7156 | 0 | 0 | 1.540719 | -3 | 0.151938 | -3 | 0.13271 | 0.22958 | 0.47188 | 0.10037 | 13.18841 | 0.011 | 0 | Neutral |
| testneg_7157 | -0.05 | -0.29 | -1.421823 | -3 | 0.488803 | -3 | 0.20765 | 0.27124 | 0.14525 | 0.14744 | 18.33333 | 0.12265 | 0 | Neutral |
| testneg_7158 | 0 | 0 | 0.777519 | -3 | -0.585591 | -3 | 0.75071 | 0.63704 | 0.41193 | 0.10133 | 10 | 0.02 | 0 | Neutral |
| testneg_7159 | 0.17 | 0.38 | -1.243392 | -3 | 1.756169 | -3 | 0.09657 | 0.58605 | 0.35945 | 0.12025 | 11.25 | 0.12 | 0 | Neutral |
| testneg_7160 | 0 | 0 | 2.403686 | 2.403686 | 2.05038 | -3 | 0.48574 | 0.20473 | 0.09416 | 0.17821 | 27.66817 | 0.338194444 | 9.8 | Neutral |
| testneg_7161 | -0.03 | -0.1 | -0.76618 | -3 | 1.683626 | -3 | 0.1503 | 0.12788 | 0.25312 | 0.12788 | 13.11346 | 0.01 | 0 | Neutral |
| testneg_7162 | 0.18 | -1.05 | 6.835273 | 6.835273 | 3.806323 | -3 | 0.08412 | 0.28307 | 0.93853 | 0.93439 | 12.4505 | 0.82 | 100 | Disease_causing |
| testneg_7163 | 0.19 | -0.27 | 1.4509 | -3 | 2.75027 | -3 | 0.64566 | 0.22552 | 0.50034 | 0.07527 | 21.99918 | 0.120674242 | 0 | Neutral |
| testneg_7164 | -0.03 | -0.12 | 0.991332 | -3 | 1.181363 | -3 | 0.15954 | 0.12788 | 0.2521 | 0.1784 | 26.65569 | 0.066652907 | 0 | Neutral |
| testneg_7165 | -0.01 | -0.8 | 3.501031 | -3 | 6.258159 | -3 | 0.25421 | 0.16235 | 0.1556 | 0.11007 | 39.88188 | 0.353965812 | 11.4 | Neutral |
| testneg_7166 | -0.14 | 1.01 | 1.450684 | -3 | 1.491454 | -3 | 0.1621 | 0.1641 | 0.1078 | 0.17394 | 18.88889 | 0.190533886 | 0 | Neutral |
| testneg_7167 | 0.03 | 0.12 | 1.304269 | -3 | 1.628888 | -3 | 0.45029 | 0.12788 | 0.04047 | 0.12788 | 18.33333 | 0.010313725 | 0 | Neutral |
| testneg_7168 | 0 | 0 | 1.081416 | -3 | 2.103134 | -0.106324 | 0.59345 | 0.26484 | 0.2521 | 0.1784 | 21.38889 | 0.144584957 | 0 | Neutral |
| testneg_7169 | 0.21 | 0.15 | 5.867155 | -3 | 2.524962 | -3 | 0.13067 | 0.1005 | 0.36324 | 0.15729 | 13.51351 | 0.433784137 | 33.8 | Neutral |
| testneg_7170 | 0.07 | 0.38 | 1.846321 | -3 | 6.228264 | -3 | 0.23088 | 0.12788 | 0.35189 | 0.12788 | 29.08689 | 0.130399729 | 0 | Neutral |
| testneg_7171 | -0.05 | -0.29 | 1.38401 | -3 | 3.966623 | -3 | 0.37212 | 0.11323 | 0.16737 | 0.12788 | 18.33333 | 0.060294365 | 0 | Neutral |
| testneg_7172 | 0 | 0 | 0.388922 | -3 | 3.137305 | -3 | 0.98107 | 0.70722 | 0.12408 | 0.08828 | 5.73099 | 0.069777778 | 0 | Neutral |
| testneg_7173 | 0 | 0 | -0.409329 | -3 | 1.555121 | -3 | 0.95993 | 0.15588 | 0.08059 | 0.09576 | 13.4047 | 0.020642652 | 0 | Neutral |
| testneg_7174 | 0.01 | -1.44 | 2.093021 | -3 | 1.290195 | 1.290195 | 0.54724 | 0.12788 | 0.10761 | 0.12246 | 18.33333 | 0.041125 | 0 | Neutral |
| testneg_7175 | 0.18 | -1.05 | 4.060219 | -3 | 6.425529 | -3 | 0.1109 | 0.17295 | 0.30657 | 0.52974 | 32.36846 | 0.809871795 | 100 | Disease_causing |
| testneg_7176 | -0.04 | 0.23 | 0.421557 | 0.421557 | 1.124718 | 1.124718 | 0.82507 | 0.19615 | 0.13356 | 0.09963 | 10.46733 | 0.040252061 | 0 | Neutral |
| testneg_7177 | 0.01 | 0.8 | 3.468346 | -3 | -0.696429 | -3 | 0.51577 | 0.19561 | 0.26476 | 0.08217 | 19.82456 | 0.07 | 0 | Neutral |
| testneg_7178 | 0 | 0 | 1.65802 | -3 | 4.213187 | -3 | 0.2118 | 0.2009 | 0.08529 | 0.12376 | 14.79675 | 0.113572401 | 0 | Neutral |
| testneg_7179 | 0 | 0 | 0.585709 | -3 | -0.627995 | -3 | 0.77644 | 0.09695 | 0.13869 | 0.10313 | 13.27141 | 0.01 | 0 | Neutral |
| testneg_7180 | -0.08 | -0.03 | 1.455667 | -3 | -2.543636 | -3 | 0.14341 | 0.10134 | 0.00487 | 0.0143 | 18.33333 | 0.01 | 0 | Neutral |
| testneg_7181 | 0 | 0 | 0.988618 | -3 | 5.283558 | -3 | 0.15019 | 0.10342 | 0.85482 | 0.1072 | 12.29167 | 0.06 | 0 | Neutral |
| testneg_7182 | 0.04 | 2.01 | 0.958784 | -3 | 2.692724 | -3 | 0.08224 | 0.29179 | 0.52765 | 0.20417 | 9.77325 | 0.435520566 | 24.2 | Neutral |
| testneg_7183 | 0.04 | -0.23 | 1.625356 | -3 | 3.322059 | -3 | 0.5065 | 0.20697 | 0.1231 | 0.3662 | 70.67478 | 0.517832533 | 65.4 | Disease_causing |
| testneg_7184 | 0.03 | -1.03 | 3.319838 | 2.403241 | 1.577566 | -3 | 0.19898 | 0.12788 | 0.68423 | 0.08163 | 13.51351 | 0.221458333 | 2 | Neutral |
| testneg_7185 | -0.07 | -0.38 | 4.53569 | -3 | 2.272856 | -3 | 0.13559 | 0.1946 | 0.0925 | 0.08222 | 18.6768 | 0.123024133 | 0 | Neutral |
| testneg_7186 | 0.09 | -0.67 | 3.099902 | -3 | 2.439632 | -3 | 0.15822 | 0.12788 | 0.28326 | 0.13944 | 15.46432 | 0.194079437 | 0 | Neutral |
| testneg_7187 | -0.02 | -0.45 | 2.957366 | -3 | 0.660384 | -3 | 0.98442 | 0.44966 | 0.1834 | 0.12788 | 37.46364 | 0.118104167 | 0 | Neutral |
| testneg_7188 | 0.01 | 0.8 | 1.69518 | -3 | 3.716565 | -3 | 0.07995 | 0.12788 | 0.1594 | 0.22002 | 13.51351 | 0.245792423 | 0.4 | Neutral |
| testneg_7189 | 0 | -0.32 | 1.00433 | -3 | 7.034309 | -3 | 0.06254 | 0.23561 | 0.049 | 0.11324 | 43.87143 | 0.41 | 19.2 | Neutral |
| testneg_7190 | 0 | 0 | 0.467241 | -3 | 1.200491 | -3 | 0.14948 | 0.12788 | 0.34198 | 0.1113 | 20.20217 | 0 | 0 | Neutral |
| testneg_7191 | 0 | 0 | -0.470651 | -3 | 6.154212 | -3 | 0.28063 | 0.40063 | 0.09288 | 0.16617 | 18.04026 | 0.532568107 | 66.6 | Disease_causing |
| testneg_7192 | 0 | 0 | -0.535508 | -0.785566 | 0.439979 | -3 | 0.01196 | 0.12788 | 0.0811 | 0.09449 | 19.44444 | 0.04 | 0 | Neutral |
| testneg_7193 | 0.21 | 0.15 | 2.896973 | -3 | 6.070697 | -3 | 0.47335 | 0.12788 | 0.08027 | 0.12788 | 10 | 0.306666667 | 3.4 | Neutral |
| testneg_7194 | -0.05 | -0.29 | -0.746943 | -3 | 3.016624 | -3 | 0.14525 | 0.14744 | 0.33022 | 0.13818 | 18.86584 | 0.199610391 | 0.2 | Neutral |
| testneg_7195 | 0.03 | -1.03 | 4.326488 | -3 | -0.247306 | -3 | 0.10604 | 0.10785 | 0.35945 | 0.12025 | 27.95796 | 0.04 | 0 | Neutral |
| testneg_7196 | 0.06 | -0.57 | 3.760456 | -3 | 6.231595 | 1.823528 | 0.96704 | 0.16878 | 0.17016 | 0.12748 | 19.55465 | 0.432583333 | 31.8 | Neutral |
| testneg_7197 | 0 | -0.73 | 1.84849 | -3 | 3.349863 | -3 | 0.64608 | 0.35524 | 0.1472 | 0.12788 | 11.25 | 0.142128187 | 0 | Neutral |

| | | | | | | | | | | | | | | |
|---|---|---|---|---|---|---|---|---|---|---|---|---|---|---|
| testneg_7198 | -0.08 | -0.03 | 4.826887 | -3 | 1.700368 | -3 | 0.1631 | 0.40344 | 0.09211 | 0.08973 | 21.38889 | 0.159454942 | 0 | Neutral |
| testneg_7199 | 0.21 | 0.15 | 3.345724 | -3 | 5.747049 | 0.304305 | 0.30275 | 0.12788 | 0.50134 | 0.15488 | 38.08363 | 0.489161124 | 50 | Neutral |
| testneg_7200 | 0 | 0 | 0.002341 | -3 | -0.803439 | -3 | 0.10799 | 0.10754 | 0.50276 | 0.09638 | 20.89902 | 0.01 | 0 | Neutral |
| testneg_7201 | 0.03 | 0.12 | 1.249184 | -3 | -0.44217 | -3 | 0.40592 | 0.22108 | 0.33148 | 0.11968 | 13.77727 | 0.021666667 | 0 | Neutral |
| testneg_7202 | 0.15 | -0.28 | 0.59012 | -3 | 6.108445 | -3 | 0.605 | 0.10748 | 0.10723 | 0.11163 | 13.51351 | 0.120949169 | 0 | Neutral |
| testneg_7203 | 0 | 0 | 0.470493 | -3 | -0.262132 | -3 | 0.42802 | 0.12096 | 0.11788 | 0.12788 | 19.22222 | 0.000693022 | 0 | Neutral |
| testneg_7204 | 0 | 0 | 0.387542 | -3 | -2.421511 | -3 | 0.76693 | 0.19444 | 0.37464 | 0.12788 | 20.39295 | 0.02 | 0 | Neutral |
| testneg_7205 | 0.09 | -0.3 | 3.200815 | -3 | 0.385855 | 0.385855 | 0.98312 | 0.12788 | 0.10153 | 0.13343 | 9.1693 | 0.13 | 0 | Neutral |
| testneg_7206 | 0.00127273 | -0.0109091 | 3.063227 | -3 | 2.421333 | -3 | 0.05153 | 0.12788 | 0.12173 | 0.12788 | 15.55791 | 0.091214286 | 0 | Neutral |
| testneg_7207 | 0.07 | 0.38 | -0.415438 | -3 | 0.425569 | -3 | 0.19898 | 0.12788 | 0.12879 | 0.12788 | 36.81442 | 0.01 | 0 | Neutral |
| testneg_7208 | 0 | 0 | 2.138956 | -3 | 3.408336 | -3 | 0.27942 | 0.73513 | 0.41647 | 0.11239 | 11.57603 | 0.110583333 | 0 | Neutral |
| testneg_7209 | -0.14 | -0.54 | 0.83037 | -3 | 1.759623 | 1.759623 | 0.16929 | 0.12788 | 0.16475 | 0.26069 | 17.03707 | 0.130104167 | 0 | Neutral |
| testneg_7210 | 0.05 | -0.81 | 3.118508 | -3 | 6.022895 | 6.022895 | 0.26971 | 0.08509 | 0.09583 | 0.09376 | 17.4373 | 0.43 | 22.4 | Neutral |
| testneg_7211 | 0.04 | -0.23 | 6.319089 | -3 | 2.712843 | -3 | 0.19898 | 0.12788 | 0.34826 | 0.49454 | 23.33159 | 0.584050651 | 83.4 | Disease_causing |
| testneg_7212 | 0 | 0 | -0.142698 | -3 | 1.269375 | -3 | 0.11133 | 0.12788 | 0.54489 | 0.09921 | 21.23983 | 0.01 | 0 | Neutral |
| testneg_7213 | 0.004 | -0.4576364 | 7.731489 | -3 | -0.235499 | -3 | 0.03378 | 0.12788 | 0.1203 | 0.13872 | 29.64429 | 0.39 | 17.4 | Neutral |
| testneg_7214 | 0 | 0 | -0.151164 | -0.151164 | 0.713568 | -3 | 0.65708 | 0.0936 | 0.11072 | 0.38259 | 19.34837 | 0.10930323 | 0 | Neutral |
| testneg_7215 | 0.03 | 0.12 | 0.199359 | 0.199359 | 0.357284 | -3 | 0.07626 | 0.12788 | 0.08973 | 0.0992 | 23.13792 | 0.02 | 0 | Neutral |
| testneg_7216 | 0 | 0 | 0.885296 | -3 | 6.214458 | -3 | 0.6992 | 0.1804 | 0.22234 | 0.09474 | 29.66438 | 0.13434318 | 0.2 | Neutral |
| testneg_7217 | 0 | 0 | -0.044046 | -3 | 0.389489 | -3 | 0.47544 | 0.17658 | 0.11858 | 0.11551 | 22.38311 | 0.03 | 0 | Neutral |
| testneg_7218 | 0.04 | 2.01 | 4.022878 | -3 | 2.247345 | -3 | 0.44923 | 0.12788 | 0.09334 | 0.10027 | 18.25571 | 0.12 | 0 | Neutral |
| testneg_7219 | 0 | 0 | 0.011632 | -3 | -1.040235 | -3 | 0.32406 | 0.10985 | 0.14007 | 0.10647 | 20.83333 | 0.01 | 0 | Neutral |
| testneg_7220 | 0.01 | 0.8 | -1.158834 | -3 | 7.114639 | -3 | 0.12739 | 0.12788 | 0.8433 | 0.13128 | 22.47743 | 0.309722222 | 2.8 | Neutral |
| testneg_7221 | 0 | 0 | -0.055895 | -3 | 1.526737 | -3 | 0.31694 | 0.13475 | 0.78536 | 0.12709 | 5.34722 | 0.01 | 0 | Neutral |
| testneg_7222 | -0.03 | -0.1 | 4.235802 | -3 | 2.261306 | -3 | 0.9989 | 0.8746 | 0.76924 | 0.15588 | 13.25269 | 0.414126984 | 21 | Neutral |
| testneg_7223 | 0.05 | 0.29 | 2.437307 | -3 | 2.148373 | -3 | 0.3047 | 0.11929 | 0.21376 | 0.12708 | 11.25 | 0.010203968 | 0 | Neutral |
| testneg_7224 | 0 | 0 | 0.16922 | -3 | 6.411258 | -3 | 0.77196 | 0.14089 | 0.12545 | 0.08963 | 16.04971 | 0.132792208 | 0 | Neutral |
| testneg_7225 | 0 | -0.01 | 0.981481 | -3 | 2.164115 | -3 | 0.24516 | 0.11809 | 0.05476 | 0.12788 | 32.73654 | 0.01 | 0 | Neutral |
| testneg_7226 | 0 | 0 | 0.227306 | -3 | 4.576491 | -3 | 0.30412 | 0.54626 | 0.09488 | 0.10426 | 4.72222 | 0.110889356 | 0 | Neutral |
| testneg_7227 | -0.19 | -0.51 | -2.016047 | -3 | 0.654716 | -3 | 0.31584 | 0.11996 | 0.76942 | 0.13267 | 22.77778 | 0.11 | 0 | Neutral |
| testneg_7228 | 0.06 | 0.18 | -2.795124 | -3 | -2.410599 | -3 | 0.03502 | 0.08119 | 0.59691 | 0.11049 | 17.72113 | 0.040662879 | 0 | Neutral |
| testneg_7229 | 0.03 | 0.1 | 4.732985 | -3 | 1.283706 | -3 | 0.23506 | 0.24555 | 0.16856 | 0.19221 | 31.86802 | 0.371394346 | 11.6 | Neutral |
| testneg_7230 | 0 | 0 | 0.048895 | -3 | 6.881359 | -3 | 0.11301 | 0.10716 | 0.07406 | 0.08059 | 42.29432 | 0.19 | 0.2 | Neutral |
| testneg_7231 | 0.07 | 0.38 | 2.43609 | -3 | 3.225783 | 3.225783 | 0.02146 | 0.10351 | 0.75978 | 0.11423 | 12.79101 | 0.2 | 0 | Neutral |
| testneg_7232 | -0.03 | -0.1 | -0.38356 | -3 | 0.021158 | -3 | 0.0811 | 0.09449 | 0.11133 | 0.12788 | 20.78035 | 0.02 | 0 | Neutral |
| testneg_7233 | -0.01 | -0.79 | -0.201762 | -3 | 0.457163 | -3 | 0.4811 | 0.13398 | 0.57517 | 0.10873 | 18.6768 | 0.020356329 | 0 | Neutral |
| testneg_7234 | 0.11 | -1.95 | 3.250824 | -3 | 4.106182 | -3 | 0.94807 | 0.14977 | 0.06394 | 0.24464 | 9.79167 | 0.603600629 | 87.8 | Disease_causing |
| testneg_7235 | 0 | 0 | 1.108803 | -3 | 1.436839 | 1.436839 | 0.85092 | 0.58871 | 0.19898 | 0.12788 | 17.71302 | 0.042549267 | 0 | Neutral |
| testneg_7236 | 0.16 | 0.08 | 0.221162 | -3 | 1.751693 | -3 | 0.01369 | 0.12788 | 0.17418 | 0.1871 | 29.2735 | 0.134844633 | 0 | Neutral |
| testneg_7237 | 0 | 0 | 0.507327 | -3 | 1.249184 | -3 | 0.11172 | 0.10034 | 0.40592 | 0.22108 | 15.18432 | 0.026893579 | 0 | Neutral |
| testneg_7238 | 0 | 0 | -0.043138 | -3 | 0.16618 | -3 | 0.17737 | 0.54971 | 0.12403 | 0.12788 | 18.88889 | 0.050151138 | 0 | Neutral |
| testneg_7239 | 0.02 | -0.63 | 1.351912 | -3 | -0.465282 | -3 | 0.18584 | 0.11114 | 0.42872 | 0.12788 | 21.97787 | 0.01 | 0 | Neutral |
| testneg_7240 | -0.02 | 0.36 | 1.284836 | -3 | 2.512097 | -3 | 0.1214 | 0.0899 | 0.06517 | 0.15107 | 19.44444 | 0.13 | 0 | Neutral |
| testneg_7241 | 0 | 0 | 1.178684 | -3 | 3.851585 | -3 | 0.12974 | 0.11886 | 0.06552 | 0.07762 | 10 | 0.040236766 | 0 | Neutral |
| testneg_7242 | 0.04 | -0.23 | 2.522879 | -3 | 5.79991 | -3 | 0.99998 | 0.86471 | 0.37212 | 0.11323 | 12.37434 | 0.280367397 | 1.2 | Neutral |
| testneg_7243 | -0.08 | -0.03 | -0.070402 | -3 | 0.637019 | -3 | 0.0513 | 0.33622 | 0.16392 | 0.12788 | 12.77241 | 0.13 | 0 | Neutral |
| testneg_7244 | 0.09 | -0.3 | -1.680478 | -3 | 0.994041 | -3 | 0.1664 | 0.10113 | 0.13413 | 0.12788 | 21.38889 | 0.010645833 | 0 | Neutral |
| testneg_7245 | -0.19 | -0.51 | 0.918079 | 0.918079 | -1.378191 | -3 | 0.83039 | 0.1727 | 0.19898 | 0.12788 | 8.97898 | 0.08 | 0 | Neutral |
| testneg_7246 | 0 | 0 | 0.172526 | -3 | 6.645041 | -3 | 0.22162 | 0.11997 | 0.13433 | 0.09578 | 19.15385 | 0.148455882 | 0 | Neutral |
| testneg_7247 | 0 | 0 | 1.983447 | -3 | 0.90777 | -3 | 0.60812 | 0.19286 | 0.0961 | 0.21256 | 13.51351 | 0.072746722 | 0 | Neutral |
| testneg_7248 | 0.14 | -1.01 | 2.301074 | -3 | 1.515569 | -3 | 0.5065 | 0.20697 | 0.10554 | 0.12788 | 19.94848 | 0.08 | 0 | Neutral |
| testneg_7249 | 0 | 0 | 0.81674 | -3 | 0.777519 | -3 | 0.84238 | 0.12559 | 0.75071 | 0.63704 | 10.41667 | 0.02 | 0 | Neutral |
| testneg_7250 | 0.21 | 0.15 | 1.819355 | -3 | 3.262273 | -3 | 0.31124 | 0.08875 | 0.24736 | 0.12777 | 16.8018 | 0.095113636 | 0 | Neutral |
| testneg_7251 | 0.03 | -1.03 | 2.17416 | -3 | 5.456932 | -3 | 0.14021 | 0.43531 | 0.19969 | 0.1163 | 10.83333 | 0.298680876 | 2.6 | Neutral |
| testneg_7252 | 0.03 | 0.12 | 1.075884 | 1.075884 | -2.410599 | -3 | 0.16497 | 0.74918 | 0.59691 | 0.11049 | 26.34921 | 0.11 | 0 | Neutral |
| testneg_7253 | 0 | 0 | 0.178815 | -3 | -0.681729 | -3 | 0.5471 | 0.11262 | 0.09531 | 0.09236 | 13.51351 | 0.01011236 | 0 | Neutral |
| testneg_7254 | 0 | -0.01 | 0.494045 | -3 | 4.192605 | -3 | 0.1304 | 0.12788 | 0.08454 | 0.22908 | 18.33333 | 0.264689863 | 1.4 | Neutral |
| testneg_7255 | -0.05 | 0.81 | 3.899481 | -3 | 4.70798 | -3 | 0.83205 | 0.40904 | 0.17527 | 0.13941 | 19.22222 | 0.669939394 | 97.2 | Disease_causing |
| testneg_7256 | 0.03 | 1.22 | 2.679679 | -3 | 5.580492 | -3 | 0.17697 | 0.12974 | 0.07347 | 0.08817 | 25.17953 | 0.19 | 0 | Neutral |
| testneg_7257 | 0 | -0.32 | 3.044361 | -3 | 0.985043 | -3 | 0.12766 | 0.14626 | 0.0611 | 0.12135 | 12.34649 | 0.05 | 0 | Neutral |
| testneg_7258 | 0.15 | 0.06 | -0.364205 | -3 | 1.651197 | -3 | 0.45628 | 0.15819 | 0.76918 | 0.13588 | 10.19537 | 0.111583333 | 0 | Neutral |
| testneg_7259 | -0.14 | 0.37 | -0.694905 | -3 | 0.196098 | -3 | 0.4132 | 0.12788 | 0.12682 | 0.11357 | 33.99647 | 0.031482335 | 0 | Neutral |
| testneg_7260 | -0.03 | -0.1 | 4.958638 | -3 | 5.667234 | -3 | 0.66711 | 0.31094 | 0.71803 | 0.95763 | 20.74074 | 0.7725 | 100 | Disease_causing |
| testneg_7261 | 0.21 | 0.15 | 0.996592 | -3 | 0.497688 | -3 | 0.7257 | 0.64394 | 0.34467 | 0.13076 | 17.01711 | 0.12 | 0 | Neutral |
| testneg_7262 | 0.02 | -0.36 | -0.250215 | -3 | 0.731991 | -3 | 0.14705 | 0.08702 | 0.13141 | 0.1123 | 17.18434 | 0.00050641 | 0 | Neutral |
| testneg_7263 | 0 | 0 | -0.613522 | -3 | 2.696296 | -3 | 0.15989 | 0.12788 | 0.48801 | 0.09118 | 16.29475 | 0.012502904 | 0 | Neutral |
| testneg_7264 | 0 | 0 | -1.73577 | -3 | 4.435214 | -3 | 0.0931 | 0.12788 | 0.84431 | 0.65315 | 15.73011 | 0.230666667 | 0.2 | Neutral |
| testneg_7265 | 0.08 | 0.03 | -2.363283 | -2.363283 | 1.411079 | -3 | 0.05846 | 0.12788 | 0.19969 | 0.1163 | 33.31168 | 0.04 | 0 | Neutral |
| testneg_7266 | -0.04 | 0.23 | -0.274997 | -3 | 2.216856 | -3 | 0.37501 | 0.0997 | 0.16057 | 0.12305 | 17.08333 | 0.02 | 0 | Neutral |
| testneg_7267 | 0 | 0 | 1.618033 | -3 | 5.295382 | -3 | 0.35092 | 0.10596 | 0.19898 | 0.09584 | 18.12633 | 0.040981939 | 0 | Neutral |
| testneg_7268 | -0.06 | -0.18 | 1.556626 | -3 | -0.024667 | -3 | 0.46919 | 0.60081 | 0.59599 | 0.12477 | 20.27778 | 0.07 | 0 | Neutral |
| testneg_7269 | 0.05 | 0.29 | 2.54094 | -3 | 0.982814 | -3 | 0.32484 | 0.11621 | 0.06395 | 0.0759 | 11.74281 | 0.01 | 0 | Neutral |
| testneg_7270 | 0 | -0.01 | 0.736606 | -3 | 1.619888 | -3 | 0.27187 | 0.41985 | 0.19477 | 0.12788 | 10.22898 | 0.015314716 | 0 | Neutral |
| testneg_7271 | 0.06 | 0.18 | 2.275026 | -3 | 1.022212 | 0.433904 | 0.16777 | 0.1749 | 0.09093 | 0.28643 | 12.29167 | 0.152133987 | 0 | Neutral |
| testneg_7272 | 0.18 | -1.05 | 3.389178 | -3 | -3.264658 | -3 | 0.34931 | 0.11906 | 0.12366 | 0.32056 | 10.83333 | 0.244375 | 0 | Neutral |
| testneg_7273 | 0.14 | -0.37 | 4.646741 | -3 | 0.142039 | -3 | 0.53447 | 0.10345 | 0.82148 | 0.14847 | 16.8018 | 0.113738095 | 0 | Neutral |
| testneg_7274 | 0.14 | -1.01 | 2.621366 | -3 | -0.808449 | -0.808449 | 0.81899 | 0.12788 | 0.13258 | 0.9258 | 4.72222 | 0.16 | 0 | Neutral |
| testneg_7275 | 0 | 0 | 0.067126 | -3 | 0.410924 | -3 | 0.18869 | 0.23984 | 0.18431 | 0.13863 | 12.29167 | 0.08 | 0 | Neutral |
| testneg_7276 | 0 | 0 | 1.87611 | -3 | 0.72503 | -3 | 0.15107 | 0.10457 | 0.56862 | 0.12788 | 19.31241 | 0.000106395 | 0 | Neutral |
| testneg_7277 | -0.04 | 0.23 | 1.498072 | -3 | -0.35186 | -3 | 0.22113 | 0.10673 | 0.89845 | 0.25991 | 15.84247 | 0.04 | 0 | Neutral |
| testneg_7278 | -0.03 | -0.1 | 2.935592 | -3 | 0.387956 | -3 | 0.1909 | 0.14279 | 0.19898 | 0.12788 | 26.28933 | 0.04 | 0 | Neutral |
| testneg_7279 | 0.03 | 1.22 | 0.780247 | -3 | 0.280808 | -3 | 0.26524 | 0.11154 | 0.81659 | 0.10424 | 10.82519 | 0.055054945 | 0 | Neutral |
| testneg_7280 | 0 | 0 | 1.708897 | -3 | -1.286656 | -1.286656 | 0.5202 | 0.16306 | 0.10208 | 0.08867 | 17.73153 | 0.03 | 0 | Neutral |
| testneg_7281 | 0.02 | 0.45 | 5.680826 | -3 | 0.147043 | -3 | 0.18013 | 0.25346 | 0.14668 | 0.12788 | 20 | 0.24 | 0.8 | Neutral |
| testneg_7282 | 0.18 | -1.05 | 4.901007 | -3 | 4.187482 | -3 | 0.227 | 0.14389 | 0.08857 | 0.83499 | 19.28571 | 0.682028104 | 97.8 | Disease_causing |
| testneg_7283 | -0.02 | 0.63 | 3.191277 | -3 | -1.333487 | -3 | 0.37074 | 0.11532 | 0.07598 | 0.08456 | 30.03718 | 0.04 | 0 | Neutral |
| testneg_7284 | 0.19 | 0.51 | 3.907412 | -3 | 5.359975 | -3 | 0.31626 | 0.12936 | 0.12196 | 0.12788 | 17.44444 | 0.321536319 | 5.6 | Neutral |
| testneg_7285 | 0.21 | 0.15 | 1.098308 | -3 | 1.69701 | -3 | 0.09346 | 0.12788 | 0.27197 | 0.12346 | 19.04744 | 0.020109163 | 0 | Neutral |
| testneg_7286 | 0.21 | 0.15 | 5.407311 | -3 | 5.438117 | -3 | 0.08975 | 0.07399 | 0.12463 | 0.12788 | 28.33579 | 0.300576497 | 6.2 | Neutral |
| testneg_7287 | 0.07 | 0.01 | -0.38551 | -3 | 1.624942 | 1.624942 | 0.16797 | 0.12788 | 0.16208 | 0.10597 | 27.46106 | 0.020139513 | 0 | Neutral |
| testneg_7288 | 0.07 | 0.38 | -0.919669 | -3 | 4.153004 | -0.416274 | 0.83936 | 0.42313 | 0.08184 | 0.10206 | 11.95767 | 0.172792208 | 0 | Neutral |
| testneg_7289 | 0 | 0 | -0.553742 | -0.553742 | 3.352237 | -3 | 0.1424 | 0.12788 | 0.17672 | 0.14229 | 18.36272 | 0.157357217 | 0 | Neutral |
| testneg_7290 | 0 | 0 | -0.049164 | -0.049164 | 1.633871 | -3 | 0.12322 | 0.1246 | 0.11849 | 0.20101 | 31.59722 | 0.0907762 | 0 | Neutral |
| testneg_7291 | 0.01 | 0.8 | 0.600693 | -3 | 3.966623 | -3 | 0.07611 | 0.11551 | 0.16737 | 0.12788 | 12.80035 | 0.09 | 0 | Neutral |
| testneg_7292 | 0 | -0.01 | 1.315543 | -3 | 1.513879 | 0.606649 | 0.28358 | 0.38765 | 0.62199 | 0.14032 | 14.68494 | 0.108802972 | 0 | Neutral |
| testneg_7293 | 0.18 | -1.05 | 3.036954 | -3 | 0.618993 | 0.618993 | 0.36541 | 0.27749 | 0.04207 | 0.12788 | 34.51624 | 0.11 | 0 | Neutral |
| testneg_7294 | 0.22 | -2.66 | 7.971027 | -3 | -3.595695 | -3 | 0.11858 | 0.11551 | 0.13615 | 0.12788 | 17.84816 | 0.29 | 2.4 | Neutral |
| testneg_7295 | 0.00436364 | 0.32418182 | 8.814673 | -3 | 0.259095 | -3 | 0.27308 | 0.12788 | 0.99994 | 0.51636 | 11.25 | 0.408039813 | 23.4 | Neutral |
| testneg_7296 | 0 | 0 | 1.067933 | -3 | 5.564611 | -3 | 0.08775 | 0.12788 | 0.54015 | 0.13176 | 19.09347 | 0.215423611 | 0.2 | Neutral |
| testneg_7297 | -0.03 | -0.1 | 7.349616 | -3 | 5.47117 | -3 | 0.19948 | 0.12788 | 0.19698 | 0.1459 | 17.88746 | 0.737424994 | 99.4 | Disease_causing |
| testneg_7298 | 0 | 0 | 0.740177 | -3 | -1.439996 | -3 | 0.40612 | 0.45135 | 0.04484 | 0.95426 | 10 | 0.130883459 | 0 | Neutral |
| testneg_7299 | 0 | 0 | 0.828595 | -3 | 6.258456 | -3 | 0.93284 | 0.71489 | 0.44923 | 0.12788 | 11.97094 | 0.165659983 | 0 | Neutral |
| testneg_7300 | 0 | 0 | 0.371766 | -3 | 4.959501 | -3 | 0.3095 | 0.13502 | 0.14301 | 0.12788 | 13.27141 | 0.065845142 | 0 | Neutral |
| testneg_7301 | 0.04 | -1.61 | 3.074697 | -3 | 0.676746 | -3 | 0.37282 | 0.07943 | 0.61306 | 0.15258 | 24.26961 | 0.14 | 0 | Neutral |
| testneg_7302 | 0 | 0 | 0.379738 | -3 | 4.6126 | -3 | 0.11731 | 0.06348 | 0.10809 | 0.09057 | 23.49305 | 0.06 | 0 | Neutral |
| testneg_7303 | 0.12 | -1.16 | 4.454599 | -3 | -0.534952 | -3 | 0.29284 | 0.11262 | 0.12203 | 0.13515 | 10 | 0.13 | 0 | Neutral |
| testneg_7304 | 0 | -0.01 | 1.125724 | -3 | 4.888628 | -3 | 0.6185 | 0.09316 | 0.40076 | 0.11341 | 4.63186 | 0.041403941 | 0 | Neutral |
| testneg_7305 | -0.21 | -0.15 | -2.747914 | -3 | 5.180034 | -3 | 0.03651 | 0.06141 | 0.1499 | 0.09632 | 21.85743 | 0.21 | 0 | Neutral |
| testneg_7306 | 0.22 | -2.66 | 6.542128 | -3 | 0.2532 | -3 | 0.38171 | 0.15977 | 0.42872 | 0.12788 | 32.97591 | 0.290055556 | 2.2 | Neutral |
| testneg_7307 | 0.07 | -0.31 | -0.565166 | -3 | 2.794319 | -3 | 0.9832 | 0.39904 | 0.60812 | 0.19286 | 21.38889 | 0.289319665 | 0.6 | Neutral |
| testneg_7308 | 0.14 | -1.01 | 7.12057 | -3 | -0.449162 | -3 | 0.46712 | 0.93559 | 0.09004 | 0.07829 | 20.83333 | 0.398642857 | 18.6 | Neutral |
| testneg_7309 | 0 | -0.01 | 2.426894 | -3 | -0.884409 | -3 | 0.06513 | 0.12353 | 0.08857 | 0.83499 | 44.58333 | 0.14 | 0 | Neutral |
| testneg_7310 | -0.14 | -0.54 | 6.019214 | -3 | 6.169765 | -3 | 0.42585 | 0.18081 | 0.09998 | 0.09022 | 16.60044 | 0.619375 | 88.8 | Disease_causing |
| testneg_7311 | -0.02 | -0.13 | 1.290429 | -1.019571 | 1.744987 | -3 | 0.19898 | 0.20319 | 0.06552 | 0.07762 | 18.6339 | 0.03 | 0 | Neutral |
| testneg_7312 | -0.02 | -0.45 | 6.712641 | -3 | -1.752013 | -3 | 0.25731 | 0.12662 | 0.0609 | 0.47188 | 20.83333 | 0.46 | 37.2 | Neutral |
| testneg_7313 | 0.21 | 0.15 | -1.015091 | -3 | -0.406057 | -3 | 0.04525 | 0.12788 | 0.7413 | 0.18974 | 12.66993 | 0.112916667 | 0 | Neutral |
| testneg_7314 | -0.07 | -0.01 | 0.920319 | -3 | -0.474499 | -3 | 0.41102 | 0.0974 | 0.1 | 0.22633 | 14.17187 | 0.050093608 | 0 | Neutral |
| testneg_7315 | 0.02 | 0.45 | 5.736474 | -3 | -1.914326 | -3 | 0.24327 | 0.12192 | 0.2124 | 0.17288 | 29.35728 | 0.348958333 | 7.4 | Neutral |
| testneg_7316 | 0.07 | 0.38 | -3.138593 | -3 | 6.096653 | -3 | 0.06254 | 0.23561 | 0.07961 | 0.11474 | 27.96262 | 0.2955 | 0.6 | Neutral |
| testneg_7317 | -0.19 | 0.25 | 4.217228 | -3 | 1.891702 | -3 | 0.23468 | 0.10874 | 0.26489 | 0.10549 | 35.4743 | 0.089084178 | 0 | Neutral |
| testneg_7318 | 0.16 | -1.37 | 6.097554 | -3 | -0.510252 | -3 | 0.13794 | 0.40936 | 0.30311 | 0.11121 | 11.21556 | 0.259444444 | 1.2 | Neutral |
| testneg_7319 | -0.07 | -0.33 | 1.690244 | -3 | -0.257167 | -3 | 0.19898 | 0.12788 | 0.1215 | 0.15825 | 10.41667 | 0.067962963 | 0 | Neutral |

| | | | | | | | | | | | | | | |
|---|---|---|---|---|---|---|---|---|---|---|---|---|---|---|
| testneg_7320 | 0 | 0 | 1.429927 | -3 | 0.188212 | -3 | 0.40612 | 0.45135 | 0.34859 | 0.09799 | 12.38369 | 0.01 | 0 | Neutral |
| testneg_7321 | -0.02 | -0.45 | 5.23066 | -3 | -2.637354 | -3 | 0.07219 | 0.12788 | 0.2698 | 0.18815 | 27.58695 | 0.347916667 | 7.2 | Neutral |
| testneg_7322 | 0.08 | 0.03 | -0.412359 | -3 | -0.272804 | -0.272804 | 0.4003 | 0.10884 | 0.17839 | 0.12009 | 25.60228 | 0.03 | 0 | Neutral |
| testneg_7323 | 0.05 | 0.29 | 1.539242 | -3 | -2.464297 | -2.464297 | 0.14477 | 0.18932 | 0.11524 | 0.12788 | 20 | 0.03 | 0 | Neutral |
| testneg_7324 | 0.07 | 0.38 | 2.315884 | -3 | 0.692563 | -3 | 0.22443 | 0.10657 | 0.39648 | 0.97288 | 4.44444 | 0.092817687 | 0 | Neutral |
| testneg_7325 | 0 | -0.32 | 3.805191 | -3 | 1.341885 | -3 | 0.56307 | 0.10536 | 0.06134 | 0.08207 | 18.91311 | 0.04 | 0 | Neutral |
| testneg_7326 | -0.000303 | -0.0069697 | 0.145016 | -3 | 5.312389 | -3 | 0.42621 | 0.49093 | 0.07293 | 0.12989 | 10.41667 | 0.15322681 | 0 | Neutral |
| testneg_7327 | 0.07 | 0.38 | -1.249264 | -3 | 0.919607 | -3 | 0.05904 | 0.09282 | 0.09719 | 0.09576 | 35.14905 | 0.04 | 0 | Neutral |
| testneg_7328 | -0.03 | -0.12 | 0.194478 | -3 | 0.334215 | -3 | 0.11006 | 0.14352 | 0.33314 | 0.47999 | 29.81852 | 0.130247689 | 0 | Neutral |
| testneg_7329 | -0.02 | -0.13 | 1.69747 | -3 | -2.248143 | -3 | 0.30942 | 0.13105 | 0.53424 | 0.08484 | 18.33333 | 0.01 | 0 | Neutral |
| testneg_7330 | 0.02 | -0.68 | 6.639247 | -3 | 0.207967 | -3 | 0.27407 | 0.1136 | 0.13295 | 0.12914 | 26.92625 | 0.197083333 | 0 | Neutral |
| testneg_7331 | 0 | 0 | 2.918397 | -3 | 0.892974 | 0.892974 | 0.63378 | 0.09451 | 0.06986 | 0.12788 | 17.08586 | 0.04 | 0 | Neutral |
| testneg_7332 | -0.06 | -0.18 | 2.052658 | -3 | 2.548086 | -3 | 0.25905 | 0.23728 | 0.18964 | 0.34036 | 11.75623 | 0.334687128 | 6 | Neutral |
| testneg_7333 | -0.02 | 0.36 | 1.311291 | -3 | 0.65067 | -3 | 0.07577 | 0.09128 | 0.25312 | 0.12788 | 22.30994 | 0.01 | 0 | Neutral |
| testneg_7334 | 0.00666667 | 0.08708333 | 0.896944 | -3 | 2.579716 | -3 | 0.11941 | 0.10144 | 0.41383 | 0.119 | 9.94779 | 0.025717391 | 0 | Neutral |
| testneg_7335 | 0 | 0 | 2.064873 | -3 | 1.498072 | -3 | 0.65564 | 0.11468 | 0.22113 | 0.10673 | 23.6233 | 0.01 | 0 | Neutral |
| testneg_7336 | 0.14 | -1.01 | 3.283596 | -3 | 1.243447 | -3 | 0.72501 | 0.7625 | 0.54895 | 0.15082 | 5.69444 | 0.28 | 2 | Neutral |
| testneg_7337 | 0 | 0 | 1.765739 | -3 | 4.894604 | -3 | 0.76186 | 0.10052 | 0.21726 | 0.23214 | 12.66667 | 0.183008985 | 0.2 | Neutral |
| testneg_7338 | 0.05 | -0.81 | 7.120988 | -3 | 0.24062 | -3 | 0.31025 | 0.09491 | 0.18351 | 0.11213 | 12.58766 | 0.25 | 2.2 | Neutral |
| testneg_7339 | 0.15 | 0.06 | 0.213595 | -3 | 4.553087 | -3 | 0.42306 | 0.86891 | 0.50403 | 0.21568 | 16.73378 | 0.520608466 | 67.2 | Disease_causing |
| testneg_7340 | 0 | 0 | -0.064878 | -3 | 0.182748 | -3 | 0.99668 | 0.30319 | 0.64188 | 0.09494 | 4.44444 | 0.04 | 0 | Neutral |
| testneg_7341 | 0.07 | -1.12 | -0.401524 | -3 | 4.220552 | -3 | 0.14185 | 0.12788 | 0.59864 | 0.14106 | 35.56492 | 0.232541667 | 0 | Neutral |
| testneg_7342 | 0.03 | -1.03 | 1.624942 | 1.624942 | -0.219149 | -3 | 0.16208 | 0.10597 | 0.16409 | 0.12788 | 11.875 | 0.05 | 0 | Neutral |
| testneg_7343 | 0 | 0 | -0.05829 | -3 | 3.191277 | 3.191277 | 0.38115 | 0.21061 | 0.37074 | 0.11532 | 26.83387 | 0.2 | 0 | Neutral |
| testneg_7344 | -0.03 | -0.1 | 2.735353 | -3 | 3.797218 | -3 | 0.29478 | 0.12788 | 0.09871 | 0.09286 | 10.30286 | 0.09869246 | 0 | Neutral |
| testneg_7345 | 0.14 | -1.01 | 2.843674 | -3 | 1.955212 | -3 | 0.84596 | 0.10633 | 0.29804 | 0.12768 | 23.47379 | 0.040833333 | 0 | Neutral |
| testneg_7346 | 0 | -0.32 | -0.864716 | -3 | 1.054082 | -3 | 0.16734 | 0.15176 | 0.23141 | 0.12788 | 26.9055 | 0.020235588 | 0 | Neutral |
| testneg_7347 | 0 | 0 | 0.919505 | -3 | 1.373367 | -3 | 0.19898 | 0.12788 | 0.34314 | 0.18668 | 16.84579 | 0.021027778 | 0 | Neutral |
| testneg_7348 | 0.02 | 0.13 | 5.072403 | -3 | 2.686668 | 2.686668 | 0.17301 | 0.14089 | 0.10517 | 0.09803 | 12.29167 | 0.262291667 | 0.6 | Neutral |
| testneg_7349 | 0 | 0 | 1.430205 | -3 | 1.749347 | -3 | 0.13884 | 0.12788 | 0.04829 | 0.09848 | 20.7596 | 0.01 | 0 | Neutral |
| testneg_7350 | 0.14 | -1.01 | 3.46287 | -3 | 4.797405 | -3 | 0.49512 | 0.08696 | 0.17451 | 0.11227 | 12.38369 | 0.19 | 0 | Neutral |
| testneg_7351 | 0 | -0.01 | 2.663664 | -3 | -3.692308 | -3 | 0.19898 | 0.26604 | 0.19898 | 0.12788 | 31.46572 | 0.04 | 0 | Neutral |
| testneg_7352 | -0.0138182 | -0.2629091 | 4.820901 | -3 | 0.292616 | -3 | 0.06358 | 0.5364 | 0.04142 | 0.09564 | 20.85753 | 0.2 | 0 | Neutral |
| testneg_7353 | -0.15 | 1.06 | -0.750656 | -3 | 4.993112 | -3 | 0.14367 | 0.95422 | 0.2852 | 0.12788 | 4.44444 | 0.409565217 | 16.2 | Neutral |
| testneg_7354 | 0.03 | 0.12 | 5.185173 | -3 | 0.194184 | -3 | 0.06118 | 0.08724 | 0.18744 | 0.09392 | 10.75027 | 0.1 | 0 | Neutral |
| testneg_7355 | 0 | 0 | 0.572685 | -3 | 2.830718 | -3 | 0.14632 | 0.12662 | 0.11229 | 0.08897 | 18.57759 | 0.01068906 | 0 | Neutral |
| testneg_7356 | 0 | 0 | 0.708274 | -3 | -0.169987 | -3 | 0.21928 | 0.22085 | 0.99445 | 0.10396 | 10.41667 | 0.040206186 | 0 | Neutral |
| testneg_7357 | 0.05 | 0.29 | 1.966589 | -3 | 2.729607 | -3 | 0.99317 | 0.60884 | 0.29739 | 0.10893 | 6.94444 | 0.12 | 0 | Neutral |
| testneg_7358 | 0.16 | 0.08 | 4.560265 | -3 | 5.955249 | -3 | 0.43459 | 0.12595 | 0.26842 | 0.09683 | 16.16034 | 0.33 | 8 | Neutral |
| testneg_7359 | -0.09 | 0.25 | 2.611714 | -3 | 1.187301 | -3 | 0.99543 | 0.12788 | 0.11746 | 0.1471 | 32.46067 | 0.114166667 | 0 | Neutral |
| testneg_7360 | -0.03 | -0.1 | 1.820578 | -3 | 3.730638 | 0.015616 | 0.31507 | 0.61057 | 0.05999 | 0.12788 | 10.22898 | 0.160082645 | 0 | Neutral |
| testneg_7361 | 0.07 | 0.7 | 4.681701 | -3 | 7.833649 | -3 | 0.10571 | 0.12788 | 0.20951 | 0.16278 | 24.38306 | 0.693759259 | 97.4 | Disease_causing |
| testneg_7362 | 0 | 0 | 0.094005 | -3 | -0.680715 | -3 | 0.64302 | 0.11262 | 0.40264 | 0.12788 | 18.12633 | 0.000318523 | 0 | Neutral |
| testneg_7363 | -0.18 | -0.16 | -1.333487 | -3 | 0.021357 | 0.021357 | 0.07598 | 0.08456 | 0.17935 | 0.36953 | 12.31231 | 0.16 | 0 | Neutral |
| testneg_7364 | 0.14 | -1.01 | 0.302883 | -3 | -0.395207 | -3 | 0.10486 | 0.08083 | 0.2098 | 0.44672 | 19.09347 | 0.1 | 0 | Neutral |
| testneg_7365 | -0.01 | -0.79 | 1.854262 | -3 | 1.241234 | -3 | 0.07701 | 0.19824 | 0.6258 | 0.16461 | 17.88514 | 0.18 | 0 | Neutral |
| testneg_7366 | 0.05 | -0.81 | 0.331709 | 0.331709 | 1.040567 | -3 | 0.16534 | 0.26726 | 0.10869 | 0.1503 | 17.21847 | 0.167471154 | 0 | Neutral |
| testneg_7367 | -0.02 | -0.41 | -0.390123 | -3 | 2.996727 | -3 | 0.09657 | 0.58605 | 0.19898 | 0.17518 | 18.88889 | 0.383403908 | 15.4 | Neutral |
| testneg_7368 | -0.15 | -0.04 | 6.878134 | -3 | -2.410599 | -3 | 0.26731 | 0.16415 | 0.59691 | 0.11049 | 19.14863 | 0.33 | 5.4 | Neutral |
| testneg_7369 | 0.21 | 0.15 | 8.09625 | -3 | 0.558914 | -3 | 0.26434 | 0.12788 | 0.90308 | 0.28373 | 19.09347 | 0.37225 | 13.6 | Neutral |
| testneg_7370 | 0.02 | -0.28 | -1.781198 | -3 | 0.479266 | -3 | 0.43897 | 0.12441 | 0.07225 | 0.12788 | 12.87484 | 0.012 | 0 | Neutral |
| testneg_7371 | 0.02 | 0.41 | -2.259592 | -3 | 3.178104 | -3 | 0.06311 | 0.14802 | 0.18559 | 0.09173 | 29.5935 | 0.134162338 | 0 | Neutral |
| testneg_7372 | 0.03 | 1.22 | 6.59818 | -3 | 6.178136 | -3 | 0.20152 | 0.09895 | 0.19898 | 0.12788 | 8.97898 | 0.64 | 93.4 | Disease_causing |
| testneg_7373 | 0.14 | -0.37 | 0.198379 | -3 | 4.37839 | -3 | 0.03991 | 0.08342 | 0.30026 | 0.23021 | 12.29167 | 0.272940171 | 0.8 | Neutral |
| testneg_7374 | -0.19 | -0.51 | 1.521336 | -3 | 6.416133 | -3 | 0.39553 | 0.11991 | 0.18499 | 0.10583 | 13.86987 | 0.221894841 | 0 | Neutral |
| testneg_7375 | 0.14 | -1.85 | 1.159993 | -3 | 1.084198 | -3 | 0.22255 | 0.10057 | 0.1434 | 0.09092 | 30.2158 | 0.04 | 0 | Neutral |
| testneg_7376 | -0.08 | -0.03 | 1.192851 | -3 | 0.96383 | -3 | 0.26181 | 0.12875 | 0.13195 | 0.57267 | 18.33333 | 0.073768939 | 0 | Neutral |
| testneg_7377 | 0 | -0.32 | -2.498179 | -3 | 6.329482 | 0.983063 | 0.50682 | 0.12788 | 0.2755 | 0.12788 | 13.27141 | 0.17 | 0 | Neutral |
| testneg_7378 | 0 | 0 | 0.56241 | -3 | -0.593986 | -3 | 0.9511 | 0.54626 | 0.31732 | 0.27963 | 26.02151 | 0.141365546 | 0 | Neutral |
| testneg_7379 | 0.21 | 0.15 | 1.941255 | -3 | -0.707221 | -3 | 0.35401 | 0.08438 | 0.11115 | 0.09814 | 18.50536 | 0.04 | 0 | Neutral |
| testneg_7380 | -0.03 | -1.22 | 2.92942 | -3 | 1.557993 | -3 | 0.15107 | 0.10457 | 0.19898 | 0.12788 | 18.33333 | 0.04 | 0 | Neutral |
| testneg_7381 | 0 | 0 | 0.704701 | -3 | 7.077282 | -3 | 0.39576 | 0.10761 | 0.80314 | 0.42344 | 13.49076 | 0.307807252 | 7 | Neutral |
| testneg_7382 | -0.05 | 0.81 | 4.203418 | -3 | 1.15789 | 1.15789 | 0.42919 | 0.1092 | 0.68439 | 0.09371 | 10.41667 | 0.1 | 0 | Neutral |
| testneg_7383 | 0 | 0 | 0.223823 | -3 | -1.422138 | -3 | 0.32484 | 0.11621 | 0.16091 | 0.90971 | 21.83934 | 0.092432213 | 0 | Neutral |
| testneg_7384 | 0.03 | 0.12 | 0.114579 | -3 | 3.642837 | -3 | 0.49745 | 0.1103 | 0.0501 | 0.12788 | 20.51588 | 0.101712066 | 0 | Neutral |
| testneg_7385 | -0.15 | -0.04 | -1.165102 | -3 | 1.441342 | -3 | 0.37237 | 0.36829 | 0.11072 | 0.38259 | 10 | 0.23366129 | 0 | Neutral |
| testneg_7386 | 0.09 | -0.3 | 4.393532 | -3 | 4.129128 | -3 | 0.58677 | 0.11579 | 0.25872 | 0.12209 | 19.66638 | 0.190677496 | 0 | Neutral |
| testneg_7387 | 0.07 | -0.31 | 2.84362 | -3 | 5.29853 | -3 | 0.2903 | 0.11176 | 0.16636 | 0.12384 | 20.42484 | 0.157317316 | 0 | Neutral |
| testneg_7388 | -0.02 | 0.68 | 0.250053 | -3 | 0.980928 | -3 | 0.36541 | 0.27749 | 0.50275 | 0.38957 | 12.30072 | 0.149647998 | 0 | Neutral |
| testneg_7389 | 0 | 0 | 0.053529 | -3 | 1.45834 | -3 | 0.46318 | 0.14229 | 0.64701 | 0.94571 | 11.64365 | 0.060037037 | 0 | Neutral |
| testneg_7390 | 0.18 | -1.05 | 5.334127 | -3 | 3.548514 | -3 | 0.38171 | 0.15977 | 0.19623 | 0.08363 | 20.60663 | 0.34270202 | 5.6 | Neutral |
| testneg_7391 | 0 | 0 | -0.264755 | -3 | -0.457064 | -3 | 0.09024 | 0.08559 | 0.21383 | 0.11171 | 12.4359 | 0.000274936 | 0 | Neutral |
| testneg_7392 | -0.1 | -1.9 | 6.033158 | -3 | 4.751068 | -3 | 0.09997 | 0.1797 | 0.9449 | 0.9378 | 14.4832 | 0.743723125 | 99.6 | Disease_causing |
| testneg_7393 | 0 | 0 | -0.32829 | -3 | 2.401476 | -3 | 0.18509 | 0.12831 | 0.28532 | 0.10663 | 20.39199 | 0.014250102 | 0 | Neutral |
| testneg_7394 | 0 | 0 | 1.154268 | 1.154268 | -0.19293 | -3 | 0.23071 | 0.09582 | 0.08332 | 0.09046 | 6.95654 | 0.021909091 | 0 | Neutral |
| testneg_7395 | 0.01 | -0.57 | 1.950103 | 1.950103 | 1.545256 | -3 | 0.20527 | 0.10053 | 0.20547 | 0.12178 | 19.20428 | 0.07 | 0 | Neutral |
| testneg_7396 | 0 | 0 | 0.274438 | -3 | 5.911847 | -3 | 0.12786 | 0.72066 | 0.40179 | 0.12112 | 11.21902 | 0.230012771 | 0.4 | Neutral |
| testneg_7397 | 0 | 0 | 1.608136 | -3 | 2.409244 | -3 | 0.19948 | 0.12788 | 0.99535 | 0.38424 | 17.26246 | 0.11725 | 0 | Neutral |
| testneg_7398 | -0.19 | -0.51 | 2.183162 | -3 | 2.798182 | 2.798182 | 0.22078 | 0.12788 | 0.31043 | 0.09688 | 18.64268 | 0.16 | 0 | Neutral |
| testneg_7399 | 0.00127273 | -0.0109091 | 3.063227 | -3 | 2.144477 | -3 | 0.05153 | 0.12788 | 0.12943 | 0.14391 | 18.88889 | 0.198881119 | 0 | Neutral |
| testneg_7400 | 0.09 | -0.67 | 2.624257 | -3 | 0.009388 | -3 | 0.19043 | 0.07855 | 0.47188 | 0.10037 | 19.85862 | 0.02 | 0 | Neutral |
| testneg_7401 | 0.04 | -0.23 | 1.940606 | -3 | 1.384899 | 1.384899 | 0.03991 | 0.08342 | 0.56315 | 0.13761 | 6.35445 | 0.100066667 | 0 | Neutral |
| testneg_7402 | 0 | 0 | 1.678033 | -3 | 3.350472 | -0.969128 | 0.33957 | 0.12788 | 0.09577 | 0.2436 | 10.40266 | 0.161997872 | 0 | Neutral |
| testneg_7403 | 0 | -0.32 | -2.345951 | -3 | 4.512673 | -3 | 0.13199 | 0.14263 | 0.14731 | 0.12788 | 22.95764 | 0.14 | 0 | Neutral |
| testneg_7404 | 0.02 | 0.45 | 3.50895 | -3 | 1.89485 | -3 | 0.28165 | 0.15912 | 0.19898 | 0.10116 | 17.21847 | 0.07 | 0 | Neutral |
| testneg_7405 | 0 | -0.01 | 0.601639 | -3 | 5.036729 | -3 | 0.08373 | 0.09191 | 0.17793 | 0.10516 | 29.16667 | 0.050361705 | 0 | Neutral |
| testneg_7406 | 0 | 0 | 0.425569 | 0.425569 | 6.742936 | -3 | 0.12879 | 0.12788 | 0.23962 | 0.10029 | 28.52027 | 0.193666667 | 0.4 | Neutral |
| testneg_7407 | 0.16 | -0.37 | 5.430369 | -3 | -0.467116 | -3 | 0.2578 | 0.09796 | 0.18673 | 0.18881 | 11.25 | 0.329642089 | 6.8 | Neutral |
| testneg_7408 | -0.05 | 0.81 | 2.250533 | -3 | 0.08175 | -3 | 0.14021 | 0.43531 | 0.32334 | 0.10525 | 17.77778 | 0.07 | 0 | Neutral |
| testneg_7409 | 0.22 | -2.66 | 3.810217 | -3 | 3.709266 | -3 | 0.10562 | 0.09329 | 0.08067 | 0.34914 | 42.31244 | 0.52 | 61 | Disease_causing |
| testneg_7410 | 0 | -0.32 | -1.995969 | -3 | 3.535649 | 3.003095 | 0.12949 | 0.44208 | 0.09284 | 0.26053 | 10 | 0.471318452 | 42.8 | Neutral |
| testneg_7411 | -0.15 | -0.04 | 3.431501 | -3 | 0.556181 | 0.556181 | 0.1289 | 0.14878 | 0.32406 | 0.28376 | 12.91516 | 0.3 | 0.8 | Neutral |
| testneg_7412 | 0.07 | 0.38 | -1.41619 | -3 | 0.011632 | -3 | 0.31124 | 0.08875 | 0.32406 | 0.10985 | 12.70833 | 0.010216721 | 0 | Neutral |
| testneg_7413 | 0 | 0 | 1.636749 | -3 | -0.357872 | -3 | 0.1109 | 0.17295 | 0.13825 | 0.0985 | 23.39477 | 0.0205 | 0 | Neutral |
| testneg_7414 | 0 | 0 | 1.376454 | -3 | 1.690495 | -3 | 0.1679 | 0.24232 | 0.11891 | 0.11186 | 12.58766 | 0.010512821 | 0 | Neutral |
| testneg_7415 | 0 | -0.01 | 0.558245 | -3 | -2.479355 | -3 | 0.19898 | 0.10362 | 0.17853 | 0.32085 | 11.56063 | 0.040185759 | 0 | Neutral |
| testneg_7416 | 0 | 0 | 1.317673 | -3 | 3.441233 | -3 | 0.45415 | 0.12971 | 0.22046 | 0.12788 | 10.83333 | 0.04 | 0 | Neutral |
| testneg_7417 | 0.04 | -0.23 | 6.279292 | -3 | 1.539468 | -3 | 0.09488 | 0.10426 | 0.19898 | 0.12788 | 28.37793 | 0.15 | 0.2 | Neutral |
| testneg_7418 | 0.16 | 0.08 | 4.643832 | -3 | 3.095089 | -3 | 0.47019 | 0.12788 | 0.22582 | 0.12788 | 19.10253 | 0.188713694 | 0 | Neutral |
| testneg_7419 | 0 | 0 | 0.484541 | -3 | 0.778541 | -3 | 0.48023 | 0.12788 | 0.226 | 0.28784 | 20.39742 | 0.05 | 0 | Neutral |
| testneg_7420 | 0 | 0 | 0.535401 | -3 | 3.103955 | -3 | 0.605 | 0.10748 | 0.06667 | 0.08666 | 19.63251 | 0.06 | 0 | Neutral |
| testneg_7421 | -0.08 | -0.03 | 6.638158 | -3 | -1.352736 | -3 | 0.07995 | 0.12788 | 0.27976 | 0.12613 | 13.51351 | 0.23 | 0.2 | Neutral |
| testneg_7422 | 0.06 | -0.57 | 3.612387 | -3 | 0.577305 | -3 | 0.17075 | 0.12788 | 0.17725 | 0.12788 | 29.8811 | 0.02 | 0 | Neutral |
| testneg_7423 | 0 | -0.32 | -0.556879 | -3 | 0.704598 | -3 | 0.07007 | 0.12788 | 0.09657 | 0.58605 | 10.22898 | 0.080968879 | 0 | Neutral |
| testneg_7424 | 0 | 0 | -0.064795 | -3 | 1.394643 | -3 | 0.4566 | 0.14201 | 0.49629 | 0.11989 | 20.89731 | 0.02 | 0 | Neutral |
| testneg_7425 | 0.07 | 0.38 | -1.962835 | -3 | 0.787669 | -3 | 0.40981 | 0.21807 | 0.09332 | 0.53331 | 16.84579 | 0.180265482 | 0 | Neutral |
| testneg_7426 | 0.15 | -0.28 | 1.899889 | -3 | 2.278399 | -3 | 0.18165 | 0.49827 | 0.26489 | 0.10549 | 26.40523 | 0.11 | 0 | Neutral |
| testneg_7427 | -0.03 | 1.03 | 4.336137 | -3 | 0.521728 | -3 | 0.19898 | 0.12788 | 0.05079 | 0.0924 | 59.93728 | 0.10198418 | 0 | Neutral |
| testneg_7428 | 0 | 0 | 2.676827 | -3 | 2.632596 | -3 | 0.29362 | 0.22177 | 0.9662 | 0.28104 | 11.83017 | 0.292879855 | 2.6 | Neutral |
| testneg_7429 | -0.02 | 0.68 | 3.109082 | -3 | 1.30203 | -3 | 0.31244 | 0.4615 | 0.20084 | 0.0851 | 17.36842 | 0.11 | 0 | Neutral |
| testneg_7430 | 0 | 0 | -0.882198 | -3 | 2.290365 | -3 | 0.11458 | 0.16496 | 0.24087 | 0.27985 | 10.3567 | 0.129460784 | 0 | Neutral |
| testneg_7431 | 0 | 0 | 2.100416 | -3 | 0.727793 | -3 | 0.22255 | 0.10057 | 0.38184 | 0.10478 | 18.99301 | 0 | 0 | Neutral |
| testneg_7432 | 0.07 | -0.31 | 1.843162 | -3 | 3.592336 | -3 | 0.30869 | 0.11948 | 0.03297 | 0.16627 | 10.83333 | 0.266925032 | 1 | Neutral |
| testneg_7433 | 0.18 | -1.05 | 5.645081 | -3 | 1.891537 | -3 | 0.30739 | 0.16306 | 0.12077 | 0.11244 | 30.10345 | 0.220496975 | 0 | Neutral |
| testneg_7434 | 0 | 0 | 1.377039 | -3 | 1.334446 | 1.334446 | 0.81838 | 0.09725 | 0.15514 | 0.14852 | 18.62552 | 0.050923295 | 0 | Neutral |
| testneg_7435 | 0.09 | -0.3 | 3.748574 | -3 | 3.519671 | 3.519671 | 0.19898 | 0.12788 | 0.18122 | 0.16895 | 19.05755 | 0.604226302 | 88.6 | Disease_causing |
| testneg_7436 | 0 | 0 | 0.953769 | -3 | 9.170362 | -3 | 0.15041 | 0.43468 | 0.1144 | 0.12788 | 15.56452 | 0.51 | 59.6 | Disease_causing |
| testneg_7437 | 0.04 | -0.23 | 1.479006 | -3 | 2.75642 | -3 | 0.19898 | 0.09322 | 0.08723 | 0.09503 | 53.57357 | 0.040555556 | 0 | Neutral |
| testneg_7438 | 0 | 0 | 0.740874 | 0.740874 | 5.831261 | -3 | 0.1424 | 0.12788 | 0.22645 | 0.1704 | 17.18626 | 0.44794697 | 35.4 | Neutral |
| testneg_7439 | 0.04 | -0.23 | 7.386822 | -3 | -2.765311 | -3 | 0.20249 | 0.10327 | 0.09902 | 0.12788 | 18.33333 | 0.26 | 2.8 | Neutral |
| testneg_7440 | -0.14 | 0.69 | 2.798182 | 2.798182 | -0.132461 | -3 | 0.31043 | 0.09688 | 0.16002 | 0.10481 | 18.64268 | 0.3 | 3.4 | Neutral |
| testneg_7441 | 0 | 0 | 0.308908 | -3 | -0.640729 | -0.640729 | 0.39305 | 0.18672 | 0.10107 | 0.09297 | 11.875 | 0.02 | 0 | Neutral |

| | | | | | | | | | | | | | | |
|---|---|---|---|---|---|---|---|---|---|---|---|---|---|---|
| testneg_7442 | 0.05 | -0.81 | 3.374642 | -3 | 3.124639 | -3 | 0.33264 | 0.10493 | 0.30585 | 0.10254 | 33.13136 | 0.097897272 | 0 | Neutral |
| testneg_7443 | 0 | -0.01 | 2.462741 | -3 | 6.381515 | -3 | 0.11858 | 0.11551 | 0.1078 | 0.17394 | 22.38311 | 0.482633877 | 46.6 | Neutral |
| testneg_7444 | -0.07 | -0.33 | 4.046311 | -3 | 1.27566 | -3 | 0.46242 | 0.39055 | 0.18535 | 0.12788 | 12.29167 | 0.129733947 | 0 | Neutral |
| testneg_7445 | -0.1 | -1.9 | 0.050482 | -3 | 3.108369 | -3 | 0.37065 | 0.12788 | 0.16424 | 0.11262 | 17.97975 | 0.08 | 0 | Neutral |
| testneg_7446 | 0.01 | -0.57 | 1.464871 | -3 | -2.166097 | -3 | 0.23809 | 0.18937 | 0.41269 | 0.19135 | 19.44444 | 0.121998028 | 0 | Neutral |
| testneg_7447 | -0.02 | -0.41 | 1.310214 | -3 | 4.259881 | -3 | 0.65012 | 0.21208 | 0.38756 | 0.08351 | 12.59789 | 0.13 | 0 | Neutral |
| testneg_7448 | 0.14 | -1.01 | 6.557595 | -3 | 6.42687 | -3 | 0.44816 | 0.56655 | 0.16098 | 0.12788 | 14.22491 | 0.818976496 | 100 | Disease_causing |
| testneg_7449 | -0.03 | -0.1 | 0.968512 | -3 | 0.838864 | -3 | 0.40074 | 0.36296 | 0.10285 | 0.10186 | 28.00377 | 0.052335498 | 0 | Neutral |
| testneg_7450 | 0.01 | 0.79 | 0.082922 | -3 | -1.760419 | -3 | 0.81384 | 0.12094 | 0.07071 | 0.12788 | 18.18061 | 0.03 | 0 | Neutral |
| testneg_7451 | 0 | 0 | 0.759983 | -3 | 2.11243 | -3 | 0.13505 | 0.09019 | 0.12782 | 0.08502 | 9.74181 | 0.01 | 0 | Neutral |
| testneg_7452 | 0.05 | 0.29 | 0.722422 | -3 | 0.741645 | -3 | 0.32669 | 0.10659 | 0.04606 | 0.20522 | 18.88889 | 0.066659141 | 0 | Neutral |
| testneg_7453 | 0.16 | -0.37 | -2.713571 | -2.713571 | -0.676534 | -0.676534 | 0.07506 | 0.09112 | 0.16262 | 0.15786 | 46.32434 | 0.1 | 0 | Neutral |
| testneg_7454 | 0 | 0 | 1.860726 | -3 | -3.291659 | -3 | 0.5901 | 0.19444 | 0.11957 | 0.08975 | 10.66756 | 0.02 | 0 | Neutral |
| testneg_7455 | 0 | 0 | 1.883707 | -3 | 4.231687 | -3 | 0.07427 | 0.09451 | 0.09616 | 0.11651 | 20.90229 | 0.07 | 0 | Neutral |
| testneg_7456 | -0.02 | 0.28 | 2.201231 | -3 | 0.133669 | -3 | 0.20231 | 0.12788 | 0.08353 | 0.21479 | 12.11131 | 0.078306452 | 0 | Neutral |
| testneg_7457 | 0 | -0.01 | 2.125818 | -3 | 1.865393 | -3 | 0.15713 | 0.3507 | 0.19898 | 0.10482 | 59.66883 | 0.110043103 | 0 | Neutral |
| testneg_7458 | 0 | 0 | 1.545946 | -3 | 0.192071 | -3 | 0.03844 | 0.12788 | 0.18312 | 0.12788 | 13.51351 | 0.020161332 | 0 | Neutral |
| testneg_7459 | 0 | 0 | 2.25618 | -3 | -1.652996 | -3 | 0.19898 | 0.09316 | 0.80264 | 0.11567 | 10.41667 | 0.01 | 0 | Neutral |
| testneg_7460 | 0 | -0.32 | 2.54814 | -3 | 1.367101 | -3 | 0.85592 | 0.20297 | 0.99765 | 0.95064 | 10.41667 | 0.202837147 | 0 | Neutral |
| testneg_7461 | -0.03 | -0.1 | 5.108849 | -3 | -1.249686 | -3 | 0.49064 | 0.12294 | 0.15685 | 0.17839 | 12.71686 | 0.269932432 | 1.2 | Neutral |
| testneg_7462 | 0.18 | -1.05 | 0.965486 | -3 | 3.992072 | -3 | 0.11114 | 0.09582 | 0.05767 | 0.07325 | 18.6768 | 0.1 | 0 | Neutral |
| testneg_7463 | -0.05 | -0.29 | -0.623218 | -3 | 0.002903 | -3 | 0.26997 | 0.63944 | 0.28162 | 0.12788 | 27.95796 | 0.075 | 0 | Neutral |
| testneg_7464 | 0.16 | -0.37 | 5.06041 | -3 | 0.647985 | -3 | 0.81054 | 0.28104 | 0.21728 | 0.1132 | 18.33333 | 0.12 | 0 | Neutral |
| testneg_7465 | 0 | 0 | 1.260562 | -3 | -0.157598 | -3 | 0.04393 | 0.12788 | 0.14432 | 0.22579 | 16.56191 | 0.080708658 | 0 | Neutral |
| testneg_7466 | 0.18 | -1.05 | 4.454666 | -3 | 1.533148 | -3 | 0.09462 | 0.11807 | 0.19898 | 0.13249 | 23.81804 | 0.13 | 0 | Neutral |
| testneg_7467 | 0.23 | -1.04 | -1.468734 | -3 | 0.820792 | -3 | 0.06789 | 0.09707 | 0.11301 | 0.10716 | 20.22059 | 0.041116939 | 0 | Neutral |
| testneg_7468 | 0.07 | -0.31 | -0.344853 | -3 | 1.867073 | -3 | 0.50276 | 0.09638 | 0.07999 | 0.23927 | 21.2963 | 0.12 | 0 | Neutral |
| testneg_7469 | 0 | -0.32 | 0.878488 | -3 | 5.669528 | -3 | 0.11359 | 0.08631 | 0.28817 | 0.10531 | 13.2927 | 0.08 | 0 | Neutral |
| testneg_7470 | 0 | -0.01 | 1.132093 | -3 | 3.877522 | -3 | 0.26372 | 0.12788 | 0.6854 | 0.12563 | 16.85192 | 0.046388889 | 0 | Neutral |
| testneg_7471 | -0.01 | 0.57 | 5.88281 | -3 | 0.161355 | -3 | 0.17322 | 0.10311 | 0.28227 | 0.24183 | 22.08736 | 0.39 | 15.4 | Neutral |
| testneg_7472 | 0.01 | -0.57 | 2.700261 | -3 | 0.977149 | -3 | 0.76957 | 0.13588 | 0.31043 | 0.09688 | 10.41667 | 0.03 | 0 | Neutral |
| testneg_7473 | 0 | 0 | -0.009858 | -3 | 0.976803 | -3 | 0.31664 | 0.14181 | 0.50687 | 0.11283 | 11.25 | 0.01 | 0 | Neutral |
| testneg_7474 | 0.12 | -1.16 | 4.936092 | -3 | 3.729393 | -3 | 0.2118 | 0.2009 | 0.85592 | 0.20297 | 18.33333 | 0.672658009 | 95.6 | Disease_causing |
| testneg_7475 | 0 | 0 | 1.431446 | -3 | 0.240038 | -3 | 0.23393 | 0.12567 | 0.19898 | 0.08111 | 10.41667 | 0 | 0 | Neutral |
| testneg_7476 | 0 | 0 | 2.100416 | -3 | 6.639974 | -3 | 0.22255 | 0.10057 | 0.58985 | 0.10504 | 21.95775 | 0.15 | 0.2 | Neutral |
| testneg_7477 | 0 | 0.32 | -0.420859 | -3 | -0.106366 | -3 | 0.0597 | 0.18092 | 0.19898 | 0.12788 | 10 | 0.05 | 0 | Neutral |
| testneg_7478 | 0 | 0 | 0.063311 | -3 | 4.488254 | -3 | 0.17404 | 0.12788 | 0.19898 | 0.08855 | 13.75 | 0.03 | 0 | Neutral |
| testneg_7479 | 0 | 0 | 0.819832 | -3 | -0.229991 | -3 | 0.19898 | 0.12788 | 0.19698 | 0.1459 | 18.36243 | 0.020633803 | 0 | Neutral |
| testneg_7480 | -0.01 | -0.8 | 2.675128 | -3 | 2.031061 | -3 | 0.19898 | 0.12788 | 0.36676 | 0.37749 | 18.21456 | 0.177673388 | 0 | Neutral |
| testneg_7481 | -0.02 | -0.13 | 1.214257 | -3 | 2.02554 | -3 | 0.43459 | 0.12595 | 0.21331 | 0.09513 | 4.28195 | 0.010084656 | 0 | Neutral |
| testneg_7482 | 0 | 0 | -0.018631 | -3 | -0.116772 | -3 | 0.12596 | 0.18924 | 0.16741 | 0.10762 | 22.8648 | 0.028452381 | 0 | Neutral |
| testneg_7483 | 0.14 | -0.37 | 5.710272 | -3 | 0.982052 | -3 | 0.508 | 0.1281 | 0.09211 | 0.08973 | 18.88889 | 0.1575 | 0 | Neutral |
| testneg_7484 | -0.18 | -0.16 | 0.910161 | -3 | -1.330673 | -3 | 0.93468 | 0.12788 | 0.03137 | 0.12788 | 28.59649 | 0.06 | 0 | Neutral |
| testneg_7485 | 0.22 | -2.66 | 2.450036 | -3 | -0.106366 | -3 | 0.09997 | 0.1797 | 0.19898 | 0.12788 | 21.81093 | 0.1 | 0 | Neutral |
| testneg_7486 | -0.05 | -0.29 | 1.310845 | -3 | 0.514472 | -3 | 0.19898 | 0.12788 | 0.19898 | 0.11185 | 28.851 | 0 | 0 | Neutral |
| testneg_7487 | 0 | 0 | 1.074253 | -3 | 6.71239 | -3 | 0.18233 | 0.14781 | 0.68568 | 0.65219 | 18.88889 | 0.524924477 | 64.4 | Disease_causing |
| testneg_7488 | 0.09 | -0.3 | 5.573517 | -3 | 0.072482 | -3 | 0.09416 | 0.17821 | 0.07394 | 0.09408 | 11.83858 | 0.2 | 0.2 | Neutral |
| testneg_7489 | 0 | 0 | 0.007077 | -3 | 1.01324 | -3 | 0.75008 | 0.70645 | 0.45354 | 0.09957 | 12.29167 | 0.021325328 | 0 | Neutral |
| testneg_7490 | -0.02 | 0.36 | 2.655425 | -3 | 1.614723 | 1.614723 | 0.19883 | 0.09434 | 0.74944 | 0.1263 | 9.862 | 0.04052662 | 0 | Neutral |
| testneg_7491 | -0.07 | -0.01 | -0.474074 | -3 | 4.236143 | -3 | 0.06419 | 0.35525 | 0.02723 | 0.04872 | 18.33333 | 0.20042507 | 0 | Neutral |
| testneg_7492 | 0 | 0 | 0.452231 | -3 | 0.054796 | -3 | 0.46691 | 0.07886 | 0.32531 | 0.12351 | 19.68594 | 0.01 | 0 | Neutral |
| testneg_7493 | 0 | 0 | 1.394643 | -3 | 0.402508 | 0.402508 | 0.49629 | 0.11989 | 0.29794 | 0.11106 | 22.1732 | 0.010458333 | 0 | Neutral |
| testneg_7494 | 0.04 | -0.23 | -0.806907 | -3 | 5.955249 | -3 | 0.09967 | 0.15417 | 0.26842 | 0.09683 | 16.18123 | 0.234398268 | 0.2 | Neutral |
| testneg_7495 | 0 | 0 | 1.626934 | -3 | 2.272008 | -3 | 0.20942 | 0.14229 | 0.23913 | 0.12788 | 18.69495 | 0.01 | 0 | Neutral |
| testneg_7496 | 0 | 0 | 1.108803 | -3 | 2.996727 | -3 | 0.85092 | 0.58871 | 0.19898 | 0.17518 | 20 | 0.238312125 | 0.2 | Neutral |
| testneg_7497 | 0.05 | -0.81 | 6.33079 | -3 | 2.97409 | -3 | 0.31906 | 0.1705 | 0.16544 | 0.43246 | 13.10079 | 0.784034264 | 99.6 | Disease_causing |
| testneg_7498 | -0.06 | -0.18 | -0.549376 | -3 | 6.179448 | -3 | 0.05904 | 0.09282 | 0.17296 | 0.09548 | 43.93631 | 0.153333333 | 0 | Neutral |
| testneg_7499 | -0.03 | -1.22 | 6.280508 | -3 | 2.08992 | -3 | 0.47166 | 0.13548 | 0.69254 | 0.10021 | 19.19724 | 0.343666667 | 5.8 | Neutral |
| testneg_7500 | 0 | 0 | 0.864288 | -3 | 5.603216 | -3 | 0.21014 | 0.12378 | 0.06112 | 0.06901 | 10.83333 | 0.08 | 0 | Neutral |
| testneg_7501 | 0.01 | 0.79 | 0.585649 | 0.585649 | 8.488711 | -3 | 0.07185 | 0.68732 | 0.19102 | 0.12788 | 13.51351 | 0.55 | 73 | Disease_causing |
| testneg_7502 | -0.03 | 0.85 | -0.309243 | -3 | 1.159886 | -3 | 0.03137 | 0.12788 | 0.19898 | 0.12788 | 49.06907 | 0.05 | 0 | Neutral |
| testneg_7503 | 0.15 | -1.06 | 5.940594 | -3 | 1.397648 | -3 | 0.71803 | 0.95763 | 0.11006 | 0.20781 | 10.83333 | 0.598610119 | 85.4 | Disease_causing |
| testneg_7504 | -0.05 | -0.29 | -1.010566 | -3 | 5.967322 | 5.967322 | 0.39192 | 0.09135 | 0.25312 | 0.12788 | 19.98461 | 0.390954545 | 12.8 | Neutral |
| testneg_7505 | 0 | 0 | 1.537848 | -3 | 0.649412 | -3 | 0.11391 | 0.12788 | 0.3394 | 0.5826 | 10.22898 | 0.030353714 | 0 | Neutral |
| testneg_7506 | 0.14 | 0.54 | 3.022433 | -3 | 3.376618 | -3 | 0.09179 | 0.12788 | 0.21884 | 0.16743 | 10.22898 | 0.425654762 | 24.6 | Neutral |
| testneg_7507 | 0.21 | 0.15 | 6.458942 | -3 | 1.75958 | -3 | 0.0978 | 0.26338 | 0.65897 | 0.67602 | 10.41667 | 0.574272727 | 79.4 | Disease_causing |
| testneg_7508 | -0.02 | 0.63 | 2.293151 | -3 | 4.228879 | -3 | 0.47997 | 0.10506 | 0.32458 | 0.1016 | 11.90227 | 0.117083333 | 0 | Neutral |
| testneg_7509 | 0.05 | -0.81 | 0.592404 | -3 | -2.289069 | -3 | 0.19494 | 0.14451 | 0.75906 | 0.09702 | 19.25467 | 0.03 | 0 | Neutral |
| testneg_7510 | 0 | 0 | 0.419995 | -3 | -0.496432 | -3 | 0.7174 | 0.15815 | 0.06247 | 0.14184 | 10.41667 | 0.060050505 | 0 | Neutral |
| testneg_7511 | 0 | 0 | 0.014242 | 0.014242 | -2.012697 | -3 | 0.07387 | 0.12788 | 0.08027 | 0.12788 | 16.03856 | 0.01 | 0 | Neutral |
| testneg_7512 | 0 | -0.32 | -0.748689 | -3 | -1.73577 | -3 | 0.07007 | 0.12788 | 0.0931 | 0.12788 | 34.48699 | 0.02 | 0 | Neutral |
| testneg_7513 | -0.02 | 0.63 | 0.385855 | -3 | 6.65036 | -3 | 0.10153 | 0.13343 | 0.30367 | 0.41964 | 17.21847 | 0.519254762 | 61.8 | Disease_causing |
| testneg_7514 | 0 | 0 | 2.594251 | -3 | 6.156103 | -3 | 0.2185 | 0.20593 | 0.19898 | 0.10335 | 11.98582 | 0.252800871 | 1.4 | Neutral |
| testneg_7515 | 0.05 | 0.29 | 3.485507 | -3 | 0.336988 | -3 | 0.29417 | 0.10965 | 0.1193 | 0.5592 | 13.25026 | 0.159551708 | 0 | Neutral |
| testneg_7516 | -0.03 | -0.1 | 2.059294 | -3 | 0.551288 | -3 | 0.66711 | 0.31094 | 0.83421 | 0.18817 | 21.85185 | 0.135 | 0 | Neutral |
| testneg_7517 | 0.02 | -0.36 | 2.305765 | -3 | 5.851071 | -3 | 0.11055 | 0.1127 | 0.32022 | 0.09529 | 22.63566 | 0.131270012 | 0 | Neutral |
| testneg_7518 | -0.03 | -0.1 | 2.260957 | 2.260957 | 2.549873 | -3 | 0.36054 | 0.12788 | 0.04484 | 0.95426 | 19.98891 | 0.326571429 | 7.4 | Neutral |
| testneg_7519 | 0.07 | 0.38 | 0.348921 | -3 | 1.775106 | -3 | 0.47379 | 0.18145 | 0.88976 | 0.22648 | 16.668 | 0.10775 | 0 | Neutral |
| testneg_7520 | 0 | -0.32 | 0.641608 | -3 | 2.925716 | -3 | 0.32458 | 0.1016 | 0.13906 | 0.09925 | 20.57907 | 0.021423266 | 0 | Neutral |
| testneg_7521 | 0.21 | 0.15 | 0.571501 | -3 | -0.782144 | -3 | 0.087 | 0.12788 | 0.34946 | 0.19092 | 25.89685 | 0.095595238 | 0 | Neutral |
| testneg_7522 | -0.07 | -0.01 | 2.629838 | -3 | 1.849894 | 1.653477 | 0.09099 | 0.2051 | 0.13458 | 0.17634 | 27.10267 | 0.270524367 | 0.2 | Neutral |
| testneg_7523 | 0 | 0 | 0.781134 | -3 | 2.629838 | -3 | 0.92237 | 0.11252 | 0.09099 | 0.2051 | 18.2504 | 0.113214286 | 0 | Neutral |
| testneg_7524 | 0.04 | -0.23 | 3.599958 | -3 | 0.860716 | -3 | 0.40491 | 0.17596 | 0.57729 | 0.10831 | 10.83333 | 0.057083333 | 0 | Neutral |
| testneg_7525 | 0 | 0 | 1.657251 | -3 | 1.712042 | -3 | 0.17973 | 0.12278 | 0.31004 | 0.09727 | 12.29167 | 0 | 0 | Neutral |
| testneg_7526 | 0.07 | 0.7 | 1.384923 | -3 | 0.058581 | -3 | 0.12875 | 0.10115 | 0.16791 | 0.22163 | 19.44444 | 0.06 | 0 | Neutral |
| testneg_7527 | 0 | 0 | 1.165089 | -3 | 0.480843 | -3 | 0.32989 | 0.13525 | 0.19898 | 0.12788 | 9.47972 | 3.50E-05 | 0 | Neutral |
| testneg_7528 | 0.07 | -0.31 | 4.193501 | -3 | 4.100398 | -3 | 0.20737 | 0.11262 | 0.19898 | 0.09395 | 22.93219 | 0.130892424 | 0 | Neutral |
| testneg_7529 | -0.02 | 0.68 | 1.913159 | -3 | 0.728537 | -3 | 0.91921 | 0.32331 | 0.08521 | 0.11239 | 11.25 | 0.045393116 | 0 | Neutral |
| testneg_7530 | -0.03 | -0.12 | -0.214687 | -3 | 0.818598 | -3 | 0.19668 | 0.16042 | 0.12739 | 0.12788 | 13.51351 | 0.021163618 | 0 | Neutral |
| testneg_7531 | 0 | 0 | -0.285925 | -3 | -1.427787 | -3 | 0.09645 | 0.13239 | 0.11261 | 0.13673 | 18.46739 | 0.08 | 0 | Neutral |
| testneg_7532 | 0.09 | -0.3 | 2.911406 | -3 | 2.125139 | -3 | 0.25421 | 0.16235 | 0.25119 | 0.13626 | 40.503 | 0.250356385 | 1 | Neutral |
| testneg_7533 | 0 | 0 | 0.616989 | -3 | 0.057569 | -3 | 0.42802 | 0.12096 | 0.05315 | 0.12788 | 28.61071 | 0.015263158 | 0 | Neutral |
| testneg_7534 | 0 | 0 | 0.907583 | -3 | 1.671519 | -3 | 0.9327 | 0.12788 | 0.07837 | 0.12788 | 8.86674 | 0.01 | 0 | Neutral |
| testneg_7535 | 0.19 | 0.51 | 4.472767 | -3 | 0.278169 | -3 | 0.12853 | 0.08565 | 0.10963 | 0.12788 | 15.23133 | 0.06 | 0 | Neutral |
| testneg_7536 | 0.03 | 0.1 | 1.113931 | -3 | 0.345146 | -3 | 0.16134 | 0.1847 | 0.20344 | 0.1705 | 6.03908 | 0.081810932 | 0 | Neutral |
| testneg_7537 | 0.00136364 | -0.048 | 1.894277 | -3 | 3.39754 | -3 | 0.53216 | 0.57092 | 0.62203 | 0.15583 | 4.44444 | 0.331253227 | 7.2 | Neutral |
| testneg_7538 | 0.09 | -2.17 | 0.344836 | -3 | 5.955249 | -3 | 0.22575 | 0.37247 | 0.26842 | 0.09683 | 18.03534 | 0.253533333 | 0.4 | Neutral |
| testneg_7539 | 0.14 | -0.69 | 2.950487 | -3 | 2.084003 | -3 | 0.14495 | 0.37127 | 0.10285 | 0.10186 | 40.15723 | 0.17 | 0 | Neutral |
| testneg_7540 | 0 | 0 | -0.39724 | -3 | 5.966703 | -3 | 0.09749 | 0.82924 | 0.19898 | 0.12788 | 20.83333 | 0.270151094 | 0.8 | Neutral |
| testneg_7541 | -0.07 | -0.38 | 5.028164 | -3 | -0.354552 | -3 | 0.29209 | 0.09791 | 0.09815 | 0.12788 | 22.89111 | 0.070453634 | 0 | Neutral |
| testneg_7542 | 0.16 | 0.08 | -0.169987 | -3 | 3.97741 | -3 | 0.99445 | 0.10396 | 0.65981 | 0.47389 | 17.90495 | 0.293513889 | 1 | Neutral |
| testneg_7543 | -0.04 | 0.23 | 4.21286 | -3 | -0.360768 | -0.360768 | 0.07447 | 0.23492 | 0.47147 | 0.09503 | 14.39044 | 0.12 | 0 | Neutral |
| testneg_7544 | 0 | 0 | 1.833275 | -3 | -0.392743 | -3 | 0.40981 | 0.21807 | 0.16275 | 0.12788 | 17.77778 | 0.011666667 | 0 | Neutral |
| testneg_7545 | 0 | 0 | 0.628811 | -3 | 0.196354 | -3 | 0.07874 | 0.12788 | 0.18067 | 0.14759 | 9.32594 | 0.023947085 | 0 | Neutral |
| testneg_7546 | -0.02 | -0.45 | -0.208207 | -3 | 3.40369 | 3.381638 | 0.07219 | 0.12788 | 0.07552 | 0.12788 | 35.5603 | 0.19 | 0 | Neutral |
| testneg_7547 | 0 | 0 | 0.968047 | -3 | 0.851258 | -0.771274 | 0.30823 | 0.17188 | 0.09306 | 0.12788 | 18.33333 | 0.01 | 0 | Neutral |
| testneg_7548 | 0 | 0 | 0.71683 | -3 | 0.947918 | -3 | 0.42583 | 0.10901 | 0.25236 | 0.11809 | 10.30286 | 9.86E-05 | 0 | Neutral |
| testneg_7549 | 0 | 0 | 1.797648 | -3 | 4.404174 | -3 | 0.30275 | 0.12788 | 0.09828 | 0.07733 | 11.01208 | 0.049016486 | 0 | Neutral |
| testneg_7550 | -0.05 | -0.29 | -0.695099 | -3 | 0.700986 | -3 | 0.84318 | 0.11313 | 0.694 | 0.09974 | 13.51351 | 0.010235327 | 0 | Neutral |
| testneg_7551 | 0.14 | -0.69 | 5.926215 | -3 | 6.244312 | 2.300553 | 0.48027 | 0.52799 | 0.06552 | 0.07762 | 10.41667 | 0.61 | 89 | Disease_causing |
| testneg_7552 | 0 | 0.32 | 5.974356 | -3 | 4.457412 | -3 | 0.11187 | 0.10744 | 0.19898 | 0.11691 | 15.34045 | 0.357501299 | 13.4 | Neutral |
| testneg_7553 | -0.14 | 1.01 | 6.502902 | -3 | 0.573225 | -3 | 0.37056 | 0.12309 | 0.73947 | 0.2058 | 25.56326 | 0.378287037 | 11 | Neutral |
| testneg_7554 | 0.01 | 0.79 | -2.775113 | -2.775113 | 4.092943 | -3 | 0.13295 | 0.12914 | 0.83421 | 0.18817 | 26.92625 | 0.255152987 | 0.6 | Neutral |
| testneg_7555 | 0.21 | 0.15 | 6.256857 | -3 | 1.674048 | -3 | 0.65674 | 0.10572 | 0.25987 | 0.31819 | 29.1844 | 0.410041667 | 22.2 | Neutral |
| testneg_7556 | 0 | 0 | 1.081416 | -3 | 0.695636 | 0.695636 | 0.59345 | 0.26484 | 0.12003 | 0.12788 | 41.63155 | 0.036325188 | 0 | Neutral |
| testneg_7557 | 0 | 0 | 1.145768 | -3 | 1.135659 | -3 | 0.22658 | 0.20646 | 0.21303 | 0.09348 | 10 | 0.011867036 | 0 | Neutral |
| testneg_7558 | -0.19 | -0.51 | 3.156196 | -3 | 5.815666 | -3 | 0.34225 | 0.12788 | 0.34109 | 0.2167 | 11.25 | 0.628293799 | 90.6 | Disease_causing |
| testneg_7559 | 0.15 | -0.28 | 1.393737 | -3 | 0.581346 | -3 | 0.34288 | 0.24605 | 0.20222 | 0.12788 | 11.98582 | 0.030294372 | 0 | Neutral |
| testneg_7560 | -0.03 | -0.1 | 1.738251 | -3 | 1.232806 | -3 | 0.17303 | 0.18733 | 0.49641 | 0.12614 | 12.29167 | 0.03 | 0 | Neutral |
| testneg_7561 | -0.23 | 1.04 | -1.164424 | -3 | 4.076957 | 4.076957 | 0.12073 | 0.10969 | 0.38324 | 0.25193 | 10 | 0.51 | 60 | Disease_causing |
| testneg_7562 | 0 | 0 | 0.308908 | -3 | 1.016479 | 1.016479 | 0.39305 | 0.18672 | 0.12649 | 0.12788 | 5.54414 | 0.02 | 0 | Neutral |
| testneg_7563 | -0.03 | -0.12 | 1.051313 | -3 | 1.1545 | -3 | 0.45029 | 0.12788 | 0.09602 | 0.10726 | 25.90643 | 0.011125 | 0 | Neutral |

| | | | | | | | | | | | | | | |
|---|---|---|---|---|---|---|---|---|---|---|---|---|---|---|
| testneg_7564 | 0 | 0 | 0.905163 | -3 | 1.810934 | -3 | 0.15267 | 0.12788 | 0.5184 | 0.11889 | 26.83387 | 0.000555556 | 0 | Neutral |
| testneg_7565 | -0.07 | -0.38 | 1.102328 | -3 | 3.437388 | -3 | 0.74271 | 0.10951 | 0.22645 | 0.1704 | 14.47772 | 0.181663551 | 0 | Neutral |
| testneg_7566 | 0 | 0 | 0.12874 | -3 | 0.258063 | -3 | 0.79692 | 0.51979 | 0.99941 | 0.50288 | 10.41667 | 0.111547619 | 0 | Neutral |
| testneg_7567 | 0 | 0 | 0.628811 | -3 | -0.072035 | -3 | 0.07874 | 0.12788 | 0.28227 | 0.24183 | 20.93194 | 0.05465873 | 0 | Neutral |
| testneg_7568 | 0 | -0.01 | 1.806927 | -3 | -0.341501 | -3 | 0.17973 | 0.12278 | 0.75008 | 0.70645 | 11.66689 | 0.031062271 | 0 | Neutral |
| testneg_7569 | 0.09 | -0.3 | 2.387342 | -3 | 5.494664 | -3 | 0.07428 | 0.52467 | 0.19898 | 0.12788 | 11.99102 | 0.361217949 | 6.8 | Neutral |
| testneg_7570 | 0 | 0 | -0.4932 | -3 | -0.058596 | -3 | 0.20379 | 0.43705 | 0.1144 | 0.09798 | 17.21847 | 0.023337852 | 0 | Neutral |
| testneg_7571 | 0.15 | -0.28 | 2.534333 | -3 | 3.832241 | 3.832241 | 0.19898 | 0.12788 | 0.213 | 0.09535 | 19.30471 | 0.270511509 | 0.4 | Neutral |
| testneg_7572 | 0 | 0 | -0.180819 | -3 | 1.294927 | -3 | 0.15155 | 0.09923 | 0.35486 | 0.11584 | 10.35388 | 0.000160256 | 0 | Neutral |
| testneg_7573 | 0 | 0 | -1.173562 | -3 | 2.135735 | -3 | 0.64661 | 0.49283 | 0.62273 | 0.10215 | 17.32224 | 0.04 | 0 | Neutral |
| testneg_7574 | 0 | 0 | 0.916236 | -3 | 0.299382 | -3 | 0.37499 | 0.17803 | 0.75728 | 0.34838 | 4.72222 | 0.040187063 | 0 | Neutral |
| testneg_7575 | 0 | 0 | 0.027672 | -3 | 5.775288 | -3 | 0.16091 | 0.90971 | 0.16663 | 0.12788 | 22.77778 | 0.239666667 | 0.2 | Neutral |
| testneg_7576 | -0.05 | -0.29 | 3.327164 | -3 | 0.108057 | -3 | 0.31124 | 0.08875 | 0.30367 | 0.41964 | 10.41667 | 0.149754943 | 0 | Neutral |
| testneg_7577 | 0 | -0.01 | 1.129155 | -3 | 4.894604 | -3 | 0.16835 | 0.09282 | 0.21726 | 0.23214 | 13.75 | 0.189013882 | 0.4 | Neutral |
| testneg_7578 | -0.05 | -0.29 | -0.695099 | -3 | 1.464574 | 1.464574 | 0.84318 | 0.11313 | 0.60603 | 0.12788 | 25.60245 | 0.03 | 0 | Neutral |
| testneg_7579 | 0 | 0 | 0.572592 | -3 | 2.598788 | -3 | 0.68918 | 0.16819 | 0.19898 | 0.10307 | 29.58333 | 0.050215196 | 0 | Neutral |
| testneg_7580 | -0.06 | -0.18 | 0.603239 | -3 | 3.009639 | -3 | 0.08684 | 0.08917 | 0.19452 | 0.79779 | 10.22898 | 0.188422626 | 0 | Neutral |
| testneg_7581 | -0.03 | -1.22 | 6.195353 | -3 | 1.140696 | -3 | 0.28532 | 0.10663 | 0.52768 | 0.12788 | 13.14653 | 0.22 | 0.6 | Neutral |
| testneg_7582 | 0.03 | 0.1 | -0.187467 | -0.187467 | -0.002747 | -3 | 0.12314 | 0.11709 | 0.07394 | 0.09408 | 16.82896 | 0.01 | 0 | Neutral |
| testneg_7583 | 0 | 0 | 1.891407 | -3 | 2.947434 | -3 | 0.08136 | 0.10647 | 0.27915 | 0.27706 | 18.47729 | 0.132460122 | 0 | Neutral |
| testneg_7584 | 0.07 | 0.38 | -1.829281 | -3 | 4.820901 | 4.820901 | 0.0984 | 0.1489 | 0.06358 | 0.5364 | 22.0041 | 0.59253252 | 87 | Disease_causing |
| testneg_7585 | 0.15 | -0.82 | 3.191701 | -3 | 1.467396 | -3 | 0.67038 | 0.1794 | 0.96995 | 0.66487 | 11.25 | 0.29 | 1.6 | Neutral |
| testneg_7586 | -0.05 | -0.29 | 2.477566 | -3 | 2.061627 | -3 | 0.29451 | 0.13582 | 0.13271 | 0.22958 | 20.04742 | 0.142329719 | 0 | Neutral |
| testneg_7587 | 0.18 | 0.16 | 1.885526 | -3 | 4.243038 | -3 | 0.2141 | 0.19929 | 0.4719 | 0.33344 | 10 | 0.629463264 | 95.2 | Disease_causing |
| testneg_7588 | 0.04 | -0.23 | 2.52259 | -3 | 4.259293 | 3.852699 | 0.5127 | 0.12263 | 0.15848 | 0.12754 | 11.25 | 0.310964286 | 0.8 | Neutral |
| testneg_7589 | 0.15 | 0.06 | 1.20059 | -3 | 1.26451 | -3 | 0.24 | 0.2063 | 0.11439 | 0.21573 | 20.64599 | 0.206650327 | 0 | Neutral |
| testneg_7590 | 0 | 0 | 1.015389 | -3 | 2.445411 | -3 | 0.37711 | 0.12788 | 0.22179 | 0.09831 | 18.6768 | 0.01 | 0 | Neutral |
| testneg_7591 | 0.09 | -0.3 | 1.031921 | -3 | 5.7095 | -3 | 0.96704 | 0.20571 | 0.93064 | 0.10933 | 5 | 0.21 | 0.2 | Neutral |
| testneg_7592 | 0 | -0.01 | 1.202769 | -3 | 1.094525 | -3 | 0.20231 | 0.12788 | 0.09838 | 0.11813 | 26.39195 | 0.001428571 | 0 | Neutral |
| testneg_7593 | 0.01 | 0.79 | 1.266152 | -3 | 0.165986 | -3 | 0.1846 | 0.375 | 0.07999 | 0.10413 | 21.49912 | 0.06 | 0 | Neutral |
| testneg_7594 | 0 | 0 | 1.123147 | -3 | 2.188986 | 1.50975 | 0.59947 | 0.14339 | 0.25209 | 0.10029 | 17.06049 | 0.02 | 0 | Neutral |
| testneg_7595 | -0.15 | 0.28 | -0.84394 | -3 | -2.464297 | -2.464297 | 0.54895 | 0.15082 | 0.11524 | 0.12788 | 10.41667 | 0.08 | 0 | Neutral |
| testneg_7596 | 0 | 0 | 0.231295 | -3 | 1.729635 | 1.729635 | 0.19898 | 0.12788 | 0.57588 | 0.09682 | 56.73177 | 0.06625 | 0 | Neutral |
| testneg_7597 | 0 | 0 | 0.961823 | -3 | 0.55724 | 0.55724 | 0.19898 | 0.37148 | 0.12937 | 0.23004 | 21.2963 | 0.200533428 | 0 | Neutral |
| testneg_7598 | 0.07 | 0.01 | -0.878719 | -3 | 0.246106 | -3 | 0.99976 | 0.28689 | 0.07297 | 0.12788 | 13.14488 | 0.07 | 0 | Neutral |
| testneg_7599 | 0.14 | -0.37 | 4.972003 | -3 | -0.197629 | -3 | 0.27843 | 0.54325 | 0.10803 | 0.14652 | 10.83333 | 0.36 | 9 | Neutral |
| testneg_7600 | -0.02 | -0.45 | 3.807977 | -3 | 3.793394 | -3 | 0.27924 | 0.12788 | 0.23955 | 0.97212 | 11.35604 | 0.487466346 | 49.6 | Neutral |
| testneg_7601 | 0 | 0 | -0.061342 | -3 | 6.520105 | -3 | 0.07626 | 0.12788 | 0.81937 | 0.22945 | 16.88659 | 0.304166667 | 6 | Neutral |
| testneg_7602 | 0.02 | 0.45 | 5.685428 | -3 | 1.447389 | 1.447389 | 0.22877 | 0.14229 | 0.14419 | 0.1668 | 13.02635 | 0.495426587 | 52.6 | Disease_causing |
| testneg_7603 | 0.09 | -2.17 | 4.797483 | -3 | 3.301794 | -3 | 0.10228 | 0.22182 | 0.75481 | 0.12788 | 34.74697 | 0.319023423 | 2.8 | Neutral |
| testneg_7604 | 0.22 | -2.66 | 4.591469 | -3 | 7.117744 | -3 | 0.18256 | 0.13499 | 0.39774 | 0.10813 | 16.85532 | 0.4925 | 53.4 | Disease_causing |
| testneg_7605 | 0 | 0 | 1.975201 | -3 | 1.669702 | -3 | 0.34495 | 0.16861 | 0.09026 | 0.09447 | 6.99956 | 0.010882218 | 0 | Neutral |
| testneg_7606 | 0.05 | -0.81 | 3.686845 | -3 | 1.737373 | -3 | 0.04508 | 0.12788 | 0.13873 | 0.12788 | 31.46572 | 0.08 | 0 | Neutral |
| testneg_7607 | 0.01 | 0.8 | 6.718863 | -3 | 2.079972 | -3 | 0.18022 | 0.12788 | 0.25549 | 0.12378 | 23.22086 | 0.3 | 6.4 | Neutral |
| testneg_7608 | 0.02 | -0.63 | 1.681173 | -3 | 3.180336 | -3 | 0.89726 | 0.10286 | 0.16361 | 0.13388 | 9.32594 | 0.193166667 | 0 | Neutral |
| testneg_7609 | -0.06 | -0.18 | 4.107901 | -3 | 2.038329 | -3 | 0.16797 | 0.11809 | 0.25159 | 0.12788 | 13.92374 | 0.040720758 | 0 | Neutral |
| testneg_7610 | 0.09 | -0.3 | 6.191692 | -3 | 2.495202 | -3 | 0.45927 | 0.74955 | 0.20327 | 0.11174 | 14.75193 | 0.47 | 43 | Neutral |
| testneg_7611 | 0 | -0.01 | 0.36151 | -3 | 0.454724 | -3 | 0.28878 | 0.16765 | 0.33022 | 0.13818 | 10.41667 | 0.05 | 0 | Neutral |
| testneg_7612 | 0 | 0 | -0.399895 | -3 | 3.109082 | -3 | 0.5632 | 0.28442 | 0.31244 | 0.4615 | 4.72222 | 0.203649999 | 0.2 | Neutral |
| testneg_7613 | 0.03 | -1.03 | 2.432661 | -3 | -0.519414 | -1.10605 | 0.1541 | 0.1077 | 0.23401 | 0.12374 | 20.42553 | 0.01 | 0 | Neutral |
| testneg_7614 | -0.04 | -2.01 | 4.558544 | -3 | -0.022585 | -3 | 0.08753 | 0.09666 | 0.2543 | 0.12929 | 8.97898 | 0.08 | 0 | Neutral |
| testneg_7615 | 0 | 0 | 0.112623 | 0.112623 | -0.255058 | -3 | 0.45152 | 0.34457 | 0.13796 | 0.27869 | 15.25194 | 0.129868421 | 0 | Neutral |
| testneg_7616 | 0.01 | -1.44 | 0.574267 | -3 | 0.169483 | -3 | 0.30367 | 0.41964 | 0.80314 | 0.42344 | 18.88889 | 0.190111111 | 0 | Neutral |
| testneg_7617 | 0.05 | -0.81 | 5.752104 | 5.752104 | 4.754501 | -3 | 0.17322 | 0.10311 | 0.1837 | 0.27351 | 17.8149 | 0.87 | 100 | Disease_causing |
| testneg_7618 | 0 | 0 | -0.492744 | -3 | 1.389503 | -3 | 0.21848 | 0.11049 | 0.13997 | 0.19598 | 20.84541 | 0.061081871 | 0 | Neutral |
| testneg_7619 | -0.03 | -0.12 | 1.627305 | -3 | -0.172833 | -3 | 0.08448 | 0.17914 | 0.09515 | 0.07784 | 21.52529 | 0.05 | 0 | Neutral |
| testneg_7620 | -0.09 | 0.3 | 0.782249 | -3 | -0.27139 | -3 | 0.5127 | 0.12263 | 0.37637 | 0.12788 | 5.27778 | 0.020853888 | 0 | Neutral |
| testneg_7621 | -0.02 | 0.36 | 4.645547 | -3 | -0.474175 | -3 | 0.56416 | 0.12788 | 0.3414 | 0.13575 | 17.46678 | 0.130662563 | 0 | Neutral |
| testneg_7622 | -0.08 | -0.03 | 5.931499 | -3 | -2.304927 | -2.304927 | 0.11019 | 0.08132 | 0.02665 | 0.05082 | 20.83333 | 0.14 | 0 | Neutral |
| testneg_7623 | 0 | 0 | 3.835304 | -3 | 0.722971 | -3 | 0.28227 | 0.24183 | 0.19898 | 0.12788 | 19.74592 | 0.065 | 0 | Neutral |
| testneg_7624 | -0.08 | -0.03 | 1.27395 | -3 | 6.245625 | -3 | 0.85252 | 0.83576 | 0.05535 | 0.12788 | 19.44444 | 0.259070513 | 0.4 | Neutral |
| testneg_7625 | -0.04 | 0.23 | 0.421557 | 0.421557 | 2.380524 | -3 | 0.82507 | 0.19615 | 0.20701 | 0.10345 | 18.32183 | 0.066530303 | 0 | Neutral |
| testneg_7626 | 0.05 | 0.29 | -1.171244 | -1.171244 | -1.599196 | -3 | 0.19898 | 0.12788 | 0.37711 | 0.12788 | 13.51351 | 0.01 | 0 | Neutral |
| testneg_7627 | -0.03 | -0.1 | 0.815782 | -3 | -1.932308 | -3 | 0.45029 | 0.12788 | 0.06474 | 0.23607 | 12.29167 | 0.061628394 | 0 | Neutral |
| testneg_7628 | 0.02 | 0.45 | -0.031677 | -3 | 0.752432 | -3 | 0.30669 | 0.11489 | 0.19898 | 0.1347 | 13.97176 | 0.06 | 0 | Neutral |
| testneg_7629 | -0.08 | -0.03 | 3.600262 | -3 | 1.018783 | -3 | 0.22944 | 0.09039 | 0.07189 | 0.11325 | 21.38889 | 0.04 | 0 | Neutral |
| testneg_7630 | 0.22 | -2.66 | 7.378253 | -3 | 3.668266 | -3 | 0.16544 | 0.1313 | 0.63645 | 0.14596 | 11.25 | 0.592100449 | 84 | Disease_causing |
| testneg_7631 | 0 | 0 | 0.504888 | -3 | 0.365748 | -3 | 0.06476 | 0.07491 | 0.55441 | 0.08704 | 20.39231 | 0.010234505 | 0 | Neutral |
| testneg_7632 | 0 | 0 | 1.309697 | -3 | 2.24087 | -3 | 0.26884 | 0.64999 | 0.39224 | 0.09664 | 10.83333 | 0.045 | 0 | Neutral |
| testneg_7633 | 0.02 | 0.45 | 5.906217 | -3 | 2.937571 | -3 | 0.72163 | 0.10596 | 0.05332 | 0.12788 | 16.8018 | 0.323541667 | 4.4 | Neutral |
| testneg_7634 | 0 | 0 | 1.095746 | -3 | 5.029108 | -3 | 0.24274 | 0.13034 | 0.10395 | 0.08737 | 13.51351 | 0.064722222 | 0 | Neutral |
| testneg_7635 | 0.15 | -0.28 | 1.144976 | -3 | 3.735934 | -3 | 0.53592 | 0.11293 | 0.20047 | 0.11827 | 26.80135 | 0.086464646 | 0 | Neutral |
| testneg_7636 | 0 | 0 | 0.288621 | -3 | 5.282618 | -3 | 0.09269 | 0.12788 | 0.4676 | 0.11257 | 29.44043 | 0.06 | 0 | Neutral |
| testneg_7637 | -0.02 | -0.41 | 2.303477 | -3 | 1.125228 | -3 | 0.53216 | 0.84775 | 0.15808 | 0.14548 | 11.72569 | 0.158711779 | 0 | Neutral |
| testneg_7638 | 0.04 | -0.23 | 4.018098 | 4.018098 | 1.66726 | -3 | 0.0606 | 0.39625 | 0.1215 | 0.12788 | 18.24178 | 0.510454545 | 60.2 | Disease_causing |
| testneg_7639 | -0.02 | -0.41 | -0.086855 | -3 | 0.061858 | -3 | 0.59827 | 0.11479 | 0.28474 | 0.12788 | 60.41783 | 0.070384615 | 0 | Neutral |
| testneg_7640 | 0.21 | 0.15 | 0.856634 | -3 | 4.42032 | -3 | 0.1214 | 0.0899 | 0.44722 | 0.11641 | 13.84202 | 0.083121693 | 0 | Neutral |
| testneg_7641 | 0 | 0 | 0.781134 | -3 | 2.597539 | -3 | 0.92237 | 0.11252 | 0.08775 | 0.12788 | 12.06801 | 0.020382653 | 0 | Neutral |
| testneg_7642 | 0 | 0 | 1.917158 | 1.917158 | 1.158382 | -3 | 0.11618 | 0.20784 | 0.19087 | 0.44337 | 10.41667 | 0.252525222 | 1.8 | Neutral |
| testneg_7643 | 0 | 0 | 0.693679 | -3 | 3.172437 | -3 | 0.13331 | 0.13638 | 0.11522 | 0.11925 | 18.69658 | 0.05 | 0 | Neutral |
| testneg_7644 | 0 | 0 | 1.067571 | -3 | 3.428327 | -3 | 0.72124 | 0.11262 | 0.11897 | 0.3919 | 10.41667 | 0.163942882 | 0 | Neutral |
| testneg_7645 | 0 | 0 | 3.296898 | -3 | 0.440748 | -3 | 0.13559 | 0.1946 | 0.14735 | 0.12788 | 19.09347 | 0.05 | 0 | Neutral |
| testneg_7646 | -0.03 | -0.1 | 6.425529 | 6.425529 | 3.305653 | -3 | 0.30657 | 0.52974 | 0.47274 | 0.11928 | 4.44444 | 0.75 | 100 | Disease_causing |
| testneg_7647 | 0.05 | -0.81 | -0.021519 | -3 | 4.98919 | 4.98919 | 0.30096 | 0.17203 | 0.18512 | 0.12788 | 23.43525 | 0.422121212 | 24.8 | Neutral |
| testneg_7648 | 0.01 | 0.79 | 4.326431 | -3 | 2.759319 | -3 | 0.13195 | 0.57267 | 0.19623 | 0.08363 | 8.97898 | 0.266515568 | 0 | Neutral |
| testneg_7649 | 0 | 0 | 0.200125 | -3 | 4.61145 | -3 | 0.13863 | 0.2579 | 0.19898 | 0.12788 | 10.83333 | 0.142294831 | 0 | Neutral |
| testneg_7650 | 0.21 | 0.15 | 6.51955 | -3 | 5.844424 | -3 | 0.90811 | 0.17448 | 0.12121 | 0.4142 | 10.41667 | 0.848143297 | 100 | Disease_causing |
| testneg_7651 | -0.15 | 1.06 | 3.424957 | -3 | 1.850619 | -3 | 0.559 | 0.2286 | 0.18498 | 0.12788 | 11.54145 | 0.200217391 | 0 | Neutral |
| testneg_7652 | 0.21 | 0.15 | 8.082349 | -3 | 0.259282 | -3 | 0.9989 | 0.8746 | 0.16468 | 0.11275 | 10.41667 | 0.49 | 54 | Disease_causing |
| testneg_7653 | 0.08 | 0.03 | 0.846733 | -3 | -1.056572 | -3 | 0.9874 | 0.15588 | 0.1109 | 0.17295 | 33.06291 | 0.17 | 0 | Neutral |
| testneg_7654 | -0.02 | -0.13 | 1.214257 | -3 | 0.508101 | -3 | 0.43459 | 0.12595 | 0.98712 | 0.10969 | 10.30744 | 0.02 | 0 | Neutral |
| testneg_7655 | 0 | 0 | 1.040406 | -3 | -0.151164 | -0.151164 | 0.41987 | 0.67373 | 0.65708 | 0.0936 | 19.11866 | 0.050262796 | 0 | Neutral |
| testneg_7656 | -0.08 | 1.2 | 1.189976 | -3 | 3.749849 | -3 | 0.17303 | 0.18733 | 0.19898 | 0.12788 | 12.29167 | 0.250087719 | 0.4 | Neutral |
| testneg_7657 | 0.00036364 | -0.0270909 | 2.070789 | -3 | 4.697774 | -3 | 0.91139 | 0.65138 | 0.22225 | 0.11186 | 18.33333 | 0.159294456 | 0 | Neutral |
| testneg_7658 | -0.07 | -0.38 | 0.816148 | -3 | 1.119261 | -3 | 0.26347 | 0.16383 | 0.11858 | 0.11551 | 20.00442 | 0.03 | 0 | Neutral |
| testneg_7659 | 0.06 | -0.57 | 2.083519 | -3 | 2.071548 | -3 | 0.14375 | 0.12788 | 0.19898 | 0.10116 | 18.33333 | 0.01 | 0 | Neutral |
| testneg_7660 | 0.22 | -2.66 | 5.527307 | -3 | 3.716419 | -3 | 0.1589 | 0.17874 | 0.14954 | 0.12788 | 18.88889 | 0.43 | 30 | Neutral |
| testneg_7661 | 0.02 | 0.41 | 3.942193 | 0.147043 | 0.235742 | -3 | 0.14668 | 0.12788 | 0.3718 | 0.2248 | 10.8046 | 0.24 | 0.2 | Neutral |
| testneg_7662 | 0.04 | 2.01 | 3.781815 | -3 | 1.428514 | -3 | 0.01704 | 0.05734 | 0.52118 | 0.36073 | 15.37043 | 0.27 | 0.6 | Neutral |
| testneg_7663 | 0 | 0 | 0.872561 | -3 | 4.699217 | -3 | 0.85984 | 0.40044 | 0.06729 | 0.17051 | 18.88889 | 0.358572939 | 9.6 | Neutral |
| testneg_7664 | 0.03 | -1.03 | 0.840482 | -3 | 1.651197 | -3 | 0.1294 | 0.10964 | 0.76918 | 0.13588 | 11.59175 | 0.05 | 0 | Neutral |
| testneg_7665 | 0.14 | -0.69 | 5.242793 | -3 | 0.663866 | 0.663866 | 0.30942 | 0.13105 | 0.08283 | 0.11797 | 20.46784 | 0.11 | 0 | Neutral |
| testneg_7666 | 0 | 0 | 1.887682 | -3 | 0.911193 | -3 | 0.18282 | 0.12022 | 0.19898 | 0.12788 | 32.87037 | 0 | 0 | Neutral |
| testneg_7667 | -0.0163636 | 0.03181818 | 12.519035 | -3 | 6.490439 | -3 | 0.03646 | 0.09863 | 0.02498 | 0.08483 | 20.13398 | 0.53 | 68 | Disease_causing |
| testneg_7668 | -0.03 | -0.1 | 6.245625 | -3 | 3.238247 | 3.238247 | 0.05535 | 0.12788 | 0.19898 | 0.12788 | 58.22139 | 0.39 | 14.6 | Neutral |
| testneg_7669 | 0 | 0 | 4.103101 | -3 | 2.177568 | -3 | 0.20011 | 0.27949 | 0.08876 | 0.09037 | 10.41667 | 0.107212349 | 0 | Neutral |
| testneg_7670 | 0 | 0 | 1.283706 | -3 | -1.868425 | -3 | 0.16856 | 0.19221 | 0.00971 | 0.0791 | 23.88889 | 0.020280338 | 0 | Neutral |
| testneg_7671 | 0 | 0 | 0.256955 | -3 | 1.336135 | -3 | 0.1846 | 0.08682 | 0.17935 | 0.36953 | 24.52338 | 0.08 | 0 | Neutral |
| testneg_7672 | 0 | -0.01 | 0.238703 | -3 | 0.941706 | -3 | 0.0887 | 0.10047 | 0.15294 | 0.12788 | 32.42945 | 0.000199966 | 0 | Neutral |
| testneg_7673 | 0.17 | 0.38 | 1.691672 | 1.691672 | 0.580057 | -3 | 0.30789 | 0.15135 | 0.10058 | 0.13882 | 16.66667 | 0.170254183 | 0 | Neutral |
| testneg_7674 | 0 | 0 | 2.437503 | -3 | 1.534492 | -3 | 0.11337 | 0.09363 | 0.10183 | 0.11847 | 13.08333 | 0.02 | 0 | Neutral |
| testneg_7675 | 0.18 | -1.05 | 6.835273 | 6.835273 | 4.396862 | -3 | 0.08412 | 0.28307 | 0.1072 | 0.63446 | 19.56324 | 0.91 | 100 | Disease_causing |
| testneg_7676 | 0.07 | 0.38 | -0.999113 | -3 | 1.986841 | 1.986841 | 0.25961 | 0.07098 | 0.19898 | 0.12788 | 25.94732 | 0.05 | 0 | Neutral |
| testneg_7677 | 0 | 0 | -0.37295 | -3 | 1.476163 | -3 | 0.07237 | 0.12788 | 0.19898 | 0.09395 | 36.8277 | 0.01 | 0 | Neutral |
| testneg_7678 | -0.01 | -0.79 | 1.315391 | -3 | 0.774689 | 0.774689 | 0.99417 | 0.17365 | 0.24219 | 0.17458 | 19.44444 | 0.12 | 0 | Neutral |
| testneg_7679 | -0.02 | 0.68 | 5.845448 | -3 | -0.284617 | -3 | 0.06926 | 0.08399 | 0.6975 | 0.53794 | 16.09234 | 0.358927661 | 8 | Neutral |
| testneg_7680 | -0.03 | -0.1 | 0.271296 | -3 | 6.71239 | -3 | 0.0978 | 0.26338 | 0.68568 | 0.65219 | 10.83333 | 0.790693978 | 99.8 | Disease_causing |
| testneg_7681 | -0.07 | -0.38 | 2.173191 | -3 | 2.66819 | -3 | 0.18233 | 0.14781 | 0.06922 | 0.07026 | 19.09347 | 0.050488095 | 0 | Neutral |
| testneg_7682 | 0 | 0 | 0.097356 | -3 | 6.6845 | -3 | 0.14485 | 0.16945 | 0.32423 | 0.08104 | 22.15924 | 0.272048611 | 2.8 | Neutral |
| testneg_7683 | 0 | 0 | 0.039637 | -3 | 1.772574 | -3 | 0.20304 | 0.31462 | 0.0887 | 0.10047 | 31.73501 | 0.03 | 0 | Neutral |
| testneg_7684 | -0.1 | -1.9 | -0.234016 | -3 | 0.695182 | -3 | 0.04436 | 0.12788 | 0.11587 | 0.12001 | 11.41859 | 0.08 | 0 | Neutral |
| testneg_7685 | 0 | 0 | 1.391383 | -3 | -0.035874 | -3 | 0.21863 | 0.16306 | 0.19898 | 0.12788 | 10 | 0.01 | 0 | Neutral |

| | | | | | | | | | | | | | | |
|---|---|---|---|---|---|---|---|---|---|---|---|---|---|---|
| testneg_7686 | 0.15 | -0.28 | 6.108445 | -3 | 1.38979 | -3 | 0.10723 | 0.11163 | 0.09059 | 0.12032 | 20.3268 | 0.144714286 | 0 | Neutral |
| testneg_7687 | 0.17 | 0.38 | 4.86313 | -3 | -0.122196 | -0.122196 | 0.24516 | 0.11809 | 0.31961 | 0.17571 | 20.22287 | 0.214647887 | 0 | Neutral |
| testneg_7688 | -0.07 | -0.38 | 4.583643 | -3 | 0.084612 | -3 | 0.62708 | 0.24932 | 0.99994 | 0.51636 | 10 | 0.311041667 | 3 | Neutral |
| testneg_7689 | 0 | 0 | 1.519976 | -3 | 1.50021 | -3 | 0.19898 | 0.12788 | 0.27618 | 0.11809 | 17.90495 | 0 | 0 | Neutral |
| testneg_7690 | 0 | 0 | 0.467241 | -3 | 6.791006 | -3 | 0.14948 | 0.12788 | 0.80264 | 0.11567 | 19.95974 | 0.16 | 0 | Neutral |
| testneg_7691 | 0 | 0 | 2.336102 | -3 | 0.988618 | 0.988618 | 0.31062 | 0.1222 | 0.15019 | 0.10342 | 21.52904 | 0.01 | 0 | Neutral |
| testneg_7692 | -0.02 | -0.45 | 0.018913 | -3 | -0.072198 | -0.072198 | 0.65141 | 0.18046 | 0.1025 | 0.3268 | 10 | 0.140274927 | 0 | Neutral |
| testneg_7693 | 0.14 | -0.37 | 1.458113 | -3 | 2.296124 | -3 | 0.08911 | 0.09298 | 0.22258 | 0.10471 | 20.90643 | 0.02 | 0 | Neutral |
| testneg_7694 | 0.07 | 0.7 | 2.665776 | -3 | 0.097957 | -3 | 0.3954 | 0.17605 | 0.14403 | 0.30507 | 10.41667 | 0.171909091 | 0 | Neutral |
| testneg_7695 | -0.18 | -0.16 | -0.566382 | -3 | 1.348057 | -3 | 0.23152 | 0.09962 | 0.9807 | 0.2062 | 14.55369 | 0.092354167 | 0 | Neutral |
| testneg_7696 | 0.16 | -0.37 | 5.827202 | -3 | 2.499788 | -3 | 0.13275 | 0.12788 | 0.18765 | 0.27794 | 11.22943 | 0.559799277 | 75.2 | Disease_causing |
| testneg_7697 | 0.14 | -1.01 | 0.460152 | -3 | 5.744002 | -3 | 0.21033 | 0.35953 | 0.15016 | 0.08826 | 18.88889 | 0.24 | 0.2 | Neutral |
| testneg_7698 | 0 | 0 | 0.439979 | -3 | 7.731489 | -3 | 0.0811 | 0.09449 | 0.03378 | 0.12788 | 40.27778 | 0.26 | 1.2 | Neutral |
| testneg_7699 | 0.04 | -0.23 | 2.522879 | -3 | 0.131996 | -3 | 0.99998 | 0.86471 | 0.19821 | 0.20954 | 11.25 | 0.198381119 | 0.2 | Neutral |
| testneg_7700 | 0.17 | -1.22 | 5.736168 | -3 | 5.859532 | -3 | 0.33245 | 0.45921 | 0.04935 | 0.06599 | 9.32889 | 0.5355 | 65.8 | Disease_causing |
| testneg_7701 | 0 | 0 | 0.728127 | -3 | -1.068385 | -3 | 0.07681 | 0.22865 | 0.07552 | 0.12788 | 17.1181 | 0.03 | 0 | Neutral |
| testneg_7702 | 0.08 | -1.2 | -0.288259 | -3 | 4.178117 | -3 | 0.04346 | 0.10785 | 0.16424 | 0.11262 | 16.8018 | 0.091339286 | 0 | Neutral |
| testneg_7703 | 0.18 | -1.05 | 4.656578 | -3 | 5.82456 | -3 | 0.31062 | 0.1222 | 0.33166 | 0.11755 | 17.21847 | 0.29 | 4.8 | Neutral |
| testneg_7704 | 0.04 | 2.01 | -0.074864 | -3 | 7.659081 | -3 | 0.16468 | 0.11275 | 0.7022 | 0.35308 | 23.57794 | 0.443351852 | 30.4 | Neutral |
| testneg_7705 | 0.18 | -1.05 | 2.987486 | -3 | 0.600136 | -3 | 0.08896 | 0.12788 | 0.26842 | 0.09683 | 26.54712 | 0.04 | 0 | Neutral |
| testneg_7706 | -0.06 | 1.29 | 4.344772 | -3 | 0.341478 | -3 | 0.34404 | 0.13209 | 0.1144 | 0.09798 | 13.02635 | 0.1 | 0 | Neutral |
| testneg_7707 | -0.06 | -0.18 | 1.398739 | -3 | 4.647205 | -3 | 0.54113 | 0.11017 | 0.2741 | 0.10478 | 18.91311 | 0.06010904 | 0 | Neutral |
| testneg_7708 | 0.02 | -0.63 | -0.055258 | -3 | 0.069364 | -3 | 0.16139 | 0.09116 | 0.17418 | 0.1871 | 20.85816 | 0.060714286 | 0 | Neutral |
| testneg_7709 | 0 | 0 | 1.466727 | -3 | 3.014014 | -3 | 0.21848 | 0.11049 | 0.88456 | 0.11674 | 10.83333 | 0.03 | 0 | Neutral |
| testneg_7710 | 0 | 0 | -0.398253 | -3 | 3.003558 | -3 | 0.04668 | 0.12788 | 0.19898 | 0.12788 | 26.66726 | 0.054313725 | 0 | Neutral |
| testneg_7711 | 0.07 | -0.31 | 1.843162 | -3 | 6.188044 | -3 | 0.30869 | 0.11948 | 0.19898 | 0.12788 | 10.41667 | 0.14 | 0 | Neutral |
| testneg_7712 | 0.14 | -1.01 | 0.341374 | -3 | 6.89099 | -3 | 0.10646 | 0.1069 | 0.33418 | 0.09979 | 21.48917 | 0.22 | 0.4 | Neutral |
| testneg_7713 | 0.22 | -2.66 | 2.765044 | -3 | 0.849086 | -3 | 0.26337 | 0.12771 | 0.30241 | 0.104 | 10.41667 | 0.07 | 0 | Neutral |
| testneg_7714 | 0 | 0 | 1.56019 | -3 | 0.246616 | -3 | 0.25487 | 0.13098 | 0.06955 | 0.1748 | 15.39358 | 0.06 | 0 | Neutral |
| testneg_7715 | 0.07 | 0.38 | 1.331486 | -3 | -0.187363 | -3 | 0.08401 | 0.16123 | 0.90649 | 0.3543 | 9.79167 | 0.120564428 | 0 | Neutral |
| testneg_7716 | 0.04 | -0.23 | -0.229525 | -3 | 1.398027 | -3 | 0.19898 | 0.12788 | 0.38934 | 0.15403 | 21.94444 | 0.06171965 | 0 | Neutral |
| testneg_7717 | 0.23 | -1.04 | -1.468734 | -3 | 0.53236 | -3 | 0.06789 | 0.09707 | 0.46056 | 0.12785 | 6.94444 | 0.044166667 | 0 | Neutral |
| testneg_7718 | -0.09 | 0.3 | -0.522638 | -3 | 1.96279 | -3 | 0.20604 | 0.10821 | 0.82126 | 0.17937 | 14.50908 | 0.06 | 0 | Neutral |
| testneg_7719 | -0.15 | 0.28 | 3.39754 | -3 | 2.670295 | -3 | 0.62203 | 0.15583 | 0.09807 | 0.24901 | 10 | 0.451409812 | 35.2 | Neutral |
| testneg_7720 | 0 | 0 | 0.159372 | 0.159372 | 2.277356 | -3 | 0.05228 | 0.12788 | 0.52957 | 0.18687 | 25.58159 | 0.158204545 | 0 | Neutral |
| testneg_7721 | 0 | -0.01 | 2.148855 | -3 | 3.027149 | -3 | 0.56904 | 0.14839 | 0.10812 | 0.12788 | 18.33333 | 0.07 | 0 | Neutral |
| testneg_7722 | -0.07 | -0.38 | 0.37014 | -3 | -0.47975 | -3 | 0.12048 | 0.2381 | 0.4719 | 0.33344 | 10.83333 | 0.194633099 | 0 | Neutral |
| testneg_7723 | 0 | 0 | -0.032453 | -3 | 0.151938 | -3 | 0.31664 | 0.14181 | 0.47188 | 0.10037 | 5 | 0.01 | 0 | Neutral |
| testneg_7724 | -0.07 | -0.38 | 2.393948 | 2.393948 | -0.932495 | -3 | 0.70819 | 0.5975 | 0.0819 | 0.1823 | 12.29167 | 0.31 | 7.8 | Neutral |
| testneg_7725 | 0 | 0 | 0.24381 | -3 | 1.674048 | -3 | 0.29624 | 0.1067 | 0.25987 | 0.31819 | 20.40541 | 0.051899542 | 0 | Neutral |
| testneg_7726 | -0.05 | -0.29 | 1.124718 | -3 | -2.611796 | -3 | 0.55043 | 0.12112 | 0.19898 | 0.12788 | 27.3569 | 0.010714286 | 0 | Neutral |
| testneg_7727 | 0 | -0.32 | -0.692188 | -3 | 3.703925 | -3 | 0.12269 | 0.08105 | 0.22901 | 0.12788 | 33.63961 | 0.05 | 0 | Neutral |
| testneg_7728 | 0 | 0 | 0.179322 | -3 | 3.490489 | -3 | 0.54349 | 0.60885 | 0.70964 | 0.12663 | 10.41667 | 0.11044765 | 0 | Neutral |
| testneg_7729 | 0.01 | -0.57 | 5.075109 | -3 | 1.243494 | -3 | 0.68863 | 0.09663 | 0.19898 | 0.10269 | 34.48332 | 0.1 | 0 | Neutral |
| testneg_7730 | 0 | 0 | 1.221853 | 1.221853 | 1.173845 | -3 | 0.92659 | 0.13588 | 0.88689 | 0.41514 | 16.42913 | 0.091475541 | 0 | Neutral |
| testneg_7731 | -0.09 | 0.25 | 4.068872 | -3 | 4.351389 | -3 | 0.19898 | 0.12788 | 0.1653 | 0.0977 | 15.29221 | 0.220203418 | 0 | Neutral |
| testneg_7732 | 0.01 | 0.8 | 3.671807 | -3 | 5.041891 | -3 | 0.17222 | 0.12788 | 0.16621 | 0.49845 | 11.10144 | 0.581932494 | 77.8 | Disease_causing |
| testneg_7733 | 0.02 | -0.36 | 1.715391 | -3 | -2.259041 | -3 | 0.60959 | 0.76078 | 0.21723 | 0.10332 | 10 | 0.04 | 0 | Neutral |
| testneg_7734 | 0 | 0 | 0.774193 | -3 | 0.313774 | -3 | 0.32963 | 0.1035 | 0.1462 | 0.18896 | 15.28736 | 0.020360665 | 0 | Neutral |
| testneg_7735 | 0.03 | 1.22 | -0.307704 | -3 | 0.641608 | -3 | 0.12497 | 0.09805 | 0.32458 | 0.1016 | 21.42525 | 0.04 | 0 | Neutral |
| testneg_7736 | 0 | 0 | 0.774504 | -3 | 0.055743 | -3 | 0.99338 | 0.14229 | 0.40116 | 0.43982 | 5.62818 | 0.057308615 | 0 | Neutral |
| testneg_7737 | 0.15 | 0.06 | 2.738091 | -3 | 5.835639 | -3 | 0.34495 | 0.16861 | 0.19898 | 0.12788 | 9.79167 | 0.331560897 | 7.8 | Neutral |
| testneg_7738 | 0 | 0 | 1.829958 | -3 | -0.154534 | -3 | 0.47775 | 0.10103 | 0.34946 | 0.19092 | 18.88889 | 0.030471491 | 0 | Neutral |
| testneg_7739 | 0 | 0 | 1.856091 | -3 | 1.995608 | -3 | 0.94822 | 0.14005 | 0.39262 | 0.12788 | 11.25 | 0.02 | 0 | Neutral |
| testneg_7740 | -0.07 | -0.38 | 1.216755 | -3 | 3.087586 | -3 | 0.54049 | 0.33876 | 0.3828 | 0.5569 | 10.83333 | 0.348740618 | 9.2 | Neutral |
| testneg_7741 | 0.09 | -2.17 | 5.174314 | -3 | 1.175993 | -3 | 0.73311 | 0.10866 | 0.06781 | 0.09009 | 10.22898 | 0.131714286 | 0 | Neutral |
| testneg_7742 | 0.02 | -0.63 | 0.567529 | -3 | 0.734765 | -3 | 0.56554 | 0.76978 | 0.07204 | 0.07708 | 10.3125 | 0.050606618 | 0 | Neutral |
| testneg_7743 | 0 | 0 | -1.230837 | -1.230837 | 0.768207 | -3 | 0.14821 | 0.11262 | 0.30669 | 0.11489 | 7.00579 | 0 | 0 | Neutral |
| testneg_7744 | -0.18 | -0.16 | 2.830718 | -3 | 1.871905 | -3 | 0.11229 | 0.08897 | 0.51852 | 0.11202 | 18.57759 | 0.070258938 | 0 | Neutral |
| testneg_7745 | 0.04 | 2.01 | -0.172559 | -3 | 2.44605 | -3 | 0.22155 | 0.1062 | 0.19898 | 0.06726 | 45.41667 | 0.1 | 0 | Neutral |
| testneg_7746 | 0 | -0.01 | 2.187535 | -3 | 2.238523 | -0.534525 | 0.93654 | 0.38765 | 0.59947 | 0.10764 | 19.8494 | 0.05 | 0 | Neutral |
| testneg_7747 | 0 | -0.01 | 1.315543 | -3 | 0.464102 | -3 | 0.28358 | 0.38765 | 0.14668 | 0.25103 | 17.77778 | 0.122458219 | 0 | Neutral |
| testneg_7748 | 0.11 | -0.19 | 0.034536 | -3 | 3.017506 | -3 | 0.2049 | 0.12678 | 0.14862 | 0.12788 | 30.04935 | 0.05 | 0 | Neutral |
| testneg_7749 | -0.02 | 0.63 | 4.675931 | -3 | 2.821548 | -3 | 0.961 | 0.25991 | 0.18578 | 0.27854 | 4.72222 | 0.577927165 | 79.4 | Disease_causing |
| testneg_7750 | 0 | 0 | 0.097083 | -3 | 4.038181 | 4.038181 | 0.12943 | 0.14391 | 0.13815 | 0.40001 | 4.72222 | 0.466261333 | 43 | Neutral |
| testneg_7751 | 0.07 | -0.31 | 2.110113 | -3 | 4.147656 | -3 | 0.68029 | 0.12788 | 0.33245 | 0.45921 | 18.33333 | 0.356256341 | 7.8 | Neutral |
| testneg_7752 | 0 | 0 | 0.840063 | -3 | 2.493678 | -3 | 0.08683 | 0.26961 | 0.50057 | 0.10408 | 38.93278 | 0.06459596 | 0 | Neutral |
| testneg_7753 | 0.01 | 0.79 | 0.266225 | -3 | -0.249888 | -3 | 0.12943 | 0.14391 | 0.47462 | 0.10826 | 12.79101 | 0.020377493 | 0 | Neutral |
| testneg_7754 | 0 | -0.01 | 0.334215 | -3 | 4.130555 | -3 | 0.33314 | 0.47999 | 0.10891 | 0.12788 | 18.87021 | 0.143337852 | 0 | Neutral |
| testneg_7755 | -0.12 | 1.16 | 3.67024 | -3 | 4.558319 | -3 | 0.15509 | 0.10665 | 0.19898 | 0.12788 | 30.55556 | 0.270833234 | 0.8 | Neutral |
| testneg_7756 | -0.02 | 0.28 | 2.201231 | -3 | 0.202818 | -3 | 0.20231 | 0.12788 | 0.07118 | 0.08861 | 23.64842 | 0.01 | 0 | Neutral |
| testneg_7757 | 0 | 0 | 0.598467 | -3 | 1.242354 | -3 | 0.52836 | 0.12785 | 0.66381 | 0.13039 | 14.51616 | 0.02 | 0 | Neutral |
| testneg_7758 | 0 | 0 | -0.161926 | -0.161926 | 2.756541 | -3 | 0.42033 | 0.17821 | 0.25236 | 0.11809 | 11.98582 | 0.05013299 | 0 | Neutral |
| testneg_7759 | 0.03 | 0.1 | 3.044807 | -3 | 4.54534 | -3 | 0.25905 | 0.23728 | 0.06713 | 0.11112 | 19.12611 | 0.277183783 | 1.4 | Neutral |
| testneg_7760 | -0.14 | 0.69 | 2.798182 | 2.798182 | -1.124893 | -3 | 0.31043 | 0.09688 | 0.07923 | 0.12788 | 19.44444 | 0.3 | 4 | Neutral |
| testneg_7761 | 0 | 0 | 0.279445 | -3 | 1.849259 | -3 | 0.25179 | 0.09355 | 0.41261 | 0.12096 | 13.51351 | 0.000752976 | 0 | Neutral |
| testneg_7762 | -0.08 | -0.03 | -0.125177 | -3 | 1.461115 | -3 | 0.27308 | 0.12788 | 0.22953 | 0.0919 | 19.14829 | 0.01 | 0 | Neutral |
| testneg_7763 | 0 | 0 | -0.88087 | -3 | 1.006301 | -3 | 0.38135 | 0.22085 | 0.21941 | 0.46348 | 10 | 0.071666544 | 0 | Neutral |
| testneg_7764 | 0 | 0 | 0.004869 | -3 | 2.849519 | -3 | 0.16134 | 0.1847 | 0.14486 | 0.33725 | 10.41667 | 0.203042212 | 0.2 | Neutral |
| testneg_7765 | 0 | 0 | 1.317673 | -3 | 6.667802 | -3 | 0.45415 | 0.12971 | 0.81838 | 0.09725 | 11.25 | 0.160289916 | 0 | Neutral |
| testneg_7766 | 0 | 0 | 0.358416 | -3 | -0.347591 | -3 | 0.37292 | 0.11979 | 0.32094 | 0.36254 | 18.88889 | 0.05 | 0 | Neutral |
| testneg_7767 | 0 | 0 | 1.445184 | 1.445184 | 5.295171 | -3 | 0.33469 | 0.17432 | 0.098 | 0.08791 | 21.90492 | 0.172916667 | 0 | Neutral |
| testneg_7768 | 0.22 | -2.66 | 4.250235 | -3 | 6.109579 | 4.284949 | 0.60412 | 0.13303 | 0.15267 | 0.12788 | 15.62189 | 0.550003374 | 73.4 | Disease_causing |
| testneg_7769 | 0 | 0 | 0.191318 | -3 | 2.918397 | -3 | 0.96113 | 0.24974 | 0.63378 | 0.09451 | 18.88889 | 0.050878395 | 0 | Neutral |
| testneg_7770 | 0 | 0 | 1.499426 | -3 | -0.4837 | -3 | 0.46791 | 0.7316 | 0.23276 | 0.31401 | 12.38782 | 0.11 | 0 | Neutral |
| testneg_7771 | -0.01 | -0.79 | 1.377684 | -3 | 2.177517 | -3 | 0.09406 | 0.19858 | 0.39262 | 0.12788 | 10.83333 | 0.0745 | 0 | Neutral |
| testneg_7772 | -0.05 | -0.29 | -1.641768 | -3 | 6.533819 | -3 | 0.31648 | 0.12788 | 0.50303 | 0.23508 | 22.36324 | 0.418730159 | 23.8 | Neutral |
| testneg_7773 | 0.15 | -0.28 | 4.207254 | -3 | 0.359803 | -3 | 0.99665 | 0.12884 | 0.01635 | 0.12788 | 13.02635 | 0.096666667 | 0 | Neutral |
| testneg_7774 | 0 | -0.32 | 0.219522 | -3 | 3.764453 | 0.434159 | 0.19898 | 0.09533 | 0.10168 | 0.12788 | 28.46801 | 0.09 | 0 | Neutral |
| testneg_7775 | 0 | 0 | 0.905769 | -3 | 0.169559 | -3 | 0.08932 | 0.32882 | 0.86166 | 0.14254 | 19.09347 | 0.111710486 | 0 | Neutral |
| testneg_7776 | 0.04 | -0.23 | 0.407117 | -3 | 2.054242 | -3 | 0.17728 | 0.46704 | 0.17016 | 0.12748 | 20.91248 | 0.070187548 | 0 | Neutral |
| testneg_7777 | 0 | 0 | 0.419995 | -3 | 2.856561 | -3 | 0.7174 | 0.15815 | 0.53798 | 0.10163 | 18.77814 | 0.044935897 | 0 | Neutral |
| testneg_7778 | 0.14 | -0.37 | 1.658258 | -3 | 0.272659 | 0.272659 | 0.3414 | 0.13575 | 0.17093 | 0.23281 | 26.58984 | 0.113287546 | 0 | Neutral |
| testneg_7779 | -0.03 | -0.1 | 0.212913 | -3 | 2.317273 | -0.186375 | 0.11391 | 0.19434 | 0.85887 | 0.1489 | 8.35719 | 0.172717796 | 0 | Neutral |
| testneg_7780 | 0 | 0 | 0.461408 | -3 | -0.196861 | -3 | 0.16797 | 0.11809 | 0.34859 | 0.09799 | 9.52684 | 0 | 0 | Neutral |
| testneg_7781 | 0 | 0 | 1.019772 | -3 | 0.193332 | -3 | 0.95076 | 0.17388 | 0.17264 | 0.09426 | 17.3494 | 0.010435606 | 0 | Neutral |
| testneg_7782 | 0 | 0 | -0.464766 | -3 | 0.332327 | -3 | 0.62197 | 0.12788 | 0.83955 | 0.10333 | 16.11675 | 0.01 | 0 | Neutral |
| testneg_7783 | -0.06 | -0.18 | 1.759866 | -3 | 0.715993 | -3 | 0.7249 | 0.41559 | 0.04047 | 0.12788 | 10 | 0.022804552 | 0 | Neutral |
| testneg_7784 | 0.08 | 0.03 | -2.112814 | -3 | 2.327743 | -3 | 0.26982 | 0.12788 | 0.0538 | 0.0823 | 18.6768 | 0.05 | 0 | Neutral |
| testneg_7785 | 0 | 0 | 0.862479 | -3 | 2.628269 | -3 | 0.06708 | 0.09234 | 0.18584 | 0.11114 | 19.11368 | 0.02 | 0 | Neutral |
| testneg_7786 | -0.08 | -0.03 | 0.950149 | -3 | 0.815861 | -3 | 0.31318 | 0.11262 | 0.11564 | 0.09588 | 20.05594 | 0.01 | 0 | Neutral |
| testneg_7787 | -0.06 | 0.57 | 0.172291 | -3 | 4.781408 | -3 | 0.0138 | 0.06769 | 0.19898 | 0.12788 | 20.11075 | 0.122 | 0 | Neutral |
| testneg_7788 | 0 | 0 | 2.987273 | -3 | 2.478795 | -3 | 0.03502 | 0.08119 | 0.1327 | 0.09885 | 12.98396 | 0.083361111 | 0 | Neutral |
| testneg_7789 | 0 | 0 | 1.690602 | -3 | 1.121685 | -3 | 0.18206 | 0.10727 | 0.33418 | 0.09979 | 19.2377 | 0 | 0 | Neutral |
| testneg_7790 | 0 | 0 | 1.114336 | -3 | 5.896833 | -3 | 0.16835 | 0.09282 | 0.23152 | 0.09962 | 23.19982 | 0.060660173 | 0 | Neutral |
| testneg_7791 | -0.16 | -0.08 | -2.628134 | -3 | 5.125466 | -3 | 0.26971 | 0.08509 | 0.04615 | 0.12788 | 38.82925 | 0.18 | 0 | Neutral |
| testneg_7792 | -0.16 | 0.37 | 5.039547 | -3 | 2.495357 | -3 | 0.42512 | 0.29333 | 0.29548 | 0.10732 | 24.13736 | 0.308560875 | 2 | Neutral |
| testneg_7793 | -0.02 | 0.36 | 2.994942 | -3 | 4.332753 | -3 | 0.2477 | 0.11928 | 0.1087 | 0.08523 | 20.42484 | 0.13 | 0 | Neutral |
| testneg_7794 | -0.09 | 0.67 | -1.392375 | -3 | 1.937283 | -3 | 0.16741 | 0.10762 | 0.3112 | 0.33703 | 6.94444 | 0.09005 | 0 | Neutral |
| testneg_7795 | 0.12 | -1.16 | 4.5703 | -3 | 6.243256 | -3 | 0.20072 | 0.08606 | 0.29804 | 0.12768 | 25.38314 | 0.261269231 | 2.2 | Neutral |
| testneg_7796 | 0 | 0 | 0.606984 | -3 | -2.855052 | -3 | 0.08932 | 0.12788 | 0.16545 | 0.10002 | 19.44444 | 0.01 | 0 | Neutral |
| testneg_7797 | 0.1 | 1.9 | 1.15713 | -3 | 2.445411 | -3 | 0.36167 | 0.15588 | 0.22179 | 0.09831 | 18.88889 | 0.140669643 | 0 | Neutral |
| testneg_7798 | 0 | 0 | 2.064931 | -3 | 0.010179 | -3 | 0.30122 | 0.09165 | 0.19898 | 0.12788 | 20.58822 | 0.000600649 | 0 | Neutral |
| testneg_7799 | -0.15 | -0.04 | 0.724899 | -3 | 4.886271 | -3 | 0.19898 | 0.13057 | 0.19898 | 0.12788 | 18.33333 | 0.189782936 | 0 | Neutral |
| testneg_7800 | 0.06 | 0.18 | -0.134916 | -0.134916 | 6.453651 | -3 | 0.16475 | 0.26069 | 0.07889 | 0.08835 | 15.89461 | 0.300561224 | 1.4 | Neutral |
| testneg_7801 | -0.07 | -0.01 | -0.23732 | -0.23732 | 2.686945 | -3 | 0.07473 | 0.51812 | 0.08499 | 0.08678 | 10 | 0.108125 | 0 | Neutral |
| testneg_7802 | 0 | 0 | 0.036223 | -3 | 2.254535 | 1.806647 | 0.9355 | 0.40418 | 0.21728 | 0.1132 | 18.33333 | 0.05 | 0 | Neutral |
| testneg_7803 | -0.02 | -0.13 | 0.687351 | -3 | 6.373102 | -3 | 0.51675 | 0.11055 | 0.05754 | 0.10881 | 18.37773 | 0.167361111 | 0 | Neutral |
| testneg_7804 | 0 | 0 | 2.047211 | -3 | 0.173288 | -3 | 0.16823 | 0.10796 | 0.61211 | 0.11059 | 14.23074 | 0 | 0 | Neutral |
| testneg_7805 | 0.18 | -1.05 | 5.614334 | -3 | 3.329851 | -3 | 0.75033 | 0.64204 | 0.5184 | 0.11889 | 10 | 0.4880625 | 49.8 | Neutral |
| testneg_7806 | 0.22 | -2.66 | 7.225361 | -3 | 1.472855 | 0.852085 | 0.19898 | 0.15588 | 0.56554 | 0.76978 | 10 | 0.536666667 | 65.8 | Disease_causing |
| testneg_7807 | 0.02 | 0.45 | -0.031677 | -3 | 0.331991 | -3 | 0.30669 | 0.11489 | 0.11571 | 0.17324 | 7.00579 | 0.06 | 0 | Neutral |

| | | | | | | | | | | | | | | |
|---|---|---|---|---|---|---|---|---|---|---|---|---|---|---|
| testneg_7808 | 0 | 0 | 0.435323 | -3 | 0.452441 | 0.452441 | 0.20527 | 0.10053 | 0.48437 | 0.09986 | 20.89902 | 0.01 | 0 | Neutral |
| testneg_7809 | 0 | 0 | 0.764567 | -3 | -0.929745 | -3 | 0.06027 | 0.11574 | 0.09284 | 0.26053 | 4.44444 | 0.07 | 0 | Neutral |
| testneg_7810 | -0.01 | 0.57 | 0.083768 | -3 | 0.989458 | -3 | 0.08041 | 0.21802 | 0.1481 | 0.1117 | 18.6768 | 0.05 | 0 | Neutral |
| testneg_7811 | 0 | 0 | 1.468723 | -3 | 2.53141 | 2.53141 | 0.45152 | 0.34457 | 0.1123 | 0.1568 | 18.33333 | 0.284976732 | 1.8 | Neutral |
| testneg_7812 | 0 | 0 | 1.394643 | -3 | 5.540938 | -3 | 0.49629 | 0.11989 | 0.19898 | 0.12788 | 19.11765 | 0.078219697 | 0 | Neutral |
| testneg_7813 | -0.03 | 1.03 | 2.97112 | -3 | 2.151559 | -3 | 0.5901 | 0.19444 | 0.82591 | 0.12207 | 10.20833 | 0.17 | 0 | Neutral |
| testneg_7814 | 0 | 0 | -0.076285 | -3 | 0.105479 | -0.439393 | 0.23624 | 0.12788 | 0.35779 | 0.12788 | 33.56481 | 0.01 | 0 | Neutral |
| testneg_7815 | -0.02 | 0.63 | 2.813311 | -3 | 3.716565 | -3 | 0.10809 | 0.09057 | 0.1594 | 0.22002 | 13.02635 | 0.35785766 | 9.6 | Neutral |
| testneg_7816 | -0.01 | 0.57 | 1.162064 | -3 | 14.078442 | -3 | 0.64608 | 0.35524 | 0.10385 | 0.09775 | 9.79167 | 0.458771368 | 38.2 | Neutral |
| testneg_7817 | 0.02 | -0.28 | 5.210854 | -3 | -0.438666 | -3 | 0.46387 | 0.17039 | 0.0669 | 0.10997 | 32.29167 | 0.15 | 0 | Neutral |
| testneg_7818 | 0 | 0 | 2.165106 | -3 | -0.13961 | -3 | 0.30257 | 0.17298 | 0.65734 | 0.1013 | 30.01634 | 0.020094747 | 0 | Neutral |
| testneg_7819 | 0 | 0 | 0.47864 | -3 | -0.21206 | -3 | 0.17597 | 0.12788 | 0.4722 | 0.13321 | 20.83333 | 0.050733333 | 0 | Neutral |
| testneg_7820 | 0.02 | 0.45 | 5.903349 | -3 | -0.062057 | -3 | 0.49745 | 0.1259 | 0.26489 | 0.10549 | 31.99714 | 0.154411765 | 0 | Neutral |
| testneg_7821 | 0 | 0 | 1.051927 | -3 | 2.455596 | -3 | 0.9516 | 0.12788 | 0.19898 | 0.12788 | 9.64693 | 0.02 | 0 | Neutral |
| testneg_7822 | 0 | 0 | 0.051943 | -3 | -0.969128 | -3 | 0.48027 | 0.52799 | 0.09577 | 0.2436 | 11.25 | 0.118289298 | 0 | Neutral |
| testneg_7823 | 0.17 | 0.38 | 0.439346 | -3 | 4.429067 | -3 | 0.06072 | 0.09873 | 0.19898 | 0.09824 | 24.20314 | 0.093666667 | 0 | Neutral |
| testneg_7824 | 0.16 | -1.37 | 2.791385 | -3 | 4.996914 | 1.62118 | 0.59853 | 0.75237 | 0.09972 | 0.2187 | 19.44444 | 0.727187383 | 99.8 | Disease_causing |
| testneg_7825 | 0 | -0.32 | 3.107095 | -3 | -0.605051 | -3 | 0.19898 | 0.40608 | 0.48144 | 0.46642 | 6.21424 | 0.330425887 | 5.2 | Neutral |
| testneg_7826 | -0.07 | -0.38 | -0.007494 | -3 | 0.62394 | -3 | 0.46919 | 0.60081 | 0.90813 | 0.10329 | 8.91119 | 0.050135076 | 0 | Neutral |
| testneg_7827 | 0.06 | 0.18 | 5.333191 | -3 | 4.819528 | -3 | 0.19898 | 0.12788 | 0.1211 | 0.1526 | 20.87531 | 0.632157987 | 90.4 | Disease_causing |
| testneg_7828 | 0 | 0 | 0.301487 | -3 | 6.733244 | -3 | 0.93942 | 0.23984 | 0.07961 | 0.11474 | 20.20688 | 0.235 | 0.8 | Neutral |
| testneg_7829 | 0.08 | -1.2 | 0.227533 | -3 | 5.933886 | -3 | 0.46723 | 0.08832 | 0.09099 | 0.2051 | 9.82562 | 0.342861472 | 5.4 | Neutral |
| testneg_7830 | 0.05 | -0.81 | 1.441797 | -3 | 3.757908 | -3 | 0.14 | 0.12151 | 0.19898 | 0.0989 | 30.23582 | 0.05 | 0 | Neutral |
| testneg_7831 | 0.02 | 0.45 | 2.528558 | -3 | 1.026417 | -3 | 0.27004 | 0.17052 | 0.2543 | 0.12929 | 5.69444 | 0.03 | 0 | Neutral |
| testneg_7832 | -0.18 | -0.16 | 0.60982 | 0.60982 | -0.483923 | -3 | 0.23844 | 0.27772 | 0.23574 | 0.12788 | 10.26553 | 0.09 | 0 | Neutral |
| testneg_7833 | 0.03 | 1.22 | 0.949523 | 0.949523 | -0.889855 | -0.889855 | 0.10056 | 0.12608 | 0.042 | 0.07218 | 13.14029 | 0.07 | 0 | Neutral |
| testneg_7834 | 0.18 | 0.16 | 0.409204 | -3 | 6.916269 | -3 | 0.29417 | 0.10965 | 0.57188 | 0.34038 | 19.34591 | 0.426009091 | 28.2 | Neutral |
| testneg_7835 | 0 | -0.01 | 1.129155 | -3 | 9.146604 | -3 | 0.16835 | 0.09282 | 0.44923 | 0.12788 | 21.97068 | 0.28 | 1 | Neutral |
| testneg_7836 | 0.07 | -0.31 | 0.469669 | -3 | 2.367948 | -3 | 0.09556 | 0.10109 | 0.85781 | 0.22949 | 28.84995 | 0.11 | 0 | Neutral |
| testneg_7837 | 0.05 | 0.29 | 2.698795 | -3 | 5.481335 | -3 | 0.17806 | 0.12788 | 0.7482 | 0.18812 | 19.07749 | 0.412831349 | 26 | Neutral |
| testneg_7838 | 0.14 | -1.01 | 0.94469 | -3 | 3.353086 | -3 | 0.08629 | 0.12788 | 0.83205 | 0.40904 | 12.77665 | 0.23 | 0.2 | Neutral |
| testneg_7839 | 0 | 0 | 0.979853 | -3 | -0.119638 | -3 | 0.34859 | 0.09799 | 0.16379 | 0.31462 | 11.96702 | 0.030260135 | 0 | Neutral |
| testneg_7840 | -0.02 | 0.36 | 2.655425 | -3 | -0.823621 | -0.823621 | 0.19883 | 0.09434 | 0.27382 | 0.16898 | 13.14653 | 0.1 | 0 | Neutral |
| testneg_7841 | -0.08 | -0.03 | 6.09382 | -3 | -0.317708 | -3 | 0.8169 | 0.23301 | 0.25077 | 0.06261 | 10.41667 | 0.27 | 0.6 | Neutral |
| testneg_7842 | 0 | -0.01 | 0.36151 | 0.36151 | 6.206483 | -3 | 0.28878 | 0.16765 | 0.18861 | 0.10842 | 5.69444 | 0.183672014 | 0.4 | Neutral |
| testneg_7843 | 0.03 | 0.1 | -2.80816 | -3 | 1.255802 | -3 | 0.19898 | 0.0989 | 0.20546 | 0.10734 | 32.67189 | 0.013333333 | 0 | Neutral |
| testneg_7844 | -0.01 | -0.8 | 2.340871 | -3 | 6.315071 | -3 | 0.60152 | 0.12166 | 0.22225 | 0.11186 | 10 | 0.17 | 0 | Neutral |
| testneg_7845 | 0.11 | -0.19 | 3.662322 | -3 | 2.61677 | -3 | 0.49745 | 0.1103 | 0.60285 | 0.15179 | 16.8018 | 0.276120702 | 1.8 | Neutral |
| testneg_7846 | -0.19 | -0.51 | -0.445993 | -0.445993 | 2.091567 | -3 | 0.18324 | 0.13041 | 0.19898 | 0.12788 | 17.21847 | 0.06 | 0 | Neutral |
| testneg_7847 | 0.09 | -0.3 | -1.840654 | -3 | 1.266152 | -3 | 0.19898 | 0.12788 | 0.1846 | 0.375 | 18.88889 | 0.1 | 0 | Neutral |
| testneg_7848 | -0.01 | -0.8 | 1.212943 | -3 | 1.343895 | -3 | 0.22944 | 0.09039 | 0.35137 | 0.09161 | 28.62818 | 0.02 | 0 | Neutral |
| testneg_7849 | -0.01 | -0.79 | 1.351518 | -3 | -0.082314 | -3 | 0.58984 | 0.15588 | 0.98712 | 0.10969 | 5.69444 | 0.04 | 0 | Neutral |
| testneg_7850 | -0.01 | -0.8 | 3.942334 | -3 | 1.557993 | -3 | 0.09445 | 0.12788 | 0.19898 | 0.12788 | 14.16667 | 0.042361111 | 0 | Neutral |
| testneg_7851 | 0.15 | 0.06 | -0.364205 | -3 | -0.945156 | -3 | 0.45628 | 0.15819 | 0.29209 | 0.09791 | 19.09347 | 0.042643049 | 0 | Neutral |
| testneg_7852 | 0.02 | 0.45 | 0.215947 | -3 | 3.369606 | -3 | 0.0418 | 0.12788 | 0.15704 | 0.72174 | 18.6768 | 0.270636509 | 1 | Neutral |
| testneg_7853 | 0 | 0 | 2.819152 | -3 | 2.090774 | -3 | 0.06877 | 0.17777 | 0.35421 | 0.10694 | 13.51351 | 0.07 | 0 | Neutral |
| testneg_7854 | 0.03 | 0.12 | 7.128653 | -3 | 0.763116 | -3 | 0.31859 | 0.11262 | 0.33026 | 0.30576 | 12.29167 | 0.471808003 | 43.6 | Neutral |
| testneg_7855 | -0.03 | -1.22 | 0.713058 | -3 | 0.148561 | -3 | 0.14375 | 0.12788 | 0.2559 | 0.10848 | 21.63869 | 0.02 | 0 | Neutral |
| testneg_7856 | 0 | 0 | 0.495852 | -3 | 4.406099 | -3 | 0.21029 | 0.27753 | 0.93064 | 0.10933 | 18.88889 | 0.119047315 | 0 | Neutral |
| testneg_7857 | 0.04 | 2.01 | 0.977834 | -3 | 0.970904 | -3 | 0.67676 | 0.12788 | 0.06313 | 0.12788 | 23.65777 | 0.08 | 0 | Neutral |
| testneg_7858 | 0 | -0.01 | 0.545018 | -3 | 4.019935 | -3 | 0.11818 | 0.10561 | 0.13869 | 0.10313 | 20.84541 | 0.05 | 0 | Neutral |
| testneg_7859 | -0.15 | -0.04 | 0.179678 | -3 | 3.650653 | -3 | 0.31168 | 0.12788 | 0.07558 | 0.12788 | 19.44444 | 0.12 | 0 | Neutral |
| testneg_7860 | 0 | 0 | -0.272739 | -3 | 4.082785 | -3 | 0.10286 | 0.26819 | 0.07204 | 0.07708 | 17.52506 | 0.1485 | 0 | Neutral |
| testneg_7861 | 0 | -0.01 | 4.102872 | -3 | 5.468956 | -3 | 0.37463 | 0.23021 | 0.10604 | 0.10785 | 17.21847 | 0.355809255 | 10.4 | Neutral |
| testneg_7862 | -0.05 | -1.52 | 0.83408 | -3 | 0.09433 | -3 | 0.95857 | 0.12137 | 0.18925 | 0.32479 | 11.25 | 0.100794733 | 0 | Neutral |
| testneg_7863 | 0.15 | -0.28 | 4.506412 | -3 | -0.267215 | -3 | 0.15954 | 0.12788 | 0.89178 | 0.23115 | 20.95238 | 0.169424298 | 0 | Neutral |
| testneg_7864 | -0.04 | 1.36 | -1.284127 | -3 | -0.137475 | -3 | 0.11096 | 0.10074 | 0.19898 | 0.12788 | 52.77778 | 0.08 | 0 | Neutral |
| testneg_7865 | 0 | 0 | 1.526737 | -3 | 5.68574 | -3 | 0.78536 | 0.12709 | 0.17947 | 0.12797 | 18.88889 | 0.07175299 | 0 | Neutral |
| testneg_7866 | -0.06 | 0.57 | 0.686594 | -3 | 4.083472 | -3 | 0.12377 | 0.12788 | 0.16394 | 0.08512 | 29.08136 | 0.1 | 0 | Neutral |
| testneg_7867 | -0.03 | -0.1 | 0.212913 | -3 | 0.900845 | -3 | 0.11391 | 0.19434 | 0.94674 | 0.12413 | 17.21847 | 0.06 | 0 | Neutral |
| testneg_7868 | 0 | 0 | 3.506817 | -3 | 1.194108 | -3 | 0.23088 | 0.21602 | 0.55618 | 0.11352 | 18.22331 | 0.06 | 0 | Neutral |
| testneg_7869 | 0.06 | -0.57 | 0.297516 | 0.297516 | 5.146768 | -3 | 0.19294 | 0.10464 | 0.10379 | 0.34799 | 20.42553 | 0.39881685 | 17.6 | Neutral |
| testneg_7870 | 0 | 0 | 0.438443 | 0.438443 | 0.044424 | -3 | 0.36857 | 0.36135 | 0.06526 | 0.12788 | 19.37853 | 0.04 | 0 | Neutral |
| testneg_7871 | -0.09 | 0.3 | 1.16938 | -3 | 3.303858 | -3 | 0.32974 | 0.12245 | 0.4558 | 0.9277 | 17.77778 | 0.235467997 | 0.2 | Neutral |
| testneg_7872 | 0.15 | -0.28 | 0.994694 | -3 | -0.187363 | -3 | 0.70964 | 0.12663 | 0.90649 | 0.3543 | 12.29167 | 0.07833908 | 0 | Neutral |
| testneg_7873 | 0.14 | -1.01 | 2.081126 | -3 | 1.795418 | -3 | 0.2663 | 0.32806 | 0.11055 | 0.19829 | 8.50247 | 0.217656783 | 0 | Neutral |
| testneg_7874 | 0.01 | -0.57 | 2.700261 | -3 | 2.88545 | -3 | 0.76957 | 0.13588 | 0.09353 | 0.12788 | 8.97898 | 0.08 | 0 | Neutral |
| testneg_7875 | 0.05 | -0.81 | 0.331709 | 0.331709 | 1.663039 | -3 | 0.16534 | 0.26726 | 0.11395 | 0.26806 | 17.21847 | 0.210744828 | 0 | Neutral |
| testneg_7876 | 0.22 | -2.66 | 7.297508 | -3 | 2.997184 | -3 | 0.15848 | 0.12754 | 0.17394 | 0.15823 | 11.25 | 0.596782407 | 84.2 | Disease_causing |
| testneg_7877 | 0 | 0 | 0.021671 | -3 | 0.312568 | -3 | 0.13156 | 0.10621 | 0.18264 | 0.10206 | 13.51351 | 0 | 0 | Neutral |
| testneg_7878 | 0 | 0 | 0.126016 | -3 | -1.410124 | -3 | 0.90955 | 0.10096 | 0.33952 | 0.12788 | 26.84685 | 0.02 | 0 | Neutral |
| testneg_7879 | 0.18 | 0.16 | 0.416167 | -3 | 0.818598 | 0.818598 | 0.1224 | 0.11899 | 0.12739 | 0.12788 | 34.85447 | 0.03 | 0 | Neutral |
| testneg_7880 | 0.18 | -1.05 | 0.192071 | -3 | 2.644462 | -3 | 0.18312 | 0.12788 | 0.10803 | 0.14652 | 17.21847 | 0.160361111 | 0 | Neutral |
| testneg_7881 | 0.22 | -2.66 | 3.346037 | -3 | 1.036843 | -3 | 0.04916 | 0.14944 | 0.18102 | 0.6537 | 18.88889 | 0.37875 | 10 | Neutral |
| testneg_7882 | 0.09 | -0.67 | 2.624257 | -3 | 1.156857 | -3 | 0.19043 | 0.07855 | 0.19898 | 0.12788 | 23.68839 | 0.02 | 0 | Neutral |
| testneg_7883 | 0 | -0.01 | 1.920609 | -3 | -0.142841 | -3 | 0.24331 | 0.11911 | 0.40081 | 0.15418 | 11.02713 | 0.030931818 | 0 | Neutral |
| testneg_7884 | 0.03 | 0.12 | 1.249184 | -3 | 0.897808 | 0.897808 | 0.40592 | 0.22108 | 0.17155 | 0.18585 | 10.41667 | 0.112462444 | 0 | Neutral |
| testneg_7885 | 0.02 | -0.36 | -0.682977 | -3 | 2.656728 | -3 | 0.05444 | 0.12788 | 0.16966 | 0.61198 | 16.01246 | 0.190717949 | 0 | Neutral |
| testneg_7886 | -0.04 | 0.23 | -0.77433 | -3 | 1.418339 | -3 | 0.51167 | 0.13142 | 0.24185 | 0.12788 | 15.65227 | 0.03 | 0 | Neutral |
| testneg_7887 | -0.07 | -0.01 | 2.087185 | -3 | -0.274976 | -3 | 0.06392 | 0.12788 | 0.26767 | 0.25506 | 17.21847 | 0.09671875 | 0 | Neutral |
| testneg_7888 | 0.18 | -1.05 | 2.305068 | -3 | 3.022813 | -3 | 0.19898 | 0.12788 | 0.17301 | 0.10048 | 33.68056 | 0.08 | 0 | Neutral |
| testneg_7889 | 0.02 | 0.45 | 2.195573 | -3 | 5.840704 | -3 | 0.10774 | 0.13117 | 0.37909 | 0.4568 | 11.875 | 0.482486493 | 46.4 | Neutral |
| testneg_7890 | 0 | 0 | 1.362887 | -3 | 2.426709 | -3 | 0.38527 | 0.24812 | 0.13271 | 0.22958 | 13.60507 | 0.160557022 | 0 | Neutral |
| testneg_7891 | -0.08 | -0.03 | 3.415166 | -3 | 0.853858 | -3 | 0.67796 | 0.10874 | 0.35513 | 0.21966 | 20 | 0.149333261 | 0 | Neutral |
| testneg_7892 | 0.01 | 0.8 | 2.76764 | -3 | -0.341214 | -3 | 0.22795 | 0.12788 | 0.36813 | 0.09414 | 73.51648 | 0.089166667 | 0 | Neutral |
| testneg_7893 | 0.02 | 0.45 | 4.877158 | -3 | 3.398658 | -3 | 0.78211 | 0.35815 | 0.06971 | 0.09918 | 13.77727 | 0.291666667 | 3 | Neutral |
| testneg_7894 | -0.03 | -0.1 | 0.129998 | -3 | 0.282676 | -3 | 0.03137 | 0.12788 | 0.09576 | 0.06903 | 43.67192 | 0.05 | 0 | Neutral |
| testneg_7895 | 0.07 | 0.33 | 2.134433 | -3 | 3.271853 | -3 | 0.27887 | 0.19826 | 0.60152 | 0.12166 | 10 | 0.153809223 | 0 | Neutral |
| testneg_7896 | 0 | 0 | 0.383952 | -3 | 0.626243 | 0.626243 | 0.28629 | 0.77234 | 0.34412 | 0.08672 | 13.75 | 0.06 | 0 | Neutral |
| testneg_7897 | 0 | 0 | -0.287761 | -3 | -0.660027 | -3 | 0.85345 | 0.13067 | 0.15432 | 0.095 | 18.33333 | 0.013362905 | 0 | Neutral |
| testneg_7898 | -0.07 | -0.38 | -1.803731 | -3 | 1.182833 | -3 | 0.07963 | 0.23713 | 0.89661 | 0.17382 | 12.04793 | 0.16 | 0 | Neutral |
| testneg_7899 | -0.03 | -0.1 | 3.084101 | -3 | 1.440837 | -3 | 0.39015 | 0.23372 | 0.20219 | 0.10986 | 13.40796 | 0.074062749 | 0 | Neutral |
| testneg_7900 | 0 | 0.32 | -0.741073 | -3 | 1.604498 | -3 | 0.20191 | 0.13449 | 0.6992 | 0.1804 | 10.83333 | 0.050228334 | 0 | Neutral |
| testneg_7901 | 0.22 | -2.66 | 4.95856 | -3 | 1.078547 | -3 | 0.36541 | 0.27749 | 0.13749 | 0.17824 | 19.09347 | 0.38 | 12.6 | Neutral |
| testneg_7902 | 0 | 0 | 0.488617 | -3 | 2.91359 | 2.91359 | 0.23088 | 0.21602 | 0.19898 | 0.12788 | 13.43868 | 0.13 | 0 | Neutral |
| testneg_7903 | -0.02 | 0.63 | 5.119073 | -3 | 2.087185 | -3 | 0.29679 | 0.12788 | 0.06392 | 0.12788 | 16.8018 | 0.14 | 0 | Neutral |
| testneg_7904 | 0 | 0 | 1.243121 | -3 | 1.085971 | -3 | 0.15958 | 0.41628 | 0.14941 | 0.0838 | 27.08333 | 0.033385471 | 0 | Neutral |
| testneg_7905 | 0.07 | 0.01 | 1.714571 | -3 | 3.518471 | 3.518471 | 0.32022 | 0.40347 | 0.19821 | 0.20954 | 18.33333 | 0.548028936 | 75.4 | Disease_causing |
| testneg_7906 | 0 | 0 | -0.626667 | -3 | -0.55851 | -3 | 0.21933 | 0.18619 | 0.06874 | 0.12788 | 11.58784 | 0.011428571 | 0 | Neutral |
| testneg_7907 | 0 | 0 | 2.124149 | -3 | 0.828771 | -3 | 0.14482 | 0.08592 | 0.71844 | 0.20204 | 12.96895 | 0.03 | 0 | Neutral |
| testneg_7908 | 0 | 0 | 0.383044 | -3 | -0.140634 | -3 | 0.63819 | 0.32258 | 0.19898 | 0.12788 | 10.60885 | 0.01028629 | 0 | Neutral |
| testneg_7909 | 0.07 | 0.7 | 5.847469 | -3 | 7.215815 | -3 | 0.28341 | 0.1005 | 0.36858 | 0.09681 | 10.02983 | 0.64 | 93 | Disease_causing |
| testneg_7910 | 0 | 0 | 1.169064 | -3 | 0.480758 | -3 | 0.51319 | 0.12971 | 0.11584 | 0.19704 | 10 | 0.04 | 0 | Neutral |
| testneg_7911 | 0 | 0 | -0.481968 | -3 | 1.885526 | -3 | 0.67331 | 0.10118 | 0.11233 | 0.28224 | 10.41667 | 0.058969016 | 0 | Neutral |
| testneg_7912 | -0.07 | -0.38 | 0.72503 | -3 | 3.136838 | -3 | 0.56862 | 0.12788 | 0.07999 | 0.10413 | 9.79693 | 0.06 | 0 | Neutral |
| testneg_7913 | -0.01 | -0.8 | -0.808449 | -0.808449 | 3.009853 | -3 | 0.13258 | 0.9258 | 0.50741 | 0.17915 | 16.92155 | 0.350959355 | 8 | Neutral |
| testneg_7914 | -0.08 | -0.03 | 1.562114 | -3 | 0.84675 | -3 | 0.78356 | 0.13377 | 0.4358 | 0.1222 | 8.8786 | 0.010506884 | 0 | Neutral |
| testneg_7915 | 0 | 0 | 0.080104 | -3 | 1.902977 | 1.902977 | 0.73947 | 0.2058 | 0.18504 | 0.0822 | 26.64954 | 0.06 | 0 | Neutral |
| testneg_7916 | 0 | 0 | 1.636782 | -3 | 0.877962 | -3 | 0.15107 | 0.10457 | 0.5043 | 0.3225 | 18.33333 | 0.032467949 | 0 | Neutral |
| testneg_7917 | 0.14 | -0.37 | 4.085645 | -3 | 6.150547 | -3 | 0.45299 | 0.16306 | 0.16121 | 0.10081 | 44.24616 | 0.394471821 | 20.4 | Neutral |
| testneg_7918 | 0.01 | 0.8 | 2.947434 | -3 | 6.89 | -3 | 0.27915 | 0.27706 | 0.25473 | 0.07863 | 10.16391 | 0.469789561 | 41.4 | Neutral |
| testneg_7919 | -0.03 | 0.85 | 2.004649 | -3 | 5.560629 | -3 | 0.10822 | 0.10942 | 0.19898 | 0.09515 | 40.39304 | 0.148365915 | 0 | Neutral |
| testneg_7920 | 0 | 0 | 1.540756 | -3 | 4.182878 | -3 | 0.10249 | 0.1399 | 0.22998 | 0.08738 | 16.03108 | 0.070595169 | 0 | Neutral |
| testneg_7921 | 0 | 0 | 2.53141 | 2.53141 | 0.36854 | -3 | 0.1123 | 0.1568 | 0.58292 | 0.87725 | 20.83333 | 0.39625 | 23.6 | Neutral |
| testneg_7922 | 0.07 | 0.01 | -0.285825 | -3 | 5.393665 | 5.393665 | 0.15016 | 0.08826 | 0.03809 | 0.16044 | 11.25 | 0.448333333 | 32.4 | Neutral |
| testneg_7923 | 0 | 0 | 0.984412 | -3 | -0.100872 | -3 | 0.34931 | 0.11906 | 0.11891 | 0.11186 | 17.70469 | 0 | 0 | Neutral |
| testneg_7924 | 0 | 0 | 0.202655 | -3 | 1.342014 | -3 | 0.18834 | 0.12788 | 0.21288 | 0.10049 | 11.00467 | 0.000266714 | 0 | Neutral |
| testneg_7925 | 0 | -0.32 | 2.305189 | -3 | 5.182475 | -3 | 0.505 | 0.12788 | 0.18122 | 0.16895 | 33.56481 | 0.406787135 | 19.6 | Neutral |
| testneg_7926 | -0.06 | -0.18 | 2.578731 | -3 | 5.907423 | -3 | 0.18283 | 0.11376 | 0.00451 | 0.2022 | 12.80035 | 0.490849967 | 51.4 | Disease_causing |
| testneg_7927 | 0.05 | -0.81 | -0.021519 | -3 | -0.072198 | -0.072198 | 0.30096 | 0.17203 | 0.1025 | 0.3268 | 8.86674 | 0.154916667 | 0 | Neutral |
| testneg_7928 | 0.03 | 0.12 | 0.367581 | -3 | 2.435234 | -3 | 0.13203 | 0.12092 | 0.56025 | 0.12788 | 18.6768 | 0.02 | 0 | Neutral |
| testneg_7929 | 0 | 0 | 0.802919 | 0.802919 | 2.643915 | -3 | 0.12042 | 0.11067 | 0.29548 | 0.10732 | 22.08193 | 0.040322917 | 0 | Neutral |

| | | | | | | | | | | | | | | |
|---|---|---|---|---|---|---|---|---|---|---|---|---|---|---|
| testneg_7930 | -0.07 | -0.38 | 1.366318 | -3 | -0.509288 | -3 | 0.78536 | 0.12709 | 0.178 | 0.7168 | 10.41667 | 0.061 | 0 | Neutral |
| testneg_7931 | 0.18 | 0.16 | 1.782077 | -3 | 3.046889 | -3 | 0.60793 | 0.18264 | 0.1066 | 0.12788 | 10.83333 | 0.1825 | 0 | Neutral |
| testneg_7932 | -0.01 | 0.57 | 2.71072 | -3 | -0.280358 | -0.280358 | 0.09719 | 0.09576 | 0.30657 | 0.52974 | 11.875 | 0.14 | 0 | Neutral |
| testneg_7933 | 0 | 0 | 1.716798 | -3 | 3.441233 | -3 | 0.70349 | 0.12788 | 0.22046 | 0.12788 | 10.83333 | 0.039805556 | 0 | Neutral |
| testneg_7934 | 0 | 0 | -0.001616 | -3 | -0.200198 | -3 | 0.90767 | 0.12788 | 0.2771 | 0.14453 | 10.41667 | 0.028802836 | 0 | Neutral |
| testneg_7935 | 0 | 0 | 1.221853 | -3 | 0.331709 | 0.331709 | 0.92659 | 0.13588 | 0.16534 | 0.26726 | 23.86511 | 0.119401786 | 0 | Neutral |
| testneg_7936 | 0 | 0 | 1.91859 | -3 | 3.381673 | 3.381673 | 0.53861 | 0.18604 | 0.23578 | 0.12788 | 10.41667 | 0.212981754 | 0.4 | Neutral |
| testneg_7937 | -0.01 | 0.57 | 5.981424 | -3 | 5.007683 | -3 | 0.1196 | 0.12788 | 0.07415 | 0.12788 | 24.68926 | 0.44 | 32.8 | Neutral |
| testneg_7938 | 0 | 0 | -0.961702 | -0.961702 | 0.744084 | -3 | 0.46979 | 0.1705 | 0.1311 | 0.12667 | 18.36272 | 0.02 | 0 | Neutral |
| testneg_7939 | 0.03 | 0.1 | 0.323324 | -3 | 2.200422 | -3 | 0.84106 | 0.63405 | 0.1662 | 0.16982 | 10.41667 | 0.17641533 | 0 | Neutral |
| testneg_7940 | 0 | 0 | -0.100514 | -3 | 4.654999 | -3 | 0.16469 | 0.16042 | 0.10183 | 0.11847 | 21.66786 | 0.152132035 | 0 | Neutral |
| testneg_7941 | 0 | 0 | 0.342408 | -3 | 0.876243 | -3 | 0.5255 | 0.15968 | 0.694 | 0.09974 | 19.90375 | 0.022 | 0 | Neutral |
| testneg_7942 | -0.02 | -0.45 | 3.615951 | -3 | 4.751408 | -3 | 0.73808 | 0.15588 | 0.06301 | 0.32138 | 24.09745 | 0.649731746 | 95.8 | Disease_causing |
| testneg_7943 | 0 | 0 | 1.861867 | -3 | 0.281426 | -3 | 0.32963 | 0.1035 | 0.23844 | 0.27772 | 13.03571 | 0.023511905 | 0 | Neutral |
| testneg_7944 | 0 | 0.32 | 2.156711 | -3 | 2.098021 | -3 | 0.16965 | 0.25991 | 0.08184 | 0.10206 | 12.37434 | 0.040294118 | 0 | Neutral |
| testneg_7945 | 0 | 0 | -0.090459 | -3 | 0.410336 | -3 | 0.15581 | 0.19351 | 0.096 | 0.09851 | 10.83333 | 0.02 | 0 | Neutral |
| testneg_7946 | 0 | 0 | 1.200491 | -3 | 3.570813 | -3 | 0.34198 | 0.1113 | 0.68054 | 0.09208 | 18.38327 | 0.031010738 | 0 | Neutral |
| testneg_7947 | -0.14 | 0.37 | 2.640854 | -3 | 5.728641 | -3 | 0.20072 | 0.08606 | 0.27806 | 0.09713 | 19.44444 | 0.206515152 | 0 | Neutral |
| testneg_7948 | -0.06 | 0.57 | 1.785637 | -3 | 0.60135 | 0.147975 | 0.21811 | 0.18489 | 0.06898 | 0.11659 | 28.72155 | 0.05 | 0 | Neutral |
| testneg_7949 | -0.16 | -0.08 | -1.975003 | -1.975003 | 2.173968 | -3 | 0.08465 | 0.15428 | 0.19898 | 0.12788 | 20.95603 | 0.150283859 | 0 | Neutral |
| testneg_7950 | 0 | -0.01 | 0.765653 | -3 | 1.73925 | -3 | 0.74099 | 0.44443 | 0.12971 | 0.11434 | 6.33755 | 0.01 | 0 | Neutral |
| testneg_7951 | 0 | 0 | 0.717822 | -3 | -0.214756 | -3 | 0.29044 | 0.12142 | 0.07428 | 0.07645 | 20.85305 | 0.01 | 0 | Neutral |
| testneg_7952 | 0 | 0 | 0.132481 | -3 | 0.287736 | -3 | 0.33026 | 0.30576 | 0.16116 | 0.08314 | 12.75394 | 0.01 | 0 | Neutral |
| testneg_7953 | -0.07 | -0.01 | 0.439914 | -3 | 0.65067 | -3 | 0.09275 | 0.14911 | 0.25312 | 0.12788 | 18.8735 | 0.03 | 0 | Neutral |
| testneg_7954 | 0.01 | 0.8 | 7.121247 | -3 | 1.668583 | -3 | 0.48075 | 0.20313 | 0.04869 | 0.05922 | 10.83333 | 0.412794794 | 21.6 | Neutral |
| testneg_7955 | 0 | 0 | 1.639442 | -3 | 2.199122 | -3 | 0.25905 | 0.23728 | 0.46058 | 0.12903 | 13.91548 | 0.020996026 | 0 | Neutral |
| testneg_7956 | 0 | 0 | 0.867971 | -3 | 1.661324 | -3 | 0.24966 | 0.12788 | 0.16524 | 0.12788 | 8.91119 | 0 | 0 | Neutral |
| testneg_7957 | -0.03 | -0.1 | 0.545839 | -3 | 5.475217 | -3 | 0.58545 | 0.14414 | 0.54625 | 0.11856 | 19.31102 | 0.133333333 | 0 | Neutral |
| testneg_7958 | 0 | 0 | 1.075324 | 1.075324 | -0.513734 | -3 | 0.49934 | 0.11069 | 0.256 | 0.20036 | 7.10199 | 0.081946614 | 0 | Neutral |
| testneg_7959 | 0 | 0 | 0.305495 | -3 | -1.13933 | -3 | 0.98255 | 0.98129 | 0.11703 | 0.09364 | 10.22898 | 0.070221579 | 0 | Neutral |
| testneg_7960 | -0.05 | 0.81 | 1.410504 | -3 | 3.646993 | -3 | 0.1947 | 0.11413 | 0.7062 | 0.83002 | 10.73625 | 0.256791056 | 0.8 | Neutral |
| testneg_7961 | 0 | 0 | 0.29615 | -3 | -0.562184 | -3 | 0.17325 | 0.17964 | 0.03466 | 0.19518 | 19.2404 | 0.113274156 | 0 | Neutral |
| testneg_7962 | 0 | 0 | 1.052386 | -3 | 1.086534 | -3 | 0.87038 | 0.43956 | 0.17006 | 0.14718 | 17.56627 | 0.073516384 | 0 | Neutral |
| testneg_7963 | 0 | 0 | -0.681556 | -3 | 5.797326 | -3 | 0.42585 | 0.18081 | 0.76942 | 0.13267 | 20.83333 | 0.319067943 | 2.8 | Neutral |
| testneg_7964 | 0 | -0.01 | 3.021892 | -3 | -3.299404 | -3 | 0.1163 | 0.20863 | 0.20849 | 0.12788 | 20.83333 | 0.088333333 | 0 | Neutral |
| testneg_7965 | -0.14 | 0.69 | 0.437223 | -3 | 0.742409 | -3 | 0.0684 | 0.08803 | 0.12949 | 0.44208 | 21.19188 | 0.140188912 | 0 | Neutral |
| testneg_7966 | 0.16 | 0.08 | 2.869124 | -3 | -1.184577 | -3 | 0.07837 | 0.12788 | 0.34826 | 0.49454 | 17.05011 | 0.179111722 | 0 | Neutral |
| testneg_7967 | 0.21 | 0.15 | 3.847333 | 3.659652 | 3.846151 | -3 | 0.53627 | 0.12788 | 0.46735 | 0.10674 | 34.29705 | 0.448376623 | 33.2 | Neutral |
| testneg_7968 | 0.01 | 0.8 | 7.39468 | -3 | 5.877627 | -3 | 0.1417 | 0.1398 | 0.2559 | 0.10848 | 20.52758 | 0.649357143 | 95.6 | Disease_causing |
| testneg_7969 | 0.19 | -0.27 | 5.724264 | -3 | 3.58202 | -3 | 0.39964 | 0.22293 | 0.09633 | 0.12788 | 18.59477 | 0.458684211 | 38.6 | Neutral |
| testneg_7970 | 0 | 0 | 0.297064 | -3 | 6.206483 | -3 | 0.2029 | 0.12788 | 0.18861 | 0.10842 | 20.5518 | 0.078303571 | 0 | Neutral |
| testneg_7971 | -0.01 | -0.79 | 3.399462 | -3 | 0.680291 | -3 | 0.05472 | 0.12788 | 0.23574 | 0.12788 | 37.4774 | 0.070346947 | 0 | Neutral |
| testneg_7972 | 0 | 0 | 1.193153 | -3 | 0.579456 | 0.579456 | 0.12259 | 0.1041 | 0.47678 | 0.38085 | 9.84405 | 0.060933626 | 0 | Neutral |
| testneg_7973 | 0 | 0 | 0.453556 | -3 | 0.410561 | -3 | 0.28191 | 0.20693 | 0.19898 | 0.19444 | 10.41667 | 0.064435541 | 0 | Neutral |
| testneg_7974 | 0.05 | -0.81 | 2.225911 | -3 | 6.526897 | -3 | 0.91619 | 0.10042 | 0.16692 | 0.09243 | 12.75292 | 0.180238095 | 0.2 | Neutral |
| testneg_7975 | -0.04 | -2.01 | 2.355419 | -3 | 2.815771 | -3 | 0.82595 | 0.52905 | 0.03558 | 0.05202 | 19.20245 | 0.138407383 | 0 | Neutral |
| testneg_7976 | 0.07 | -0.31 | 0.040505 | -3 | 0.292074 | -3 | 0.16926 | 0.11871 | 0.65952 | 0.49708 | 17.77778 | 0.050536183 | 0 | Neutral |
| testneg_7977 | -0.07 | -0.38 | -1.63682 | -3 | 14.612558 | -3 | 0.13505 | 0.09019 | 0.96704 | 0.20571 | 8.49181 | 0.46 | 37 | Neutral |
| testneg_7978 | 0.05 | -0.81 | 0.668367 | -3 | 8.044918 | -3 | 0.03656 | 0.07074 | 0.06301 | 0.32138 | 38.23099 | 0.436625 | 29.8 | Neutral |
| testneg_7979 | 0.09 | -0.3 | 5.573517 | -3 | 2.442902 | -3 | 0.09416 | 0.17821 | 0.19898 | 0.12788 | 29.9024 | 0.295208333 | 3.4 | Neutral |
| testneg_7980 | 0.07 | -0.31 | 2.014715 | -3 | 0.106535 | -3 | 0.99859 | 0.12788 | 0.44429 | 0.20468 | 10.41667 | 0.08 | 0 | Neutral |
| testneg_7981 | 0.16 | -1.37 | 2.859591 | -3 | 5.054815 | 5.054815 | 0.1576 | 0.12788 | 0.69563 | 0.12788 | 10.83333 | 0.45 | 35.4 | Neutral |
| testneg_7982 | -0.14 | 0.69 | 3.370031 | -3 | 5.87251 | -3 | 0.71814 | 0.72461 | 0.11827 | 0.23034 | 4.72222 | 0.740066427 | 98.6 | Disease_causing |
| testneg_7983 | -0.07 | -0.38 | 2.424425 | -3 | 0.052371 | 0.052371 | 0.54211 | 0.12592 | 0.82107 | 0.27926 | 6.80941 | 0.073358586 | 0 | Neutral |
| testneg_7984 | -0.07 | -0.38 | -0.872044 | -3 | -3.265819 | -3 | 0.04792 | 0.12788 | 0.04519 | 0.12788 | 25.6804 | 0.06 | 0 | Neutral |
| testneg_7985 | -0.01 | 1.44 | 0.734765 | -3 | 1.183533 | -3 | 0.07204 | 0.07708 | 0.11042 | 0.09881 | 28.22366 | 0.08 | 0 | Neutral |
| testneg_7986 | -0.09 | 0.25 | -2.500749 | -3 | 6.982059 | -3 | 0.35422 | 0.11185 | 0.23691 | 0.1015 | 48.55104 | 0.28 | 1.8 | Neutral |
| testneg_7987 | 0.09 | -0.67 | 0.066913 | -3 | 0.513578 | -3 | 0.53239 | 0.17509 | 0.19898 | 0.07409 | 15.46432 | 0.04 | 0 | Neutral |
| testneg_7988 | 0.15 | -0.28 | 1.567816 | -3 | -0.867132 | -3 | 0.80979 | 0.5595 | 0.08353 | 0.21479 | 10.41667 | 0.15 | 0 | Neutral |
| testneg_7989 | -0.06 | -0.18 | 2.70043 | -3 | -1.914326 | -3 | 0.33082 | 0.12788 | 0.2124 | 0.17288 | 28.34879 | 0.102092803 | 0 | Neutral |
| testneg_7990 | 0 | 0 | 0.190753 | -3 | -0.142939 | -3 | 0.85984 | 0.40044 | 0.11254 | 0.19973 | 12.23844 | 0.089944139 | 0 | Neutral |
| testneg_7991 | 0 | 0 | -0.344792 | -3 | 7.35694 | -3 | 0.19898 | 0.58572 | 0.32669 | 0.10659 | 6.80556 | 0.371992063 | 13.8 | Neutral |
| testneg_7992 | -0.21 | -0.15 | 0.197741 | -3 | 5.632523 | -3 | 0.31923 | 0.17492 | 0.17764 | 0.09681 | 6.23829 | 0.292819039 | 2.8 | Neutral |
| testneg_7993 | 0 | -0.01 | 6.62337 | -3 | 0.098665 | 0.098665 | 0.04208 | 0.12788 | 0.19898 | 0.12788 | 18.6768 | 0.223333333 | 0.6 | Neutral |
| testneg_7994 | 0.03 | 0.12 | 3.356959 | -3 | 0.939929 | -3 | 0.3922 | 0.65186 | 0.18701 | 0.12788 | 19.09347 | 0.120330393 | 0 | Neutral |
| testneg_7995 | 0 | 0 | 0.419995 | -3 | -1.561199 | -3 | 0.7174 | 0.15815 | 0.07617 | 0.07182 | 12.35574 | 0.01141129 | 0 | Neutral |
| testneg_7996 | 0.06 | -0.57 | 0.297516 | 0.297516 | 2.44605 | -3 | 0.19294 | 0.10464 | 0.19898 | 0.06726 | 49.3883 | 0.03 | 0 | Neutral |
| testneg_7997 | -0.07 | -0.38 | 0.158286 | -3 | 0.489487 | 0.489487 | 0.10133 | 0.32155 | 0.1462 | 0.18896 | 10 | 0.191553175 | 0 | Neutral |
| testneg_7998 | 0.05 | -0.81 | 5.789217 | -3 | -0.092233 | -3 | 0.26668 | 0.11598 | 0.213 | 0.09535 | 21.42525 | 0.11 | 0 | Neutral |
| testneg_7999 | 0.02 | -0.63 | -0.055258 | -3 | 3.817144 | -3 | 0.16139 | 0.09116 | 0.12321 | 0.08084 | 29.9024 | 0.06578373 | 0 | Neutral |
| testneg_8000 | 0.05 | -0.81 | 0.660523 | -3 | 1.621831 | -3 | 0.06897 | 0.08445 | 0.31434 | 0.11315 | 25.02646 | 0.010133714 | 0 | Neutral |
| testneg_8001 | 0 | 0 | -0.32829 | -3 | 0.02851 | -3 | 0.18509 | 0.12831 | 0.44594 | 0.13793 | 18.88883 | 0.032352041 | 0 | Neutral |
| testneg_8002 | -0.18 | 1.05 | 3.906443 | -3 | 0.222999 | -3 | 0.19898 | 0.12788 | 0.1201 | 0.1062 | 11.56063 | 0.13 | 0 | Neutral |
| testneg_8003 | 0.07 | 0.38 | -2.242853 | -3 | 4.126674 | -3 | 0.06358 | 0.5364 | 0.08579 | 0.6068 | 13.76815 | 0.591519841 | 85 | Disease_causing |
| testneg_8004 | 0 | -0.01 | 0.639661 | -3 | -2.092635 | -3 | 0.16263 | 0.53493 | 0.06704 | 0.1068 | 23.68839 | 0.05 | 0 | Neutral |
| testneg_8005 | 0 | 0 | 2.197785 | -3 | 0.66275 | -3 | 0.39863 | 0.44944 | 0.30429 | 0.29938 | 4.72222 | 0.095034776 | 0 | Neutral |
| testneg_8006 | -0.18 | -0.16 | 5.532186 | -3 | 1.550276 | -3 | 0.77944 | 0.1858 | 0.18142 | 0.21216 | 10.83333 | 0.559449405 | 74.6 | Disease_causing |
| testneg_8007 | -0.01 | 0.57 | -0.767729 | -3 | 7.667553 | -3 | 0.1289 | 0.14878 | 0.17697 | 0.12974 | 27.5592 | 0.46 | 39.4 | Neutral |
| testneg_8008 | 0.003 | -0.1380909 | 2.674709 | -3 | 2.690946 | -3 | 0.6185 | 0.09316 | 0.06723 | 0.10296 | 10.69779 | 0.079206183 | 0 | Neutral |
| testneg_8009 | 0.14 | -1.01 | 5.471755 | -3 | 0.90228 | -3 | 0.1784 | 0.12788 | 0.10058 | 0.13882 | 10.13889 | 0.252989899 | 1.2 | Neutral |
| testneg_8010 | 0.05 | -0.81 | 1.909741 | -3 | 6.127805 | -3 | 0.05898 | 0.12788 | 0.67549 | 0.1273 | 17.42866 | 0.19 | 0 | Neutral |
| testneg_8011 | -0.07 | -0.33 | 6.123714 | -3 | -0.126067 | -3 | 0.0893 | 0.11207 | 0.19898 | 0.09171 | 32.30375 | 0.14 | 0 | Neutral |
| testneg_8012 | -0.08 | -0.03 | 4.036486 | -3 | 4.51671 | -3 | 0.35422 | 0.11185 | 0.12073 | 0.10969 | 19.73833 | 0.179635756 | 0 | Neutral |
| testneg_8013 | 0.08 | 0.03 | 2.365497 | -3 | 1.623152 | -3 | 0.61491 | 0.11954 | 0.19898 | 0.12788 | 18.60587 | 0.023143939 | 0 | Neutral |
| testneg_8014 | 0 | 0 | 1.836976 | -3 | 4.478975 | -3 | 0.84491 | 0.30945 | 0.85781 | 0.22949 | 8.05313 | 0.283781555 | 3.2 | Neutral |
| testneg_8015 | -0.18 | -0.16 | 0.455154 | -3 | 1.628872 | 1.26926 | 0.39648 | 0.97288 | 0.38934 | 0.15403 | 5.69444 | 0.25 | 0 | Neutral |
| testneg_8016 | 0 | 0 | 1.27406 | -3 | -0.814463 | -3 | 0.37855 | 0.47307 | 0.28334 | 0.10368 | 18.6768 | 0.02 | 0 | Neutral |
| testneg_8017 | 0 | 0 | 0.498058 | -3 | 12.427766 | -3 | 0.10581 | 0.11631 | 0.12871 | 0.29252 | 20.8819 | 0.475565018 | 45.4 | Neutral |
| testneg_8018 | 0 | -0.01 | 0.912506 | -3 | -0.330967 | -0.330967 | 0.38429 | 0.11956 | 0.20657 | 0.12788 | 10.41667 | 8.82E-05 | 0 | Neutral |
| testneg_8019 | -0.0033333 | -0.1675 | 0.051852 | -3 | -0.144702 | -3 | 0.06311 | 0.14802 | 0.28162 | 0.12788 | 27.62126 | 0.05 | 0 | Neutral |
| testneg_8020 | 0 | 0 | 0.123542 | -3 | -0.139258 | -3 | 0.99915 | 0.96333 | 0.31025 | 0.09491 | 8.05313 | 0.1 | 0 | Neutral |
| testneg_8021 | -0.02 | 0.68 | -0.009485 | -3 | 3.866074 | -3 | 0.08186 | 0.11813 | 0.04829 | 0.09848 | 32.23427 | 0.100292869 | 0 | Neutral |
| testneg_8022 | 0 | 0 | -0.299858 | -3 | 2.550881 | -3 | 0.29284 | 0.11262 | 0.19898 | 0.12788 | 17.62359 | 0.01 | 0 | Neutral |
| testneg_8023 | -0.02 | 0.68 | 5.845448 | -3 | 1.123926 | -0.179933 | 0.06926 | 0.08399 | 0.45281 | 0.21621 | 25.9009 | 0.380459831 | 13 | Neutral |
| testneg_8024 | 0.02 | -0.63 | -2.196989 | -3 | -0.223591 | -0.223591 | 0.02759 | 0.12458 | 0.48705 | 0.09993 | 13.51351 | 0.040325407 | 0 | Neutral |
| testneg_8025 | 0 | 0 | 0.193332 | -3 | 1.300657 | -3 | 0.17264 | 0.09426 | 0.12317 | 0.12345 | 28.47222 | 0.01 | 0 | Neutral |
| testneg_8026 | -0.03 | 1.03 | 0.958989 | -3 | 7.308333 | -3 | 0.39436 | 0.16446 | 0.10822 | 0.10942 | 23.47355 | 0.414279614 | 22.8 | Neutral |
| testneg_8027 | 0.07 | 0.38 | 2.405373 | -3 | 0.748878 | -3 | 0.19898 | 0.13495 | 0.93064 | 0.10933 | 12.29167 | 0.02 | 0 | Neutral |
| testneg_8028 | 0 | 0 | 3.466764 | -3 | -1.556127 | -3 | 0.34205 | 0.11195 | 0.98339 | 0.18221 | 18.88889 | 0.110434856 | 0 | Neutral |
| testneg_8029 | -0.14 | -0.54 | -0.251883 | -3 | 6.913969 | -3 | 0.39258 | 0.12788 | 0.15267 | 0.12788 | 32.48176 | 0.291428571 | 2.2 | Neutral |
| testneg_8030 | 0 | 0 | -0.961702 | -0.961702 | 0.886934 | -3 | 0.46979 | 0.1705 | 0.64749 | 0.14763 | 30.94249 | 0.08 | 0 | Neutral |
| testneg_8031 | 0 | 0 | 1.687932 | -3 | -1.973513 | -3 | 0.16299 | 0.10454 | 0.67796 | 0.10874 | 11.65512 | 0.01 | 0 | Neutral |
| testneg_8032 | 0.14 | -1.01 | 0.877962 | -3 | 0.284309 | -3 | 0.5043 | 0.3225 | 0.18882 | 0.14637 | 22.2049 | 0.1620244 | 0 | Neutral |
| testneg_8033 | -0.07 | -0.7 | 3.861511 | -3 | 3.780286 | -3 | 0.09018 | 0.16175 | 0.19721 | 0.12485 | 10 | 0.344123016 | 6.2 | Neutral |
| testneg_8034 | 0.2 | -1.26 | 1.467073 | -3 | 7.005219 | -3 | 0.50562 | 0.2196 | 0.43977 | 0.17284 | 10.22898 | 0.650576979 | 97.4 | Disease_causing |
| testneg_8035 | 0 | 0 | 1.122479 | -3 | 3.487162 | -3 | 0.53654 | 0.22867 | 0.75906 | 0.09702 | 18.9586 | 0.09094335 | 0 | Neutral |
| testneg_8036 | -0.08 | -0.03 | 1.50021 | -3 | 1.517974 | -3 | 0.27618 | 0.11809 | 0.11373 | 0.10739 | 18.33333 | 0.01 | 0 | Neutral |
| testneg_8037 | -0.02 | 0.36 | 3.14455 | -3 | 0.925915 | -3 | 0.31434 | 0.11315 | 0.18142 | 0.21216 | 11.25 | 0.140652244 | 0 | Neutral |
| testneg_8038 | 0 | -0.01 | -0.496432 | -3 | 1.397648 | -3 | 0.06247 | 0.14184 | 0.11006 | 0.20781 | 10.41667 | 0.090300179 | 0 | Neutral |
| testneg_8039 | 0 | 0 | 0.494101 | -3 | 0.55478 | -3 | 0.28038 | 0.11814 | 0.22258 | 0.10471 | 24.35185 | 9.17E-05 | 0 | Neutral |
| testneg_8040 | -0.08 | -0.03 | 1.50021 | -3 | 11.796446 | -3 | 0.27618 | 0.11809 | 0.13626 | 0.12788 | 22.11623 | 0.37125 | 11.4 | Neutral |
| testneg_8041 | -0.02 | 0.28 | 1.48034 | -3 | -1.050714 | -3 | 0.18055 | 0.12564 | 0.09515 | 0.07784 | 20.41417 | 0.01 | 0 | Neutral |
| testneg_8042 | 0 | 0 | 0.299785 | -3 | -0.113262 | -2.712847 | 0.64152 | 0.15239 | 0.56154 | 0.0939 | 21.95402 | 0.02 | 0 | Neutral |
| testneg_8043 | 0 | 0 | 1.255675 | -3 | -0.076459 | -3 | 0.20675 | 0.13709 | 0.96377 | 0.1167 | 4.72222 | 0.01 | 0 | Neutral |
| testneg_8044 | -0.14 | 1.85 | 3.890081 | -3 | -0.061002 | -0.061002 | 0.19898 | 0.58778 | 0.35561 | 0.08845 | 23.72519 | 0.23 | 0 | Neutral |
| testneg_8045 | 0 | 0 | 2.165106 | -3 | 2.637768 | -3 | 0.30257 | 0.17298 | 0.30209 | 0.14229 | 22.0922 | 0.173124008 | 0 | Neutral |
| testneg_8046 | 0 | 0 | -1.230837 | -1.230837 | 2.614937 | -3 | 0.14821 | 0.11262 | 0.14185 | 0.12788 | 31.35135 | 0.02 | 0 | Neutral |
| testneg_8047 | 0 | 0 | 3.637193 | -3 | -0.454393 | -3 | 0.99976 | 0.28689 | 0.03443 | 0.1032 | 10.41667 | 0.090678733 | 0 | Neutral |
| testneg_8048 | 0.02 | -0.68 | 2.497761 | -3 | 6.282934 | 2.009811 | 0.16728 | 0.11868 | 0.05087 | 0.09682 | 11.875 | 0.24 | 0 | Neutral |
| testneg_8049 | 0 | 0 | 0.002341 | -3 | 0.148971 | -3 | 0.10799 | 0.10754 | 0.17007 | 0.12771 | 30.45796 | 0.01 | 0 | Neutral |
| testneg_8050 | 0 | 0 | -0.171144 | -3 | 0.277014 | 0.277014 | 0.3095 | 0.13502 | 0.18305 | 0.12743 | 35.08171 | 0.020230159 | 0 | Neutral |
| testneg_8051 | 0 | 0 | 4.801067 | -3 | 0.085914 | -3 | 0.5103 | 0.10923 | 0.32531 | 0.12351 | 23.77138 | 0.05 | 0 | Neutral |

| | | | | | | | | | | | | | | |
|---|---|---|---|---|---|---|---|---|---|---|---|---|---|---|
| testneg_8052 | 0.16 | -0.37 | 4.700219 | -3 | 3.63027 | -3 | 0.20011 | 0.27949 | 0.15025 | 0.4872 | 20.31414 | 0.774579354 | 99.6 | Disease_causing |
| testneg_8053 | 0.04 | -0.23 | 6.629524 | -3 | 0.66227 | -3 | 0.50687 | 0.11283 | 0.23141 | 0.12788 | 16.84579 | 0.206279762 | 1 | Neutral |
| testneg_8054 | 0 | 0 | -0.549073 | -3 | 4.80576 | -3 | 0.38016 | 0.14503 | 0.99564 | 0.25794 | 10 | 0.266116764 | 1.4 | Neutral |
| testneg_8055 | 0.08 | -0.03 | 1.514563 | -3 | 1.318968 | -3 | 0.60152 | 0.12166 | 0.18438 | 0.13272 | 10.41667 | 0.060290404 | 0 | Neutral |
| testneg_8056 | -0.04 | 1.61 | 2.25571 | -3 | 6.395815 | -3 | 0.02518 | 0.07165 | 0.52005 | 0.12788 | 23.98724 | 0.253095238 | 0 | Neutral |
| testneg_8057 | 0 | 0 | 6.373062 | -3 | 3.130587 | -3 | 0.36793 | 0.15588 | 0.05027 | 0.10981 | 7.77787 | 0.47 | 43.2 | Neutral |
| testneg_8058 | 0.14 | -1.01 | 4.688089 | -3 | 1.555382 | 1.555382 | 0.49745 | 0.1103 | 0.80134 | 0.08685 | 18.6768 | 0.091302966 | 0 | Neutral |
| testneg_8059 | 0.03 | -1.03 | 2.660158 | -3 | 0.049172 | -3 | 0.18243 | 0.11102 | 0.22034 | 0.12788 | 18.12633 | 0.011510965 | 0 | Neutral |
| testneg_8060 | 0 | -0.01 | -0.474469 | -3 | 5.537244 | -3 | 0.35422 | 0.11185 | 0.19898 | 0.09972 | 28.84083 | 0.061514286 | 0 | Neutral |
| testneg_8061 | 0.18 | 0.16 | -0.580188 | -3 | 2.651637 | -3 | 0.1675 | 0.09413 | 0.11439 | 0.21573 | 33.85257 | 0.200398293 | 0 | Neutral |
| testneg_8062 | -0.11 | 0.19 | 3.787927 | -3 | 0.028094 | -3 | 0.12839 | 0.09503 | 0.46712 | 0.93559 | 10.41667 | 0.194083333 | 0 | Neutral |
| testneg_8063 | 0 | -0.01 | 1.587876 | -3 | -0.995205 | -3 | 0.57019 | 0.38533 | 0.18673 | 0.18881 | 19.44444 | 0.11 | 0 | Neutral |
| testneg_8064 | -0.19 | 0.25 | -1.729462 | -3 | 0.082842 | -3 | 0.16155 | 0.20638 | 0.00971 | 0.0791 | 10.83333 | 0.1105 | 0 | Neutral |
| testneg_8065 | -0.06 | 0.57 | 2.921538 | -3 | 2.724086 | -3 | 0.38428 | 0.38566 | 0.71697 | 0.19162 | 10.83333 | 0.540687775 | 68.2 | Disease_causing |
| testneg_8066 | 0 | 0 | 0.339282 | -3 | 2.611714 | -3 | 0.2547 | 0.07589 | 0.99543 | 0.12788 | 9.94358 | 0.05 | 0 | Neutral |
| testneg_8067 | -0.09 | 0.67 | 0.082232 | -3 | 1.202839 | -3 | 0.08475 | 0.19889 | 0.11144 | 0.08199 | 13.02635 | 0.07 | 0 | Neutral |
| testneg_8068 | 0 | 0 | -0.961702 | -0.961702 | 1.036843 | -3 | 0.46979 | 0.1705 | 0.18102 | 0.6537 | 19.44444 | 0.105379592 | 0 | Neutral |
| testneg_8069 | 0.22 | -2.66 | 5.803249 | -3 | 0.393827 | -3 | 0.19898 | 0.08029 | 0.07516 | 0.15472 | 18.6768 | 0.2935 | 2 | Neutral |
| testneg_8070 | -0.04 | 0.23 | 4.21286 | -3 | 1.913341 | -3 | 0.07447 | 0.23492 | 0.67549 | 0.1273 | 17.47057 | 0.188406318 | 0 | Neutral |
| testneg_8071 | 0.17 | 0.38 | 1.318863 | -3 | 4.972003 | 4.768418 | 0.19898 | 0.12788 | 0.27843 | 0.54325 | 18.88889 | 0.512745986 | 57.2 | Disease_causing |
| testneg_8072 | 0 | 0 | 0.327712 | -3 | 2.76764 | -3 | 0.57517 | 0.10873 | 0.22795 | 0.12788 | 65.38226 | 0.090441729 | 0 | Neutral |
| testneg_8073 | 0 | 0 | 0.864288 | -3 | 4.474669 | 4.047733 | 0.21014 | 0.12378 | 0.12446 | 0.08908 | 25.79439 | 0.26 | 0.4 | Neutral |
| testneg_8074 | 0.21 | 0.15 | 1.704228 | -3 | 0.635234 | -3 | 0.27062 | 0.12378 | 0.61872 | 0.10479 | 22.77778 | 0.02 | 0 | Neutral |
| testneg_8075 | 0 | 0 | 0.628811 | -3 | 6.07853 | -3 | 0.07874 | 0.12788 | 0.0968 | 0.15592 | 23.98876 | 0.357333333 | 10 | Neutral |
| testneg_8076 | 0 | 0 | 1.718581 | -3 | 8.01783 | -3 | 0.7003 | 0.12788 | 0.38695 | 0.12579 | 11.09296 | 0.27 | 1.8 | Neutral |
| testneg_8077 | -0.01 | 1.44 | 1.762821 | 1.762821 | 5.50061 | -3 | 0.1375 | 0.112 | 0.09972 | 0.2187 | 21.53664 | 0.509759804 | 57.8 | Disease_causing |
| testneg_8078 | -0.02 | -0.13 | -0.203425 | -3 | 2.74177 | -3 | 0.1276 | 0.30104 | 0.04935 | 0.06599 | 11.98582 | 0.08 | 0 | Neutral |
| testneg_8079 | 0.21 | 0.15 | 1.179788 | -3 | 0.015616 | -3 | 0.0571 | 0.10747 | 0.05999 | 0.12788 | 13.02635 | 0.05 | 0 | Neutral |
| testneg_8080 | -0.02 | 0.36 | 4.55191 | -3 | 1.699863 | -3 | 0.10822 | 0.10942 | 0.06048 | 0.12788 | 39.46913 | 0.061994624 | 0 | Neutral |
| testneg_8081 | 0 | 0 | 1.019099 | -3 | 4.839828 | -3 | 0.19898 | 0.12788 | 0.05095 | 0.08353 | 11.57603 | 0.046919643 | 0 | Neutral |
| testneg_8082 | -0.18 | -0.16 | -0.566382 | -3 | 4.31851 | -3 | 0.23152 | 0.09962 | 0.04694 | 0.44682 | 25.38779 | 0.35 | 4.2 | Neutral |
| testneg_8083 | 0.22 | -2.66 | 6.248531 | -3 | 1.417622 | -3 | 0.12839 | 0.09503 | 0.50392 | 0.12788 | 10.83333 | 0.191041667 | 0.2 | Neutral |
| testneg_8084 | 0.05 | -0.81 | 5.140271 | -3 | 0.081615 | 0.081615 | 0.09542 | 0.14806 | 0.14491 | 0.10508 | 17.15045 | 0.1103125 | 0 | Neutral |
| testneg_8085 | 0.04 | -0.23 | 4.461467 | -3 | 3.233481 | -0.947523 | 0.20547 | 0.12178 | 0.10638 | 0.66554 | 13.75 | 0.467366443 | 41 | Neutral |
| testneg_8086 | -0.19 | -0.51 | 4.930424 | -3 | 4.035824 | -3 | 0.11796 | 0.12788 | 0.65141 | 0.18046 | 13.75 | 0.604884134 | 86 | Disease_causing |
| testneg_8087 | 0 | 0 | -0.001679 | -3 | -0.184879 | -3 | 0.70964 | 0.12663 | 0.19898 | 0.12788 | 28.07101 | 0.010606618 | 0 | Neutral |
| testneg_8088 | 0.14 | -1.01 | 0.207897 | -3 | 5.205515 | -3 | 0.20616 | 0.29531 | 0.11613 | 0.1109 | 19.44444 | 0.213333333 | 0.2 | Neutral |
| testneg_8089 | 0 | 0 | 0.707491 | -3 | 6.959549 | -3 | 0.43762 | 0.12788 | 0.13967 | 0.15696 | 22.02418 | 0.426130013 | 28.8 | Neutral |
| testneg_8090 | 0 | 0 | -0.032453 | -0.032453 | 3.847949 | -3 | 0.31664 | 0.14181 | 0.41655 | 0.49485 | 11.25 | 0.297410808 | 2.6 | Neutral |
| testneg_8091 | -0.07 | -0.38 | 3.479092 | -3 | 6.384552 | -3 | 0.22113 | 0.10673 | 0.05873 | 0.09792 | 23.46591 | 0.26 | 1.2 | Neutral |
| testneg_8092 | -0.07 | -0.38 | 2.184344 | -3 | 0.334215 | -3 | 0.05472 | 0.12788 | 0.33314 | 0.47999 | 42.35819 | 0.17 | 0 | Neutral |
| testneg_8093 | 0 | 0 | 0.475124 | -3 | 1.064842 | -3 | 0.17677 | 0.67006 | 0.19898 | 0.12788 | 20.37554 | 0.05 | 0 | Neutral |
| testneg_8094 | 0 | 0 | 2.259254 | -3 | 8.085184 | -3 | 0.56285 | 0.18619 | 0.15767 | 0.12788 | 13.2927 | 0.431555556 | 30.6 | Neutral |
| testneg_8095 | 0.004 | -0.4576364 | 7.731489 | -3 | 0.297516 | 0.297516 | 0.03378 | 0.12788 | 0.19294 | 0.10464 | 20.98976 | 0.336875 | 7.8 | Neutral |
| testneg_8096 | 0 | 0 | -0.674729 | -3 | 1.006031 | -3 | 0.24765 | 0.32858 | 0.78537 | 0.33817 | 11.875 | 0.092212065 | 0 | Neutral |
| testneg_8097 | -0.06 | 0.57 | 1.183447 | -3 | 5.0589 | -3 | 0.0146 | 0.03045 | 0.09658 | 0.11286 | 36.05565 | 0.13 | 0 | Neutral |
| testneg_8098 | -0.08 | -0.03 | -0.949463 | -3 | 1.549087 | -3 | 0.09018 | 0.16175 | 0.3025 | 0.2499 | 10.41667 | 0.150164706 | 0 | Neutral |
| testneg_8099 | 0 | 0 | 1.012321 | -3 | 1.866705 | -3 | 0.21805 | 0.12777 | 0.14938 | 0.12747 | 13.08587 | 0 | 0 | Neutral |
| testneg_8100 | 0 | 0 | 0.298529 | -3 | -0.782194 | -3 | 0.26264 | 0.13812 | 0.20011 | 0.27949 | 10.41667 | 0.033782615 | 0 | Neutral |
| testneg_8101 | 0 | 0 | 0.82819 | 0.82819 | 0.116245 | -3 | 0.19898 | 0.12788 | 0.8238 | 0.38836 | 10.41667 | 0.091750575 | 0 | Neutral |
| testneg_8102 | 0.14 | -1.01 | 2.843639 | 1.643871 | 0.896683 | -3 | 0.19898 | 0.12788 | 0.16002 | 0.10481 | 23.32524 | 0.08 | 0 | Neutral |
| testneg_8103 | -0.06 | -0.18 | 0.815968 | -3 | 2.988243 | -3 | 0.11587 | 0.12001 | 0.54015 | 0.13176 | 11.25 | 0.127339125 | 0 | Neutral |
| testneg_8104 | 0.03 | -1.03 | 4.768167 | -3 | 7.197642 | -3 | 0.91587 | 0.10818 | 0.15708 | 0.12788 | 13.01219 | 0.49 | 51.2 | Disease_causing |
| testneg_8105 | 0 | 0 | -0.32247 | -3 | 1.361739 | -3 | 0.35092 | 0.10596 | 0.25556 | 0.09342 | 18.6786 | 8.82E-05 | 0 | Neutral |
| testneg_8106 | 0 | 0 | -0.433937 | -0.433937 | 6.349976 | -3 | 0.84572 | 0.78546 | 0.12403 | 0.12788 | 10.83333 | 0.201721491 | 0 | Neutral |
| testneg_8107 | -0.2 | 1.26 | 6.127935 | -3 | 4.542847 | -3 | 0.13918 | 0.12788 | 0.3405 | 0.1816 | 14.90842 | 0.711458333 | 99.6 | Disease_causing |
| testneg_8108 | 0.07 | -0.31 | 1.60347 | -3 | 5.594118 | -3 | 0.11261 | 0.14664 | 0.17808 | 0.12788 | 17.1972 | 0.202613636 | 0 | Neutral |
| testneg_8109 | 0 | 0 | 1.050504 | -3 | 0.734935 | -3 | 0.46792 | 0.11262 | 0.2877 | 0.18291 | 17.77778 | 0.02 | 0 | Neutral |
| testneg_8110 | -0.03 | -0.1 | 1.820578 | -3 | 1.101447 | -3 | 0.31507 | 0.61057 | 0.2306 | 0.0952 | 10.83333 | 0.050801282 | 0 | Neutral |
| testneg_8111 | 0.02 | 0.45 | 3.429131 | -3 | 3.974134 | -3 | 0.03499 | 0.12971 | 0.16468 | 0.11275 | 13.08587 | 0.191547619 | 0 | Neutral |
| testneg_8112 | 0 | -0.01 | 0.505843 | -3 | 1.342384 | -3 | 0.64188 | 0.09494 | 0.36813 | 0.09414 | 10.83333 | 0.010121969 | 0 | Neutral |
| testneg_8113 | -0.03 | -0.1 | 6.734048 | -3 | -1.970817 | -3 | 0.19898 | 0.48545 | 0.2058 | 0.28174 | 7.08233 | 0.711833333 | 99 | Disease_causing |
| testneg_8114 | -0.07 | -0.38 | 1.181002 | -3 | -0.036235 | -3 | 0.19898 | 0.58778 | 0.21118 | 0.10074 | 16.8018 | 0.050801282 | 0 | Neutral |
| testneg_8115 | 0.07 | 0.38 | 3.047706 | -3 | 1.268416 | 1.268416 | 0.0695 | 0.12788 | 0.43977 | 0.17284 | 24.68926 | 0.210142157 | 0 | Neutral |
| testneg_8116 | 0 | 0 | 0.29615 | -3 | 0.807877 | -3 | 0.17325 | 0.17964 | 0.10751 | 0.08512 | 23.04078 | 0.020833333 | 0 | Neutral |
| testneg_8117 | 0 | 0 | 0.307719 | -3 | -1.638874 | -3 | 0.16091 | 0.11571 | 0.22078 | 0.12788 | 26.07729 | 0.000129958 | 0 | Neutral |
| testneg_8118 | 0 | 0 | 0.114574 | -3 | 0.416167 | 0.416167 | 0.08922 | 0.10539 | 0.1224 | 0.11899 | 19.48718 | 0.01 | 0 | Neutral |
| testneg_8119 | 0.02 | -0.28 | 0.69881 | -3 | 0.655763 | 0.655763 | 0.20692 | 0.70649 | 0.09087 | 0.12788 | 10 | 0.060086207 | 0 | Neutral |
| testneg_8120 | 0.16 | -0.37 | 5.827202 | -3 | 0.775551 | -3 | 0.13275 | 0.12788 | 0.0903 | 0.09372 | 12.8741 | 0.12 | 0 | Neutral |
| testneg_8121 | -0.03 | -0.1 | 0.815782 | -3 | 3.467641 | -3 | 0.45029 | 0.12788 | 0.13779 | 0.0945 | 20.91877 | 0.058465909 | 0 | Neutral |
| testneg_8122 | 0.06 | 0.18 | 1.825183 | -3 | -0.76618 | -3 | 0.204 | 0.15424 | 0.1503 | 0.12788 | 12.29167 | 0.02 | 0 | Neutral |
| testneg_8123 | 0 | 0 | 0.584441 | -3 | -2.345951 | -3 | 0.97678 | 0.09885 | 0.13199 | 0.14263 | 16.07106 | 0.050285714 | 0 | Neutral |
| testneg_8124 | 0.08 | -1.2 | 2.652042 | -3 | 5.961755 | -3 | 0.83798 | 0.14433 | 0.27783 | 0.62384 | 14.14796 | 0.574737964 | 83.2 | Disease_causing |
| testneg_8125 | 0 | 0 | 2.153274 | -3 | -0.244741 | -3 | 0.835 | 0.13588 | 0.09645 | 0.13239 | 16.98213 | 0.07020979 | 0 | Neutral |
| testneg_8126 | 0.05 | -0.81 | 2.541406 | -3 | 1.211461 | -3 | 0.18283 | 0.11376 | 0.38552 | 0.14172 | 10 | 0.04225 | 0 | Neutral |
| testneg_8127 | -0.06 | -0.18 | 1.350332 | -3 | 0.082158 | -3 | 0.55179 | 0.29199 | 0.07394 | 0.09408 | 8.99708 | 0.02 | 0 | Neutral |
| testneg_8128 | 0 | 0 | -0.633568 | -3 | 0.839368 | 0.839368 | 0.23027 | 0.37171 | 0.18785 | 0.14723 | 16.2309 | 0.122583333 | 0 | Neutral |
| testneg_8129 | 0 | 0 | 0.565761 | -3 | 1.134092 | -3 | 0.10383 | 0.08853 | 0.17808 | 0.12788 | 14.45783 | 0.000247606 | 0 | Neutral |
| testneg_8130 | 0 | 0 | 1.314085 | -3 | 1.989667 | -3 | 0.10236 | 0.09622 | 0.06517 | 0.15107 | 39.97067 | 0.070407754 | 0 | Neutral |
| testneg_8131 | 0 | -0.01 | 1.028355 | -3 | -0.002309 | -3 | 0.87835 | 0.13522 | 0.06729 | 0.17051 | 12.80035 | 0.07 | 0 | Neutral |
| testneg_8132 | 0.18 | -1.05 | 3.036954 | -3 | 1.638182 | -3 | 0.36541 | 0.27749 | 0.62203 | 0.12788 | 23.96199 | 0.142161836 | 0 | Neutral |
| testneg_8133 | 0 | -0.01 | 1.609618 | -3 | 1.678033 | -3 | 0.67038 | 0.1794 | 0.33957 | 0.12788 | 11.25 | 0.010315136 | 0 | Neutral |
| testneg_8134 | 0 | -0.32 | 2.337712 | -3 | -1.32005 | -3 | 0.14273 | 0.1162 | 0.16345 | 0.11716 | 20.31414 | 0.01 | 0 | Neutral |
| testneg_8135 | -0.01 | 0.57 | 0.120012 | -3 | 4.450954 | -3 | 0.20491 | 0.22087 | 0.19898 | 0.12788 | 12.25575 | 0.191501891 | 0 | Neutral |
| testneg_8136 | 0 | 0 | 0.611732 | -3 | 1.772012 | -3 | 0.79692 | 0.51979 | 0.26731 | 0.16415 | 10.83333 | 0.08 | 0 | Neutral |
| testneg_8137 | 0 | 0 | 0.458753 | -3 | 1.119912 | -3 | 0.7413 | 0.18974 | 0.09724 | 0.09172 | 16.22079 | 0.01 | 0 | Neutral |
| testneg_8138 | -0.16 | -0.08 | 6.205256 | -3 | 0.945821 | -3 | 0.12377 | 0.12788 | 0.19898 | 0.12788 | 45.34722 | 0.19 | 0.6 | Neutral |
| testneg_8139 | 0 | -0.32 | 6.40665 | -3 | 5.508045 | -3 | 0.29828 | 0.12788 | 0.26884 | 0.64999 | 19.44444 | 0.77 | 99.6 | Disease_causing |
| testneg_8140 | 0 | 0 | -0.4932 | -3 | 9.146604 | -3 | 0.20379 | 0.43705 | 0.44923 | 0.12788 | 18.21138 | 0.45 | 35 | Neutral |
| testneg_8141 | 0 | 0 | 0.917232 | -3 | 2.140243 | -3 | 0.25556 | 0.09342 | 0.03558 | 0.05202 | 33.78439 | 0.02 | 0 | Neutral |
| testneg_8142 | -0.07 | -0.38 | 5.544856 | -3 | 4.617328 | -3 | 0.51675 | 0.11055 | 0.19898 | 0.12788 | 27.00364 | 0.349984432 | 10.4 | Neutral |
| testneg_8143 | -0.04 | 0.23 | 0.2182 | -3 | 6.721356 | -3 | 0.19898 | 0.12788 | 0.31773 | 0.10816 | 50.54802 | 0.19 | 0 | Neutral |
| testneg_8144 | 0.05 | 0.29 | 3.391965 | -3 | 0.450255 | -3 | 0.19898 | 0.12788 | 0.1601 | 0.12546 | 18.6768 | 0.02 | 0 | Neutral |
| testneg_8145 | 0.03 | -1.03 | 2.514378 | -3 | 3.92229 | -3 | 0.3793 | 0.24105 | 0.09184 | 0.38254 | 10.83333 | 0.602667132 | 90.4 | Disease_causing |
| testneg_8146 | -0.02 | -0.45 | 5.135114 | -3 | 0.565031 | -3 | 0.28629 | 0.77234 | 0.19898 | 0.12788 | 10.80591 | 0.228759398 | 0.6 | Neutral |
| testneg_8147 | 0.15 | -0.28 | -0.964308 | -3 | 3.095581 | 3.095581 | 0.32989 | 0.13525 | 0.41269 | 0.19135 | 21.49912 | 0.360053366 | 8.8 | Neutral |
| testneg_8148 | 0 | 0 | 2.100416 | -3 | -2.609079 | -3 | 0.22255 | 0.10057 | 0.19898 | 0.09533 | 30.38941 | 0.01 | 0 | Neutral |
| testneg_8149 | 0.05 | -0.81 | 3.204485 | -3 | 1.499766 | -3 | 0.16728 | 0.11868 | 0.08836 | 0.0911 | 19.45901 | 0.03 | 0 | Neutral |
| testneg_8150 | 0 | 0 | 1.269835 | -3 | 3.449691 | -3 | 0.94662 | 0.11262 | 0.08531 | 0.12788 | 10.41667 | 0.04375 | 0 | Neutral |
| testneg_8151 | -0.04 | -2.01 | 0.424648 | -3 | 1.893207 | -3 | 0.3769 | 0.12788 | 0.35908 | 0.18099 | 18.88889 | 0.100828187 | 0 | Neutral |
| testneg_8152 | 0.18 | 0.16 | -0.110087 | -3 | 0.30563 | -3 | 0.82126 | 0.17937 | 0.1417 | 0.1398 | 10 | 0.137496408 | 0 | Neutral |
| testneg_8153 | -0.01 | -0.79 | -0.6503 | -3 | -2.664317 | -3 | 0.13994 | 0.11181 | 0.16321 | 0.09621 | 18.33333 | 0.010284389 | 0 | Neutral |
| testneg_8154 | 0.01 | 0.8 | 0.78234 | -3 | 0.002159 | -3 | 0.10176 | 0.12788 | 0.10231 | 0.1 | 10.22898 | 0.02 | 0 | Neutral |
| testneg_8155 | 0 | 0 | 0.332253 | -3 | 1.077291 | -3 | 0.23461 | 0.11809 | 0.15612 | 0.11465 | 15.07524 | 0 | 0 | Neutral |
| testneg_8156 | 0.04 | -0.23 | 6.924944 | -3 | -0.030885 | -3 | 0.19979 | 0.09328 | 0.18578 | 0.27854 | 16.8018 | 0.4 | 21 | Neutral |
| testneg_8157 | 0.19 | -0.27 | 6.416509 | -3 | -0.437663 | -3 | 0.08814 | 0.42357 | 0.34946 | 0.19092 | 10.41667 | 0.584792398 | 84.2 | Disease_causing |
| testneg_8158 | -0.02 | 0.63 | 2.293151 | -3 | 3.544108 | -3 | 0.47997 | 0.10506 | 0.13974 | 0.10225 | 16.8018 | 0.091883513 | 0 | Neutral |
| testneg_8159 | -0.01 | -0.8 | 3.501031 | -3 | 0.231015 | -3 | 0.25421 | 0.16235 | 0.19898 | 0.12788 | 40.503 | 0.062637029 | 0 | Neutral |
| testneg_8160 | 0.09 | -0.3 | -0.408308 | -3 | -0.006272 | -0.006272 | 0.28165 | 0.15912 | 0.07335 | 0.12788 | 13.51351 | 0.04 | 0 | Neutral |
| testneg_8161 | -0.004965 | 0.03867133 | 1.375958 | -3 | 12.915528 | -3 | 0.0538 | 0.23912 | 0.15856 | 0.10994 | 26.96533 | 0.53 | 69.2 | Disease_causing |
| testneg_8162 | -0.03 | -1.22 | 4.348114 | -3 | 0.512138 | -3 | 0.12079 | 0.62023 | 0.89661 | 0.17382 | 13.07136 | 0.377819767 | 11.8 | Neutral |
| testneg_8163 | 0 | 0 | 0.302733 | -3 | 5.361142 | 0.297064 | 0.19898 | 0.12788 | 0.2029 | 0.12788 | 29.14274 | 0.061666667 | 0 | Neutral |
| testneg_8164 | 0.18 | 0.16 | 0.416167 | -3 | 3.936847 | -3 | 0.1224 | 0.11899 | 0.12036 | 0.0873 | 33.51926 | 0.09 | 0 | Neutral |
| testneg_8165 | 0.08 | 0.03 | -1.290562 | -3 | -2.237532 | -3 | 0.19898 | 0.12895 | 0.42836 | 0.12788 | 21.51158 | 0.03 | 0 | Neutral |
| testneg_8166 | -0.08 | -0.03 | 3.093558 | -3 | 1.319914 | 1.319914 | 0.28499 | 0.10637 | 0.08896 | 0.12788 | 25.2849 | 0.04 | 0 | Neutral |
| testneg_8167 | 0.21 | 0.15 | 6.418378 | -3 | 0.22446 | 0.22446 | 0.38552 | 0.14172 | 0.32144 | 0.11774 | 12.29167 | 0.2 | 0.8 | Neutral |
| testneg_8168 | 0 | 0 | 1.550971 | -3 | 1.810934 | -3 | 0.73311 | 0.10866 | 0.5184 | 0.11889 | 5.69444 | 0 | 0 | Neutral |
| testneg_8169 | 0 | 0 | 0.555194 | -3 | 2.935592 | -3 | 0.91921 | 0.32331 | 0.1909 | 0.14279 | 18.36272 | 0.183573627 | 0 | Neutral |
| testneg_8170 | 0.03 | -1.03 | 0.896883 | -3 | 2.393948 | 2.393948 | 0.02124 | 0.05186 | 0.70819 | 0.5975 | 20.41298 | 0.25 | 0.2 | Neutral |
| testneg_8171 | 0.22 | -2.66 | 4.259451 | -3 | 0.35929 | -3 | 0.05027 | 0.10981 | 0.19898 | 0.12788 | 25.6274 | 0.11 | 0 | Neutral |
| testneg_8172 | 0.18 | -1.05 | 5.955324 | -3 | -0.592626 | -3 | 0.24287 | 0.11626 | 0.143 | 0.11236 | 12.28162 | 0.116035714 | 0 | Neutral |
| testneg_8173 | 0.21 | 0.15 | 3.267547 | -3 | 0.581582 | -3 | 0.06027 | 0.11574 | 0.06636 | 0.1079 | 4.72222 | 0.07 | 0 | Neutral |

| | | | | | | | | | | | | | | |
|---|---|---|---|---|---|---|---|---|---|---|---|---|---|---|
| testneg_8174 | 0.06 | -0.57 | 6.407239 | -3 | -0.748995 | -3 | 0.81838 | 0.09725 | 0.19898 | 0.12788 | 11.25 | 0.2 | 0.6 | Neutral |
| testneg_8175 | -0.03 | -0.1 | 2.935592 | -3 | 0.39745 | 0.39745 | 0.1909 | 0.14279 | 0.11404 | 0.12788 | 9.80316 | 0.049791667 | 0 | Neutral |
| testneg_8176 | 0 | 0 | -0.00629 | -3 | 6.431621 | 6.431621 | 0.1324 | 0.09503 | 0.26676 | 0.76568 | 16.32654 | 0.5 | 55.8 | Disease_causing |
| testneg_8177 | 0 | 0 | -1.056572 | -3 | 5.812276 | -3 | 0.1109 | 0.17295 | 0.19898 | 0.12788 | 38.03819 | 0.201428571 | 0.2 | Neutral |
| testneg_8178 | 0.16 | 0.08 | -0.169987 | -3 | 1.170456 | -3 | 0.99445 | 0.10396 | 0.339 | 0.13169 | 10.83333 | 0.090177778 | 0 | Neutral |
| testneg_8179 | 0 | 0 | 1.609501 | -3 | -0.112406 | -3 | 0.16404 | 0.09378 | 0.94807 | 0.14977 | 7.77778 | 0.010353594 | 0 | Neutral |
| testneg_8180 | 0 | 0 | -1.230837 | -1.230837 | 0.500506 | -3 | 0.14821 | 0.11262 | 0.68233 | 0.58998 | 10 | 0.03 | 0 | Neutral |
| testneg_8181 | -0.07 | 0.31 | -0.564575 | -3 | 1.3012 | -3 | 0.68029 | 0.12788 | 0.10893 | 0.12072 | 12.39396 | 0.020833333 | 0 | Neutral |
| testneg_8182 | 0.06 | 0.18 | -0.134916 | -0.134916 | 2.009531 | -3 | 0.16475 | 0.26069 | 0.47322 | 0.11118 | 15.54911 | 0.04 | 0 | Neutral |
| testneg_8183 | -0.06 | -0.18 | -0.439044 | -3 | 2.207668 | -3 | 0.3797 | 0.1257 | 0.26618 | 0.2057 | 10.83333 | 0.063796592 | 0 | Neutral |
| testneg_8184 | -0.02 | 0.63 | 4.4929 | -3 | 1.076529 | -3 | 0.1205 | 0.23508 | 0.06711 | 0.09765 | 10.38862 | 0.13 | 0 | Neutral |
| testneg_8185 | 0 | 0 | 0.275842 | -3 | 0.438158 | -3 | 0.14462 | 0.11239 | 0.21241 | 0.08684 | 10.24716 | 0 | 0 | Neutral |
| testneg_8186 | -0.07 | -0.33 | -2.373259 | -3 | 3.54624 | -3 | 0.04525 | 0.12788 | 0.05476 | 0.12788 | 24.49409 | 0.108333333 | 0 | Neutral |
| testneg_8187 | 0.21 | 0.15 | 0.856696 | -3 | -4.511894 | -3 | 0.07237 | 0.12788 | 0.10011 | 0.08014 | 18.97884 | 0.04 | 0 | Neutral |
| testneg_8188 | 0 | 0 | -0.002747 | -3 | 3.218787 | -3 | 0.07394 | 0.09408 | 0.3405 | 0.1816 | 15.25808 | 0.128204991 | 0 | Neutral |
| testneg_8189 | -0.01 | -0.79 | 1.351518 | -3 | 0.223803 | -3 | 0.58984 | 0.15588 | 0.10026 | 0.12788 | 17.88889 | 0.03 | 0 | Neutral |
| testneg_8190 | -0.04 | 0.23 | -0.274997 | -3 | -0.199859 | -3 | 0.37501 | 0.0997 | 0.11006 | 0.14352 | 29.84187 | 0.080600358 | 0 | Neutral |
| testneg_8191 | 0.17 | 0.38 | 3.117033 | -3 | 1.641419 | -3 | 0.13203 | 0.12092 | 0.11891 | 0.11186 | 17.28802 | 0.033095238 | 0 | Neutral |
| testneg_8192 | 0 | 0 | 1.335711 | -3 | 3.393414 | -0.641115 | 0.2741 | 0.10478 | 0.62381 | 0.12788 | 13.14653 | 0.020215909 | 0 | Neutral |
| testneg_8193 | -0.02 | 0.63 | 1.73624 | -3 | 2.541405 | -3 | 0.74488 | 0.12788 | 0.03235 | 0.0948 | 15.28736 | 0.051138889 | 0 | Neutral |
| testneg_8194 | -0.07 | 0.31 | -0.564575 | -3 | -0.002309 | -3 | 0.68029 | 0.12788 | 0.06729 | 0.17051 | 18.33333 | 0.101483516 | 0 | Neutral |
| testneg_8195 | -0.15 | -0.04 | 2.470276 | -3 | 5.893689 | -3 | 0.06841 | 0.12788 | 0.21383 | 0.11171 | 13.35742 | 0.25 | 0 | Neutral |
| testneg_8196 | 0.02 | -0.36 | 1.555382 | 1.555382 | 1.986323 | -3 | 0.80134 | 0.08685 | 0.19898 | 0.12788 | 10 | 0.05 | 0 | Neutral |
| testneg_8197 | 0 | 0 | 0.97071 | -3 | 7.144915 | -3 | 0.17855 | 0.11262 | 0.07415 | 0.12788 | 16.8018 | 0.23075 | 1.4 | Neutral |
| testneg_8198 | 0.06 | -0.57 | 0.297516 | 0.297516 | 0.478518 | -3 | 0.19294 | 0.10464 | 0.89959 | 0.55078 | 12.40248 | 0.08 | 0 | Neutral |
| testneg_8199 | 0.09 | -0.3 | 1.5447 | -3 | 2.055296 | -3 | 0.08932 | 0.32882 | 0.15092 | 0.09074 | 12.32809 | 0.08 | 0 | Neutral |
| testneg_8200 | 0.06 | -1.29 | 2.692696 | -3 | 5.414675 | -3 | 0.0903 | 0.09372 | 0.10385 | 0.09775 | 20.83333 | 0.137904423 | 0 | Neutral |
| testneg_8201 | -0.03 | -0.1 | 2.83954 | -3 | 1.29341 | -3 | 0.09432 | 0.15749 | 0.11773 | 0.11669 | 32.20126 | 0.07225 | 0 | Neutral |
| testneg_8202 | 0 | -0.73 | 1.128181 | -3 | 1.489203 | -3 | 0.92235 | 0.62242 | 0.26928 | 0.12774 | 8.73053 | 0.05 | 0 | Neutral |
| testneg_8203 | 0.05 | -0.81 | 5.789217 | -3 | 0.892974 | 0.892974 | 0.26668 | 0.11598 | 0.06986 | 0.12788 | 17.48419 | 0.12 | 0 | Neutral |
| testneg_8204 | 0.01 | -0.57 | 6.366826 | -3 | 1.029427 | -3 | 0.06781 | 0.09009 | 0.80314 | 0.42344 | 24.61676 | 0.320083333 | 5.6 | Neutral |
| testneg_8205 | -0.07 | -0.33 | -1.001041 | -3 | 2.440516 | -3 | 0.08136 | 0.10647 | 0.92776 | 0.09677 | 20.86969 | 0.05 | 0 | Neutral |
| testneg_8206 | -0.08 | -0.03 | 5.746826 | -3 | 0.36991 | -3 | 0.36541 | 0.27749 | 0.31621 | 0.12788 | 16.20187 | 0.27 | 1.2 | Neutral |
| testneg_8207 | 0 | 0.32 | -0.420859 | -3 | 0.71504 | -3 | 0.0597 | 0.18092 | 0.21894 | 0.09041 | 26.40523 | 0.058079205 | 0 | Neutral |
| testneg_8208 | 0.07 | 0.38 | 0.200034 | -3 | 2.25618 | -3 | 0.84318 | 0.11313 | 0.19898 | 0.09316 | 17.21847 | 0.021277174 | 0 | Neutral |
| testneg_8209 | -0.07 | -0.38 | 0.339731 | -3 | 4.488254 | -3 | 0.26192 | 0.15452 | 0.19898 | 0.08855 | 18.33333 | 0.130925697 | 0 | Neutral |
| testneg_8210 | 0 | 0 | -0.211085 | -0.211085 | 1.480614 | -3 | 0.04543 | 0.28103 | 0.11849 | 0.20101 | 10.41667 | 0.190655932 | 0 | Neutral |
| testneg_8211 | 0 | 0 | 0.51635 | -3 | 1.373367 | 1.373367 | 0.19898 | 0.82835 | 0.34314 | 0.18668 | 10.83333 | 0.174672414 | 0 | Neutral |
| testneg_8212 | 0.14 | -1.01 | 0.849231 | -3 | 1.855162 | -3 | 0.08629 | 0.12788 | 0.33294 | 0.16155 | 10 | 0.085545455 | 0 | Neutral |
| testneg_8213 | -0.17 | -0.38 | 3.270943 | -3 | 5.289856 | -3 | 0.44526 | 0.09404 | 0.06955 | 0.1748 | 31.10669 | 0.499479798 | 52.8 | Disease_causing |
| testneg_8214 | -0.02 | 0.68 | 1.797759 | -3 | 0.654388 | -3 | 0.39553 | 0.11991 | 0.85887 | 0.1489 | 20.28986 | 0.050039063 | 0 | Neutral |
| testneg_8215 | 0.01 | 0.79 | -0.188304 | -3 | 0.570579 | -3 | 0.16418 | 0.12788 | 0.56204 | 0.10762 | 10.15504 | 0.01 | 0 | Neutral |
| testneg_8216 | 0.21 | 0.15 | 1.098308 | -3 | 3.47892 | -3 | 0.09346 | 0.12788 | 0.18423 | 0.10808 | 30.02093 | 0.075631845 | 0 | Neutral |
| testneg_8217 | 0.04 | -0.23 | 3.570813 | -3 | 2.864999 | -3 | 0.68054 | 0.09208 | 0.82269 | 0.15732 | 28.82782 | 0.240100733 | 0.4 | Neutral |
| testneg_8218 | -0.08 | -0.03 | 1.580417 | -3 | 3.051125 | -3 | 0.29417 | 0.10965 | 0.0709 | 0.07843 | 13.51351 | 0.04 | 0 | Neutral |
| testneg_8219 | 0 | 0 | 0.08585 | -3 | 0.178228 | -3 | 0.5471 | 0.11262 | 0.18964 | 0.34036 | 19.09347 | 0.05324446 | 0 | Neutral |
| testneg_8220 | 0 | 0 | -0.041068 | -3 | -1.243237 | -3 | 0.19898 | 0.07875 | 0.3823 | 0.37294 | 28.43434 | 0.089228395 | 0 | Neutral |
| testneg_8221 | 0.02 | -0.63 | -0.730881 | -3 | 6.194646 | -3 | 0.26476 | 0.08217 | 0.19898 | 0.09584 | 28.04291 | 0.09 | 0 | Neutral |
| testneg_8222 | -0.07 | -0.38 | 3.920732 | -3 | 0.279354 | -3 | 0.0811 | 0.09449 | 0.81212 | 0.14915 | 18.88889 | 0.131380475 | 0 | Neutral |
| testneg_8223 | 0 | 0 | 1.774559 | -3 | 1.160968 | -3 | 0.08628 | 0.11498 | 0.98565 | 0.23629 | 18.94626 | 0.040285646 | 0 | Neutral |
| testneg_8224 | 0.01 | 0.8 | 2.217361 | -3 | 5.458389 | -3 | 0.11829 | 0.12788 | 0.15125 | 0.08976 | 20.48701 | 0.138857573 | 0 | Neutral |
| testneg_8225 | 0.19 | -0.25 | 1.267038 | -3 | 5.752104 | 5.752104 | 0.19898 | 0.12788 | 0.17322 | 0.10311 | 16.43865 | 0.392142857 | 13.4 | Neutral |
| testneg_8226 | 0 | 0 | 0.101787 | -3 | 0.199359 | -3 | 0.13782 | 0.13794 | 0.07626 | 0.12788 | 23.68839 | 0.02 | 0 | Neutral |
| testneg_8227 | 0 | 0 | 0.701413 | -3 | 1.291344 | -3 | 0.1214 | 0.0899 | 0.08876 | 0.09037 | 12.31894 | 0.01 | 0 | Neutral |
| testneg_8228 | 0.04 | -0.23 | 7.332994 | -3 | 6.06046 | -3 | 0.1434 | 0.09092 | 0.00487 | 0.0143 | 32.91748 | 0.42 | 24.2 | Neutral |
| testneg_8229 | 0.11 | -0.19 | 4.882145 | -3 | 12.873188 | -3 | 0.16068 | 0.12788 | 0.16158 | 0.3836 | 10.41667 | 0.790589962 | 100 | Disease_causing |
| testneg_8230 | 0 | 0 | 1.205908 | -3 | 5.071893 | -3 | 0.32333 | 0.23356 | 0.28817 | 0.10531 | 18.76225 | 0.100514347 | 0 | Neutral |
| testneg_8231 | 0 | 0 | 1.861867 | -3 | 3.908744 | -3 | 0.32963 | 0.1035 | 0.47379 | 0.18145 | 18.87038 | 0.242876057 | 0 | Neutral |
| testneg_8232 | 0.03 | 1.22 | 0.647985 | -3 | 7.114639 | -3 | 0.21728 | 0.1132 | 0.8433 | 0.13128 | 28.55956 | 0.29 | 2.2 | Neutral |
| testneg_8233 | 0.21 | 0.15 | 1.631249 | -3 | 4.149716 | -3 | 0.47166 | 0.13548 | 0.19322 | 0.12788 | 38.07491 | 0.141428571 | 0 | Neutral |
| testneg_8234 | 0.02 | 0.13 | 1.648546 | -3 | 0.923434 | -3 | 0.16734 | 0.15176 | 0.16436 | 0.19109 | 26.9055 | 0.096563782 | 0 | Neutral |
| testneg_8235 | 0.02 | -0.63 | 0.686794 | 0.665096 | 2.251984 | -3 | 0.15132 | 0.11724 | 0.15685 | 0.17839 | 10.83333 | 0.145462413 | 0 | Neutral |
| testneg_8236 | 0.01 | -1.44 | 2.093021 | -3 | -0.360768 | -0.360768 | 0.54724 | 0.12788 | 0.47147 | 0.09503 | 19.66924 | 0.04 | 0 | Neutral |
| testneg_8237 | 0 | 0 | 0.123542 | -3 | 1.429927 | -3 | 0.99915 | 0.96333 | 0.40612 | 0.45135 | 4.44444 | 0.16 | 0 | Neutral |
| testneg_8238 | 0 | 0 | -0.553742 | -3 | 5.143935 | -3 | 0.1424 | 0.12788 | 0.12511 | 0.11897 | 29.01351 | 0.06 | 0 | Neutral |
| testneg_8239 | -0.14 | 1.01 | -1.162831 | -1.162831 | -0.052442 | -3 | 0.13398 | 0.12788 | 0.19898 | 0.09201 | 10.28409 | 0.06 | 0 | Neutral |
| testneg_8240 | -0.23 | 1.04 | 5.857535 | -3 | 4.047618 | -3 | 0.17532 | 0.36561 | 0.59302 | 0.10897 | 14.21141 | 0.639878049 | 94 | Disease_causing |
| testneg_8241 | 0.01 | 0.8 | 1.447055 | -3 | 2.565617 | -3 | 0.60959 | 0.76078 | 0.24011 | 0.28559 | 4.72222 | 0.251452857 | 0.6 | Neutral |
| testneg_8242 | -0.08 | -0.03 | 0.950149 | -3 | 4.952298 | -3 | 0.31318 | 0.11262 | 0.10042 | 0.09296 | 18.00078 | 0.085387311 | 0 | Neutral |
| testneg_8243 | 0 | -0.01 | 1.315543 | -3 | 2.767501 | -3 | 0.28358 | 0.38765 | 0.06781 | 0.09009 | 16.42913 | 0.05 | 0 | Neutral |
| testneg_8244 | -0.01 | 1.44 | 0.734765 | -3 | 3.429969 | 1.0263 | 0.07204 | 0.07708 | 0.08183 | 0.11603 | 9.95238 | 0.16 | 0 | Neutral |
| testneg_8245 | -0.21 | -0.15 | -2.734447 | -3 | 1.14709 | -3 | 0.48023 | 0.12788 | 0.42544 | 0.11809 | 27.32716 | 0.060144673 | 0 | Neutral |
| testneg_8246 | 0.04 | -0.23 | 1.583169 | -3 | 2.733966 | -3 | 0.31626 | 0.12936 | 0.34205 | 0.11195 | 10.5 | 0.020709124 | 0 | Neutral |
| testneg_8247 | 0 | 0 | 0.103447 | 0.103447 | -0.024667 | -0.024667 | 0.15813 | 0.61197 | 0.59599 | 0.12477 | 16.04167 | 0.06 | 0 | Neutral |
| testneg_8248 | 0.09 | -0.25 | -0.383584 | -3 | 3.021368 | -3 | 0.25236 | 0.11809 | 0.31025 | 0.09491 | 9.377 | 0.049984869 | 0 | Neutral |
| testneg_8249 | 0 | 0 | 1.446081 | -3 | 0.931654 | -3 | 0.48585 | 0.66989 | 0.2074 | 0.09696 | 19.09347 | 0.03865377 | 0 | Neutral |
| testneg_8250 | -0.03 | -0.1 | 0.224925 | -3 | 1.912827 | -3 | 0.03676 | 0.12788 | 0.1203 | 0.13872 | 10.83333 | 0.13 | 0 | Neutral |
| testneg_8251 | 0 | 0 | -0.566651 | -3 | 5.438117 | -3 | 0.38016 | 0.14503 | 0.12463 | 0.12788 | 17.68589 | 0.111686508 | 0 | Neutral |
| testneg_8252 | 0 | 0 | -0.216528 | -3 | 1.445184 | 1.445184 | 0.40612 | 0.45135 | 0.33469 | 0.17432 | 9.46862 | 0.096406122 | 0 | Neutral |
| testneg_8253 | 0.09 | -0.3 | 1.743348 | -3 | 0.612491 | -3 | 0.69563 | 0.12788 | 0.52768 | 0.12788 | 18.1536 | 0.010241279 | 0 | Neutral |
| testneg_8254 | -0.0430909 | 0.216 | 9.272843 | -3 | 0.976803 | -3 | 0.24544 | 0.0894 | 0.50687 | 0.11283 | 20.36321 | 0.37 | 8.8 | Neutral |
| testneg_8255 | -0.03 | -0.1 | 4.716547 | -3 | -0.727719 | -3 | 0.57239 | 0.16622 | 0.14248 | 0.12788 | 42.27361 | 0.093146084 | 0 | Neutral |
| testneg_8256 | 0 | 0 | 0.346053 | -3 | 1.764772 | -3 | 0.3047 | 0.11929 | 0.32016 | 0.11638 | 10.83333 | 0.000183502 | 0 | Neutral |
| testneg_8257 | 0.01 | 0.79 | 1.277395 | -3 | 2.450124 | -3 | 0.35967 | 0.09418 | 0.24823 | 0.12788 | 10.22898 | 0.040204167 | 0 | Neutral |
| testneg_8258 | -0.06 | -0.18 | 0.491194 | 0.491194 | 1.681199 | -3 | 0.72124 | 0.09906 | 0.52932 | 0.65749 | 16.8018 | 0.100090229 | 0 | Neutral |
| testneg_8259 | 0 | 0 | 1.002333 | -3 | 4.455256 | -3 | 0.13918 | 0.12788 | 0.13615 | 0.12788 | 16.51743 | 0.032969515 | 0 | Neutral |
| testneg_8260 | -0.02 | 0.68 | 0.243834 | -3 | 1.369352 | -3 | 0.58545 | 0.14414 | 0.21331 | 0.09513 | 18.47769 | 0.03 | 0 | Neutral |
| testneg_8261 | 0.07 | 0.38 | 0.034534 | 0.034534 | 2.298112 | -3 | 0.62381 | 0.9477 | 0.25159 | 0.12788 | 21.49912 | 0.15034912 | 0 | Neutral |
| testneg_8262 | 0 | -0.01 | 0.505705 | -3 | -0.4837 | -3 | 0.73395 | 0.1705 | 0.23276 | 0.31401 | 24.33414 | 0.097704416 | 0 | Neutral |
| testneg_8263 | 0.18 | -1.05 | 7.32748 | -3 | 0.009894 | -3 | 0.08465 | 0.1932 | 0.07996 | 0.08354 | 22.72479 | 0.300227273 | 4.8 | Neutral |
| testneg_8264 | 0.09 | -0.3 | 0.037077 | -3 | 2.470276 | 2.470276 | 0.09004 | 0.07829 | 0.06841 | 0.12788 | 13.02635 | 0.09 | 0 | Neutral |
| testneg_8265 | -0.01 | -0.8 | 2.565801 | -3 | 0.675786 | -3 | 0.99538 | 0.40044 | 0.09531 | 0.09236 | 13.09388 | 0.07 | 0 | Neutral |
| testneg_8266 | 0 | 0 | 1.66597 | -3 | 2.300987 | -3 | 0.66027 | 0.23984 | 0.24931 | 0.78312 | 10.41667 | 0.151105344 | 0 | Neutral |
| testneg_8267 | -0.07 | -0.38 | 5.93813 | -3 | 0.171876 | -3 | 0.14484 | 0.14928 | 0.19969 | 0.1163 | 11.25 | 0.22 | 0.2 | Neutral |
| testneg_8268 | 0 | 0 | 2.064931 | -3 | -0.21059 | -3 | 0.30122 | 0.09165 | 0.03443 | 0.1032 | 20.83333 | 0.02 | 0 | Neutral |
| testneg_8269 | 0 | 0 | 2.100416 | -3 | 0.994999 | -3 | 0.22255 | 0.10057 | 0.51282 | 0.12075 | 14.22149 | 0 | 0 | Neutral |
| testneg_8270 | -0.02 | 0.28 | -0.931321 | -3 | 0.551379 | -3 | 0.06743 | 0.20381 | 0.08265 | 0.12788 | 36.11994 | 0.062123947 | 0 | Neutral |
| testneg_8271 | 0 | 0 | 1.441246 | -3 | 1.625411 | -3 | 0.33292 | 0.17111 | 0.12857 | 0.09376 | 19.78389 | 0.010437406 | 0 | Neutral |
| testneg_8272 | 0.14 | -1.01 | 0.207897 | -3 | 0.332787 | -3 | 0.20616 | 0.29531 | 0.08011 | 0.12788 | 27.95796 | 0.053349774 | 0 | Neutral |
| testneg_8273 | 0 | -0.01 | 1.028355 | -3 | -1.39588 | -3 | 0.87835 | 0.13522 | 0.0478 | 0.12788 | 23.06546 | 0.030136164 | 0 | Neutral |
| testneg_8274 | 0.09 | -0.25 | 3.985725 | -3 | 4.841102 | -3 | 0.72946 | 0.28603 | 0.39932 | 0.10693 | 10.75974 | 0.358718724 | 12.8 | Neutral |
| testneg_8275 | 0.14 | -0.37 | -0.41063 | -3 | 2.70354 | -3 | 0.09611 | 0.07772 | 0.46735 | 0.10674 | 40.63132 | 0.04 | 0 | Neutral |
| testneg_8276 | 0 | 0 | 0.129868 | -3 | 3.798457 | -3 | 0.9874 | 0.15588 | 0.25462 | 0.12788 | 16.8018 | 0.096916667 | 0 | Neutral |
| testneg_8277 | 0 | 0 | 0.850542 | -3 | 0.385771 | -3 | 0.13029 | 0.12788 | 0.11077 | 0.10076 | 10 | 0 | 0 | Neutral |
| testneg_8278 | 0 | 0 | 1.313175 | -3 | 3.273498 | -0.77772 | 0.03953 | 0.05763 | 0.13433 | 0.09578 | 54.39558 | 0.08 | 0 | Neutral |
| testneg_8279 | 0.18 | -1.05 | 7.032758 | -3 | 0.166429 | -3 | 0.07428 | 0.07645 | 0.81659 | 0.10424 | 24.11357 | 0.19 | 0 | Neutral |
| testneg_8280 | 0 | 0 | 0.067126 | -3 | 4.39798 | -3 | 0.18869 | 0.23984 | 0.12689 | 0.13376 | 17.15045 | 0.419219578 | 23.8 | Neutral |
| testneg_8281 | 0 | -0.01 | 0.995506 | -3 | 2.84362 | -3 | 0.39087 | 0.10381 | 0.2903 | 0.11176 | 10.41667 | 0.010533428 | 0 | Neutral |
| testneg_8282 | -0.16 | -0.08 | -1.975003 | -1.975003 | -1.712174 | -3 | 0.08465 | 0.15428 | 0.07604 | 0.35375 | 10 | 0.23 | 0 | Neutral |
| testneg_8283 | 0 | 0 | -0.074827 | -3 | 1.739542 | -3 | 0.30429 | 0.29938 | 0.15443 | 0.10913 | 5.69444 | 0.01 | 0 | Neutral |
| testneg_8284 | -0.1 | -1.9 | 1.885758 | -3 | 4.815849 | -3 | 0.6854 | 0.12563 | 0.16709 | 0.08659 | 12.29167 | 0.132620879 | 0 | Neutral |
| testneg_8285 | 0 | 0 | -0.696429 | -3 | 3.44149 | -3 | 0.26476 | 0.08217 | 0.44103 | 0.10698 | 46.29241 | 0.040669643 | 0 | Neutral |
| testneg_8286 | 0.14 | -0.37 | 1.764776 | -3 | 0.483601 | -3 | 0.6975 | 0.53794 | 0.1 | 0.22633 | 4.44444 | 0.14 | 0 | Neutral |
| testneg_8287 | 0 | 0 | 1.048155 | -3 | 5.697436 | -3 | 0.42523 | 0.13467 | 0.0832 | 0.09323 | 23.21563 | 0.121022072 | 0 | Neutral |
| testneg_8288 | 0 | 0 | 0.991757 | -3 | 3.097852 | -3 | 0.19896 | 0.11604 | 0.16361 | 0.13388 | 33.10982 | 0.15 | 0 | Neutral |
| testneg_8289 | 0 | 0 | 2.988133 | -3 | 0.912939 | -3 | 0.99338 | 0.14229 | 0.49425 | 0.25538 | 10 | 0.182161128 | 0 | Neutral |
| testneg_8290 | 0.14 | -0.37 | -0.153476 | -3 | 0.43445 | -3 | 0.23126 | 0.16717 | 0.09522 | 0.20323 | 10.22898 | 0.112186847 | 0 | Neutral |
| testneg_8291 | 0.08 | 0.03 | -1.290562 | -3 | -1.410124 | -3 | 0.19898 | 0.12895 | 0.33952 | 0.12788 | 13.51351 | 0.02 | 0 | Neutral |
| testneg_8292 | 0 | 0 | 2.099193 | -3 | 1.249535 | -3 | 0.97252 | 0.8859 | 0.75123 | 0.20667 | 10.41667 | 0.11 | 0 | Neutral |
| testneg_8293 | 0 | 0 | 1.861867 | -3 | 0.084517 | -3 | 0.32963 | 0.1035 | 0.25179 | 0.09355 | 19.42085 | 0 | 0 | Neutral |
| testneg_8294 | 0.14 | -0.37 | -0.153476 | -3 | 3.867182 | -3 | 0.23126 | 0.16717 | 0.16777 | 0.1749 | 10 | 0.454656603 | 39.4 | Neutral |
| testneg_8295 | 0.01485714 | -0.0434286 | 1.535215 | -3 | 0.569699 | -3 | 0.26323 | 0.2399 | 0.06344 | 0.12788 | 12.29167 | 0.020195669 | 0 | Neutral |

| | | | | | | | | | | | | | | |
|---|---|---|---|---|---|---|---|---|---|---|---|---|---|---|
| testneg_8296 | 0 | -0.32 | -2.498179 | -3 | 0.340025 | -3 | 0.50682 | 0.12788 | 0.16309 | 0.12788 | 23.32524 | 0.025053763 | 0 | Neutral |
| testneg_8297 | 0.1 | 1.9 | 5.899674 | -3 | 4.323647 | -3 | 0.11329 | 0.12116 | 0.178 | 0.7168 | 23.86584 | 0.6 | 85.2 | Disease_causing |
| testneg_8298 | 0.05 | 0.29 | -0.614223 | -3 | 2.391857 | -3 | 0.20502 | 0.10447 | 0.50687 | 0.11283 | 19.37842 | 0.020177455 | 0 | Neutral |
| testneg_8299 | -0.18 | -0.16 | 0.078335 | -3 | 3.132136 | -3 | 0.20918 | 0.10825 | 0.0105 | 0.12788 | 32.98657 | 0.090136912 | 0 | Neutral |
| testneg_8300 | -0.06 | -0.18 | -0.439044 | -3 | 7.407053 | -3 | 0.3797 | 0.1257 | 0.19883 | 0.09434 | 18.2527 | 0.285076087 | 2.2 | Neutral |
| testneg_8301 | -0.01 | -0.8 | 3.329851 | -3 | 0.468996 | -3 | 0.5184 | 0.11889 | 0.38072 | 0.15993 | 18.88889 | 0.142839938 | 0 | Neutral |
| testneg_8302 | 0.21 | 0.15 | 1.478329 | -3 | 2.155576 | -3 | 0.27614 | 0.12085 | 0.50034 | 0.07527 | 20.69013 | 0.030697115 | 0 | Neutral |
| testneg_8303 | -0.08 | -0.03 | 0.579763 | -3 | 5.351052 | -3 | 0.09978 | 0.12788 | 0.87877 | 0.60823 | 11.25 | 0.413613445 | 21.4 | Neutral |
| testneg_8304 | -0.07 | -0.38 | 0.816148 | -3 | 4.652483 | 4.652483 | 0.26347 | 0.16383 | 0.06841 | 0.10927 | 15.3478 | 0.38 | 13.6 | Neutral |
| testneg_8305 | 0.03 | 0.12 | -0.432053 | -3 | -0.898086 | -3 | 0.10472 | 0.10014 | 0.12314 | 0.11709 | 17.77778 | 0.01 | 0 | Neutral |
| testneg_8306 | 0 | 0 | -0.179321 | -3 | 0.148971 | -3 | 0.14101 | 0.14318 | 0.17007 | 0.12771 | 18.88889 | 0.01 | 0 | Neutral |
| testneg_8307 | 0 | 0 | 0.152024 | -3 | 1.68118 | -3 | 0.30257 | 0.17298 | 0.17412 | 0.10595 | 11.719 | 0.010187325 | 0 | Neutral |
| testneg_8308 | 0.01 | 0.8 | 3.18418 | -3 | -0.360768 | -0.360768 | 0.52412 | 0.21177 | 0.47147 | 0.09503 | 20.22479 | 0.083333333 | 0 | Neutral |
| testneg_8309 | -0.1 | -1.9 | 0.700131 | -3 | -0.931321 | -3 | 0.46056 | 0.12785 | 0.06743 | 0.20381 | 10 | 0.110053191 | 0 | Neutral |
| testneg_8310 | 0.14 | -1.01 | 4.119739 | -3 | -1.089481 | -3 | 0.31664 | 0.14181 | 0.06027 | 0.11574 | 10.83333 | 0.068869048 | 0 | Neutral |
| testneg_8311 | 0 | 0 | 0.390004 | -3 | 1.674048 | -3 | 0.85252 | 0.83576 | 0.25987 | 0.31819 | 10.83333 | 0.12068794 | 0 | Neutral |
| testneg_8312 | 0 | 0 | 0.407242 | -3 | 4.341427 | -3 | 0.77364 | 0.12676 | 0.08143 | 0.44857 | 11.875 | 0.224404637 | 0.2 | Neutral |
| testneg_8313 | 0 | -0.01 | 0.655486 | -3 | -0.274997 | -3 | 0.20772 | 0.16918 | 0.37501 | 0.0997 | 20.83333 | 0.012476302 | 0 | Neutral |
| testneg_8314 | 0 | 0 | 0.717822 | -3 | 1.156713 | 1.156713 | 0.29044 | 0.12142 | 0.36458 | 0.16397 | 17.18434 | 0.050070423 | 0 | Neutral |
| testneg_8315 | 0 | 0 | 0.507327 | -3 | 1.631249 | -3 | 0.11172 | 0.10034 | 0.47166 | 0.13548 | 20.5518 | 0.047083333 | 0 | Neutral |
| testneg_8316 | 0 | 0 | -0.48561 | -3 | 1.360315 | -3 | 0.09432 | 0.15749 | 0.29482 | 0.15726 | 11.92204 | 0.08 | 0 | Neutral |
| testneg_8317 | 0.05 | 0.29 | -0.214978 | -3 | 2.031708 | -3 | 0.39192 | 0.09135 | 0.13061 | 0.21023 | 20 | 0.090142155 | 0 | Neutral |
| testneg_8318 | 0.06 | -0.57 | -0.383439 | -3 | 5.896021 | -3 | 0.23152 | 0.09962 | 0.13601 | 0.19952 | 13.02486 | 0.3493173 | 7 | Neutral |
| testneg_8319 | 0 | 0 | 1.525825 | -3 | 6.248556 | -3 | 0.96651 | 0.28104 | 0.46037 | 0.1343 | 9.1693 | 0.309604608 | 4 | Neutral |
| testneg_8320 | -0.09 | 0.3 | 1.202839 | -3 | 0.220357 | -3 | 0.11144 | 0.08199 | 0.08521 | 0.11239 | 10.22898 | 0.03 | 0 | Neutral |
| testneg_8321 | -0.16 | -0.08 | 5.960242 | -3 | 2.240684 | -3 | 0.1304 | 0.12788 | 0.22686 | 0.24974 | 10.83333 | 0.579354167 | 84 | Disease_causing |
| testneg_8322 | 0 | 0 | 0.227306 | -3 | 0.708831 | -3 | 0.30412 | 0.54626 | 0.40803 | 0.47186 | 4.44444 | 0.1 | 0 | Neutral |
| testneg_8323 | 0 | 0 | 0.067806 | -3 | 2.849519 | 0.059493 | 0.19898 | 0.19444 | 0.14486 | 0.33725 | 12.38369 | 0.238316787 | 0.2 | Neutral |
| testneg_8324 | 0 | 0.32 | 1.479722 | -3 | 2.598788 | -3 | 0.57564 | 0.12788 | 0.19898 | 0.10307 | 20.33333 | 0.04 | 0 | Neutral |
| testneg_8325 | 0 | 0 | 0.079033 | -3 | 4.688426 | -3 | 0.25827 | 0.21386 | 0.37424 | 0.12788 | 16.42913 | 0.111474747 | 0 | Neutral |
| testneg_8326 | 0.04 | -0.23 | 1.5079 | -3 | 0.112631 | -3 | 0.34694 | 0.20397 | 0.10084 | 0.12788 | 18.88889 | 0.03 | 0 | Neutral |
| testneg_8327 | -0.16 | -0.08 | 3.747066 | 3.747066 | 0.774689 | 0.774689 | 0.1535 | 0.12382 | 0.24219 | 0.17458 | 31.66667 | 0.64 | 95.8 | Disease_causing |
| testneg_8328 | 0 | 0 | 1.259391 | -3 | 0.951477 | 0.951477 | 0.27528 | 0.12788 | 0.27083 | 0.14313 | 18.33333 | 0.031102488 | 0 | Neutral |
| testneg_8329 | 0 | -0.01 | 1.828167 | -3 | 0.341464 | 0.341464 | 0.25474 | 0.11262 | 0.1535 | 0.12382 | 39.62462 | 0.01 | 0 | Neutral |
| testneg_8330 | 0 | 0 | 1.302273 | -3 | 1.334446 | 1.334446 | 0.1947 | 0.11413 | 0.15514 | 0.14852 | 18.24258 | 0.046767071 | 0 | Neutral |
| testneg_8331 | 0.21 | 0.15 | 3.567896 | -3 | 1.542294 | -3 | 0.22443 | 0.10657 | 0.18165 | 0.49827 | 5 | 0.265426136 | 0.4 | Neutral |
| testneg_8332 | 0.15 | 0.06 | 6.577474 | -3 | 8.183189 | -3 | 0.08646 | 0.24855 | 0.05767 | 0.07325 | 18.6768 | 0.729933333 | 99.4 | Disease_causing |
| testneg_8333 | 0.07 | -0.31 | 1.374903 | -3 | 6.221201 | -3 | 0.50649 | 0.11394 | 0.22582 | 0.12788 | 13.51351 | 0.12256727 | 0 | Neutral |
| testneg_8334 | 0 | 0 | 0.327712 | 0.327712 | 0.708321 | -3 | 0.57517 | 0.10873 | 0.13869 | 0.10313 | 13.27141 | 0.01003876 | 0 | Neutral |
| testneg_8335 | 0.18 | -1.05 | 6.524735 | -3 | 1.550276 | -3 | 0.20116 | 0.10008 | 0.18142 | 0.21216 | 17.21847 | 0.41734386 | 27.2 | Neutral |
| testneg_8336 | 0 | 0 | 1.826458 | -3 | 2.203067 | -3 | 0.46401 | 0.24608 | 0.27887 | 0.18663 | 10.83333 | 0.12110404 | 0 | Neutral |
| testneg_8337 | -0.07 | -0.38 | 0.517289 | -3 | 2.151559 | -3 | 0.8433 | 0.13128 | 0.82591 | 0.12207 | 10.61346 | 0.030294219 | 0 | Neutral |
| testneg_8338 | -0.19 | -0.51 | 5.496814 | -3 | 6.538148 | -3 | 0.18224 | 0.10419 | 0.16054 | 0.11683 | 4.44444 | 0.58 | 85.2 | Disease_causing |
| testneg_8339 | -0.08 | -0.03 | -0.514498 | -3 | -0.418378 | -3 | 0.81305 | 0.17045 | 0.31744 | 0.10986 | 12.46106 | 0.02 | 0 | Neutral |
| testneg_8340 | 0.18 | -1.05 | 1.792345 | -3 | 2.652503 | -3 | 0.14806 | 0.09574 | 0.08059 | 0.09576 | 50.76058 | 0.06 | 0 | Neutral |
| testneg_8341 | -0.15 | 0.28 | 3.6541 | -3 | 6.111474 | -3 | 0.19898 | 0.37148 | 0.54147 | 0.20818 | 12.98246 | 0.882505656 | 100 | Disease_causing |
| testneg_8342 | 0.02 | 0.45 | -1.097382 | -3 | 0.440748 | -3 | 0.17471 | 0.12788 | 0.14735 | 0.12788 | 11.25 | 0.01 | 0 | Neutral |
| testneg_8343 | -0.07 | 0.31 | 4.767597 | -3 | 2.139035 | -3 | 0.31924 | 0.11145 | 0.85829 | 0.4292 | 11.22104 | 0.279477444 | 1.2 | Neutral |
| testneg_8344 | -0.02 | 0.68 | 2.752736 | -3 | 2.648194 | -3 | 0.15556 | 0.12841 | 0.04655 | 0.10225 | 11.92982 | 0.070405513 | 0 | Neutral |
| testneg_8345 | 0 | 0 | -0.133279 | -3 | -0.121817 | -3 | 0.14948 | 0.12788 | 0.33821 | 0.39417 | 19.06369 | 0.050526877 | 0 | Neutral |
| testneg_8346 | 0.07 | 0.33 | 5.347264 | -3 | 1.233889 | -3 | 0.38911 | 0.12779 | 0.10228 | 0.22182 | 29.86111 | 0.366547619 | 8 | Neutral |
| testneg_8347 | 0.00666667 | 0.08708333 | 0.896944 | -3 | -0.151951 | -3 | 0.11941 | 0.10144 | 0.09918 | 0.11061 | 24.04751 | 0.01 | 0 | Neutral |
| testneg_8348 | 0 | 0 | 0.900502 | -0.566651 | 3.128301 | -3 | 0.38016 | 0.14503 | 0.13458 | 0.17634 | 12.71739 | 0.184347106 | 0 | Neutral |
| testneg_8349 | -0.02 | -0.45 | 1.35797 | -3 | -0.676534 | -0.676534 | 0.51994 | 0.37283 | 0.16262 | 0.15786 | 12.30981 | 0.134686439 | 0 | Neutral |
| testneg_8350 | 0 | 0 | 0.671384 | -3 | 5.21592 | -3 | 0.1798 | 0.0954 | 0.83798 | 0.14433 | 16.0119 | 0.166013302 | 0 | Neutral |
| testneg_8351 | 0.15 | -0.28 | 4.875592 | -3 | 3.373436 | -3 | 0.12269 | 0.08105 | 0.20657 | 0.12788 | 20.89414 | 0.15 | 0 | Neutral |
| testneg_8352 | 0 | 0 | 1.36412 | -3 | 0.584676 | -3 | 0.91691 | 0.25567 | 0.31701 | 0.09986 | 10.83333 | 0.010088975 | 0 | Neutral |
| testneg_8353 | 0.22 | -2.66 | 6.776625 | -3 | 0.05475 | -3 | 0.17039 | 0.12788 | 0.13468 | 0.09235 | 58.89543 | 0.21 | 0.4 | Neutral |
| testneg_8354 | 0 | 0 | 0.387274 | -3 | 4.740439 | -3 | 0.35412 | 0.10819 | 0.28156 | 0.11579 | 17.72163 | 0.020582579 | 0 | Neutral |
| testneg_8355 | 0.22 | -2.66 | 6.542128 | -3 | 3.642505 | -3 | 0.38171 | 0.15977 | 0.39774 | 0.10813 | 20.24348 | 0.5 | 55.2 | Disease_causing |
| testneg_8356 | 0.16 | 0.08 | 1.765317 | -3 | 2.843058 | -3 | 0.072 | 0.3038 | 0.04787 | 0.12788 | 11.78 | 0.14 | 0 | Neutral |
| testneg_8357 | 0.2 | -1.26 | 5.986437 | -3 | 2.229283 | -3 | 0.12325 | 0.09383 | 0.20701 | 0.10345 | 25.12484 | 0.16 | 0 | Neutral |
| testneg_8358 | 0.03 | -1.03 | 3.202376 | -3 | 4.83533 | -3 | 0.25854 | 0.18247 | 0.23975 | 0.13861 | 29.16667 | 0.637485077 | 92.2 | Disease_causing |
| testneg_8359 | 0 | 0 | 1.095746 | -3 | 1.334446 | 1.334446 | 0.24274 | 0.13034 | 0.15514 | 0.14852 | 13.51351 | 0.050809883 | 0 | Neutral |
| testneg_8360 | 0.21 | 0.15 | 6.924355 | -3 | -2.340461 | -3 | 0.38002 | 0.22508 | 0.19898 | 0.12788 | 16.42857 | 0.309258621 | 3.6 | Neutral |
| testneg_8361 | -0.14 | 1.01 | 0.263803 | -3 | 0.104192 | -3 | 0.19898 | 0.12788 | 0.67644 | 0.11323 | 22.11633 | 0.07 | 0 | Neutral |
| testneg_8362 | 0.2 | -1.26 | 5.986437 | -3 | 2.277375 | -3 | 0.12325 | 0.09383 | 0.19898 | 0.09584 | 22.44879 | 0.1555 | 0 | Neutral |
| testneg_8363 | 0.01 | 0.8 | 7.923469 | -3 | 1.802906 | -3 | 0.46792 | 0.11262 | 0.33886 | 0.15644 | 18.19459 | 0.47 | 42 | Neutral |
| testneg_8364 | -0.04 | 0.23 | -1.033933 | -3 | 4.857402 | -3 | 0.16692 | 0.09243 | 0.31648 | 0.12788 | 32.67733 | 0.09 | 0 | Neutral |
| testneg_8365 | -0.15 | 0.28 | -0.668638 | -3 | 5.066909 | -3 | 0.37131 | 0.10994 | 0.0895 | 0.12788 | 29.58554 | 0.199402373 | 0 | Neutral |
| testneg_8366 | 0 | 0 | 0.614162 | -3 | 3.410287 | -3 | 0.54211 | 0.12592 | 0.38942 | 0.345 | 13.62968 | 0.176840049 | 0 | Neutral |
| testneg_8367 | 0 | 0 | -0.072696 | -0.072696 | 0.81674 | 0.81674 | 0.30326 | 0.21562 | 0.84238 | 0.12559 | 10.41667 | 0.03 | 0 | Neutral |
| testneg_8368 | -0.04 | 0.23 | 1.519913 | -3 | 0.430003 | -3 | 0.16436 | 0.19109 | 0.12161 | 0.24178 | 26.34995 | 0.149422261 | 0 | Neutral |
| testneg_8369 | 0.05 | -0.81 | 3.064064 | -3 | -0.857046 | -3 | 0.5481 | 0.10207 | 0.27924 | 0.12788 | 30.26191 | 0.04 | 0 | Neutral |
| testneg_8370 | 0.03 | 1.22 | 0.647985 | -3 | -0.151164 | -0.151164 | 0.21728 | 0.1132 | 0.65708 | 0.0936 | 19.90393 | 0.05 | 0 | Neutral |
| testneg_8371 | 0 | 0 | 1.519976 | -3 | 2.954586 | -3 | 0.19898 | 0.12788 | 0.10646 | 0.1069 | 5.20285 | 0.019210423 | 0 | Neutral |
| testneg_8372 | -0.02 | 0.63 | 4.675931 | -3 | 0.035498 | -3 | 0.961 | 0.25991 | 0.7639 | 0.10408 | 11.5016 | 0.14 | 0 | Neutral |
| testneg_8373 | 0.04 | -0.23 | 1.605283 | -3 | 3.179644 | -3 | 0.7257 | 0.64394 | 0.12871 | 0.29252 | 10.41667 | 0.348818859 | 7 | Neutral |
| testneg_8374 | 0 | -0.01 | 1.887787 | -3 | 1.534507 | -3 | 0.51398 | 0.38821 | 0.98118 | 0.10229 | 4.44444 | 0.025576923 | 0 | Neutral |
| testneg_8375 | -0.0032143 | 0.04654762 | 3.528007 | -3 | 6.395236 | -3 | 0.14785 | 0.09084 | 0.14214 | 0.09456 | 23.78269 | 0.24 | 0.4 | Neutral |
| testneg_8376 | -0.04 | 0.23 | 0.465086 | -3 | 7.611717 | -3 | 0.19898 | 0.12788 | 0.28201 | 0.33127 | 20.37554 | 0.493494709 | 52 | Disease_causing |
| testneg_8377 | 0.04 | -0.23 | 0.872833 | -3 | 6.628365 | -1.271425 | 0.27924 | 0.12788 | 0.10511 | 0.12788 | 11.71717 | 0.21 | 0.2 | Neutral |
| testneg_8378 | 0.23 | -1.86 | 3.254411 | -3 | 2.378984 | -3 | 0.12121 | 0.4142 | 0.11891 | 0.11186 | 15.41302 | 0.24 | 0 | Neutral |
| testneg_8379 | 0.21 | 0.15 | 3.345724 | -3 | 3.6451 | -3 | 0.30275 | 0.12788 | 0.18998 | 0.26095 | 14.36011 | 0.494062549 | 52.2 | Disease_causing |
| testneg_8380 | -0.05 | -0.29 | 0.980231 | -3 | 1.917416 | -3 | 0.53203 | 0.12788 | 0.17697 | 0.12974 | 41.74066 | 0.017424242 | 0 | Neutral |
| testneg_8381 | 0.04 | -0.23 | 6.252157 | -3 | 1.704228 | -3 | 0.21848 | 0.11049 | 0.27062 | 0.12378 | 22.78986 | 0.170551282 | 0.4 | Neutral |
| testneg_8382 | 0.01 | 0.8 | 2.76764 | -3 | 1.294111 | -3 | 0.22795 | 0.12788 | 0.22575 | 0.37247 | 72.82204 | 0.240549786 | 0 | Neutral |
| testneg_8383 | 0 | 0 | 2.044451 | -3 | -0.19389 | -3 | 0.6513 | 0.15254 | 0.12642 | 0.11218 | 18.65054 | 0.01 | 0 | Neutral |
| testneg_8384 | 0.14 | -0.37 | -0.742649 | -3 | 1.424612 | -3 | 0.08476 | 0.12788 | 0.65857 | 0.26294 | 12.40248 | 0.07 | 0 | Neutral |
| testneg_8385 | 0 | 0 | 0.585939 | -3 | 2.478795 | -3 | 0.56904 | 0.14839 | 0.1327 | 0.09885 | 19.31499 | 0.04 | 0 | Neutral |
| testneg_8386 | 0 | 0 | 1.075324 | 1.075324 | -1.162831 | -1.162831 | 0.49934 | 0.11069 | 0.13398 | 0.12788 | 10.38652 | 0.03 | 0 | Neutral |
| testneg_8387 | 0.14 | -0.37 | -0.57366 | -3 | -1.653555 | -1.653555 | 0.12633 | 0.19996 | 0.14588 | 0.12156 | 18.73163 | 0.048651515 | 0 | Neutral |
| testneg_8388 | 0 | 0 | -0.334686 | -3 | 0.86808 | -3 | 0.19898 | 0.10269 | 0.46242 | 0.39055 | 27.25845 | 0.07 | 0 | Neutral |
| testneg_8389 | 0 | 0 | 0.51635 | -3 | 1.986841 | 1.986841 | 0.19898 | 0.82835 | 0.19898 | 0.12788 | 19.46657 | 0.110521077 | 0 | Neutral |
| testneg_8390 | -0.06 | 0.57 | 1.295804 | -3 | 0.203322 | -3 | 0.64795 | 0.21581 | 0.25312 | 0.12788 | 18.8735 | 0.04 | 0 | Neutral |
| testneg_8391 | -0.03 | -1.22 | 5.833395 | -3 | 3.5216 | -3 | 0.2663 | 0.32806 | 0.08646 | 0.24855 | 10.2717 | 0.847995681 | 100 | Disease_causing |
| testneg_8392 | 0 | 0 | 1.103204 | -3 | 1.115221 | -3 | 0.57199 | 0.11262 | 0.64188 | 0.09494 | 14.16105 | 0.01 | 0 | Neutral |
| testneg_8393 | -0.05 | 0.81 | -1.8866 | -3 | 6.649549 | -3 | 0.21811 | 0.18489 | 0.07973 | 0.09208 | 32.70928 | 0.35 | 6 | Neutral |
| testneg_8394 | 0.18 | -1.05 | 3.992521 | -3 | 0.752432 | -3 | 0.19898 | 0.12788 | 0.19898 | 0.1347 | 29.35511 | 0.12 | 0 | Neutral |
| testneg_8395 | 0.22 | -2.66 | 6.137657 | -3 | 4.184741 | -3 | 0.40232 | 0.13342 | 0.89178 | 0.23115 | 10 | 0.69 | 96.6 | Disease_causing |
| testneg_8396 | 0.01 | 0.8 | 3.905173 | -3 | 0.722971 | -3 | 0.16057 | 0.12305 | 0.19898 | 0.12788 | 20.76389 | 0.039411765 | 0 | Neutral |
| testneg_8397 | 0.14 | -0.37 | 2.505643 | -3 | 2.51727 | -3 | 0.96995 | 0.66487 | 0.06196 | 0.10881 | 10.83333 | 0.13 | 0 | Neutral |
| testneg_8398 | 0 | 0 | 1.511019 | -3 | -0.870191 | -3 | 0.33795 | 0.17455 | 0.12831 | 0.10983 | 17.21847 | 0.001974178 | 0 | Neutral |
| testneg_8399 | 0.07 | -0.31 | 2.110113 | -3 | 2.12021 | 2.12021 | 0.68029 | 0.12788 | 0.07698 | 0.12788 | 18.6768 | 0.06 | 0 | Neutral |
| testneg_8400 | 0.07 | 0.38 | 5.286886 | -3 | 0.601255 | -3 | 0.25528 | 0.24339 | 0.14628 | 0.09488 | 12.31231 | 0.171036866 | 0 | Neutral |
| testneg_8401 | 0.02 | 0.41 | 3.730704 | -3 | 1.417365 | 1.417365 | 0.37951 | 0.19435 | 0.61184 | 0.55597 | 17.88889 | 0.312605263 | 3.2 | Neutral |
| testneg_8402 | 0 | 0 | 0.819832 | -3 | 2.235084 | -3 | 0.19898 | 0.12788 | 0.03942 | 0.12788 | 45.49196 | 0.01 | 0 | Neutral |
| testneg_8403 | 0.1 | 1.9 | 4.724695 | -3 | 0.501495 | -3 | 0.25989 | 0.11651 | 0.89357 | 0.22348 | 24.76165 | 0.179778758 | 0 | Neutral |
| testneg_8404 | 0 | 0 | -0.063417 | -3 | 4.614268 | -3 | 0.12554 | 0.10854 | 0.13974 | 0.10225 | 17.21847 | 0.04 | 0 | Neutral |
| testneg_8405 | -0.02 | -0.41 | 0.579456 | 0.579456 | 0.290985 | -3 | 0.47678 | 0.38085 | 0.20986 | 0.10633 | 12.59871 | 0.043333333 | 0 | Neutral |
| testneg_8406 | -0.08 | -0.03 | 3.206814 | -3 | 1.590262 | -3 | 0.7022 | 0.35308 | 0.44385 | 0.1311 | 5 | 0.19 | 0.2 | Neutral |
| testneg_8407 | 0 | 0 | 0.435323 | -3 | -1.873728 | -1.873728 | 0.20527 | 0.10053 | 0.19898 | 0.12788 | 29.34685 | 3.70E-05 | 0 | Neutral |
| testneg_8408 | -0.02 | -0.41 | -1.730578 | -3 | 0.32205 | -3 | 0.24949 | 0.29709 | 0.71276 | 0.88182 | 10.41667 | 0.15 | 0 | Neutral |
| testneg_8409 | 0 | 0 | 1.486765 | -3 | 1.040333 | -3 | 0.41698 | 0.10384 | 0.06793 | 0.12788 | 14.65165 | 0 | 0 | Neutral |
| testneg_8410 | 0.17 | 0.38 | 1.703954 | -3 | 4.69301 | -3 | 0.30823 | 0.17188 | 0.11479 | 0.12788 | 16.32353 | 0.209345238 | 0 | Neutral |
| testneg_8411 | 0 | 0 | 0.477808 | -3 | 3.263002 | -3 | 0.36461 | 0.11015 | 0.33992 | 0.15484 | 14.24157 | 0.118831169 | 0 | Neutral |
| testneg_8412 | 0.15 | -0.28 | 2.017059 | -3 | 3.384241 | -3 | 0.27275 | 0.11752 | 0.1014 | 0.12788 | 29.83991 | 0.07 | 0 | Neutral |
| testneg_8413 | 0 | 0 | -0.104866 | -3 | -0.201571 | -3 | 0.2698 | 0.18815 | 0.09332 | 0.53331 | 11.25 | 0.110603961 | 0 | Neutral |
| testneg_8414 | -0.01 | -0.79 | -3.816017 | -3 | 1.790281 | -3 | 0.06095 | 0.08179 | 0.02059 | 0.066 | 12.29167 | 0.05 | 0 | Neutral |
| testneg_8415 | 0.00916667 | -0.1625 | 0.801954 | -3 | 2.547548 | -3 | 0.09661 | 0.09981 | 0.41647 | 0.11239 | 13.51351 | 0.02 | 0 | Neutral |
| testneg_8416 | 0.07 | 0.38 | 5.12683 | -3 | 5.710272 | -3 | 0.70616 | 0.09323 | 0.508 | 0.1281 | 23.31601 | 0.361977941 | 13 | Neutral |
| testneg_8417 | 0 | 0 | 0.55724 | 0.55724 | -0.490467 | -3 | 0.12937 | 0.23004 | 0.0931 | 0.12788 | 28.93644 | 0.043483516 | 0 | Neutral |

| | | | | | | | | | | | | | | |
|---|---|---|---|---|---|---|---|---|---|---|---|---|---|---|
| testneg_8418 | 0.22 | -2.66 | 5.261584 | -3 | 6.945851 | -3 | 0.49425 | 0.25538 | 0.58292 | 0.87725 | 9.90291 | 0.85 | 100 | Disease_causing |
| testneg_8419 | 0 | 0 | 0.461563 | -3 | 1.673428 | 1.673428 | 0.01573 | 0.10031 | 0.32352 | 0.10243 | 13.40128 | 0.04 | 0 | Neutral |
| testneg_8420 | 0 | 0 | 1.591831 | -3 | 1.003673 | -3 | 0.99668 | 0.30319 | 0.18142 | 0.21216 | 4.72222 | 0.04263571 | 0 | Neutral |
| testneg_8421 | 0 | 0 | 0.040441 | -3 | 3.585728 | -3 | 0.54777 | 0.25991 | 0.17947 | 0.12797 | 11.25 | 0.096140387 | 0 | Neutral |
| testneg_8422 | -0.06 | -0.18 | 0.101783 | -3 | 1.963259 | -3 | 0.06196 | 0.10881 | 0.18964 | 0.34036 | 11.25 | 0.11 | 0 | Neutral |
| testneg_8423 | -0.04 | 0.23 | -0.654233 | -3 | 5.67286 | -3 | 0.04795 | 0.07008 | 0.12161 | 0.24178 | 26.95989 | 0.353020833 | 4.6 | Neutral |
| testneg_8424 | 0.18 | 0.16 | 0.131914 | 0.131914 | 9.272843 | -3 | 0.23879 | 0.12788 | 0.24544 | 0.0894 | 23.30685 | 0.334142857 | 4 | Neutral |
| testneg_8425 | -0.01 | -0.79 | 0.577464 | -3 | 2.065261 | -3 | 0.06366 | 0.12788 | 0.09967 | 0.15417 | 26.12035 | 0.131323656 | 0 | Neutral |
| testneg_8426 | -0.08 | -0.03 | 1.395184 | -3 | 0.061061 | -3 | 0.11335 | 0.14225 | 0.8433 | 0.13128 | 18.05031 | 0.062004902 | 0 | Neutral |
| testneg_8427 | 0 | 0 | 0.044073 | -3 | -0.720759 | -0.720759 | 0.05663 | 0.12788 | 0.17758 | 0.15983 | 27.04107 | 0.14 | 0 | Neutral |
| testneg_8428 | 0.07 | 0.7 | 5.816768 | -3 | 2.72063 | -3 | 0.204 | 0.15424 | 0.65141 | 0.18046 | 10 | 0.669148551 | 96.8 | Disease_causing |
| testneg_8429 | 0 | 0 | -0.02198 | -0.02198 | 2.138507 | -3 | 0.15509 | 0.1326 | 0.95992 | 0.10958 | 19.67631 | 0.03 | 0 | Neutral |
| testneg_8430 | -0.03 | -0.1 | 0.545839 | -3 | 2.456744 | -3 | 0.58545 | 0.14414 | 0.17737 | 0.54971 | 20.69991 | 0.195681211 | 0 | Neutral |
| testneg_8431 | 0 | 0 | 0.096056 | -3 | 1.668921 | -1.582259 | 0.41655 | 0.49485 | 0.15994 | 0.11186 | 10.41667 | 0.02 | 0 | Neutral |
| testneg_8432 | -0.02 | 0.36 | -0.000915 | -3 | 6.835273 | -3 | 0.10349 | 0.12788 | 0.08412 | 0.28307 | 23.5451 | 0.48 | 46 | Neutral |
| testneg_8433 | 0.15 | 0.06 | 5.868312 | -3 | -1.249423 | -3 | 0.19898 | 0.13057 | 0.07225 | 0.12788 | 18.33333 | 0.17 | 0 | Neutral |
| testneg_8434 | -0.21 | -0.15 | -0.68836 | -3 | 6.211821 | -3 | 0.19898 | 0.28128 | 0.1215 | 0.12788 | 11.25 | 0.358738022 | 8.8 | Neutral |
| testneg_8435 | 0 | 0 | 0.894121 | -3 | 1.334627 | -3 | 0.02146 | 0.10351 | 0.10213 | 0.17266 | 13.05945 | 0.11 | 0 | Neutral |
| testneg_8436 | 0.14 | -1.01 | 0.207897 | -3 | 0.991447 | -3 | 0.20616 | 0.29531 | 0.08729 | 0.10515 | 10.41667 | 0.040351261 | 0 | Neutral |
| testneg_8437 | 0 | 0 | 5.815666 | 5.815666 | 0.429523 | -3 | 0.34109 | 0.2167 | 0.18328 | 0.12788 | 10.64628 | 0.62 | 95.6 | Disease_causing |
| testneg_8438 | 0 | -0.32 | 3.779532 | -3 | -0.290973 | -3 | 0.11373 | 0.10739 | 0.1594 | 0.22002 | 10.22898 | 0.160577078 | 0 | Neutral |
| testneg_8439 | -0.04 | 0.89 | 1.989667 | -3 | 2.686945 | -3 | 0.06517 | 0.15107 | 0.08499 | 0.08678 | 18.33333 | 0.100813492 | 0 | Neutral |
| testneg_8440 | 0 | 0 | -0.064795 | -3 | 7.096872 | -3 | 0.4566 | 0.14201 | 0.14486 | 0.33725 | 11.90227 | 0.457245706 | 39.2 | Neutral |
| testneg_8441 | 0.02 | 0.45 | 5.685428 | -3 | 0.897808 | 0.897808 | 0.22877 | 0.14229 | 0.17155 | 0.18585 | 17.20874 | 0.43822496 | 32.6 | Neutral |
| testneg_8442 | -0.19 | 0.25 | -2.608658 | -3 | 5.807705 | -3 | 0.19898 | 0.12788 | 0.89649 | 0.23208 | 36.35443 | 0.360422414 | 7.6 | Neutral |
| testneg_8443 | 0 | 0 | 2.097948 | -3 | -0.44217 | -3 | 0.19989 | 0.06568 | 0.33148 | 0.11968 | 23.23685 | 0.000225996 | 0 | Neutral |
| testneg_8444 | 0 | 0 | 1.599689 | -3 | 1.080175 | -3 | 0.17057 | 0.12788 | 0.18504 | 0.0822 | 11.25 | 0.00015889 | 0 | Neutral |
| testneg_8445 | 0 | 0 | -0.161926 | -3 | 5.824473 | -3 | 0.42033 | 0.17821 | 0.43707 | 0.10923 | 39.94626 | 0.130592157 | 0 | Neutral |
| testneg_8446 | 0.15 | 0.06 | 3.129189 | -3 | 1.996983 | 1.996983 | 0.51137 | 0.19235 | 0.09553 | 0.11637 | 12.89568 | 0.132461124 | 0 | Neutral |
| testneg_8447 | 0.15 | -1.06 | 5.940594 | -3 | 6.509691 | -3 | 0.71803 | 0.95763 | 0.26586 | 0.10825 | 11.00193 | 0.69900432 | 98.4 | Disease_causing |
| testneg_8448 | 0.02 | 0.45 | -1.097382 | -3 | -0.763372 | -3 | 0.17471 | 0.12788 | 0.19979 | 0.09328 | 19.56496 | 0.01 | 0 | Neutral |
| testneg_8449 | 0.15 | -0.28 | 1.45686 | -3 | 3.441233 | -3 | 0.38552 | 0.14172 | 0.22046 | 0.12788 | 12.29167 | 0.085687754 | 0 | Neutral |
| testneg_8450 | -0.1 | -1.9 | 0.261865 | -3 | 1.00433 | -3 | 0.15874 | 0.13137 | 0.06254 | 0.23561 | 18.47544 | 0.13 | 0 | Neutral |
| testneg_8451 | -0.09 | 0.3 | 2.432995 | -3 | 3.316479 | -3 | 0.11941 | 0.10144 | 0.87806 | 0.38966 | 11.92982 | 0.257338034 | 0.2 | Neutral |
| testneg_8452 | 0.16 | 0.08 | 5.361311 | -3 | 0.73492 | -3 | 0.10901 | 0.09549 | 0.67899 | 0.16647 | 13.75 | 0.285044643 | 2.4 | Neutral |
| testneg_8453 | 0 | 0 | -0.027704 | -3 | 1.86207 | -3 | 0.45299 | 0.16306 | 0.96804 | 0.99301 | 5 | 0.090253205 | 0 | Neutral |
| testneg_8454 | 0 | 0 | 0.182861 | -3 | 5.267807 | -3 | 0.25474 | 0.11262 | 0.15685 | 0.17839 | 17.11441 | 0.25130584 | 0.2 | Neutral |
| testneg_8455 | 0.02 | 0.13 | 4.492963 | -3 | 2.297873 | -3 | 0.54648 | 0.19042 | 0.31168 | 0.12788 | 6.99956 | 0.159892611 | 0 | Neutral |
| testneg_8456 | 0 | 0 | 0.861365 | -3 | -0.427621 | -3 | 0.37953 | 0.11262 | 0.40803 | 0.47186 | 10.22898 | 0.020535363 | 0 | Neutral |
| testneg_8457 | 0.01 | 0.8 | 2.08777 | -3 | 0.265638 | -3 | 0.56605 | 0.15893 | 0.17443 | 0.19181 | 14.64741 | 0.080621986 | 0 | Neutral |
| testneg_8458 | -0.01 | -0.8 | 0.369076 | -3 | 1.849894 | 1.653477 | 0.11703 | 0.09364 | 0.13458 | 0.17634 | 27.95796 | 0.12 | 0 | Neutral |
| testneg_8459 | -0.02 | -0.41 | 2.303477 | -3 | 1.290195 | 1.290195 | 0.53216 | 0.84775 | 0.10761 | 0.12246 | 19.44444 | 0.115 | 0 | Neutral |
| testneg_8460 | -0.04 | 0.23 | 2.348169 | -3 | 6.022895 | -3 | 0.60793 | 0.18264 | 0.09583 | 0.09376 | 10.83333 | 0.262291667 | 0.4 | Neutral |
| testneg_8461 | -0.21 | -0.15 | 3.200926 | -3 | 7.496918 | -3 | 0.16052 | 0.12788 | 0.30521 | 0.36067 | 29.31784 | 0.620979167 | 92.4 | Disease_causing |
| testneg_8462 | 0 | 0 | -0.708217 | -3 | 2.950487 | -3 | 0.11533 | 0.12788 | 0.14495 | 0.37127 | 39.04612 | 0.160260135 | 0 | Neutral |
| testneg_8463 | 0.09 | -1.07 | 5.078819 | 5.078819 | 1.983943 | -3 | 0.1228 | 0.10482 | 0.19898 | 0.12788 | 21.5133 | 0.5 | 59 | Disease_causing |
| testneg_8464 | 0.02 | 0.45 | 0.807497 | -3 | 7.372651 | -3 | 0.13395 | 0.29754 | 0.24966 | 0.12788 | 43.06018 | 0.44 | 30.4 | Neutral |
| testneg_8465 | 0 | 0 | 0.029606 | -3 | 2.740758 | -3 | 0.14635 | 0.13588 | 0.17325 | 0.17964 | 8.04888 | 0.098889112 | 0 | Neutral |
| testneg_8466 | -0.05 | -0.29 | -0.695099 | -3 | 1.290195 | 1.290195 | 0.84318 | 0.11313 | 0.10761 | 0.12246 | 13.51351 | 0.02 | 0 | Neutral |
| testneg_8467 | 0 | 0 | 1.13018 | -3 | 0.626243 | 0.626243 | 0.50505 | 0.11563 | 0.34412 | 0.08672 | 13.02635 | 0.01 | 0 | Neutral |
| testneg_8468 | 0.22 | -2.66 | 5.605976 | -3 | 0.727995 | -3 | 0.40897 | 0.13253 | 0.06215 | 0.08575 | 30.01316 | 0.174861111 | 0 | Neutral |
| testneg_8469 | 0.04 | 2.01 | 1.240851 | -3 | 1.873147 | 1.873147 | 0.23859 | 0.1202 | 0.1499 | 0.09632 | 18.15261 | 0.1 | 0 | Neutral |
| testneg_8470 | 0.05 | 0.29 | 5.917332 | -3 | 3.423345 | -3 | 0.32089 | 0.1035 | 0.24038 | 0.15296 | 12.70833 | 0.579504426 | 80.6 | Disease_causing |
| testneg_8471 | 0.03 | 0.12 | 6.140208 | -3 | 0.013725 | -3 | 0.40179 | 0.12112 | 0.19898 | 0.12788 | 14.64915 | 0.15 | 0 | Neutral |
| testneg_8472 | -0.01 | -0.79 | 1.315391 | -3 | 6.760768 | -3 | 0.99417 | 0.17365 | 0.35532 | 0.24703 | 11.25 | 0.504722222 | 56.4 | Disease_causing |
| testneg_8473 | -0.05 | -1.52 | -1.75663 | -3 | 6.717504 | -3 | 0.19898 | 0.12788 | 0.19898 | 0.12788 | 30.91017 | 0.21 | 0.2 | Neutral |
| testneg_8474 | -0.1 | -1.9 | 0.865544 | -3 | -0.808449 | -0.808449 | 0.99448 | 0.67288 | 0.13258 | 0.9258 | 10.41667 | 0.221675559 | 0.2 | Neutral |
| testneg_8475 | 0 | -0.01 | 1.693273 | -3 | 2.022518 | -3 | 0.47788 | 0.10817 | 0.10891 | 0.12788 | 18.88889 | 0.01 | 0 | Neutral |
| testneg_8476 | 0.02 | -0.68 | 1.359285 | -3 | 2.396949 | -3 | 0.85075 | 0.16646 | 0.29268 | 0.16012 | 14.27748 | 0.1700337 | 0 | Neutral |
| testneg_8477 | 0 | 0 | 0.938815 | -3 | 0.436295 | -3 | 0.33292 | 0.17111 | 0.32677 | 0.12788 | 11.54167 | 0.01 | 0 | Neutral |
| testneg_8478 | 0 | 0 | 1.132066 | -3 | 1.746837 | -3 | 0.99317 | 0.60884 | 0.07912 | 0.09055 | 12.7356 | 0.04 | 0 | Neutral |
| testneg_8479 | 0.14 | -0.37 | 0.198379 | -3 | -0.136916 | -3 | 0.03991 | 0.08342 | 0.04853 | 0.4186 | 15 | 0.144166667 | 0 | Neutral |
| testneg_8480 | -0.005035 | 0.0134965 | 0.615258 | -3 | 0.192939 | -3 | 0.59345 | 0.26484 | 0.17725 | 0.12788 | 19.43262 | 0.033196658 | 0 | Neutral |
| testneg_8481 | 0 | 0 | 1.639442 | -3 | 0.470746 | -3 | 0.25905 | 0.23728 | 0.99543 | 0.12788 | 6.45329 | 0.05 | 0 | Neutral |
| testneg_8482 | 0 | 0 | 0.300937 | -3 | -0.262132 | -3 | 0.46318 | 0.14229 | 0.11788 | 0.12788 | 18.88889 | 0.01 | 0 | Neutral |
| testneg_8483 | 0.06 | 0.18 | 2.169317 | -3 | 9.808835 | -3 | 0.17083 | 0.31043 | 0.41193 | 0.10133 | 10.41667 | 0.51 | 60.2 | Disease_causing |
| testneg_8484 | 0 | 0 | 0.097566 | -3 | 5.900567 | -3 | 0.24452 | 0.21203 | 0.07275 | 0.09562 | 22.29376 | 0.180691303 | 0.2 | Neutral |
| testneg_8485 | 0 | 0 | 0.663777 | -3 | -0.055258 | -3 | 0.68055 | 0.12788 | 0.16139 | 0.09116 | 21.38889 | 0.013858974 | 0 | Neutral |
| testneg_8486 | 0 | 0 | 1.15789 | -3 | 0.196098 | -3 | 0.68439 | 0.09371 | 0.12682 | 0.11357 | 9.98893 | 0.01 | 0 | Neutral |
| testneg_8487 | -0.03 | 1.03 | -1.689038 | -3 | 2.030516 | -3 | 0.64256 | 0.12241 | 0.15056 | 0.40109 | 19.09347 | 0.120072464 | 0 | Neutral |
| testneg_8488 | 0 | -0.01 | 0.494045 | -3 | 1.178544 | -3 | 0.1304 | 0.12788 | 0.19898 | 0.12788 | 30.55556 | 0 | 0 | Neutral |
| testneg_8489 | 0.14 | -1.01 | 3.778793 | -3 | 4.8749 | -3 | 0.91921 | 0.32331 | 0.22225 | 0.11186 | 12.70833 | 0.355871327 | 10.6 | Neutral |
| testneg_8490 | 0.09 | -0.3 | -0.213037 | -3 | 3.063227 | -3 | 0.54113 | 0.11017 | 0.05153 | 0.12788 | 21.92108 | 0.106666667 | 0 | Neutral |
| testneg_8491 | 0 | 0 | 0.29846 | -3 | -0.203981 | -3 | 0.07543 | 0.10211 | 0.1417 | 0.1398 | 18.33333 | 0.04 | 0 | Neutral |
| testneg_8492 | -0.21 | -0.15 | 1.014556 | -3 | 0.343601 | -3 | 0.09734 | 0.08962 | 0.22874 | 0.13202 | 20 | 0.11 | 0 | Neutral |
| testneg_8493 | 0.07 | 0.01 | 0.631605 | -3 | 6.018396 | -3 | 0.14524 | 0.10162 | 0.07571 | 0.09432 | 18.33333 | 0.107933333 | 0 | Neutral |
| testneg_8494 | 0.03 | 0.12 | 3.628292 | -3 | 0.771752 | -3 | 0.15562 | 0.10384 | 0.13702 | 0.12788 | 19.44444 | 0.020180006 | 0 | Neutral |
| testneg_8495 | 0 | 0 | -0.179818 | -3 | -1.638874 | -3 | 0.21014 | 0.12378 | 0.22078 | 0.12788 | 18.76159 | 0 | 0 | Neutral |
| testneg_8496 | 0.03 | 1.22 | 5.911847 | -3 | 5.713281 | -3 | 0.40179 | 0.12112 | 0.28066 | 0.18241 | 18.8191 | 0.739232955 | 99.6 | Disease_causing |
| testneg_8497 | 0.05 | -0.81 | 0.660523 | -3 | -0.690909 | -3 | 0.06897 | 0.08445 | 0.62381 | 0.12788 | 28.03459 | 0.02 | 0 | Neutral |
| testneg_8498 | -0.03 | -0.1 | 6.945269 | -3 | 0.746089 | -3 | 0.09972 | 0.13992 | 0.08454 | 0.22908 | 10.83333 | 0.568819444 | 79.6 | Disease_causing |
| testneg_8499 | -0.14 | 1.01 | 2.72498 | -3 | -0.380842 | -3 | 0.34001 | 0.12788 | 0.62381 | 0.12788 | 12.8129 | 0.07 | 0 | Neutral |
| testneg_8500 | 0.09 | -0.3 | 3.536595 | -3 | 2.655425 | -3 | 0.33795 | 0.17455 | 0.19883 | 0.09434 | 16.668 | 0.149895833 | 0 | Neutral |
| testneg_8501 | 0.03 | 0.12 | 3.086284 | 0.262307 | 1.377333 | -3 | 0.13099 | 0.17453 | 0.06215 | 0.08575 | 18.88889 | 0.098571429 | 0 | Neutral |
| testneg_8502 | 0 | 0 | 1.928458 | -3 | 1.795817 | -3 | 0.45152 | 0.34457 | 0.16053 | 0.15049 | 20.90842 | 0.143356983 | 0 | Neutral |
| testneg_8503 | 0 | -0.01 | 2.245398 | -3 | 0.997782 | -3 | 0.20675 | 0.13709 | 0.48817 | 0.10601 | 8.97898 | 0.000196154 | 0 | Neutral |
| testneg_8504 | -0.02 | 0.68 | 5.845448 | -3 | 3.701009 | -3 | 0.06926 | 0.08399 | 0.12642 | 0.11218 | 25.9009 | 0.300081585 | 3.8 | Neutral |
| testneg_8505 | 0.17 | 0.38 | 5.080349 | -3 | 0.341374 | 0.341374 | 0.20219 | 0.10986 | 0.10646 | 0.1069 | 14.68708 | 0.07 | 0 | Neutral |
| testneg_8506 | 0 | 0 | 1.943537 | 1.943537 | 6.195353 | -3 | 0.19237 | 0.23658 | 0.28532 | 0.10663 | 17.21847 | 0.293333333 | 2.2 | Neutral |
| testneg_8507 | 0.04 | -0.23 | -0.122196 | -0.122196 | 1.294129 | -3 | 0.31961 | 0.17571 | 0.14498 | 0.33896 | 5.69444 | 0.123990661 | 0 | Neutral |
| testneg_8508 | 0.17 | 0.38 | 3.17007 | -3 | -2.044408 | -3 | 0.12377 | 0.12788 | 0.10084 | 0.12788 | 29.28571 | 0.04 | 0 | Neutral |
| testneg_8509 | 0 | 0 | 1.975201 | -3 | 6.314281 | -3 | 0.34495 | 0.16861 | 0.21811 | 0.18489 | 10.41667 | 0.533044311 | 65.8 | Disease_causing |
| testneg_8510 | 0 | -0.01 | 0.265894 | -3 | 6.399925 | -3 | 0.10723 | 0.11163 | 0.92262 | 0.12859 | 12.83951 | 0.14 | 0 | Neutral |
| testneg_8511 | -0.08 | -0.03 | 1.02493 | -3 | 2.025162 | -3 | 0.17301 | 0.14089 | 0.32016 | 0.11638 | 8.612 | 0.02 | 0 | Neutral |
| testneg_8512 | 0 | -0.01 | 2.087209 | -3 | 3.314525 | -3 | 0.06841 | 0.10927 | 0.0359 | 0.09574 | 12.32809 | 0.040267794 | 0 | Neutral |
| testneg_8513 | 0 | 0 | 0.101625 | -3 | 0.921828 | -3 | 0.6323 | 0.09928 | 0.31646 | 0.09844 | 12.35574 | 0.00036039 | 0 | Neutral |
| testneg_8514 | 0.02 | -0.36 | 2.239256 | -3 | 3.162469 | -3 | 0.07175 | 0.29293 | 0.17412 | 0.10595 | 11.78141 | 0.14470064 | 0 | Neutral |
| testneg_8515 | -0.15 | 0.28 | 3.03067 | -3 | 0.933262 | -3 | 0.08723 | 0.09503 | 0.49831 | 0.29427 | 56.94444 | 0.258035714 | 0 | Neutral |
| testneg_8516 | 0.21 | 0.15 | 6.051557 | -3 | -0.431211 | -3 | 0.13482 | 0.12788 | 0.36676 | 0.37749 | 10.22898 | 0.408996337 | 21 | Neutral |
| testneg_8517 | 0.02 | -0.68 | 5.953248 | -3 | -0.247306 | -3 | 0.81169 | 0.11096 | 0.35945 | 0.12025 | 18.88889 | 0.164666667 | 0 | Neutral |
| testneg_8518 | 0.04 | -0.23 | 5.589671 | -3 | 3.952275 | -3 | 0.15556 | 0.12841 | 0.12036 | 0.18651 | 19.93118 | 0.69002264 | 98.2 | Disease_causing |
| testneg_8519 | 0.02 | -0.36 | 1.715391 | -3 | 0.031503 | -3 | 0.60959 | 0.76078 | 0.49504 | 0.08191 | 10 | 0.041330121 | 0 | Neutral |
| testneg_8520 | 0 | -0.32 | 2.665731 | -3 | 1.942918 | -3 | 0.54826 | 0.24974 | 0.11187 | 0.10744 | 17.62359 | 0.05 | 0 | Neutral |
| testneg_8521 | 0 | 0 | 0.984412 | -3 | 5.56944 | -3 | 0.34931 | 0.11906 | 0.1664 | 0.10113 | 20.83333 | 0.074944655 | 0 | Neutral |
| testneg_8522 | 0.05 | 0.29 | 2.385227 | -3 | 1.357664 | 1.357664 | 0.21265 | 0.1284 | 0.1798 | 0.0954 | 18.33333 | 0.02 | 0 | Neutral |
| testneg_8523 | 0 | 0 | -0.055177 | -3 | 0.530222 | -3 | 0.32558 | 0.32796 | 0.19898 | 0.12788 | 10.83333 | 0.020113085 | 0 | Neutral |
| testneg_8524 | 0.21 | 0.15 | 3.051827 | -3 | 0.280115 | -3 | 0.113 | 0.12788 | 0.19898 | 0.08111 | 14.40495 | 0.04 | 0 | Neutral |
| testneg_8525 | 0 | 0 | 0.358339 | 0.358339 | 3.427531 | -3 | 0.13682 | 0.11872 | 0.09527 | 0.09424 | 42.88607 | 0.05 | 0 | Neutral |
| testneg_8526 | 0 | 0 | 1.333395 | -3 | 5.728537 | -3 | 0.12259 | 0.1041 | 0.28199 | 0.10688 | 13.81407 | 0.06 | 0 | Neutral |
| testneg_8527 | -0.04 | 0.23 | -0.28312 | -3 | 6.448066 | -3 | 0.06513 | 0.12353 | 0.40707 | 0.37988 | 45.00049 | 0.433333333 | 26.6 | Neutral |
| testneg_8528 | -0.19 | 0.25 | -1.729462 | -3 | 2.019386 | -3 | 0.16155 | 0.20638 | 0.13751 | 0.13274 | 21.38889 | 0.25 | 0 | Neutral |
| testneg_8529 | 0.14 | -0.69 | 3.846708 | -3 | 0.740476 | -3 | 0.50403 | 0.21568 | 0.49562 | 0.11809 | 18.45005 | 0.093148148 | 0 | Neutral |
| testneg_8530 | 0.08 | 0.03 | 0.574162 | -3 | 3.089818 | -3 | 0.33292 | 0.17111 | 0.09026 | 0.09447 | 11.54167 | 0.141813899 | 0 | Neutral |
| testneg_8531 | 0.04 | -0.23 | 4.987949 | -3 | 5.359975 | -3 | 0.19898 | 0.48545 | 0.12196 | 0.12788 | 10.41667 | 0.57264148 | 81 | Disease_causing |
| testneg_8532 | 0 | 0 | 1.73925 | -3 | 0.43445 | -3 | 0.12971 | 0.11434 | 0.09522 | 0.20323 | 9.62209 | 0.03197561 | 0 | Neutral |
| testneg_8533 | 0 | 0 | -0.384176 | -3 | 12.061237 | -3 | 0.31238 | 0.14393 | 0.19898 | 0.12788 | 42.33561 | 0.37 | 11.2 | Neutral |
| testneg_8534 | 0 | 0 | 0.327712 | -3 | 5.091023 | -3 | 0.57517 | 0.10873 | 0.09028 | 0.10643 | 13.51351 | 0.055714204 | 0 | Neutral |
| testneg_8535 | 0.02 | -0.63 | 0.686794 | -3 | 4.065196 | -3 | 0.15132 | 0.11724 | 0.10286 | 0.26819 | 10.83333 | 0.302607559 | 3.4 | Neutral |
| testneg_8536 | -0.16 | -0.08 | 1.821379 | -3 | 0.411221 | -3 | 0.24026 | 0.12788 | 0.05851 | 0.0724 | 18.72908 | 0.05 | 0 | Neutral |
| testneg_8537 | 0 | 0 | 1.264197 | 1.264197 | 1.455667 | -3 | 0.37542 | 0.19444 | 0.14341 | 0.10134 | 19.44444 | 0.030208424 | 0 | Neutral |
| testneg_8538 | 0.09 | -0.3 | 6.191692 | -3 | -0.999741 | -3 | 0.45927 | 0.74955 | 0.21587 | 0.11701 | 10.22898 | 0.3295 | 3.8 | Neutral |
| testneg_8539 | 0.08 | 0.03 | -2.363283 | -2.363283 | 3.314525 | -3 | 0.05846 | 0.12788 | 0.0359 | 0.09574 | 26.84566 | 0.11 | 0 | Neutral |

| | | | | | | | | | | | | | | |
|---|---|---|---|---|---|---|---|---|---|---|---|---|---|---|
| testneg_8540 | 0.05 | 0.29 | -1.569633 | -3 | 1.285186 | 1.285186 | 0.02759 | 0.12458 | 0.35957 | 0.11476 | 17.21847 | 0.04 | 0 | Neutral |
| testneg_8541 | 0 | 0 | -0.034419 | -0.034419 | 2.080531 | -3 | 0.62203 | 0.12788 | 0.42235 | 0.12474 | 13.14653 | 0.02 | 0 | Neutral |
| testneg_8542 | -0.07 | -0.38 | 2.988243 | -3 | 5.047397 | -3 | 0.54015 | 0.13176 | 0.09439 | 0.1884 | 20.98109 | 0.649480437 | 95.6 | Disease_causing |
| testneg_8543 | 0.07 | 0.33 | 2.134433 | -3 | 5.02555 | -3 | 0.27887 | 0.19826 | 0.17725 | 0.12788 | 18.81641 | 0.243794486 | 0.2 | Neutral |
| testneg_8544 | -0.0078182 | -0.0090909 | 10.86776 | -3 | 1.624378 | -3 | 0.12357 | 0.14417 | 0.2016 | 0.13956 | 19.16667 | 0.507984375 | 57.4 | Disease_causing |
| testneg_8545 | 0 | 0 | 1.916985 | -3 | 0.127079 | -3 | 0.84031 | 0.16749 | 0.3012 | 0.10317 | 10.73171 | 0.02 | 0 | Neutral |
| testneg_8546 | 0 | 0 | 0.800388 | -3 | 0.009894 | -3 | 0.47816 | 0.66526 | 0.07996 | 0.08354 | 22.72479 | 0.052140845 | 0 | Neutral |
| testneg_8547 | -0.04 | -2.01 | 4.558544 | -3 | 1.241126 | -3 | 0.08753 | 0.09666 | 0.89959 | 0.55078 | 21.42735 | 0.265479437 | 0.8 | Neutral |
| testneg_8548 | 0.05 | 0.29 | 2.040423 | -3 | 1.835259 | -3 | 0.835 | 0.13588 | 0.14767 | 0.09559 | 11.58727 | 0.020345238 | 0 | Neutral |
| testneg_8549 | 0.06 | 0.18 | 3.024681 | -3 | 4.54141 | -3 | 0.40557 | 0.11865 | 0.20431 | 0.17304 | 13.02635 | 0.478640375 | 45.2 | Neutral |
| testneg_8550 | 0.07 | 0.38 | -1.485747 | -3 | 2.327225 | -3 | 0.13029 | 0.12788 | 0.48504 | 0.12788 | 10 | 0.03 | 0 | Neutral |
| testneg_8551 | 0 | 0 | 0.240863 | -3 | 1.649381 | -3 | 0.22816 | 0.10878 | 0.22686 | 0.24974 | 11.92061 | 0.020833333 | 0 | Neutral |
| testneg_8552 | 0.02 | -0.36 | 2.305765 | -3 | 2.275026 | -3 | 0.11055 | 0.1127 | 0.16777 | 0.1749 | 14.06008 | 0.109813084 | 0 | Neutral |
| testneg_8553 | 0 | 0 | 0.044073 | -3 | 2.165318 | -3 | 0.05663 | 0.12788 | 0.19989 | 0.16777 | 44.08717 | 0.13 | 0 | Neutral |
| testneg_8554 | -0.06 | -0.18 | 4.193908 | -3 | -0.269735 | -0.269735 | 0.23088 | 0.12788 | 0.12461 | 0.1241 | 33.28229 | 0.04 | 0 | Neutral |
| testneg_8555 | 0 | 0 | 0.004541 | -1.007938 | 4.30647 | -3 | 0.35984 | 0.11129 | 0.19898 | 0.1347 | 18.80655 | 0.203047619 | 0 | Neutral |
| testneg_8556 | 0 | 0 | 0.068014 | 0.068014 | 3.309253 | -3 | 0.20869 | 0.13493 | 0.47335 | 0.12788 | 11.90227 | 0.045077433 | 0 | Neutral |
| testneg_8557 | 0 | 0 | 1.436812 | -3 | 1.723834 | -3 | 0.29302 | 0.17394 | 0.06958 | 0.16086 | 12.80909 | 0.090249956 | 0 | Neutral |
| testneg_8558 | 0 | 0 | 2.286408 | -3 | 0.134977 | -3 | 0.99997 | 0.53296 | 0.0925 | 0.08222 | 10.41667 | 0.05267885 | 0 | Neutral |
| testneg_8559 | -0.08 | -0.03 | 2.38382 | -3 | 1.140918 | -3 | 0.13083 | 0.23231 | 0.1846 | 0.08682 | 16.8018 | 0.042232561 | 0 | Neutral |
| testneg_8560 | 0 | -0.01 | 0.418093 | -3 | 1.577566 | -3 | 0.26524 | 0.11154 | 0.68423 | 0.08163 | 13.51351 | 0.001042781 | 0 | Neutral |
| testneg_8561 | -0.19 | -0.51 | 2.183162 | -3 | 1.621929 | -3 | 0.22078 | 0.12788 | 0.95247 | 0.11285 | 16.76768 | 0.06 | 0 | Neutral |
| testneg_8562 | 0 | 0 | -0.113096 | -0.113096 | -0.765143 | -0.765143 | 0.04436 | 0.12788 | 0.31627 | 0.23288 | 11.41859 | 0.090152778 | 0 | Neutral |
| testneg_8563 | 0.08 | 0.03 | -1.587151 | -3 | -0.022376 | -3 | 0.15486 | 0.28768 | 0.05817 | 0.09497 | 18.88889 | 0.067991071 | 0 | Neutral |
| testneg_8564 | 0 | 0 | -0.76928 | -3 | 7.300397 | -3 | 0.19898 | 0.12788 | 0.22162 | 0.09445 | 28.50211 | 0.228541667 | 1.2 | Neutral |
| testneg_8565 | 0.04 | 2.01 | -1.062451 | -1.062451 | 8.837798 | -3 | 0.03072 | 0.27473 | 0.20222 | 0.12788 | 20.98109 | 0.53 | 68.8 | Disease_causing |
| testneg_8566 | 0 | 0 | 0.011632 | -3 | 4.231853 | -3 | 0.32406 | 0.10985 | 0.47751 | 0.2062 | 12.70833 | 0.179800595 | 0.2 | Neutral |
| testneg_8567 | 0 | 0 | 2.587234 | 2.587234 | -3.330542 | -3 | 0.34931 | 0.11906 | 0.02723 | 0.04872 | 18.88889 | 0.2 | 0.4 | Neutral |
| testneg_8568 | 0.02 | 0.45 | 1.83509 | -3 | 1.8745 | -3 | 0.57199 | 0.11262 | 0.17597 | 0.12788 | 22.13012 | 0.020392308 | 0 | Neutral |
| testneg_8569 | -0.07 | -0.01 | 1.176476 | 1.176476 | 1.040422 | -3 | 0.64432 | 0.10409 | 0.13413 | 0.12788 | 4.72222 | 0.03 | 0 | Neutral |
| testneg_8570 | -0.08 | -0.03 | 0.885873 | -3 | 3.63274 | -3 | 0.06474 | 0.23607 | 0.25987 | 0.31819 | 10.83333 | 0.510056548 | 57.8 | Disease_causing |
| testneg_8571 | 0 | 0 | -0.920517 | -3 | 1.227969 | -3 | 0.6975 | 0.53794 | 0.27924 | 0.12788 | 11.83526 | 0.022263158 | 0 | Neutral |
| testneg_8572 | 0.07 | 0.38 | 1.157706 | -3 | 6.431695 | -3 | 0.09951 | 0.14299 | 0.17764 | 0.09681 | 18.41698 | 0.259166667 | 0.2 | Neutral |
| testneg_8573 | 0.16 | -1.37 | 2.859591 | -3 | 6.431621 | 6.431621 | 0.1576 | 0.12788 | 0.26676 | 0.76568 | 11.22698 | 0.685546685 | 95 | Disease_causing |
| testneg_8574 | -0.08 | -0.03 | 5.054259 | -3 | 3.775158 | -3 | 0.29478 | 0.12788 | 0.07297 | 0.12788 | 13.2557 | 0.246436275 | 0.6 | Neutral |
| testneg_8575 | 0 | 0 | 0.261866 | -3 | 1.01324 | -3 | 0.35584 | 0.10874 | 0.45354 | 0.09957 | 13.44572 | 0 | 0 | Neutral |
| testneg_8576 | 0 | 0 | 1.431043 | -3 | 5.388804 | -3 | 0.08489 | 0.12788 | 0.11613 | 0.1109 | 14.25869 | 0.06118118 | 0 | Neutral |
| testneg_8577 | 0.16 | 0.08 | -2.804234 | -3 | 0.049172 | -3 | 0.57349 | 0.09764 | 0.22034 | 0.12788 | 16.25133 | 0.031589992 | 0 | Neutral |
| testneg_8578 | 0.21 | 0.15 | 5.850445 | -3 | 2.716468 | -3 | 0.25236 | 0.11809 | 0.28066 | 0.18241 | 11.64327 | 0.566820175 | 75 | Disease_causing |
| testneg_8579 | 0.04 | -0.23 | 3.770404 | -3 | 2.597966 | -3 | 0.27942 | 0.73513 | 0.19898 | 0.12788 | 23.3887 | 0.249698068 | 0.6 | Neutral |
| testneg_8580 | 0.07 | -0.31 | 2.678817 | -3 | 0.505411 | -3 | 0.12977 | 0.1128 | 0.19898 | 0.09431 | 14.27748 | 0.01 | 0 | Neutral |
| testneg_8581 | 0.16 | -1.37 | 4.304011 | 4.304011 | 5.365154 | -3 | 0.23401 | 0.12374 | 0.10213 | 0.22954 | 10.83333 | 0.870901973 | 100 | Disease_causing |
| testneg_8582 | -0.02 | 0.68 | 4.605406 | -3 | 1.159987 | 1.159987 | 0.22113 | 0.10673 | 0.42748 | 0.10231 | 19.44444 | 0.07 | 0 | Neutral |
| testneg_8583 | 0.21 | 0.15 | 0.789411 | -3 | 7.845205 | 2.840674 | 0.13067 | 0.1005 | 0.12965 | 0.75487 | 18.6768 | 0.56 | 80.6 | Disease_causing |
| testneg_8584 | 0.15 | 0.06 | 5.566047 | -3 | -0.78482 | -3 | 0.13331 | 0.13638 | 0.1273 | 0.12788 | 11.25 | 0.16 | 0 | Neutral |
| testneg_8585 | 0 | 0 | -0.404338 | -3 | 1.856947 | -3 | 0.17631 | 0.14648 | 0.56862 | 0.12788 | 34.28675 | 0.02 | 0 | Neutral |
| testneg_8586 | 0.16 | -1.37 | -0.647508 | -3 | -0.513734 | -3 | 0.14628 | 0.09488 | 0.256 | 0.20036 | 10.22898 | 0.080330601 | 0 | Neutral |
| testneg_8587 | 0.17 | 0.38 | 5.080349 | -3 | 2.297873 | -3 | 0.20219 | 0.10986 | 0.31168 | 0.12788 | 13.41359 | 0.12 | 0 | Neutral |
| testneg_8588 | 0 | 0 | 1.431043 | -3 | 6.000031 | -3 | 0.08489 | 0.12788 | 0.13173 | 0.12788 | 16.06424 | 0.1 | 0 | Neutral |
| testneg_8589 | -0.02 | 0.36 | 2.178935 | -3 | 1.110558 | -3 | 0.77847 | 0.10751 | 0.42666 | 0.12788 | 10.83333 | 0.02 | 0 | Neutral |
| testneg_8590 | 0.00454545 | -0.1357273 | 0.208107 | -3 | -0.35214 | -3 | 0.10863 | 0.098 | 0.11134 | 0.13818 | 28.85698 | 0.06 | 0 | Neutral |
| testneg_8591 | 0.15 | -0.28 | 0.00589 | -3 | -0.729977 | -0.729977 | 0.38503 | 0.11286 | 0.17974 | 0.12788 | 11.69163 | 0.010253205 | 0 | Neutral |
| testneg_8592 | 0 | 0 | 0.861365 | -3 | 0.473397 | -3 | 0.37953 | 0.11262 | 0.07064 | 0.11699 | 19.09347 | 0 | 0 | Neutral |
| testneg_8593 | 0 | 0 | 0.371766 | -3 | 1.551152 | -3 | 0.3095 | 0.13502 | 0.22982 | 0.11215 | 21.30293 | 0.01 | 0 | Neutral |
| testneg_8594 | 0 | -0.01 | 0.507765 | -3 | 0.272659 | 0.272659 | 0.65407 | 0.50749 | 0.17093 | 0.23281 | 17.56627 | 0.129659665 | 0 | Neutral |
| testneg_8595 | 0.08 | -1.2 | -0.474484 | -3 | -1.73577 | -3 | 0.45075 | 0.12232 | 0.0931 | 0.12788 | 25.97348 | 0.040428571 | 0 | Neutral |
| testneg_8596 | 0 | 0 | -0.491414 | -3 | 3.003558 | -3 | 0.61027 | 0.14229 | 0.19898 | 0.12788 | 28.68357 | 0.066048244 | 0 | Neutral |
| testneg_8597 | 0.2 | -1.26 | -1.784808 | -3 | -1.310227 | -3 | 0.11133 | 0.12788 | 0.08688 | 0.08809 | 32.47259 | 0.05 | 0 | Neutral |
| testneg_8598 | -0.15 | -0.04 | -1.165102 | -1.165102 | -0.152553 | -3 | 0.37237 | 0.36829 | 0.05087 | 0.09682 | 10.41667 | 0.08 | 0 | Neutral |
| testneg_8599 | -0.02 | -0.13 | 3.206825 | -3 | 1.214875 | -3 | 0.55197 | 0.11305 | 0.80889 | 0.40016 | 5.76832 | 0.14855489 | 0 | Neutral |
| testneg_8600 | 0 | 0 | -0.342253 | -3 | 1.669334 | -3 | 0.63363 | 0.14795 | 0.19898 | 0.12788 | 13.03436 | 0.01 | 0 | Neutral |
| testneg_8601 | 0.15 | -0.28 | 5.739294 | -3 | 1.018818 | 1.018818 | 0.07701 | 0.19824 | 0.13398 | 0.12788 | 13.51351 | 0.23 | 0.8 | Neutral |
| testneg_8602 | -0.18 | -0.16 | 4.070803 | -3 | 3.735934 | -3 | 0.37951 | 0.19435 | 0.20047 | 0.11827 | 27.3569 | 0.282353239 | 1.4 | Neutral |
| testneg_8603 | 0.22 | -2.66 | 7.225361 | -3 | 2.195109 | -3 | 0.19898 | 0.15588 | 0.43502 | 0.14712 | 11.875 | 0.537526145 | 67 | Disease_causing |
| testneg_8604 | -0.18 | -0.16 | 1.369414 | -3 | 4.245 | -3 | 0.13997 | 0.19598 | 0.33057 | 0.10549 | 13.16376 | 0.260260417 | 0.6 | Neutral |
| testneg_8605 | 0 | -0.01 | 1.077123 | -3 | 5.486225 | -3 | 0.18485 | 0.40044 | 0.10963 | 0.12788 | 16.8018 | 0.19619906 | 0 | Neutral |
| testneg_8606 | 0.09 | -1.07 | 1.734212 | -3 | -0.601278 | -3 | 0.18502 | 0.21329 | 0.03466 | 0.19518 | 28.03118 | 0.2 | 0 | Neutral |
| testneg_8607 | 0.04 | -0.23 | 3.709266 | -3 | 0.010391 | -3 | 0.08067 | 0.34914 | 0.03558 | 0.05202 | 58.95202 | 0.150545906 | 0 | Neutral |
| testneg_8608 | 0.08 | 0.03 | -2.363283 | -3 | 3.77321 | -3 | 0.05846 | 0.12788 | 0.33528 | 0.29463 | 22.76045 | 0.319599206 | 3.4 | Neutral |
| testneg_8609 | 0 | 0 | 0.509246 | -3 | 0.892238 | -3 | 0.08714 | 0.08945 | 0.20655 | 0.15317 | 18.36272 | 0.030260135 | 0 | Neutral |
| testneg_8610 | -0.05 | -0.29 | 0.531209 | -3 | -1.844931 | -3 | 0.10774 | 0.13117 | 0.12957 | 0.12788 | 10.28409 | 0.02 | 0 | Neutral |
| testneg_8611 | -0.15 | 0.28 | -1.223654 | -3 | 3.29285 | -3 | 0.09353 | 0.12788 | 0.42214 | 0.21008 | 26.34995 | 0.278452381 | 0.6 | Neutral |
| testneg_8612 | 0.14 | -0.37 | 4.741413 | -3 | 0.879763 | -3 | 0.21115 | 0.17723 | 0.68569 | 0.12463 | 16.07106 | 0.097928571 | 0 | Neutral |
| testneg_8613 | -0.16 | -0.08 | 2.356205 | -3 | 5.420665 | -3 | 0.13023 | 0.29692 | 0.50303 | 0.23508 | 5 | 0.777151757 | 100 | Disease_causing |
| testneg_8614 | -0.03 | 0.85 | 1.432399 | -3 | 1.624378 | -3 | 0.93284 | 0.71489 | 0.2016 | 0.13956 | 11.25 | 0.148345645 | 0 | Neutral |
| testneg_8615 | 0.07 | 0.7 | 5.31634 | -3 | 3.066208 | -3 | 0.11261 | 0.14664 | 0.2049 | 0.12678 | 13.51351 | 0.34 | 7.2 | Neutral |
| testneg_8616 | 0.02 | 0.13 | 1.091299 | -3 | 0.185181 | -3 | 0.1163 | 0.20863 | 0.82156 | 0.14621 | 10.41667 | 0.100799729 | 0 | Neutral |
| testneg_8617 | 0.04 | -0.23 | 2.208687 | -3 | 1.477151 | -3 | 0.19898 | 0.12788 | 0.04515 | 0.14318 | 27.9624 | 0.07 | 0 | Neutral |
| testneg_8618 | -0.02 | -0.45 | 1.618588 | -3 | 5.263161 | -3 | 0.14378 | 0.41915 | 0.96377 | 0.1167 | 13.08333 | 0.249087302 | 0.2 | Neutral |
| testneg_8619 | -0.07 | -0.38 | 2.424425 | -3 | 3.781577 | -3 | 0.54211 | 0.12592 | 0.19821 | 0.20954 | 19.44444 | 0.369069043 | 8.2 | Neutral |
| testneg_8620 | 0.02 | 0.45 | 5.906217 | -3 | -0.248361 | -3 | 0.72163 | 0.10596 | 0.66381 | 0.13039 | 10.83333 | 0.21 | 0.2 | Neutral |
| testneg_8621 | 0 | 0 | 0.884796 | -3 | -0.094519 | -1.231064 | 0.26337 | 0.12771 | 0.0774 | 0.1702 | 17.34649 | 0.07 | 0 | Neutral |
| testneg_8622 | -0.08 | -0.03 | 1.88068 | -3 | 0.759801 | -3 | 0.40803 | 0.47186 | 0.19898 | 0.12788 | 15.83802 | 0.03144905 | 0 | Neutral |
| testneg_8623 | 0 | -0.01 | 1.132093 | -3 | 6.617346 | -3 | 0.26372 | 0.12788 | 0.30275 | 0.12788 | 14.92465 | 0.13 | 0 | Neutral |
| testneg_8624 | -0.08 | -0.03 | 1.588775 | -3 | -1.628421 | -1.628421 | 0.24 | 0.2063 | 0.54441 | 0.08891 | 12.70833 | 0.04 | 0 | Neutral |
| testneg_8625 | 0.05 | 0.29 | 0.668985 | -1.085568 | 0.105498 | -3 | 0.22901 | 0.12788 | 0.09918 | 0.11061 | 22.50437 | 0.010101634 | 0 | Neutral |
| testneg_8626 | -0.16 | 1.37 | 4.782612 | -3 | -1.280796 | -3 | 0.1475 | 0.4323 | 0.06743 | 0.20381 | 10.41667 | 0.440039683 | 29.6 | Neutral |
| testneg_8627 | 0.21 | 0.15 | 3.540195 | -3 | 1.27566 | -3 | 0.20655 | 0.15317 | 0.18535 | 0.12788 | 22.29193 | 0.078970659 | 0 | Neutral |
| testneg_8628 | -0.03 | 1.03 | 3.933884 | -3 | 1.808824 | 1.808824 | 0.08684 | 0.08917 | 0.2445 | 0.53069 | 16.8018 | 0.289976016 | 0.6 | Neutral |
| testneg_8629 | 0 | 0 | 0.223823 | -3 | -0.905147 | -3 | 0.32484 | 0.11621 | 0.13169 | 0.06895 | 12.22032 | 0 | 0 | Neutral |
| testneg_8630 | 0.05 | -0.81 | 4.628168 | -3 | 7.068053 | -3 | 0.16379 | 0.31462 | 0.82156 | 0.14621 | 10 | 0.775136986 | 99.8 | Disease_causing |
| testneg_8631 | -0.09 | 0.3 | -2.446934 | -3 | 1.712042 | -3 | 0.46664 | 0.09008 | 0.31004 | 0.09727 | 19.44444 | 0.04 | 0 | Neutral |
| testneg_8632 | 0.16 | -0.37 | 5.864236 | -3 | -0.488738 | -3 | 0.23939 | 0.25284 | 0.46142 | 0.11057 | 22.94358 | 0.210884615 | 0.4 | Neutral |
| testneg_8633 | 0 | 0 | 2.064931 | -3 | -0.157625 | -1.308597 | 0.30122 | 0.09165 | 0.14668 | 0.12788 | 14.09794 | 0.001319244 | 0 | Neutral |
| testneg_8634 | 0 | 0.32 | 0.66894 | -3 | 4.106182 | -3 | 0.81212 | 0.14915 | 0.06394 | 0.24464 | 18.33333 | 0.332160973 | 5.2 | Neutral |
| testneg_8635 | 0 | -0.01 | 2.187535 | -3 | 1.628587 | -3 | 0.93654 | 0.38765 | 0.67644 | 0.11323 | 20.24133 | 0.04 | 0 | Neutral |
| testneg_8636 | 0.07 | 0.7 | 5.311585 | -3 | 1.214192 | -3 | 0.24026 | 0.12788 | 0.19898 | 0.12788 | 22.61823 | 0.12 | 0 | Neutral |
| testneg_8637 | 0.07 | 0.7 | 3.275701 | -3 | 4.106182 | -3 | 0.19102 | 0.12788 | 0.06394 | 0.24464 | 16.8018 | 0.475303609 | 45.8 | Neutral |
| testneg_8638 | 0 | -0.01 | 6.151767 | -3 | 0.890388 | -3 | 0.08002 | 0.08792 | 0.21138 | 0.11067 | 31.5768 | 0.13 | 0 | Neutral |
| testneg_8639 | -0.14 | 0.69 | 4.542574 | -3 | 0.083451 | -3 | 0.11827 | 0.23034 | 0.15994 | 0.11186 | 9.84463 | 0.17 | 0 | Neutral |
| testneg_8640 | 0 | 0 | 0.573548 | -3 | 1.594121 | -3 | 0.99267 | 0.1069 | 0.09259 | 0.11165 | 10.20833 | 0.02 | 0 | Neutral |
| testneg_8641 | 0 | 0 | 1.031973 | -3 | 1.280927 | -3 | 0.18256 | 0.13499 | 0.47217 | 0.15544 | 10.83333 | 0.04 | 0 | Neutral |
| testneg_8642 | -0.16 | -0.08 | 2.129082 | -3 | -0.545857 | -3 | 0.38719 | 0.15548 | 0.12488 | 0.12788 | 28.9372 | 0.08 | 0 | Neutral |
| testneg_8643 | 0.02 | 0.45 | 1.791247 | -3 | 1.200776 | -3 | 0.39193 | 0.11262 | 0.25092 | 0.10843 | 5.70654 | 0.01 | 0 | Neutral |
| testneg_8644 | 0.07 | 0.7 | 4.781408 | -3 | -1.97127 | -3 | 0.19898 | 0.12788 | 0.1211 | 0.1526 | 28.75878 | 0.190954887 | 0 | Neutral |
| testneg_8645 | 0 | 0 | 0.161519 | -3 | 1.225871 | -3 | 0.16544 | 0.1313 | 0.07571 | 0.09432 | 18.33333 | 0.01 | 0 | Neutral |
| testneg_8646 | -0.07 | -0.33 | 2.723699 | -3 | 2.507265 | -3 | 0.19898 | 0.12788 | 0.98581 | 0.27035 | 35.26843 | 0.196798845 | 0 | Neutral |
| testneg_8647 | 0.05 | -0.81 | 3.374642 | -3 | 2.750883 | -3 | 0.33264 | 0.10493 | 0.19898 | 0.12788 | 21.59547 | 0.081259615 | 0 | Neutral |
| testneg_8648 | -0.21 | -0.15 | -2.496516 | -3 | 6.490439 | -3 | 0.40076 | 0.11341 | 0.02498 | 0.08483 | 18.86179 | 0.244011164 | 0.2 | Neutral |
| testneg_8649 | 0 | 0 | 0.100811 | -3 | 3.391485 | -3 | 0.23027 | 0.37171 | 0.86306 | 0.10962 | 10.625 | 0.089757232 | 0 | Neutral |
| testneg_8650 | 0.09 | -0.3 | 3.302206 | -3 | 2.261013 | -3 | 0.28156 | 0.11579 | 0.19237 | 0.53191 | 9.5922 | 0.280958213 | 2 | Neutral |
| testneg_8651 | 0 | -0.01 | 0.505705 | -3 | 1.326802 | -3 | 0.73395 | 0.1705 | 0.05183 | 0.18779 | 23.37245 | 0.114844444 | 0 | Neutral |
| testneg_8652 | -0.04 | -2.01 | 5.42516 | -3 | 0.433941 | 0.433941 | 0.2141 | 0.19929 | 0.30475 | 0.21064 | 5.73716 | 0.541875 | 69.6 | Disease_causing |
| testneg_8653 | -0.0073485 | -0.0268939 | 1.974616 | -3 | 2.157695 | -3 | 0.30951 | 0.12788 | 0.05216 | 0.23851 | 6.00877 | 0.1 | 0 | Neutral |
| testneg_8654 | -0.17 | -0.38 | -0.92068 | -3 | 0.714207 | -3 | 0.28038 | 0.11814 | 0.04346 | 0.10785 | 16.42913 | 0.040231407 | 0 | Neutral |
| testneg_8655 | -0.09 | 0.25 | 3.307948 | -3 | -1.428127 | -3 | 0.06789 | 0.09707 | 0.11564 | 0.1184 | 14.01736 | 0.05 | 0 | Neutral |
| testneg_8656 | 0.03 | 0.12 | 1.304269 | -3 | -0.022585 | -3 | 0.45029 | 0.12788 | 0.2543 | 0.12929 | 12.77665 | 0.01 | 0 | Neutral |
| testneg_8657 | 0.03 | -1.03 | 1.335597 | -3 | 6.334007 | -3 | 0.11687 | 0.09069 | 0.19861 | 0.12057 | 29.40012 | 0.126978022 | 0 | Neutral |
| testneg_8658 | 0.07 | 0.38 | -0.902914 | -3 | 6.356297 | -3 | 0.40398 | 0.12788 | 0.24966 | 0.12788 | 27.82237 | 0.148845238 | 0 | Neutral |
| testneg_8659 | 0 | -0.01 | 2.507328 | -3 | 0.88775 | 0.88775 | 0.22364 | 0.12788 | 0.32649 | 0.15357 | 10.22898 | 0.044745329 | 0 | Neutral |
| testneg_8660 | -0.05 | -0.29 | 1.952213 | -3 | 0.278089 | -3 | 0.49504 | 0.08191 | 0.22233 | 0.31758 | 18.33333 | 0.070729076 | 0 | Neutral |
| testneg_8661 | -0.01 | -0.79 | 1.345014 | -3 | -0.249221 | -3 | 0.3437 | 0.5169 | 0.31651 | 0.12788 | 9.74181 | 0.036952381 | 0 | Neutral |

| | | | | | | | | | | | | | | |
|---|---|---|---|---|---|---|---|---|---|---|---|---|---|---|
| testneg_8662 | 0.18 | -1.05 | 4.092733 | -3 | 5.229818 | -3 | 0.08283 | 0.11797 | 0.0961 | 0.09786 | 20.46784 | 0.21 | 0.2 | Neutral |
| testneg_8663 | 0.16 | 0.08 | 1.755566 | -3 | 1.806835 | -3 | 0.161 | 0.11597 | 0.44488 | 0.31694 | 16.84579 | 0.085377778 | 0 | Neutral |
| testneg_8664 | 0.04 | -0.23 | -0.186131 | -3 | 1.311291 | -3 | 0.11802 | 0.09304 | 0.07577 | 0.09128 | 20 | 0.02 | 0 | Neutral |
| testneg_8665 | -0.15 | -0.04 | -1.165102 | -3 | -0.168925 | -3 | 0.37237 | 0.36829 | 0.19898 | 0.12788 | 16.8018 | 0.08 | 0 | Neutral |
| testneg_8666 | 0 | 0 | 2.403686 | 2.403686 | 2.410627 | -3 | 0.48574 | 0.20473 | 0.07275 | 0.12788 | 30.0903 | 0.18 | 0.2 | Neutral |
| testneg_8667 | 0.04 | -0.23 | 1.305896 | -3 | 1.300657 | -3 | 0.1871 | 0.10907 | 0.12317 | 0.12345 | 19.44444 | 0.01 | 0 | Neutral |
| testneg_8668 | 0 | 0 | 0.100154 | -3 | -0.676534 | -0.676534 | 0.21122 | 0.12686 | 0.16262 | 0.15786 | 18.33333 | 0.051621148 | 0 | Neutral |
| testneg_8669 | 0.02 | -0.63 | 0.539564 | -3 | -0.113262 | -3 | 0.85759 | 0.1031 | 0.56154 | 0.0939 | 27.00364 | 0.02 | 0 | Neutral |
| testneg_8670 | 0 | 0 | -0.172086 | -3 | 0.063137 | -3 | 0.12325 | 0.09383 | 0.24058 | 0.14928 | 21.38889 | 0.05 | 0 | Neutral |
| testneg_8671 | 0.02 | 0.13 | 7.041902 | -3 | 1.854082 | -3 | 0.34467 | 0.13076 | 0.20084 | 0.0851 | 35.84737 | 0.308690476 | 3.8 | Neutral |
| testneg_8672 | 0.07 | 0.38 | 2.530625 | -3 | 1.871099 | -3 | 0.2852 | 0.12788 | 0.87611 | 0.16759 | 10.19608 | 0.06 | 0 | Neutral |
| testneg_8673 | 0 | 0 | 0.988418 | -3 | -0.303489 | -3 | 0.1004 | 0.11409 | 0.508 | 0.1281 | 22.01754 | 0.01 | 0 | Neutral |
| testneg_8674 | 0.01 | 0.8 | 6.431918 | -3 | 3.510198 | -3 | 0.19364 | 0.14027 | 0.2521 | 0.1784 | 19.09347 | 0.722105556 | 99.2 | Disease_causing |
| testneg_8675 | 0.08 | 0.03 | 1.082238 | -3 | 2.640218 | -3 | 0.27062 | 0.12378 | 0.16621 | 0.49845 | 14.52969 | 0.160805892 | 0 | Neutral |
| testneg_8676 | -0.01 | -0.8 | 4.547515 | -3 | 1.288901 | -3 | 0.20491 | 0.22087 | 0.19898 | 0.20319 | 10.83333 | 0.386853535 | 16.4 | Neutral |
| testneg_8677 | -0.0105455 | 0.10618182 | 13.949479 | -0.065016 | 5.399087 | -3 | 0.38695 | 0.12579 | 0.1594 | 0.22002 | 17.14941 | 0.797416667 | 100 | Disease_causing |
| testneg_8678 | 0.06 | 0.18 | -1.19222 | -3 | 3.256775 | -3 | 0.02971 | 0.33983 | 0.16309 | 0.12788 | 48.69853 | 0.22 | 0 | Neutral |
| testneg_8679 | 0.15 | -0.28 | -0.437316 | -3 | 6.678547 | 0.069349 | 0.20231 | 0.12788 | 0.19898 | 0.12788 | 22.25953 | 0.182028571 | 0.2 | Neutral |
| testneg_8680 | -0.05 | -1.52 | 4.861941 | -3 | 0.66455 | -3 | 0.25195 | 0.09584 | 0.18067 | 0.14759 | 12.12084 | 0.19 | 0 | Neutral |
| testneg_8681 | 0 | 0 | -0.061795 | -3 | 6.571813 | 6.279292 | 0.34698 | 0.12788 | 0.09488 | 0.10426 | 18.33333 | 0.443214286 | 30.4 | Neutral |
| testneg_8682 | 0.03 | 0.12 | 6.692068 | -3 | 6.094021 | -3 | 0.20616 | 0.29531 | 0.20502 | 0.12788 | 18.88889 | 0.738439993 | 98.6 | Disease_causing |
| testneg_8683 | 0.03 | 0.1 | 0.713336 | -3 | 2.639631 | -3 | 0.25009 | 0.11017 | 0.25151 | 0.12788 | 14.58384 | 0.02 | 0 | Neutral |
| testneg_8684 | 0 | 0 | -0.47975 | -3 | 1.986841 | 1.986841 | 0.4719 | 0.33344 | 0.19898 | 0.12788 | 17.53755 | 0.06 | 0 | Neutral |
| testneg_8685 | 0.01 | -0.57 | 5.075109 | -3 | 4.74441 | -3 | 0.68863 | 0.09663 | 0.42306 | 0.86891 | 22.29927 | 0.61846745 | 89.4 | Disease_causing |
| testneg_8686 | -0.02 | 0.63 | 4.675931 | -3 | 1.80664 | -3 | 0.961 | 0.25991 | 0.23385 | 0.12788 | 10.23791 | 0.15 | 0 | Neutral |
| testneg_8687 | 0.03 | 0.1 | 0.500506 | -3 | -0.573377 | -3 | 0.68233 | 0.58998 | 0.42366 | 0.14372 | 10.41667 | 0.110663383 | 0 | Neutral |
| testneg_8688 | 0 | 0 | 2.004172 | -3 | -1.367472 | -3 | 0.18175 | 0.13527 | 0.049 | 0.11324 | 48.62495 | 0.02 | 0 | Neutral |
| testneg_8689 | 0.04 | -0.23 | 4.680114 | -3 | 1.315669 | -3 | 0.31124 | 0.08875 | 0.13331 | 0.09729 | 18.33333 | 0.04 | 0 | Neutral |
| testneg_8690 | 0.07 | 0.01 | 2.947718 | -3 | -0.072198 | -0.072198 | 0.5068 | 0.12276 | 0.1025 | 0.3268 | 4.44444 | 0.186727354 | 0 | Neutral |
| testneg_8691 | 0.03 | 0.12 | 2.323415 | -3 | 1.516949 | -3 | 0.03942 | 0.12788 | 0.87202 | 0.42207 | 14.87554 | 0.11 | 0 | Neutral |
| testneg_8692 | -0.03 | -0.1 | -0.369846 | -3 | 1.161725 | -3 | 0.33467 | 0.12788 | 0.35486 | 0.11584 | 18.94894 | 0.01 | 0 | Neutral |
| testneg_8693 | 0 | 0 | -0.402189 | -3 | -0.551863 | -3 | 0.22155 | 0.1062 | 0.51037 | 0.10999 | 20.89606 | 0.000837456 | 0 | Neutral |
| testneg_8694 | 0.07 | 0.33 | 2.323649 | -3 | 5.713281 | 1.51257 | 0.10822 | 0.10942 | 0.28066 | 0.18241 | 25.48416 | 0.428676097 | 25.6 | Neutral |
| testneg_8695 | 0 | 0 | 1.840966 | -3 | 0.157379 | -3 | 0.85793 | 0.80177 | 0.45997 | 0.15162 | 19.44444 | 0.122100188 | 0 | Neutral |
| testneg_8696 | -0.16 | -0.08 | 1.439827 | -3 | 3.54624 | -3 | 0.09577 | 0.2436 | 0.05476 | 0.12788 | 31.16278 | 0.235833333 | 0.4 | Neutral |
| testneg_8697 | 0 | -0.32 | 1.047467 | -3 | 0.473397 | -3 | 0.26972 | 0.12484 | 0.07064 | 0.11699 | 12.64837 | 0.000223471 | 0 | Neutral |
| testneg_8698 | -0.16 | 1.37 | 3.207028 | -3 | -0.743188 | -0.743188 | 0.03169 | 0.10615 | 0.19898 | 0.535 | 14.25869 | 0.27 | 0 | Neutral |
| testneg_8699 | -0.03 | -0.1 | 0.314664 | -3 | 5.270445 | -3 | 0.08224 | 0.29179 | 0.25179 | 0.09355 | 8.97898 | 0.23 | 0.4 | Neutral |
| testneg_8700 | 0 | 0 | 2.10021 | -3 | 0.775573 | -3 | 0.15526 | 0.08511 | 0.13325 | 0.12788 | 30.01545 | 0.004697802 | 0 | Neutral |
| testneg_8701 | 0 | 0 | 0.598467 | -3 | 5.149802 | -3 | 0.52836 | 0.12785 | 0.06376 | 0.0935 | 27.42689 | 0.092130952 | 0 | Neutral |
| testneg_8702 | 0.02 | -0.36 | 2.56113 | -3 | 1.671747 | -3 | 0.06304 | 0.12788 | 0.12643 | 0.12266 | 16.42913 | 0.02 | 0 | Neutral |
| testneg_8703 | 0 | 0 | 1.172534 | -3 | 0.730901 | -3 | 0.04056 | 0.55024 | 0.18705 | 0.43522 | 4.72222 | 0.213136992 | 0 | Neutral |
| testneg_8704 | -0.03 | -0.1 | 6.981338 | -3 | -0.10563 | -3 | 0.52278 | 0.41207 | 0.09028 | 0.10643 | 17.5501 | 0.4 | 18 | Neutral |
| testneg_8705 | 0 | 0 | 0.432603 | -3 | 3.864374 | -3 | 0.34546 | 0.0932 | 0.14021 | 0.43531 | 10.41667 | 0.209772241 | 0 | Neutral |
| testneg_8706 | -0.07 | -0.7 | 1.84182 | -3 | 4.926965 | -3 | 0.53239 | 0.17509 | 0.16286 | 0.15786 | 5.75579 | 0.511458453 | 59 | Disease_causing |
| testneg_8707 | -0.02 | 0.63 | 1.217652 | -3 | -0.597203 | -3 | 0.19898 | 0.12788 | 0.0925 | 0.08222 | 26.80135 | 0.010091326 | 0 | Neutral |
| testneg_8708 | 0 | 0 | 0.606984 | -3 | -0.41063 | -3 | 0.08932 | 0.12788 | 0.09611 | 0.07772 | 16.8018 | 0.000412415 | 0 | Neutral |
| testneg_8709 | -0.08 | -0.03 | -0.070402 | -3 | 3.319567 | 3.146686 | 0.0513 | 0.33622 | 0.0634 | 0.0824 | 11.25 | 0.28 | 0.2 | Neutral |
| testneg_8710 | -0.02 | -0.45 | 3.26904 | -3 | -0.230704 | -3 | 0.18554 | 0.12299 | 0.16361 | 0.13388 | 32.0259 | 0.1 | 0 | Neutral |
| testneg_8711 | -0.02 | 0.63 | 5.119073 | -3 | 3.17078 | -3 | 0.29679 | 0.12788 | 0.17818 | 0.15728 | 17.26246 | 0.494249903 | 52.8 | Disease_causing |
| testneg_8712 | -0.09 | 0.67 | 0.082232 | -3 | 1.495199 | -3 | 0.08475 | 0.19889 | 0.33049 | 0.12788 | 19.86666 | 0.08 | 0 | Neutral |
| testneg_8713 | -0.08 | -0.03 | 0.390383 | -3 | 0.994423 | -3 | 0.08213 | 0.08913 | 0.1072 | 0.63446 | 19.09347 | 0.11 | 0 | Neutral |
| testneg_8714 | -0.06 | 0.57 | 1.272946 | -3 | -0.221228 | -3 | 0.12977 | 0.1128 | 0.11153 | 0.10767 | 20.997 | 0.02 | 0 | Neutral |
| testneg_8715 | -0.18 | 1.05 | 1.43145 | -3 | 0.175778 | -3 | 0.2477 | 0.11928 | 0.19898 | 0.12788 | 27.91246 | 0.08 | 0 | Neutral |
| testneg_8716 | 0 | 0 | 1.897319 | -3 | 0.734765 | -3 | 0.39225 | 0.10625 | 0.07204 | 0.07708 | 16.0119 | 0 | 0 | Neutral |
| testneg_8717 | 0 | -0.01 | 1.221125 | -3 | 6.562934 | -3 | 0.36793 | 0.15588 | 0.12965 | 0.75487 | 12.70833 | 0.515199859 | 60 | Disease_causing |
| testneg_8718 | 0.01 | 0.79 | 1.560893 | 1.560893 | 2.151559 | -3 | 0.16524 | 0.12788 | 0.82591 | 0.12207 | 16.8018 | 0.06 | 0 | Neutral |
| testneg_8719 | 0.05 | -0.81 | 1.595101 | -3 | 5.471207 | -3 | 0.13156 | 0.10621 | 0.19821 | 0.20954 | 19.44444 | 0.35921303 | 9.4 | Neutral |
| testneg_8720 | -0.02 | -0.45 | 1.632161 | -3 | 3.206305 | -3 | 0.15107 | 0.10457 | 0.38503 | 0.11286 | 12.51462 | 0.033252976 | 0 | Neutral |
| testneg_8721 | -0.2 | 1.26 | -0.70557 | -3 | 1.135324 | -3 | 0.29008 | 0.19268 | 0.79692 | 0.51979 | 7.01832 | 0.214852941 | 0 | Neutral |
| testneg_8722 | 0 | 0 | 0.501525 | -3 | -1.172819 | -3 | 0.20457 | 0.12378 | 0.31238 | 0.14393 | 32.25789 | 0.032677418 | 0 | Neutral |
| testneg_8723 | 0.15 | -0.28 | 2.58225 | 1.012728 | 2.821548 | -3 | 0.19883 | 0.09434 | 0.18578 | 0.27854 | 16.25133 | 0.3138 | 2.4 | Neutral |
| testneg_8724 | 0 | -0.32 | -0.761734 | -3 | -2.720928 | -3 | 0.2578 | 0.09796 | 0.55367 | 0.57993 | 6.00877 | 0.060408333 | 0 | Neutral |
| testneg_8725 | 0 | 0 | 2.193905 | -3 | 1.885526 | -3 | 0.16952 | 0.10613 | 0.11233 | 0.28224 | 15.20594 | 0.059843623 | 0 | Neutral |
| testneg_8726 | 0 | 0 | 0.560537 | -3 | 0.010135 | -3 | 0.32804 | 0.10857 | 0.12951 | 0.08688 | 22.10244 | 0.00097619 | 0 | Neutral |
| testneg_8727 | -0.05 | -0.29 | 2.605779 | -3 | 6.390861 | -3 | 0.22817 | 0.10634 | 0.87877 | 0.60823 | 10.41667 | 0.518150686 | 61.4 | Disease_causing |
| testneg_8728 | 0 | -0.01 | 0.306573 | -3 | 5.969277 | -3 | 0.03898 | 0.09943 | 0.31067 | 0.35414 | 6.28502 | 0.320580297 | 6.6 | Neutral |
| testneg_8729 | 0.02 | 0.45 | -0.784242 | -3 | 0.293949 | -3 | 0.18153 | 0.10845 | 0.16616 | 0.11073 | 19.44444 | 0.01 | 0 | Neutral |
| testneg_8730 | 0 | 0 | 1.9846 | -3 | 0.552815 | -3 | 0.64701 | 0.94571 | 0.20417 | 0.10681 | 12.70833 | 0.06 | 0 | Neutral |
| testneg_8731 | -0.02 | -0.13 | -2.297489 | -3 | 6.394805 | -3 | 0.21236 | 0.12788 | 0.10285 | 0.10186 | 18.88889 | 0.186190476 | 0.2 | Neutral |
| testneg_8732 | 0 | 0 | 0.730138 | -3 | -0.457394 | -3 | 0.16952 | 0.10613 | 0.19898 | 0.46867 | 21.61491 | 0.06 | 0 | Neutral |
| testneg_8733 | 0 | 0 | 1.006031 | -3 | 7.191968 | -3 | 0.78537 | 0.33817 | 0.07665 | 0.31408 | 14.04625 | 0.473472222 | 45.8 | Neutral |
| testneg_8734 | 0.05 | -0.81 | 2.699324 | -3 | 0.212096 | -3 | 0.10723 | 0.11163 | 0.88456 | 0.11674 | 19.21569 | 0.020446429 | 0 | Neutral |
| testneg_8735 | 0.22 | -2.66 | 6.203124 | -3 | 1.579942 | -3 | 0.10581 | 0.11631 | 0.15767 | 0.12788 | 13.51351 | 0.16875 | 0.2 | Neutral |
| testneg_8736 | 0.09 | -1.07 | 0.484682 | -3 | 0.861976 | -3 | 0.19898 | 0.12788 | 0.81659 | 0.10424 | 17.44991 | 0.03 | 0 | Neutral |
| testneg_8737 | 0 | 0 | 1.707083 | -3 | 3.155825 | -3 | 0.90296 | 0.28521 | 0.13708 | 0.81545 | 24.41367 | 0.308768942 | 3.2 | Neutral |
| testneg_8738 | 0.02 | 0.13 | 5.790455 | -3 | 6.984226 | -3 | 0.10895 | 0.10442 | 0.87877 | 0.60823 | 4.44444 | 0.719166667 | 99.4 | Disease_causing |
| testneg_8739 | 0.22 | -2.66 | 5.950905 | -3 | 0.452231 | -3 | 0.19898 | 0.12788 | 0.46691 | 0.07886 | 18.61111 | 0.15 | 0 | Neutral |
| testneg_8740 | -0.15 | -0.06 | -1.712174 | -3 | 2.542983 | -3 | 0.07604 | 0.35375 | 0.19898 | 0.30319 | 16.8018 | 0.420104167 | 22.2 | Neutral |
| testneg_8741 | -0.14 | 1.01 | 2.036499 | -3 | -0.78092 | -3 | 0.43897 | 0.12441 | 0.14432 | 0.22579 | 7.56982 | 0.091780436 | 0 | Neutral |
| testneg_8742 | 0.00583333 | -0.0200758 | -1.007569 | -3 | -2.103676 | -3 | 0.19898 | 0.12788 | 0.23975 | 0.13861 | 24.13209 | 0.050715198 | 0 | Neutral |
| testneg_8743 | -0.05 | -0.29 | -1.375282 | -3 | -0.053274 | -3 | 0.19898 | 0.12788 | 0.31621 | 0.12788 | 19.44444 | 0.01 | 0 | Neutral |
| testneg_8744 | 0.16 | -0.37 | 2.851691 | -3 | 1.372699 | -3 | 0.21623 | 0.42074 | 0.06257 | 0.12788 | 10.41667 | 0.099772727 | 0 | Neutral |
| testneg_8745 | 0 | 0 | 0.347529 | -3 | -0.494899 | -3 | 0.22443 | 0.10657 | 0.1834 | 0.12788 | 25.38961 | 0.000192379 | 0 | Neutral |
| testneg_8746 | -0.05 | -0.29 | 0.110557 | -3 | -0.557277 | -3 | 0.11703 | 0.09364 | 0.559 | 0.2286 | 11.54145 | 0.051652098 | 0 | Neutral |
| testneg_8747 | -0.05 | -0.29 | -2.677475 | -3 | -0.416655 | -0.416655 | 0.03664 | 0.06179 | 0.05453 | 0.12788 | 65.41447 | 0.12 | 0 | Neutral |
| testneg_8748 | 0.09 | -0.3 | 6.314583 | -3 | -0.03109 | -3 | 0.42583 | 0.10901 | 0.0626 | 0.1393 | 19.51014 | 0.259065934 | 1.2 | Neutral |
| testneg_8749 | 0.09 | -0.3 | 3.200815 | -3 | 1.091417 | -3 | 0.98312 | 0.12788 | 0.06617 | 0.12788 | 12.42141 | 0.05 | 0 | Neutral |
| testneg_8750 | 0.05 | 0.29 | -1.171244 | -1.171244 | 0.634938 | -3 | 0.19898 | 0.12788 | 0.14458 | 0.14062 | 13.51351 | 0.04679261 | 0 | Neutral |
| testneg_8751 | -0.11 | 0.19 | 5.217851 | -3 | 0.604056 | -3 | 0.34295 | 0.12788 | 0.75906 | 0.09702 | 10.46895 | 0.131396104 | 0 | Neutral |
| testneg_8752 | 0.03 | -1.03 | 1.284996 | -3 | 1.568815 | -3 | 0.52028 | 0.47239 | 0.14379 | 0.12788 | 18.33333 | 0.07 | 0 | Neutral |
| testneg_8753 | 0 | 0 | 0.310149 | -3 | -0.451132 | -3 | 0.0626 | 0.5373 | 0.12048 | 0.2381 | 21.49912 | 0.279885135 | 0.2 | Neutral |
| testneg_8754 | 0.04 | -0.23 | 7.79718 | -3 | 3.761879 | -3 | 0.21814 | 0.13855 | 0.58985 | 0.10504 | 19.44444 | 0.537408043 | 67 | Disease_causing |
| testneg_8755 | 0 | -0.01 | 0.736606 | -3 | 4.681701 | 4.681701 | 0.27187 | 0.41985 | 0.10571 | 0.12788 | 24.93861 | 0.389058442 | 15.8 | Neutral |
| testneg_8756 | 0 | 0 | 2.263609 | -3 | 6.226262 | -3 | 0.4811 | 0.13398 | 0.19898 | 0.12788 | 18.6768 | 0.16644697 | 0 | Neutral |
| testneg_8757 | 0 | 0 | 0.840063 | -3 | 4.623974 | -3 | 0.08683 | 0.26961 | 0.09866 | 0.104 | 19.86666 | 0.15927305 | 0 | Neutral |
| testneg_8758 | 0.2 | -1.26 | -1.784808 | -3 | 3.467861 | -3 | 0.11133 | 0.12788 | 0.37293 | 0.25159 | 20.22479 | 0.290192308 | 1 | Neutral |
| testneg_8759 | 0.01 | 0.8 | 4.304984 | -3 | 0.635023 | -3 | 0.19898 | 0.14645 | 0.12136 | 0.27707 | 45.00704 | 0.265758644 | 0.6 | Neutral |
| testneg_8760 | 0.07 | 0.38 | 1.107256 | -3 | 5.796244 | -3 | 0.3889 | 0.10831 | 0.0538 | 0.0823 | 13.98631 | 0.11 | 0 | Neutral |
| testneg_8761 | 0 | 0 | 0.777181 | -3 | -0.491414 | -3 | 0.19948 | 0.12788 | 0.61027 | 0.14229 | 28.24141 | 0.030690476 | 0 | Neutral |
| testneg_8762 | 0 | -0.32 | 0.562804 | -3 | 2.55089 | -3 | 0.11692 | 0.08324 | 0.08027 | 0.12788 | 17.21847 | 0.020435606 | 0 | Neutral |
| testneg_8763 | 0.01 | 0.8 | 0.586242 | -3 | 2.629087 | -3 | 0.559 | 0.2286 | 0.28162 | 0.12788 | 11.54145 | 0.113428571 | 0 | Neutral |
| testneg_8764 | -0.11 | 1.95 | 2.355101 | -3 | 0.811043 | -3 | 0.18312 | 0.12788 | 0.82148 | 0.14847 | 16.8018 | 0.13 | 0 | Neutral |
| testneg_8765 | 0.01 | 0.8 | -0.725111 | -0.725111 | 5.86661 | -3 | 0.05116 | 0.08996 | 0.15581 | 0.10782 | 23.70738 | 0.151180556 | 0 | Neutral |
| testneg_8766 | 0.14 | -0.37 | 0.864504 | -3 | 0.707625 | -3 | 0.67216 | 0.17808 | 0.07297 | 0.12788 | 11.15097 | 0.04 | 0 | Neutral |
| testneg_8767 | -0.02 | 0.68 | 5.845448 | 5.845448 | 6.638158 | -3 | 0.06926 | 0.08399 | 0.07995 | 0.12788 | 29.86822 | 0.64 | 95.8 | Disease_causing |
| testneg_8768 | 0 | 0 | 0.943238 | -3 | 0.552629 | -3 | 0.19898 | 0.58572 | 0.48615 | 0.14369 | 11.25 | 0.102019246 | 0 | Neutral |
| testneg_8769 | 0 | -0.01 | 1.315543 | 1.315543 | 0.148971 | -3 | 0.28358 | 0.38765 | 0.17007 | 0.12771 | 15 | 0.04 | 0 | Neutral |
| testneg_8770 | -0.03 | -0.12 | 1.627305 | -3 | -0.496606 | -3 | 0.08448 | 0.17914 | 0.57564 | 0.12788 | 10 | 0.05 | 0 | Neutral |
| testneg_8771 | 0 | 0 | 1.537848 | -3 | -0.664442 | -3 | 0.11391 | 0.12788 | 0.1592 | 0.12788 | 18.76159 | 0 | 0 | Neutral |
| testneg_8772 | 0 | 0 | 1.600017 | -3 | 1.873147 | 1.873147 | 0.07994 | 0.1101 | 0.1499 | 0.09632 | 13.08701 | 0.03 | 0 | Neutral |
| testneg_8773 | 0.07 | 0.7 | 6.215755 | -3 | 5.034335 | -3 | 0.10459 | 0.12783 | 0.06704 | 0.1068 | 23.68839 | 0.397997835 | 20.8 | Neutral |
| testneg_8774 | 0 | 0 | 0.777519 | -3 | -1.543536 | -1.543536 | 0.75071 | 0.63704 | 0.06744 | 0.12788 | 4.72222 | 0.011498663 | 0 | Neutral |
| testneg_8775 | 0.03072727 | -0.4116364 | 2.905355 | -3 | -0.285825 | -3 | 0.04305 | 0.06354 | 0.15016 | 0.08826 | 29.9024 | 0.06 | 0 | Neutral |
| testneg_8776 | 0.03 | 0.12 | 0.199359 | -3 | 0.194184 | -3 | 0.07626 | 0.12788 | 0.18744 | 0.09392 | 19.45844 | 0.01 | 0 | Neutral |
| testneg_8777 | 0 | 0 | 0.759983 | -3 | 0.784892 | -3 | 0.13505 | 0.09019 | 0.47192 | 0.11305 | 12.53918 | 8.00E-05 | 0 | Neutral |
| testneg_8778 | -0.01 | -0.79 | 2.141385 | -3 | 4.762824 | -3 | 0.1017 | 0.11809 | 0.32089 | 0.1035 | 17.18434 | 0.090615836 | 0 | Neutral |
| testneg_8779 | 0 | 0 | -1.581669 | -3 | -0.201778 | -3 | 0.68029 | 0.12788 | 0.02658 | 0.18737 | 10.41667 | 0.060450297 | 0 | Neutral |
| testneg_8780 | 0.12 | -1.16 | 4.936092 | -3 | 0.574992 | -3 | 0.2118 | 0.2009 | 0.13624 | 0.09458 | 17.77778 | 0.1 | 0 | Neutral |
| testneg_8781 | 0.22 | -2.66 | 7.86876 | -3 | 0.70994 | -0.230704 | 0.40632 | 0.1094 | 0.16361 | 0.13388 | 16.8018 | 0.32 | 4.2 | Neutral |
| testneg_8782 | -0.07 | -0.38 | 1.280741 | -3 | 0.590862 | -3 | 0.2852 | 0.12788 | 0.19898 | 0.20319 | 10.41667 | 0.04 | 0 | Neutral |
| testneg_8783 | 0.02 | -0.63 | 0.686794 | 0.665096 | 0.620269 | -3 | 0.15132 | 0.11724 | 0.1287 | 0.10423 | 20.15873 | 0.03 | 0 | Neutral |

| | | | | | | | | | | | | | |
|---|---|---|---|---|---|---|---|---|---|---|---|---|---|
| testneg_8784 | 0 | 0 | 0.864288 | -3 | 3.47892 | -3 | 0.21014 | 0.12378 | 0.18423 | 0.10808 | 19.41375 | 0.021760936 | 0 Neutral |
| testneg_8785 | -0.21 | -0.15 | -1.902903 | -3 | 0.082365 | -3 | 0.1254 | 0.08697 | 0.10011 | 0.08014 | 11.875 | 0.07 | 0 Neutral |
| testneg_8786 | 0.22 | -2.66 | 7.972239 | -3 | -0.673818 | -3 | 0.13099 | 0.17453 | 0.76138 | 0.16258 | 18.88889 | 0.45 | 36.6 Neutral |
| testneg_8787 | 0.04 | 2.01 | 3.781815 | -3 | 3.236399 | 0.826872 | 0.01704 | 0.05734 | 0.22953 | 0.0919 | 21.01455 | 0.2 | 0 Neutral |
| testneg_8788 | 0.21 | 0.15 | 3.945475 | -3 | -2.248026 | -3 | 0.27115 | 0.14766 | 0.25179 | 0.09355 | 19.09347 | 0.07 | 0 Neutral |
| testneg_8789 | 0 | 0 | 0.828595 | -3 | 1.417365 | 1.417365 | 0.93284 | 0.71489 | 0.61184 | 0.55597 | 4.72222 | 0.1 | 0 Neutral |
| testneg_8790 | 0 | 0 | 1.538591 | -3 | 3.319567 | -3 | 0.64302 | 0.11262 | 0.0634 | 0.0824 | 13.14653 | 0.046539584 | 0 Neutral |
| testneg_8791 | 0.07 | 0.7 | 5.661416 | -3 | 1.464526 | -3 | 0.9352 | 0.14229 | 0.12513 | 0.12171 | 19.72881 | 0.26 | 1.2 Neutral |
| testneg_8792 | 0.21 | 0.15 | 5.407311 | -3 | 2.077024 | -3 | 0.08975 | 0.07399 | 0.08973 | 0.07662 | 24.45963 | 0.13225 | 0 Neutral |
| testneg_8793 | -0.21 | -0.15 | -2.723139 | -3 | -0.131159 | -3 | 0.08186 | 0.11813 | 0.2218 | 0.11533 | 28.4009 | 0.07 | 0 Neutral |
| testneg_8794 | 0 | -0.01 | 0.130932 | -3 | 0.132481 | -3 | 0.19898 | 0.12788 | 0.33026 | 0.30576 | 23.92312 | 0.06 | 0 Neutral |
| testneg_8795 | 0.02 | -0.63 | 2.81231 | -3 | 0.230409 | 0.230409 | 0.06647 | 0.09129 | 0.16116 | 0.08314 | 11.92061 | 0.03 | 0 Neutral |
| testneg_8796 | 0 | 0 | 0.936162 | -3 | 0.205041 | -3 | 0.08999 | 0.0989 | 0.20084 | 0.0851 | 17.36842 | 0 | 0 Neutral |
| testneg_8797 | 0.01 | -0.57 | 2.700261 | -3 | -0.042671 | -3 | 0.76957 | 0.13588 | 0.20951 | 0.16278 | 10.41667 | 0.110358974 | 0 Neutral |
| testneg_8798 | 0 | 0 | 1.304345 | -3 | 3.157227 | -3 | 0.3889 | 0.10831 | 0.1977 | 0.09728 | 10.01954 | 0.028219842 | 0 Neutral |
| testneg_8799 | 0.01 | -0.57 | 2.700261 | -3 | -0.548012 | -3 | 0.76957 | 0.13588 | 0.10565 | 0.08739 | 10.41667 | 0.03 | 0 Neutral |
| testneg_8800 | 0 | 0 | 0.482222 | -3 | 2.251286 | -3 | 0.95557 | 0.38833 | 0.31125 | 0.20061 | 14.51389 | 0.112297129 | 0 Neutral |
| testneg_8801 | -0.02 | -0.13 | 3.206825 | -3 | 2.575019 | -3 | 0.55197 | 0.11305 | 0.28269 | 0.12505 | 21.38889 | 0.070378788 | 0 Neutral |
| testneg_8802 | -0.07 | -0.38 | 1.211461 | -3 | 2.227386 | -3 | 0.38552 | 0.14172 | 0.20047 | 0.11827 | 16.76768 | 0.011111111 | 0 Neutral |
| testneg_8803 | 0 | -0.01 | 0.636505 | -3 | 3.413304 | -3 | 0.84238 | 0.12559 | 0.31626 | 0.12936 | 17.44444 | 0.035351571 | 0 Neutral |
| testneg_8804 | -0.07 | -0.38 | 3.116757 | -3 | 5.950583 | -3 | 0.24949 | 0.29709 | 0.19898 | 0.12788 | 18.33333 | 0.415378552 | 25.8 Neutral |
| testneg_8805 | 0.04 | -0.23 | 2.208687 | -3 | 6.215755 | -3 | 0.19898 | 0.12788 | 0.10459 | 0.12783 | 23.68839 | 0.16 | 0 Neutral |
| testneg_8806 | -0.01 | -0.79 | -3.816017 | -3 | 3.513863 | -3 | 0.06095 | 0.08179 | 0.19024 | 0.1132 | 12.69679 | 0.073904762 | 0 Neutral |
| testneg_8807 | 0.12 | -1.14 | 1.528223 | 1.528223 | 3.93025 | -3 | 0.35957 | 0.11476 | 0.19898 | 0.08486 | 18.88889 | 0.122033745 | 0 Neutral |
| testneg_8808 | 0 | 0 | 1.511019 | 0.030464 | 9.272843 | -3 | 0.33795 | 0.17455 | 0.24544 | 0.0894 | 20.40047 | 0.4 | 18.6 Neutral |
| testneg_8809 | 0.14 | -1.01 | 3.224374 | -3 | 5.360167 | -3 | 0.44329 | 0.12788 | 0.09918 | 0.11061 | 45.81879 | 0.196333333 | 0.2 Neutral |
| testneg_8810 | 0.14 | -1.01 | -0.007811 | -3 | 7.116856 | -3 | 0.49629 | 0.11989 | 0.25831 | 0.10988 | 21.53791 | 0.235 | 0.4 Neutral |
| testneg_8811 | 0.1 | 1.9 | 5.687294 | -3 | 5.30476 | 5.30476 | 0.19898 | 0.12788 | 0.20547 | 0.12178 | 26.95652 | 0.557977238 | 78.2 Disease_causing |
| testneg_8812 | 0 | 0 | -0.961702 | -0.961702 | 2.752111 | -3 | 0.46979 | 0.1705 | 0.10556 | 0.28423 | 10.41667 | 0.186588584 | 0.2 Neutral |
| testneg_8813 | 0 | 0 | 1.504538 | -3 | 2.13556 | -3 | 0.19898 | 0.12788 | 0.14668 | 0.12788 | 13.32006 | 0 | 0 Neutral |
| testneg_8814 | 0.16 | -1.37 | 0.303661 | -3 | 4.334907 | -3 | 0.03788 | 0.10996 | 0.15263 | 0.12788 | 10 | 0.11 | 0 Neutral |
| testneg_8815 | 0.14 | -1.01 | 3.043761 | -3 | 0.998327 | -1.951993 | 0.08224 | 0.29179 | 0.19898 | 0.12788 | 8.97898 | 0.106488095 | 0 Neutral |
| testneg_8816 | -0.11 | 0.19 | 5.217851 | -3 | 3.381673 | 3.381673 | 0.34295 | 0.12788 | 0.23578 | 0.12788 | 9.75627 | 0.427685897 | 27.2 Neutral |
| testneg_8817 | -0.09 | 0.3 | 2.309209 | -3 | 3.233677 | -3 | 0.04863 | 0.12149 | 0.13356 | 0.09963 | 14.94243 | 0.110147059 | 0 Neutral |
| testneg_8818 | 0 | 0 | 0.990948 | -3 | 1.341806 | -3 | 0.31859 | 0.11262 | 0.50876 | 0.10038 | 18.50844 | 0 | 0 Neutral |
| testneg_8819 | 0.18 | 0.16 | 0.416167 | -3 | 2.238223 | -3 | 0.1224 | 0.11899 | 0.32606 | 0.10251 | 21.46358 | 0.03 | 0 Neutral |
| testneg_8820 | 0.01 | 0.8 | 2.947434 | -3 | 1.202078 | -3 | 0.27915 | 0.27706 | 0.16286 | 0.15786 | 10.50298 | 0.261438874 | 1 Neutral |
| testneg_8821 | 0.15 | -0.28 | -1.024835 | -3 | 1.999625 | -3 | 0.08628 | 0.11498 | 0.58587 | 0.09866 | 15.82026 | 0.02 | 0 Neutral |
| testneg_8822 | 0 | 0 | 0.524955 | -3 | 1.105669 | -3 | 0.2734 | 0.2578 | 0.44385 | 0.1311 | 4.72222 | 0.038214938 | 0 Neutral |
| testneg_8823 | 0.04 | 2.01 | 5.095532 | -3 | -0.569149 | -3 | 0.01573 | 0.10031 | 0.27528 | 0.12788 | 18.88889 | 0.14 | 0 Neutral |
| testneg_8824 | -0.07 | -0.38 | 1.681972 | -3 | 1.445184 | 1.445184 | 0.4563 | 0.12971 | 0.33469 | 0.17432 | 11.83526 | 0.070741758 | 0 Neutral |
| testneg_8825 | 0 | 0 | 0.991757 | -3 | 2.542983 | -3 | 0.19896 | 0.11604 | 0.19898 | 0.30319 | 13.51351 | 0.071468863 | 0 Neutral |
| testneg_8826 | -0.06 | 0.57 | 2.929719 | -3 | 3.036045 | -3 | 0.34288 | 0.24605 | 0.16098 | 0.12788 | 23.61749 | 0.220168852 | 0.4 Neutral |
| testneg_8827 | 0.14 | -0.37 | 5.499025 | -3 | -0.823621 | -0.823621 | 0.19898 | 0.10269 | 0.27382 | 0.16898 | 25.75 | 0.328243735 | 4.8 Neutral |
| testneg_8828 | 0 | -0.01 | 2.889805 | -3 | 1.365197 | -3 | 0.80979 | 0.5595 | 0.27246 | 0.12788 | 12.10882 | 0.099818182 | 0 Neutral |
| testneg_8829 | -0.09 | 0.67 | 1.95706 | -3 | -1.457001 | -3 | 0.14948 | 0.12788 | 0.25119 | 0.13626 | 19.06369 | 0.073205128 | 0 Neutral |
| testneg_8830 | 0 | 0 | 1.194099 | -3 | -2.770894 | -3 | 0.14643 | 0.12788 | 0.54648 | 0.19042 | 5.9904 | 0.02779386 | 0 Neutral |
| testneg_8831 | 0.06 | -0.57 | 5.440283 | -3 | 0.100154 | 0.100154 | 0.32406 | 0.28376 | 0.21122 | 0.12686 | 10 | 0.22 | 0.4 Neutral |
| testneg_8832 | 0.18 | -1.05 | 7.147075 | -3 | 0.193292 | -3 | 0.25989 | 0.11651 | 0.09633 | 0.12788 | 25.55556 | 0.216845238 | 0.6 Neutral |
| testneg_8833 | 0.18 | -1.05 | 1.057128 | -3 | 0.1648 | -3 | 0.11358 | 0.12788 | 0.04869 | 0.05922 | 24.83405 | 0.03 | 0 Neutral |
| testneg_8834 | -0.07 | -0.38 | 2.393948 | 2.393948 | 6.149134 | -3 | 0.70819 | 0.5975 | 0.29372 | 0.07609 | 8.1746 | 0.34 | 5.4 Neutral |
| testneg_8835 | 0 | 0 | -0.76928 | -3 | 2.620924 | -3 | 0.19898 | 0.12788 | 0.19898 | 0.10307 | 63 | 0.070501253 | 0 Neutral |
| testneg_8836 | 0.05 | -0.81 | 5.683077 | -3 | 0.162051 | -3 | 0.18964 | 0.34036 | 0.20047 | 0.11827 | 19.05934 | 0.25 | 0.6 Neutral |
| testneg_8837 | 0.14 | -0.37 | 4.514811 | -3 | 4.076957 | 2.768711 | 0.48662 | 0.23781 | 0.38324 | 0.25193 | 10.22898 | 0.86355736 | 100 Disease_causing |
| testneg_8838 | 0 | -0.01 | 1.961035 | -3 | 2.864843 | -3 | 0.55105 | 0.43208 | 0.63363 | 0.14795 | 9.85476 | 0.18010101 | 0 Neutral |
| testneg_8839 | 0 | 0 | -0.475892 | -3 | 2.956376 | -3 | 0.85508 | 0.12634 | 0.12176 | 0.09356 | 16.88062 | 0.04 | 0 Neutral |
| testneg_8840 | 0.21 | 0.15 | 1.478329 | -3 | -0.009242 | -3 | 0.27614 | 0.12085 | 0.11945 | 0.10135 | 32.9735 | 0.03 | 0 Neutral |
| testneg_8841 | 0.03 | 0.12 | 2.210824 | -3 | 0.402508 | 0.402508 | 0.38171 | 0.15977 | 0.29794 | 0.11106 | 22.39796 | 0.03 | 0 Neutral |
| testneg_8842 | 0 | 0 | 1.683026 | -3 | -0.974137 | -3 | 0.84106 | 0.63405 | 0.62273 | 0.10215 | 16.53509 | 0.03 | 0 Neutral |
| testneg_8843 | 0 | 0 | 0.263551 | -3 | -0.699961 | -3 | 0.42523 | 0.13467 | 0.0472 | 0.12788 | 10.83333 | 0.011379808 | 0 Neutral |
| testneg_8844 | 0.07 | 0.38 | 2.031453 | -3 | 3.460958 | -3 | 0.04519 | 0.12788 | 0.22998 | 0.08738 | 27.31171 | 0.08 | 0 Neutral |
| testneg_8845 | 0 | 0 | 1.028069 | -3 | 3.489496 | -3 | 0.2771 | 0.14453 | 0.69254 | 0.10021 | 20.8819 | 0.063295455 | 0 Neutral |
| testneg_8846 | 0 | 0 | 1.39106 | -3 | 5.597055 | -3 | 0.74136 | 0.1049 | 0.53496 | 0.13586 | 18.88889 | 0.200165913 | 0 Neutral |
| testneg_8847 | 0 | -0.01 | 1.16328 | -3 | 5.393665 | 5.393665 | 0.42583 | 0.10901 | 0.03809 | 0.16044 | 13.51351 | 0.451940805 | 33.4 Neutral |
| testneg_8848 | 0 | 0 | 1.205908 | -3 | 4.915232 | -3 | 0.32333 | 0.23356 | 0.33418 | 0.09979 | 21.63113 | 0.111665146 | 0 Neutral |
| testneg_8849 | -0.15 | -0.04 | 0.591299 | -3 | -0.133882 | -3 | 0.07293 | 0.12989 | 0.1056 | 0.12788 | 14.8058 | 0.06 | 0 Neutral |
| testneg_8850 | 0.04 | -0.23 | 0.007971 | -3 | 0.432421 | -3 | 0.12591 | 0.09845 | 0.0922 | 0.07873 | 17.917 | 0.000817308 | 0 Neutral |
| testneg_8851 | 0.14 | -0.69 | 3.53144 | -3 | 1.167115 | -3 | 0.45405 | 0.10896 | 0.03558 | 0.05202 | 40.44613 | 0.05 | 0 Neutral |
| testneg_8852 | 0 | 0 | 0.910599 | 0.910599 | 5.002476 | -3 | 0.19898 | 0.12788 | 0.22162 | 0.09445 | 30.76728 | 0.110162907 | 0 Neutral |
| testneg_8853 | 0.14 | -0.69 | 4.727194 | -3 | 1.216081 | -3 | 0.09997 | 0.1797 | 0.08973 | 0.07662 | 29.46915 | 0.11 | 0 Neutral |
| testneg_8854 | 0 | -0.01 | 1.149331 | -3 | 6.179448 | -3 | 0.15582 | 0.14216 | 0.17296 | 0.09548 | 18.33333 | 0.149666667 | 0 Neutral |
| testneg_8855 | 0.22 | -2.66 | 6.703583 | -3 | 0.024257 | -3 | 0.09346 | 0.12788 | 0.10052 | 0.10074 | 31.44568 | 0.2 | 0.4 Neutral |
| testneg_8856 | -0.07 | -0.38 | -1.653555 | -1.653555 | 0.468996 | -3 | 0.14588 | 0.12156 | 0.38072 | 0.15993 | 18.33333 | 0.07 | 0 Neutral |
| testneg_8857 | 0 | 0 | 0.565761 | -3 | 0.05039 | -3 | 0.10383 | 0.08853 | 0.19898 | 0.12788 | 22.49104 | 0.000205134 | 0 Neutral |
| testneg_8858 | 0.06 | 0.18 | -0.650796 | -3 | -0.435286 | -0.435286 | 0.10921 | 0.34026 | 0.18341 | 0.13901 | 19.12989 | 0.171351852 | 0 Neutral |
| testneg_8859 | 0 | 0 | 0.714605 | -3 | 1.176476 | 1.176476 | 0.82126 | 0.17937 | 0.64432 | 0.10409 | 4.44444 | 0.010950092 | 0 Neutral |
| testneg_8860 | 0.01 | 0.79 | 4.139348 | -3 | 0.376826 | -3 | 0.20344 | 0.1705 | 0.41571 | 0.11118 | 12.38369 | 0.080192379 | 0 Neutral |
| testneg_8861 | -0.07 | -0.38 | 2.580867 | -3 | 1.367341 | -3 | 0.17786 | 0.15314 | 0.04694 | 0.44682 | 9.97525 | 0.167408425 | 0 Neutral |
| testneg_8862 | -0.07 | -0.38 | -0.094721 | -3 | 0.602917 | -3 | 0.16778 | 0.82322 | 0.16125 | 0.08406 | 16.02296 | 0.073245614 | 0 Neutral |
| testneg_8863 | 0 | 0 | -0.168453 | -3 | 1.687316 | -3 | 0.99689 | 0.28104 | 0.26434 | 0.12788 | 9.74183 | 0.04 | 0 Neutral |
| testneg_8864 | -0.02 | 0.36 | 6.611139 | -3 | 3.817144 | -3 | 0.36324 | 0.15729 | 0.12321 | 0.08084 | 13.51351 | 0.549723872 | 72.2 Disease_causing |
| testneg_8865 | 0.03 | 0.12 | 1.249184 | -3 | 3.299396 | 3.299396 | 0.40592 | 0.22108 | 0.12547 | 0.09267 | 21.53005 | 0.240704446 | 0.2 Neutral |
| testneg_8866 | -0.15 | 0.28 | 1.664306 | -3 | 0.501607 | -3 | 0.17301 | 0.14089 | 0.19898 | 0.40608 | 10.23791 | 0.112551997 | 0 Neutral |
| testneg_8867 | -0.15 | -0.04 | -1.165102 | -3 | 7.121043 | -3 | 0.37237 | 0.36829 | 0.35945 | 0.12025 | 12.01426 | 0.46 | 41.2 Neutral |
| testneg_8868 | -0.02 | 0.68 | 0.381493 | 0.381493 | 3.270074 | -3 | 0.14432 | 0.16919 | 0.1837 | 0.27351 | 10 | 0.349906174 | 8 Neutral |
| testneg_8869 | 0.04 | -0.23 | 7.572239 | -3 | 3.355696 | -3 | 0.7342 | 0.11023 | 0.16098 | 0.12788 | 17.14246 | 0.514764706 | 59.2 Disease_causing |
| testneg_8870 | -0.03 | -0.12 | 3.470123 | -3 | -0.691406 | -3 | 0.32532 | 0.09651 | 0.04046 | 0.06599 | 18.33333 | 0.03 | 0 Neutral |
| testneg_8871 | 0.03 | 1.22 | 7.358991 | -3 | 3.907702 | -3 | 0.03109 | 0.10705 | 0.14498 | 0.33896 | 10.09474 | 0.69597619 | 98.8 Disease_causing |
| testneg_8872 | 0.05 | -0.81 | 5.724036 | -3 | 3.753987 | -3 | 0.54231 | 0.15254 | 0.06736 | 0.06344 | 12.52604 | 0.381763779 | 15.6 Neutral |
| testneg_8873 | 0 | 0 | 1.040333 | -3 | 2.470789 | 0.057057 | 0.06793 | 0.12788 | 0.61098 | 0.11179 | 24.53553 | 0.03 | 0 Neutral |
| testneg_8874 | 0.01 | 0.79 | 0.643195 | -3 | 4.834631 | -3 | 0.54777 | 0.25991 | 0.24058 | 0.14928 | 19.44444 | 0.462679616 | 39.2 Neutral |
| testneg_8875 | -0.14 | 1.01 | 2.72498 | -3 | -0.825559 | -3 | 0.34001 | 0.12788 | 0.14349 | 0.27976 | 14.29208 | 0.13076457 | 0 Neutral |
| testneg_8876 | 0 | 0 | 0.507327 | -3 | 3.576466 | -0.697584 | 0.11172 | 0.10034 | 0.19898 | 0.12788 | 23.68839 | 0.031509491 | 0 Neutral |
| testneg_8877 | 0.15 | 0.06 | -3.376225 | -3 | 5.237542 | -3 | 0.04123 | 0.06896 | 0.12689 | 0.13376 | 23.45218 | 0.280176282 | 0.6 Neutral |
| testneg_8878 | 0 | 0 | 0.919505 | -3 | 3.635519 | -3 | 0.19898 | 0.12788 | 0.07995 | 0.12788 | 20.15462 | 0.043392857 | 0 Neutral |
| testneg_8879 | 0 | 0 | -0.586784 | -3 | -0.260879 | -3 | 0.43314 | 0.67252 | 0.09527 | 0.12788 | 18.33333 | 0.05 | 0 Neutral |
| testneg_8880 | 0.07 | -0.31 | 3.54624 | 3.54624 | 0.989156 | -3 | 0.05476 | 0.12788 | 0.14941 | 0.0838 | 42.43023 | 0.43 | 27.8 Neutral |
| testneg_8881 | 0 | 0 | 1.002333 | -3 | 1.418708 | -3 | 0.13918 | 0.12788 | 0.6532 | 0.12788 | 19.70009 | 0.000392387 | 0 Neutral |
| testneg_8882 | -0.15 | -0.04 | 2.11574 | -3 | 5.803421 | -3 | 0.27407 | 0.1136 | 0.04515 | 0.14318 | 11.25 | 0.438505661 | 25.2 Neutral |
| testneg_8883 | 0.21 | 0.15 | 1.937963 | -3 | 3.074697 | -3 | 0.19898 | 0.37148 | 0.37282 | 0.07943 | 23.2019 | 0.18 | 0 Neutral |
| testneg_8884 | 0 | 0 | 2.004172 | -3 | -2.464297 | -2.464297 | 0.18175 | 0.13527 | 0.11524 | 0.12788 | 18.09499 | 0.01 | 0 Neutral |
| testneg_8885 | 0 | 0 | 1.316654 | -3 | -1.122075 | -3 | 0.16613 | 0.12378 | 0.34184 | 0.1964 | 10.88899 | 0.020731451 | 0 Neutral |
| testneg_8886 | 0 | 0 | 0.062047 | -3 | 2.493168 | -3 | 0.21928 | 0.22085 | 0.14897 | 0.19122 | 11.95767 | 0.17681308 | 0 Neutral |
| testneg_8887 | 0.14 | -1.01 | 0.302883 | -3 | -2.855052 | -3 | 0.10486 | 0.08083 | 0.16545 | 0.10002 | 19.12074 | 0.02 | 0 Neutral |
| testneg_8888 | 0.08 | 0.03 | -1.432545 | -3 | 1.606772 | 1.606772 | 0.18416 | 0.13292 | 0.35401 | 0.08438 | 9.71235 | 0.04 | 0 Neutral |
| testneg_8889 | 0 | 0 | 0.279883 | -3 | -0.11259 | -3 | 0.48906 | 0.11262 | 0.14806 | 0.09574 | 33.98632 | 0.010313982 | 0 Neutral |
| testneg_8890 | 0 | 0 | -0.078585 | -3 | 1.334446 | 1.334446 | 0.52457 | 0.38957 | 0.15514 | 0.14852 | 13.51351 | 0.124345513 | 0 Neutral |
| testneg_8891 | 0.07 | 0.38 | 0.342677 | -3 | 0.188594 | -3 | 0.03526 | 0.06855 | 0.35499 | 0.12052 | 46.84387 | 0.030510128 | 0 Neutral |
| testneg_8892 | 0 | 0 | 1.349695 | -3 | 0.717863 | -3 | 0.20016 | 0.20159 | 0.19832 | 0.19622 | 14.34433 | 0.061080136 | 0 Neutral |
| testneg_8893 | 0.02 | 0.41 | -2.259592 | -3 | 0.190526 | -3 | 0.06311 | 0.14802 | 0.1834 | 0.12788 | 51.36777 | 0.060736656 | 0 Neutral |
| testneg_8894 | -0.21 | -0.15 | 2.147578 | -3 | -1.989097 | -3 | 0.47335 | 0.12788 | 0.11094 | 0.12788 | 10 | 0.05 | 0 Neutral |
| testneg_8895 | 0 | 0 | 0.634312 | -3 | 2.428489 | -3 | 0.07681 | 0.22865 | 0.05027 | 0.10981 | 19.47429 | 0.0602 | 0 Neutral |
| testneg_8896 | 0 | -0.01 | 0.494045 | -3 | 3.319567 | -3 | 0.1304 | 0.12788 | 0.0634 | 0.0824 | 18.88889 | 0.03 | 0 Neutral |
| testneg_8897 | 0.07 | 0.01 | 3.189025 | -3 | 3.934428 | -3 | 0.7639 | 0.10408 | 0.19898 | 0.12788 | 18.03715 | 0.136114719 | 0 Neutral |
| testneg_8898 | 0.09 | -0.3 | -1.575461 | -3 | 6.041443 | -3 | 0.21241 | 0.08684 | 0.10703 | 0.11611 | 30.95458 | 0.110697654 | 0 Neutral |
| testneg_8899 | 0.14 | -0.37 | 3.020834 | -3 | 3.138075 | 0.185181 | 0.19894 | 0.27411 | 0.82156 | 0.14621 | 10 | 0.467807853 | 43 Neutral |
| testneg_8900 | 0.14 | -1.01 | 2.521088 | -3 | 3.461522 | -3 | 0.10021 | 0.1105 | 0.07923 | 0.12788 | 22.14286 | 0.086904762 | 0 Neutral |
| testneg_8901 | 0.04 | 2.01 | 5.095532 | -3 | 0.681978 | -3 | 0.01573 | 0.10031 | 0.12543 | 0.12788 | 29.8569 | 0.14 | 0 Neutral |
| testneg_8902 | 0.05 | -0.81 | 5.265577 | -3 | 3.379422 | -3 | 0.55438 | 0.11671 | 0.15848 | 0.12754 | 21.94444 | 0.26749183 | 1.6 Neutral |
| testneg_8903 | -0.03 | -0.12 | -1.612426 | -3 | 0.284309 | -3 | 0.32989 | 0.13525 | 0.18882 | 0.14637 | 14.86134 | 0.05 | 0 Neutral |
| testneg_8904 | 0.01 | 0.8 | -0.725111 | -0.725111 | -0.214687 | -3 | 0.05116 | 0.08996 | 0.19668 | 0.16042 | 13.51351 | 0.13 | 0 Neutral |
| testneg_8905 | 0 | -0.01 | 6.857724 | 6.857724 | 0.278169 | -3 | 0.18224 | 0.10419 | 0.10963 | 0.12788 | 16.8018 | 0.629565217 | 94 Disease_causing |

| | | | | | | | | | | | | | | |
|---|---|---|---|---|---|---|---|---|---|---|---|---|---|---|
| testneg_8906 | 0.06 | 0.18 | 0.526851 | -3 | 0.955677 | 0.955677 | 0.57188 | 0.34038 | 0.10034 | 0.10906 | 10.41667 | 0.050748837 | 0 | Neutral |
| testneg_8907 | 0 | 0 | 0.71683 | -3 | 5.88281 | -3 | 0.42583 | 0.10901 | 0.17322 | 0.10311 | 23.24046 | 0.081168831 | 0 | Neutral |
| testneg_8908 | 0 | 0 | -0.044046 | -3 | 1.84534 | 1.84534 | 0.47544 | 0.17658 | 0.40996 | 0.57507 | 10 | 0.1246865 | 0 | Neutral |
| testneg_8909 | 0 | 0 | -0.031285 | -3 | -0.337044 | -3 | 0.64661 | 0.49283 | 0.02396 | 0.12703 | 4.98847 | 0.02 | 0 | Neutral |
| testneg_8910 | 0.15 | 0.06 | 4.245447 | -3 | 7.672083 | -3 | 0.19898 | 0.12788 | 0.16355 | 0.12788 | 18.76267 | 0.536945804 | 67 | Disease_causing |
| testneg_8911 | 0.23 | -1.86 | 3.479077 | -3 | -2.732287 | -3 | 0.17487 | 0.1672 | 0.32558 | 0.32796 | 10.12902 | 0.373779215 | 10 | Neutral |
| testneg_8912 | 0.07 | 0.7 | 3.89415 | -3 | -0.133236 | -3 | 0.16671 | 0.18311 | 0.18328 | 0.12788 | 10.64628 | 0.09 | 0 | Neutral |
| testneg_8913 | 0 | -0.32 | 0.785937 | -3 | 3.781815 | -3 | 0.39459 | 0.12788 | 0.01704 | 0.05734 | 23.95033 | 0.08 | 0 | Neutral |
| testneg_8914 | 0 | -0.32 | 5.503592 | -3 | 3.86389 | -3 | 0.38552 | 0.14172 | 0.19898 | 0.12788 | 8.97898 | 0.459807316 | 37 | Neutral |
| testneg_8915 | 0.11 | -0.19 | 1.794771 | 1.794771 | 6.6361 | -3 | 0.00755 | 0.12788 | 0.26618 | 0.2057 | 26.34995 | 0.566605392 | 76.6 | Disease_causing |
| testneg_8916 | 0 | 0 | 0.645167 | -3 | 1.344217 | -3 | 0.03169 | 0.10615 | 0.28128 | 0.12883 | 21.51158 | 0.03 | 0 | Neutral |
| testneg_8917 | 0 | 0 | 1.312473 | -3 | 3.803348 | -3 | 0.45152 | 0.34457 | 0.67263 | 0.09781 | 12.63889 | 0.110603044 | 0 | Neutral |
| testneg_8918 | 0.15 | -0.28 | 0.118932 | 0.118932 | 1.678033 | -3 | 0.19567 | 0.09606 | 0.33957 | 0.12788 | 10.65539 | 0.02 | 0 | Neutral |
| testneg_8919 | 0.05 | 0.29 | 4.610666 | -3 | 2.846083 | -3 | 0.16139 | 0.09116 | 0.1059 | 0.1265 | 30.90278 | 0.12 | 0 | Neutral |
| testneg_8920 | 0.05 | 0.29 | 0.403916 | -3 | 1.927035 | -3 | 0.0978 | 0.26338 | 0.71814 | 0.72461 | 5 | 0.190333333 | 0 | Neutral |
| testneg_8921 | -0.02 | 0.36 | 4.584749 | -3 | 4.433629 | -3 | 0.86306 | 0.10962 | 0.05087 | 0.09682 | 12.34649 | 0.240326923 | 0.2 | Neutral |
| testneg_8922 | 0 | -0.73 | 5.679645 | -3 | 0.916536 | -3 | 0.3954 | 0.17605 | 0.66863 | 0.10981 | 24.67462 | 0.26 | 0.4 | Neutral |
| testneg_8923 | 0 | 0 | 1.371909 | -3 | 0.21945 | -3 | 0.47733 | 0.08388 | 0.81566 | 0.22678 | 26.469 | 0.04 | 0 | Neutral |
| testneg_8924 | 0.18 | -1.05 | 5.931475 | -3 | 1.200667 | -3 | 0.19494 | 0.14451 | 0.44774 | 0.15015 | 14.53974 | 0.399869565 | 21.6 | Neutral |
| testneg_8925 | 0 | -0.32 | 2.54814 | -3 | 0.665575 | -3 | 0.85592 | 0.20297 | 0.11613 | 0.1109 | 18.88889 | 0.025 | 0 | Neutral |
| testneg_8926 | 0 | 0 | -1.231395 | -3 | -0.235365 | -3 | 0.46757 | 0.14122 | 0.99535 | 0.38424 | 20.27778 | 0.120326802 | 0 | Neutral |
| testneg_8927 | 0.14 | -0.37 | 6.124865 | 2.291577 | 4.532035 | -3 | 0.15406 | 0.1454 | 0.0968 | 0.15592 | 20 | 0.828285714 | 100 | Disease_causing |
| testneg_8928 | 0 | 0 | -1.231395 | -3 | -0.062562 | -3 | 0.46757 | 0.14122 | 0.48437 | 0.09986 | 25.74074 | 0.020196154 | 0 | Neutral |
| testneg_8929 | -0.08 | -0.03 | 4.945871 | -3 | 2.73719 | -3 | 0.055 | 0.20858 | 0.1454 | 0.10285 | 18.6768 | 0.292175373 | 1 | Neutral |
| testneg_8930 | -0.09 | 0.3 | -0.671472 | -3 | 0.238135 | -3 | 0.08878 | 0.0974 | 0.27528 | 0.12788 | 24.83405 | 0.020194193 | 0 | Neutral |
| testneg_8931 | 0 | 0 | 0.865958 | -3 | 2.161297 | -3 | 0.9874 | 0.15588 | 0.07071 | 0.12788 | 10.41667 | 0.021165773 | 0 | Neutral |
| testneg_8932 | 0.19 | -0.25 | 1.146824 | -3 | 1.539176 | 1.539176 | 0.77364 | 0.12676 | 0.91208 | 0.52258 | 10.41667 | 0.12 | 0 | Neutral |
| testneg_8933 | 0 | 0 | 0.82819 | 0.82819 | 0.734956 | -3 | 0.19898 | 0.12788 | 0.14954 | 0.12788 | 27.77778 | 0.03 | 0 | Neutral |
| testneg_8934 | 0.18 | -1.05 | 4.521011 | -3 | 5.743656 | -3 | 0.04436 | 0.12788 | 0.85345 | 0.13067 | 22.24087 | 0.358181818 | 11.2 | Neutral |
| testneg_8935 | -0.07 | -0.38 | 3.513469 | -3 | 1.902059 | -3 | 0.99445 | 0.10396 | 0.60755 | 0.41186 | 10.83333 | 0.219455882 | 0 | Neutral |
| testneg_8936 | 0 | 0 | -0.296644 | -0.71253 | 1.575308 | -3 | 0.03563 | 0.10455 | 0.20655 | 0.15317 | 21.46199 | 0.11 | 0 | Neutral |
| testneg_8937 | -0.05 | -0.29 | -2.95337 | -3 | 0.002159 | -3 | 0.19898 | 0.12788 | 0.10231 | 0.1 | 18.76159 | 0.02 | 0 | Neutral |
| testneg_8938 | 0.14 | -1.85 | 2.534211 | -3 | 1.792362 | -3 | 0.17487 | 0.1672 | 0.23427 | 0.10875 | 18.50536 | 0.062151316 | 0 | Neutral |
| testneg_8939 | 0 | 0 | 0.053529 | -3 | -0.01248 | -3 | 0.46318 | 0.14229 | 0.61231 | 0.13362 | 13.69835 | 0.060302135 | 0 | Neutral |
| testneg_8940 | 0 | 0 | 0.002341 | -3 | 1.52981 | -3 | 0.10799 | 0.10754 | 0.34001 | 0.12788 | 19.15481 | 0 | 0 | Neutral |
| testneg_8941 | 0 | 0 | -0.101683 | -3 | 2.54434 | -3 | 0.72657 | 0.11262 | 0.29425 | 0.50236 | 7.01832 | 0.077995608 | 0 | Neutral |
| testneg_8942 | -0.15 | -0.04 | -1.47103 | -3 | 0.330829 | -3 | 0.50682 | 0.12788 | 0.06392 | 0.12788 | 13.27141 | 0.05 | 0 | Neutral |
| testneg_8943 | -0.01 | -0.8 | 7.038327 | -3 | -0.620486 | -3 | 0.30367 | 0.41964 | 0.03942 | 0.12788 | 15.29221 | 0.38 | 13 | Neutral |
| testneg_8944 | -0.03 | -1.22 | 1.025171 | -3 | 3.809317 | -3 | 0.46664 | 0.09008 | 0.2442 | 0.11401 | 13.51351 | 0.071371374 | 0 | Neutral |
| testneg_8945 | 0.16 | -1.37 | 5.18093 | -3 | 6.543992 | -3 | 0.13203 | 0.12092 | 0.19898 | 0.2621 | 17.77778 | 0.741057018 | 99.4 | Disease_causing |
| testneg_8946 | -0.23 | 1.86 | -0.389748 | -3 | 4.319571 | -3 | 0.08891 | 0.12788 | 0.36324 | 0.15729 | 29.0094 | 0.42 | 21 | Neutral |
| testneg_8947 | 0 | 0 | 0.432421 | -3 | 4.350853 | -3 | 0.0922 | 0.07873 | 0.12161 | 0.24178 | 19.26471 | 0.226653382 | 0.4 | Neutral |
| testneg_8948 | 0 | 0 | 1.975201 | -3 | 1.205837 | -3 | 0.34495 | 0.16861 | 0.07071 | 0.11499 | 10.41667 | 0.010536398 | 0 | Neutral |
| testneg_8949 | 0 | 0 | 0.546525 | -3 | 1.018783 | -3 | 0.12545 | 0.08963 | 0.07189 | 0.11325 | 16.84579 | 0.000521328 | 0 | Neutral |
| testneg_8950 | 0 | -0.01 | 0.507765 | -3 | 2.716952 | -3 | 0.65407 | 0.50749 | 0.8433 | 0.13128 | 12.46689 | 0.126464547 | 0 | Neutral |
| testneg_8951 | 0 | 0 | 0.093296 | 0.093296 | 5.91019 | -3 | 0.19241 | 0.15665 | 0.16462 | 0.18467 | 12.70833 | 0.458030108 | 39.8 | Neutral |
| testneg_8952 | 0.06 | 0.18 | 4.598509 | -3 | 5.24199 | -3 | 0.11351 | 0.10667 | 0.10566 | 0.12788 | 20.20726 | 0.274939815 | 2.2 | Neutral |
| testneg_8953 | 0.08 | 0.03 | -0.196592 | -0.196592 | -1.427787 | -3 | 0.32733 | 0.16869 | 0.11261 | 0.13673 | 20.76828 | 0.164800187 | 0 | Neutral |
| testneg_8954 | 0 | 0 | -0.029976 | -3 | 2.72498 | -3 | 0.11731 | 0.06348 | 0.34001 | 0.12788 | 35.97495 | 0.03 | 0 | Neutral |
| testneg_8955 | 0.09 | -2.17 | 5.174314 | -3 | 1.751991 | 1.157706 | 0.73311 | 0.10866 | 0.09951 | 0.14299 | 10.67325 | 0.27 | 0.8 | Neutral |
| testneg_8956 | 0.14 | -1.01 | 6.804639 | -3 | 4.271499 | 0.486085 | 0.07577 | 0.09128 | 0.71844 | 0.20204 | 19.44444 | 0.584166667 | 82.8 | Disease_causing |
| testneg_8957 | -0.02 | 0.63 | -0.552746 | -3 | -0.756067 | -3 | 0.02971 | 0.33983 | 0.58124 | 0.31208 | 30.95448 | 0.28 | 0.6 | Neutral |
| testneg_8958 | 0 | -0.01 | -0.474469 | -3 | 2.36406 | -3 | 0.35422 | 0.11185 | 0.10395 | 0.08737 | 30.22972 | 0.03 | 0 | Neutral |
| testneg_8959 | -0.05 | -0.29 | -0.343428 | -3 | -0.459078 | -3 | 0.13794 | 0.40936 | 0.19214 | 0.12788 | 20.20694 | 0.06 | 0 | Neutral |
| testneg_8960 | 0.09 | -0.3 | 3.872922 | -3 | 0.897808 | 0.897808 | 0.1675 | 0.09413 | 0.17155 | 0.18585 | 21.43606 | 0.191825475 | 0 | Neutral |
| testneg_8961 | 0.11 | -1.95 | 4.730643 | -3 | -1.919458 | -3 | 0.94662 | 0.11262 | 0.09918 | 0.11061 | 20.4497 | 0.11 | 0 | Neutral |
| testneg_8962 | 0 | 0 | 1.62409 | -3 | 0.083768 | -3 | 0.20457 | 0.12378 | 0.08041 | 0.21802 | 18.88889 | 0.050201288 | 0 | Neutral |
| testneg_8963 | 0.02 | 0.41 | 3.953483 | -3 | 0.812392 | -3 | 0.97273 | 0.27711 | 0.19898 | 0.10512 | 12.6598 | 0.08 | 0 | Neutral |
| testneg_8964 | -0.15 | -0.04 | 3.934875 | -3 | -0.058596 | -3 | 0.1224 | 0.11899 | 0.1144 | 0.09798 | 26.41358 | 0.061004902 | 0 | Neutral |
| testneg_8965 | 0 | 0 | -0.029976 | -3 | 0.92793 | -3 | 0.11731 | 0.06348 | 0.0925 | 0.08222 | 27.67377 | 0.02 | 0 | Neutral |
| testneg_8966 | 0 | 0 | -0.174435 | -3 | 8.984539 | -3 | 0.31542 | 0.26739 | 0.18697 | 0.12232 | 35.05388 | 0.415 | 24.6 | Neutral |
| testneg_8967 | 0.16 | -0.37 | 0.701088 | -3 | 6.776625 | -3 | 0.99538 | 0.40044 | 0.17039 | 0.12788 | 12.32809 | 0.362680403 | 10.6 | Neutral |
| testneg_8968 | 0.09 | -0.3 | 4.334032 | -3 | -0.660504 | -3 | 0.18022 | 0.12788 | 0.19898 | 0.12788 | 13.46801 | 0.020741758 | 0 | Neutral |
| testneg_8969 | 0 | 0 | 1.548509 | -3 | -0.550575 | -3 | 0.06711 | 0.09765 | 0.09445 | 0.12788 | 14.44718 | 0.000235327 | 0 | Neutral |
| testneg_8970 | 0 | 0 | 0.202655 | -3 | 6.645041 | -3 | 0.18834 | 0.12788 | 0.13433 | 0.09578 | 21.38813 | 0.160501253 | 0.2 | Neutral |
| testneg_8971 | 0.02 | 0.45 | 5.933886 | -3 | 2.628269 | -3 | 0.09099 | 0.2051 | 0.18584 | 0.11114 | 19.03726 | 0.384758621 | 14.2 | Neutral |
| testneg_8972 | -0.08 | -0.03 | 1.417345 | -3 | -0.01214 | -3 | 0.04615 | 0.12788 | 0.16966 | 0.61198 | 11.0208 | 0.130283242 | 0 | Neutral |
| testneg_8973 | 0.2 | -1.26 | 2.481337 | -3 | -0.04628 | -3 | 0.12408 | 0.08828 | 0.19898 | 0.12788 | 11.4135 | 0.03015239 | 0 | Neutral |
| testneg_8974 | 0.02 | 0.13 | 1.091299 | -3 | 3.730704 | -3 | 0.1163 | 0.20863 | 0.37951 | 0.19435 | 10.83333 | 0.488870248 | 49.8 | Neutral |
| testneg_8975 | 0 | 0 | 3.506817 | -3 | -2.660026 | -2.660026 | 0.23088 | 0.21602 | 0.37293 | 0.25159 | 10.83333 | 0.261857769 | 0.6 | Neutral |
| testneg_8976 | 0.16 | 0.08 | -0.28306 | -3 | 2.943194 | -3 | 0.44067 | 0.11201 | 0.05817 | 0.09497 | 26.34995 | 0.082778184 | 0 | Neutral |
| testneg_8977 | 0.14 | -1.01 | 0.023757 | -3 | 1.081088 | -3 | 0.3221 | 0.3367 | 0.21201 | 0.20962 | 7.00579 | 0.15021934 | 0 | Neutral |
| testneg_8978 | 0.14 | -1.01 | 7.095637 | -3 | -2.323467 | -3 | 0.11797 | 0.12788 | 0.05877 | 0.12788 | 18.33333 | 0.23 | 1.2 | Neutral |
| testneg_8979 | 0.16 | -0.37 | 4.574218 | -3 | 0.080104 | -3 | 0.26451 | 0.09729 | 0.73947 | 0.2058 | 30.95732 | 0.172583333 | 0 | Neutral |
| testneg_8980 | -0.07 | -0.7 | 2.642734 | -3 | 3.392586 | -3 | 0.19898 | 0.12788 | 0.10084 | 0.12788 | 31.04675 | 0.07 | 0 | Neutral |
| testneg_8981 | 0 | 0 | 2.064931 | -3 | 7.530767 | -3 | 0.30122 | 0.09165 | 0.1897 | 0.12459 | 17.88746 | 0.251127413 | 1.2 | Neutral |
| testneg_8982 | -0.21 | -0.15 | -2.327557 | -3 | 3.861511 | -3 | 0.34694 | 0.20397 | 0.09018 | 0.16175 | 11.92982 | 0.48959277 | 50.2 | Disease_causing |
| testneg_8983 | 0.01 | 0.79 | 3.682003 | -3 | 0.295768 | -3 | 0.51319 | 0.12971 | 0.16741 | 0.10762 | 7.01832 | 0.05003876 | 0 | Neutral |
| testneg_8984 | 0.14 | -0.69 | 1.909906 | -3 | -1.367259 | -3 | 0.29523 | 0.27623 | 0.06099 | 0.12788 | 20.31414 | 0.050384615 | 0 | Neutral |
| testneg_8985 | -0.03 | -0.1 | 2.928572 | -3 | 0.080284 | -3 | 0.60285 | 0.15179 | 0.63818 | 0.12788 | 10.41667 | 0.06 | 0 | Neutral |
| testneg_8986 | 0.01 | 0.8 | 0.307203 | -0.508098 | 0.810517 | -3 | 0.07872 | 0.12788 | 0.08729 | 0.10515 | 16.8018 | 0.010435606 | 0 | Neutral |
| testneg_8987 | 0.04 | 2.01 | -1.062451 | -3 | 5.202788 | -3 | 0.03072 | 0.27473 | 0.89845 | 0.25991 | 10.41667 | 0.556515152 | 79 | Disease_causing |
| testneg_8988 | 0.17 | 0.38 | 1.534359 | -3 | 3.445512 | -3 | 0.07275 | 0.09562 | 0.38126 | 0.27764 | 13.02635 | 0.263482143 | 0.8 | Neutral |
| testneg_8989 | 0.03 | 1.22 | 0.362146 | -3 | 0.438443 | 0.438443 | 0.16386 | 0.12788 | 0.36857 | 0.36135 | 16.48772 | 0.120236291 | 0 | Neutral |
| testneg_8990 | 0 | 0 | 1.519976 | -3 | 2.60676 | -3 | 0.19898 | 0.12788 | 0.09439 | 0.07415 | 16.8018 | 0.01 | 0 | Neutral |
| testneg_8991 | 0 | 0 | 1.626934 | -3 | 2.454179 | -3 | 0.20942 | 0.14229 | 0.11788 | 0.12788 | 12.72648 | 0.02 | 0 | Neutral |
| testneg_8992 | -0.08 | -0.03 | 3.206814 | -3 | 2.724707 | -3 | 0.7022 | 0.35308 | 0.49012 | 0.12138 | 18.77814 | 0.2218125 | 0 | Neutral |
| testneg_8993 | -0.03 | -0.12 | -0.132037 | -3 | 5.824607 | -3 | 0.22794 | 0.10688 | 0.03391 | 0.09745 | 41.05931 | 0.10039283 | 0 | Neutral |
| testneg_8994 | 0 | 0 | 1.943537 | 1.943537 | 7.240809 | -3 | 0.19237 | 0.23658 | 0.19898 | 0.10593 | 19.77034 | 0.440357143 | 33 | Neutral |
| testneg_8995 | 0.08 | -1.2 | 5.073628 | 3.672069 | 2.987486 | -3 | 0.02146 | 0.10351 | 0.08896 | 0.12788 | 18.88889 | 0.48 | 48.6 | Neutral |
| testneg_8996 | 0 | 0 | 1.992408 | -3 | 0.985586 | -3 | 0.19898 | 0.25991 | 0.12077 | 0.11244 | 9.88506 | 0.01 | 0 | Neutral |
| testneg_8997 | 0.19 | 0.51 | 1.770771 | -3 | 1.853291 | -3 | 0.71406 | 0.2012 | 0.13468 | 0.09235 | 27.27062 | 0.09 | 0 | Neutral |
| testneg_8998 | 0.19 | -0.27 | 5.193514 | -3 | 0.222955 | -3 | 0.57188 | 0.34038 | 0.0538 | 0.0823 | 10 | 0.16 | 0 | Neutral |
| testneg_8999 | 0 | 0 | 1.52981 | -3 | 4.092031 | -3 | 0.34001 | 0.12788 | 0.50134 | 0.15488 | 51.99903 | 0.218668197 | 0.4 | Neutral |
| testneg_9000 | 0.01 | 0.79 | 0.841519 | -3 | 1.118815 | -3 | 0.81054 | 0.28104 | 0.13199 | 0.14263 | 16.07106 | 0.108042408 | 0 | Neutral |
| testneg_9001 | 0 | 0 | -0.583572 | -3 | 6.519516 | -3 | 0.04555 | 0.06327 | 0.28211 | 0.25332 | 26.84685 | 0.322791667 | 4.4 | Neutral |
| testneg_9002 | 0 | 0 | 1.241234 | -3 | 1.365197 | -3 | 0.6258 | 0.16461 | 0.27246 | 0.12788 | 12.00993 | 0.01 | 0 | Neutral |
| testneg_9003 | 0 | 0 | 0.760575 | -3 | 0.6838 | 0.6838 | 0.1878 | 0.08946 | 0.67327 | 0.23543 | 14.34932 | 0.030689705 | 0 | Neutral |
| testneg_9004 | 0 | 0 | -0.011929 | -3 | 0.469254 | -3 | 0.15192 | 0.14302 | 0.16052 | 0.12788 | 18.07092 | 0.01 | 0 | Neutral |
| testneg_9005 | 0 | 0 | 0.811666 | -3 | 3.854582 | -3 | 0.2674 | 0.28725 | 0.44953 | 0.14403 | 10.83333 | 0.321766676 | 5.4 | Neutral |
| testneg_9006 | 0 | 0 | 0.169215 | -3 | 1.638668 | -3 | 0.14335 | 0.14499 | 0.14199 | 0.1327 | 19.98891 | 0.0751 | 0 | Neutral |
| testneg_9007 | 0 | 0 | -0.054636 | -3 | 0.2136 | -3 | 0.5632 | 0.28442 | 0.04063 | 0.05848 | 28.2825 | 0.04 | 0 | Neutral |
| testneg_9008 | -0.02 | -0.41 | 1.480614 | -3 | -2.289069 | -3 | 0.11849 | 0.20101 | 0.75906 | 0.09702 | 11.30228 | 0.06 | 0 | Neutral |
| testneg_9009 | -0.04 | 1.61 | 4.905457 | -3 | 0.06448 | -3 | 0.21236 | 0.12788 | 0.30684 | 0.12788 | 32.54399 | 0.108381038 | 0 | Neutral |
| testneg_9010 | 0.05 | -0.81 | 5.140271 | -3 | -0.006282 | -0.006282 | 0.09542 | 0.14806 | 0.50287 | 0.15426 | 20.83333 | 0.327059309 | 6.6 | Neutral |
| testneg_9011 | -0.03 | -0.1 | 1.820578 | -3 | 1.603037 | -3 | 0.31507 | 0.61057 | 0.09522 | 0.20323 | 10.22898 | 0.200072464 | 0.2 | Neutral |
| testneg_9012 | 0 | 0 | -0.390566 | -3 | 1.009692 | -3 | 0.08283 | 0.11797 | 0.10571 | 0.12788 | 38.70251 | 0.01 | 0 | Neutral |
| testneg_9013 | 0 | -0.32 | -2.498179 | -3 | 0.468237 | -3 | 0.50682 | 0.12788 | 0.32334 | 0.10525 | 18.31365 | 0.02 | 0 | Neutral |
| testneg_9014 | 0.02 | 0.13 | -0.947523 | -3 | 2.908555 | -3 | 0.10638 | 0.66554 | 0.14175 | 0.12788 | 21.315 | 0.16 | 0 | Neutral |
| testneg_9015 | 0 | 0 | 4.936806 | -3 | 8.009008 | -3 | 0.54826 | 0.24974 | 0.64188 | 0.09494 | 5.89195 | 0.616128788 | 90.6 | Disease_causing |
| testneg_9016 | 0.21 | 0.15 | 1.212904 | -3 | 6.011632 | -3 | 0.08475 | 0.19889 | 0.16968 | 0.12788 | 17.4473 | 0.317485294 | 2 | Neutral |
| testneg_9017 | -0.18 | 1.05 | 2.40944 | -3 | 6.109579 | -3 | 0.17075 | 0.12788 | 0.15267 | 0.12788 | 41.47949 | 0.288 | 0 | Neutral |
| testneg_9018 | 0.04 | -0.23 | 0.007971 | -3 | 1.539176 | 1.539176 | 0.12591 | 0.09845 | 0.91208 | 0.52258 | 10.22898 | 0.09 | 0 | Neutral |
| testneg_9019 | 0.02 | 0.45 | 6.097514 | -3 | -0.811454 | -0.811454 | 0.25905 | 0.23728 | 0.72912 | 0.18218 | 11.42857 | 0.481134615 | 47.2 | Neutral |
| testneg_9020 | 0.01 | 0.8 | 0.586242 | -3 | 2.399305 | -3 | 0.559 | 0.2286 | 0.1324 | 0.09503 | 20.80137 | 0.094761905 | 0 | Neutral |
| testneg_9021 | 0.14 | -1.01 | 5.0214 | -3 | 5.295933 | -3 | 0.16646 | 0.16629 | 0.0669 | 0.10997 | 26.93805 | 0.400268895 | 18.8 | Neutral |
| testneg_9022 | -0.05 | -0.29 | 1.124718 | -3 | 1.53322 | -3 | 0.55043 | 0.12112 | 0.78537 | 0.33817 | 11.875 | 0.05 | 0 | Neutral |
| testneg_9023 | 0.18 | -1.05 | 6.294594 | -3 | -0.242431 | -3 | 0.21033 | 0.35953 | 0.55367 | 0.57993 | 10.41667 | 0.55 | 72.8 | Disease_causing |
| testneg_9024 | 0.22 | -2.66 | 7.081701 | -3 | 0.433941 | -3 | 0.34826 | 0.49454 | 0.30475 | 0.21064 | 10.83333 | 0.554166667 | 71.4 | Disease_causing |
| testneg_9025 | 0 | 0 | 1.684928 | -3 | 1.648359 | -3 | 0.33795 | 0.17455 | 0.07665 | 0.31408 | 12.33727 | 0.078183094 | 0 | Neutral |
| testneg_9026 | 0.07 | 0.33 | 2.581601 | -3 | -1.763617 | -3 | 0.3548 | 0.10717 | 0.19433 | 0.12788 | 18.40941 | 0.01 | 0 | Neutral |
| testneg_9027 | -0.2 | 1.26 | -0.70557 | -3 | 3.907702 | -3 | 0.29008 | 0.19268 | 0.14498 | 0.33896 | 5.97222 | 0.565665987 | 80.8 | Disease_causing |

| | | | | | | | | | | | | | | |
|---|---|---|---|---|---|---|---|---|---|---|---|---|---|---|
| testneg_9028 | 0.03 | 0.12 | 0.681978 | -3 | 1.225871 | -3 | 0.12543 | 0.12788 | 0.07571 | 0.09432 | 16.88659 | 0.01 | 0 | Neutral |
| testneg_9029 | 0 | 0 | 0.587716 | -3 | 0.822036 | -3 | 0.60285 | 0.15179 | 0.13575 | 0.12788 | 13.19331 | 0.01 | 0 | Neutral |
| testneg_9030 | 0.03 | 0.12 | -0.805405 | -3 | 0.88775 | 0.88775 | 0.99293 | 0.79067 | 0.32649 | 0.15357 | 4.44444 | 0.120183575 | 0 | Neutral |
| testneg_9031 | 0.17 | -1.22 | 4.581463 | -3 | -2.304927 | -2.304927 | 0.16741 | 0.10762 | 0.02665 | 0.05082 | 9.74181 | 0.06 | 0 | Neutral |
| testneg_9032 | -0.16 | -0.08 | 1.821379 | -3 | 0.643588 | -3 | 0.24026 | 0.12788 | 0.30471 | 0.11609 | 28.86444 | 0.05 | 0 | Neutral |
| testneg_9033 | 0 | 0 | 0.930011 | -3 | 2.124149 | -3 | 0.12786 | 0.72066 | 0.14482 | 0.08592 | 19.11368 | 0.067898268 | 0 | Neutral |
| testneg_9034 | 0.05 | -0.81 | 4.124389 | -3 | 7.384828 | -3 | 0.52412 | 0.21177 | 0.26323 | 0.2399 | 10.83333 | 0.90225487 | 100 | Disease_causing |
| testneg_9035 | 0.03 | 1.22 | 2.297873 | -3 | 7.749981 | -3 | 0.31168 | 0.12788 | 0.10278 | 0.08667 | 23.33333 | 0.344551282 | 5.6 | Neutral |
| testneg_9036 | 0 | 0 | 0.90039 | -3 | 0.561005 | -3 | 0.33332 | 0.12106 | 0.09527 | 0.09424 | 28.91549 | 0.01 | 0 | Neutral |
| testneg_9037 | -0.18 | -0.16 | 3.226919 | -3 | -0.68836 | -3 | 0.30042 | 0.13517 | 0.19898 | 0.28128 | 19.44444 | 0.229889393 | 0 | Neutral |
| testneg_9038 | 0 | -0.01 | 0.730013 | -3 | 5.350273 | -3 | 0.14785 | 0.09084 | 0.09284 | 0.26053 | 10.22898 | 0.260402462 | 1.8 | Neutral |
| testneg_9039 | 0 | 0 | 2.403686 | 2.403686 | 7.214082 | -3 | 0.48574 | 0.20473 | 0.09815 | 0.12788 | 22.89111 | 0.48 | 45.4 | Neutral |
| testneg_9040 | -0.14 | 1.01 | 6.153381 | -3 | 5.996034 | -3 | 0.08489 | 0.12788 | 0.08306 | 0.12788 | 10.29033 | 0.65552134 | 96.2 | Disease_causing |
| testneg_9041 | -0.03 | -0.1 | -0.76618 | -3 | 0.021671 | -3 | 0.1503 | 0.12788 | 0.13156 | 0.10621 | 12.29167 | 0.01 | 0 | Neutral |
| testneg_9042 | -0.12 | 1.16 | 6.464782 | -3 | 0.322721 | -3 | 0.98833 | 0.48581 | 0.19898 | 0.12788 | 14.31225 | 0.41 | 19 | Neutral |
| testneg_9043 | 0 | 0 | -0.68826 | -3 | 0.976803 | -3 | 0.32076 | 0.14229 | 0.50687 | 0.11283 | 10 | 0.001031123 | 0 | Neutral |
| testneg_9044 | 0 | -0.01 | 1.102315 | -3 | 0.955677 | 0.955677 | 0.31168 | 0.12788 | 0.10034 | 0.10906 | 12.29167 | 0.01 | 0 | Neutral |
| testneg_9045 | 0.21 | 0.15 | 2.998118 | -3 | 2.198339 | -3 | 0.143 | 0.11236 | 0.12176 | 0.09356 | 12.16938 | 0.068831169 | 0 | Neutral |
| testneg_9046 | 0.05 | -0.81 | 4.368284 | -3 | 2.299534 | -3 | 0.78253 | 0.1114 | 0.2074 | 0.09696 | 18.6768 | 0.07375 | 0 | Neutral |
| testneg_9047 | 0.17 | 0.38 | 1.603588 | -3 | 0.283085 | -3 | 0.10496 | 0.21491 | 0.09527 | 0.12788 | 12.70833 | 0.061666667 | 0 | Neutral |
| testneg_9048 | 0.06 | 0.18 | 4.598509 | -3 | 2.028242 | -3 | 0.11351 | 0.10667 | 0.27058 | 0.13363 | 13.45608 | 0.160130952 | 0 | Neutral |
| testneg_9049 | 0.01 | -1.3 | 6.324545 | -3 | 5.0589 | -3 | 0.24971 | 0.12971 | 0.09658 | 0.11286 | 35.43176 | 0.421971226 | 27.6 | Neutral |
| testneg_9050 | -0.09 | 0.3 | 1.440086 | -3 | 1.594197 | -3 | 0.78356 | 0.13377 | 0.03942 | 0.12788 | 28.70562 | 0.040772118 | 0 | Neutral |
| testneg_9051 | 0 | -0.01 | 0.602039 | -3 | 4.205494 | -3 | 0.126 | 0.14076 | 0.05095 | 0.08353 | 29.93204 | 0.091767045 | 0 | Neutral |
| testneg_9052 | 0.07 | 0.38 | -1.06095 | -3 | 4.637381 | -0.78887 | 0.23393 | 0.12567 | 0.47188 | 0.10037 | 10.41667 | 0.071321429 | 0 | Neutral |
| testneg_9053 | 0.15 | -0.28 | 2.534333 | -3 | 1.706136 | 0.229872 | 0.19898 | 0.12788 | 0.98853 | 0.88685 | 10.72853 | 0.131958333 | 0 | Neutral |
| testneg_9054 | -0.05 | -0.29 | 0.483862 | -3 | 3.618842 | -3 | 0.21551 | 0.12788 | 0.30241 | 0.104 | 17.77778 | 0.030741758 | 0 | Neutral |
| testneg_9055 | 0 | 0 | 1.093055 | -3 | 5.526065 | -3 | 0.99448 | 0.67288 | 0.19898 | 0.12788 | 5.69444 | 0.139705882 | 0 | Neutral |
| testneg_9056 | -0.07 | -0.33 | 4.400708 | -3 | 1.377684 | -3 | 0.12974 | 0.11886 | 0.09406 | 0.19858 | 10.83333 | 0.240697645 | 0.4 | Neutral |
| testneg_9057 | 0.09 | -2.17 | 3.834388 | -3 | -0.468724 | -3 | 0.03109 | 0.10705 | 0.02617 | 0.12788 | 45.14115 | 0.103333333 | 0 | Neutral |
| testneg_9058 | 0.03 | 0.1 | 0.477823 | -3 | 0.723491 | -3 | 0.05663 | 0.12788 | 0.55315 | 0.09195 | 27.77182 | 0.04 | 0 | Neutral |
| testneg_9059 | 0.01 | -1.44 | -2.044558 | -3 | 0.774689 | 0.774689 | 0.14511 | 0.12788 | 0.24219 | 0.17458 | 14.16667 | 0.081 | 0 | Neutral |
| testneg_9060 | 0.04 | -0.23 | -2.488973 | -3 | 0.066913 | -3 | 0.0513 | 0.33622 | 0.53239 | 0.17509 | 17.77778 | 0.26967033 | 0 | Neutral |
| testneg_9061 | 0.08 | 0.03 | 1.830459 | -3 | 0.471907 | -3 | 0.9827 | 0.16837 | 0.53424 | 0.08484 | 18.86584 | 0.06 | 0 | Neutral |
| testneg_9062 | -0.15 | 0.28 | 4.749136 | -3 | 6.085767 | -3 | 0.15767 | 0.12788 | 0.19898 | 0.12788 | 19.83449 | 0.429945652 | 28.8 | Neutral |
| testneg_9063 | 0 | 0 | 5.815666 | -3 | 0.341374 | 0.341374 | 0.34109 | 0.2167 | 0.10646 | 0.1069 | 12.87484 | 0.234804198 | 0.6 | Neutral |
| testneg_9064 | 0 | 0 | -0.258437 | -3 | 0.923463 | -3 | 0.07044 | 0.11679 | 0.06313 | 0.12788 | 13.73522 | 0.005 | 0 | Neutral |
| testneg_9065 | 0.21 | 0.15 | 6.6845 | -3 | -0.635986 | -3 | 0.32423 | 0.08104 | 0.10782 | 0.14795 | 15.78729 | 0.298647887 | 1.6 | Neutral |
| testneg_9066 | 0.17 | 0.38 | 4.08479 | -3 | 2.104789 | -3 | 0.37074 | 0.10627 | 0.54792 | 0.54626 | 4.54869 | 0.359887179 | 10 | Neutral |
| testneg_9067 | 0.01 | 0.8 | 6.01046 | -3 | 1.720206 | -3 | 0.12853 | 0.08565 | 0.6992 | 0.1804 | 18.90684 | 0.33196374 | 8.2 | Neutral |
| testneg_9068 | 0.04 | -0.23 | 7.39573 | -3 | 0.209628 | -3 | 0.8426 | 0.47836 | 0.19898 | 0.08486 | 10.82179 | 0.4 | 18.2 | Neutral |
| testneg_9069 | -0.15 | -0.04 | 2.846922 | -3 | 2.290907 | -3 | 0.18081 | 0.1258 | 0.30789 | 0.15135 | 10 | 0.193268605 | 0 | Neutral |
| testneg_9070 | 0 | 0 | -1.008848 | -3 | 5.224064 | -3 | 0.38873 | 0.11686 | 0.22278 | 0.85665 | 23.88889 | 0.291056373 | 1.2 | Neutral |
| testneg_9071 | 0 | -0.01 | 1.116052 | -3 | 0.933869 | -3 | 0.1631 | 0.40344 | 0.9862 | 0.4528 | 9.18689 | 0.094404762 | 0 | Neutral |
| testneg_9072 | -0.03 | 1.03 | 0.958989 | -3 | 0.335227 | -3 | 0.39436 | 0.16446 | 0.04346 | 0.10785 | 16.8018 | 0.05 | 0 | Neutral |
| testneg_9073 | 0.09 | -0.3 | 6.223377 | -3 | 1.273835 | -3 | 0.143 | 0.11236 | 0.21876 | 0.12373 | 16.03856 | 0.15 | 0 | Neutral |
| testneg_9074 | -0.07 | -0.38 | -0.008187 | -3 | 2.497761 | -3 | 0.16054 | 0.11683 | 0.16728 | 0.11868 | 10 | 0.02 | 0 | Neutral |
| testneg_9075 | 0 | 0 | 1.610282 | -3 | 0.382894 | -3 | 0.2663 | 0.32806 | 0.84236 | 0.14093 | 11.93907 | 0.06828213 | 0 | Neutral |
| testneg_9076 | 0 | 0 | 0.276668 | -3 | 2.331947 | -3 | 0.35376 | 0.09478 | 0.0406 | 0.12788 | 20.27778 | 0.020327957 | 0 | Neutral |
| testneg_9077 | 0 | 0 | 0.159876 | -3 | -0.360768 | -0.360768 | 0.10462 | 0.1278 | 0.47147 | 0.09503 | 15.62364 | 0.01 | 0 | Neutral |
| testneg_9078 | 0.09 | -0.3 | 6.573032 | -3 | -3.124511 | -3 | 0.48906 | 0.11262 | 0.41643 | 0.09692 | 24.1335 | 0.22 | 0.2 | Neutral |
| testneg_9079 | 0 | 0 | 3.466764 | -3 | 4.149716 | -3 | 0.34205 | 0.11195 | 0.19322 | 0.12788 | 35.64435 | 0.120490221 | 0 | Neutral |
| testneg_9080 | 0 | 0 | 0.955737 | -3 | 1.995233 | -3 | 0.24528 | 0.37575 | 0.48627 | 0.09509 | 14.40298 | 0.03 | 0 | Neutral |
| testneg_9081 | 0 | 0 | -0.272804 | -0.272804 | 3.39754 | -3 | 0.17839 | 0.12009 | 0.62203 | 0.15583 | 9.74181 | 0.131423199 | 0 | Neutral |
| testneg_9082 | 0.04 | -0.23 | 1.062269 | -3 | -2.611796 | -3 | 0.24184 | 0.06921 | 0.19898 | 0.12788 | 19.5422 | 0.01 | 0 | Neutral |
| testneg_9083 | 0 | 0 | 0.743431 | -3 | 6.112375 | -3 | 0.20942 | 0.14229 | 0.07342 | 0.0661 | 11.69461 | 0.120484982 | 0 | Neutral |
| testneg_9084 | 0 | 0 | 0.302733 | -3 | 0.082683 | -3 | 0.19898 | 0.12788 | 0.19898 | 0.12788 | 30.57129 | 9.86E-05 | 0 | Neutral |
| testneg_9085 | 0.05 | 0.29 | -1.171244 | -1.171244 | 1.516343 | -3 | 0.19898 | 0.12788 | 0.19898 | 0.12788 | 22.64794 | 0.01 | 0 | Neutral |
| testneg_9086 | 0 | 0 | 2.123947 | -3 | 1.973746 | -3 | 0.12867 | 0.11159 | 0.0594 | 0.16845 | 18.86584 | 0.086736111 | 0 | Neutral |
| testneg_9087 | -0.09 | 0.67 | 0.998175 | -3 | 2.643019 | 0.168099 | 0.34248 | 0.15472 | 0.10891 | 0.12788 | 11.25 | 0.100289916 | 0 | Neutral |
| testneg_9088 | 0.06 | -0.57 | 0.237935 | -3 | -2.550688 | -3 | 0.16362 | 0.11879 | 0.19898 | 0.12788 | 20.56995 | 0.01 | 0 | Neutral |
| testneg_9089 | 0 | 0 | 0.351623 | -3 | 2.116202 | -3 | 0.2771 | 0.14453 | 0.21978 | 0.16522 | 11.875 | 0.067244318 | 0 | Neutral |
| testneg_9090 | 0 | 0 | -0.060754 | -3 | 0.116519 | 0.116519 | 0.70946 | 0.11809 | 0.2207 | 0.09998 | 10.83333 | 0.02 | 0 | Neutral |
| testneg_9091 | -0.03 | 1.03 | 5.710421 | -3 | 1.643466 | -3 | 0.43736 | 0.10017 | 0.19343 | 0.11347 | 10.81468 | 0.254333333 | 1.6 | Neutral |
| testneg_9092 | -0.01 | 0.57 | -0.565908 | -3 | 4.435284 | -3 | 0.19898 | 0.12788 | 0.24791 | 0.09675 | 18.76159 | 0.080039063 | 0 | Neutral |
| testneg_9093 | 0.18 | -1.05 | 6.781678 | -3 | 4.820901 | -3 | 0.98565 | 0.23629 | 0.06358 | 0.5364 | 11.89315 | 0.815482143 | 100 | Disease_causing |
| testneg_9094 | 0 | 0 | -0.471382 | -3 | 1.018818 | 1.018818 | 0.21933 | 0.18619 | 0.13398 | 0.12788 | 10.29033 | 0.020188948 | 0 | Neutral |
| testneg_9095 | 0.18 | -1.05 | 3.975046 | 1.216755 | 3.064064 | -3 | 0.54049 | 0.33876 | 0.5481 | 0.10207 | 7.80841 | 0.34 | 5.2 | Neutral |
| testneg_9096 | 0.18 | 0.16 | 0.600136 | -3 | 0.520281 | -3 | 0.26842 | 0.09683 | 0.1336 | 0.15669 | 26.54712 | 0.103229167 | 0 | Neutral |
| testneg_9097 | 0.21 | 0.15 | 4.319063 | -3 | 0.781229 | 0.781229 | 0.1647 | 0.12076 | 0.07321 | 0.12788 | 40.06972 | 0.06 | 0 | Neutral |
| testneg_9098 | -0.02 | -0.45 | 2.568042 | -3 | 0.158774 | -3 | 0.84236 | 0.14093 | 0.05315 | 0.12788 | 17.45803 | 0.03 | 0 | Neutral |
| testneg_9099 | 0 | 0 | 0.223823 | -3 | 3.506817 | -3 | 0.32484 | 0.11621 | 0.23088 | 0.21602 | 11.92982 | 0.137654473 | 0 | Neutral |
| testneg_9100 | 0.14 | -0.37 | 2.363756 | 1.574468 | 2.151559 | -3 | 0.12121 | 0.4142 | 0.82591 | 0.12207 | 5.73716 | 0.15 | 0 | Neutral |
| testneg_9101 | -0.09 | 0.3 | -0.671472 | -3 | 0.233063 | 0.233063 | 0.08878 | 0.0974 | 0.12937 | 0.23004 | 27.24146 | 0.12 | 0 | Neutral |
| testneg_9102 | 0 | -0.32 | 3.955787 | -3 | 3.291156 | -3 | 0.04988 | 0.25503 | 0.19199 | 0.16616 | 12.29167 | 0.677164579 | 97.2 | Disease_causing |
| testneg_9103 | -0.03 | -0.12 | 1.988279 | -3 | -0.159822 | -3 | 0.10083 | 0.17511 | 0.27783 | 0.62384 | 10.41667 | 0.180344634 | 0 | Neutral |
| testneg_9104 | -0.15 | 1.06 | 0.996068 | -3 | 2.481102 | -3 | 0.30085 | 0.18374 | 0.25804 | 0.11876 | 18.33333 | 0.186666667 | 0 | Neutral |
| testneg_9105 | 0 | 0 | 0.899443 | -3 | 0.90039 | -3 | 0.26928 | 0.12774 | 0.33332 | 0.12106 | 10.74877 | 3.50E-05 | 0 | Neutral |
| testneg_9106 | 0.15 | -0.28 | 0.59012 | -3 | 1.075324 | 1.075324 | 0.605 | 0.10748 | 0.49934 | 0.11069 | 10.38652 | 0.02 | 0 | Neutral |
| testneg_9107 | 0.02 | -0.63 | 2.631274 | -3 | 4.730038 | 2.367211 | 0.14458 | 0.14062 | 0.19772 | 0.09261 | 22.64794 | 0.22 | 0 | Neutral |
| testneg_9108 | 0 | 0 | -0.091112 | -3 | 0.161355 | -3 | 0.93035 | 0.31722 | 0.28227 | 0.24183 | 9.46025 | 0.04257437 | 0 | Neutral |
| testneg_9109 | -0.07 | -0.38 | 1.275959 | -3 | 5.564089 | -3 | 0.2058 | 0.28174 | 0.21654 | 0.24468 | 20 | 0.608607245 | 88.4 | Disease_causing |
| testneg_9110 | -0.03 | -1.22 | 6.056888 | -3 | 1.283706 | -3 | 0.32804 | 0.10857 | 0.16856 | 0.19221 | 24.38403 | 0.454750305 | 37.4 | Neutral |
| testneg_9111 | 0.08 | 0.03 | 3.001665 | -3 | 6.066721 | -3 | 0.05846 | 0.12788 | 0.15041 | 0.43468 | 22.76045 | 0.58 | 81.6 | Disease_causing |
| testneg_9112 | 0.1 | 1.9 | 2.193084 | -3 | 1.873147 | 1.873147 | 0.95076 | 0.17388 | 0.1499 | 0.09632 | 11.18474 | 0.140437192 | 0 | Neutral |
| testneg_9113 | -0.05 | -0.29 | 1.952213 | -3 | -1.653555 | -1.653555 | 0.49504 | 0.08191 | 0.14588 | 0.12156 | 10.41667 | 0.02 | 0 | Neutral |
| testneg_9114 | 0.18 | -1.05 | 4.164795 | -3 | 0.710328 | -3 | 0.16091 | 0.90971 | 0.13007 | 0.12788 | 13.51351 | 0.18 | 0 | Neutral |
| testneg_9115 | 0 | -0.01 | 0.494045 | -3 | 0.811217 | -3 | 0.1304 | 0.12788 | 0.56416 | 0.12788 | 19.44444 | 0 | 0 | Neutral |
| testneg_9116 | -0.21 | -0.15 | -1.902903 | -3 | 0.265756 | -3 | 0.1254 | 0.08697 | 0.96396 | 0.7628 | 10.41667 | 0.13021055 | 0 | Neutral |
| testneg_9117 | 0 | 0 | 5.501633 | -3 | 7.071894 | -3 | 0.12582 | 0.12788 | 0.28341 | 0.1005 | 13.51351 | 0.579682659 | 82.2 | Disease_causing |
| testneg_9118 | 0 | -0.01 | 1.795817 | -3 | -0.193992 | -3 | 0.16053 | 0.15049 | 0.1336 | 0.15669 | 30.99664 | 0.094093766 | 0 | Neutral |
| testneg_9119 | -0.15 | -0.04 | 4.218719 | -3 | 4.343105 | -3 | 0.17258 | 0.12788 | 0.13698 | 0.12788 | 13.49076 | 0.275153469 | 0.8 | Neutral |
| testneg_9120 | -0.09 | 0.3 | 0.850367 | -3 | 1.142068 | -3 | 0.29024 | 0.1472 | 0.38428 | 0.38566 | 23.91309 | 0.132251263 | 0 | Neutral |
| testneg_9121 | -0.08 | -0.03 | 4.142037 | -3 | 0.256741 | -3 | 0.95535 | 0.12788 | 0.3012 | 0.10317 | 11.14837 | 0.05 | 0 | Neutral |
| testneg_9122 | 0.17 | 0.38 | -0.644713 | -3 | 7.320532 | -3 | 0.14511 | 0.12788 | 0.19898 | 0.12788 | 19.03093 | 0.29 | 2.6 | Neutral |
| testneg_9123 | -0.05 | -0.29 | 0.055751 | -3 | 6.785733 | -3 | 0.19898 | 0.12788 | 0.16524 | 0.12788 | 5.69444 | 0.266008826 | 2.6 | Neutral |
| testneg_9124 | 0.03 | -1.03 | -0.561373 | -3 | 0.590862 | -3 | 0.08332 | 0.09046 | 0.19898 | 0.20319 | 6.03357 | 0.050956044 | 0 | Neutral |
| testneg_9125 | 0.18 | -1.05 | 3.769006 | -3 | 1.205691 | -3 | 0.44479 | 0.20004 | 0.18559 | 0.09173 | 18.33333 | 0.096214286 | 0 | Neutral |
| testneg_9126 | -0.04 | 1.61 | 13.172408 | -3 | 7.325251 | -3 | 0.13433 | 0.09578 | 0.16601 | 0.14015 | 36.74254 | 0.680651261 | 97.4 | Disease_causing |
| testneg_9127 | 0.09 | -0.67 | -0.874528 | -3 | 5.595391 | -3 | 0.1592 | 0.12788 | 0.16709 | 0.08659 | 19.95887 | 0.11 | 0 | Neutral |
| testneg_9128 | 0 | 0 | 0.635932 | -3 | 2.067861 | 2.067861 | 0.24516 | 0.11809 | 0.28537 | 0.12788 | 19.1869 | 0.03 | 0 | Neutral |
| testneg_9129 | -0.02 | -0.13 | -0.586534 | -0.586534 | 1.272902 | -3 | 0.09439 | 0.1884 | 0.19898 | 0.12788 | 44.98717 | 0.041088235 | 0 | Neutral |
| testneg_9130 | 0 | 0 | 1.284685 | -3 | 1.967428 | -3 | 0.53496 | 0.13586 | 0.42797 | 0.17974 | 10 | 0.041073749 | 0 | Neutral |
| testneg_9131 | -0.07 | -0.38 | 2.948419 | -3 | 1.016702 | -3 | 0.04863 | 0.12149 | 0.09306 | 0.09084 | 21.33768 | 0.07 | 0 | Neutral |
| testneg_9132 | -0.08 | -0.03 | 0.579763 | -3 | 0.495531 | 0.330554 | 0.09978 | 0.12788 | 0.06431 | 0.12788 | 13.5 | 0.02 | 0 | Neutral |
| testneg_9133 | 0.03 | 0.12 | 6.409576 | -3 | 1.291726 | -3 | 0.10241 | 0.08165 | 0.29525 | 0.12788 | 27.5 | 0.16525 | 0 | Neutral |
| testneg_9134 | 0.05 | 0.29 | 1.87439 | -3 | 0.725745 | -3 | 0.07189 | 0.11325 | 0.06603 | 0.08071 | 13.51351 | 0.01 | 0 | Neutral |
| testneg_9135 | 0.22 | -2.66 | 6.743317 | -3 | 1.275456 | -3 | 0.50649 | 0.11394 | 0.30103 | 0.1462 | 17.78631 | 0.32 | 4.8 | Neutral |
| testneg_9136 | 0 | 0 | 2.036514 | -3 | 1.326969 | -3 | 0.16544 | 0.1313 | 0.12642 | 0.11218 | 11.25 | 0.01 | 0 | Neutral |
| testneg_9137 | 0.06 | 0.18 | 2.716241 | -3 | -0.253658 | -3 | 0.12591 | 0.09845 | 0.07293 | 0.12989 | 10.28409 | 0.02 | 0 | Neutral |
| testneg_9138 | 0 | 0 | -0.002747 | -3 | 4.548656 | -3 | 0.07394 | 0.09408 | 0.18559 | 0.13755 | 14.95396 | 0.174004945 | 0 | Neutral |
| testneg_9139 | 0.09 | -0.67 | 4.020687 | -3 | 1.866705 | -3 | 0.34001 | 0.12788 | 0.14938 | 0.12747 | 20.92724 | 0.040190374 | 0 | Neutral |
| testneg_9140 | 0 | 0 | 1.371909 | -3 | -0.560973 | -3 | 0.47733 | 0.08388 | 0.10485 | 0.09565 | 13.27141 | 0.010526316 | 0 | Neutral |
| testneg_9141 | 0 | 0 | 1.686834 | -3 | 3.388454 | -3 | 0.29828 | 0.12788 | 0.05291 | 0.08633 | 19.13908 | 0.030089568 | 0 | Neutral |
| testneg_9142 | 0.05 | 0.29 | 3.783684 | -3 | 0.828771 | -3 | 0.39399 | 0.12788 | 0.71844 | 0.20204 | 15.50666 | 0.122881481 | 0 | Neutral |
| testneg_9143 | 0.01 | 0.8 | 4.81138 | -3 | 6.247828 | -3 | 0.284 | 0.15339 | 0.15856 | 0.10994 | 20.90643 | 0.469208993 | 42.8 | Neutral |
| testneg_9144 | 0 | -0.01 | 1.961035 | -3 | 1.823474 | -3 | 0.55105 | 0.43208 | 0.16355 | 0.20781 | 10 | 0.092166063 | 0 | Neutral |
| testneg_9145 | 0 | 0 | 0.348328 | -3 | 1.810934 | -3 | 0.28878 | 0.16765 | 0.5184 | 0.11889 | 10 | 0.010034965 | 0 | Neutral |
| testneg_9146 | -0.01 | -0.79 | -0.267483 | -3 | 0.630219 | 0.630219 | 0.47887 | 0.09483 | 0.09442 | 0.1153 | 16.62115 | 0.03 | 0 | Neutral |
| testneg_9147 | 0.14 | -1.01 | 3.061839 | -3 | 1.850753 | -3 | 0.78537 | 0.33817 | 0.16057 | 0.12305 | 18.54167 | 0.141625 | 0 | Neutral |
| testneg_9148 | 0.07 | 0.38 | 0.829687 | -3 | 4.717965 | -3 | 0.06841 | 0.12788 | 0.10895 | 0.10442 | 8.97898 | 0.08468092 | 0 | Neutral |
| testneg_9149 | 0.02654545 | -0.3807273 | 14.874927 | -3 | 3.993033 | -3 | 0.06845 | 0.13843 | 0.19898 | 0.12788 | 14.16667 | 0.583261555 | 84.8 | Disease_causing |

| | | | | | | | | | | | | | | |
|---|---|---|---|---|---|---|---|---|---|---|---|---|---|---|
| testneg_9150 | 0.22 | -2.66 | 7.635846 | -3 | 2.585384 | -3 | 0.19898 | 0.16603 | 0.17039 | 0.12788 | 21.02328 | 0.508289627 | 58.2 | Disease_causing |
| testneg_9151 | 0.1 | 1.9 | 6.059259 | -3 | 6.830695 | -3 | 0.19898 | 0.12788 | 0.53016 | 0.10373 | 20.16957 | 0.581214286 | 84 | Disease_causing |
| testneg_9152 | 0 | -0.32 | -0.590938 | -3 | -1.428127 | -3 | 0.06513 | 0.12353 | 0.11564 | 0.1184 | 46.25 | 0.02 | 0 | Neutral |
| testneg_9153 | -0.15 | -0.04 | 0.888493 | -3 | 0.379365 | -3 | 0.89357 | 0.22348 | 0.06547 | 0.09123 | 17.21847 | 0.055 | 0 | Neutral |
| testneg_9154 | 0.03 | -1.03 | 5.357825 | -3 | -1.159974 | -3 | 0.04305 | 0.06354 | 0.09515 | 0.07784 | 13.51351 | 0.14 | 0 | Neutral |
| testneg_9155 | 0 | 0 | 1.383951 | -3 | 3.684842 | -3 | 0.08683 | 0.26961 | 0.16601 | 0.14015 | 32.18771 | 0.366614789 | 11.6 | Neutral |
| testneg_9156 | 0.17 | 0.38 | 3.544414 | -3 | 2.804418 | -3 | 0.18153 | 0.10845 | 0.32416 | 0.09959 | 16.07106 | 0.107536569 | 0 | Neutral |
| testneg_9157 | 0 | 0 | 1.474437 | -3 | 2.661909 | -3 | 0.54648 | 0.19042 | 0.92776 | 0.09677 | 10 | 0.043734127 | 0 | Neutral |
| testneg_9158 | 0.14 | -1.85 | 3.31974 | -3 | 5.241235 | -3 | 0.19898 | 0.12788 | 0.89178 | 0.23115 | 19.5422 | 0.543828833 | 71.4 | Disease_causing |
| testneg_9159 | 0 | 0 | 1.66597 | -3 | 0.046064 | -3 | 0.66027 | 0.23984 | 0.2543 | 0.15588 | 11.875 | 0.053699187 | 0 | Neutral |
| testneg_9160 | 0 | 0 | 1.205908 | -3 | 0.200393 | -3 | 0.32333 | 0.23356 | 0.19898 | 0.08714 | 11.875 | 0.01 | 0 | Neutral |
| testneg_9161 | 0.21 | 0.15 | 4.635288 | -3 | 2.833872 | -3 | 0.38749 | 0.1076 | 0.19898 | 0.12788 | 27.61749 | 0.14 | 0 | Neutral |
| testneg_9162 | 0.22 | -2.66 | 3.810217 | -3 | 2.658618 | -3 | 0.10562 | 0.09329 | 0.113 | 0.12788 | 29.90985 | 0.15 | 0 | Neutral |
| testneg_9163 | 0.15 | 0.04 | -0.180109 | -3 | 1.795418 | -3 | 0.11458 | 0.16496 | 0.11055 | 0.19829 | 14.56463 | 0.152440476 | 0 | Neutral |
| testneg_9164 | 0 | -0.32 | 5.213254 | -3 | 0.312359 | -3 | 0.62172 | 0.18135 | 0.11261 | 0.13673 | 14.18205 | 0.27 | 0.6 | Neutral |
| testneg_9165 | 0.07 | 0.38 | -0.919669 | -3 | 0.955677 | 0.955677 | 0.83936 | 0.42313 | 0.10034 | 0.10906 | 14.24934 | 0.060253442 | 0 | Neutral |
| testneg_9166 | 0.01 | 0.8 | 3.997628 | -3 | 1.349086 | -3 | 0.66799 | 0.12788 | 0.17199 | 0.10586 | 13.20767 | 0.07 | 0 | Neutral |
| testneg_9167 | -0.04 | 1.36 | -3.143764 | -3.143764 | 4.486783 | -3 | 0.03137 | 0.12788 | 0.19898 | 0.12788 | 55.55556 | 0.21 | 0 | Neutral |
| testneg_9168 | -0.03 | -0.1 | 3.350172 | -3 | 3.83753 | -3 | 0.13331 | 0.13638 | 0.30209 | 0.14229 | 21.53664 | 0.4148187 | 23.6 | Neutral |
| testneg_9169 | 0.2 | -1.26 | 3.203428 | -3 | 0.22996 | -1.285414 | 0.35325 | 0.12614 | 0.0693 | 0.09293 | 10.41667 | 0.05 | 0 | Neutral |
| testneg_9170 | 0 | 0 | 0.995978 | -3 | 0.725853 | -3 | 0.96341 | 0.18404 | 0.16736 | 0.10113 | 18.03052 | 0.011864629 | 0 | Neutral |
| testneg_9171 | 0.03 | 0.1 | 0.500506 | -3 | 0.019844 | -3 | 0.68233 | 0.58998 | 0.16737 | 0.12788 | 10 | 0.05 | 0 | Neutral |
| testneg_9172 | -0.03 | -0.1 | 4.889785 | -3 | -0.187467 | -0.187467 | 0.09048 | 0.62605 | 0.12314 | 0.11709 | 17.77778 | 0.22 | 0 | Neutral |
| testneg_9173 | 0.15 | -0.28 | 0.676746 | -3 | 0.196098 | -3 | 0.61306 | 0.15258 | 0.12682 | 0.11357 | 18.29495 | 0.03 | 0 | Neutral |
| testneg_9174 | 0.11 | -0.19 | 4.344872 | -3 | 1.056615 | -3 | 0.28732 | 0.11223 | 0.09471 | 0.12788 | 27.64437 | 0.03 | 0 | Neutral |
| testneg_9175 | 0.17 | 0.38 | 0.212654 | -3 | -0.354552 | -3 | 0.17173 | 0.15363 | 0.09815 | 0.12788 | 22.89111 | 0.05 | 0 | Neutral |
| testneg_9176 | -0.01 | 0.57 | 0.811998 | -3 | -1.898366 | -1.898366 | 0.21848 | 0.11049 | 0.11983 | 0.24162 | 14.54757 | 0.060555556 | 0 | Neutral |
| testneg_9177 | 0 | -0.01 | 0.302881 | -3 | 2.140684 | -3 | 0.56285 | 0.18619 | 0.30823 | 0.17188 | 12.40248 | 0.081178605 | 0 | Neutral |
| testneg_9178 | 0.21 | 0.15 | 7.822266 | -3 | 0.651852 | -3 | 0.23005 | 0.37216 | 0.55618 | 0.11352 | 11.67232 | 0.41 | 19.2 | Neutral |
| testneg_9179 | 0 | 0 | 1.323164 | -3 | 6.597532 | -3 | 0.22761 | 0.14229 | 0.27887 | 0.18663 | 19.44444 | 0.443049129 | 32.8 | Neutral |
| testneg_9180 | 0 | 0 | 0.097356 | -3 | -2.062376 | -2.062376 | 0.14485 | 0.16945 | 0.11315 | 0.12788 | 24.68926 | 0.03 | 0 | Neutral |
| testneg_9181 | 0.07 | 0.7 | 5.516867 | -3 | -2.175808 | -3 | 0.22816 | 0.10878 | 0.13704 | 0.10054 | 18.33333 | 0.12 | 0 | Neutral |
| testneg_9182 | 0.02 | -0.63 | 2.326926 | -3 | 6.956883 | -3 | 0.07558 | 0.12788 | 0.12131 | 0.12533 | 49.88402 | 0.28 | 1.6 | Neutral |
| testneg_9183 | 0.02 | -0.63 | 3.104093 | -3 | 0.922605 | -3 | 0.65564 | 0.11468 | 0.2152 | 0.12788 | 18.33333 | 0.021125 | 0 | Neutral |
| testneg_9184 | -0.04 | 0.89 | 5.832243 | -3 | 1.406345 | -3 | 0.19898 | 0.12788 | 0.7062 | 0.83002 | 10.90717 | 0.45 | 36 | Neutral |
| testneg_9185 | 0 | 0 | 1.040406 | -3 | 0.655763 | 0.655763 | 0.41987 | 0.67373 | 0.09087 | 0.12788 | 10 | 0.04 | 0 | Neutral |
| testneg_9186 | 0.03 | 0.12 | 2.593624 | -3 | -1.686029 | -3 | 0.22364 | 0.12788 | 0.09964 | 0.10384 | 13.51351 | 0.01 | 0 | Neutral |
| testneg_9187 | 0 | 0 | 0.795894 | -3 | 3.546645 | -3 | 0.99448 | 0.67288 | 0.18502 | 0.21329 | 10.83333 | 0.276338964 | 1 | Neutral |
| testneg_9188 | 0 | 0 | -0.076686 | -3 | 0.081615 | 0.081615 | 0.84596 | 0.10633 | 0.14491 | 0.10508 | 30.55556 | 0.03 | 0 | Neutral |
| testneg_9189 | 0 | 0 | 0.64648 | 0.64648 | 1.307704 | -3 | 0.18055 | 0.12564 | 0.64749 | 0.14763 | 34.84448 | 0.084038954 | 0 | Neutral |
| testneg_9190 | 0 | 0 | 0.123542 | -3 | 2.268054 | -3 | 0.99915 | 0.96333 | 0.55155 | 0.10709 | 5.16667 | 0.110444664 | 0 | Neutral |
| testneg_9191 | 0.14 | -0.69 | 1.912827 | -3 | 5.957971 | -3 | 0.1203 | 0.13872 | 0.20701 | 0.10345 | 15.09363 | 0.193254762 | 0 | Neutral |
| testneg_9192 | 0 | -0.01 | 7.854373 | -3 | 0.195289 | -3 | 0.4113 | 0.2303 | 0.09972 | 0.2187 | 33.726 | 0.55815 | 77 | Disease_causing |
| testneg_9193 | 0.04 | -0.23 | 0.526783 | -3 | 1.247016 | -3 | 0.31507 | 0.61057 | 0.31025 | 0.09491 | 9.30313 | 0.04 | 0 | Neutral |
| testneg_9194 | 0.03 | 0.12 | 3.408834 | -3 | 4.256646 | -3 | 0.19479 | 0.12788 | 0.08723 | 0.09503 | 35.34347 | 0.123333333 | 0 | Neutral |
| testneg_9195 | 0 | 0 | -0.784713 | -0.784713 | 1.570658 | -3 | 0.54899 | 0.31462 | 0.51622 | 0.11809 | 14.78773 | 0.015 | 0 | Neutral |
| testneg_9196 | 0.01 | 0.8 | 7.50868 | -3 | -0.018797 | -3 | 0.40283 | 0.34309 | 0.11891 | 0.11186 | 19.24569 | 0.439282051 | 28.2 | Neutral |
| testneg_9197 | 0.02 | 0.13 | 4.492963 | -3 | 5.180313 | -3 | 0.54648 | 0.19042 | 0.19898 | 0.12788 | 12.29167 | 0.433303571 | 32.4 | Neutral |
| testneg_9198 | 0 | 0 | 1.229821 | -3 | 0.067806 | -3 | 0.46091 | 0.13852 | 0.19898 | 0.19444 | 22.77778 | 0.071661255 | 0 | Neutral |
| testneg_9199 | 0 | 0 | 0.720647 | -3 | 0.082232 | -3 | 0.161 | 0.11597 | 0.08475 | 0.19889 | 13.02635 | 0.04 | 0 | Neutral |
| testneg_9200 | -0.02 | 0.63 | 3.574998 | -3 | 6.36911 | -3 | 0.32974 | 0.12245 | 0.17412 | 0.10595 | 7.02384 | 0.366869048 | 15.6 | Neutral |
| testneg_9201 | -0.15 | -0.06 | 5.384158 | -3 | 0.726355 | -3 | 0.15 | 0.18963 | 0.09439 | 0.1884 | 20.98109 | 0.563306805 | 78.4 | Disease_causing |
| testneg_9202 | 0.16 | 0.08 | 1.986437 | -3 | 1.795415 | -3 | 0.69666 | 0.11055 | 0.65717 | 0.16156 | 19.85904 | 0.102088278 | 0 | Neutral |
| testneg_9203 | 0 | 0 | 0.64288 | -3 | 2.737712 | -3 | 0.57188 | 0.34038 | 0.44385 | 0.1311 | 4.44444 | 0.103646864 | 0 | Neutral |
| testneg_9204 | -0.07 | -0.38 | 2.997184 | -3 | 3.254411 | 0.336354 | 0.17394 | 0.15823 | 0.12121 | 0.4142 | 10.41667 | 0.569690149 | 78.2 | Disease_causing |
| testneg_9205 | -0.17 | -0.38 | 1.19692 | -3 | -2.351627 | -3 | 0.08224 | 0.29179 | 0.16646 | 0.16629 | 24.35262 | 0.26 | 0 | Neutral |
| testneg_9206 | 0.09 | -0.3 | 6.316011 | -3 | -0.658155 | -3 | 0.3038 | 0.11278 | 0.05087 | 0.09682 | 17.21847 | 0.14 | 0.2 | Neutral |
| testneg_9207 | -0.04 | 1.36 | -1.284127 | -3 | 2.364808 | -3 | 0.11096 | 0.10074 | 0.16621 | 0.49845 | 39.6507 | 0.203625 | 0 | Neutral |
| testneg_9208 | 0 | 0 | 1.433601 | -3 | 1.119912 | -3 | 0.85984 | 0.40044 | 0.09724 | 0.09172 | 8.59957 | 0.01 | 0 | Neutral |
| testneg_9209 | -0.02 | 0.63 | -1.27031 | -3 | -2.576045 | -2.576045 | 0.10977 | 0.31695 | 0.25275 | 0.12788 | 5.69444 | 0.06 | 0 | Neutral |
| testneg_9210 | -0.01 | -0.8 | 1.016479 | 1.016479 | -0.682977 | -3 | 0.12649 | 0.12788 | 0.05444 | 0.12788 | 26.34995 | 0.04 | 0 | Neutral |
| testneg_9211 | 0 | 0 | 0.707491 | -3 | 3.88874 | -3 | 0.43762 | 0.12788 | 0.09618 | 0.10294 | 17.21847 | 0.06059127 | 0 | Neutral |
| testneg_9212 | 0.01 | 0.8 | 2.217361 | -3 | 0.095102 | -3 | 0.11829 | 0.12788 | 0.19898 | 0.13495 | 21.57429 | 0.062747199 | 0 | Neutral |
| testneg_9213 | 0.03 | 1.22 | 0.949523 | 0.949523 | 5.710272 | -3 | 0.10056 | 0.12608 | 0.508 | 0.1281 | 9.01337 | 0.259761905 | 0 | Neutral |
| testneg_9214 | 0 | 0 | -0.216528 | -3 | 1.590262 | -3 | 0.40612 | 0.45135 | 0.44385 | 0.1311 | 4.72222 | 0.040408333 | 0 | Neutral |
| testneg_9215 | 0.03 | 1.22 | 4.663109 | -3 | 0.032284 | -3 | 0.23292 | 0.15913 | 0.23393 | 0.12567 | 10.83333 | 0.130052632 | 0 | Neutral |
| testneg_9216 | 0 | 0 | 0.463587 | -3 | 1.712042 | -3 | 0.36494 | 0.12763 | 0.31004 | 0.09727 | 18.09499 | 0 | 0 | Neutral |
| testneg_9217 | 0.14 | -0.37 | 3.24283 | -3 | 3.749392 | -3 | 0.20986 | 0.10633 | 0.10317 | 0.14954 | 13.02635 | 0.390792237 | 17.8 | Neutral |
| testneg_9218 | 0.14 | -1.01 | 2.16064 | -3 | 0.955677 | 0.955677 | 0.16734 | 0.15176 | 0.10034 | 0.10906 | 11.25 | 0.04 | 0 | Neutral |
| testneg_9219 | 0 | -0.32 | -2.076593 | -3 | 6.279292 | -3 | 0.1276 | 0.30104 | 0.09488 | 0.10426 | 13.93346 | 0.3 | 1.8 | Neutral |
| testneg_9220 | 0.09 | -0.3 | 3.181713 | -3 | 5.296637 | -3 | 0.44329 | 0.12788 | 0.84365 | 0.12788 | 21.42455 | 0.220229703 | 0.4 | Neutral |
| testneg_9221 | 0 | 0 | -0.376647 | -3 | 5.917396 | -3 | 0.05087 | 0.09682 | 0.12933 | 0.12788 | 18.88889 | 0.111180556 | 0 | Neutral |
| testneg_9222 | 0.16 | -1.37 | 7.137326 | -3 | -0.456172 | -0.63117 | 0.48936 | 0.10963 | 0.19898 | 0.1054 | 10.22898 | 0.260279503 | 1.8 | Neutral |
| testneg_9223 | 0 | 0.32 | 0.733606 | -3 | 2.378984 | -3 | 0.51398 | 0.38821 | 0.11891 | 0.11186 | 16.23481 | 0.080707143 | 0 | Neutral |
| testneg_9224 | 0 | -0.01 | 0.611606 | -3 | 0.6838 | 0.6838 | 0.9149 | 0.27793 | 0.67327 | 0.23543 | 11.875 | 0.062727202 | 0 | Neutral |
| testneg_9225 | -0.15 | -0.06 | 2.266671 | -3 | 1.658258 | -3 | 0.09058 | 0.50106 | 0.3414 | 0.13575 | 17.46678 | 0.283644231 | 0.2 | Neutral |
| testneg_9226 | 0.05 | -0.81 | 4.114832 | -3 | 4.920318 | -3 | 0.20655 | 0.15317 | 0.14668 | 0.17962 | 12.6183 | 0.795303152 | 100 | Disease_causing |
| testneg_9227 | 0.18 | 0.16 | 5.910362 | -3 | 0.675511 | -3 | 0.73395 | 0.1705 | 0.23723 | 0.1339 | 13.02635 | 0.32213014 | 3.8 | Neutral |
| testneg_9228 | 0 | 0 | 2.175318 | -3 | 0.172291 | -3 | 0.41698 | 0.10384 | 0.0138 | 0.06769 | 19.13479 | 0.01 | 0 | Neutral |
| testneg_9229 | 0.03 | 0.1 | -0.187467 | -0.187467 | 0.239025 | -3 | 0.12314 | 0.11709 | 0.1215 | 0.15825 | 18.33333 | 0.074207207 | 0 | Neutral |
| testneg_9230 | 0 | 0 | -0.02198 | -0.02198 | 5.50519 | -3 | 0.15509 | 0.1326 | 0.35984 | 0.11129 | 17.21847 | 0.12 | 0 | Neutral |
| testneg_9231 | 0.03 | 1.22 | 2.244475 | -3 | 2.767022 | -3 | 0.34406 | 0.09894 | 0.46791 | 0.7316 | 9.55396 | 0.228547619 | 0 | Neutral |
| testneg_9232 | 0 | 0 | 1.659032 | 1.659032 | 0.506986 | -3 | 0.755 | 0.12788 | 0.65857 | 0.26294 | 23.90097 | 0.121547619 | 0 | Neutral |
| testneg_9233 | -0.07 | -0.38 | 6.939074 | -3 | 2.51125 | -3 | 0.07467 | 0.09224 | 0.31344 | 0.12788 | 13.51351 | 0.334166667 | 7.8 | Neutral |
| testneg_9234 | -0.01 | 0.57 | 1.162064 | -3 | 2.846841 | -3 | 0.64608 | 0.35524 | 0.2852 | 0.12788 | 10.41667 | 0.116353268 | 0 | Neutral |
| testneg_9235 | -0.16 | 0.37 | 5.039547 | -3 | 5.093198 | -3 | 0.42512 | 0.29333 | 0.99859 | 0.12788 | 12.19153 | 0.598939519 | 89 | Disease_causing |
| testneg_9236 | 0.15 | 0.04 | -0.962385 | -3 | 0.006704 | -3 | 0.60285 | 0.15179 | 0.30925 | 0.10335 | 12.30981 | 0.043080565 | 0 | Neutral |
| testneg_9237 | -0.1 | -1.9 | 1.091771 | -3 | 6.707128 | -3 | 0.7174 | 0.15815 | 0.31025 | 0.09491 | 15.90395 | 0.31 | 2 | Neutral |
| testneg_9238 | -0.14 | 1.01 | 6.055805 | -3 | 6.184485 | -3 | 0.96396 | 0.7628 | 0.11134 | 0.13818 | 20.59685 | 0.84 | 100 | Disease_causing |
| testneg_9239 | -0.05 | -0.29 | 1.376936 | 0.165609 | 4.260553 | -3 | 0.89992 | 0.10231 | 0.13624 | 0.09458 | 10 | 0.07875 | 0 | Neutral |
| testneg_9240 | 0.03 | -1.03 | 2.478795 | -3 | -0.139258 | -3 | 0.1327 | 0.09885 | 0.31025 | 0.09491 | 21.47432 | 0.02 | 0 | Neutral |
| testneg_9241 | 0 | 0 | 0.28875 | -3 | 0.357284 | -3 | 0.40179 | 0.12112 | 0.08973 | 0.0992 | 17.79605 | 0.000185249 | 0 | Neutral |
| testneg_9242 | 0 | 0 | 1.941287 | -3 | 1.678575 | -3 | 0.24977 | 0.11262 | 0.5958 | 0.0929 | 11.12646 | 0.000549402 | 0 | Neutral |
| testneg_9243 | 0 | -0.01 | -0.3595 | -3 | -0.209426 | -3 | 0.19898 | 0.12788 | 0.99531 | 0.86203 | 15.41667 | 0.095 | 0 | Neutral |
| testneg_9244 | 0.21 | 0.15 | -0.20606 | -3 | 0.299785 | -3 | 0.32299 | 0.12788 | 0.64152 | 0.15239 | 25.94604 | 0.084594988 | 0 | Neutral |
| testneg_9245 | -0.1 | -1.9 | 0.470705 | -3 | 1.313372 | -3 | 0.62858 | 0.16645 | 0.12836 | 0.09736 | 14.27748 | 0.07 | 0 | Neutral |
| testneg_9246 | 0.00027273 | -0.0447273 | 0.427324 | -3 | 1.794179 | -3 | 0.68863 | 0.09663 | 0.4722 | 0.13321 | 36.69636 | 0.070750661 | 0 | Neutral |
| testneg_9247 | -0.03 | -0.1 | 2.078666 | -3 | 4.268449 | 3.112174 | 0.17576 | 0.12193 | 0.4069 | 0.3841 | 21.15585 | 0.520280176 | 62 | Disease_causing |
| testneg_9248 | 0.03 | -1.03 | 3.488962 | -3 | 6.913969 | -3 | 0.59245 | 0.12788 | 0.15267 | 0.12788 | 34.82436 | 0.379375 | 15.6 | Neutral |
| testneg_9249 | 0.03 | -1.03 | 6.478196 | -3 | 0.722971 | -3 | 0.32188 | 0.08974 | 0.19898 | 0.12788 | 25.74074 | 0.19 | 0.4 | Neutral |
| testneg_9250 | 0 | 0 | 1.1308 | -3 | -2.747914 | -3 | 0.19898 | 0.11185 | 0.03651 | 0.06141 | 41.57379 | 0.01 | 0 | Neutral |
| testneg_9251 | -0.01 | 1.44 | 1.762821 | 1.762821 | 1.795418 | -3 | 0.1375 | 0.112 | 0.11055 | 0.19829 | 11.74429 | 0.212114286 | 0.4 | Neutral |
| testneg_9252 | 0.01 | 0.8 | 6.431918 | -3 | 6.139578 | -3 | 0.19364 | 0.14027 | 0.15406 | 0.1454 | 30.55556 | 0.715794118 | 98.6 | Disease_causing |
| testneg_9253 | 0 | 0 | -0.098829 | -3 | 1.979502 | -3 | 0.1602 | 0.11262 | 0.18498 | 0.12788 | 25.10109 | 0.01 | 0 | Neutral |
| testneg_9254 | 0.2 | -1.26 | 3.203428 | -3 | 1.755966 | -3 | 0.35325 | 0.12614 | 0.17231 | 0.1358 | 10.83333 | 0.15 | 0 | Neutral |
| testneg_9255 | -0.14 | 1.01 | 5.525252 | -3 | 2.166075 | -3 | 0.37212 | 0.11323 | 0.86552 | 0.33462 | 17.42316 | 0.412416667 | 20.4 | Neutral |
| testneg_9256 | 0.22 | -2.66 | 4.102892 | -3 | 5.804724 | -3 | 0.96704 | 0.16878 | 0.23624 | 0.12788 | 9.26617 | 0.448333333 | 34.8 | Neutral |
| testneg_9257 | 0.03 | 0.12 | 4.41565 | -3 | 6.245625 | -3 | 0.08337 | 0.18096 | 0.05535 | 0.12788 | 40.11276 | 0.44 | 32.8 | Neutral |
| testneg_9258 | 0.12 | -1.16 | 2.187673 | -3 | 2.246903 | -3 | 0.29828 | 0.12788 | 0.1019 | 0.17526 | 30.55556 | 0.179188596 | 0 | Neutral |
| testneg_9259 | 0 | 0 | 0.60253 | -3 | 4.085318 | -3 | 0.50687 | 0.11283 | 0.3828 | 0.5569 | 10.41667 | 0.205501865 | 0 | Neutral |
| testneg_9260 | -0.07 | -0.38 | 2.948419 | -3 | -0.462963 | -0.462963 | 0.04863 | 0.12149 | 0.1946 | 0.14591 | 14.16667 | 0.16 | 0 | Neutral |
| testneg_9261 | 0 | 0 | -0.163107 | -3 | 3.009853 | -3 | 0.21289 | 0.08709 | 0.50741 | 0.17915 | 15.87989 | 0.103215278 | 0 | Neutral |
| testneg_9262 | -0.21 | -0.15 | -1.808697 | -3 | 4.833511 | -3 | 0.14386 | 0.12788 | 0.19898 | 0.07312 | 37.57733 | 0.17 | 0 | Neutral |
| testneg_9263 | 0.05 | 1.52 | 3.689006 | -3 | 2.094119 | -3 | 0.1755 | 0.10455 | 0.64152 | 0.15239 | 38.25843 | 0.208 | 0.2 | Neutral |
| testneg_9264 | -0.15 | -0.04 | 1.43553 | -3 | 0.188097 | -0.833744 | 0.13565 | 0.46572 | 0.31646 | 0.09844 | 6.29554 | 0.090277825 | 0 | Neutral |
| testneg_9265 | 0.02 | 0.45 | 2.796708 | -3 | 1.842694 | -3 | 0.11966 | 0.16464 | 0.19898 | 0.12788 | 29.63785 | 0.07 | 0 | Neutral |
| testneg_9266 | 0 | -0.01 | 0.64147 | -3 | 0.944088 | -3 | 0.35967 | 0.09418 | 0.71025 | 0.1632 | 7.78257 | 0.020593809 | 0 | Neutral |
| testneg_9267 | -0.01 | -0.8 | 7.657206 | -3 | 3.05843 | 3.05843 | 0.19898 | 0.12788 | 0.74136 | 0.1049 | 10.83333 | 0.58 | 86.8 | Disease_causing |
| testneg_9268 | -0.19 | -0.51 | 3.156196 | -3 | 0.550262 | -3 | 0.34225 | 0.12788 | 0.31701 | 0.09986 | 10.83333 | 0.06 | 0 | Neutral |
| testneg_9269 | 0 | 0 | 0.537223 | -3 | -2.323867 | -3 | 0.08186 | 0.11813 | 0.07595 | 0.11822 | 35.24977 | 0.01 | 0 | Neutral |
| testneg_9270 | -0.04 | -2.01 | 5.968963 | -3 | 2.846083 | -3 | 0.33256 | 0.14909 | 0.1059 | 0.1265 | 19.44444 | 0.409827586 | 21 | Neutral |
| testneg_9271 | 0 | -0.01 | 1.708082 | -3 | 0.554274 | -3 | 0.64721 | 0.8758 | 0.32963 | 0.1035 | 12.61905 | 0.05 | 0 | Neutral |

| | | | | | | | | | | | | | | |
|---|---|---|---|---|---|---|---|---|---|---|---|---|---|---|
| testneg_9272 | -0.02 | -0.45 | 6.897588 | -3 | 6.365761 | -3 | 0.20914 | 0.10089 | 0.21348 | 0.10616 | 23.19718 | 0.54 | 65.4 | Disease_causing |
| testneg_9273 | 0.22 | -2.66 | 4.376841 | -3 | 1.162821 | -3 | 0.34859 | 0.09799 | 0.75481 | 0.12788 | 25.42027 | 0.11 | 0 | Neutral |
| testneg_9274 | 0 | 0 | 2.047211 | -3 | 3.905173 | -3 | 0.16823 | 0.10796 | 0.16057 | 0.12305 | 18.95833 | 0.051176471 | 0 | Neutral |
| testneg_9275 | 0.03 | -1.03 | 2.707374 | -3 | 3.163542 | -3 | 0.33127 | 0.49952 | 0.10751 | 0.08512 | 14.30802 | 0.179761772 | 0 | Neutral |
| testneg_9276 | 0 | 0 | 1.347 | -3 | 0.054796 | -3 | 0.16252 | 0.11134 | 0.32531 | 0.12351 | 13.08587 | 0 | 0 | Neutral |
| testneg_9277 | 0 | -0.01 | 0.855893 | -3 | 1.613091 | -3 | 0.26916 | 0.13285 | 0.53922 | 0.12788 | 12.63889 | 0.01 | 0 | Neutral |
| testneg_9278 | 0.18 | -1.05 | 3.274763 | -3 | 0.018583 | -3 | 0.49512 | 0.08696 | 0.37293 | 0.25159 | 12.81915 | 0.180757615 | 0 | Neutral |
| testneg_9279 | 0 | 0 | 2.297055 | -3 | 1.847923 | -3 | 0.30739 | 0.16306 | 0.13118 | 0.08646 | 18.04499 | 0.02 | 0 | Neutral |
| testneg_9280 | -0.18 | -0.16 | 0.369791 | -3 | 2.185335 | -3 | 0.66227 | 0.11002 | 0.19898 | 0.12788 | 20.218 | 0.06 | 0 | Neutral |
| testneg_9281 | 0 | 0 | -0.43021 | -3 | 0.694077 | -3 | 0.17041 | 0.12378 | 0.21235 | 0.1449 | 20.83333 | 0.035855357 | 0 | Neutral |
| testneg_9282 | 0 | -0.01 | 8.609409 | -3 | 3.181713 | -3 | 0.49012 | 0.12138 | 0.44329 | 0.12788 | 10.75027 | 0.575939394 | 85.2 | Disease_causing |
| testneg_9283 | 0.06 | 0.18 | 2.453732 | -3 | 0.467505 | -3 | 0.18744 | 0.09392 | 0.0676 | 0.12788 | 10.73298 | 0.015 | 0 | Neutral |
| testneg_9284 | -0.06 | -0.18 | 1.437183 | -3 | -0.020693 | -3 | 0.19898 | 0.12788 | 0.80034 | 0.10518 | 18.6768 | 0.01 | 0 | Neutral |
| testneg_9285 | 0 | 0 | 1.076963 | -3 | 5.3866 | -3 | 0.20011 | 0.27949 | 0.05716 | 0.12491 | 38.11308 | 0.145166667 | 0 | Neutral |
| testneg_9286 | 0 | 0 | -0.001756 | -3 | -0.136533 | -3 | 0.16662 | 0.11306 | 0.05932 | 0.09551 | 18.82092 | 0.01 | 0 | Neutral |
| testneg_9287 | -0.19 | 0.25 | 0.456275 | 0.456275 | 0.71227 | -3 | 0.21153 | 0.11357 | 0.86437 | 0.69972 | 10.83333 | 0.14 | 0 | Neutral |
| testneg_9288 | 0.22 | -2.66 | 0.385189 | -0.708217 | 1.336135 | -3 | 0.11533 | 0.12788 | 0.17935 | 0.36953 | 20.83333 | 0.187211538 | 0 | Neutral |
| testneg_9289 | 0 | 0 | -0.397788 | -3 | 2.939296 | -3 | 0.15911 | 0.11022 | 0.04519 | 0.12788 | 28.37793 | 0.03 | 0 | Neutral |
| testneg_9290 | 0.03 | -1.03 | 2.514378 | -3 | -0.272804 | -0.272804 | 0.3793 | 0.24105 | 0.17839 | 0.12009 | 17.78823 | 0.05 | 0 | Neutral |
| testneg_9291 | -0.14 | 1.01 | -0.146531 | -3 | -0.059162 | -3 | 0.23005 | 0.37216 | 0.8433 | 0.13128 | 11.23846 | 0.180355478 | 0 | Neutral |
| testneg_9292 | 0 | 0 | 0.024952 | -3 | -0.416655 | -0.416655 | 0.19423 | 0.14002 | 0.05453 | 0.12788 | 29.05903 | 0.025 | 0 | Neutral |
| testneg_9293 | 0 | 0 | 0.986599 | -3 | 0.423127 | -3 | 0.59066 | 0.10874 | 0.25643 | 0.12788 | 23.3819 | 0.01 | 0 | Neutral |
| testneg_9294 | -0.02 | 0.36 | -0.367202 | -3 | 4.552776 | -3 | 0.82846 | 0.13042 | 0.47975 | 0.18568 | 12.89568 | 0.264503064 | 0.4 | Neutral |
| testneg_9295 | 0.04 | -0.23 | 5.483519 | -3 | 3.546645 | -3 | 0.42583 | 0.10901 | 0.18502 | 0.21329 | 17.21847 | 0.604236111 | 88.2 | Disease_causing |
| testneg_9296 | -0.03 | -0.1 | 1.55494 | -3 | -0.280583 | -3 | 0.12819 | 0.1229 | 0.19898 | 0.09201 | 18.88889 | 0.01 | 0 | Neutral |
| testneg_9297 | 0.22 | -2.66 | 5.312389 | 5.312389 | 3.64719 | -3 | 0.07293 | 0.12989 | 0.82595 | 0.52905 | 6.77409 | 0.76 | 100 | Disease_causing |
| testneg_9298 | -0.05 | -0.29 | -1.389008 | -3 | 1.131706 | -3 | 0.19989 | 0.16777 | 0.43921 | 0.153 | 20.30303 | 0.115170424 | 0 | Neutral |
| testneg_9299 | 0 | 0 | 1.678033 | -3 | -0.034419 | -0.034419 | 0.33957 | 0.12788 | 0.62203 | 0.12788 | 10.28841 | 0.00117849 | 0 | Neutral |
| testneg_9300 | 0.2 | -1.26 | 3.785602 | -3 | 0.626243 | -3 | 0.3922 | 0.65186 | 0.34412 | 0.08672 | 20.83333 | 0.150166667 | 0 | Neutral |
| testneg_9301 | 0 | 0 | 2.508934 | -3 | -0.231245 | -3 | 0.10895 | 0.10442 | 0.1077 | 0.12788 | 4.44444 | 0.01 | 0 | Neutral |
| testneg_9302 | 0.05 | -0.81 | 1.039181 | -3 | -0.122196 | -0.122196 | 0.1551 | 0.09276 | 0.31961 | 0.17571 | 10.91235 | 0.05 | 0 | Neutral |
| testneg_9303 | -0.02 | -0.41 | 6.266856 | -3 | 0.92463 | -3 | 0.45997 | 0.15162 | 0.66693 | 0.0986 | 19.44444 | 0.291217391 | 1.6 | Neutral |
| testneg_9304 | -0.04 | 0.23 | 0.695292 | -3 | -0.301372 | -3 | 0.65674 | 0.10572 | 0.11019 | 0.08132 | 42.03605 | 0.021213235 | 0 | Neutral |
| testneg_9305 | 0 | 0 | 0.707491 | -3 | -0.142841 | -3 | 0.43762 | 0.12788 | 0.40081 | 0.15418 | 11.66667 | 0.022138578 | 0 | Neutral |
| testneg_9306 | 0.15 | 0.06 | 1.596806 | -3 | -0.33477 | -3 | 0.11796 | 0.12788 | 0.06431 | 0.12788 | 17.57509 | 0.015 | 0 | Neutral |
| testneg_9307 | 0 | 0 | 1.218246 | -3 | 0.224648 | -3 | 0.20448 | 0.12788 | 0.09067 | 0.10671 | 18.32431 | 0 | 0 | Neutral |
| testneg_9308 | 0 | 0 | 1.683026 | -3 | -0.22016 | -0.22016 | 0.84106 | 0.63405 | 0.17911 | 0.58893 | 21.71748 | 0.193333333 | 0 | Neutral |
| testneg_9309 | -0.08 | -0.03 | 5.10743 | -3 | -1.635844 | -3 | 0.13067 | 0.1005 | 0.15562 | 0.10384 | 13.51351 | 0.07 | 0 | Neutral |
| testneg_9310 | 0 | 0 | 0.29014 | -3 | 0.608491 | -3 | 0.16134 | 0.1847 | 0.15713 | 0.3507 | 10.1805 | 0.074660447 | 0 | Neutral |
| testneg_9311 | 0 | 0 | 0.461408 | -3 | 1.750774 | -3 | 0.16797 | 0.11809 | 0.19898 | 0.12788 | 9.52684 | 0 | 0 | Neutral |
| testneg_9312 | 0 | 0 | 2.307137 | -3 | 0.460694 | -3 | 0.91587 | 0.10818 | 0.24327 | 0.12192 | 12.4028 | 0.000672696 | 0 | Neutral |
| testneg_9313 | 0.08 | -1.2 | 0.387956 | -3 | 3.256775 | -3 | 0.19898 | 0.12788 | 0.16309 | 0.12788 | 33.61654 | 0.077145468 | 0 | Neutral |
| testneg_9314 | 0.22 | -2.66 | 0.385189 | -3 | 0.194184 | -3 | 0.11533 | 0.12788 | 0.18744 | 0.09392 | 18.22258 | 0.06 | 0 | Neutral |
| testneg_9315 | -0.05 | -0.29 | 2.254133 | -3 | -1.841925 | -3 | 0.24971 | 0.12971 | 0.44479 | 0.20004 | 14.16667 | 0.0749 | 0 | Neutral |
| testneg_9316 | 0.21 | 0.15 | 1.48266 | -3 | 3.879651 | -3 | 0.07427 | 0.09451 | 0.18705 | 0.43522 | 12.42141 | 0.30755 | 2.2 | Neutral |
| testneg_9317 | -0.01 | 0.57 | -2.581564 | -3 | 3.566467 | -3 | 0.19898 | 0.12788 | 0.17548 | 0.11384 | 13.51351 | 0.082974693 | 0 | Neutral |
| testneg_9318 | -0.16 | -0.08 | 2.243022 | -3 | 2.493678 | -3 | 0.07064 | 0.11699 | 0.50057 | 0.10408 | 31.94915 | 0.110115316 | 0 | Neutral |
| testneg_9319 | 0.02 | 0.41 | 3.953483 | -3 | 0.758366 | -3 | 0.97273 | 0.27711 | 0.16053 | 0.15049 | 19.79731 | 0.226165574 | 0 | Neutral |
| testneg_9320 | -0.07 | -0.38 | 4.428125 | -3 | 3.618173 | -3 | 0.76693 | 0.19444 | 0.78515 | 0.35093 | 10.41667 | 0.612351577 | 88.2 | Disease_causing |
| testneg_9321 | 0.03 | -1.03 | 2.800909 | -3 | 1.26259 | 1.26259 | 0.61491 | 0.11954 | 0.44722 | 0.11641 | 6.24727 | 0.04 | 0 | Neutral |
| testneg_9322 | 0 | 0 | 4.941822 | -3 | 6.955427 | 6.955427 | 0.08789 | 0.6361 | 0.23401 | 0.12374 | 11.875 | 0.79 | 99.8 | Disease_causing |
| testneg_9323 | 0 | -0.01 | 1.609618 | -3 | 2.901134 | -3 | 0.67038 | 0.1794 | 0.82014 | 0.14054 | 10.83333 | 0.130575855 | 0 | Neutral |
| testneg_9324 | -0.09 | 0.3 | -2.556301 | -3 | -0.202498 | -3 | 0.12379 | 0.12788 | 0.30789 | 0.15135 | 12.20279 | 0.07 | 0 | Neutral |
| testneg_9325 | 0 | 0 | 0.122969 | -3 | -1.117253 | -1.117253 | 0.1215 | 0.12788 | 0.84682 | 0.35105 | 6.0461 | 0.05 | 0 | Neutral |
| testneg_9326 | 0 | 0 | 1.413285 | -3 | 2.041175 | -3 | 0.90811 | 0.17448 | 0.04988 | 0.25503 | 10.41667 | 0.07 | 0 | Neutral |
| testneg_9327 | 0 | 0 | -0.843404 | -3 | 5.071172 | -3 | 0.11587 | 0.12001 | 0.25195 | 0.09584 | 7.04228 | 0.07 | 0 | Neutral |
| testneg_9328 | 0.22 | -2.66 | 7.490177 | -3 | 4.69301 | -3 | 0.34205 | 0.11195 | 0.11479 | 0.12788 | 21.8254 | 0.511929619 | 58.6 | Disease_causing |
| testneg_9329 | 0 | 0 | 1.541318 | -3 | 2.418801 | -3 | 0.11564 | 0.09588 | 0.05318 | 0.12788 | 29.89616 | 0.02 | 0 | Neutral |
| testneg_9330 | 0 | 0 | 0.547246 | -3 | 1.078945 | -3 | 0.81169 | 0.11096 | 0.12099 | 0.11502 | 12.67985 | 0.002 | 0 | Neutral |
| testneg_9331 | 0 | -0.01 | 3.021892 | -3 | 6.453651 | -3 | 0.1163 | 0.20863 | 0.07889 | 0.08835 | 10.83333 | 0.420518519 | 24 | Neutral |
| testneg_9332 | 0.06 | 0.18 | 0.256159 | -3 | 1.147993 | -3 | 0.08954 | 0.63401 | 0.46386 | 0.16229 | 18.33333 | 0.22499183 | 0.2 | Neutral |
| testneg_9333 | -0.02 | -0.45 | -0.240957 | -3 | 0.551288 | -3 | 0.06526 | 0.12788 | 0.83421 | 0.18817 | 20.37554 | 0.08 | 0 | Neutral |
| testneg_9334 | 0.01666667 | -0.1875 | 0.933869 | -3 | 0.432868 | -3 | 0.9862 | 0.4528 | 0.37292 | 0.11979 | 11.89874 | 0.037575188 | 0 | Neutral |
| testneg_9335 | -0.03 | -0.12 | -1.033749 | -3 | 4.018957 | 2.663229 | 0.09964 | 0.10384 | 0.25275 | 0.12788 | 10.22898 | 0.16025 | 0 | Neutral |
| testneg_9336 | 0 | -0.01 | -0.337044 | -3 | 2.726693 | -3 | 0.02396 | 0.12703 | 0.28227 | 0.10211 | 10.41667 | 0.040559468 | 0 | Neutral |
| testneg_9337 | 0 | 0 | -0.39724 | -3 | 2.449599 | -3 | 0.09749 | 0.82924 | 0.18578 | 0.27854 | 21.38889 | 0.327907066 | 2.8 | Neutral |
| testneg_9338 | 0.15 | -0.28 | -1.251158 | -3 | -0.269735 | -0.269735 | 0.20127 | 0.13478 | 0.12461 | 0.1241 | 31.55193 | 0.03 | 0 | Neutral |
| testneg_9339 | -0.15 | 0.28 | -0.668638 | -0.668638 | 0.550914 | -3 | 0.37131 | 0.10994 | 0.17322 | 0.10311 | 28.98534 | 0.05 | 0 | Neutral |
| testneg_9340 | 0.04 | -0.23 | 4.118776 | -3 | 4.536943 | -3 | 0.35082 | 0.12788 | 0.08982 | 0.15279 | 18.33333 | 0.505685067 | 56.2 | Disease_causing |
| testneg_9341 | 0.05 | 0.29 | -1.171244 | -1.171244 | 0.157379 | -3 | 0.19898 | 0.12788 | 0.45997 | 0.15162 | 27.95796 | 0.057816667 | 0 | Neutral |
| testneg_9342 | 0.06 | -0.57 | 2.077607 | -3 | 1.782871 | -3 | 0.53203 | 0.12788 | 0.61834 | 0.10784 | 35.21069 | 0.02 | 0 | Neutral |
| testneg_9343 | 0.00545455 | -0.0303636 | 5.729308 | -3 | 0.616062 | -3 | 0.06793 | 0.12788 | 0.54711 | 0.11809 | 14.65165 | 0.1375 | 0.2 | Neutral |
| testneg_9344 | 0.18 | -1.05 | 6.918655 | -3 | 2.433708 | -3 | 0.15155 | 0.09923 | 0.21399 | 0.10708 | 22.61823 | 0.2575 | 2.2 | Neutral |
| testneg_9345 | 0 | 0 | 2.044018 | -3 | 0.459406 | -3 | 0.63431 | 0.09837 | 0.03981 | 0.09432 | 13.51351 | 0.010194002 | 0 | Neutral |
| testneg_9346 | 0 | 0 | 1.1308 | -3 | -1.062235 | -3 | 0.19898 | 0.11185 | 0.29172 | 0.08143 | 46.53346 | 0 | 0 | Neutral |
| testneg_9347 | 0.04 | -1.36 | -0.054368 | -3 | 5.803451 | -3 | 0.09488 | 0.10426 | 0.16692 | 0.09243 | 27.51545 | 0.110455466 | 0 | Neutral |
| testneg_9348 | 0 | 0 | -0.09322 | -0.09322 | 3.355715 | -3 | 0.14341 | 0.10134 | 0.35026 | 0.09817 | 18.88889 | 0.03 | 0 | Neutral |
| testneg_9349 | 0.04 | -0.23 | 5.204302 | -3 | 1.799258 | 1.799258 | 0.1947 | 0.11413 | 0.204 | 0.15424 | 10.83333 | 0.328648183 | 6.8 | Neutral |
| testneg_9350 | 0.06 | -1.29 | 6.321709 | -3 | 5.587743 | -3 | 0.96879 | 0.38765 | 0.21983 | 0.10146 | 16.42913 | 0.621458333 | 92.6 | Disease_causing |
| testneg_9351 | -0.04 | 0.23 | -1.561199 | -3 | 6.298478 | -3 | 0.07617 | 0.07182 | 0.19898 | 0.12788 | 20.17482 | 0.165727273 | 0 | Neutral |
| testneg_9352 | 0 | 0 | 1.468723 | -3 | 3.35171 | -3 | 0.45152 | 0.34457 | 0.20112 | 0.17491 | 11.25 | 0.28752324 | 2.4 | Neutral |
| testneg_9353 | 0.07 | 0.38 | 0.667855 | -3 | 0.55724 | 0.55724 | 0.53592 | 0.11293 | 0.12937 | 0.23004 | 21.2963 | 0.112542494 | 0 | Neutral |
| testneg_9354 | 0.03 | 1.22 | 0.949523 | 0.949523 | 1.069555 | -3 | 0.10056 | 0.12608 | 0.19898 | 0.12788 | 20.77357 | 0.07 | 0 | Neutral |
| testneg_9355 | 0.21 | 0.15 | 6.016475 | -3 | 5.524661 | -3 | 0.16926 | 0.11871 | 0.19898 | 0.12788 | 35.83242 | 0.38 | 20.8 | Neutral |
| testneg_9356 | 0 | 0 | 0.28875 | -3 | -0.133213 | -3 | 0.40179 | 0.12112 | 0.39163 | 0.2681 | 13.04529 | 0.02086553 | 0 | Neutral |
| testneg_9357 | 0.04 | 2.01 | -0.573377 | -1.816133 | -0.474499 | -3 | 0.42366 | 0.14372 | 0.1 | 0.22633 | 18.33333 | 0.180496661 | 0 | Neutral |
| testneg_9358 | -0.15 | -0.04 | 2.269407 | -3 | 0.916588 | -3 | 0.98442 | 0.44966 | 0.06603 | 0.08071 | 13.02635 | 0.070231383 | 0 | Neutral |
| testneg_9359 | 0 | 0 | 1.813816 | -3 | 1.806927 | -3 | 0.13794 | 0.40936 | 0.17973 | 0.12278 | 14.16667 | 0.030185759 | 0 | Neutral |
| testneg_9360 | -0.02 | -0.45 | 1.073382 | -3 | 2.800312 | -3 | 0.31542 | 0.26739 | 0.09259 | 0.11165 | 35.05388 | 0.0645 | 0 | Neutral |
| testneg_9361 | 0 | 0 | 0.501525 | -3 | -0.171687 | -3 | 0.20457 | 0.12378 | 0.60079 | 0.11198 | 23.04078 | 0 | 0 | Neutral |
| testneg_9362 | 0 | -0.01 | 0.611606 | -3 | 3.769006 | -3 | 0.9149 | 0.27793 | 0.44479 | 0.20004 | 10.83333 | 0.282471902 | 1.8 | Neutral |
| testneg_9363 | 0 | -0.01 | 1.961035 | -3 | 0.654716 | -3 | 0.55105 | 0.43208 | 0.76942 | 0.13267 | 20.40495 | 0.100849191 | 0 | Neutral |
| testneg_9364 | -0.06 | 0.57 | 0.883365 | -3 | 3.333939 | -3 | 0.10004 | 0.12788 | 0.06749 | 0.1776 | 11.54167 | 0.229945276 | 0.2 | Neutral |
| testneg_9365 | 0 | -0.01 | 0.465839 | -3 | -0.187363 | -3 | 0.07875 | 0.11082 | 0.90649 | 0.3543 | 9.25676 | 0.030666229 | 0 | Neutral |
| testneg_9366 | -0.05 | -0.29 | -1.161569 | -3 | 0.085787 | -3 | 0.16974 | 0.80184 | 0.40996 | 0.57507 | 10.83333 | 0.212027582 | 0.2 | Neutral |
| testneg_9367 | 0 | 0 | 0.084018 | -3 | -0.559847 | -3 | 0.08365 | 0.34122 | 0.08891 | 0.12788 | 20.49588 | 0.050080645 | 0 | Neutral |
| testneg_9368 | 0 | 0 | -0.009172 | -3 | 3.178384 | -3 | 0.20055 | 0.12788 | 0.07923 | 0.08035 | 39.62997 | 0.051485186 | 0 | Neutral |
| testneg_9369 | -0.23 | 1.04 | 1.436839 | -3 | 1.479184 | -3 | 0.19898 | 0.12788 | 0.26909 | 0.10159 | 41.56132 | 0.11 | 0 | Neutral |
| testneg_9370 | -0.03 | -1.22 | 6.195353 | -3 | 0.641246 | -3 | 0.28532 | 0.10663 | 0.21118 | 0.10074 | 13.51351 | 0.19 | 0.2 | Neutral |
| testneg_9371 | 0 | 0 | 1.283931 | -3 | 0.069349 | -3 | 0.03499 | 0.12971 | 0.19898 | 0.12788 | 29.86111 | 0.03 | 0 | Neutral |
| testneg_9372 | 0 | 0 | -0.299858 | -3 | 0.66275 | -3 | 0.29284 | 0.11262 | 0.30429 | 0.29938 | 9.67871 | 0.020834058 | 0 | Neutral |
| testneg_9373 | -0.14 | 1.01 | 4.151556 | -3 | 0.119587 | -3 | 0.13563 | 0.12378 | 0.13177 | 0.12788 | 24.41767 | 0.090169115 | 0 | Neutral |
| testneg_9374 | -0.05 | -0.29 | 1.652664 | -3 | 5.091084 | -3 | 0.62708 | 0.24932 | 0.03502 | 0.08119 | 13.2365 | 0.138375228 | 0 | Neutral |
| testneg_9375 | -0.02 | 0.63 | 4.759697 | -3 | 0.97098 | -3 | 0.19898 | 0.10335 | 0.75415 | 0.29199 | 20.27778 | 0.230690476 | 0 | Neutral |
| testneg_9376 | 0 | 0 | 0.425569 | 0.425569 | 3.701009 | -3 | 0.12879 | 0.12788 | 0.12642 | 0.11218 | 12.81915 | 0.06 | 0 | Neutral |
| testneg_9377 | -0.03 | -1.22 | 1.187592 | -3 | -0.925939 | -3 | 0.07864 | 0.12788 | 0.39192 | 0.09135 | 10.41667 | 0.03 | 0 | Neutral |
| testneg_9378 | 0 | 0 | 0.052371 | 0.052371 | 0.475023 | -3 | 0.82107 | 0.27926 | 0.1926 | 0.12788 | 9.93641 | 0.01 | 0 | Neutral |
| testneg_9379 | 0 | 0 | 1.436812 | -3 | 4.637381 | -0.78887 | 0.29302 | 0.17394 | 0.47188 | 0.10037 | 13.53166 | 0.081717881 | 0 | Neutral |
| testneg_9380 | 0.18 | 0.16 | -0.13961 | -3 | 2.711498 | -3 | 0.65734 | 0.1013 | 0.15406 | 0.1454 | 32.72367 | 0.21 | 0 | Neutral |
| testneg_9381 | 0.16 | -0.37 | -2.713571 | -2.713571 | 2.947718 | -3 | 0.07506 | 0.09112 | 0.5068 | 0.12276 | 27.65823 | 0.06 | 0 | Neutral |
| testneg_9382 | 0 | 0 | 1.716346 | -3 | 3.088745 | -3 | 0.09645 | 0.13239 | 0.17412 | 0.10595 | 9.82562 | 0.045119048 | 0 | Neutral |
| testneg_9383 | 0.11 | -1.95 | 3.52493 | -3 | 1.189051 | -3 | 0.04916 | 0.14944 | 0.19227 | 0.1096 | 12.30045 | 0.14 | 0 | Neutral |
| testneg_9384 | 0.02 | -0.68 | 6.639247 | -3 | 4.341836 | -3 | 0.27407 | 0.1136 | 0.10187 | 0.08499 | 18.33333 | 0.4 | 21.8 | Neutral |
| testneg_9385 | 0 | 0 | 0.32262 | -3 | 5.60807 | -3 | 0.32089 | 0.1035 | 0.25171 | 0.13527 | 20.5518 | 0.239340909 | 0.2 | Neutral |
| testneg_9386 | 0.16 | -1.37 | -0.103493 | -3 | 3.057504 | -3 | 0.12835 | 0.15143 | 0.09893 | 0.1266 | 22.91479 | 0.136458333 | 0 | Neutral |
| testneg_9387 | 0 | 0.32 | 1.687645 | -3 | 2.908555 | -3 | 0.04519 | 0.12788 | 0.14175 | 0.12788 | 18.88889 | 0.06 | 0 | Neutral |
| testneg_9388 | -0.08 | -0.03 | 5.721411 | -3 | 2.149049 | -3 | 0.19494 | 0.14451 | 0.43502 | 0.14712 | 21.38889 | 0.466468797 | 41.4 | Neutral |
| testneg_9389 | 0 | -0.32 | 2.305189 | -3 | 1.030888 | -3 | 0.505 | 0.12788 | 0.2439 | 0.08848 | 22.39953 | 0.010122704 | 0 | Neutral |
| testneg_9390 | 0 | 0 | -0.470651 | -3 | 3.105034 | -3 | 0.28063 | 0.40063 | 0.15267 | 0.12788 | 28.04989 | 0.095702381 | 0 | Neutral |
| testneg_9391 | -0.02 | 0.63 | 2.556329 | -3 | -0.086743 | -3 | 0.33256 | 0.14909 | 0.86851 | 0.17715 | 5.997 | 0.091033489 | 0 | Neutral |
| testneg_9392 | 0 | 0 | 1.504538 | -3 | 1.632466 | -3 | 0.19898 | 0.12788 | 0.19214 | 0.12788 | 20.05873 | 0 | 0 | Neutral |
| testneg_9393 | -0.09 | 0.25 | 3.048378 | -3 | 1.225058 | -3 | 0.32861 | 0.09592 | 0.30209 | 0.14229 | 27.0922 | 0.11528169 | 0 | Neutral |

| | | | | | | | | | | | | | | |
|---|---|---|---|---|---|---|---|---|---|---|---|---|---|---|
| testneg_9394 | 0.22 | -2.66 | 6.956883 | -3 | 4.2288 | -3 | 0.12131 | 0.12533 | 0.18706 | 0.10259 | 44.76775 | 0.38 | 15 | Neutral |
| testneg_9395 | 0.04 | -0.23 | 1.519913 | -3 | 3.784732 | -3 | 0.03008 | 0.12788 | 0.98578 | 0.91075 | 8.97898 | 0.323798363 | 3 | Neutral |
| testneg_9396 | 0 | 0 | 2.259254 | -3 | 1.389503 | -3 | 0.56285 | 0.18619 | 0.13997 | 0.19598 | 11.98582 | 0.07462136 | 0 | Neutral |
| testneg_9397 | 0.22 | -2.66 | 6.248531 | -3 | -0.354855 | -3 | 0.12839 | 0.09503 | 0.14524 | 0.10162 | 18.88889 | 0.14 | 0 | Neutral |
| testneg_9398 | -0.1 | -1.9 | 3.7566 | -3 | 1.52844 | -3 | 0.54489 | 0.09921 | 0.37065 | 0.12788 | 6.01504 | 0.090122222 | 0 | Neutral |
| testneg_9399 | -0.07 | -0.38 | 1.275959 | -3 | 5.787827 | -3 | 0.2058 | 0.28174 | 0.59691 | 0.11049 | 17.92796 | 0.221672384 | 0.2 | Neutral |
| testneg_9400 | -0.03 | -0.1 | 1.538056 | 0.552009 | 0.12577 | -3 | 0.53172 | 0.12788 | 0.07872 | 0.12788 | 13.46801 | 0.03 | 0 | Neutral |
| testneg_9401 | 0 | 0 | 1.31141 | -3 | 3.664538 | -3 | 0.9449 | 0.9378 | 0.10395 | 0.08737 | 18.33333 | 0.140755054 | 0 | Neutral |
| testneg_9402 | 0.21 | 0.15 | 5.657785 | 5.657785 | 0.452231 | -3 | 0.16497 | 0.74918 | 0.46691 | 0.07886 | 18.88889 | 0.55 | 78.6 | Disease_causing |
| testneg_9403 | -0.07 | -0.38 | -0.872044 | -3 | 2.479467 | -3 | 0.04792 | 0.12788 | 0.1601 | 0.12546 | 22.89111 | 0.068149254 | 0 | Neutral |
| testneg_9404 | 0 | 0 | -0.02198 | -0.02198 | -0.061342 | -3 | 0.15509 | 0.1326 | 0.07626 | 0.12788 | 20.26596 | 0.021039673 | 0 | Neutral |
| testneg_9405 | 0.22 | -2.66 | 2.520208 | -3 | 5.486225 | -3 | 0.08684 | 0.08917 | 0.10963 | 0.12788 | 13.51351 | 0.171195286 | 0 | Neutral |
| testneg_9406 | 0.19 | 0.51 | 4.472767 | -3 | 1.420779 | -3 | 0.12853 | 0.08565 | 0.14386 | 0.12788 | 22.12292 | 0.063095238 | 0 | Neutral |
| testneg_9407 | -0.02 | -0.41 | -0.390123 | -3 | -0.68836 | -3 | 0.09657 | 0.58605 | 0.19898 | 0.28128 | 10.41667 | 0.211177812 | 0 | Neutral |
| testneg_9408 | 0.05 | -0.81 | 1.61682 | -3 | 11.796446 | -3 | 0.21596 | 0.08969 | 0.13626 | 0.12788 | 54.51463 | 0.32 | 3.8 | Neutral |
| testneg_9409 | -0.15 | -0.04 | 0.012434 | -3 | -0.694905 | -3 | 0.08629 | 0.12788 | 0.4132 | 0.12788 | 34.73889 | 0.06 | 0 | Neutral |
| testneg_9410 | -0.11 | 0.19 | 4.451137 | -3 | 2.993925 | -3 | 0.20675 | 0.13709 | 0.12848 | 0.65153 | 8.612 | 0.508854167 | 55.4 | Disease_causing |
| testneg_9411 | -0.14 | 0.37 | 0.9707 | -3 | 6.366587 | -3 | 0.82131 | 0.12788 | 0.64072 | 0.12788 | 39.79929 | 0.22 | 0.2 | Neutral |
| testneg_9412 | 0 | 0 | 1.436812 | -3 | -0.586534 | -0.586534 | 0.29302 | 0.17394 | 0.09439 | 0.1884 | 24.24555 | 0.122644231 | 0 | Neutral |
| testneg_9413 | 0 | 0 | 1.103204 | -3 | 5.00875 | -3 | 0.57199 | 0.11262 | 0.68054 | 0.09208 | 20.5134 | 0.061437766 | 0 | Neutral |
| testneg_9414 | -0.04 | 0.23 | 0.98325 | -3 | -0.720759 | -0.720759 | 0.16054 | 0.11683 | 0.17758 | 0.15983 | 8.49181 | 0.07 | 0 | Neutral |
| testneg_9415 | 0.02 | -0.68 | -0.416108 | -3 | -0.780137 | -3 | 0.1621 | 0.1641 | 0.06958 | 0.16086 | 4.72222 | 0.113702957 | 0 | Neutral |
| testneg_9416 | 0 | 0 | -0.399895 | -3 | 1.267171 | -3 | 0.5632 | 0.28442 | 0.27915 | 0.27706 | 11.90227 | 0.070215829 | 0 | Neutral |
| testneg_9417 | -0.21 | -0.15 | 1.169097 | -3 | -0.151164 | -0.151164 | 0.10011 | 0.08014 | 0.65708 | 0.0936 | 8.59165 | 0.08 | 0 | Neutral |
| testneg_9418 | 0 | 0 | 0.27864 | -3 | 0.252532 | -3 | 0.17737 | 0.54971 | 0.06215 | 0.08575 | 19.64428 | 0.050427196 | 0 | Neutral |
| testneg_9419 | 0 | 0 | 1.15789 | 1.15789 | 1.018673 | -3 | 0.68439 | 0.09371 | 0.18147 | 0.12788 | 16.47059 | 0.03 | 0 | Neutral |
| testneg_9420 | 0.008 | 0.24890909 | 4.814261 | -3 | 1.827547 | -3 | 0.08775 | 0.12788 | 0.35584 | 0.10874 | 13.51351 | 0.06 | 0 | Neutral |
| testneg_9421 | 0 | 0 | -1.056572 | -3 | -0.574737 | -3 | 0.1109 | 0.17295 | 0.56678 | 0.09955 | 38.77183 | 0.04 | 0 | Neutral |
| testneg_9422 | 0.05 | -0.81 | 2.670334 | -3 | -0.39694 | -3 | 0.31906 | 0.1705 | 0.21807 | 0.12788 | 11.25 | 0.04 | 0 | Neutral |
| testneg_9423 | 0 | 0 | 1.191933 | -3 | 1.242354 | -3 | 0.54876 | 0.22335 | 0.66381 | 0.13039 | 10.9845 | 0.039949495 | 0 | Neutral |
| testneg_9424 | -0.1 | -1.9 | -2.176445 | -3 | 2.370221 | -3 | 0.226 | 0.28784 | 0.19898 | 0.13249 | 24.7633 | 0.242208333 | 0.2 | Neutral |
| testneg_9425 | 0.02 | -0.68 | 4.942272 | -3 | 0.224881 | -3 | 0.36566 | 0.15312 | 0.1653 | 0.0977 | 8.97898 | 0.1 | 0 | Neutral |
| testneg_9426 | 0.02 | 0.13 | 7.041902 | -3 | -0.650443 | -3 | 0.34467 | 0.13076 | 0.53447 | 0.10345 | 31.65147 | 0.26 | 1.4 | Neutral |
| testneg_9427 | 0.07 | 0.01 | 1.365447 | -3 | -0.730881 | -3 | 0.07701 | 0.19824 | 0.26476 | 0.08217 | 28.59338 | 0.04 | 0 | Neutral |
| testneg_9428 | 0 | 0 | 0.619175 | 0.619175 | 0.383334 | -3 | 0.68138 | 0.20944 | 0.04935 | 0.06599 | 10 | 0.030356193 | 0 | Neutral |
| testneg_9429 | 0.03 | 0.1 | -0.187467 | -0.187467 | 0.075565 | -3 | 0.12314 | 0.11709 | 0.03788 | 0.10996 | 11.875 | 0.01 | 0 | Neutral |
| testneg_9430 | 0 | 0 | 1.302665 | 1.302665 | -0.025439 | -3 | 0.71803 | 0.95763 | 0.21355 | 0.12788 | 19.45242 | 0.1 | 0 | Neutral |
| testneg_9431 | 0.04 | 2.01 | -0.172559 | -3 | 3.878129 | -3 | 0.22155 | 0.1062 | 0.06744 | 0.12788 | 18.88889 | 0.16 | 0 | Neutral |
| testneg_9432 | 0 | 0 | -0.090459 | -0.090459 | 5.036276 | -3 | 0.15581 | 0.19351 | 0.22944 | 0.09039 | 20.83333 | 0.147236953 | 0 | Neutral |
| testneg_9433 | 0 | 0 | 0.628299 | -3 | 9.146604 | -3 | 0.40612 | 0.45135 | 0.44923 | 0.12788 | 13.12342 | 0.418375 | 26 | Neutral |
| testneg_9434 | 0 | -0.01 | 0.181227 | -3 | 0.976401 | -3 | 0.15132 | 0.11724 | 0.10498 | 0.08421 | 18.33333 | 0.00051458 | 0 | Neutral |
| testneg_9435 | 0.2 | -1.26 | 4.70798 | -3 | 5.969277 | -3 | 0.17527 | 0.13941 | 0.31067 | 0.35414 | 11.5 | 0.811448523 | 99.8 | Disease_causing |
| testneg_9436 | 0 | 0 | 0.81674 | -3 | 5.481335 | -3 | 0.84238 | 0.12559 | 0.7482 | 0.18812 | 18.88889 | 0.217989068 | 0 | Neutral |
| testneg_9437 | -0.08 | -0.03 | 2.090774 | -3 | -0.416655 | -0.416655 | 0.35421 | 0.10694 | 0.05453 | 0.12788 | 28.67527 | 0.02 | 0 | Neutral |
| testneg_9438 | 0 | 0 | 1.618033 | -3 | 0.982788 | -3 | 0.35092 | 0.10596 | 0.1481 | 0.1117 | 18.6768 | 0 | 0 | Neutral |
| testneg_9439 | -0.07 | -0.38 | 5.176715 | -3 | 1.214875 | -3 | 0.206 | 0.14631 | 0.80889 | 0.40016 | 10.7989 | 0.355 | 9.6 | Neutral |
| testneg_9440 | -0.02 | 0.28 | 2.768215 | -3 | 1.354044 | -3 | 0.26928 | 0.12774 | 0.64608 | 0.35524 | 16.42913 | 0.133011742 | 0 | Neutral |
| testneg_9441 | 0 | 0 | 0.338971 | -3 | 0.804817 | -3 | 0.0752 | 0.74467 | 0.12831 | 0.10983 | 13.08587 | 0.06 | 0 | Neutral |
| testneg_9442 | 0.15 | -0.28 | 0.666695 | -3 | 0.288264 | -3 | 0.13278 | 0.20513 | 0.68004 | 0.12345 | 7.00579 | 0.051055556 | 0 | Neutral |
| testneg_9443 | 0 | -0.01 | 0.169949 | -3 | 1.784024 | -3 | 0.38445 | 0.58278 | 0.15152 | 0.12645 | 10 | 0.03 | 0 | Neutral |
| testneg_9444 | 0 | 0 | 0.598487 | -3 | -0.772413 | -3 | 0.25011 | 0.12092 | 0.10231 | 0.1 | 5.69444 | 0.01 | 0 | Neutral |
| testneg_9445 | -0.03 | -0.1 | -1.512759 | -3 | 1.846205 | -3 | 0.32335 | 0.10604 | 0.23078 | 0.10588 | 10.36687 | 0.01 | 0 | Neutral |
| testneg_9446 | 0 | -0.01 | 0.597852 | -3 | 4.209007 | -3 | 0.06027 | 0.11574 | 0.10724 | 0.49624 | 4.72222 | 0.27817051 | 2 | Neutral |
| testneg_9447 | -0.01 | 0.57 | 2.185335 | -3 | 0.467995 | -3 | 0.19898 | 0.12788 | 0.11237 | 0.08916 | 20.63101 | 0.02 | 0 | Neutral |
| testneg_9448 | 0.01 | -0.57 | 1.867073 | -3 | 1.75958 | -3 | 0.07999 | 0.23927 | 0.65897 | 0.67602 | 10 | 0.217734311 | 0 | Neutral |
| testneg_9449 | 0.08 | 0.03 | 3.533518 | -3 | 3.79031 | -3 | 0.32406 | 0.28376 | 0.10042 | 0.09296 | 10.16725 | 0.251558767 | 0.2 | Neutral |
| testneg_9450 | -0.06 | -0.18 | 0.491194 | 0.491194 | -0.513734 | -3 | 0.72124 | 0.09906 | 0.256 | 0.20036 | 10.22898 | 0.071054459 | 0 | Neutral |
| testneg_9451 | -0.04 | 0.23 | 0.421557 | 0.421557 | 0.213037 | -3 | 0.82507 | 0.19615 | 0.25077 | 0.06261 | 14.5242 | 0.03 | 0 | Neutral |
| testneg_9452 | -0.01 | -0.8 | 5.577235 | -3 | 9.935182 | -3 | 0.31062 | 0.1222 | 0.08393 | 0.1123 | 29.07519 | 0.586071429 | 84 | Disease_causing |
| testneg_9453 | 0.05 | -0.81 | 3.455713 | -3 | 0.523312 | -3 | 0.12175 | 0.21177 | 0.41987 | 0.67373 | 10 | 0.327140015 | 5 | Neutral |
| testneg_9454 | -0.14 | 1.01 | 5.258974 | -3 | -0.923371 | -3 | 0.19898 | 0.28128 | 0.07611 | 0.11551 | 10.41667 | 0.249444444 | 1 | Neutral |
| testneg_9455 | -0.07 | -0.01 | -1.321361 | -3 | -0.960298 | -3 | 0.21843 | 0.85214 | 0.27924 | 0.12788 | 19.66924 | 0.09 | 0 | Neutral |
| testneg_9456 | -0.19 | 0.25 | 1.864528 | 1.864528 | 3.730673 | -3 | 0.21153 | 0.11357 | 0.09426 | 0.1658 | 19.44444 | 0.44907197 | 33 | Neutral |
| testneg_9457 | 0.02 | 0.45 | 2.338504 | -3 | 0.088732 | -3 | 0.53496 | 0.13586 | 0.24524 | 0.10803 | 18.12633 | 0.02 | 0 | Neutral |
| testneg_9458 | 0.03 | -0.85 | -1.765145 | -3 | 4.941921 | -3 | 0.19898 | 0.12788 | 0.1725 | 0.09643 | 23.0391 | 0.090772909 | 0 | Neutral |
| testneg_9459 | 0 | -0.01 | 3.849198 | -3 | 0.019411 | -3 | 0.08337 | 0.18096 | 0.10863 | 0.098 | 26.359 | 0.08 | 0 | Neutral |
| testneg_9460 | 0 | -0.01 | 0.729709 | -3 | 1.052456 | -3 | 0.40557 | 0.11865 | 0.19898 | 0.1347 | 21.00633 | 0.057411616 | 0 | Neutral |
| testneg_9461 | 0 | 0 | -0.334686 | -3 | 3.122928 | 2.114299 | 0.19898 | 0.10269 | 0.37637 | 0.12788 | 25.73753 | 0.08 | 0 | Neutral |
| testneg_9462 | -0.08 | -0.03 | 2.314112 | -3 | 7.140458 | -3 | 0.10774 | 0.13117 | 0.24823 | 0.12788 | 10.28409 | 0.4 | 20 | Neutral |
| testneg_9463 | -0.14 | 1.01 | -0.889855 | -0.889855 | 7.507173 | -3 | 0.042 | 0.07218 | 0.13967 | 0.15696 | 23.21544 | 0.479409439 | 45.6 | Neutral |
| testneg_9464 | 0.14 | -1.01 | 1.865393 | -3 | -0.093394 | -0.093394 | 0.19898 | 0.10482 | 0.34587 | 0.29822 | 71.46337 | 0.21 | 0 | Neutral |
| testneg_9465 | -0.15 | -0.04 | 5.146748 | -3 | -0.484804 | -3 | 0.16778 | 0.82322 | 0.14668 | 0.17962 | 16.13039 | 0.527551471 | 63.6 | Disease_causing |
| testneg_9466 | 0.16 | -0.37 | 3.233391 | -3 | -0.844127 | -3 | 0.2674 | 0.28725 | 0.22817 | 0.10634 | 12.29167 | 0.06 | 0 | Neutral |
| testneg_9467 | -0.05 | -0.29 | 1.981611 | -3 | 3.717487 | -3 | 0.24977 | 0.11262 | 0.41643 | 0.09692 | 12.58479 | 0.06 | 0 | Neutral |
| testneg_9468 | -0.04 | 0.23 | 0.760013 | -3 | 1.308546 | -3 | 0.19898 | 0.14896 | 0.12545 | 0.08963 | 17.71829 | 0.03 | 0 | Neutral |
| testneg_9469 | 0.03 | 0.12 | 6.140208 | -3 | 6.380091 | -3 | 0.40179 | 0.12112 | 0.14704 | 0.12788 | 18.34652 | 0.59 | 83.4 | Disease_causing |
| testneg_9470 | 0 | 0 | 0.268006 | -3 | -0.975203 | -3 | 0.68568 | 0.65219 | 0.37292 | 0.11979 | 11.25 | 0.028831169 | 0 | Neutral |
| testneg_9471 | 0.14 | -0.69 | 3.700286 | -3 | 2.149681 | -3 | 0.11687 | 0.09069 | 0.19052 | 0.12788 | 18.12875 | 0.06 | 0 | Neutral |
| testneg_9472 | 0 | 0 | 0.297064 | -3 | 5.730226 | -3 | 0.2029 | 0.12788 | 0.13704 | 0.10054 | 18.6768 | 0.061952381 | 0 | Neutral |
| testneg_9473 | 0.02 | -0.63 | 2.525878 | -3 | -0.01248 | -3 | 0.19898 | 0.08275 | 0.61231 | 0.13362 | 13.33989 | 0.050303571 | 0 | Neutral |
| testneg_9474 | 0.07 | 0.38 | -0.291949 | -3 | -0.131159 | -3 | 0.08999 | 0.0989 | 0.2218 | 0.11533 | 18.77103 | 0.000711279 | 0 | Neutral |
| testneg_9475 | -0.01 | -0.8 | 6.193509 | -3 | 1.023884 | -3 | 0.31817 | 0.17348 | 0.19898 | 0.12788 | 16.19072 | 0.299704185 | 3 | Neutral |
| testneg_9476 | -0.07 | -0.7 | 0.044039 | -3 | 2.300638 | -3 | 0.36486 | 0.14268 | 0.3405 | 0.1816 | 18.6768 | 0.12163474 | 0 | Neutral |
| testneg_9477 | 0 | 0 | 0.815861 | -3 | 0.900933 | -3 | 0.11564 | 0.09588 | 0.07611 | 0.11551 | 20 | 0.01 | 0 | Neutral |
| testneg_9478 | 0 | 0 | 1.916985 | -3 | -0.266524 | -3 | 0.84031 | 0.16749 | 0.23962 | 0.10029 | 8.49181 | 0.010184746 | 0 | Neutral |
| testneg_9479 | 0.02 | 0.45 | -0.887241 | -3 | 0.390383 | -3 | 0.19898 | 0.12788 | 0.08213 | 0.08913 | 19.09347 | 0.010142886 | 0 | Neutral |
| testneg_9480 | -0.01 | -0.79 | -3.816017 | -3 | 1.51268 | -3 | 0.06095 | 0.08179 | 0.1592 | 0.12788 | 18.76159 | 0.035649254 | 0 | Neutral |
| testneg_9481 | 0.04 | -0.23 | 1.5079 | -3 | -0.216792 | -3 | 0.34694 | 0.20397 | 0.47192 | 0.11305 | 19 | 0.026666667 | 0 | Neutral |
| testneg_9482 | 0.08 | -1.2 | -0.273849 | -3 | -0.120714 | -3 | 0.024 | 0.07726 | 0.99665 | 0.12884 | 22.62994 | 0.08 | 0 | Neutral |
| testneg_9483 | 0 | 0 | 0.395562 | -3 | 1.148993 | -3 | 0.2674 | 0.28725 | 0.32531 | 0.12351 | 15.29221 | 0.01 | 0 | Neutral |
| testneg_9484 | 0 | 0 | 2.181363 | -3 | 9.678287 | -3 | 0.04441 | 0.13603 | 0.09997 | 0.1797 | 22.36648 | 0.605078947 | 92.6 | Disease_causing |
| testneg_9485 | -0.08 | -0.03 | 3.219123 | -3 | 0.440748 | -3 | 0.31924 | 0.11145 | 0.14735 | 0.12788 | 19.09347 | 0.02 | 0 | Neutral |
| testneg_9486 | -0.06 | 0.57 | 1.042833 | -3 | 0.664079 | -3 | 0.19898 | 0.12788 | 0.24 | 0.2063 | 6.87339 | 0.050389267 | 0 | Neutral |
| testneg_9487 | 0.15 | 0.06 | 1.596806 | -3 | 13.89061 | 0.140148 | 0.11796 | 0.12788 | 0.19898 | 0.12788 | 20.27778 | 0.37 | 9.4 | Neutral |
| testneg_9488 | 0 | 0 | 0.173288 | -3 | 4.450954 | -3 | 0.61211 | 0.11059 | 0.19898 | 0.12788 | 23.37087 | 0.054617555 | 0 | Neutral |
| testneg_9489 | 0 | 0 | 1.333395 | -3 | -1.379881 | -3 | 0.12259 | 0.1041 | 0.18106 | 0.12788 | 18.83729 | 0 | 0 | Neutral |
| testneg_9490 | 0 | 0 | 0.339282 | -3 | -0.237401 | -3 | 0.2547 | 0.07589 | 0.07537 | 0.1399 | 12.29167 | 0.05 | 0 | Neutral |
| testneg_9491 | -0.08 | -0.03 | 0.6992 | -3 | 1.452692 | -3 | 0.20772 | 0.16918 | 0.0406 | 0.12788 | 20.27778 | 0.030296425 | 0 | Neutral |
| testneg_9492 | 0.04 | -0.23 | 8.129904 | -3 | 2.309456 | -3 | 0.14668 | 0.25103 | 0.8433 | 0.13128 | 18.8735 | 0.55 | 75.6 | Disease_causing |
| testneg_9493 | -0.08 | -0.03 | -0.657715 | -3 | 0.803253 | -3 | 0.07684 | 0.09138 | 0.05637 | 0.06544 | 20.11299 | 0.03 | 0 | Neutral |
| testneg_9494 | 0.09 | -0.3 | 0.035498 | -3 | 1.24588 | 1.24588 | 0.7639 | 0.10408 | 0.13535 | 0.13562 | 12.18244 | 0.091222148 | 0 | Neutral |
| testneg_9495 | -0.01 | 0.57 | 4.126674 | -3 | -0.388631 | -3 | 0.08579 | 0.6068 | 0.19227 | 0.1096 | 12.1131 | 0.17 | 0 | Neutral |
| testneg_9496 | 0 | 0 | 0.227758 | -3 | 2.947434 | -3 | 0.07874 | 0.12788 | 0.27915 | 0.27706 | 17.06064 | 0.096584901 | 0 | Neutral |
| testneg_9497 | 0.16 | -1.37 | 2.614667 | -3 | 1.672333 | 1.672333 | 0.53654 | 0.22867 | 0.15338 | 0.13614 | 12.40248 | 0.229646932 | 0 | Neutral |
| testneg_9498 | -0.15 | 0.28 | 0.075246 | -3 | 4.737572 | -3 | 0.19898 | 0.12788 | 0.0421 | 0.17669 | 22.17807 | 0.417105191 | 18.6 | Neutral |
| testneg_9499 | 0 | 0 | 0.919505 | -3 | 2.063991 | -3 | 0.19898 | 0.12788 | 0.1134 | 0.12788 | 26.9055 | 0.001140683 | 0 | Neutral |
| testneg_9500 | -0.01 | -0.79 | 4.513115 | -3 | 3.505042 | -3 | 0.03526 | 0.06855 | 0.11858 | 0.11551 | 19.09347 | 0.16 | 0 | Neutral |
| testneg_9501 | 0 | 0 | 0.391204 | -3 | 1.496811 | -3 | 0.19898 | 0.13162 | 0.22582 | 0.12788 | 19.09347 | 0.01 | 0 | Neutral |
| testneg_9502 | 0.14 | -1.01 | 2.658618 | -3 | 0.506959 | -3 | 0.113 | 0.12788 | 0.84236 | 0.14093 | 13.57162 | 0.06 | 0 | Neutral |
| testneg_9503 | 0 | -0.01 | 1.811936 | -3 | 2.811366 | -3 | 0.11447 | 0.12378 | 0.61718 | 0.1648 | 10.41667 | 0.092790687 | 0 | Neutral |
| testneg_9504 | -0.07 | -0.38 | 6.101041 | -3 | -1.278679 | -3 | 0.28732 | 0.11223 | 0.04574 | 0.05824 | 30.45428 | 0.15 | 0 | Neutral |
| testneg_9505 | 0 | 0 | -0.785566 | -3 | 1.217734 | -3 | 0.01196 | 0.12788 | 0.18404 | 0.14494 | 12.7356 | 0.07 | 0 | Neutral |
| testneg_9506 | -0.14 | 1.01 | -0.55255 | -3 | 2.508934 | -3 | 0.06793 | 0.12788 | 0.10895 | 0.10442 | 12.59789 | 0.11 | 0 | Neutral |
| testneg_9507 | 0.08 | 0.03 | -1.726521 | -3 | -1.052595 | -3 | 0.03502 | 0.08119 | 0.19898 | 0.12788 | 18.76159 | 0.04 | 0 | Neutral |
| testneg_9508 | 0.17 | 0.38 | 1.675638 | -3 | 6.92218 | -3 | 0.39163 | 0.2681 | 0.15767 | 0.12788 | 17.21847 | 0.451209615 | 35.4 | Neutral |
| testneg_9509 | 0 | -0.01 | 0.389904 | -3 | -1.653555 | -1.653555 | 0.95998 | 0.11262 | 0.14588 | 0.12156 | 10.41667 | 0.01 | 0 | Neutral |
| testneg_9510 | 0 | 0 | 0.942465 | -3 | 3.287571 | -3 | 0.24536 | 0.38555 | 0.2852 | 0.12788 | 10 | 0.072180556 | 0 | Neutral |
| testneg_9511 | 0 | 0 | 1.108803 | -3 | 0.411846 | -3 | 0.85092 | 0.58871 | 0.10133 | 0.32155 | 10.41667 | 0.108740801 | 0 | Neutral |
| testneg_9512 | -0.11 | 0.19 | 5.824607 | -3 | -1.224645 | -3 | 0.03391 | 0.09745 | 0.10187 | 0.08499 | 33.40108 | 0.19 | 0 | Neutral |
| testneg_9513 | -0.06 | 0.57 | 2.959696 | -3 | 0.827687 | -3 | 0.19898 | 0.15588 | 0.39576 | 0.10761 | 13.08587 | 0.07 | 0 | Neutral |
| testneg_9514 | 0 | 0 | 4.037148 | -3 | -1.833303 | -3 | 0.92832 | 0.16306 | 0.19898 | 0.12788 | 12.6824 | 0.07 | 0 | Neutral |
| testneg_9515 | -0.14 | 0.37 | -0.45388 | -3 | 0.349562 | -3 | 0.39964 | 0.22293 | 0.22686 | 0.24974 | 18.33333 | 0.151356345 | 0 | Neutral |

| | | | | | | | | | | | | | | |
|---|---|---|---|---|---|---|---|---|---|---|---|---|---|---|
| testneg_9516 | 0 | 0 | -0.926281 | -0.926281 | -0.173726 | -3 | 0.77196 | 0.14089 | 0.12543 | 0.12788 | 26.84914 | 0.030763376 | 0 | Neutral |
| testneg_9517 | 0.06 | -0.57 | -0.383439 | -3 | 0.052034 | -3 | 0.23152 | 0.09962 | 0.21807 | 0.12788 | 13.44153 | 0.000101382 | 0 | Neutral |
| testneg_9518 | 0.07 | -0.31 | 4.644707 | -3 | 1.795415 | -3 | 0.27614 | 0.12085 | 0.65717 | 0.16156 | 19.56877 | 0.217154727 | 0.4 | Neutral |
| testneg_9519 | -0.08 | -0.03 | 1.588775 | -3 | -0.285825 | -3 | 0.24 | 0.2063 | 0.15016 | 0.08826 | 19.44444 | 0.04 | 0 | Neutral |
| testneg_9520 | -0.02 | 0.63 | 0.385855 | 0.385855 | 7.216753 | -3 | 0.10153 | 0.13343 | 0.07297 | 0.12788 | 18.07168 | 0.37 | 9.8 | Neutral |
| testneg_9521 | 0.02 | 0.45 | 6.053701 | -3 | -0.35214 | -0.35214 | 0.43918 | 0.46357 | 0.11134 | 0.13818 | 22.47185 | 0.439615385 | 30.4 | Neutral |
| testneg_9522 | 0 | 0 | 1.626934 | -3 | 2.178935 | -3 | 0.20942 | 0.14229 | 0.77847 | 0.10751 | 12.30981 | 0.01 | 0 | Neutral |
| testneg_9523 | 0.02 | 0.45 | 2.390589 | -3 | 0.586936 | -3 | 0.19002 | 0.12138 | 0.3405 | 0.1816 | 18.87903 | 0.06 | 0 | Neutral |
| testneg_9524 | 0 | 0 | -0.064878 | -3 | 0.852085 | -3 | 0.99668 | 0.30319 | 0.56554 | 0.76978 | 5.69444 | 0.086433722 | 0 | Neutral |
| testneg_9525 | 0 | -0.01 | 1.116052 | -3 | 4.591087 | -3 | 0.1631 | 0.40344 | 0.15116 | 0.13759 | 51.31523 | 0.437817619 | 31.6 | Neutral |
| testneg_9526 | 0 | 0 | -0.480058 | -3 | 4.259881 | -3 | 0.2029 | 0.12788 | 0.38756 | 0.08351 | 19.57845 | 0.037122396 | 0 | Neutral |
| testneg_9527 | 0 | 0 | 0.606984 | -3 | 2.537219 | -3 | 0.08932 | 0.12788 | 0.0328 | 0.0691 | 16.42913 | 0.01036747 | 0 | Neutral |
| testneg_9528 | -0.08 | -0.03 | -0.741718 | -3 | 1.435364 | -3 | 0.06218 | 0.10661 | 0.20357 | 0.12788 | 30.48962 | 0.03 | 0 | Neutral |
| testneg_9529 | 0.08 | 0.03 | 2.365497 | -3 | 0.994999 | -3 | 0.61491 | 0.11954 | 0.51282 | 0.12075 | 12.0794 | 0.020402576 | 0 | Neutral |
| testneg_9530 | 0.09 | -0.3 | -1.680478 | -3 | 4.34233 | -3 | 0.1664 | 0.10113 | 0.31648 | 0.12788 | 34.22114 | 0.07 | 0 | Neutral |
| testneg_9531 | -0.03 | 1.03 | 2.366503 | -3 | 0.251655 | -3 | 0.96879 | 0.38765 | 0.27058 | 0.13363 | 4.69594 | 0.143210784 | 0 | Neutral |
| testneg_9532 | 0 | 0 | 0.818501 | -3 | 2.273291 | -3 | 0.19898 | 0.12788 | 0.19898 | 0.12788 | 21.94444 | 0.001003715 | 0 | Neutral |
| testneg_9533 | 0 | -0.01 | 1.057977 | -3 | 0.374605 | -3 | 0.15623 | 0.1447 | 0.45112 | 0.12083 | 13.07204 | 0.00036039 | 0 | Neutral |
| testneg_9534 | -0.06 | 0.57 | -0.170377 | -3 | -0.284048 | -3 | 0.10278 | 0.08667 | 0.16709 | 0.08659 | 19.44444 | 0.03 | 0 | Neutral |
| testneg_9535 | -0.02 | -0.41 | 2.303477 | -3 | 0.699729 | -0.925939 | 0.53216 | 0.84775 | 0.39192 | 0.09135 | 10.83333 | 0.07 | 0 | Neutral |
| testneg_9536 | -0.01 | -0.79 | 0.119573 | -3 | 5.867155 | -3 | 0.29451 | 0.13582 | 0.13067 | 0.1005 | 13.51351 | 0.130741758 | 0 | Neutral |
| testneg_9537 | -0.14 | 1.85 | 5.509658 | -3 | 4.027424 | -3 | 0.16485 | 0.13045 | 0.16926 | 0.11871 | 22.86278 | 0.44 | 30.4 | Neutral |
| testneg_9538 | -0.05 | -0.29 | 2.605779 | -3 | -1.505295 | -3 | 0.22817 | 0.10634 | 0.29525 | 0.12788 | 27.5 | 0.01 | 0 | Neutral |
| testneg_9539 | 0.19 | 0.51 | 3.28427 | -3 | 3.136188 | -3 | 0.08982 | 0.15279 | 0.06617 | 0.12788 | 21.56188 | 0.21 | 0.2 | Neutral |
| testneg_9540 | -0.18 | -0.16 | 2.826632 | 1.614823 | 0.273493 | -3 | 0.52388 | 0.25831 | 0.09353 | 0.12788 | 10.41976 | 0.14 | 0 | Neutral |
| testneg_9541 | 0.14 | -1.85 | 5.761185 | -3 | 3.227121 | -3 | 0.47217 | 0.15544 | 0.47462 | 0.10826 | 11.95767 | 0.435620173 | 29.8 | Neutral |
| testneg_9542 | 0.09 | -0.3 | 0.035498 | -3 | 0.918679 | -3 | 0.7639 | 0.10408 | 0.43314 | 0.67252 | 13.12884 | 0.088 | 0 | Neutral |
| testneg_9543 | 0 | 0 | 0.618993 | 0.618993 | -1.097876 | -3 | 0.04207 | 0.12788 | 0.12317 | 0.12345 | 48.08616 | 0.06 | 0 | Neutral |
| testneg_9544 | 0 | 0 | 0.546525 | -3 | 2.829858 | -3 | 0.12545 | 0.08963 | 0.10895 | 0.10442 | 16.60044 | 0.020327957 | 0 | Neutral |
| testneg_9545 | 0 | 0 | 1.01324 | -3 | 1.252341 | -3 | 0.45354 | 0.09957 | 0.38072 | 0.15993 | 20.83333 | 0.07 | 0 | Neutral |
| testneg_9546 | -0.06 | 0.57 | 2.133607 | -3 | 0.200393 | -3 | 0.46837 | 0.14896 | 0.19898 | 0.08714 | 18.88889 | 0.031 | 0 | Neutral |
| testneg_9547 | 0 | -0.01 | 2.237213 | -3 | 4.592525 | -3 | 0.31244 | 0.4615 | 0.1056 | 0.12788 | 13.08333 | 0.180070513 | 0 | Neutral |
| testneg_9548 | 0.18 | -1.05 | -0.806649 | -3 | 1.350468 | -3 | 0.37065 | 0.12788 | 0.7413 | 0.18974 | 13.78431 | 0.079309524 | 0 | Neutral |
| testneg_9549 | 0 | 0 | 1.261253 | -3 | 1.849259 | -3 | 0.09028 | 0.10643 | 0.41261 | 0.12096 | 13.51351 | 0 | 0 | Neutral |
| testneg_9550 | 0 | 0 | 1.323079 | -3 | 7.525025 | -3 | 0.5103 | 0.10923 | 0.12943 | 0.14391 | 21.25203 | 0.413963027 | 21.4 | Neutral |
| testneg_9551 | -0.03 | 1.03 | 6.067908 | -3 | 4.027424 | -3 | 0.3047 | 0.11929 | 0.16926 | 0.11871 | 18.33333 | 0.453534795 | 36 | Neutral |
| testneg_9552 | 0.19 | 0.51 | 6.664454 | -3 | 2.018667 | -3 | 0.9237 | 0.55859 | 0.39344 | 0.12788 | 10.41667 | 0.486668362 | 49 | Neutral |
| testneg_9553 | 0 | 0 | 0.261866 | -3 | 5.894587 | -3 | 0.35584 | 0.10874 | 0.15364 | 0.08789 | 21.2963 | 0.074677193 | 0 | Neutral |
| testneg_9554 | 0.06 | -0.57 | 5.676784 | -3 | 3.889468 | -3 | 0.19898 | 0.12788 | 0.12867 | 0.11159 | 14.16667 | 0.30656853 | 4.2 | Neutral |
| testneg_9555 | -0.07 | -0.38 | 2.28341 | -3 | -0.433937 | -3 | 0.27723 | 0.13229 | 0.84572 | 0.78546 | 16.84579 | 0.112168534 | 0 | Neutral |
| testneg_9556 | 0.21 | 0.15 | 5.338361 | -3 | 4.061859 | -3 | 0.21587 | 0.11701 | 0.04047 | 0.12788 | 17.21847 | 0.3 | 4.4 | Neutral |
| testneg_9557 | -0.1 | -1.9 | 1.643871 | -3 | 3.650561 | -3 | 0.19898 | 0.12788 | 0.0961 | 0.21256 | 13.27141 | 0.286856927 | 0.8 | Neutral |
| testneg_9558 | 0 | 0 | -0.211085 | -3 | 1.464574 | 1.464574 | 0.04543 | 0.28103 | 0.60603 | 0.12788 | 18.33333 | 0.09 | 0 | Neutral |
| testneg_9559 | -0.08 | -0.03 | 1.483649 | 1.483649 | 2.499111 | -3 | 0.47678 | 0.38085 | 0.58994 | 0.11602 | 32.46216 | 0.119666667 | 0 | Neutral |
| testneg_9560 | 0.01 | 0.8 | 2.947434 | -3 | 5.829899 | -3 | 0.27915 | 0.27706 | 0.06196 | 0.10881 | 18.88889 | 0.360050505 | 10 | Neutral |
| testneg_9561 | 0.01 | 0.8 | 4.915244 | -3 | 0.730644 | -3 | 0.47975 | 0.18568 | 0.16545 | 0.10002 | 18.88889 | 0.12 | 0 | Neutral |
| testneg_9562 | 0.09 | -0.3 | 5.519487 | -3 | 5.817235 | 5.817235 | 0.74919 | 0.85508 | 0.19898 | 0.12788 | 26.72149 | 0.71 | 99.6 | Disease_causing |
| testneg_9563 | 0 | 0 | 3.637193 | -3 | 0.425569 | 0.425569 | 0.99976 | 0.28689 | 0.12879 | 0.12788 | 11.98582 | 0.1 | 0 | Neutral |
| testneg_9564 | 0.01 | -1.44 | 5.296637 | -3 | 3.910833 | -3 | 0.84365 | 0.12788 | 0.1201 | 0.1062 | 5.70654 | 0.354104167 | 6.6 | Neutral |
| testneg_9565 | 0 | -0.01 | -0.099884 | -3 | 1.239593 | -3 | 0.0887 | 0.10047 | 0.60755 | 0.41186 | 31.73501 | 0.07 | 0 | Neutral |
| testneg_9566 | 0.07 | 0.38 | 3.047706 | -3 | 1.436812 | -3 | 0.0695 | 0.12788 | 0.29302 | 0.17394 | 21.81672 | 0.19 | 0 | Neutral |
| testneg_9567 | -0.12 | 1.14 | 3.68988 | -3 | 0.583229 | -3 | 0.62864 | 0.11103 | 0.19898 | 0.12788 | 8.97898 | 0.090342491 | 0 | Neutral |
| testneg_9568 | -0.05 | -0.29 | -1.527311 | -3 | 3.765273 | -3 | 0.64254 | 0.10729 | 0.48817 | 0.10601 | 10.26553 | 0.078119048 | 0 | Neutral |
| testneg_9569 | 0 | -0.01 | 4.094824 | -3 | 4.922988 | -3 | 0.16468 | 0.11275 | 0.33264 | 0.10493 | 34.90691 | 0.157272232 | 0 | Neutral |
| testneg_9570 | -0.15 | -0.04 | 2.203067 | -3 | 5.984999 | -3 | 0.27887 | 0.18663 | 0.59245 | 0.12788 | 27.4024 | 0.34 | 4 | Neutral |
| testneg_9571 | 0.09 | -0.3 | 4.862066 | -3 | 1.181316 | -3 | 0.19898 | 0.12788 | 0.2293 | 0.12788 | 33.66132 | 0.041742424 | 0 | Neutral |
| testneg_9572 | 0 | 0 | -0.32247 | -3 | 4.970317 | -3 | 0.35092 | 0.10596 | 0.05887 | 0.10046 | 29.98054 | 0.07 | 0 | Neutral |
| testneg_9573 | 0 | 0 | -0.465388 | -3 | 3.163542 | -3 | 0.29838 | 0.63658 | 0.10751 | 0.08512 | 22.62411 | 0.121533412 | 0 | Neutral |
| testneg_9574 | 0.21 | 0.15 | 1.623618 | -3 | 1.632331 | -3 | 0.10083 | 0.17511 | 0.098 | 0.08791 | 19.09347 | 0.06 | 0 | Neutral |
| testneg_9575 | 0 | -0.01 | 0.522695 | -3 | 5.829822 | -3 | 0.84616 | 0.1471 | 0.06895 | 0.06354 | 17.77778 | 0.101612146 | 0 | Neutral |
| testneg_9576 | 0 | 0 | 1.115221 | -3 | 2.931463 | -3 | 0.64188 | 0.09494 | 0.27585 | 0.20571 | 5 | 0.086501032 | 0 | Neutral |
| testneg_9577 | -0.2 | 1.26 | 3.523664 | -3 | 1.296281 | -3 | 0.1247 | 0.10331 | 0.63484 | 0.16728 | 15.77058 | 0.230086207 | 0 | Neutral |
| testneg_9578 | 0.03 | 0.12 | 0.235146 | -3 | 1.040567 | -3 | 0.16856 | 0.19221 | 0.10869 | 0.1503 | 22.36648 | 0.14826462 | 0 | Neutral |
| testneg_9579 | 0.09 | -0.3 | 3.637231 | -3 | 4.228814 | -3 | 0.30042 | 0.13517 | 0.07675 | 0.11464 | 12.46106 | 0.202936508 | 0 | Neutral |
| testneg_9580 | 0 | 0 | 1.170021 | -3 | 1.019206 | -3 | 0.68568 | 0.65219 | 0.50388 | 0.12295 | 14.49301 | 0.020738262 | 0 | Neutral |
| testneg_9581 | -0.03 | -0.12 | -0.107462 | -3 | -0.034419 | -0.034419 | 0.07982 | 0.12788 | 0.62203 | 0.12788 | 28.07676 | 0.02 | 0 | Neutral |
| testneg_9582 | 0 | 0 | 1.323164 | -3 | -0.707182 | -3 | 0.22761 | 0.14229 | 0.06789 | 0.09707 | 9.38938 | 0.000521351 | 0 | Neutral |
| testneg_9583 | 0 | 0 | 0.105719 | -3 | 2.996727 | -3 | 0.98442 | 0.44966 | 0.19898 | 0.17518 | 30.46043 | 0.243366748 | 0.2 | Neutral |
| testneg_9584 | 0.18 | 0.16 | 5.112733 | -3 | 2.091567 | -3 | 0.03676 | 0.12788 | 0.19898 | 0.12788 | 13.14653 | 0.134331984 | 0 | Neutral |
| testneg_9585 | 0.04 | -0.23 | 5.929272 | -3 | 3.236399 | -3 | 0.79983 | 0.1105 | 0.22953 | 0.0919 | 18.95079 | 0.28 | 2.4 | Neutral |
| testneg_9586 | 0.14 | -0.37 | 3.266214 | -3 | -0.100872 | -3 | 0.1 | 0.22633 | 0.11891 | 0.11186 | 9.377 | 0.08 | 0 | Neutral |
| testneg_9587 | 0.09 | -0.3 | 5.344639 | -3 | 0.148762 | 0.148762 | 0.6992 | 0.1804 | 0.44644 | 0.10813 | 12.70833 | 0.19 | 0 | Neutral |
| testneg_9588 | -0.05 | -0.29 | -1.117253 | -3 | -0.81151 | -3 | 0.84682 | 0.35105 | 0.01635 | 0.12788 | 12.31231 | 0.04 | 0 | Neutral |
| testneg_9589 | -0.03 | -0.12 | 0.550914 | -3 | -0.518868 | -3 | 0.17322 | 0.10311 | 0.08386 | 0.12788 | 26.9182 | 0.01 | 0 | Neutral |
| testneg_9590 | 0.05 | -0.81 | 7.149184 | -3 | 0.573644 | -0.38834 | 0.52278 | 0.41207 | 0.8433 | 0.13128 | 13.06467 | 0.414166667 | 20.6 | Neutral |
| testneg_9591 | -0.03 | -1.22 | 0.713058 | -3 | 3.547766 | -3 | 0.14375 | 0.12788 | 0.05124 | 0.07099 | 18.75015 | 0.06003876 | 0 | Neutral |
| testneg_9592 | 0.14 | -0.37 | 0.198379 | -3 | 3.225783 | -3 | 0.03991 | 0.08342 | 0.75978 | 0.11423 | 12.37434 | 0.090371774 | 0 | Neutral |
| testneg_9593 | 0.15 | 0.04 | 1.205837 | -3 | 4.280436 | -3 | 0.07071 | 0.11499 | 0.40116 | 0.43982 | 18.22258 | 0.312118789 | 2.8 | Neutral |
| testneg_9594 | 0.14 | -1.01 | 2.081126 | -3 | -0.40166 | -3 | 0.2663 | 0.32806 | 0.08683 | 0.26961 | 9.78453 | 0.200818247 | 0 | Neutral |
| testneg_9595 | 0.14 | -0.37 | 3.348627 | -3 | 0.472156 | -3 | 0.04808 | 0.10018 | 0.15446 | 0.08618 | 13.2927 | 0.05 | 0 | Neutral |
| testneg_9596 | 0.04 | -0.23 | 0.007971 | -3 | -0.060754 | -3 | 0.12591 | 0.09845 | 0.70946 | 0.11809 | 8.9812 | 0.01 | 0 | Neutral |
| testneg_9597 | 0 | 0 | 0.503836 | -3 | 1.476163 | 1.476163 | 0.19898 | 0.07875 | 0.19898 | 0.09395 | 24.12281 | 0.02 | 0 | Neutral |
| testneg_9598 | 0.02 | 0.45 | 5.507024 | -3 | 3.206825 | 3.206825 | 0.21587 | 0.11701 | 0.55197 | 0.11305 | 17.01711 | 0.347511111 | 8.6 | Neutral |
| testneg_9599 | -0.07 | -0.33 | 0.222047 | -3 | 2.901134 | -3 | 0.19898 | 0.12788 | 0.82014 | 0.14054 | 11.1452 | 0.1 | 0 | Neutral |
| testneg_9600 | 0 | 0 | 0.676994 | -3 | 0.865658 | -3 | 0.09356 | 0.2448 | 0.64108 | 0.51829 | 10.55958 | 0.111204365 | 0 | Neutral |
| testneg_9601 | 0 | -0.01 | 1.519608 | -3 | 5.86772 | -3 | 0.18206 | 0.10727 | 0.25209 | 0.10029 | 13.02635 | 0.060346476 | 0 | Neutral |
| testneg_9602 | 0.17 | 0.38 | 1.603588 | -3 | 5.907423 | -3 | 0.10496 | 0.21491 | 0.00451 | 0.2022 | 12.80035 | 0.595034848 | 85.8 | Disease_causing |
| testneg_9603 | -0.07 | -0.38 | -1.066068 | -3 | 6.717504 | -3 | 0.04515 | 0.14318 | 0.19898 | 0.12788 | 18.88889 | 0.340575092 | 6.6 | Neutral |
| testneg_9604 | 0 | -0.01 | 2.148855 | -3 | 1.959161 | -3 | 0.56904 | 0.14839 | 0.18312 | 0.12788 | 13.02635 | 0.012069742 | 0 | Neutral |
| testneg_9605 | 0.01 | 0.8 | 0.851144 | -3 | -3.346595 | -3 | 0.10004 | 0.12788 | 0.24949 | 0.29709 | 10.08333 | 0.07 | 0 | Neutral |
| testneg_9606 | -0.02 | 0.68 | 5.284085 | -3 | 1.001127 | -3 | 0.08509 | 0.12788 | 0.14 | 0.12151 | 26.70811 | 0.14 | 0 | Neutral |
| testneg_9607 | 0 | 0 | 1.323079 | -3 | -0.751147 | -3 | 0.5103 | 0.10923 | 0.09972 | 0.2187 | 31.06426 | 0.085 | 0 | Neutral |
| testneg_9608 | 0.05 | 0.29 | 3.186451 | -3 | 5.312389 | -3 | 0.17141 | 0.1104 | 0.07293 | 0.12989 | 11.35684 | 0.23 | 0.4 | Neutral |
| testneg_9609 | 0 | 0 | 1.185651 | -3 | 5.788486 | -3 | 0.87813 | 0.23021 | 0.15025 | 0.4872 | 13.77727 | 0.351719105 | 10.4 | Neutral |
| testneg_9610 | 0 | 0 | 1.51996 | -3 | 0.445956 | -3 | 0.09105 | 0.27434 | 0.2172 | 0.10891 | 51.12724 | 0.04 | 0 | Neutral |
| testneg_9611 | 0.02 | -0.63 | -0.055258 | -3 | 1.557244 | -3 | 0.16139 | 0.09116 | 0.0538 | 0.0823 | 23.24074 | 0.02 | 0 | Neutral |
| testneg_9612 | 0 | 0 | 0.824678 | -3 | 1.744987 | -3 | 0.6854 | 0.12563 | 0.06552 | 0.07762 | 15.9293 | 0.01 | 0 | Neutral |
| testneg_9613 | 0 | 0 | -0.137475 | -3 | 1.782913 | -3 | 0.19898 | 0.12788 | 0.1327 | 0.09885 | 21.52125 | 0.01 | 0 | Neutral |
| testneg_9614 | -0.01 | -0.79 | 1.842441 | -3 | -0.687024 | -3 | 0.05904 | 0.09282 | 0.19898 | 0.20319 | 30.33382 | 0.111414052 | 0 | Neutral |
| testneg_9615 | 0.14 | -0.37 | 3.506128 | -3 | 4.679417 | -3 | 0.27605 | 0.12788 | 0.06939 | 0.08981 | 24.52381 | 0.152269841 | 0 | Neutral |
| testneg_9616 | -0.01 | -0.79 | 0.443004 | -3 | 4.894604 | -3 | 0.1551 | 0.09276 | 0.21726 | 0.23214 | 13.11773 | 0.266225026 | 1.4 | Neutral |
| testneg_9617 | 0 | 0 | 0.648536 | -3 | 2.532517 | -3 | 0.31124 | 0.08875 | 0.0538 | 0.0823 | 12.37434 | 0.02 | 0 | Neutral |
| testneg_9618 | 0 | 0 | 0.185804 | -3 | 0.288206 | -3 | 0.08306 | 0.12788 | 0.25462 | 0.12788 | 13.51351 | 0 | 0 | Neutral |
| testneg_9619 | 0.14 | -0.69 | 4.914058 | -3 | 0.705249 | -3 | 0.44594 | 0.13793 | 0.1324 | 0.09503 | 21.91163 | 0.08525 | 0 | Neutral |
| testneg_9620 | -0.04 | 0.23 | 0.729199 | -3 | 4.319571 | -3 | 0.11373 | 0.10739 | 0.36324 | 0.15729 | 10.22898 | 0.277106215 | 1.6 | Neutral |
| testneg_9621 | 0.01 | 0.8 | 1.937875 | -3 | -0.03839 | -3 | 0.16393 | 0.08897 | 0.0895 | 0.12788 | 38.0826 | 0.02 | 0 | Neutral |
| testneg_9622 | 0 | 0 | -0.734901 | -3 | 5.050499 | -3 | 0.64721 | 0.8758 | 0.12819 | 0.1229 | 10.41667 | 0.14 | 0 | Neutral |
| testneg_9623 | -0.08 | -0.03 | -0.038604 | -3 | 1.635811 | -3 | 0.493 | 0.35835 | 0.11187 | 0.10744 | 16.8018 | 0.05 | 0 | Neutral |
| testneg_9624 | 0.14 | -0.69 | 7.221815 | -3 | 0.970366 | -3 | 0.81305 | 0.17045 | 0.19898 | 0.12788 | 16.48772 | 0.370833333 | 14 | Neutral |
| testneg_9625 | 0.09 | -0.3 | 5.756209 | -3 | 0.464102 | -3 | 0.06845 | 0.13843 | 0.14668 | 0.25103 | 20.27778 | 0.492629605 | 52.6 | Disease_causing |
| testneg_9626 | 0.01545455 | -0.092 | 6.776197 | -3 | 1.643466 | -3 | 0.7219 | 0.41509 | 0.19343 | 0.11347 | 18.63732 | 0.4505 | 38 | Neutral |
| testneg_9627 | -0.05 | 0.44 | 3.587885 | -1.918971 | 5.349526 | -3 | 0.77644 | 0.09695 | 0.06397 | 0.12788 | 30.80739 | 0.231286569 | 0 | Neutral |
| testneg_9628 | 0 | 0 | 1.799258 | -3 | -0.803634 | -3 | 0.204 | 0.15424 | 0.16392 | 0.12788 | 10.83333 | 0.010421473 | 0 | Neutral |
| testneg_9629 | 0.05 | -0.81 | 1.432298 | 1.432298 | 5.362959 | -3 | 0.49512 | 0.08696 | 0.31507 | 0.61057 | 7.00579 | 0.411400463 | 21.2 | Neutral |
| testneg_9630 | -0.15 | -0.06 | -1.278679 | -3 | 2.653525 | -3 | 0.04574 | 0.05824 | 0.20546 | 0.10734 | 30.45428 | 0.12 | 0 | Neutral |
| testneg_9631 | -0.08 | -0.03 | 3.894533 | -3 | 1.162064 | -3 | 0.11187 | 0.10744 | 0.64608 | 0.35524 | 17.44725 | 0.22 | 0.2 | Neutral |
| testneg_9632 | -0.08 | -0.03 | 5.050499 | -3 | 1.250385 | 0.888376 | 0.12819 | 0.1229 | 0.21908 | 0.12788 | 10.3336 | 0.1 | 0 | Neutral |
| testneg_9633 | -0.08 | -0.03 | 0.950149 | -3 | 3.660063 | -3 | 0.31318 | 0.11262 | 0.22162 | 0.09445 | 20.6326 | 0.04185008 | 0 | Neutral |
| testneg_9634 | 0 | 0 | -0.090737 | -3 | 0.359116 | -3 | 0.09734 | 0.08962 | 0.19898 | 0.12788 | 5.92593 | 0.000257734 | 0 | Neutral |
| testneg_9635 | -0.05 | 0.81 | 2.250533 | -3 | 0.461408 | -3 | 0.14021 | 0.43531 | 0.01642 | 0.12788 | 12.11131 | 0.07 | 0 | Neutral |
| testneg_9636 | 0.04 | -0.23 | -0.624462 | -3 | 6.087637 | -3 | 0.24038 | 0.15296 | 0.098 | 0.08791 | 17.21847 | 0.190857143 | 0.2 | Neutral |
| testneg_9637 | -0.04 | 0.23 | -2.175398 | -3 | -0.985486 | -3 | 0.10228 | 0.22182 | 0.2755 | 0.12788 | 22.76087 | 0.062435185 | 0 | Neutral |

| | | | | | | | | | | | | | | |
|---|---|---|---|---|---|---|---|---|---|---|---|---|---|---|
| testneg_9638 | -0.01 | -0.79 | -0.307406 | -3 | 0.143249 | -3 | 0.62858 | 0.16645 | 0.43691 | 0.12794 | 13.84202 | 0.03 | 0 | Neutral |
| testneg_9639 | 0.01 | 0.8 | 2.217361 | -3 | 2.322245 | -3 | 0.11829 | 0.12788 | 0.30734 | 0.12788 | 33.14935 | 0.03 | 0 | Neutral |
| testneg_9640 | 0 | 0 | 3.466764 | -3 | 1.748821 | -3 | 0.34205 | 0.11195 | 0.47816 | 0.66526 | 10.41667 | 0.15293633 | 0 | Neutral |
| testneg_9641 | 0.02 | -0.63 | 1.636375 | -3 | 1.345014 | -3 | 0.11905 | 0.23747 | 0.3437 | 0.5169 | 10.41667 | 0.18237521 | 0 | Neutral |
| testneg_9642 | 0.23 | -1.04 | 3.675216 | -3 | 1.847342 | -3 | 0.16468 | 0.11275 | 0.18206 | 0.10727 | 13.51351 | 0.050455487 | 0 | Neutral |
| testneg_9643 | 0 | 0 | 0.462587 | -3 | 0.774689 | 0.774689 | 0.26014 | 0.12378 | 0.24219 | 0.17458 | 13.2073 | 0.04 | 0 | Neutral |
| testneg_9644 | 0.06 | 0.18 | -1.722922 | -3 | -0.279308 | -3 | 0.06704 | 0.1068 | 0.12885 | 0.61089 | 27.51545 | 0.11070197 | 0 | Neutral |
| testneg_9645 | 0 | 0 | 0.29846 | -3 | 2.272958 | -3 | 0.07543 | 0.10211 | 0.13001 | 0.13176 | 21.53664 | 0.07 | 0 | Neutral |
| testneg_9646 | -0.03 | -0.1 | 2.735353 | -3 | 1.644806 | -3 | 0.29478 | 0.12788 | 0.11172 | 0.10034 | 10.30286 | 0.02 | 0 | Neutral |
| testneg_9647 | 0.02 | 0.45 | -2.670084 | -3 | -0.348662 | -3 | 0.19898 | 0.09171 | 0.10569 | 0.09059 | 32.30375 | 0.030106113 | 0 | Neutral |
| testneg_9648 | -0.17 | -0.38 | 3.887632 | -3 | 0.699729 | -0.925939 | 0.45112 | 0.12083 | 0.39192 | 0.09135 | 14.21227 | 0.06855 | 0 | Neutral |
| testneg_9649 | -0.05 | -0.29 | 3.952579 | -3 | -0.236104 | -3 | 0.23088 | 0.21602 | 0.48075 | 0.20313 | 4.72222 | 0.312142595 | 5 | Neutral |
| testneg_9650 | 0 | 0 | 1.663039 | -3 | 0.196475 | -3 | 0.11395 | 0.26806 | 0.15562 | 0.10384 | 12.7356 | 0.02 | 0 | Neutral |
| testneg_9651 | 0.05 | -0.81 | 0.237851 | -3 | 1.620568 | -3 | 0.1072 | 0.63446 | 0.30471 | 0.11609 | 28.86444 | 0.110350985 | 0 | Neutral |
| testneg_9652 | -0.08 | -0.03 | 0.355585 | -3 | -1.989097 | -3 | 0.13195 | 0.57267 | 0.11094 | 0.12788 | 10.41667 | 0.081339286 | 0 | Neutral |
| testneg_9653 | 0 | 0 | 1.416689 | 1.416689 | 0.79796 | 0.79796 | 0.33163 | 0.26601 | 0.57466 | 0.12788 | 4.44444 | 0.04 | 0 | Neutral |
| testneg_9654 | 0.01 | 0.79 | -0.623999 | -3 | -0.11312 | -3 | 0.19301 | 0.10828 | 0.19894 | 0.27411 | 10 | 0.05 | 0 | Neutral |
| testneg_9655 | -0.08 | -0.03 | 2.022784 | -3 | 1.587828 | -3 | 0.84616 | 0.1471 | 0.41743 | 0.78545 | 10 | 0.122810428 | 0 | Neutral |
| testneg_9656 | 0 | 0 | 1.848117 | -3 | 0.304305 | -3 | 0.10249 | 0.1399 | 0.50134 | 0.15488 | 48.8214 | 0.082108696 | 0 | Neutral |
| testneg_9657 | 0.09 | -0.3 | 1.845442 | -3 | 1.347794 | 1.08973 | 0.99293 | 0.79067 | 0.15344 | 0.16141 | 10.83333 | 0.211092469 | 0 | Neutral |
| testneg_9658 | 0 | 0 | 0.175869 | -3 | 6.034791 | -3 | 0.43897 | 0.12441 | 0.10422 | 0.0994 | 11.21556 | 0.1 | 0 | Neutral |
| testneg_9659 | -0.2 | 1.26 | 4.283648 | -3 | 0.822036 | -3 | 0.19898 | 0.11691 | 0.13575 | 0.12788 | 22.14009 | 0.13 | 0 | Neutral |
| testneg_9660 | 0 | 0 | 1.034225 | -3 | 0.774689 | 0.774689 | 0.23913 | 0.12788 | 0.24219 | 0.17458 | 18.6768 | 0.05 | 0 | Neutral |
| testneg_9661 | 0 | 0 | 1.470632 | -3 | 1.473658 | -3 | 0.67549 | 0.1273 | 0.59302 | 0.10897 | 17.42866 | 0.000282194 | 0 | Neutral |
| testneg_9662 | -0.04 | 0.23 | 2.344511 | -3 | -1.367472 | -3 | 0.42585 | 0.18081 | 0.049 | 0.11324 | 33.45973 | 0.040494505 | 0 | Neutral |
| testneg_9663 | -0.14 | 1.85 | 1.695811 | -3 | 4.666155 | -3 | 0.10371 | 0.08655 | 0.15152 | 0.12645 | 17.21847 | 0.239761905 | 0 | Neutral |
| testneg_9664 | 0.18 | -1.05 | 8.410694 | -3 | -1.401819 | -3 | 0.08646 | 0.24855 | 0.19898 | 0.12788 | 26.64043 | 0.39 | 15.4 | Neutral |
| testneg_9665 | 0 | 0 | -0.035358 | -3 | 1.5447 | 1.5447 | 0.19898 | 0.07312 | 0.08932 | 0.32882 | 27.51545 | 0.147252798 | 0 | Neutral |
| testneg_9666 | 0.17 | -1.22 | 5.736168 | -3 | 0.309305 | -3 | 0.33245 | 0.45921 | 0.16973 | 0.17555 | 23.19121 | 0.54 | 73.2 | Disease_causing |
| testneg_9667 | 0.03 | -1.03 | 1.284996 | -3 | 0.100154 | 0.100154 | 0.52028 | 0.47239 | 0.21122 | 0.12686 | 11.875 | 0.06 | 0 | Neutral |
| testneg_9668 | -0.05 | -0.29 | 5.247936 | -3 | 5.075013 | -3 | 0.0832 | 0.09323 | 0.47019 | 0.12788 | 19.10253 | 0.35 | 11 | Neutral |
| testneg_9669 | 0 | 0 | 0.140815 | -3 | 5.393665 | 5.393665 | 0.51278 | 0.35859 | 0.03809 | 0.16044 | 10.41667 | 0.54 | 70.8 | Disease_causing |
| testneg_9670 | 0.2 | -1.26 | 1.668583 | -3 | 3.553199 | -3 | 0.04869 | 0.05922 | 0.16601 | 0.14015 | 34.07441 | 0.25 | 0.2 | Neutral |
| testneg_9671 | 0.03 | 0.12 | 0.235146 | -3 | 4.60551 | -3 | 0.16856 | 0.19221 | 0.29958 | 0.11203 | 22.36648 | 0.152743305 | 0 | Neutral |
| testneg_9672 | 0 | 0 | 0.275913 | -3 | 0.425597 | -3 | 0.61027 | 0.14229 | 0.13023 | 0.29692 | 12.86501 | 0.060346861 | 0 | Neutral |
| testneg_9673 | 0 | 0 | 0.788897 | -3 | 4.932005 | -3 | 0.80849 | 0.21975 | 0.8433 | 0.13128 | 10.45898 | 0.178165188 | 0.2 | Neutral |
| testneg_9674 | -0.14 | 1.85 | 0.310587 | 0.310587 | 4.556229 | -3 | 0.2132 | 0.12788 | 0.14735 | 0.12788 | 11.25 | 0.26 | 0.2 | Neutral |
| testneg_9675 | -0.07 | -0.38 | -0.094721 | -3 | 0.968512 | -3 | 0.16778 | 0.82322 | 0.40074 | 0.36296 | 16.9194 | 0.237792869 | 0.2 | Neutral |
| testneg_9676 | 0.18 | -1.05 | 6.835273 | -3 | 1.167256 | -3 | 0.08412 | 0.28307 | 0.13601 | 0.19952 | 12.4505 | 0.584619048 | 82.6 | Disease_causing |
| testneg_9677 | 0.02 | 0.45 | 1.791247 | -3 | 2.395534 | -3 | 0.39193 | 0.11262 | 0.13967 | 0.15696 | 21.58145 | 0.130896916 | 0 | Neutral |
| testneg_9678 | -0.06 | -0.18 | -1.158658 | -3 | -1.557278 | -3 | 0.13696 | 0.09566 | 0.27077 | 0.10337 | 18.03808 | 0.01 | 0 | Neutral |
| testneg_9679 | 0.05 | -0.81 | 2.346503 | -3 | 3.376108 | -3 | 0.09618 | 0.10294 | 0.16474 | 0.41578 | 21.60066 | 0.304041999 | 2.4 | Neutral |
| testneg_9680 | -0.02 | -0.45 | -1.184577 | -3 | 3.199571 | 0.882369 | 0.34826 | 0.49454 | 0.30474 | 0.12788 | 17.21847 | 0.1305625 | 0 | Neutral |
| testneg_9681 | -0.1 | -1.9 | -1.487847 | -3 | 6.332228 | -3 | 0.06728 | 0.20664 | 0.18964 | 0.34036 | 24.93958 | 0.68 | 98.6 | Disease_causing |
| testneg_9682 | 0 | 0 | 0.655162 | -3 | 4.350853 | -3 | 0.26756 | 0.15588 | 0.12161 | 0.24178 | 14.14796 | 0.350499084 | 9.2 | Neutral |
| testneg_9683 | -0.0025714 | -0.1897143 | 2.053049 | -3 | 2.374058 | -3 | 0.26586 | 0.10825 | 0.70872 | 0.09884 | 15.3478 | 0.02153402 | 0 | Neutral |
| testneg_9684 | 0 | -0.32 | 0.23782 | -3 | 5.902081 | -3 | 0.3425 | 0.11597 | 0.09067 | 0.10671 | 16.54167 | 0.097488739 | 0 | Neutral |
| testneg_9685 | 0.02181818 | 0.11090909 | 9.146604 | -3 | 2.598788 | -3 | 0.44923 | 0.12788 | 0.19898 | 0.10307 | 30.81978 | 0.479514411 | 47 | Neutral |
| testneg_9686 | 0.18 | -1.05 | 3.730021 | -3 | 5.456507 | -3 | 0.39087 | 0.10381 | 0.19898 | 0.12788 | 9.11877 | 0.291254545 | 3.8 | Neutral |
| testneg_9687 | 0.02 | -0.63 | 4.65436 | -3 | -2.160669 | -3 | 0.10472 | 0.10014 | 0.12051 | 0.08503 | 20.41417 | 0.040047619 | 0 | Neutral |
| testneg_9688 | 0 | -0.01 | 0.389904 | -3 | -2.069741 | -3 | 0.95998 | 0.11262 | 0.16259 | 0.102 | 19.09347 | 0.010188248 | 0 | Neutral |
| testneg_9689 | 0.15 | -1.06 | 1.112437 | -3 | -2.237532 | -3 | 0.62172 | 0.18135 | 0.42836 | 0.12788 | 11.875 | 0.062761905 | 0 | Neutral |
| testneg_9690 | -0.02 | -0.45 | 4.684618 | -3 | 5.030268 | -3 | 0.51577 | 0.19561 | 0.39932 | 0.10693 | 9.50166 | 0.399424451 | 20.6 | Neutral |
| testneg_9691 | 0 | 0 | 0.807195 | -3 | 3.928636 | -3 | 0.6513 | 0.15254 | 0.05851 | 0.0724 | 12.05846 | 0.087045043 | 0 | Neutral |
| testneg_9692 | 0.16 | -1.37 | -0.103493 | -3 | -0.384446 | -3 | 0.12835 | 0.15143 | 0.36829 | 0.11457 | 18.33333 | 0.05 | 0 | Neutral |
| testneg_9693 | 0 | 0 | 0.774504 | -3 | 1.621602 | -3 | 0.99338 | 0.14229 | 0.0676 | 0.12788 | 10.81604 | 0.03 | 0 | Neutral |
| testneg_9694 | -0.04 | 0.23 | 2.526107 | -3 | 1.865118 | -3 | 0.64721 | 0.8758 | 0.16616 | 0.11073 | 18.88889 | 0.100211676 | 0 | Neutral |
| testneg_9695 | 0 | 0 | 1.54969 | -3 | -0.617826 | -3 | 0.46723 | 0.08832 | 0.06395 | 0.0759 | 16.82252 | 0.01 | 0 | Neutral |
| testneg_9696 | -0.14 | 0.37 | 0.83713 | -3 | 6.151216 | -3 | 0.06027 | 0.11574 | 0.34324 | 0.085 | 10.22898 | 0.26 | 0.2 | Neutral |
| testneg_9697 | 0 | -0.32 | 0.23782 | -3 | 4.801044 | -3 | 0.3425 | 0.11597 | 0.19355 | 0.18675 | 20.27778 | 0.27192102 | 1.4 | Neutral |
| testneg_9698 | 0 | 0.32 | -0.741073 | -3 | 2.69034 | -3 | 0.20191 | 0.13449 | 0.07225 | 0.12788 | 10 | 0.04046131 | 0 | Neutral |
| testneg_9699 | 0.03 | -1.03 | 4.438967 | -3 | 5.195379 | 4.200111 | 0.13565 | 0.46572 | 0.10168 | 0.12788 | 17.77778 | 0.6 | 86 | Disease_causing |
| testneg_9700 | -0.03 | -0.1 | 0.545839 | -3 | -0.061002 | -0.061002 | 0.58545 | 0.14414 | 0.35561 | 0.08845 | 37.83824 | 0.03 | 0 | Neutral |
| testneg_9701 | 0.07 | 0.7 | 4.02114 | -3 | 0.557877 | -3 | 0.1481 | 0.1117 | 0.25209 | 0.10029 | 13.02635 | 0.030337535 | 0 | Neutral |
| testneg_9702 | 0.01 | 0.8 | 7.50868 | -3 | 0.576122 | -3 | 0.40283 | 0.34309 | 0.18341 | 0.13901 | 21.38889 | 0.527573529 | 67.8 | Disease_causing |
| testneg_9703 | 0.21 | 0.15 | -0.402147 | -3 | 0.80883 | -3 | 0.0606 | 0.39625 | 0.19898 | 0.12788 | 9.848 | 0.1 | 0 | Neutral |
| testneg_9704 | 0 | -0.01 | 0.655486 | -3 | 2.820155 | -3 | 0.20772 | 0.16918 | 0.06254 | 0.23561 | 20.35044 | 0.203512821 | 0 | Neutral |
| testneg_9705 | -0.09 | 0.25 | -1.1177 | -3 | 0.032284 | -3 | 0.29965 | 0.26486 | 0.23393 | 0.12567 | 5.76832 | 0.044374768 | 0 | Neutral |
| testneg_9706 | 0.02 | -0.36 | 1.958686 | -3 | 2.080531 | -3 | 0.15736 | 0.5002 | 0.42235 | 0.12474 | 16.8018 | 0.080473802 | 0 | Neutral |
| testneg_9707 | -0.07 | -0.38 | -1.480749 | -3 | 6.717504 | -3 | 0.19898 | 0.12788 | 0.19898 | 0.12788 | 21.341 | 0.23 | 0.4 | Neutral |
| testneg_9708 | 0 | 0 | 1.720206 | -3 | -0.39724 | -0.39724 | 0.6992 | 0.1804 | 0.09749 | 0.82924 | 32.29167 | 0.170895722 | 0 | Neutral |
| testneg_9709 | 0.08 | 0.03 | -0.129461 | -3 | -1.897549 | -3 | 0.18416 | 0.13292 | 0.11775 | 0.10812 | 11.75141 | 0.03 | 0 | Neutral |
| testneg_9710 | 0.01 | -0.57 | 4.956408 | -3 | 0.88775 | 0.88775 | 0.28341 | 0.1005 | 0.32649 | 0.15357 | 20.11299 | 0.195565476 | 0 | Neutral |
| testneg_9711 | 0 | 0 | 1.901205 | -3 | 1.547478 | -3 | 0.20914 | 0.10089 | 0.25884 | 0.12788 | 13.51351 | 0 | 0 | Neutral |
| testneg_9712 | 0.21 | 0.15 | 1.098308 | -3 | -0.172833 | -3 | 0.09346 | 0.12788 | 0.09515 | 0.07784 | 33.56358 | 0.03 | 0 | Neutral |
| testneg_9713 | 0.22 | -2.66 | 5.243219 | -3 | 4.022598 | -3 | 0.15874 | 0.13137 | 0.12253 | 0.10814 | 6.32123 | 0.352527417 | 9.4 | Neutral |
| testneg_9714 | 0 | 0 | 0.507327 | -3 | -0.445162 | -1.511687 | 0.11172 | 0.10034 | 0.13173 | 0.12788 | 27.47685 | 0.00011838 | 0 | Neutral |
| testneg_9715 | 0.19 | -0.27 | 4.660061 | -3 | 1.795418 | -3 | 0.1897 | 0.12459 | 0.11055 | 0.19829 | 11.37731 | 0.268440028 | 1 | Neutral |
| testneg_9716 | 0.08 | 0.03 | 0.559785 | -3 | 0.632434 | -3 | 0.63431 | 0.09837 | 0.17451 | 0.11227 | 15.06206 | 0.03 | 0 | Neutral |
| testneg_9717 | 0 | 0 | 0.457027 | -3 | 3.301264 | -3 | 0.26434 | 0.12788 | 0.30951 | 0.12788 | 20.60663 | 0.025 | 0 | Neutral |
| testneg_9718 | 0.01 | -0.57 | 4.060516 | 4.060516 | 1.480614 | -3 | 0.92659 | 0.13588 | 0.11849 | 0.20101 | 16.84579 | 0.663666667 | 95.2 | Disease_causing |
| testneg_9719 | -0.16 | -0.08 | 5.861007 | -3 | 4.187482 | -3 | 0.23506 | 0.24555 | 0.08857 | 0.83499 | 17.21847 | 0.796778361 | 100 | Disease_causing |
| testneg_9720 | 0 | 0 | 0.193192 | -3 | 3.749392 | -3 | 0.1215 | 0.15825 | 0.10317 | 0.14954 | 19.44444 | 0.399545313 | 23.2 | Neutral |
| testneg_9721 | -0.05 | -0.29 | 5.247936 | -3 | 2.449752 | 2.449752 | 0.0832 | 0.09323 | 0.06736 | 0.06344 | 18.6768 | 0.21 | 0 | Neutral |
| testneg_9722 | 0 | 0 | 2.364247 | -3 | 2.081252 | 2.081252 | 0.09532 | 0.35459 | 0.27783 | 0.62384 | 18.12633 | 0.31975 | 1.6 | Neutral |
| testneg_9723 | 0.14 | -1.01 | 3.224374 | -3 | 5.481679 | -3 | 0.44329 | 0.12788 | 0.48627 | 0.09509 | 32.27587 | 0.190977884 | 0.2 | Neutral |
| testneg_9724 | 0 | 0.73 | 3.376127 | -3 | 4.518999 | -3 | 0.43918 | 0.46357 | 0.90955 | 0.10096 | 18.33333 | 0.368741965 | 7.2 | Neutral |
| testneg_9725 | 0 | 0 | 0.237368 | -3 | 2.058438 | -3 | 0.1602 | 0.11262 | 0.0749 | 0.12788 | 24.37034 | 0.01 | 0 | Neutral |
| testneg_9726 | -0.21 | -0.15 | -0.83994 | -3 | 1.461115 | -3 | 0.95319 | 0.20297 | 0.22953 | 0.0919 | 10.24716 | 0.08 | 0 | Neutral |
| testneg_9727 | -0.05 | -1.52 | -1.197202 | -1.197202 | 3.107095 | -3 | 0.08688 | 0.08809 | 0.19898 | 0.40608 | 16.89285 | 0.21 | 0 | Neutral |
| testneg_9728 | -0.16 | -0.08 | 2.94383 | 2.94383 | -0.361103 | -3 | 0.51707 | 0.45685 | 0.26982 | 0.12788 | 13.97697 | 0.42 | 22 | Neutral |
| testneg_9729 | 0.05 | -0.81 | 3.173176 | -3 | 1.997964 | -3 | 0.17226 | 0.13001 | 0.0959 | 0.12788 | 26.93217 | 0.05 | 0 | Neutral |
| testneg_9730 | 0.04 | -0.23 | 3.962157 | -3 | 2.892408 | -3 | 0.06898 | 0.11659 | 0.2288 | 0.15849 | 28.92245 | 0.316042365 | 5.4 | Neutral |
| testneg_9731 | -0.05 | -0.29 | 0.79176 | -3 | 0.253071 | -3 | 0.2141 | 0.19929 | 0.0418 | 0.12788 | 16.8018 | 0.020382928 | 0 | Neutral |
| testneg_9732 | -0.08 | -0.03 | -0.514498 | -3 | 4.382959 | -3 | 0.81305 | 0.17045 | 0.86513 | 0.31355 | 16.48772 | 0.353356613 | 6 | Neutral |
| testneg_9733 | -0.0043357 | 0.02363636 | -0.760295 | -3 | -0.674759 | -0.674759 | 0.33082 | 0.12788 | 0.02782 | 0.06218 | 19.09347 | 0.02 | 0 | Neutral |
| testneg_9734 | 0 | 0 | 0.556564 | -3 | -0.50288 | -3 | 0.18834 | 0.12788 | 0.1294 | 0.10964 | 9.34598 | 0 | 0 | Neutral |
| testneg_9735 | 0 | 0 | 0.079919 | -0.590049 | -0.100872 | -3 | 0.50063 | 0.154 | 0.11891 | 0.11186 | 17.34285 | 0.02 | 0 | Neutral |
| testneg_9736 | -0.02 | 0.63 | 0.11685 | -3 | 5.969277 | -3 | 0.21989 | 0.10981 | 0.31067 | 0.35414 | 13.54885 | 0.339345694 | 8.2 | Neutral |
| testneg_9737 | 0 | 0 | 0.221415 | -3 | 0.034544 | -3 | 0.89972 | 0.10766 | 0.13023 | 0.29692 | 11.14078 | 0.050357418 | 0 | Neutral |
| testneg_9738 | 0.07 | 0.33 | 3.241723 | -3 | 1.01004 | -3 | 0.61343 | 0.11838 | 0.19898 | 0.09533 | 27.07912 | 0.03 | 0 | Neutral |
| testneg_9739 | 0.16 | -0.37 | 5.42275 | -3 | 5.328896 | -3 | 0.44816 | 0.56655 | 0.08529 | 0.12376 | 13.15432 | 0.631458333 | 94.4 | Disease_causing |
| testneg_9740 | 0 | 0 | 0.556564 | -3 | 0.93478 | -3 | 0.18834 | 0.12788 | 0.57239 | 0.16622 | 5.22523 | 0.020229915 | 0 | Neutral |
| testneg_9741 | 0 | 0 | 1.626934 | -3 | 6.678689 | -3 | 0.20942 | 0.14229 | 0.20765 | 0.27124 | 12.72648 | 0.443130914 | 32.4 | Neutral |
| testneg_9742 | 0.03 | 0.12 | 1.249184 | -3 | -1.375282 | -3 | 0.40592 | 0.22108 | 0.19898 | 0.12788 | 4.72222 | 0.021310069 | 0 | Neutral |
| testneg_9743 | 0.02 | 0.41 | -1.253946 | -3 | 0.790139 | -3 | 0.36566 | 0.15312 | 0.26482 | 0.12788 | 4.98847 | 0.020115492 | 0 | Neutral |
| testneg_9744 | 0 | 0 | 0.162648 | -3 | 1.472855 | -3 | 0.38002 | 0.22508 | 0.56554 | 0.76978 | 10.41667 | 0.095526316 | 0 | Neutral |
| testneg_9745 | 0 | 0 | 0.028094 | -3 | 0.070966 | -3 | 0.46712 | 0.93559 | 0.53424 | 0.08484 | 10.41667 | 0.070202206 | 0 | Neutral |
| testneg_9746 | 0 | 0 | 1.454524 | -3 | 13.536888 | -3 | 0.09577 | 0.2436 | 0.19898 | 0.12788 | 27.95796 | 0.505833333 | 57.8 | Disease_causing |
| testneg_9747 | 0.03 | -1.03 | 4.859478 | -3 | -2.337287 | -3 | 0.11501 | 0.13748 | 0.04946 | 0.12788 | 29.87756 | 0.09 | 0 | Neutral |
| testneg_9748 | 0 | 0 | 0.27864 | -3 | 0.821439 | -3 | 0.17737 | 0.54971 | 0.16424 | 0.11262 | 10.22898 | 0.021339286 | 0 | Neutral |
| testneg_9749 | 0.18 | -1.05 | 3.992521 | -3 | -1.989097 | -3 | 0.19898 | 0.12788 | 0.11094 | 0.12788 | 27.51545 | 0.05 | 0 | Neutral |
| testneg_9750 | 0 | -0.32 | 1.824862 | -3 | 5.495658 | -3 | 0.44816 | 0.56655 | 0.15294 | 0.12788 | 12.29167 | 0.24740942 | 0.6 | Neutral |
| testneg_9751 | 0 | 0 | 1.56019 | -3 | -2.12413 | -3 | 0.25487 | 0.13098 | 0.42836 | 0.12788 | 19.44444 | 0.01 | 0 | Neutral |
| testneg_9752 | 0 | 0 | 0.829195 | -3 | 2.292445 | -3 | 0.68423 | 0.08163 | 0.10362 | 0.09649 | 17.21847 | 0.03 | 0 | Neutral |
| testneg_9753 | 0 | 0 | 0.254114 | -3 | 2.70354 | -3 | 0.19002 | 0.12138 | 0.46735 | 0.10674 | 34.84275 | 0.012591099 | 0 | Neutral |
| testneg_9754 | -0.01 | 1.44 | -1.970817 | -3 | -0.81151 | -3 | 0.2058 | 0.28174 | 0.01635 | 0.12788 | 10.36687 | 0.12 | 0 | Neutral |
| testneg_9755 | 0.07 | 0.38 | 2.246546 | -3 | 2.246903 | -3 | 0.32022 | 0.09529 | 0.1019 | 0.17526 | 11.25 | 0.130831498 | 0 | Neutral |
| testneg_9756 | 0 | 0 | 1.774202 | -3 | 1.705087 | -3 | 0.29356 | 0.47581 | 0.22225 | 0.11186 | 18.88889 | 0.021591781 | 0 | Neutral |
| testneg_9757 | 0.03 | 0.12 | 5.153522 | -3 | 2.12021 | 2.12021 | 0.1205 | 0.23508 | 0.07698 | 0.12788 | 13.20837 | 0.255767899 | 0.2 | Neutral |
| testneg_9758 | 0 | 0 | 0.039637 | -3 | 0.104989 | -3 | 0.20304 | 0.31462 | 0.35026 | 0.09817 | 10.83333 | 0.010944129 | 0 | Neutral |
| testneg_9759 | -0.03 | -0.1 | 5.429346 | -3 | 0.641608 | -3 | 0.95998 | 0.11262 | 0.32458 | 0.1016 | 14.28596 | 0.17 | 0 | Neutral |

| | | | | | | | | | | | | | | |
|---|---|---|---|---|---|---|---|---|---|---|---|---|---|---|
| testneg_9760 | 0 | 0 | -0.633568 | -3 | 1.112726 | -1.033749 | 0.23027 | 0.37171 | 0.09964 | 0.10384 | 10.22898 | 0.03 | 0 | Neutral |
| testneg_9761 | 0 | 0 | 0.191318 | -3 | -1.599196 | -3 | 0.96113 | 0.24974 | 0.37711 | 0.12788 | 19.09347 | 0.02 | 0 | Neutral |
| testneg_9762 | 0.01 | 0.8 | -0.725111 | -0.725111 | 1.554221 | -3 | 0.05116 | 0.08996 | 0.15341 | 0.12272 | 16.8018 | 0.05 | 0 | Neutral |
| testneg_9763 | 0.09 | -0.3 | 0.037077 | -3 | 0.775551 | -3 | 0.09004 | 0.07829 | 0.0903 | 0.09372 | 25.87252 | 0.02 | 0 | Neutral |
| testneg_9764 | 0.14 | -0.37 | 2.102067 | -3 | 3.579296 | 0.549099 | 0.95557 | 0.38833 | 0.98641 | 0.83983 | 5.69444 | 0.479705285 | 45.6 | Neutral |
| testneg_9765 | 0.02 | -0.36 | 3.235017 | -3 | -0.61032 | -3 | 0.30329 | 0.14582 | 0.09522 | 0.20323 | 17.43148 | 0.199591667 | 0 | Neutral |
| testneg_9766 | -0.08 | -0.03 | 6.695303 | 6.695303 | 2.732818 | -3 | 0.09871 | 0.09286 | 0.16204 | 0.12788 | 18.6768 | 0.65 | 98 | Disease_causing |
| testneg_9767 | 0.07 | 0.7 | 3.889256 | -3 | 0.232205 | -3 | 0.59345 | 0.26484 | 0.09473 | 0.08327 | 30.6092 | 0.082014286 | 0 | Neutral |
| testneg_9768 | 0.18 | 0.16 | -0.13961 | -3 | 5.604435 | -3 | 0.65734 | 0.1013 | 0.10261 | 0.0933 | 29.64606 | 0.151577847 | 0 | Neutral |
| testneg_9769 | 0 | 0 | 1.108803 | -3 | 0.099948 | -3 | 0.85092 | 0.58871 | 0.19898 | 0.56316 | 12.29167 | 0.101992063 | 0 | Neutral |
| testneg_9770 | 0.03 | -1.03 | 5.824661 | -3 | 2.944773 | -3 | 0.13458 | 0.17634 | 0.42666 | 0.12788 | 18.95155 | 0.41 | 22.4 | Neutral |
| testneg_9771 | 0 | -0.01 | 6.431621 | 6.431621 | 1.15899 | -3 | 0.26676 | 0.76568 | 0.17969 | 0.11473 | 22.0615 | 0.66 | 96.4 | Disease_causing |
| testneg_9772 | 0.09 | -0.67 | 2.208846 | -3 | -1.139419 | -3 | 0.1496 | 0.16692 | 0.36852 | 0.0903 | 50.51283 | 0.06 | 0 | Neutral |
| testneg_9773 | 0.07 | 0.7 | 4.781408 | -3 | 2.950487 | -3 | 0.19898 | 0.12788 | 0.14495 | 0.37127 | 35.22867 | 0.461852998 | 40.6 | Neutral |
| testneg_9774 | 0 | 0 | 1.594568 | -3 | 7.077697 | -3 | 0.31318 | 0.11262 | 0.36324 | 0.15729 | 18.6768 | 0.353262821 | 11 | Neutral |
| testneg_9775 | 0.01 | 0.79 | 2.037729 | -3 | 4.13176 | -3 | 0.58984 | 0.15588 | 0.226 | 0.28784 | 11.59649 | 0.426137105 | 24.8 | Neutral |
| testneg_9776 | 0.21 | 0.15 | 1.775814 | -3 | -0.195051 | -3 | 0.18499 | 0.10583 | 0.36417 | 0.21506 | 12.57614 | 0.075135135 | 0 | Neutral |
| testneg_9777 | 0.14 | -1.01 | 7.093027 | -3 | 1.460979 | -3 | 0.41953 | 0.44852 | 0.10385 | 0.09775 | 29.86111 | 0.449375 | 32.6 | Neutral |
| testneg_9778 | 0.21 | 0.15 | 0.833236 | -3 | 4.566551 | -3 | 0.4172 | 0.11643 | 0.28227 | 0.24183 | 15.6487 | 0.300416606 | 1.8 | Neutral |
| testneg_9779 | 0.03 | -0.85 | 0.02572 | -3 | 0.400695 | -3 | 0.0138 | 0.06769 | 0.02146 | 0.10351 | 10.83333 | 0.04 | 0 | Neutral |
| testneg_9780 | 0 | 0 | -0.090737 | -3 | -0.168532 | -3 | 0.09734 | 0.08962 | 0.16388 | 0.14439 | 21.75792 | 0.05 | 0 | Neutral |
| testneg_9781 | 0 | 0 | -0.071864 | -0.071864 | 1.925327 | -3 | 0.97005 | 0.31407 | 0.23385 | 0.12788 | 5.69444 | 0.01 | 0 | Neutral |
| testneg_9782 | -0.15 | -0.06 | -1.712174 | -3 | 0.878368 | -3 | 0.07604 | 0.35375 | 0.31648 | 0.12788 | 15.37063 | 0.13 | 0 | Neutral |
| testneg_9783 | 0 | -0.01 | 0.684698 | -3 | 6.645026 | -3 | 0.09661 | 0.09981 | 0.59245 | 0.12788 | 13.51351 | 0.16 | 0 | Neutral |
| testneg_9784 | -0.08 | -0.03 | 3.763357 | -3 | 1.456728 | -3 | 0.08631 | 0.10194 | 0.04063 | 0.05848 | 34.13176 | 0.05 | 0 | Neutral |
| testneg_9785 | 0.12 | -1.14 | -0.706205 | -3 | 0.626243 | 0.626243 | 0.37074 | 0.10627 | 0.34412 | 0.08672 | 7.01832 | 0.03 | 0 | Neutral |
| testneg_9786 | 0.14 | -1.01 | 0.599729 | -3 | 2.257953 | -3 | 0.38762 | 0.12788 | 0.18697 | 0.12232 | 33.95189 | 0.03 | 0 | Neutral |
| testneg_9787 | -0.2 | 1.26 | -2.106364 | -3 | 1.721437 | -3 | 0.10383 | 0.08853 | 0.13873 | 0.12788 | 22.38265 | 0.14 | 0 | Neutral |
| testneg_9788 | -0.02 | 0.68 | 1.913159 | -3 | -0.659954 | -3 | 0.91921 | 0.32331 | 0.19898 | 0.12788 | 19.09347 | 0.04256435 | 0 | Neutral |
| testneg_9789 | 0.15 | -0.28 | -1.251158 | -3 | 0.781337 | -3 | 0.20127 | 0.13478 | 0.67899 | 0.16647 | 12.14396 | 0.063083333 | 0 | Neutral |
| testneg_9790 | -0.14 | 0.37 | 0.175857 | -3 | 0.567562 | -3 | 0.30042 | 0.13517 | 0.17046 | 0.21736 | 21.38889 | 0.122505097 | 0 | Neutral |
| testneg_9791 | 0.14 | -1.01 | -0.176493 | -3 | 5.134112 | -3 | 0.95978 | 0.09493 | 0.50448 | 0.50327 | 9.27711 | 0.303123217 | 1.2 | Neutral |
| testneg_9792 | 0.21 | 0.15 | 5.407311 | -3 | 2.696752 | -3 | 0.08975 | 0.07399 | 0.51319 | 0.12971 | 20.2851 | 0.181041667 | 0 | Neutral |
| testneg_9793 | 0.04 | 2.01 | 5.876399 | -3 | 0.972685 | -3 | 0.7823 | 0.5949 | 0.25697 | 0.13883 | 20.208 | 0.461385507 | 38 | Neutral |
| testneg_9794 | 0 | 0 | 0.974808 | 0.974808 | 3.997628 | -3 | 0.32992 | 0.33227 | 0.66799 | 0.12788 | 10.83333 | 0.171324813 | 0 | Neutral |
| testneg_9795 | -0.15 | -0.04 | 3.97614 | -3 | 0.199587 | -3 | 0.33135 | 0.73068 | 0.26524 | 0.11154 | 12.90858 | 0.16 | 0 | Neutral |
| testneg_9796 | -0.05 | -0.29 | 1.012343 | -3 | 1.07442 | -3 | 0.93035 | 0.31722 | 0.25422 | 0.10102 | 10.83333 | 0.0225 | 0 | Neutral |
| testneg_9797 | -0.07 | -0.33 | 2.384666 | -3 | 2.60676 | -3 | 0.17006 | 0.14718 | 0.09439 | 0.07415 | 26.84685 | 0.05 | 0 | Neutral |
| testneg_9798 | 0.04 | -0.23 | 3.28103 | -3 | 2.665942 | -3 | 0.27077 | 0.10337 | 0.2755 | 0.12788 | 18.76882 | 0.061259615 | 0 | Neutral |
| testneg_9799 | 0.03 | -1.03 | 5.016685 | -3 | 5.542651 | -3 | 0.49629 | 0.11989 | 0.13708 | 0.81545 | 28.32663 | 0.645817241 | 92.6 | Disease_causing |
| testneg_9800 | -0.03 | -0.1 | 2.059294 | -3 | -0.274976 | -3 | 0.66711 | 0.31094 | 0.26767 | 0.25506 | 20.85185 | 0.171261364 | 0 | Neutral |
| testneg_9801 | 0 | 0 | -0.43021 | -3 | -1.214088 | -3 | 0.17041 | 0.12378 | 0.04192 | 0.06306 | 20.5518 | 0.01 | 0 | Neutral |
| testneg_9802 | -0.02 | 0.68 | 1.913159 | -3 | 0.093296 | 0.093296 | 0.91921 | 0.32331 | 0.19241 | 0.15665 | 11.25 | 0.142549031 | 0 | Neutral |
| testneg_9803 | -0.14 | 1.01 | 5.832346 | -3 | 2.493168 | 2.493168 | 0.43736 | 0.10017 | 0.14897 | 0.19122 | 5.69444 | 0.618907345 | 91 | Disease_causing |
| testneg_9804 | 0.21 | 0.15 | 8.082349 | -3 | 4.959501 | -3 | 0.9989 | 0.8746 | 0.14301 | 0.12788 | 17.21847 | 0.78087538 | 100 | Disease_causing |
| testneg_9805 | 0.03 | 1.22 | 2.934646 | -3 | -0.077697 | -3 | 0.26972 | 0.12484 | 0.5043 | 0.3225 | 19.14347 | 0.18 | 0 | Neutral |
| testneg_9806 | 0.18 | -1.05 | 4.646906 | -3 | 0.620269 | -3 | 0.09508 | 0.09834 | 0.1287 | 0.10423 | 19.15873 | 0.05 | 0 | Neutral |
| testneg_9807 | 0 | 0 | 1.846412 | -3 | -0.339896 | -3 | 0.35421 | 0.10694 | 0.10385 | 0.09775 | 18.15261 | 0.000897895 | 0 | Neutral |
| testneg_9808 | 0 | 0 | 0.430658 | -3 | -2.242853 | -3 | 0.0752 | 0.74467 | 0.06358 | 0.5364 | 18.05349 | 0.23804539 | 0 | Neutral |
| testneg_9809 | -0.03 | -0.1 | -0.877392 | -3 | 0.482446 | -3 | 0.1311 | 0.12667 | 0.4118 | 0.1154 | 18.36272 | 0.01 | 0 | Neutral |
| testneg_9810 | 0 | 0 | 0.470493 | -3 | 0.297516 | 0.297516 | 0.42802 | 0.12096 | 0.19294 | 0.10464 | 14.44649 | 0.01 | 0 | Neutral |
| testneg_9811 | 0.09 | -0.3 | 1.845442 | -3 | -0.136533 | -3 | 0.99293 | 0.79067 | 0.05932 | 0.09551 | 9.95504 | 0.09 | 0 | Neutral |
| testneg_9812 | 0.22 | -2.66 | 7.446509 | 3.054372 | 1.562114 | -3 | 0.10869 | 0.1503 | 0.78356 | 0.13377 | 21.24367 | 0.659931507 | 95.8 | Disease_causing |
| testneg_9813 | -0.0018939 | 0.10159091 | 2.238223 | -3 | 5.961755 | -3 | 0.32606 | 0.10251 | 0.27783 | 0.62384 | 10.83333 | 0.441158493 | 33.2 | Neutral |
| testneg_9814 | 0.03 | 1.22 | 2.118816 | -3 | -1.304747 | -3 | 0.6781 | 0.1291 | 0.10383 | 0.08853 | 22.49681 | 0.06 | 0 | Neutral |
| testneg_9815 | 0 | -0.01 | 1.609618 | -3 | 0.157379 | -3 | 0.67038 | 0.1794 | 0.45997 | 0.15162 | 12.72648 | 0.040096703 | 0 | Neutral |
| testneg_9816 | -0.01 | -0.8 | -0.697584 | -3 | 2.681065 | -1.511687 | 0.19898 | 0.12788 | 0.13173 | 0.12788 | 31.2634 | 0.03 | 0 | Neutral |
| testneg_9817 | 0 | 0.32 | 1.852415 | -3 | 3.865715 | -3 | 0.90767 | 0.12788 | 0.17639 | 0.12919 | 10.83333 | 0.098437229 | 0 | Neutral |
| testneg_9818 | 0.16 | 0.08 | 3.022813 | -3 | 2.511594 | -3 | 0.17301 | 0.10048 | 0.15432 | 0.095 | 14.16667 | 0.070979437 | 0 | Neutral |
| testneg_9819 | 0 | 0 | 0.416565 | -3 | 2.882534 | -3 | 0.05535 | 0.12788 | 0.23385 | 0.12788 | 19.44444 | 0.051024306 | 0 | Neutral |
| testneg_9820 | -0.02 | 0.28 | 1.849543 | -3 | 1.714629 | -3 | 0.19898 | 0.1054 | 0.03235 | 0.0948 | 10 | 0.02 | 0 | Neutral |
| testneg_9821 | 0 | 0 | -0.090459 | -0.090459 | -0.878612 | -3 | 0.15581 | 0.19351 | 0.11802 | 0.09304 | 7.58876 | 0.013333333 | 0 | Neutral |
| testneg_9822 | 0.03 | 0.1 | 6.254981 | -3 | 3.684371 | 3.684371 | 0.95993 | 0.15588 | 0.10751 | 0.08512 | 22.62411 | 0.498675214 | 53.8 | Disease_causing |
| testneg_9823 | 0.04 | -0.23 | 7.16104 | -3 | 2.851691 | -3 | 0.25961 | 0.07098 | 0.21623 | 0.42074 | 26.13392 | 0.531018669 | 66.2 | Disease_causing |
| testneg_9824 | 0 | 0 | 1.436812 | -3 | -0.553129 | -3 | 0.29302 | 0.17394 | 0.1601 | 0.12546 | 19.68216 | 0.01 | 0 | Neutral |
| testneg_9825 | -0.21 | -0.15 | -0.83994 | -3 | 7.720593 | -3 | 0.95319 | 0.20297 | 0.08386 | 0.12788 | 10.20833 | 0.442678571 | 33.6 | Neutral |
| testneg_9826 | 0.07 | 0.38 | -2.188525 | -2.188525 | 6.070697 | -3 | 0.07282 | 0.12788 | 0.08027 | 0.12788 | 19.51724 | 0.16875 | 0 | Neutral |
| testneg_9827 | -0.07 | -0.38 | 4.209007 | -3 | 1.237552 | -3 | 0.10724 | 0.49624 | 0.21399 | 0.10708 | 16.3108 | 0.15 | 0 | Neutral |
| testneg_9828 | 0 | -0.32 | 5.684554 | -3 | 2.648595 | -3 | 0.9449 | 0.9378 | 0.27843 | 0.54325 | 10 | 0.727034848 | 99.2 | Disease_causing |
| testneg_9829 | -0.04 | -2.01 | 4.558544 | -3 | 1.973746 | -3 | 0.08753 | 0.09666 | 0.0594 | 0.16845 | 20.59766 | 0.33 | 3.4 | Neutral |
| testneg_9830 | 0 | 0 | 1.649381 | -3 | 0.415464 | -3 | 0.22686 | 0.24974 | 0.18671 | 0.12788 | 19.44444 | 0.010886133 | 0 | Neutral |
| testneg_9831 | -0.07 | -0.38 | 2.097779 | -3 | 0.617218 | -3 | 0.81054 | 0.28104 | 0.10458 | 0.10499 | 21.43606 | 0.04 | 0 | Neutral |
| testneg_9832 | 0 | 0 | -0.22016 | -0.22016 | -0.062057 | -3 | 0.17911 | 0.58893 | 0.26489 | 0.10549 | 57.39678 | 0.13 | 0 | Neutral |
| testneg_9833 | 0.08 | -1.2 | 3.426369 | -3 | 8.162991 | 6.797249 | 0.42585 | 0.18081 | 0.13985 | 0.08349 | 11.92982 | 0.705333333 | 99.8 | Disease_causing |
| testneg_9834 | -0.18 | 1.05 | -0.907393 | -3 | 0.543778 | 0.543778 | 0.05332 | 0.12788 | 0.09749 | 0.82924 | 29.9024 | 0.27 | 0.4 | Neutral |
| testneg_9835 | 0.02 | 0.45 | 3.429131 | -3 | 1.912827 | -3 | 0.03499 | 0.12971 | 0.1203 | 0.13872 | 18.88889 | 0.192916667 | 0 | Neutral |
| testneg_9836 | 0.07 | 0.01 | -0.38551 | -3 | 0.370077 | -3 | 0.16797 | 0.12788 | 0.27083 | 0.14313 | 26.34995 | 0.04422619 | 0 | Neutral |
| testneg_9837 | 0 | -0.01 | 1.149331 | -3 | 0.502731 | -3 | 0.15582 | 0.14216 | 0.75033 | 0.64204 | 10.41667 | 0.03057266 | 0 | Neutral |
| testneg_9838 | 0.14 | -1.01 | 1.010678 | -3 | 2.080809 | -3 | 0.1109 | 0.17295 | 0.06636 | 0.1079 | 24.65364 | 0.061845513 | 0 | Neutral |
| testneg_9839 | 0.21 | 0.15 | 2.671154 | -3 | -0.276963 | -3 | 0.08775 | 0.12788 | 0.6001 | 0.90845 | 10.04623 | 0.150912698 | 0 | Neutral |
| testneg_9840 | -0.07 | -0.01 | 0.439914 | -3 | 1.430219 | -3 | 0.09275 | 0.14911 | 0.21731 | 0.11795 | 10 | 0.030887489 | 0 | Neutral |
| testneg_9841 | 0.01 | -0.57 | 0.14373 | -3 | 5.33097 | -3 | 0.17728 | 0.46704 | 0.08814 | 0.42357 | 18.88889 | 0.62606685 | 93.2 | Disease_causing |
| testneg_9842 | 0 | 0 | -0.076898 | -3 | 0.294506 | -3 | 0.07973 | 0.09208 | 0.0961 | 0.09786 | 32.07516 | 0.02 | 0 | Neutral |
| testneg_9843 | 0 | 0 | -0.342253 | -0.342253 | -1.178484 | -3 | 0.63363 | 0.14795 | 0.13669 | 0.09263 | 21.00919 | 0.030425725 | 0 | Neutral |
| testneg_9844 | 0.06 | -0.57 | 0.297516 | 0.297516 | 4.689762 | -3 | 0.19294 | 0.10464 | 0.08268 | 0.09939 | 12.40248 | 0.086545181 | 0 | Neutral |
| testneg_9845 | 0.02 | 0.41 | 8.488711 | -3 | 2.652042 | -3 | 0.19102 | 0.12788 | 0.83798 | 0.14433 | 11.74429 | 0.51 | 59.2 | Disease_causing |
| testneg_9846 | 0.03 | 0.12 | 2.433198 | -3 | 0.90434 | -3 | 0.19989 | 0.06568 | 0.53424 | 0.08484 | 29.16667 | 0.01 | 0 | Neutral |
| testneg_9847 | 0 | 0 | 0.717822 | -3 | -2.556301 | -3 | 0.29044 | 0.12142 | 0.12379 | 0.12788 | 28.49438 | 0.01 | 0 | Neutral |
| testneg_9848 | 0 | 0 | 0.77406 | -3 | 1.237616 | -3 | 0.1192 | 0.1508 | 0.10395 | 0.08737 | 20.83333 | 0.03 | 0 | Neutral |
| testneg_9849 | 0.01 | 0.8 | 1.074849 | -3 | 4.187482 | -3 | 0.06775 | 0.09511 | 0.08857 | 0.83499 | 10.83333 | 0.32 | 4.2 | Neutral |
| testneg_9850 | -0.04 | 0.23 | 1.327969 | -3 | -0.410022 | -3 | 0.60812 | 0.19286 | 0.17419 | 0.33842 | 11.25 | 0.101525232 | 0 | Neutral |
| testneg_9851 | 0 | 0 | -0.384176 | -3 | 0.849086 | -3 | 0.31238 | 0.14393 | 0.30241 | 0.104 | 26.80135 | 0.010098553 | 0 | Neutral |
| testneg_9852 | 0 | 0 | 0.585709 | -3 | 1.370202 | -3 | 0.77644 | 0.09695 | 0.47335 | 0.12788 | 16.8018 | 0.00070197 | 0 | Neutral |
| testneg_9853 | 0.19 | -0.27 | 3.647477 | -3 | 0.980675 | -3 | 0.25236 | 0.11809 | 0.35137 | 0.09161 | 10.76632 | 0.03 | 0 | Neutral |
| testneg_9854 | 0.21 | 0.15 | 2.671154 | -3 | 2.577846 | -3 | 0.08775 | 0.12788 | 0.12819 | 0.1229 | 13.02635 | 0.07 | 0 | Neutral |
| testneg_9855 | 0.05 | 0.29 | 2.036657 | -3 | 0.104835 | -3 | 0.91691 | 0.25567 | 0.91208 | 0.52258 | 10.41667 | 0.11 | 0 | Neutral |
| testneg_9856 | 0.08 | 0.03 | -1.709673 | -3 | 0.604429 | -3 | 0.16708 | 0.11594 | 0.08973 | 0.07662 | 24.0282 | 0.020606618 | 0 | Neutral |
| testneg_9857 | -0.09 | 0.3 | 0.156321 | -3 | -0.725111 | -0.725111 | 0.21566 | 0.15056 | 0.05116 | 0.08996 | 17.89614 | 0.03875 | 0 | Neutral |
| testneg_9858 | 0.06 | -0.57 | 0.297516 | 0.297516 | -1.601251 | -1.601251 | 0.19294 | 0.10464 | 0.19898 | 0.20319 | 13.95284 | 0.083128863 | 0 | Neutral |
| testneg_9859 | 0.22 | -2.66 | 7.225361 | -3 | -0.190524 | -3 | 0.19898 | 0.15588 | 0.13322 | 0.11297 | 34.45035 | 0.297857143 | 4 | Neutral |
| testneg_9860 | -0.09 | 0.3 | 0.850367 | -3 | 5.617032 | -3 | 0.29024 | 0.1472 | 0.18328 | 0.12788 | 23.66368 | 0.230666667 | 0 | Neutral |
| testneg_9861 | 0 | 0 | 0.520351 | -0.0906 | 5.704027 | -3 | 0.05732 | 0.1098 | 0.09504 | 0.38939 | 61.73161 | 0.394621212 | 16.4 | Neutral |
| testneg_9862 | -0.15 | 1.06 | 2.299586 | -3 | 0.085787 | -3 | 0.19898 | 0.12788 | 0.40996 | 0.57507 | 28.90523 | 0.141657509 | 0 | Neutral |
| testneg_9863 | 0 | 0 | 2.995532 | -3 | 3.095105 | -3 | 0.96879 | 0.38765 | 0.52914 | 0.25278 | 4.86652 | 0.53009712 | 65.6 | Disease_causing |
| testneg_9864 | 0.05 | -0.81 | 0.475752 | -3 | 0.211586 | 0.211586 | 0.59871 | 0.34719 | 0.1119 | 0.24875 | 9.36773 | 0.15875 | 0 | Neutral |
| testneg_9865 | -0.1 | -1.9 | 2.550514 | -3 | 7.297508 | -3 | 0.47887 | 0.09483 | 0.15848 | 0.12754 | 19.6749 | 0.33 | 4.8 | Neutral |
| testneg_9866 | -0.18 | -0.16 | 5.75006 | -3 | 0.426905 | -3 | 0.12867 | 0.11159 | 0.19531 | 0.18996 | 20.83333 | 0.388075397 | 16.6 | Neutral |
| testneg_9867 | 0.14 | -1.01 | 2.621366 | -3 | 3.812103 | -3 | 0.81899 | 0.12788 | 0.30311 | 0.11121 | 11.21556 | 0.111822917 | 0 | Neutral |
| testneg_9868 | 0 | 0 | 0.254114 | -3 | 0.269085 | -3 | 0.19002 | 0.12138 | 0.09281 | 0.13929 | 19.43459 | 0.06 | 0 | Neutral |
| testneg_9869 | 0.21 | 0.15 | 1.523309 | -3 | 1.856425 | -3 | 0.05378 | 0.12788 | 0.09806 | 0.12788 | 27.51545 | 0.07 | 0 | Neutral |
| testneg_9870 | 0 | 0 | 0.223823 | -3 | 1.894277 | -3 | 0.32484 | 0.11621 | 0.53216 | 0.57092 | 9.81132 | 0.022928571 | 0 | Neutral |
| testneg_9871 | 0.04 | -0.23 | 1.711588 | -3 | 1.746837 | -3 | 0.44615 | 0.09968 | 0.07912 | 0.09055 | 20.86969 | 0.010741758 | 0 | Neutral |
| testneg_9872 | -0.01 | 1.44 | 0.099948 | -3 | 0.396977 | -3 | 0.19898 | 0.56316 | 0.12848 | 0.65153 | 21.22834 | 0.34 | 3 | Neutral |
| testneg_9873 | 0 | 0 | -0.117483 | -3 | 6.420723 | -3 | 0.06793 | 0.12788 | 0.42666 | 0.12788 | 12.98941 | 0.140444444 | 0 | Neutral |
| testneg_9874 | 0 | 0 | 1.107552 | -3 | 0.655162 | 0.655162 | 0.11687 | 0.09069 | 0.26756 | 0.15588 | 18.14743 | 0.043813725 | 0 | Neutral |
| testneg_9875 | 0.19 | -0.27 | 6.416509 | -3 | 0.615335 | -3 | 0.08814 | 0.42357 | 0.70872 | 0.09884 | 23.37302 | 0.286666667 | 2.2 | Neutral |
| testneg_9876 | 0 | 0 | -0.396403 | -3 | 3.103812 | -3 | 0.42585 | 0.18081 | 0.0693 | 0.09293 | 10 | 0.07 | 0 | Neutral |
| testneg_9877 | 0.05 | -0.81 | 1.160689 | -3 | -0.304634 | -3 | 0.18842 | 0.12378 | 0.25957 | 0.06969 | 11.13095 | 0.01 | 0 | Neutral |
| testneg_9878 | 0 | 0 | 1.39998 | -3 | 3.089818 | -3 | 0.161 | 0.11597 | 0.09026 | 0.09447 | 13.51351 | 0.020572632 | 0 | Neutral |
| testneg_9879 | -0.02 | 0.63 | 2.537077 | -3 | -0.260132 | -3 | 0.33256 | 0.14909 | 0.18502 | 0.21329 | 11.25 | 0.111292563 | 0 | Neutral |
| testneg_9880 | 0 | 0 | -0.258437 | -3 | 1.089314 | -3 | 0.07044 | 0.11679 | 0.3783 | 0.08224 | 11.25 | 0.000544848 | 0 | Neutral |
| testneg_9881 | -0.05 | 0.81 | -1.267005 | -3 | 1.826257 | -3 | 0.16469 | 0.16042 | 0.29356 | 0.46298 | 7.23475 | 0.137725379 | 0 | Neutral |

| | | | | | | | | | | | | | | |
|---|---|---|---|---|---|---|---|---|---|---|---|---|---|---|
| testneg_9882 | 0.02 | 0.45 | -0.784242 | -3 | 2.050194 | -3 | 0.18153 | 0.10845 | 0.1897 | 0.12459 | 16.25133 | 0.01 | 0 | Neutral |
| testneg_9883 | -0.03 | -0.1 | 1.673524 | -3 | 0.259735 | -3 | 0.28156 | 0.11579 | 0.16468 | 0.11275 | 15.57171 | 0.000801282 | 0 | Neutral |
| testneg_9884 | -0.03 | -0.1 | 2.625574 | -3 | 1.053751 | -3 | 0.284 | 0.15339 | 0.0917 | 0.13124 | 19.11747 | 0.079354167 | 0 | Neutral |
| testneg_9885 | 0.04 | -0.23 | 0.145821 | -3 | 3.596868 | -3 | 0.21265 | 0.1284 | 0.19898 | 0.12788 | 7.14037 | 0.040824867 | 0 | Neutral |
| testneg_9886 | -0.03 | 1.03 | 2.996727 | -3 | 6.182448 | -3 | 0.19898 | 0.17518 | 0.1977 | 0.09728 | 27.77955 | 0.37 | 11.8 | Neutral |
| testneg_9887 | 0 | -0.01 | 0.834318 | -3 | 5.809418 | -3 | 0.31726 | 0.17249 | 0.13083 | 0.23231 | 11.875 | 0.426091362 | 27.4 | Neutral |
| testneg_9888 | 0 | 0 | 1.011117 | -3 | 0.310965 | -3 | 0.15524 | 0.10607 | 0.05087 | 0.09682 | 23.40643 | 0.01 | 0 | Neutral |
| testneg_9889 | 0.16 | -0.37 | 2.39997 | -3 | 2.970791 | -3 | 0.08011 | 0.12788 | 0.15685 | 0.17839 | 28.51351 | 0.229347854 | 0 | Neutral |
| testneg_9890 | 0 | 0 | 0.432421 | -3 | 3.032676 | -3 | 0.0922 | 0.07873 | 0.10895 | 0.11082 | 22.4104 | 0.04 | 0 | Neutral |
| testneg_9891 | -0.06 | 0.57 | 1.920471 | -3 | 1.120506 | -3 | 0.23809 | 0.18937 | 0.16547 | 0.16092 | 12.29518 | 0.158978228 | 0 | Neutral |
| testneg_9892 | -0.08 | -0.03 | 1.582915 | -3 | 0.021671 | -3 | 0.09493 | 0.104 | 0.13156 | 0.10621 | 39.98223 | 0.01 | 0 | Neutral |
| testneg_9893 | 0.02 | -0.63 | 2.813857 | -3 | 1.052366 | -3 | 0.1017 | 0.11809 | 0.1205 | 0.23508 | 32.18069 | 0.13 | 0 | Neutral |
| testneg_9894 | 0.03 | 0.1 | -1.665414 | -3 | -0.398253 | -3 | 0.19898 | 0.12788 | 0.04668 | 0.12788 | 20.81729 | 0.02 | 0 | Neutral |
| testneg_9895 | 0.14 | -1.01 | 6.193403 | -3 | 0.840482 | -3 | 0.04508 | 0.12788 | 0.1294 | 0.10964 | 22.15335 | 0.17 | 0 | Neutral |
| testneg_9896 | 0 | 0 | 1.23569 | -3 | 5.542651 | -3 | 0.64914 | 0.12481 | 0.13708 | 0.81545 | 24.14343 | 0.335257937 | 4.2 | Neutral |
| testneg_9897 | -0.08 | -0.03 | 0.177819 | -3 | 0.4795 | -3 | 0.18753 | 0.09983 | 0.0935 | 0.0819 | 29.4055 | 0.01 | 0 | Neutral |
| testneg_9898 | -0.02 | 0.36 | -0.000915 | -3 | 1.104931 | -3 | 0.10349 | 0.12788 | 0.06708 | 0.09112 | 22.77778 | 0.02 | 0 | Neutral |
| testneg_9899 | 0 | 0 | 1.495836 | -3 | 0.169559 | -3 | 0.24977 | 0.11262 | 0.86166 | 0.14254 | 17.5116 | 0.02 | 0 | Neutral |
| testneg_9900 | -0.08 | -0.03 | 1.50021 | -3 | 0.368677 | -0.054702 | 0.27618 | 0.11809 | 0.02518 | 0.07165 | 21.9697 | 0.02 | 0 | Neutral |
| testneg_9901 | 0.04 | -0.23 | 5.956747 | -3 | 0.460446 | -3 | 0.19898 | 0.15588 | 0.25077 | 0.06261 | 10.41667 | 0.191666667 | 0.2 | Neutral |
| testneg_9902 | -0.01 | 1.44 | -1.970817 | -3 | 1.336135 | -3 | 0.2058 | 0.28174 | 0.17935 | 0.36953 | 9.16566 | 0.24 | 0.2 | Neutral |
| testneg_9903 | 0.03 | 0.1 | 3.638601 | -3 | 0.500297 | -3 | 0.9149 | 0.27793 | 0.86437 | 0.69972 | 10.41667 | 0.228873626 | 0.4 | Neutral |
| testneg_9904 | -0.04 | 0.23 | 0.6838 | 0.6838 | 2.031708 | -3 | 0.67327 | 0.23543 | 0.13061 | 0.21023 | 20 | 0.21 | 0.2 | Neutral |
| testneg_9905 | 0.05 | 0.29 | 2.437307 | -3 | 1.092413 | -3 | 0.3047 | 0.11929 | 0.96396 | 0.7628 | 5 | 0.070565757 | 0 | Neutral |
| testneg_9906 | 0 | 0 | 1.848117 | -3 | 5.528516 | -3 | 0.10249 | 0.1399 | 0.21399 | 0.10708 | 21.02339 | 0.134060458 | 0 | Neutral |
| testneg_9907 | 0.08 | 0.03 | 3.085025 | -3 | 1.475091 | -3 | 0.83067 | 0.10583 | 0.3414 | 0.13575 | 28.80628 | 0.12 | 0 | Neutral |
| testneg_9908 | 0 | 0 | 0.386087 | 0.386087 | 3.805663 | -3 | 0.26927 | 0.10752 | 0.24519 | 0.12788 | 15.37063 | 0.05061553 | 0 | Neutral |
| testneg_9909 | -0.07 | -0.7 | 3.861511 | -3 | 2.326926 | -3 | 0.09018 | 0.16175 | 0.07558 | 0.12788 | 10.83333 | 0.185611111 | 0 | Neutral |
| testneg_9910 | 0.09 | -0.3 | 1.031921 | -3 | 2.53141 | 2.53141 | 0.96704 | 0.20571 | 0.1123 | 0.1568 | 7.77778 | 0.290034965 | 1.6 | Neutral |
| testneg_9911 | 0 | 0 | 0.759983 | -3 | 4.821873 | -3 | 0.13505 | 0.09019 | 0.78664 | 0.11725 | 16.22204 | 0.05 | 0 | Neutral |
| testneg_9912 | 0 | -0.01 | 1.668898 | -3 | 0.31188 | -3 | 0.48027 | 0.52799 | 0.14349 | 0.27976 | 6.98716 | 0.081362745 | 0 | Neutral |
| testneg_9913 | 0 | 0.32 | -0.420859 | -3 | 2.959696 | -3 | 0.0597 | 0.18092 | 0.19898 | 0.15588 | 18.03534 | 0.289639355 | 2 | Neutral |
| testneg_9914 | 0.07 | 0.38 | -1.249264 | -3 | 0.331991 | -3 | 0.05904 | 0.09282 | 0.11571 | 0.17324 | 25.62301 | 0.12 | 0 | Neutral |
| testneg_9915 | 0 | -0.01 | 1.82373 | -3 | 2.005102 | -3 | 0.07428 | 0.52467 | 0.12949 | 0.44208 | 10 | 0.220473579 | 0 | Neutral |
| testneg_9916 | -0.03 | 1.03 | 0.93405 | -3 | 1.885526 | 1.885526 | 0.05677 | 0.06758 | 0.11233 | 0.28224 | 10 | 0.19 | 0 | Neutral |
| testneg_9917 | -0.03 | 1.03 | 4.336137 | -3 | -1.08606 | -3 | 0.19898 | 0.12788 | 0.28269 | 0.12505 | 22.24612 | 0.070434028 | 0 | Neutral |
| testneg_9918 | 0.02072727 | 0.05618182 | 10.04388 | -3 | 4.686021 | -3 | 0.10617 | 0.15182 | 0.57564 | 0.12788 | 11.25 | 0.703743386 | 98.6 | Disease_causing |
| testneg_9919 | 0.16 | -0.37 | 6.462793 | -3 | 1.588676 | -3 | 0.16646 | 0.16629 | 0.19894 | 0.27411 | 17.31482 | 0.582483333 | 80 | Disease_causing |
| testneg_9920 | 0.11 | -0.19 | 5.286102 | -3 | 2.055296 | -3 | 0.83067 | 0.10583 | 0.15092 | 0.09074 | 24.29414 | 0.14 | 0 | Neutral |
| testneg_9921 | -0.16 | -0.08 | 2.815622 | -3 | 3.355715 | -3 | 0.46037 | 0.1343 | 0.35026 | 0.09817 | 24.90817 | 0.155645833 | 0 | Neutral |
| testneg_9922 | 0 | 0 | 1.38626 | -3 | 0.051943 | -3 | 0.50248 | 0.10939 | 0.48027 | 0.52799 | 5 | 0.022470519 | 0 | Neutral |
| testneg_9923 | 0.09 | -0.3 | 4.85311 | -3 | -0.020693 | -3 | 0.47379 | 0.18145 | 0.80034 | 0.10518 | 13.14653 | 0.099940476 | 0 | Neutral |
| testneg_9924 | 0 | -0.01 | 7.512937 | 7.512937 | 0.905656 | -3 | 0.56315 | 0.13761 | 0.30585 | 0.10254 | 20.1182 | 0.68 | 98.2 | Disease_causing |
| testneg_9925 | -0.05 | -0.29 | -1.131173 | -3 | -0.176493 | -3 | 0.99448 | 0.67288 | 0.95978 | 0.09493 | 14.21227 | 0.0735 | 0 | Neutral |
| testneg_9926 | 0.03 | 1.22 | 1.790281 | -3 | 0.196098 | -3 | 0.02059 | 0.066 | 0.12682 | 0.11357 | 17.91815 | 0.075090909 | 0 | Neutral |
| testneg_9927 | 0.04 | 2.01 | 0.838864 | -3 | 0.27289 | -3 | 0.56307 | 0.10536 | 0.14668 | 0.25103 | 19.77778 | 0.14 | 0 | Neutral |
| testneg_9928 | -0.14 | 0.37 | 3.214942 | -3 | -0.480501 | -3 | 0.10236 | 0.09622 | 0.14704 | 0.12788 | 23.70087 | 0.06 | 0 | Neutral |
| testneg_9929 | 0 | 0 | 1.786425 | -3 | 1.970217 | -3 | 0.27942 | 0.73513 | 0.27246 | 0.12788 | 7.79105 | 0.031483516 | 0 | Neutral |
| testneg_9930 | -0.15 | 0.28 | 3.6541 | -3 | 5.180313 | 5.180313 | 0.19898 | 0.37148 | 0.19898 | 0.12788 | 17.77778 | 0.639703947 | 94.4 | Disease_causing |
| testneg_9931 | -0.04 | 0.23 | 4.755795 | -3 | 0.172519 | -3 | 0.06877 | 0.17777 | 0.85887 | 0.1489 | 20.2381 | 0.265777778 | 0.4 | Neutral |
| testneg_9932 | 0 | 0 | 0.082683 | -3 | 0.815573 | -3 | 0.19898 | 0.12788 | 0.19898 | 0.12788 | 20.04433 | 0 | 0 | Neutral |
| testneg_9933 | -0.06 | 0.57 | 0.172291 | -3 | -0.481271 | -0.481271 | 0.0138 | 0.06769 | 0.09332 | 0.53331 | 10.08333 | 0.145362408 | 0 | Neutral |
| testneg_9934 | 0.14 | -0.37 | 2.51438 | -3 | 1.146824 | -3 | 0.06118 | 0.08724 | 0.77364 | 0.12676 | 10.41667 | 0.030400641 | 0 | Neutral |
| testneg_9935 | 0 | 0 | 1.485335 | -3 | 6.860523 | -3 | 0.19894 | 0.27411 | 0.10084 | 0.12788 | 12.34649 | 0.345013889 | 8.8 | Neutral |
| testneg_9936 | 0.2 | -1.26 | 2.305488 | -3 | 0.54521 | -3 | 0.18979 | 0.07948 | 0.3718 | 0.2248 | 18.33333 | 0.12 | 0 | Neutral |
| testneg_9937 | -0.03 | -0.1 | 1.750131 | -3 | 3.103812 | -3 | 0.85075 | 0.16646 | 0.0693 | 0.09293 | 13.75 | 0.077762238 | 0 | Neutral |
| testneg_9938 | 0 | 0 | 4.656753 | -3 | 4.377214 | 0.562475 | 0.52388 | 0.25831 | 0.11657 | 0.21661 | 10.30752 | 0.716275808 | 98.4 | Disease_causing |
| testneg_9939 | 0.09 | -0.3 | 6.809894 | -3 | 3.849548 | -3 | 0.43918 | 0.46357 | 0.36257 | 0.26115 | 18.88889 | 0.920570085 | 100 | Disease_causing |
| testneg_9940 | 0 | 0 | 0.185804 | -3 | 3.381673 | 3.381673 | 0.08306 | 0.12788 | 0.23578 | 0.12788 | 16.8018 | 0.150291986 | 0 | Neutral |
| testneg_9941 | -0.07 | -0.38 | 5.043975 | -3 | -1.359951 | -3 | 0.1296 | 0.164 | 0.47816 | 0.66526 | 20 | 0.399496337 | 17.8 | Neutral |
| testneg_9942 | 0 | 0 | 1.173845 | -3 | 0.854923 | -3 | 0.88689 | 0.41514 | 0.11458 | 0.16496 | 5 | 0.056632475 | 0 | Neutral |
| testneg_9943 | -0.21 | -0.15 | 5.077905 | -3 | -1.327088 | -3 | 0.18022 | 0.12788 | 0.22225 | 0.11186 | 19.5977 | 0.1 | 0 | Neutral |
| testneg_9944 | 0.18 | -1.05 | 8.297965 | -3 | 1.334446 | 1.334446 | 0.71762 | 0.14417 | 0.15514 | 0.14852 | 13.51351 | 0.490166667 | 52.4 | Disease_causing |
| testneg_9945 | 0 | 0 | 2.209811 | -3 | 1.8794 | -3 | 0.1647 | 0.12076 | 0.54711 | 0.11809 | 14.1885 | 0 | 0 | Neutral |
| testneg_9946 | 0.15 | -0.28 | -0.431211 | -3 | -0.58291 | -3 | 0.36676 | 0.37749 | 0.64608 | 0.35524 | 5 | 0.180702381 | 0 | Neutral |
| testneg_9947 | 0 | 0 | 2.53141 | 2.53141 | 0.009894 | -3 | 0.1123 | 0.1568 | 0.07996 | 0.08354 | 33.2671 | 0.180540936 | 0.2 | Neutral |
| testneg_9948 | 0 | 0 | 0.922494 | -3 | 0.095976 | -3 | 0.53203 | 0.12788 | 0.11522 | 0.11925 | 21.35057 | 0.02 | 0 | Neutral |
| testneg_9949 | 0.01 | 0.79 | 2.324267 | -3 | 2.94636 | -3 | 0.19898 | 0.94603 | 0.5958 | 0.0929 | 10.83333 | 0.19 | 0 | Neutral |
| testneg_9950 | 0.06 | -0.57 | 2.053078 | -3 | -0.4932 | -0.4932 | 0.07014 | 0.53339 | 0.20379 | 0.43705 | 10 | 0.270517979 | 0.2 | Neutral |
| testneg_9951 | -0.02 | -0.45 | 1.767422 | -3 | 0.432868 | -3 | 0.41698 | 0.10384 | 0.37292 | 0.11979 | 15.15809 | 0.000606061 | 0 | Neutral |
| testneg_9952 | -0.07 | -0.7 | 0.232874 | -3 | 5.021764 | -3 | 0.14901 | 0.12788 | 0.30684 | 0.12788 | 22.87281 | 0.090424448 | 0 | Neutral |
| testneg_9953 | 0.09 | -0.3 | -0.213037 | -3 | 0.034534 | 0.034534 | 0.54113 | 0.11017 | 0.62381 | 0.9477 | 19.06273 | 0.14 | 0 | Neutral |
| testneg_9954 | 0.21 | 0.15 | 1.305251 | -3 | 0.488199 | -3 | 0.06304 | 0.12788 | 0.23088 | 0.12788 | 29.08689 | 0.03 | 0 | Neutral |
| testneg_9955 | 0 | 0 | -0.450207 | -3 | 0.686233 | -3 | 0.17083 | 0.31043 | 0.17969 | 0.11473 | 13.15023 | 0.01153568 | 0 | Neutral |
| testneg_9956 | 0.04 | -0.23 | 4.017255 | -3 | -3.138593 | -3 | 0.19898 | 0.16603 | 0.06254 | 0.23561 | 11.61005 | 0.261506738 | 0.4 | Neutral |
| testneg_9957 | 0.05 | 0.29 | 3.783684 | -3 | 0.975564 | -3 | 0.39399 | 0.12788 | 0.16734 | 0.15176 | 23.65801 | 0.123956583 | 0 | Neutral |
| testneg_9958 | 0 | 0 | 0.707638 | -3 | 6.628365 | -1.271425 | 0.62339 | 0.19592 | 0.10511 | 0.12788 | 10.83333 | 0.240452863 | 1.6 | Neutral |
| testneg_9959 | 0 | 0 | 0.755726 | -3 | -1.27031 | -3 | 0.08454 | 0.22908 | 0.10977 | 0.31695 | 10 | 0.13034493 | 0 | Neutral |
| testneg_9960 | 0 | 0 | 1.713577 | -3 | 1.305054 | -3 | 0.7174 | 0.15815 | 0.16248 | 0.14352 | 10.81031 | 0.05 | 0 | Neutral |
| testneg_9961 | -0.04 | 0.23 | 3.778972 | -3 | 0.275288 | -3 | 0.48574 | 0.20473 | 0.6667 | 0.20993 | 17.21847 | 0.237090376 | 0.2 | Neutral |
| testneg_9962 | 0.15 | -1.06 | 4.643757 | -3 | 1.241234 | -3 | 0.113 | 0.12788 | 0.6258 | 0.16461 | 17.79882 | 0.239486492 | 0.4 | Neutral |
| testneg_9963 | -0.2 | 1.26 | -0.70557 | -0.70557 | -0.270969 | -3 | 0.29008 | 0.19268 | 0.04441 | 0.13603 | 12.29167 | 0.23 | 0.2 | Neutral |
| testneg_9964 | 0.19 | 0.51 | 1.770771 | -3 | 0.82266 | -3 | 0.71406 | 0.2012 | 0.07923 | 0.12788 | 10.41667 | 0.0725 | 0 | Neutral |
| testneg_9965 | -0.03 | -0.12 | -0.107462 | -3 | -0.044628 | -3 | 0.07982 | 0.12788 | 0.62339 | 0.19592 | 29.14047 | 0.080215278 | 0 | Neutral |
| testneg_9966 | 0.02 | 0.41 | -2.259592 | -3 | -1.290562 | -3 | 0.06311 | 0.14802 | 0.19898 | 0.12895 | 39.93056 | 0.06 | 0 | Neutral |
| testneg_9967 | 0 | 0 | 1.095746 | -3 | 1.1308 | -3 | 0.24274 | 0.13034 | 0.19898 | 0.11185 | 46.82901 | 0.01 | 0 | Neutral |
| testneg_9968 | 0 | 0 | 1.726432 | -3 | 5.815338 | -3 | 0.15192 | 0.14302 | 0.05932 | 0.09551 | 14.12826 | 0.131585498 | 0 | Neutral |
| testneg_9969 | -0.02 | -0.45 | 2.813678 | -3 | -0.305503 | -3 | 0.755 | 0.12788 | 0.55197 | 0.11305 | 19.44444 | 0.02 | 0 | Neutral |
| testneg_9970 | 0 | 0 | 0.870722 | -3 | -2.083041 | -3 | 0.1193 | 0.5592 | 0.10383 | 0.08853 | 19.52521 | 0.07 | 0 | Neutral |
| testneg_9971 | 0 | 0 | -1.056572 | -3 | 2.906447 | -3 | 0.1109 | 0.17295 | 0.15839 | 0.10522 | 32.52827 | 0.07 | 0 | Neutral |
| testneg_9972 | -0.01 | 0.57 | -2.581564 | -3 | -0.121102 | -0.121102 | 0.19898 | 0.12788 | 0.78935 | 0.17351 | 10.22898 | 0.050743467 | 0 | Neutral |
| testneg_9973 | 0 | 0 | 2.047211 | -3 | 2.30425 | -3 | 0.16823 | 0.10796 | 0.46792 | 0.11262 | 17.73615 | 0.01 | 0 | Neutral |
| testneg_9974 | 0.01755556 | 0.21888889 | 7.812451 | -3 | -2.464297 | -2.464297 | 0.7219 | 0.41509 | 0.11524 | 0.12788 | 18.08176 | 0.46 | 38.6 | Neutral |
| testneg_9975 | 0.07 | 0.7 | 5.435134 | -3 | -0.493952 | -3 | 0.85508 | 0.12634 | 0.05444 | 0.12788 | 16.84579 | 0.16 | 0 | Neutral |
| testneg_9976 | 0 | -0.01 | 1.587876 | -3 | -0.01248 | -3 | 0.57019 | 0.38533 | 0.61231 | 0.13362 | 19.44444 | 0.104745872 | 0 | Neutral |
| testneg_9977 | 0.06 | -0.57 | 4.974068 | -3 | 7.30631 | -3 | 0.18423 | 0.10808 | 0.19898 | 0.12788 | 27.92726 | 0.47 | 45.2 | Neutral |
| testneg_9978 | 0.03 | 1.22 | 2.753966 | -3 | 1.44947 | -3 | 0.1551 | 0.09276 | 0.18459 | 0.33541 | 19.24501 | 0.160884354 | 0 | Neutral |
| testneg_9979 | 0 | 0 | -0.034419 | -0.034419 | 3.447686 | -3 | 0.62203 | 0.12788 | 0.19898 | 0.1054 | 7.01754 | 0.051510211 | 0 | Neutral |
| testneg_9980 | 0 | 0 | 0.528019 | -3 | 2.60676 | -3 | 0.42621 | 0.49093 | 0.09439 | 0.07415 | 17.21847 | 0.057467949 | 0 | Neutral |
| testneg_9981 | -0.04 | 0.23 | 3.260847 | -3 | -0.354552 | -3 | 0.27605 | 0.12788 | 0.09815 | 0.12788 | 22.89111 | 0.03 | 0 | Neutral |
| testneg_9982 | -0.05 | -0.29 | 4.5578 | -3 | 4.617328 | -3 | 0.77364 | 0.12676 | 0.19898 | 0.12788 | 10.83333 | 0.269946237 | 1.8 | Neutral |
| testneg_9983 | -0.03 | -0.1 | 1.177705 | -3 | 3.584282 | -3 | 0.0501 | 0.12788 | 0.18559 | 0.09173 | 21.341 | 0.08 | 0 | Neutral |
| testneg_9984 | -0.06 | 0.57 | 1.042833 | -3 | 3.302975 | -3 | 0.19898 | 0.12788 | 0.098 | 0.08791 | 16.8018 | 0.060602467 | 0 | Neutral |
| testneg_9985 | -0.15 | 1.06 | 5.350836 | -3 | 2.670295 | -3 | 0.30367 | 0.41964 | 0.09807 | 0.24901 | 10.41667 | 0.786399849 | 100 | Disease_causing |
| testneg_9986 | 0.16 | -0.37 | 3.291397 | -3 | 3.321188 | -3 | 0.46387 | 0.17039 | 0.82131 | 0.12788 | 20.6181 | 0.234935897 | 0.6 | Neutral |
| testneg_9987 | -0.03 | -0.1 | 3.544108 | 2.688968 | 5.582498 | -3 | 0.13974 | 0.10225 | 0.85829 | 0.4292 | 11.22104 | 0.729232143 | 99.8 | Disease_causing |
| testneg_9988 | 0 | -0.73 | 2.726423 | -3 | 2.248402 | -3 | 0.14273 | 0.1162 | 0.37424 | 0.12788 | 5.69444 | 0.030574534 | 0 | Neutral |
| testneg_9989 | 0 | 0 | 1.38626 | -3 | 1.738935 | -3 | 0.50248 | 0.10939 | 0.07571 | 0.09432 | 10.23791 | 0.01 | 0 | Neutral |
| testneg_9990 | 0.21 | 0.15 | 4.953102 | -3 | 5.892448 | -3 | 0.05904 | 0.09282 | 0.10385 | 0.09775 | 33.53535 | 0.290083333 | 3 | Neutral |
| testneg_9991 | 0 | 0 | -0.117483 | -3 | 2.11694 | -3 | 0.06793 | 0.12788 | 0.40403 | 0.12788 | 18.8477 | 0.01 | 0 | Neutral |
| testneg_9992 | 0.21 | 0.15 | 3.757742 | -3 | 1.43948 | -3 | 0.25528 | 0.24339 | 0.40116 | 0.43982 | 9.12105 | 0.407577409 | 21.4 | Neutral |
| testneg_9993 | 0.04 | -0.23 | 4.579603 | -3 | 0.710328 | -3 | 0.19898 | 0.09927 | 0.13007 | 0.12788 | 13.51351 | 0.03 | 0 | Neutral |
| testneg_9994 | 0 | 0 | 0.216807 | -3 | 6.418366 | -3 | 0.16475 | 0.26069 | 0.20513 | 0.09801 | 11.65872 | 0.233748747 | 1.2 | Neutral |
| testneg_9995 | 0.02 | -0.36 | 1.555382 | 1.555382 | 6.994643 | -3 | 0.80134 | 0.08685 | 0.22234 | 0.09474 | 28.96994 | 0.25 | 0.4 | Neutral |
| testneg_9996 | -0.03 | -0.1 | -0.432565 | -3 | -0.335332 | -3 | 0.1105 | 0.44945 | 0.50388 | 0.12295 | 14.6265 | 0.06125 | 0 | Neutral |
| testneg_9997 | -0.01 | -0.8 | 1.016479 | 1.016479 | 3.170156 | -3 | 0.12649 | 0.12788 | 0.10084 | 0.12788 | 10 | 0.09 | 0 | Neutral |
| testneg_9998 | 0 | 0 | 1.941287 | -3 | -0.502305 | -1.422138 | 0.24977 | 0.11262 | 0.16091 | 0.90971 | 22.88801 | 0.1 | 0 | Neutral |
| testneg_10001 | 0.04 | -0.23 | 4.680114 | -3 | 2.653525 | -3 | 0.31124 | 0.08875 | 0.20546 | 0.10734 | 16.8018 | 0.088798077 | 0 | Neutral |
| testneg_10002 | 0 | 0 | 1.186882 | -3 | 0.144637 | -3 | 0.37463 | 0.23021 | 0.20657 | 0.12788 | 10.83333 | 0.010336348 | 0 | Neutral |
