## Supplementary material for "Predicting disease-causing variant combinations": Dataset S5

| Posey_combination_ID | Flex1 | Hydr1 | CADD1 | CADD2 | CADD3 | CADD4 | HI_A | RecA | HI_B | RecB | Biol_Dist | Classification_score | Support_score | Predicted_class |
|---|---|---|---|---|---|---|---|---|---|---|---|---|---|---|
| 1 | 0.01 | 0.8 | 7.466843 | -3 | 8.857964 | -3 | 0.1228 | 0.23648 | 0.45035 | 0.88273 | 10 | 0.88 | 100 | Disease_causing |
| 2 | 0 | -0.01 | 6.939009 | -3 | 3.645827 | -3 | 0.2494 | 0.10736 | 0.08975 | 0.12788 | 11.25 | 0.47775641 | 45.2 | Neutral |
| 4 | 0 | -0.01 | 5.1686335 | -3 | 3.742048 | -3 | 0.58012 | 0.48908 | 0.15152 | 0.12645 | 6.2588 | 0.422079151 | 23.8 | Neutral |
| 5 | 0 | 0 | 2.819812 | -3 | 11.366191 | -3 | 0.15044 | 0.76645 | 0.10816 | 0.12788 | 19.35743 | 0.590927098 | 87.4 | Disease_causing |
| 6 | -0.16 | 1.37 | 3.705321 | -3 | 7.08488 | -3 | 0.15798 | 0.17331 | 0.58636 | 0.55905 | 10.17809 | 0.810016938 | 100 | Disease_causing |
| 8 | 0 | -0.01 | 5.063158 | -3 | 13.116067 | -3 | 0.69297 | 0.12788 | 0.71112 | 0.12031 | 10 | 0.575784884 | 83.8 | Disease_causing |
| 9 | -0.0087273 | 0.00581818 | 10.530907 | -3 | 1.402203 | -3 | 0.29024 | 0.1472 | 0.85262 | 0.49245 | 22.81414 | 0.520762393 | 66.4 | Disease_causing |
| 10 | 0 | -0.01 | 8.374283 | -3 | 2.532432 | -3 | 0.19738 | 0.13472 | 0.99999 | 0.5507 | 9.03073 | 0.54 | 70.4 | Disease_causing |
| 11 | 0.00145455 | 0.43818182 | 12.82818 | -3 | 2.028319 | -3 | 0.19898 | 0.94603 | 0.21825 | 0.67914 | 4.72222 | 0.72 | 99.6 | Disease_causing |
| 12 | 0.04 | -0.23 | 3.713686 | -3 | 4.532255 | -3 | 0.37121 | 0.20079 | 0.8338 | 0.11376 | 17.77778 | 0.315218275 | 5 | Neutral |
| 13 | -0.06 | -0.18 | 5.616767 | -3 | 3.645827 | -3 | 0.97825 | 0.09412 | 0.08975 | 0.12788 | 10.41667 | 0.371505435 | 11 | Neutral |
| 14 | 0.22 | -2.66 | 7.635711 | -3 | 3.645827 | -3 | 0.26293 | 0.1149 | 0.08975 | 0.12788 | 12.42293 | 0.49 | 51.8 | Disease_causing |
| 15 | -0.01 | -0.8 | 5.658066 | -3 | 1.320816 | -3 | 0.98629 | 0.45051 | 0.53569 | 0.21387 | 11.25 | 0.611403941 | 91 | Disease_causing |
| 17 | -0.07 | -0.01 | 2.092924 | -3 | 7.899259 | -3 | 0.97757 | 0.34098 | 0.6458 | 0.34375 | 11.66667 | 0.577736111 | 80 | Disease_causing |
| 18 | 0.01509091 | -0.3487273 | 10.775339 | -3 | 6.987202 | -3 | 0.19898 | 0.08201 | 0.60193 | 0.13678 | 12.5836 | 0.678455882 | 96 | Disease_causing |
| 19 | 0.15 | 0.06 | 6.536498 | -3 | 4.87919 | -3 | 0.26245 | 0.7289 | 0.49106 | 0.89143 | 4.44444 | 0.852906236 | 100 | Disease_causing |
| 20 | 0.14 | -1.01 | 4.297717 | -3 | 6.597932 | -3 | 0.25753 | 0.21408 | 0.99994 | 0.51636 | 12.29167 | 0.805719355 | 100 | Disease_causing |
| 23 | 0 | -0.01 | 6.363192 | -3 | 9.145308 | -3 | 0.30697 | 0.08082 | 0.49147 | 0.15128 | 20.27778 | 0.64 | 91.6 | Disease_causing |
| 24 | 0.04 | -0.23 | 7.134717 | -3 | 8.776734 | -3 | 0.18869 | 0.23984 | 0.2494 | 0.10736 | 11.25 | 0.871949301 | 100 | Disease_causing |
| 25 | -0.0243636 | 0.02509091 | 13.092159 | -3 | 1.215018 | -3 | 0.7413 | 0.18974 | 0.5402 | 0.78677 | 10.20833 | 0.61875652 | 94.2 | Disease_causing |
| 26 | 0 | -0.01 | 6.680055 | -3 | 2.774072 | -3 | 0.15958 | 0.41628 | 0.35429 | 0.34658 | 10.83333 | 0.837321419 | 100 | Disease_causing |
| 27 | -0.05 | -0.29 | 3.526437 | -3 | 5.485505 | -3 | 0.6031 | 0.25451 | 0.49147 | 0.15128 | 11.66667 | 0.728044932 | 99.6 | Disease_causing |
| 28 | -0.14 | 0.37 | 3.04257 | -3 | 3.989005 | -3 | 0.85252 | 0.83576 | 0.6001 | 0.90845 | 5 | 0.611575692 | 89.2 | Disease_causing |
| 29 | -0.0467273 | 0.06472727 | 10.77899 | -3 | 2.540059 | -3 | 0.95805 | 0.25081 | 0.89178 | 0.23115 | 12.72872 | 0.684908964 | 99 | Disease_causing |
| 33 | 0.00927273 | -0.0987273 | 10.137813 | 7.155611 | 6.078869 | -3 | 0.42478 | 0.12788 | 0.7551 | 0.29962 | 15.97222 | 0.869527778 | 100 | Disease_causing |
| 34 | -0.08 | -0.03 | 3.99134 | -3 | 22.161135 | 1.414477 | 0.10242 | 0.10521 | 0.40699 | 0.38418 | 18.16 | 0.745095238 | 100 | Disease_causing |
| 36 | -0.1 | -1.9 | 0.782101 | -3 | 4.908936 | -0.032748 | 0.92981 | 0.12788 | 0.14612 | 0.10979 | 6.94444 | 0.131017241 | 0 | Neutral |
| 37 | 0.2 | -1.26 | 4.908739 | -3 | 6.626073 | 2.467515 | 0.78691 | 0.12202 | 0.42478 | 0.12788 | 20.74074 | 0.48 | 47.8 | Neutral |
| 38 | -0.0018182 | -0.166 | 11.850573 | -3 | 2.156893 | -0.19769 | 0.76491 | 0.10651 | 0.48891 | 0.98177 | 14.16667 | 0.529723485 | 67.8 | Disease_causing |
| 39 | 0.07 | 0.7 | 6.240871 | -3 | 5.072541 | 2.975482 | 0.24762 | 0.17859 | 0.15704 | 0.72174 | 10 | 0.888103318 | 100 | Disease_causing |
| 40 | 0 | 0 | 0.77228 | -3 | 8.416626 | 8.416626 | 0.19898 | 0.08201 | 0.30256 | 0.24991 | 10.41667 | 0.528301282 | 66.4 | Disease_causing |
| 41 | 0.07 | 0.7 | 6.28779 | -3 | 4.268765 | 4.268765 | 0.755 | 0.12788 | 0.09022 | 0.12788 | 29.16667 | 0.527619048 | 71.2 | Disease_causing |
| 42 | 0 | 0 | 6.270361 | 4.812251 | -0.05658 | -3 | 0.11138 | 0.59493 | 0.69282 | 0.10507 | 17.08254 | 0.59 | 88.8 | Disease_causing |
| 44 | 0.19 | -0.25 | 5.969171 | -3 | 11.179158 | 11.179158 | 0.6001 | 0.90845 | 0.22774 | 0.12788 | 10.83333 | 0.86 | 100 | Disease_causing |
| 45 | 0 | 0 | 3.417808 | -3 | 11.394229 | 6.242526 | 0.24714 | 0.14229 | 0.17569 | 0.10035 | 18.16 | 0.716904762 | 99.8 | Disease_causing |
| 46 | -0.15 | -0.06 | 2.328162 | -3 | 7.905851 | 2.495704 | 0.6031 | 0.25451 | 0.06035 | 0.22441 | 17.63514 | 0.682942152 | 98.2 | Disease_causing |
| 48 | 0.02036364 | -0.2292727 | 12.858377 | -3 | 5.674633 | 5.674633 | 0.44902 | 0.09206 | 0.09064 | 0.12411 | 17.01711 | 0.679669082 | 99.6 | Disease_causing |
| 49 | 0.14 | -0.69 | 7.040864 | 5.724946 | -0.005019 | -3 | 0.82422 | 0.12092 | 0.44861 | 0.11578 | 29.51573 | 0.58 | 81.4 | Disease_causing |
| 50 | 0.21 | 0.15 | 7.222104 | -3 | 6.750037 | 5.689983 | 0.64721 | 0.8758 | 0.27943 | 0.89315 | 10.41667 | 0.867883721 | 100 | Disease_causing |
| 52 | -0.0254545 | 0.31818182 | 13.86265 | 13.86265 | -1.130517 | -3 | 0.12989 | 0.20455 | 0.21968 | 0.12788 | 10.83333 | 0.760839161 | 100 | Disease_causing |
| 53 | 0 | -0.01 | 4.634264 | 4.634264 | 0.193503 | -3 | 0.23813 | 0.12788 | 0.08975 | 0.12788 | 17.21847 | 0.484642857 | 49.8 | Neutral |
| 54 | 0.03 | -1.03 | 5.704354 | -3 | 7.184713 | 7.184713 | 0.19898 | 0.70074 | 0.13274 | 0.12788 | 19.89489 | 0.85 | 100 | Disease_causing |
| 55 | 0.2 | -1.26 | 5.413374 | -3 | 5.396431 | 4.99961 | 0.12366 | 0.32056 | 0.19452 | 0.79779 | 6.94444 | 0.879659155 | 100 | Disease_causing |
| 57 | 0.18 | -1.05 | 7.613165 | -3 | 6.047161 | 6.047161 | 0.54329 | 0.14172 | 0.65897 | 0.67602 | 18.21114 | 0.86004329 | 100 | Disease_causing |
| 58 | 0 | -0.01 | 4.71357 | -3 | 2.264129 | -1.131836 | 0.97757 | 0.34098 | 0.75964 | 0.61682 | 10.41667 | 0.413609391 | 20.2 | Neutral |
| 59 | 0.07236364 | -0.1945455 | 10.726534 | -3 | 6.272062 | -3 | 0.72124 | 0.09906 | 0.71112 | 0.12031 | 8.1746 | 0.638090909 | 95.2 | Disease_causing |
| 61 | 0.2 | -1.26 | 4.427217 | -3 | 5.200633 | -3 | 0.37858 | 0.98951 | 0.15152 | 0.12645 | 14.26037 | 0.465246212 | 39.8 | Neutral |
| 62 | -0.07 | 1.12 | 6.611733 | -3 | 7.406018 | 7.406018 | 0.14151 | 0.13965 | 0.3401 | 0.7637 | 17.77778 | 0.853397436 | 100 | Disease_causing |
| 64 | 0.03 | -1.03 | 3.568276 | -3 | 6.203487 | -3 | 0.40663 | 0.24974 | 0.27813 | 0.5595 | 14.16667 | 0.863690384 | 100 | Disease_causing |
| 65 | 0.01 | 0.8 | 3.717966 | -3 | 10.564686 | -3 | 0.95805 | 0.25081 | 0.35867 | 0.15462 | 22.12579 | 0.711662018 | 99.2 | Disease_causing |
| 66 | 0.14 | -1.01 | 6.177742 | 6.177742 | 5.810561 | -3 | 0.23488 | 0.19342 | 0.97757 | 0.34098 | 16.29902 | 0.88 | 100 | Disease_causing |
| 67 | 0.22 | -2.66 | 6.806716 | 6.806716 | 8.465588 | -3 | 0.10697 | 0.0807 | 0.9964 | 0.10708 | 10 | 0.71 | 100 | Disease_causing |
| 68 | -0.04 | 1.61 | 6.162695 | -3 | 4.938791 | 4.938791 | 0.44258 | 0.47751 | 0.19898 | 0.12788 | 18.16 | 0.78 | 100 | Disease_causing |
| 69 | 0.03 | 0.12 | 7.135263 | -3 | 3.56366 | 3.56366 | 0.80941 | 0.50577 | 0.32307 | 0.28713 | 4.44444 | 0.859122459 | 99.8 | Disease_causing |
| 70 | 0.05 | 0.29 | 5.70919 | -3 | 4.502333 | 4.502333 | 0.25753 | 0.21408 | 0.27813 | 0.5595 | 18.88889 | 0.895114286 | 100 | Disease_causing |
| 72 | 0 | 0 | 5.068284 | -3 | 7.00558 | -3 | 0.80264 | 0.11567 | 0.84682 | 0.35105 | 10 | 0.623583333 | 90 | Disease_causing |
| 73 | 0.04 | -0.23 | 4.997978 | 4.997978 | 0.958319 | -3 | 0.75981 | 0.33817 | 0.14584 | 0.25273 | 10.41667 | 0.75 | 99.4 | Disease_causing |
| 74 | 0.14 | -1.01 | 6.534411 | 6.534411 | 3.43081 | -3 | 0.19898 | 0.56316 | 0.49846 | 0.17535 | 12.17125 | 0.937113095 | 100 | Disease_causing |
| 75 | 0.03 | 1.22 | 6.67069 | 6.67069 | 7.095051 | 7.095051 | 0.67467 | 0.42741 | 0.44349 | 0.24792 | 13.14394 | 0.939537037 | 100 | Disease_causing |
| 76 | 0.09 | -0.3 | 6.99527 | 6.99527 | 4.827067 | 4.827067 | 0.03572 | 0.18035 | 0.20473 | 0.08046 | 17.71961 | 0.74 | 100 | Disease_causing |
| 77 | 0.02 | -0.63 | 6.351519 | 6.351519 | 7.091883 | 1.395558 | 0.0821 | 0.07339 | 0.19658 | 0.22977 | 15.22436 | 0.79 | 100 | Disease_causing |
| 78 | 0 | 0 | 4.743094 | 4.743094 | 12.068674 | 12.068674 | 0.28185 | 0.48408 | 0.06646 | 0.17226 | 18.16 | 0.906212121 | 100 | Disease_causing |
| 79 | 0.12 | -1.16 | 5.387121 | 5.387121 | 7.009446 | 7.009446 | 0.87847 | 0.96194 | 0.18628 | 0.48162 | 5 | 0.87 | 100 | Disease_causing |
| 80 | 0.01 | 0.8 | 4.70155 | 4.70155 | 5.778517 | 5.778517 | 0.21341 | 0.10005 | 0.44349 | 0.24792 | 13.14394 | 0.903570707 | 100 | Disease_causing |
| 81 | 0 | -0.01 | 9.5087 | 9.5087 | 4.832137 | 4.832137 | 0.21677 | 0.1681 | 0.18235 | 0.15914 | 29.86111 | 0.869399351 | 100 | Disease_causing |
| 83 | 0.01018182 | 0.37709091 | 14.044668 | 4.938791 | 11.90521 | 7.728962 | 0.1025 | 0.16015 | 0.18328 | 0.12788 | 24.22595 | 0.89 | 100 | Disease_causing |
| 85 | 0.06 | 0.18 | 6.168946 | -3 | 5.596815 | 5.596815 | 0.43937 | 0.69428 | 0.10286 | 0.26819 | 18.33333 | 0.928379027 | 100 | Disease_causing |
| 86 | 0.2 | -1.26 | 4.427217 | -3 | 7.921575 | 7.921575 | 0.37858 | 0.98951 | 0.19898 | 0.40608 | 7.00691 | 0.837428634 | 100 | Disease_causing |
| 87 | 0 | -0.01 | 7.596737 | 7.596737 | 4.372054 | 4.372054 | 0.22224 | 0.4222 | 0.7413 | 0.18974 | 12.30981 | 0.88 | 100 | Disease_causing |
| 90 | -0.04 | 0.89 | 4.387764 | -3 | 2.653977 | 2.653977 | 0.95805 | 0.25081 | 0.76593 | 0.87681 | 11.875 | 0.501428571 | 55.8 | Disease_causing |
| 91 | 0.04 | -0.89 | 5.029824 | 5.029824 | 7.252694 | 7.252694 | 0.17911 | 0.58893 | 0.10957 | 0.58549 | 33.95664 | 0.92 | 100 | Disease_causing |
| 92 | 0.18 | -1.05 | 5.1686335 | -3 | 3.749683 | 2.4392 | 0.10149 | 0.12788 | 0.30252 | 0.17567 | 24.58333 | 0.619897959 | 90.6 | Disease_causing |
| 93 | -0.03 | -1.22 | 6.209004 | -3 | 8.193296 | 2.537395 | 0.18055 | 0.12564 | 0.66381 | 0.13039 | 18.33333 | 0.71 | 99.8 | Disease_causing |
| 94 | -0.0038182 | -0.2923636 | 6.622757 | 6.622757 | 11.057244 | 7.131784 | 0.81566 | 0.22678 | 0.18102 | 0.6537 | 28.14183 | 0.879259398 | 100 | Disease_causing |
| 95 | 0 | -0.01 | 6.998064 | 6.998064 | 6.747932 | 6.747932 | 0.4809 | 0.10114 | 0.50276 | 0.09638 | 29.94245 | 0.76 | 100 | Disease_causing |
